# Comparative gene regulatory network analysis in Alzheimer’s disease and major depressive disorder identifies shared core regulatory circuits

**DOI:** 10.1101/2023.02.23.529626

**Authors:** Hanne Puype, Joke Deschildre, Vanessa Vermeirssen

## Abstract

Alzheimer’s disease and major depressive disorder are prevalent, devastating conditions with limited treatment options. Recent insights suggest that despite distinct phenotypes, these disorders share similar processes such as neuroinflammation. However, the extent of overlapping biological processes and their underlying molecular mechanisms remain to be elucidated. Therefore, we adopted a computational systems biology approach to compare regulatory programs in the prefrontal cortex of both disorders. Leveraging publicly available RNA sequencing data on different human cohorts, both at bulk and single-cell level, and using diverse computational methodologies, we inferred gene regulatory networks, which model the molecular interactions between transcription factors and their target genes, and characterized cell-type-specific regulatory programs and biological pathways. We identified core regulatory circuits shared in both disorders, including transcription factors that play a pivotal role in microglial activation such as IKZF1, IRF8, RUNX1 and SPI1. Most of these transcription factors had a reported role in Alzheimer’s, but not in depression. We found several common pathways such as microglial activation, but also more disease-specific pathways. Through orthogonal data analysis, we were able to validate several of the predicted regulatory interactions in Alzheimer’s disease and major depressive disorder. In summary, our work revealed neuroinflammation and microglial activation in both diseases, under the control of shared core regulatory circuits. The potential relevance of these transcription factors and genes warrants additional investigation, especially in depression, offering possible novel therapeutic opportunities.

## 1. Introduction

Neurodegenerative and neuropsychiatric disorders like Alzheimer’s disease (AD) and major depressive disorder (MDD) have a high burden on global health and patients still lack effective treatment. Recent insights suggest that despite distinct phenotypes, these disorders share similar processes involved in neuroinflammation^1^. Neuroinflammation is defined by infiltrating leukocytes in the central nervous system (CNS) and activation of microglia^2^, brain-residing macrophages that are crucial for the development and functioning of the brain^1^. However, microglia can also release pro-inflammatory mediators, such as cytokines, causing neuronal and brain barriers damage, and recruiting pro-inflammatory immune cells, hence exacerbating this neuroinflammation^3^. Whether neuroinflammation is causative or consequential, and to what extent it plays a role in these disorders, is not yet elucidated. We define neuroinflammatory disorders here as neurodegenerative and neuropsychiatric disorders that portray neuroinflammation.

AD, which is mainly characterized by extracellular β-amyloid (Aβ) plaques and intracellular neurofibrillary tangles, is the most prevalent cause of dementia^4^. Microglia play a considerable role in protecting against AD development by clearing Aβ^5^. Nevertheless, activated microglia can induce a neurotoxic phenotype in astrocytes, leading to neurodegeneration, and the majority of identified risk genes for AD are preferentially expressed in microglia^5,6^. Notably, neurodegenerative disorders are accompanied by psychiatric comorbidities: in AD, depression is frequently seen^4^.

MDD is a neuropsychiatric disorder with a still largely unknown pathophysiology and is influenced by both genetic and environmental factors. MDD is highly heterogeneous in symptoms, physiological changes, and treatment response, and can be characterized by decreased concentrations of the neurotransmitters serotonin, noradrenaline and dopamine in synaptic clefts^7^. In addition, the hypothalamic-pituitary-adrenal axis can be defective^8^. Moreover, there can be significant reductions in the number, density and size of astrocytes in several brain regions^9^. There is also mounting evidence for the implication of the immune system in MDD^7^. Cytokines such as IL-1, IL-6 and TNF-α are overexpressed and there are higher levels of T-cells and neutrophils in the CNS and periphery of MDD patients^7,10^. Furthermore, activated microglia lead to chronic inflammation. Consequently, a neuroinflammation hypothesis is rising. However, as depression is highly heterogeneous in different patients, so is the neuroinflammation phenotype, with higher inflammatory levels correlating with treatment resistance or a more severe phenotype^10^.

Remarkably, three recent studies investigated the shared genetic etiology between AD and MDD using GWAS datasets^11–13^. In the first study, the researchers found a moderate level of polygenic overlap between the two disorders^11^. An enrichment of SNPs was found of which the proximal genes were involved in two major pathways; immune response and regulation of endocytosis, indicating that MDD and AD have some genetic overlap in immune aberrations. In the second study, they found 98 causal variants that overlapped between the two disorders and found that MDD was a predictor of AD, while this was less the case the other way around^12^. In the third study, the researchers again found a genetic overlap between AD and MDD^13^. Additionally, they discovered that DNA methylation, transcripts and proteins underly the contribution of MDD to AD.

Network inference on omics data retrieves systems-wide insights into disease mechanisms. Gene regulatory networks (GRNs) map directed, molecular interactions between transcription factors (TFs) and target genes. We and others have shown that integrating different omics data provides a more accurate, multi-modal view of gene regulation^14,15^. In this respect, adding TF-binding information to transcriptome data in network inference allows going beyond statistical association^16^. There are several methods to infer GRNs, which are based on different underlying assumptions such as correlation, information theory, Bayesian networks, probabilistic models, regression, differential equations, deep learning, and multi-omics data integration methods^16^. Typically, inferring GRNs from bulk data requires a sufficient number of samples. Despite high-dimensionality, sparsity and overdispersion, single-cell omics of a single sample can be leveraged for single-cell network inference. We and others have found that distinct network inference methods reveal complementary aspects of the underlying GRNs and that a consensus solution provides a more robust and accurate representation of the underlying regulatory programs^14,17^.

Network inference has already been conducted on several neuroinflammatory disorders^18–27^. Hitherto, the popular method WGCNA^28^ has often been used, which builds co-expression networks of undirected interactions, and often the focus is on differentially expressed genes (DEGs), losing a substantial part of the data. Two studies have executed GRN inference of several neurological disorders from bulk transcriptomics data^19,20^. In the first study, microarray data from AD, Parkinson’s disease, Huntington’s disease, schizophrenia, amyotrophic lateral sclerosis, multiple sclerosis, and control samples were analyzed for DEGs, which were utilized for STRING protein-protein interaction network construction and for TF-binding targets enrichment from the ChEA database, to predict core GRNs^19^. In this study the number of samples was limited: around ten patients and ten controls for every disease. In the second study, researchers constructed GRNs for schizophrenia, bipolar disorder, MDD, AD, and autism integrating multiple brain-specific (epi)genomics and transcriptomics data^20^. Firstly, a TF-gene binding network was created by TF binding site enrichment using ENCODE DNase I footprints. This network was further filtered for TF-gene co-expression using gene expression profiles from healthy individuals of the Allen Human Brain Atlas. Finally, key regulatory TFs of the different diseases were identified by calculating the enrichment of their network target genes in disease-specific DEGs^20^. The major limitation of this study is the fact that a healthy brain GRN was created and the disease-specificity only resulted from DEGs. Hence, this study found no results for MDD, as no DEGs were found in the MDD dataset. Considering network inference on single-cell studies of neuroinflammatory disorders, these tend to focus on one disorder only and DEGs between different cell types and conditions^29–32^. However, some studies have already inferred GRNs at the single-cell level on AD samples^22,33–35^. In addition, researchers have investigated regulatory programs of AD, Parkinson’s disease and multiple sclerosis by applying SCENIC on an integrated single-nucleus RNA-seq (snRNA-seq) dataset from several distinct brain regions^21^. By integrating all datasets and running SCENIC on all samples together, they lost part of the disease- and brain region-specific signals. In this study, we overcame the limitations of previous studies and compared GRNs of AD and MDD, which were inferred unbiasedly from sufficient samples from bulk and single-cell data.

Our overarching hypothesis postulates that AD and MDD exhibit common and distinct characteristics of neuroinflammation, which will be reflected in their respective GRNs. Hence, in this study, we aimed to comprehensively unravel shared and distinct regulatory programs in these neuroinflammatory conditions, by leveraging diverse computational approaches to infer GRNs, both at bulk and single-cell level. Using rank aggregation, a consensus network per disorder was constructed from three different methods’ GRNs (CLR, GENIE3 and Lemon-Tree) inferred from bulk RNA-seq data. These AD and MDD networks were compared to each other at the functional and regulatory levels. Moreover, single-cell GRNs were inferred using SCENIC on snRNA-seq data, pointing to specific brain cell types implicated in the disorders. Consequently, neuroinflammation and activation of microglia were found in both AD and MDD, while our investigations also revealed unique disrupted pathways specific to each disorder. Importantly, common core regulatory circuits were uncovered, including TFs that play a pivotal role in microglial activation (IKZF1, IRF8, NFATC2, RUNX1, SPI1, and TAL1).

## 2. Methods

### 2.1. Retrieval and preprocessing of bulk data

The publicly available RNA-seq datasets GSE174367/syn22130832^33^, GSE101521^36^, and GSE80655^37^ were retrieved. Different datasets of the same disease were merged to make a compendium per disease. We selected high-quality and biologically relevant datasets for our study as follows: human post-mortem brain samples from the prefrontal cortex, with each dataset containing at least 20 diseased and 20 control samples, not enriched for certain cell types (e.g. for neurons or glial cells), and with a similar number of samples for each disorder (Supplemental Table 1). Raw counts were preprocessed using the R package *edgeR*^38^ (R, v4.1.3) by filtering genes on a counts per million (cpm) value greater than one in at least five samples, normalizing by library size^39^, logarithmic transformation with a prior count of one, batch effect and outlier detection by multidimensional scaling and hierarchical clustering, and batch correction with the *removeBatchEffects* function. In the AD dataset, six outliers out of 95 samples were omitted: four AD samples and two controls. In the MDD dataset, one MDD outlier sample was deleted out of 106 samples. Next, protein-coding genes and highly variable genes were selected, after which TFs with expression data that were left out were added again^40^. This ultimately resulted in 9724 and 9217 genes in the AD and MDD datasets, respectively. Lastly, scaling and centering by gene were performed.

### 2.2. Bulk network inference

For all network inference methods, the input regulators were TFs as defined by Lovering et al.^40^ (1455 TFs in total). We ran GENIE3^41^ (R package *GENIE3*) with random forest and default parameters, CLR^42^ (R package *minet*) with the Miller-Madow estimator and the equal frequency discretization, with edges filtered to contain at least one TF, and Lemon-Tree^43^ (in Java as a command-line program (v3.1.1), https://github.com/erbon7/lemon-tree) with 100 permutations for the clustering and a minimum of ten genes per module.

### 2.3. Bulk consensus networks

From the networks inferred by the different methods above, a consensus network was created by average rank aggregation using the R package *TopKLists*^44^ with the *Borda* function. The overlap in predicted TFs was calculated using the Jaccard similarity index, on the base of which k-medoids clustering grouped the genes into modules of coregulated genes using the R package *cluster*^45^ with the *pam* function. The optimal number of clusters was calculated using the average silhouette width and Calinski-Harabasz index, with the *NbClust*^46^ R package and *NbClust* function. However, in the *NbClust* function, it was not possible to use k-medoids and the Jaccard similarity index, thus we approximated this by using the median and k-means methods and the default dissimilarity metric on the scaled counts. As in all cases the optimal number of clusters was close to the number of clusters in the Lemon-Tree solution, i.e. 155 and 156 modules for AD and MDD respectively, we selected those numbers. There was for both AD and MDD networks one cluster with more than 1000 genes, which both were briefly inspected and then omitted. They contained genes of which the significant (adjusted p-value ≤ 0.05) functional enrichment terms indicated regulation of transcription and DNA binding, since the module contained mostly TFs and cofactors. Finally, TFs were assigned as regulators to each module as follows: TFs were ordered by the number of genes they regulated in the module, TFs had to regulate at least half of the module genes, and a maximum of ten TFs were allocated to each module. Hubs were defined as the TFs with the highest out-degree.

### 2.4. Retrieval and preprocessing of single-cell datasets and single-cell network inference

snRNA-seq datasets from post-mortem human brain samples were from the prefrontal cortex as well. Similarly, we chose datasets with both neurons and glial cells, control samples, and a similar number of samples (cells) for each disorder. The publicly available files (AD: GSE174367/syn22130832^33^, MDD: GSE144136^29^) were preprocessed with the R package *Seurat*^47^. Cells with too low (<200) or too high (>5000 MDD, >7500 AD) UMI (unique molecular identifier) counts were omitted, resulting in 59968 cells in the AD dataset and 73371 cells in the MDD dataset. In addition, only protein-coding genes were selected. Moreover, the dataset was logarithmically transformed (*NormalizeData* function) and highly variable genes (8000 genes) were selected. Next, TFs that had been omitted by the filtering, were added again. The cell annotations were taken from the original authors. The ‘mix’ cell type from the MDD dataset was excluded from most analyses because of limited biological interpretation. Subsequently, GRNs were inferred using SCENIC^48,49^ (*pySCENIC* (v.0.10.4) with GRNBoost2^50^) from the single-cell data, which was run through a Nextflow pipeline available at https://github.com/vib-singlecell-nf/vsn-pipelines with default parameters. Firstly, GRNBoost2 is used to infer GRNs, which are then further refined to direct target genes by using TF motif data, from RcisTarget, reducing the number of false positives. AUCell is then used to score the regulon activity in each cell type. Differences in cell type proportions were calculated with the R package *speckle* and the function *propeller* (logit transformation).

### 2.5. Ground truth datasets

We retrieved ground truth datasets from different sources. The gene regulatory interactions from OmniPath^51^ were retrieved, consisting of three main collections: CollecTRI^52^, DoRothEA^53^ (confidence levels A-D), and additional literature-curated interactions. After filtering for TFs defined by Lovering et al.^40^, this resulted in 111376 interactions and 862 TFs. Marbach and colleagues have constructed 394 cell- or tissue-specific regulatory networks by integrating TF motifs with promoter and enhancer activity data^54^. The frontal lobe region network was chosen (Synapse syn4956655), as the expression datasets in our study were from the prefrontal cortex. This network contained 1086698 edges and 567 TFs. Furthermore, we downloaded (on May 5^th^, 2023) all perturbation datasets of all available TFs of KnockTF^55^, consisting of 308 TFs and 22132 unique DEGs. A KnockTF network was created by forming regulatory edges between perturbed TFs and DEGs. Lastly, UniBind^56^ was used as a ground-truth ChIP-seq database. The robust collection of TFBSs was downloaded for Homo sapiens (on July 4^th^, 2023). For each TFBS in the collection, the closest gene was retrieved using the transcription start sites from all protein-coding genes, using BedTools closest. Next, the distance between each gene and TFBS was filtered to be under 10kb, based on the distance used by SCENIC. This resulted in 1463247 edges, with 265 TFs and 19069 target genes. Next, for all ground truth collections, precision and recall were calculated after filtering our networks and ground truth for the TFs and genes that were present in both^14^.

### 2.6. Functional characterization

The *enrichR*^57^ package in R was used for functional enrichment analysis, with the databases Gene Ontology^58,59^ (GO) Biological Process (2021), GO Molecular Function (2021), KEGG^60^ (2021), Reactome^61^ (2016) and WikiPathways^62^ Human (2021). Immune-related modules were defined as functional enrichment terms containing ‘immune’/’inflammation’ and terms related to cytokines, the complement system, innate and adaptive immune cells, phagocytosis, pattern recognition receptors such as Toll-like receptors, and NFκB signaling. Additionally, we used *enrichR* to look at the enrichment (adjusted p-value ≤ 0.05) of target genes of certain TFs, using the libraries of ChEA (2016 and 2022) and ENCODE TF ChIP-seq (2015). Moreover, motif enrichment analysis was executed for each module, using the R package RcisTarget. The files were downloaded from https://resources.aertslab.org/cistarget/: we selected version 10 and considered 10 kb up- and downstream of the transcription start site. Further, to see if the target genes of TFs were enriched for disease-associated genes, we used DisGeNET^63^ through *enrichR*. For the TFs in the AD networks, we looked at whether their target genes were enriched in the gene sets of AD terms (‘Alzheimer’s disease’, ‘Alzheimer Disease, Late Onset’, ‘Familial Alzheimer Disease (FAD)’, ‘Alzheimer Disease, Early Onset’). Similarly, for the MDD networks, we looked at whether the target genes were enriched for genes previously associated with MDD (‘Major Depressive Disorder’, ‘Unipolar Depression’, ‘Mental Depression’, ‘Depressive disorder’, ‘Severe depression’, ‘Major depression, single episode’).

### 2.7. Visualization

Venn diagrams were made with the R package *VennDiagram*^64^. The regulatory signs of the regulatory TFs in Figure 2 were inferred with SIREN^65^ in R, using a cut-off of ±0.15 (activation, repression, or not definable). The modules from the consensus networks were visualized with Module Viewer^14^ (https://github.com/thpar/ModuleViewer), a Java program in which the expression of different modules can be visualized, together with its TFs and annotation data. HumanNet (v3)^66^ interactions were downloaded for integrative visualization with the modules, more specifically physical protein-protein interactions, co-expression links, co-functional links by pathway database, co-functional links by protein domain profile associations, co-functional links by genetic interaction, and co-functional links by gene neighborhood. Physical protein-protein interactions and co-expression links were kept separately, while the other interactions were merged into ‘functional’ interactions. Additionally, GO Biological Process terms were pruned with the R package rrvgo^67^ (threshold of 0.7 if there were less than 100 terms, threshold of 0.5 if there were more than 100 terms: the more terms, the more pruning), and the top five of the pruned terms were kept (according to adjusted p-value). Together with these, the top five terms of Reactome and WikiPathways were retrieved for visualization. Network visualizations were created with Cytoscape^68^.

### 2.8. hdWGCNA

The R package *hdWGCNA*^26^ was utilized for module projection of the consensus population-level modules onto the single-cell data with the function *ProjectModules* to investigate in which cell types the modules were most active. We selected the modules that were prioritized with either of the two prioritization strategies and visualized the results with *DotPlot* (*Seurat*) figures. The AD modules were projected into the AD single-cell data, and the MDD modules were projected into the MDD single-cell data. Additionally, hdWGCNA was utilized similarly to look for different microglial states in patients vs control samples. Microglial states were taken from Olah and colleagues^69^ and the gene signatures for each state were downloaded from their Supplemental Table 5. Here, the microglial states were seen as modules and the differences in diagnosis were investigated instead of cell types. Only microglia were used for this analysis.

### 2.9. Connection specificity index (CSI)

The CSI was calculated between pairs of regulons in the AD and MDD networks separately, using the AUC scores from SCENIC. This was done with the R package *scFunctions*^70^ (https://github.com/FloWuenne/scFunctions).

### 2.10. GWAS SNPs

Genome-wide association studies (GWAS) summary statistics were downloaded from the NHGRI-EBI GWAS Catalog^71^ for ‘Alzheimer disease’ (MONDO_0004975) and ‘unipolar depression’ (EFO_0003761) on December 6^th^, 2022. The summary statistics were inspected for SNPs associated with genes contained in the prioritized modules. More specifically, the SNPs had to be located intragenically or a maximum of 3 kb upstream of the gene’s transcription start site.

### 2.11. Differential edge analysis

Edges were defined as differential if the edge was present in one of the diseases, and not in the other. Firstly, target genes and TFs were filtered based on their presence in both AD and MDD single-cell networks. Next, we selected genes with more than five edges and with more than 20% differential edges. Differentially targeted genes were then divided into ‘AD-biased’ (more than 90% of differential edges present in AD), ‘MDD-biased’ (more than 90% of differential edges present in MDD), and just differential genes.

### 2.12. Validation of IRF8 regulons

scRNA-seq counts of the Irf8 conditional KO and wildtype mice were downloaded from https://www.brainimmuneatlas.org/download.php^72^. The matrix was processed with *Seurat* and *scran*, based on the script on their GitHub (https://github.com/saeyslab/brainimmuneatlas/blob/master/script_scRNAseq.R). The differentially expressed genes between Irf8 KO and wildtype mouse microglia were calculated using the *FindMarkers* function from *Seurat* with the default Wilcoxon rank sum test. Using a cut-off of the adjusted p-value of < 0.01 and an absolute log fold change of > 0.5, these DEGs (2442 genes) were used as a gene set to calculate the enrichment of the IRF8 regulon genes of our analysis. The *phyper* function of the *stats* R package was used to calculate the overrepresentation. As the genes are from a different species, homologs from Ensembl^73^ and the function *convert_mouse_to_human_symbols* from the R package *nichenetr*^74^ were both used to convert the mouse genes to human genes. As such, 2171 human genes were used as a final gene set.

### 2.13. Validation dataset single-cell MDD

The raw counts from Maitra and colleagues^75^ were downloaded from GEO GSE213982. They were preprocessed with *Seurat*^47^. The counts contained both new female samples and the male samples that we utilized, hence, the female samples were selected before filtering. Cells with too low (<500) or too high (>10000) UMI counts were omitted, as well as cells with too many gene counts (>50000), resulting in 73780 cells. Protein-coding genes were selected, and the cell type annotations of the authors were used. *FindMarkers* was used to find the DEGs in microglia between cases and controls.

### 2.14. Statistical analysis

Multiple hypothesis testing was corrected for by the Benjamini-Hochberg method with FDR ≤ 0.05, unless mentioned otherwise.

## 3. Results

### 3.1. Population-level network inference on AD and MDD cohorts

To compare regulatory programs in AD and MDD, we first ran different GRN inference algorithms on bulk transcriptomics data (Figure 1A). We utilized publicly available RNA-seq data from human post-mortem brain (prefrontal cortex) samples: 43 AD and 46 healthy control samples (GSE174367/syn22130832^33^) and 52 MDD and 53 healthy control samples (GSE101521^36^ and GSE80655^37^) (Methods, Supplemental Table 1). The prefrontal cortex was the favored brain region, as it has been implicated in several neuropsychiatric, neurodevelopmental and neurodegenerative disorders^20^. Three methods that rely on different algorithmic principles were used to infer GRNs (Methods): the random forest regression method GENIE3^41^, the mutual-information-based relevance network method CLR^76^, and the integrative multi-omics module network inference method Lemon-Tree^43^, which combines an ensemble clustering of genes into modules with an ensemble of decision trees to infer the regulators of each module. Two networks were made with every method, one for each disorder, after which the top 100000 edges were selected from each network, and compared. For both disorders, about 10000 edges were shared by all three GRN inference methods (Figure 1B). To confirm that the different methods gave complementary and biologically relevant results, we looked at the overlap between the inferred interactions and multi-evidence-based ground truth regulatory interactions retrieved from four sources (Methods). The first ground truth GRNs consisted of interactions from OmniPath^51,52^, which includes known regulatory interactions combined from different resources. The second ground truth contained the frontal lobe GRNs from Marbach and colleagues^54^, which were inferred by integrating TF binding sites with promoter and enhancer activity data from the FANTOM5 project^54^. The third ground truth network consisted of TF perturbations in KnockTF^55^, and the last ground truth contained ChIP-seq TF binding interactions from UniBind^56^. After filtering and combining these ground truths, there were 1878921 and 1830357 ground truth interactions for AD and MDD, respectively. The overlap with the ground truth networks was low for all GRN methods and both disorders (around 2% for ADD and 2.3% for MDD, Figure 1B), but according to the state of the art^17^. There were more edges in common between every method separately and the ground truth (+/-13000 edges every time) than between the intersection of all three methods and the ground truth (1643 edges for AD, 3639 edges for MDD, Figure S1), hinting at complementarity of the different GRN inference methods.

**Figure 1.**
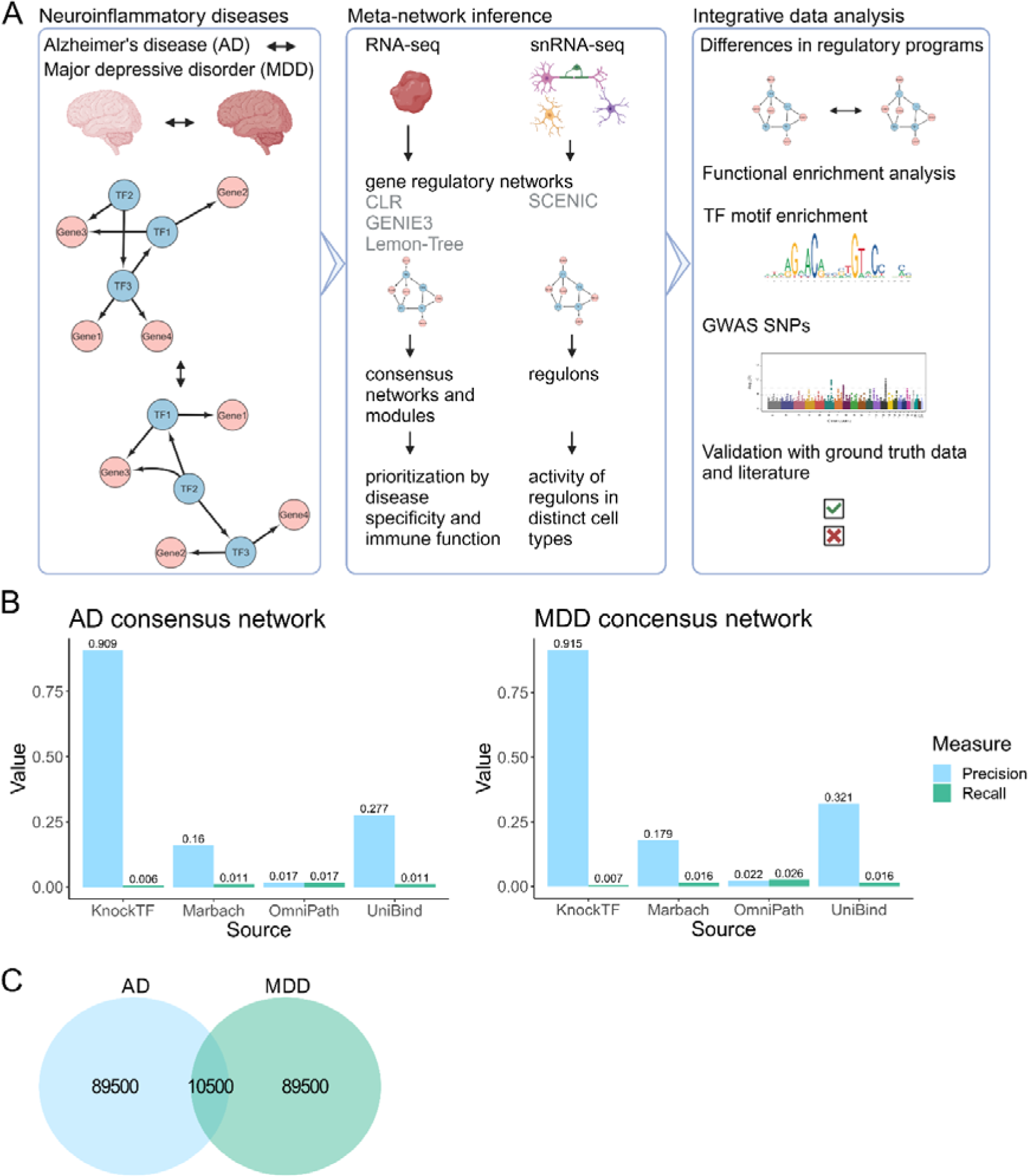
Population-level network inference on Alzheimer’s disease (AD) and major depressive disorder (MDD) cohorts. A) The neuroinflammatory disorders AD and MDD were compared through meta-network inference both at population and single-cell level, using different transcriptomics data sets and network inference methods. At the population level, modules of coregulated genes were prioritized for disease differences and immune system function. At the single-cell level, the activity of regulatory programs was further investigated in the specific cell types of the brain. At both levels, the networks were validated in silico and integrated with other biologically relevant data. Figure created with BioRender. B) Precision and recall (AD, left; MDD, right) of the consensus networks depicted with ground truth interactions from OmniPath, Marbach et al., KnockTF (TF perturbation) and UniBind (TF binding). TF perturbation data resulted in the highest precision, but lowest recall. C) The Venn diagram shows that there were 10500 edges (10.5%) in common between the 100000 edges in the consensus networks of AD and MDD.

To obtain more robustly predicted regulatory programs, we constructed consensus regulatory networks from the three different methods for both diseases, through average rank aggregation (Methods). As before, the top 100000 edges were retained, with interactions between 1041 TFs and 9675 target genes (9685 nodes in total) for AD and between 1101 TFs and 9006 target genes (9021 nodes in total) for MDD. Next, precision and recall were calculated for the consensus networks, using the four distinct ground truth datasets (Methods, Figure 1B). Due to incomplete experimental GRN mapping in the field, precision was higher than recall, except for the OmniPath interactions. KnockTF perturbation data resulted in the highest precisions. Between the AD and MDD consensus networks, there were 7042 nodes (72.42% for AD, 76.40% for MDD) and 10500 edges (10.5%) in common (Figure 1C). Of the top 100 hub TFs of each consensus network, there were 34 in common, including several zinc finger and SOX family TFs (Supplemental Table 2).

### 3.2. Intertwined regulation of neuroinflammation in AD and MDD

To investigate the biological implications of the inferred networks, we disentangled the hairball structure of the GRNs by decomposing them into modules, here clusters of coregulated genes. The overlap in predicted TFs of all genes was calculated using the Jaccard similarity index, which was then used to group the genes into modules of coregulated genes through k-medoids clustering (Methods). The number of clusters to choose with k-medoids can be hard, especially for a large dataset. We chose the numbers of clusters that were retrieved with Lemon-Tree, i.e. 155 for AD and 156 for MDD, as this was similar to the optimal number of clusters according to the average silhouette width and Calinski-Harabasz index (Methods). Next, a maximum of ten TFs were assigned to the modules, and they had to be predicted to regulate at least half of the genes in the module.

Coregulated genes tend to function in similar biological processes. Therefore, functional enrichment analysis was performed on all modules using Gene Ontology (GO) Biological Process, GO Molecular Function, KEGG, Reactome, and WikiPathways annotations. Additionally, TF binding motif enrichment analysis using RcisTarget^48^ was executed and we investigated whether the modules’ genes were enriched for target genes of TFs, using enrichR, which we further refer to as TF target gene enrichment (Methods). For both disorders, there were some modules enriched for the ‘Alzheimer’s disease’ term from KEGG (11 for AD, five for MDD), indicating an etiological overlap between AD and MDD (Supplemental Table 3). We prioritized modules in two ways (Figure 2, Supplemental Table 3): firstly by functional annotation, namely module genes enriched for immune-system-related pathways (Methods), as both disorders are characterized by neuroinflammation, and secondly, according to their differential expression between control and disease phenotypes (two-sided Wilcoxon Rank sum test). Hence, we selected for AD and MDD respectively 10 and 12 modules functioning in immune pathways and/or 15 and 11 modules top-ranked by differential expression between disease and control (Supplemental Table 3). An overview of all prioritized modules can be found in the Supplemental material. We found several disease-specific, non-immune modules. As an example, module 60 of the AD network contained the tau gene (*MAPT*) and was involved in mitochondrial pathways, such as mitochondrial protein import and the citric acid cycle, and metabolism of proteins, with a lower expression in AD patients and was regulated by *HINFP, STOX2, ZNF26, ZNF480, ZNF552, ZNF587B,* and *ZNF814*. *HINFP* was found as a potential biomarker for AD^77^. There were seven module genes enriched in the ‘Alzheimer’s disease’ term of the KEGG database (*PSMB7, PSMC6, MAPT, CYC1, COX6A1, GAPDH, RTN4*). MDD-module 93 was functionally enriched for terms related to oligodendrocytes, regulation of nervous system development, GABA receptor signaling, and mBDNF and proBDNF regulation of GABA neurotransmission. As it has been observed that the balance between proBDNF and BDNF is disturbed in MDD and that GABA levels are decreased in MDD patients^7^, the expression of this module was indeed decreased in patients compared to controls. The predicted regulators were *TRPS, 1NR2E1, PPARA, ZBTB10, RFX4, GLI3, STOX1, NPAS3, PAX6,* and *SOX21*. PPARA, NPAS3, and PAX6 have been associated with MDD before^78–80^. Below, we further elaborate upon the top-ranked modules by differential expression that also functioned in immune pathways (AD: modules 22, 39, 51 and 153; MDD: modules 24, 36, 110 and 115), which will be called ‘disease-specific neuroinflammation modules’ throughout. These modules had regulatory TFs and target genes overlapping between the two disorders (Figure 2). Modules were functionally coherent, as evidenced by overlap with HumanNet^66^ co-expression, protein-protein, and other functional interactions (Figures 3, 5 and Supplemental material).

**Figure 2.**
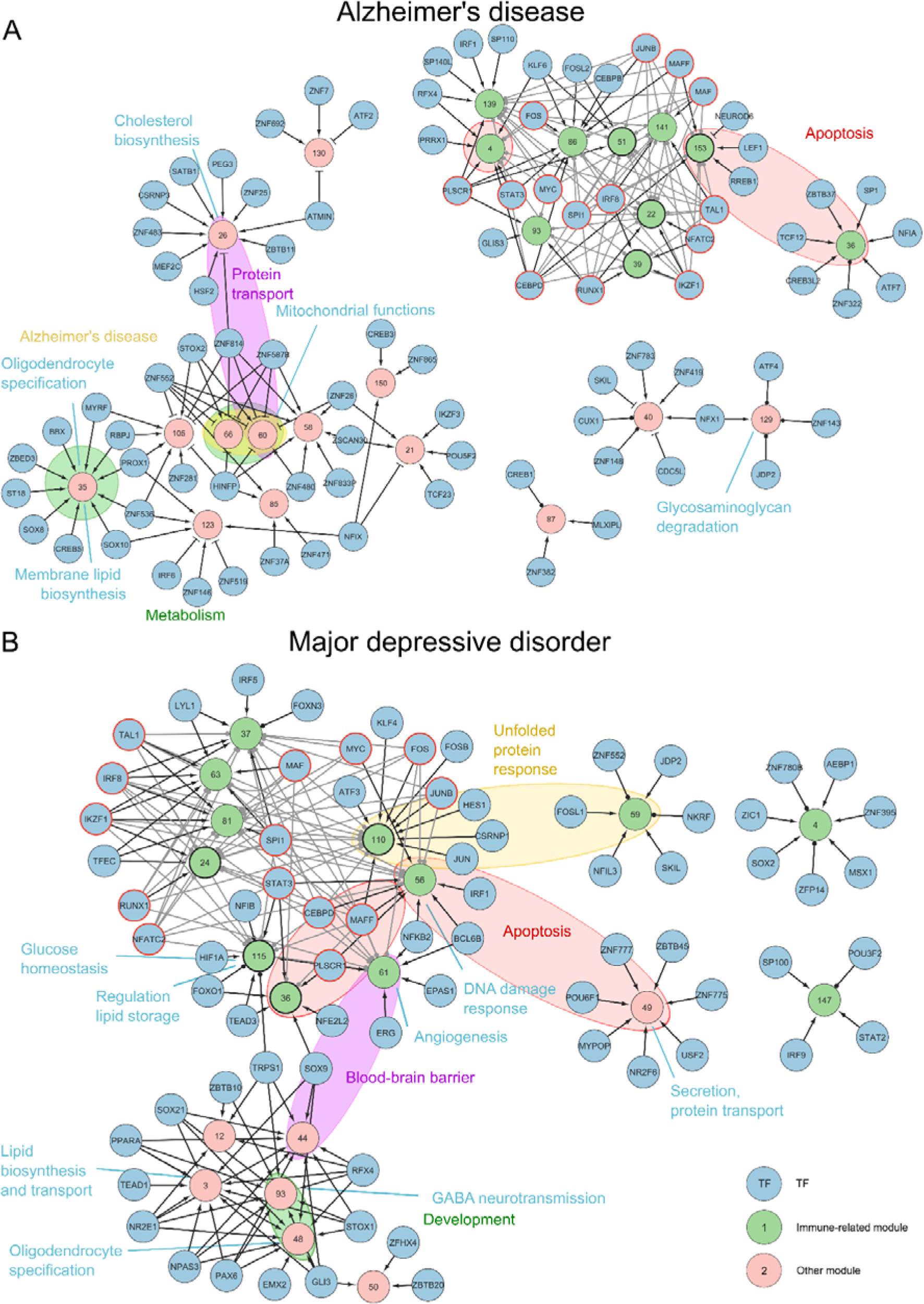
Intertwined regulation of neuroinflammation in Alzheimer’s disease (AD) and major depressive disorder (MDD). Modules of the consensus networks were prioritized through differential expression between disease and control, and functional enrichment for immune-related annotations. A) Prioritized AD modules and their regulatory transcription factors (TFs). B) Prioritized MDD modules and their regulatory TFs. Many TFs co-regulated several modules and immune-related modules tended to cluster together because they were regulated by similar TFs. TFs are depicted in blue, immune-related modules in green, and other modules in pink. Nodes with red borders represent TFs co-regulating multiple immune modules that were shared between AD and MDD. Grey edges indicate additional interactions detected by motif enrichment analysis. Colored ovals represent functional annotation terms or pathways (non-immune-related) that are shared between several modules, while module-specific functional annotations are indicated in blue text.

**Figure 3.**
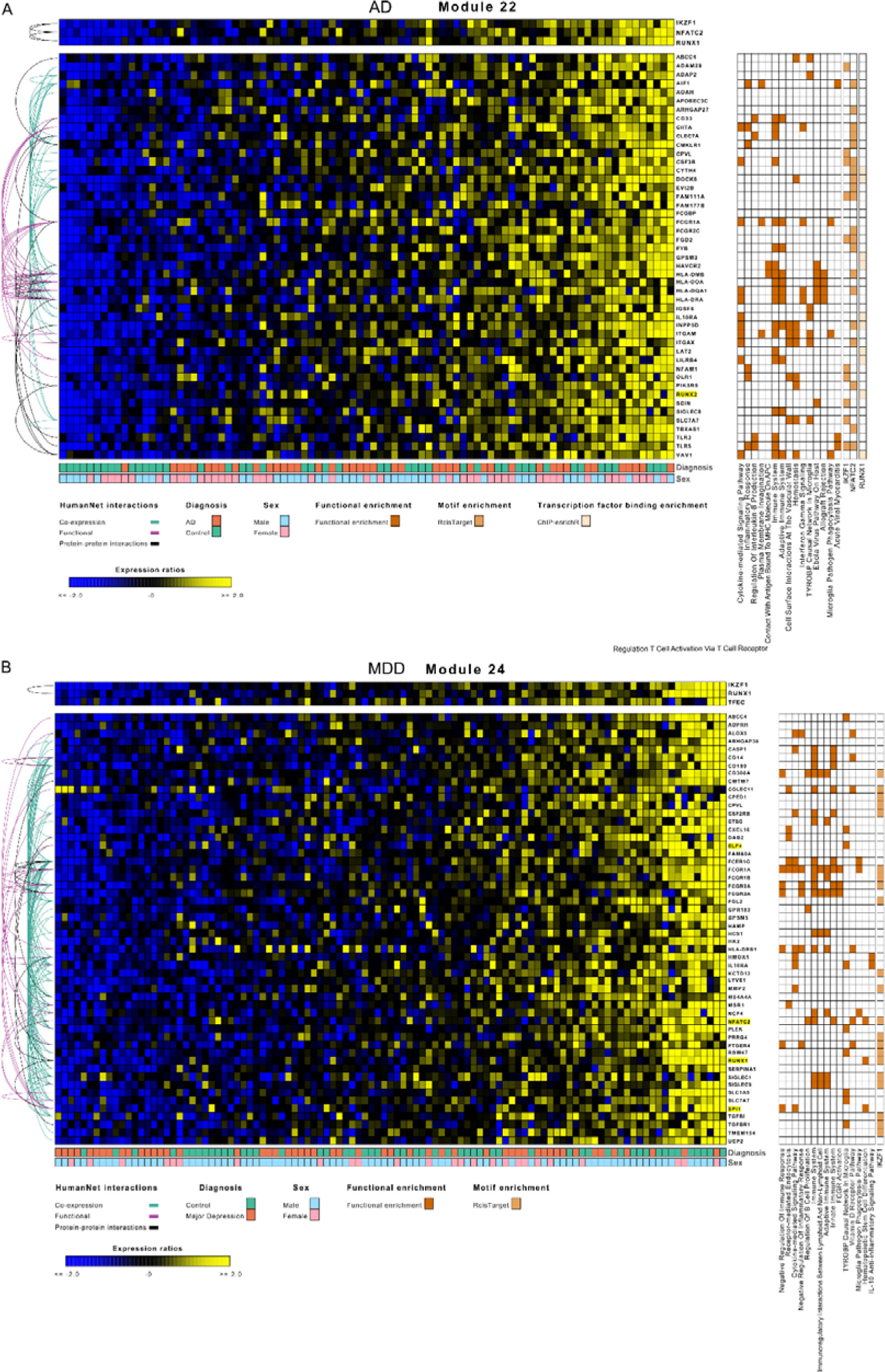
Modules 22 of the Alzheimer’s disease (AD) network and 24 of the major depressive disorder (MDD) network highlighted functioning in neuroinflammation and shared IKZF1 and RUNX1 as regulatory transcription factors (TFs) and *ABCC4*, *CPVL*, *FCGR1A*, *GPSM3*, *IL10RA*, and *SLC7A7* as target genes. A) AD module 22 contained 47 genes being upregulated in AD as compared to controls and functioned in cytokine signaling, antigen presentation and phagocytosis. Its regulators IKZF1, NFATC2 and RUNX1 were not only predicted through network inference, but also by TF binding motif or TF target gene enrichment, suggesting a direct regulatory and binding effect of these TFs on the module genes. B) MDD module 24 contained 54 genes being downregulated in MDD and functioned in neutrophil activation, cytokine release and phagocytosis. The predicted regulators were IKZF1, RUNX1 and TFEC, with IKZF1 also found by motif enrichment. The ModuleViewer figures include the sample diagnosis, sex, functional enrichment terms, TF binding motif enrichment (RcisTarget), TF target gene enrichment (enrichR), and interactions from HumanNet (see legends). Highlighted module genes are TFs.

### 3.3. Similar TFs target immune-related modules in AD and MDD

Several inferred regulatory TFs were shared between neuroinflammatory modules of AD and MDD (Figure 2). In addition to TFs inferred through the consensus network approach, TF binding motif enrichment analysis on the module genes identified several other TFs (grey edges in Figure 2; only indicated for immune modules with shared TFs). *IKZF1*, *IRF8*, *MAF*, *NFATC2*, *RUNX1*, *SPI1*, and *TAL1* together regulated several modules in both AD and MDD. All these TFs are regulators of immune functions: *NFATC2* is a regulator of cytokine release, *IKZF1* and *TAL1* play a role in hematopoietic cell differentiation, *MAF*, *NFATC2* and *RUNX1* are all involved in T-cell functioning and *SPI1* controls hematopoietic cell fate and can recruit IRFs^81–86^. In a recent study in knockout mice, Ikzf1 was found to specifically alter microglial state and function, and microglial Ikzf1 levels were increased in neuroinflammation models, pointing to Ikzf1 as a novel regulator of microglial homeostasis^87^. Additionally, mutations in the TF binding sites of IKFZ1 and TAL1, and mutations in the genes of *RUNX1* and *SPI1* have been associated with AD risk, while *IKZF1*, *NFATC2*, *SPI1*, *IRF8* have reported roles in microglial activation in AD^88–92^. Only *TAL1* was found to be implicated in MDD^93,94^ and was found to function as a master regulator in microglial homeostatis^95^. Interestingly, RUNX1 was found to bind the promoter of *SPI1* in hematopoietic cells^96^ and the other way around in microglia^97^, both in mice. Moreover, the protein product of *SPI1* (PU.1) was found to bind to the promoter of *IRF8* and *SPI1* itself in microglia in mice^97^. This indicates that these regulatory TFs play significant roles in microglial functioning.

Another group of regulatory TFs shared between the two diseases consisted of *CEBPD*, *FOS*, *JUNB*, *MAFF*, *MYC*, *PLSCR1*, and *STAT3* (Figure 2). *CEBPD* is an important regulator of immune and inflammatory responses^98^ and is highly expressed in astrocytes during neuroinflammation^99^. TFs from the FOS, JUN and MAF families can dimerize to form the AP-1 complex, which supports higher cognitive functions and regulates inflammatory processes^100–102^. *MYC* plays a role in microglia activation^103^ and *PLSCR1* is involved in apoptosis and cytokine-regulated cell proliferation and differentiation^104^. Furthermore, STAT3 phosphorylation is critical for the production of the cytokines IL-1β and IL-6^105^. Further, AP-1, *CEBPD*, *MAFF*, *MYC,* and *STAT3* have been implicated in AD before^99,106–110^, while AP-1, *MYC*, *PLSCR1* and *STAT3* are known to play a role in MDD^111–115^.

Several modules with common regulators also shared biological pathways between AD and MDD. AD-module 22 and MDD-module 24 were predicted to be regulated by *IKZF1*, *NFATC2* and *RUNX1*, and *IKZF1*, *RUNX1* and *TFEC*, respectively, and had six genes in common: *ABCC4*, *CPVL*, *FCGR1A*, *GPSM3*, *IL10RA* (IL-10 receptor alfa), and *SLC7A7* (Figure 3). *ABCC4*, *FCGR1A, and CPVL* are linked to AD through DisGeNET^63^, while IL-10 has been associated with a worsening plaque load and reduced Aβ phagocytosis by microglia in mouse models of AD^116^. Functional enrichment analysis associated both modules to activated microglia functions such as WikiPathways’ ‘TYROPB causal network in microglia’ and ‘Microglia Pathogen Phagocytosis Pathway’. However, we also found different biological processes in these modules. The 47 genes in AD-module 22 were functionally enriched for terms related to cytokines (IFN-γ, IL-1β, IL-2, IL-3, IL-6, IL-8, IL-10, IL-12, and TNF), phagocytosis and T-cells (Figure 3A). Classically activated microglia are activated by IFN-γ or LPS and secrete the cytokines TNF, IL-1β, IL-2, IL-6, and IL-8, amongst others^117,118^. These microglia attract and activate T-cells, more specifically T_H_1 and T_H_17 T-cells. Phagocytosis is decreased in classically activated microglia^118^. Higher levels of T-cell infiltration are observed in AD patients, however, their exact role remains unclear^119^. Alzheimer’s patients tended to have a higher expression of the module genes compared to control individuals. The module gene *INPP5D* contains altered binding sites for IKZF1 in AD patients^119^ *INPP5D* functions in the regulation of immune cell homeostasis, chemotaxis, macrophage activation, and neutrophil migration^120^, and has been found to control microglia activation in a mouse model of AD^121^. The 54 genes in MDD-module 24 were functionally enriched for pathways related to the cytokine response as well (IFN-γ, IL-4, IL-10), but also neutrophil activation and B- and T-cells (Figure 3B). This agrees with what has been reported in the literature: in depression neutrophils and IFN-γ are increased, while IL-10 is decreased^10^. IFN-γ is produced by T_H_1 cells, while IL-10 is mostly secreted by T_reg_ cells. The α-subunit of the IL-10 receptor (*IL10RA*) was one of the genes shared by the two modules. Furthermore, it is seen that B, T_H_2 and T_reg_ cells are decreased, while T_H_1 and T_H_17 cells are increased in depression^10^. Here depressive patients tended to have a lower expression of the module genes compared to controls. As both modules were enriched for genes functioning in the immune system and microglia activation, these modules were likely active in microglia. However, while module 22 was upregulated in AD, module 24 was downregulated in MDD.

Module 39 of the AD network (103 genes) was involved in neutrophil activation, cytokine production, B-cell activation, amyloid-beta binding, and TYROBP causal network in microglia, with *TYROBP* being part of this module. Similar to module 22, the expression of this module seemed to be higher in AD patients. TYROBP, which is an adaptor protein that associates with activating receptors of immune cells, such as the pattern-recognition receptor TREM2, was found to be significantly upregulated in the brains of patients with AD^122,123^. Both TYROBP and TREM2 are required for phagocytosis in microglia. Microglia lacking TREM2 are not able to fulfill phagocytosis, chemotaxis, proliferation, and inflammatory response, resulting in the exacerbation of AD^5,124^. In addition, TYROBP can suppress cytokine production and secretion. Researchers have found that TYROBP could play a key role in the first stage of the disease-associated microglia phenotype, which is TREM2 independent^122^. The protein product of *SPI1* (PU.1) also binds *TYROBP, TREM2* and *INPP5D*^97,125,126^. Modules 39 and 141 in the AD network shared several regulatory TFs with AD-module 22, similar to MDD-modules 37, 63 and 81 with module 24 (Figure 2). These modules also had immune-related functional enrichment terms. Furthermore, genome-wide association studies (GWAS) summary statistics from the GWAS Catalog were downloaded to see whether SNPs associated with AD or MDD were present intragenic or upstream (max 3 kb) of genes in the prioritized population-level modules (Methods). Indeed, intragenic SNPs associated with AD were found in module 22, module 39, module 153, and module 141 genes (Supplemental Table 4). Additionally, SNPs upstream of *CD33* (module 22) and *MS4A6A* (module 39) were found to be associated with AD. For MDD, intragenic SNPs were found in module 36, module 37, module 63, and module 81 genes, while a SNP was found upstream of module 115 gene *ANXA1* (Figure 4, Supplemental Table 4).

**Figure 4.**
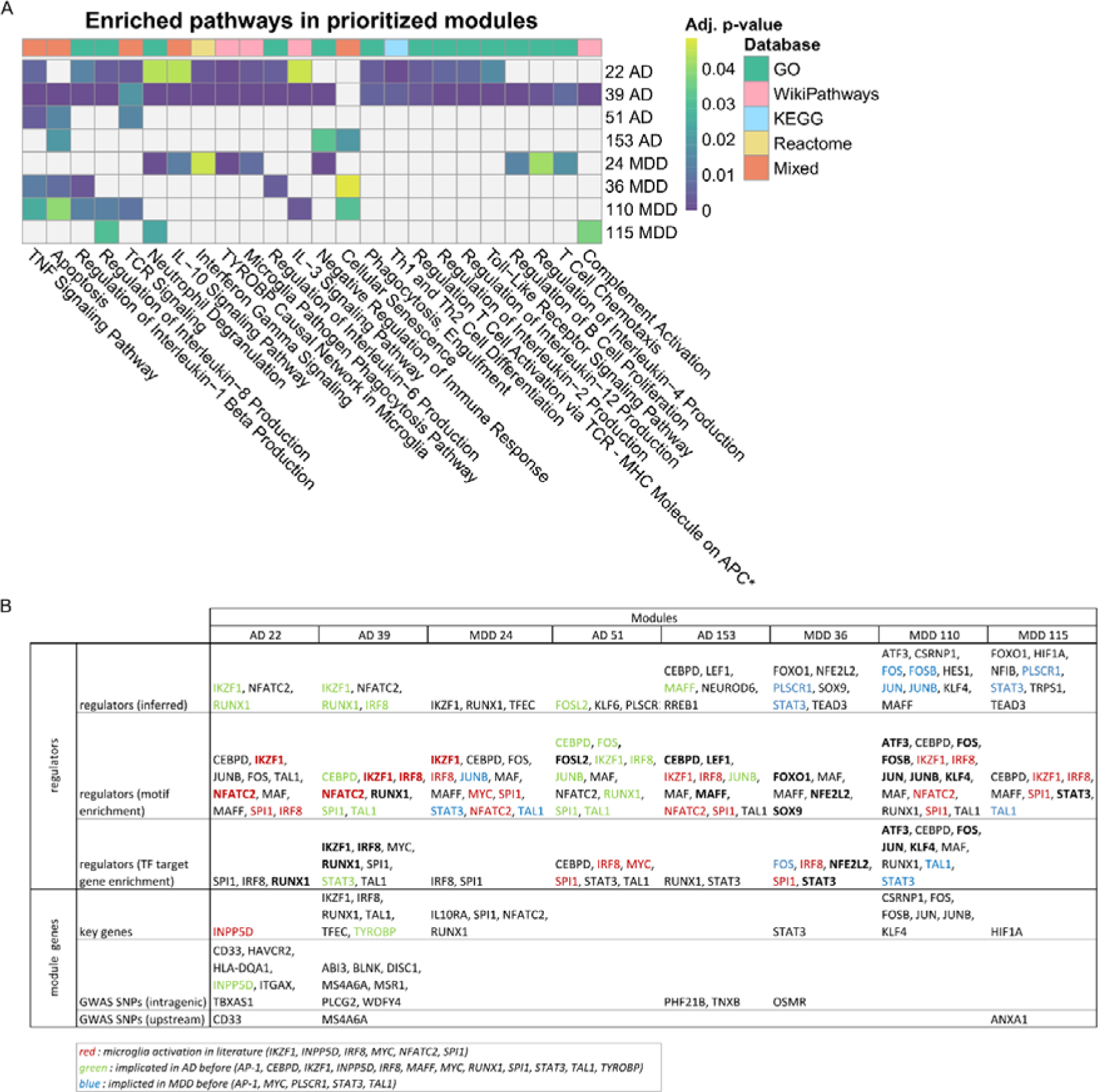
Overview of the pathways, regulators and module genes of the AD and MDD disease-specific neuroinflammation modules. A) Overview of shared pathways, quantified with the adjusted p-value and annotated with the pathway database. * ‘Regulation of T cell activation via T cell receptor contact with antigen bound to MHC molecule on antigen presenting cell’. B) Overview of the regulators and module genes of the modules. Regulators in bold were both found by the TF binding motif or TF target gene enrichment and through network inference. Colored genes have been found in the literature to be involved in microglial activation (red), AD (green) or MDD (blue). GWAS SNPs are specific SNPs for the disease of the module i.e. AD or MDD.

Additionally, two other interesting modules had an overlap in genes and TFs. Module 153 from AD and module 110 from MDD were both regulated by MAFF and shared the module genes *RGS1* and *TNFAIP3* (Figure 5). *RGS1* has previously been found to be upregulated in AD patients^127^. Interestingly, TNFAIP3 is involved in NFκB signaling^128^, has been proposed as a predictive biomarker of response to antidepressant treatment in MDD^129^, and a mutation in this gene has been associated with AD risk as well^130^. Module 153 tended to be upregulated in AD patients and its functional enrichment pointed to apoptosis, regulation of inflammatory response and cellular senescence (Figure 5A). One of the predicted regulatory TFs, *NEUROD6*, was downregulated in AD patients, which has been described before^131^. MDD-module 110 contained genes involved in the unfolded protein response, response to cellular stress, and cytokine signaling (Figure 5B). Surprisingly, this module tended to be downregulated in MDD patients compared to control individuals. Further, the TF *PLSCR1* regulated several modules (AD-module 51 and MDD-modules 36 and 115) that were functionally enriched in apoptosis. Next to apoptosis, the genes of AD module 51 were involved in TNF and T-cell receptor signaling (Figure 4). The module genes seemed higher expressed in AD compared to healthy control samples. Similarly in MDD, in addition to apoptosis, module 36 functioned in cytokine signaling (IL-1β, IL-6, IFN-β, TNF), while module 115 was involved in complement activation, neutrophil activation, and cytokine response (IL-1 and IL-8). Here, both modules seemed to have a lower expression of module genes in MDD compared to controls. Overall, for both AD and MDD, dysregulation in immune functions, such as cytokine release, microglial activation, and phagocytosis, were found, but with only a partial overlap in affected pathways. Notably, we identified several master regulators shared between both disorders. To further investigate the specific cell types involved in these immunological changes, we turned to single-cell transcriptomics data.

**Figure 5.**
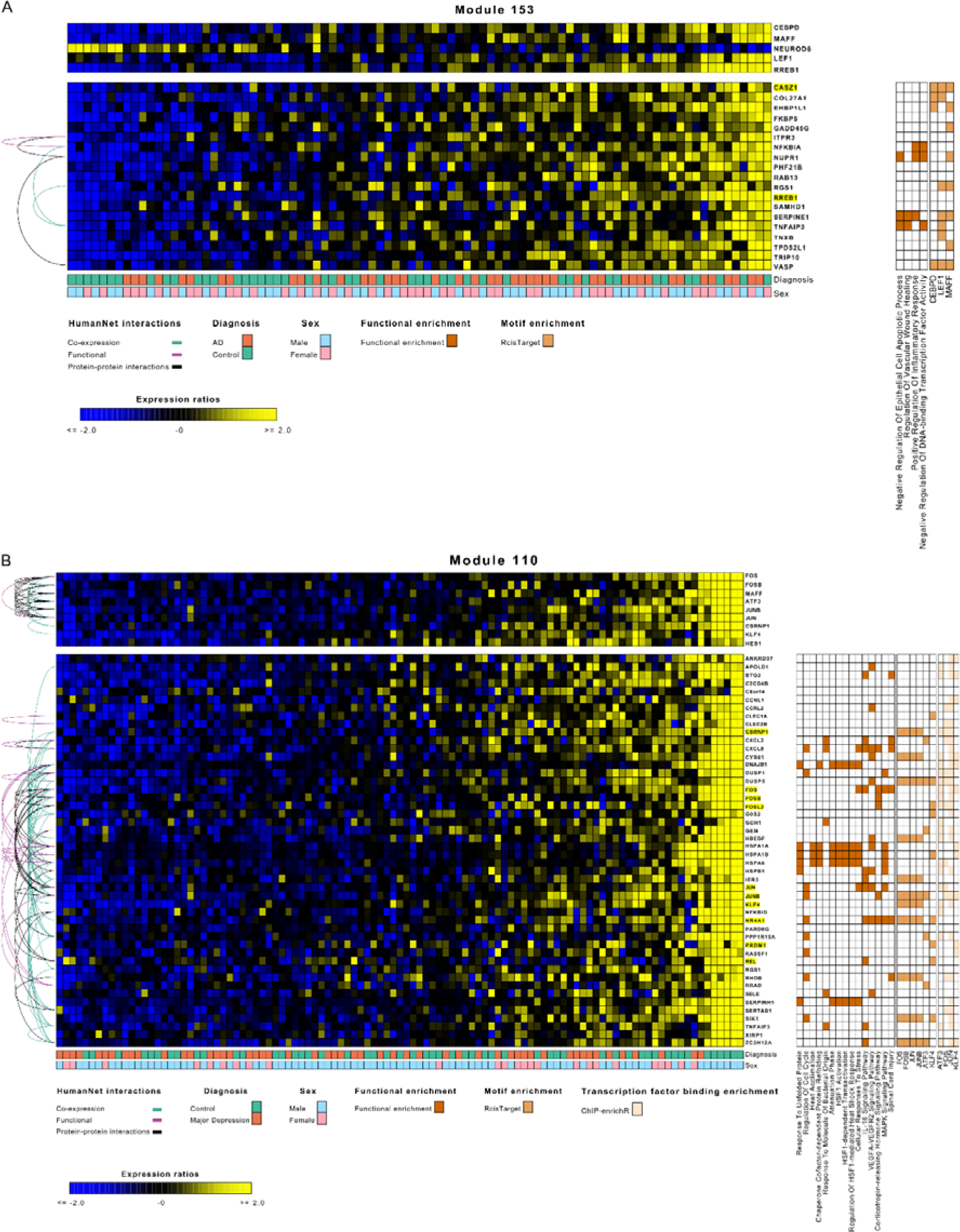
Modules 153 of the Alzheimer’s disease (AD) network and 110 of the major depressive disorder (MDD) network highlighted functioning in neuroinflammation and shared MAFF as regulatory transcription factor (TF) and RGS1 and TNFAIP3 as target genes. A) Module 153 of the AD network contained 19 genes being upregulated in AD as compared to controls, and functioned in apoptosis and regulation of inflammatory response. Its regulators CEBPD, LEF1, and MAFF were not only predicted through network inference, but also TF motif enrichment, suggesting a direct regulatory and binding effect of these TFs on the module genes. B) Module 110 from the MDD network contained 48 genes being downregulated in MDD as compared to controls, and functioned in the unfolded protein response and response to cellular stress, regulation of angiogenesis, and cytokine signaling. Its regulators ATF3, FOS, FOSB, JUN, JUNB, and KLF4 were not only predicted through network inference, but also through TF binding motif or TF target gene enrichment, suggesting a direct regulatory and binding effect of these TFs on the module genes. The ModuleViewer figures include the sample diagnosis, functional enrichment terms, TF motif enrichment (RcisTarget), TF target gene enrichment (enrichR), and interactions from HumanNet (see legends). Highlighted module genes are TFs.

### 3.4. Single-cell network inference on AD and MDD cohorts

We downloaded and analyzed snRNA-seq data from the prefrontal cortex (MDD: GSE144136^29^, AD: GSE174367/syn22130834^33^) (Figure 6A, Supplemental Table 1, Supplemental Figure S2). To bridge the bulk and single-cell data, we projected each population-level module for both AD and MDD onto the single-cell expression datasets using hdWGCNA^26^. Many projected modules were cell-type-specific (Figure 6B). Several projected disease-specific neuroinflammation modules of the population-level networks were indeed most specifically expressed in microglia, which confirms the hypothesis that these modules (AD-modules 22, 39, 51 and 153; MDD-module 24) represented regulatory programs active in microglia. Interestingly, in AD, most prioritized modules were active in microglia, while for MDD modules, most module genes were expressed in astrocytes. Some modules were not specific to a certain cell type (AD modules 21, 60 and 150; MDD modules 59, 115 and 147). As has been previously reported, we observed a reduction of astrocytes in MDD compared to controls (t-test, adjusted p-value 0.0055 and proportion ratio 2.642). The other cell types of MDD, or AD, were not significantly different in proportion (Supplemental Figure S3).

**Figure 6.**
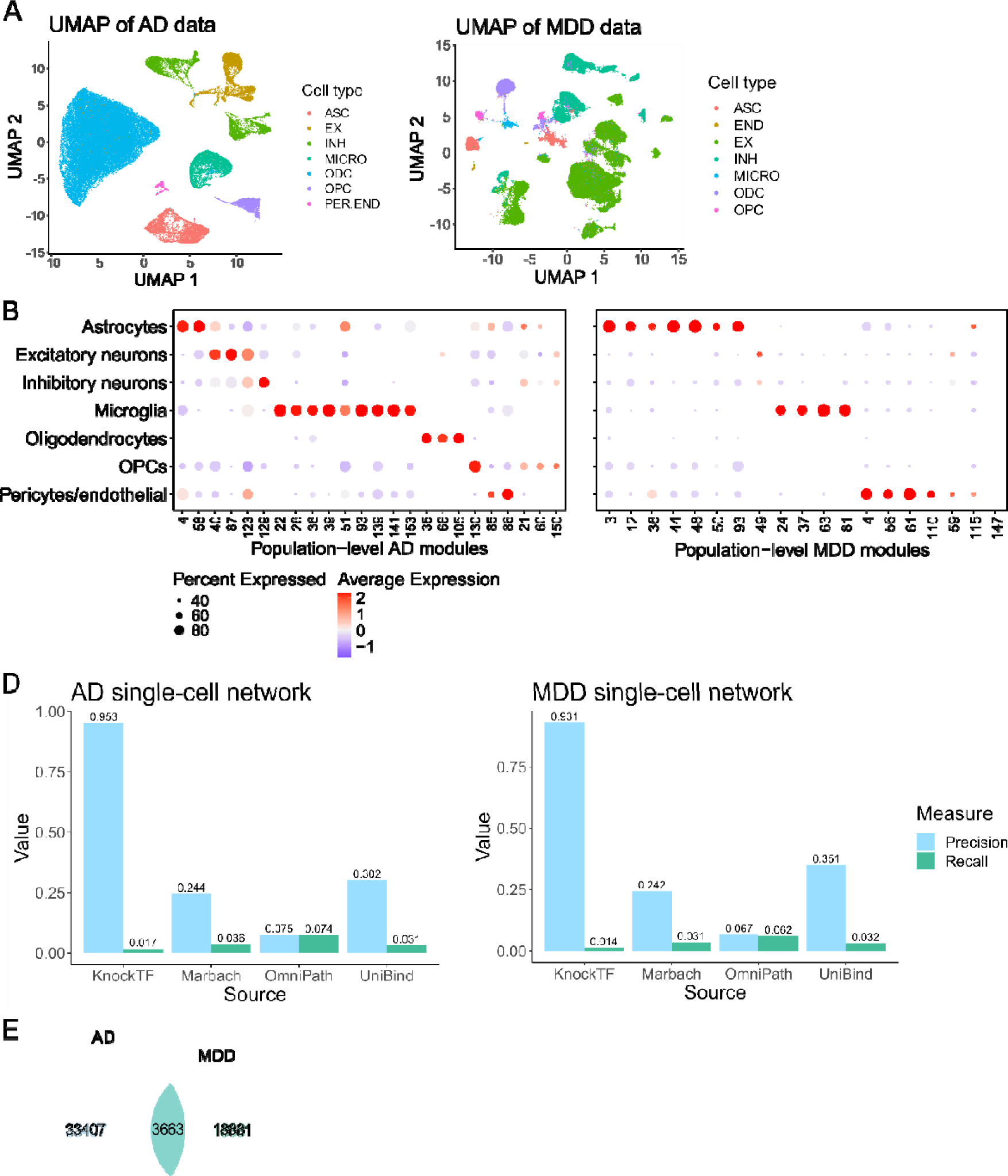
Single-cell network inference on Alzheimer’s disease (AD) and major depressive disorder (MDD) cohorts. A) UMAP plots of the AD and MDD datasets show the clusters and cell types of the datasets. For the AD dataset, the first 13 principal components were used, while for the MDD dataset, the first 50 principal components were used (based on elbow plots). B) Dotplot of population-level AD module projection into the single-cell AD data (left) and dotplot of population-level MDD module projection into the single-cell MDD data (right) show many modules with activity in specific cell types. Percent expressed indicates the number of cells from that cell type that express the module genes. The depicted modules are the prioritized ones also in Figure 2. In AD, most prioritized modules were active in microglia, while in MDD most prioritized modules were active in astrocytes. C) Precision and recall (AD, left; MDD, right) of the single-cell networks with all inferred edges with ground truth interactions from OmniPath and Marbach et al., KnockTF, and UniBind. The TF perturbation ground truth shows the highest precision and lowest recall. D) The Venn diagram shows that there were 3663 edges in common between the edges of the single-cell networks of AD and MDD. ASC: astrocytes, EX: excitatory neurons, INH: inhibitory neurons, MICRO: microglia, ODC: oligodendrocytes, OPCs: oligodendrocyte precursor cells, PER: pericytes, END: endothelial cells.

Next, to investigate gene regulation at the cell-type level, single-cell GRNs were inferred using SCENIC^48,49^, which outputs regulons, sets of an activating TF with its direct target genes, and their activity scores in specific cell types (Methods). The single-cell networks of MDD (22544 edges, 4814 nodes, 265 regulatory TFs) had fewer edges and nodes than the single-cell network of AD (37070 edges, 6360 nodes, 359 regulatory TFs, Supplemental Table 5). In an in silico validation against the ground truths defined above, the highest precision was obtained for the TF perturbations, similar to the population-level networks (Figure 6D).

Comparing the single-cell to the population-level consensus networks, 1.8-5% of the edges and 258 (AD) and 238 (MDD) TFs were shared (Supplemental Figure S4). As SCENIC makes use of the TF binding motif enrichment method RcisTarget, it identifies the direct binding of TFs to target genes, resulting in fewer predicted regulators and edges in the single-cell networks. There were 3663 edges in common between the AD and MDD single-cell networks, which was about 10% of the total edges of the networks, similar to the population-level networks (Figure 6E).

### 3.5. Regulon activity in the cell types of the brain

Next, we investigated the activity and specificity of the regulons in the different cell types of the brain. Different cell types were characterized by distinct regulatory programs, with some overlap in regulons between AD and MDD, for instance, PAX6 for astrocytes, LEF1 for endothelial cells, and ZMAT4 for excitatory neurons (Figure 7A, Supplemental Table 6). Some regulons were highly specific to a certain cell type, as indicated by a high regulon specificity score^49^, such as PAX6 in astrocytes in AD (Figure 7A, Supplemental Figure S5).

**Figure 7.**
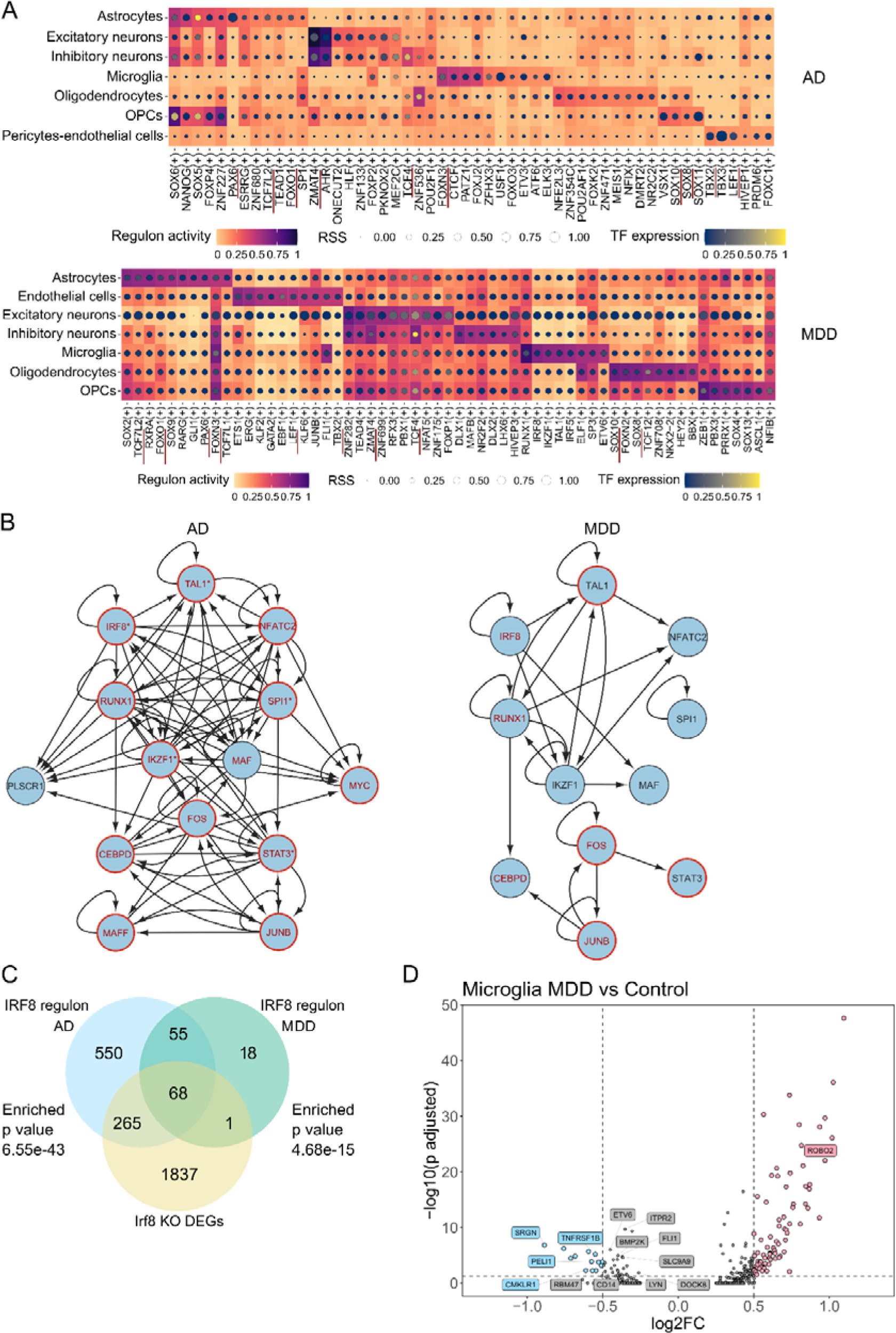
Implication of microglia, astrocytes, and oligodendrocytes in Alzheimer’s disease (AD) and major depressive disorder (MDD). A) Heatmap-dot-plots depicting the regulon activity (AUC values from SCENIC), RSS (regulon specificity score) and TF expression of selected regulons (top AUC scores) in the seven cell types of the brain. All values were pseudo-bulked, normalized for counts per cell type, and scaled over all TFs between zero and one for visualization. Top: AD, bottom: MDD. Different cell types were characterized by different regulatory programs, with some overlap between AD and MDD (red underlined). B) Single-cell core gene regulatory networks with 14 transcription factors (TFs) for AD (left) and MDD (right). The AD core network was more highly connected than the MDD network. TF nodes with a red frame have been implicated in the respective disease before through literature, while red letters indicate their target genes were enriched for DisGeNET genes involved in the respective disease. The TFs with * had differential edges (>90%) in AD compared to MDD. TFs not present in the MDD network were either not present in the full network, had no regulon, or were not regulated by the other TFs. C) Venn diagram of the overlap between the IRF8 regulons of the AD and MDD single-cell networks with the Irf8 conditional knock-out (KO) differentially expressed genes in microglia is depicted. The AD IRF8 regulon was larger and had a larger overlap with the DEGs. Besides the diagram, the p-values of the enrichment tests are indicated. E) Volcano plot of the differentially expressed genes in female microglia of MDD patients compared to controls indicated that most genes were found to be upregulated in MDD. Labeled genes were also part of the regulons of IKZF1, IRF8, RUNX, SPI1, and/or TAL1.

Furthermore, we investigated in which cell types the regulons controlled by the TFs steering the disease-specific neuroinflammation modules of the population-level networks, were active (Supplemental Figure S6). The regulons of *IKZF1, IRF8, MAF, NFATC2, RUNX1, SPI1,* and *TAL1* showed specifically high activity in the microglia in the single-cell AD networks, while they were most active in the microglia in the single-cell MDD networks. The regulons of *MAF* and *NFATC2* were not present in the single-cell MDD networks. The other group of TFs (CEBPD, FOS, JUNB, MAFF, MYC, and STAT3) was not specific to microglia, but their regulons were active in several cell types (Supplemental Figure S6). The overlap between population-level module genes and these single-cell-level regulon target genes is high for the AD regulons (± 69% of module genes found in regulons), but lower for the MDD regulons (± 27% of module genes found in regulons) (Supplemental Table 7). This could be explained by the fact that there were fewer genes in the regulons of the single-cell MDD networks. For the other prioritized population-level modules, there was only a small overlap in genes between the regulons of the TFs and the modules steered by these TFs. Thus, functional enrichment analysis was performed on these regulons to assess convergence at the pathway level. For most prioritized modules, their regulators had regulons in the single-cell networks of which their genes were involved in similar pathways as the population-level modules (Supplemental material). As an example, AD-module 35 genes functioned in lipid metabolism, oligodendrocyte specification and differentiation and myelin formation, was regulated by *BBX, CREB5, MYRF, PROX1, RBPJ, SOX8, SOX10, ST18, ZBED3* and *ZNF536,* and had a lower expression in AD patients compared to controls, indicating a dysfunction in myelination by oligodendrocytes in AD (Supplemental Figure S7). The regulons of BBX, SOX10, SOX8, RBPJ, and ZNF536 were most active in oligodendrocytes and oligodendrocyte precursor cells, and the SOX8 and SOX10 regulons contained genes enriched for oligodendrocyte differentiation and myelination. In conclusion, by inferring single-cell GRNs, we were able to implicate microglia and oligodendrocytes in both disorders and astrocytes in MDD.

To further investigate the coregulation of these TFs, we looked into the regulatory interactions between them in the inferred single-cell networks. Indeed, core regulatory circuits could be retrieved containing these TFs, highly regulating one another in both neuroinflammatory disorders with more interconnecting edges in the AD than in the MDD network (Figure 7B). Both core networks contained several feed-forward and feedback regulatory loops, as well as some self-regulating feedback loops, where TFs bind upstream of their own gene, as supported by TF motif information from SCENIC. Most genes of the IKZF1, IRF8, SPI1, and TAL1 MDD regulons were contained in the AD regulon of the same TF, but not the other way around because of the differences in size (Supplemental Table 8). The single-cell MDD network had fewer edges and smaller regulons compared to the single-cell AD network, which could result from a less informative dataset. This was also seen in the core regulatory circuit, with fewer TF-TF interactions in MDD. Next, to investigate regulon coregulation, we clustered the regulons based on the connection specificity index (CSI), a measure for the specificity of connections between TFs and target genes^132^ (Supplemental Figures S9B-E). Hence, a high CSI indicates the interaction is highly specific. Regulons in the same CSI module tended to coregulate target genes (Supplemental Figures 9B, 7D). In addition, the regulon clusters’ activity was visualized for each cell type, with differences in cell-type specificity for the distinct clusters (Supplemental Figures S9C, E). The regulons of *IKZF1, IRF8, MAF, NFATC2, RUNX1, SPI1,* and *TAL1* clustered together in both diseases, indicating that the shared immune TFs indeed worked together and steered similar target genes. As an example, *SOX8*, *SOX10* and *ZNF536* were, amongst others, regulatory TFs of myelination module 35, and their regulons appeared together in a CSI cluster (cluster 8) that was highly active in oligodendrocytes. To see whether there were differences in microglia cell states between control individuals and patients, we used hdWGCNA to project healthy microglia states from literature^69^. Fewer difference were found when comparing the MDD patients to controls than when comparing the AD patients to control individuals (Supplemental Figure S8).

### 3.6. Validation of TF-gene interactions in independent datasets

As further in silico validation, we re-analyzed scRNA-seq data of the conditional knock-out (KO) of Irf8 in a mouse model^72^. Based on differential expression between Irf8 KO and wildtype microglia, we demonstrated that the target genes of the AD and MDD IRF8 regulon were enriched in the human orthologs of these DEGs (2442 mouse genes, corresponding to 2171 human genes) with a p-value for AD of 6.55e-43, and a p-value for MDD of 4.68e-15, indicating the single-cell networks were biologically valid (Figure 7C). Additionally, re-analyzing preprocessed single-cell MDD data from female individuals from Maitra and colleagues^75^, we found 14 DEGs (*ROBO2*, *SRGN*, *ETV6*, *ITPR2*, *TNFRSF1B*, *BMP2K*, *PELI1*, *SLC9A9*, *RBM47*, *CMKLR1*, *FLI1, DOCK8, CD14* and *LYN*) between MDD and control samples in microglia that were also inferred target genes of the core regulatory circuits active in microglia (IKZF1, IRF8, RUNX1, SPI1, and TAL1) (175 DEGs (adj. p-value ≤ 0.05) in total in microglia, figure 7D). More specifically, 11 targets of IKFZ1, eight targets of IRF8, ten targets of RUNX1, three targets of SPI1, and five targets of TAL1 were differentially expressed in the validation dataset. This indicates that the core circuit is biologically relevant and altered in MDD compared to control individuals in both sexes.

### 3.7. Disease specificity of the GRNs

To investigate differences in regulatory programs between AD and MDD at the single-cell level, we performed differential edge analysis between the single-cell GRNs (Methods). Target genes and TFs were retained of which more than 20% of the edges were differential (i.e. only present in one of the two networks) compared to all edges of that gene. This resulted in 2308 target genes and 168 TFs. Next, we looked at whether these differential edges were gained in the AD network or the MDD network, using a cut-off of 90% gained edges. Overall, 665 target genes had gained edges in AD (hereafter AD-biased genes), 94 in MDD (hereafter MDD-biased genes) and 1549 target genes had differential edges that were neither more than 90% gained in AD nor MDD. Similarly, for the TFs, 26 TFs had over 90% gained edges in AD, 11 TFs in MDD, and 131 in neither. A probable reason for the larger number of genes with differential edges in the AD network is because of the larger number of edges and larger regulons inferred in this network compared to the MDD network. The AD-biased TFs include *IKZF1*, *IRF8*, *MEF2D*, *RUNX3*, *SPI1*, and *TAL1,* indicating rewiring of the target genes of these TFs in AD compared to MDD. The MDD-biased TFs include *FOXN3*, *MAFB*, and *SMAD1* (Supplemental Table 9). This does not mean that these regulons are not important in the other disease, but rather that they had distinct target genes, and this was partially driven by the differences in regulon sizes. Interestingly, the functional enrichment analysis of the biased target genes revealed terms related to the respective disease (Figure 8). Terms enriched for the AD-biased genes include cellular response to cytokine stimulus, regulation of immune response, TNF signaling pathway, TYROBP causal network in microglia, and others. On the other hand, the MDD-biased genes were enriched for terms related to sodium ion channel transport, exocytosis, and action potential, thus, synapse-related functions.

**Figure 8.**
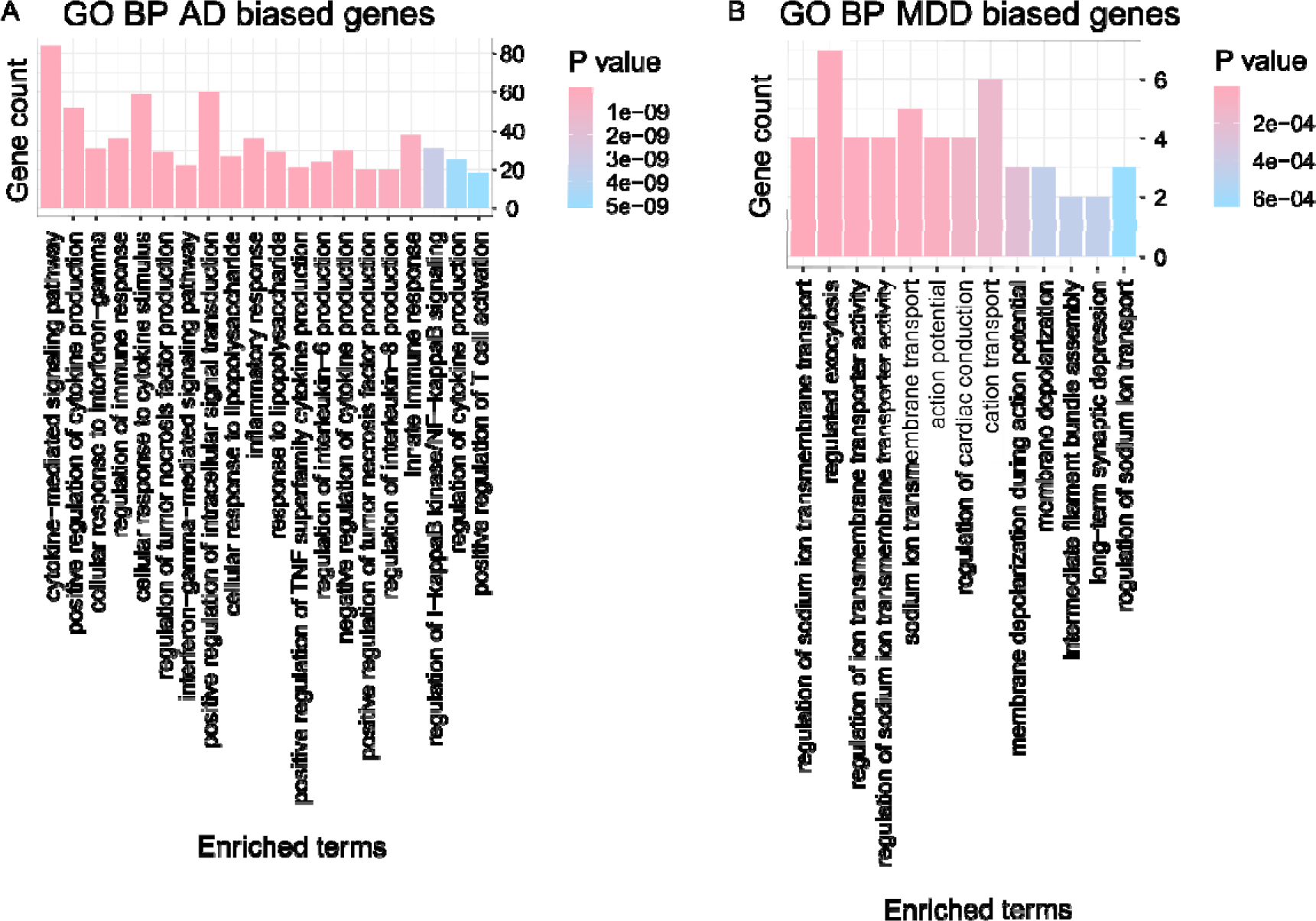
Functional enrichment analysis of disease-biased genes. A, B) The graphs for the differential node targeting functional enrichment analysis depict the top 20 terms from GO Biological Process for the AD-biased (A) and MDD-biased (B) genes. The AD-biased genes were involved in immune pathways, while the MDD-biased genes were mainly involved in synapse-related functions. TNF: tumor necrosis factor.

To further characterize the disease-specificity of the TFs of the core circuit, we did an enrichment analysis with DisGeNET^63^ (database of gene-disease associations) of the target genes of each TF in the single-cell and population-level networks of both disorders. For each TF in the AD/MDD networks, we looked at whether their target genes were enriched in the gene sets of AD/MDD in DisGeNET (Methods, Supplemental Table 10). TFs of which target genes were enriched in both single-cell and population-level networks for the same disease were, amongst others, ATF3, CEBPD, FOS, IRF8, KLF6, and NFATC2 in the AD networks (12 in total), and JUNB, SOX2, SOX9, SOX21, and RUNX1 in the MDD networks (14 in total). Hence, with this analysis, we again link several TFs of the core circuit to the diseases (Figure 6B). There were three TFs of which the target genes were enriched for disease genes in all four networks (population- and single-cell level and both disorders): FOSB, FOSL2, and NFIL3, in agreement with literature^109,133,134^. Hence, our results suggest that the regulatory rewiring observed in AD and MDD may be driven by a combination of shared and disease-specific factors, necessitating further investigation into their interplay.

## 4. Discussion

In this study, we unraveled common and distinct regulatory programs of neuroinflammation and other pathophysiological processes in AD and MDD, thereby discovering novel underlying molecular mechanisms that could facilitate the development of novel treatments for neuroinflammatory disorders in the long term. The field of GRN mapping in neuroinflammatory disorders has grown quickly over the last few years. However, limitations remain by using bulk data, only a single network inference method^17^, a single disease, or only DEGs^135^ to comprehensively and cell-type specifically infer GRNs. In the first part of this work, population-level GRNs were inferred with different methods (GENIE3, CLR and Lemon-Tree). The overlap between the networks retrieved from the different methods and with the ground truth interactions highlights the methods’ complementarities in retrieving biologically relevant regulatory interactions. Therefore, more robust consensus networks were made for each disorder, which were decomposed into coregulated modules. We prioritized several modules based on differential expression between patients and controls, and functional enrichment related to the immune system, to find disease-specific neuroinflammation modules. Our investigation revealed shared coregulatory TFs orchestrating the dysregulation of immune functions, including cytokine response, microglia activation, and phagocytosis, in both disorders.

In addition, networks were inferred with the single-cell GRN inference method SCENIC^48^ on snRNA-seq data, which retrieved cell-type-specific regulons for both AD and MDD. Even though bulk RNA-seq data have more reads per sample and contain more sample heterogeneity as typically more samples are sequenced, all reads are summed over the cells, and the results are influenced by the number of cells of each cell type present in the samples. Additionally, while our methods on population-level were only based on transcriptomics data, SCENIC uses TF-binding information to refine direct regulatory interactions. Most of the TFs predicted to regulate several population-level, disease-specific neuroinflammation modules had regulons that were highly active in microglia, in both AD and MDD. The IKZF1, IRF8, SPI1 and TAL1 regulons were specific to microglia in both diseases, while the RUNX1 regulon in MDD was also active in the other cell types, but was specific to microglia in AD. Thus, a large part of the genes in the immune-related modules are probably highly expressed in microglia. It is already known that microglia play a pivotal role in both AD and MDD^136,137^. Additionally, our findings implicated astrocytes in MDD and revealed an enrichment of apoptosis genes, implying elevated astrocytic apoptosis in depression. A reduction in the number of astrocytes can be caused by chronic stress and several older studies support that there are significant reductions in the number and density of astrocytes in several brain regions in MDD^9^, which we also showed by investigating the cell type proportions in scRNA-seq data and which was confirmed by Maitra and colleagues^75^.

Interestingly, we found core regulatory circuits in both disorders that regulated each other and similar target genes (both at module and regulon level). Besides, we could also demonstrate that the identified TFs and their regulons in the shared core regulatory circuits were disease-specific a.o. with the differential edge targeting, the DisGeNet enrichment, and the differential expression in the external MDD dataset. *SPI1* was already reported to be implicated in AD, where it was regulated by *NFKB1* and its expression strongly correlated with AD/Braak progression in the hippocampus^138^. In addition, upon transcriptome and chromatin accessibility profiling in primary human microglia, *SPI1* was identified as a key putative regulator of microglia gene expression and AD risk^139^. In another study, *SPI1* and *STAT3* were central regulators of AD-associated microglia^24^. Furthermore, in AD brains, *SPI1* was found as a common master regulator. Additionally, they discovered *TAL1* to be implicated in AD, and similar to our work, they found modules with immune-related processes and myelination functionalities^22^. Lastly, in a recent preprint of an AD atlas by the Seattle Alzheimer’s Disease Cell Atlas (SEA-AD) consortium, microglial GRNs were inferred using SCENIC+ and they identified RUNX1, IKZF1, NFATC2, and MAF as TFs driving up-regulation of pro-inflammatory and plaque-induced genes, with specific expression in microglia^140^, confirming our core regulatory circuits in AD.

We found SPI1 as a master regulator and a downregulated module 35 in AD involved in myelination, similar to the original publication of the AD bulk and single-cell data, where they mostly focused on oligodendrocytes^33^. Furthermore, we discovered immune system alterations in MDD similar to the original publication of the single-cell MDD data, where they focused on differential gene expression between MDD and healthy brain samples, mainly identified in immature oligodendrocyte-precursor cells and deep layer excitatory neurons, but implicating also other neuronal and non-neuronal cell types^29^.

Interestingly, of the top 100 TFs in our AD and MDD population-level networks (Supplemental table 2), MEF2C and TFEB were two shared TFs between AD and MDD, which other researchers identified as hub TFs in AD^20^. TFEB has been put forward as a promising candidate for AD treatment^141^ and MEF2C was discovered to be highly involved in regulatory interactions in AD^142^. MEF2C plays an essential role in hippocampal-dependent learning and memory and is crucial for normal neuronal development and electrical activity^143^. It is also involved in some immune functions^144^ and has been linked to AD and MDD before^145,146^. The regulon of MEF2C was highly active in both the excitatory and inhibitory neurons in our single-cell network of AD. In addition, they discovered upregulated microglia-enriched TF networks in AD and found IKZF1, IRF8, and RUNX1 target genes enriched in microglia^20^. However, no results for MDD were reported since no DEGs were found in their dataset^20^.

In summary, our investigation revealed shared dysregulation of neuroinflammation and microglial activation in both AD and MDD, under the control of common coregulatory TFs. In addition, distinct dysregulated pathways and cell types were identified, specific to each disorder. Notably, this study has highlighted pivotal TFs and genes, primarily associated with AD and with limited prior associations with MDD. The potential relevance of these key TFs and genes in MDD warrants additional investigation, offering promising avenues for therapeutic intervention.

## 5. Data availability

Data is available with the original authors.

## 6. Acknowledgments

Funding: This work was supported by the Ghent University Special Research Fund [grant number BOF/STA/201909/030]. We thank Jane Foster and Boris Vandemoortele for the critical reading of this work and their helpful suggestions.

## 7. Author contributions

Contributions according to the CRediT system (https://www.ucl.ac.uk/library/research-support/open-access/credit-taxonomy), HP = Hanne Puype, JD = Joke Deschildre, VV = Vanessa Vermeirssen): Conceptualization: HP, VV, Data curation: HP, Formal analysis: HP, Funding acquisition: VV, Investigation: HP, Methodology: HP, VV, Project administration: VV, Software: HP, JD, Supervision: VV, Validation: HP, Visualization: HP, Writing— original draft: HP, VV, Writing—rewriting and editing HP, VV. All authors read and approved the final manuscript.

