## Supplemental material for "Comparative gene regulatory network analysis in Alzheimer’s disease and major depressive disorder identifies shared core regulatory circuits"

1. **Supplemental text**
   1.1. **Functional annotation of additional modules implicated disruption in mitochondrial, myelination and GABA receptor signaling processes**

All figures of the prioritized modules can be found in the Supplemental material. AD-module 26 was involved in cholesterol biosynthesis and intracellular protein transport, with a lower expression in AD patients. The TFs of module 26 were *ATMIN, CSRNP3, HSF2, MEF2C, PEG3, SATB1, ZBTB11, ZNF25, ZNF384* and *ZNF814*. Module 35 from the AD network had functional terms related to membrane lipid biosynthesis, actin reorganization, cell movement, and the term Oligodendrocyte specification and differentiation, leading to myelin components for CNS. The expression of the AD patients was lower compared to the controls, indicating a dysfunction in myelination by oligodendrocytes in AD. Interestingly, researchers found that myelination can be impaired by cholesterol dysregulation in oligodendrocytes^1^. The TFs of this module were *BBX, CREB5, MYRF, PROX1, RBPJ, SOX8, SOX10, ST18, ZBED3* and *ZNF536*. Module 66 contained genes with functions related to GTP binding, pathways of neurodegeneration (*COX8A, TUBA1A, TUBB, CHMP2B, NDUFC1, RAC1, TUBB4A*, and *COX6B1*), ‘Alzheimer disease’ (*COX8A, TUBA1A, TUBB, NDUFC1, TUBB4A, COX6B1*) and metabolism. The genes in this module had an overall lower expression in AD patients compared to controls. The TFs of this module were *HINFP, STOX2, ZNF552, ZNF587B* and *ZNF814*. MDD module 44 was involved in the regulation of macrophage activation, transport across the blood-brain barrier, and aminoglycan biosynthetic process. The expression was lower in MDD patients compared to control individuals. The TFs were *GLI3, NPAS3, NR2E1, PAX6, PPARA, RFX4, SOX9, SOX21, STOX1* and *TRPS1*. MDD-module 49 contained genes related to the regulation of neuron death, secretion, and protein transport. The expression was higher in MDD patients. *MYPOP, NR2F6, POU6F1, USF2, ZBTB45, ZNF775,* and *ZNF777* were the TFs of this module.

1.2. **Overlap between population-level modules and single-cell regulons on pathway level**

AD-module 153 (apoptosis and inflammatory response) was steered by TFs of which the regulons were most active in microglia (*CEBPD*), and endothelial cells (*CEBPD* and *LEF1*). The AD regulon of CEBPD contained genes involved in cytokine response, apoptosis, T_H_ cell differentiation, and cellular senescence, while the regulon of *LEF1* also contained genes involved in cytokine response. MDD-module 36 (cytokine response and apoptosis) had TFs most active in astrocytes (*FOXO1, SOX9, STAT3*). The MDD regulon of *FOXO1* had genes enriched in positive regulation of apoptotic process, regulation of cell differentiation, and response to cytokine stimulus. MDD-module 93 had some TFs (*PAX6, PPARA, SOX21*) of which the regulons were most active in astrocytes in the single-cell networks, while the module had functional enrichment terms related to oligodendrocytes, regulation of nervous system development, GABA receptor signaling, and BDNF. The MDD regulons of *PAX6* and *SOX21* both contained genes involved in nervous system development, while the PPARA regulon was associated with the regulation of lipid transport. MDD-module 110 (cytokine signaling and unfolded protein response) contained TFs most active in endothelial cells in the MDD single-cell networks (*FOS, JUNB, KLF4*), as well as inhibitory neurons (*ATF3*), and mixed cell types (*JUN, JUNB*). The MDD regulons of *FOS*, *JUN*, and *JUNB* contained genes involved in response to cytokine stimulus, stress and unfolded protein, and apoptosis, while the *KLF4* regulon was involved in cytokine response. Lastly, MDD-module 115 (inflammatory response, apoptosis, glucose homeostasis, and regulation of lipid storage) had TFs of which the regulons were most active in astrocytes (*FOXO1, STAT3*) and inhibitory neurons (*NFIB*). The MDD *FOXO1* regulon contained genes related to cytokine response, while the *NFIB* regulon contained terms involved in brain development and chemotaxis.

Module 35 from the AD network, with terms involving lipids and oligodendrocytes, had TFs (*BBX, SOX10, SOX8, RBPJ, ZNF536*) of which the regulons were most active in oligodendrocytes and oligodendrocyte precursor cells. The AD regulons of *SOX8* and *SOX10* had genes enriched for oligodendrocyte differentiation, myelination, amoeboidal-type cell migration, and oligodendrocyte specification and differentiation, leading to myelin components for CNS. Module 44, containing terms related to transport across the blood-brain barrier, had TFs of which the regulons were most active in astrocytes (*PAX6, SOX9, SOX21, PPARA*). The *PAX6* regulon of the single-cell MDD network contained genes related to the maintenance of the blood-brain barrier, and macrophage chemotaxis, amongst others. The *SOX9* regulon was also enriched for the term transport across the blood-brain barrier. In modules 36 and 44, the expression was lower in the patients. This analysis further implicated oligodendrocytes, oligodendrocyte precursor cells, and excitatory neurons in AD. In contrast, in MDD, astrocytes and inhibitory neurons were found to play an important role.

Beyond the immune-related processes, the AD network also revealed perturbed pathways encompassing mitochondrial functions, the proteasome system, and myelination by oligodendrocytes. Myelin damage has been implicated before in AD, but it is not yet known whether the damage and death of oligodendrocytes is a result of the Aβ plaques and neurofibrillary tangles, or whether it is more a primary event^2^. On the other hand, our analysis highlighted GABA (γ-amino butyric acid) signaling, BDNF (brain-derived neurotrophic factor) and dysregulation of nervous system development in MDD. Additionally, our findings implicated astrocytes and revealed an enrichment of genes linked to apoptosis, implying elevated astrocytic apoptosis in depression. In conclusion, by inferring single-cell GRNs, we were able to implicate microglia in both disorders, oligodendrocytes in AD and astrocytes in MDD.

Several methodological limitations require consideration in our study. Firstly, only TFs were included as regulators, without incorporating chromatin modifiers, co-factors, or regulatory RNA molecules like lncRNA and miRNA. Furthermore, our population-level network inference relied solely on transcriptomics data, potentially introducing indirect interactions. Nevertheless, SCENIC utilizes additional motif data, resulting in the inference of direct interactions. Additionally, by using publicly available data, there is less control over the used samples. For instance, it is not always known if the patients took medication, batch effects had to be accounted for regarding the MDD datasets and the single-cell MDD dataset consisted of only male samples. Furthermore, the predominant focus on individuals of European descent within the research landscape translates to a limited genetic diversity, restricting the generalizability of our findings to broader populations. Lastly, post-mortem brain samples are prone to the degradation of RNA molecules, and these samples only represent one snapshot of the patient’s lifetime^3^. Our focus was on one specific brain region, underscoring the need for further exploration across other brain regions. As described in the review by Poletti and colleagues, the inflammatory markers found in MDD are highly dependent on the phase of the disease and are influenced by treatment and age^4^. Hence, in future studies, stratification according to these factors, as well as sex, will be important.

**2. Supplemental Figures**

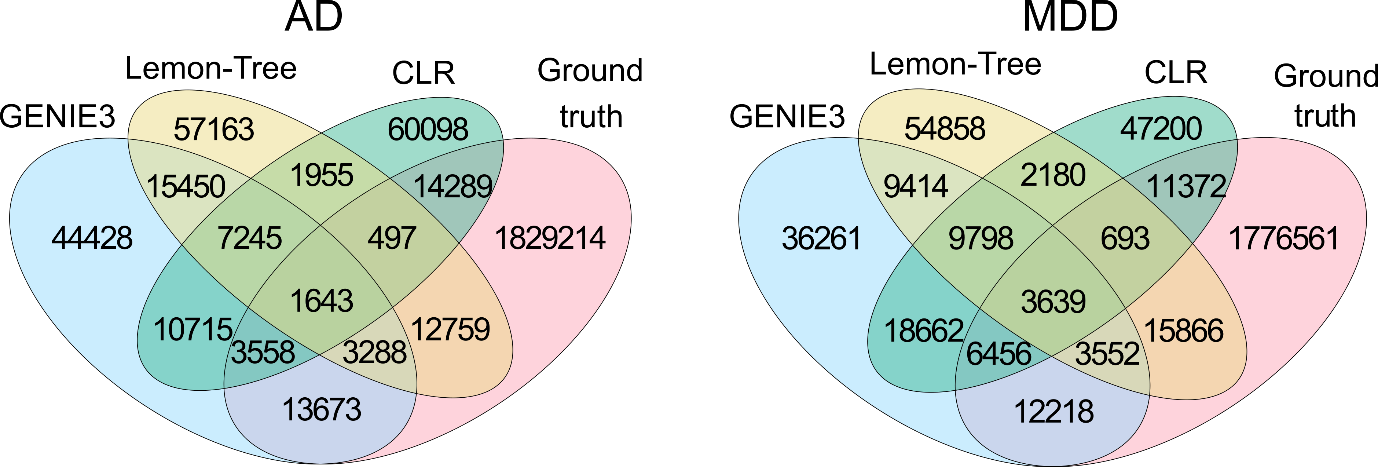

**Supplemental Figure S1.** Venn diagrams of the overlap between the three network inference methods (GENIE3, CLR and Lemon-Tree) and the regulatory interactions from the ground truths for AD (left) and MDD (right) point to the complementarity of the different methods in predicting the ground truth.

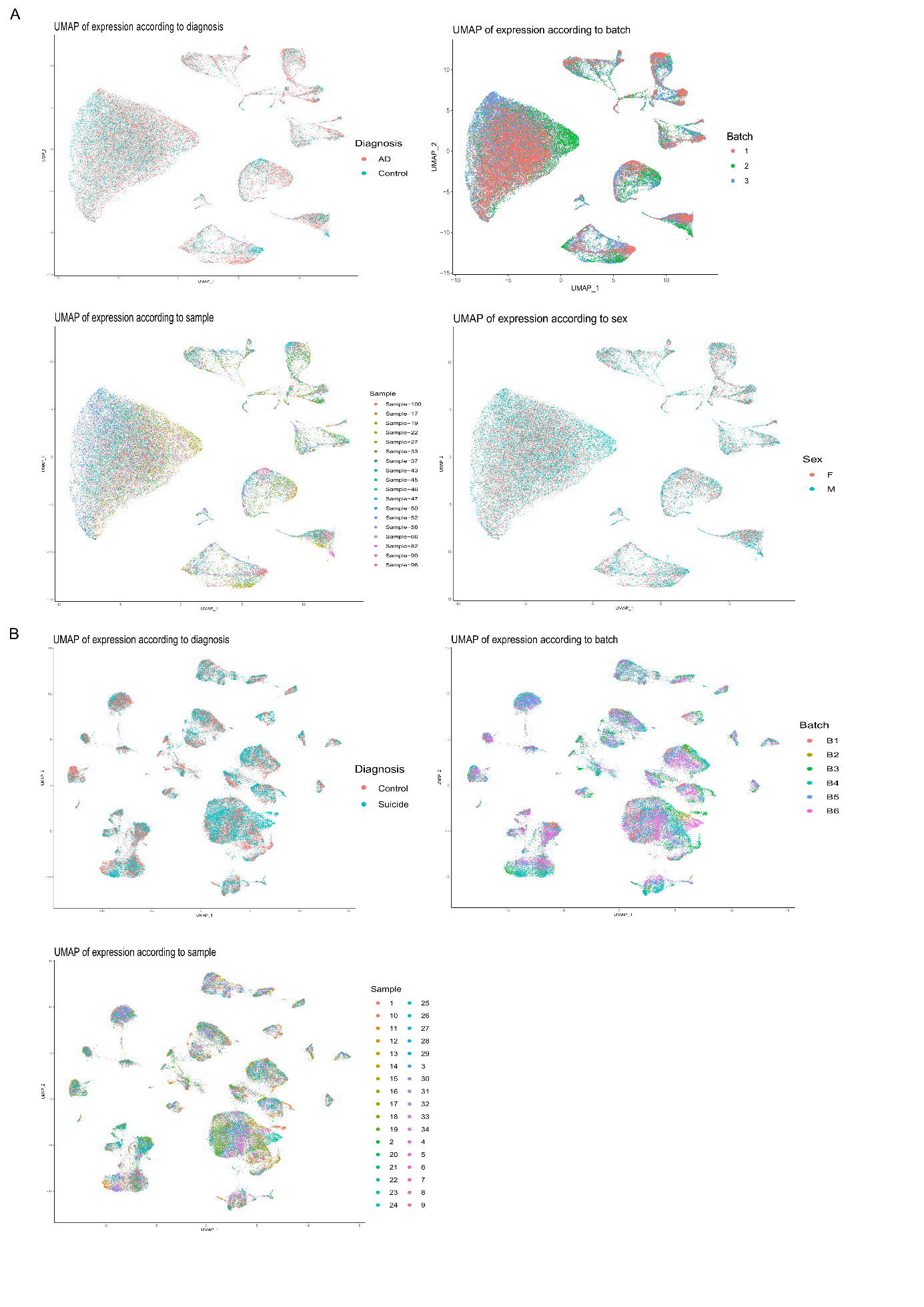

**Supplemental Figure S2. Uniform Manifold** **Approximation and Projection (UMAP) plots annotated with the metadata of the single-cell Alzheimer’s disease (AD) and major depressive disorder (MDD) datasets.** A) UMAPs of the single-cell AD dataset with the plots annotated by diagnosis, batch, sample or sex are depicted. Metadata comes from the original publication. UMAPs were plotted using the first 13 principle components. B) UMAPs of the single-cell MDD dataset with the plots annotated by diagnosis, batch or sample are depicted. All samples were male and the metadata is from the original publication. UMAPs were plotted using the first 50 principle components. None of the UMAPs show clustering according to the metadata annotation.

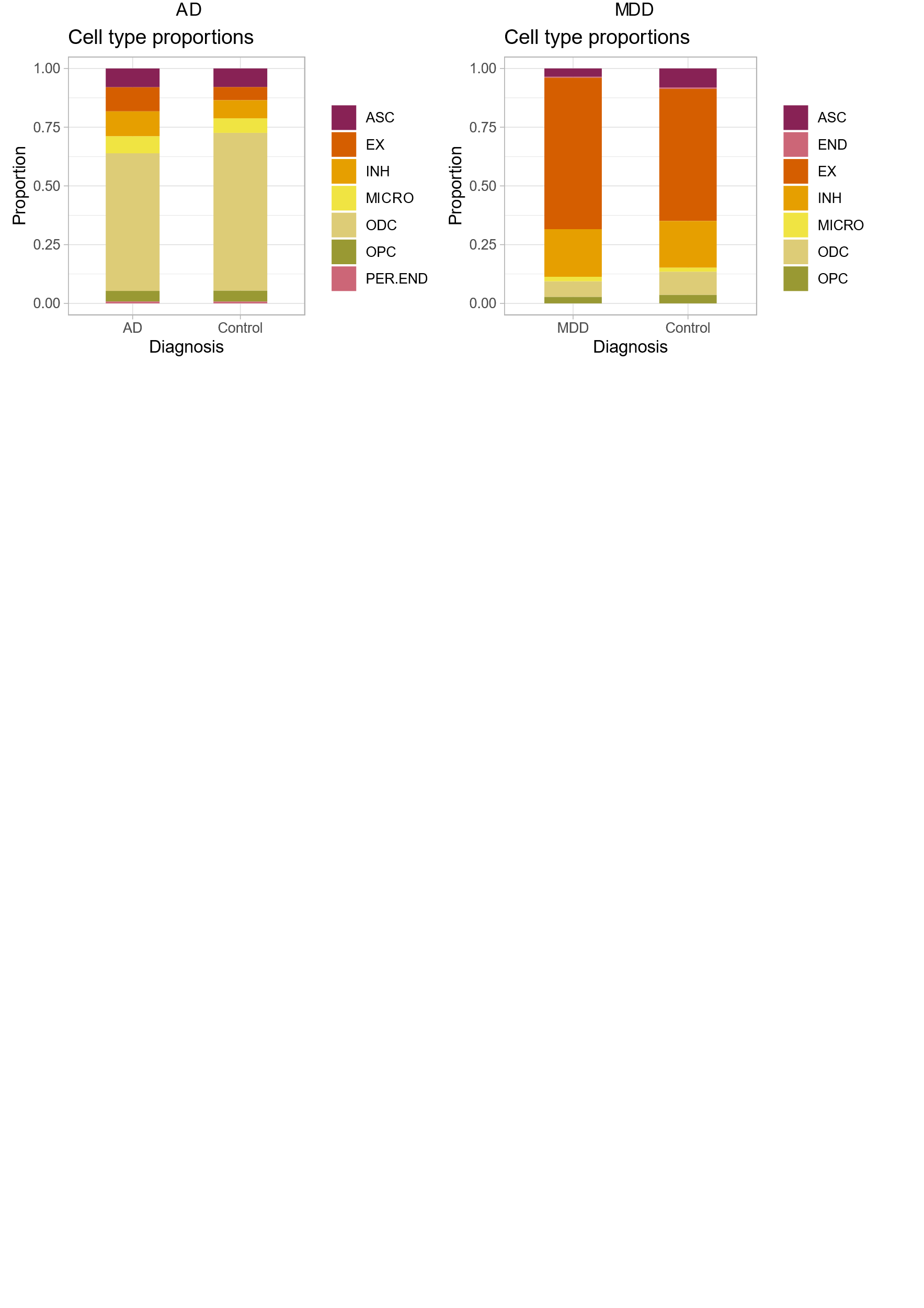

**Supplemental Figure S3. Cell type proportions of the AD and MDD single-cell dataset split by diagnosis.** Astrocytes were significantly different in MDD compared to control. ASC: astrocytes, EX: excitatory neurons, INH: inhibitory neurons, MICRO: microglia, ODC: oligodendrocytes, OPC: oligodendrocyte-precursor cells, PER: pericytes, END: endothelial cells.

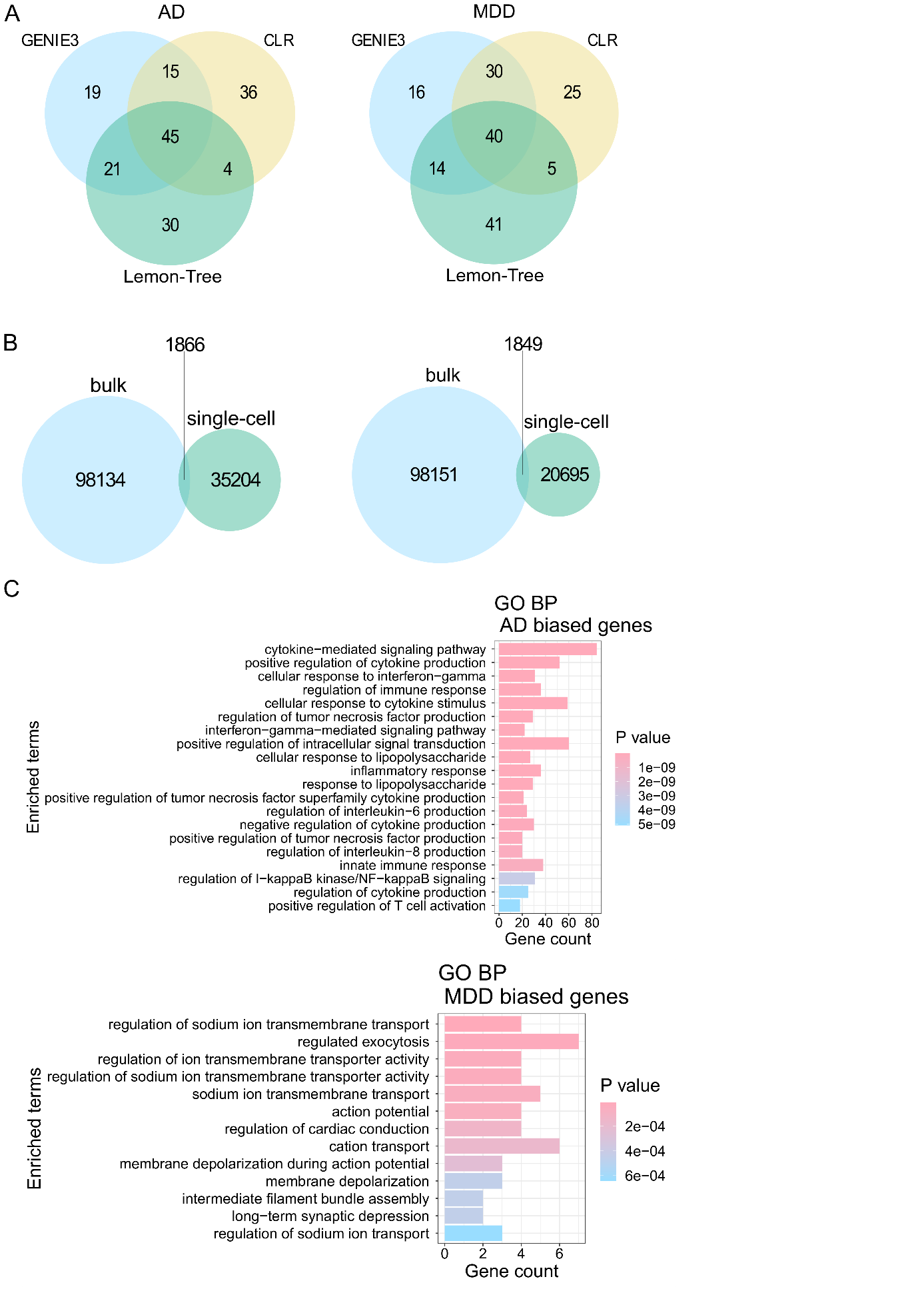

**Supplemental Figure S4.** **Venn diagrams of the overlap between population-level (bulk) networks and single-cell networks for AD (left) and MDD (right****).** A similar number of edges were common between the bulk and single-cell networks for both diseases, showing the impact of using a different method for GRN inference.

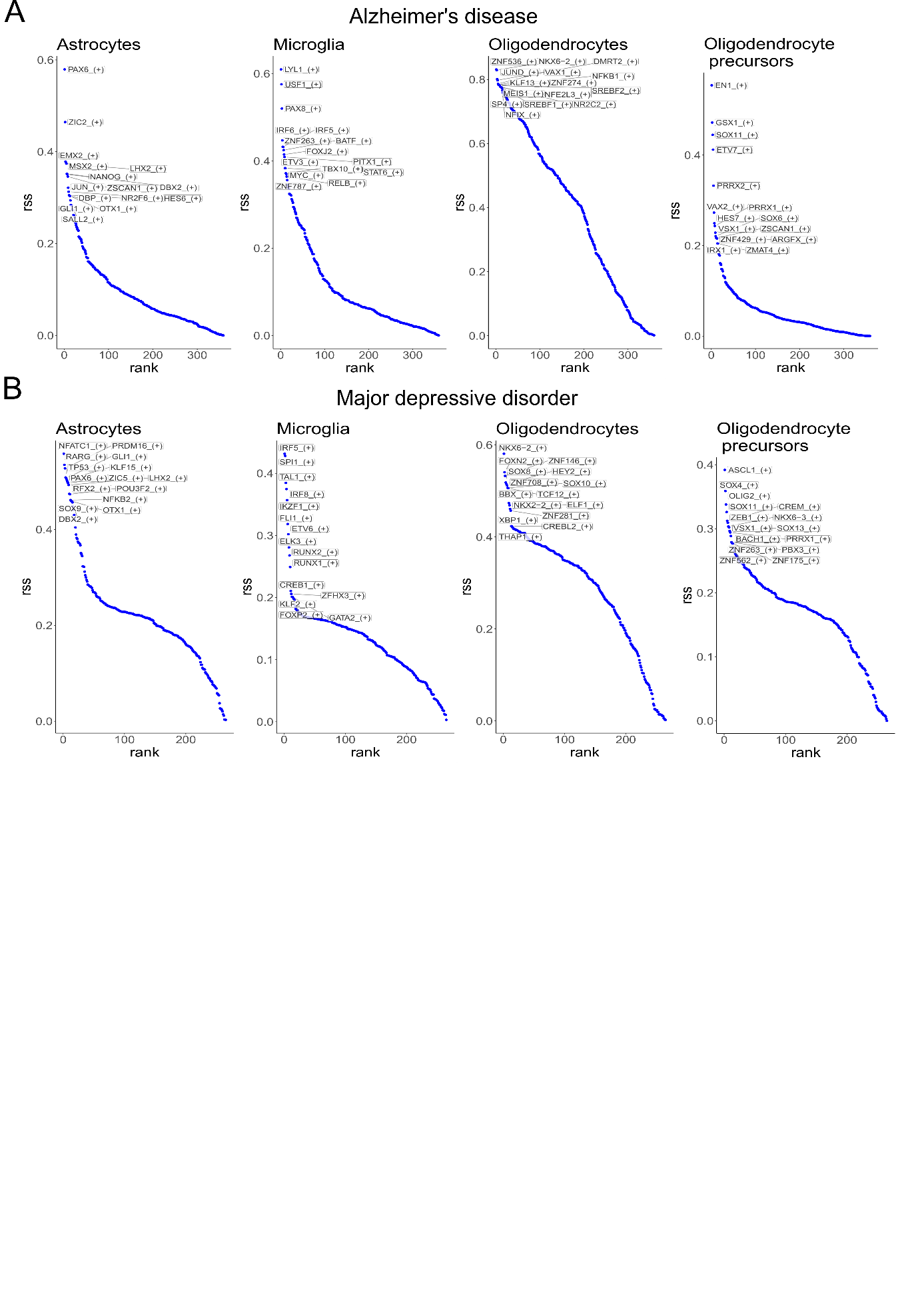

**Supplemental Figure S5. Regulon specificity scores (RSS).** The RSS is depicted on the y-axis, the rank of the regulons is depicted on the x-axis. The top fifteen regulons are highlighted on the plots. A) RSS plots from the single-cell Alzheimer’s networks for each glial cell type. B) RSS plots from the single-cell depressive networks for each glial cell type. The ZSCAN1 regulon was in both the astrocytes’ and oligodendrocyte precursor cells’ top regulons in the single-cell AD networks. The regulons of DBX2, GLI1, LHX2, OTX1, and PAX6 were in the top regulons of the astrocytes for both AD and MDD single-cell networks. The IRF5 regulon was in the top in microglia in both networks, and the NKX6-2 regulon was a shared top regulon of the oligodendrocytes. Lastly, the regulons of SOX11, PRRX1, and VSX1 were shared top regulons in both diseases in the oligodendrocyte precursor cells.

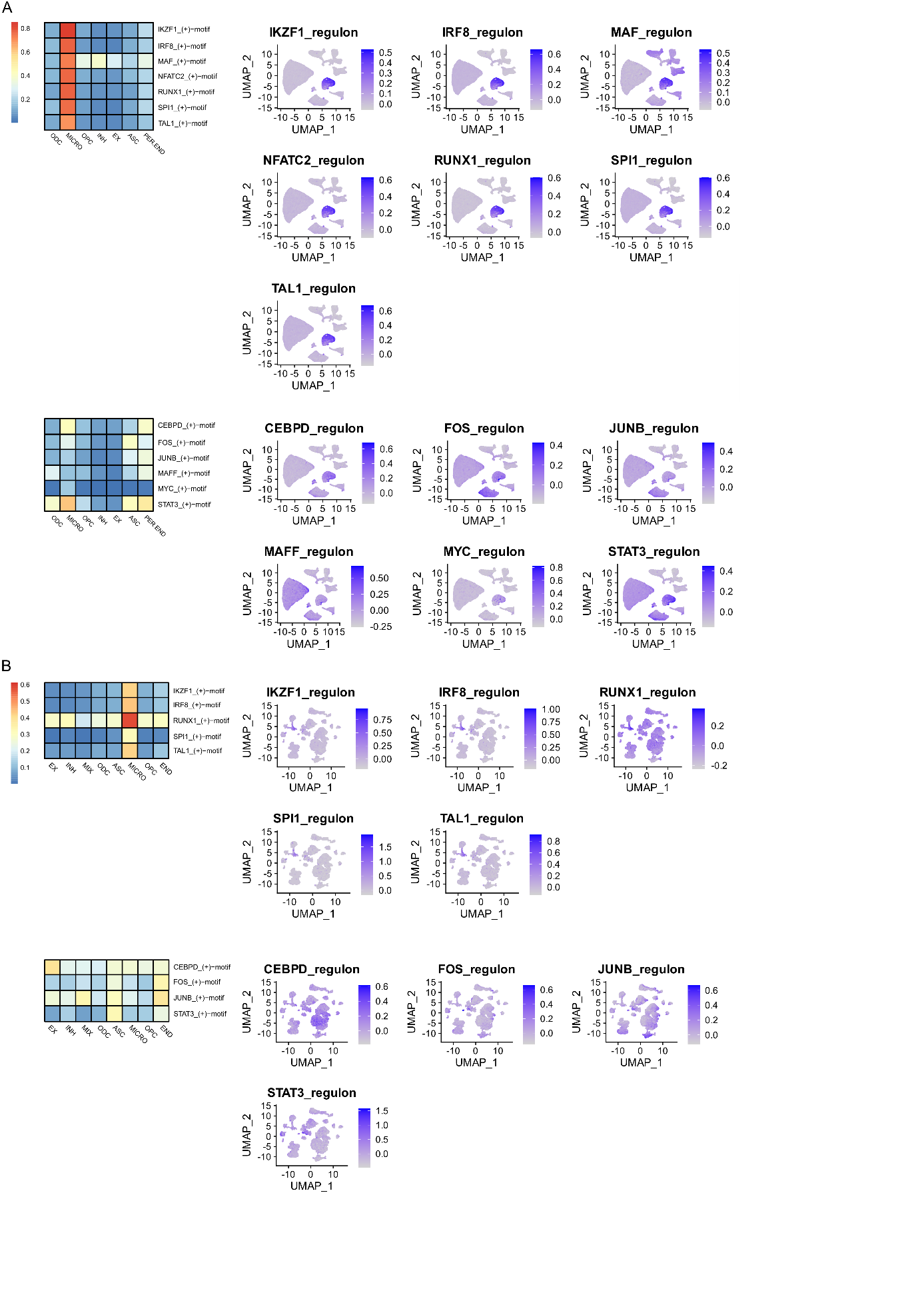

**Supplemental Figure S6. Regulon activity and expression of immune regulons.** On the left, the regulon activity (AUC values) from SCENIC is depicted for each cell type (EX: excitatory neurons, INH: inhibitory neurons, MICRO: microglia, ASC: astrocytes, ODC: oligodendrocytes, OPC: Oligodendrocyte precursor cells, END: endothelial cells, PER: pericytes). On the right, UMAPs are depicted with the expression of the regulon genes for each regulon. A) AD, B) MDD. The first block of regulons contains regulons that are most active in microglia, while the second block of regulons are active in several cell types.

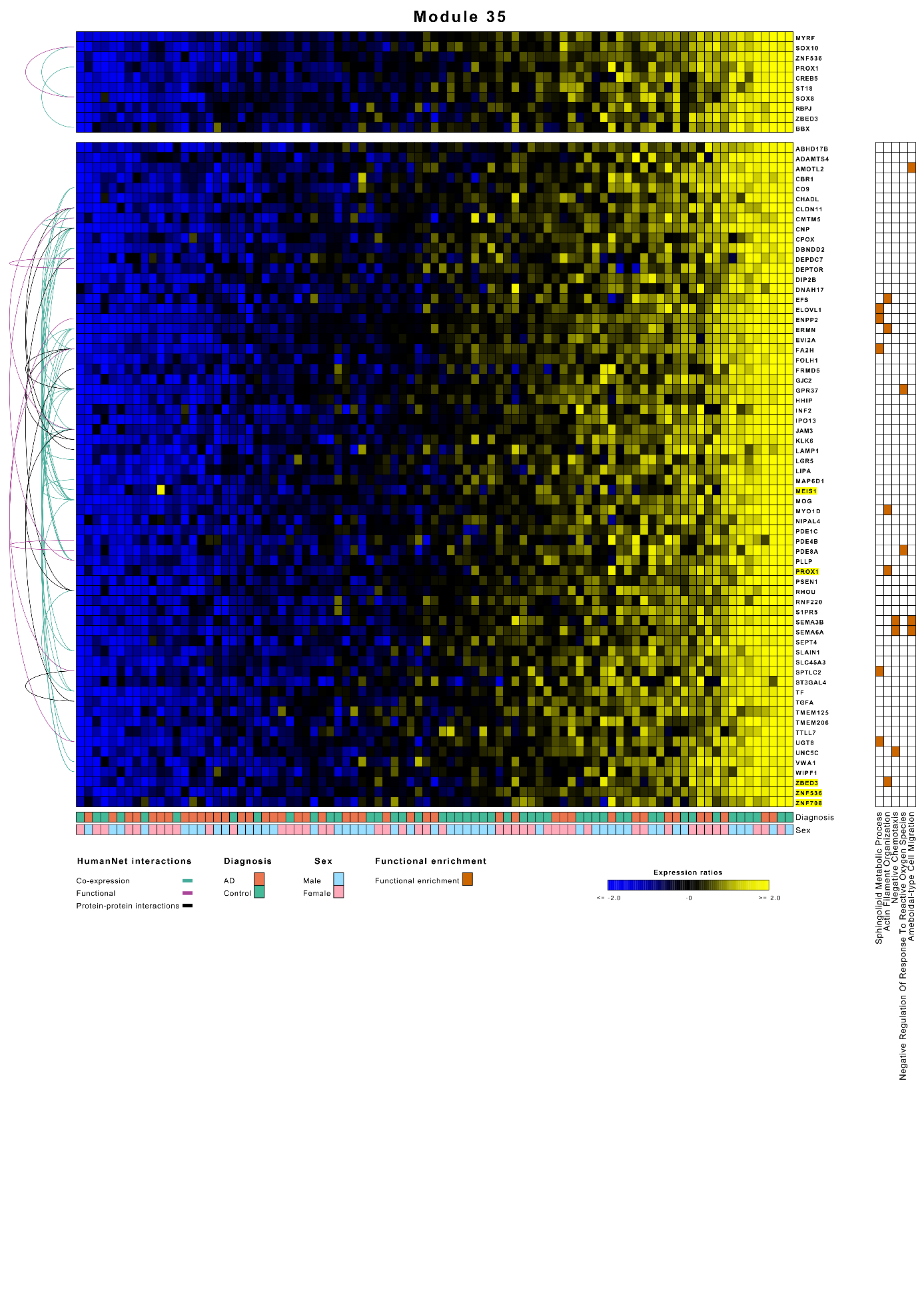

**Supplemental figure S7.** **Module 35 of the AD network contained 66 genes and was regulated by BBX, CREB5, MYRF, PROX1, RBPJ, SOX10, SOX8, ST18, ZBED3, and ZNF536.** It was involved in lipid metabolism, actin reorganization, cell movement, and oligodendrocyte specification and differentiation. The ModuleViewer figure includes the sample diagnosis, functional enrichment terms, and interactions from HumanNet (see legends). Highlighted module genes are TFs.

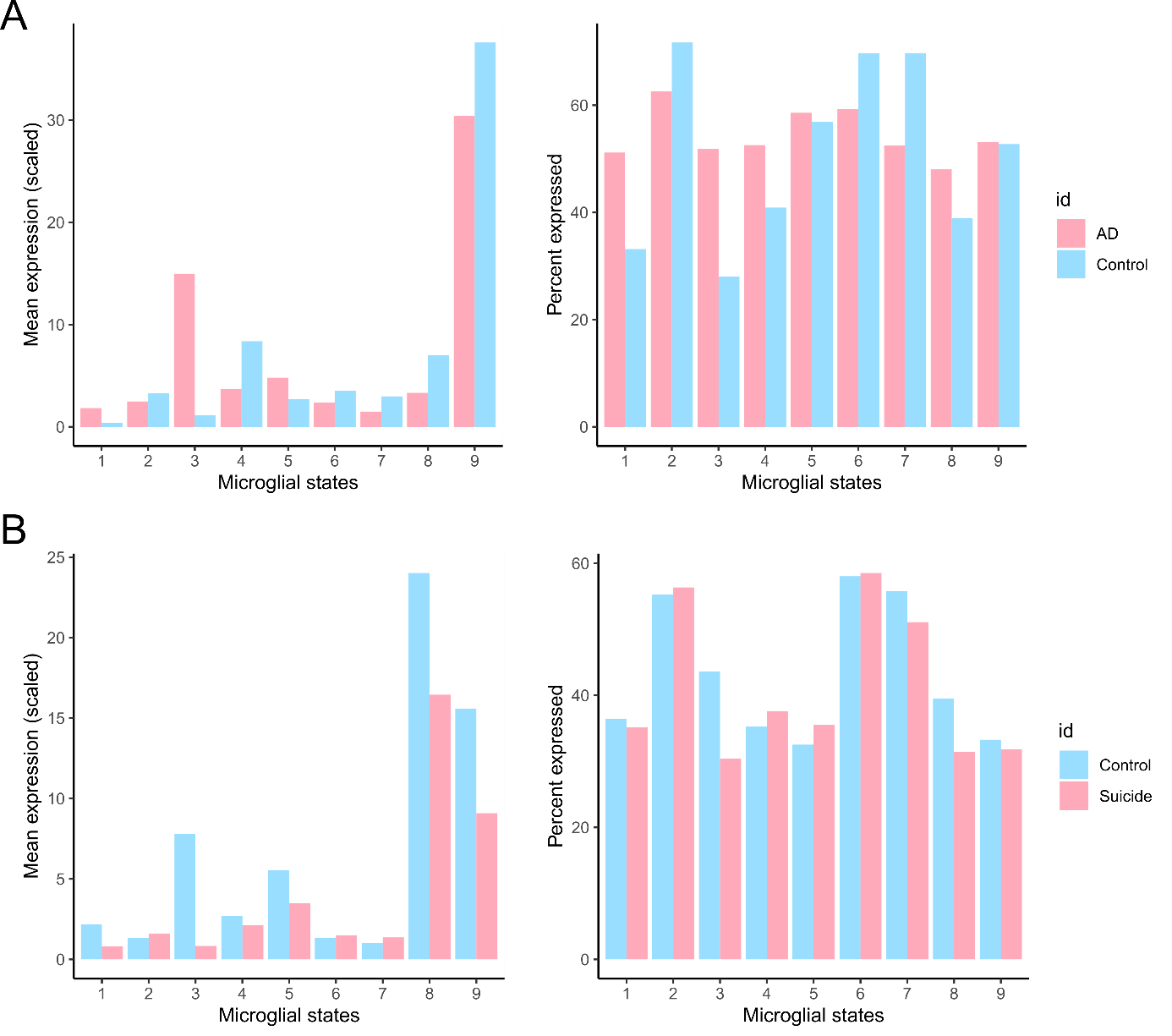

**Supplemental Figure S8. Bar plots of microglia states projection from Olah and colleagues healthy scRNA-seq data with hdWGCNA. They found clusters 5, 6 and 7 enriched for AD-associated genes, clusters 4 and 7 having altered expression of AD susceptibility genes and cluster 7 downregulated in AD compared to control individuals.** A) AD versus control cells. Microglial states were different between AD and control individuals, with the biggest changes in states 1 (homeostatic), 3 (cellular stress) and 7 (antigen presentation). B) MDD versus control cells. There were fewer differences between the microglial states, with the biggest changes in states 3 and 8 (ion transport). The percent expressed indicates how many cells expressed the gene signature.

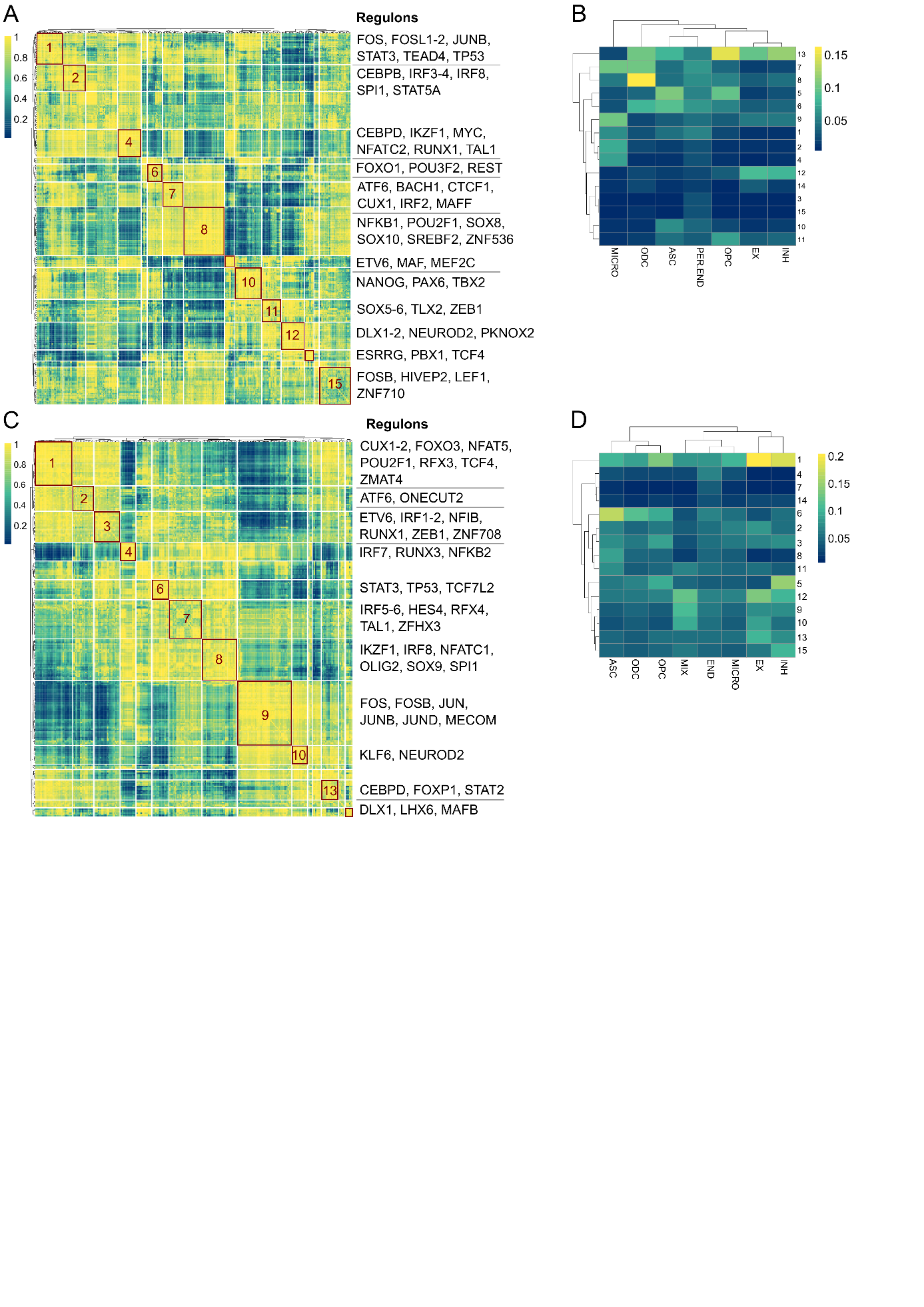
**Supplemental Figure S9. Co-regulation of regulons with the connection specificity index.** A, C) Connection specificity index (CSI) heatmap of AD (A) and MDD (C) with representative regulons in the clusters, depicted as the TF. There are three overarching large clusters, with fifteen smaller subclusters. Regulons with similar functions tend to cluster together. B, D) Mean AUC values of the regulon clusters per CSI module and cell type of AD (B) and MDD (D). Distinct clusters were active in distinct cell types, with some clusters more cell-type-specific than others.

**Supplemental Figure S10. ModuleViewer figures of all AD modules.** The ModuleViewer figures include the sample diagnosis, sex, functional enrichment terms, TF binding motif enrichment (RcisTarget), TF target gene enrichment (enrichR), and interactions from HumanNet (see legends). Highlighted module genes are TFs. See extra pdf file ModuleViewer_AD.pdf

**Supplemental Figure S11. ModuleViewer figures of all MDD modules.** The ModuleViewer figures include the sample diagnosis, sex, functional enrichment terms, TF binding motif enrichment (RcisTarget), TF target gene enrichment (enrichR), and interactions from HumanNet (see legends). Highlighted module genes are TFs. See extra pdf file ModuleViewer_MDD.pdf

### 3. **Supplemental Tables**

**Supplemental Table 1**. Overview of the bulk and single-cell datasets. The number of female and male individuals was similar in the AD dataset^5^, in contrast to the bulk MDD datasets^6,7^, where there were more men than women (26/106). The single-cell MDD dataset consisted of only male samples. The overall age of the control individuals was higher in the AD dataset (mean 80.90, sd 8.42) compared to the MDD datasets (mean 50.25, sd 12.50 and mean 43.50, sd 21.30). In the bulk AD dataset, other clinical scores were checked as well, such as plaque and tangle stage and other neuropathologies. This was not the case for the MDD datasets.

| Disorder | Accession | Data type | Brain region | Number of disease samples | | Number of control samples |
| --- | --- | --- | --- | --- | --- | --- |
| AD | GSE174367  syn22130832 | RNA-seq | Prefrontal cortex | | 47 | 48 |
| MDD | GSE101521 | RNA-seq | Dorsolateral prefrontal cortex | | 30 | 29 |
| MDD | GSE80655 | RNA-seq | Dorsolateral prefrontal cortex | | 23 | 24 |
| AD | GSE174367  syn22130834 | snRNA-seq | Prefrontal cortex | | 8 | 11 |
| MDD | GSE144136 | snRNA-seq | Prefrontal cortex | | 17 | 19 |

**Supplemental Table 2.** Top 100 (highest out-degree) regulating transcription factors of the consensus Alzheimer’s disease and depression networks. The transcription factors are ordered from high to low out-degree (range 1865 - 207). There are 34 shared transcription factors from the top 100.

| Alzheimer’s disease network | CSRNP3, MEF2C, PEG3, HLF, SATB1, PRDM2, THRB, ZNF25, NKRF, MYT1L, ZNF814, DACH2, ZNF483, STOX2, ZNF26, MEF2D, ZNF480, EGR3, ZBTB11, HIVEP2, CDC5L, ZSCAN30, ZNF777, HINFP, ZNF552, ZNF512, STAT4, BCL11A, POU5F2 TCF7, ZNF280B, ZNF621, ATMIN, ZNF587B, ZNF204P, ZBTB37, ZNF471, ZNF711, ZNF184, ATF2, MEF2A, CREB5, SOX10, PROX1, SOX8, ST18, NFIX, MYRF, ATF7, ZNF528, ZNF536, NFIA, LHX6, ZNF345, CTCF, BBX, TBR1, ARNT2, NEUROD6, ZBTB45, NFE2L3, STAT3, TCF12, RBPJ, NKX6-2, ZNF833P, POU2F1, ZNF317, PLSCR1, FOXJ3, TFEB, CREB3L2, ZBTB4, FOXN2, ZNF652, RFX4, ZBED3, OLIG1, SOX9, ZNF382, ZNF99, EPAS1, ZNF322, VEZF1, ELF1, ZNF217, KLF6, RUNX1, SP1, PPARA, ZGPAT, ZNF692, CREB3, ZBTB17, NR2C2, ZFP1, ZNF562, ZBED6, ZNF492, IKZF3 |
| --- | --- |
| Major depressive disorder network | THAP11, USF1, MLXIP, ATF2, MEF2C, CREB3, HEY1, HLF, CSRNP3, ZFHX2, ZNF576, ZBTB18, HSF1, NR2F6, ZNF787, ENO1, ZNF316, ZBTB22, SOX9, ATF7, ZNF25, HIVEP2, ATF4, ZNF711, PEG3, THRB, ST18, CIC, IRF3, MESP1, WIZ, RXRB, MAFG, MYRF, NPAS3, PPARA, SALL1, GLI3, RFX1, MEF2A, PLAGL2, SOX10, RXRA, NR2E1, ZHX2, MLXIPL, NKX6-2, DEAF1, KLF15, CC2D1A, ZNF532, SKI, TFEB, NR1H2, ZNF552, IKZF4, PAX6, SOX21, ZBTB11, ZNF518B, ZNF660, RFX4, NFKB2, ZNF536, POU5F2, TRPS1, KLF16, ZFP57, TEAD1, FOXO1, ZNF579, ATMIN, SREBF1, PLSCR1, RREB1, SOX2, SOX8, STOX1, SATB1, GLI4, VEZF1, USF2, ZFP1, ZNF204P, ZNF322P1, ZNF282, ZNF512B, MEIS3, RELA, ZNF444, CDC5L, MAZ, OLIG2, NKRF, ZNF783, ZNF865, HIVEP3, RARB, TCF12, HSF4 |
| Shared transcription factors | CSRNP3, MEF2C, PEG3, HLF, SATB1, THRB, ZNF25, NKRF, ZBTB11, HIVEP2, CDC5L, ZNF552, POU5F2, ATMIN, ZNF204P, ZNF711, ATF2, MEF2A, SOX10, SOX8, ST18, MYRF, ATF7, ZNF536, TCF12, NKX6-2, PLSCR1, TFEB, RFX4, SOX9, VEZF1, PPARA, CREB3, ZFP1 |

**Supplemental Table 3.** Overview of modules in the consensus networks prioritized through distinct criteria. AD: Alzheimer’s disease, MDD: major depressive disorder. Modules in bold are prioritized in several ways.

| **Module** | **Prioritization criteria** |
| --- | --- |
| AD modules 5, 7, 9, 27, **60**, 64, **66**, 77, 83, 140, 144 | Enriched in ‘Alzheimer’s disease’ term from KEGG (not used for prioritization) |
| MDD modules 30, 40, 91, 94, 117 | Enriched in ‘Alzheimer’s disease’ term from KEGG (not used for prioritization) |
| AD modules 4, 22, 36, 39, 51, 86, 93, 139, 141, **153** | Immune-system-related functional enrichment terms |
| MDD modules 4, **24**, **36**, 37, 56, 59, 61, 63, 81, **110**, **115**, 147 | Immune-system-related functional enrichment terms |
| AD modules 150, 85, 87, **153**, 40, 105, 130, **60**, **66**, 123, 26, 129, 21, 58, 35 | Differential expression between control and disease phenotypes |
| MDD modules 12, 44, 93, 48, 50, 3, **115**, **36**, **110**, 49, **24** | Differential expression between control and disease phenotypes |

**Supplemental Table 4.** Overview of SNPs from the GWAS Catalog intragenic or < 3kb upstream of genes (promoter region) in prioritized modules. Upstream is depicted in bold to visually indicate the difference with intragenic.

| SNP | Trait | Module | Associated gene | Upstream/intragenic |
| --- | --- | --- | --- | --- |
| rs1354106; rs7245846 | Alzheimer’s disease | 22 | CD33 | Intragenic |
| rs3826656; rs3865444 | Alzheimer’s disease | 22 | CD33 | **Upstream** |
| rs6891966 | Alzheimer’s disease | 22 | HAVCR2 | Intragenic |
| rs7142 | Alzheimer’s disease | 22 | HLA-DQA1 | Intragenic |
| rs7597763; rs36133610; rs6741388; rs36165346; rs7577360; rs35349669; rs10933431; rs10929072; | Alzheimer’s disease | 22 | INPP5D | Intragenic |
| rs7190997 | Alzheimer’s disease | 22 | ITGAX | Intragenic |
| rs192232892 | Alzheimer’s disease | 22 | TBXAS1 | Intragenic |
| rs286043; rs117627917; rs188255 | Alzheimer’s disease | 26 | DLG2 | Intragenic |
| rs11218343; rs4559697; rs74685827 | Alzheimer’s disease | 26 | SORL1 | Intragenic |
| rs4575098 | Alzheimer’s disease | 35 | ADAMTS4 | Intragenic |
| rs533143 | Alzheimer’s disease | 35 | FRMD5 | Intragenic |
| rs146168866 | Alzheimer’s disease | 35 | LAMP1 | Intragenic |
| rs3009868; rs7531270; rs12058296; rs2503185; rs4255357; rs2840677; rs35021135; rs12744291; rs6695557 | Alzheimer’s disease | 35 | PDE4B | Intragenic |
| rs34743976 | Alzheimer’s disease | 35 | PDE8A | Intragenic |
| rs9896800; rs616338 | Alzheimer’s disease | 39 | ABI3 | Intragenic |
| rs6584063 | Alzheimer’s disease | 39 | BLNK | Intragenic |
| rs12044355 | Alzheimer’s disease | 39 | DISC1 | Intragenic |
| rs7935829; rs7232 | Alzheimer’s disease | 39 | MS4A6A | Intragenic |
| rs610932 | Alzheimer’s disease | 39 | MS4A6A | **Upstream** |
| rs6985143 | Alzheimer’s disease | 39 | MSR1 | Intragenic |
| rs72824905; rs12446759; rs7342692; rs12444183 | Alzheimer’s disease | 39 | PLCG2 | Intragenic |
| rs778217695 | Alzheimer’s disease | 39 | WDFY4 | Intragenic |
| rs1969355 | Alzheimer’s disease | 60 | C12orf65 | Intragenic |
| rs6850546 | Alzheimer’s disease | 60 | ENOPH1 | Intragenic |
| rs113568679 | Alzheimer’s disease | 60 | MAPT | Intragenic |
| rs1065712 | Alzheimer’s disease | 66 | CTSB | Intragenic |
| rs785129 | Alzheimer’s disease | 66 | HS3ST5 | Intragenic |
| rs3824874 | Alzheimer’s disease | 66 | MTMR2 | Intragenic |
| rs117780815 | Alzheimer’s disease | 66 | NKAIN2 | Intragenic |
| rs10631987 | Alzheimer’s disease | 66 | RAC1 | Intragenic |
| rs62177277 | Alzheimer’s disease | 66 | SESTD1 | Intragenic |
| rs9269853 | Alzheimer’s disease | 141 | HLA-DRB1 | Intragenic |
| rs9614981 | Alzheimer’s disease | 153 | PHF21B | Intragenic |
| rs11969759 | Alzheimer’s disease | 153 | TNXB | Intragenic |
| rs74378198 | Unipolar depression | 36 | OSMR | Intragenic |
| rs116949162 | Unipolar depression | 37 | KCNQ | Intragenic |
| rs2042772 | Unipolar depression | 37 | LY75 | Intragenic |
| rs7288411 | Unipolar depression | 37 | NFAM1 | Intragenic |
| rs11835646 | Unipolar depression | 44 | ACSS3 | Intragenic |
| rs9589468 | Unipolar depression | 44 | GPC5 | Intragenic |
| rs913930 | Unipolar depression | 44 | TLR4 | Intragenic |
| rs138256067 | Unipolar depression | 63 | DOCK8 | Intragenic |
| rs201203751; rs76400352 | Unipolar depression | 63 | FYB | Intragenic |
| rs544475121 | Unipolar depression | 63 | LPAR6 | Intragenic |
| rs7534271 | Unipolar depression | 63 | TAL1 | Intragenic |
| rs7254215 | Unipolar depression | 81 | CLEC17A | Intragenic |
| rs201483250 | Unipolar depression | 81 | LILRA1 | Intragenic |
| rs1965523 | Unipolar depression | 81 | TFEC | Intragenic |
| rs162181; rs56390503 | Unipolar depression | 93 | FAT1 | Intragenic |
| rs115767724 | Unipolar depression | 93 | MCC | Intragenic |
| rs11140773 | Unipolar depression | 93 | NTRK2 | Intragenic |
| rs139747326 | Unipolar depression | 93 | PTAR1 | Intragenic |
| rs3758354 | Unipolar depression | 115 | ANXA1 | **Upstream** |

**Supplemental Table 5.** Top 100 (highest out-degree) regulating transcription factors of the Alzheimer and depression single-cell networks. The transcription factors are ordered from high to low out-degree (range 1826 - 44). There were 52 transcription factors shared.

| Alzheimer’s disease network | ETV6, FLI1, IKZF1, ETS2, ZEB1, SPI1, RUNX1, IRF8, TCF7L1, RFX2, STAT3, ELK3, REL, CEBPB, ELF1, RFX3, NFATC2, ERG, MAF, ETS1, EGR3, HLF, IRF5, STAT6, TAL1, FOS, PRDM1, RUNX3, CUX1, TBX15, GATA2, MXI1, IRF1, ATF2, FOXO1, TCF7L2, MEF2C, EGR1, FOSL2, JUN, BHLHE40, SREBF2, JUNB, CEBPD, EGR4, DLX6, RFX1, JUND, FOSB, DLX1, LHX6, RELB, NFIX, PRRX2, NFE2L1, ESRRA, NRF1, STAT1, FOXO3, PBX3, ZNF821, NFIC, BCL6, FOSL1, PRRX1, NKX6.2, EOMES, KLF10, KLF6, MAFF, IRF7, DLX5, NFKB2, ATF3, HIC1, DLX2, VSX1, HIF1A, JDP2, FOXF2, NR2C1, IRF4, LHX2, KLF16, FOXC2, HES5, NFIL3, IRF2, MSX2, NFKB1, SOX5, SREBF1, NR1D1, MAX, STAT5A, GLIS1, SOX8, SP3, TBX21, TEAD4 |
| --- | --- |
| Major depressive disorder networks | EGR1, ETS2, MAZ, JUND, EGR3, BHLHE40, RFX3, ZMAT4, YY1, FOSL2, ZBTB7A, FLI1, TCF7L2, JUN, HLF, JUNB, LHX6, KLF9, DLX1, NFE2L1, RUNX1, SREBF2, KLF16, FOXO1, ATF4, MAX, SOX2, CEBPB, FOXN3, IKZF1, TEF, NFAT5, NFIB, FOS, PRRX1, SOX10, USF2, KLF10, SOX8, ERG, GATA2, DBP, XBP1, HIF1A, LEF1, ATF2, EGR4, IRF8, NR2F2, ESRRA, SOX9, POU2F2, FOXO3, SOX4, KLF2, CREB3, KLF6, KLF13, SNAI3, IRF5, KLF4, ZEB1, DEAF1, ELF1, CEBPD, DLX2, FOXP2, TAL1, ETS1, HIVEP3, IRF7, MAFB, BCL6, CREBL2, THAP11, FOSB, DLX5, BACH1, SMAD1, CREM, ETV6, FOXJ3, DLX6, RXRA, SOX21, THRA, SOX15, NR2C1, POU2F1, TCF7L1, NKX2.2, EGR2, FOXP1, GZF1, PPARA, RFX2, ZNF281, PRDM16, VAX1, HIC1 |
| Shared transcription factors | ETV6, FLI1, IKZF1, ETS2, ZEB1, RUNX1, IRF8, TCF7L1, RFX2, CEBPB, ELF1, RFX3, ERG, ETS1, EGR3, HLF, IRF5, TAL1, FOS, GATA2, ATF2, FOXO1, TCF7L2, EGR1, FOSL2, JUN, BHLHE40, SREBF2, JUNB, CEBPD, EGR4, DLX6, JUND, FOSB, DLX1, LHX6, NFE2L1, ESRRA, FOXO3, BCL6, PRRX1, KLF10, KLF6, IRF7, DLX5, HIC1, DLX2, HIF1A, NR2C1, KLF16, MAX, SOX8 |

**Supplemental Table 6.** Overview of the highly active regulons (depicted as the TF, i.e. regulatory programs highly active, AUC scores from SCENIC) in different cell types of the brain in the single-cell networks of Alzheimer’s disease and major depressive disorder. Regulons in red were also inferred and active in the other network, in the same cell type.

| Cell type | Alzheimer’s disease | Major depressive disorder |
| --- | --- | --- |
| Excitatory neurons | AHR, HLF, MEF2C, ZEB1, ZMAT4 | FOXP1, NFAT5, PBX1, RFX3, TEAD4, ZMAT4, ZNF282, ZNF699 |
| Inhibitory neurons | HLF, MEF2C, PKNOX2, ZEB1, ZMAT4 | DLX1, DLX2, DLX5, FOXN3, HIVEP3, LHX6, MAFB, NR2F2, TCF4, ZMAT4, ZNF282 |
| Microglia | ELF1, ELK3, ETS2, ETV6, FLI1, IKZF1, IRF8, MAF, NFATC2, RUNX1, SPI1, STAT6 | FOXN3, IRF8, RUNX1 |
| Oligodendrocytes | MXI1, NFIX, SREBF2, ZNF536 | FOXN2, SOX10 |
| Oligodendrocyte precursor cells | PRRX1, PRRX2, SOX6, VSX1, ZEB1, ZNF227 | FOXN3, PBX3, PRRX1, SOX13, SOX4, ZEB1 |
| Astrocytes | FOXO1, RFX2, SOX5, TCF7L1, TCF7L2 | FOXO1, PAX6, RARG, RXRA, SOX2, SOX9, TCF7L2 |

**Supplemental table 7.** Overlap between population-level modules and single-cell regulons. AD Alzheimer’s disease, MDD major depressive disorder. The regulons of IKZF1 had the highest overlap for the AD-modules 22 and 39 and the MDD-module 24, but was also the largest regulon.

| Module | Number of module genes | Regulon | Number of regulon genes | Overlap |
| --- | --- | --- | --- | --- |
| 22 AD | 47 | IKZF1 AD | 1520 | 41 |
| 22 AD | 47 | IRF8 AD | 938 | 41 |
| 22 AD | 47 | NFATC2 AD | 541 | 16 |
| 22 AD | 47 | RUNX1 AD | 1016 | 30 |
| 22 AD | 47 | SPI1 AD | 1018 | 41 |
| 22 AD | 47 | TAL1 AD | 401 | 15 |
| 24 MDD | 52 | IKZF1 MDD | 227 | 20 |
| 24 MDD | 52 | IRF8 MDD | 142 | 10 |
| 24 MDD | 52 | RUNX1 MDD | 314 | 7 |
| 24 MDD | 52 | SPI1 MDD | 19 | 2 |
| 24 MDD | 52 | TAL1 MDD | 89 | 8 |
| 36 MDD | 35 | STAT3 MDD | 23 | 1 |
| 39 AD | 103 | IKZF1 AD | 1520 | 91 |
| 39 AD | 103 | IRF8 AD | 938 | 90 |
| 39 AD | 103 | NFATC2 AD | 541 | 51 |
| 39 AD | 103 | RUNX1 AD | 1016 | 79 |
| 39 AD | 103 | SPI1 AD | 1018 | 91 |
| 39 AD | 103 | TAL1 AD | 401 | 50 |
| 51 AD | 35 | KLF6 AD | 107 | 1 |
| 51 AD | 35 | FOSL2 AD | 274 | 6 |
| 110 MDD | 48 | FOS MDD | 191 | 20 |
| 110 MDD | 48 | JUNB MDD | 340 | 16 |
| 115 MDD | 38 | STAT3 MDD | 23 | 0 |
| 153 AD | 19 | CEBPD AD | 217 | 4 |
| 153 AD | 19 | MAFF AD | 106 | 2 |

**Supplemental table 8.** Overlap between Alzheimer (AD) and depression (MDD) regulons. The AD regulons were larger than the MDD regulons. Especially for the first five regulons, most of the MDD regulon genes were also in the AD regulon.

| Regulons | Number of genes AD | Number of genes MDD | Overlap |
| --- | --- | --- | --- |
| IKZF1 | 1520 | 227 | 211 |
| IRF8 | 938 | 142 | 123 |
| RUNX1 | 1016 | 314 | 138 |
| SPI1 | 1018 | 19 | 18 |
| TAL1 | 401 | 89 | 73 |
| CEBPD | 217 | 93 | 12 |
| FOS | 374 | 191 | 79 |
| JUNB | 229 | 340 | 65 |
| STAT3 | 679 | 23 | 7 |

**Supplemental Table 9.** Biased regulatory transcription factors and target genes from the differential edge analysis, with a cut-off of 90% gained edges for the respective disorder. The number of genes and transcription factors was higher for AD than MDD, probably because of the larger single-cell AD network.

|  | **Transcription factors** | **Target genes** |
| --- | --- | --- |
| AD-biased | ATF3, DDIT3, ELF1, ELK3, ELK4, ETV6, FLI1, IKZF1, IRF1, IRF8, MEF2D, MSX1, MSX2, NKX6.2, NR2C2, NRF1, PKNOX2, REL, RFX2, RUNX3, SPI1, STAT3, TAL1, TCF7L1, ZEB1, ZNF821 | COL5A1, ATG5, NFYC, KHDRBS3, IL1RAP, GRM4, LEPR, P4HA1, THUMPD2, SPECC1, ATF6, ZNF148, ADNP, MRC1, GPR183, ADM, RAB20, ARID5A, CXCR4, JAML, C3AR1, FAM126A, MAFF, HEY2, NKX6-2, KCTD12, CDKN1A, VEZF1, FKBP5, ZBTB7B, LMO2, TPRG1, HAMP, STAB1, ANKRD22, HRH2, S100A8, S100A9, GPR65, KCNG2, SIGLEC9, ZNF618, LYZ, SECTM1, SERPINA1, ATF5, SLAMF8, TSPAN9, MDGA1, FZD7, IL1B, GBP2, GEM, LCP2, KIAA0040, DPYSL3, BOC, ABCA1, APLNR, COL27A1, FAS, DRAM1, MYBPH, FCGR1A, PELI2, LDLR, CCR1, IRF1, TIFA, APOL6, LIMS1, PDLIM4, USP3, HERC5, ATOH8, IRF2, IRF7, STAT3, NFIC, IGFBP4, NPNT, PDLIM1, COL8A1, EPSTI1, ANXA2, BARX2, MSX2, SMAD6, KCNG3, ICAM1, SLC16A6, DLL4, TEAD4, SMAD1, NOS3, KCNJ8, KCTD15, SEMA3F, A4GALT, METTL21A, HSPA12B, PNP, RFX5, NFKB2, TGIF1, CCL3, TNF, ITGAX, ABCC3, HLA-DRA, TNFSF15, RGS18, TNFAIP2, ITGB2, CLEC17A, SPI1, ADAMTSL2, NOD2, TREM2, HOXC4, DYRK4, ATP10D, PLD4, SLC15A3, DOCK8, RBM47, MARCO, PTPRC, SCIN, NRP1, TBX15, SYK, LYN, MS4A6A, RIN3, LPL, PPP1R3C, KYNU, HCK, HMOX1, ALPK1, IL1A, MS4A4A, SLA, TFEC, PROS1, ZC3HAV1, PYGL, TGFBI, ANO6, LYVE1, CLEC5A, LY96, POU2F2, IER3, SYT17, TNFAIP3, TLR5, CD58, SIPA1L2, FCGR3A, APOC1, HK2, GPAT3, LUZP1, TUBA1C, TNFRSF10B, CD14, CCL20, TMEM182, FNDC3B, ABHD3, FGD4, IL21R, RELL1, LCP1, OGFRL1, COTL1, ALOX5AP, WNT5A, HLA-DRB1, ITGAL, SIGLEC10, CLEC1A, MYEOV, TNFSF13B, DHRS9, ENG, NR2E1, IL15RA, PARVG, ARHGDIB, STX11, DAPP1, OLFML2B, SMCO4, C1QA, FOSL1, ELK3, GLRX, CYTL1, TP53, SP140L, SMIM3, PLAUR, IKBKE, ZNF124, CD93, GALNT10, FAM89A, HELB, XIRP1, RELB, MAP3K14, NABP1, TP53I11, HLA-DRB5, NRIP1, CD180, RCOR1, CASP7, FGL2, STAT6, SLC25A37, PRKD3, CTSZ, CASP1, DENND2D, MYO1G, RNASE2, WNT6, BATF, NMI, RALB, TMEM119, REL, LSP1, ST14, APOBR, FCGRT, SNX20, MMP2, STAT1, FFAR2, IL4I1, TSPAN33, ADAM9, GPR84, PSTPIP1, FOLR2, CAPG, ZNF217, RAC2, LAIR2, C16orf54, FAM72B, SKI, PPARD, ZNF143, SNAI3, MNT, STAT5A, MAFG, ADAMTS9, SERPINE1, CP, CPA6, ADGRD1, KCNE4, RGS1, CXCL11, CPVL, ANXA3, ARHGAP26, LRRK2, PDPN, PLXDC1, PLEKHA2, GYPC, PELI1, ITGA5, PLAU, ATF3, HLX, IL10, S100A16, TMEM52B, TCAF2, MYC, MLXIP, GCNT1, PBX3, LACTB, VAV1, RHOH, CDCP1, TM6SF1, PAQR5, GBE1, GAS2L3, MOB1B, MGMT, KCNJ5, FMNL3, MALT1, FFAR4, RASGRP2, ZNF641, GTF2IRD1, MANBA, CD274, TNFRSF13C, FBXW8, COL8A2, ZNF362, KLF3, TFEB, GTF2I, CREB3L2, ZNF33A, CREBL2, NFATC3, PYGM, GADD45A, SAMD11, ZNF385D, GPRIN3, NAAA, TSPAN2, SLC9A2, FGD5, LAMA3, SCML4, ELFN1, WDR49, SLCO2A1, VWCE, SPHK1, CRYBB1, CCL4, ARL5C, SLC37A2, SEL1L3, LIPC, ABCB4, ST3GAL1, IL1RN, MSR1, INPP5D, ATP8B4, DOCK2, CD69, LY86, PIK3AP1, PIK3R5, RHBDF2, KCNK13, NFATC2, FAM149A, ALOX5, MYO1F, IL18, CD53, CTSC, CMTM7, RGS10, RUNX3, TRIM22, CD84, ADCY7, MX2, SP140, CYTH4, TRAF3IP3, LILRB1, CYTIP, IL10RA, PRAM1, TEC, WNT2B, TNFRSF11A, CEACAM21, CASP5, SAMD9L, GALM, TMEM71, HPSE, XRCC4, CLEC2D, MS4A14, PTGS1, HLA-DQB1, RAI14, PLCB2, TMC8, SRBD1, PTGER4, SP110, KCNMB1, OAS1, GPR132, LILRB4, SLFN11, BVES, ACSM5, B2M, IFIH1, TYROBP, SUCLG2, ERAP2, LRRC69, TMEM236, GREM1, TBX19, LILRB3, PDCD1LG2, LRRC25, METTL7A, SIGLEC7, LGALS9, BHLHE41, UNC93B1, CD37, HLA-DOA, SLFN12, NCF4, PTPN6, DDX58, LYL1, SPATA5, HAVCR2, RNASE6, CCDC88B, SAMD9, SLC17A9, PYCARD, REST, FOXK1, CUX1, ZNF267, ZNF562, ZNF100, PPARA, ZBED4, FCGBP, HS3ST4, SLCO2B1, ADGRL4, TBXAS1, ADAM28, LRRK1, SAMSN1, BLNK, PLA1A, CLCF1, CD86, CD74, IL4R, WDFY4, TNFAIP8, FGD2, PLEK, IL15, PLCG2, C12orf42, LAPTM5, CLEC2B, HLA-DQA1, RYR1, CC2D2B, SYT6, GCNT2, EDN3, HPGDS, MRC2, MEP1A, CTSS, GIMAP8, LAT2, ALPK3, PATL2, TRIM38, KCNQ1, MFSD2A, FGD3, IFNLR1, BRCA2, TNFSF8, GPATCH2, TMCO4, CLEC9A, HNMT, FCGR2A, SIGLEC8, HLA-DMB, PIK3CG, P2RY6, PADI1, TREML4, CLEC19A, SUSD3, RAB38, TMEM176B, LOXL3, KIF20B, VNN2, MILR1, TMIGD3, FGF20, NFATC1, RASAL3, NCF1, ACY3, APOL3, GNA15, CD300C, BTN3A2, ZNRF2, MPEG1, EHBP1L1, IGSF6, IRF5, GAL3ST4, CD33, TREML1, BCO2, RAB32, ELF1, ATP2A3, BIK, TOR4A, GIMAP5, PAX8, ZKSCAN4, ZNF787, MYT1, HAS2, RBM20, NXN, TNIP3, JAM2, ALPL, PRTG, NEDD4, RPH3AL, TES, BCL6B, GRAPL, ARHGEF15, YES1, APOL1, SHE, TMEM255B, TEAD2, PXDC1, PODN, MASP1, CA12, RXRG, ESRRB, BATF3, BEST3, GFRA1, SP100, PRF1, SLFN12L, RAB31, ITGA4, EOMES, ACAN, VWA2, GATA3, ICOS, SPSB4, ITK, IKZF3, MAP3K1, ICAM2, MYCT1, TC2N, PTPN22, SOX13, FCRL6, GZMB, CST7, S1PR4, GZMA, SH2D2A, FZD10, CD244, ROBO4, SLAMF1, TSPAN32, VSX1, VSIG2, PLEKHG2, ZAP70, TRAT1, TMEM163, CALCR, KBTBD12, TNFRSF11B, TG, SRGAP1, GBP4, RAB39A, TREM1, DDX60, C5AR2, TLR10, S100A4, SUCNR1, STEAP3, CEP135, CEACAM4, KCNJ1, ZYX, OAS2, TNNI2, GAPT, CEBPA, HAVCR1, FCGR3B, LILRA4, ASGR2, CD300LF, CCR5, VDR, USF1, PBX2, SPTLC3, ARHGAP20, VNN1, GBP1, HOXB3, MS4A6E, SPATA18, GBP3, OAS3, LILRA6, CD80, OLFML3, PARP12, ZNF438, CXCL16, SPTBN5, GNGT2, ROR2, GPRC5C, SORBS3, ZNF592, MSX1, SPX, TRIM47, PRDM6, CCDC80, TFAP2C, CYP11A1, SMAD5, POU3F2, EPB41L4A, GNLY, RAD51B, KIF19, ZIC1 |
| MDD-biased | ATF4, FOXN3, KLF4, MAFB, NKX2.1, NKX6.1, SMAD1, TBR1, TEF, ZBTB7A, ZMAT4 | EGR4, JPH1, SYT13, NPAS4, PRDM8, ZBTB11, NGEF, STMN2, ONECUT2, SERPINI1, YWHAH, NEUROD6, GPR22, TUBB2A, IFT57, MLF2, KCNMB2, NRIP3, SNAP25, CPE, ELAVL4, HLF, SCG2, SCN3B, THY1, GALNT9, NRN1, MAP1B, HMGCS1, PEG10, SLC7A14, PLK2, EEF1A2, FBXL16, CYP26B1, MAP9, KCNF1, MOAP1, GSKIP, RNF7, LRTM2, COX6A1, ZNF25, ZFHX4, SERPINE2, GFRA2, SLC6A17, CPLX2, TMEM130, GABRD, RASD1, NEFM, INA, NEFH, NECAP1, RNF6, TSPYL4, KCNA2, CCDC184, SCN4B, FXYD1, ATP1A3, UQCRB, C14orf132, TOMM20, NDUFA5, ZNF184, MAZ, VSNL1, TIMP3, ADAMTS3, NPTXR, HPCAL4, NBL1, PRRT1, RPH3A, SCG3, PODXL2, ALDOA, CDH9, ATP1B2, GAD2, NPPC, CKS2, MLLT11, CACNB3, FBXO2, TEF, EGR2, DLC1, SYNM, PHF24, CD36, SOX5 |

**Supplemental table 10**. TFs of which their target genes were enriched in DisGeNET disease-specific gene sets (‘Alzheimer’s disease’, ‘Alzheimer Disease, Late Onset’, ‘Familial Alzheimer Disease (FAD)’, ‘Alzheimer Disease, Early Onset’, ‘Major Depressive Disorder’, ‘Unipolar Depression’, ‘Mental Depression’, ‘Depressive disorder’, ‘Severe depression’, ‘Major depression, single episode’). There was quite some overlap between the TFs for the different networks, including the TFs in the core regulatory circuit (red).

|  | **Alzheimer’s disease** | **Depression** | **Overlap** |
| --- | --- | --- | --- |
| **Population-level networks** | ATF3, ATF4, BCL11A, CEBPD, CSRNP3, EPAS1, ETV6, FEZF2, FOS, FOSB, FOSL2, IRF8, IRF9, KLF6, MEF2C, MYC, MYT1L, NEUROD6, NFATC2, NFIA, NFIL3, NFIX, NKRF, PEG3, SATB1, SOX9, SP100, STAT4, STOX2, TFEC, TGIF1, THRB, ZBTB11, ZNF204P, ZNF483 | ATF2, ATF3, ATF4, CLOCK, CREM, CSRNP1, CSRNP2, DDIT3, DEAF1, EGR4, EHF, EMX2, ETV1, ETV3, FOSB, FOSL1, FOSL2, FOXC1, FOXP2, GLI2, GLI3, HLF, JUNB, KLF10, KLF11, MAFF, MEF2C, MEOX2, MESP1, MXD1, MXD3, MYT1L, NFATC4, NFE2L1, NFE2L2, NFIL3, NFIX, NPAS3, NPAS4, NR3C2, NR4A1, OTX1, OVOL2, PBX1, PBX2, PEG3, PLSCR1, PPARA, RFX2, RFX4, RUNX1, SALL2, SALL3, SATB2, SCAND2P, SNAI1, SNAI2, SOX2, SOX21, SOX9, STAT5A, STAT5B, STOX2, TEAD4, TGIF1, ZBTB18, ZNF184, ZNF331, ZNF333, ZNF436, ZNF438, ZNF518B, ZNF680, ZNF681, ZNF697, ZNF699, ZSCAN31, ZSCAN32 | ATF3, ATF4, FOSB, FOSL2, MEF2C, MYT1L, NFIL3, NFIX, PEG3, SOX9, STOX2, TGIF1 |
| **Single-cell networks** | ATF3, ATF4, BACH1, BCL6, BHLHE40, CEBPA, CEBPB, CEBPD, CREB1, DDIT3, EGR1, EGR2, EGR3, EGR4, ELF1, ELK3, ELK4, ERG, ETS1, ETS2, ETV6, FLI1, FOS, FOSB, FOSL1, FOSL2, FOXJ3, FOXO1, FOXO3, GATA2, HIF1A, HLF, IKZF1, IRF1, IRF4, IRF5, IRF7, IRF8, JUN, JUNB, JUND, KLF10, KLF6, MAF, MAFB, MAFF, MAX, MEF2D, MXI1, NFATC2, NFE2L1, NFIC, NFIL3, NFIX, NFKB2, PRDM1, PRRX1, REL, RELB, RUNX1, RUNX3, SOX18, SP1, SP3, SPI1, SREBF1, STAT1, STAT3, STAT5A, STAT6, TAL1, TBX15, TCF7L1, TCF7L2, TEAD4, TP53, VDR, XBP1, YY1, ZBTB7A, ZEB1 | BACH1, BBX, BCL6, BHLHE40, CEBPD, CREM, DLX1, DLX2, DLX5, DLX6, EGR1, EGR2, EGR3, EGR4, ELF1, ERG, ESRRA, ETS1, FLI1, FOS, FOSB, FOSL2, FOXN3, FOXO1, FOXO3, FOXP1, FOXP2, HIF1A, IRF5, IRF7, IRF8, JDP2, JUN, JUNB, JUND, KLF10, KLF16, KLF6, KLF9, LHX2, LHX6, MAFB, MAZ, MSX1, NFE2L1, NFIB, NFIL3, NFYB, NKX2.2, NKX6.3, NR2F2, PAX6, POU3F2, POU3F3, PRRX1, RUNX1, RXRA, SOX10, SOX11, SOX13, SOX2, SOX21, SOX9, SREBF2, TCF7L1, TCF7L2, VAX1, ZBTB7A, ZMAT4, ZNF331 | BACH1, BHLHE40, CEBPD, EGR1, EGR2, EGR3, EGR4, ERG, ETS1, FLI1, FOS, FOSB, FOSL2, FOXO1, FOXO3, HIF1A, IRF5, IRF7, IRF8, JUN, JUNB, JUND, KLF10, KLF6, MAFB, NFE2L1, NFIL3, PRRX1, RUNX1, TCF7L1, TCF7L2, ZBTB7A |
| **Overlap** | ATF3, ATF4, CEBPD, ETV6, FOS, FOSB, FOSL2, IRF8, KLF6, NFATC2, NFIL3, NFIX | CREM, EGR4, FOSB, FOSL2, FOXP2, JUNB, KLF10, NFE2L1, NFIL3, RUNX1, SOX2, SOX21, SOX9, ZNF331 | FOSB, FOSL2, NFIL3 |

**Supplemental table 11. Characteristics of population-level modules of AD and MDD.** Excel table.
