## Supplemental Figure 10 for "Comparative gene regulatory network analysis in Alzheimer’s disease and major depressive disorder identifies shared core regulatory circuits"

Module 1

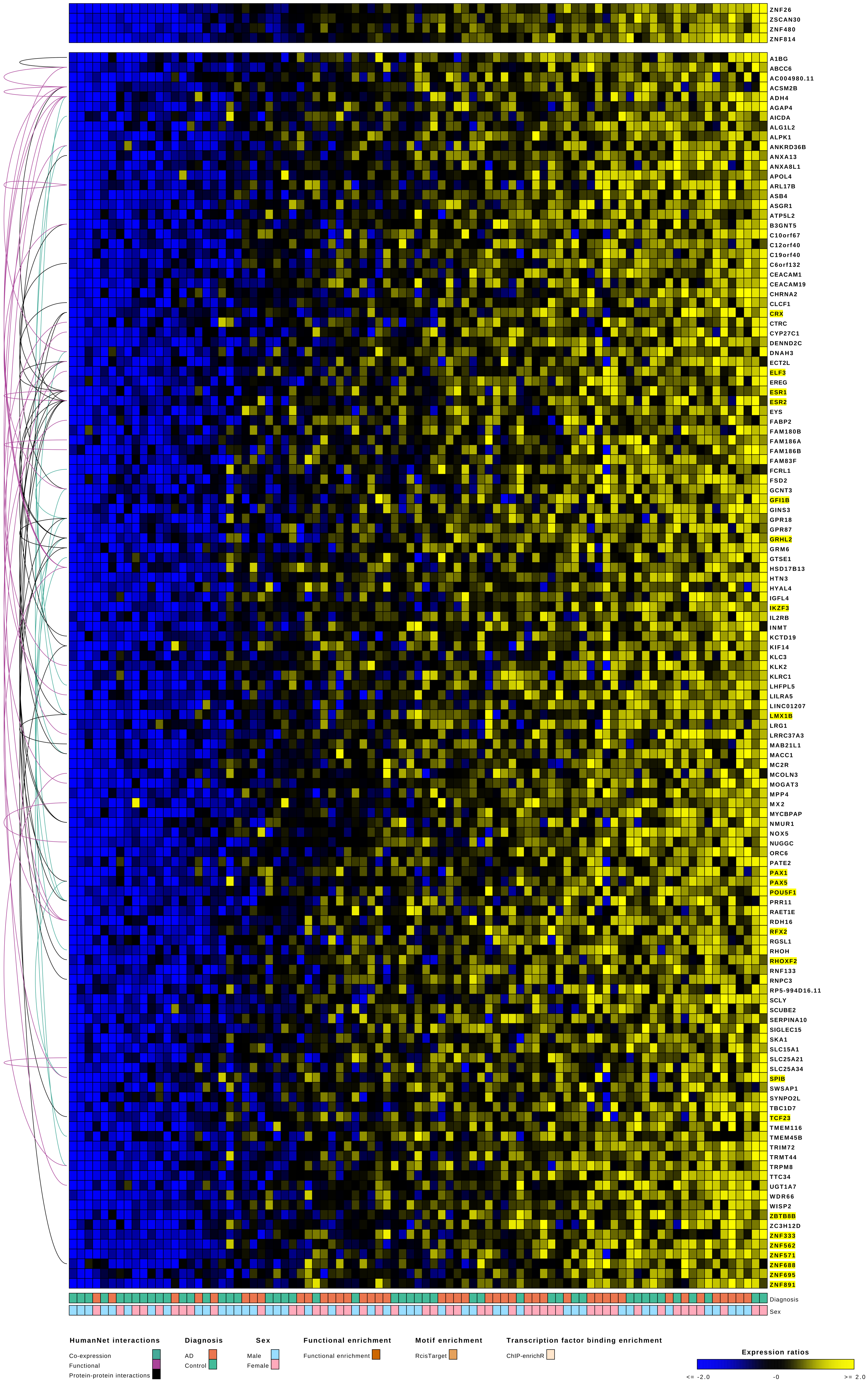
ZNF26
ZSCAN30
ZNF480
ZNF814
A1BG
ABCC6
AC004980.11
ACSM2B
ADH4
AGAP4
AICDA
ALG1L2
ALPK1
ANKRD36B
ANXA13
ANXA8L1
APOL4
ARL17B
ASB4
ASGR1
ATP5L2
B3GNT5
C10orf67
C12orf40
C19orf40
C6orf132
CEACAM1
CEACAM19
CHRNA2
CLCF1
CRX
CTRC
CYP27C1
DENND2C
DNAH3
ECT2L
ELF3
EREG
ESR1
ESR2
EYS
FABP2
FAM180B
FAM186A
FAM186B
FAM83F
FCRL1
FSD2
GCNT3
GFI1B
GINS3
GPR18
GPR87
GRHL2
GRM6
GTSE1
HSD17B13
HTN3
HYAL4
IGFL4
IKZF3
IL2RB
INMT
KCTD19
KIF14
KLC3
KLK2
KLRC1
LHFPL5
LILRA5
LINC01207
LMX1B
LRG1
LRRC37A3
MAB21L1
MACC1
MC2R
MCOLN3
MOGAT3
MPP4
MX2
MYCBPAP
NMUR1
NOX5
NUGGC
ORC6
PATE2
PAX1
PAX5
POU5F1
PRR11
RAET1E
RDH16
RFX2
RGSL1
RHOH
RHOXF2
RNF133
RNPC3
RP5-994D16.11
SCLY
SCUBE2
SERPINA10
SIGLEC15
SKA1
SLC15A1
SLC25A21
SLC25A34
SPIB
SWSAP1
SYNPO2L
TBC1D7
TCF23
TMEM116
TMEM45B
TRIM72
TRMT44
TRPM8
TTC34
UGT1A7
WDR66
WISP2
ZBTB8B
ZC3H12D
ZNF333
ZNF562
ZNF571
ZNF688
ZNF695
ZNF891
Diagnosis
Sex
HumanNet interactions
Co-expression
Functional
Protein-protein interactions
Diagnosis
AD
Control
Sex
Male
Female
Functional enrichment
Functional enrichment
Motif enrichment
RcisTarget
Transcription factor binding enrichment
ChIP-enrichR
Expression ratios
<= -2.0
-0
>= 2.0

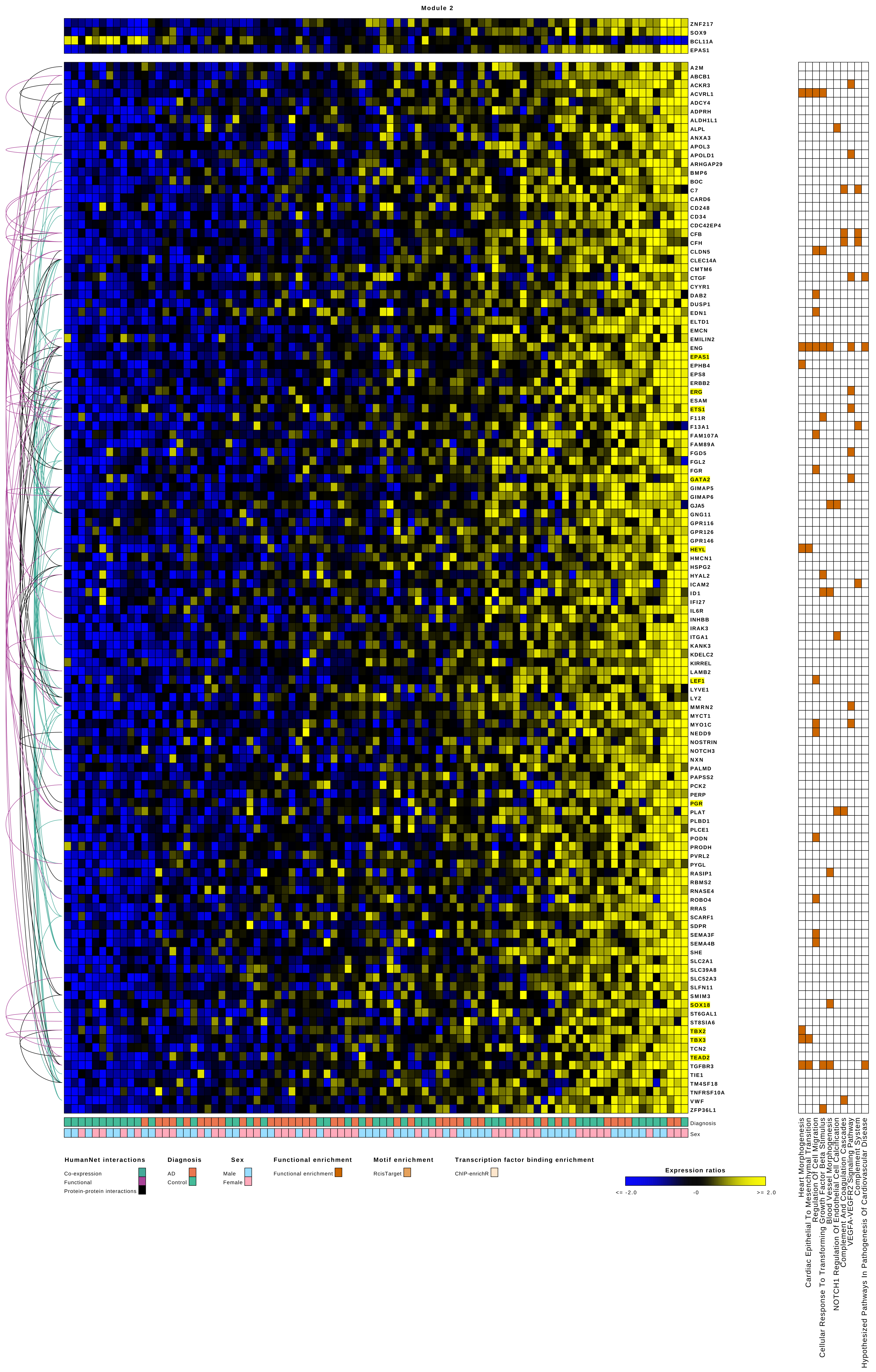

Module 2
ZNF217
SOX9
BCL11A
EPAS1
A2M
ABCB1
ACKR3
ACVRL1
ADCY4
ADPRH
ALDH1L1
ALPL
ANXA3
APOL3
APOLD1
ARHGAP29
BMP6
BOC
C7
CARD6
CD248
CD34
CDC42EP4
CFB
CFH
CLDN5
CLEC14A
CMTM6
CTGF
CYYR1
DAB2
DUSP1
EDN1
ELTD1
EMCN
EMILIN2
ENG
EPAS1
EPHB4
EPS8
ERBB2
ERG
ESAM
ETS1
F11R
F13A1
FAM107A
FAM89A
FGD5
FGL2
FGR
GATA2
GIMAP5
GIMAP6
GJA5
GNG11
GPR116
GPR126
GPR146
HEYL
HMCN1
HSPG2
HYAL2
ICAM2
ID1
IFI27
IL6R
INHBB
IRAK3
ITGA1
KANK3
KDELC2
KIRREL
LAMB2
LEF1
LYVE1
LYZ
MMRN2
MYCT1
MYO1C
NEDD9
NOSTRIN
NOTCH3
NXN
PALMD
PAPSS2
PCK2
PERP
PGR
PLAT
PLBD1
PLCE1
PODN
PRODH
PVRL2
PYGL
RASIP1
RBMS2
RNASE4
ROBO4
RRAS
SCARF1
SDPR
SEMA3F
SEMA4B
SHE
SLC2A1
SLC39A8
SLC52A3
SLFN11
SMIM3
SOX18
ST6GAL1
ST8SIA6
TBX2
TBX3
TCN2
TEAD2
TGFBR3
TIE1
TM4SF18
TNFRSF10A
VWF
ZFP36L1
Diagnosis
Sex
Heart Morphogenesis
Cardiac Epithelial To Mesenchymal Transition
Regulation Of Cell Migration
Cellular Response To Transforming Growth Factor Beta Stimulus
Blood Vessel Morphogenesis
NOTCH1 Regulation Of Endothelial Cell Calcification
Complement And Coagulation Cascades
VEGFA-VEGFR2 Signaling Pathway
Complement System
Hypothesized Pathways In Pathogenesis Of Cardiovascular Disease
HumanNet interactions
Co-expression
Functional
Protein-protein interactions
Diagnosis
AD
Control
Sex
Male
Female
Functional enrichment
Functional enrichment
Motif enrichment
RcisTarget
Transcription factor binding enrichment
ChIP-enrichR
Expression ratios
<= -2.0
-0
>= 2.0

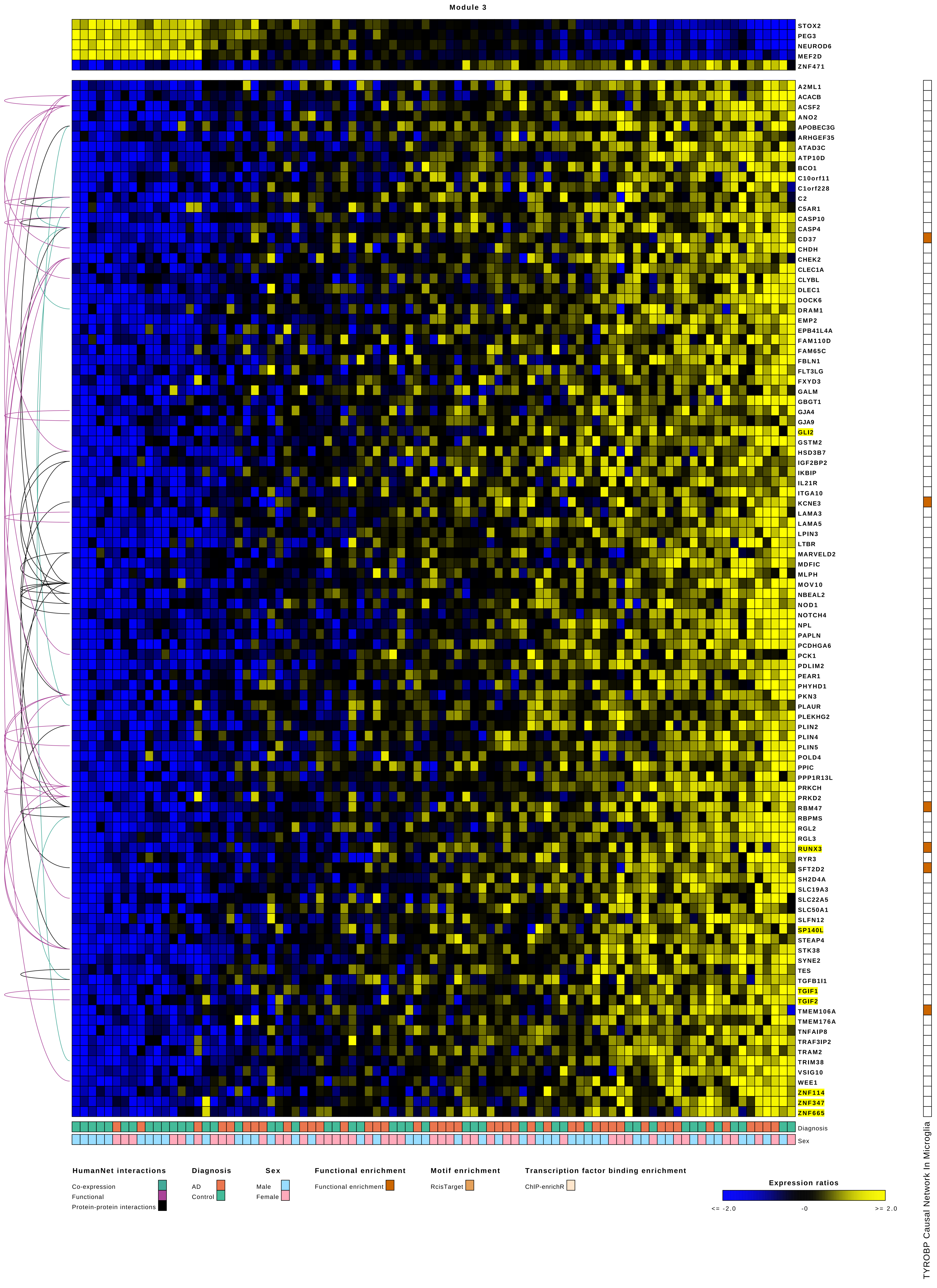
Module 3
STOX2
PEG3
NEUROD6
MEF2D
ZNF471
A2ML1
ACACB
ACSF2
ANO2
APOBEC3G
ARHGEF35
ATAD3C
ATP10D
BCO1
C10orf11
C1orf228
C2
C5AR1
CASP10
CASP4
CD37
CHDH
CHEK2
CLEC1A
CLYBL
DLEC1
DOCK6
DRAM1
EMP2
EPB41L4A
FAM110D
FAM65C
FBLN1
FLT3LG
FXYD3
GALM
GBGT1
GJA4
GJA9
GLI2
GSTM2
HSD3B7
IGF2BP2
IKBIP
IL21R
ITGA10
KCNE3
LAMA3
LAMA5
LPIN3
LTBR
MARVELD2
MDFIC
MLPH
MOV10
NBEAL2
NOD1
NOTCH4
NPL
PAPLN
PCDHGA6
PCK1
PDLIM2
PEAR1
PHYHD1
PKN3
PLAUR
PLEKHG2
PLIN2
PLIN4
PLIN5
POLD4
PPIC
PPP1R13L
PRKCH
PRKD2
RBM47
RBPMS
RGL2
RGL3
RUNX3
RYR3
SFT2D2
SH2D4A
SLC19A3
SLC22A5
SLC50A1
SLFN12
SP140L
STEAP4
STK38
SYNE2
TES
TGFB1I1
TGIF1
TGIF2
TMEM106A
TMEM176A
TNFAIP8
TRAF3IP2
TRAM2
TRIM38
VSIG10
WEE1
ZNF114
ZNF347
ZNF665
Diagnosis
Sex
HumanNet interactions
Co-expression
Functional
Protein-protein interactions
Diagnosis
AD
Control
Sex
Male
Female
Functional enrichment
Functional enrichment
Motif enrichment
RcisTarget
Transcription factor binding enrichment
ChIP-enrichR
Expression ratios
<= -2.0
-0
>= 2.0
TYROBP Causal Network In Microglia

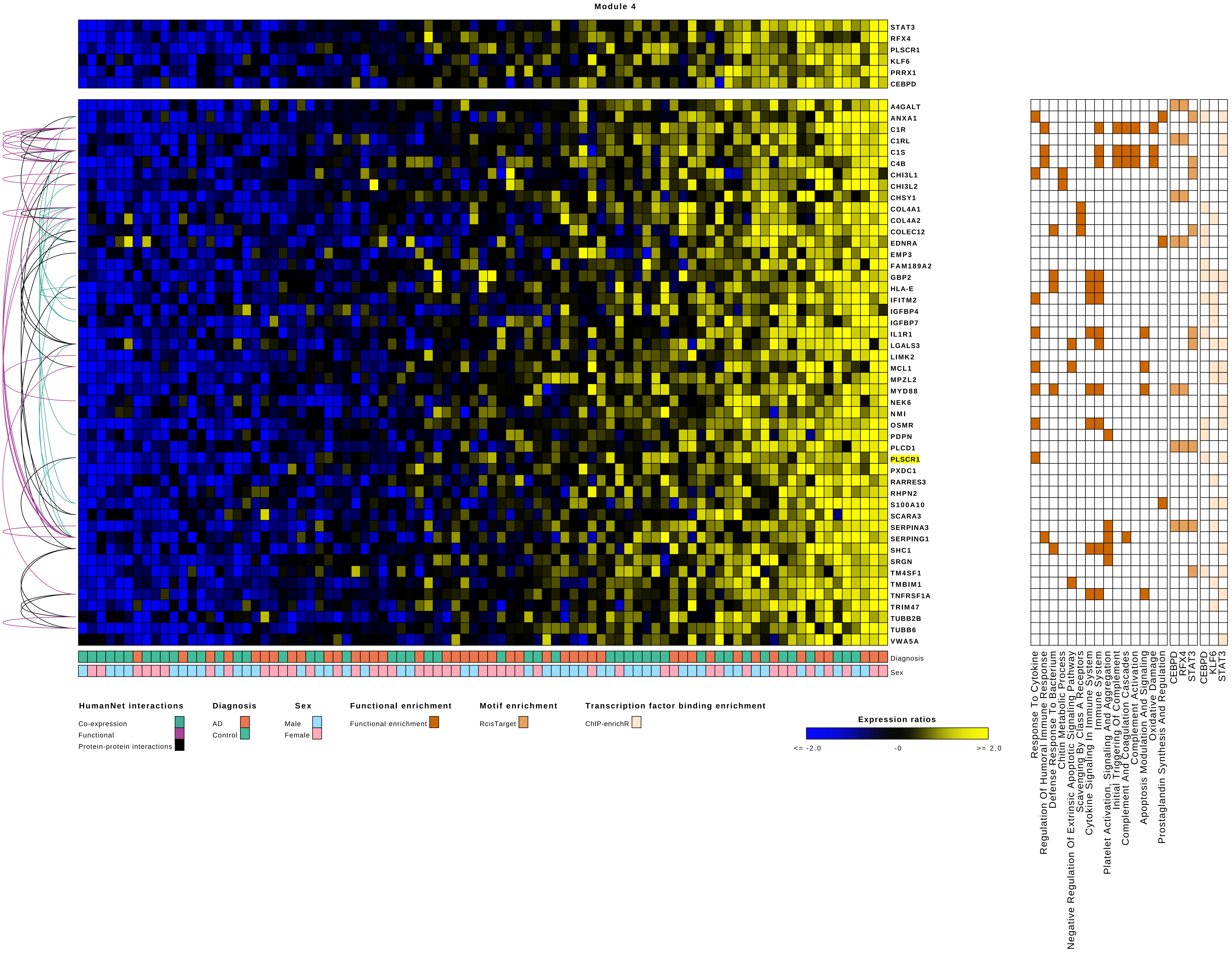
Module 4
STAT3
RFX4
PLSCR1
KLF6
PRRX1
CEBPD
A4GALT
ANXA1
C1R
C1RL
C1S
C4B
CHI3L1
CHI3L2
CHSY1
COL4A1
COL4A2
COLEC12
EDNRA
EMP3
FAM189A2
GBP2
HLA-E
IFITM2
IGFBP4
IGFBP7
IL1R1
LGALS3
LIMK2
MCL1
MPZL2
MYD88
NEK6
NMI
OSMR
PDPN
PLCD1
PLSCR1
PXDC1
RARRES3
RHPN2
S100A10
SCARA3
SERPINA3
SERPING1
SHC1
SRGN
TM4SF1
TMBIM1
TNFRSF1A
TRIM47
TUBB2B
TUBB6
VWA5A
Diagnosis
Sex
HumanNet interactions
Co-expression
Functional
Protein-protein interactions
Diagnosis
AD
Control
Sex
Male
Female
Functional enrichment
Functional enrichment
Motif enrichment
RcisTarget
Transcription factor binding enrichment
ChIP-enrichR
Expression ratios
<= -2.0
-0
>= 2.0
Response To Cytokine
Regulation Of Humoral Immune Response
Defense Response To Bacterium
Chitin Metabolic Process
Negative Regulation Of Extrinsic Apoptotic Signaling Pathway
Scavenging By Class A Receptors
Cytokine Signaling In Immune System
Immune System
Platelet Activation, Signaling And Aggregation
Initial Triggering Of Complement
Complement And Coagulation Cascades
Complement Activation
Apoptosis Modulation And Signaling
Oxidative Damage
Prostaglandin Synthesis And Regulation
CEBPD
RFX4
STAT3
CEBPD
KLF6
STAT3

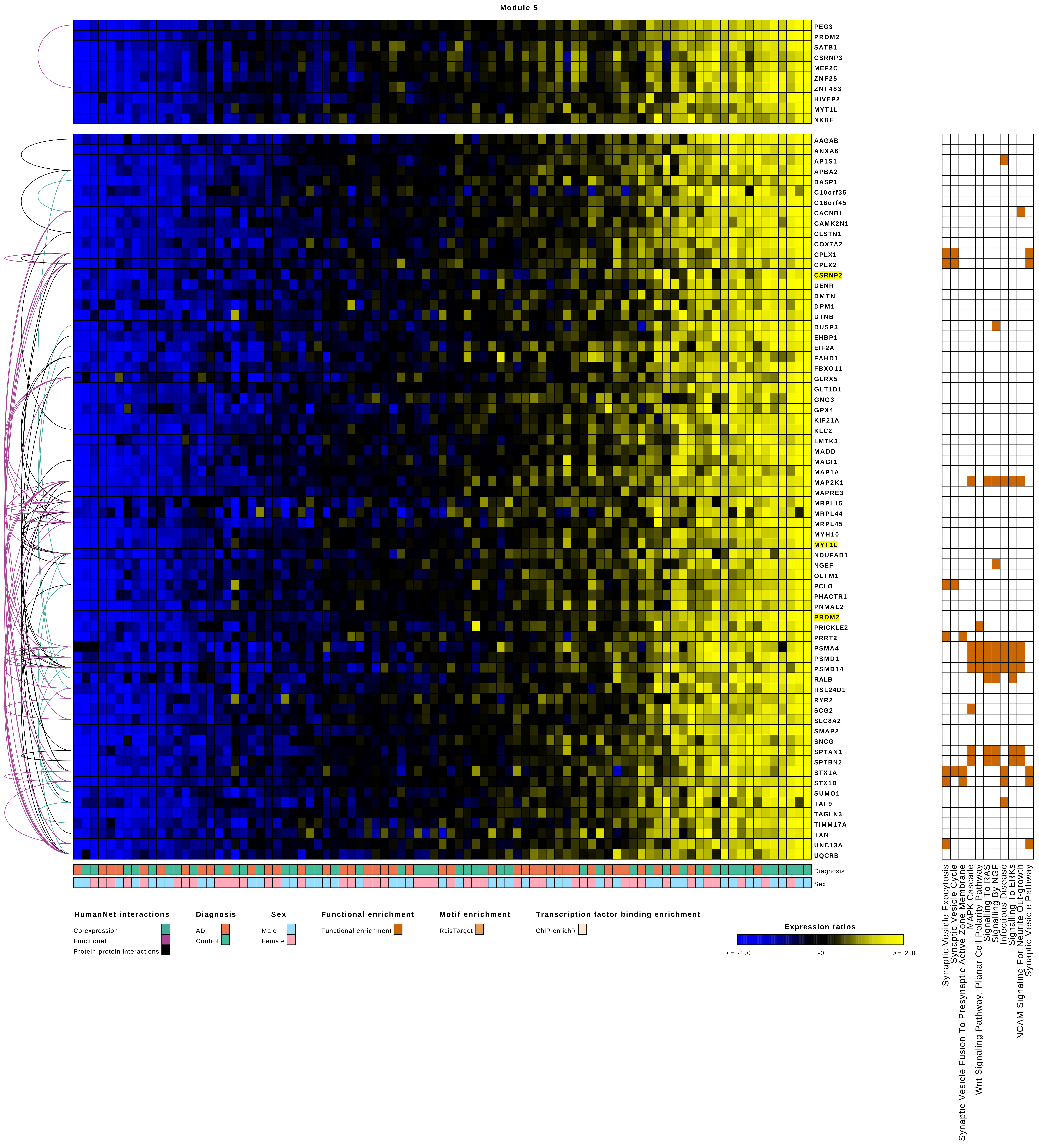
Module 5
PEG3
PRDM2
SATB1
CSRNP3
MEF2C
ZNF25
ZNF483
HIVEP2
MYT1L
NKRF
AAGAB
ANXA6
AP1S1
APBA2
BASP1
C10orf35
C16orf45
CACNB1
CAMK2N1
CLSTN1
COX7A2
CPLX1
CPLX2
CSRNP2
DENR
DMTN
DPM1
DTNB
DUSP3
EHBP1
EIF2A
FAHD1
FBXO11
GLRX5
GLT1D1
GNG3
GPX4
KIF21A
KLC2
LMTK3
MADD
MAGI1
MAP1A
MAP2K1
MAPRE3
MRPL15
MRPL44
MRPL45
MYH10
MYT1L
NDUFAB1
NGEF
OLFM1
PCLO
PHACTR1
PNMAL2
PRDM2
PRICKLE2
PRRT2
PSMA4
PSMD1
PSMD14
RALB
RSL24D1
RYR2
SCG2
SLC8A2
SMAP2
SNCG
SPTAN1
SPTBN2
STX1A
STX1B
SUMO1
TAF9
TAGLN3
TIMM17A
TXN
UNC13A
UQCRB
Diagnosis
Sex
Synaptic Vesicle Exocytosis
Synaptic Vesicle Cycle
Synaptic Vesicle Fusion To Presynaptic Active Zone Membrane
MAPK Cascade
Wnt Signaling Pathway, Planar Cell Polarity Pathway
Signalling To RAS
Signalling By NGF
Infectious Disease
Signalling To ERKs
NCAM Signaling For Neurite Out-growth
Synaptic Vesicle Pathway
HumanNet interactions
Co-expression
Functional
Protein-protein interactions
Diagnosis
AD
Control
Sex
Male
Female
Functional enrichment
Functional enrichment
Motif enrichment
RcisTarget
Transcription factor binding enrichment
ChIP-enrichR
Expression ratios
<= -2.0
-0
>= 2.0

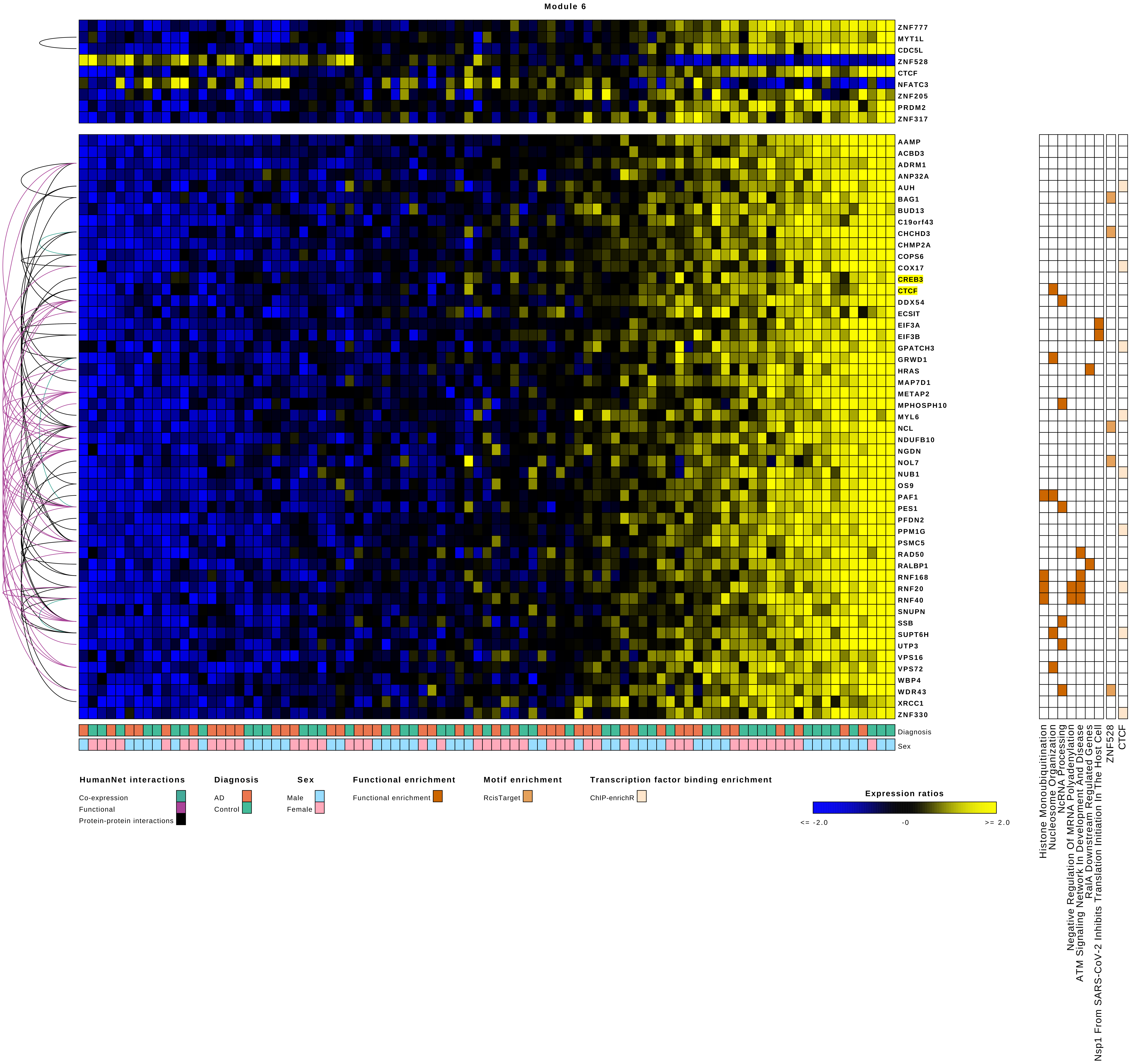
Module 6
ZNF777
MYT1L
CDC5L
ZNF528
CTCF
NFATC3
ZNF205
PRDM2
ZNF317
AAMP
ACBD3
ADRM1
ANP32A
AUH
BAG1
BUD13
C19orf43
CHCHD3
CHMP2A
COPS6
COX17
CREB3
CTCF
DDX54
ECSIT
EIF3A
EIF3B
GPATCH3
GRWD1
HRAS
MAP7D1
METAP2
MPHOSPH10
MYL6
NCL
NDUFB10
NGDN
NOL7
NUB1
OS9
PAF1
PES1
PFDN2
PPM1G
PSMC5
RAD50
RALBP1
RNF168
RNF20
RNF40
SNUPN
SSB
SUPT6H
UTP3
VPS16
VPS72
WBP4
WDR43
XRCC1
ZNF330
Diagnosis
Sex
Histone Monoubiquitination
Nucleosome Organization
NcRNA Processing
Negative Regulation Of MRNA Polyadenylation
ATM Signaling Network In Development And Disease
RalA Downstream Regulated Genes
Nsp1 From SARS-CoV-2 Inhibits Translation Initiation In The Host Cell
ZNF528
CTCF
HumanNet interactions
Co-expression
Functional
Protein-protein interactions
Diagnosis
AD
Control
Sex
Male
Female
Functional enrichment
Functional enrichment
Motif enrichment
RcisTarget
Transcription factor binding enrichment
ChIP-enrichR
Expression ratios
<= -2.0
-0
>= 2.0

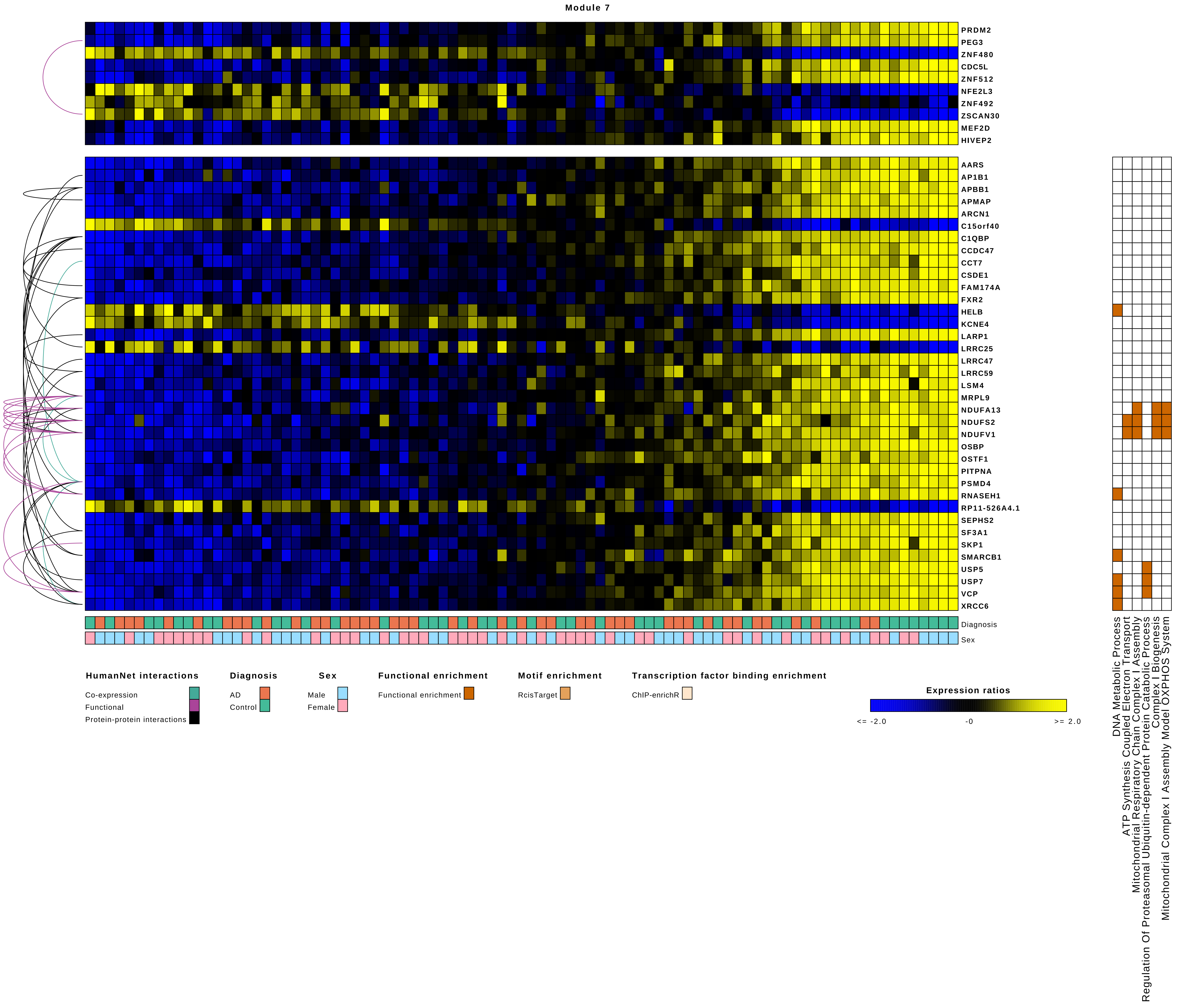
Module 7
PRDM2
PEG3
ZNF480
CDC5L
ZNF512
NFE2L3
ZNF492
ZSCAN30
MEF2D
HIVEP2
AARS
AP1B1
APBB1
APMAP
ARCN1
C15orf40
C1QBP
CCDC47
CCT7
CSDE1
FAM174A
FXR2
HELB
KCNE4
LARP1
LRRC25
LRRC47
LRRC59
LSM4
MRPL9
NDUFA13
NDUFS2
NDUFV1
OSBP
OSTF1
PITPNA
PSMD4
RNASEH1
RP11-526A4.1
SEPHS2
SF3A1
SKP1
SMARCB1
USP5
USP7
VCP
XRCC6
Diagnosis
Sex
DNA Metabolic Process
ATP Synthesis Coupled Electron Transport
Mitochondrial Respiratory Chain Complex I Assembly
Regulation Of Proteasomal Ubiquitin-dependent Protein Catabolic Process
Complex I Biogenesis
Mitochondrial Complex I Assembly Model OXPHOS System
HumanNet interactions
Co-expression
Functional
Protein-protein interactions
Diagnosis
AD
Control
Sex
Male
Female
Functional enrichment
Functional enrichment
Motif enrichment
RcisTarget
Transcription factor binding enrichment
ChIP-enrichR
Expression ratios
<= -2.0
-0
>= 2.0

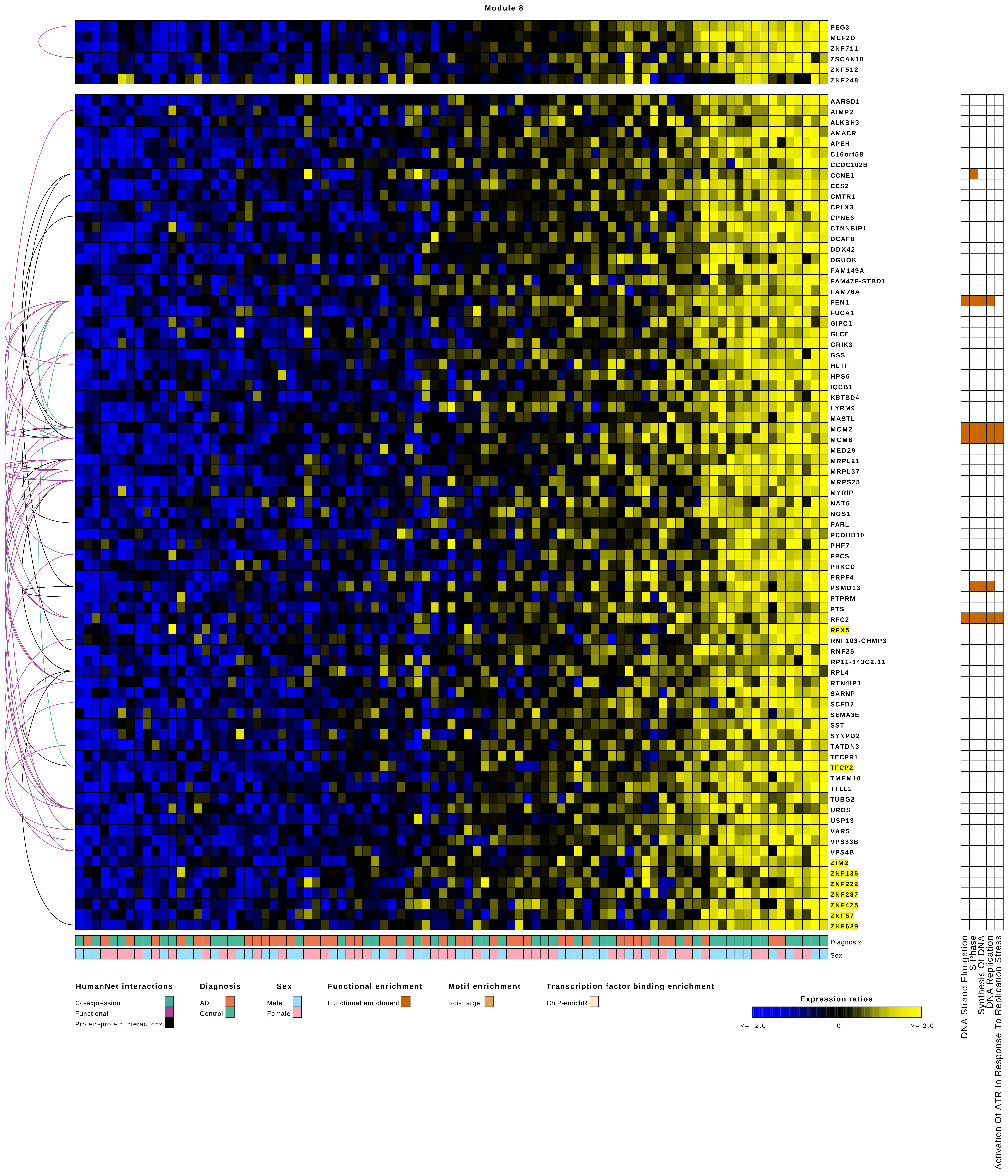
Module 8
PEG3
MEF2D
ZNF711
ZSCAN18
ZNF512
ZNF248
AARSD1
AIMP2
ALKBH3
AMACR
APEH
C16orf58
CCDC102B
CCNE1
CES2
CMTR1
CPLX3
CPNE6
CTNNBIP1
DCAF8
DDX42
DGUOK
FAM149A
FAM47E-STBD1
FAM76A
FEN1
FUCA1
GIPC1
GLCE
GRIK3
GSS
HLTF
HPS6
IQCB1
KBTBD4
LYRM9
MASTL
MCM2
MCM6
MED29
MRPL21
MRPL37
MRPS25
MYRIP
NAT6
NOS1
PARL
PCDHB10
PHF7
PPCS
PRKCD
PRPF4
PSMD13
PTPRM
PTS
RFC2
RFX5
RNF103-CHMP3
RNF25
RP11-343C2.11
RPL4
RTN4IP1
SARNP
SCFD2
SEMA3E
SST
SYNPO2
TATDN3
TECPR1
TFCP2
TMEM18
TTLL1
TUBG2
UROS
USP13
VARS
VPS33B
VPS4B
ZIM2
ZNF136
ZNF222
ZNF287
ZNF425
ZNF57
ZNF629
Diagnosis
Sex
DNA Strand Elongation
S Phase
Synthesis Of DNA
DNA Replication
Activation Of ATR In Response To Replication Stress
HumanNet interactions
Co-expression
Functional
Protein-protein interactions
Diagnosis
AD
Control
Sex
Male
Female
Functional enrichment
Functional enrichment
Motif enrichment
RcisTarget
Transcription factor binding enrichment
ChIP-enrichR
Expression ratios
<= -2.0
-0
>= 2.0

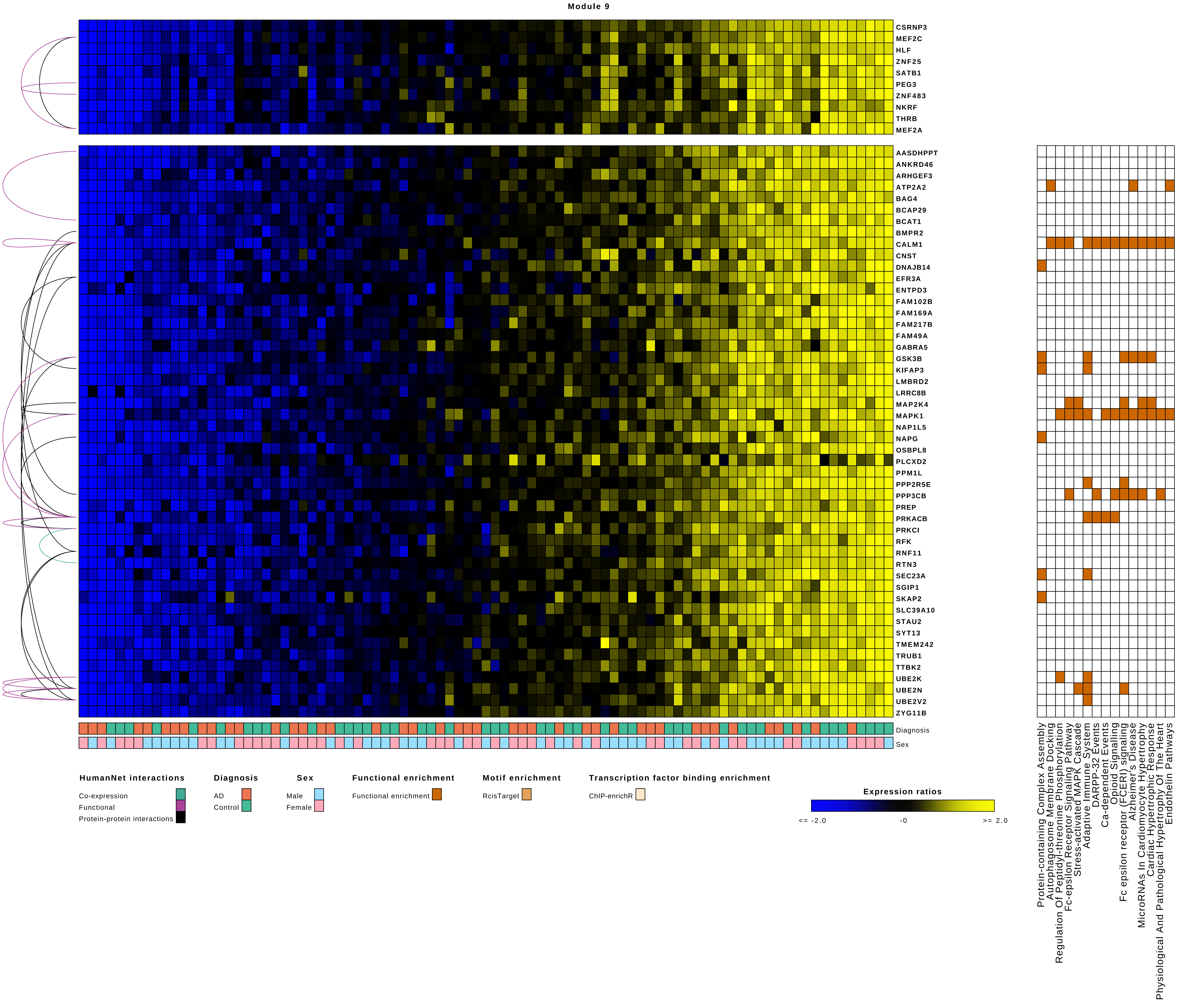
Module 9
CSRNP3
MEF2C
HLF
ZNF25
SATB1
PEG3
ZNF483
NKRF
THRB
MEF2A
AASDHPPT
ANKRD46
ARHGEF3
ATP2A2
BAG4
BCAP29
BCAT1
BMPR2
CALM1
CNST
DNAJB14
EFR3A
ENTPD3
FAM102B
FAM169A
FAM217B
FAM49A
GABRA5
GSK3B
KIFAP3
LMBRD2
LRRC8B
MAP2K4
MAPK1
NAP1L5
NAPG
OSBPL8
PLCXD2
PPM1L
PPP2R5E
PPP3CB
PREP
PRKACB
PRKCI
RFK
RNF11
RTN3
SEC23A
SGIP1
SKAP2
SLC39A10
STAU2
SYT13
TMEM242
TRUB1
TTBK2
UBE2K
UBE2N
UBE2V2
ZYG11B
Diagnosis
Sex
Protein-containing Complex Assembly
Autophagosome Membrane Docking
Regulation Of Peptidyl-threonine Phosphorylation
Fc-epsilon Receptor Signaling Pathway
Stress-activated MAPK Cascade
Adaptive Immune System
DARPP-32 Events
Ca-dependent Events
Opioid Signalling
Fc epsilon receptor (FCERI) signaling
Alzheimer's Disease
MicroRNAs In Cardiomyocyte Hypertrophy
Cardiac Hypertrophic Response
Physiological And Pathological Hypertrophy Of The Heart
Endothelin Pathways
HumanNet interactions
Co-expression
Functional
Protein-protein interactions
Diagnosis
AD
Control
Sex
Male
Female
Functional enrichment
Functional enrichment
Motif enrichment
RcisTarget
Transcription factor binding enrichment
ChIP-enrichR
Expression ratios
<= -2.0
-0
>= 2.0

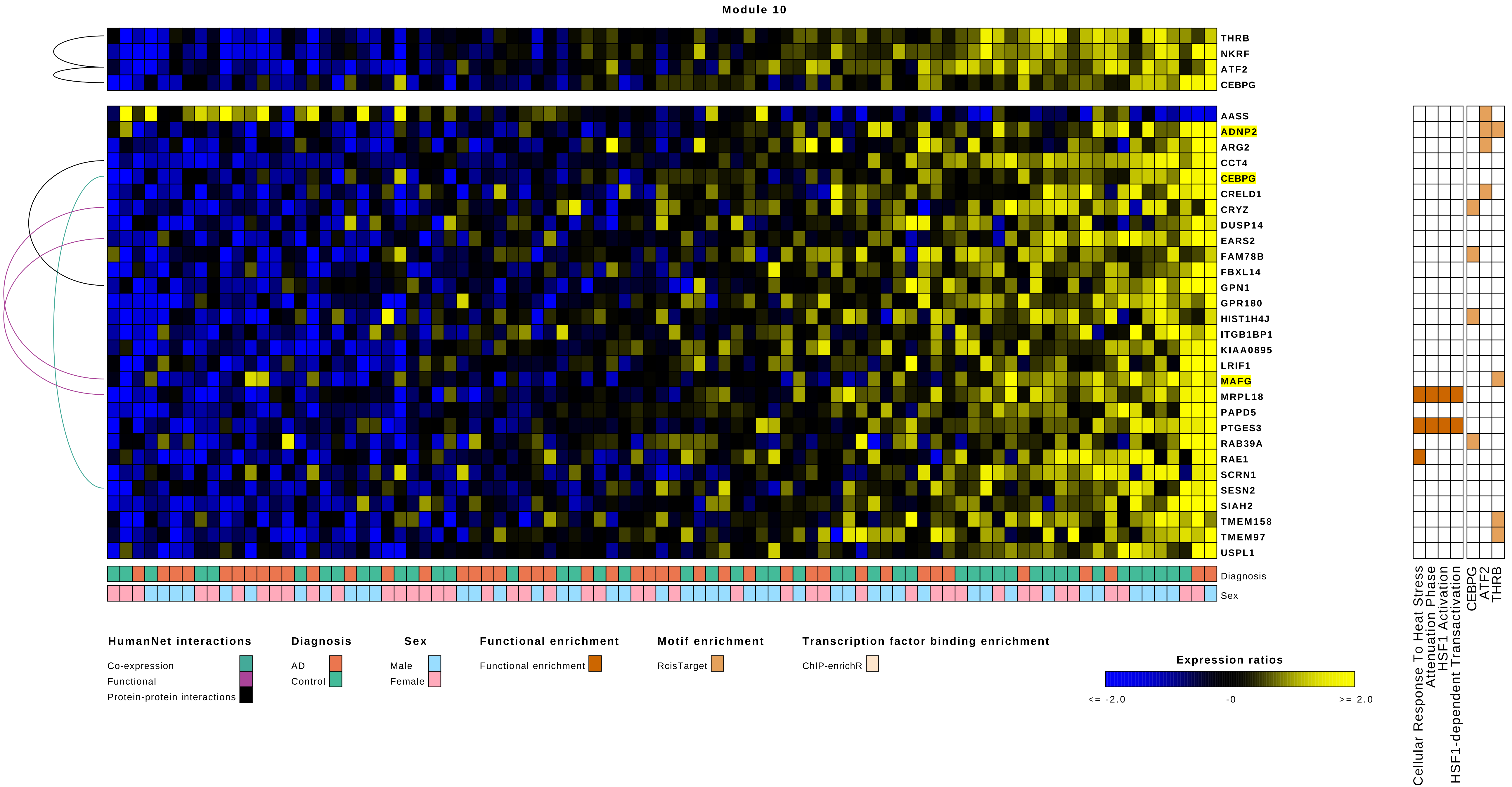

Module 10
THRB
NKRF
ATF2
CEBPG
AASS
ADNP2
ARG2
CCT4
CEBPG
CRELD1
CRYZ
DUSP14
EARS2
FAM78B
FBXL14
GPN1
GPR180
HIST1H4J
ITGB1BP1
KIAA0895
LRIF1
MAFG
MRPL18
PAPD5
PTGES3
RAB39A
RAE1
SCRN1
SESN2
SIAH2
TMEM158
TMEM97
USPL1
Diagnosis
Sex
Cellular Response To Heat Stress
Attenuation Phase
HSF1 Activation
HSF1-dependent Transactivation
CEBPG
ATF2
THRB
HumanNet interactions
Co-expression
Functional
Protein-protein interactions
Diagnosis
AD
Control
Sex
Male
Female
Functional enrichment
Functional enrichment
Motif enrichment
RcisTarget
Transcription factor binding enrichment
ChIP-enrichR
Expression ratios
<= -2.0
-0
>= 2.0

Module 11

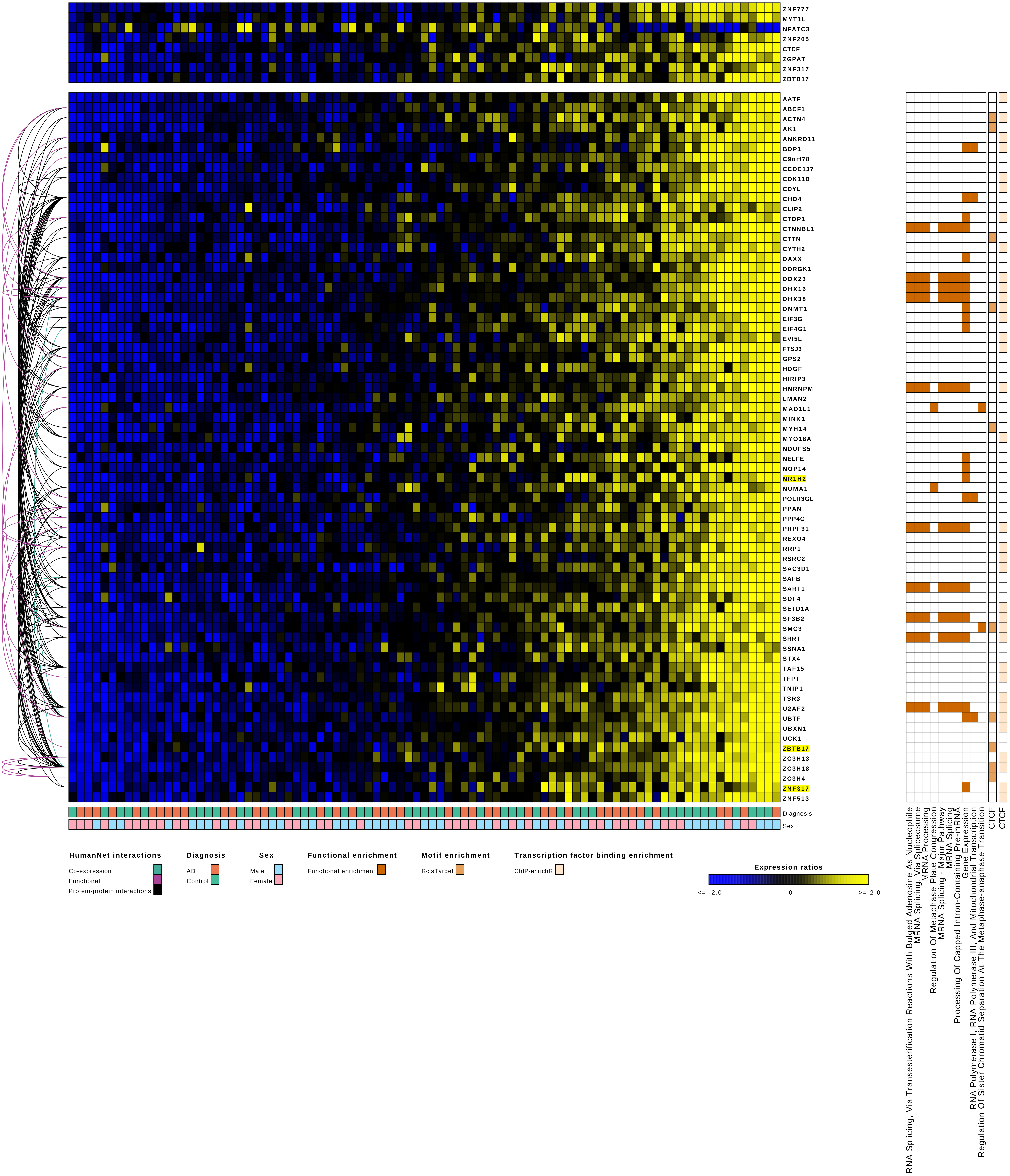
ZNF777
MYT1L
NFATC3
ZNF205
CTCF
ZGPAT
ZNF317
ZBTB17
AATF
ABCF1
ACTN4
AK1
ANKRD11
BDP1
C9orf78
CCDC137
CDK11B
CDYL
CHD4
CLIP2
CTDP1
CTNNBL1
CTTN
CYTH2
DAXX
DDRGK1
DDX23
DHX16
DHX38
DNMT1
EIF3G
EIF4G1
EVI5L
FTSJ3
GPS2
HDGF
HIRIP3
HNRNPM
LMAN2
MAD1L1
MINK1
MYH14
MYO18A
NDUFS5
NELFE
NOP14
NR1H2
NUMA1
POLR3GL
PPAN
PPP4C
PRPF31
REXO4
RRP1
RSRC2
SAC3D1
SAFB
SART1
SDF4
SETD1A
SF3B2
SMC3
SRRT
SSNA1
STX4
TAF15
TFPT
TNIP1
TSR3
U2AF2
UBTF
UBXN1
UCK1
ZBTB17
ZC3H13
ZC3H18
ZC3H4
ZNF317
ZNF513
Diagnosis
Sex
RNA Splicing, Via Transesterification Reactions With Bulged Adenosine As Nucleophile
MRNA Splicing, Via Spliceosome
MRNA Processing
Regulation Of Metaphase Plate Congression
MRNA Splicing - Major Pathway
MRNA Splicing
Processing Of Capped Intron-Containing Pre-mRNA
Gene Expression
RNA Polymerase I, RNA Polymerase III, And Mitochondrial Transcription
Regulation Of Sister Chromatid Separation At The Metaphase-anaphase Transition
CTCF
CTCF
HumanNet interactions
Co-expression
Functional
Protein-protein interactions
Diagnosis
AD
Control
Sex
Male
Female
Functional enrichment
Functional enrichment
Motif enrichment
RcisTarget
Transcription factor binding enrichment
ChIP-enrichR
Expression ratios
<= -2.0
-0
>= 2.0

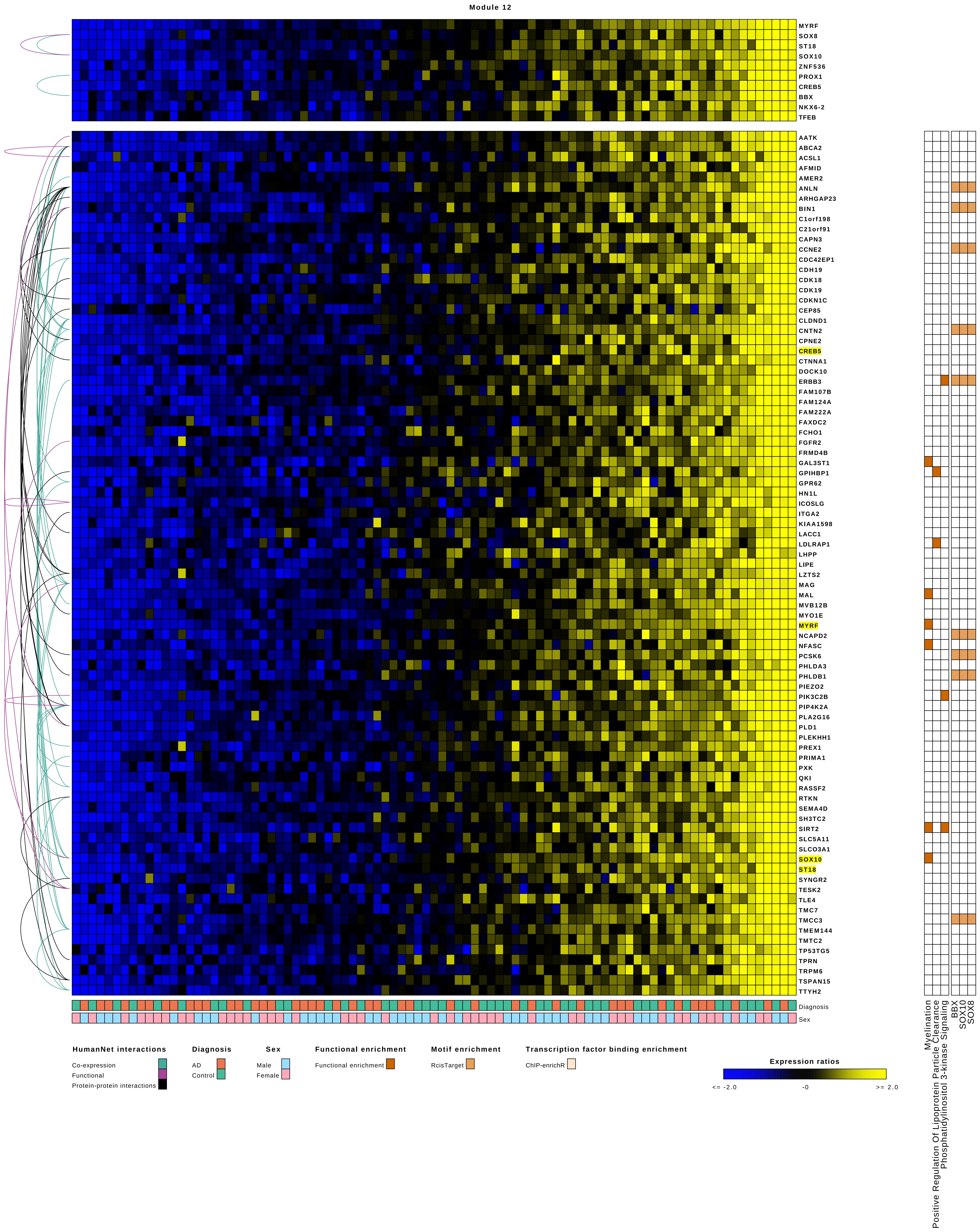
Module 12
MYRF
SOX8
ST18
SOX10
ZNF536
PROX1
CREB5
BBX
NKX6-2
TFEB
AATK
ABCA2
ACSL1
AFMID
AMER2
ANLN
ARHGAP23
BIN1
C1orf198
C21orf91
CAPN3
CCNE2
CDC42EP1
CDH19
CDK18
CDK19
CDKN1C
CEP85
CLDND1
CNTN2
CPNE2
CREB5
CTNNA1
DOCK10
ERBB3
FAM107B
FAM124A
FAM222A
FAXDC2
FCHO1
FGFR2
FRMD4B
GAL3ST1
GPIHBP1
GPR62
HN1L
ICOSLG
ITGA2
KIAA1598
LACC1
LDLRAP1
LHPP
LIPE
LZTS2
MAG
MAL
MVB12B
MYO1E
MYRF
NCAPD2
NFASC
PCSK6
PHLDA3
PHLDB1
PIEZO2
PIK3C2B
PIP4K2A
PLA2G16
PLD1
PLEKHH1
PREX1
PRIMA1
PXK
QKI
RASSF2
RTKN
SEMA4D
SH3TC2
SIRT2
SLC5A11
SLCO3A1
SOX10
ST18
SYNGR2
TESK2
TLE4
TMC7
TMCC3
TMEM144
TMTC2
TP53TG5
TPRN
TRPM6
TSPAN15
TTYH2
Diagnosis
Sex
Myelination
Positive Regulation Of Lipoprotein Particle Clearance
Phosphatidylinositol 3-kinase Signaling
BBX
SOX10
SOX8
HumanNet interactions
Co-expression
Functional
Protein-protein interactions
Diagnosis
AD
Control
Sex
Male
Female
Functional enrichment
Functional enrichment
Motif enrichment
RcisTarget
Transcription factor binding enrichment
ChIP-enrichR
Expression ratios
<= -2.0
-0
>= 2.0

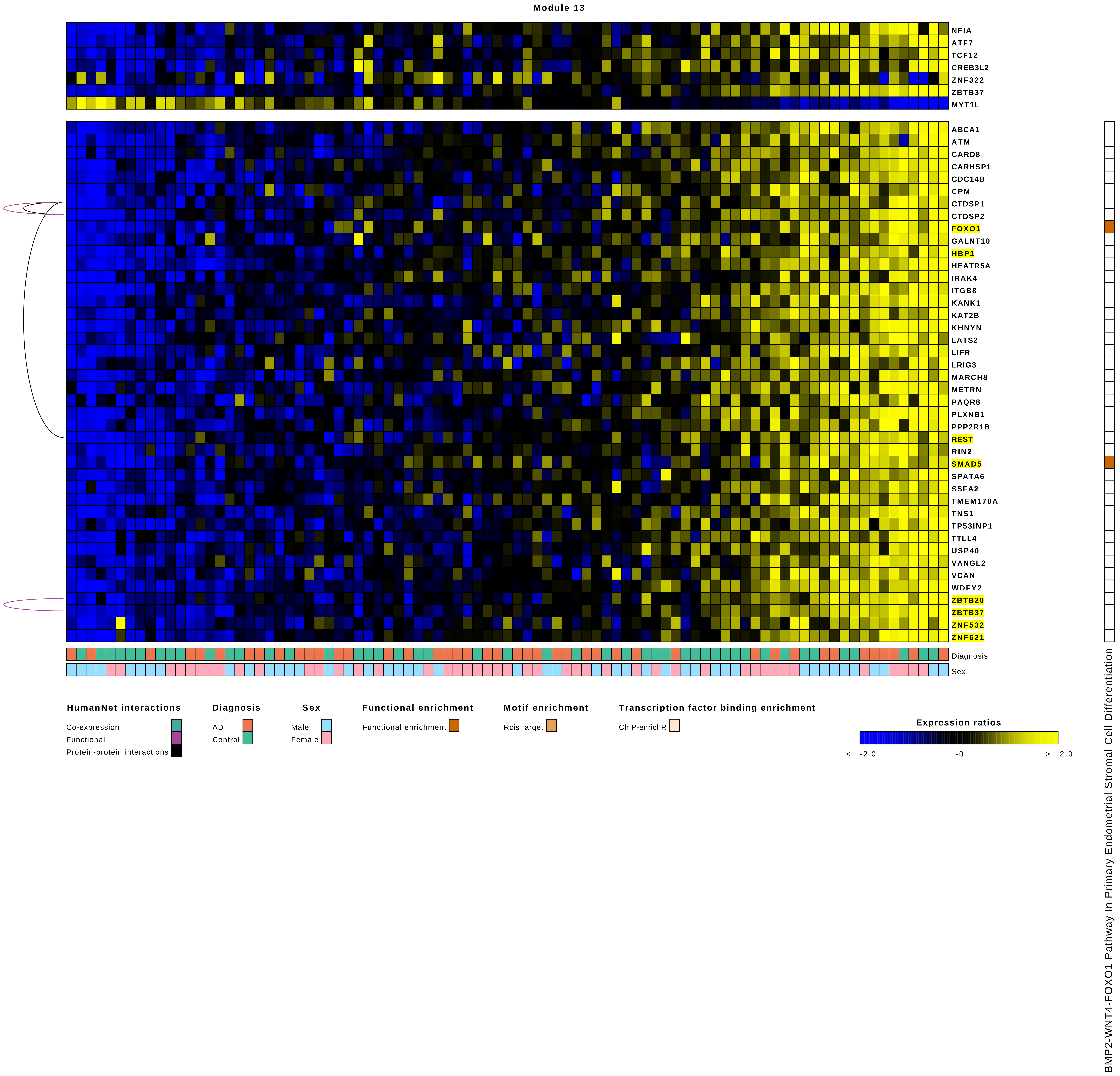

Module 13
NFIA
ATF7
TCF12
CREB3L2
ZNF322
ZBTB37
MYT1L
ABCA1
ATM
CARD8
CARHSP1
CDC14B
CPM
CTDSP1
CTDSP2
FOXO1
GALNT10
HBP1
HEATR5A
IRAK4
ITGB8
KANK1
KAT2B
KHNYN
LATS2
LIFR
LRIG3
MARCH8
METRN
PAQR8
PLXNB1
PPP2R1B
REST
RIN2
SMAD5
SPATA6
SSFA2
TMEM170A
TNS1
TP53INP1
TTLL4
USP40
VANGL2
VCAN
WDFY2
ZBTB20
ZBTB37
ZNF532
ZNF621
Diagnosis
Sex
HumanNet interactions
Co-expression
Functional
Protein-protein interactions
Diagnosis
AD
Control
Sex
Male
Female
Functional enrichment
Functional enrichment
Motif enrichment
RcisTarget
Transcription factor binding enrichment
ChIP-enrichR
Expression ratios
<= -2.0
-0
>= 2.0
BMP2-WNT4-FOXO1 Pathway In Primary Endometrial Stromal Cell Differentiation

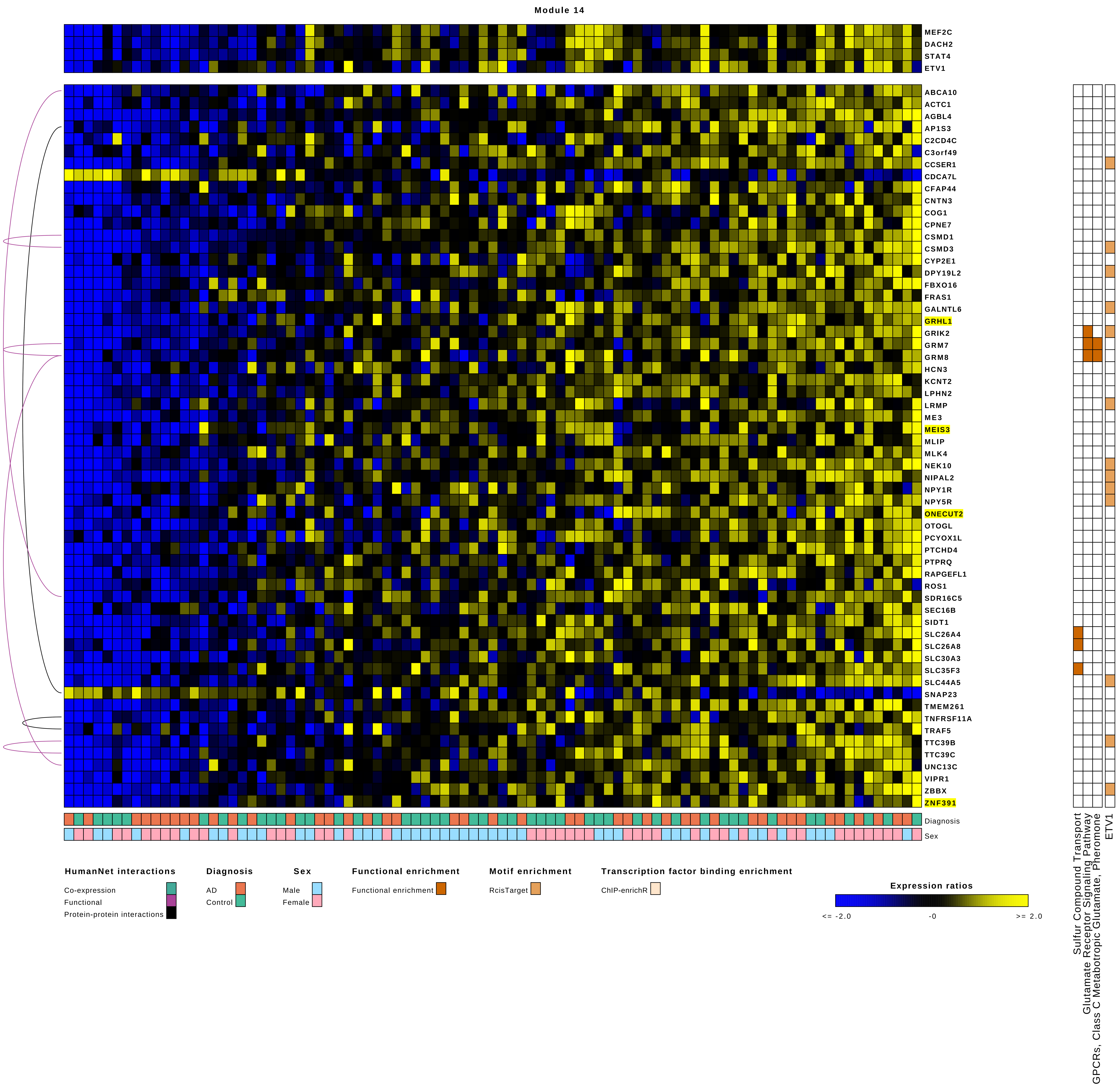
Module 14
MEF2C
DACH2
STAT4
ETV1
ABCA10
ACTC1
AGBL4
AP1S3
C2CD4C
C3orf49
CCSER1
CDCA7L
CFAP44
CNTN3
COG1
CPNE7
CSMD1
CSMD3
CYP2E1
DPY19L2
FBXO16
FRAS1
GALNTL6
GRHL1
GRIK2
GRM7
GRM8
HCN3
KCNT2
LPHN2
LRMP
ME3
MEIS3
MLIP
MLK4
NEK10
NIPAL2
NPY1R
NPY5R
ONECUT2
OTOGL
PCYOX1L
PTCHD4
PTPRQ
RAPGEFL1
ROS1
SDR16C5
SEC16B
SIDT1
SLC26A4
SLC26A8
SLC30A3
SLC35F3
SLC44A5
SNAP23
TMEM261
TNFRSF11A
TRAF5
TTC39B
TTC39C
UNC13C
VIPR1
ZBBX
ZNF391
Diagnosis
Sex
Sulfur Compound Transport
Glutamate Receptor Signaling Pathway
GPCRs, Class C Metabotropic Glutamate, Pheromone
ETV1
HumanNet interactions
Co-expression
Functional
Protein-protein interactions
Diagnosis
AD
Control
Sex
Male
Female
Functional enrichment
Functional enrichment
Motif enrichment
RcisTarget
Transcription factor binding enrichment
ChIP-enrichR
Expression ratios
<= -2.0
-0
>= 2.0

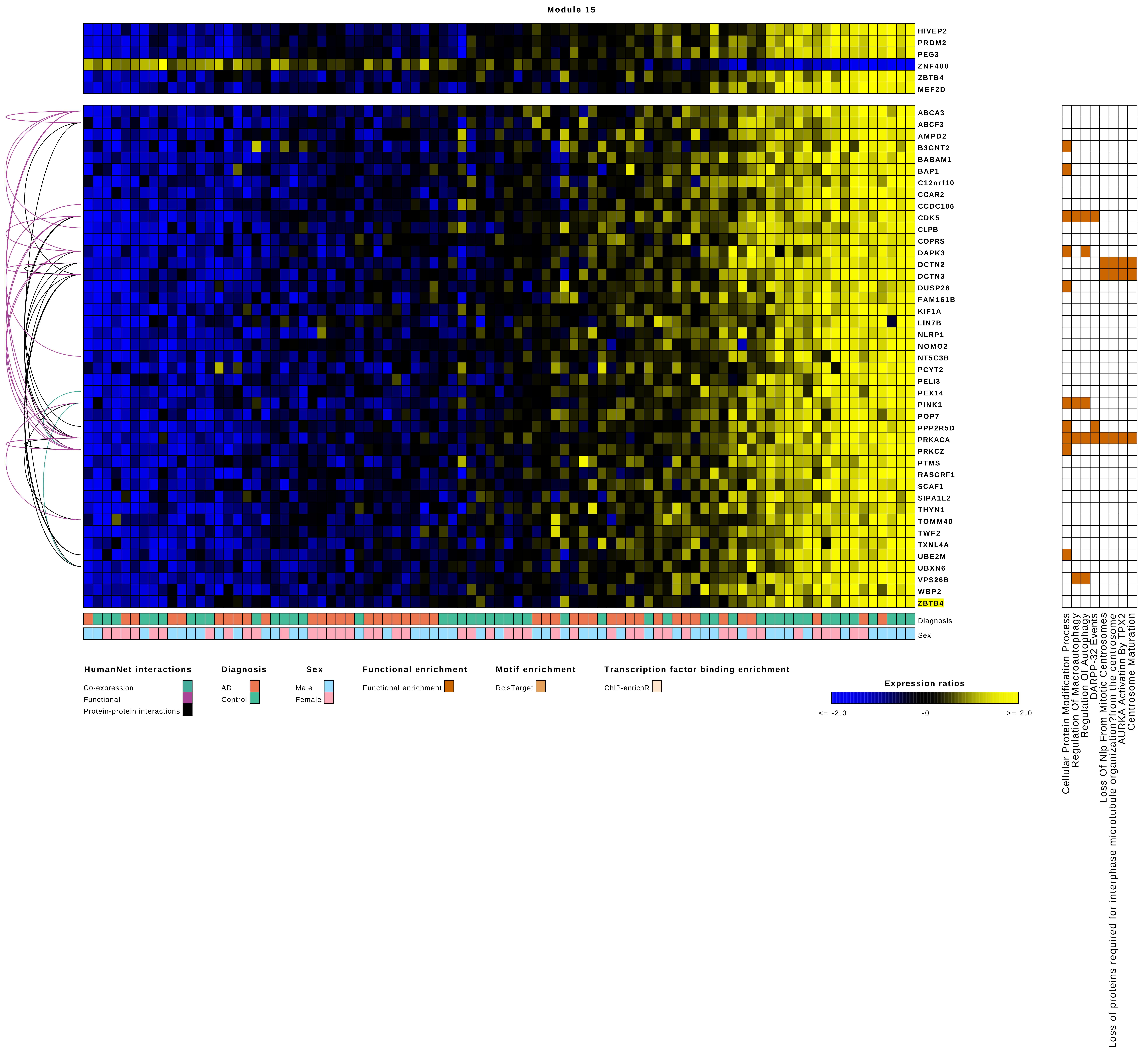
Module 15
HIVEP2
PRDM2
PEG3
ZNF480
ZBTB4
MEF2D
ABCA3
ABCF3
AMPD2
B3GNT2
BABAM1
BAP1
C12orf10
CCAR2
CCDC106
CDK5
CLPB
COPRS
DAPK3
DCTN2
DCTN3
DUSP26
FAM161B
KIF1A
LIN7B
NLRP1
NOMO2
NT5C3B
PCYT2
PELI3
PEX14
PINK1
POP7
PPP2R5D
PRKACA
PRKCZ
PTMS
RASGRF1
SCAF1
SIPA1L2
THYN1
TOMM40
TWF2
TXNL4A
UBE2M
UBXN6
VPS26B
WBP2
ZBTB4
Diagnosis
Sex
Cellular Protein Modification Process
Regulation Of Macroautophagy
Regulation Of Autophagy
DARPP-32 Events
Loss Of Nlp From Mitotic Centrosomes
Loss of proteins required for interphase microtubule organization?from the centrosome
AURKA Activation By TPX2
Centrosome Maturation
HumanNet interactions
Co-expression
Functional
Protein-protein interactions
Diagnosis
AD
Control
Sex
Male
Female
Functional enrichment
Functional enrichment
Motif enrichment
RcisTarget
Transcription factor binding enrichment
ChIP-enrichR
Expression ratios
<= -2.0
-0
>= 2.0

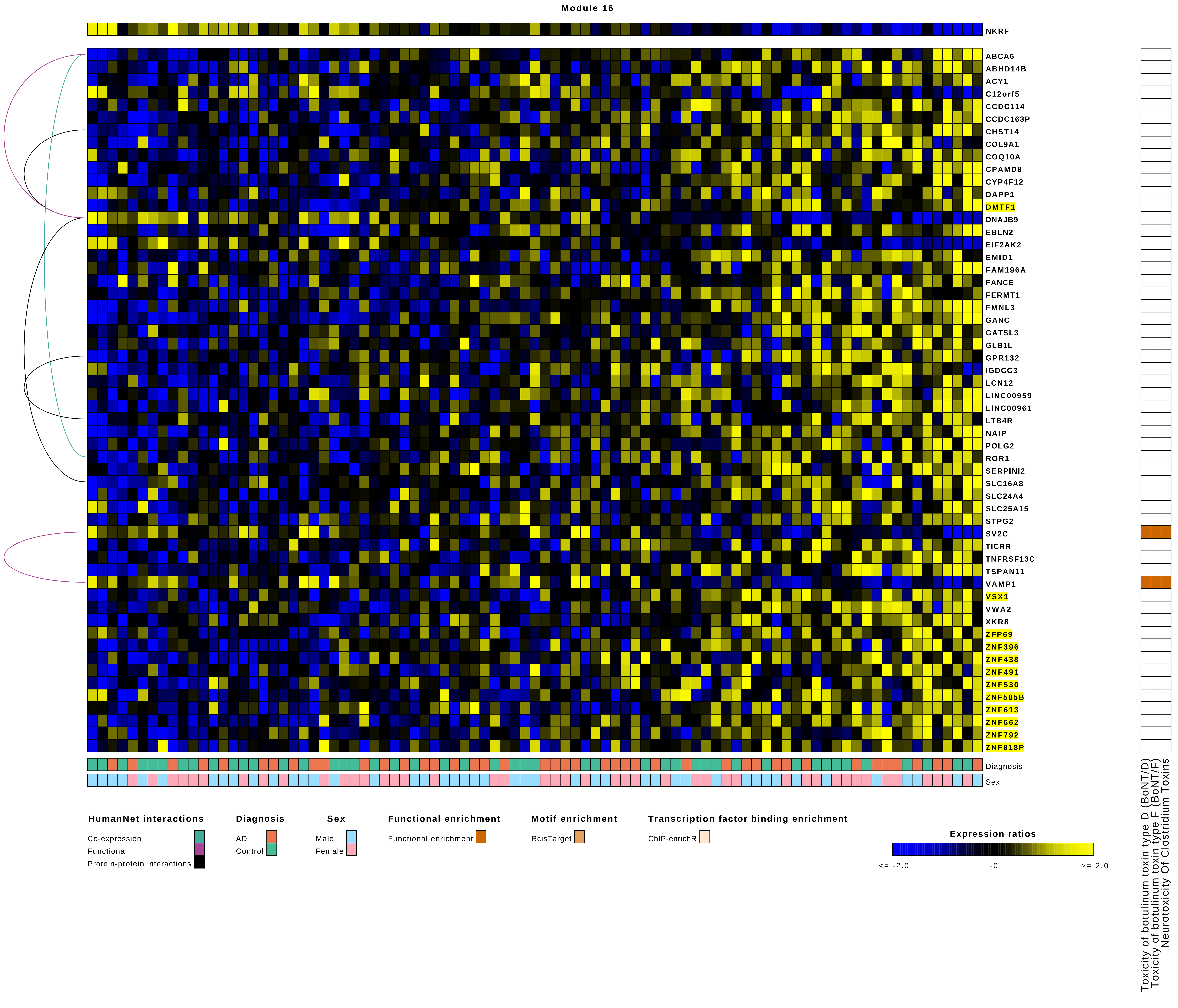
Module 16
NKRF
ABCA6
ABHD14B
ACY1
C12orf5
CCDC114
CCDC163P
CHST14
COL9A1
COQ10A
CPAMD8
CYP4F12
DAPP1
DMTF1
DNAJB9
EBLN2
EIF2AK2
EMID1
FAM196A
FANCE
FERMT1
FMNL3
GANC
GATSL3
GLB1L
GPR132
IGDCC3
LCN12
LINC00959
LINC00961
LTB4R
NAIP
POLG2
ROR1
SERPINI2
SLC16A8
SLC24A4
SLC25A15
STPG2
SV2C
TICRR
TNFRSF13C
TSPAN11
VAMP1
VSX1
VWA2
XKR8
ZFP69
ZNF396
ZNF438
ZNF491
ZNF530
ZNF585B
ZNF613
ZNF662
ZNF792
ZNF818P
Diagnosis
Sex
Toxicity of botulinum toxin type D (BoNT/D)
Toxicity of botulinum toxin type F (BoNT/F)
Neurotoxicity Of Clostridium Toxins
HumanNet interactions
Co-expression
Functional
Protein-protein interactions
Diagnosis
AD
Control
Sex
Male
Female
Functional enrichment
Functional enrichment
Motif enrichment
RcisTarget
Transcription factor binding enrichment
ChIP-enrichR
Expression ratios
<= -2.0
-0
>= 2.0

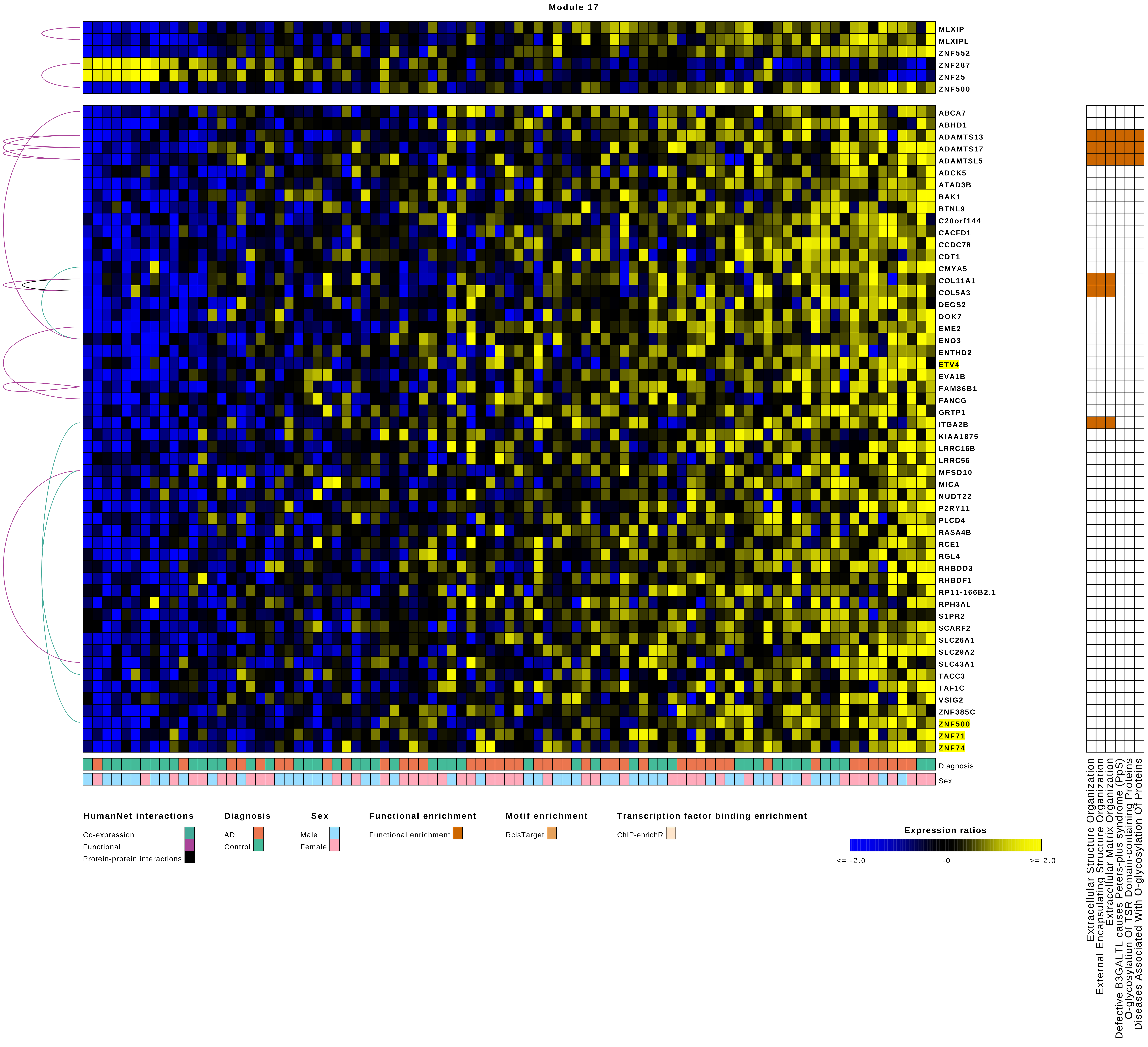
Module 17
MLXIP
MLXIPL
ZNF552
ZNF287
ZNF25
ZNF500
ABCA7
ABHD1
ADAMTS13
ADAMTS17
ADAMTSL5
ADCK5
ATAD3B
BAK1
BTNL9
C20orf144
CACFD1
CCDC78
CDT1
CMYA5
COL11A1
COL5A3
DEGS2
DOK7
EME2
ENO3
ENTHD2
ETV4
EVA1B
FAM86B1
FANCG
GRTP1
ITGA2B
KIAA1875
LRRC16B
LRRC56
MFSD10
MICA
NUDT22
P2RY11
PLCD4
RASA4B
RCE1
RGL4
RHBDD3
RHBDF1
RP11-166B2.1
RPH3AL
S1PR2
SCARF2
SLC26A1
SLC29A2
SLC43A1
TACC3
TAF1C
VSIG2
ZNF385C
ZNF500
ZNF71
ZNF74
Diagnosis
Sex
Extracellular Structure Organization
External Encapsulating Structure Organization
Extracellular Matrix Organization
Defective B3GALTL causes Peters-plus syndrome (PpS)
O-glycosylation Of TSR Domain-containing Proteins
Diseases Associated With O-glycosylation Of Proteins
HumanNet interactions
Co-expression
Functional
Protein-protein interactions
Diagnosis
AD
Control
Sex
Male
Female
Functional enrichment
Functional enrichment
Motif enrichment
RcisTarget
Transcription factor binding enrichment
ChIP-enrichR
Expression ratios
<= -2.0
-0
>= 2.0

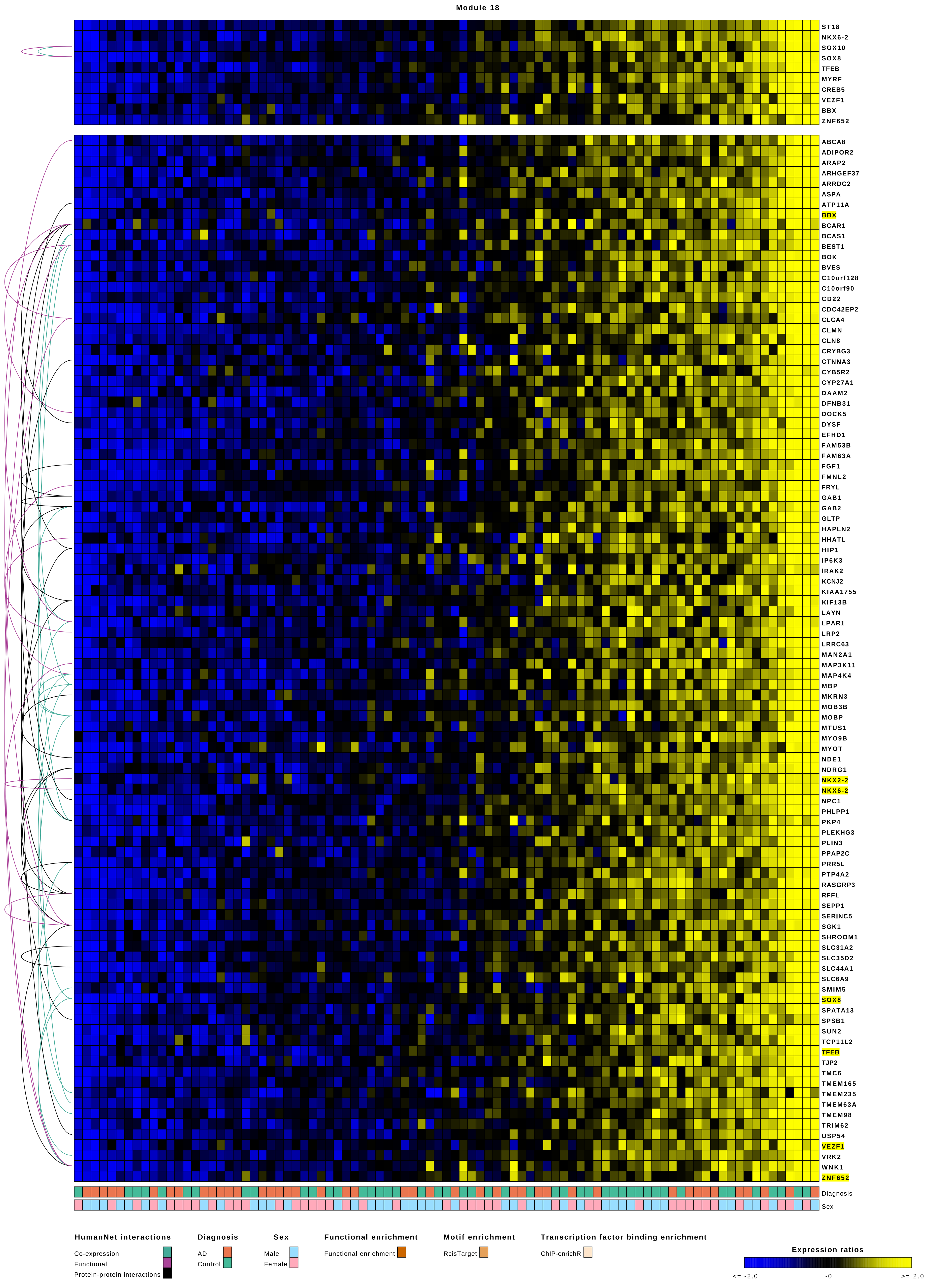
Module 18
ST18
NKX6-2
SOX10
SOX8
TFEB
MYRF
CREB5
VEZF1
BBX
ZNF652
ABCA8
ADIPOR2
ARAP2
ARHGEF37
ARRDC2
ASPA
ATP11A
BBX
BCAR1
BCAS1
BEST1
BOK
BVES
C10orf128
C10orf90
CD22
CDC42EP2
CLCA4
CLMN
CLN8
CRYBG3
CTNNA3
CYB5R2
CYP27A1
DAAM2
DFNB31
DOCK5
DYSF
EFHD1
FAM53B
FAM63A
FGF1
FMNL2
FRYL
GAB1
GAB2
GLTP
HAPLN2
HHATL
HIP1
IP6K3
IRAK2
KCNJ2
KIAA1755
KIF13B
LAYN
LPAR1
LRP2
LRRC63
MAN2A1
MAP3K11
MAP4K4
MBP
MKRN3
MOB3B
MOBP
MTUS1
MYO9B
MYOT
NDE1
NDRG1
NKX2-2
NKX6-2
NPC1
PHLPP1
PKP4
PLEKHG3
PLIN3
PPAP2C
PRR5L
PTP4A2
RASGRP3
RFFL
SEPP1
SERINC5
SGK1
SHROOM1
SLC31A2
SLC35D2
SLC44A1
SLC6A9
SMIM5
SOX8
SPATA13
SPSB1
SUN2
TCP11L2
TFEB
TJP2
TMC6
TMEM165
TMEM235
TMEM63A
TMEM98
TRIM62
USP54
VEZF1
VRK2
WNK1
ZNF652
Diagnosis
Sex
HumanNet interactions
Co-expression
Functional
Protein-protein interactions
Diagnosis
AD
Control
Sex
Male
Female
Functional enrichment
Functional enrichment
Motif enrichment
RcisTarget
Transcription factor binding enrichment
ChIP-enrichR
Expression ratios
<= -2.0
-0
>= 2.0

Module 19

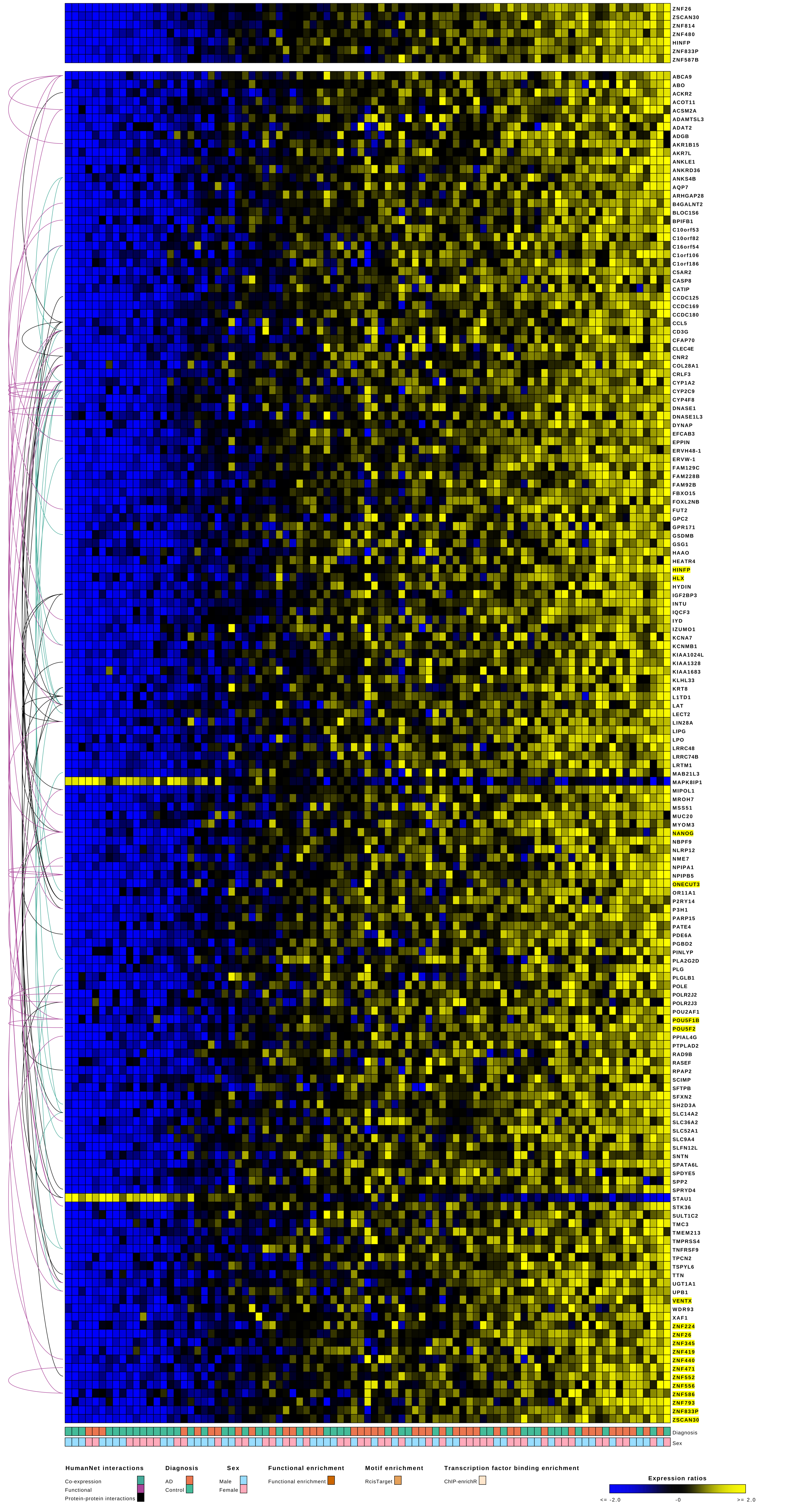
ZNF26
ZSCAN30
ZNF814
ZNF480
HINFP
ZNF833P
ZNF587B
ABCA9
ABO
ACKR2
ACOT11
ACSM2A
ADAMTSL3
ADAT2
ADGB
AKR1B15
AKR7L
ANKLE1
ANKRD36
ANKS4B
AQP7
ARHGAP28
B4GALNT2
BLOC1S6
BPIFB1
C10orf53
C10orf82
C16orf54
C1orf106
C1orf186
C5AR2
CASP8
CATIP
CCDC125
CCDC169
CCDC180
CCL5
CD3G
CFAP70
CLEC4E
CNR2
COL28A1
CRLF3
CYP1A2
CYP2C9
CYP4F8
DNASE1
DNASE1L3
DYNAP
EFCAB3
EPPIN
ERVH48-1
ERVW-1
FAM129C
FAM228B
FAM92B
FBXO15
FOXL2NB
FUT2
GPC2
GPR171
GSDMB
GSG1
HAAO
HEATR4
HINFP
HLX
HYDIN
IGF2BP3
INTU
IQCF3
IYD
IZUMO1
KCNA7
KCNMB1
KIAA1024L
KIAA1328
KIAA1683
KLHL33
KRT8
L1TD1
LAT
LECT2
LIN28A
LIPG
LPO
LRRC48
LRRC74B
LRTM1
MAB21L3
MAPK8IP1
MIPOL1
MROH7
MSS51
MUC20
MYOM3
NANOG
NBPF9
NLRP12
NME7
NPIPA1
NPIPB5
ONECUT3
OR11A1
P2RY14
P3H1
PARP15
PATE4
PDE6A
PGBD2
PINLYP
PLA2G2D
PLG
PLGLB1
POLE
POLR2J2
POLR2J3
POU2AF1
POU5F1B
POU5F2
PPIAL4G
PTPLAD2
RAD9B
RASEF
RPAP2
SCIMP
SFTPB
SFXN2
SH2D3A
SLC14A2
SLC36A2
SLC52A1
SLC9A4
SLFN12L
SNTN
SPATA6L
SPDYE5
SPP2
SPRYD4
STAU1
STK36
SULT1C2
TMC3
TMEM213
TMPRSS4
TNFRSF9
TPCN2
TSPYL6
TTN
UGT1A1
UPB1
VENTX
WDR93
XAF1
ZNF224
ZNF26
ZNF345
ZNF419
ZNF440
ZNF471
ZNF552
ZNF556
ZNF586
ZNF793
ZNF833P
ZSCAN30
Diagnosis
Sex
HumanNet interactions
Co-expression
Functional
Protein-protein interactions
Diagnosis
AD
Control
Sex
Male
Female
Functional enrichment
Functional enrichment
Motif enrichment
RcisTarget
Transcription factor binding enrichment
ChIP-enrichR
Expression ratios
<= -2.0
-0
>= 2.0

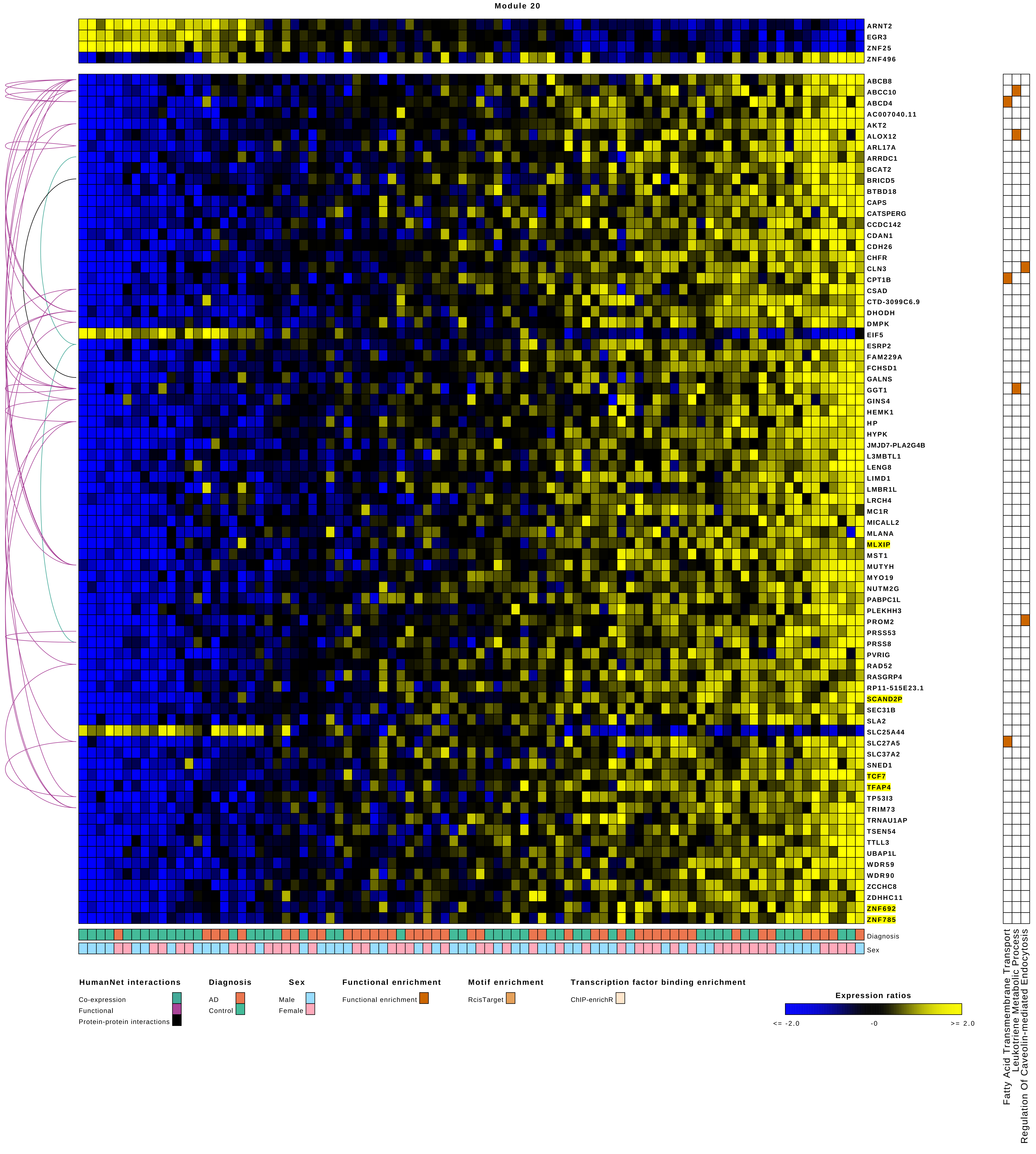

Module 20
ARNT2
EGR3
ZNF25
ZNF496
ABCB8
ABCC10
ABCD4
AC007040.11
AKT2
ALOX12
ARL17A
ARRDC1
BCAT2
BRICD5
BTBD18
CAPS
CATSPERG
CCDC142
CDAN1
CDH26
CHFR
CLN3
CPT1B
CSAD
CTD-3099C6.9
DHODH
DMPK
EIF5
ESRP2
FAM229A
FCHSD1
GALNS
GGT1
GINS4
HEMK1
HP
HYPK
JMJD7-PLA2G4B
L3MBTL1
LENG8
LIMD1
LMBR1L
LRCH4
MC1R
MICALL2
MLANA
MLXIP
MST1
MUTYH
MYO19
NUTM2G
PABPC1L
PLEKHH3
PROM2
PRSS53
PRSS8
PVRIG
RAD52
RASGRP4
RP11-515E23.1
SCAND2P
SEC31B
SLA2
SLC25A44
SLC27A5
SLC37A2
SNED1
TCF7
TFAP4
TP53I3
TRIM73
TRNAU1AP
TSEN54
TTLL3
UBAP1L
WDR59
WDR90
ZCCHC8
ZDHHC11
ZNF692
ZNF785
Diagnosis
Sex
Fatty Acid Transmembrane Transport
Leukotriene Metabolic Process
Regulation Of Caveolin-mediated Endocytosis
HumanNet interactions
Co-expression
Functional
Protein-protein interactions
Diagnosis
AD
Control
Sex
Male
Female
Functional enrichment
Functional enrichment
Motif enrichment
RcisTarget
Transcription factor binding enrichment
ChIP-enrichR
Expression ratios
<= -2.0
-0
>= 2.0

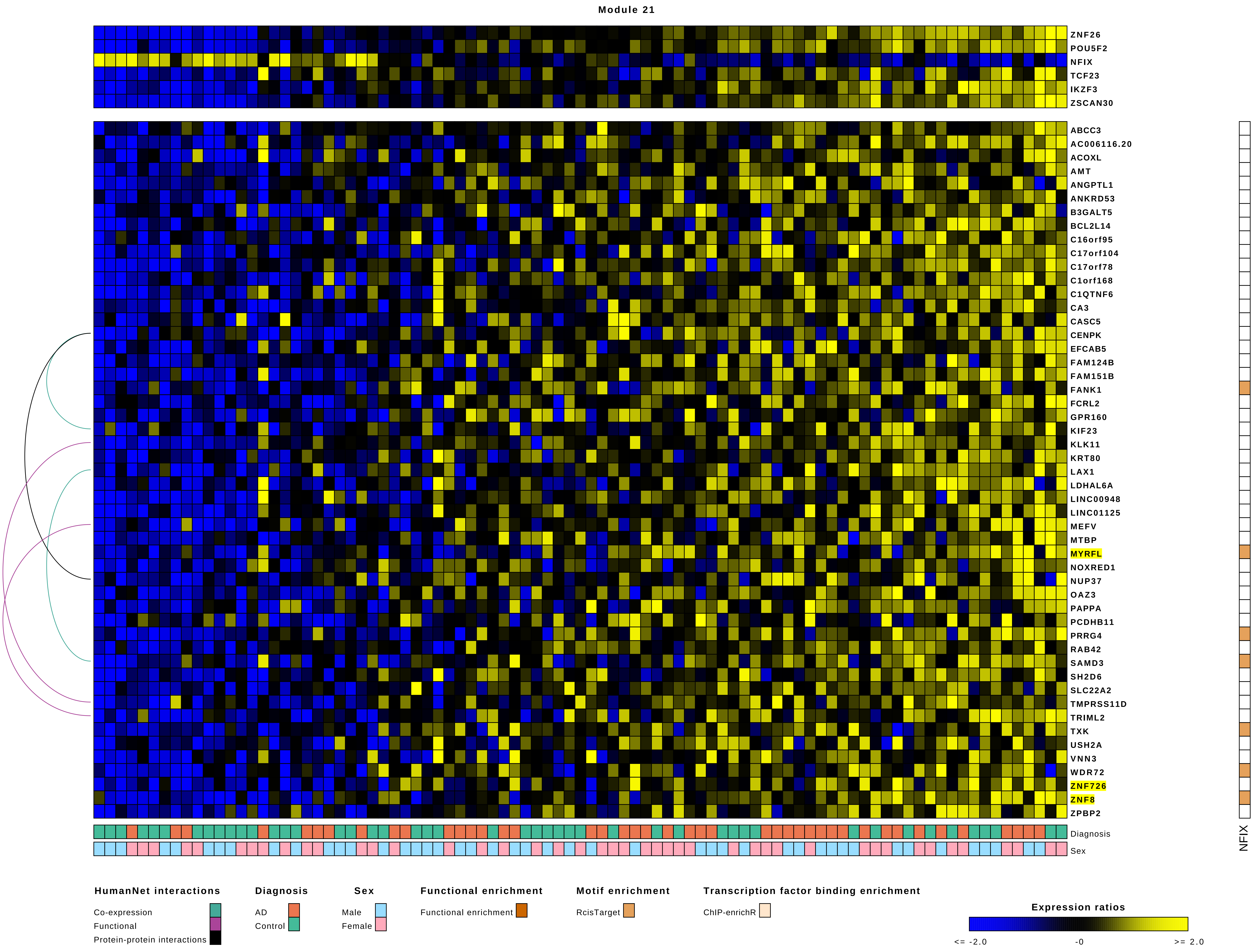
Module 21
ZNF26
POU5F2
NFIX
TCF23
IKZF3
ZSCAN30
ABCC3
AC006116.20
ACOXL
AMT
ANGPTL1
ANKRD53
B3GALT5
BCL2L14
C16orf95
C17orf104
C17orf78
C1orf168
C1QTNF6
CA3
CASC5
CENPK
EFCAB5
FAM124B
FAM151B
FANK1
FCRL2
GPR160
KIF23
KLK11
KRT80
LAX1
LDHAL6A
LINC00948
LINC01125
MEFV
MTBP
MYRFL
NOXRED1
NUP37
OAZ3
PAPPA
PCDHB11
PRRG4
RAB42
SAMD3
SH2D6
SLC22A2
TMPRSS11D
TRIML2
TXK
USH2A
VNN3
WDR72
ZNF726
ZNF8
ZPBP2
Diagnosis
Sex
NFIX
HumanNet interactions
Co-expression
Functional
Protein-protein interactions
Diagnosis
AD
Control
Sex
Male
Female
Functional enrichment
Functional enrichment
Motif enrichment
RcisTarget
Transcription factor binding enrichment
ChIP-enrichR
Expression ratios
<= -2.0
-0
>= 2.0

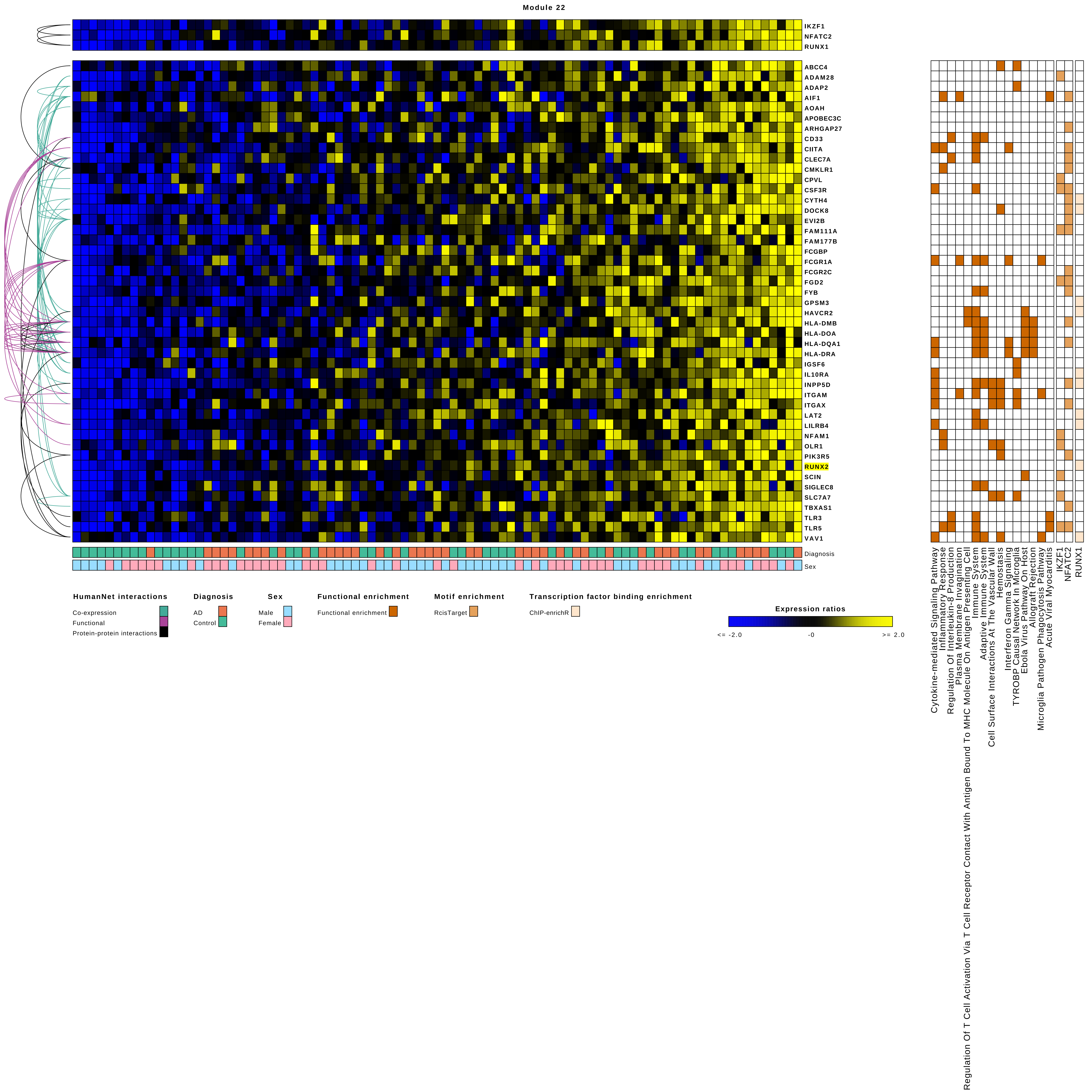
Module 22
IKZF1
NFATC2
RUNX1
ABCC4
ADAM28
ADAP2
AIF1
AOAH
APOBEC3C
ARHGAP27
CD33
CIITA
CLEC7A
CMKLR1
CPVL
CSF3R
CYTH4
DOCK8
EVI2B
FAM111A
FAM177B
FCGBP
FCGR1A
FCGR2C
FGD2
FYB
GPSM3
HAVCR2
HLA-DMB
HLA-DOA
HLA-DQA1
HLA-DRA
IGSF6
IL10RA
INPP5D
ITGAM
ITGAX
LAT2
LILRB4
NFAM1
OLR1
PIK3R5
RUNX2
SCIN
SIGLEC8
SLC7A7
TBXAS1
TLR3
TLR5
VAV1
Diagnosis
Sex
HumanNet interactions
Co-expression
Functional
Protein-protein interactions
Diagnosis
AD
Control
Sex
Male
Female
Functional enrichment
Functional enrichment
Motif enrichment
RcisTarget
Transcription factor binding enrichment
ChIP-enrichR
Expression ratios
<= -2.0
-0
>= 2.0
Cytokine-mediated Signaling Pathway
Inflammatory Response
Regulation Of Interleukin-8 Production
Plasma Membrane Invagination
Regulation Of T Cell Activation Via T Cell Receptor Contact With Antigen Bound To MHC Molecule On Antigen Presenting Cell
Immune System
Adaptive Immune System
Cell Surface Interactions At The Vascular Wall
Hemostasis
Interferon Gamma Signaling
TYROBP Causal Network In Microglia
Ebola Virus Pathway On Host
Allograft Rejection
Microglia Pathogen Phagocytosis Pathway
Acute Viral Myocarditis
IKZF1
NFATC2
RUNX1

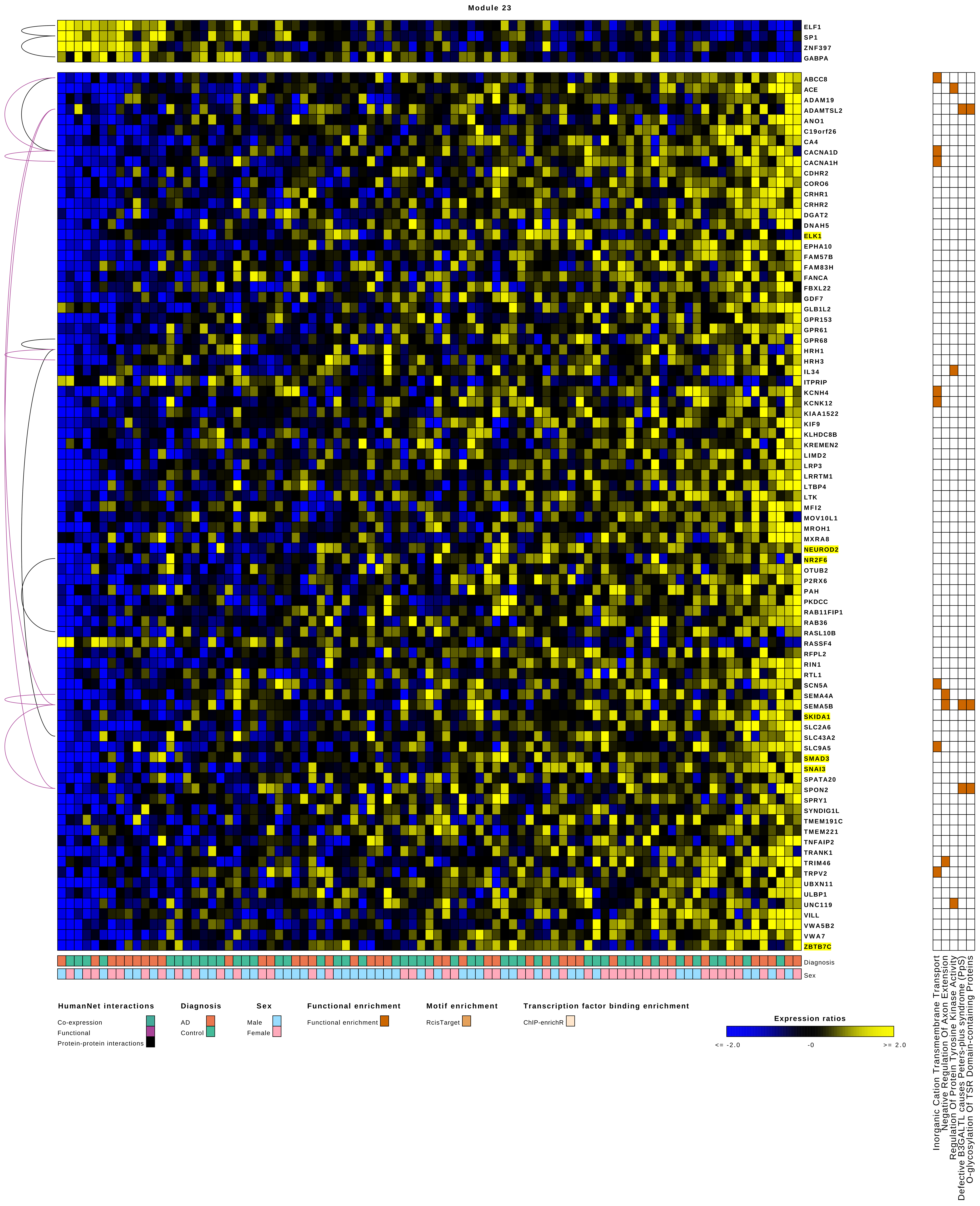
Module 23
ELF1
SP1
ZNF397
GABPA
ABCC8
ACE
ADAM19
ADAMTSL2
ANO1
C19orf26
CA4
CACNA1D
CACNA1H
CDHR2
CORO6
CRHR1
CRHR2
DGAT2
DNAH5
ELK1
EPHA10
FAM57B
FAM83H
FANCA
FBXL22
GDF7
GLB1L2
GPR153
GPR61
GPR68
HRH1
HRH3
IL34
ITPRIP
KCNH4
KCNK12
KIAA1522
KIF9
KLHDC8B
KREMEN2
LIMD2
LRP3
LRRTM1
LTBP4
LTK
MFI2
MOV10L1
MROH1
MXRA8
NEUROD2
NR2F6
OTUB2
P2RX6
PAH
PKDCC
RAB11FIP1
RAB36
RASL10B
RASSF4
RFPL2
RIN1
RTL1
SCN5A
SEMA4A
SEMA5B
SKIDA1
SLC2A6
SLC43A2
SLC9A5
SMAD3
SNAI3
SPATA20
SPON2
SPRY1
SYNDIG1L
TMEM191C
TMEM221
TNFAIP2
TRANK1
TRIM46
TRPV2
UBXN11
ULBP1
UNC119
VILL
VWA5B2
VWA7
ZBTB7C
Diagnosis
Sex
Inorganic Cation Transmembrane Transport
Negative Regulation Of Axon Extension
Regulation Of Protein Tyrosine Kinase Activity
Defective B3GALTL causes Peters-plus syndrome (PpS)
O-glycosylation Of TSR Domain-containing Proteins
HumanNet interactions
Co-expression
Functional
Protein-protein interactions
Diagnosis
AD
Control
Sex
Male
Female
Functional enrichment
Functional enrichment
Motif enrichment
RcisTarget
Transcription factor binding enrichment
ChIP-enrichR
Expression ratios
<= -2.0
-0
>= 2.0

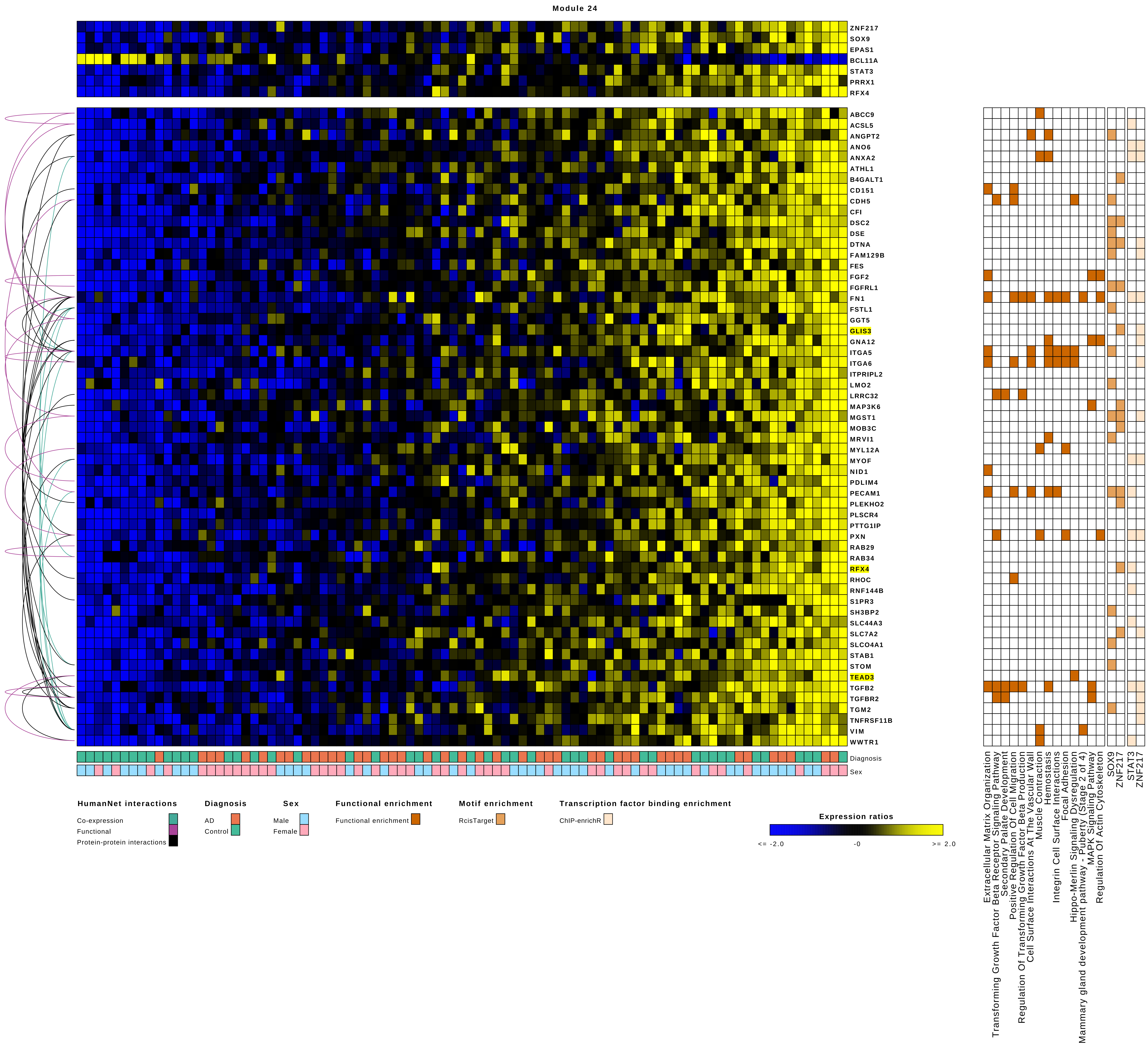

Module 24
ZNF217
SOX9
EPAS1
BCL11A
STAT3
PRRX1
RFX4
ABCC9
ACSL5
ANGPT2
ANO6
ANXA2
ATHL1
B4GALT1
CD151
CDH5
CFI
DSC2
DSE
DTNA
FAM129B
FES
FGF2
FGFRL1
FN1
FSTL1
GGT5
GLIS3
GNA12
ITGA5
ITGA6
ITPRIPL2
LMO2
LRRC32
MAP3K6
MGST1
MOB3C
MRVI1
MYL12A
MYOF
NID1
PDLIM4
PECAM1
PLEKHO2
PLSCR4
PTTG1IP
PXN
RAB29
RAB34
RFX4
RHOC
RNF144B
S1PR3
SH3BP2
SLC44A3
SLC7A2
SLCO4A1
STAB1
STOM
TEAD3
TGFB2
TGFBR2
TGM2
TNFRSF11B
VIM
WWTR1
Diagnosis
Sex
HumanNet interactions
Co-expression
Functional
Protein-protein interactions
Diagnosis
AD
Control
Sex
Male
Female
Functional enrichment
Functional enrichment
Motif enrichment
RcisTarget
Transcription factor binding enrichment
ChIP-enrichR
Expression ratios
<= -2.0
-0
>= 2.0
Extracellular Matrix Organization
Transforming Growth Factor Beta Receptor Signaling Pathway
Secondary Palate Development
Positive Regulation Of Cell Migration
Regulation Of Transforming Growth Factor Beta Production
Cell Surface Interactions At The Vascular Wall
Muscle Contraction
Hemostasis
Integrin Cell Surface Interactions
Focal Adhesion
Hippo-Merlin Signaling Dysregulation
Mammary gland development pathway - Puberty (Stage 2 of 4)
MAPK Signaling Pathway
Regulation Of Actin Cytoskeleton
SOX9
ZNF217
STAT3
ZNF217

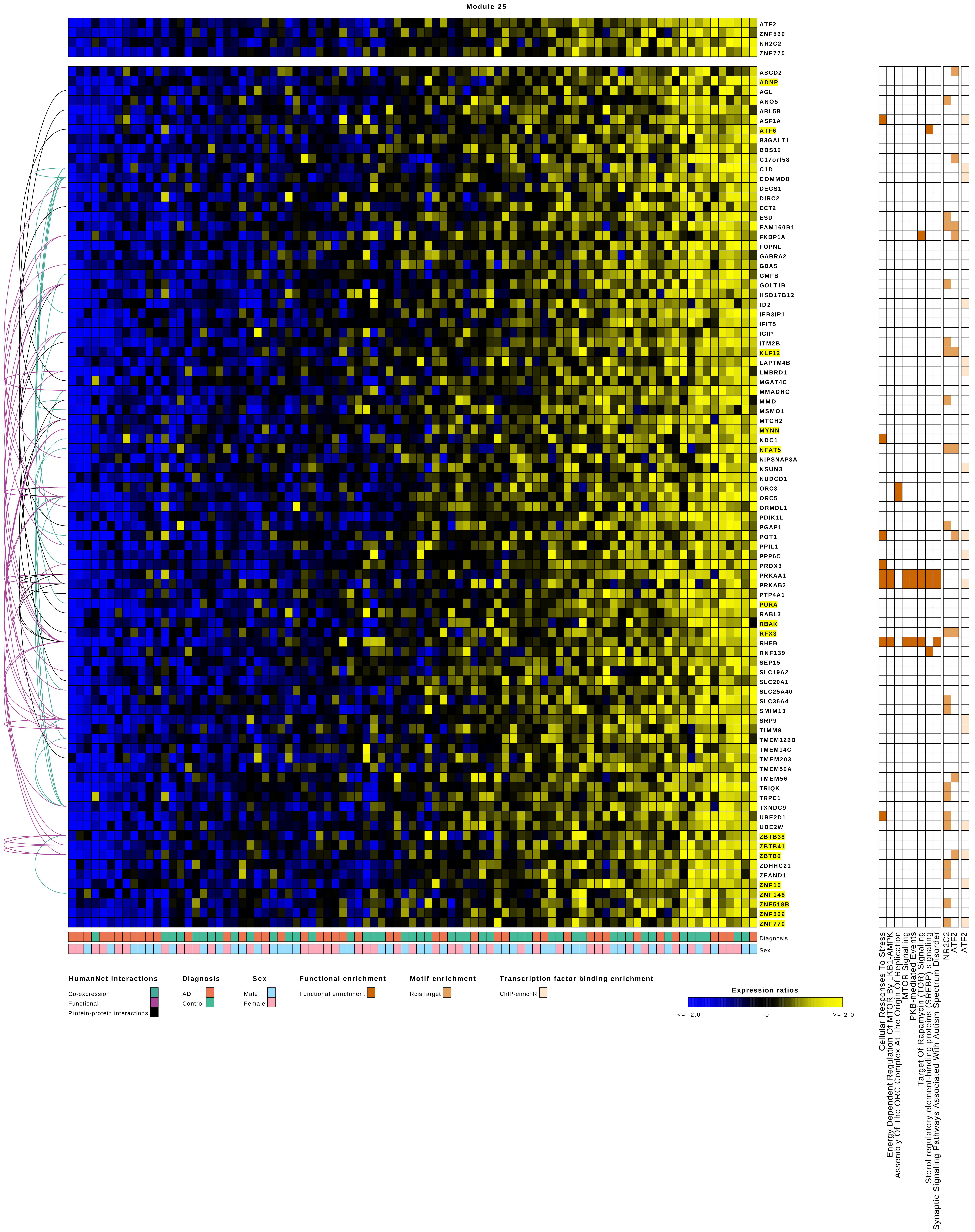

Module 25
ATF2
ZNF569
NR2C2
ZNF770
ABCD2
ADNP
AGL
ANO5
ARL5B
ASF1A
ATF6
B3GALT1
BBS10
C17orf58
C1D
COMMD8
DEGS1
DIRC2
ECT2
ESD
FAM160B1
FKBP1A
FOPNL
GABRA2
GBAS
GMFB
GOLT1B
HSD17B12
ID2
IER3IP1
IFIT5
IGIP
ITM2B
KLF12
LAPTM4B
LMBRD1
MGAT4C
MMADHC
MMD
MSMO1
MTCH2
MYNN
NDC1
NFAT5
NIPSNAP3A
NSUN3
NUDCD1
ORC3
ORC5
ORMDL1
PDIK1L
PGAP1
POT1
PPIL1
PPP6C
PRDX3
PRKAA1
PRKAB2
PTP4A1
PURA
RABL3
RBAK
RFX3
RHEB
RNF139
SEP15
SLC19A2
SLC20A1
SLC25A40
SLC36A4
SMIM13
SRP9
TIMM9
TMEM126B
TMEM14C
TMEM203
TMEM50A
TMEM56
TRIQK
TRPC1
TXNDC9
UBE2D1
UBE2W
ZBTB38
ZBTB41
ZBTB6
ZDHHC21
ZFAND1
ZNF10
ZNF148
ZNF518B
ZNF569
ZNF770
Diagnosis
Sex
Cellular Responses To Stress
Energy Dependent Regulation Of MTOR By LKB1-AMPK
Assembly Of The ORC Complex At The Origin Of Replication
MTOR Signalling
PKB-mediated Events
Target Of Rapamycin (TOR) Signaling
Sterol regulatory element-binding proteins (SREBP) signaling
Synaptic Signaling Pathways Associated With Autism Spectrum Disorder
NR2C2
ATF2
ATF2
HumanNet interactions
Co-expression
Functional
Protein-protein interactions
Diagnosis
AD
Control
Sex
Male
Female
Functional enrichment
Functional enrichment
Motif enrichment
RcisTarget
Transcription factor binding enrichment
ChIP-enrichR
Expression ratios
<= -2.0
-0
>= 2.0

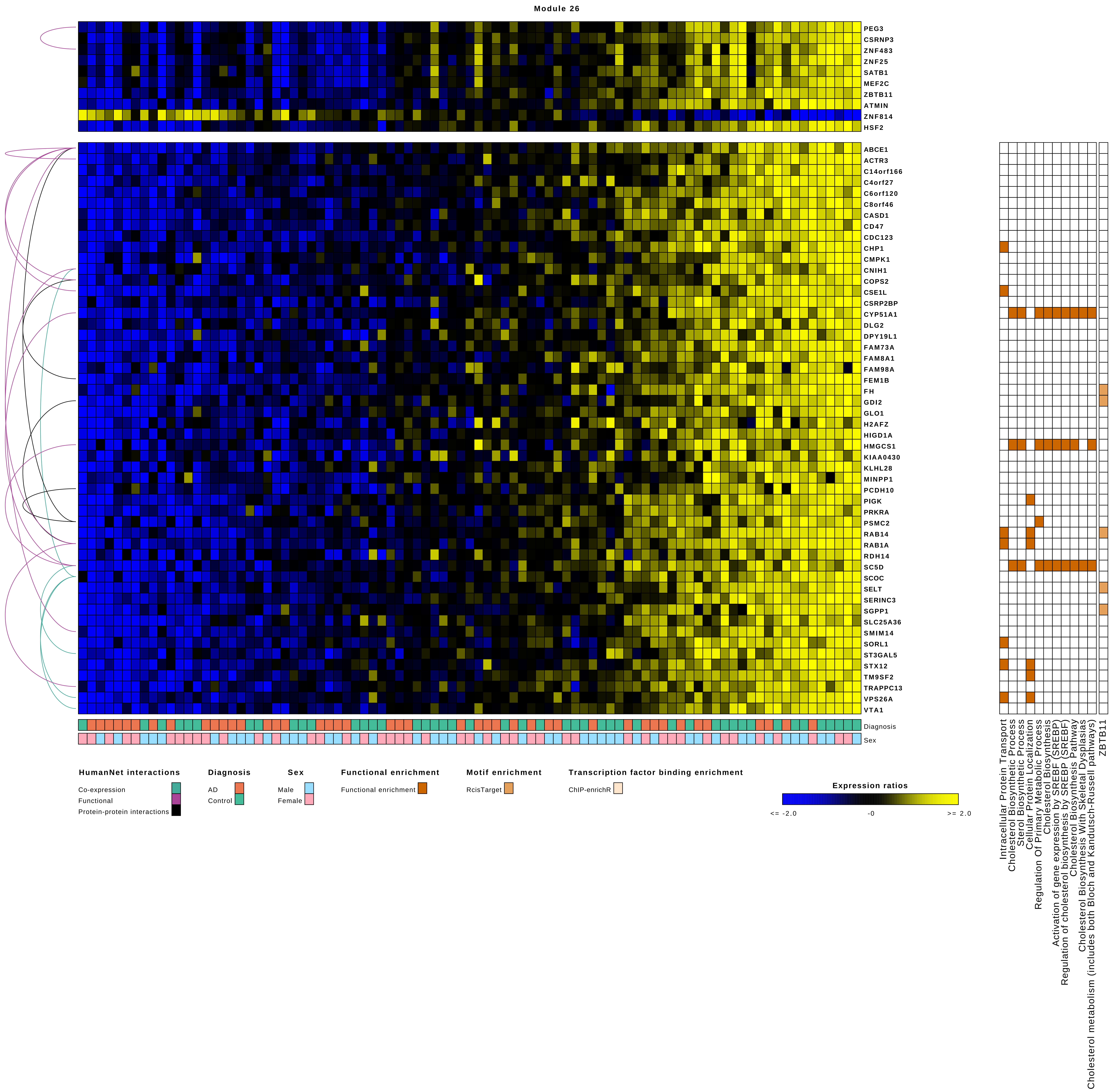
Module 26
PEG3
CSRNP3
ZNF483
ZNF25
SATB1
MEF2C
ZBTB11
ATMIN
ZNF814
HSF2
ABCE1
ACTR3
C14orf166
C4orf27
C6orf120
C8orf46
CASD1
CD47
CDC123
CHP1
CMPK1
CNIH1
COPS2
CSE1L
CSRP2BP
CYP51A1
DLG2
DPY19L1
FAM73A
FAM8A1
FAM98A
FEM1B
FH
GDI2
GLO1
H2AFZ
HIGD1A
HMGCS1
KIAA0430
KLHL28
MINPP1
PCDH10
PIGK
PRKRA
PSMC2
RAB14
RAB1A
RDH14
SC5D
SCOC
SELT
SERINC3
SGPP1
SLC25A36
SMIM14
SORL1
ST3GAL5
STX12
TM9SF2
TRAPPC13
VPS26A
VTA1
Diagnosis
Sex
HumanNet interactions
Co-expression
Functional
Protein-protein interactions
Diagnosis
AD
Control
Sex
Male
Female
Functional enrichment
Functional enrichment
Motif enrichment
RcisTarget
Transcription factor binding enrichment
ChIP-enrichR
Expression ratios
<= -2.0
-0
>= 2.0
Intracellular Protein Transport
Cholesterol Biosynthetic Process
Sterol Biosynthetic Process
Cellular Protein Localization
Regulation Of Primary Metabolic Process
Cholesterol Biosynthesis
Activation of gene expression by SREBF (SREBP)
Regulation of cholesterol biosynthesis by SREBP (SREBF)
Cholesterol Biosynthesis Pathway
Cholesterol Biosynthesis With Skeletal Dysplasias
Cholesterol metabolism (includes both Bloch and Kandutsch-Russell pathways)
ZBTB11

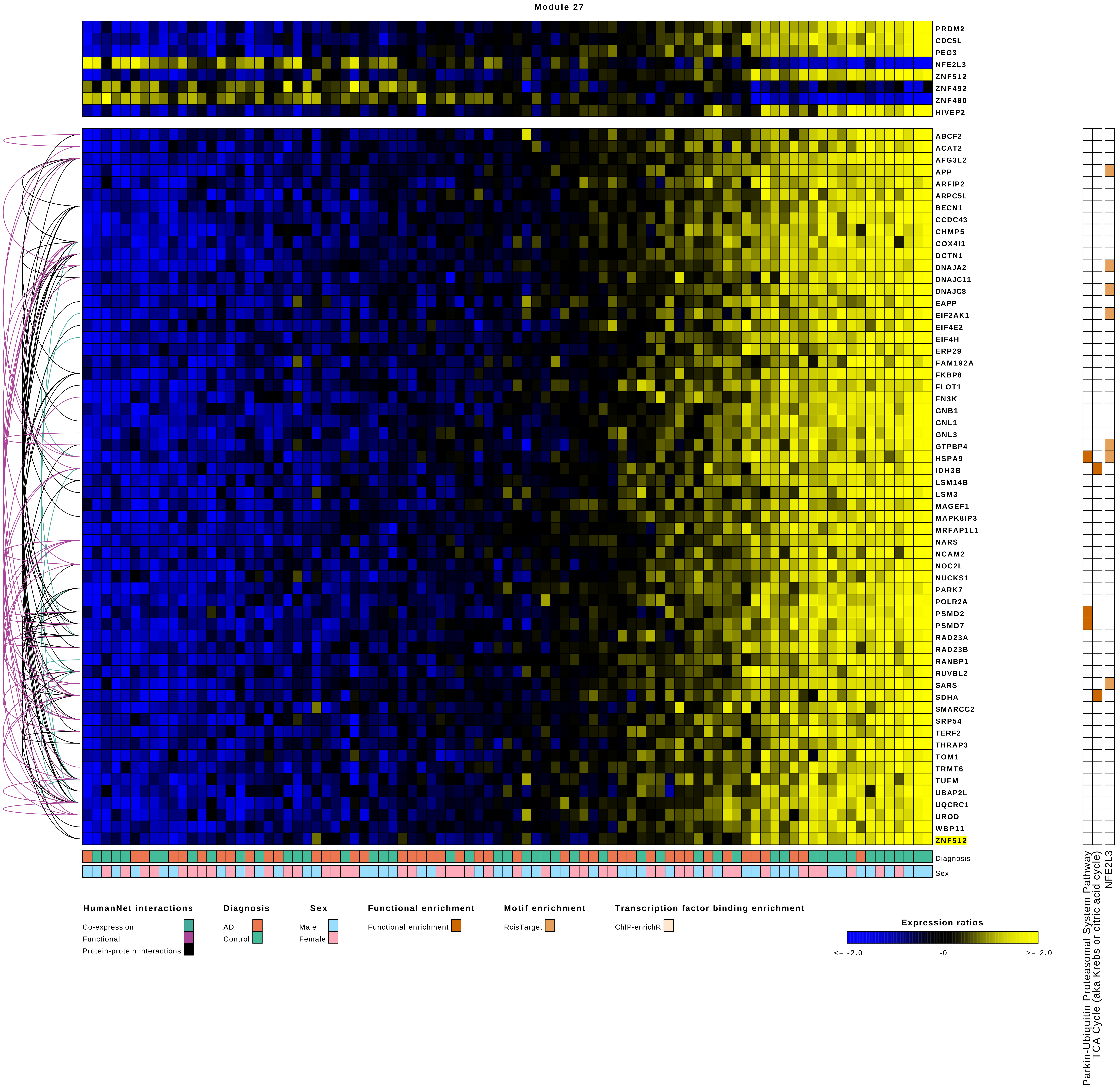

Module 27
PRDM2
CDC5L
PEG3
NFE2L3
ZNF512
ZNF492
ZNF480
HIVEP2
ABCF2
ACAT2
AFG3L2
APP
ARFIP2
ARPC5L
BECN1
CCDC43
CHMP5
COX4I1
DCTN1
DNAJA2
DNAJC11
DNAJC8
EAPP
EIF2AK1
EIF4E2
EIF4H
ERP29
FAM192A
FKBP8
FLOT1
FN3K
GNB1
GNL1
GNL3
GTPBP4
HSPA9
IDH3B
LSM14B
LSM3
MAGEF1
MAPK8IP3
MRFAP1L1
NARS
NCAM2
NOC2L
NUCKS1
PARK7
POLR2A
PSMD2
PSMD7
RAD23A
RAD23B
RANBP1
RUVBL2
SARS
SDHA
SMARCC2
SRP54
TERF2
THRAP3
TOM1
TRMT6
TUFM
UBAP2L
UQCRC1
UROD
WBP11
ZNF512
Diagnosis
Sex
Parkin-Ubiquitin Proteasomal System Pathway
TCA Cycle (aka Krebs or citric acid cycle)
NFE2L3
HumanNet interactions
Co-expression
Functional
Protein-protein interactions
Diagnosis
AD
Control
Sex
Male
Female
Functional enrichment
Functional enrichment
Motif enrichment
RcisTarget
Transcription factor binding enrichment
ChIP-enrichR
Expression ratios
<= -2.0
-0
>= 2.0

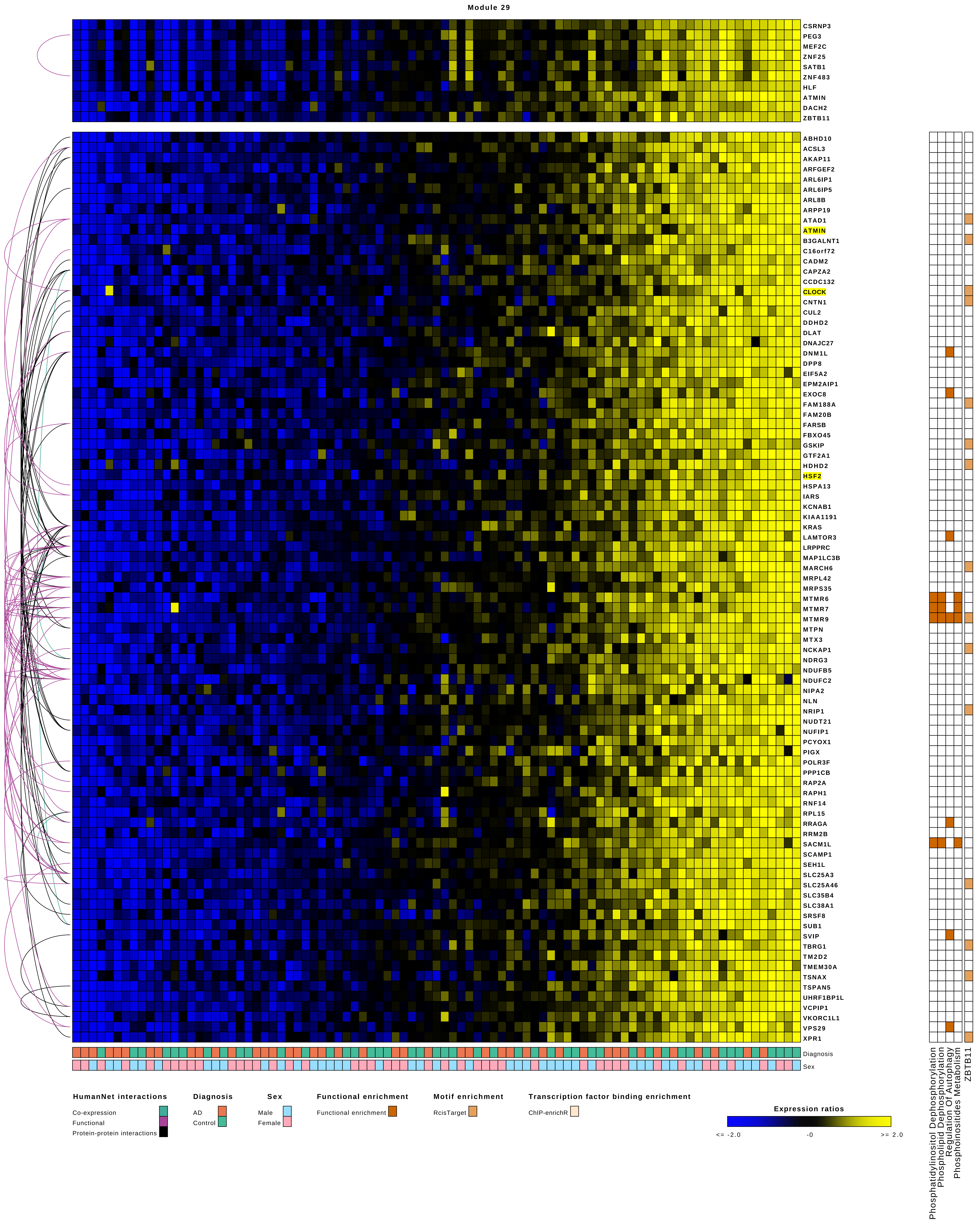
Module 29
CSRNP3
PEG3
MEF2C
ZNF25
SATB1
ZNF483
HLF
ATMIN
DACH2
ZBTB11
ABHD10
ACSL3
AKAP11
ARFGEF2
ARL6IP1
ARL6IP5
ARL8B
ARPP19
ATAD1
ATMIN
B3GALNT1
C16orf72
CADM2
CAPZA2
CCDC132
CLOCK
CNTN1
CUL2
DDHD2
DLAT
DNAJC27
DNM1L
DPP8
EIF5A2
EPM2AIP1
EXOC8
FAM188A
FAM20B
FARSB
FBXO45
GSKIP
GTF2A1
HDHD2
HSF2
HSPA13
IARS
KCNAB1
KIAA1191
KRAS
LAMTOR3
LRPPRC
MAP1LC3B
MARCH6
MRPL42
MRPS35
MTMR6
MTMR7
MTMR9
MTPN
MTX3
NCKAP1
NDRG3
NDUFB5
NDUFC2
NIPA2
NLN
NRIP1
NUDT21
NUFIP1
PCYOX1
PIGX
POLR3F
PPP1CB
RAP2A
RAPH1
RNF14
RPL15
RRAGA
RRM2B
SACM1L
SCAMP1
SEH1L
SLC25A3
SLC25A46
SLC35B4
SLC38A1
SRSF8
SUB1
SVIP
TBRG1
TM2D2
TMEM30A
TSNAX
TSPAN5
UHRF1BP1L
VCPIP1
VKORC1L1
VPS29
XPR1
Diagnosis
Sex
Phosphatidylinositol Dephosphorylation
Phospholipid Dephosphorylation
Regulation Of Autophagy
Phosphoinositides Metabolism
ZBTB11
HumanNet interactions
Co-expression
Functional
Protein-protein interactions
Diagnosis
AD
Control
Sex
Male
Female
Functional enrichment
Functional enrichment
Motif enrichment
RcisTarget
Transcription factor binding enrichment
ChIP-enrichR
Expression ratios
<= -2.0
-0
>= 2.0

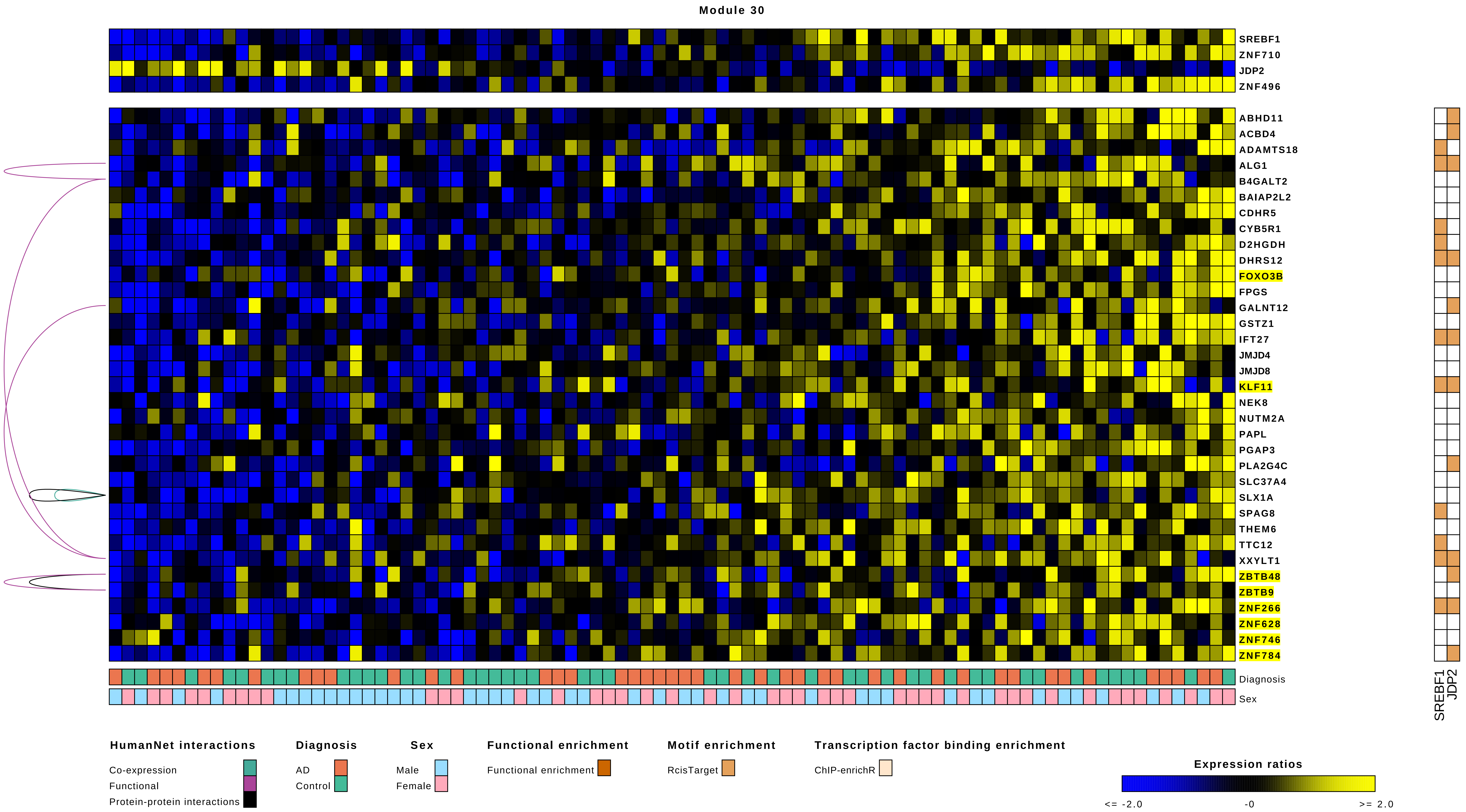
Module 30
SREBF1
ZNF710
JDP2
ZNF496
ABHD11
ACBD4
ADAMTS18
ALG1
B4GALT2
BAIAP2L2
CDHR5
CYB5R1
D2HGDH
DHRS12
FOXO3B
FPGS
GALNT12
GSTZ1
IFT27
JMJD4
JMJD8
KLF11
NEK8
NUTM2A
PAPL
PGAP3
PLA2G4C
SLC37A4
SLX1A
SPAG8
THEM6
TTC12
XXYLT1
ZBTB48
ZBTB9
ZNF266
ZNF628
ZNF746
ZNF784
Diagnosis
Sex
SREBF1
JDP2
HumanNet interactions
Co-expression
Functional
Protein-protein interactions
Diagnosis
AD
Control
Sex
Male
Female
Functional enrichment
Functional enrichment
Motif enrichment
RcisTarget
Transcription factor binding enrichment
ChIP-enrichR
Expression ratios
<= -2.0
-0
>= 2.0

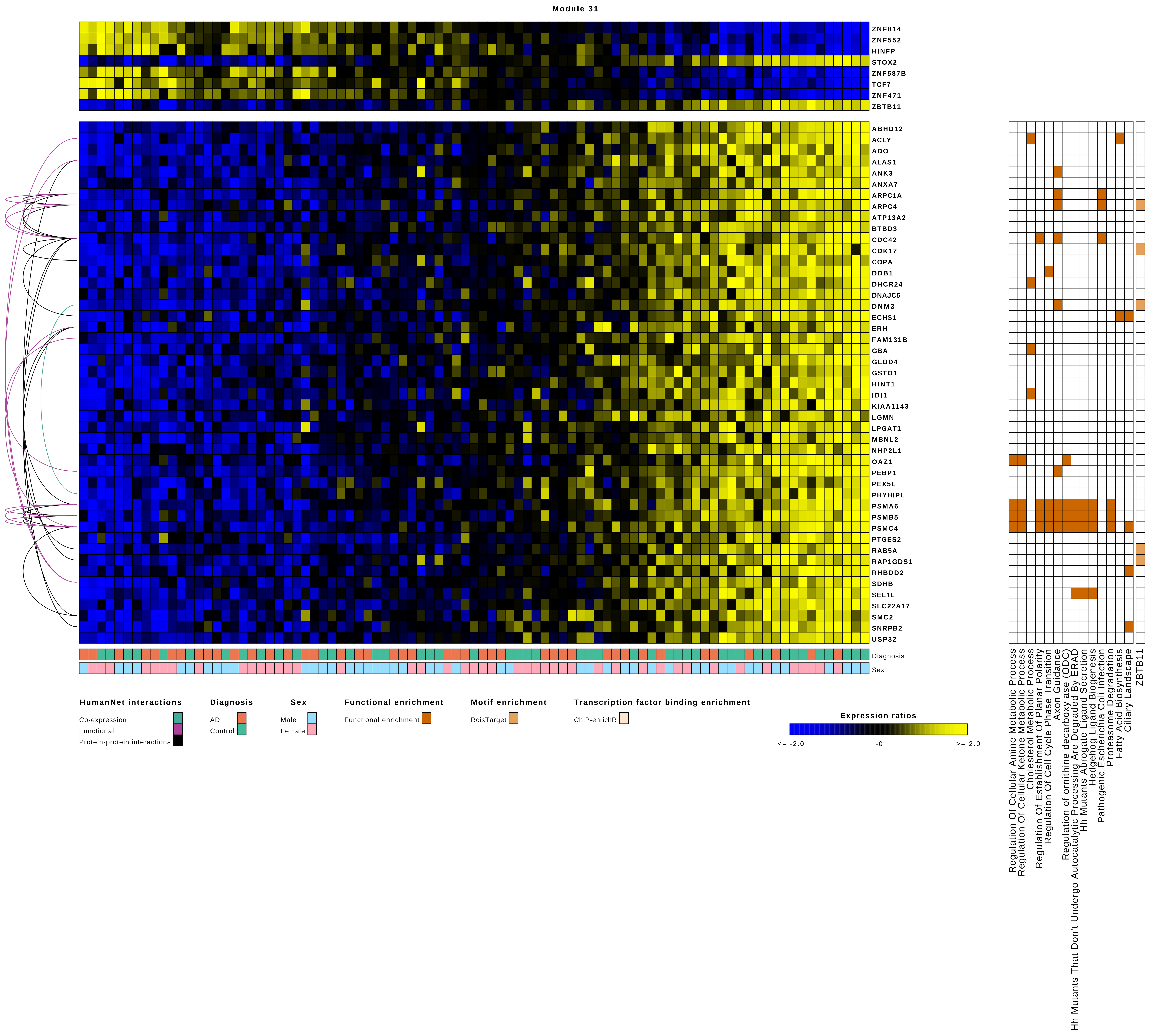
Module 31
ZNF814
ZNF552
HINFP
STOX2
ZNF587B
TCF7
ZNF471
ZBTB11
ABHD12
ACLY
ADO
ALAS1
ANK3
ANXA7
ARPC1A
ARPC4
ATP13A2
BTBD3
CDC42
CDK17
COPA
DDB1
DHCR24
DNAJC5
DNM3
ECHS1
ERH
FAM131B
GBA
GLOD4
GSTO1
HINT1
IDI1
KIAA1143
LGMN
LPGAT1
MBNL2
NHP2L1
OAZ1
PEBP1
PEX5L
PHYHIPL
PSMA6
PSMB5
PSMC4
PTGES2
RAB5A
RAP1GDS1
RHBDD2
SDHB
SEL1L
SLC22A17
SMC2
SNRPB2
USP32
Diagnosis
Sex
Regulation Of Cellular Amine Metabolic Process
Regulation Of Cellular Ketone Metabolic Process
Cholesterol Metabolic Process
Regulation Of Establishment Of Planar Polarity
Regulation Of Cell Cycle Phase Transition
Axon Guidance
Regulation of ornithine decarboxylase (ODC)
Hh Mutants That Don't Undergo Autocatalytic Processing Are Degraded By ERAD
Hh Mutants Abrogate Ligand Secretion
Hedgehog Ligand Biogenesis
Pathogenic Escherichia Coli Infection
Proteasome Degradation
Fatty Acid Biosynthesis
Ciliary Landscape
ZBTB11
HumanNet interactions
Co-expression
Functional
Protein-protein interactions
Diagnosis
AD
Control
Sex
Male
Female
Functional enrichment
Functional enrichment
Motif enrichment
RcisTarget
Transcription factor binding enrichment
ChIP-enrichR
Expression ratios
<= -2.0
-0
>= 2.0

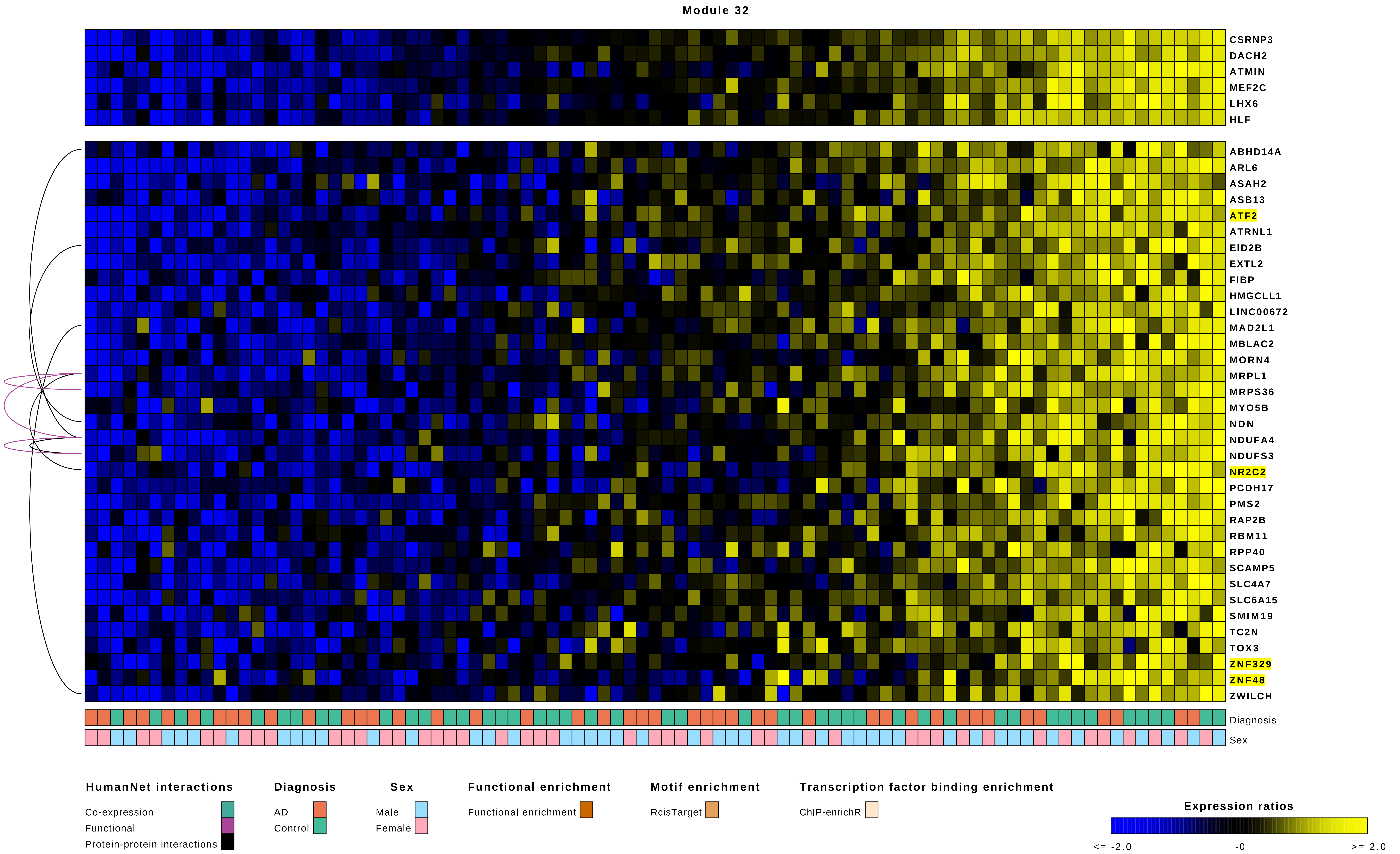

Module 32
CSRNP3
DACH2
ATMIN
MEF2C
LHX6
HLF
ABHD14A
ARL6
ASAH2
ASB13
ATF2
ATRNL1
EID2B
EXTL2
FIBP
HMGCLL1
LINC00672
MAD2L1
MBLAC2
MORN4
MRPL1
MRPS36
MYO5B
NDN
NDUFA4
NDUFS3
NR2C2
PCDH17
PMS2
RAP2B
RBM11
RPP40
SCAMP5
SLC4A7
SLC6A15
SMIM19
TC2N
TOX3
ZNF329
ZNF48
ZWILCH
Diagnosis
Sex
HumanNet interactions
Co-expression
Functional
Protein-protein interactions
Diagnosis
AD
Control
Sex
Male
Female
Functional enrichment
Functional enrichment
Motif enrichment
RcisTarget
Transcription factor binding enrichment
ChIP-enrichR
Expression ratios
<= -2.0
-0
>= 2.0

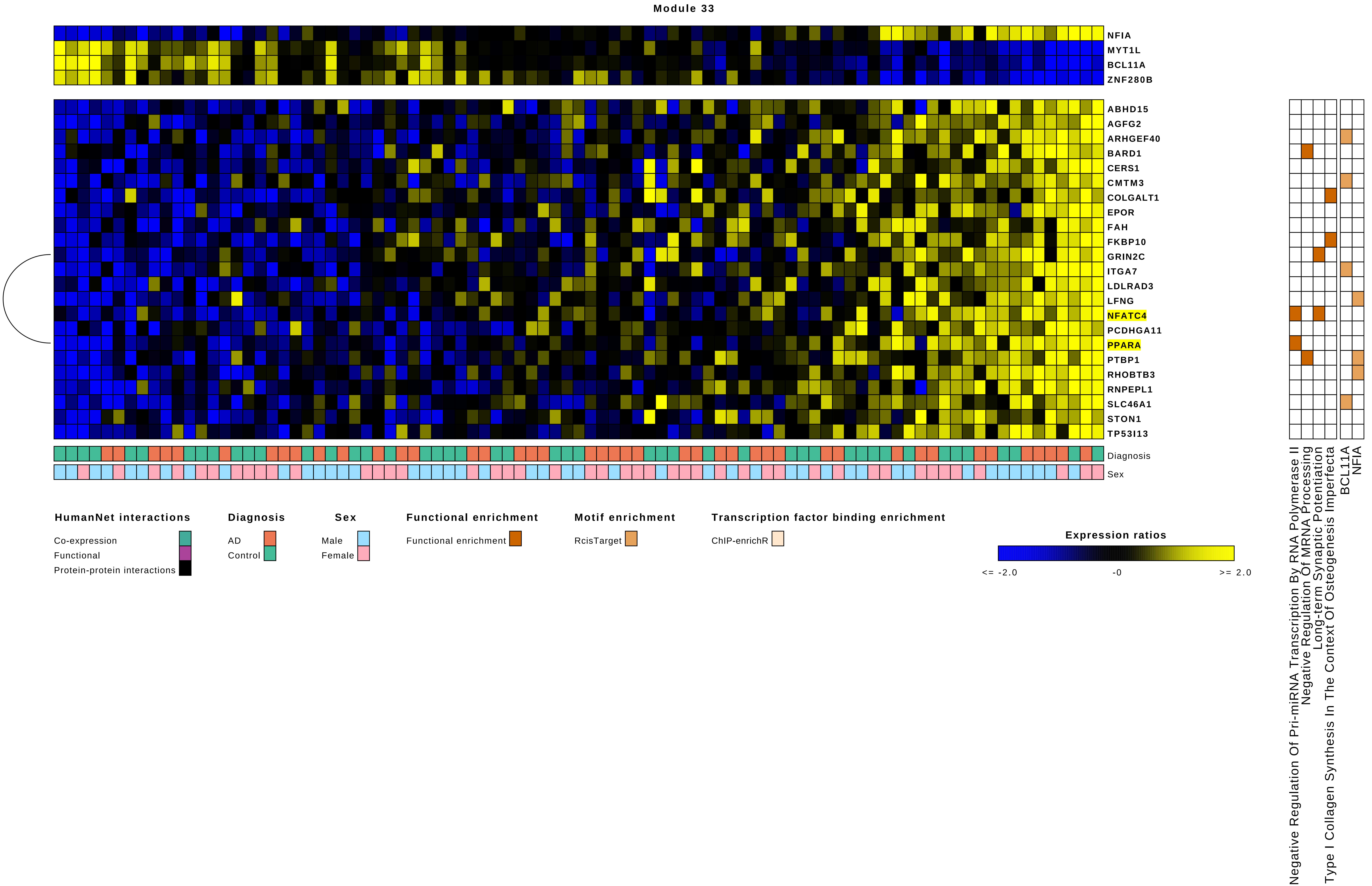

Module 33
NFIA
MYT1L
BCL11A
ZNF280B
ABHD15
AGFG2
ARHGEF40
BARD1
CERS1
CMTM3
COLGALT1
EPOR
FAH
FKBP10
GRIN2C
ITGA7
LDLRAD3
LFNG
NFATC4
PCDHGA11
PPARA
PTBP1
RHOBTB3
RNPEPL1
SLC46A1
STON1
TP53I13
Diagnosis
Sex
HumanNet interactions
Co-expression
Functional
Protein-protein interactions
Diagnosis
AD
Control
Sex
Male
Female
Functional enrichment
Functional enrichment
Motif enrichment
RcisTarget
Transcription factor binding enrichment
ChIP-enrichR
Expression ratios
<= -2.0
-0
>= 2.0
Negative Regulation Of Pri-miRNA Transcription By RNA Polymerase II
Negative Regulation Of MRNA Processing
Long-term Synaptic Potentiation
Type I Collagen Synthesis In The Context Of Osteogenesis Imperfecta
BCL11A
NFIA

Module 34

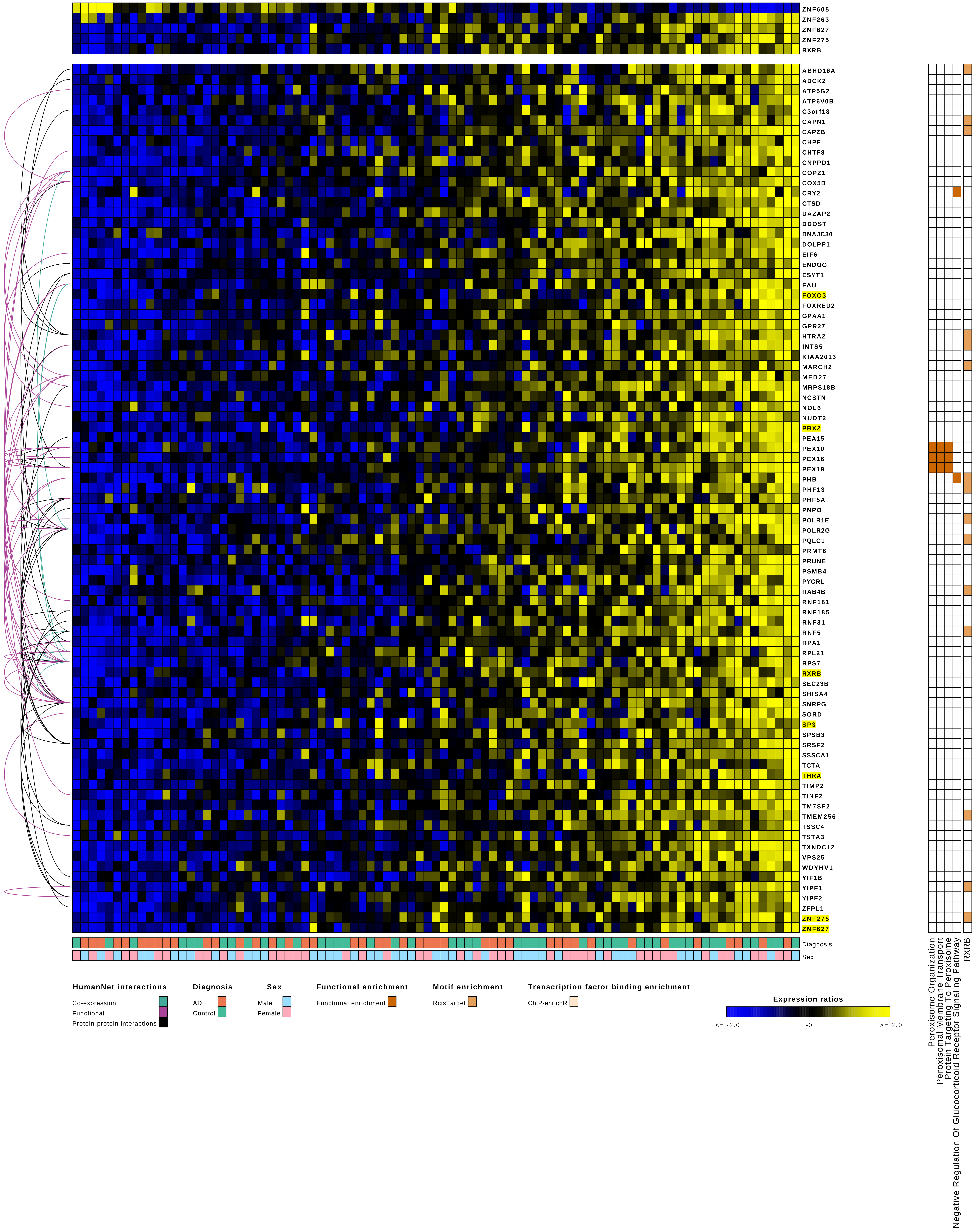

ZNF605
ZNF263
ZNF627
ZNF275
RXRB
ABHD16A
ADCK2
ATP5G2
ATP6V0B
C3orf18
CAPN1
CAPZB
CHPF
CHTF8
CNPPD1
COPZ1
COX5B
CRY2
CTSD
DAZAP2
DDOST
DNAJC30
DOLPP1
EIF6
ENDOG
ESYT1
FAU
FOXO3
FOXRED2
GPAA1
GPR27
HTRA2
INTS5
KIAA2013
MARCH2
MED27
MRPS18B
NCSTN
NOL6
NUDT2
PBX2
PEA15
PEX10
PEX16
PEX19
PHB
PHF13
PHF5A
PNPO
POLR1E
POLR2G
PQLC1
PRMT6
PRUNE
PSMB4
PYCRL
RAB4B
RNF181
RNF185
RNF31
RNF5
RPA1
RPL21
RPS7
RXRB
SEC23B
SHISA4
SNRPG
SORD
SP3
SPSB3
SRSF2
SSSCA1
TCTA
THRA
TIMP2
TINF2
TM7SF2
TMEM256
TSSC4
TSTA3
TXNDC12
VPS25
WDYHV1
YIF1B
YIPF1
YIPF2
ZFPL1
ZNF275
ZNF627
Diagnosis
Sex
Peroxisome Organization
Peroxisomal Membrane Transport
Protein Targeting To Peroxisome
Negative Regulation Of Glucocorticoid Receptor Signaling Pathway
RXRB
HumanNet interactions
Co-expression
Functional
Protein-protein interactions
Diagnosis
AD
Control
Sex
Male
Female
Functional enrichment
Functional enrichment
Motif enrichment
RcisTarget
Transcription factor binding enrichment
ChIP-enrichR
Expression ratios
<= -2.0
-0
>= 2.0

Module 35

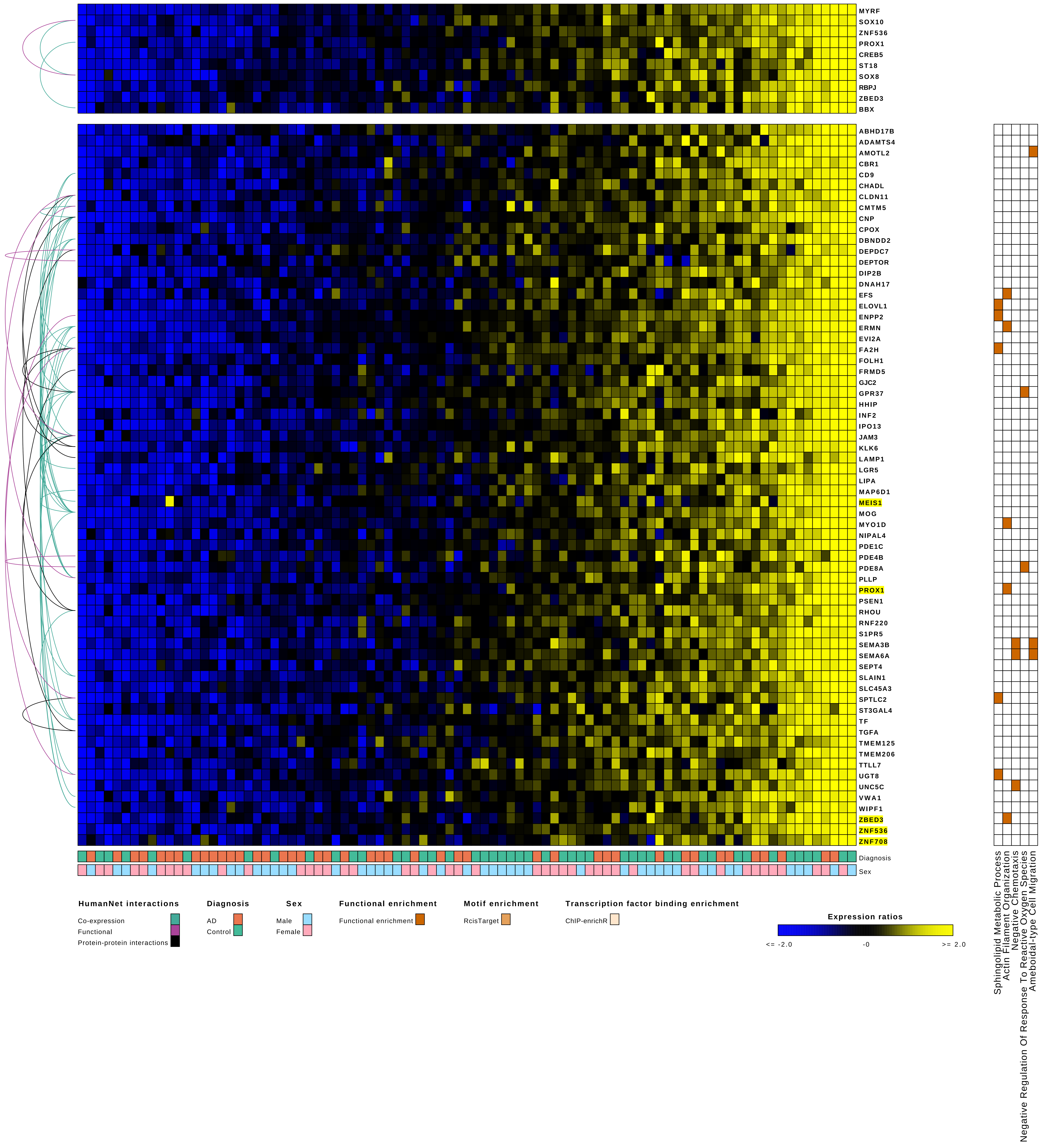
MYRF
SOX10
ZNF536
PROX1
CREB5
ST18
SOX8
RBPJ
ZBED3
BBX
ABHD17B
ADAMTS4
AMOTL2
CBR1
CD9
CHADL
CLDN11
CMTM5
CNP
CPOX
DBNDD2
DEPDC7
DEPTOR
DIP2B
DNAH17
EFS
ELOVL1
ENPP2
ERMN
EVI2A
FA2H
FOLH1
FRMD5
GJC2
GPR37
HHIP
INF2
IPO13
JAM3
KLK6
LAMP1
LGR5
LIPA
MAP6D1
MEIS1
MOG
MYO1D
NIPAL4
PDE1C
PDE4B
PDE8A
PLLP
PROX1
PSEN1
RHOU
RNF220
S1PR5
SEMA3B
SEMA6A
SEPT4
SLAIN1
SLC45A3
SPTLC2
ST3GAL4
TF
TGFA
TMEM125
TMEM206
TTLL7
UGT8
UNC5C
VWA1
WIPF1
ZBED3
ZNF536
ZNF708
Diagnosis
Sex
Sphingolipid Metabolic Process
Actin Filament Organization
Negative Chemotaxis
Negative Regulation Of Response To Reactive Oxygen Species
Ameboidal-type Cell Migration
HumanNet interactions
Co-expression
Functional
Protein-protein interactions
Diagnosis
AD
Control
Sex
Male
Female
Functional enrichment
Functional enrichment
Motif enrichment
RcisTarget
Transcription factor binding enrichment
ChIP-enrichR
Expression ratios
<= -2.0
-0
>= 2.0

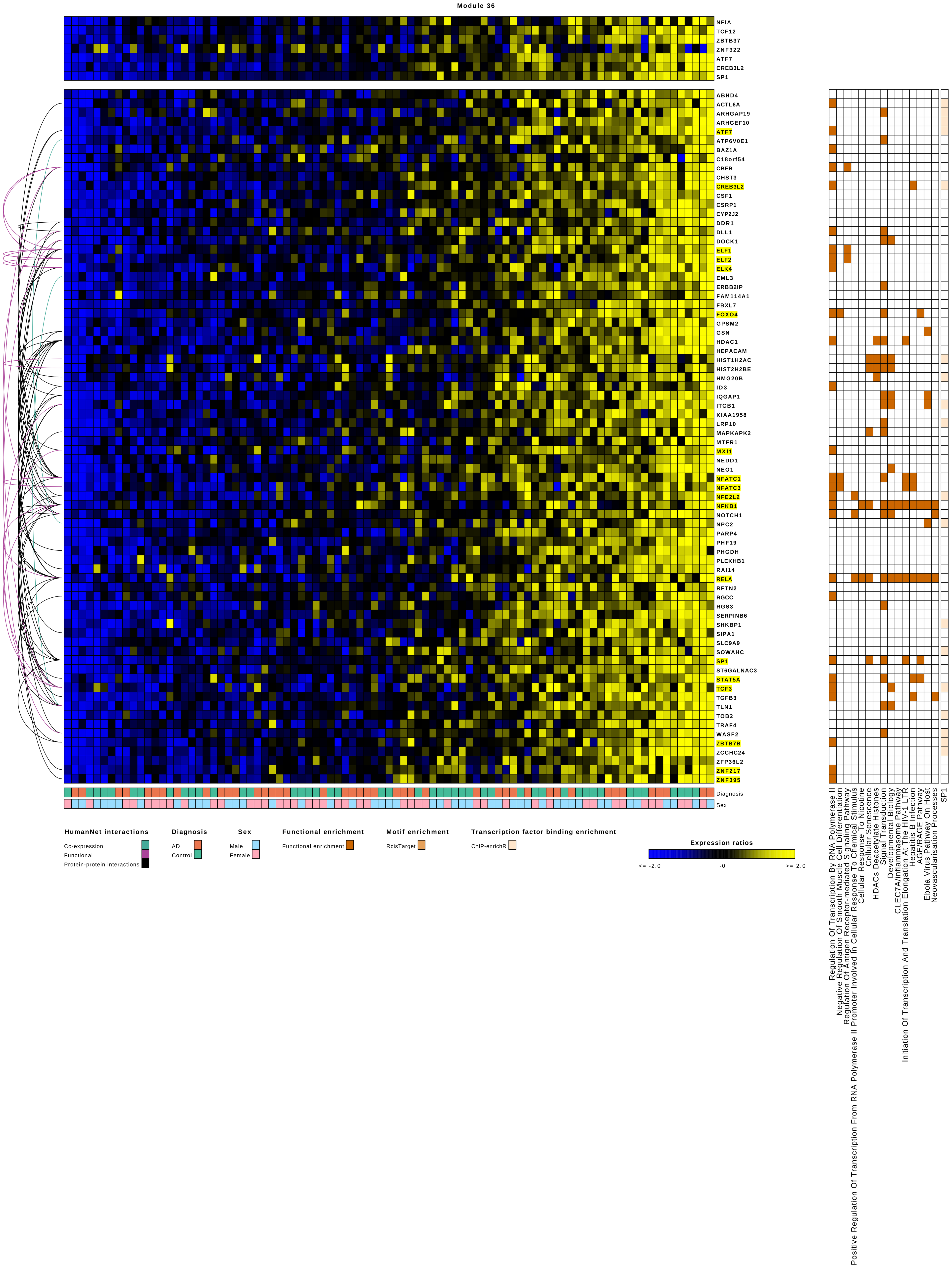
Module 36
NFIA
TCF12
ZBTB37
ZNF322
ATF7
CREB3L2
SP1
ABHD4
ACTL6A
ARHGAP19
ARHGEF10
ATF7
ATP6V0E1
BAZ1A
C18orf54
CBFB
CHST3
CREB3L2
CSF1
CSRP1
CYP2J2
DDR1
DLL1
DOCK1
ELF1
ELF2
ELK4
EML3
ERBB2IP
FAM114A1
FBXL7
FOXO4
GPSM2
GSN
HDAC1
HEPACAM
HIST1H2AC
HIST2H2BE
HMG20B
ID3
IQGAP1
ITGB1
KIAA1958
LRP10
MAPKAPK2
MTFR1
MXI1
NEDD1
NEO1
NFATC1
NFATC3
NFE2L2
NFKB1
NOTCH1
NPC2
PARP4
PHF19
PHGDH
PLEKHB1
RAI14
RELA
RFTN2
RGCC
RGS3
SERPINB6
SHKBP1
SIPA1
SLC9A9
SOWAHC
SP1
ST6GALNAC3
STAT5A
TCF3
TGFB3
TLN1
TOB2
TRAF4
WASF2
ZBTB7B
ZCCHC24
ZFP36L2
ZNF217
ZNF395
Diagnosis
Sex
HumanNet interactions
Co-expression
Functional
Protein-protein interactions
Diagnosis
AD
Control
Sex
Male
Female
Functional enrichment
Functional enrichment
Motif enrichment
RcisTarget
Transcription factor binding enrichment
ChIP-enrichR
Expression ratios
<= -2.0
-0
>= 2.0
Regulation Of Transcription By RNA Polymerase II
Negative Regulation Of Smooth Muscle Cell Differentiation
Regulation Of Antigen Receptor-mediated Signaling Pathway
Positive Regulation Of Transcription From RNA Polymerase II Promoter Involved In Cellular Response To Chemical Stimulus
Cellular Response To Nicotine
Cellular Senescence
HDACs Deacetylate Histones
Signal Transduction
Developmental Biology
CLEC7A/Inflammasome Pathway
Initiation Of Transcription And Translation Elongation At The HIV-1 LTR
Hepatitis B Infection
AGE/RAGE Pathway
Ebola Virus Pathway On Host
Neovascularisation Processes
SP1

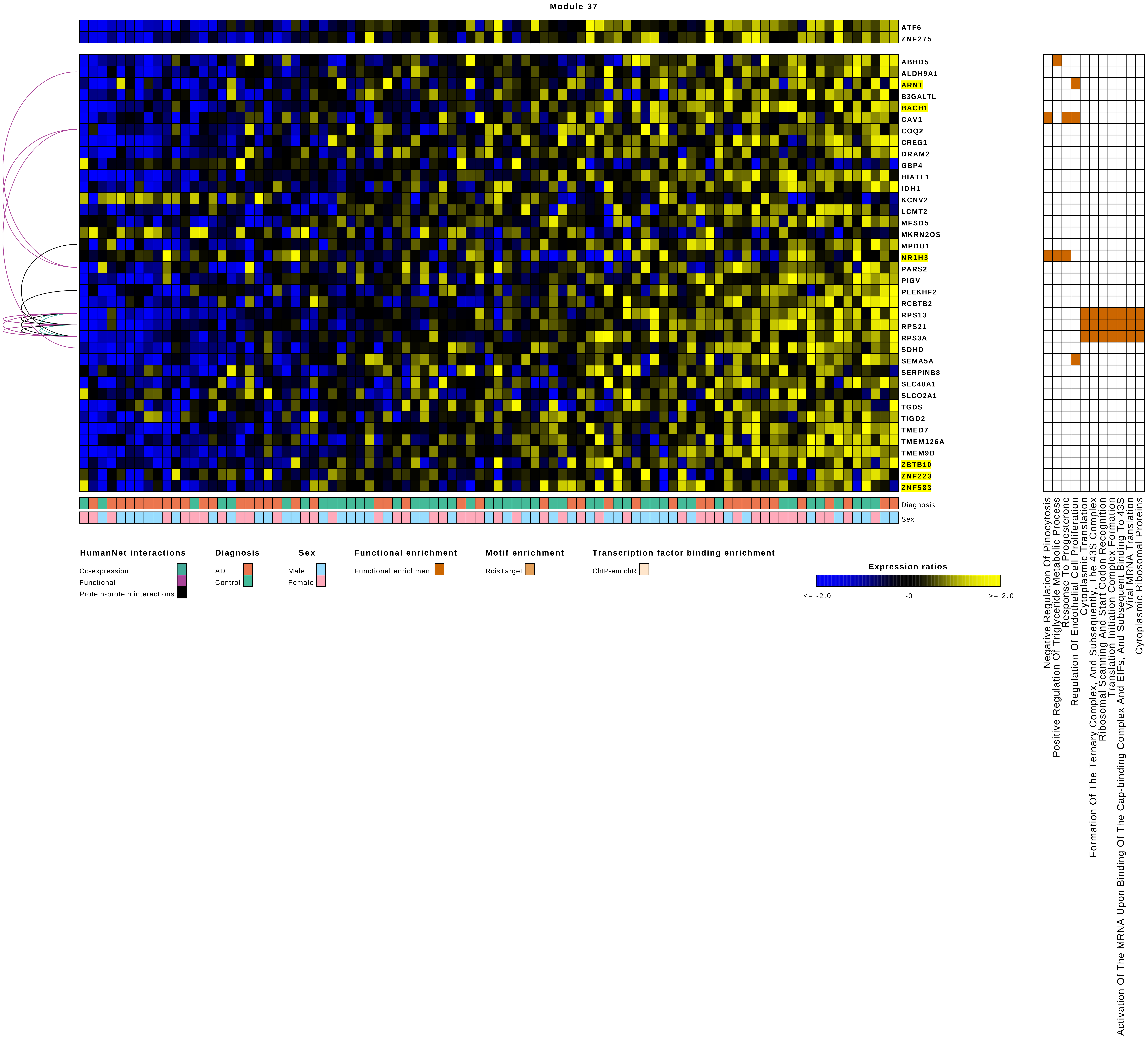
Module 37
ATF6
ZNF275
ABHD5
ALDH9A1
ARNT
B3GALTL
BACH1
CAV1
COQ2
CREG1
DRAM2
GBP4
HIATL1
IDH1
KCNV2
LCMT2
MFSD5
MKRN2OS
MPDU1
NR1H3
PARS2
PIGV
PLEKHF2
RCBTB2
RPS13
RPS21
RPS3A
SDHD
SEMA5A
SERPINB8
SLC40A1
SLCO2A1
TGDS
TIGD2
TMED7
TMEM126A
TMEM9B
ZBTB10
ZNF223
ZNF583
Diagnosis
Sex
HumanNet interactions
Co-expression
Functional
Protein-protein interactions
Diagnosis
AD
Control
Sex
Male
Female
Functional enrichment
Functional enrichment
Motif enrichment
RcisTarget
Transcription factor binding enrichment
ChIP-enrichR
Expression ratios
<= -2.0
-0
>= 2.0
Negative Regulation Of Pinocytosis
Positive Regulation Of Triglyceride Metabolic Process
Response To Progesterone
Regulation Of Endothelial Cell Proliferation
Cytoplasmic Translation
Formation Of The Ternary Complex, And Subsequently, The 43S Complex
Ribosomal Scanning And Start Codon Recognition
Translation Initiation Complex Formation
Activation Of The MRNA Upon Binding Of The Cap-binding Complex And EIFs, And Subsequent Binding To 43S
Viral MRNA Translation
Cytoplasmic Ribosomal Proteins

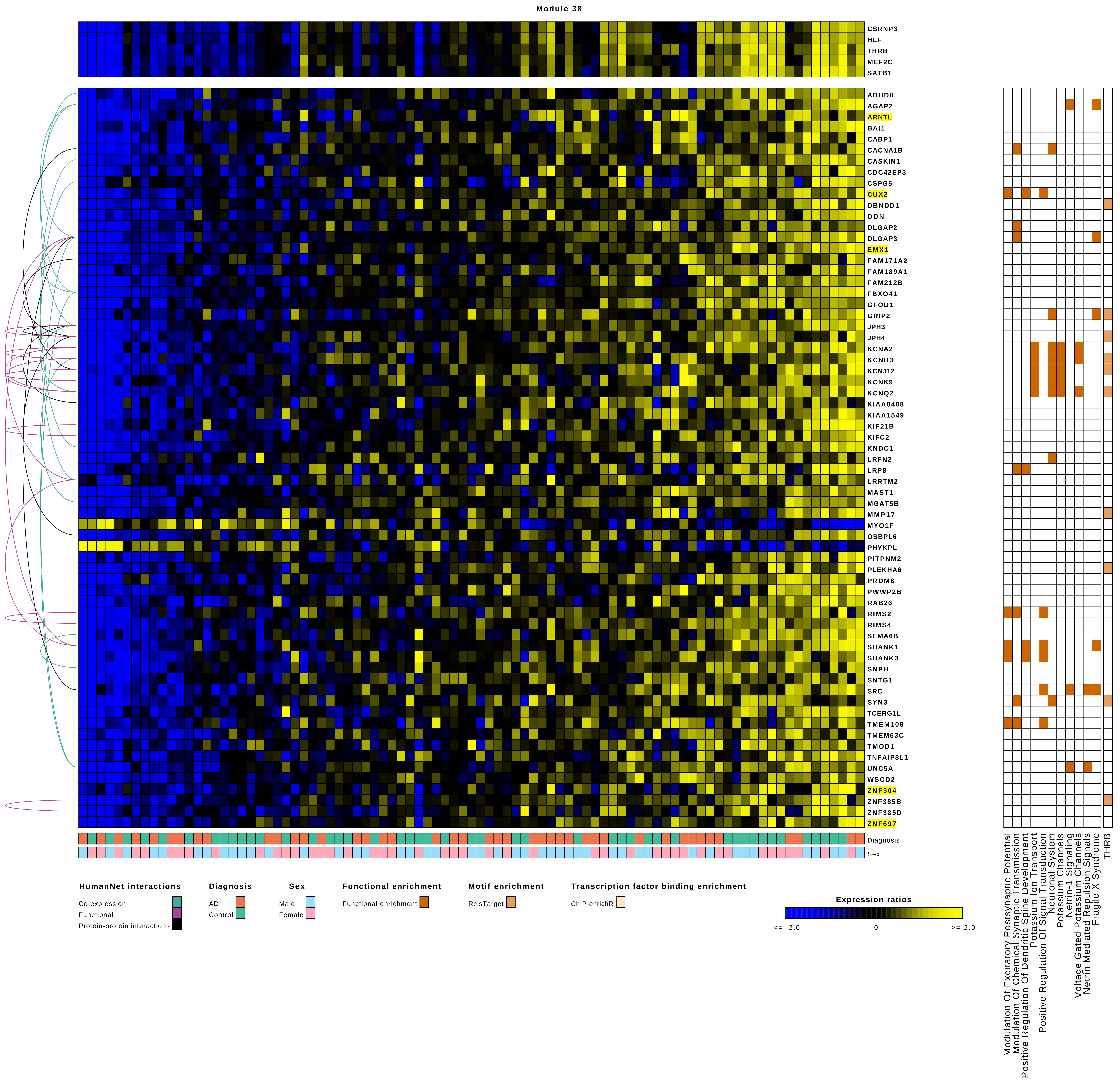

Module 38
CSRNP3
HLF
THRB
MEF2C
SATB1
ABHD8
AGAP2
ARNTL
BAI1
CABP1
CACNA1B
CASKIN1
CDC42EP3
CSPG5
CUX2
DBNDD1
DDN
DLGAP2
DLGAP3
EMX1
FAM171A2
FAM189A1
FAM212B
FBXO41
GFOD1
GRIP2
JPH3
JPH4
KCNA2
KCNH3
KCNJ12
KCNK9
KCNQ2
KIAA0408
KIAA1549
KIF21B
KIFC2
KNDC1
LRFN2
LRP8
LRRTM2
MAST1
MGAT5B
MMP17
MYO1F
OSBPL6
PHYKPL
PITPNM2
PLEKHA6
PRDM8
PWWP2B
RAB26
RIMS2
RIMS4
SEMA6B
SHANK1
SHANK3
SNPH
SNTG1
SRC
SYN3
TCERG1L
TMEM108
TMEM63C
TMOD1
TNFAIP8L1
UNC5A
WSCD2
ZNF304
ZNF385B
ZNF385D
ZNF697
Diagnosis
Sex
Modulation Of Excitatory Postsynaptic Potential
Modulation Of Chemical Synaptic Transmission
Positive Regulation Of Dendritic Spine Development
Potassium Ion Transport
Positive Regulation Of Signal Transduction
Neuronal System
Potassium Channels
Netrin-1 Signaling
Voltage Gated Potassium Channels
Netrin Mediated Repulsion Signals
Fragile X Syndrome
THRB
HumanNet interactions
Co-expression
Functional
Protein-protein interactions
Diagnosis
AD
Control
Sex
Male
Female
Functional enrichment
Functional enrichment
Motif enrichment
RcisTarget
Transcription factor binding enrichment
ChIP-enrichR
Expression ratios
<= -2.0
-0
>= 2.0

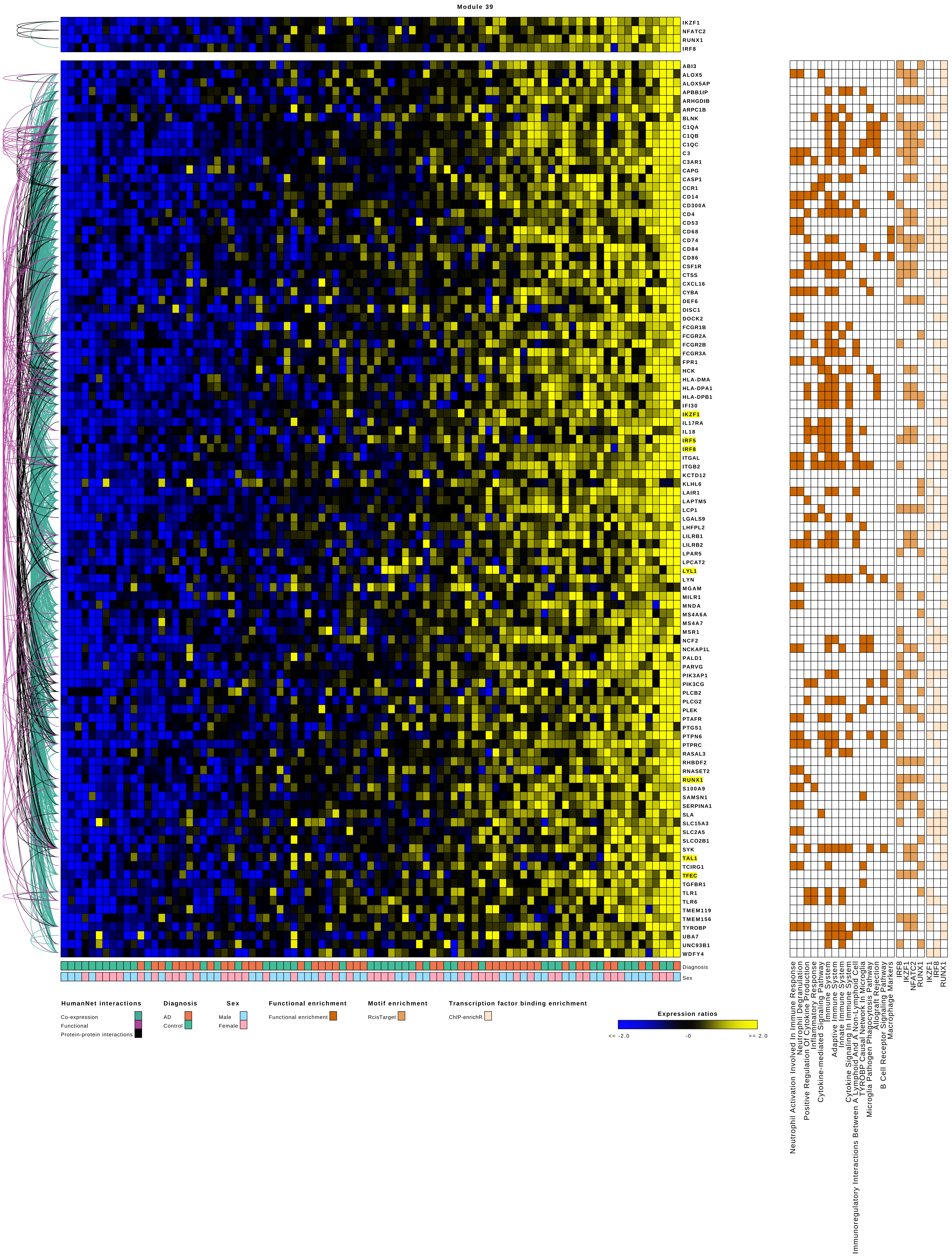

Module 39
IKZF1
NFATC2
RUNX1
IRF8
ABI3
ALOX5
ALOX5AP
APBB1IP
ARHGDIB
ARPC1B
BLNK
C1QA
C1QB
C1QC
C3
C3AR1
CAPG
CASP1
CCR1
CD14
CD300A
CD4
CD53
CD68
CD74
CD84
CD86
CSF1R
CTSS
CXCL16
CYBA
DEF6
DISC1
DOCK2
FCGR1B
FCGR2A
FCGR2B
FCGR3A
FPR1
HCK
HLA-DMA
HLA-DPA1
HLA-DPB1
IFI30
IKZF1
IL17RA
IL18
IRF5
IRF8
ITGAL
ITGB2
KCTD12
KLHL6
LAIR1
LAPTM5
LCP1
LGALS9
LHFPL2
LILRB1
LILRB2
LPAR5
LPCAT2
LYL1
LYN
MGAM
MILR1
MNDA
MS4A6A
MS4A7
MSR1
NCF2
NCKAP1L
PALD1
PARVG
PIK3AP1
PIK3CG
PLCB2
PLCG2
PLEK
PTAFR
PTGS1
PTPN6
PTPRC
RASAL3
RHBDF2
RNASET2
RUNX1
S100A9
SAMSN1
SERPINA1
SLA
SLC15A3
SLC2A5
SLCO2B1
SYK
TAL1
TCIRG1
TFEC
TGFBR1
TLR1
TLR6
TMEM119
TMEM156
TYROBP
UBA7
UNC93B1
WDFY4
Diagnosis
Sex
Neutrophil Activation Involved In Immune Response
Neutrophil Degranulation
Positive Regulation Of Cytokine Production
Inflammatory Response
Cytokine-mediated Signaling Pathway
Immune System
Adaptive Immune System
Innate Immune System
Cytokine Signaling In Immune System
Immunoregulatory Interactions Between A Lymphoid And A Non-Lymphoid Cell
TYROBP Causal Network In Microglia
Microglia Pathogen Phagocytosis Pathway
Allograft Rejection
B Cell Receptor Signaling Pathway
Macrophage Markers
IRF8
IKZF1
NFATC2
RUNX1
IKZF1
IRF8
RUNX1
HumanNet interactions
Co-expression
Functional
Protein-protein interactions
Diagnosis
AD
Control
Sex
Male
Female
Functional enrichment
Functional enrichment
Motif enrichment
RcisTarget
Transcription factor binding enrichment
ChIP-enrichR
Expression ratios
<= -2.0
-0
>= 2.0

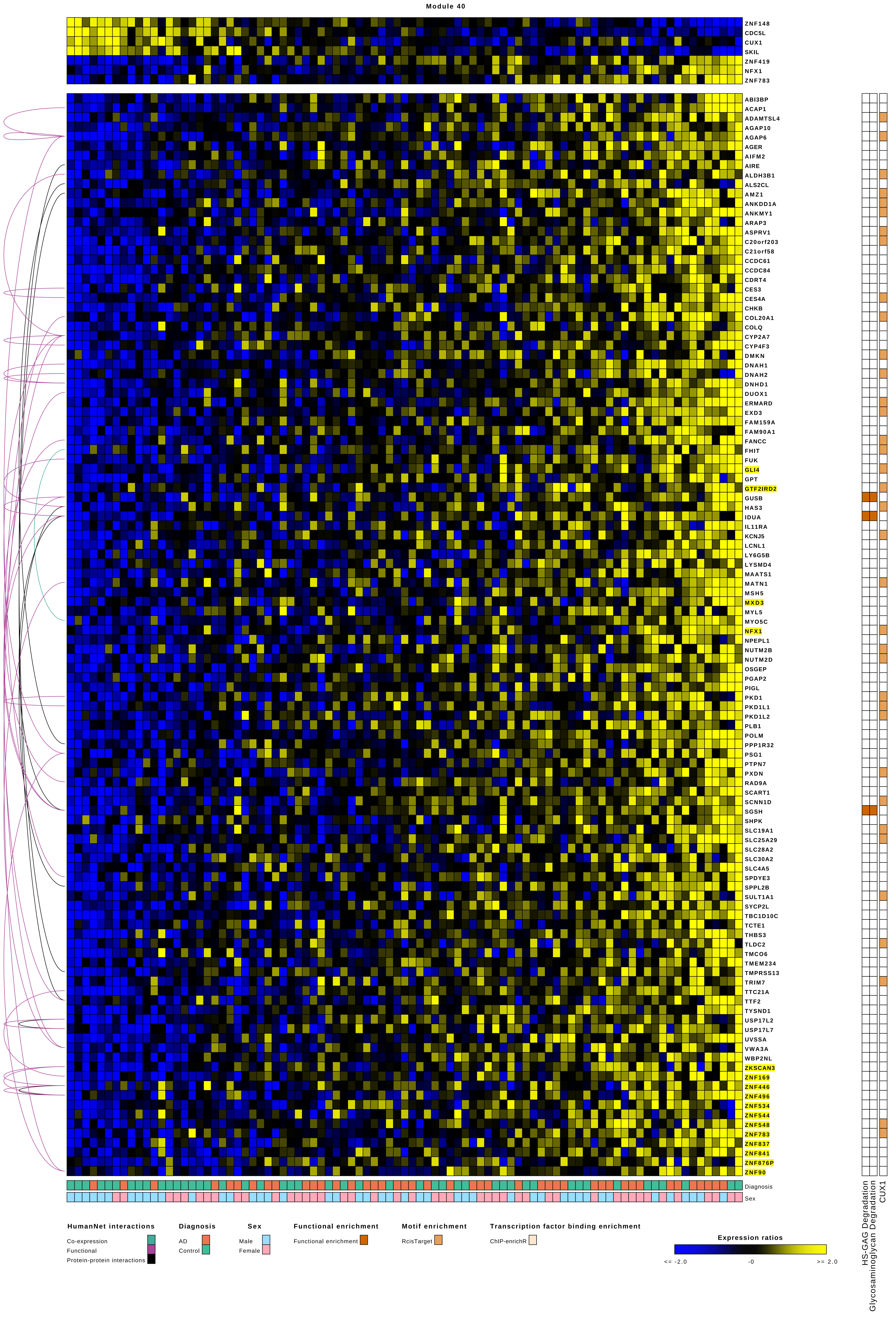

Module 40
ZNF148
CDC5L
CUX1
SKIL
ZNF419
NFX1
ZNF783
ABI3BP
ACAP1
ADAMTSL4
AGAP10
AGAP6
AGER
AIFM2
AIRE
ALDH3B1
ALS2CL
AMZ1
ANKDD1A
ANKMY1
ARAP3
ASPRV1
C20orf203
C21orf58
CCDC61
CCDC84
CDRT4
CES3
CES4A
CHKB
COL20A1
COLQ
CYP2A7
CYP4F3
DMKN
DNAH1
DNAH2
DNHD1
DUOX1
ERMARD
EXD3
FAM159A
FAM90A1
FANCC
FHIT
FUK
GLI4
GPT
GTF2IRD2
GUSB
HAS3
IDUA
IL11RA
KCNJ5
LCNL1
LY6G5B
LYSMD4
MAATS1
MATN1
MSH5
MXD3
MYL5
MYO5C
NFX1
NPEPL1
NUTM2B
NUTM2D
OSGEP
PGAP2
PIGL
PKD1
PKD1L1
PKD1L2
PLB1
POLM
PPP1R32
PSG1
PTPN7
PXDN
RAD9A
SCART1
SCNN1D
SGSH
SHPK
SLC19A1
SLC25A29
SLC28A2
SLC30A2
SLC4A5
SPDYE3
SPPL2B
SULT1A1
SYCP2L
TBC1D10C
TCTE1
THBS3
TLDC2
TMCO6
TMEM234
TMPRSS13
TRIM7
TTC21A
TTF2
TYSND1
USP17L2
USP17L7
UVSSA
VWA3A
WBP2NL
ZKSCAN3
ZNF169
ZNF446
ZNF496
ZNF534
ZNF544
ZNF548
ZNF783
ZNF837
ZNF841
ZNF876P
ZNF90
Diagnosis
Sex
HS-GAG Degradation
Glycosaminoglycan Degradation
CUX1
HumanNet interactions
Co-expression
Functional
Protein-protein interactions
Diagnosis
AD
Control
Sex
Male
Female
Functional enrichment
Functional enrichment
Motif enrichment
RcisTarget
Transcription factor binding enrichment
ChIP-enrichR
Expression ratios
<= -2.0
-0
>= 2.0

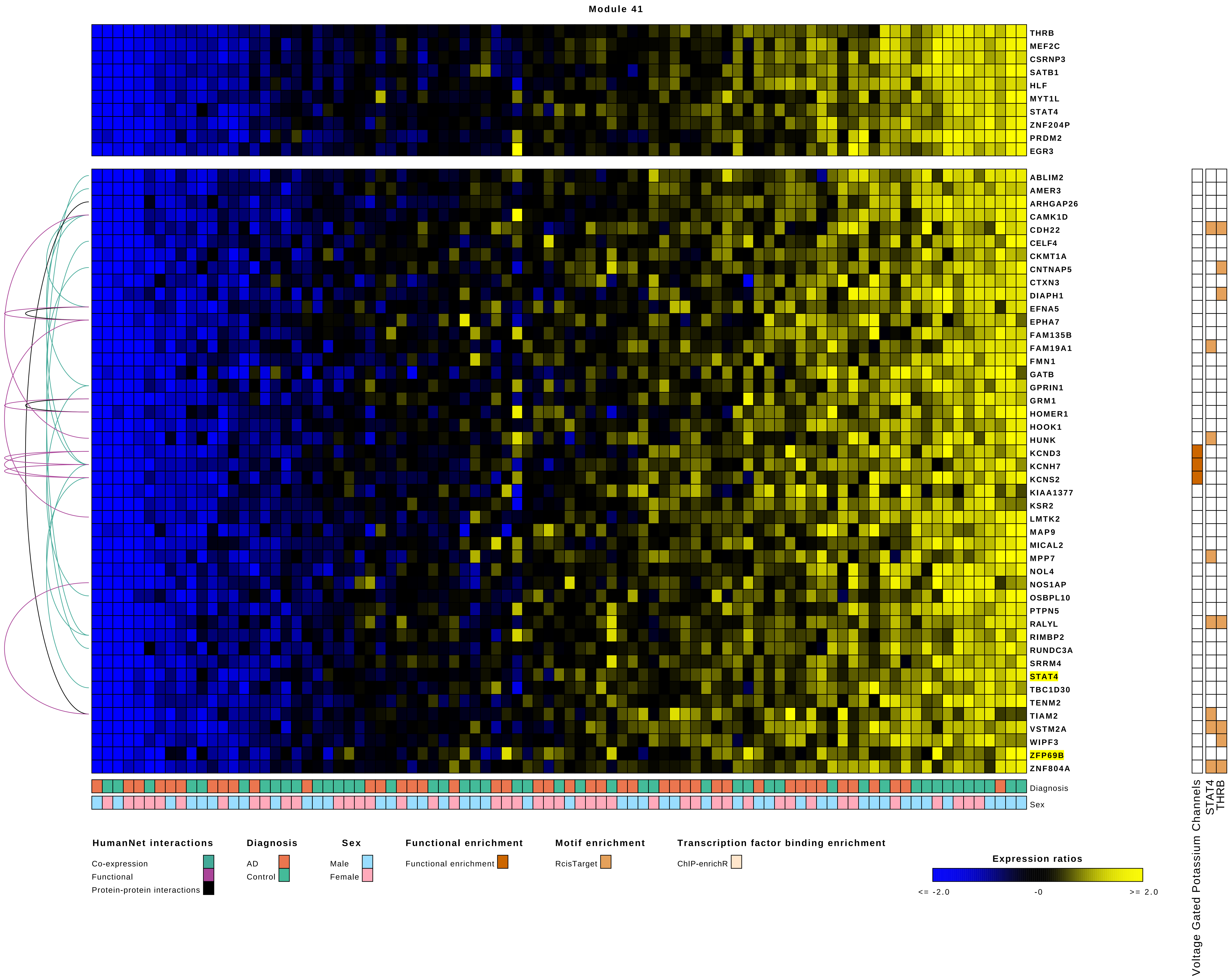
Module 41
THRB
MEF2C
CSRNP3
SATB1
HLF
MYT1L
STAT4
ZNF204P
PRDM2
EGR3
ABLIM2
AMER3
ARHGAP26
CAMK1D
CDH22
CELF4
CKMT1A
CNTNAP5
CTXN3
DIAPH1
EFNA5
EPHA7
FAM135B
FAM19A1
FMN1
GATB
GPRIN1
GRM1
HOMER1
HOOK1
HUNK
KCND3
KCNH7
KCNS2
KIAA1377
KSR2
LMTK2
MAP9
MICAL2
MPP7
NOL4
NOS1AP
OSBPL10
PTPN5
RALYL
RIMBP2
RUNDC3A
SRRM4
STAT4
TBC1D30
TENM2
TIAM2
VSTM2A
WIPF3
ZFP69B
ZNF804A
Diagnosis
Sex
Voltage Gated Potassium Channels
STAT4
THRB
HumanNet interactions
Co-expression
Functional
Protein-protein interactions
Diagnosis
AD
Control
Sex
Male
Female
Functional enrichment
Functional enrichment
Motif enrichment
RcisTarget
Transcription factor binding enrichment
ChIP-enrichR
Expression ratios
<= -2.0
-0
>= 2.0

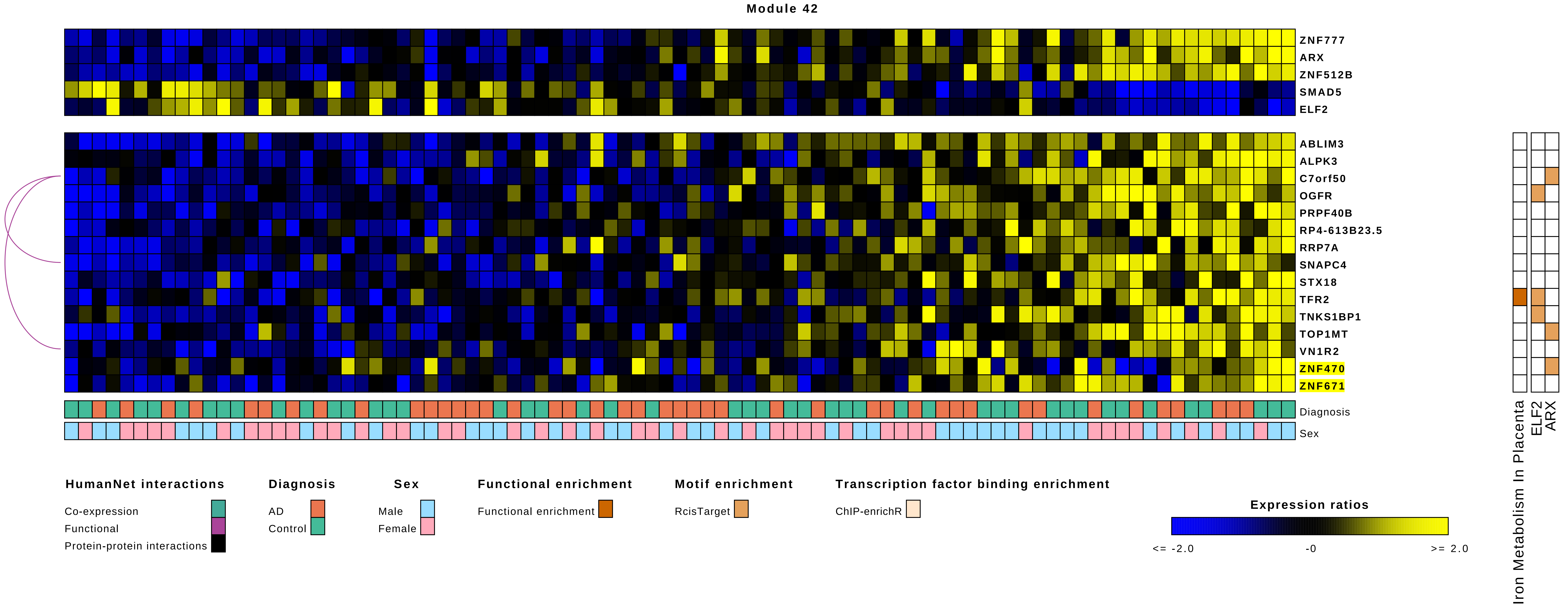

Module 42
ZNF777
ARX
ZNF512B
SMAD5
ELF2
ABLIM3
ALPK3
C7orf50
OGFR
PRPF40B
RP4-613B23.5
RRP7A
SNAPC4
STX18
TFR2
TNKS1BP1
TOP1MT
VN1R2
ZNF470
ZNF671
Diagnosis
Sex
Iron Metabolism In Placenta
ELF2
ARX
HumanNet interactions
Co-expression
Functional
Protein-protein interactions
Diagnosis
AD
Control
Sex
Male
Female
Functional enrichment
Functional enrichment
Motif enrichment
RcisTarget
Transcription factor binding enrichment
ChIP-enrichR
Expression ratios
<= -2.0
-0
>= 2.0

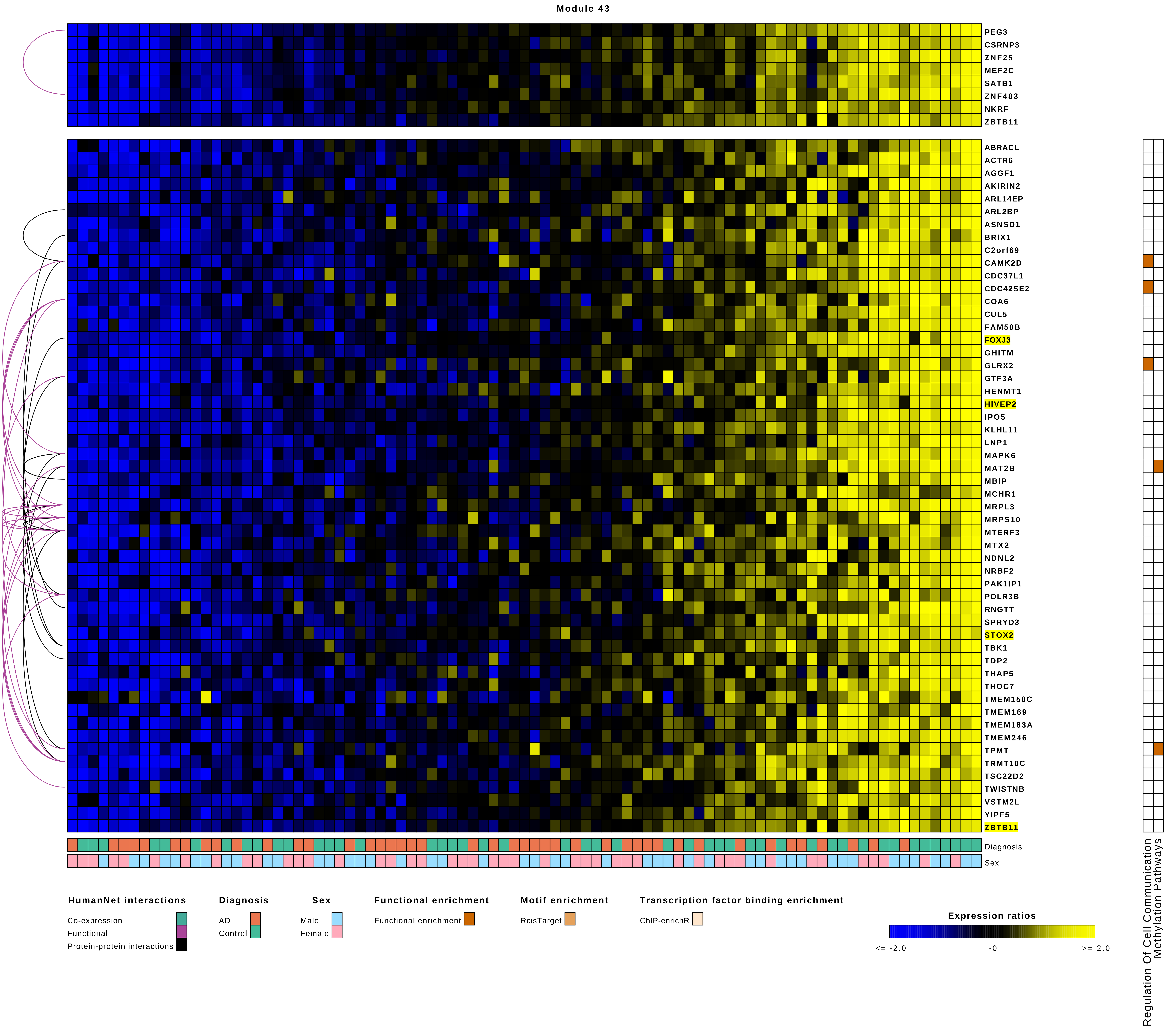
Module 43
PEG3
CSRNP3
ZNF25
MEF2C
SATB1
ZNF483
NKRF
ZBTB11
ABRACL
ACTR6
AGGF1
AKIRIN2
ARL14EP
ARL2BP
ASNSD1
BRIX1
C2orf69
CAMK2D
CDC37L1
CDC42SE2
COA6
CUL5
FAM50B
FOXJ3
GHITM
GLRX2
GTF3A
HENMT1
HIVEP2
IPO5
KLHL11
LNP1
MAPK6
MAT2B
MBIP
MCHR1
MRPL3
MRPS10
MTERF3
MTX2
NDNL2
NRBF2
PAK1IP1
POLR3B
RNGTT
SPRYD3
STOX2
TBK1
TDP2
THAP5
THOC7
TMEM150C
TMEM169
TMEM183A
TMEM246
TPMT
TRMT10C
TSC22D2
TWISTNB
VSTM2L
YIPF5
ZBTB11
Diagnosis
Sex
Regulation Of Cell Communication
Methylation Pathways
HumanNet interactions
Co-expression
Functional
Protein-protein interactions
Diagnosis
AD
Control
Sex
Male
Female
Functional enrichment
Functional enrichment
Motif enrichment
RcisTarget
Transcription factor binding enrichment
ChIP-enrichR
Expression ratios
<= -2.0
-0
>= 2.0

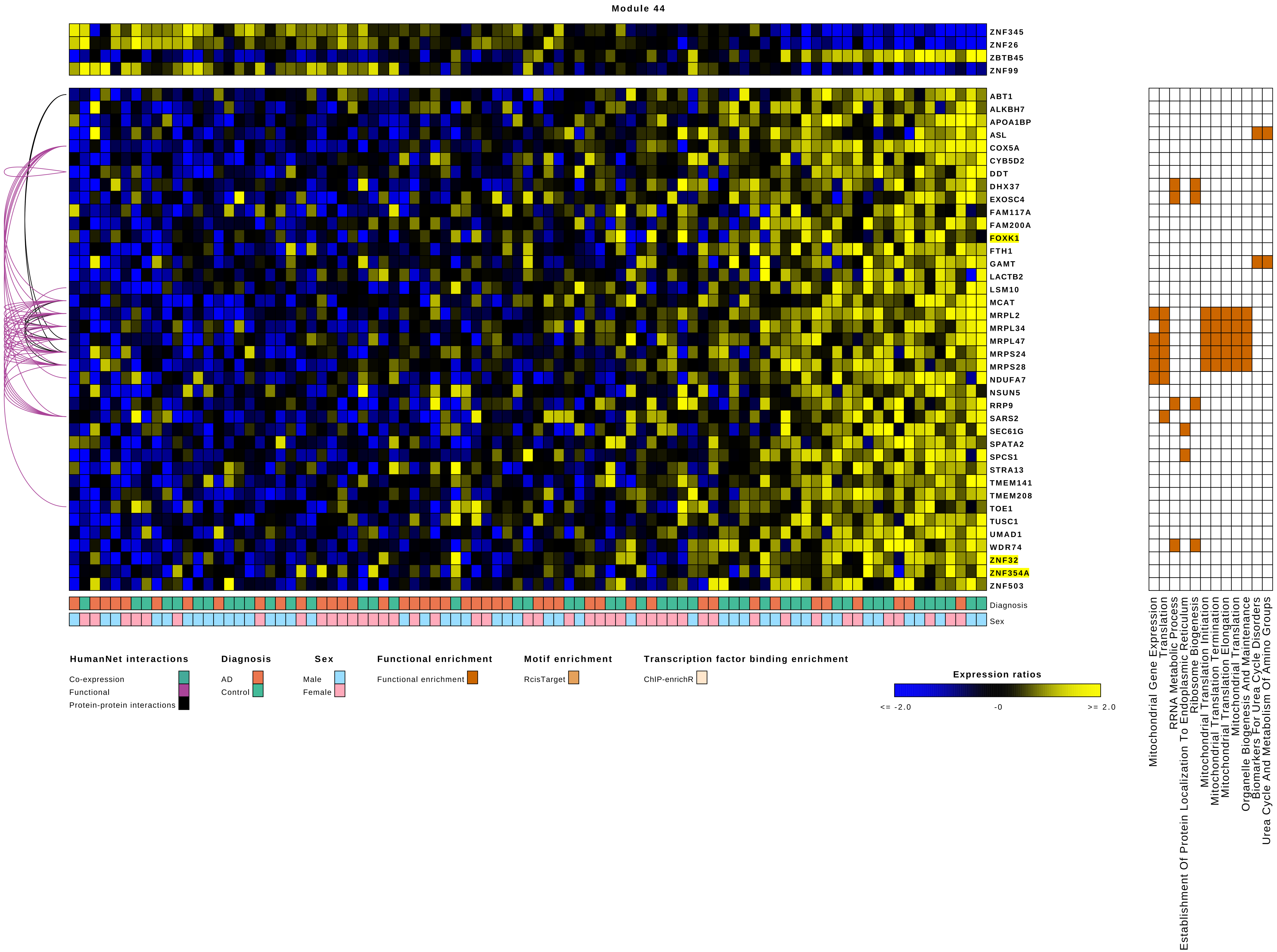
Module 44
ZNF345
ZNF26
ZBTB45
ZNF99
ABT1
ALKBH7
APOA1BP
ASL
COX5A
CYB5D2
DDT
DHX37
EXOSC4
FAM117A
FAM200A
FOXK1
FTH1
GAMT
LACTB2
LSM10
MCAT
MRPL2
MRPL34
MRPL47
MRPS24
MRPS28
NDUFA7
NSUN5
RRP9
SARS2
SEC61G
SPATA2
SPCS1
STRA13
TMEM141
TMEM208
TOE1
TUSC1
UMAD1
WDR74
ZNF32
ZNF354A
ZNF503
Diagnosis
Sex
Mitochondrial Gene Expression
Translation
RRNA Metabolic Process
Establishment Of Protein Localization To Endoplasmic Reticulum
Ribosome Biogenesis
Mitochondrial Translation Initiation
Mitochondrial Translation Termination
Mitochondrial Translation Elongation
Mitochondrial Translation
Organelle Biogenesis And Maintenance
Biomarkers For Urea Cycle Disorders
Urea Cycle And Metabolism Of Amino Groups
HumanNet interactions
Co-expression
Functional
Protein-protein interactions
Diagnosis
AD
Control
Sex
Male
Female
Functional enrichment
Functional enrichment
Motif enrichment
RcisTarget
Transcription factor binding enrichment
ChIP-enrichR
Expression ratios
<= -2.0
-0
>= 2.0

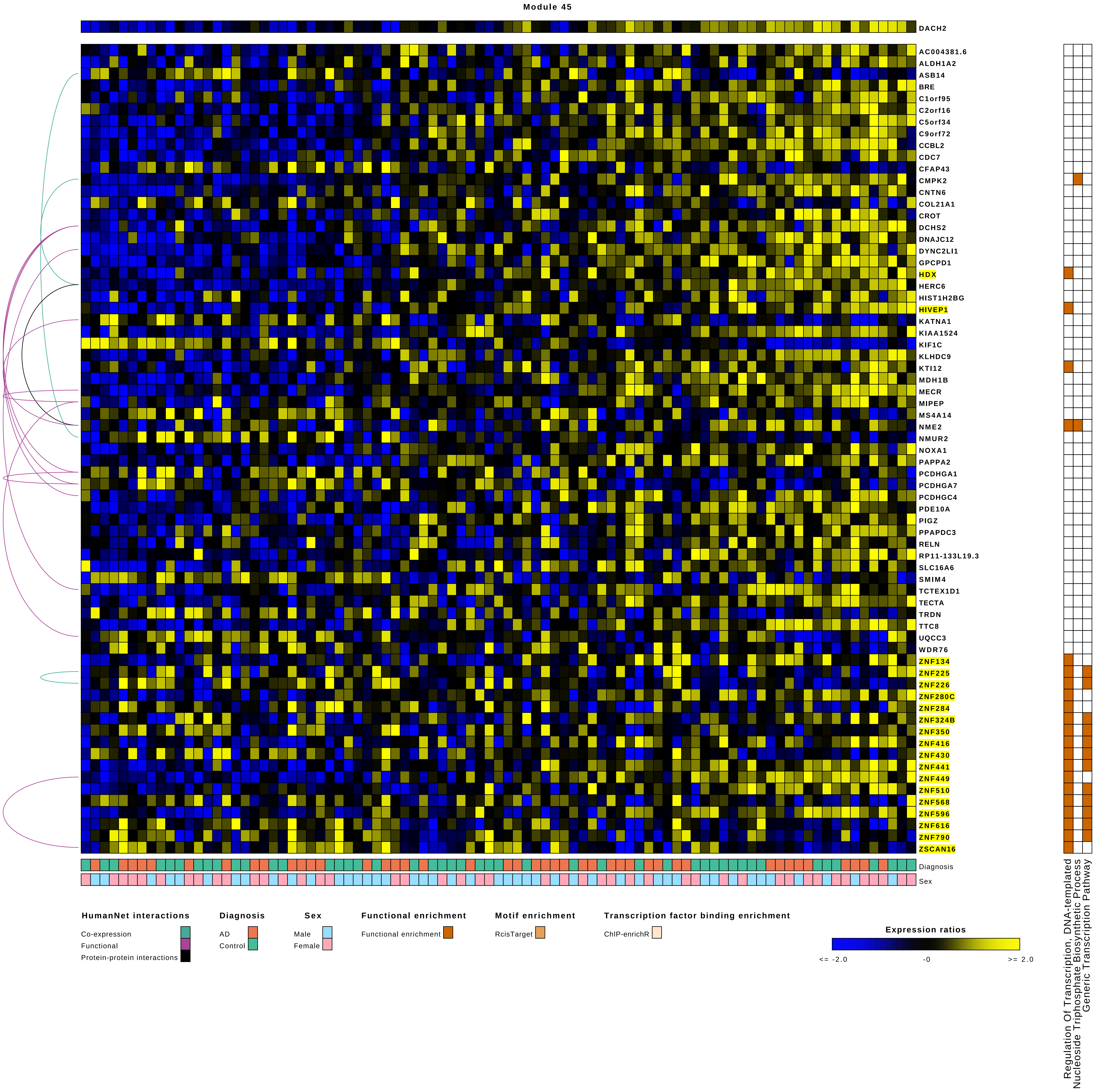
Module 45
DACH2
AC004381.6
ALDH1A2
ASB14
BRE
C1orf95
C2orf16
C5orf34
C9orf72
CCBL2
CDC7
CFAP43
CMPK2
CNTN6
COL21A1
CROT
DCHS2
DNAJC12
DYNC2LI1
GPCPD1
HDX
HERC6
HIST1H2BG
HIVEP1
KATNA1
KIAA1524
KIF1C
KLHDC9
KTI12
MDH1B
MECR
MIPEP
MS4A14
NME2
NMUR2
NOXA1
PAPPA2
PCDHGA1
PCDHGA7
PCDHGC4
PDE10A
PIGZ
PPAPDC3
RELN
RP11-133L19.3
SLC16A6
SMIM4
TCTEX1D1
TECTA
TRDN
TTC8
UQCC3
WDR76
ZNF134
ZNF225
ZNF226
ZNF280C
ZNF284
ZNF324B
ZNF350
ZNF416
ZNF430
ZNF441
ZNF449
ZNF510
ZNF568
ZNF596
ZNF616
ZNF790
ZSCAN16
Diagnosis
Sex
Regulation Of Transcription, DNA-templated
Nucleoside Triphosphate Biosynthetic Process
Generic Transcription Pathway
HumanNet interactions
Co-expression
Functional
Protein-protein interactions
Diagnosis
AD
Control
Sex
Male
Female
Functional enrichment
Functional enrichment
Motif enrichment
RcisTarget
Transcription factor binding enrichment
ChIP-enrichR
Expression ratios
<= -2.0
-0
>= 2.0

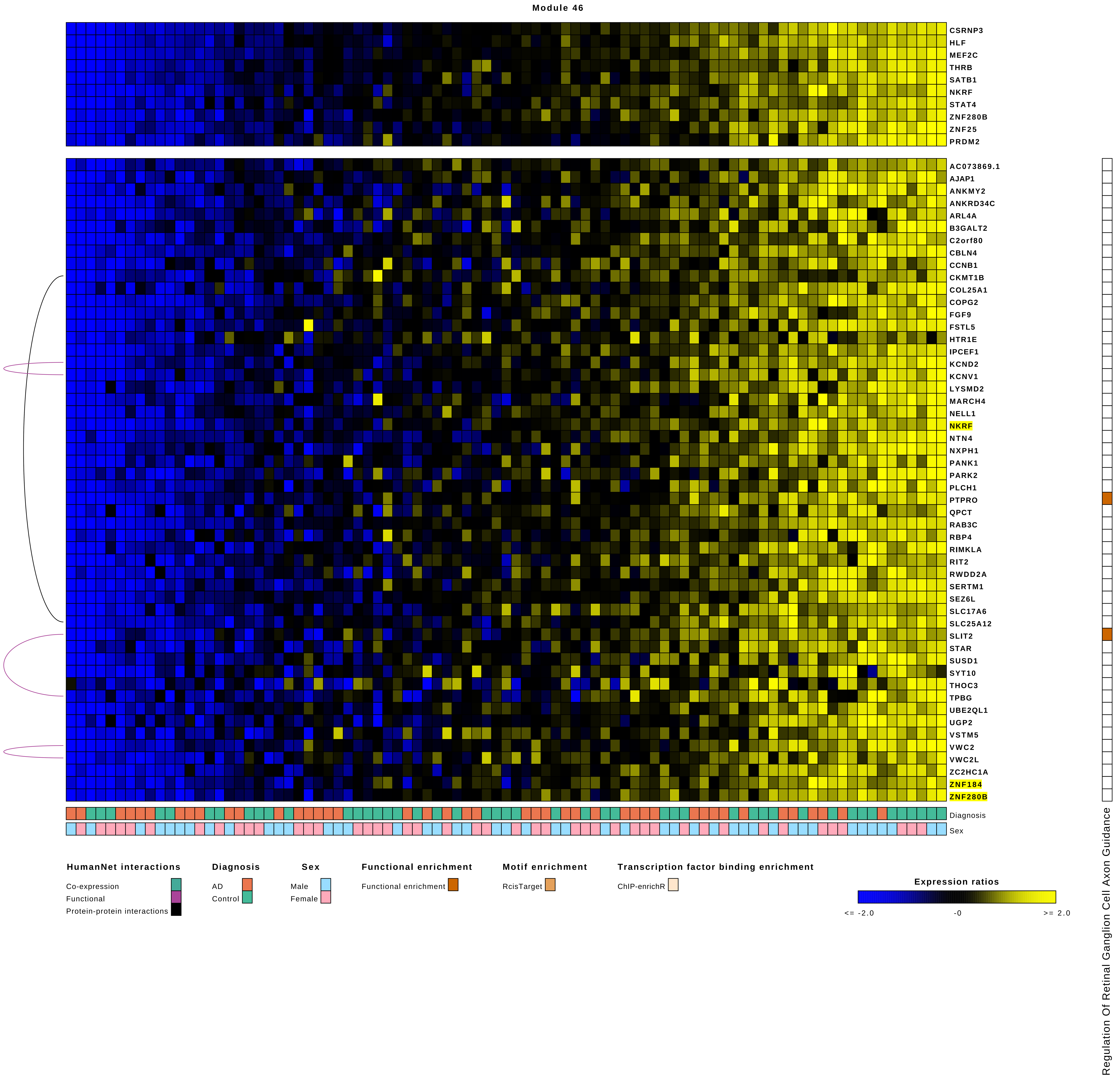
Module 46
CSRNP3
HLF
MEF2C
THRB
SATB1
NKRF
STAT4
ZNF280B
ZNF25
PRDM2
AC073869.1
AJAP1
ANKMY2
ANKRD34C
ARL4A
B3GALT2
C2orf80
CBLN4
CCNB1
CKMT1B
COL25A1
COPG2
FGF9
FSTL5
HTR1E
IPCEF1
KCND2
KCNV1
LYSMD2
MARCH4
NELL1
NKRF
NTN4
NXPH1
PANK1
PARK2
PLCH1
PTPRO
QPCT
RAB3C
RBP4
RIMKLA
RIT2
RWDD2A
SERTM1
SEZ6L
SLC17A6
SLC25A12
SLIT2
STAR
SUSD1
SYT10
THOC3
TPBG
UBE2QL1
UGP2
VSTM5
VWC2
VWC2L
ZC2HC1A
ZNF184
ZNF280B
Diagnosis
Sex
Regulation Of Retinal Ganglion Cell Axon Guidance
HumanNet interactions
Co-expression
Functional
Protein-protein interactions
Diagnosis
AD
Control
Sex
Male
Female
Functional enrichment
Functional enrichment
Motif enrichment
RcisTarget
Transcription factor binding enrichment
ChIP-enrichR
Expression ratios
<= -2.0
-0
>= 2.0

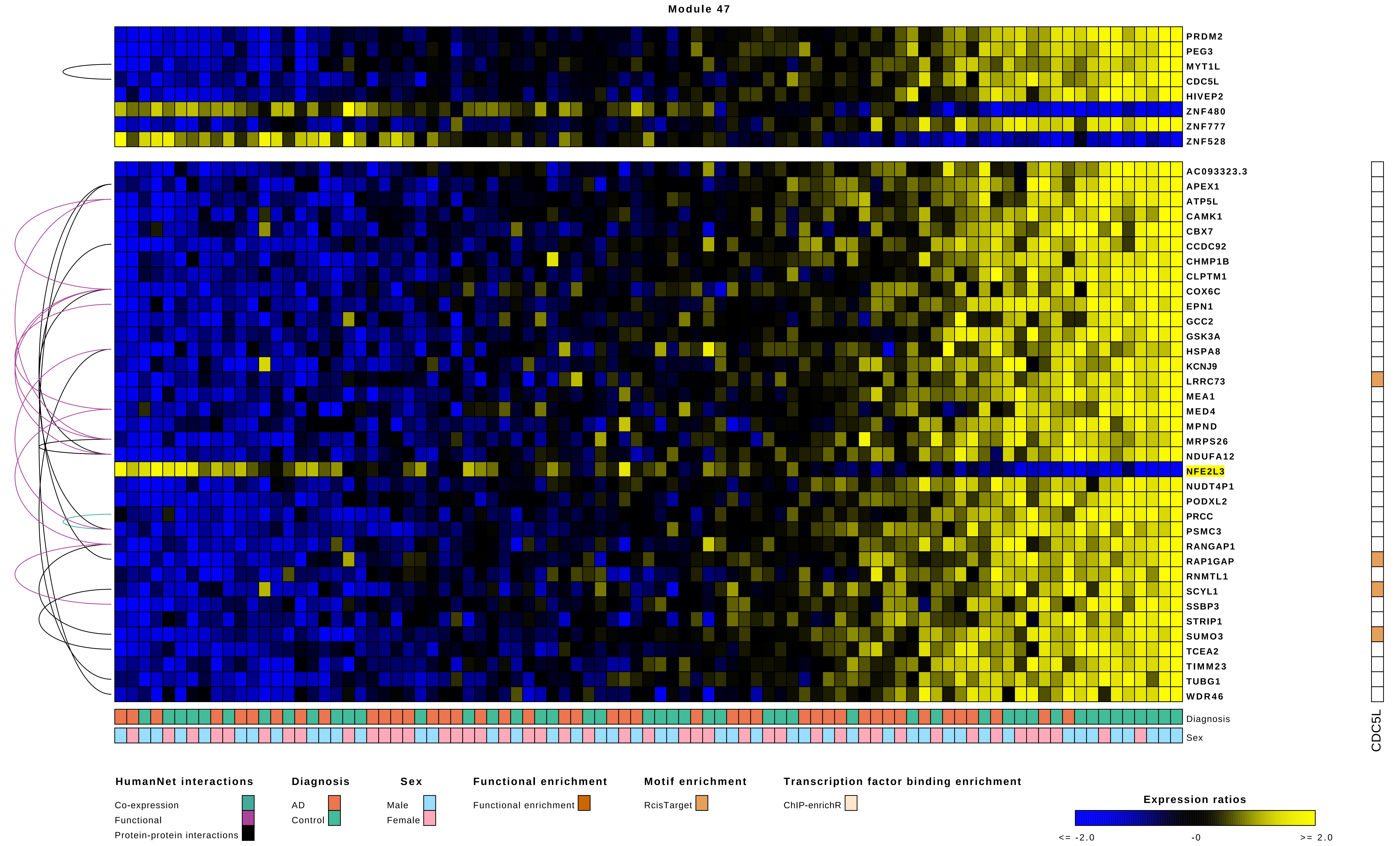

Module 47
PRDM2
PEG3
MYT1L
CDC5L
HIVEP2
ZNF480
ZNF777
ZNF528
AC093323.3
APEX1
ATP5L
CAMK1
CBX7
CCDC92
CHMP1B
CLPTM1
COX6C
EPN1
GCC2
GSK3A
HSPA8
KCNJ9
LRRC73
MEA1
MED4
MPND
MRPS26
NDUFA12
NFE2L3
NUDT4P1
PODXL2
PRCC
PSMC3
RANGAP1
RAP1GAP
RNMTL1
SCYL1
SSBP3
STRIP1
SUMO3
TCEA2
TIMM23
TUBG1
WDR46
Diagnosis
Sex
CDC5L
HumanNet interactions
Co-expression
Functional
Protein-protein interactions
Diagnosis
AD
Control
Sex
Male
Female
Functional enrichment
Functional enrichment
Motif enrichment
RcisTarget
Transcription factor binding enrichment
ChIP-enrichR
Expression ratios
<= -2.0
-0
>= 2.0

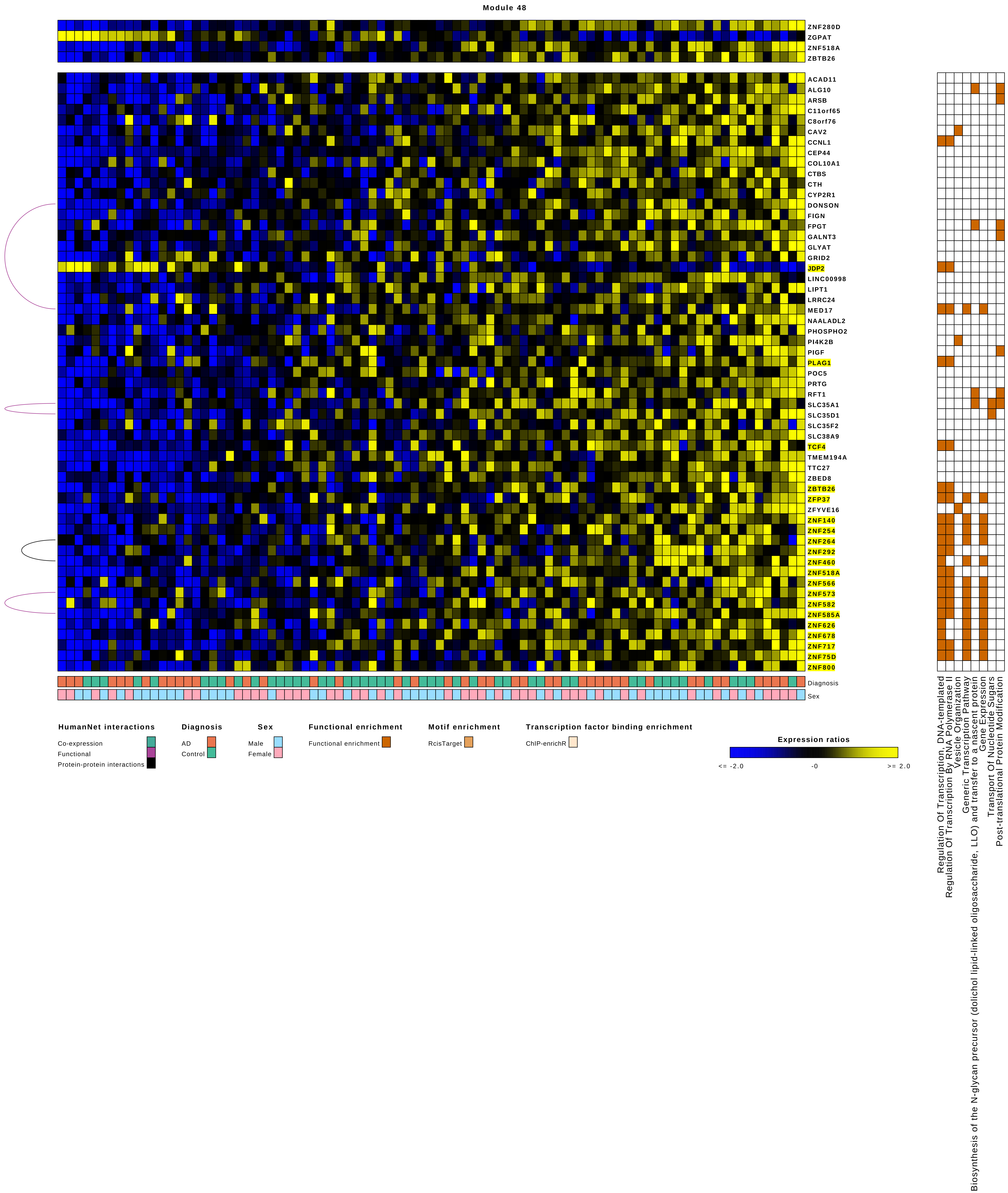
Module 48
ZNF280D
ZGPAT
ZNF518A
ZBTB26
ACAD11
ALG10
ARSB
C11orf65
C8orf76
CAV2
CCNL1
CEP44
COL10A1
CTBS
CTH
CYP2R1
DONSON
FIGN
FPGT
GALNT3
GLYAT
GRID2
JDP2
LINC00998
LIPT1
LRRC24
MED17
NAALADL2
PHOSPHO2
PI4K2B
PIGF
PLAG1
POC5
PRTG
RFT1
SLC35A1
SLC35D1
SLC35F2
SLC38A9
TCF4
TMEM194A
TTC27
ZBED8
ZBTB26
ZFP37
ZFYVE16
ZNF140
ZNF254
ZNF264
ZNF292
ZNF460
ZNF518A
ZNF566
ZNF573
ZNF582
ZNF585A
ZNF626
ZNF678
ZNF717
ZNF75D
ZNF800
Diagnosis
Sex
HumanNet interactions
Co-expression
Functional
Protein-protein interactions
Diagnosis
AD
Control
Sex
Male
Female
Functional enrichment
Functional enrichment
Motif enrichment
RcisTarget
Transcription factor binding enrichment
ChIP-enrichR
Expression ratios
<= -2.0
-0
>= 2.0
Regulation Of Transcription, DNA-templated
Regulation Of Transcription By RNA Polymerase II
Vesicle Organization
Generic Transcription Pathway
Biosynthesis of the N-glycan precursor (dolichol lipid-linked oligosaccharide, LLO) and transfer to a nascent protein
Gene Expression
Transport Of Nucleotide Sugars
Post-translational Protein Modification

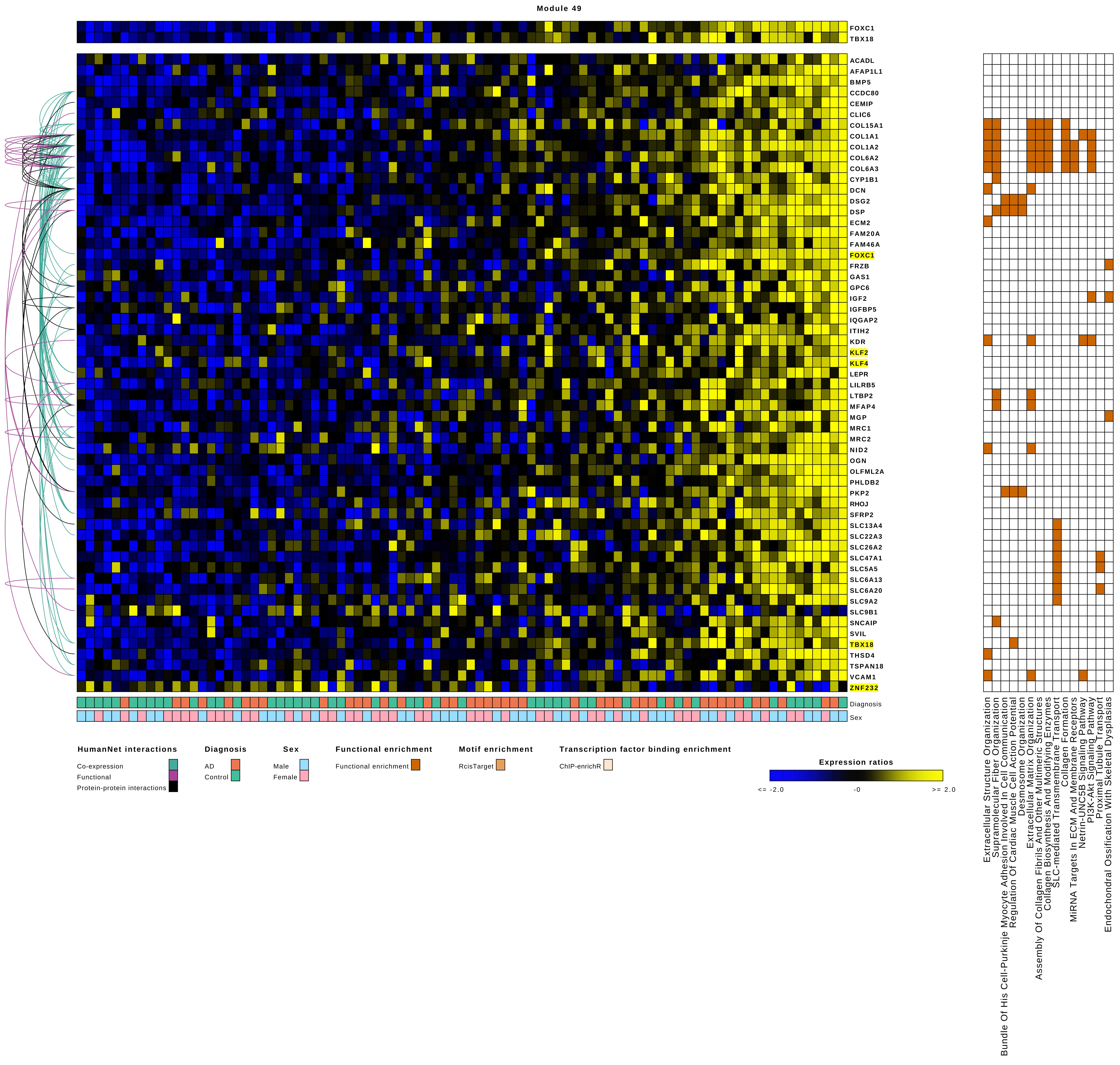
Module 49
FOXC1
TBX18
ACADL
AFAP1L1
BMP5
CCDC80
CEMIP
CLIC6
COL15A1
COL1A1
COL1A2
COL6A2
COL6A3
CYP1B1
DCN
DSG2
DSP
ECM2
FAM20A
FAM46A
FOXC1
FRZB
GAS1
GPC6
IGF2
IGFBP5
IQGAP2
ITIH2
KDR
KLF2
KLF4
LEPR
LILRB5
LTBP2
MFAP4
MGP
MRC1
MRC2
NID2
OGN
OLFML2A
PHLDB2
PKP2
RHOJ
SFRP2
SLC13A4
SLC22A3
SLC26A2
SLC47A1
SLC5A5
SLC6A13
SLC6A20
SLC9A2
SLC9B1
SNCAIP
SVIL
TBX18
THSD4
TSPAN18
VCAM1
ZNF232
Diagnosis
Sex
Extracellular Structure Organization
Supramolecular Fiber Organization
Bundle Of His Cell-Purkinje Myocyte Adhesion Involved In Cell Communication
Regulation Of Cardiac Muscle Cell Action Potential
Desmosome Organization
Extracellular Matrix Organization
Assembly Of Collagen Fibrils And Other Multimeric Structures
Collagen Biosynthesis And Modifying Enzymes
SLC-mediated Transmembrane Transport
Collagen Formation
MiRNA Targets In ECM And Membrane Receptors
Netrin-UNC5B Signaling Pathway
PI3K-Akt Signaling Pathway
Proximal Tubule Transport
Endochondral Ossification With Skeletal Dysplasias
HumanNet interactions
Co-expression
Functional
Protein-protein interactions
Diagnosis
AD
Control
Sex
Male
Female
Functional enrichment
Functional enrichment
Motif enrichment
RcisTarget
Transcription factor binding enrichment
ChIP-enrichR
Expression ratios
<= -2.0
-0
>= 2.0

Module 50

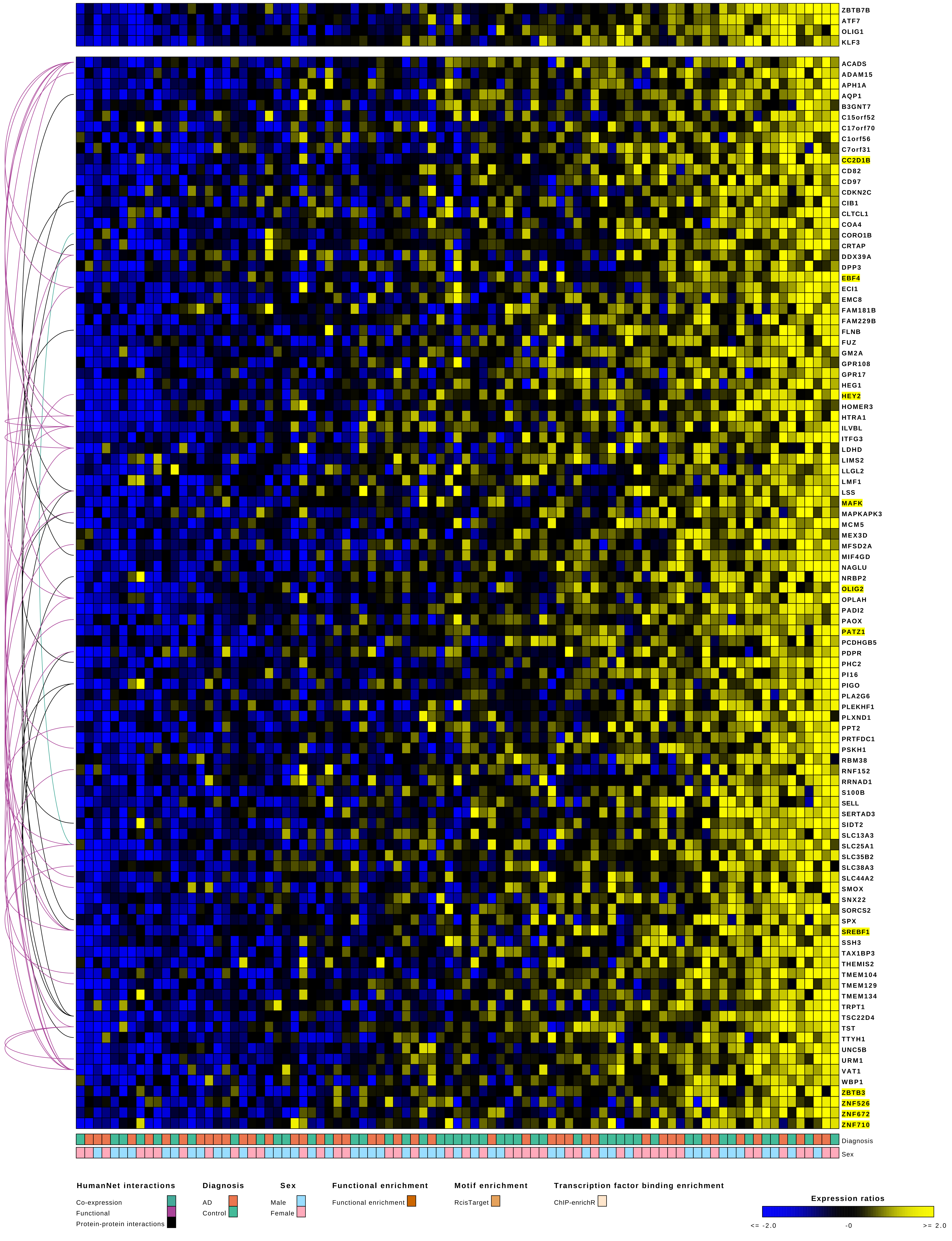
ZBTB7B
ATF7
OLIG1
KLF3
ACADS
ADAM15
APH1A
AQP1
B3GNT7
C15orf52
C17orf70
C1orf56
C7orf31
CC2D1B
CD82
CD97
CDKN2C
CIB1
CLTCL1
COA4
CORO1B
CRTAP
DDX39A
DPP3
EBF4
ECI1
EMC8
FAM181B
FAM229B
FLNB
FUZ
GM2A
GPR108
GPR17
HEG1
HEY2
HOMER3
HTRA1
ILVBL
ITFG3
LDHD
LIMS2
LLGL2
LMF1
LSS
MAFK
MAPKAPK3
MCM5
MEX3D
MFSD2A
MIF4GD
NAGLU
NRBP2
OLIG2
OPLAH
PADI2
PAOX
PATZ1
PCDHGB5
PDPR
PHC2
PI16
PIGO
PLA2G6
PLEKHF1
PLXND1
PPT2
PRTFDC1
PSKH1
RBM38
RNF152
RRNAD1
S100B
SELL
SERTAD3
SIDT2
SLC13A3
SLC25A1
SLC35B2
SLC38A3
SLC44A2
SMOX
SNX22
SORCS2
SPX
SREBF1
SSH3
TAX1BP3
THEMIS2
TMEM104
TMEM129
TMEM134
TRPT1
TSC22D4
TST
TTYH1
UNC5B
URM1
VAT1
WBP1
ZBTB3
ZNF526
ZNF672
ZNF710
Diagnosis
Sex
HumanNet interactions
Co-expression
Functional
Protein-protein interactions
Diagnosis
AD
Control
Sex
Male
Female
Functional enrichment
Functional enrichment
Motif enrichment
RcisTarget
Transcription factor binding enrichment
ChIP-enrichR
Expression ratios
<= -2.0
-0
>= 2.0

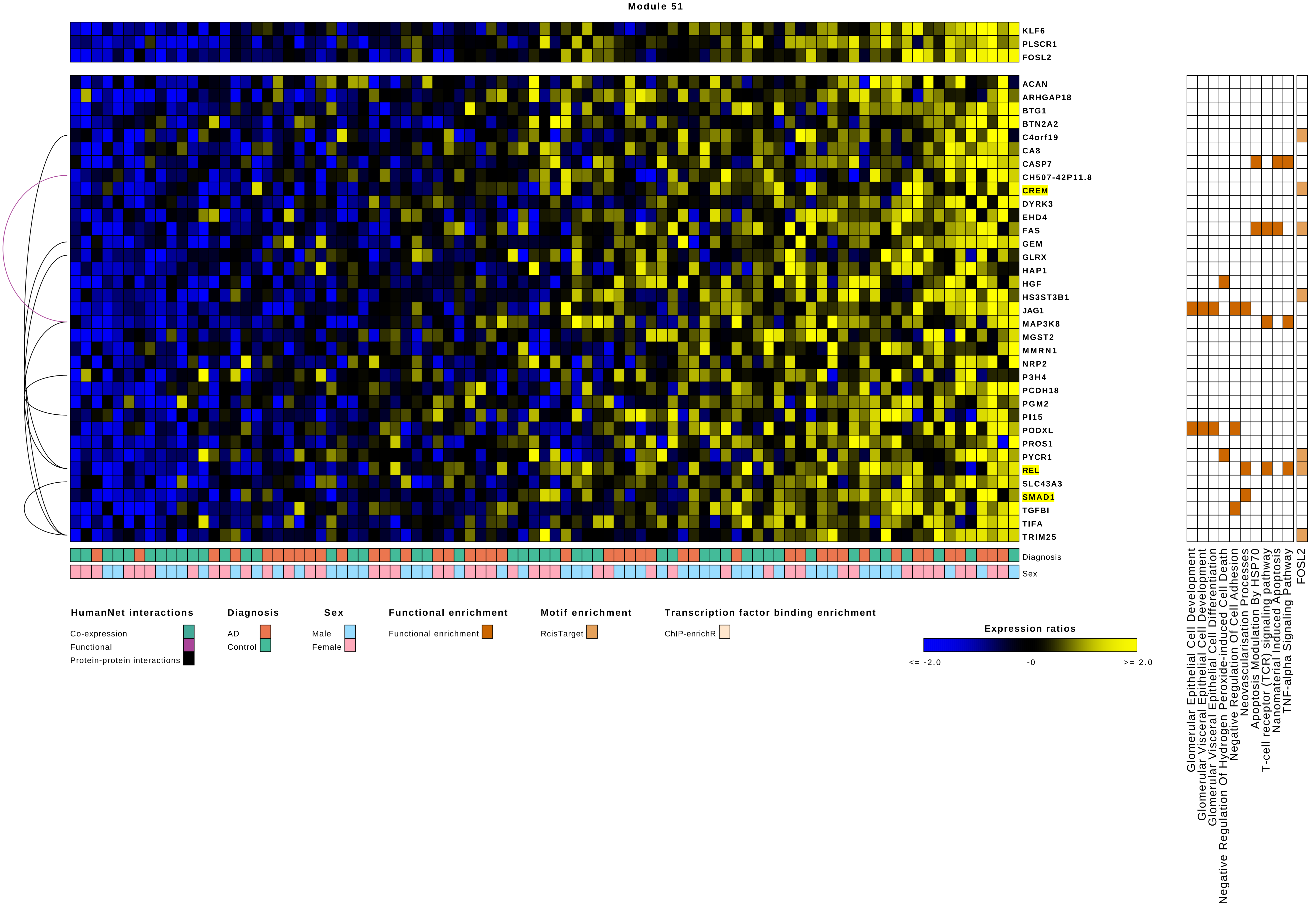
Module 51
KLF6
PLSCR1
FOSL2
ACAN
ARHGAP18
BTG1
BTN2A2
C4orf19
CA8
CASP7
CH507-42P11.8
CREM
DYRK3
EHD4
FAS
GEM
GLRX
HAP1
HGF
HS3ST3B1
JAG1
MAP3K8
MGST2
MMRN1
NRP2
P3H4
PCDH18
PGM2
PI15
PODXL
PROS1
PYCR1
REL
SLC43A3
SMAD1
TGFBI
TIFA
TRIM25
Diagnosis
Sex
Glomerular Epithelial Cell Development
Glomerular Visceral Epithelial Cell Development
Glomerular Visceral Epithelial Cell Differentiation
Negative Regulation Of Hydrogen Peroxide-induced Cell Death
Negative Regulation Of Cell Adhesion
Neovascularisation Processes
Apoptosis Modulation By HSP70
T-cell receptor (TCR) signaling pathway
Nanomaterial Induced Apoptosis
TNF-alpha Signaling Pathway
FOSL2
HumanNet interactions
Co-expression
Functional
Protein-protein interactions
Diagnosis
AD
Control
Sex
Male
Female
Functional enrichment
Functional enrichment
Motif enrichment
RcisTarget
Transcription factor binding enrichment
ChIP-enrichR
Expression ratios
<= -2.0
-0
>= 2.0

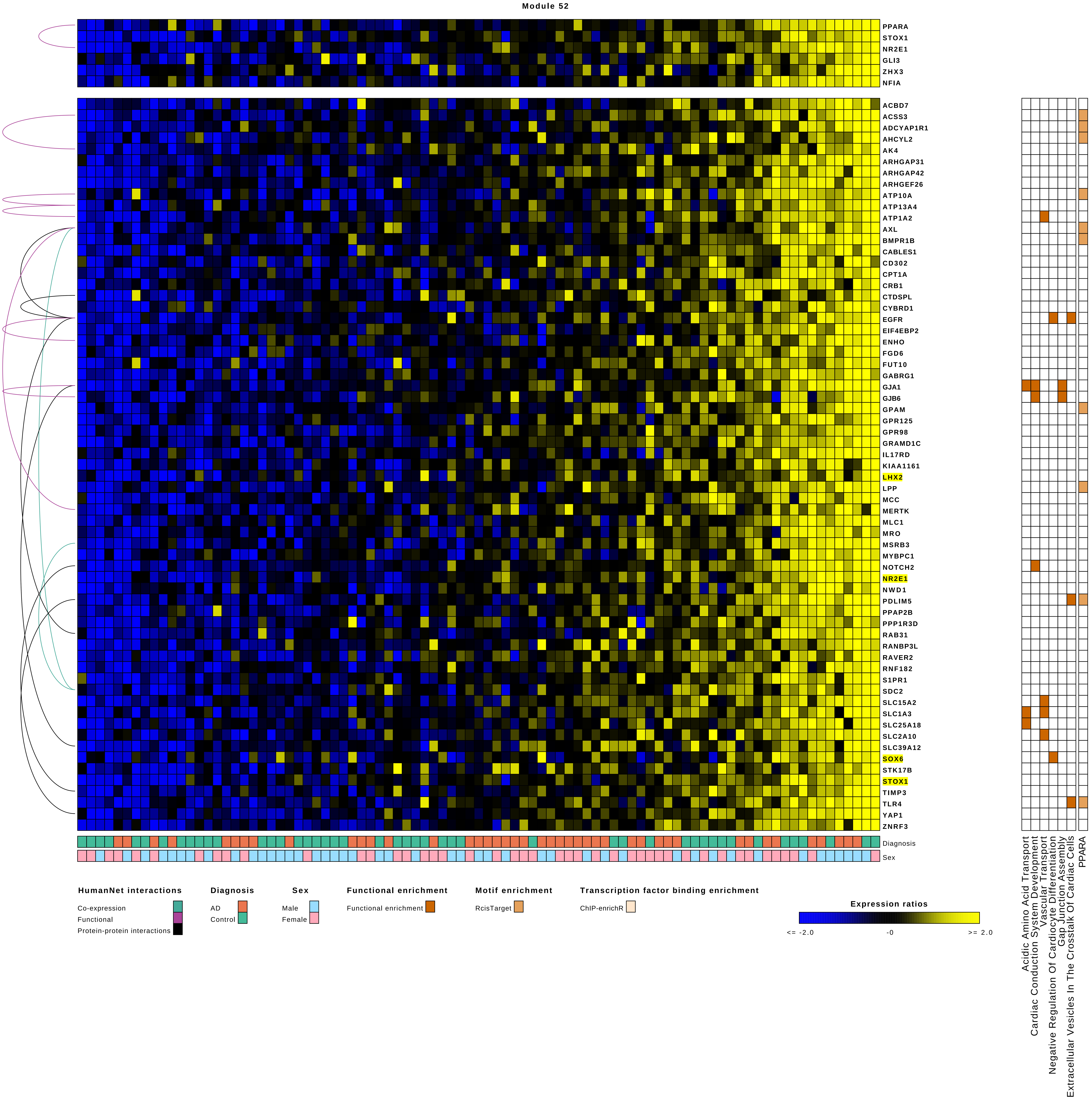
Module 52
PPARA
STOX1
NR2E1
GLI3
ZHX3
NFIA
ACBD7
ACSS3
ADCYAP1R1
AHCYL2
AK4
ARHGAP31
ARHGAP42
ARHGEF26
ATP10A
ATP13A4
ATP1A2
AXL
BMPR1B
CABLES1
CD302
CPT1A
CRB1
CTDSPL
CYBRD1
EGFR
EIF4EBP2
ENHO
FGD6
FUT10
GABRG1
GJA1
GJB6
GPAM
GPR125
GPR98
GRAMD1C
IL17RD
KIAA1161
LHX2
LPP
MCC
MERTK
MLC1
MRO
MSRB3
MYBPC1
NOTCH2
NR2E1
NWD1
PDLIM5
PPAP2B
PPP1R3D
RAB31
RANBP3L
RAVER2
RNF182
S1PR1
SDC2
SLC15A2
SLC1A3
SLC25A18
SLC2A10
SLC39A12
SOX6
STK17B
STOX1
TIMP3
TLR4
YAP1
ZNRF3
Diagnosis
Sex
Acidic Amino Acid Transport
Cardiac Conduction System Development
Vascular Transport
Negative Regulation Of Cardiocyte Differentiation
Gap Junction Assembly
Extracellular Vesicles In The Crosstalk Of Cardiac Cells
PPARA
HumanNet interactions
Co-expression
Functional
Protein-protein interactions
Diagnosis
AD
Control
Sex
Male
Female
Functional enrichment
Functional enrichment
Motif enrichment
RcisTarget
Transcription factor binding enrichment
ChIP-enrichR
Expression ratios
<= -2.0
-0
>= 2.0

Module 53

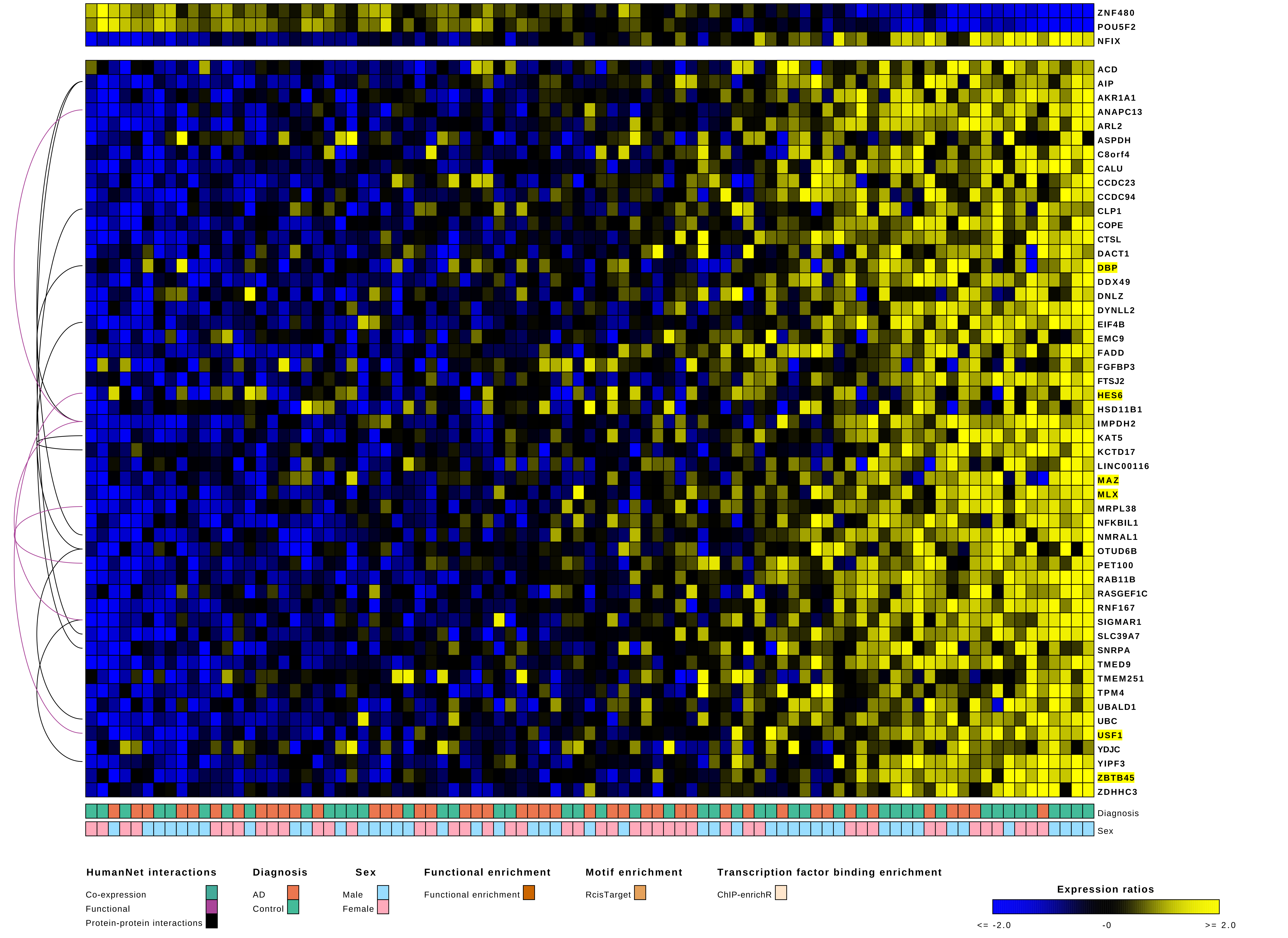
ZNF480
POU5F2
NFIX
ACD
AIP
AKR1A1
ANAPC13
ARL2
ASPDH
C8orf4
CALU
CCDC23
CCDC94
CLP1
COPE
CTSL
DACT1
DBP
DDX49
DNLZ
DYNLL2
EIF4B
EMC9
FADD
FGFBP3
FTSJ2
HES6
HSD11B1
IMPDH2
KAT5
KCTD17
LINC00116
MAZ
MLX
MRPL38
NFKBIL1
NMRAL1
OTUD6B
PET100
RAB11B
RASGEF1C
RNF167
SIGMAR1
SLC39A7
SNRPA
TMED9
TMEM251
TPM4
UBALD1
UBC
USF1
YDJC
YIPF3
ZBTB45
ZDHHC3
Diagnosis
Sex
HumanNet interactions
Co-expression
Functional
Protein-protein interactions
Diagnosis
AD
Control
Sex
Male
Female
Functional enrichment
Functional enrichment
Motif enrichment
RcisTarget
Transcription factor binding enrichment
ChIP-enrichR
Expression ratios
<= -2.0
-0
>= 2.0

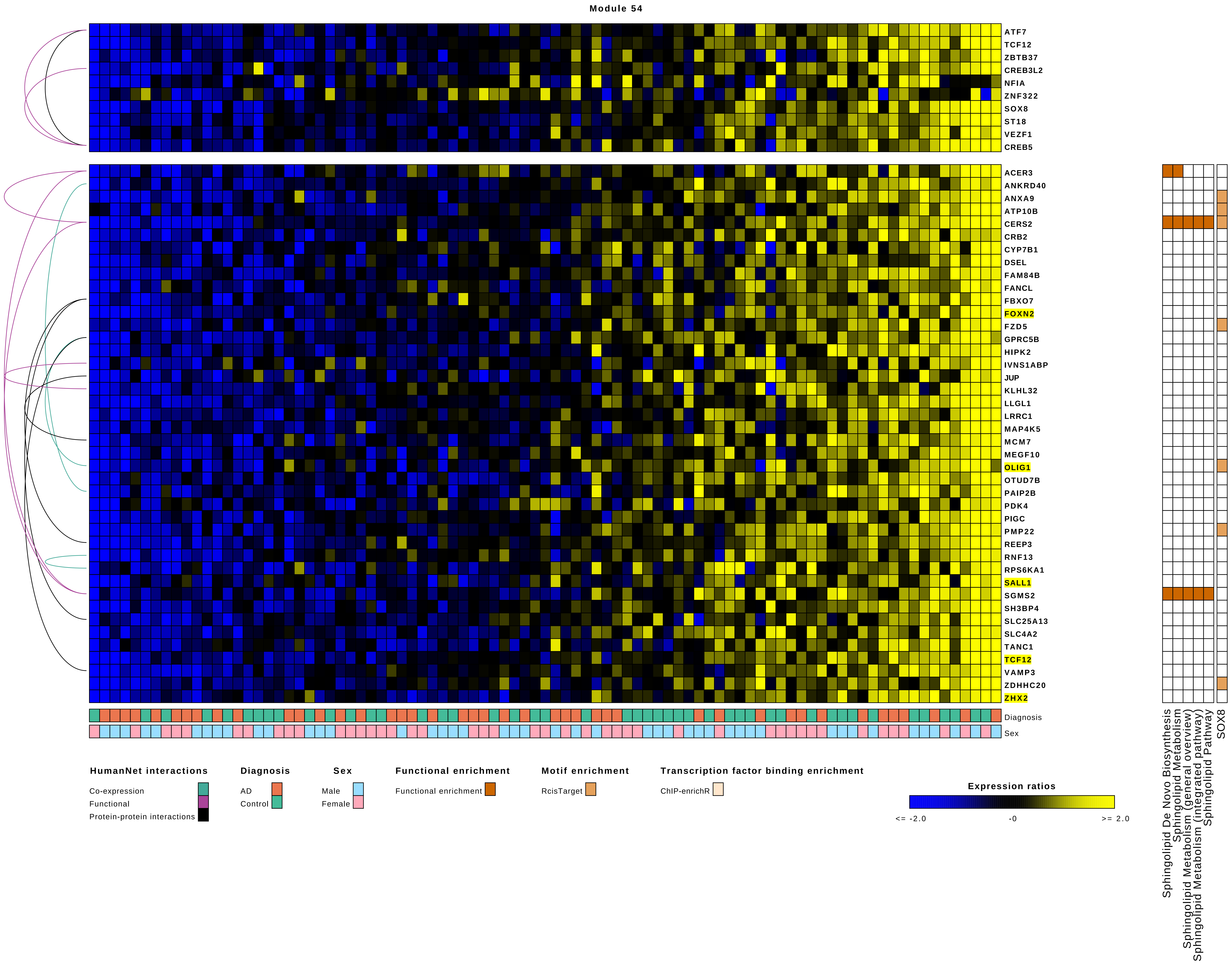

Module 54
ATF7
TCF12
ZBTB37
CREB3L2
NFIA
ZNF322
SOX8
ST18
VEZF1
CREB5
ACER3
ANKRD40
ANXA9
ATP10B
CERS2
CRB2
CYP7B1
DSEL
FAM84B
FANCL
FBXO7
FOXN2
FZD5
GPRC5B
HIPK2
IVNS1ABP
JUP
KLHL32
LLGL1
LRRC1
MAP4K5
MCM7
MEGF10
OLIG1
OTUD7B
PAIP2B
PDK4
PIGC
PMP22
REEP3
RNF13
RPS6KA1
SALL1
SGMS2
SH3BP4
SLC25A13
SLC4A2
TANC1
TCF12
VAMP3
ZDHHC20
ZHX2
Diagnosis
Sex
Sphingolipid De Novo Biosynthesis
Sphingolipid Metabolism
Sphingolipid Metabolism (general overview)
Sphingolipid Metabolism (integrated pathway)
Sphingolipid Pathway
SOX8
HumanNet interactions
Co-expression
Functional
Protein-protein interactions
Diagnosis
AD
Control
Sex
Male
Female
Functional enrichment
Functional enrichment
Motif enrichment
RcisTarget
Transcription factor binding enrichment
ChIP-enrichR
Expression ratios
<= -2.0
-0
>= 2.0

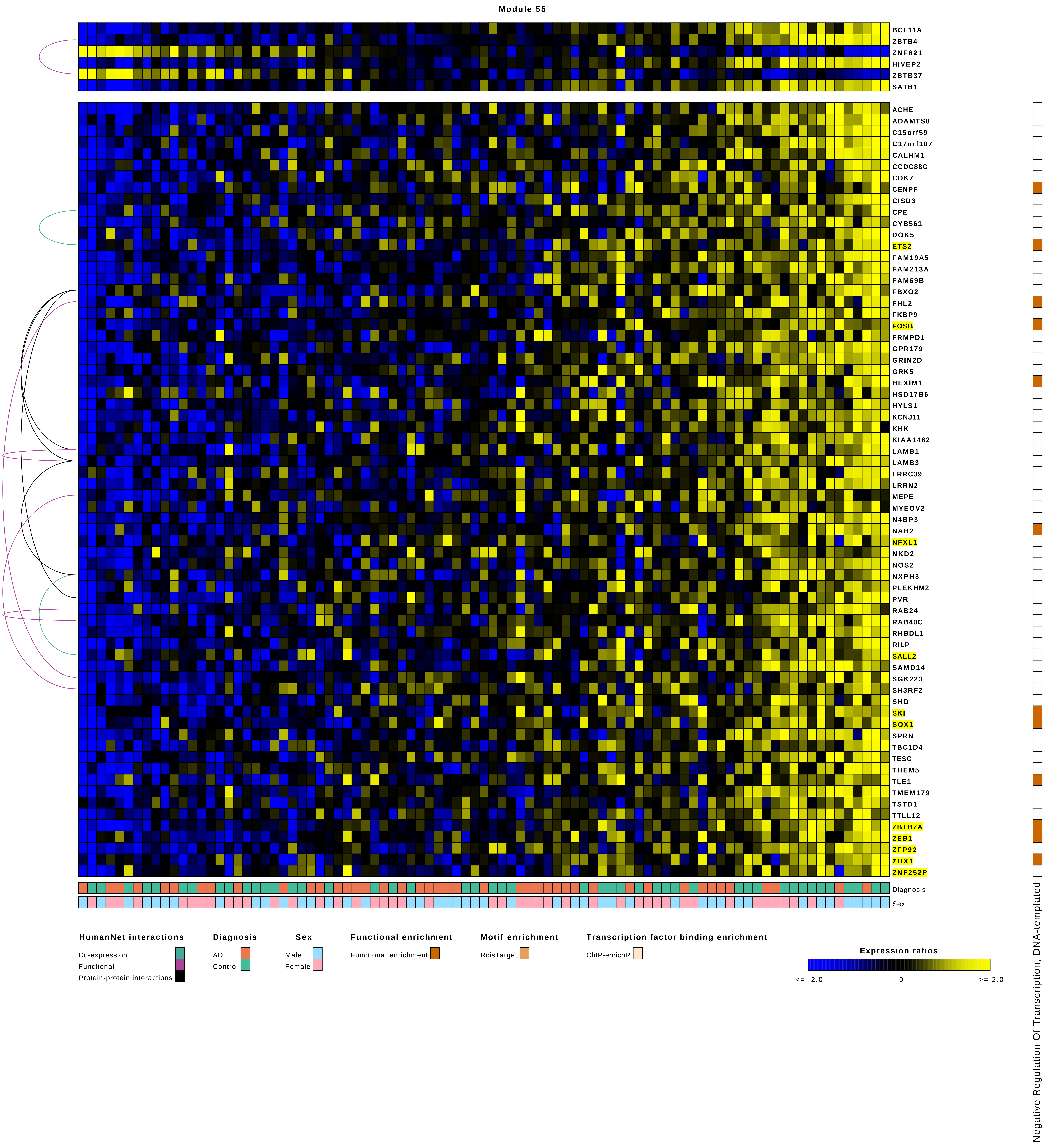
Module 55
BCL11A
ZBTB4
ZNF621
HIVEP2
ZBTB37
SATB1
ACHE
ADAMTS8
C15orf59
C17orf107
CALHM1
CCDC88C
CDK7
CENPF
CISD3
CPE
CYB561
DOK5
ETS2
FAM19A5
FAM213A
FAM69B
FBXO2
FHL2
FKBP9
FOSB
FRMPD1
GPR179
GRIN2D
GRK5
HEXIM1
HSD17B6
HYLS1
KCNJ11
KHK
KIAA1462
LAMB1
LAMB3
LRRC39
LRRN2
MEPE
MYEOV2
N4BP3
NAB2
NFXL1
NKD2
NOS2
NXPH3
PLEKHM2
PVR
RAB24
RAB40C
RHBDL1
RILP
SALL2
SAMD14
SGK223
SH3RF2
SHD
SKI
SOX1
SPRN
TBC1D4
TESC
THEM5
TLE1
TMEM179
TSTD1
TTLL12
ZBTB7A
ZEB1
ZFP92
ZHX1
ZNF252P
Diagnosis
Sex
Negative Regulation Of Transcription, DNA-templated
HumanNet interactions
Co-expression
Functional
Protein-protein interactions
Diagnosis
AD
Control
Sex
Male
Female
Functional enrichment
Functional enrichment
Motif enrichment
RcisTarget
Transcription factor binding enrichment
ChIP-enrichR
Expression ratios
<= -2.0
-0
>= 2.0

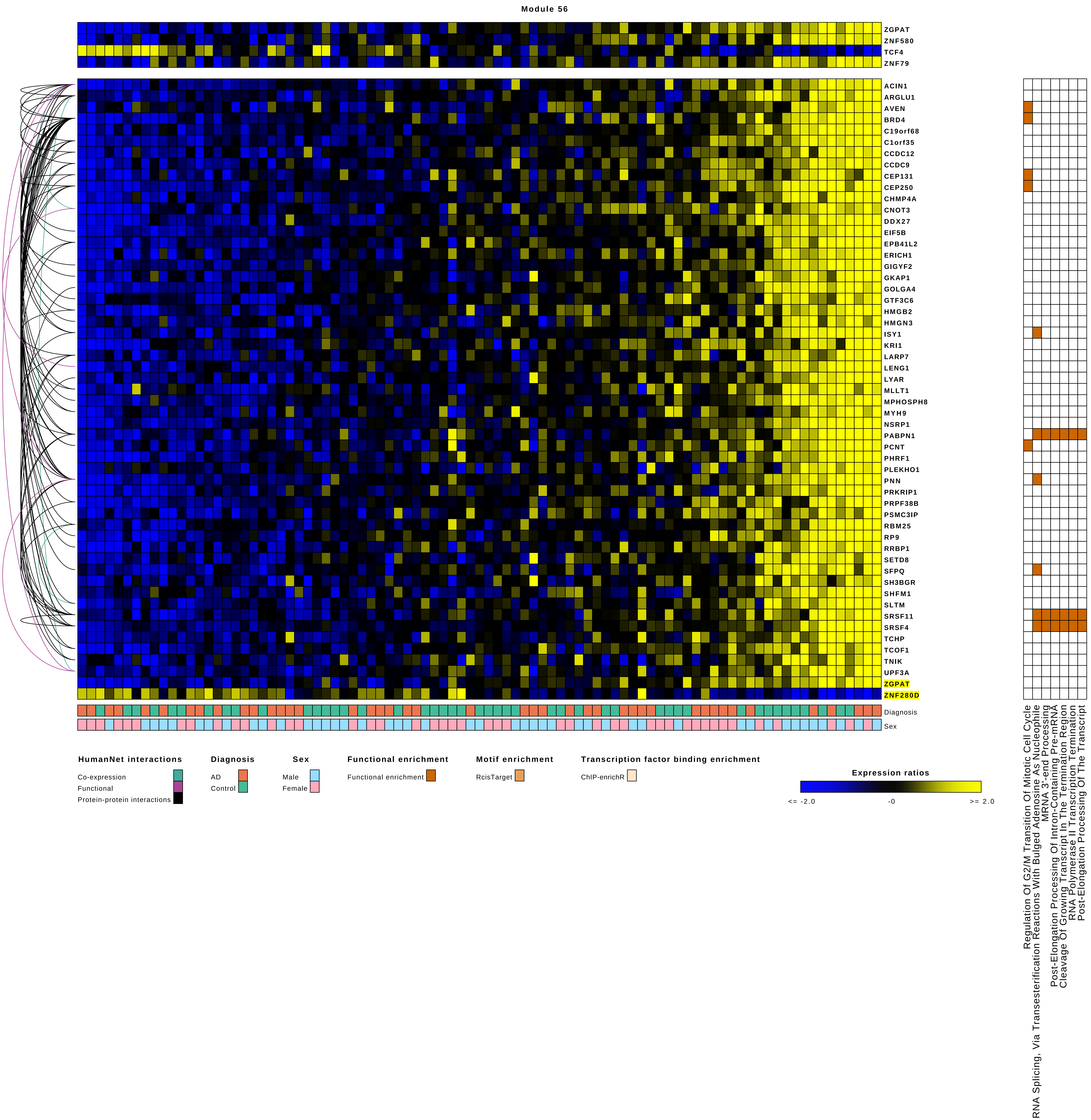

Module 56
ZGPAT
ZNF580
TCF4
ZNF79
ACIN1
ARGLU1
AVEN
BRD4
C19orf68
C1orf35
CCDC12
CCDC9
CEP131
CEP250
CHMP4A
CNOT3
DDX27
EIF5B
EPB41L2
ERICH1
GIGYF2
GKAP1
GOLGA4
GTF3C6
HMGB2
HMGN3
ISY1
KRI1
LARP7
LENG1
LYAR
MLLT1
MPHOSPH8
MYH9
NSRP1
PABPN1
PCNT
PHRF1
PLEKHO1
PNN
PRKRIP1
PRPF38B
PSMC3IP
RBM25
RP9
RRBP1
SETD8
SFPQ
SH3BGR
SHFM1
SLTM
SRSF11
SRSF4
TCHP
TCOF1
TNIK
UPF3A
ZGPAT
ZNF280D
Diagnosis
Sex
Regulation Of G2/M Transition Of Mitotic Cell Cycle
RNA Splicing, Via Transesterification Reactions With Bulged Adenosine As Nucleophile
MRNA 3'-end Processing
Post-Elongation Processing Of Intron-Containing Pre-mRNA
Cleavage Of Growing Transcript In The Termination Region
RNA Polymerase II Transcription Termination
Post-Elongation Processing Of The Transcript
HumanNet interactions
Co-expression
Functional
Protein-protein interactions
Diagnosis
AD
Control
Sex
Male
Female
Functional enrichment
Functional enrichment
Motif enrichment
RcisTarget
Transcription factor binding enrichment
ChIP-enrichR
Expression ratios
<= -2.0
-0
>= 2.0

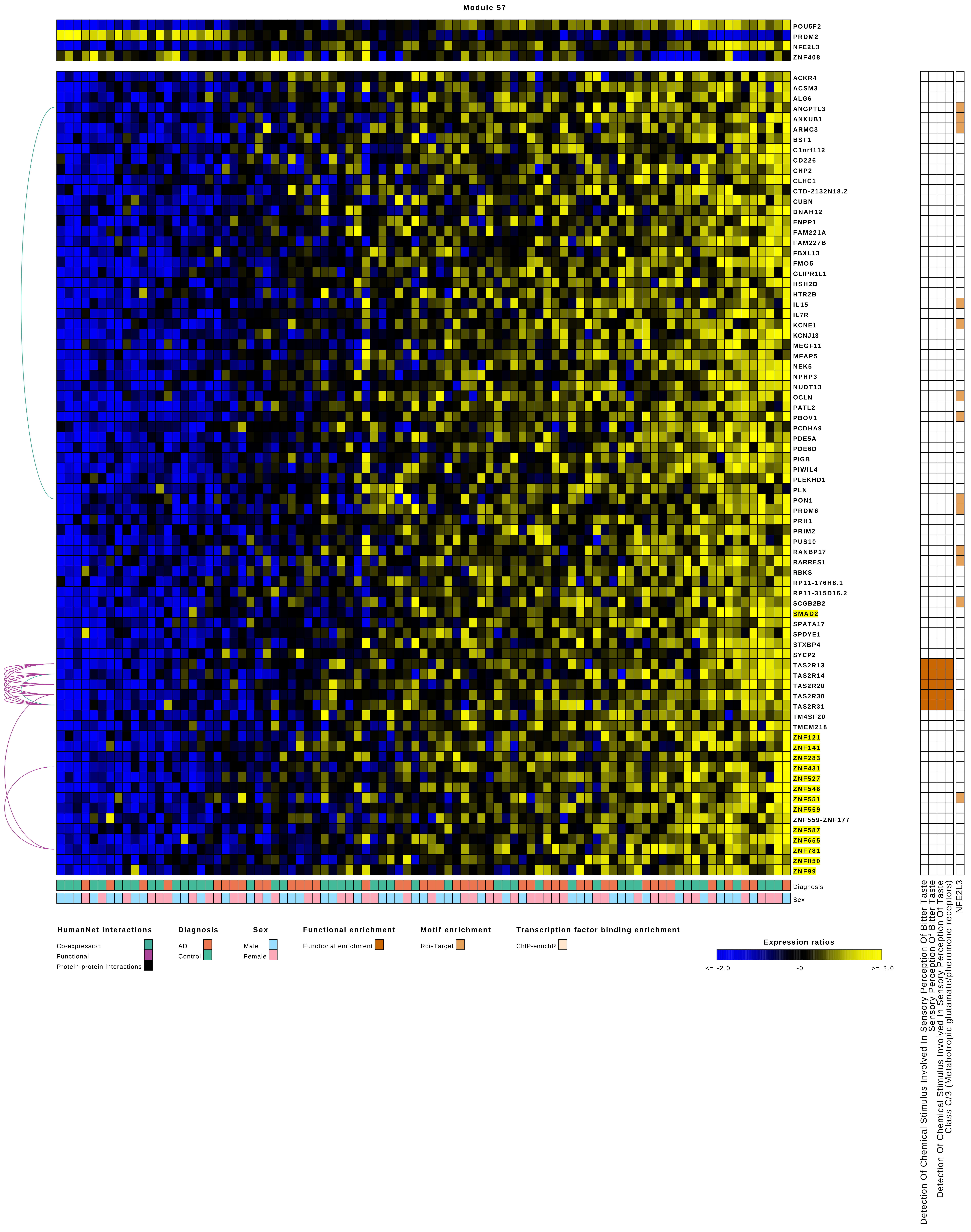

Module 57
POU5F2
PRDM2
NFE2L3
ZNF408
ACKR4
ACSM3
ALG6
ANGPTL3
ANKUB1
ARMC3
BST1
C1orf112
CD226
CHP2
CLHC1
CTD-2132N18.2
CUBN
DNAH12
ENPP1
FAM221A
FAM227B
FBXL13
FMO5
GLIPR1L1
HSH2D
HTR2B
IL15
IL7R
KCNE1
KCNJ13
MEGF11
MFAP5
NEK5
NPHP3
NUDT13
OCLN
PATL2
PBOV1
PCDHA9
PDE5A
PDE6D
PIGB
PIWIL4
PLEKHD1
PLN
PON1
PRDM6
PRH1
PRIM2
PUS10
RANBP17
RARRES1
RBKS
RP11-176H8.1
RP11-315D16.2
SCGB2B2
SMAD2
SPATA17
SPDYE1
STXBP4
SYCP2
TAS2R13
TAS2R14
TAS2R20
TAS2R30
TAS2R31
TM4SF20
TMEM218
ZNF121
ZNF141
ZNF283
ZNF431
ZNF527
ZNF546
ZNF551
ZNF559
ZNF559-ZNF177
ZNF587
ZNF655
ZNF781
ZNF850
ZNF99
Diagnosis
Sex
Detection Of Chemical Stimulus Involved In Sensory Perception Of Bitter Taste
Sensory Perception Of Bitter Taste
Detection Of Chemical Stimulus Involved In Sensory Perception Of Taste
Class C/3 (Metabotropic glutamate/pheromone receptors)
NFE2L3
HumanNet interactions
Co-expression
Functional
Protein-protein interactions
Diagnosis
AD
Control
Sex
Male
Female
Functional enrichment
Functional enrichment
Motif enrichment
RcisTarget
Transcription factor binding enrichment
ChIP-enrichR
Expression ratios
<= -2.0
-0
>= 2.0

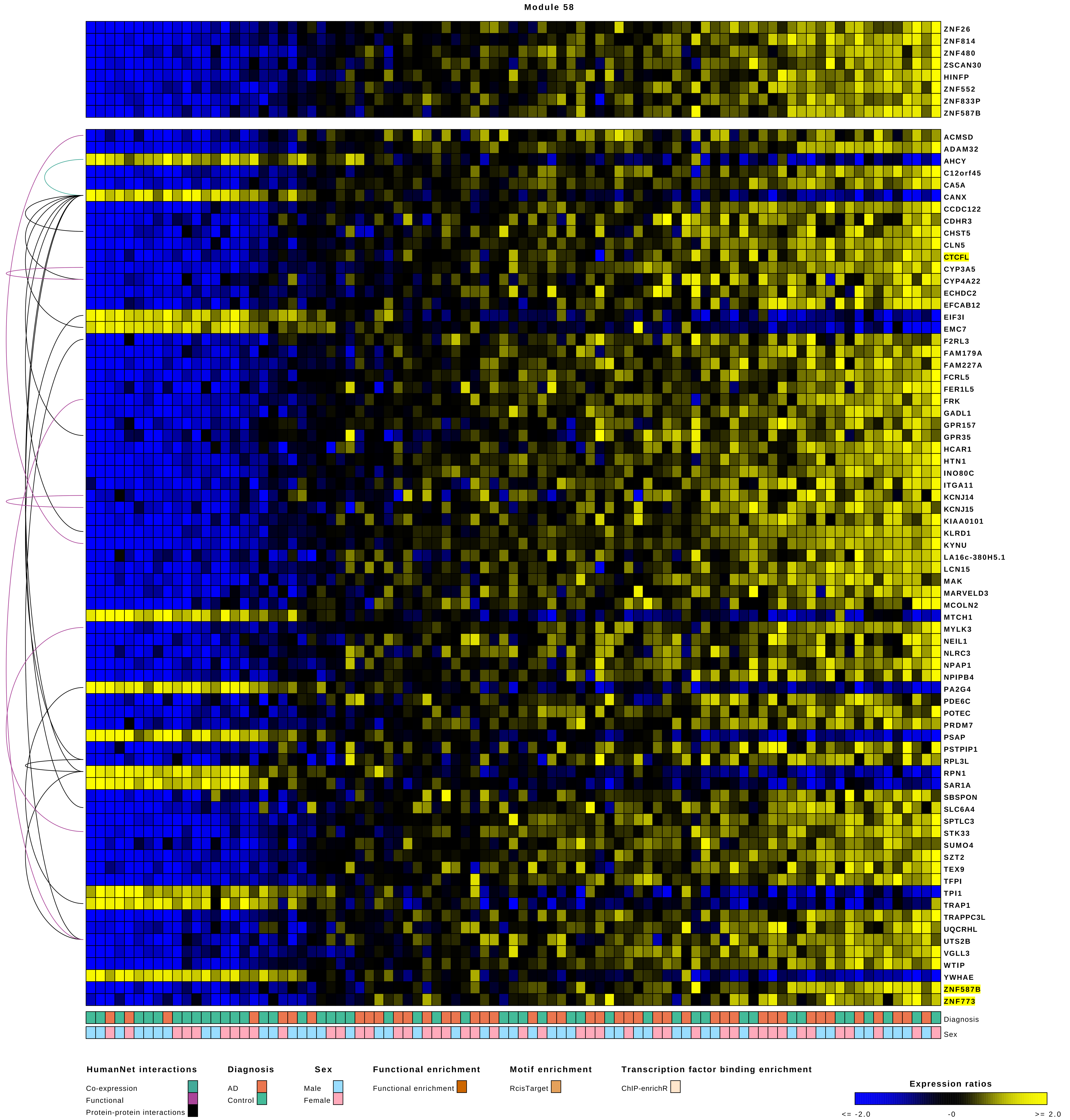
Module 58
ZNF26
ZNF814
ZNF480
ZSCAN30
HINFP
ZNF552
ZNF833P
ZNF587B
ACMSD
ADAM32
AHCY
C12orf45
CA5A
CANX
CCDC122
CDHR3
CHST5
CLN5
CTCFL
CYP3A5
CYP4A22
ECHDC2
EFCAB12
EIF3I
EMC7
F2RL3
FAM179A
FAM227A
FCRL5
FER1L5
FRK
GADL1
GPR157
GPR35
HCAR1
HTN1
INO80C
ITGA11
KCNJ14
KCNJ15
KIAA0101
KLRD1
KYNU
LA16c-380H5.1
LCN15
MAK
MARVELD3
MCOLN2
MTCH1
MYLK3
NEIL1
NLRC3
NPAP1
NPIPB4
PA2G4
PDE6C
POTEC
PRDM7
PSAP
PSTPIP1
RPL3L
RPN1
SAR1A
SBSPON
SLC6A4
SPTLC3
STK33
SUMO4
SZT2
TEX9
TFPI
TPI1
TRAP1
TRAPPC3L
UQCRHL
UTS2B
VGLL3
WTIP
YWHAE
ZNF587B
ZNF773
Diagnosis
Sex
HumanNet interactions
Co-expression
Functional
Protein-protein interactions
Diagnosis
AD
Control
Sex
Male
Female
Functional enrichment
Functional enrichment
Motif enrichment
RcisTarget
Transcription factor binding enrichment
ChIP-enrichR
Expression ratios
<= -2.0
-0
>= 2.0

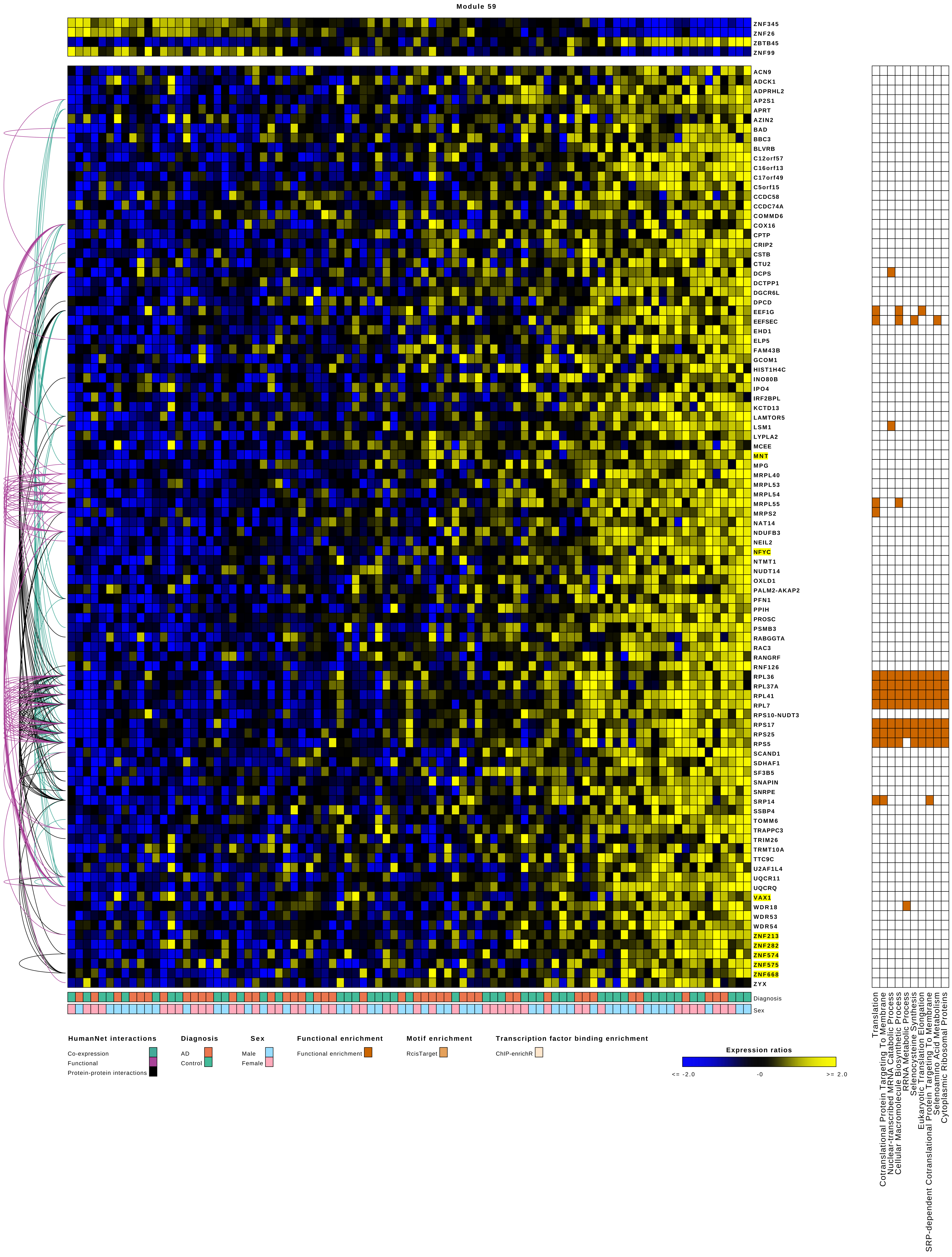

Module 59
ZNF345
ZNF26
ZBTB45
ZNF99
ACN9
ADCK1
ADPRHL2
AP2S1
APRT
AZIN2
BAD
BBC3
BLVRB
C12orf57
C16orf13
C17orf49
C5orf15
CCDC58
CCDC74A
COMMD6
COX16
CPTP
CRIP2
CSTB
CTU2
DCPS
DCTPP1
DGCR6L
DPCD
EEF1G
EEFSEC
EHD1
ELP5
FAM43B
GCOM1
HIST1H4C
INO80B
IPO4
IRF2BPL
KCTD13
LAMTOR5
LSM1
LYPLA2
MCEE
MNT
MPG
MRPL40
MRPL53
MRPL54
MRPL55
MRPS2
NAT14
NDUFB3
NEIL2
NFYC
NTMT1
NUDT14
OXLD1
PALM2-AKAP2
PFN1
PPIH
PROSC
PSMB3
RABGGTA
RAC3
RANGRF
RNF126
RPL36
RPL37A
RPL41
RPL7
RPS10-NUDT3
RPS17
RPS25
RPS5
SCAND1
SDHAF1
SF3B5
SNAPIN
SNRPE
SRP14
SSBP4
TOMM6
TRAPPC3
TRIM26
TRMT10A
TTC9C
U2AF1L4
UQCR11
UQCRQ
VAX1
WDR18
WDR53
WDR54
ZNF213
ZNF282
ZNF574
ZNF575
ZNF668
ZYX
Diagnosis
Sex
Translation
Cotranslational Protein Targeting To Membrane
Nuclear-transcribed MRNA Catabolic Process
Cellular Macromolecule Biosynthetic Process
RRNA Metabolic Process
Selenocysteine Synthesis
Eukaryotic Translation Elongation
SRP-dependent Cotranslational Protein Targeting To Membrane
Selenoamino Acid Metabolism
Cytoplasmic Ribosomal Proteins
HumanNet interactions
Co-expression
Functional
Protein-protein interactions
Diagnosis
AD
Control
Sex
Male
Female
Functional enrichment
Functional enrichment
Motif enrichment
RcisTarget
Transcription factor binding enrichment
ChIP-enrichR
Expression ratios
<= -2.0
-0
>= 2.0

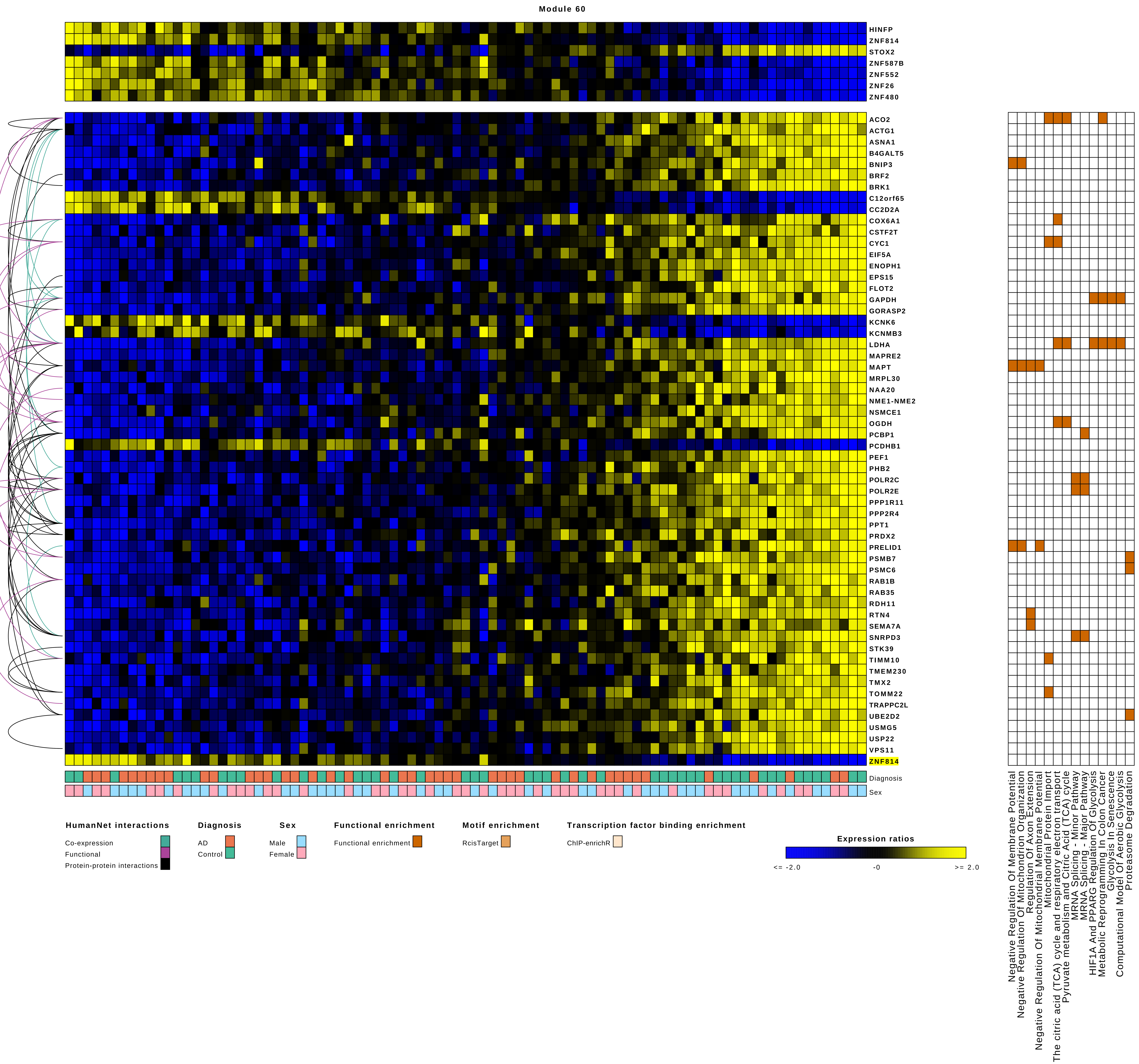
Module 60
HINFP
ZNF814
STOX2
ZNF587B
ZNF552
ZNF26
ZNF480
ACO2
ACTG1
ASNA1
B4GALT5
BNIP3
BRF2
BRK1
C12orf65
CC2D2A
COX6A1
CSTF2T
CYC1
EIF5A
ENOPH1
EPS15
FLOT2
GAPDH
GORASP2
KCNK6
KCNMB3
LDHA
MAPRE2
MAPT
MRPL30
NAA20
NME1-NME2
NSMCE1
OGDH
PCBP1
PCDHB1
PEF1
PHB2
POLR2C
POLR2E
PPP1R11
PPP2R4
PPT1
PRDX2
PRELID1
PSMB7
PSMC6
RAB1B
RAB35
RDH11
RTN4
SEMA7A
SNRPD3
STK39
TIMM10
TMEM230
TMX2
TOMM22
TRAPPC2L
UBE2D2
USMG5
USP22
VPS11
ZNF814
Diagnosis
Sex
HumanNet interactions
Co-expression
Functional
Protein-protein interactions
Diagnosis
AD
Control
Sex
Male
Female
Functional enrichment
Functional enrichment
Motif enrichment
RcisTarget
Transcription factor binding enrichment
ChIP-enrichR
Expression ratios
<= -2.0
-0
>= 2.0
Negative Regulation Of Membrane Potential
Negative Regulation Of Mitochondrion Organization
Regulation Of Axon Extension
Negative Regulation Of Mitochondrial Membrane Potential
Mitochondrial Protein Import
The citric acid (TCA) cycle and respiratory electron transport
Pyruvate metabolism and Citric Acid (TCA) cycle
MRNA Splicing - Minor Pathway
MRNA Splicing - Major Pathway
HIF1A And PPARG Regulation Of Glycolysis
Metabolic Reprogramming In Colon Cancer
Glycolysis In Senescence
Computational Model Of Aerobic Glycolysis
Proteasome Degradation

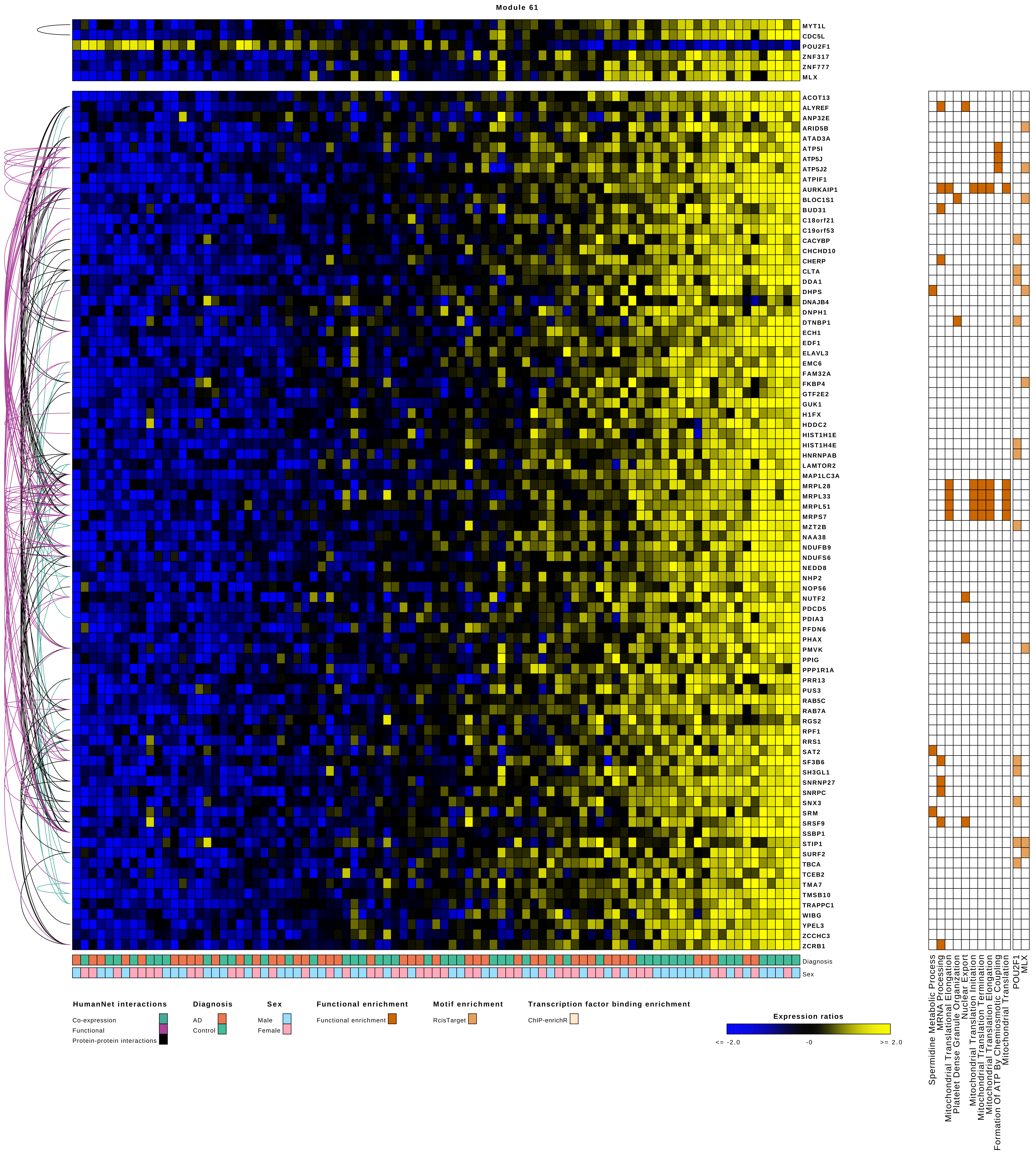

Module 61
MYT1L
CDC5L
POU2F1
ZNF317
ZNF777
MLX
ACOT13
ALYREF
ANP32E
ARID5B
ATAD3A
ATP5I
ATP5J
ATP5J2
ATPIF1
AURKAIP1
BLOC1S1
BUD31
C18orf21
C19orf53
CACYBP
CHCHD10
CHERP
CLTA
DDA1
DHPS
DNAJB4
DNPH1
DTNBP1
ECH1
EDF1
ELAVL3
EMC6
FAM32A
FKBP4
GTF2E2
GUK1
H1FX
HDDC2
HIST1H1E
HIST1H4E
HNRNPAB
LAMTOR2
MAP1LC3A
MRPL28
MRPL33
MRPL51
MRPS7
MZT2B
NAA38
NDUFB9
NDUFS6
NEDD8
NHP2
NOP56
NUTF2
PDCD5
PDIA3
PFDN6
PHAX
PMVK
PPIG
PPP1R1A
PRR13
PUS3
RAB5C
RAB7A
RGS2
RPF1
RRS1
SAT2
SF3B6
SH3GL1
SNRNP27
SNRPC
SNX3
SRM
SRSF9
SSBP1
STIP1
SURF2
TBCA
TCEB2
TMA7
TMSB10
TRAPPC1
WIBG
YPEL3
ZCCHC3
ZCRB1
Diagnosis
Sex
Spermidine Metabolic Process
MRNA Processing
Mitochondrial Translational Elongation
Platelet Dense Granule Organization
Nuclear Export
Mitochondrial Translation Initiation
Mitochondrial Translation Termination
Mitochondrial Translation Elongation
Formation Of ATP By Chemiosmotic Coupling
Mitochondrial Translation
POU2F1
MLX
HumanNet interactions
Co-expression
Functional
Protein-protein interactions
Diagnosis
AD
Control
Sex
Male
Female
Functional enrichment
Functional enrichment
Motif enrichment
RcisTarget
Transcription factor binding enrichment
ChIP-enrichR
Expression ratios
<= -2.0
-0
>= 2.0

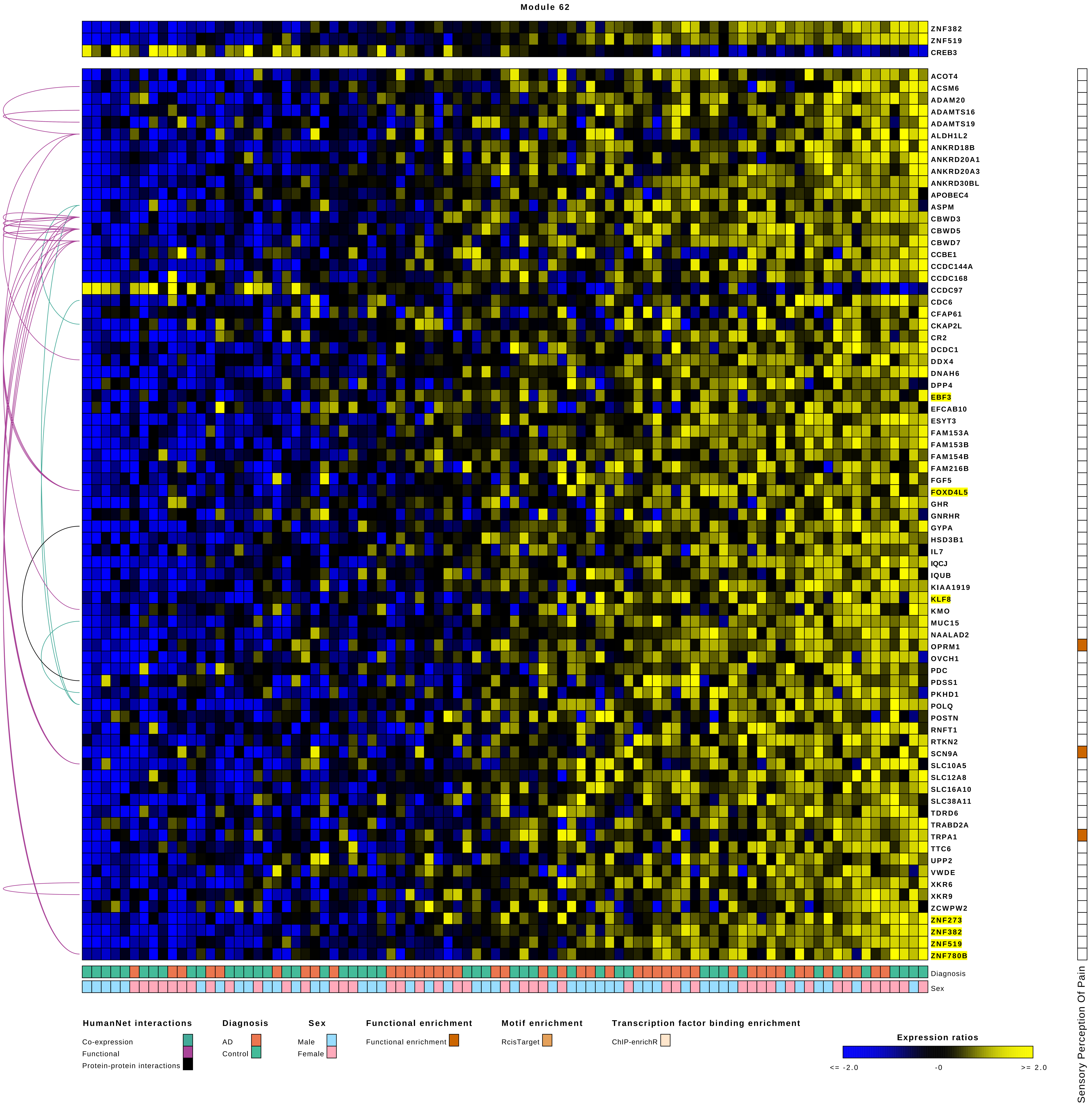
Module 62
ZNF382
ZNF519
CREB3
ACOT4
ACSM6
ADAM20
ADAMTS16
ADAMTS19
ALDH1L2
ANKRD18B
ANKRD20A1
ANKRD20A3
ANKRD30BL
APOBEC4
ASPM
CBWD3
CBWD5
CBWD7
CCBE1
CCDC144A
CCDC168
CCDC97
CDC6
CFAP61
CKAP2L
CR2
DCDC1
DDX4
DNAH6
DPP4
EBF3
EFCAB10
ESYT3
FAM153A
FAM153B
FAM154B
FAM216B
FGF5
FOXD4L5
GHR
GNRHR
GYPA
HSD3B1
IL7
IQCJ
IQUB
KIAA1919
KLF8
KMO
MUC15
NAALAD2
OPRM1
OVCH1
PDC
PDSS1
PKHD1
POLQ
POSTN
RNFT1
RTKN2
SCN9A
SLC10A5
SLC12A8
SLC16A10
SLC38A11
TDRD6
TRABD2A
TRPA1
TTC6
UPP2
VWDE
XKR6
XKR9
ZCWPW2
ZNF273
ZNF382
ZNF519
ZNF780B
Diagnosis
Sex
Sensory Perception Of Pain
HumanNet interactions
Co-expression
Functional
Protein-protein interactions
Diagnosis
AD
Control
Sex
Male
Female
Functional enrichment
Functional enrichment
Motif enrichment
RcisTarget
Transcription factor binding enrichment
ChIP-enrichR
Expression ratios
<= -2.0
-0
>= 2.0

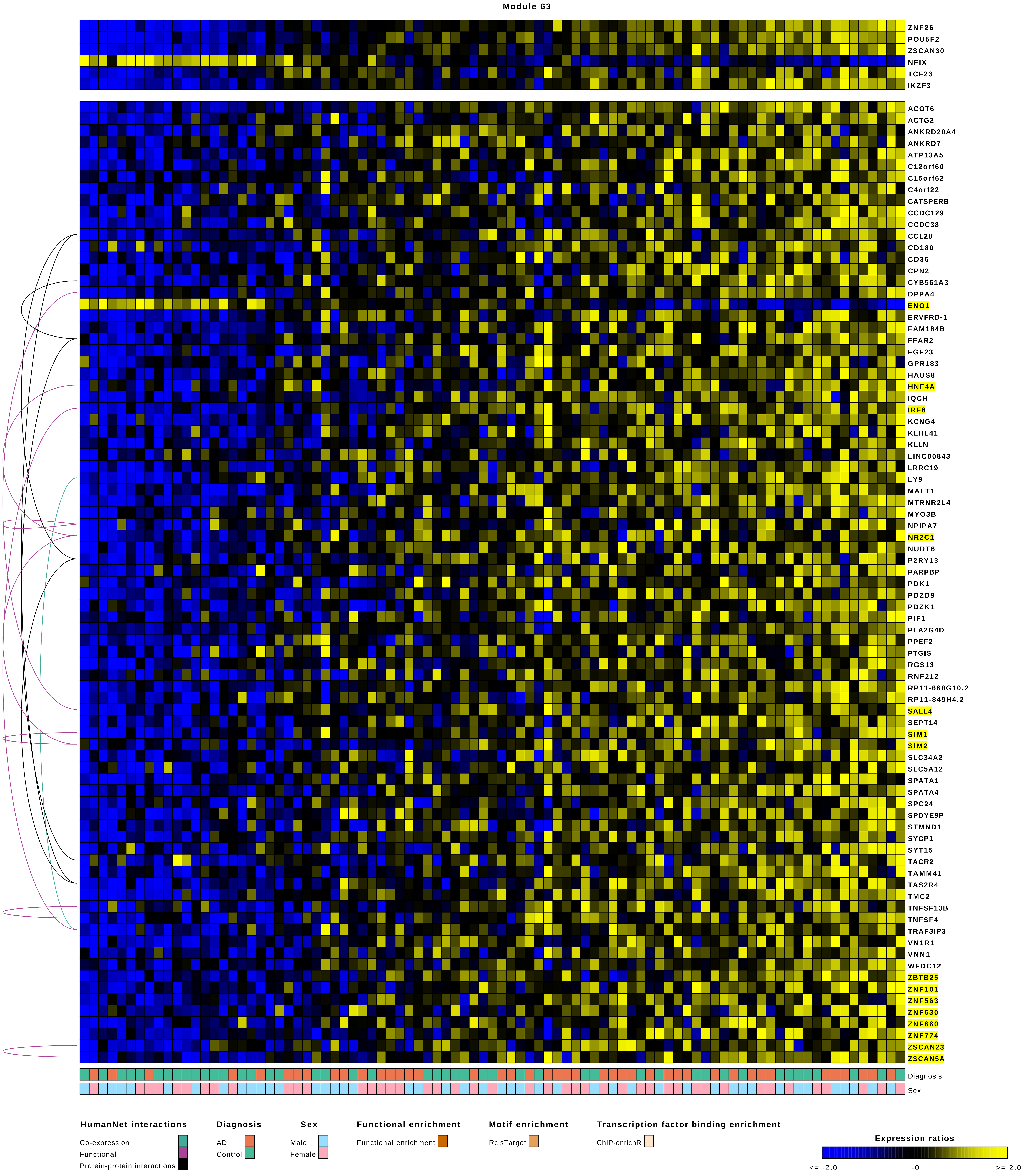
Module 63
ZNF26
POU5F2
ZSCAN30
NFIX
TCF23
IKZF3
ACOT6
ACTG2
ANKRD20A4
ANKRD7
ATP13A5
C12orf60
C15orf62
C4orf22
CATSPERB
CCDC129
CCDC38
CCL28
CD180
CD36
CPN2
CYB561A3
DPPA4
ENO1
ERVFRD-1
FAM184B
FFAR2
FGF23
GPR183
HAUS8
HNF4A
IQCH
IRF6
KCNG4
KLHL41
KLLN
LINC00843
LRRC19
LY9
MALT1
MTRNR2L4
MYO3B
NPIPA7
NR2C1
NUDT6
P2RY13
PARPBP
PDK1
PDZD9
PDZK1
PIF1
PLA2G4D
PPEF2
PTGIS
RGS13
RNF212
RP11-668G10.2
RP11-849H4.2
SALL4
SEPT14
SIM1
SIM2
SLC34A2
SLC5A12
SPATA1
SPATA4
SPC24
SPDYE9P
STMND1
SYCP1
SYT15
TACR2
TAMM41
TAS2R4
TMC2
TNFSF13B
TNFSF4
TRAF3IP3
VN1R1
VNN1
WFDC12
ZBTB25
ZNF101
ZNF563
ZNF630
ZNF660
ZNF774
ZSCAN23
ZSCAN5A
Diagnosis
Sex
HumanNet interactions
Co-expression
Functional
Protein-protein interactions
Diagnosis
AD
Control
Sex
Male
Female
Functional enrichment
Functional enrichment
Motif enrichment
RcisTarget
Transcription factor binding enrichment
ChIP-enrichR
Expression ratios
<= -2.0
-0
>= 2.0

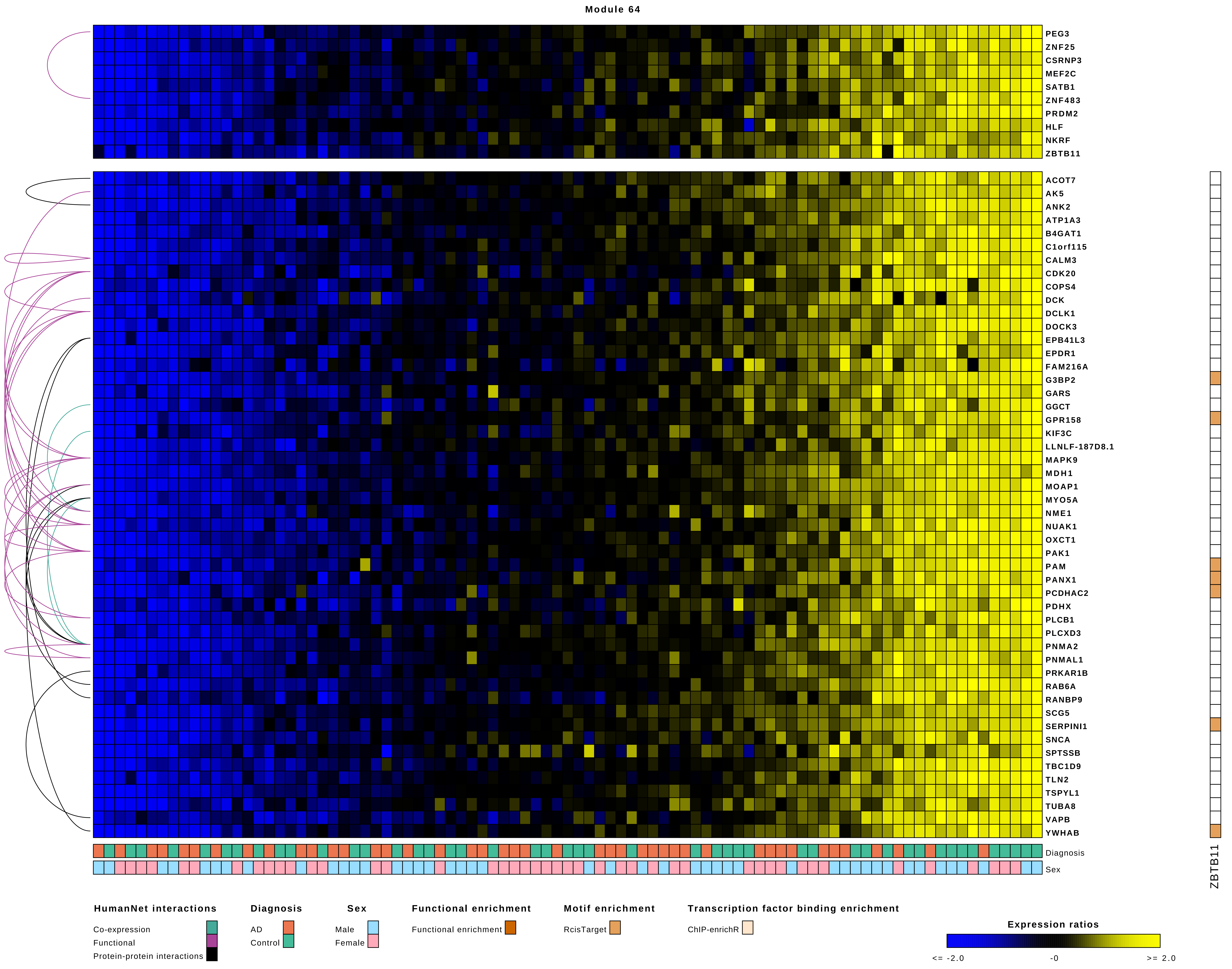
Module 64
PEG3
ZNF25
CSRNP3
MEF2C
SATB1
ZNF483
PRDM2
HLF
NKRF
ZBTB11
ACOT7
AK5
ANK2
ATP1A3
B4GAT1
C1orf115
CALM3
CDK20
COPS4
DCK
DCLK1
DOCK3
EPB41L3
EPDR1
FAM216A
G3BP2
GARS
GGCT
GPR158
KIF3C
LLNLF-187D8.1
MAPK9
MDH1
MOAP1
MYO5A
NME1
NUAK1
OXCT1
PAK1
PAM
PANX1
PCDHAC2
PDHX
PLCB1
PLCXD3
PNMA2
PNMAL1
PRKAR1B
RAB6A
RANBP9
SCG5
SERPINI1
SNCA
SPTSSB
TBC1D9
TLN2
TSPYL1
TUBA8
VAPB
YWHAB
Diagnosis
Sex
ZBTB11
HumanNet interactions
Co-expression
Functional
Protein-protein interactions
Diagnosis
AD
Control
Sex
Male
Female
Functional enrichment
Functional enrichment
Motif enrichment
RcisTarget
Transcription factor binding enrichment
ChIP-enrichR
Expression ratios
<= -2.0
-0
>= 2.0

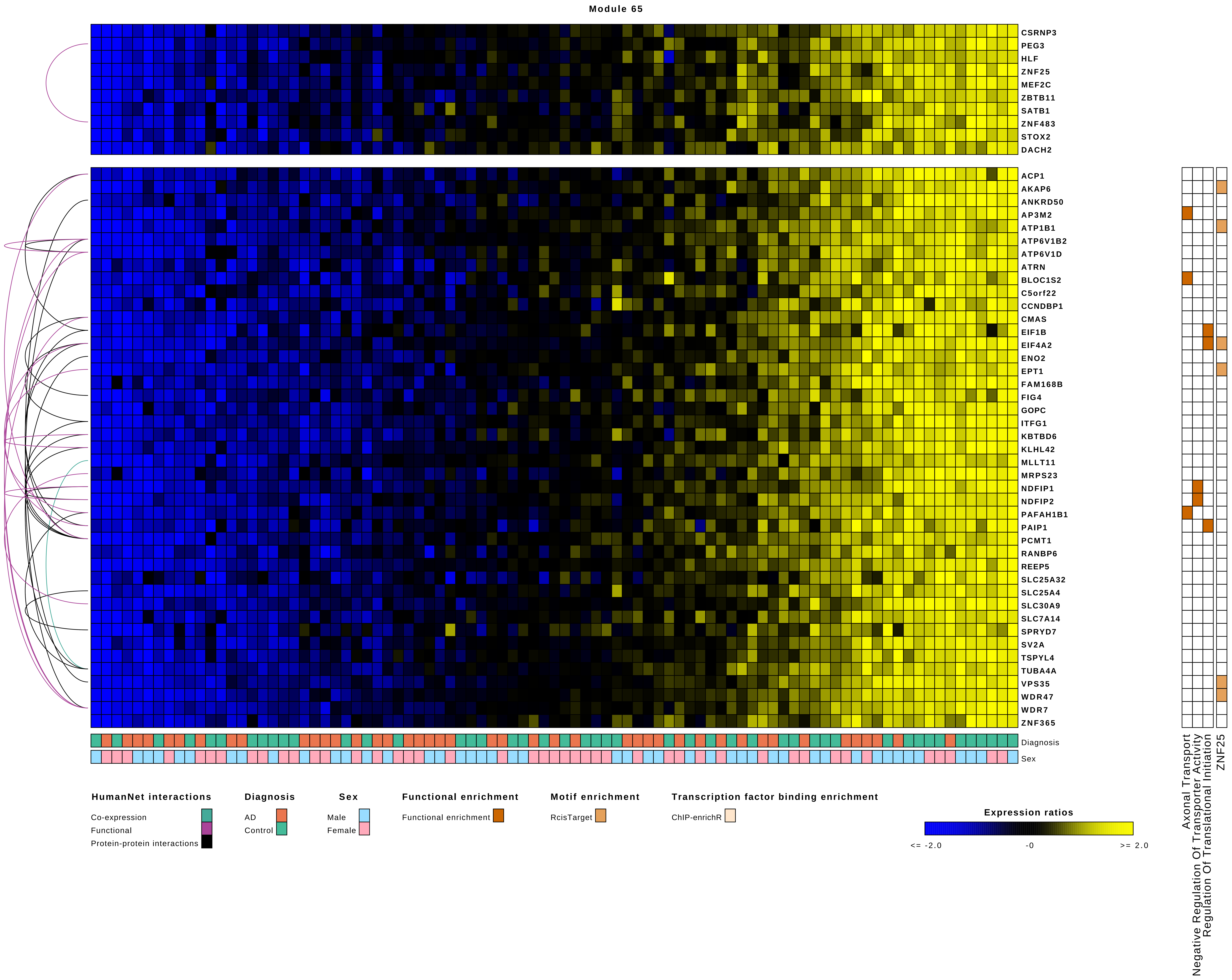
Module 65
CSRNP3
PEG3
HLF
ZNF25
MEF2C
ZBTB11
SATB1
ZNF483
STOX2
DACH2
ACP1
AKAP6
ANKRD50
AP3M2
ATP1B1
ATP6V1B2
ATP6V1D
ATRN
BLOC1S2
C5orf22
CCNDBP1
CMAS
EIF1B
EIF4A2
ENO2
EPT1
FAM168B
FIG4
GOPC
ITFG1
KBTBD6
KLHL42
MLLT11
MRPS23
NDFIP1
NDFIP2
PAFAH1B1
PAIP1
PCMT1
RANBP6
REEP5
SLC25A32
SLC25A4
SLC30A9
SLC7A14
SPRYD7
SV2A
TSPYL4
TUBA4A
VPS35
WDR47
WDR7
ZNF365
Diagnosis
Sex
Axonal Transport
Negative Regulation Of Transporter Activity
Regulation Of Translational Initiation
ZNF25
HumanNet interactions
Co-expression
Functional
Protein-protein interactions
Diagnosis
AD
Control
Sex
Male
Female
Functional enrichment
Functional enrichment
Motif enrichment
RcisTarget
Transcription factor binding enrichment
ChIP-enrichR
Expression ratios
<= -2.0
-0
>= 2.0

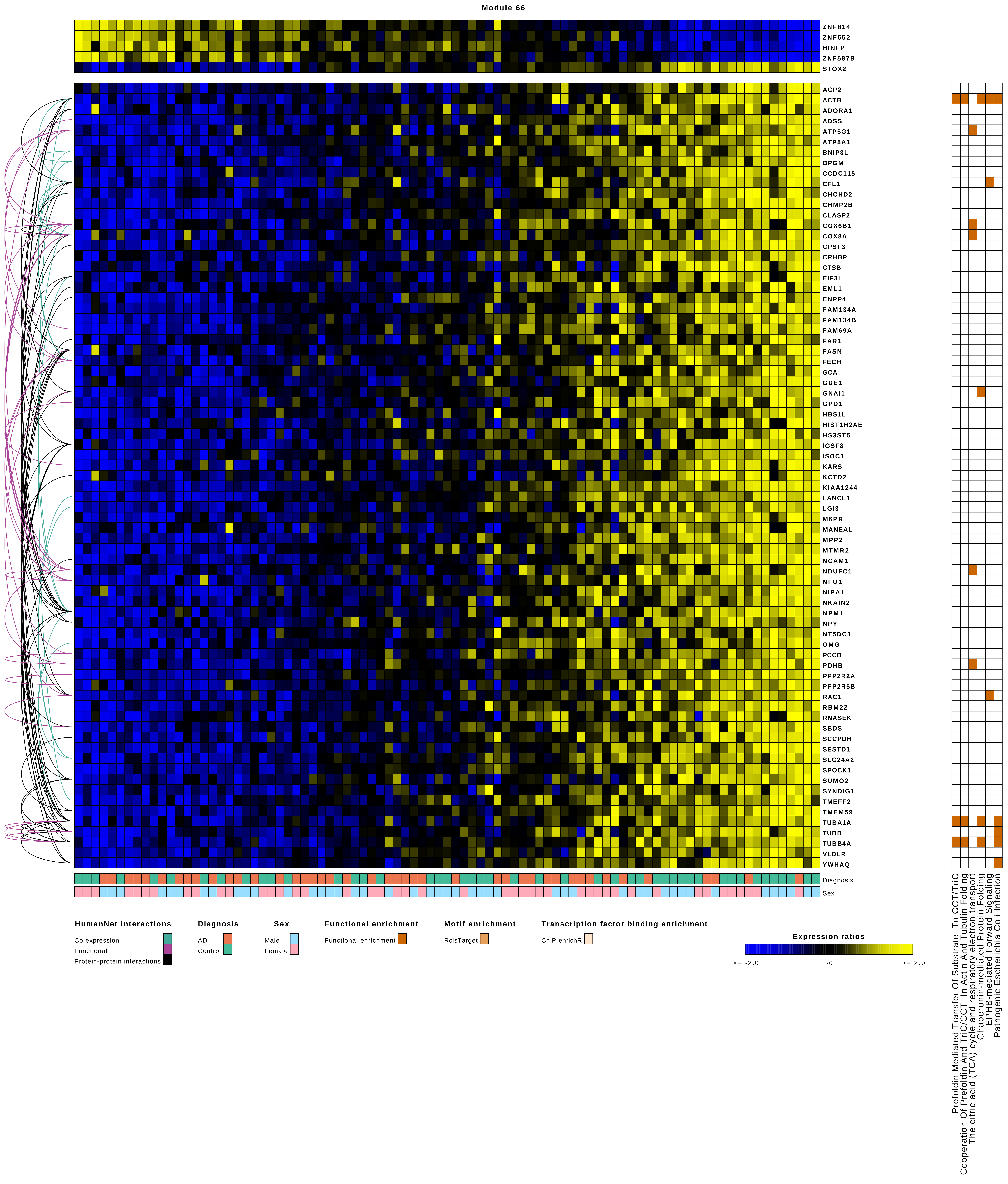
Module 66
ZNF814
ZNF552
HINFP
ZNF587B
STOX2
ACP2
ACTB
ADORA1
ADSS
ATP5G1
ATP8A1
BNIP3L
BPGM
CCDC115
CFL1
CHCHD2
CHMP2B
CLASP2
COX6B1
COX8A
CPSF3
CRHBP
CTSB
EIF3L
EML1
ENPP4
FAM134A
FAM134B
FAM69A
FAR1
FASN
FECH
GCA
GDE1
GNAI1
GPD1
HBS1L
HIST1H2AE
HS3ST5
IGSF8
ISOC1
KARS
KCTD2
KIAA1244
LANCL1
LGI3
M6PR
MANEAL
MPP2
MTMR2
NCAM1
NDUFC1
NFU1
NIPA1
NKAIN2
NPM1
NPY
NT5DC1
OMG
PCCB
PDHB
PPP2R2A
PPP2R5B
RAC1
RBM22
RNASEK
SBDS
SCCPDH
SESTD1
SLC24A2
SPOCK1
SUMO2
SYNDIG1
TMEFF2
TMEM59
TUBA1A
TUBB
TUBB4A
VLDLR
YWHAQ
Diagnosis
Sex
Prefoldin Mediated Transfer Of Substrate To CCT/TriC
Cooperation Of Prefoldin And TriC/CCT In Actin And Tubulin Folding
The citric acid (TCA) cycle and respiratory electron transport
Chaperonin-mediated Protein Folding
EPHB-mediated Forward Signaling
Pathogenic Escherichia Coli Infection
HumanNet interactions
Co-expression
Functional
Protein-protein interactions
Diagnosis
AD
Control
Sex
Male
Female
Functional enrichment
Functional enrichment
Motif enrichment
RcisTarget
Transcription factor binding enrichment
ChIP-enrichR
Expression ratios
<= -2.0
-0
>= 2.0

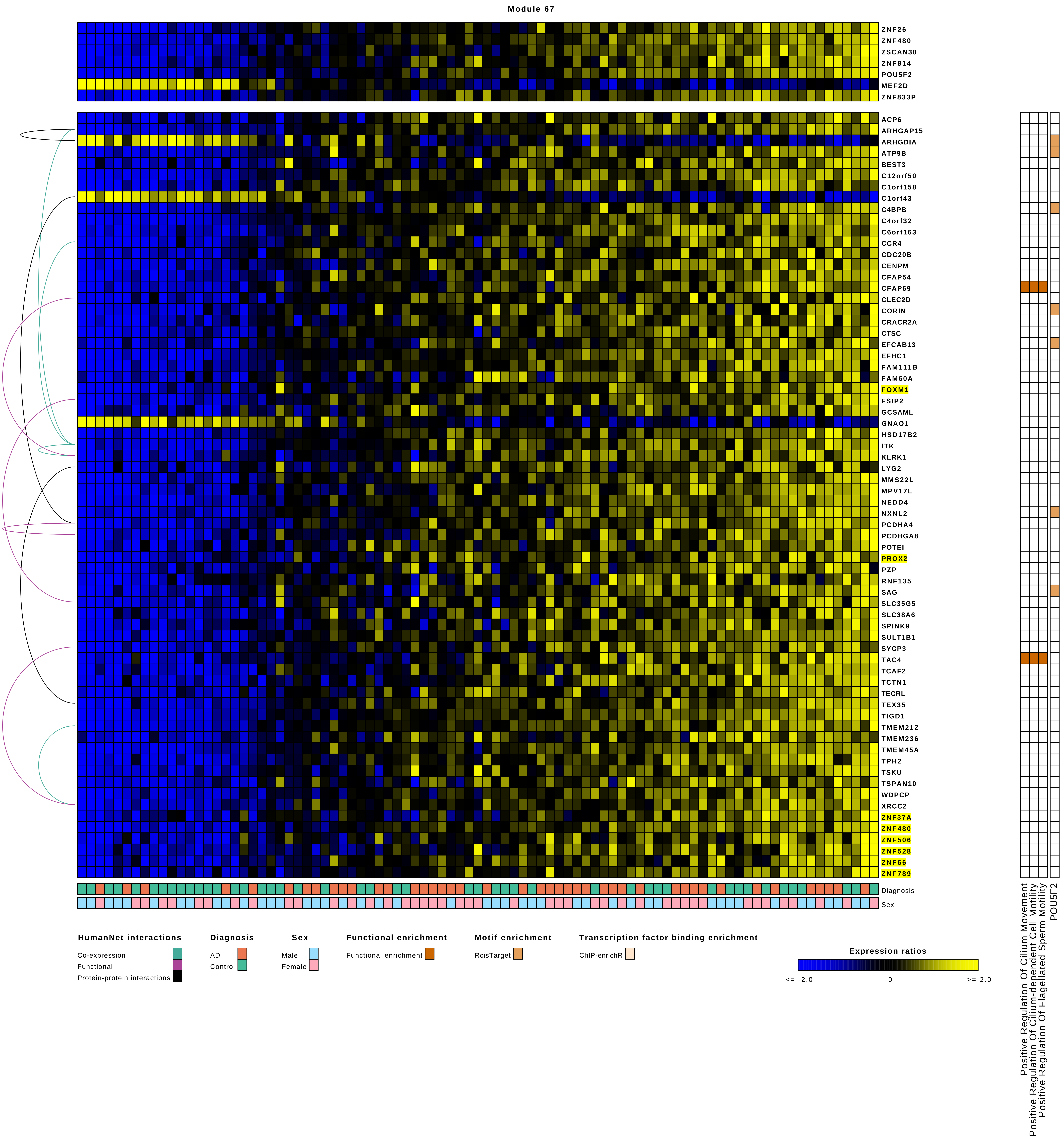

Module 67
ZNF26
ZNF480
ZSCAN30
ZNF814
POU5F2
MEF2D
ZNF833P
ACP6
ARHGAP15
ARHGDIA
ATP9B
BEST3
C12orf50
C1orf158
C1orf43
C4BPB
C4orf32
C6orf163
CCR4
CDC20B
CENPM
CFAP54
CFAP69
CLEC2D
CORIN
CRACR2A
CTSC
EFCAB13
EFHC1
FAM111B
FAM60A
FOXM1
FSIP2
GCSAML
GNAO1
HSD17B2
ITK
KLRK1
LYG2
MMS22L
MPV17L
NEDD4
NXNL2
PCDHA4
PCDHGA8
POTEI
PROX2
PZP
RNF135
SAG
SLC35G5
SLC38A6
SPINK9
SULT1B1
SYCP3
TAC4
TCAF2
TCTN1
TECRL
TEX35
TIGD1
TMEM212
TMEM236
TMEM45A
TPH2
TSKU
TSPAN10
WDPCP
XRCC2
ZNF37A
ZNF480
ZNF506
ZNF528
ZNF66
ZNF789
Diagnosis
Sex
Positive Regulation Of Cilium Movement
Positive Regulation Of Cilium-dependent Cell Motility
Positive Regulation Of Flagellated Sperm Motility
POU5F2
HumanNet interactions
Co-expression
Functional
Protein-protein interactions
Diagnosis
AD
Control
Sex
Male
Female
Functional enrichment
Functional enrichment
Motif enrichment
RcisTarget
Transcription factor binding enrichment
ChIP-enrichR
Expression ratios
<= -2.0
-0
>= 2.0

Module 68

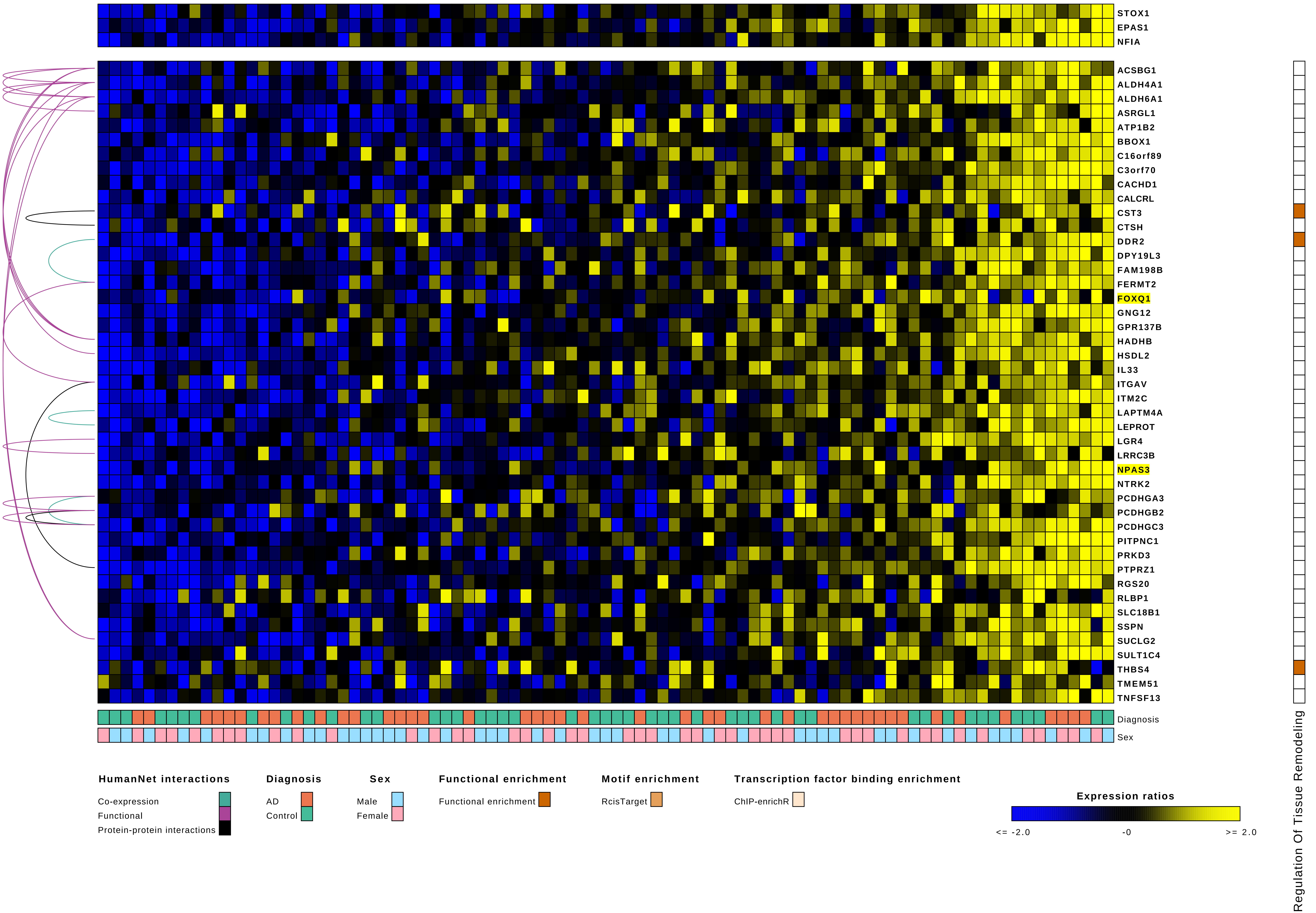

STOX1
EPAS1
NFIA
ACSBG1
ALDH4A1
ALDH6A1
ASRGL1
ATP1B2
BBOX1
C16orf89
C3orf70
CACHD1
CALCRL
CST3
CTSH
DDR2
DPY19L3
FAM198B
FERMT2
FOXQ1
GNG12
GPR137B
HADHB
HSDL2
IL33
ITGAV
ITM2C
LAPTM4A
LEPROT
LGR4
LRRC3B
NPAS3
NTRK2
PCDHGA3
PCDHGB2
PCDHGC3
PITPNC1
PRKD3
PTPRZ1
RGS20
RLBP1
SLC18B1
SSPN
SUCLG2
SULT1C4
THBS4
TMEM51
TNFSF13
Diagnosis
Sex
Regulation Of Tissue Remodeling
HumanNet interactions
Co-expression
Functional
Protein-protein interactions
Diagnosis
AD
Control
Sex
Male
Female
Functional enrichment
Functional enrichment
Motif enrichment
RcisTarget
Transcription factor binding enrichment
ChIP-enrichR
Expression ratios
<= -2.0
-0
>= 2.0

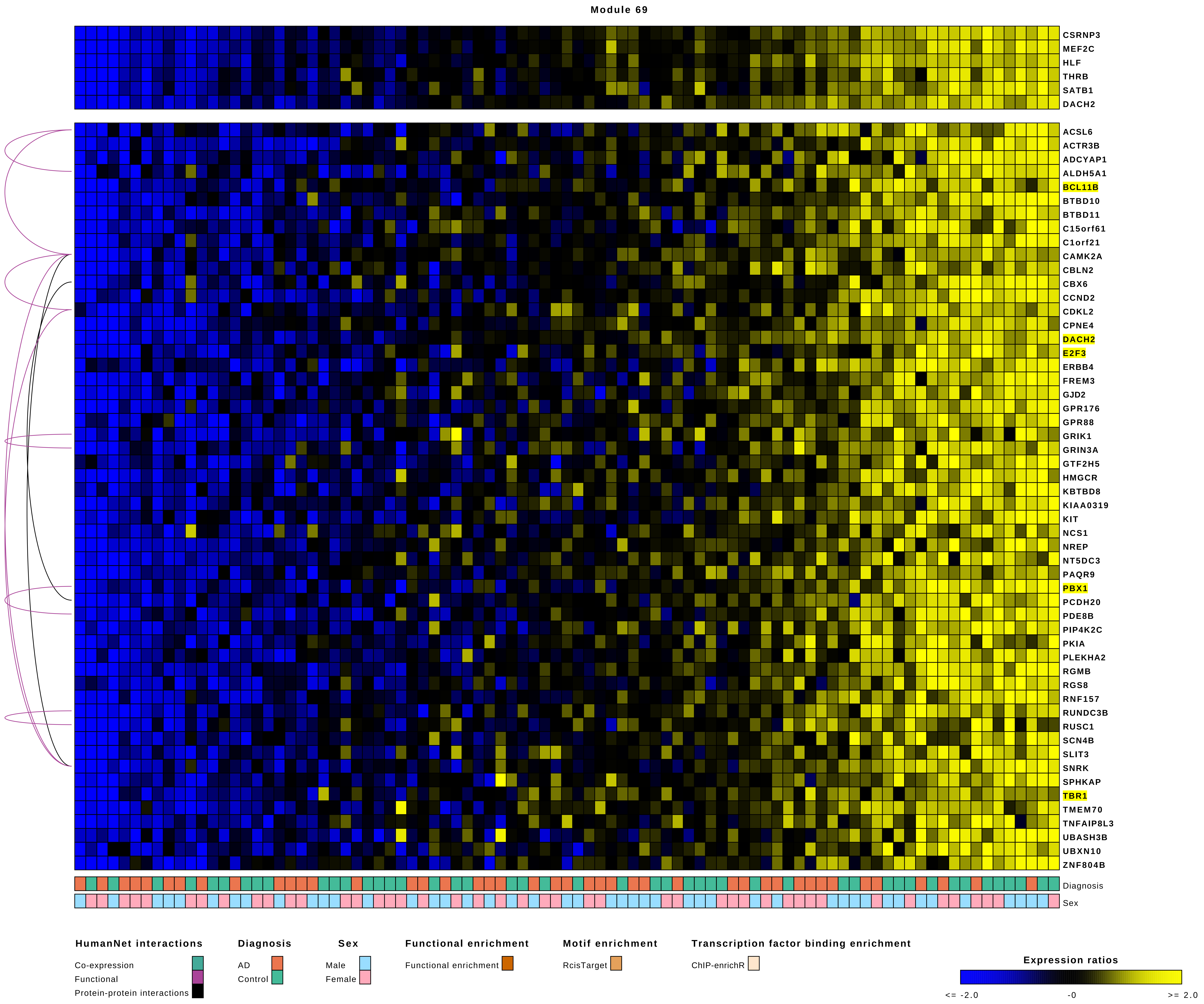
Module 69
CSRNP3
MEF2C
HLF
THRB
SATB1
DACH2
ACSL6
ACTR3B
ADCYAP1
ALDH5A1
BCL11B
BTBD10
BTBD11
C15orf61
C1orf21
CAMK2A
CBLN2
CBX6
CCND2
CDKL2
CPNE4
DACH2
E2F3
ERBB4
FREM3
GJD2
GPR176
GPR88
GRIK1
GRIN3A
GTF2H5
HMGCR
KBTBD8
KIAA0319
KIT
NCS1
NREP
NT5DC3
PAQR9
PBX1
PCDH20
PDE8B
PIP4K2C
PKIA
PLEKHA2
RGMB
RGS8
RNF157
RUNDC3B
RUSC1
SCN4B
SLIT3
SNRK
SPHKAP
TBR1
TMEM70
TNFAIP8L3
UBASH3B
UBXN10
ZNF804B
Diagnosis
Sex
HumanNet interactions
Co-expression
Functional
Protein-protein interactions
Diagnosis
AD
Control
Sex
Male
Female
Functional enrichment
Functional enrichment
Motif enrichment
RcisTarget
Transcription factor binding enrichment
ChIP-enrichR
Expression ratios
<= -2.0
-0
>= 2.0

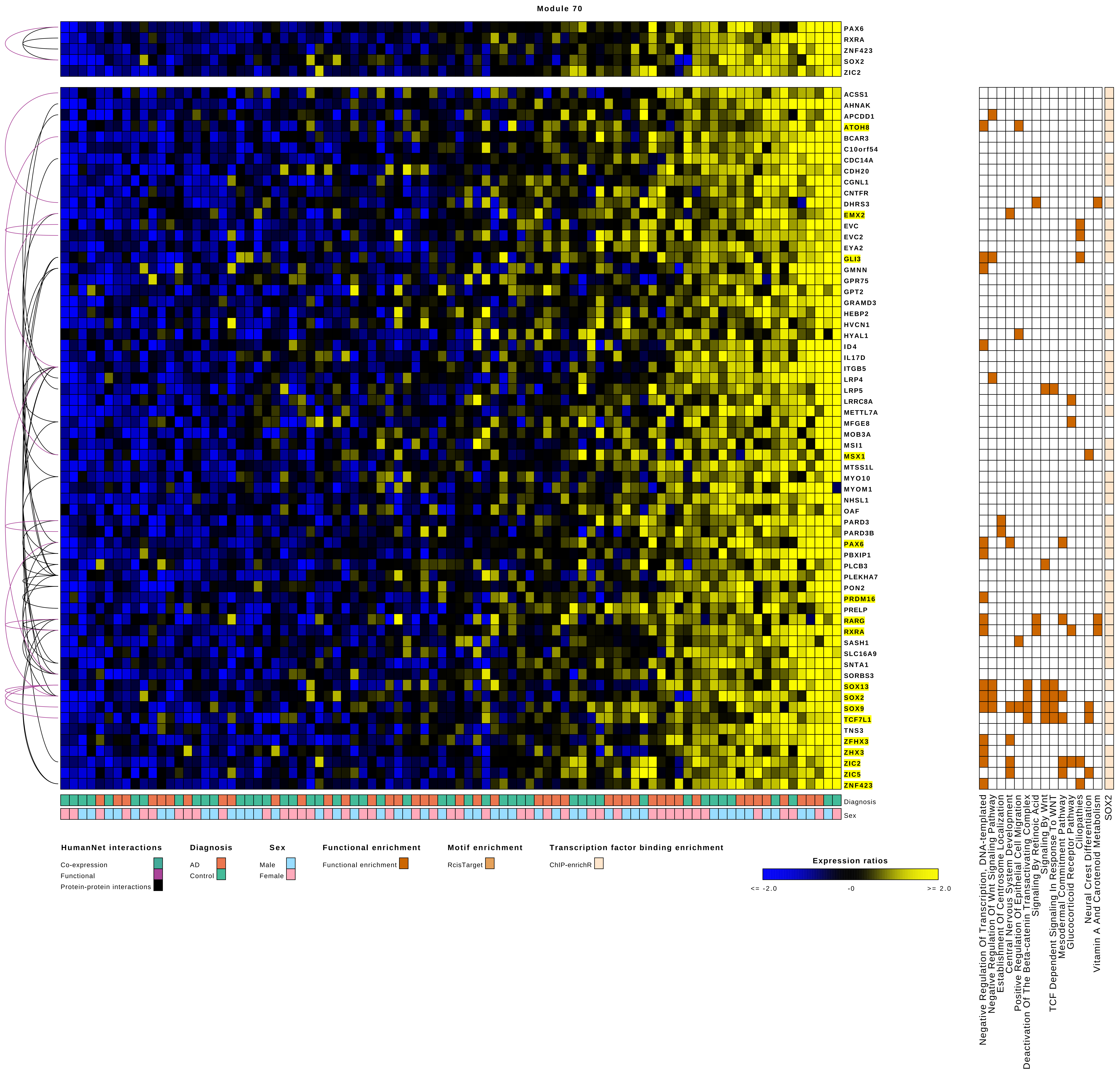

Module 70
PAX6
RXRA
ZNF423
SOX2
ZIC2
ACSS1
AHNAK
APCDD1
ATOH8
BCAR3
C10orf54
CDC14A
CDH20
CGNL1
CNTFR
DHRS3
EMX2
EVC
EVC2
EYA2
GLI3
GMNN
GPR75
GPT2
GRAMD3
HEBP2
HVCN1
HYAL1
ID4
IL17D
ITGB5
LRP4
LRP5
LRRC8A
METTL7A
MFGE8
MOB3A
MSI1
MSX1
MTSS1L
MYO10
MYOM1
NHSL1
OAF
PARD3
PARD3B
PAX6
PBXIP1
PLCB3
PLEKHA7
PON2
PRDM16
PRELP
RARG
RXRA
SASH1
SLC16A9
SNTA1
SORBS3
SOX13
SOX2
SOX9
TCF7L1
TNS3
ZFHX3
ZHX3
ZIC2
ZIC5
ZNF423
Diagnosis
Sex
Negative Regulation Of Transcription, DNA-templated
Negative Regulation Of Wnt Signaling Pathway
Establishment Of Centrosome Localization
Central Nervous System Development
Positive Regulation Of Epithelial Cell Migration
Deactivation Of The Beta-catenin Transactivating Complex
Signaling By Retinoic Acid
Signaling By WNT
TCF Dependent Signaling In Response To WNT
Mesodermal Commitment Pathway
Glucocorticoid Receptor Pathway
Ciliopathies
Neural Crest Differentiation
Vitamin A And Carotenoid Metabolism
SOX2
HumanNet interactions
Co-expression
Functional
Protein-protein interactions
Diagnosis
AD
Control
Sex
Male
Female
Functional enrichment
Functional enrichment
Motif enrichment
RcisTarget
Transcription factor binding enrichment
ChIP-enrichR
Expression ratios
<= -2.0
-0
>= 2.0

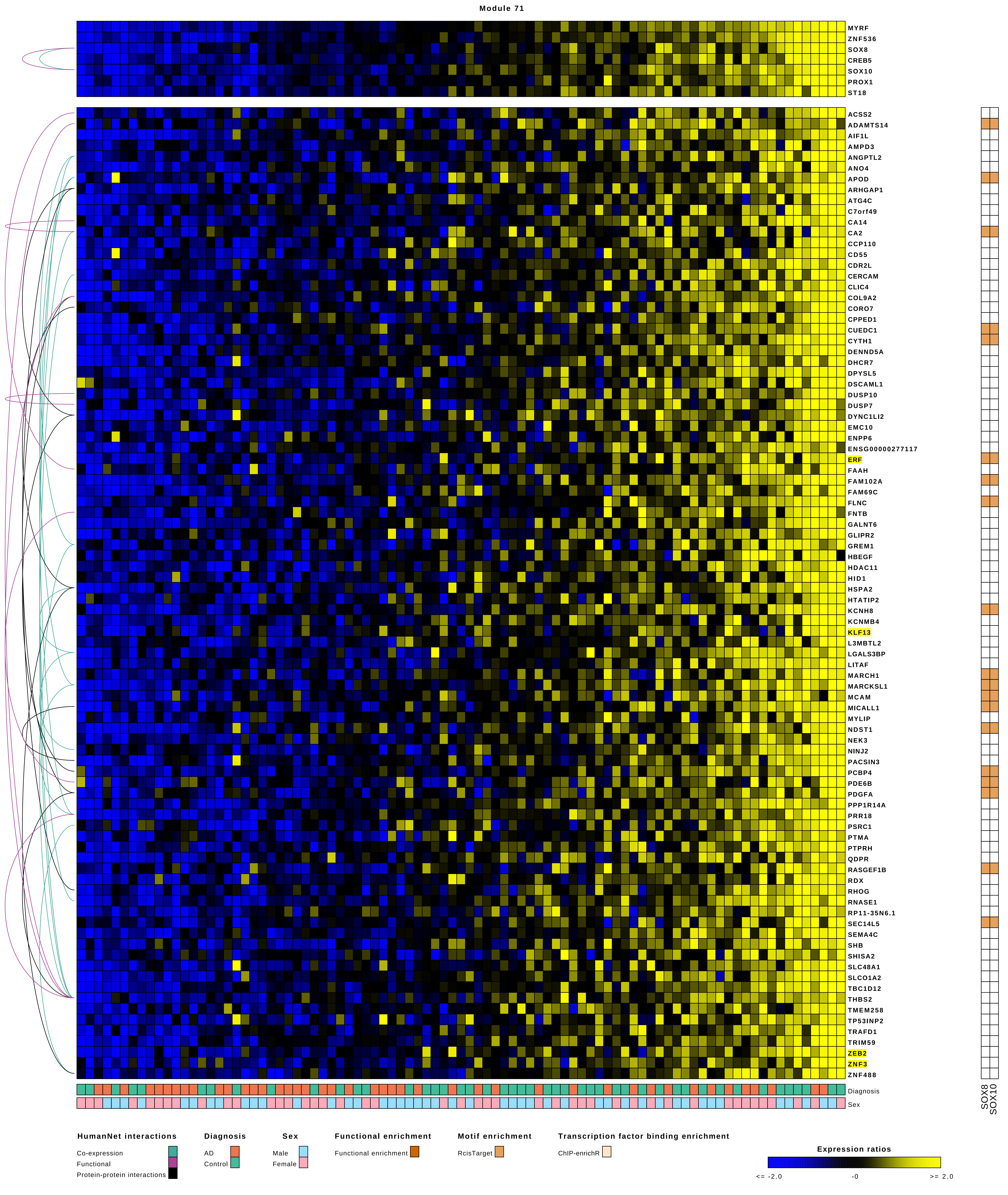

Module 71
MYRF
ZNF536
SOX8
CREB5
SOX10
PROX1
ST18
ACSS2
ADAMTS14
AIF1L
AMPD3
ANGPTL2
ANO4
APOD
ARHGAP1
ATG4C
C7orf49
CA14
CA2
CCP110
CD55
CDR2L
CERCAM
CLIC4
COL9A2
CORO7
CPPED1
CUEDC1
CYTH1
DENND5A
DHCR7
DPYSL5
DSCAML1
DUSP10
DUSP7
DYNC1LI2
EMC10
ENPP6
ENSG00000277117
ERF
FAAH
FAM102A
FAM69C
FLNC
FNTB
GALNT6
GLIPR2
GREM1
HBEGF
HDAC11
HID1
HSPA2
HTATIP2
KCNH8
KCNMB4
KLF13
L3MBTL2
LGALS3BP
LITAF
MARCH1
MARCKSL1
MCAM
MICALL1
MYLIP
NDST1
NEK3
NINJ2
PACSIN3
PCBP4
PDE6B
PDGFA
PPP1R14A
PRR18
PSRC1
PTMA
PTPRH
QDPR
RASGEF1B
RDX
RHOG
RNASE1
RP11-35N6.1
SEC14L5
SEMA4C
SHB
SHISA2
SLC48A1
SLCO1A2
TBC1D12
THBS2
TMEM258
TP53INP2
TRAFD1
TRIM59
ZEB2
ZNF3
ZNF488
Diagnosis
Sex
SOX8
SOX10
HumanNet interactions
Co-expression
Functional
Protein-protein interactions
Diagnosis
AD
Control
Sex
Male
Female
Functional enrichment
Functional enrichment
Motif enrichment
RcisTarget
Transcription factor binding enrichment
ChIP-enrichR
Expression ratios
<= -2.0
-0
>= 2.0

Module 72

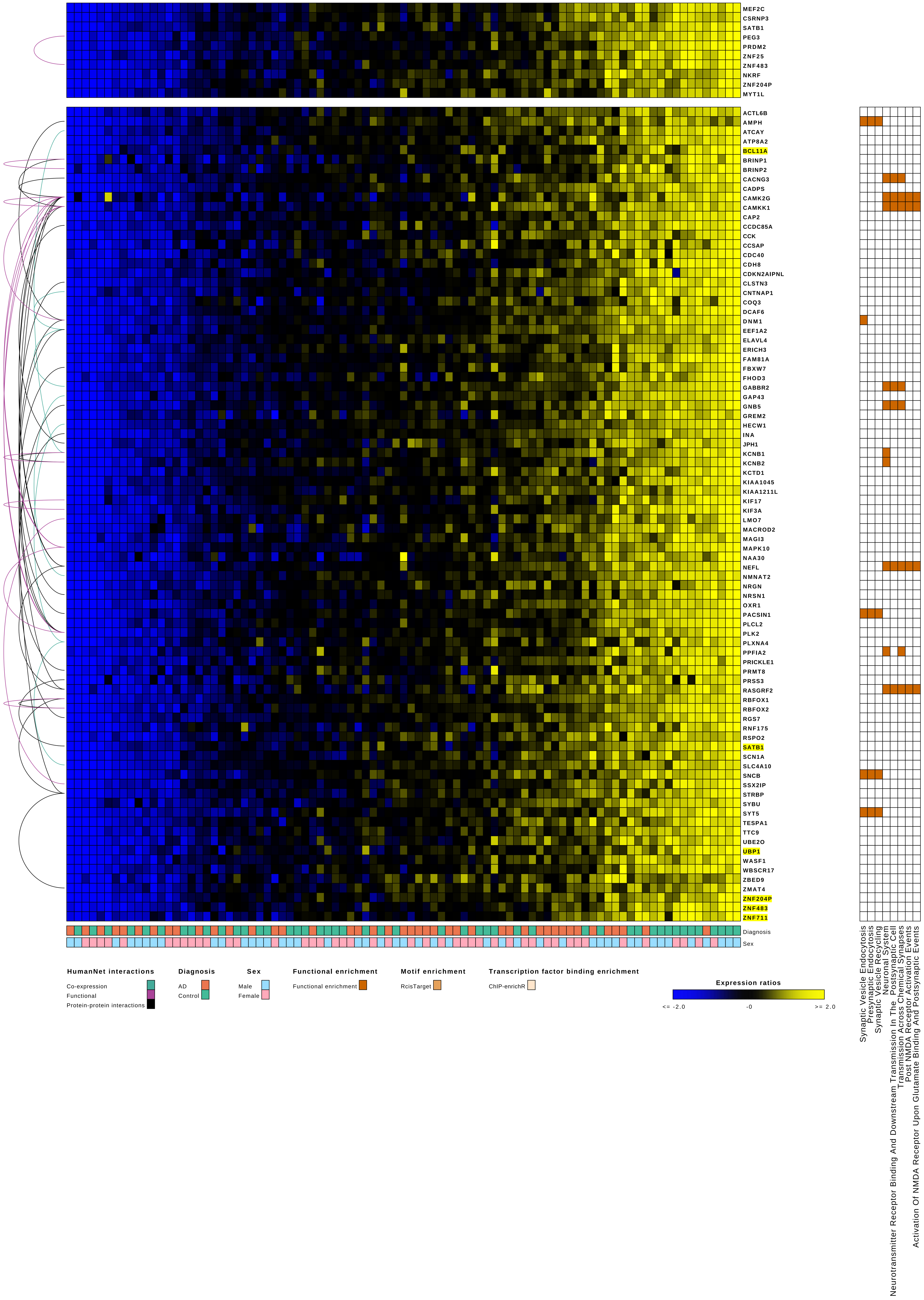
MEF2C
CSRNP3
SATB1
PEG3
PRDM2
ZNF25
ZNF483
NKRF
ZNF204P
MYT1L
ACTL6B
AMPH
ATCAY
ATP8A2
BCL11A
BRINP1
BRINP2
CACNG3
CADPS
CAMK2G
CAMKK1
CAP2
CCDC85A
CCK
CCSAP
CDC40
CDH8
CDKN2AIPNL
CLSTN3
CNTNAP1
COQ3
DCAF6
DNM1
EEF1A2
ELAVL4
ERICH3
FAM81A
FBXW7
FHOD3
GABBR2
GAP43
GNB5
GREM2
HECW1
INA
JPH1
KCNB1
KCNB2
KCTD1
KIAA1045
KIAA1211L
KIF17
KIF3A
LMO7
MACROD2
MAGI3
MAPK10
NAA30
NEFL
NMNAT2
NRGN
NRSN1
OXR1
PACSIN1
PLCL2
PLK2
PLXNA4
PPFIA2
PRICKLE1
PRMT8
PRSS3
RASGRF2
RBFOX1
RBFOX2
RGS7
RNF175
RSPO2
SATB1
SCN1A
SLC4A10
SNCB
SSX2IP
STRBP
SYBU
SYT5
TESPA1
TTC9
UBE2O
UBP1
WASF1
WBSCR17
ZBED9
ZMAT4
ZNF204P
ZNF483
ZNF711
Diagnosis
Sex
HumanNet interactions
Co-expression
Functional
Protein-protein interactions
Diagnosis
AD
Control
Sex
Male
Female
Functional enrichment
Functional enrichment
Motif enrichment
RcisTarget
Transcription factor binding enrichment
ChIP-enrichR
Expression ratios
<= -2.0
-0
>= 2.0
Synaptic Vesicle Endocytosis
Presynaptic Endocytosis
Synaptic Vesicle Recycling
Neuronal System
Neurotransmitter Receptor Binding And Downstream Transmission In The Postsynaptic Cell
Transmission Across Chemical Synapses
Post NMDA Receptor Activation Events
Activation Of NMDA Receptor Upon Glutamate Binding And Postsynaptic Events

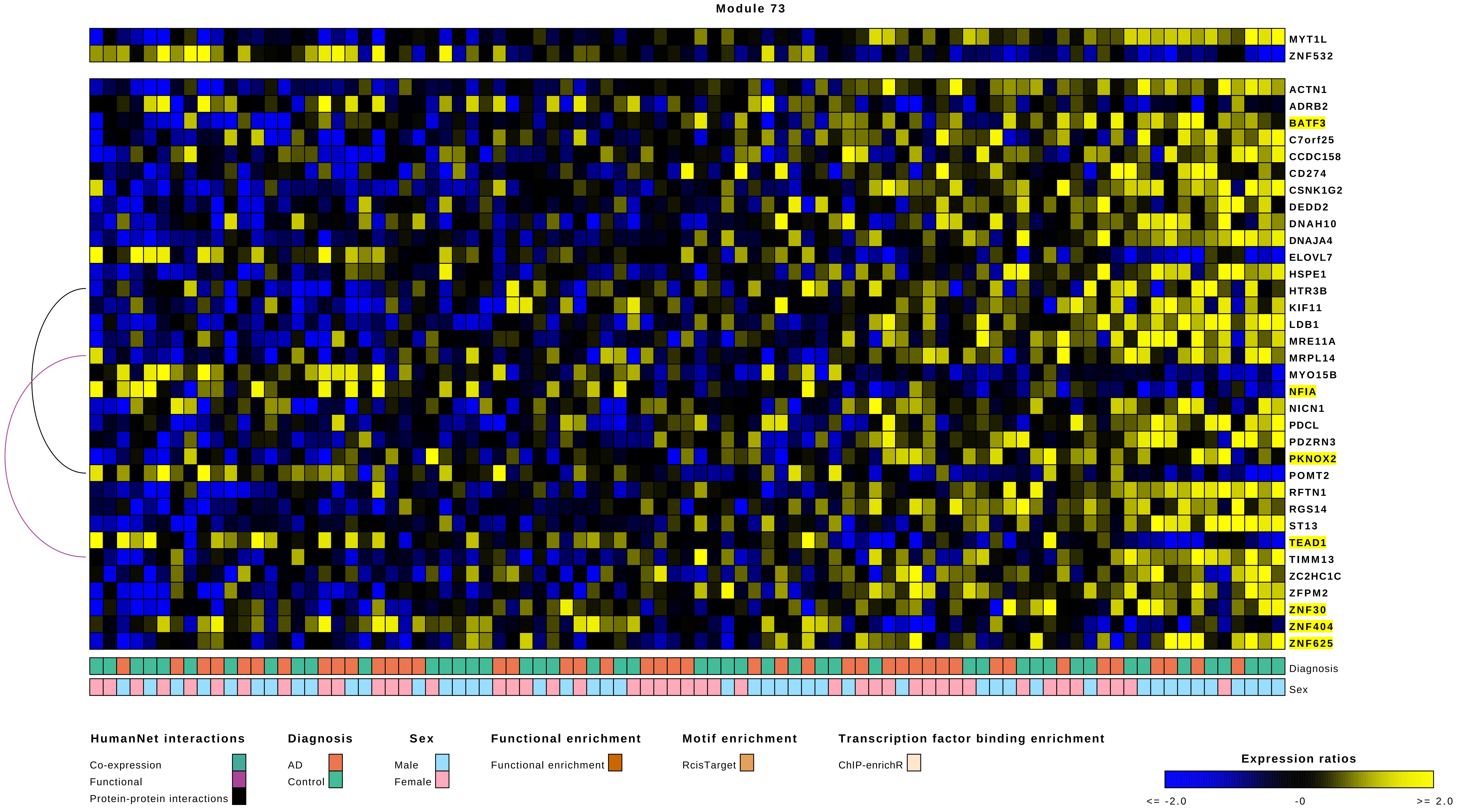

Module 73
MYT1L
ZNF532
ACTN1
ADRB2
BATF3
C7orf25
CCDC158
CD274
CSNK1G2
DEDD2
DNAH10
DNAJA4
ELOVL7
HSPE1
HTR3B
KIF11
LDB1
MRE11A
MRPL14
MYO15B
NFIA
NICN1
PDCL
PDZRN3
PKNOX2
POMT2
RFTN1
RGS14
ST13
TEAD1
TIMM13
ZC2HC1C
ZFPM2
ZNF30
ZNF404
ZNF625
Diagnosis
Sex
HumanNet interactions
Co-expression
Functional
Protein-protein interactions
Diagnosis
AD
Control
Sex
Male
Female
Functional enrichment
Functional enrichment
Motif enrichment
RcisTarget
Transcription factor binding enrichment
ChIP-enrichR
Expression ratios
<= -2.0
-0
>= 2.0

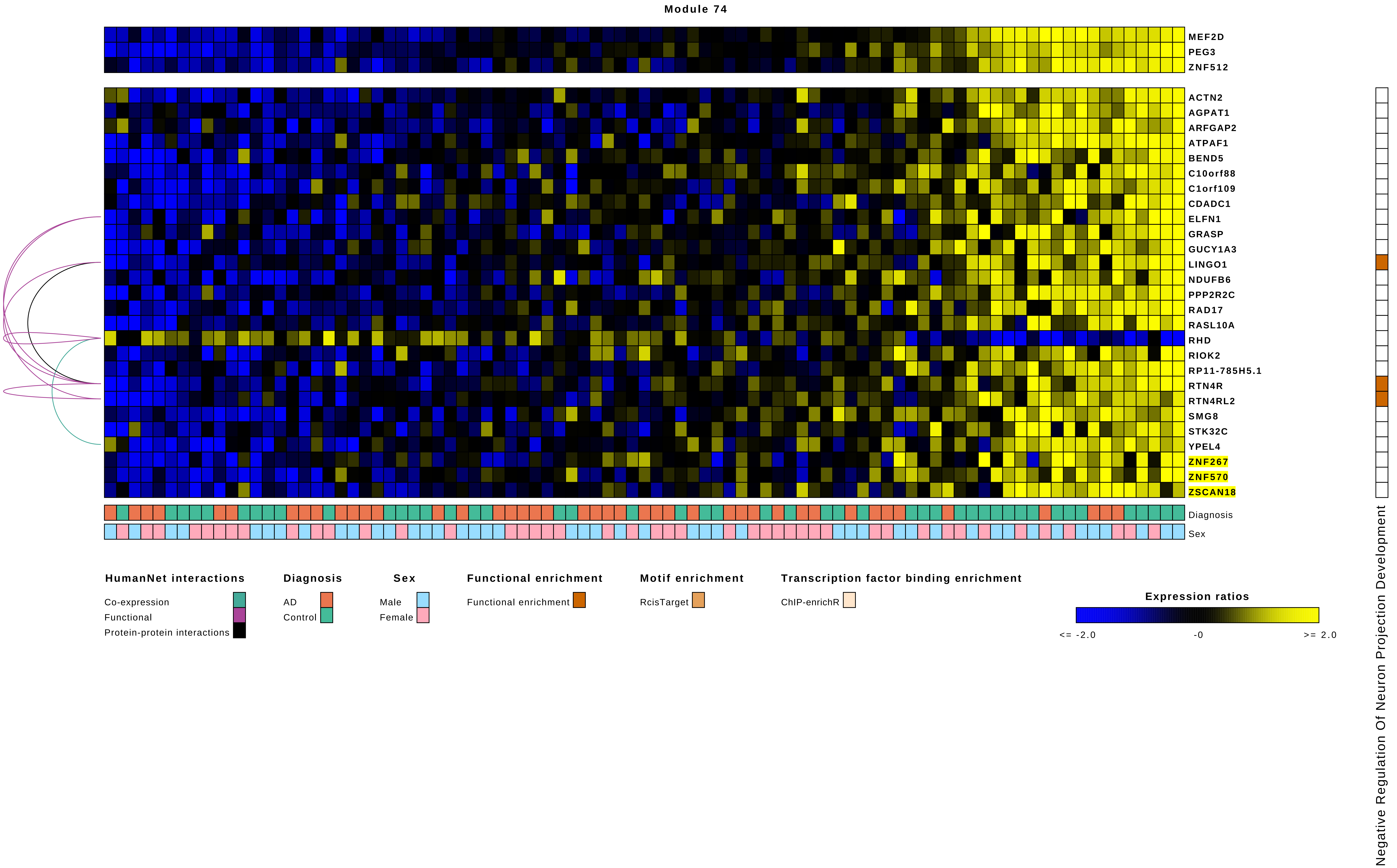
Module 74
MEF2D
PEG3
ZNF512
ACTN2
AGPAT1
ARFGAP2
ATPAF1
BEND5
C10orf88
C1orf109
CDADC1
ELFN1
GRASP
GUCY1A3
LINGO1
NDUFB6
PPP2R2C
RAD17
RASL10A
RHD
RIOK2
RP11-785H5.1
RTN4R
RTN4RL2
SMG8
STK32C
YPEL4
ZNF267
ZNF570
ZSCAN18
Diagnosis
Sex
Negative Regulation Of Neuron Projection Development
HumanNet interactions
Co-expression
Functional
Protein-protein interactions
Diagnosis
AD
Control
Sex
Male
Female
Functional enrichment
Functional enrichment
Motif enrichment
RcisTarget
Transcription factor binding enrichment
ChIP-enrichR
Expression ratios
<= -2.0
-0
>= 2.0

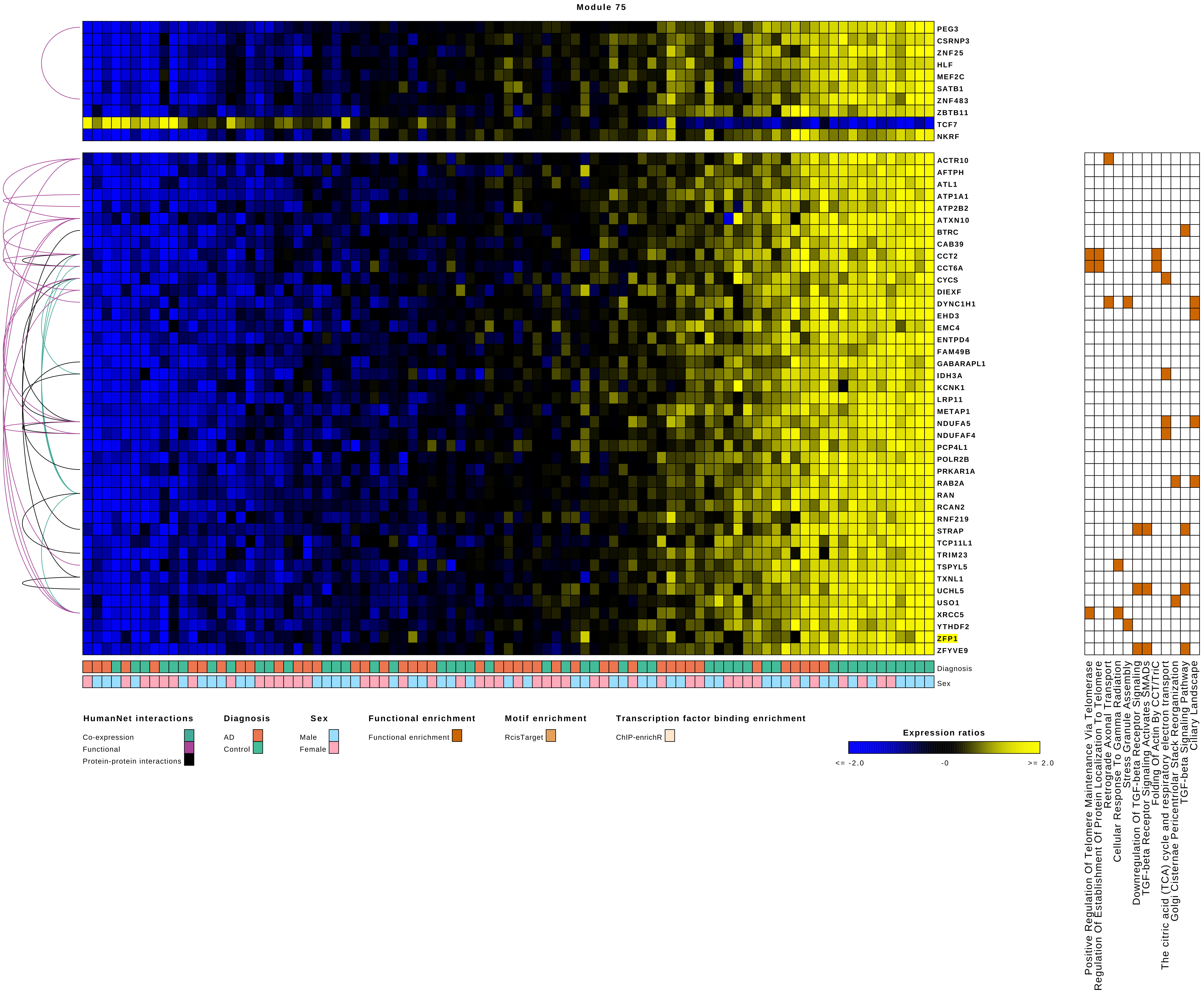
Module 75
PEG3
CSRNP3
ZNF25
HLF
MEF2C
SATB1
ZNF483
ZBTB11
TCF7
NKRF
ACTR10
AFTPH
ATL1
ATP1A1
ATP2B2
ATXN10
BTRC
CAB39
CCT2
CCT6A
CYCS
DIEXF
DYNC1H1
EHD3
EMC4
ENTPD4
FAM49B
GABARAPL1
IDH3A
KCNK1
LRP11
METAP1
NDUFA5
NDUFAF4
PCP4L1
POLR2B
PRKAR1A
RAB2A
RAN
RCAN2
RNF219
STRAP
TCP11L1
TRIM23
TSPYL5
TXNL1
UCHL5
USO1
XRCC5
YTHDF2
ZFP1
ZFYVE9
Diagnosis
Sex
HumanNet interactions
Co-expression
Functional
Protein-protein interactions
Diagnosis
AD
Control
Sex
Male
Female
Functional enrichment
Functional enrichment
Motif enrichment
RcisTarget
Transcription factor binding enrichment
ChIP-enrichR
Expression ratios
<= -2.0
-0
>= 2.0
Positive Regulation Of Telomere Maintenance Via Telomerase
Regulation Of Establishment Of Protein Localization To Telomere
Retrograde Axonal Transport
Cellular Response To Gamma Radiation
Stress Granule Assembly
Downregulation Of TGF-beta Receptor Signaling
TGF-beta Receptor Signaling Activates SMADS
Folding Of Actin By CCT/TriC
The citric acid (TCA) cycle and respiratory electron transport
Golgi Cisternae Pericentriolar Stack Reorganization
TGF-beta Signaling Pathway
Ciliary Landscape

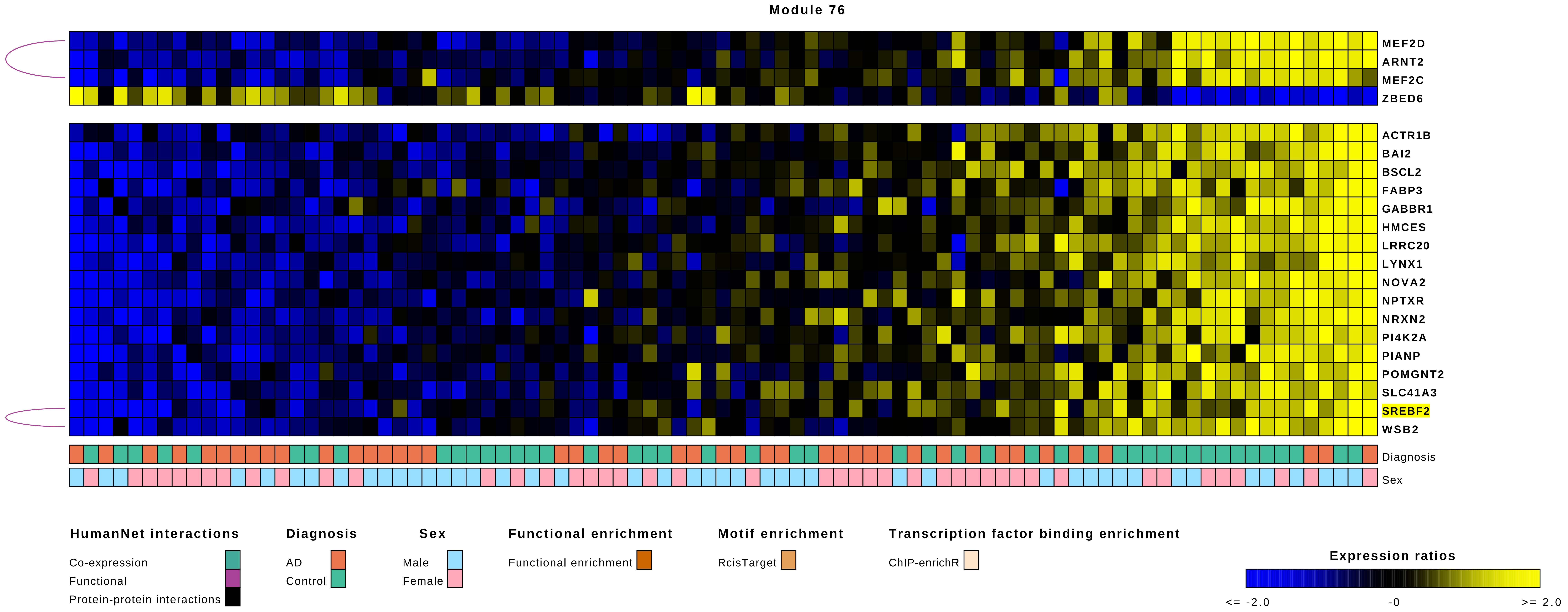

Module 76
MEF2D
ARNT2
MEF2C
ZBED6
ACTR1B
BAI2
BSCL2
FABP3
GABBR1
HMCES
LRRC20
LYNX1
NOVA2
NPTXR
NRXN2
PI4K2A
PIANP
POMGNT2
SLC41A3
SREBF2
WSB2
Diagnosis
Sex
HumanNet interactions
Co-expression
Functional
Protein-protein interactions
Diagnosis
AD
Control
Sex
Male
Female
Functional enrichment
Functional enrichment
Motif enrichment
RcisTarget
Transcription factor binding enrichment
ChIP-enrichR
Expression ratios
<= -2.0
-0
>= 2.0

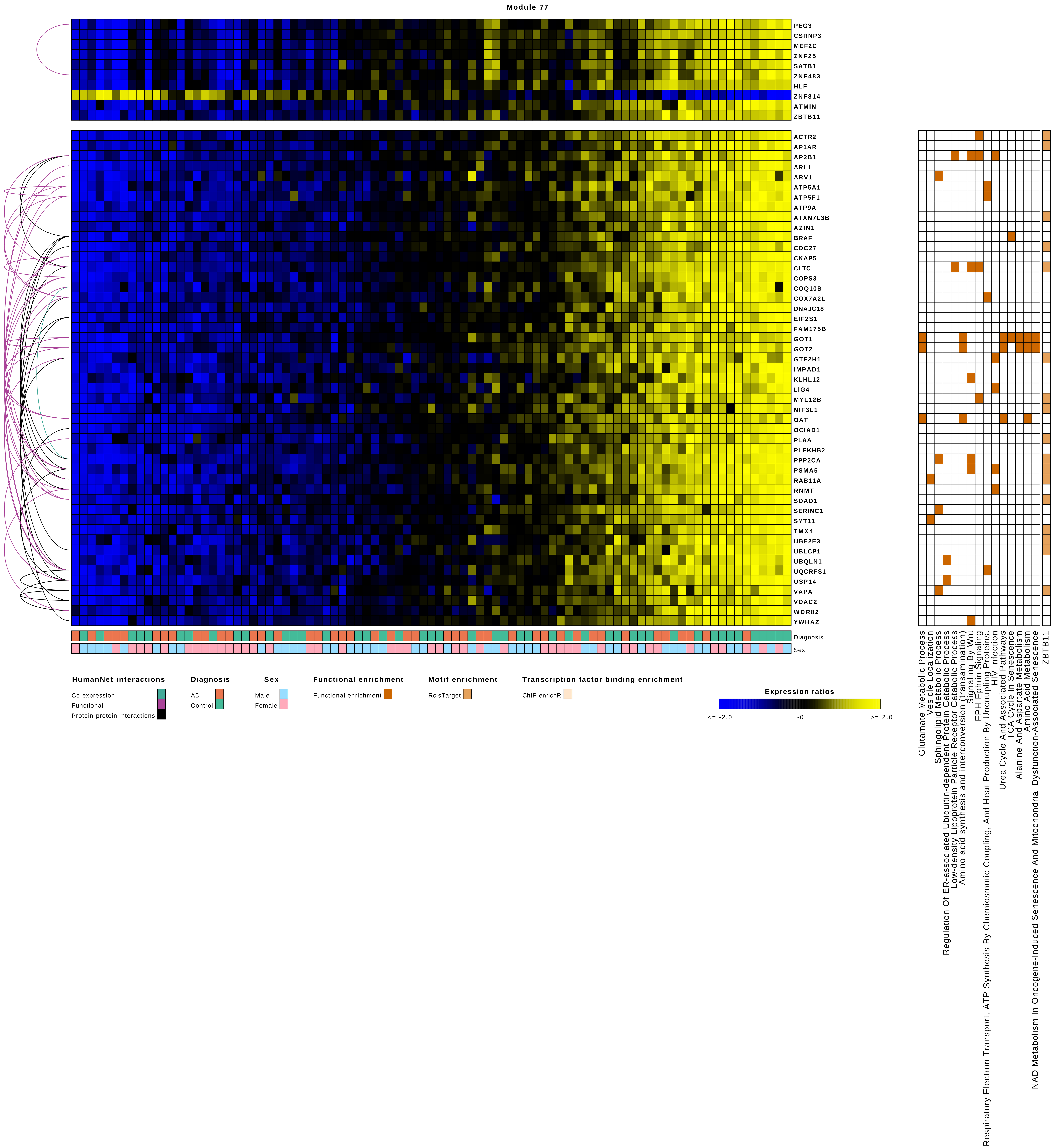

Module 77
PEG3
CSRNP3
MEF2C
ZNF25
SATB1
ZNF483
HLF
ZNF814
ATMIN
ZBTB11
ACTR2
AP1AR
AP2B1
ARL1
ARV1
ATP5A1
ATP5F1
ATP9A
ATXN7L3B
AZIN1
BRAF
CDC27
CKAP5
CLTC
COPS3
COQ10B
COX7A2L
DNAJC18
EIF2S1
FAM175B
GOT1
GOT2
GTF2H1
IMPAD1
KLHL12
LIG4
MYL12B
NIF3L1
OAT
OCIAD1
PLAA
PLEKHB2
PPP2CA
PSMA5
RAB11A
RNMT
SDAD1
SERINC1
SYT11
TMX4
UBE2E3
UBLCP1
UBQLN1
UQCRFS1
USP14
VAPA
VDAC2
WDR82
YWHAZ
Diagnosis
Sex
Glutamate Metabolic Process
Vesicle Localization
Sphingolipid Metabolic Process
Regulation Of ER-associated Ubiquitin-dependent Protein Catabolic Process
Low-density Lipoprotein Particle Receptor Catabolic Process
Amino acid synthesis and interconversion (transamination)
Signaling By Wnt
EPH-Ephrin Signaling
Respiratory Electron Transport, ATP Synthesis By Chemiosmotic Coupling, And Heat Production By Uncoupling Proteins.
HIV Infection
Urea Cycle And Associated Pathways
TCA Cycle In Senescence
Alanine And Aspartate Metabolism
Amino Acid Metabolism
NAD Metabolism In Oncogene-Induced Senescence And Mitochondrial Dysfunction-Associated Senescence
ZBTB11
HumanNet interactions
Co-expression
Functional
Protein-protein interactions
Diagnosis
AD
Control
Sex
Male
Female
Functional enrichment
Functional enrichment
Motif enrichment
RcisTarget
Transcription factor binding enrichment
ChIP-enrichR
Expression ratios
<= -2.0
-0
>= 2.0

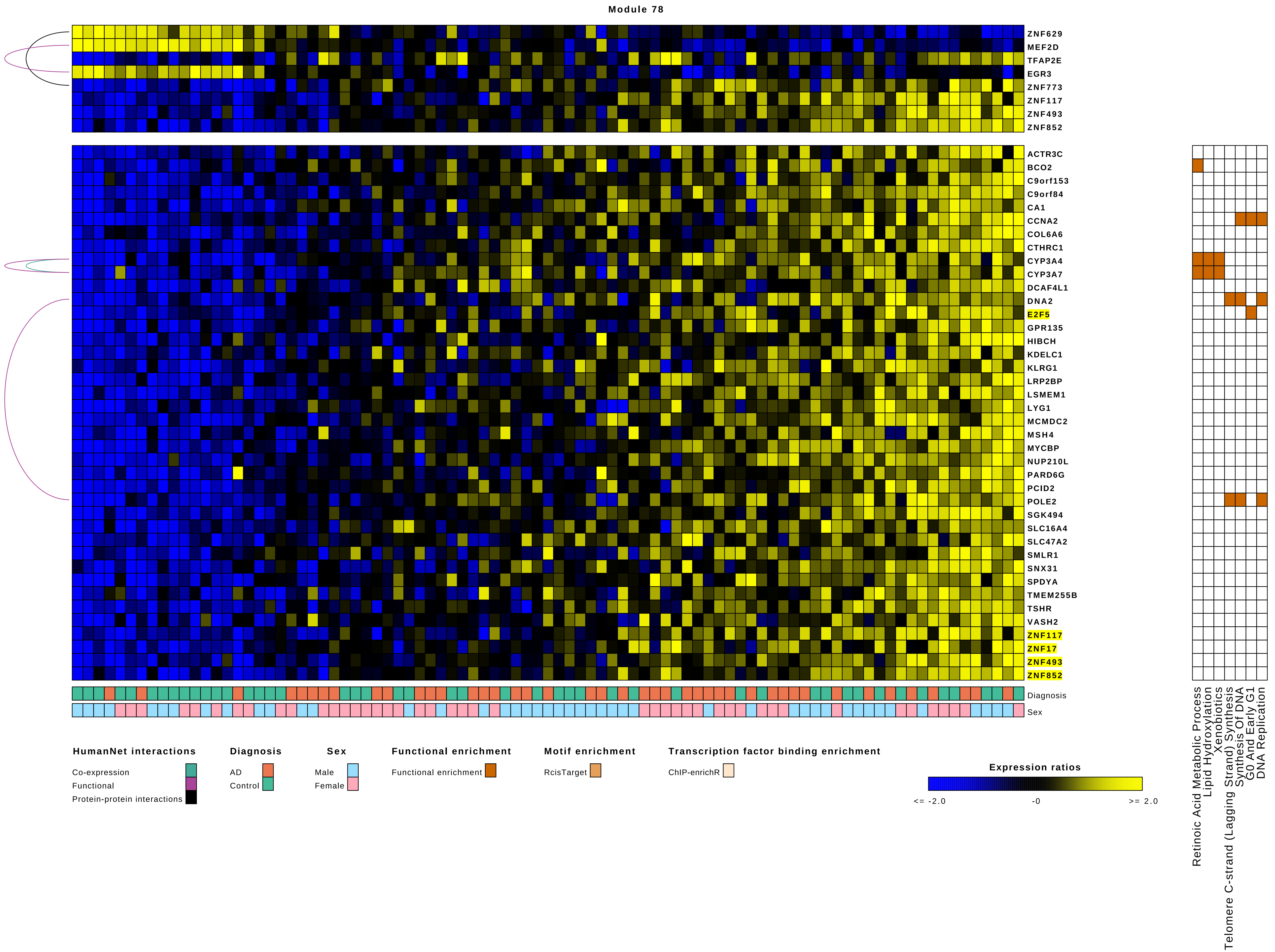

Module 78
ZNF629
MEF2D
TFAP2E
EGR3
ZNF773
ZNF117
ZNF493
ZNF852
ACTR3C
BCO2
C9orf153
C9orf84
CA1
CCNA2
COL6A6
CTHRC1
CYP3A4
CYP3A7
DCAF4L1
DNA2
E2F5
GPR135
HIBCH
KDELC1
KLRG1
LRP2BP
LSMEM1
LYG1
MCMDC2
MSH4
MYCBP
NUP210L
PARD6G
PCID2
POLE2
SGK494
SLC16A4
SLC47A2
SMLR1
SNX31
SPDYA
TMEM255B
TSHR
VASH2
ZNF117
ZNF17
ZNF493
ZNF852
Diagnosis
Sex
Retinoic Acid Metabolic Process
Lipid Hydroxylation
Xenobiotics
Telomere C-strand (Lagging Strand) Synthesis
Synthesis Of DNA
G0 And Early G1
DNA Replication
HumanNet interactions
Co-expression
Functional
Protein-protein interactions
Diagnosis
AD
Control
Sex
Male
Female
Functional enrichment
Functional enrichment
Motif enrichment
RcisTarget
Transcription factor binding enrichment
ChIP-enrichR
Expression ratios
<= -2.0
-0
>= 2.0

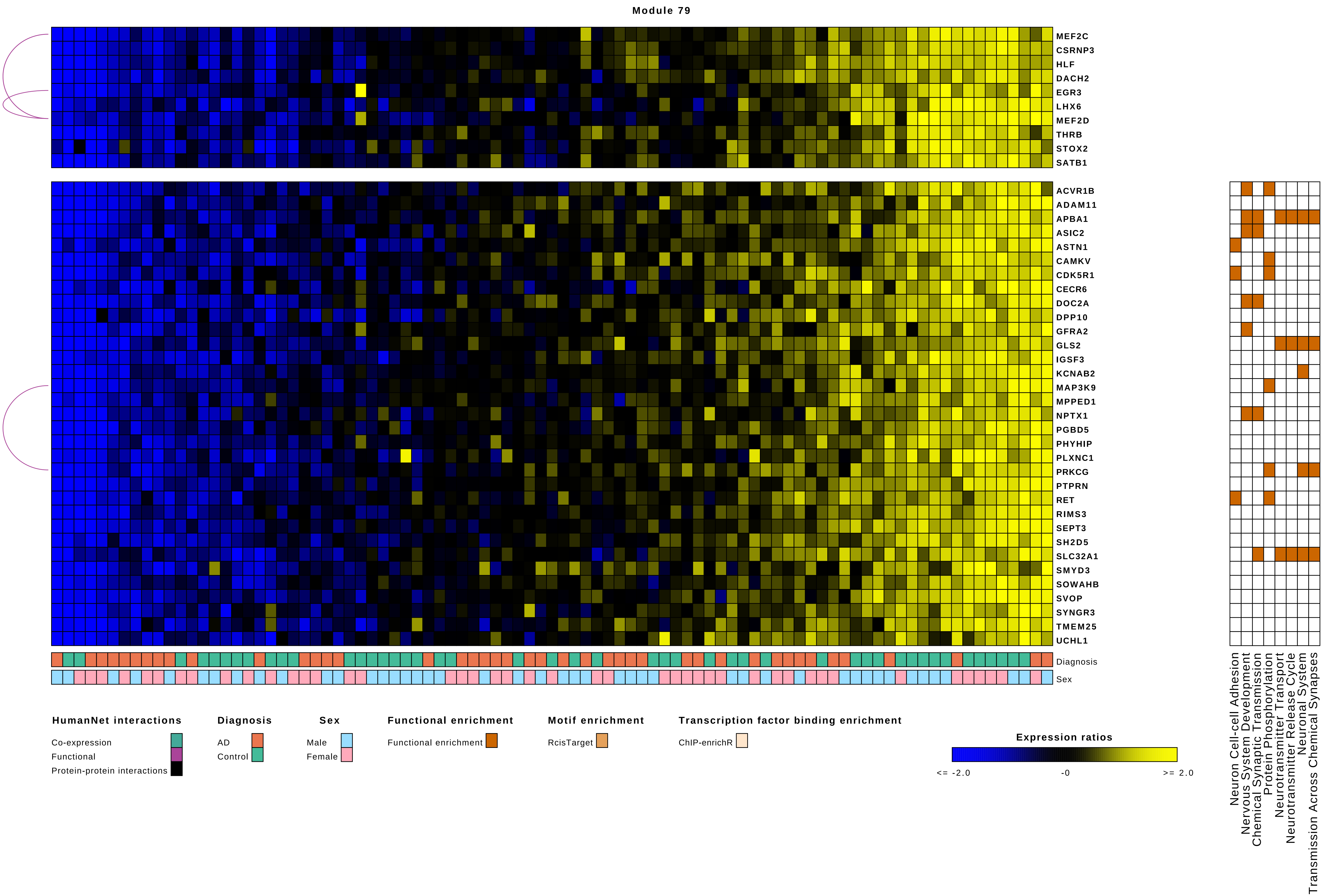

Module 79
MEF2C
CSRNP3
HLF
DACH2
EGR3
LHX6
MEF2D
THRB
STOX2
SATB1
ACVR1B
ADAM11
APBA1
ASIC2
ASTN1
CAMKV
CDK5R1
CECR6
DOC2A
DPP10
GFRA2
GLS2
IGSF3
KCNAB2
MAP3K9
MPPED1
NPTX1
PGBD5
PHYHIP
PLXNC1
PRKCG
PTPRN
RET
RIMS3
SEPT3
SH2D5
SLC32A1
SMYD3
SOWAHB
SVOP
SYNGR3
TMEM25
UCHL1
Diagnosis
Sex
Neuron Cell-cell Adhesion
Nervous System Development
Chemical Synaptic Transmission
Protein Phosphorylation
Neurotransmitter Transport
Neurotransmitter Release Cycle
Neuronal System
Transmission Across Chemical Synapses
HumanNet interactions
Co-expression
Functional
Protein-protein interactions
Diagnosis
AD
Control
Sex
Male
Female
Functional enrichment
Functional enrichment
Motif enrichment
RcisTarget
Transcription factor binding enrichment
ChIP-enrichR
Expression ratios
<= -2.0
-0
>= 2.0

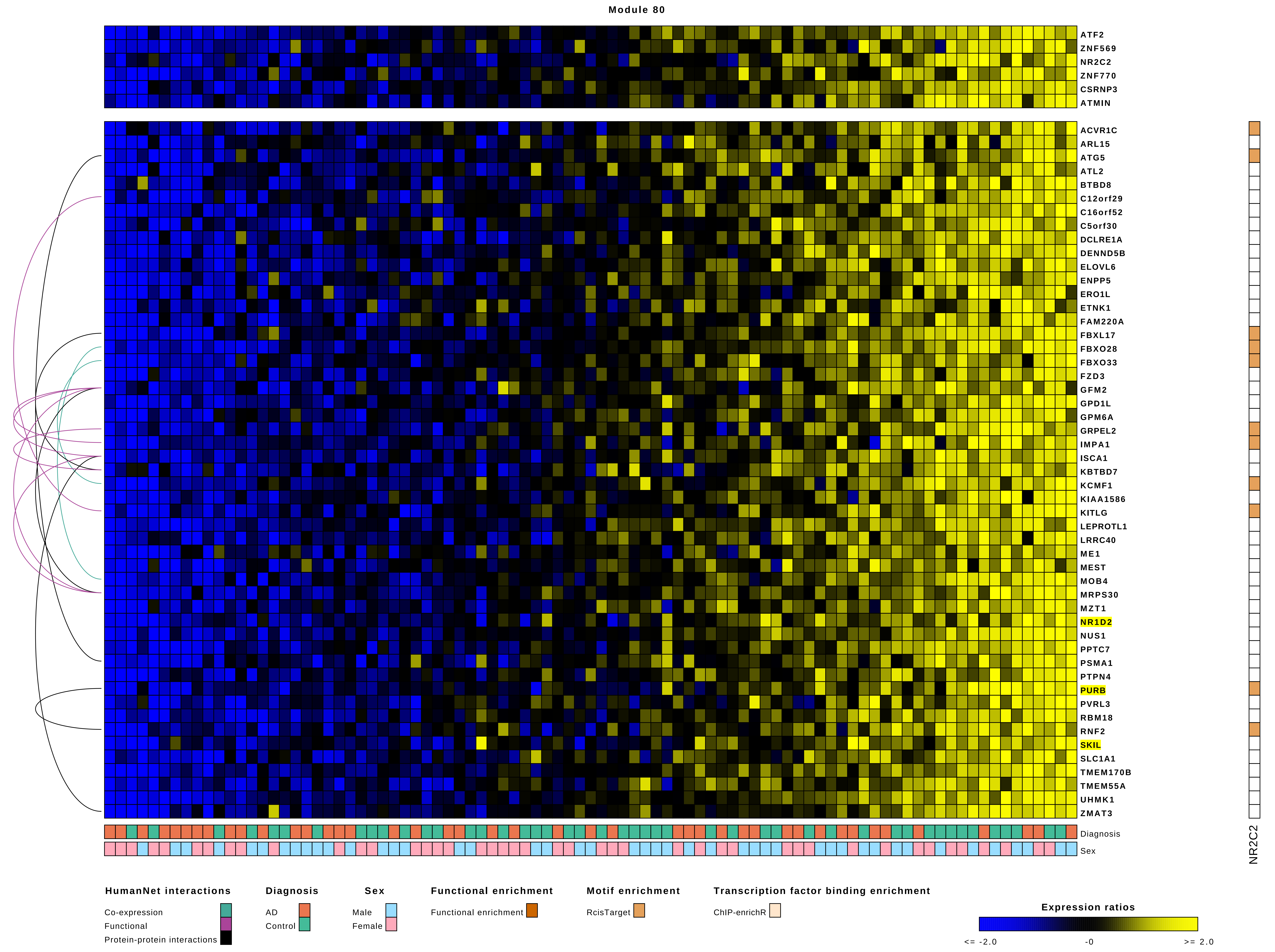

Module 80
ATF2
ZNF569
NR2C2
ZNF770
CSRNP3
ATMIN
ACVR1C
ARL15
ATG5
ATL2
BTBD8
C12orf29
C16orf52
C5orf30
DCLRE1A
DENND5B
ELOVL6
ENPP5
ERO1L
ETNK1
FAM220A
FBXL17
FBXO28
FBXO33
FZD3
GFM2
GPD1L
GPM6A
GRPEL2
IMPA1
ISCA1
KBTBD7
KCMF1
KIAA1586
KITLG
LEPROTL1
LRRC40
ME1
MEST
MOB4
MRPS30
MZT1
NR1D2
NUS1
PPTC7
PSMA1
PTPN4
PURB
PVRL3
RBM18
RNF2
SKIL
SLC1A1
TMEM170B
TMEM55A
UHMK1
ZMAT3
Diagnosis
Sex
NR2C2
HumanNet interactions
Co-expression
Functional
Protein-protein interactions
Diagnosis
AD
Control
Sex
Male
Female
Functional enrichment
Functional enrichment
Motif enrichment
RcisTarget
Transcription factor binding enrichment
ChIP-enrichR
Expression ratios
<= -2.0
-0
>= 2.0

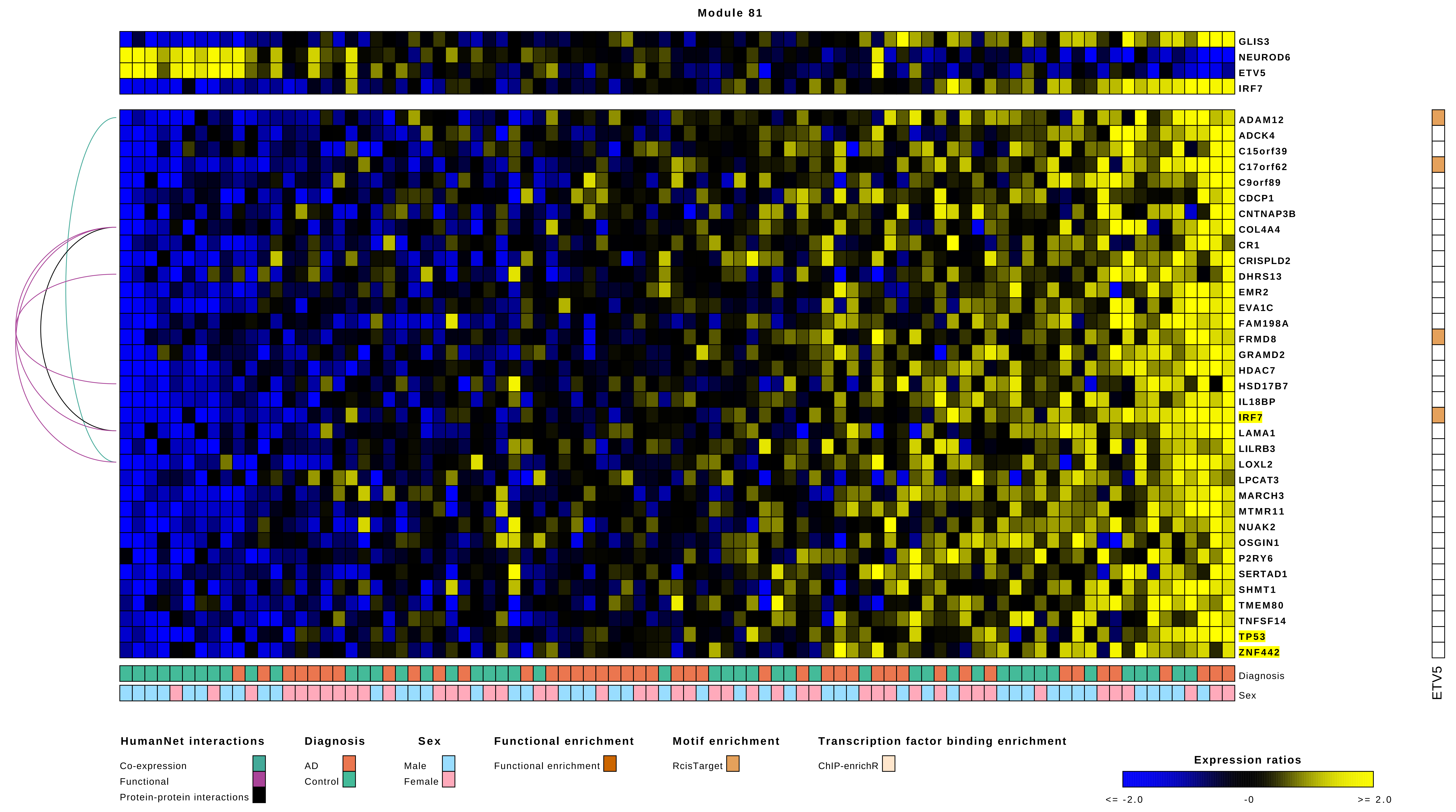

Module 81
GLIS3
NEUROD6
ETV5
IRF7
ADAM12
ADCK4
C15orf39
C17orf62
C9orf89
CDCP1
CNTNAP3B
COL4A4
CR1
CRISPLD2
DHRS13
EMR2
EVA1C
FAM198A
FRMD8
GRAMD2
HDAC7
HSD17B7
IL18BP
IRF7
LAMA1
LILRB3
LOXL2
LPCAT3
MARCH3
MTMR11
NUAK2
OSGIN1
P2RY6
SERTAD1
SHMT1
TMEM80
TNFSF14
TP53
ZNF442
Diagnosis
Sex
ETV5
HumanNet interactions
Co-expression
Functional
Protein-protein interactions
Diagnosis
AD
Control
Sex
Male
Female
Functional enrichment
Functional enrichment
Motif enrichment
RcisTarget
Transcription factor binding enrichment
ChIP-enrichR
Expression ratios
<= -2.0
-0
>= 2.0

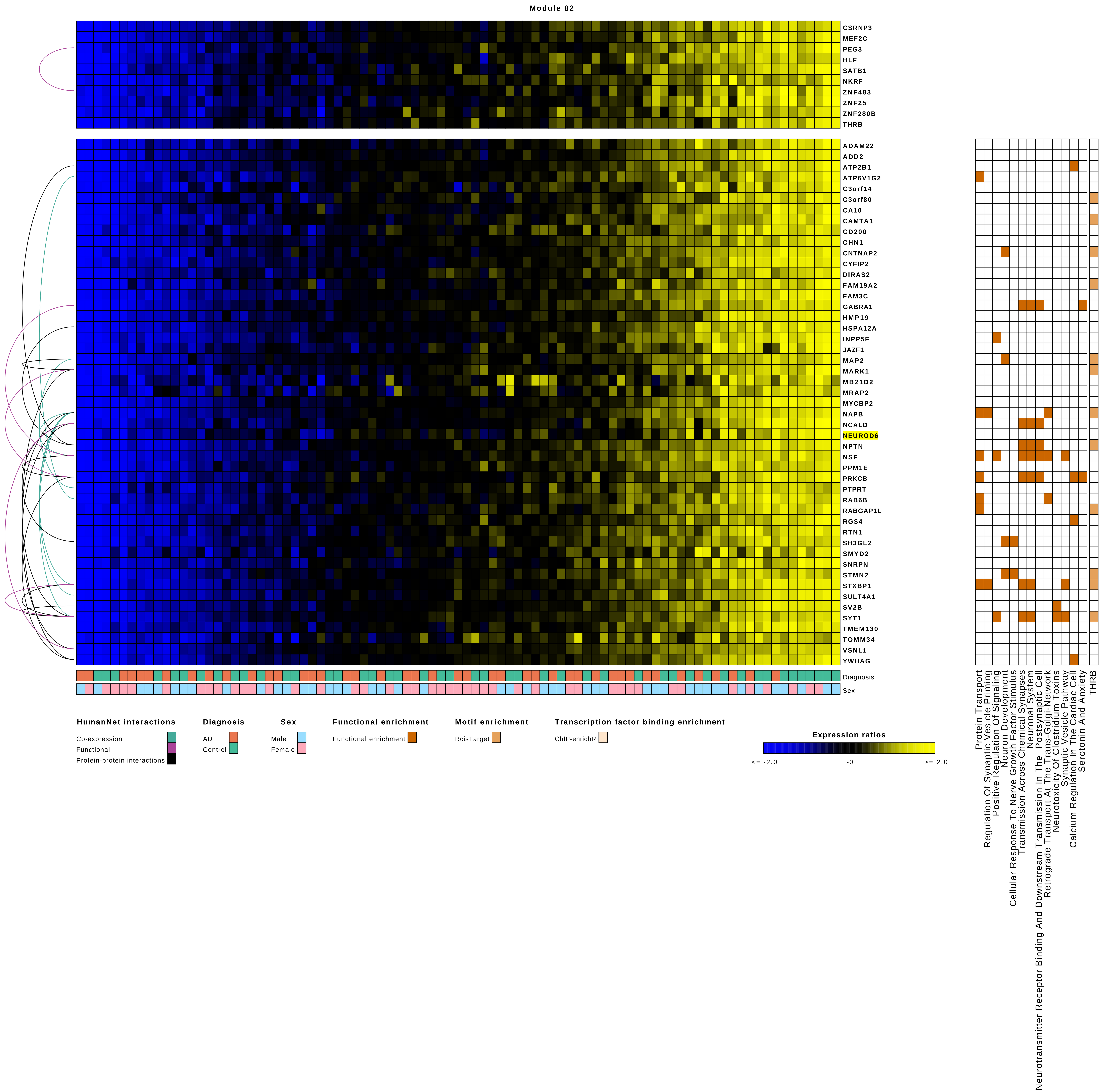

Module 82
CSRNP3
MEF2C
PEG3
HLF
SATB1
NKRF
ZNF483
ZNF25
ZNF280B
THRB
ADAM22
ADD2
ATP2B1
ATP6V1G2
C3orf14
C3orf80
CA10
CAMTA1
CD200
CHN1
CNTNAP2
CYFIP2
DIRAS2
FAM19A2
FAM3C
GABRA1
HMP19
HSPA12A
INPP5F
JAZF1
MAP2
MARK1
MB21D2
MRAP2
MYCBP2
NAPB
NCALD
NEUROD6
NPTN
NSF
PPM1E
PRKCB
PTPRT
RAB6B
RABGAP1L
RGS4
RTN1
SH3GL2
SMYD2
SNRPN
STMN2
STXBP1
SULT4A1
SV2B
SYT1
TMEM130
TOMM34
VSNL1
YWHAG
Diagnosis
Sex
Protein Transport
Regulation Of Synaptic Vesicle Priming
Positive Regulation Of Signaling
Neuron Development
Cellular Response To Nerve Growth Factor Stimulus
Transmission Across Chemical Synapses
Neuronal System
Neurotransmitter Receptor Binding And Downstream Transmission In The Postsynaptic Cell
Retrograde Transport At The Trans-Golgi-Network
Neurotoxicity Of Clostridium Toxins
Synaptic Vesicle Pathway
Calcium Regulation In The Cardiac Cell
Serotonin And Anxiety
THRB
HumanNet interactions
Co-expression
Functional
Protein-protein interactions
Diagnosis
AD
Control
Sex
Male
Female
Functional enrichment
Functional enrichment
Motif enrichment
RcisTarget
Transcription factor binding enrichment
ChIP-enrichR
Expression ratios
<= -2.0
-0
>= 2.0

Module 83

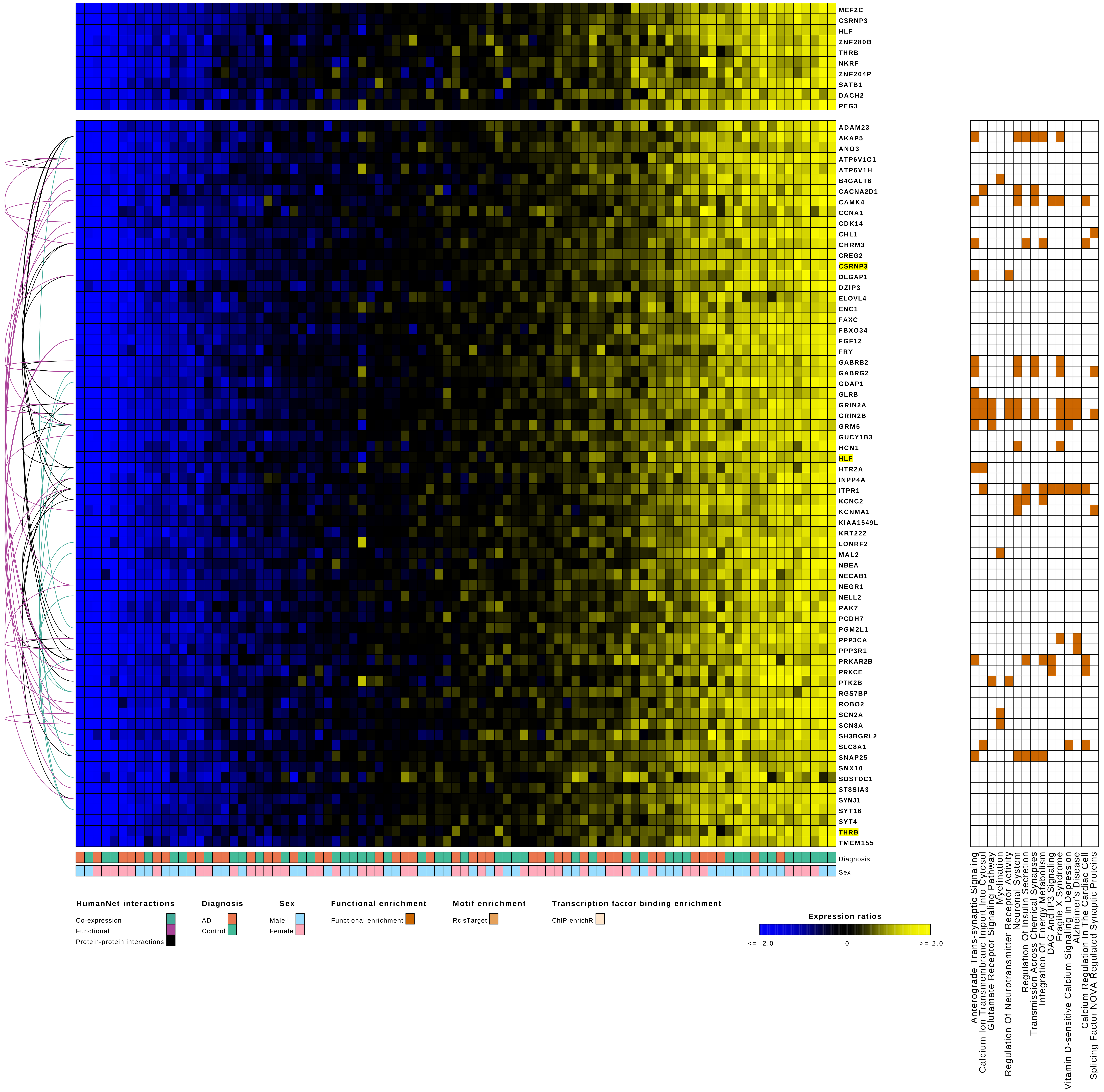
MEF2C
CSRNP3
HLF
ZNF280B
THRB
NKRF
ZNF204P
SATB1
DACH2
PEG3
ADAM23
AKAP5
ANO3
ATP6V1C1
ATP6V1H
B4GALT6
CACNA2D1
CAMK4
CCNA1
CDK14
CHL1
CHRM3
CREG2
CSRNP3
DLGAP1
DZIP3
ELOVL4
ENC1
FAXC
FBXO34
FGF12
FRY
GABRB2
GABRG2
GDAP1
GLRB
GRIN2A
GRIN2B
GRM5
GUCY1B3
HCN1
HLF
HTR2A
INPP4A
ITPR1
KCNC2
KCNMA1
KIAA1549L
KRT222
LONRF2
MAL2
NBEA
NECAB1
NEGR1
NELL2
PAK7
PCDH7
PGM2L1
PPP3CA
PPP3R1
PRKAR2B
PRKCE
PTK2B
RGS7BP
ROBO2
SCN2A
SCN8A
SH3BGRL2
SLC8A1
SNAP25
SNX10
SOSTDC1
ST8SIA3
SYNJ1
SYT16
SYT4
THRB
TMEM155
Diagnosis
Sex
Anterograde Trans-synaptic Signaling
Calcium Ion Transmembrane Import Into Cytosol
Glutamate Receptor Signaling Pathway
Myelination
Regulation Of Neurotransmitter Receptor Activity
Neuronal System
Regulation Of Insulin Secretion
Transmission Across Chemical Synapses
Integration Of Energy Metabolism
DAG And IP3 Signaling
Fragile X Syndrome
Vitamin D-sensitive Calcium Signaling In Depression
Alzheimer's Disease
Calcium Regulation In The Cardiac Cell
Splicing Factor NOVA Regulated Synaptic Proteins
HumanNet interactions
Co-expression
Functional
Protein-protein interactions
Diagnosis
AD
Control
Sex
Male
Female
Functional enrichment
Functional enrichment
Motif enrichment
RcisTarget
Transcription factor binding enrichment
ChIP-enrichR
Expression ratios
<= -2.0
-0
>= 2.0

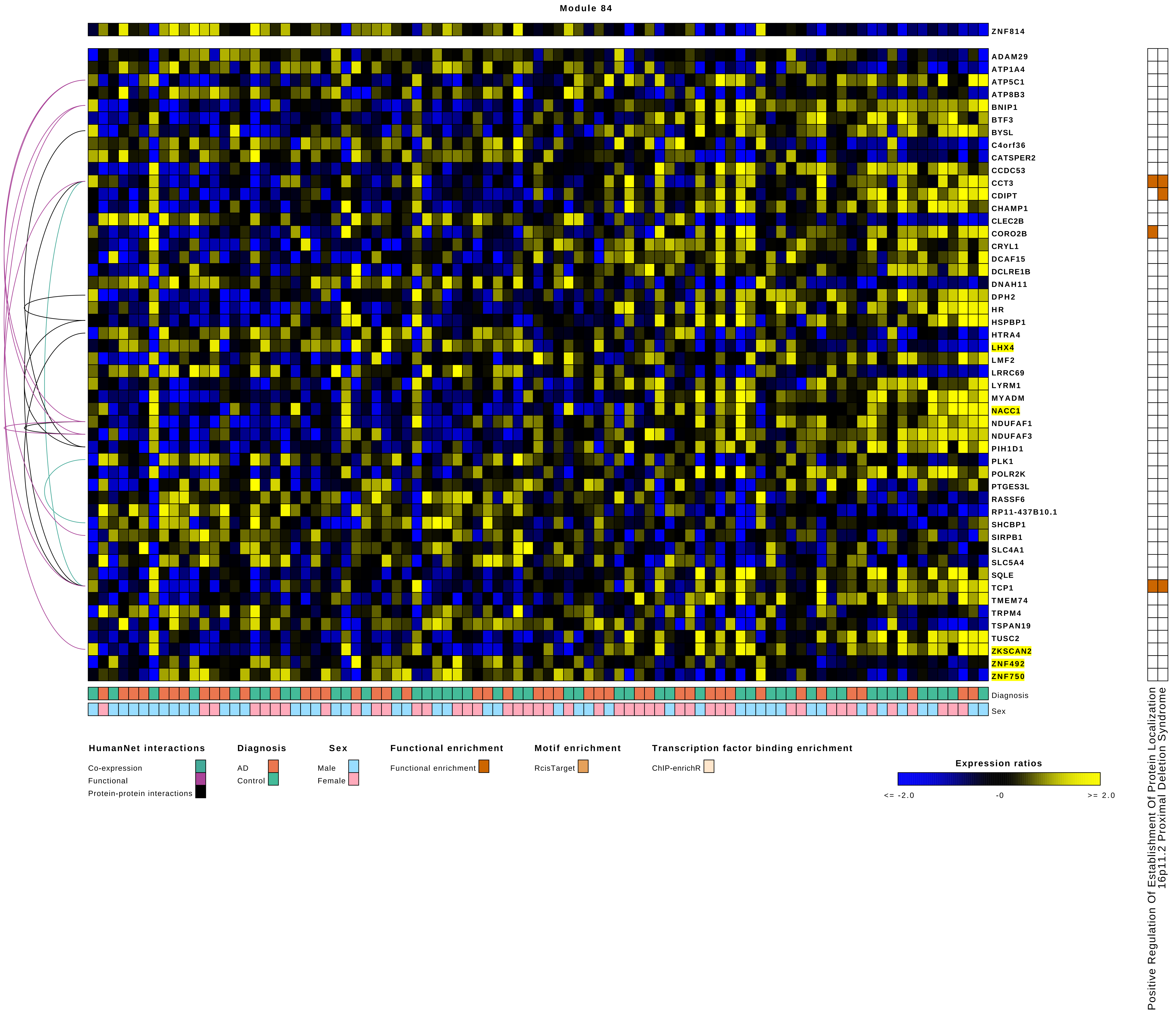

Module 84
ZNF814
ADAM29
ATP1A4
ATP5C1
ATP8B3
BNIP1
BTF3
BYSL
C4orf36
CATSPER2
CCDC53
CCT3
CDIPT
CHAMP1
CLEC2B
CORO2B
CRYL1
DCAF15
DCLRE1B
DNAH11
DPH2
HR
HSPBP1
HTRA4
LHX4
LMF2
LRRC69
LYRM1
MYADM
NACC1
NDUFAF1
NDUFAF3
PIH1D1
PLK1
POLR2K
PTGES3L
RASSF6
RP11-437B10.1
SHCBP1
SIRPB1
SLC4A1
SLC5A4
SQLE
TCP1
TMEM74
TRPM4
TSPAN19
TUSC2
ZKSCAN2
ZNF492
ZNF750
Diagnosis
Sex
Positive Regulation Of Establishment Of Protein Localization
16p11.2 Proximal Deletion Syndrome
HumanNet interactions
Co-expression
Functional
Protein-protein interactions
Diagnosis
AD
Control
Sex
Male
Female
Functional enrichment
Functional enrichment
Motif enrichment
RcisTarget
Transcription factor binding enrichment
ChIP-enrichR
Expression ratios
<= -2.0
-0
>= 2.0

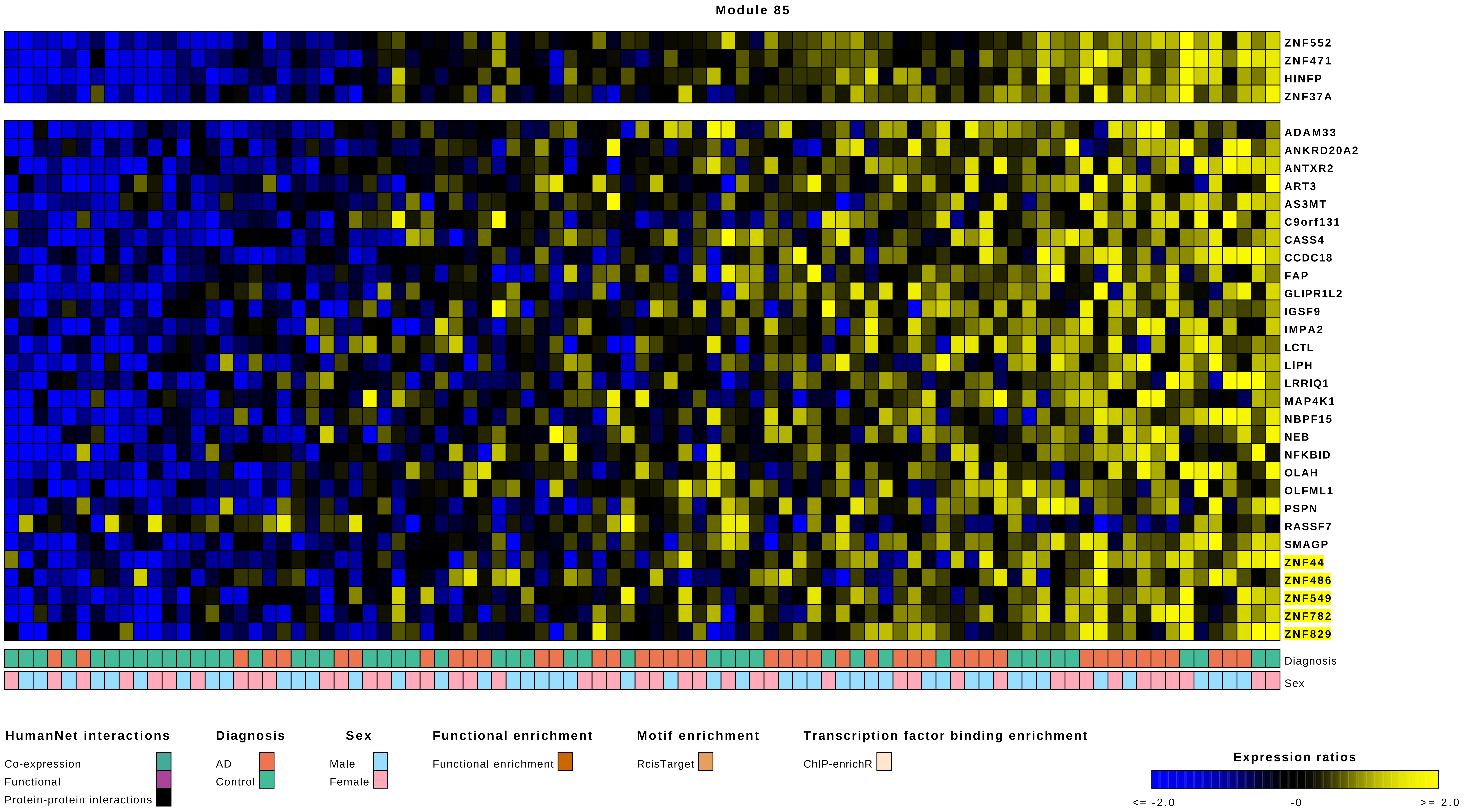

Module 85
ZNF552
ZNF471
HINFP
ZNF37A
ADAM33
ANKRD20A2
ANTXR2
ART3
AS3MT
C9orf131
CASS4
CCDC18
FAP
GLIPR1L2
IGSF9
IMPA2
LCTL
LIPH
LRRIQ1
MAP4K1
NBPF15
NEB
NFKBID
OLAH
OLFML1
PSPN
RASSF7
SMAGP
ZNF44
ZNF486
ZNF549
ZNF782
ZNF829
Diagnosis
Sex
HumanNet interactions
Co-expression
Functional
Protein-protein interactions
Diagnosis
AD
Control
Sex
Male
Female
Functional enrichment
Functional enrichment
Motif enrichment
RcisTarget
Transcription factor binding enrichment
ChIP-enrichR
Expression ratios
<= -2.0
-0
>= 2.0

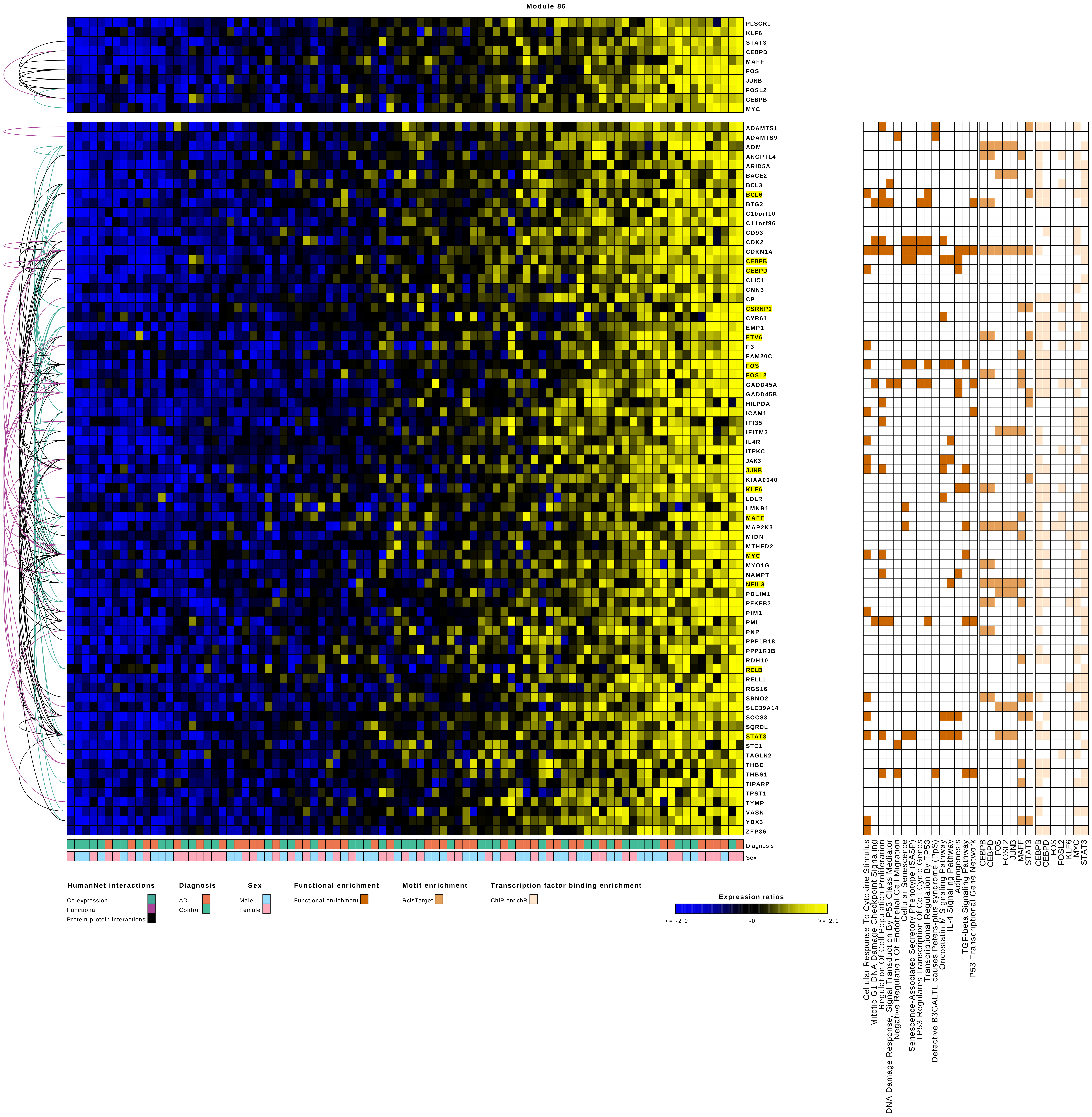

Module 86
PLSCR1
KLF6
STAT3
CEBPD
MAFF
FOS
JUNB
FOSL2
CEBPB
MYC
ADAMTS1
ADAMTS9
ADM
ANGPTL4
ARID5A
BACE2
BCL3
BCL6
BTG2
C10orf10
C11orf96
CD93
CDK2
CDKN1A
CEBPB
CEBPD
CLIC1
CNN3
CP
CSRNP1
CYR61
EMP1
ETV6
F3
FAM20C
FOS
FOSL2
GADD45A
GADD45B
HILPDA
ICAM1
IFI35
IFITM3
IL4R
ITPKC
JAK3
JUNB
KIAA0040
KLF6
LDLR
LMNB1
MAFF
MAP2K3
MIDN
MTHFD2
MYC
MYO1G
NAMPT
NFIL3
PDLIM1
PFKFB3
PIM1
PML
PNP
PPP1R18
PPP1R3B
RDH10
RELB
RELL1
RGS16
SBNO2
SLC39A14
SOCS3
SQRDL
STAT3
STC1
TAGLN2
THBD
THBS1
TIPARP
TPST1
TYMP
VASN
YBX3
ZFP36
Diagnosis
Sex
Cellular Response To Cytokine Stimulus
Mitotic G1 DNA Damage Checkpoint Signaling
Regulation Of Cell Population Proliferation
DNA Damage Response, Signal Transduction By P53 Class Mediator
Negative Regulation Of Endothelial Cell Migration
Cellular Senescence
Senescence-Associated Secretory Phenotype (SASP)
TP53 Regulates Transcription Of Cell Cycle Genes
Transcriptional Regulation By TP53
Defective B3GALTL causes Peters-plus syndrome (PpS)
Oncostatin M Signaling Pathway
IL-4 Signaling Pathway
Adipogenesis
TGF-beta Signaling Pathway
P53 Transcriptional Gene Network
CEBPB
CEBPD
FOS
FOSL2
JUNB
MAFF
STAT3
CEBPB
CEBPD
FOS
FOSL2
KLF6
MYC
STAT3
HumanNet interactions
Co-expression
Functional
Protein-protein interactions
Diagnosis
AD
Control
Sex
Male
Female
Functional enrichment
Functional enrichment
Motif enrichment
RcisTarget
Transcription factor binding enrichment
ChIP-enrichR
Expression ratios
<= -2.0
-0
>= 2.0

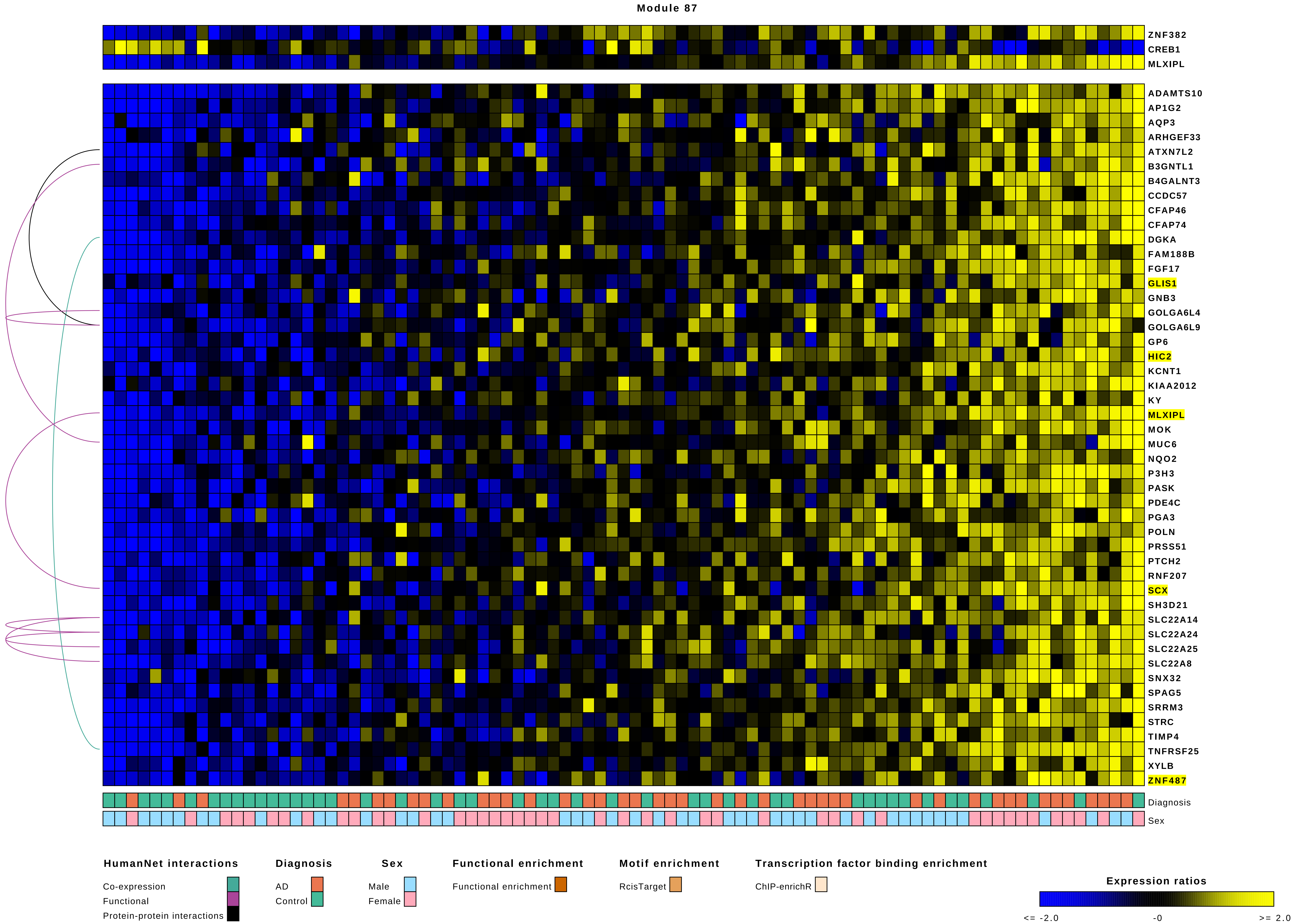
Module 87
ZNF382
CREB1
MLXIPL
ADAMTS10
AP1G2
AQP3
ARHGEF33
ATXN7L2
B3GNTL1
B4GALNT3
CCDC57
CFAP46
CFAP74
DGKA
FAM188B
FGF17
GLIS1
GNB3
GOLGA6L4
GOLGA6L9
GP6
HIC2
KCNT1
KIAA2012
KY
MLXIPL
MOK
MUC6
NQO2
P3H3
PASK
PDE4C
PGA3
POLN
PRSS51
PTCH2
RNF207
SCX
SH3D21
SLC22A14
SLC22A24
SLC22A25
SLC22A8
SNX32
SPAG5
SRRM3
STRC
TIMP4
TNFRSF25
XYLB
ZNF487
Diagnosis
Sex
HumanNet interactions
Co-expression
Functional
Protein-protein interactions
Diagnosis
AD
Control
Sex
Male
Female
Functional enrichment
Functional enrichment
Motif enrichment
RcisTarget
Transcription factor binding enrichment
ChIP-enrichR
Expression ratios
<= -2.0
-0
>= 2.0

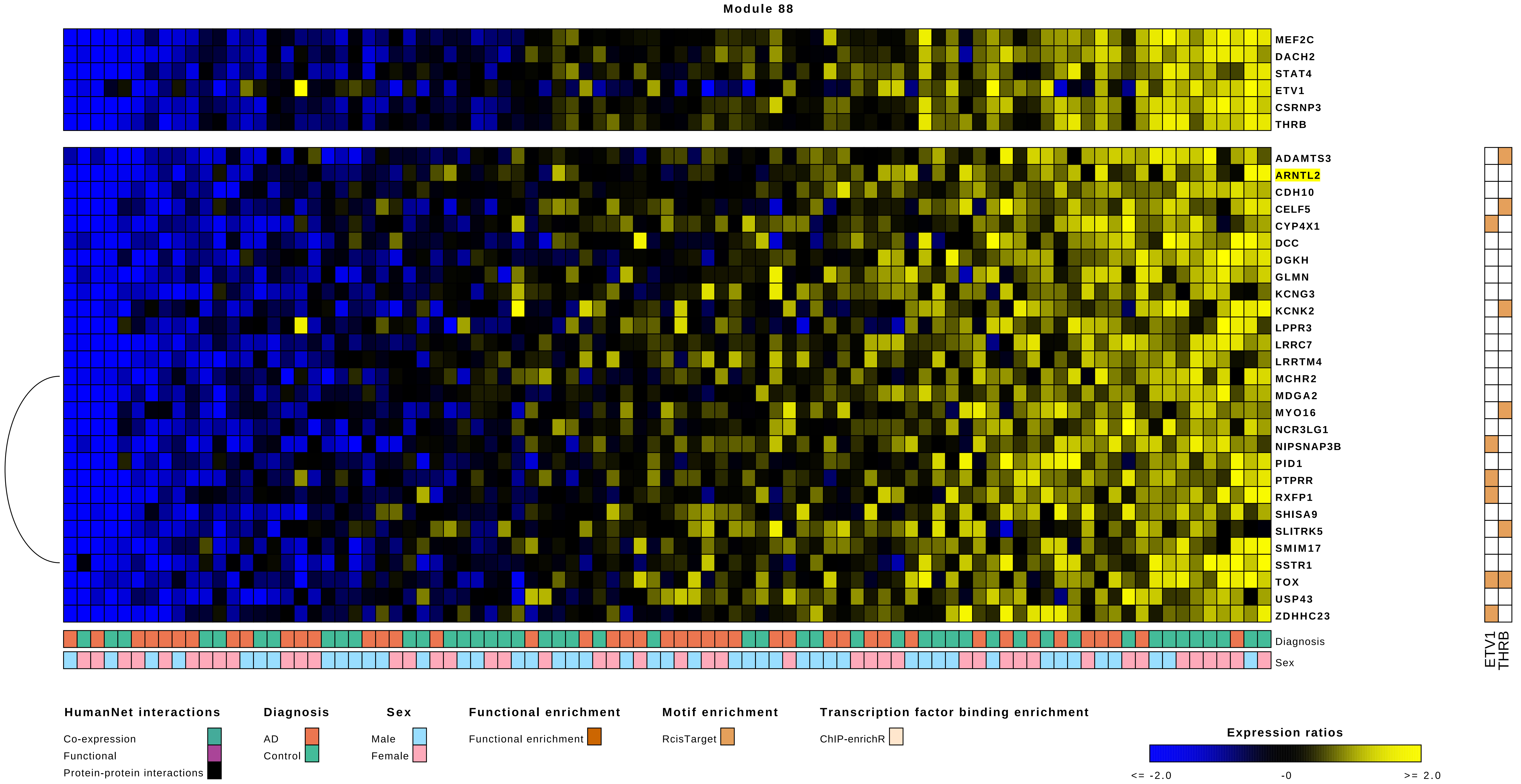

Module 88
MEF2C
DACH2
STAT4
ETV1
CSRNP3
THRB
ADAMTS3
ARNTL2
CDH10
CELF5
CYP4X1
DCC
DGKH
GLMN
KCNG3
KCNK2
LPPR3
LRRC7
LRRTM4
MCHR2
MDGA2
MYO16
NCR3LG1
NIPSNAP3B
PID1
PTPRR
RXFP1
SHISA9
SLITRK5
SMIM17
SSTR1
TOX
USP43
ZDHHC23
Diagnosis
Sex
ETV1
THRB
HumanNet interactions
Co-expression
Functional
Protein-protein interactions
Diagnosis
AD
Control
Sex
Male
Female
Functional enrichment
Functional enrichment
Motif enrichment
RcisTarget
Transcription factor binding enrichment
ChIP-enrichR
Expression ratios
<= -2.0
-0
>= 2.0

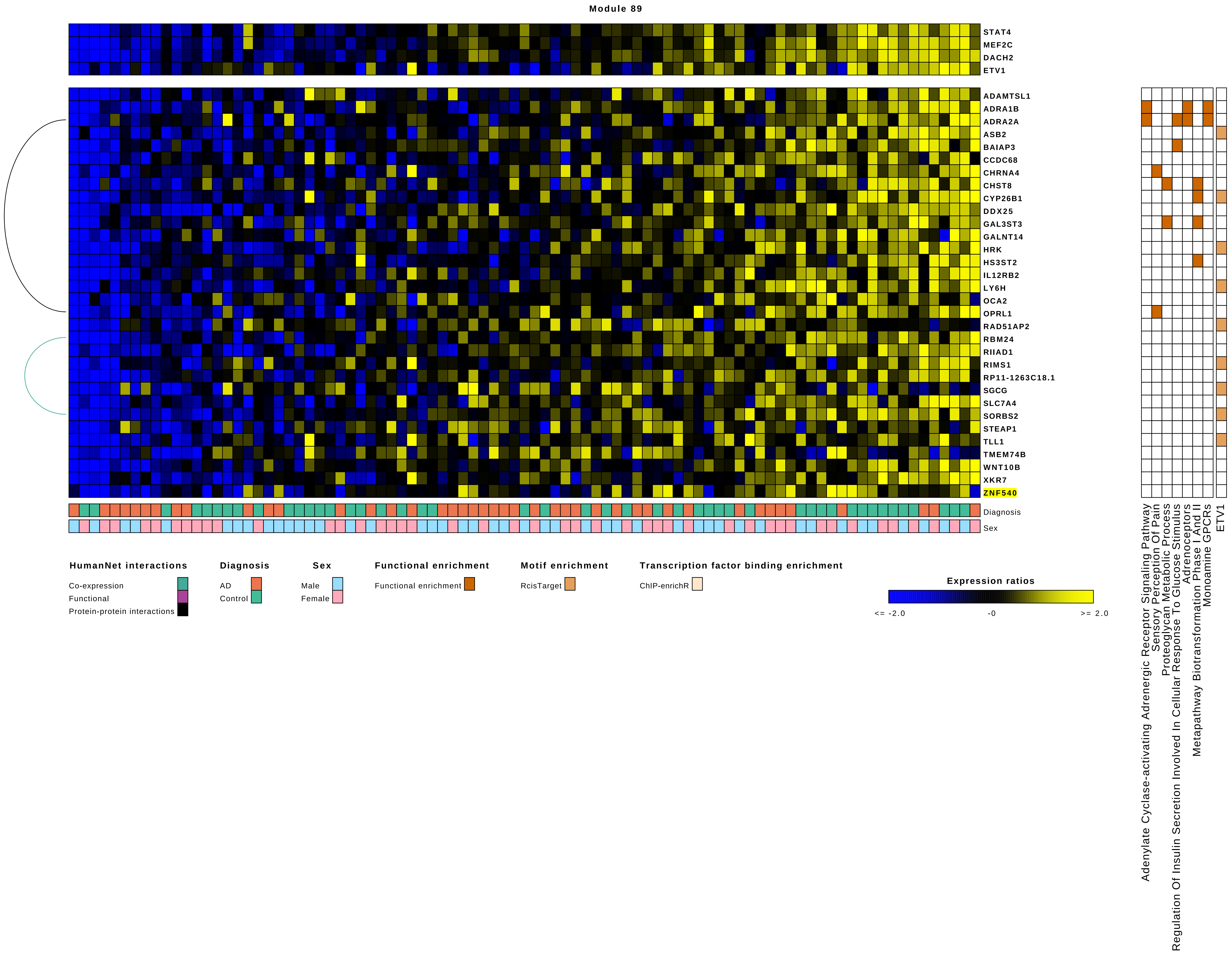

Module 89
STAT4
MEF2C
DACH2
ETV1
ADAMTSL1
ADRA1B
ADRA2A
ASB2
BAIAP3
CCDC68
CHRNA4
CHST8
CYP26B1
DDX25
GAL3ST3
GALNT14
HRK
HS3ST2
IL12RB2
LY6H
OCA2
OPRL1
RAD51AP2
RBM24
RIIAD1
RIMS1
RP11-1263C18.1
SGCG
SLC7A4
SORBS2
STEAP1
TLL1
TMEM74B
WNT10B
XKR7
ZNF540
Diagnosis
Sex
Adenylate Cyclase-activating Adrenergic Receptor Signaling Pathway
Sensory Perception Of Pain
Proteoglycan Metabolic Process
Regulation Of Insulin Secretion Involved In Cellular Response To Glucose Stimulus
Adrenoceptors
Metapathway Biotransformation Phase I And II
Monoamine GPCRs
ETV1
HumanNet interactions
Co-expression
Functional
Protein-protein interactions
Diagnosis
AD
Control
Sex
Male
Female
Functional enrichment
Functional enrichment
Motif enrichment
RcisTarget
Transcription factor binding enrichment
ChIP-enrichR
Expression ratios
<= -2.0
-0
>= 2.0

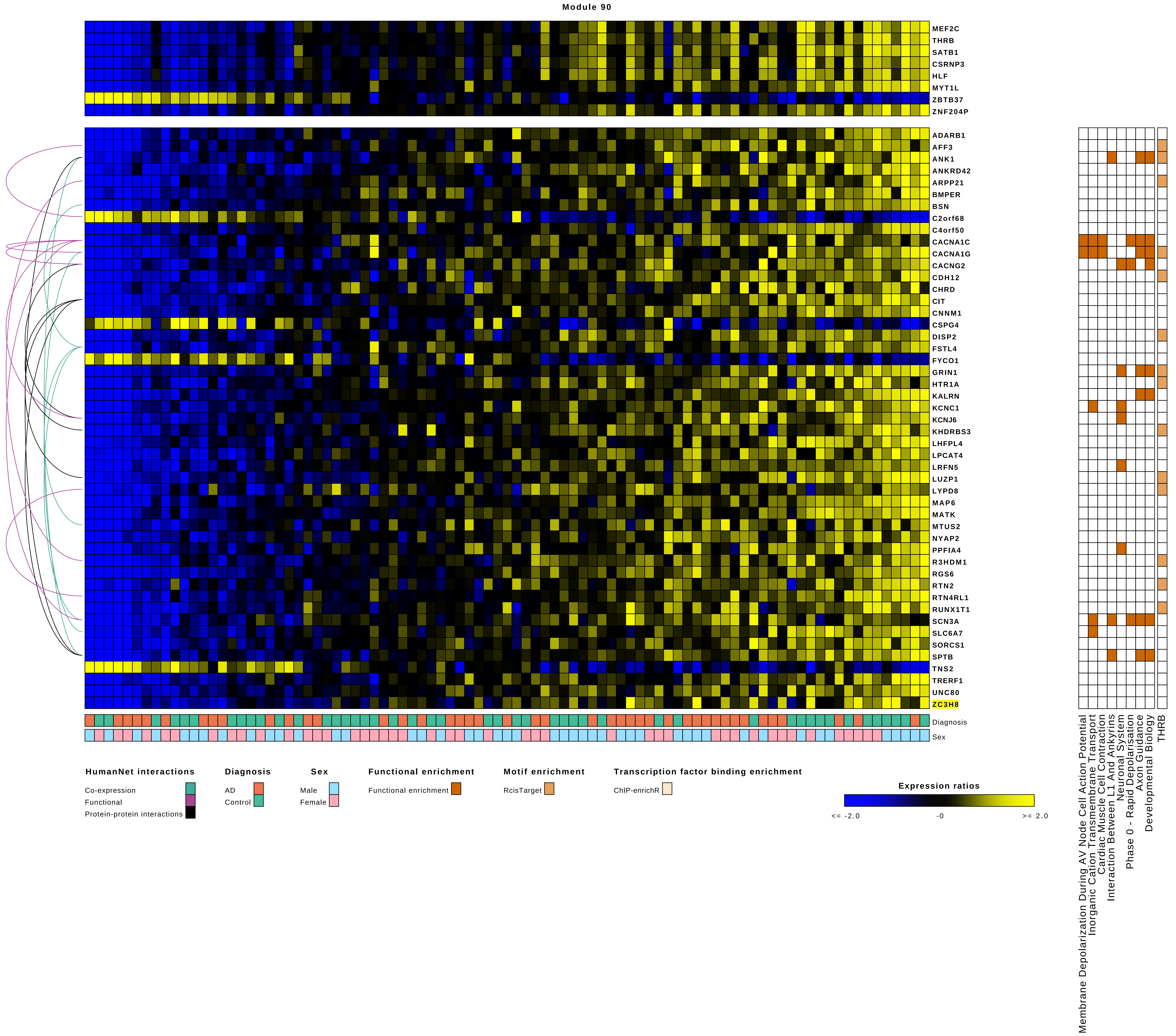

Module 90
MEF2C
THRB
SATB1
CSRNP3
HLF
MYT1L
ZBTB37
ZNF204P
ADARB1
AFF3
ANK1
ANKRD42
ARPP21
BMPER
BSN
C2orf68
C4orf50
CACNA1C
CACNA1G
CACNG2
CDH12
CHRD
CIT
CNNM1
CSPG4
DISP2
FSTL4
FYCO1
GRIN1
HTR1A
KALRN
KCNC1
KCNJ6
KHDRBS3
LHFPL4
LPCAT4
LRFN5
LUZP1
LYPD8
MAP6
MATK
MTUS2
NYAP2
PPFIA4
R3HDM1
RGS6
RTN2
RTN4RL1
RUNX1T1
SCN3A
SLC6A7
SORCS1
SPTB
TNS2
TRERF1
UNC80
ZC3H8
Diagnosis
Sex
Membrane Depolarization During AV Node Cell Action Potential
Inorganic Cation Transmembrane Transport
Cardiac Muscle Cell Contraction
Interaction Between L1 And Ankyrins
Neuronal System
Phase 0 - Rapid Depolarisation
Axon Guidance
Developmental Biology
THRB
HumanNet interactions
Co-expression
Functional
Protein-protein interactions
Diagnosis
AD
Control
Sex
Male
Female
Functional enrichment
Functional enrichment
Motif enrichment
RcisTarget
Transcription factor binding enrichment
ChIP-enrichR
Expression ratios
<= -2.0
-0
>= 2.0

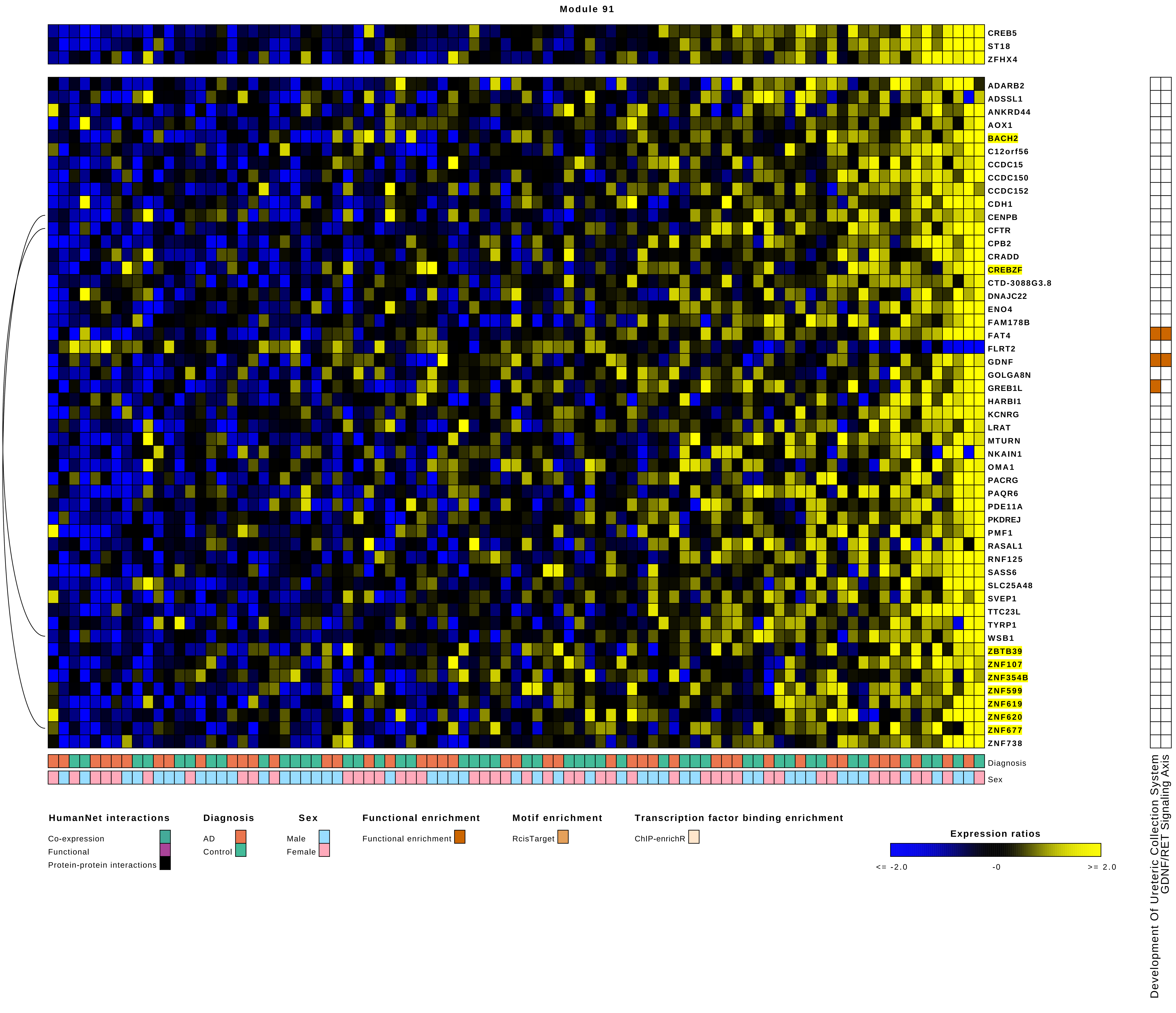
Module 91
CREB5
ST18
ZFHX4
ADARB2
ADSSL1
ANKRD44
AOX1
BACH2
C12orf56
CCDC15
CCDC150
CCDC152
CDH1
CENPB
CFTR
CPB2
CRADD
CREBZF
CTD-3088G3.8
DNAJC22
ENO4
FAM178B
FAT4
FLRT2
GDNF
GOLGA8N
GREB1L
HARBI1
KCNRG
LRAT
MTURN
NKAIN1
OMA1
PACRG
PAQR6
PDE11A
PKDREJ
PMF1
RASAL1
RNF125
SASS6
SLC25A48
SVEP1
TTC23L
TYRP1
WSB1
ZBTB39
ZNF107
ZNF354B
ZNF599
ZNF619
ZNF620
ZNF677
ZNF738
Diagnosis
Sex
Development Of Ureteric Collection System
GDNF/RET Signaling Axis
HumanNet interactions
Co-expression
Functional
Protein-protein interactions
Diagnosis
AD
Control
Sex
Male
Female
Functional enrichment
Functional enrichment
Motif enrichment
RcisTarget
Transcription factor binding enrichment
ChIP-enrichR
Expression ratios
<= -2.0
-0
>= 2.0

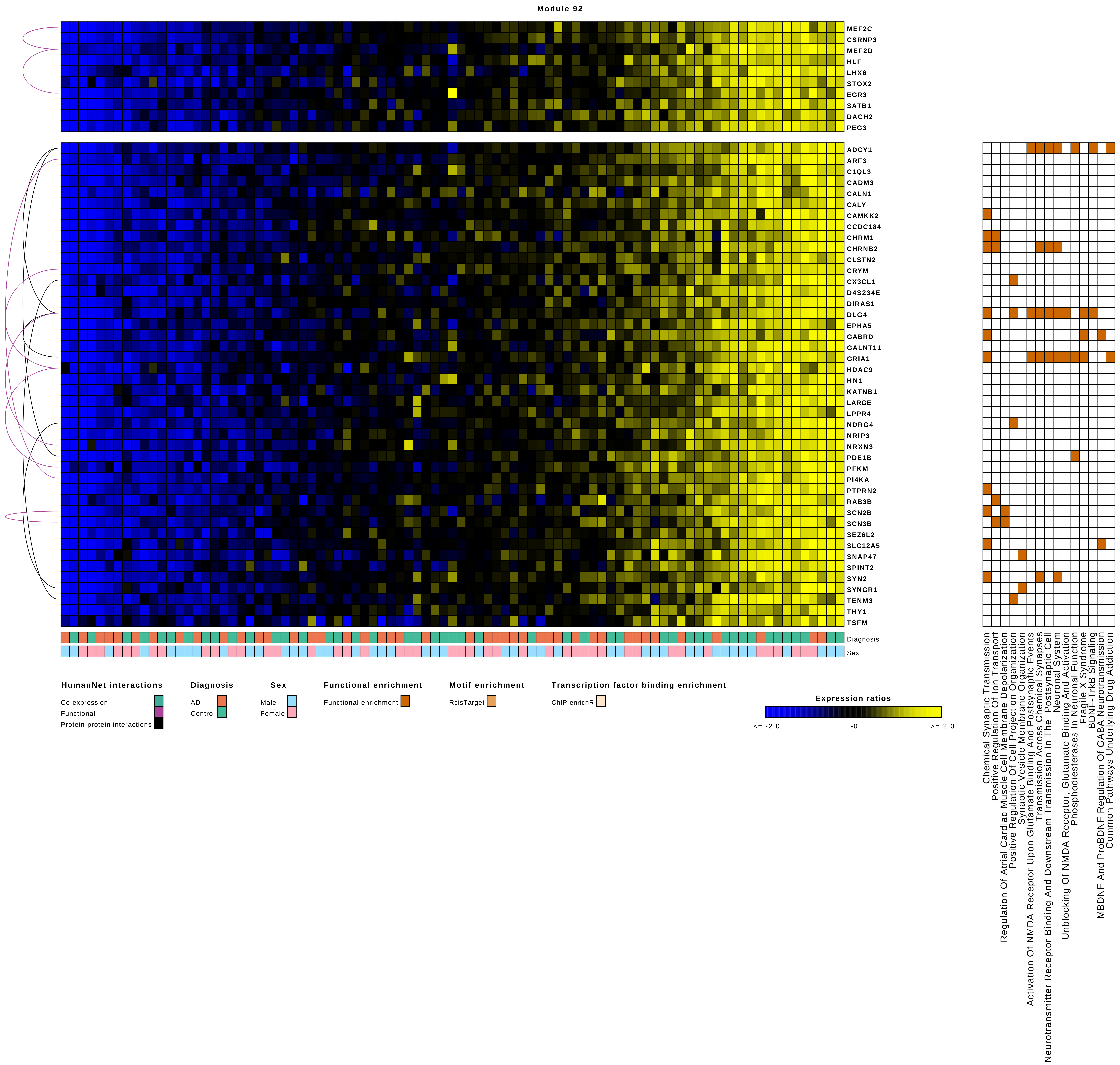
Module 92
MEF2C
CSRNP3
MEF2D
HLF
LHX6
STOX2
EGR3
SATB1
DACH2
PEG3
ADCY1
ARF3
C1QL3
CADM3
CALN1
CALY
CAMKK2
CCDC184
CHRM1
CHRNB2
CLSTN2
CRYM
CX3CL1
D4S234E
DIRAS1
DLG4
EPHA5
GABRD
GALNT11
GRIA1
HDAC9
HN1
KATNB1
LARGE
LPPR4
NDRG4
NRIP3
NRXN3
PDE1B
PFKM
PI4KA
PTPRN2
RAB3B
SCN2B
SCN3B
SEZ6L2
SLC12A5
SNAP47
SPINT2
SYN2
SYNGR1
TENM3
THY1
TSFM
Diagnosis
Sex
HumanNet interactions
Co-expression
Functional
Protein-protein interactions
Diagnosis
AD
Control
Sex
Male
Female
Functional enrichment
Functional enrichment
Motif enrichment
RcisTarget
Transcription factor binding enrichment
ChIP-enrichR
Expression ratios
<= -2.0
-0
>= 2.0
Chemical Synaptic Transmission
Positive Regulation Of Ion Transport
Regulation Of Atrial Cardiac Muscle Cell Membrane Depolarization
Positive Regulation Of Cell Projection Organization
Synaptic Vesicle Membrane Organization
Activation Of NMDA Receptor Upon Glutamate Binding And Postsynaptic Events
Transmission Across Chemical Synapses
Neurotransmitter Receptor Binding And Downstream Transmission In The Postsynaptic Cell
Neuronal System
Unblocking Of NMDA Receptor, Glutamate Binding And Activation
Phosphodiesterases In Neuronal Function
Fragile X Syndrome
BDNF-TrkB Signaling
MBDNF And ProBDNF Regulation Of GABA Neurotransmission
Common Pathways Underlying Drug Addiction

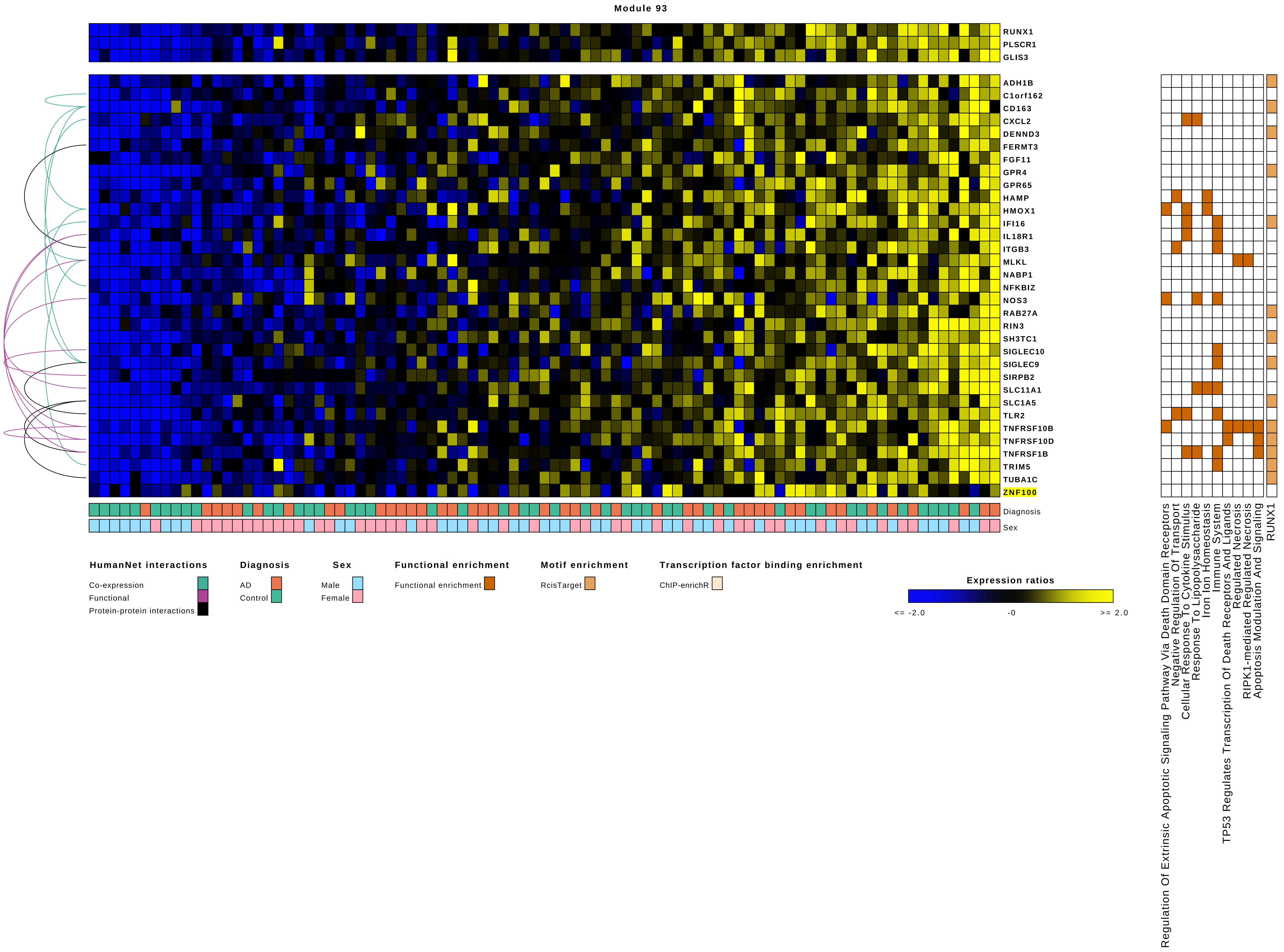

Module 93
RUNX1
PLSCR1
GLIS3
ADH1B
C1orf162
CD163
CXCL2
DENND3
FERMT3
FGF11
GPR4
GPR65
HAMP
HMOX1
IFI16
IL18R1
ITGB3
MLKL
NABP1
NFKBIZ
NOS3
RAB27A
RIN3
SH3TC1
SIGLEC10
SIGLEC9
SIRPB2
SLC11A1
SLC1A5
TLR2
TNFRSF10B
TNFRSF10D
TNFRSF1B
TRIM5
TUBA1C
ZNF100
Diagnosis
Sex
HumanNet interactions
Co-expression
Functional
Protein-protein interactions
Diagnosis
AD
Control
Sex
Male
Female
Functional enrichment
Functional enrichment
Motif enrichment
RcisTarget
Transcription factor binding enrichment
ChIP-enrichR
Expression ratios
<= -2.0
-0
>= 2.0
Regulation Of Extrinsic Apoptotic Signaling Pathway Via Death Domain Receptors
Negative Regulation Of Transport
Cellular Response To Cytokine Stimulus
Response To Lipopolysaccharide
Iron Ion Homeostasis
Immune System
TP53 Regulates Transcription Of Death Receptors And Ligands
Regulated Necrosis
RIPK1-mediated Regulated Necrosis
Apoptosis Modulation And Signaling
RUNX1

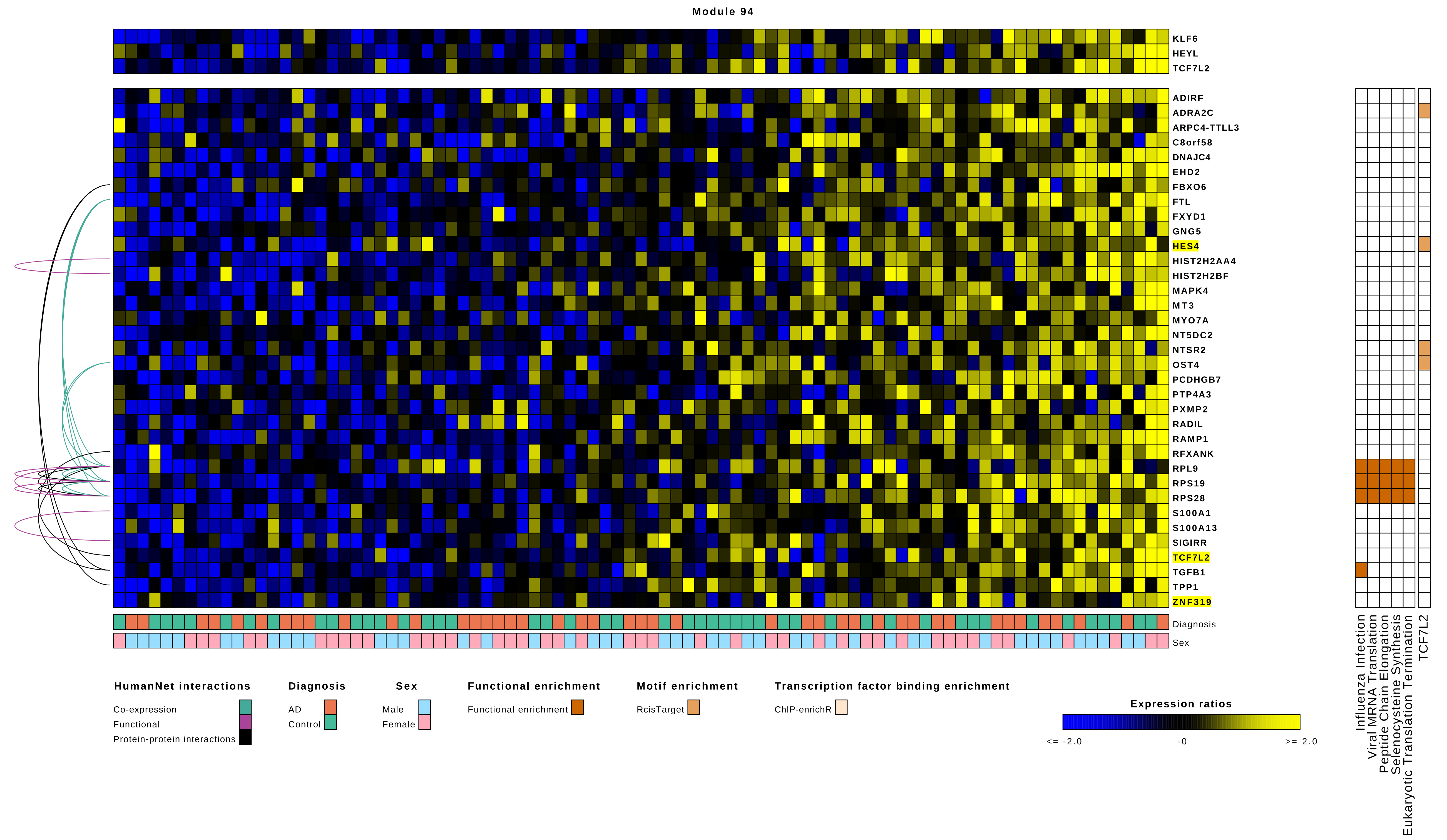

Module 94
KLF6
HEYL
TCF7L2
ADIRF
ADRA2C
ARPC4-TTLL3
C8orf58
DNAJC4
EHD2
FBXO6
FTL
FXYD1
GNG5
HES4
HIST2H2AA4
HIST2H2BF
MAPK4
MT3
MYO7A
NT5DC2
NTSR2
OST4
PCDHGB7
PTP4A3
PXMP2
RADIL
RAMP1
RFXANK
RPL9
RPS19
RPS28
S100A1
S100A13
SIGIRR
TCF7L2
TGFB1
TPP1
ZNF319
Diagnosis
Sex
Influenza Infection
Viral MRNA Translation
Peptide Chain Elongation
Selenocysteine Synthesis
Eukaryotic Translation Termination
TCF7L2
HumanNet interactions
Co-expression
Functional
Protein-protein interactions
Diagnosis
AD
Control
Sex
Male
Female
Functional enrichment
Functional enrichment
Motif enrichment
RcisTarget
Transcription factor binding enrichment
ChIP-enrichR
Expression ratios
<= -2.0
-0
>= 2.0

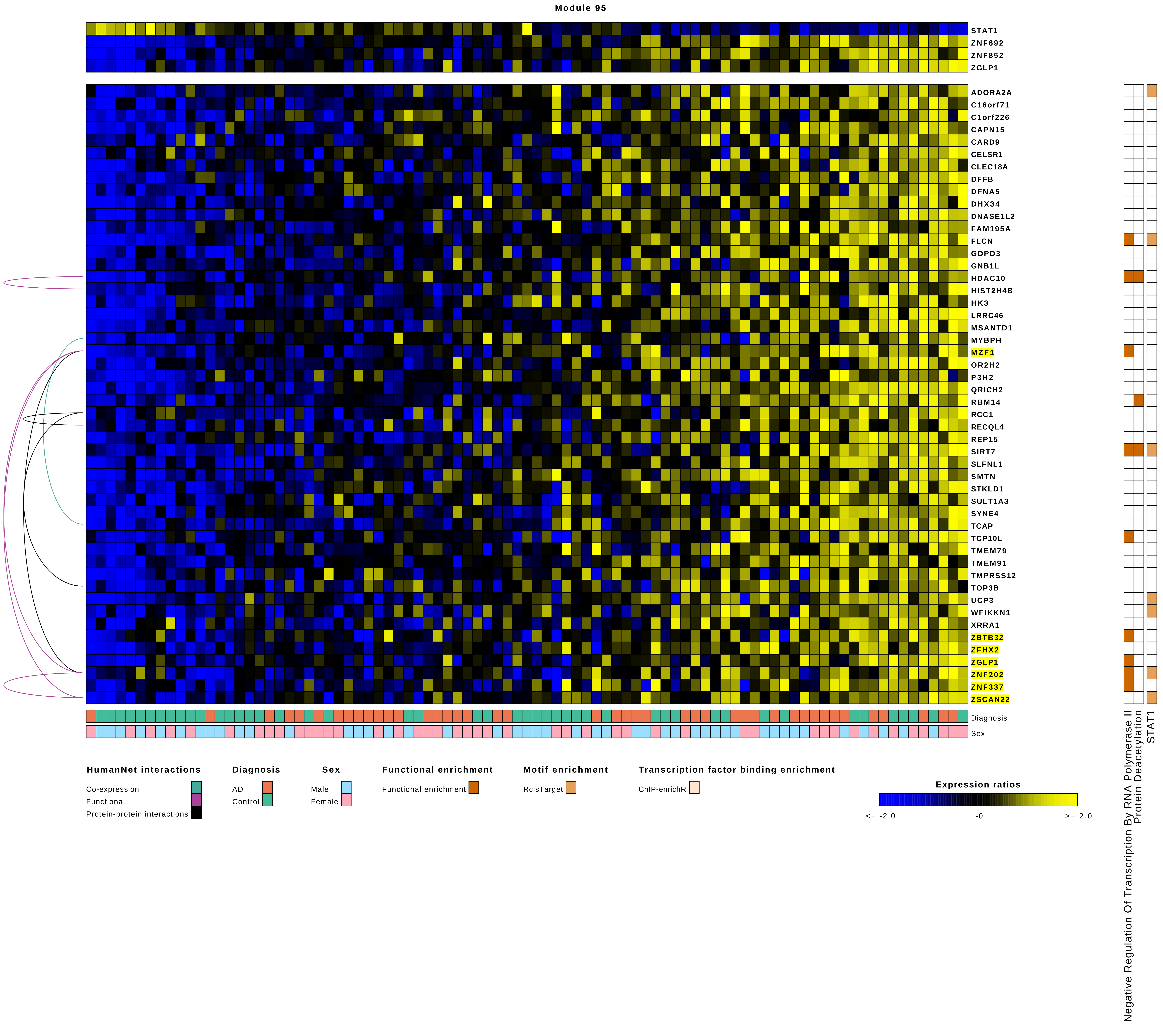

Module 95
STAT1
ZNF692
ZNF852
ZGLP1
ADORA2A
C16orf71
C1orf226
CAPN15
CARD9
CELSR1
CLEC18A
DFFB
DFNA5
DHX34
DNASE1L2
FAM195A
FLCN
GDPD3
GNB1L
HDAC10
HIST2H4B
HK3
LRRC46
MSANTD1
MYBPH
MZF1
OR2H2
P3H2
QRICH2
RBM14
RCC1
RECQL4
REP15
SIRT7
SLFNL1
SMTN
STKLD1
SULT1A3
SYNE4
TCAP
TCP10L
TMEM79
TMEM91
TMPRSS12
TOP3B
UCP3
WFIKKN1
XRRA1
ZBTB32
ZFHX2
ZGLP1
ZNF202
ZNF337
ZSCAN22
Diagnosis
Sex
Negative Regulation Of Transcription By RNA Polymerase II
Protein Deacetylation
STAT1
HumanNet interactions
Co-expression
Functional
Protein-protein interactions
Diagnosis
AD
Control
Sex
Male
Female
Functional enrichment
Functional enrichment
Motif enrichment
RcisTarget
Transcription factor binding enrichment
ChIP-enrichR
Expression ratios
<= -2.0
-0
>= 2.0

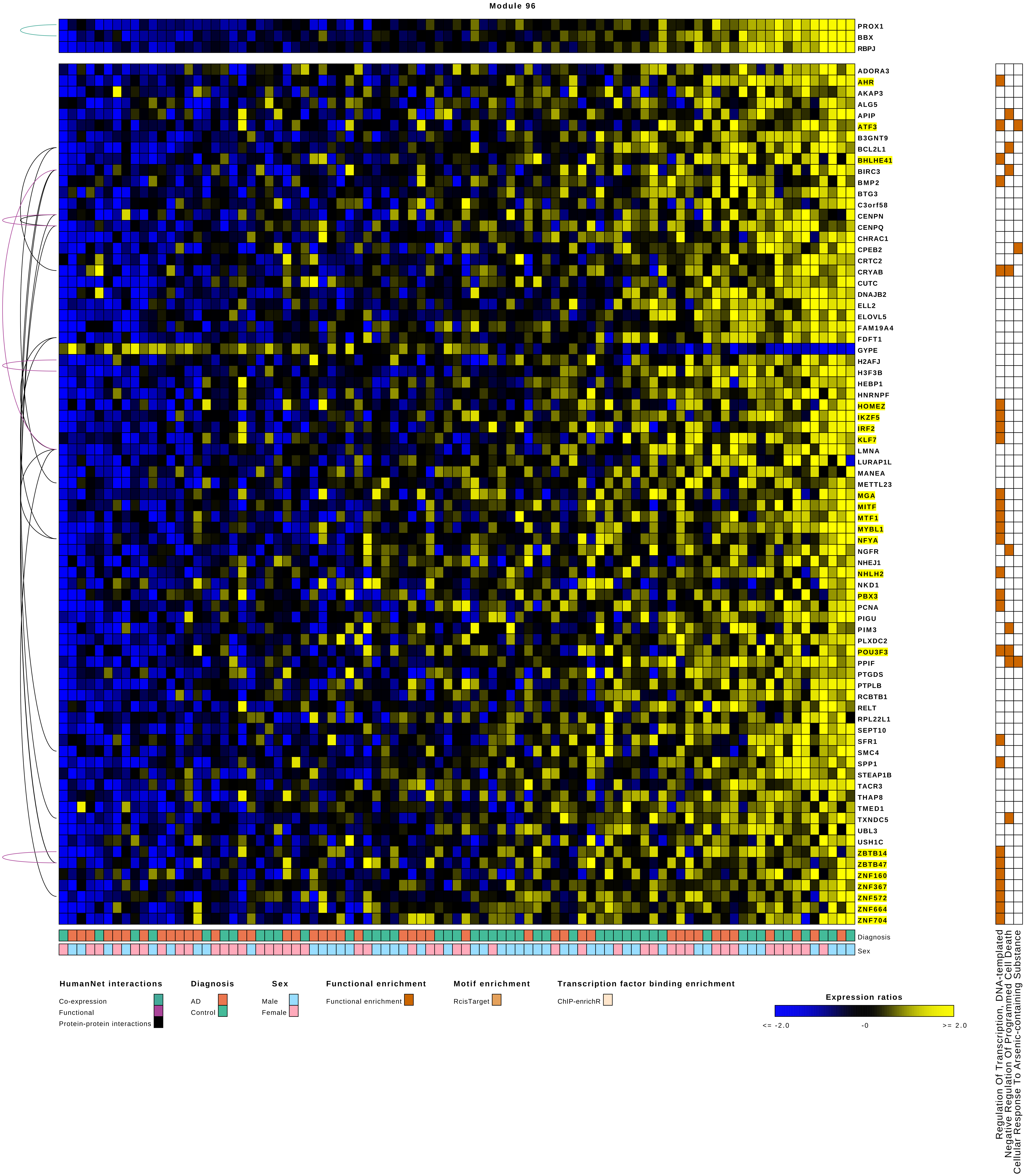
Module 96
PROX1
BBX
RBPJ
ADORA3
AHR
AKAP3
ALG5
APIP
ATF3
B3GNT9
BCL2L1
BHLHE41
BIRC3
BMP2
BTG3
C3orf58
CENPN
CENPQ
CHRAC1
CPEB2
CRTC2
CRYAB
CUTC
DNAJB2
ELL2
ELOVL5
FAM19A4
FDFT1
GYPE
H2AFJ
H3F3B
HEBP1
HNRNPF
HOMEZ
IKZF5
IRF2
KLF7
LMNA
LURAP1L
MANEA
METTL23
MGA
MITF
MTF1
MYBL1
NFYA
NGFR
NHEJ1
NHLH2
NKD1
PBX3
PCNA
PIGU
PIM3
PLXDC2
POU3F3
PPIF
PTGDS
PTPLB
RCBTB1
RELT
RPL22L1
SEPT10
SFR1
SMC4
SPP1
STEAP1B
TACR3
THAP8
TMED1
TXNDC5
UBL3
USH1C
ZBTB14
ZBTB47
ZNF160
ZNF367
ZNF572
ZNF664
ZNF704
Diagnosis
Sex
Regulation Of Transcription, DNA-templated
Negative Regulation Of Programmed Cell Death
Cellular Response To Arsenic-containing Substance
HumanNet interactions
Co-expression
Functional
Protein-protein interactions
Diagnosis
AD
Control
Sex
Male
Female
Functional enrichment
Functional enrichment
Motif enrichment
RcisTarget
Transcription factor binding enrichment
ChIP-enrichR
Expression ratios
<= -2.0
-0
>= 2.0

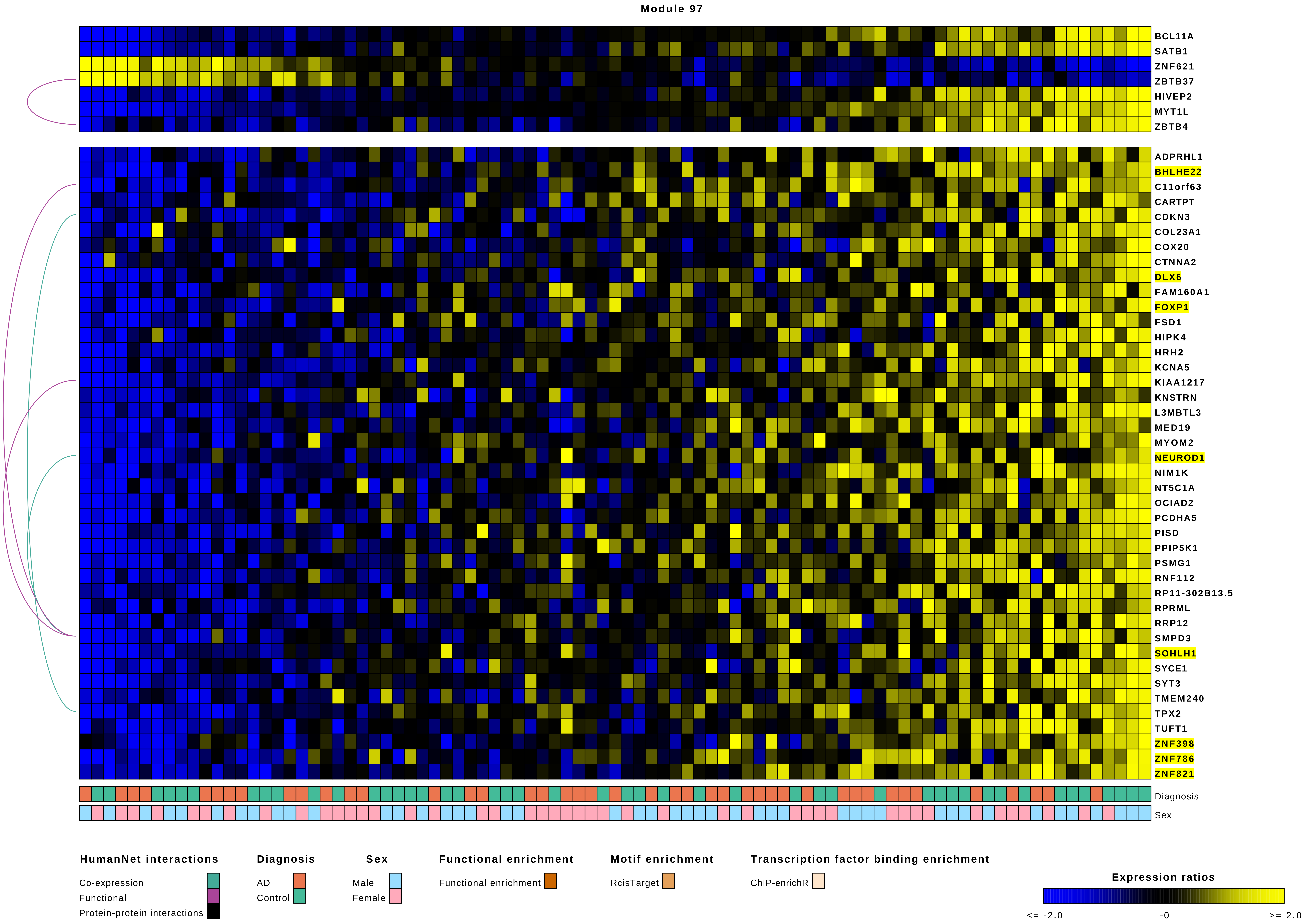
Module 97
BCL11A
SATB1
ZNF621
ZBTB37
HIVEP2
MYT1L
ZBTB4
ADPRHL1
BHLHE22
C11orf63
CARTPT
CDKN3
COL23A1
COX20
CTNNA2
DLX6
FAM160A1
FOXP1
FSD1
HIPK4
HRH2
KCNA5
KIAA1217
KNSTRN
L3MBTL3
MED19
MYOM2
NEUROD1
NIM1K
NT5C1A
OCIAD2
PCDHA5
PISD
PPIP5K1
PSMG1
RNF112
RP11-302B13.5
RPRML
RRP12
SMPD3
SOHLH1
SYCE1
SYT3
TMEM240
TPX2
TUFT1
ZNF398
ZNF786
ZNF821
Diagnosis
Sex
HumanNet interactions
Co-expression
Functional
Protein-protein interactions
Diagnosis
AD
Control
Sex
Male
Female
Functional enrichment
Functional enrichment
Motif enrichment
RcisTarget
Transcription factor binding enrichment
ChIP-enrichR
Expression ratios
<= -2.0
-0
>= 2.0

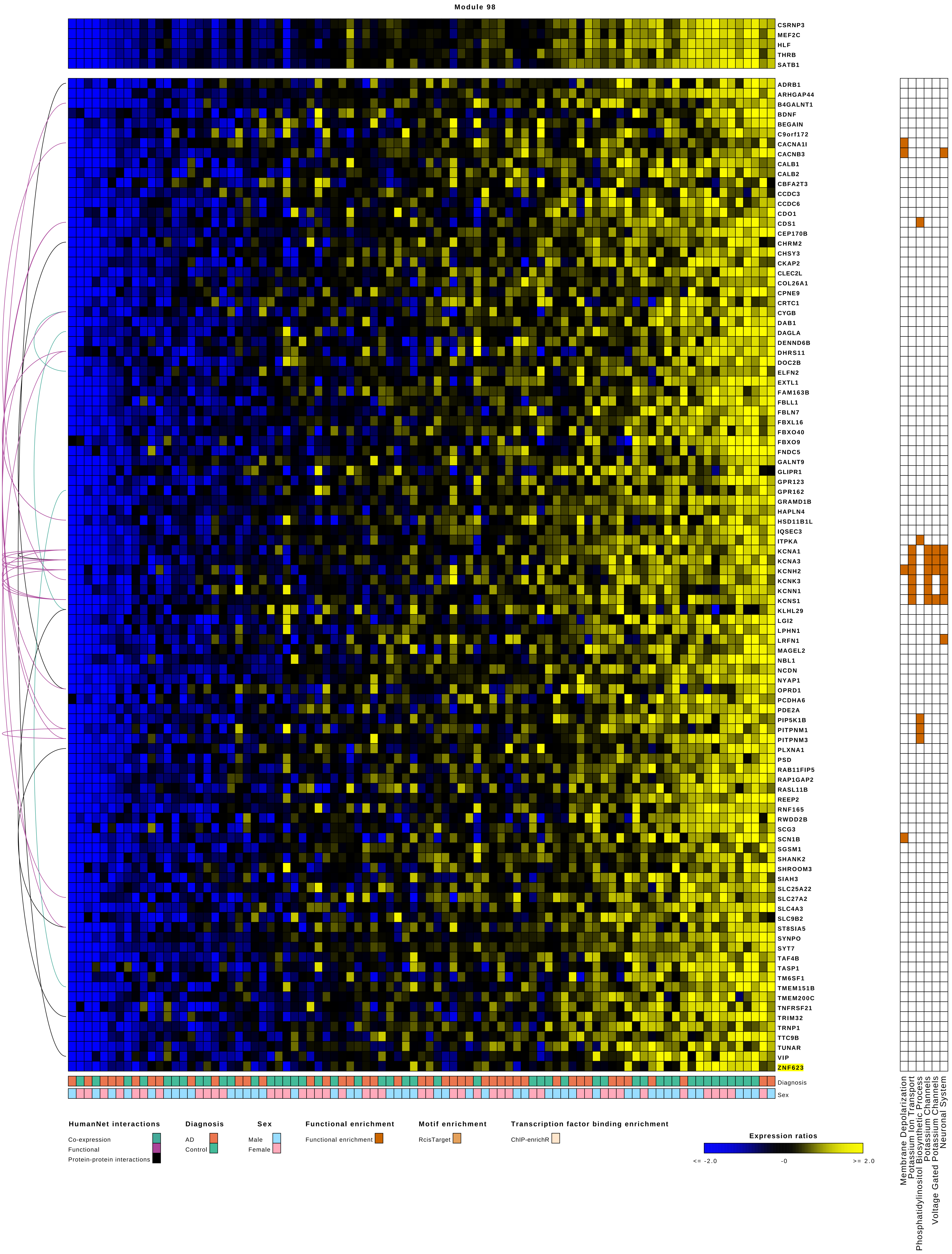
Module 98
CSRNP3
MEF2C
HLF
THRB
SATB1
ADRB1
ARHGAP44
B4GALNT1
BDNF
BEGAIN
C9orf172
CACNA1I
CACNB3
CALB1
CALB2
CBFA2T3
CCDC3
CCDC6
CDO1
CDS1
CEP170B
CHRM2
CHSY3
CKAP2
CLEC2L
COL26A1
CPNE9
CRTC1
CYGB
DAB1
DAGLA
DENND6B
DHRS11
DOC2B
ELFN2
EXTL1
FAM163B
FBLL1
FBLN7
FBXL16
FBXO40
FBXO9
FNDC5
GALNT9
GLIPR1
GPR123
GPR162
GRAMD1B
HAPLN4
HSD11B1L
IQSEC3
ITPKA
KCNA1
KCNA3
KCNH2
KCNK3
KCNN1
KCNS1
KLHL29
LGI2
LPHN1
LRFN1
MAGEL2
NBL1
NCDN
NYAP1
OPRD1
PCDHA6
PDE2A
PIP5K1B
PITPNM1
PITPNM3
PLXNA1
PSD
RAB11FIP5
RAP1GAP2
RASL11B
REEP2
RNF165
RWDD2B
SCG3
SCN1B
SGSM1
SHANK2
SHROOM3
SIAH3
SLC25A22
SLC27A2
SLC4A3
SLC9B2
ST8SIA5
SYNPO
SYT7
TAF4B
TASP1
TM6SF1
TMEM151B
TMEM200C
TNFRSF21
TRIM32
TRNP1
TTC9B
TUNAR
VIP
ZNF623
Diagnosis
Sex
Membrane Depolarization
Potassium Ion Transport
Phosphatidylinositol Biosynthetic Process
Potassium Channels
Voltage Gated Potassium Channels
Neuronal System
HumanNet interactions
Co-expression
Functional
Protein-protein interactions
Diagnosis
AD
Control
Sex
Male
Female
Functional enrichment
Functional enrichment
Motif enrichment
RcisTarget
Transcription factor binding enrichment
ChIP-enrichR
Expression ratios
<= -2.0
-0
>= 2.0

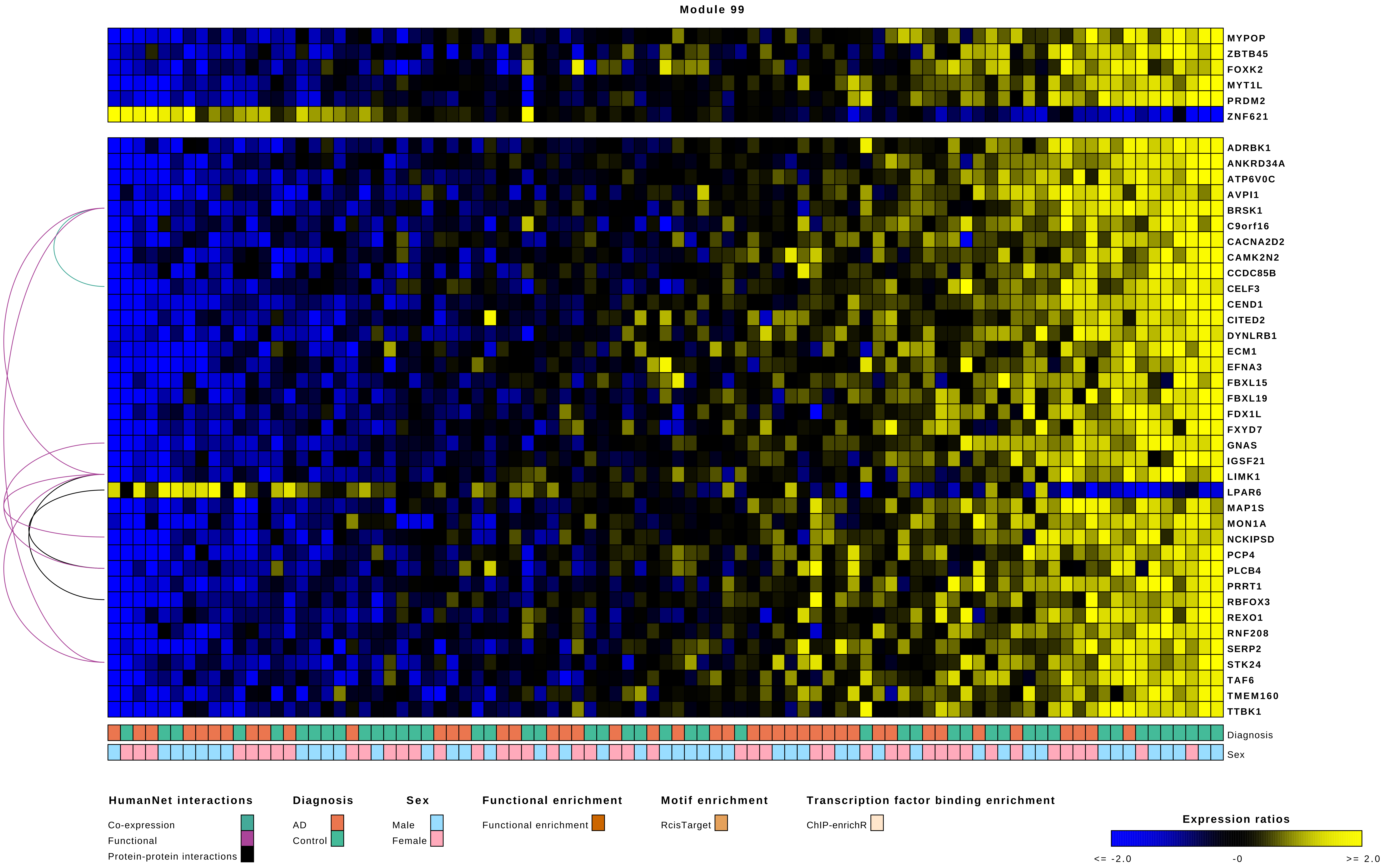
Module 99
MYPOP
ZBTB45
FOXK2
MYT1L
PRDM2
ZNF621
ADRBK1
ANKRD34A
ATP6V0C
AVPI1
BRSK1
C9orf16
CACNA2D2
CAMK2N2
CCDC85B
CELF3
CEND1
CITED2
DYNLRB1
ECM1
EFNA3
FBXL15
FBXL19
FDX1L
FXYD7
GNAS
IGSF21
LIMK1
LPAR6
MAP1S
MON1A
NCKIPSD
PCP4
PLCB4
PRRT1
RBFOX3
REXO1
RNF208
SERP2
STK24
TAF6
TMEM160
TTBK1
Diagnosis
Sex
HumanNet interactions
Co-expression
Functional
Protein-protein interactions
Diagnosis
AD
Control
Sex
Male
Female
Functional enrichment
Functional enrichment
Motif enrichment
RcisTarget
Transcription factor binding enrichment
ChIP-enrichR
Expression ratios
<= -2.0
-0
>= 2.0

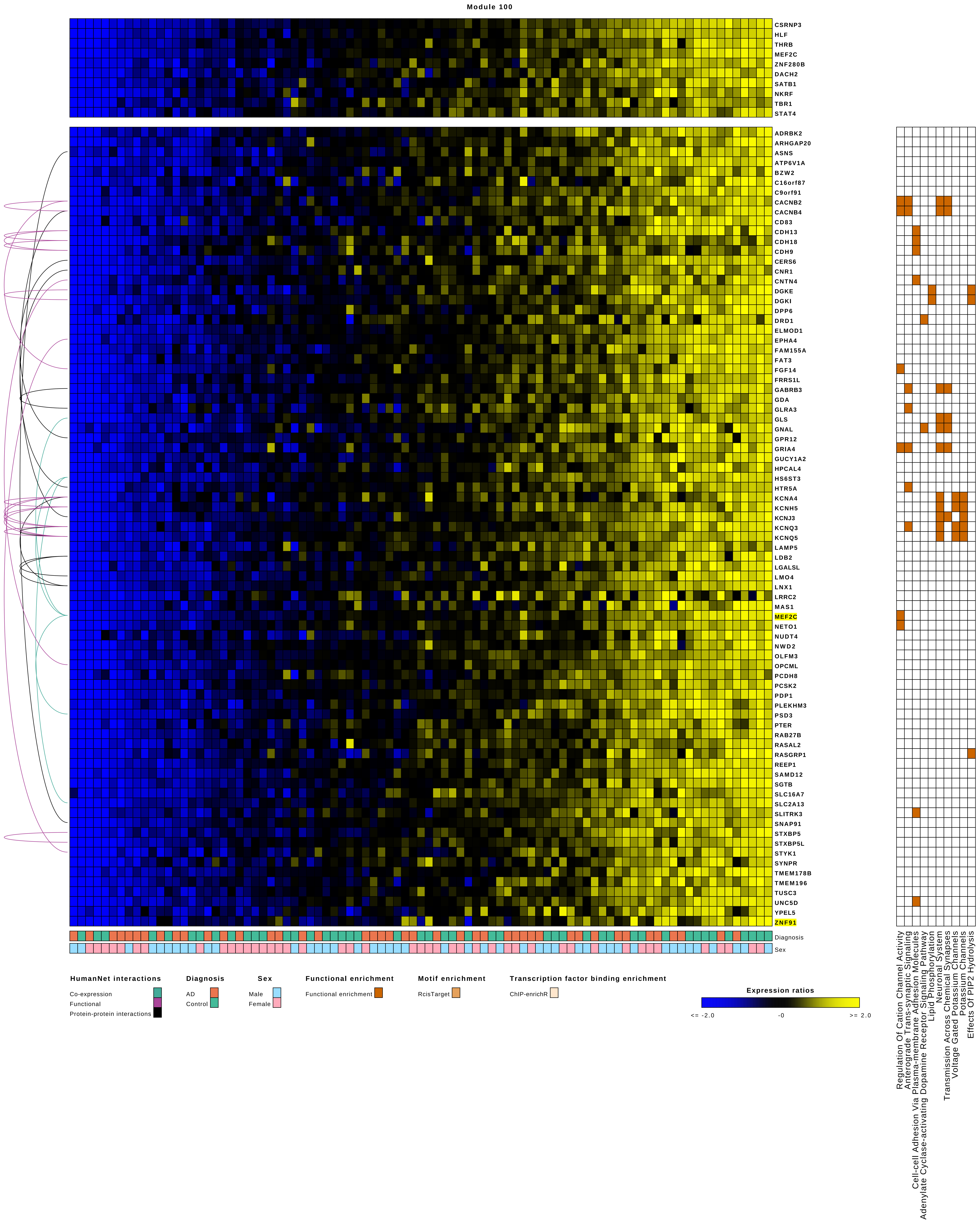

Module 100
CSRNP3
HLF
THRB
MEF2C
ZNF280B
DACH2
SATB1
NKRF
TBR1
STAT4
ADRBK2
ARHGAP20
ASNS
ATP6V1A
BZW2
C16orf87
C9orf91
CACNB2
CACNB4
CD83
CDH13
CDH18
CDH9
CERS6
CNR1
CNTN4
DGKE
DGKI
DPP6
DRD1
ELMOD1
EPHA4
FAM155A
FAT3
FGF14
FRRS1L
GABRB3
GDA
GLRA3
GLS
GNAL
GPR12
GRIA4
GUCY1A2
HPCAL4
HS6ST3
HTR5A
KCNA4
KCNH5
KCNJ3
KCNQ3
KCNQ5
LAMP5
LDB2
LGALSL
LMO4
LNX1
LRRC2
MAS1
MEF2C
NETO1
NUDT4
NWD2
OLFM3
OPCML
PCDH8
PCSK2
PDP1
PLEKHM3
PSD3
PTER
RAB27B
RASAL2
RASGRP1
REEP1
SAMD12
SGTB
SLC16A7
SLC2A13
SLITRK3
SNAP91
STXBP5
STXBP5L
STYK1
SYNPR
TMEM178B
TMEM196
TUSC3
UNC5D
YPEL5
ZNF91
Diagnosis
Sex
HumanNet interactions
Co-expression
Functional
Protein-protein interactions
Diagnosis
AD
Control
Sex
Male
Female
Functional enrichment
Functional enrichment
Motif enrichment
RcisTarget
Transcription factor binding enrichment
ChIP-enrichR
Expression ratios
<= -2.0
-0
>= 2.0
Regulation Of Cation Channel Activity
Anterograde Trans-synaptic Signaling
Cell-cell Adhesion Via Plasma-membrane Adhesion Molecules
Adenylate Cyclase-activating Dopamine Receptor Signaling Pathway
Lipid Phosphorylation
Neuronal System
Transmission Across Chemical Synapses
Voltage Gated Potassium Channels
Potassium Channels
Effects Of PIP2 Hydrolysis

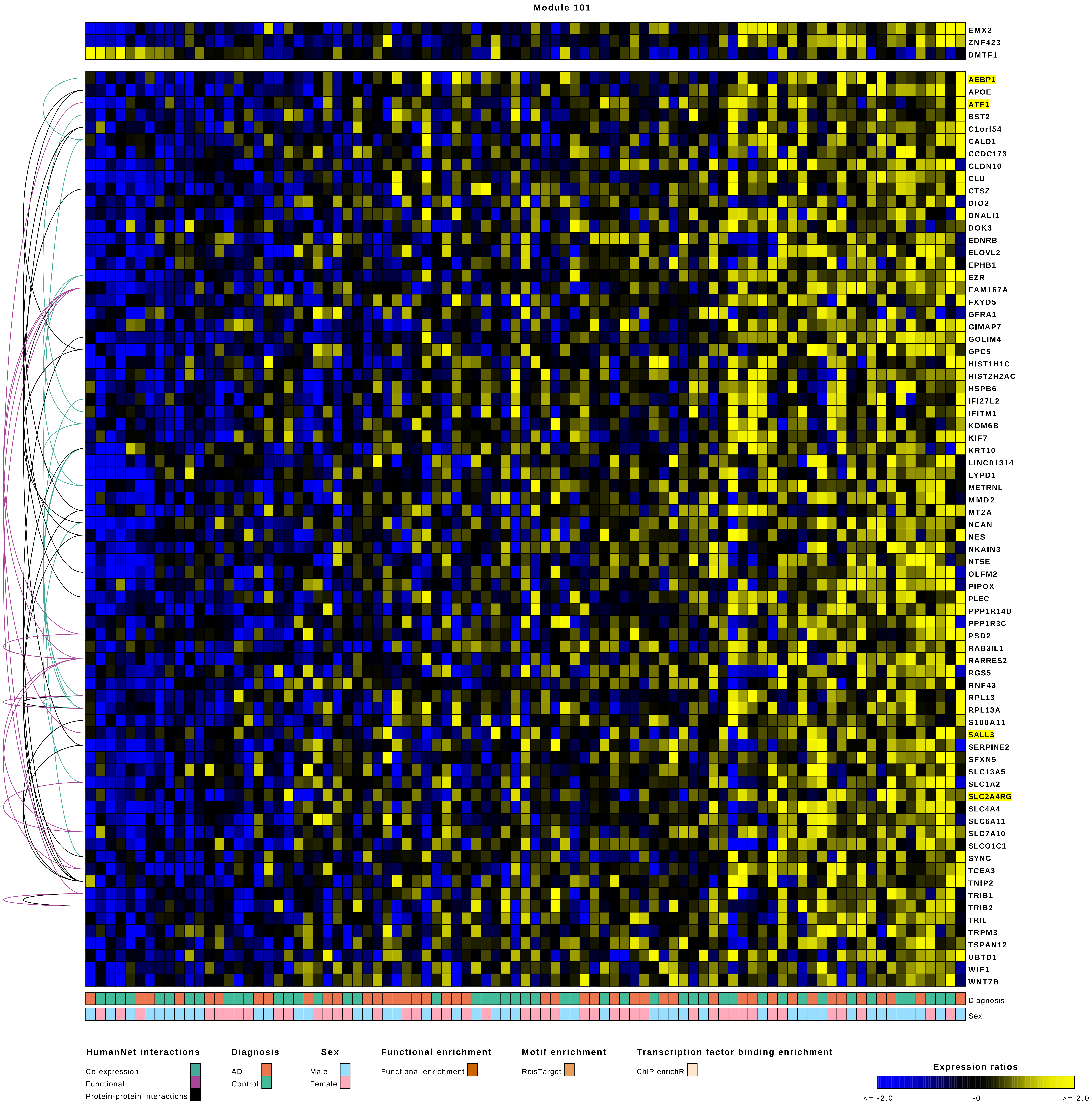
Module 101
EMX2
ZNF423
DMTF1
AEBP1
APOE
ATF1
BST2
C1orf54
CALD1
CCDC173
CLDN10
CLU
CTSZ
DIO2
DNALI1
DOK3
EDNRB
ELOVL2
EPHB1
EZR
FAM167A
FXYD5
GFRA1
GIMAP7
GOLIM4
GPC5
HIST1H1C
HIST2H2AC
HSPB6
IFI27L2
IFITM1
KDM6B
KIF7
KRT10
LINC01314
LYPD1
METRNL
MMD2
MT2A
NCAN
NES
NKAIN3
NT5E
OLFM2
PIPOX
PLEC
PPP1R14B
PPP1R3C
PSD2
RAB3IL1
RARRES2
RGS5
RNF43
RPL13
RPL13A
S100A11
SALL3
SERPINE2
SFXN5
SLC13A5
SLC1A2
SLC2A4RG
SLC4A4
SLC6A11
SLC7A10
SLCO1C1
SYNC
TCEA3
TNIP2
TRIB1
TRIB2
TRIL
TRPM3
TSPAN12
UBTD1
WIF1
WNT7B
Diagnosis
Sex
HumanNet interactions
Co-expression
Functional
Protein-protein interactions
Diagnosis
AD
Control
Sex
Male
Female
Functional enrichment
Functional enrichment
Motif enrichment
RcisTarget
Transcription factor binding enrichment
ChIP-enrichR
Expression ratios
<= -2.0
-0
>= 2.0

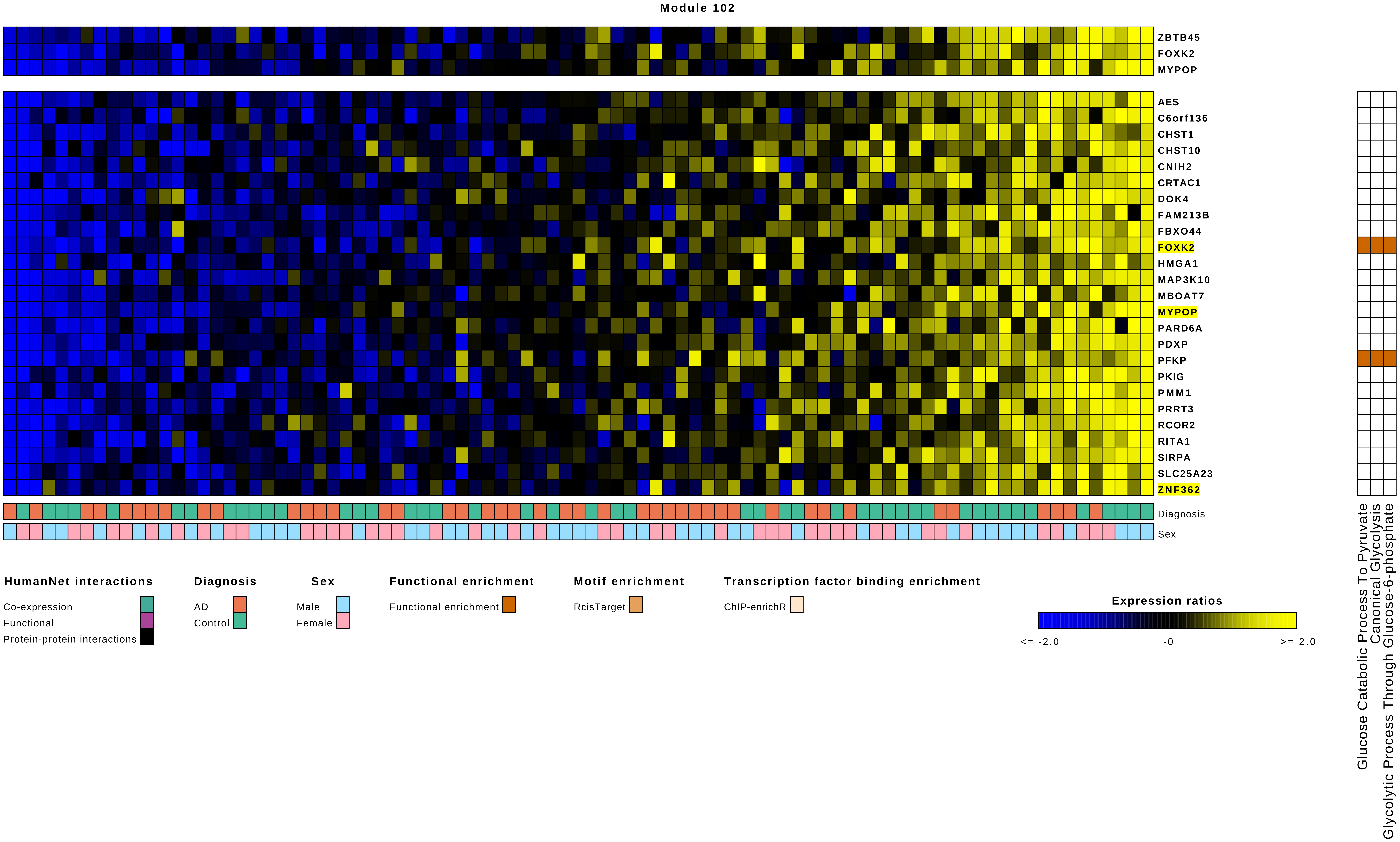
Module 102
ZBTB45
FOXK2
MYPOP
AES
C6orf136
CHST1
CHST10
CNIH2
CRTAC1
DOK4
FAM213B
FBXO44
FOXK2
HMGA1
MAP3K10
MBOAT7
MYPOP
PARD6A
PDXP
PFKP
PKIG
PMM1
PRRT3
RCOR2
RITA1
SIRPA
SLC25A23
ZNF362
Diagnosis
Sex
Glucose Catabolic Process To Pyruvate
Canonical Glycolysis
Glycolytic Process Through Glucose-6-phosphate
HumanNet interactions
Co-expression
Functional
Protein-protein interactions
Diagnosis
AD
Control
Sex
Male
Female
Functional enrichment
Functional enrichment
Motif enrichment
RcisTarget
Transcription factor binding enrichment
ChIP-enrichR
Expression ratios
<= -2.0
-0
>= 2.0

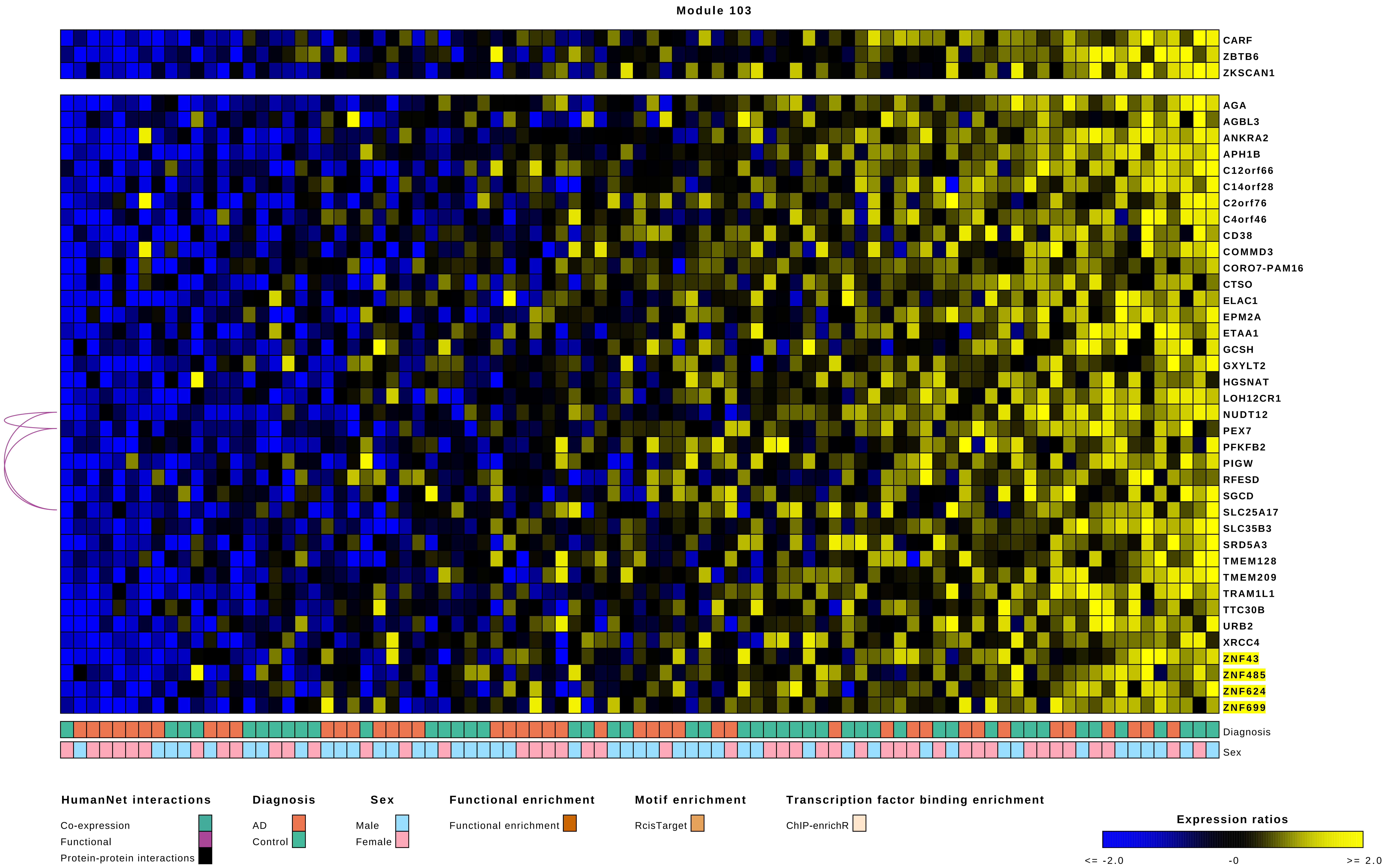

Module 103
CARF
ZBTB6
ZKSCAN1
AGA
AGBL3
ANKRA2
APH1B
C12orf66
C14orf28
C2orf76
C4orf46
CD38
COMMD3
CORO7-PAM16
CTSO
ELAC1
EPM2A
ETAA1
GCSH
GXYLT2
HGSNAT
LOH12CR1
NUDT12
PEX7
PFKFB2
PIGW
RFESD
SGCD
SLC25A17
SLC35B3
SRD5A3
TMEM128
TMEM209
TRAM1L1
TTC30B
URB2
XRCC4
ZNF43
ZNF485
ZNF624
ZNF699
Diagnosis
Sex
HumanNet interactions
Co-expression
Functional
Protein-protein interactions
Diagnosis
AD
Control
Sex
Male
Female
Functional enrichment
Functional enrichment
Motif enrichment
RcisTarget
Transcription factor binding enrichment
ChIP-enrichR
Expression ratios
<= -2.0
-0
>= 2.0

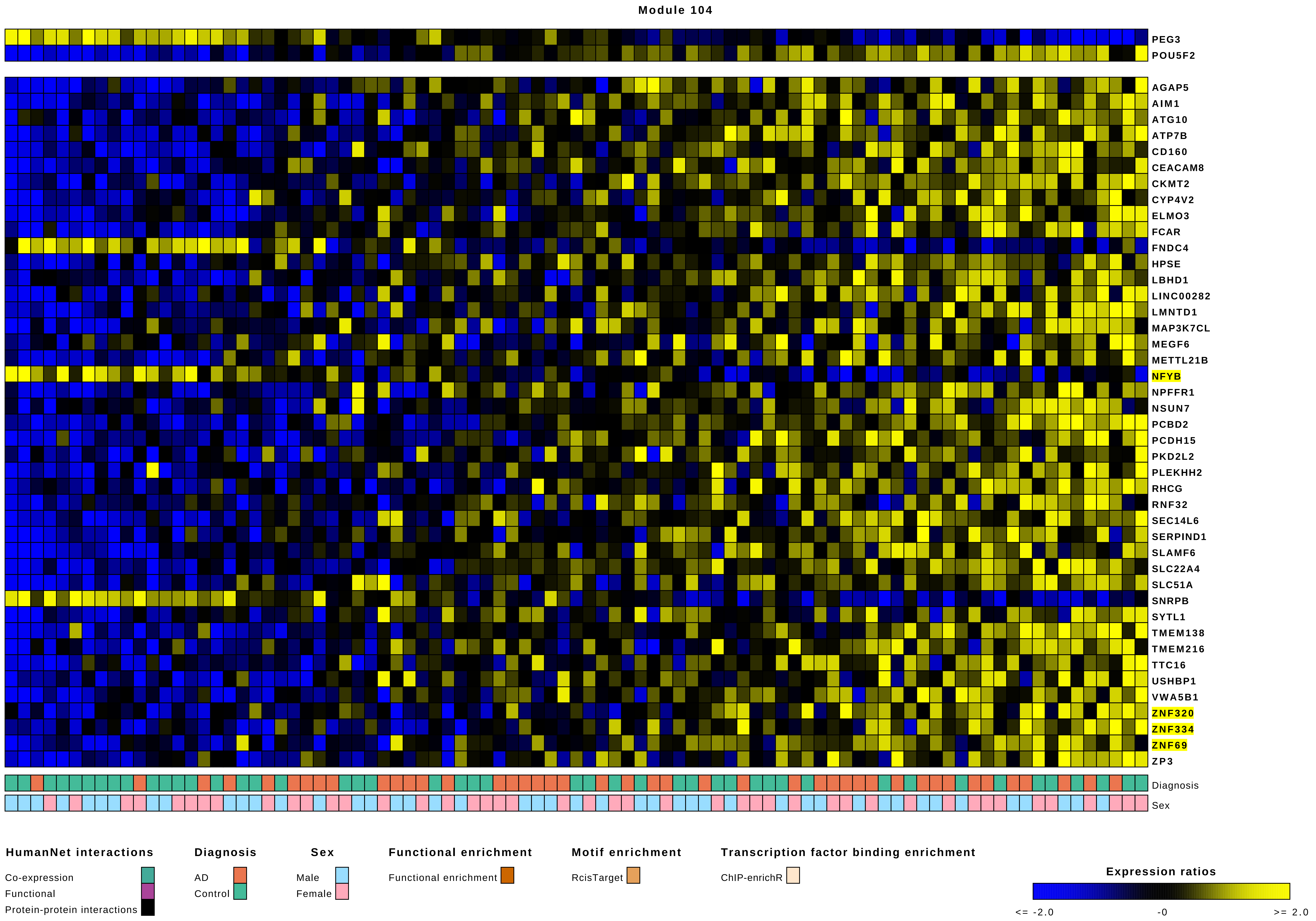
Module 104
PEG3
POU5F2
AGAP5
AIM1
ATG10
ATP7B
CD160
CEACAM8
CKMT2
CYP4V2
ELMO3
FCAR
FNDC4
HPSE
LBHD1
LINC00282
LMNTD1
MAP3K7CL
MEGF6
METTL21B
NFYB
NPFFR1
NSUN7
PCBD2
PCDH15
PKD2L2
PLEKHH2
RHCG
RNF32
SEC14L6
SERPIND1
SLAMF6
SLC22A4
SLC51A
SNRPB
SYTL1
TMEM138
TMEM216
TTC16
USHBP1
VWA5B1
ZNF320
ZNF334
ZNF69
ZP3
Diagnosis
Sex
HumanNet interactions
Co-expression
Functional
Protein-protein interactions
Diagnosis
AD
Control
Sex
Male
Female
Functional enrichment
Functional enrichment
Motif enrichment
RcisTarget
Transcription factor binding enrichment
ChIP-enrichR
Expression ratios
<= -2.0
-0
>= 2.0

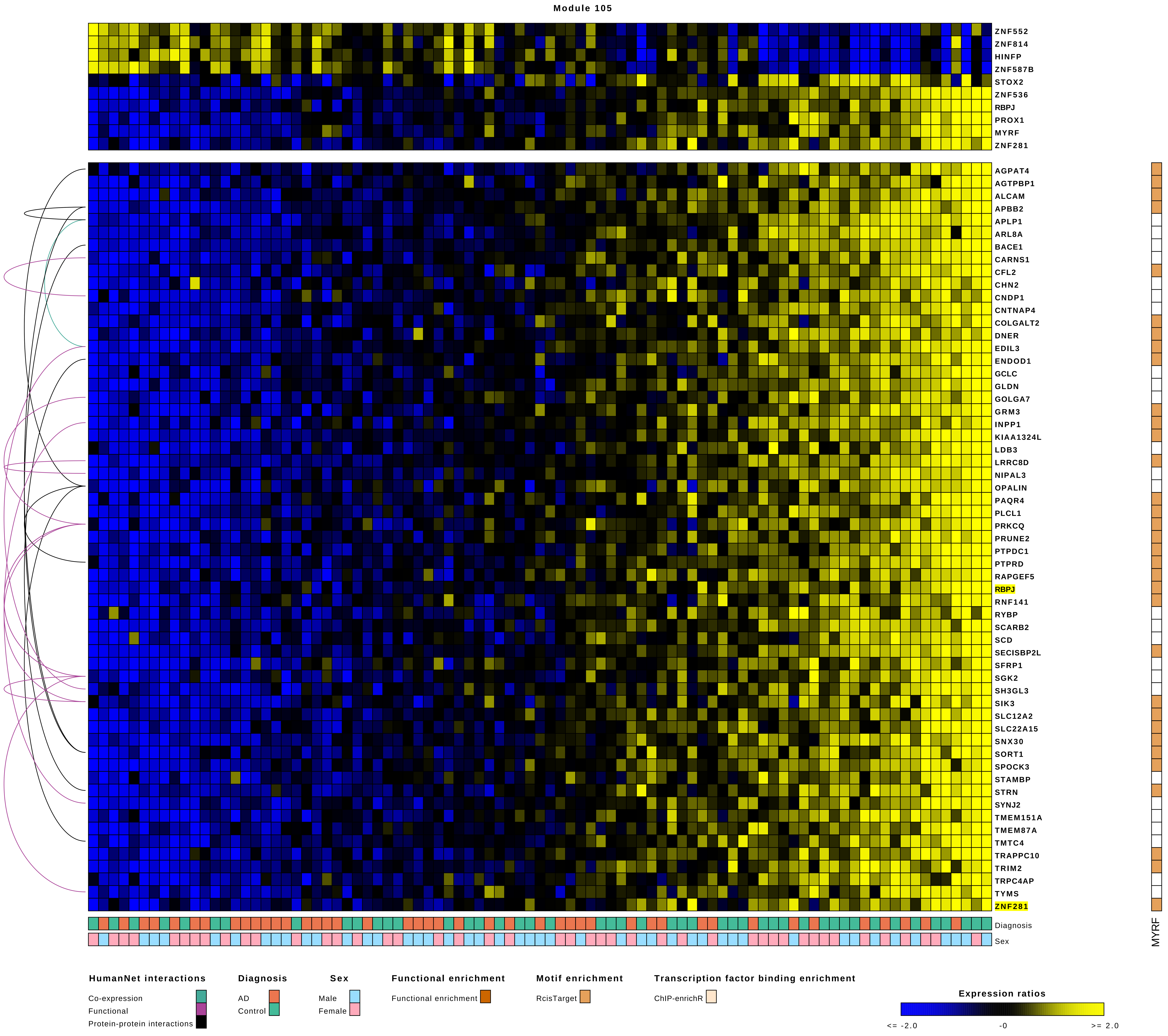
Module 105
ZNF552
ZNF814
HINFP
ZNF587B
STOX2
ZNF536
RBPJ
PROX1
MYRF
ZNF281
AGPAT4
AGTPBP1
ALCAM
APBB2
APLP1
ARL8A
BACE1
CARNS1
CFL2
CHN2
CNDP1
CNTNAP4
COLGALT2
DNER
EDIL3
ENDOD1
GCLC
GLDN
GOLGA7
GRM3
INPP1
KIAA1324L
LDB3
LRRC8D
NIPAL3
OPALIN
PAQR4
PLCL1
PRKCQ
PRUNE2
PTPDC1
PTPRD
RAPGEF5
RBPJ
RNF141
RYBP
SCARB2
SCD
SECISBP2L
SFRP1
SGK2
SH3GL3
SIK3
SLC12A2
SLC22A15
SNX30
SORT1
SPOCK3
STAMBP
STRN
SYNJ2
TMEM151A
TMEM87A
TMTC4
TRAPPC10
TRIM2
TRPC4AP
TYMS
ZNF281
Diagnosis
Sex
MYRF
HumanNet interactions
Co-expression
Functional
Protein-protein interactions
Diagnosis
AD
Control
Sex
Male
Female
Functional enrichment
Functional enrichment
Motif enrichment
RcisTarget
Transcription factor binding enrichment
ChIP-enrichR
Expression ratios
<= -2.0
-0
>= 2.0

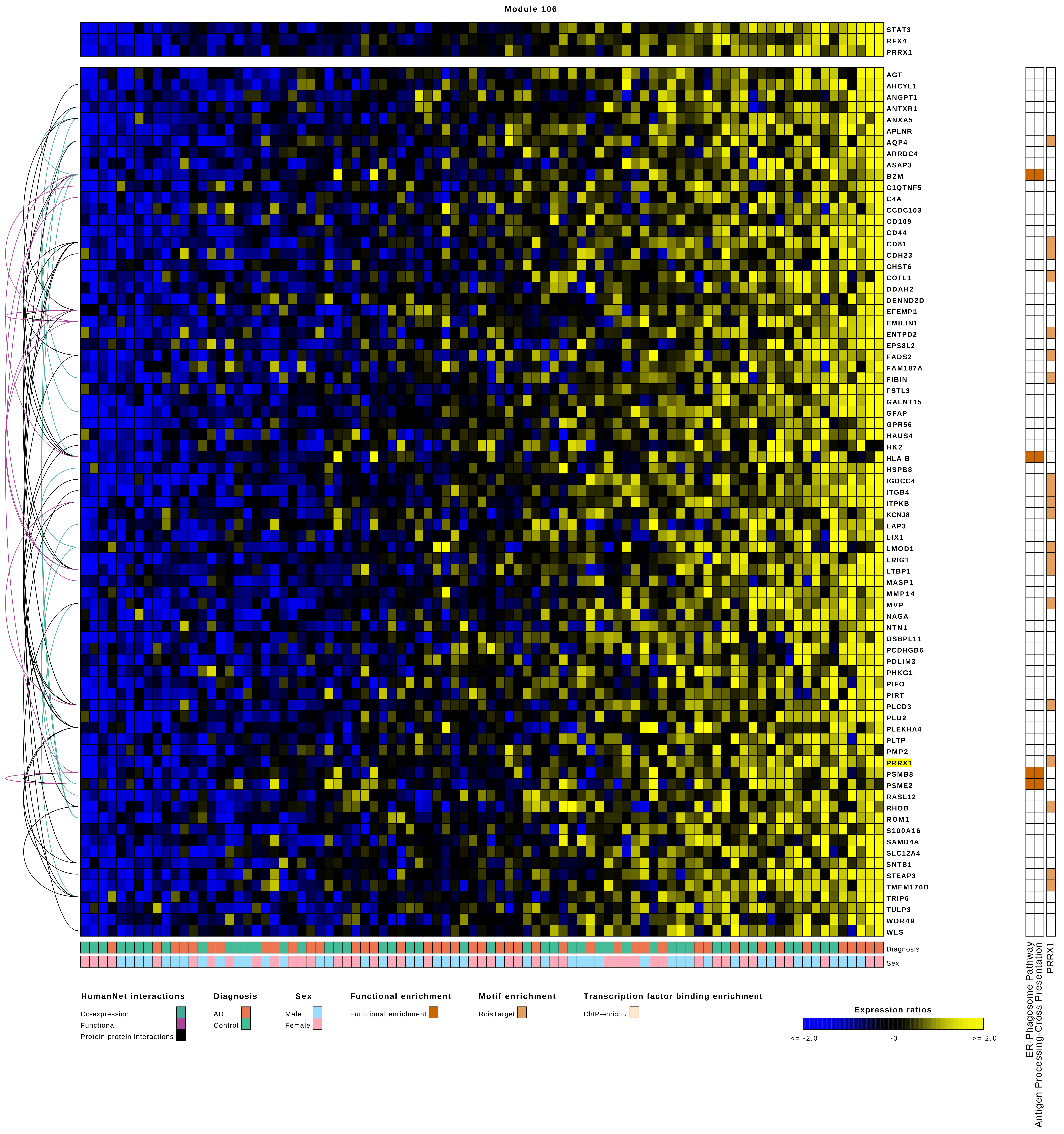
Module 106
STAT3
RFX4
PRRX1
AGT
AHCYL1
ANGPT1
ANTXR1
ANXA5
APLNR
AQP4
ARRDC4
ASAP3
B2M
C1QTNF5
C4A
CCDC103
CD109
CD44
CD81
CDH23
CHST6
COTL1
DDAH2
DENND2D
EFEMP1
EMILIN1
ENTPD2
EPS8L2
FADS2
FAM187A
FIBIN
FSTL3
GALNT15
GFAP
GPR56
HAUS4
HK2
HLA-B
HSPB8
IGDCC4
ITGB4
ITPKB
KCNJ8
LAP3
LIX1
LMOD1
LRIG1
LTBP1
MASP1
MMP14
MVP
NAGA
NTN1
OSBPL11
PCDHGB6
PDLIM3
PHKG1
PIFO
PIRT
PLCD3
PLD2
PLEKHA4
PLTP
PMP2
PRRX1
PSMB8
PSME2
RASL12
RHOB
ROM1
S100A16
SAMD4A
SLC12A4
SNTB1
STEAP3
TMEM176B
TRIP6
TULP3
WDR49
WLS
Diagnosis
Sex
ER-Phagosome Pathway
Antigen Processing-Cross Presentation
PRRX1
HumanNet interactions
Co-expression
Functional
Protein-protein interactions
Diagnosis
AD
Control
Sex
Male
Female
Functional enrichment
Functional enrichment
Motif enrichment
RcisTarget
Transcription factor binding enrichment
ChIP-enrichR
Expression ratios
<= -2.0
-0
>= 2.0

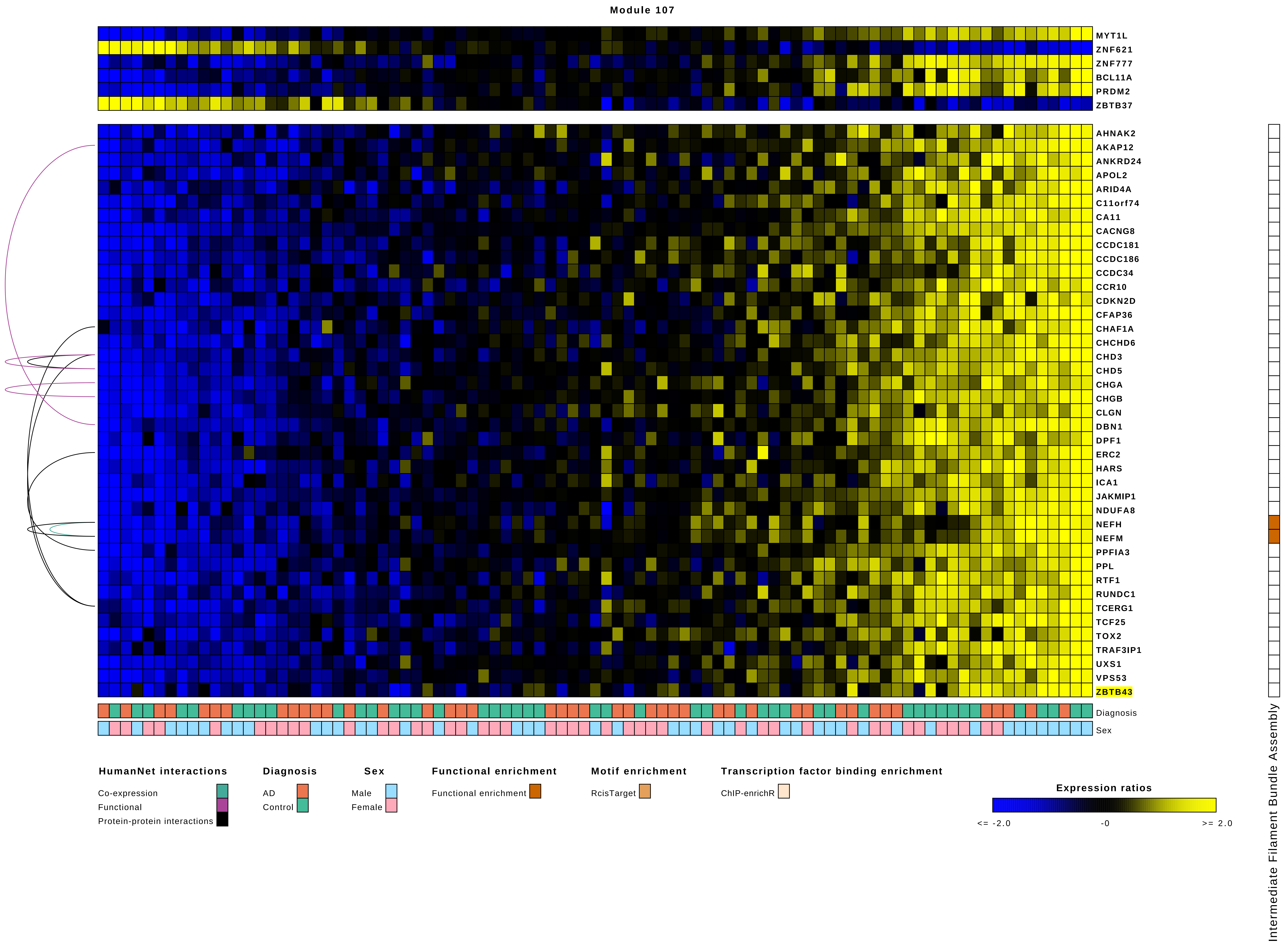

Module 107
MYT1L
ZNF621
ZNF777
BCL11A
PRDM2
ZBTB37
AHNAK2
AKAP12
ANKRD24
APOL2
ARID4A
C11orf74
CA11
CACNG8
CCDC181
CCDC186
CCDC34
CCR10
CDKN2D
CFAP36
CHAF1A
CHCHD6
CHD3
CHD5
CHGA
CHGB
CLGN
DBN1
DPF1
ERC2
HARS
ICA1
JAKMIP1
NDUFA8
NEFH
NEFM
PPFIA3
PPL
RTF1
RUNDC1
TCERG1
TCF25
TOX2
TRAF3IP1
UXS1
VPS53
ZBTB43
Diagnosis
Sex
Intermediate Filament Bundle Assembly
HumanNet interactions
Co-expression
Functional
Protein-protein interactions
Diagnosis
AD
Control
Sex
Male
Female
Functional enrichment
Functional enrichment
Motif enrichment
RcisTarget
Transcription factor binding enrichment
ChIP-enrichR
Expression ratios
<= -2.0
-0
>= 2.0

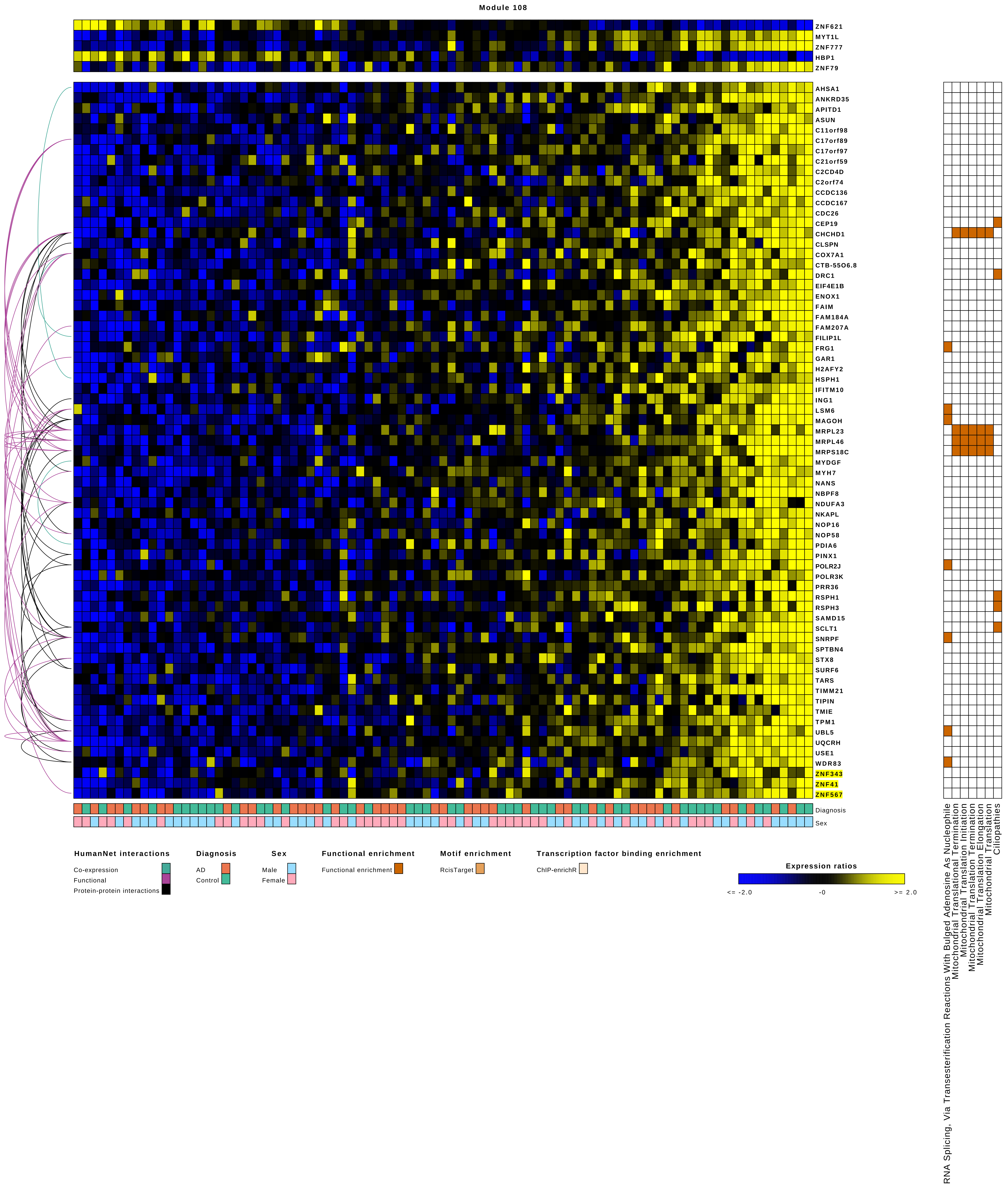
Module 108
ZNF621
MYT1L
ZNF777
HBP1
ZNF79
AHSA1
ANKRD35
APITD1
ASUN
C11orf98
C17orf89
C17orf97
C21orf59
C2CD4D
C2orf74
CCDC136
CCDC167
CDC26
CEP19
CHCHD1
CLSPN
COX7A1
CTB-55O6.8
DRC1
EIF4E1B
ENOX1
FAIM
FAM184A
FAM207A
FILIP1L
FRG1
GAR1
H2AFY2
HSPH1
IFITM10
ING1
LSM6
MAGOH
MRPL23
MRPL46
MRPS18C
MYDGF
MYH7
NANS
NBPF8
NDUFA3
NKAPL
NOP16
NOP58
PDIA6
PINX1
POLR2J
POLR3K
PRR36
RSPH1
RSPH3
SAMD15
SCLT1
SNRPF
SPTBN4
STX8
SURF6
TARS
TIMM21
TIPIN
TMIE
TPM1
UBL5
UQCRH
USE1
WDR83
ZNF343
ZNF41
ZNF567
Diagnosis
Sex
RNA Splicing, Via Transesterification Reactions With Bulged Adenosine As Nucleophile
Mitochondrial Translational Termination
Mitochondrial Translation Initiation
Mitochondrial Translation Termination
Mitochondrial Translation Elongation
Mitochondrial Translation
Ciliopathies
HumanNet interactions
Co-expression
Functional
Protein-protein interactions
Diagnosis
AD
Control
Sex
Male
Female
Functional enrichment
Functional enrichment
Motif enrichment
RcisTarget
Transcription factor binding enrichment
ChIP-enrichR
Expression ratios
<= -2.0
-0
>= 2.0

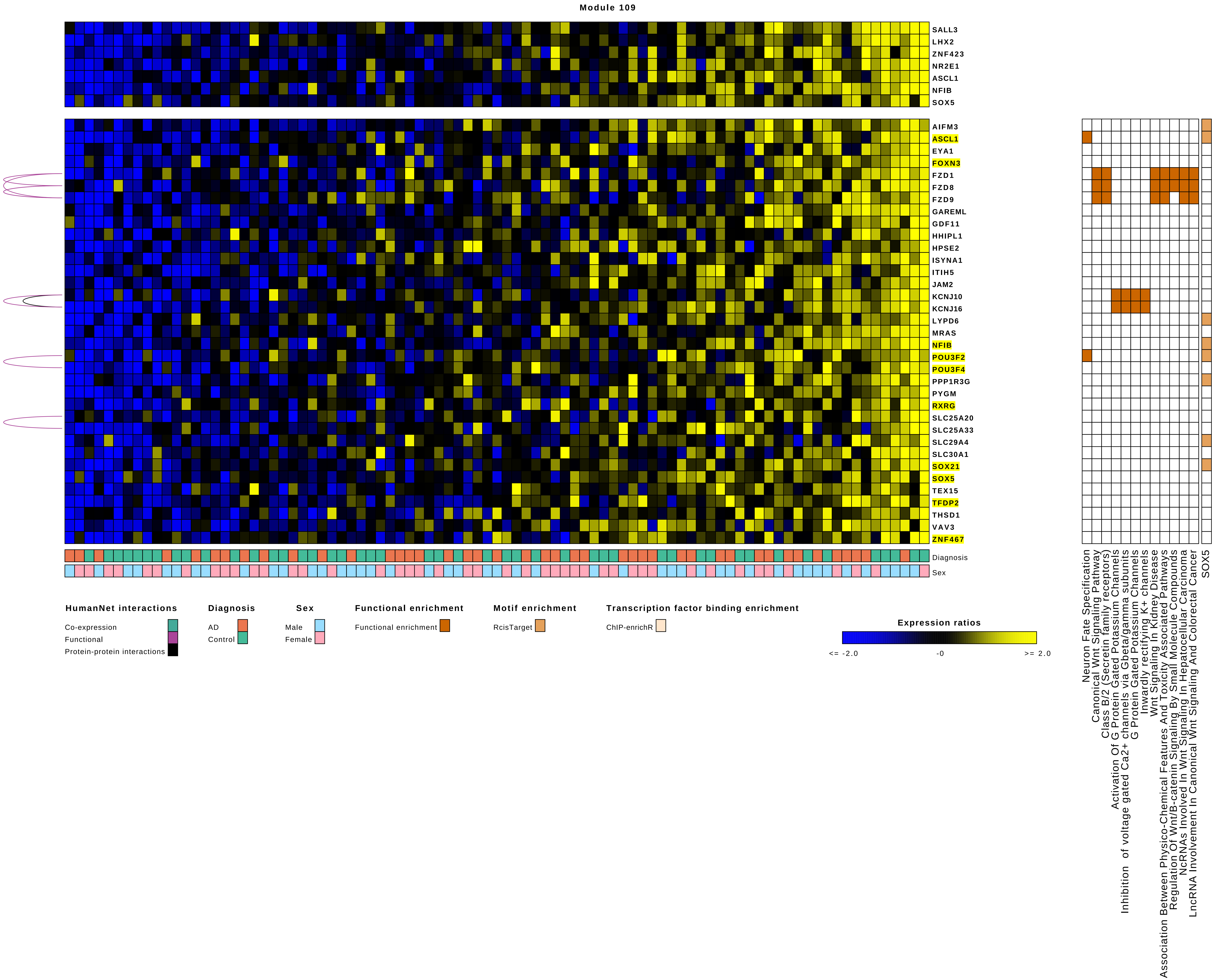

Module 109
SALL3
LHX2
ZNF423
NR2E1
ASCL1
NFIB
SOX5
AIFM3
ASCL1
EYA1
FOXN3
FZD1
FZD8
FZD9
GAREML
GDF11
HHIPL1
HPSE2
ISYNA1
ITIH5
JAM2
KCNJ10
KCNJ16
LYPD6
MRAS
NFIB
POU3F2
POU3F4
PPP1R3G
PYGM
RXRG
SLC25A20
SLC25A33
SLC29A4
SLC30A1
SOX21
SOX5
TEX15
TFDP2
THSD1
VAV3
ZNF467
Diagnosis
Sex
HumanNet interactions
Co-expression
Functional
Protein-protein interactions
Diagnosis
AD
Control
Sex
Male
Female
Functional enrichment
Functional enrichment
Motif enrichment
RcisTarget
Transcription factor binding enrichment
ChIP-enrichR
Expression ratios
<= -2.0
-0
>= 2.0
Neuron Fate Specification
Canonical Wnt Signaling Pathway
Class B/2 (Secretin family receptors)
Activation Of G Protein Gated Potassium Channels
Inhibition of voltage gated Ca2+ channels via Gbeta/gamma subunits
G Protein Gated Potassium Channels
Inwardly rectifying K+ channels
Wnt Signaling In Kidney Disease
Association Between Physico-Chemical Features And Toxicity Associated Pathways
Regulation Of Wnt/B-catenin Signaling By Small Molecule Compounds
NcRNAs Involved In Wnt Signaling In Hepatocellular Carcinoma
LncRNA Involvement In Canonical Wnt Signaling And Colorectal Cancer
SOX5

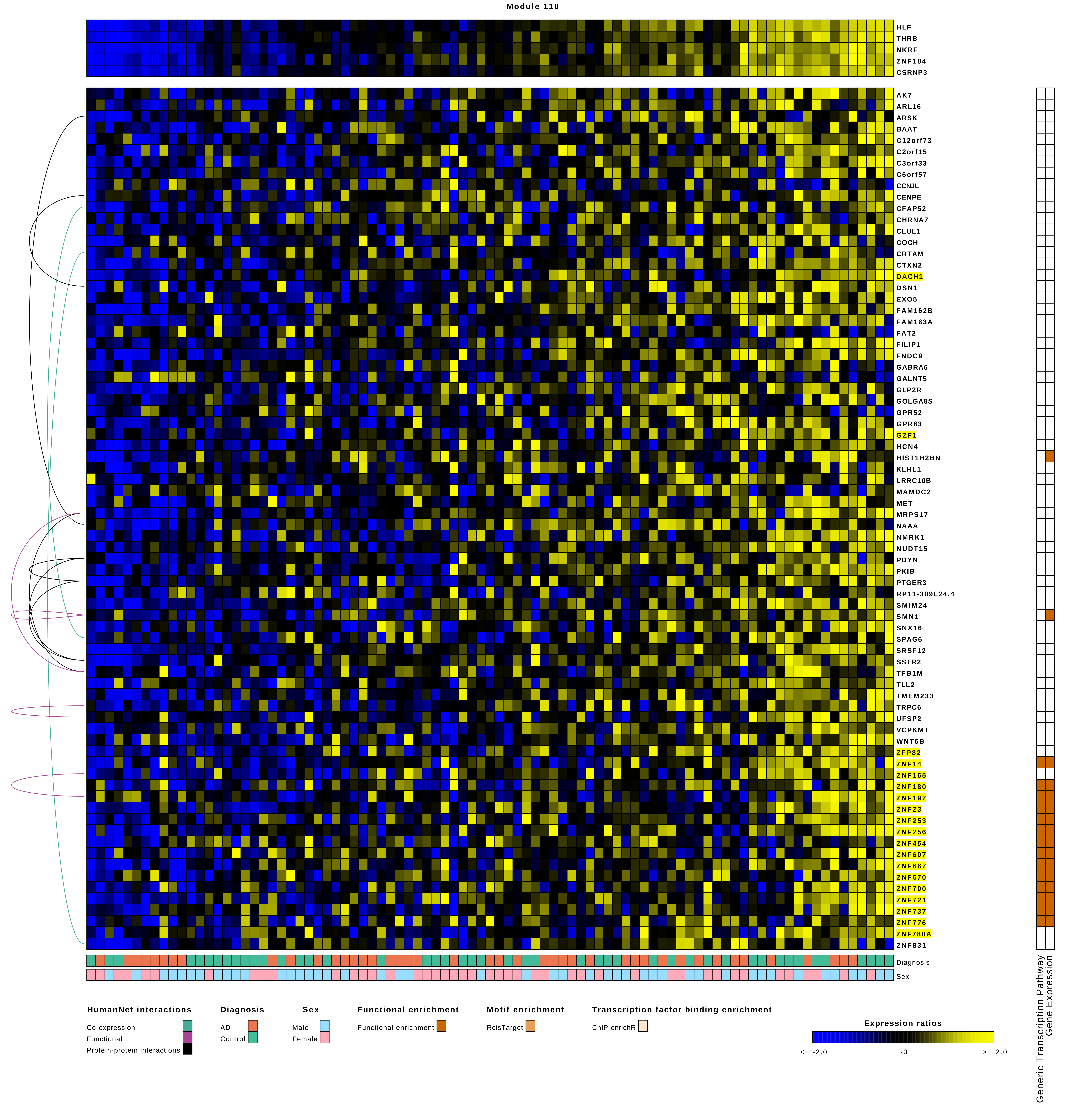

Module 110
HLF
THRB
NKRF
ZNF184
CSRNP3
AK7
ARL16
ARSK
BAAT
C12orf73
C2orf15
C3orf33
C6orf57
CCNJL
CENPE
CFAP52
CHRNA7
CLUL1
COCH
CRTAM
CTXN2
DACH1
DSN1
EXO5
FAM162B
FAM163A
FAT2
FILIP1
FNDC9
GABRA6
GALNT5
GLP2R
GOLGA8S
GPR52
GPR83
GZF1
HCN4
HIST1H2BN
KLHL1
LRRC10B
MAMDC2
MET
MRPS17
NAAA
NMRK1
NUDT15
PDYN
PKIB
PTGER3
RP11-309L24.4
SMIM24
SMN1
SNX16
SPAG6
SRSF12
SSTR2
TFB1M
TLL2
TMEM233
TRPC6
UFSP2
VCPKMT
WNT5B
ZFP82
ZNF14
ZNF165
ZNF180
ZNF197
ZNF23
ZNF253
ZNF256
ZNF454
ZNF607
ZNF667
ZNF670
ZNF700
ZNF721
ZNF737
ZNF776
ZNF780A
ZNF831
Diagnosis
Sex
Generic Transcription Pathway
Gene Expression
HumanNet interactions
Co-expression
Functional
Protein-protein interactions
Diagnosis
AD
Control
Sex
Male
Female
Functional enrichment
Functional enrichment
Motif enrichment
RcisTarget
Transcription factor binding enrichment
ChIP-enrichR
Expression ratios
<= -2.0
-0
>= 2.0

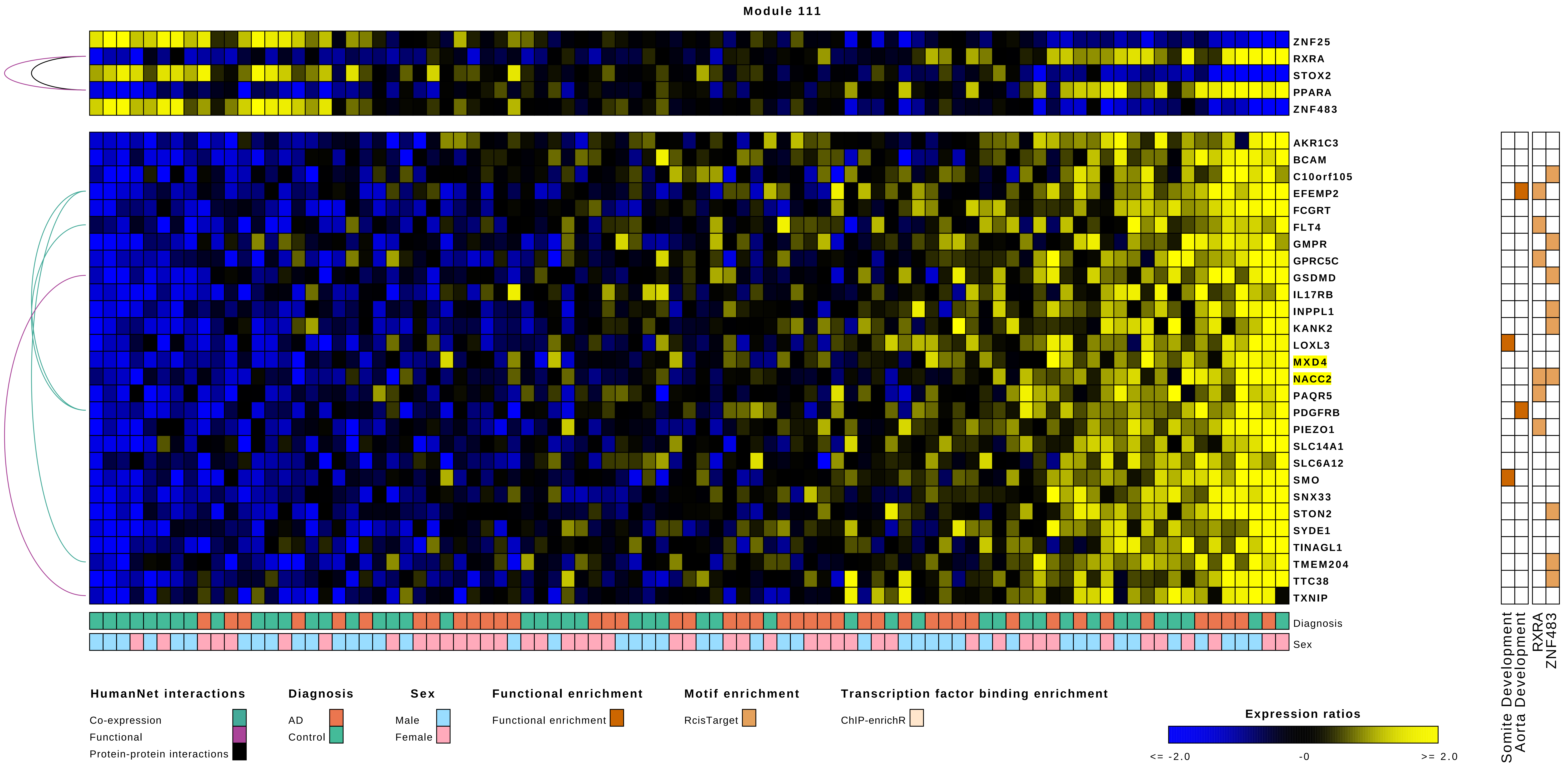
Module 111
ZNF25
RXRA
STOX2
PPARA
ZNF483
AKR1C3
BCAM
C10orf105
EFEMP2
FCGRT
FLT4
GMPR
GPRC5C
GSDMD
IL17RB
INPPL1
KANK2
LOXL3
MXD4
NACC2
PAQR5
PDGFRB
PIEZO1
SLC14A1
SLC6A12
SMO
SNX33
STON2
SYDE1
TINAGL1
TMEM204
TTC38
TXNIP
Diagnosis
Sex
Somite Development
Aorta Development
RXRA
ZNF483
HumanNet interactions
Co-expression
Functional
Protein-protein interactions
Diagnosis
AD
Control
Sex
Male
Female
Functional enrichment
Functional enrichment
Motif enrichment
RcisTarget
Transcription factor binding enrichment
ChIP-enrichR
Expression ratios
<= -2.0
-0
>= 2.0

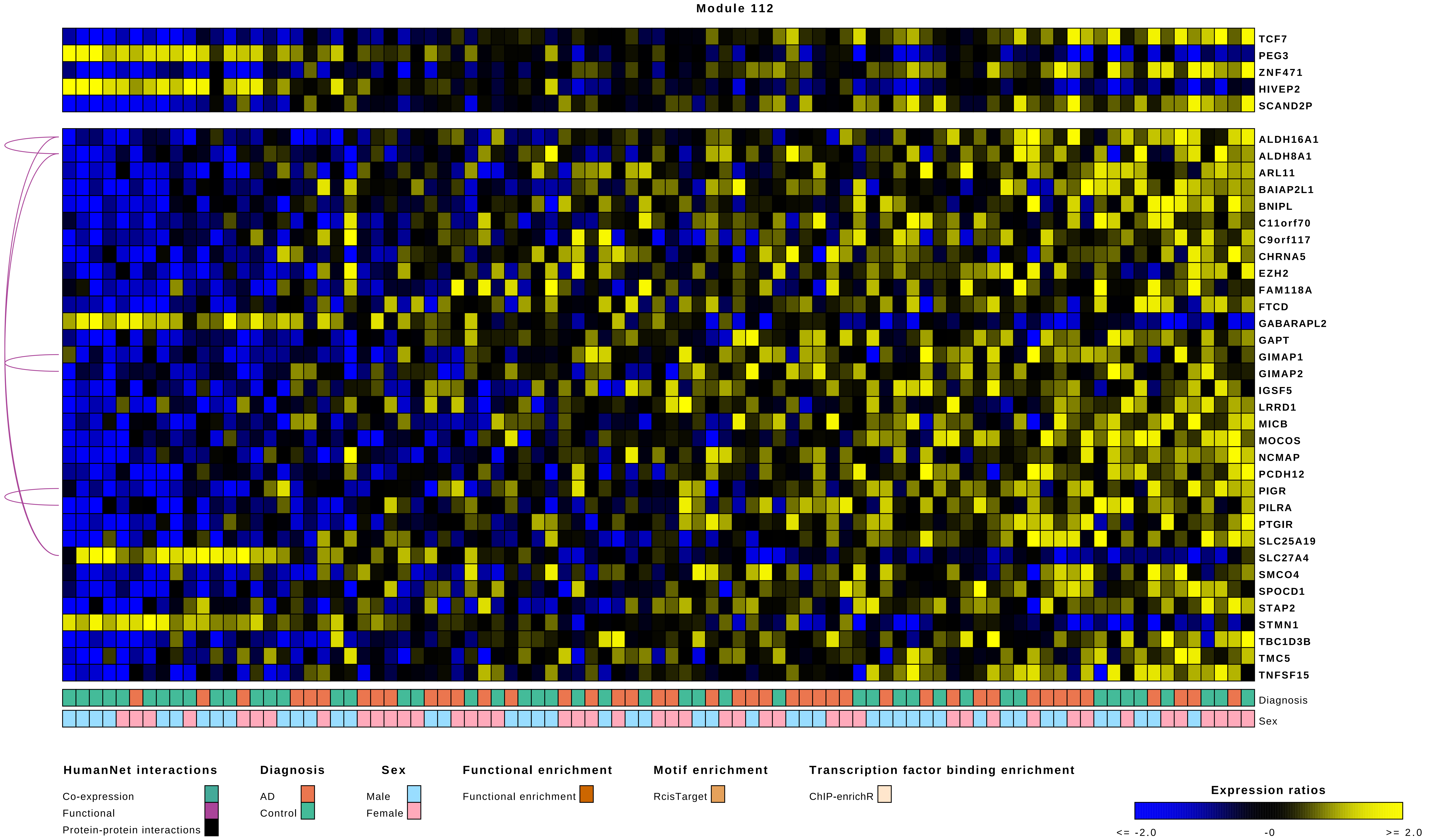

Module 112
TCF7
PEG3
ZNF471
HIVEP2
SCAND2P
ALDH16A1
ALDH8A1
ARL11
BAIAP2L1
BNIPL
C11orf70
C9orf117
CHRNA5
EZH2
FAM118A
FTCD
GABARAPL2
GAPT
GIMAP1
GIMAP2
IGSF5
LRRD1
MICB
MOCOS
NCMAP
PCDH12
PIGR
PILRA
PTGIR
SLC25A19
SLC27A4
SMCO4
SPOCD1
STAP2
STMN1
TBC1D3B
TMC5
TNFSF15
Diagnosis
Sex
HumanNet interactions
Co-expression
Functional
Protein-protein interactions
Diagnosis
AD
Control
Sex
Male
Female
Functional enrichment
Functional enrichment
Motif enrichment
RcisTarget
Transcription factor binding enrichment
ChIP-enrichR
Expression ratios
<= -2.0
-0
>= 2.0

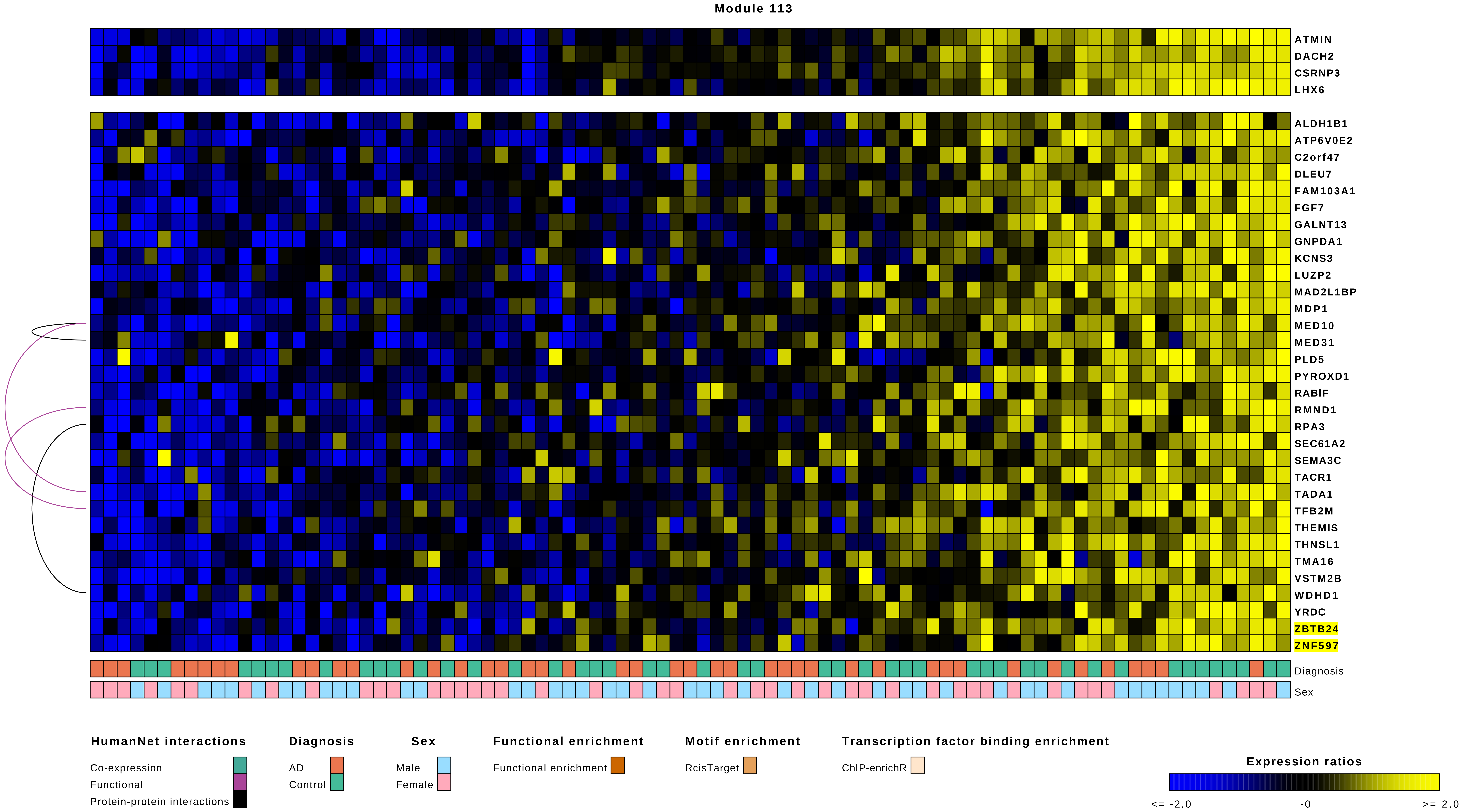
Module 113
ATMIN
DACH2
CSRNP3
LHX6
ALDH1B1
ATP6V0E2
C2orf47
DLEU7
FAM103A1
FGF7
GALNT13
GNPDA1
KCNS3
LUZP2
MAD2L1BP
MDP1
MED10
MED31
PLD5
PYROXD1
RABIF
RMND1
RPA3
SEC61A2
SEMA3C
TACR1
TADA1
TFB2M
THEMIS
THNSL1
TMA16
VSTM2B
WDHD1
YRDC
ZBTB24
ZNF597
Diagnosis
Sex
HumanNet interactions
Co-expression
Functional
Protein-protein interactions
Diagnosis
AD
Control
Sex
Male
Female
Functional enrichment
Functional enrichment
Motif enrichment
RcisTarget
Transcription factor binding enrichment
ChIP-enrichR
Expression ratios
<= -2.0
-0
>= 2.0

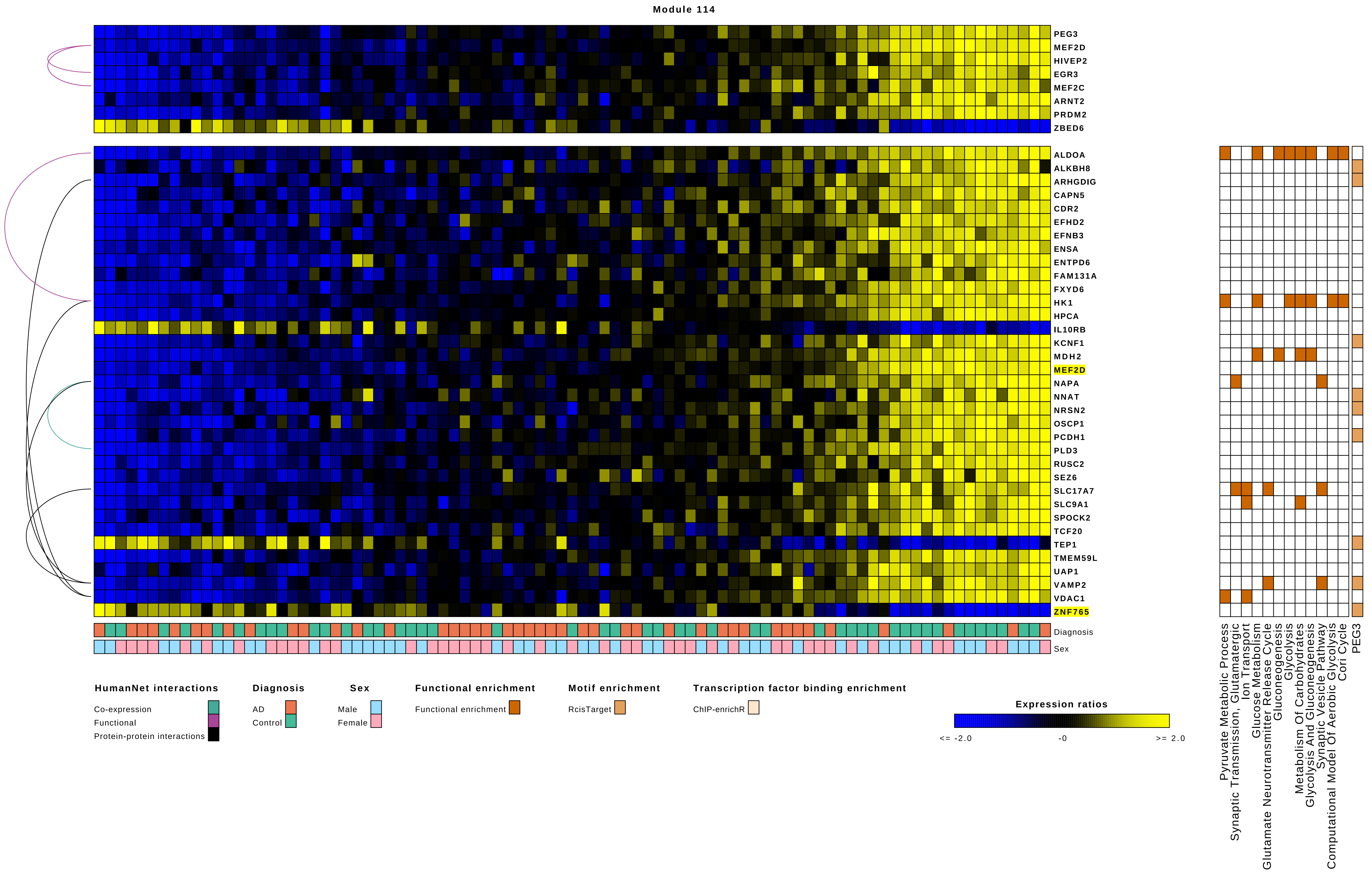

Module 114
PEG3
MEF2D
HIVEP2
EGR3
MEF2C
ARNT2
PRDM2
ZBED6
ALDOA
ALKBH8
ARHGDIG
CAPN5
CDR2
EFHD2
EFNB3
ENSA
ENTPD6
FAM131A
FXYD6
HK1
HPCA
IL10RB
KCNF1
MDH2
MEF2D
NAPA
NNAT
NRSN2
OSCP1
PCDH1
PLD3
RUSC2
SEZ6
SLC17A7
SLC9A1
SPOCK2
TCF20
TEP1
TMEM59L
UAP1
VAMP2
VDAC1
ZNF765
Diagnosis
Sex
Pyruvate Metabolic Process
Synaptic Transmission, Glutamatergic
Ion Transport
Glucose Metabolism
Glutamate Neurotransmitter Release Cycle
Gluconeogenesis
Glycolysis
Metabolism Of Carbohydrates
Glycolysis And Gluconeogenesis
Synaptic Vesicle Pathway
Computational Model Of Aerobic Glycolysis
Cori Cycle
PEG3
HumanNet interactions
Co-expression
Functional
Protein-protein interactions
Diagnosis
AD
Control
Sex
Male
Female
Functional enrichment
Functional enrichment
Motif enrichment
RcisTarget
Transcription factor binding enrichment
ChIP-enrichR
Expression ratios
<= -2.0
-0
>= 2.0

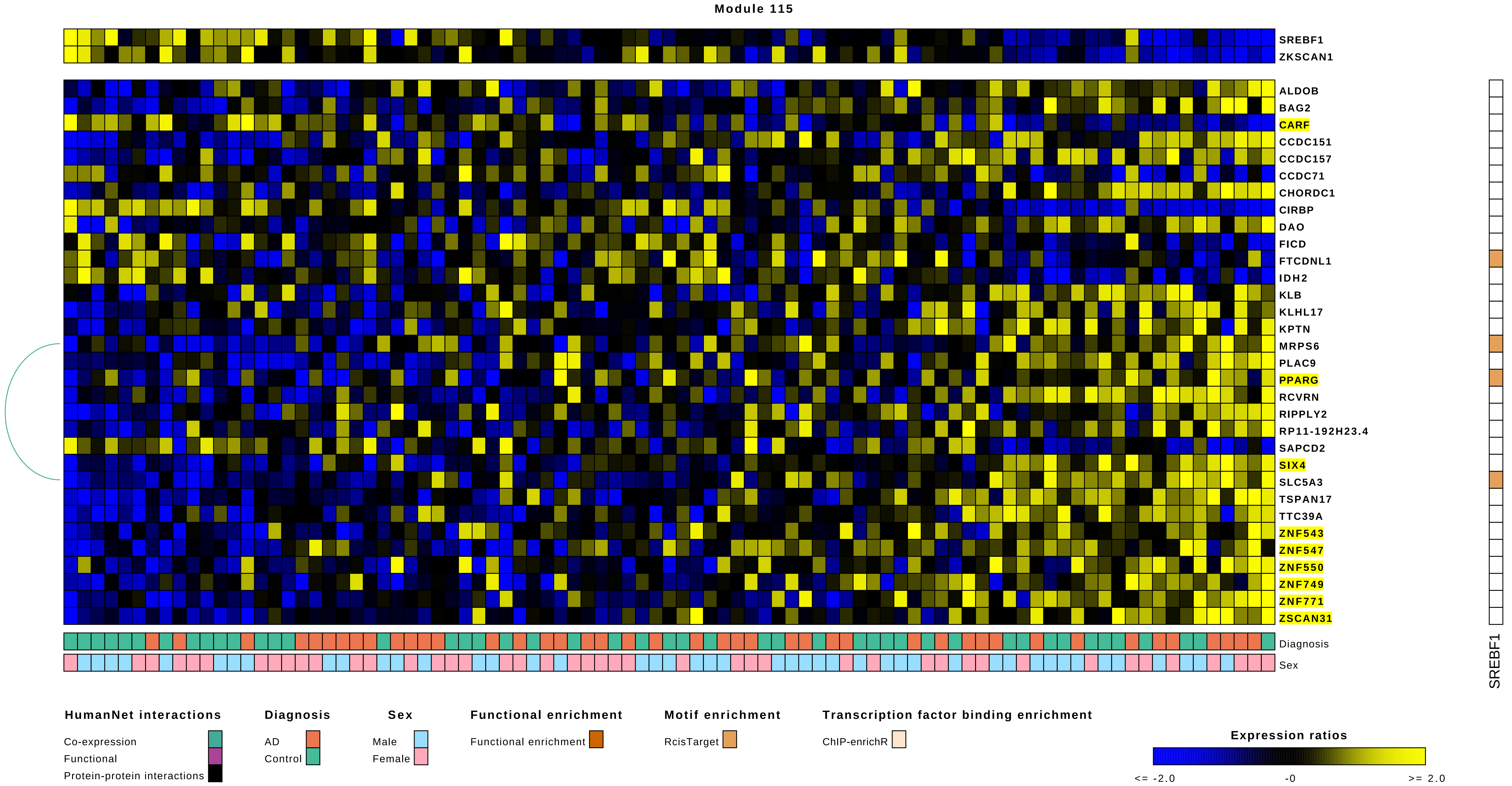
Module 115
SREBF1
ZKSCAN1
ALDOB
BAG2
CARF
CCDC151
CCDC157
CCDC71
CHORDC1
CIRBP
DAO
FICD
FTCDNL1
IDH2
KLB
KLHL17
KPTN
MRPS6
PLAC9
PPARG
RCVRN
RIPPLY2
RP11-192H23.4
SAPCD2
SIX4
SLC5A3
TSPAN17
TTC39A
ZNF543
ZNF547
ZNF550
ZNF749
ZNF771
ZSCAN31
Diagnosis
Sex
SREBF1
HumanNet interactions
Co-expression
Functional
Protein-protein interactions
Diagnosis
AD
Control
Sex
Male
Female
Functional enrichment
Functional enrichment
Motif enrichment
RcisTarget
Transcription factor binding enrichment
ChIP-enrichR
Expression ratios
<= -2.0
-0
>= 2.0

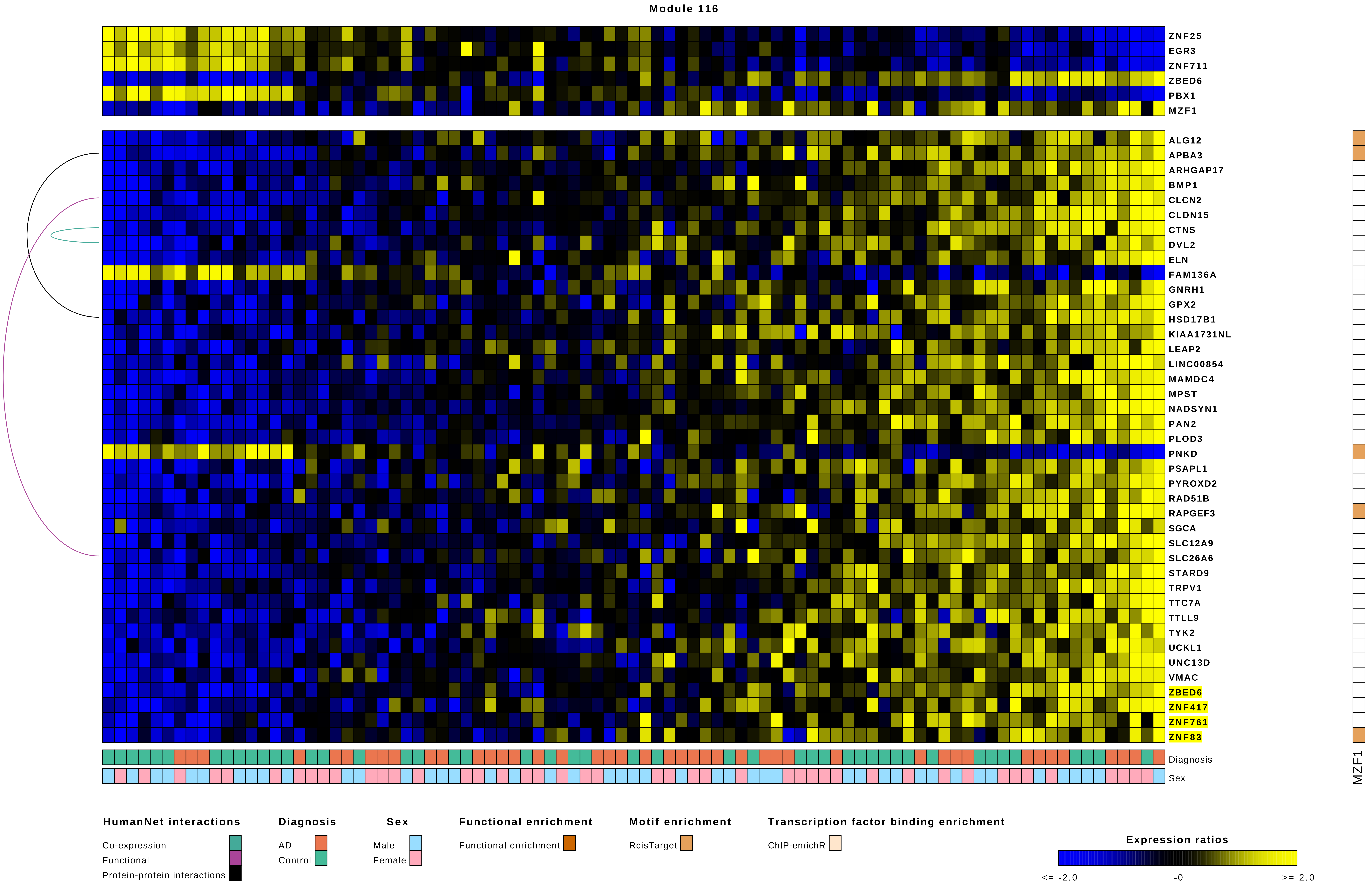

Module 116
ZNF25
EGR3
ZNF711
ZBED6
PBX1
MZF1
ALG12
APBA3
ARHGAP17
BMP1
CLCN2
CLDN15
CTNS
DVL2
ELN
FAM136A
GNRH1
GPX2
HSD17B1
KIAA1731NL
LEAP2
LINC00854
MAMDC4
MPST
NADSYN1
PAN2
PLOD3
PNKD
PSAPL1
PYROXD2
RAD51B
RAPGEF3
SGCA
SLC12A9
SLC26A6
STARD9
TRPV1
TTC7A
TTLL9
TYK2
UCKL1
UNC13D
VMAC
ZBED6
ZNF417
ZNF761
ZNF83
Diagnosis
Sex
MZF1
HumanNet interactions
Co-expression
Functional
Protein-protein interactions
Diagnosis
AD
Control
Sex
Male
Female
Functional enrichment
Functional enrichment
Motif enrichment
RcisTarget
Transcription factor binding enrichment
ChIP-enrichR
Expression ratios
<= -2.0
-0
>= 2.0

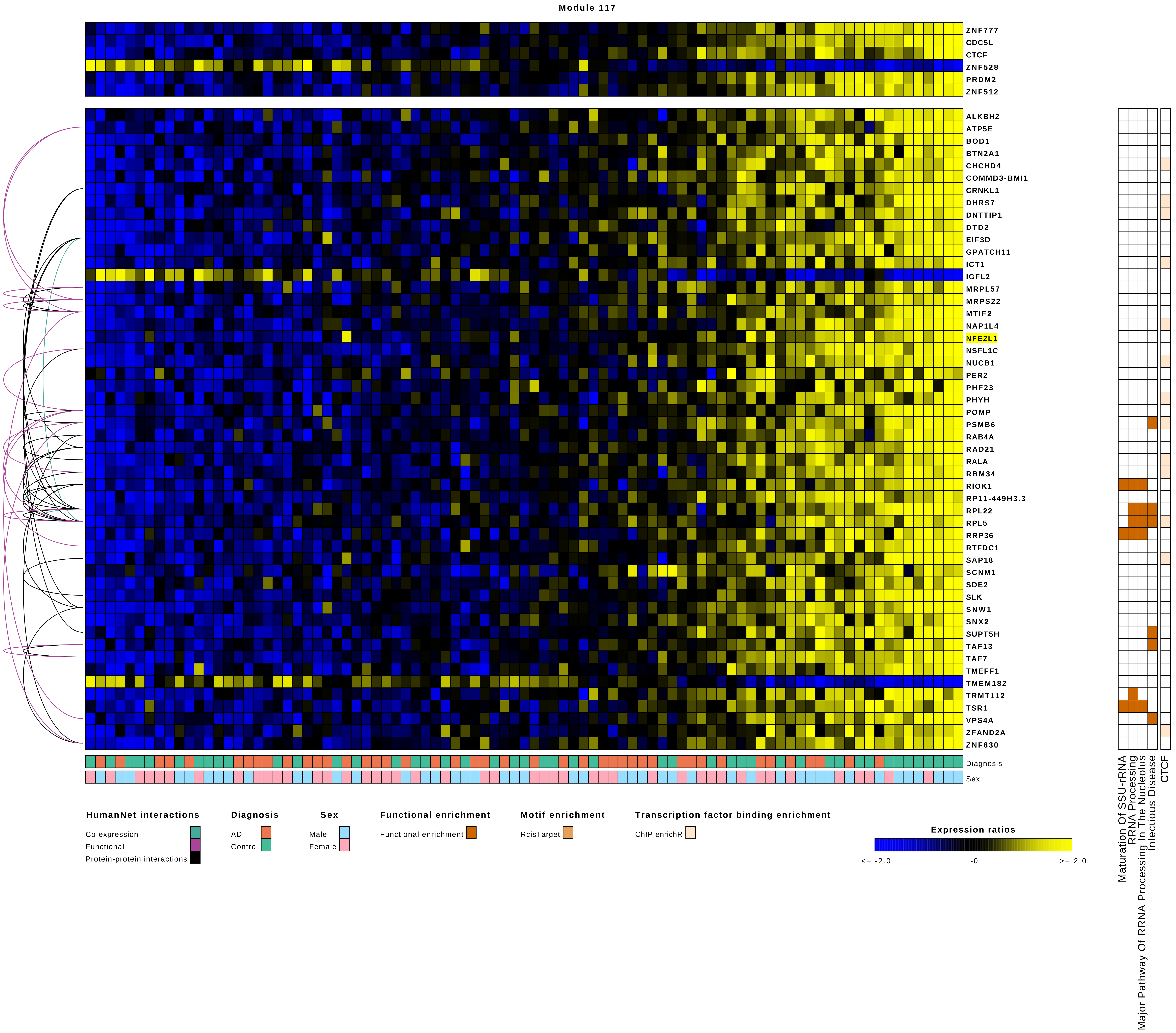
Module 117
ZNF777
CDC5L
CTCF
ZNF528
PRDM2
ZNF512
ALKBH2
ATP5E
BOD1
BTN2A1
CHCHD4
COMMD3-BMI1
CRNKL1
DHRS7
DNTTIP1
DTD2
EIF3D
GPATCH11
ICT1
IGFL2
MRPL57
MRPS22
MTIF2
NAP1L4
NFE2L1
NSFL1C
NUCB1
PER2
PHF23
PHYH
POMP
PSMB6
RAB4A
RAD21
RALA
RBM34
RIOK1
RP11-449H3.3
RPL22
RPL5
RRP36
RTFDC1
SAP18
SCNM1
SDE2
SLK
SNW1
SNX2
SUPT5H
TAF13
TAF7
TMEFF1
TMEM182
TRMT112
TSR1
VPS4A
ZFAND2A
ZNF830
Diagnosis
Sex
Maturation Of SSU-rRNA
RRNA Processing
Major Pathway Of RRNA Processing In The Nucleolus
Infectious Disease
CTCF
HumanNet interactions
Co-expression
Functional
Protein-protein interactions
Diagnosis
AD
Control
Sex
Male
Female
Functional enrichment
Functional enrichment
Motif enrichment
RcisTarget
Transcription factor binding enrichment
ChIP-enrichR
Expression ratios
<= -2.0
-0
>= 2.0

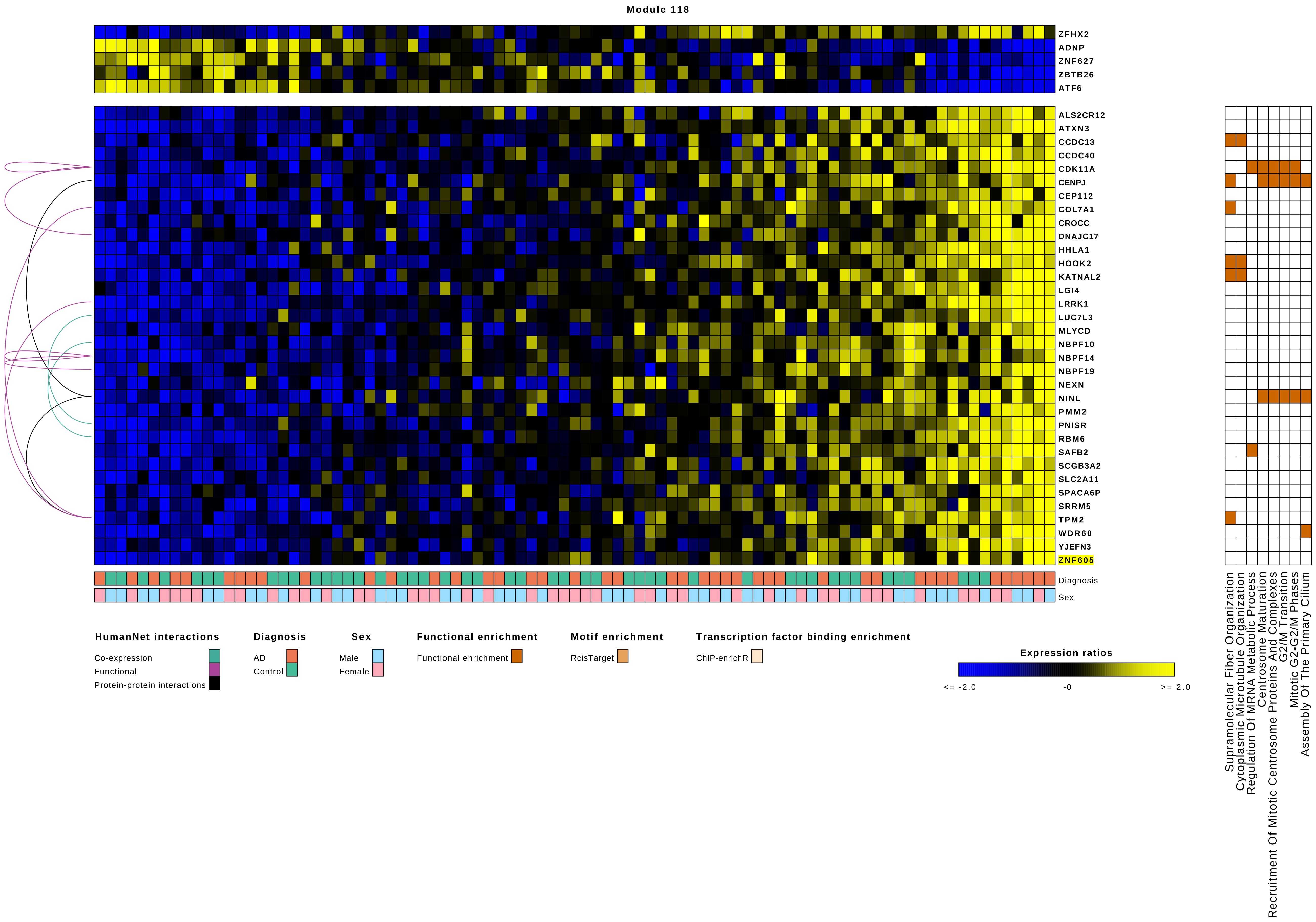
Module 118
ZFHX2
ADNP
ZNF627
ZBTB26
ATF6
ALS2CR12
ATXN3
CCDC13
CCDC40
CDK11A
CENPJ
CEP112
COL7A1
CROCC
DNAJC17
HHLA1
HOOK2
KATNAL2
LGI4
LRRK1
LUC7L3
MLYCD
NBPF10
NBPF14
NBPF19
NEXN
NINL
PMM2
PNISR
RBM6
SAFB2
SCGB3A2
SLC2A11
SPACA6P
SRRM5
TPM2
WDR60
YJEFN3
ZNF605
Diagnosis
Sex
HumanNet interactions
Co-expression
Functional
Protein-protein interactions
Diagnosis
AD
Control
Sex
Male
Female
Functional enrichment
Functional enrichment
Motif enrichment
RcisTarget
Transcription factor binding enrichment
ChIP-enrichR
Expression ratios
<= -2.0
-0
>= 2.0
Supramolecular Fiber Organization
Cytoplasmic Microtubule Organization
Regulation Of MRNA Metabolic Process
Centrosome Maturation
Recruitment Of Mitotic Centrosome Proteins And Complexes
G2/M Transition
Mitotic G2-G2/M Phases
Assembly Of The Primary Cilium

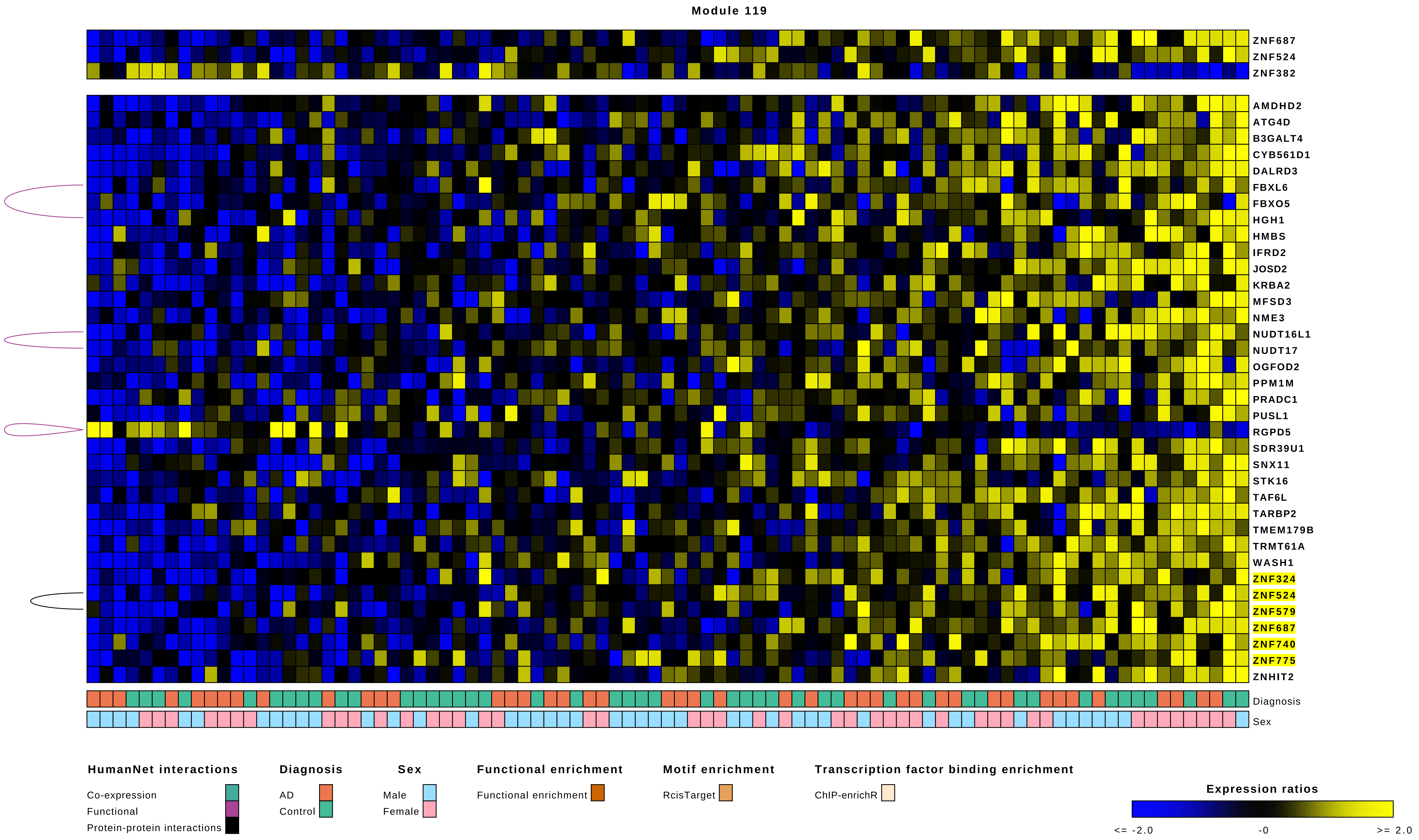
Module 119
ZNF687
ZNF524
ZNF382
AMDHD2
ATG4D
B3GALT4
CYB561D1
DALRD3
FBXL6
FBXO5
HGH1
HMBS
IFRD2
JOSD2
KRBA2
MFSD3
NME3
NUDT16L1
NUDT17
OGFOD2
PPM1M
PRADC1
PUSL1
RGPD5
SDR39U1
SNX11
STK16
TAF6L
TARBP2
TMEM179B
TRMT61A
WASH1
ZNF324
ZNF524
ZNF579
ZNF687
ZNF740
ZNF775
ZNHIT2
Diagnosis
Sex
HumanNet interactions
Co-expression
Functional
Protein-protein interactions
Diagnosis
AD
Control
Sex
Male
Female
Functional enrichment
Functional enrichment
Motif enrichment
RcisTarget
Transcription factor binding enrichment
ChIP-enrichR
Expression ratios
<= -2.0
-0
>= 2.0

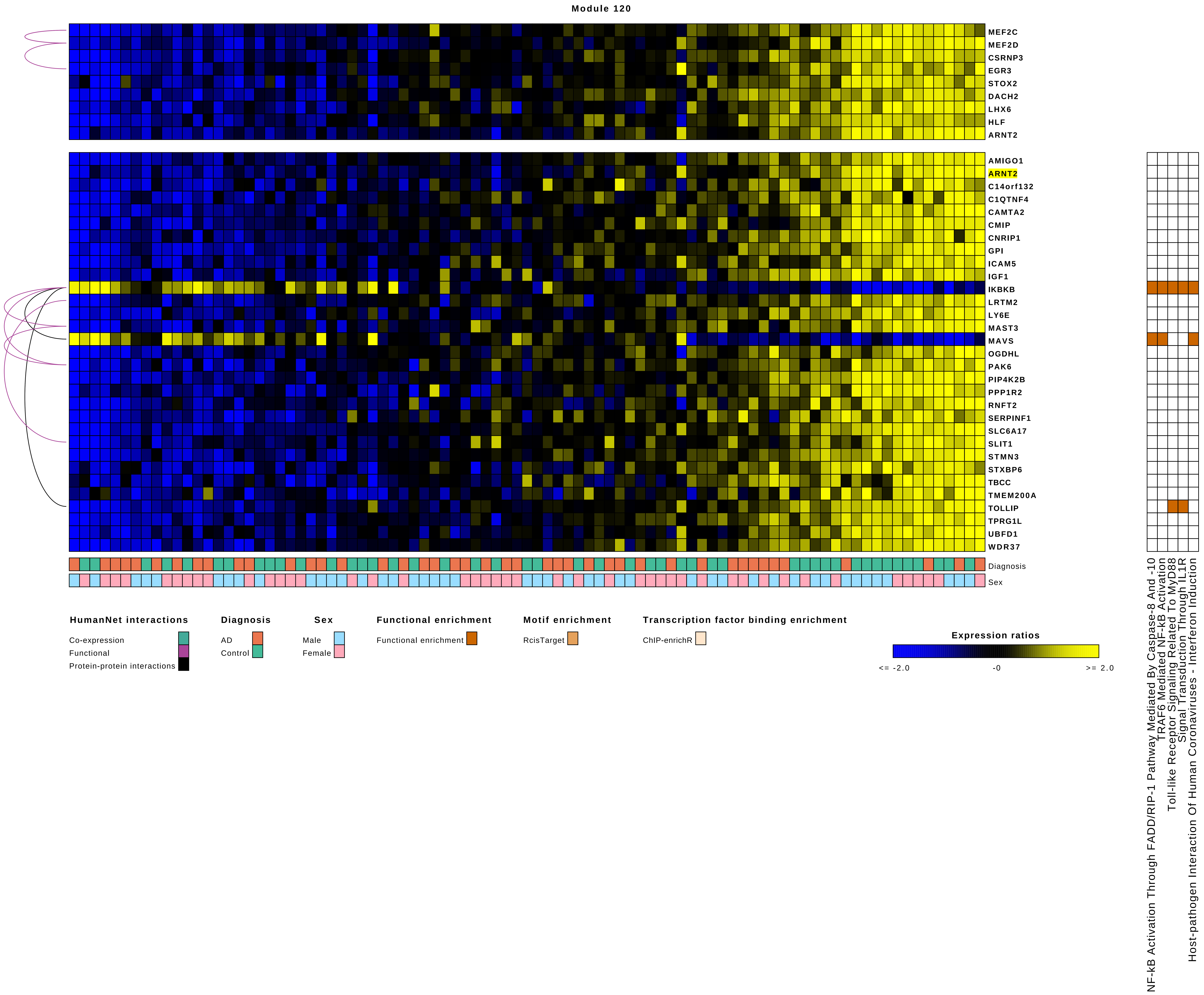
Module 120
MEF2C
MEF2D
CSRNP3
EGR3
STOX2
DACH2
LHX6
HLF
ARNT2
AMIGO1
ARNT2
C14orf132
C1QTNF4
CAMTA2
CMIP
CNRIP1
GPI
ICAM5
IGF1
IKBKB
LRTM2
LY6E
MAST3
MAVS
OGDHL
PAK6
PIP4K2B
PPP1R2
RNFT2
SERPINF1
SLC6A17
SLIT1
STMN3
STXBP6
TBCC
TMEM200A
TOLLIP
TPRG1L
UBFD1
WDR37
Diagnosis
Sex
NF-kB Activation Through FADD/RIP-1 Pathway Mediated By Caspase-8 And -10
TRAF6 Mediated NF-kB Activation
Toll-like Receptor Signaling Related To MyD88
Signal Transduction Through IL1R
Host-pathogen Interaction Of Human Coronaviruses - Interferon Induction
HumanNet interactions
Co-expression
Functional
Protein-protein interactions
Diagnosis
AD
Control
Sex
Male
Female
Functional enrichment
Functional enrichment
Motif enrichment
RcisTarget
Transcription factor binding enrichment
ChIP-enrichR
Expression ratios
<= -2.0
-0
>= 2.0

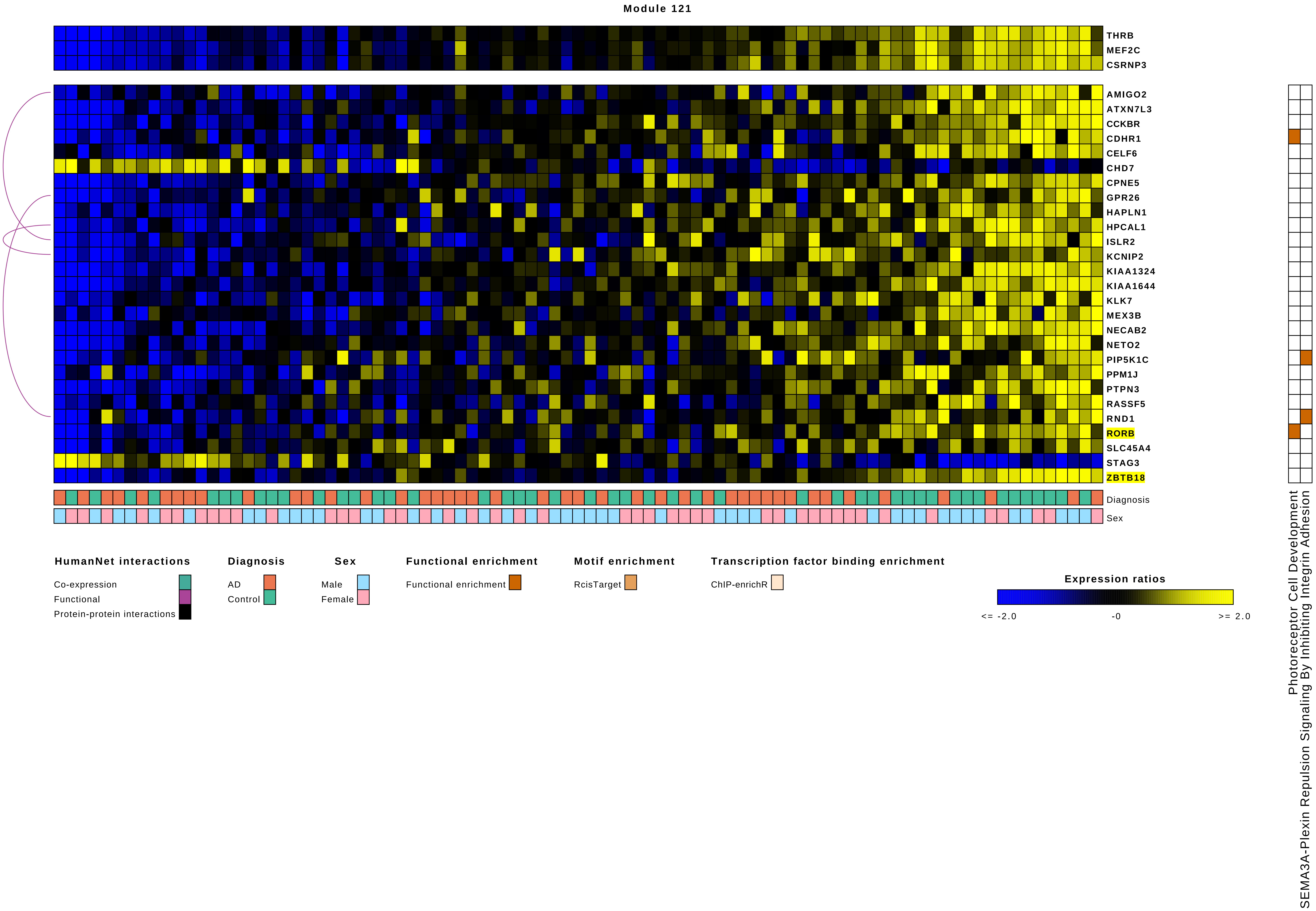
Module 121
THRB
MEF2C
CSRNP3
AMIGO2
ATXN7L3
CCKBR
CDHR1
CELF6
CHD7
CPNE5
GPR26
HAPLN1
HPCAL1
ISLR2
KCNIP2
KIAA1324
KIAA1644
KLK7
MEX3B
NECAB2
NETO2
PIP5K1C
PPM1J
PTPN3
RASSF5
RND1
RORB
SLC45A4
STAG3
ZBTB18
Diagnosis
Sex
Photoreceptor Cell Development
SEMA3A-Plexin Repulsion Signaling By Inhibiting Integrin Adhesion
HumanNet interactions
Co-expression
Functional
Protein-protein interactions
Diagnosis
AD
Control
Sex
Male
Female
Functional enrichment
Functional enrichment
Motif enrichment
RcisTarget
Transcription factor binding enrichment
ChIP-enrichR
Expression ratios
<= -2.0
-0
>= 2.0

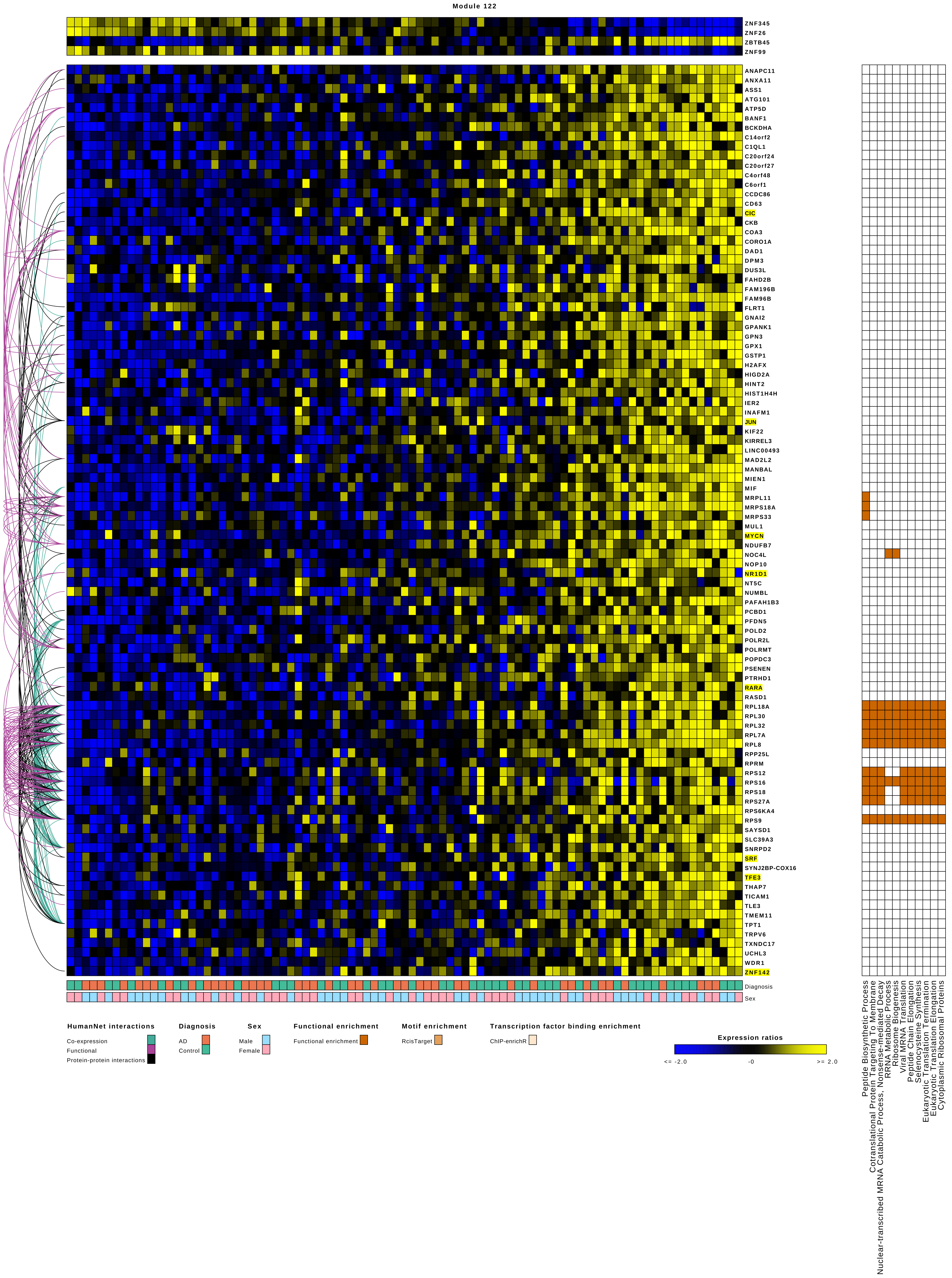
Module 122
ZNF345
ZNF26
ZBTB45
ZNF99
ANAPC11
ANXA11
ASS1
ATG101
ATP5D
BANF1
BCKDHA
C14orf2
C1QL1
C20orf24
C20orf27
C4orf48
C6orf1
CCDC86
CD63
CIC
CKB
COA3
CORO1A
DAD1
DPM3
DUS3L
FAHD2B
FAM196B
FAM96B
FLRT1
GNAI2
GPANK1
GPN3
GPX1
GSTP1
H2AFX
HIGD2A
HINT2
HIST1H4H
IER2
INAFM1
JUN
KIF22
KIRREL3
LINC00493
MAD2L2
MANBAL
MIEN1
MIF
MRPL11
MRPS18A
MRPS33
MUL1
MYCN
NDUFB7
NOC4L
NOP10
NR1D1
NT5C
NUMBL
PAFAH1B3
PCBD1
PFDN5
POLD2
POLR2L
POLRMT
POPDC3
PSENEN
PTRHD1
RARA
RASD1
RPL18A
RPL30
RPL32
RPL7A
RPL8
RPP25L
RPRM
RPS12
RPS16
RPS18
RPS27A
RPS6KA4
RPS9
SAYSD1
SLC39A3
SNRPD2
SRF
SYNJ2BP-COX16
TFE3
THAP7
TICAM1
TLE3
TMEM11
TPT1
TRPV6
TXNDC17
UCHL3
WDR1
ZNF142
Diagnosis
Sex
Peptide Biosynthetic Process
Cotranslational Protein Targeting To Membrane
Nuclear-transcribed MRNA Catabolic Process, Nonsense-mediated Decay
RRNA Metabolic Process
Ribosome Biogenesis
Viral MRNA Translation
Peptide Chain Elongation
Selenocysteine Synthesis
Eukaryotic Translation Termination
Eukaryotic Translation Elongation
Cytoplasmic Ribosomal Proteins
HumanNet interactions
Co-expression
Functional
Protein-protein interactions
Diagnosis
AD
Control
Sex
Male
Female
Functional enrichment
Functional enrichment
Motif enrichment
RcisTarget
Transcription factor binding enrichment
ChIP-enrichR
Expression ratios
<= -2.0
-0
>= 2.0

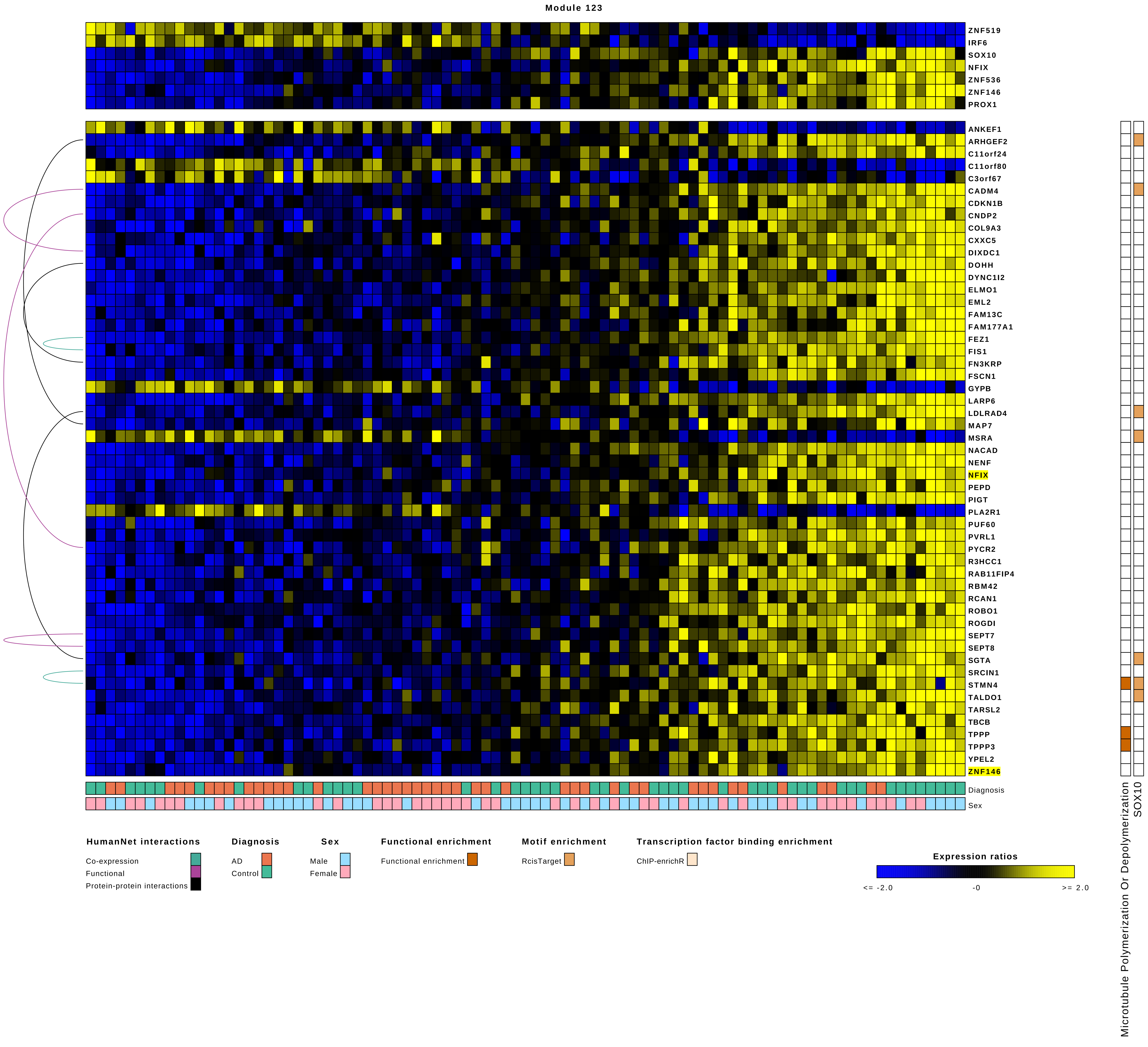

Module 123
ZNF519
IRF6
SOX10
NFIX
ZNF536
ZNF146
PROX1
ANKEF1
ARHGEF2
C11orf24
C11orf80
C3orf67
CADM4
CDKN1B
CNDP2
COL9A3
CXXC5
DIXDC1
DOHH
DYNC1I2
ELMO1
EML2
FAM13C
FAM177A1
FEZ1
FIS1
FN3KRP
FSCN1
GYPB
LARP6
LDLRAD4
MAP7
MSRA
NACAD
NENF
NFIX
PEPD
PIGT
PLA2R1
PUF60
PVRL1
PYCR2
R3HCC1
RAB11FIP4
RBM42
RCAN1
ROBO1
ROGDI
SEPT7
SEPT8
SGTA
SRCIN1
STMN4
TALDO1
TARSL2
TBCB
TPPP
TPPP3
YPEL2
ZNF146
Diagnosis
Sex
Microtubule Polymerization Or Depolymerization
SOX10
HumanNet interactions
Co-expression
Functional
Protein-protein interactions
Diagnosis
AD
Control
Sex
Male
Female
Functional enrichment
Functional enrichment
Motif enrichment
RcisTarget
Transcription factor binding enrichment
ChIP-enrichR
Expression ratios
<= -2.0
-0
>= 2.0

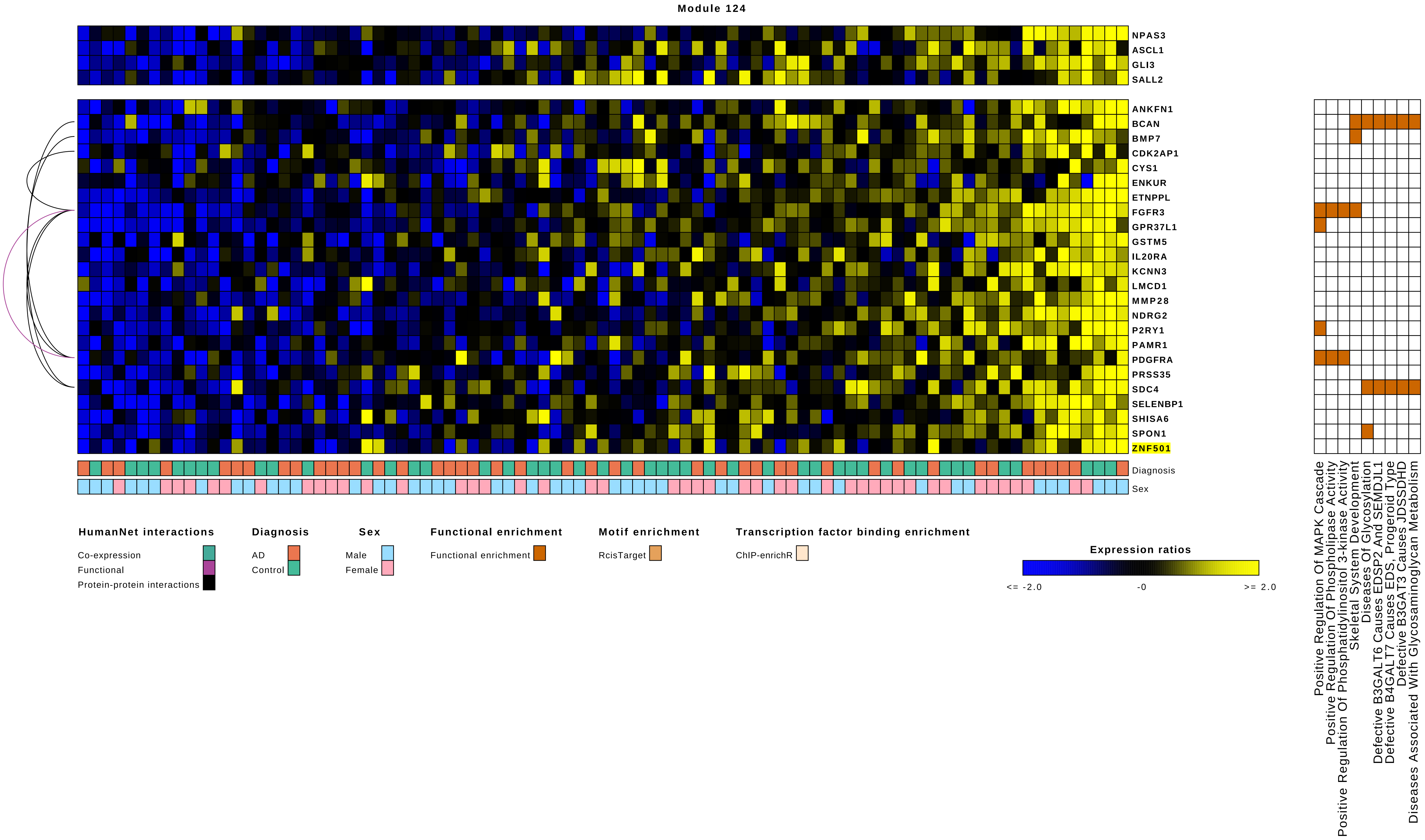

Module 124
NPAS3
ASCL1
GLI3
SALL2
ANKFN1
BCAN
BMP7
CDK2AP1
CYS1
ENKUR
ETNPPL
FGFR3
GPR37L1
GSTM5
IL20RA
KCNN3
LMCD1
MMP28
NDRG2
P2RY1
PAMR1
PDGFRA
PRSS35
SDC4
SELENBP1
SHISA6
SPON1
ZNF501
Diagnosis
Sex
Positive Regulation Of MAPK Cascade
Positive Regulation Of Phospholipase Activity
Positive Regulation Of Phosphatidylinositol 3-kinase Activity
Skeletal System Development
Diseases Of Glycosylation
Defective B3GALT6 Causes EDSP2 And SEMDJL1
Defective B4GALT7 Causes EDS, Progeroid Type
Defective B3GAT3 Causes JDSSDHD
Diseases Associated With Glycosaminoglycan Metabolism
HumanNet interactions
Co-expression
Functional
Protein-protein interactions
Diagnosis
AD
Control
Sex
Male
Female
Functional enrichment
Functional enrichment
Motif enrichment
RcisTarget
Transcription factor binding enrichment
ChIP-enrichR
Expression ratios
<= -2.0
-0
>= 2.0

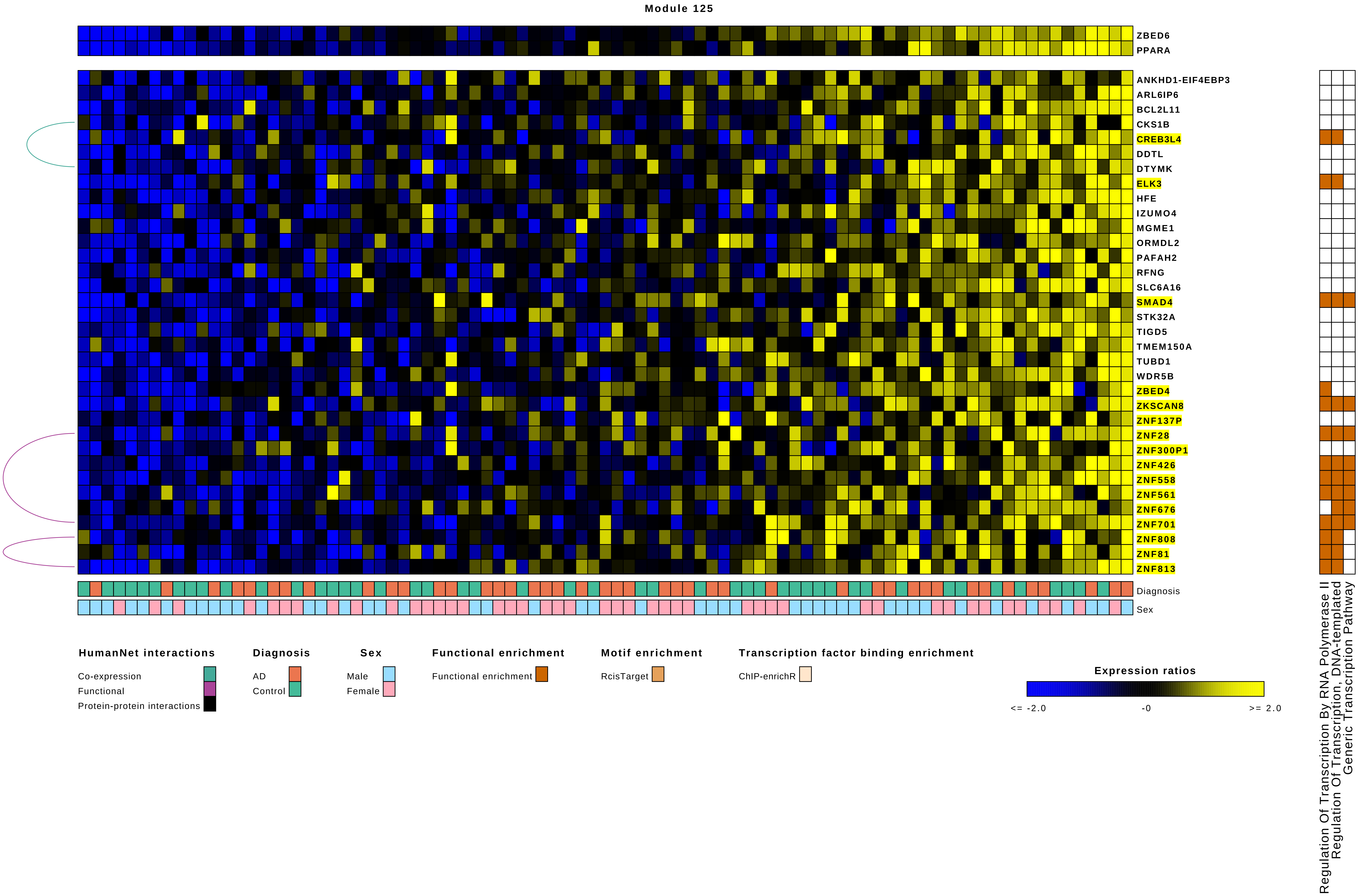

Module 125
ZBED6
PPARA
ANKHD1-EIF4EBP3
ARL6IP6
BCL2L11
CKS1B
CREB3L4
DDTL
DTYMK
ELK3
HFE
IZUMO4
MGME1
ORMDL2
PAFAH2
RFNG
SLC6A16
SMAD4
STK32A
TIGD5
TMEM150A
TUBD1
WDR5B
ZBED4
ZKSCAN8
ZNF137P
ZNF28
ZNF300P1
ZNF426
ZNF558
ZNF561
ZNF676
ZNF701
ZNF808
ZNF81
ZNF813
Diagnosis
Sex
Regulation Of Transcription By RNA Polymerase II
Regulation Of Transcription, DNA-templated
Generic Transcription Pathway
HumanNet interactions
Co-expression
Functional
Protein-protein interactions
Diagnosis
AD
Control
Sex
Male
Female
Functional enrichment
Functional enrichment
Motif enrichment
RcisTarget
Transcription factor binding enrichment
ChIP-enrichR
Expression ratios
<= -2.0
-0
>= 2.0

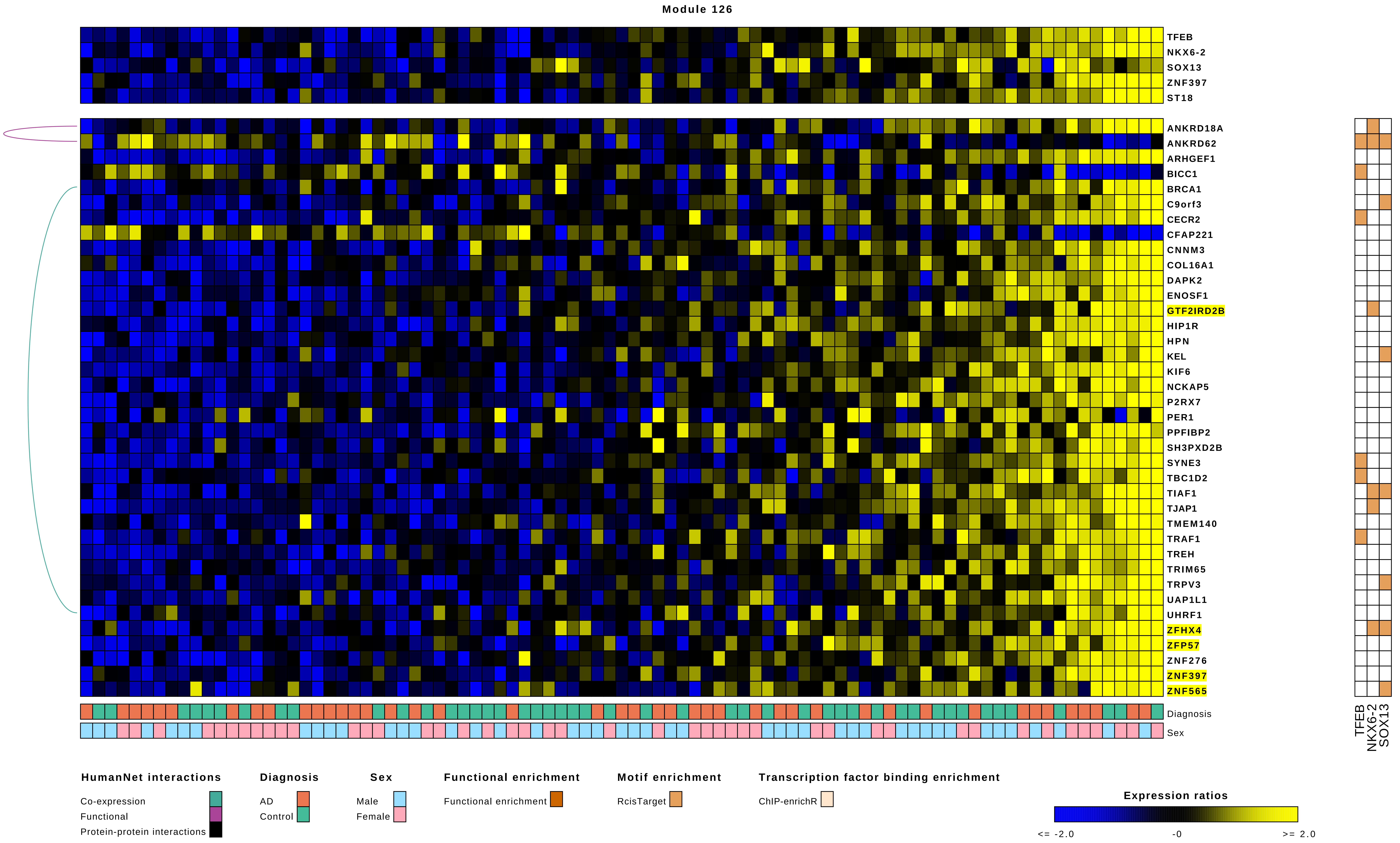

Module 126
TFEB
NKX6-2
SOX13
ZNF397
ST18
ANKRD18A
ANKRD62
ARHGEF1
BICC1
BRCA1
C9orf3
CECR2
CFAP221
CNNM3
COL16A1
DAPK2
ENOSF1
GTF2IRD2B
HIP1R
HPN
KEL
KIF6
NCKAP5
P2RX7
PER1
PPFIBP2
SH3PXD2B
SYNE3
TBC1D2
TIAF1
TJAP1
TMEM140
TRAF1
TREH
TRIM65
TRPV3
UAP1L1
UHRF1
ZFHX4
ZFP57
ZNF276
ZNF397
ZNF565
Diagnosis
Sex
TFEB
NKX6-2
SOX13
HumanNet interactions
Co-expression
Functional
Protein-protein interactions
Diagnosis
AD
Control
Sex
Male
Female
Functional enrichment
Functional enrichment
Motif enrichment
RcisTarget
Transcription factor binding enrichment
ChIP-enrichR
Expression ratios
<= -2.0
-0
>= 2.0

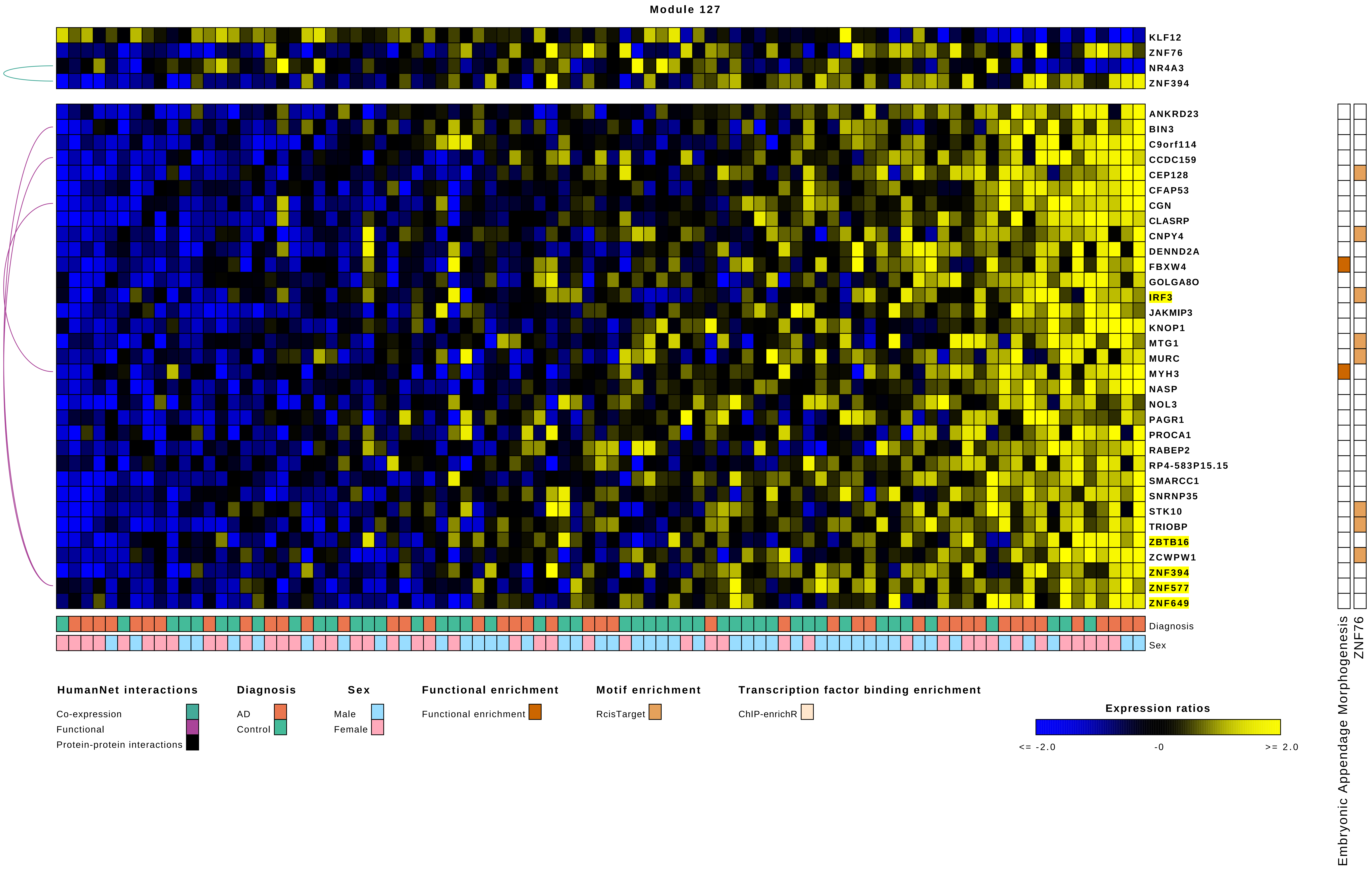

Module 127
KLF12
ZNF76
NR4A3
ZNF394
ANKRD23
BIN3
C9orf114
CCDC159
CEP128
CFAP53
CGN
CLASRP
CNPY4
DENND2A
FBXW4
GOLGA8O
IRF3
JAKMIP3
KNOP1
MTG1
MURC
MYH3
NASP
NOL3
PAGR1
PROCA1
RABEP2
RP4-583P15.15
SMARCC1
SNRNP35
STK10
TRIOBP
ZBTB16
ZCWPW1
ZNF394
ZNF577
ZNF649
Diagnosis
Sex
Embryonic Appendage Morphogenesis
ZNF76
HumanNet interactions
Co-expression
Functional
Protein-protein interactions
Diagnosis
AD
Control
Sex
Male
Female
Functional enrichment
Functional enrichment
Motif enrichment
RcisTarget
Transcription factor binding enrichment
ChIP-enrichR
Expression ratios
<= -2.0
-0
>= 2.0

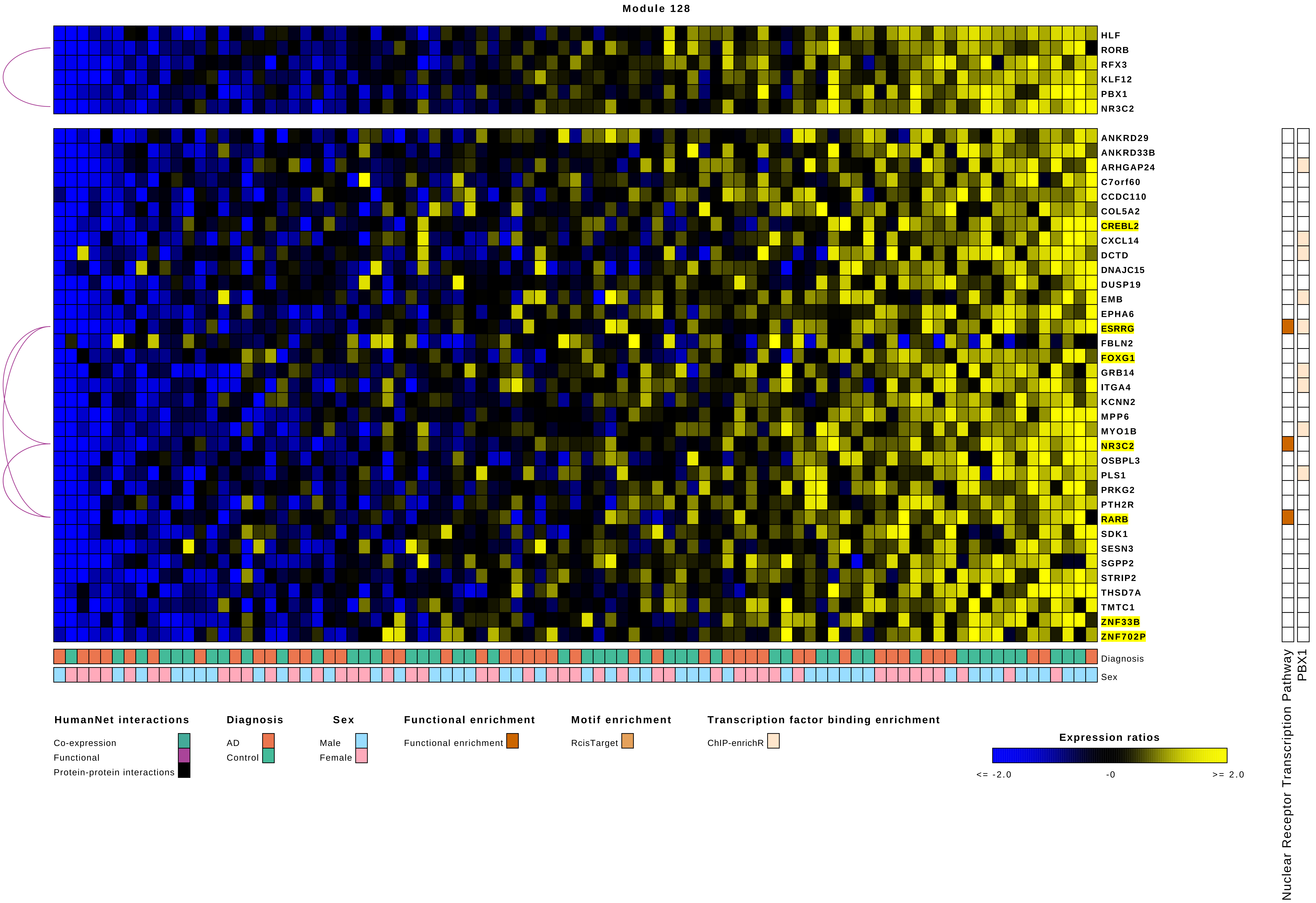

Module 128
HLF
RORB
RFX3
KLF12
PBX1
NR3C2
ANKRD29
ANKRD33B
ARHGAP24
C7orf60
CCDC110
COL5A2
CREBL2
CXCL14
DCTD
DNAJC15
DUSP19
EMB
EPHA6
ESRRG
FBLN2
FOXG1
GRB14
ITGA4
KCNN2
MPP6
MYO1B
NR3C2
OSBPL3
PLS1
PRKG2
PTH2R
RARB
SDK1
SESN3
SGPP2
STRIP2
THSD7A
TMTC1
ZNF33B
ZNF702P
Diagnosis
Sex
Nuclear Receptor Transcription Pathway
PBX1
HumanNet interactions
Co-expression
Functional
Protein-protein interactions
Diagnosis
AD
Control
Sex
Male
Female
Functional enrichment
Functional enrichment
Motif enrichment
RcisTarget
Transcription factor binding enrichment
ChIP-enrichR
Expression ratios
<= -2.0
-0
>= 2.0

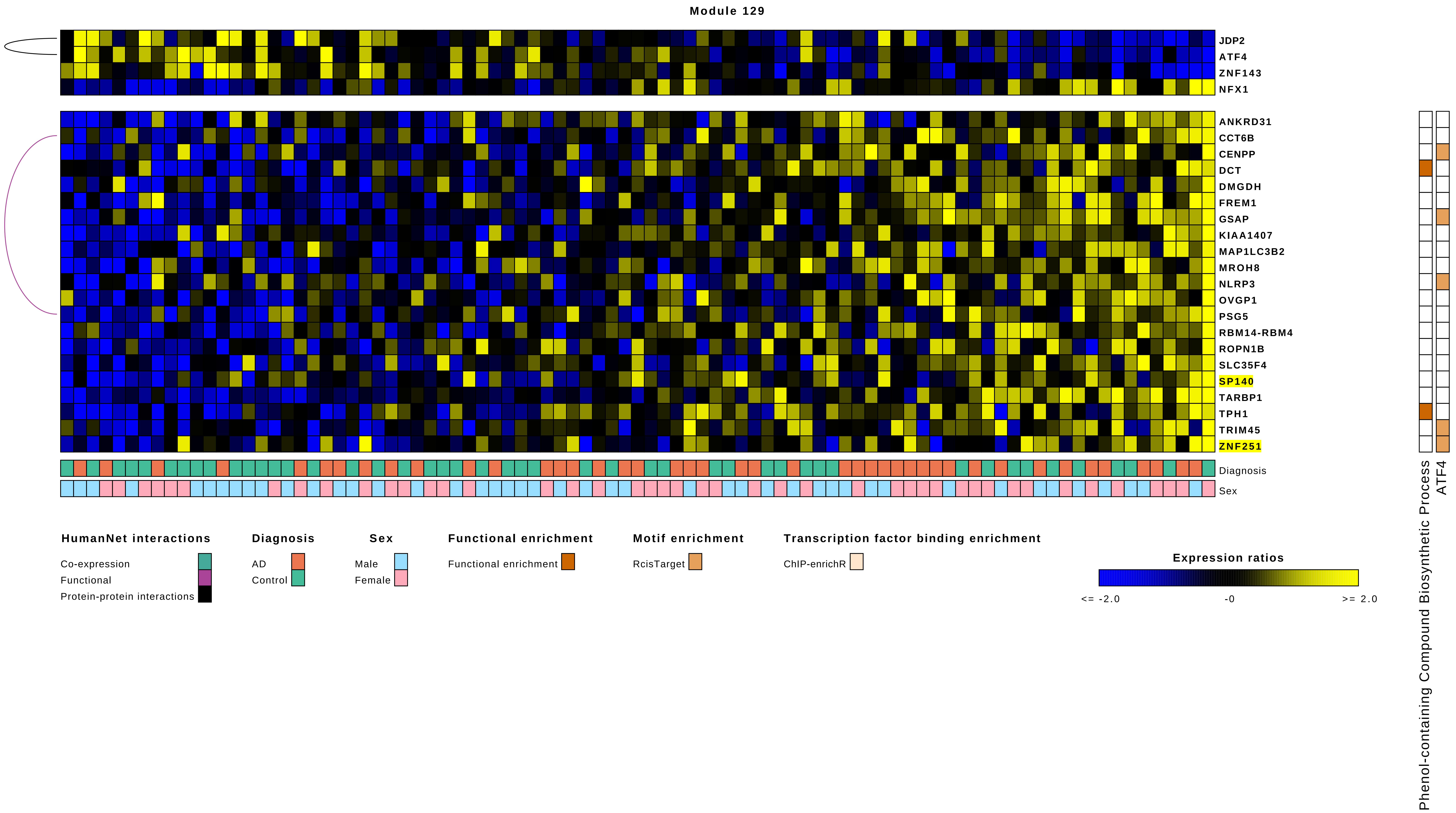
Module 129
JDP2
ATF4
ZNF143
NFX1
ANKRD31
CCT6B
CENPP
DCT
DMGDH
FREM1
GSAP
KIAA1407
MAP1LC3B2
MROH8
NLRP3
OVGP1
PSG5
RBM14-RBM4
ROPN1B
SLC35F4
SP140
TARBP1
TPH1
TRIM45
ZNF251
Diagnosis
Sex
Phenol-containing Compound Biosynthetic Process
ATF4
HumanNet interactions
Co-expression
Functional
Protein-protein interactions
Diagnosis
AD
Control
Sex
Male
Female
Functional enrichment
Functional enrichment
Motif enrichment
RcisTarget
Transcription factor binding enrichment
ChIP-enrichR
Expression ratios
<= -2.0
-0
>= 2.0

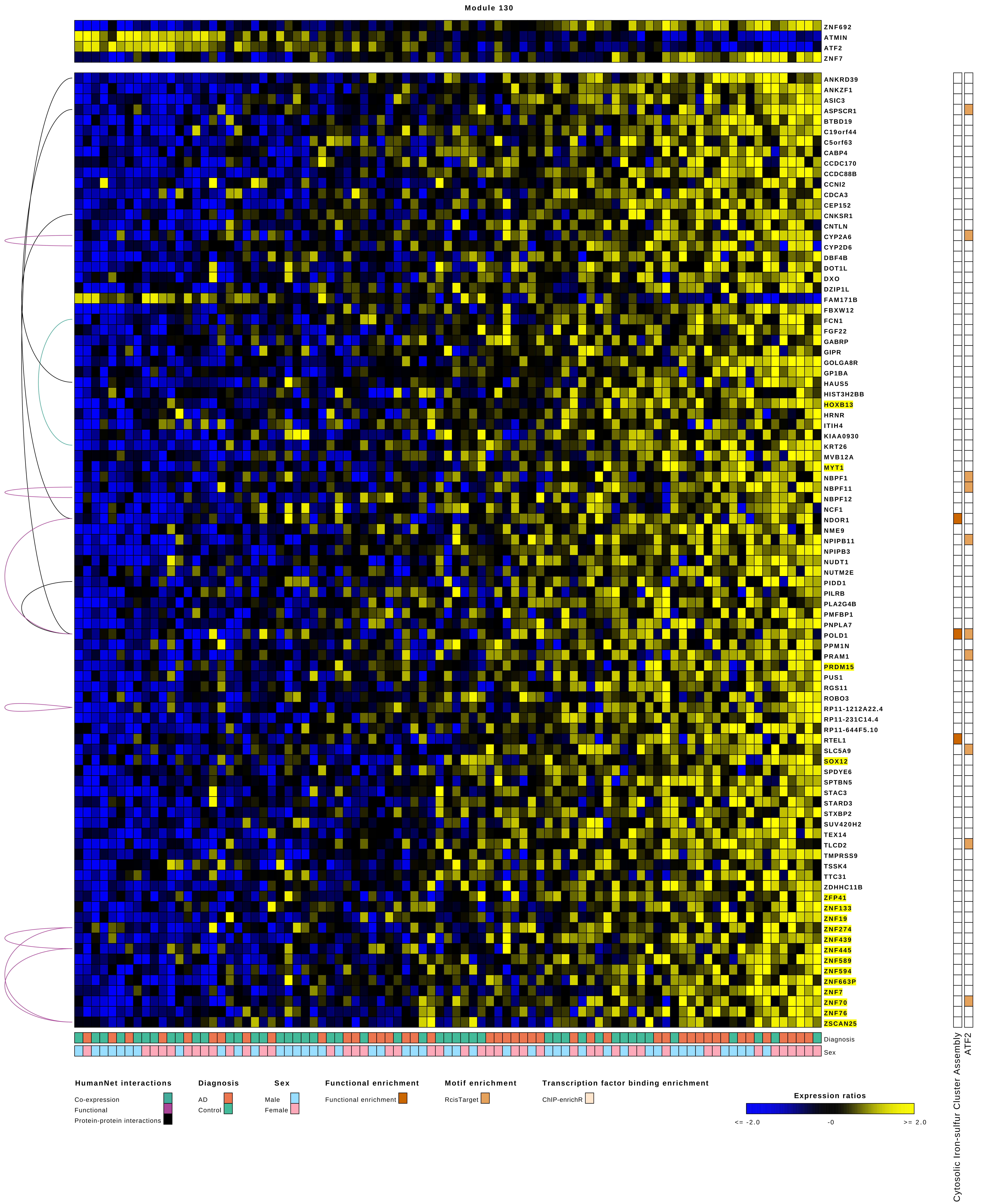

Module 130
ZNF692
ATMIN
ATF2
ZNF7
ANKRD39
ANKZF1
ASIC3
ASPSCR1
BTBD19
C19orf44
C5orf63
CABP4
CCDC170
CCDC88B
CCNI2
CDCA3
CEP152
CNKSR1
CNTLN
CYP2A6
CYP2D6
DBF4B
DOT1L
DXO
DZIP1L
FAM171B
FBXW12
FCN1
FGF22
GABRP
GIPR
GOLGA8R
GP1BA
HAUS5
HIST3H2BB
HOXB13
HRNR
ITIH4
KIAA0930
KRT26
MVB12A
MYT1
NBPF1
NBPF11
NBPF12
NCF1
NDOR1
NME9
NPIPB11
NPIPB3
NUDT1
NUTM2E
PIDD1
PILRB
PLA2G4B
PMFBP1
PNPLA7
POLD1
PPM1N
PRAM1
PRDM15
PUS1
RGS11
ROBO3
RP11-1212A22.4
RP11-231C14.4
RP11-644F5.10
RTEL1
SLC5A9
SOX12
SPDYE6
SPTBN5
STAC3
STARD3
STXBP2
SUV420H2
TEX14
TLCD2
TMPRSS9
TSSK4
TTC31
ZDHHC11B
ZFP41
ZNF133
ZNF19
ZNF274
ZNF439
ZNF445
ZNF589
ZNF594
ZNF663P
ZNF7
ZNF70
ZNF76
ZSCAN25
Diagnosis
Sex
Cytosolic Iron-sulfur Cluster Assembly
ATF2
HumanNet interactions
Co-expression
Functional
Protein-protein interactions
Diagnosis
AD
Control
Sex
Male
Female
Functional enrichment
Functional enrichment
Motif enrichment
RcisTarget
Transcription factor binding enrichment
ChIP-enrichR
Expression ratios
<= -2.0
-0
>= 2.0

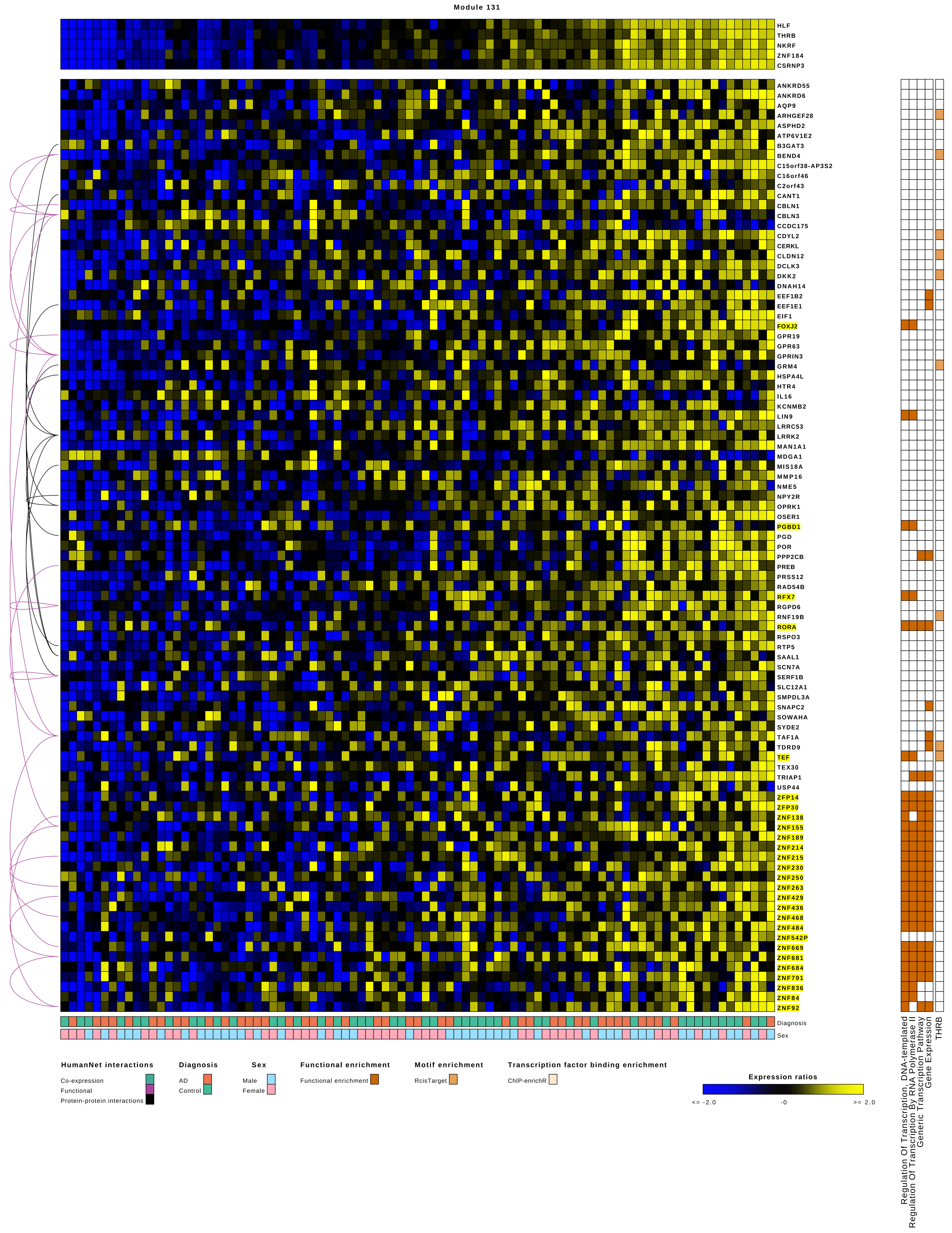
Module 131
HLF
THRB
NKRF
ZNF184
CSRNP3
ANKRD55
ANKRD6
AQP9
ARHGEF28
ASPHD2
ATP6V1E2
B3GAT3
BEND4
C15orf38-AP3S2
C16orf46
C2orf43
CANT1
CBLN1
CBLN3
CCDC175
CDYL2
CERKL
CLDN12
DCLK3
DKK2
DNAH14
EEF1B2
EEF1E1
EIF1
FOXJ2
GPR19
GPR63
GPRIN3
GRM4
HSPA4L
HTR4
IL16
KCNMB2
LIN9
LRRC53
LRRK2
MAN1A1
MDGA1
MIS18A
MMP16
NME5
NPY2R
OPRK1
OSER1
PGBD1
PGD
POR
PPP2CB
PREB
PRSS12
RAD54B
RFX7
RGPD6
RNF19B
RORA
RSPO3
RTP5
SAAL1
SCN7A
SERF1B
SLC12A1
SMPDL3A
SNAPC2
SOWAHA
SYDE2
TAF1A
TDRD9
TEF
TEX30
TRIAP1
USP44
ZFP14
ZFP30
ZNF138
ZNF155
ZNF189
ZNF214
ZNF215
ZNF230
ZNF250
ZNF263
ZNF429
ZNF436
ZNF468
ZNF484
ZNF542P
ZNF669
ZNF681
ZNF684
ZNF791
ZNF836
ZNF84
ZNF92
Diagnosis
Sex
Regulation Of Transcription, DNA-templated
Regulation Of Transcription By RNA Polymerase II
Generic Transcription Pathway
Gene Expression
THRB
HumanNet interactions
Co-expression
Functional
Protein-protein interactions
Diagnosis
AD
Control
Sex
Male
Female
Functional enrichment
Functional enrichment
Motif enrichment
RcisTarget
Transcription factor binding enrichment
ChIP-enrichR
Expression ratios
<= -2.0
-0
>= 2.0

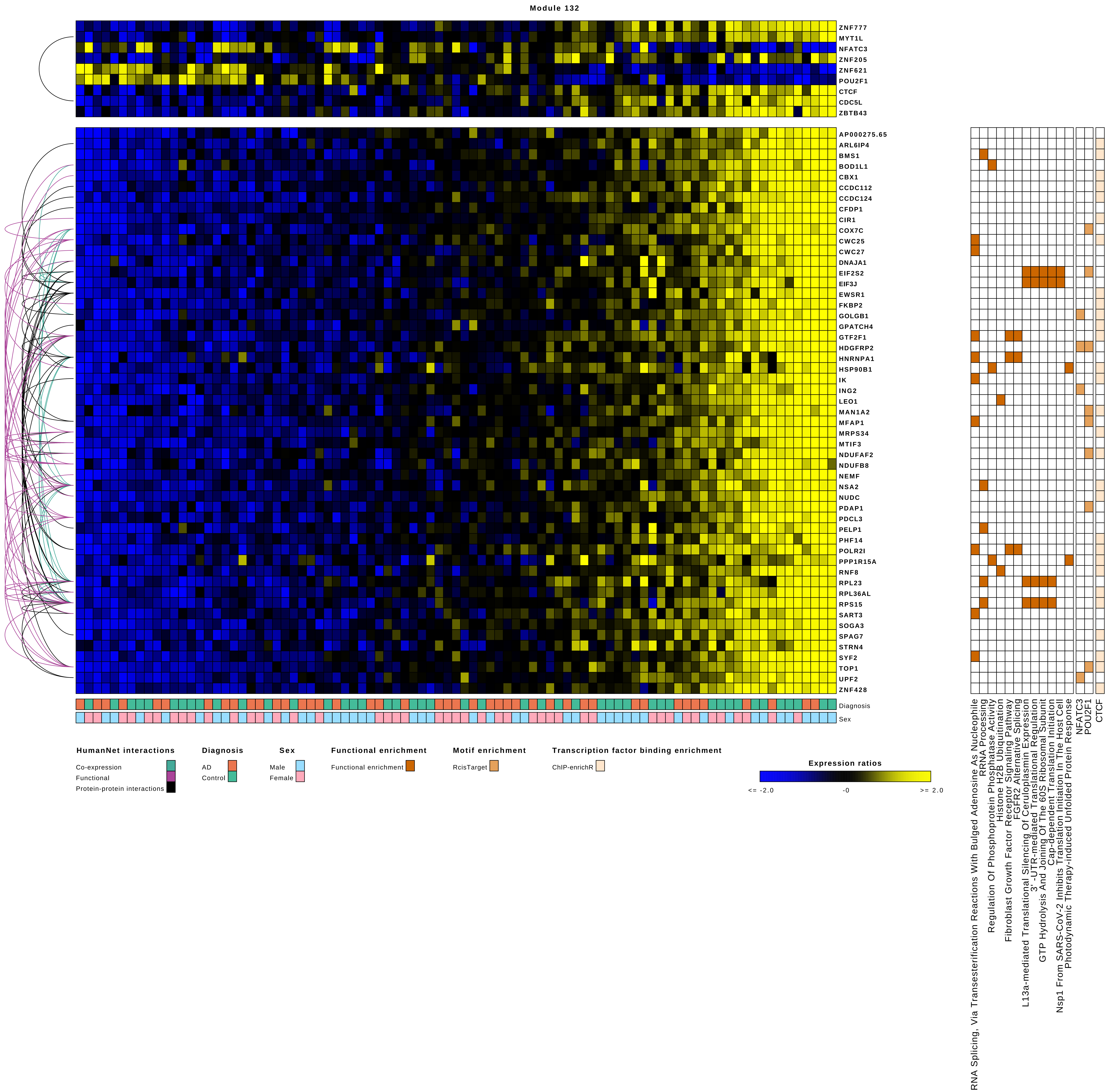
Module 132
ZNF777
MYT1L
NFATC3
ZNF205
ZNF621
POU2F1
CTCF
CDC5L
ZBTB43
AP000275.65
ARL6IP4
BMS1
BOD1L1
CBX1
CCDC112
CCDC124
CFDP1
CIR1
COX7C
CWC25
CWC27
DNAJA1
EIF2S2
EIF3J
EWSR1
FKBP2
GOLGB1
GPATCH4
GTF2F1
HDGFRP2
HNRNPA1
HSP90B1
IK
ING2
LEO1
MAN1A2
MFAP1
MRPS34
MTIF3
NDUFAF2
NDUFB8
NEMF
NSA2
NUDC
PDAP1
PDCL3
PELP1
PHF14
POLR2I
PPP1R15A
RNF8
RPL23
RPL36AL
RPS15
SART3
SOGA3
SPAG7
STRN4
SYF2
TOP1
UPF2
ZNF428
Diagnosis
Sex
RNA Splicing, Via Transesterification Reactions With Bulged Adenosine As Nucleophile
RRNA Processing
Regulation Of Phosphoprotein Phosphatase Activity
Histone H2B Ubiquitination
Fibroblast Growth Factor Receptor Signaling Pathway
FGFR2 Alternative Splicing
L13a-mediated Translational Silencing Of Ceruloplasmin Expression
3'-UTR-mediated Translational Regulation
GTP Hydrolysis And Joining Of The 60S Ribosomal Subunit
Cap-dependent Translation Initiation
Nsp1 From SARS-CoV-2 Inhibits Translation Initiation In The Host Cell
Photodynamic Therapy-induced Unfolded Protein Response
NFATC3
POU2F1
CTCF
HumanNet interactions
Co-expression
Functional
Protein-protein interactions
Diagnosis
AD
Control
Sex
Male
Female
Functional enrichment
Functional enrichment
Motif enrichment
RcisTarget
Transcription factor binding enrichment
ChIP-enrichR
Expression ratios
<= -2.0
-0
>= 2.0

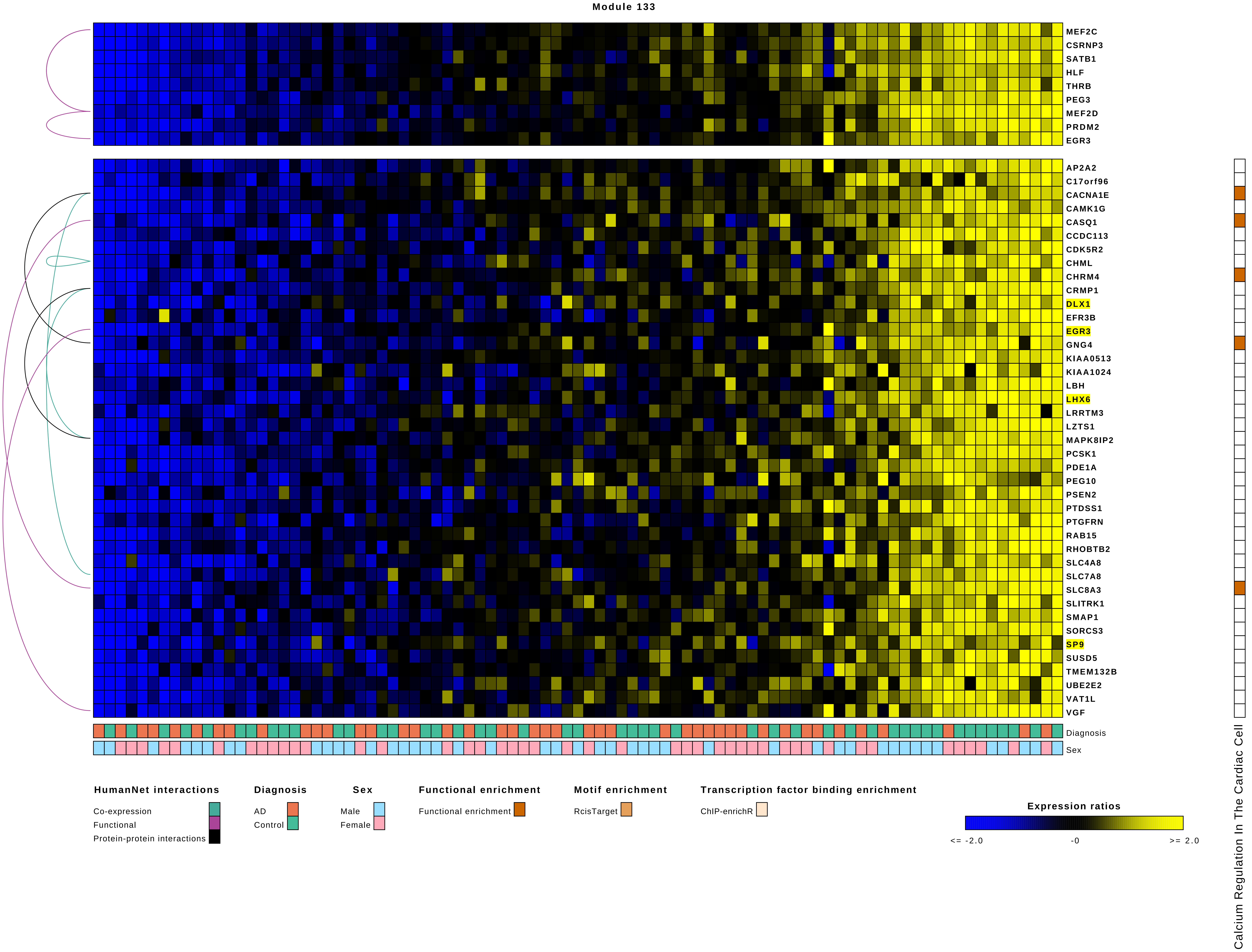
Module 133
MEF2C
CSRNP3
SATB1
HLF
THRB
PEG3
MEF2D
PRDM2
EGR3
AP2A2
C17orf96
CACNA1E
CAMK1G
CASQ1
CCDC113
CDK5R2
CHML
CHRM4
CRMP1
DLX1
EFR3B
EGR3
GNG4
KIAA0513
KIAA1024
LBH
LHX6
LRRTM3
LZTS1
MAPK8IP2
PCSK1
PDE1A
PEG10
PSEN2
PTDSS1
PTGFRN
RAB15
RHOBTB2
SLC4A8
SLC7A8
SLC8A3
SLITRK1
SMAP1
SORCS3
SP9
SUSD5
TMEM132B
UBE2E2
VAT1L
VGF
Diagnosis
Sex
Calcium Regulation In The Cardiac Cell
HumanNet interactions
Co-expression
Functional
Protein-protein interactions
Diagnosis
AD
Control
Sex
Male
Female
Functional enrichment
Functional enrichment
Motif enrichment
RcisTarget
Transcription factor binding enrichment
ChIP-enrichR
Expression ratios
<= -2.0
-0
>= 2.0

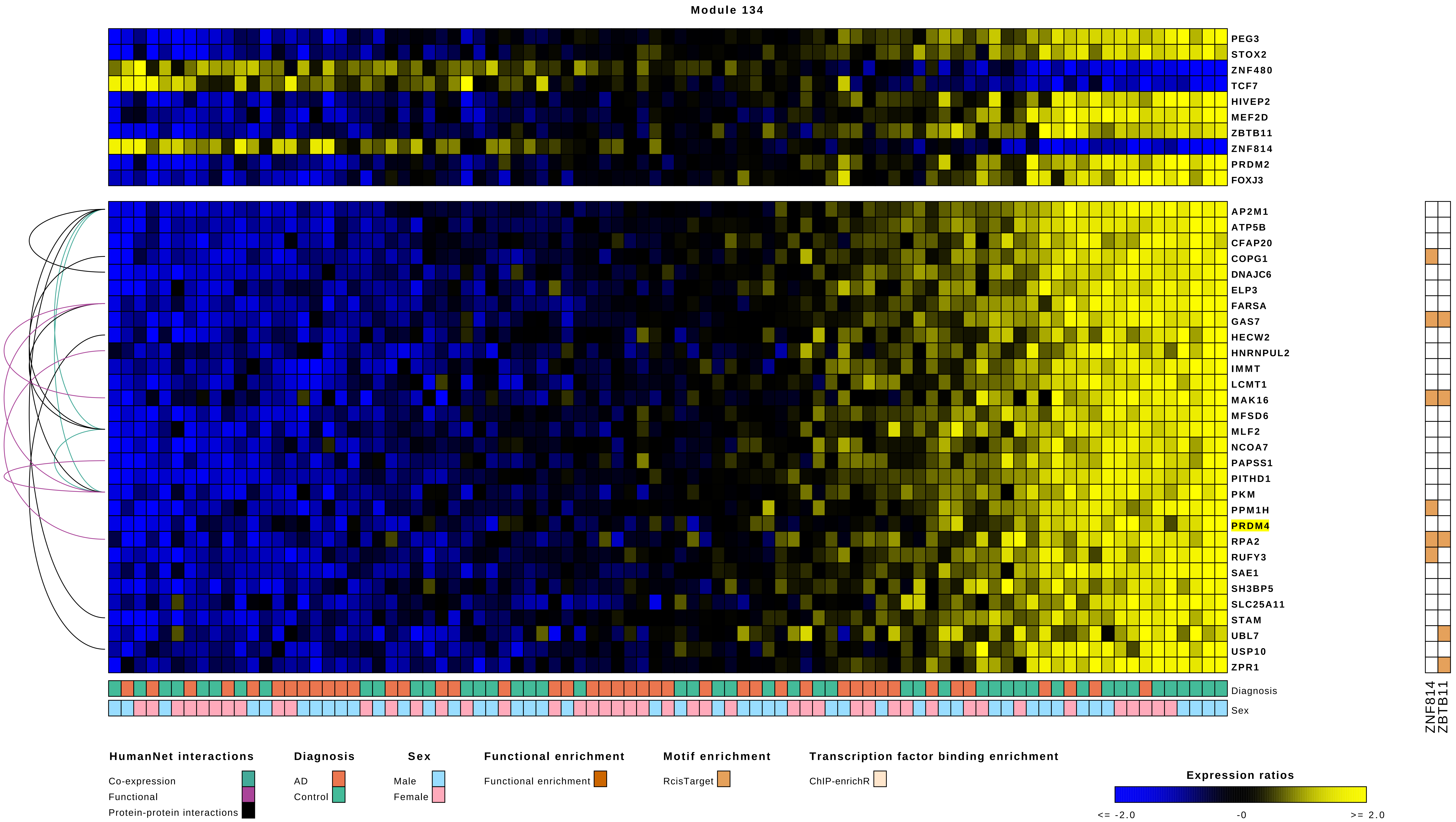

Module 134
PEG3
STOX2
ZNF480
TCF7
HIVEP2
MEF2D
ZBTB11
ZNF814
PRDM2
FOXJ3
AP2M1
ATP5B
CFAP20
COPG1
DNAJC6
ELP3
FARSA
GAS7
HECW2
HNRNPUL2
IMMT
LCMT1
MAK16
MFSD6
MLF2
NCOA7
PAPSS1
PITHD1
PKM
PPM1H
PRDM4
RPA2
RUFY3
SAE1
SH3BP5
SLC25A11
STAM
UBL7
USP10
ZPR1
Diagnosis
Sex
ZNF814
ZBTB11
HumanNet interactions
Co-expression
Functional
Protein-protein interactions
Diagnosis
AD
Control
Sex
Male
Female
Functional enrichment
Functional enrichment
Motif enrichment
RcisTarget
Transcription factor binding enrichment
ChIP-enrichR
Expression ratios
<= -2.0
-0
>= 2.0

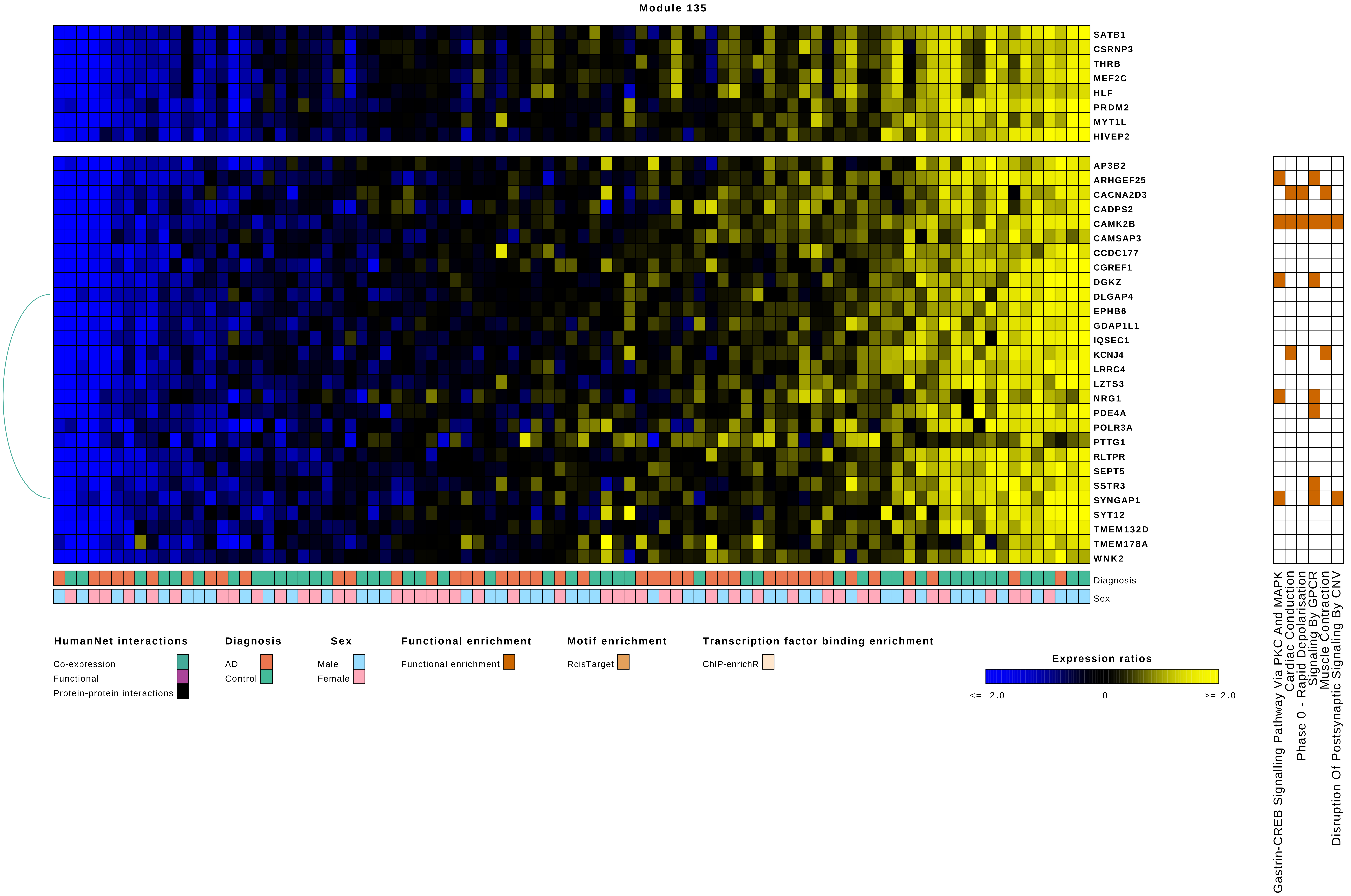
Module 135
SATB1
CSRNP3
THRB
MEF2C
HLF
PRDM2
MYT1L
HIVEP2
AP3B2
ARHGEF25
CACNA2D3
CADPS2
CAMK2B
CAMSAP3
CCDC177
CGREF1
DGKZ
DLGAP4
EPHB6
GDAP1L1
IQSEC1
KCNJ4
LRRC4
LZTS3
NRG1
PDE4A
POLR3A
PTTG1
RLTPR
SEPT5
SSTR3
SYNGAP1
SYT12
TMEM132D
TMEM178A
WNK2
Diagnosis
Sex
HumanNet interactions
Co-expression
Functional
Protein-protein interactions
Diagnosis
AD
Control
Sex
Male
Female
Functional enrichment
Functional enrichment
Motif enrichment
RcisTarget
Transcription factor binding enrichment
ChIP-enrichR
Expression ratios
<= -2.0
-0
>= 2.0
Gastrin-CREB Signalling Pathway Via PKC And MAPK
Cardiac Conduction
Phase 0 - Rapid Depolarisation
Signaling By GPCR
Muscle Contraction
Disruption Of Postsynaptic Signaling By CNV

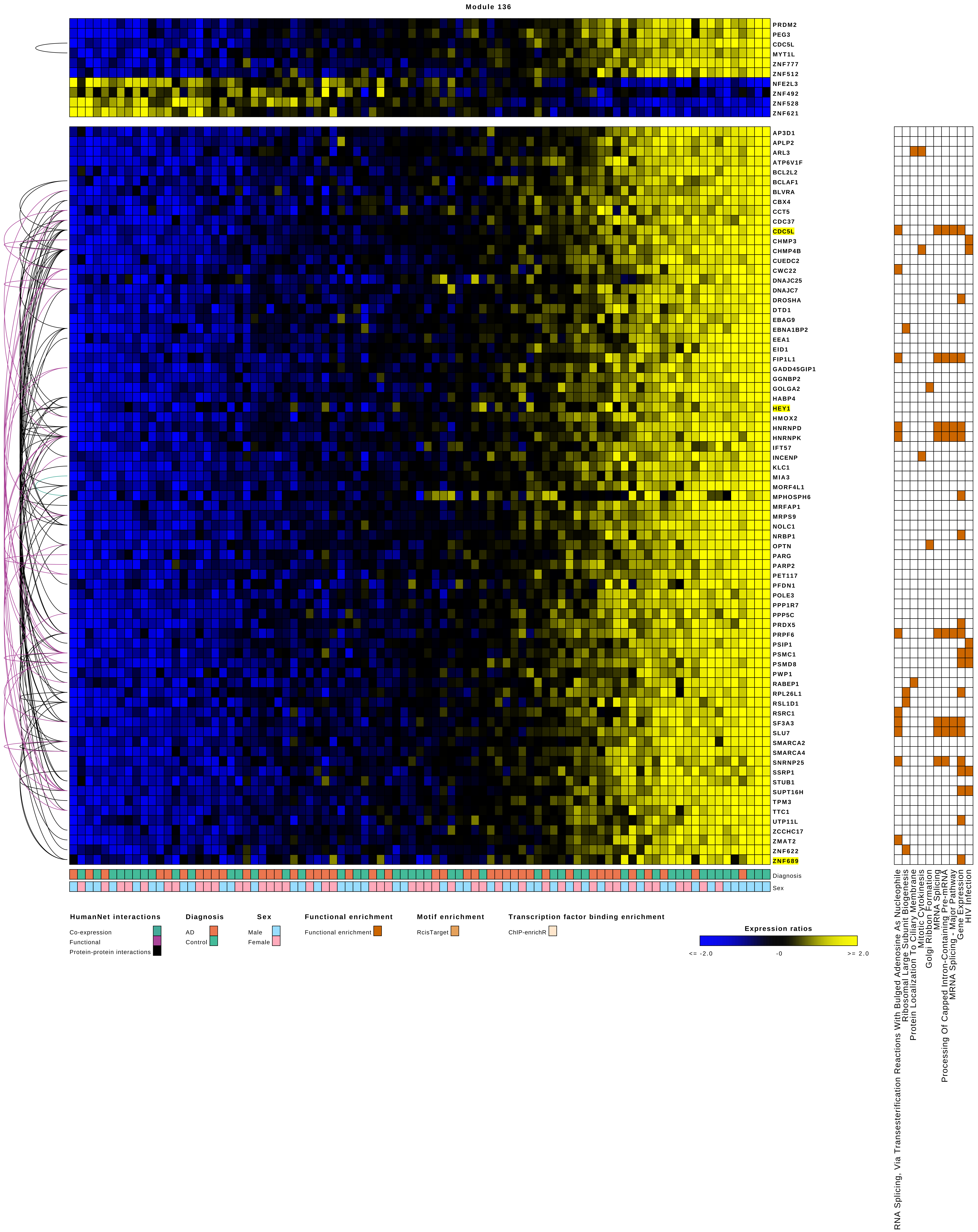

Module 136
PRDM2
PEG3
CDC5L
MYT1L
ZNF777
ZNF512
NFE2L3
ZNF492
ZNF528
ZNF621
AP3D1
APLP2
ARL3
ATP6V1F
BCL2L2
BCLAF1
BLVRA
CBX4
CCT5
CDC37
CDC5L
CHMP3
CHMP4B
CUEDC2
CWC22
DNAJC25
DNAJC7
DROSHA
DTD1
EBAG9
EBNA1BP2
EEA1
EID1
FIP1L1
GADD45GIP1
GGNBP2
GOLGA2
HABP4
HEY1
HMOX2
HNRNPD
HNRNPK
IFT57
INCENP
KLC1
MIA3
MORF4L1
MPHOSPH6
MRFAP1
MRPS9
NOLC1
NRBP1
OPTN
PARG
PARP2
PET117
PFDN1
POLE3
PPP1R7
PPP5C
PRDX5
PRPF6
PSIP1
PSMC1
PSMD8
PWP1
RABEP1
RPL26L1
RSL1D1
RSRC1
SF3A3
SLU7
SMARCA2
SMARCA4
SNRNP25
SSRP1
STUB1
SUPT16H
TPM3
TTC1
UTP11L
ZCCHC17
ZMAT2
ZNF622
ZNF689
Diagnosis
Sex
HumanNet interactions
Co-expression
Functional
Protein-protein interactions
Diagnosis
AD
Control
Sex
Male
Female
Functional enrichment
Functional enrichment
Motif enrichment
RcisTarget
Transcription factor binding enrichment
ChIP-enrichR
Expression ratios
<= -2.0
-0
>= 2.0
RNA Splicing, Via Transesterification Reactions With Bulged Adenosine As Nucleophile
Ribosomal Large Subunit Biogenesis
Protein Localization To Ciliary Membrane
Mitotic Cytokinesis
Golgi Ribbon Formation
MRNA Splicing
Processing Of Capped Intron-Containing Pre-mRNA
MRNA Splicing - Major Pathway
Gene Expression
HIV Infection

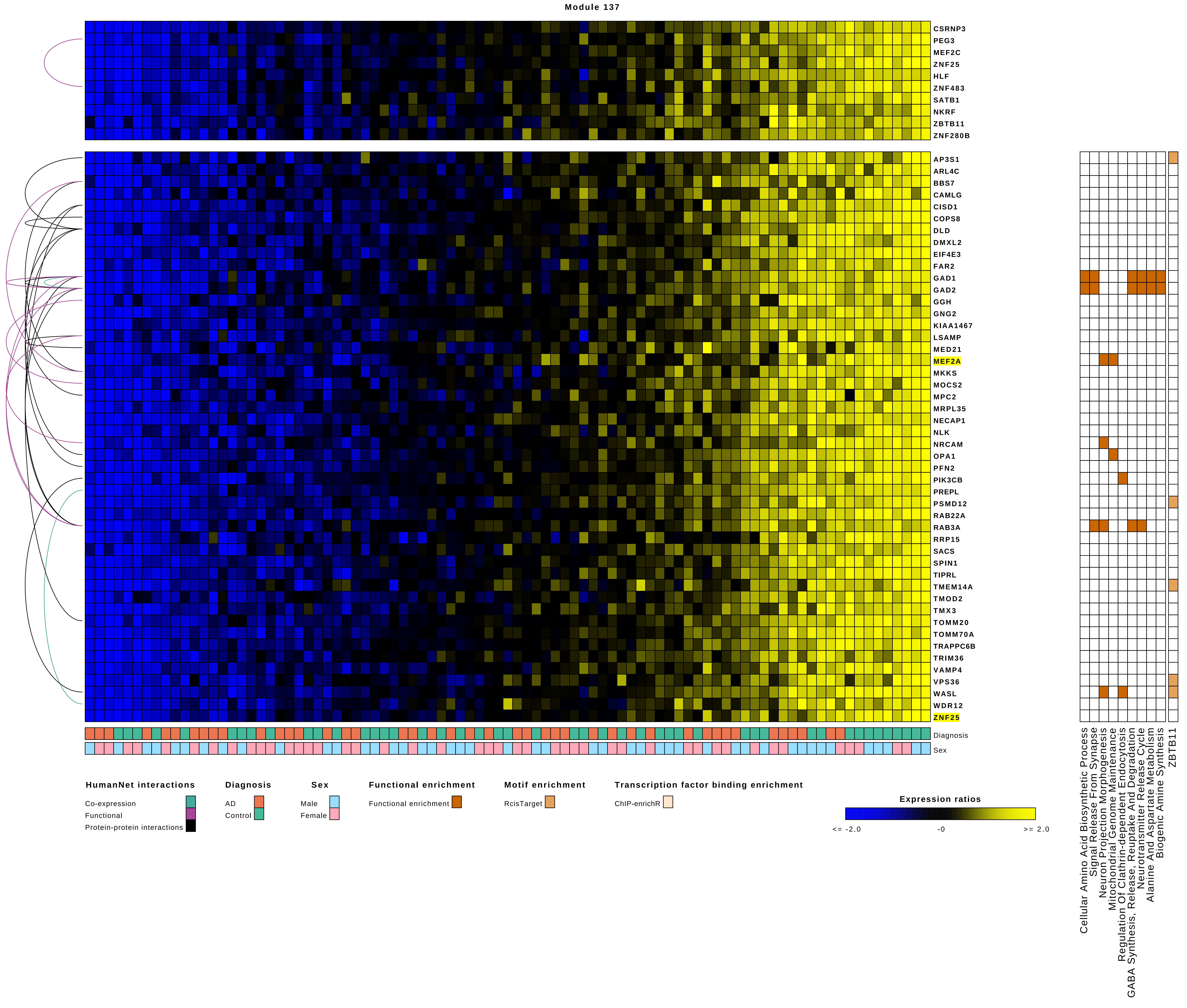
Module 137
CSRNP3
PEG3
MEF2C
ZNF25
HLF
ZNF483
SATB1
NKRF
ZBTB11
ZNF280B
AP3S1
ARL4C
BBS7
CAMLG
CISD1
COPS8
DLD
DMXL2
EIF4E3
FAR2
GAD1
GAD2
GGH
GNG2
KIAA1467
LSAMP
MED21
MEF2A
MKKS
MOCS2
MPC2
MRPL35
NECAP1
NLK
NRCAM
OPA1
PFN2
PIK3CB
PREPL
PSMD12
RAB22A
RAB3A
RRP15
SACS
SPIN1
TIPRL
TMEM14A
TMOD2
TMX3
TOMM20
TOMM70A
TRAPPC6B
TRIM36
VAMP4
VPS36
WASL
WDR12
ZNF25
Diagnosis
Sex
Cellular Amino Acid Biosynthetic Process
Signal Release From Synapse
Neuron Projection Morphogenesis
Mitochondrial Genome Maintenance
Regulation Of Clathrin-dependent Endocytosis
GABA Synthesis, Release, Reuptake And Degradation
Neurotransmitter Release Cycle
Alanine And Aspartate Metabolism
Biogenic Amine Synthesis
ZBTB11
HumanNet interactions
Co-expression
Functional
Protein-protein interactions
Diagnosis
AD
Control
Sex
Male
Female
Functional enrichment
Functional enrichment
Motif enrichment
RcisTarget
Transcription factor binding enrichment
ChIP-enrichR
Expression ratios
<= -2.0
-0
>= 2.0

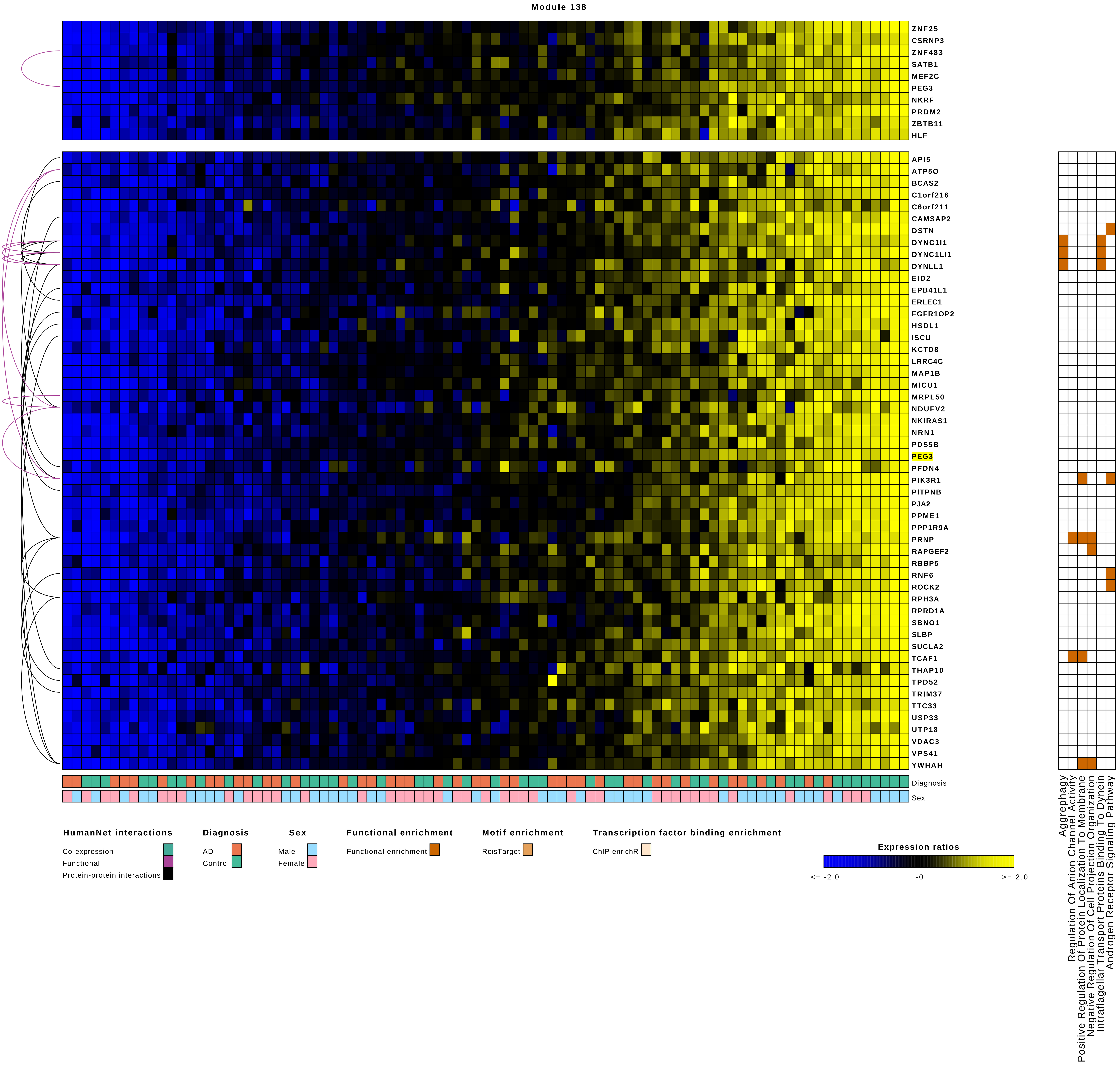
Module 138
ZNF25
CSRNP3
ZNF483
SATB1
MEF2C
PEG3
NKRF
PRDM2
ZBTB11
HLF
API5
ATP5O
BCAS2
C1orf216
C6orf211
CAMSAP2
DSTN
DYNC1I1
DYNC1LI1
DYNLL1
EID2
EPB41L1
ERLEC1
FGFR1OP2
HSDL1
ISCU
KCTD8
LRRC4C
MAP1B
MICU1
MRPL50
NDUFV2
NKIRAS1
NRN1
PDS5B
PEG3
PFDN4
PIK3R1
PITPNB
PJA2
PPME1
PPP1R9A
PRNP
RAPGEF2
RBBP5
RNF6
ROCK2
RPH3A
RPRD1A
SBNO1
SLBP
SUCLA2
TCAF1
THAP10
TPD52
TRIM37
TTC33
USP33
UTP18
VDAC3
VPS41
YWHAH
Diagnosis
Sex
Aggrephagy
Regulation Of Anion Channel Activity
Positive Regulation Of Protein Localization To Membrane
Negative Regulation Of Cell Projection Organization
Intraflagellar Transport Proteins Binding To Dynein
Androgen Receptor Signaling Pathway
HumanNet interactions
Co-expression
Functional
Protein-protein interactions
Diagnosis
AD
Control
Sex
Male
Female
Functional enrichment
Functional enrichment
Motif enrichment
RcisTarget
Transcription factor binding enrichment
ChIP-enrichR
Expression ratios
<= -2.0
-0
>= 2.0

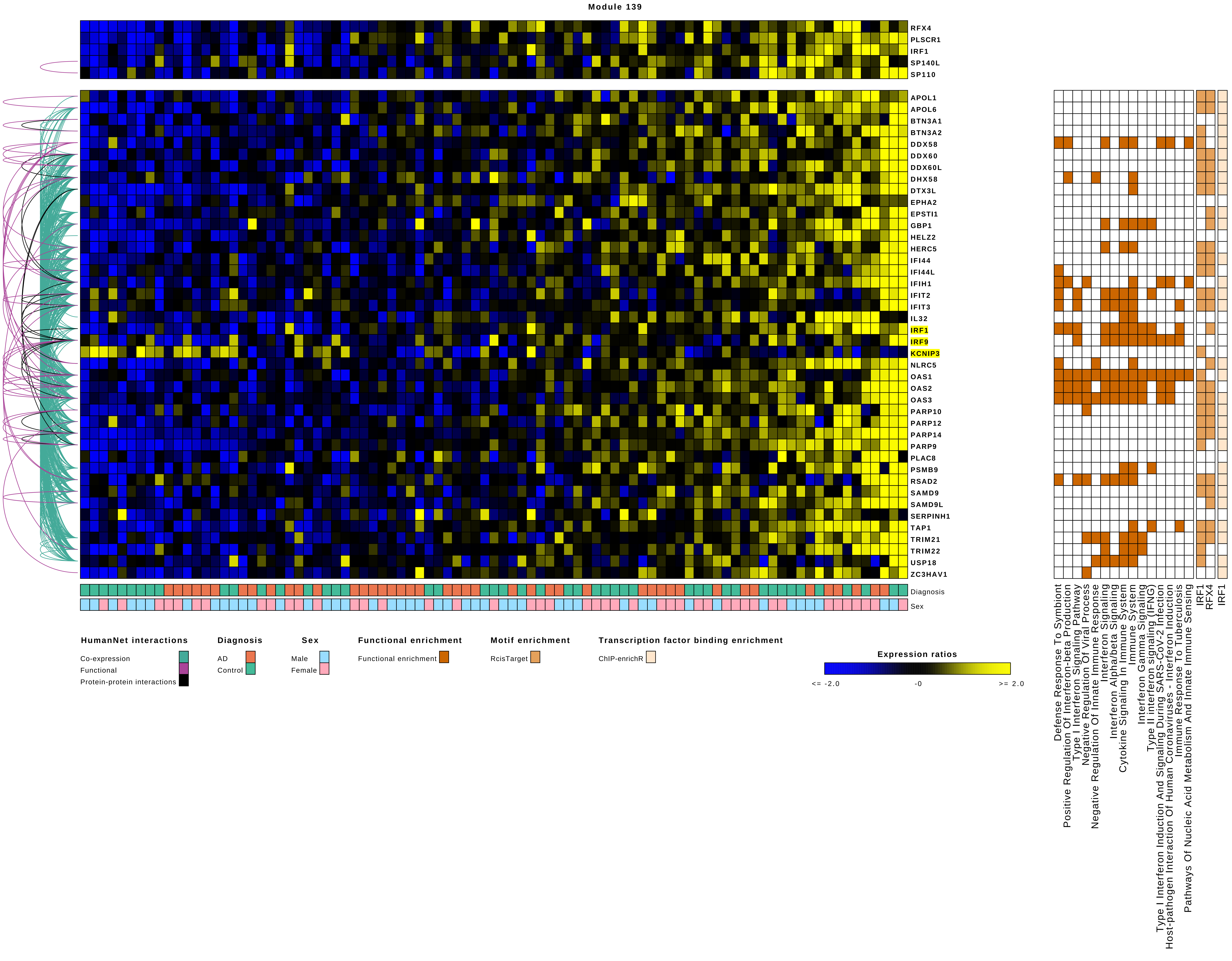
Module 139
RFX4
PLSCR1
IRF1
SP140L
SP110
APOL1
APOL6
BTN3A1
BTN3A2
DDX58
DDX60
DDX60L
DHX58
DTX3L
EPHA2
EPSTI1
GBP1
HELZ2
HERC5
IFI44
IFI44L
IFIH1
IFIT2
IFIT3
IL32
IRF1
IRF9
KCNIP3
NLRC5
OAS1
OAS2
OAS3
PARP10
PARP12
PARP14
PARP9
PLAC8
PSMB9
RSAD2
SAMD9
SAMD9L
SERPINH1
TAP1
TRIM21
TRIM22
USP18
ZC3HAV1
Diagnosis
Sex
Defense Response To Symbiont
Positive Regulation Of Interferon-beta Production
Type I Interferon Signaling Pathway
Negative Regulation Of Viral Process
Negative Regulation Of Innate Immune Response
Interferon Signaling
Interferon Alpha/beta Signaling
Cytokine Signaling In Immune System
Immune System
Interferon Gamma Signaling
Type II interferon signaling (IFNG)
Type I Interferon Induction And Signaling During SARS-CoV-2 Infection
Host-pathogen Interaction Of Human Coronaviruses - Interferon Induction
Immune Response To Tuberculosis
Pathways Of Nucleic Acid Metabolism And Innate Immune Sensing
IRF1
RFX4
IRF1
HumanNet interactions
Co-expression
Functional
Protein-protein interactions
Diagnosis
AD
Control
Sex
Male
Female
Functional enrichment
Functional enrichment
Motif enrichment
RcisTarget
Transcription factor binding enrichment
ChIP-enrichR
Expression ratios
<= -2.0
-0
>= 2.0

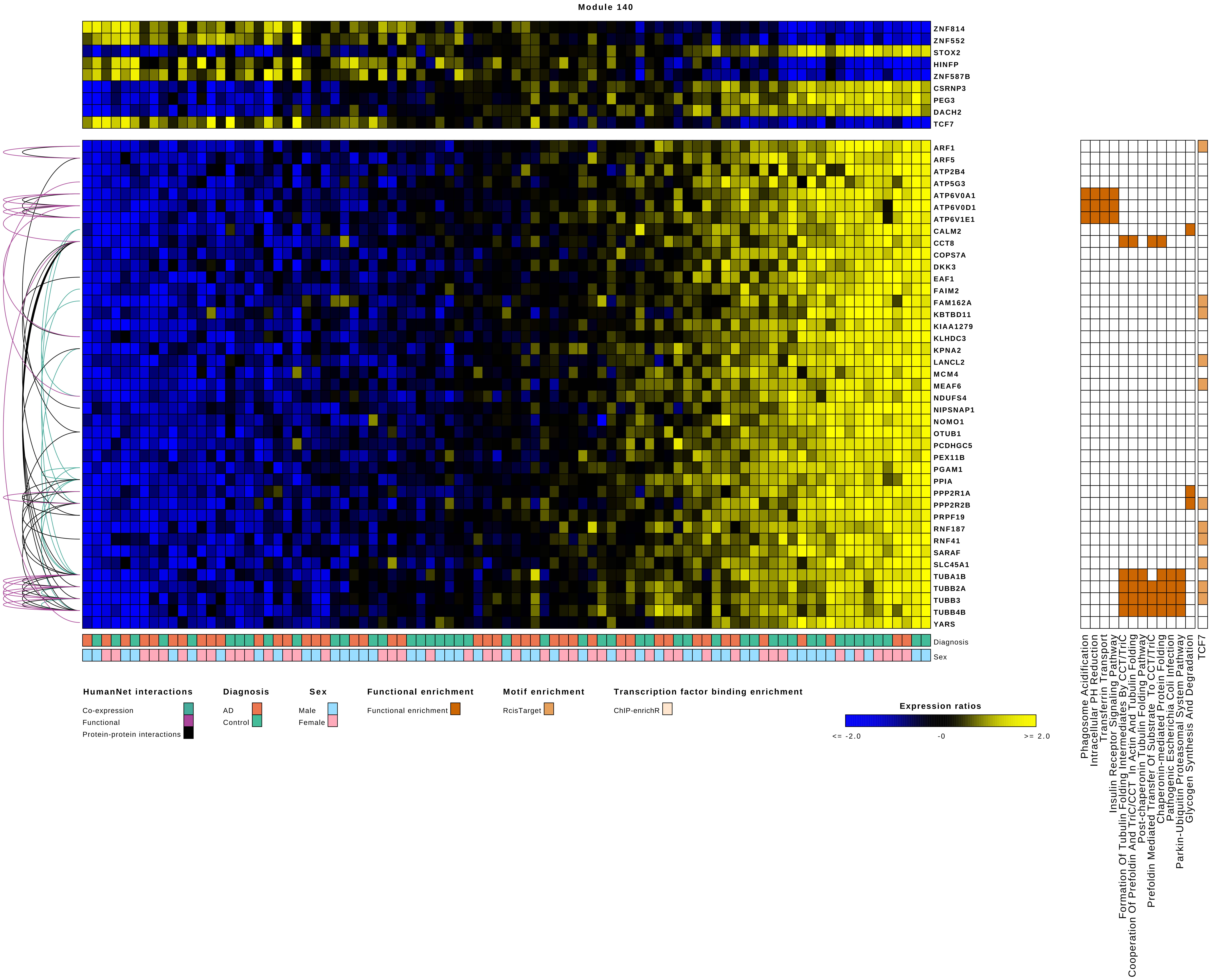

Module 140
ZNF814
ZNF552
STOX2
HINFP
ZNF587B
CSRNP3
PEG3
DACH2
TCF7
ARF1
ARF5
ATP2B4
ATP5G3
ATP6V0A1
ATP6V0D1
ATP6V1E1
CALM2
CCT8
COPS7A
DKK3
EAF1
FAIM2
FAM162A
KBTBD11
KIAA1279
KLHDC3
KPNA2
LANCL2
MCM4
MEAF6
NDUFS4
NIPSNAP1
NOMO1
OTUB1
PCDHGC5
PEX11B
PGAM1
PPIA
PPP2R1A
PPP2R2B
PRPF19
RNF187
RNF41
SARAF
SLC45A1
TUBA1B
TUBB2A
TUBB3
TUBB4B
YARS
Diagnosis
Sex
HumanNet interactions
Co-expression
Functional
Protein-protein interactions
Diagnosis
AD
Control
Sex
Male
Female
Functional enrichment
Functional enrichment
Motif enrichment
RcisTarget
Transcription factor binding enrichment
ChIP-enrichR
Expression ratios
<= -2.0
-0
>= 2.0
Phagosome Acidification
Intracellular PH Reduction
Transferrin Transport
Insulin Receptor Signaling Pathway
Formation Of Tubulin Folding Intermediates By CCT/TriC
Cooperation Of Prefoldin And TriC/CCT In Actin And Tubulin Folding
Post-chaperonin Tubulin Folding Pathway
Prefoldin Mediated Transfer Of Substrate To CCT/TriC
Chaperonin-mediated Protein Folding
Pathogenic Escherichia Coli Infection
Parkin-Ubiquitin Proteasomal System Pathway
Glycogen Synthesis And Degradation
TCF7

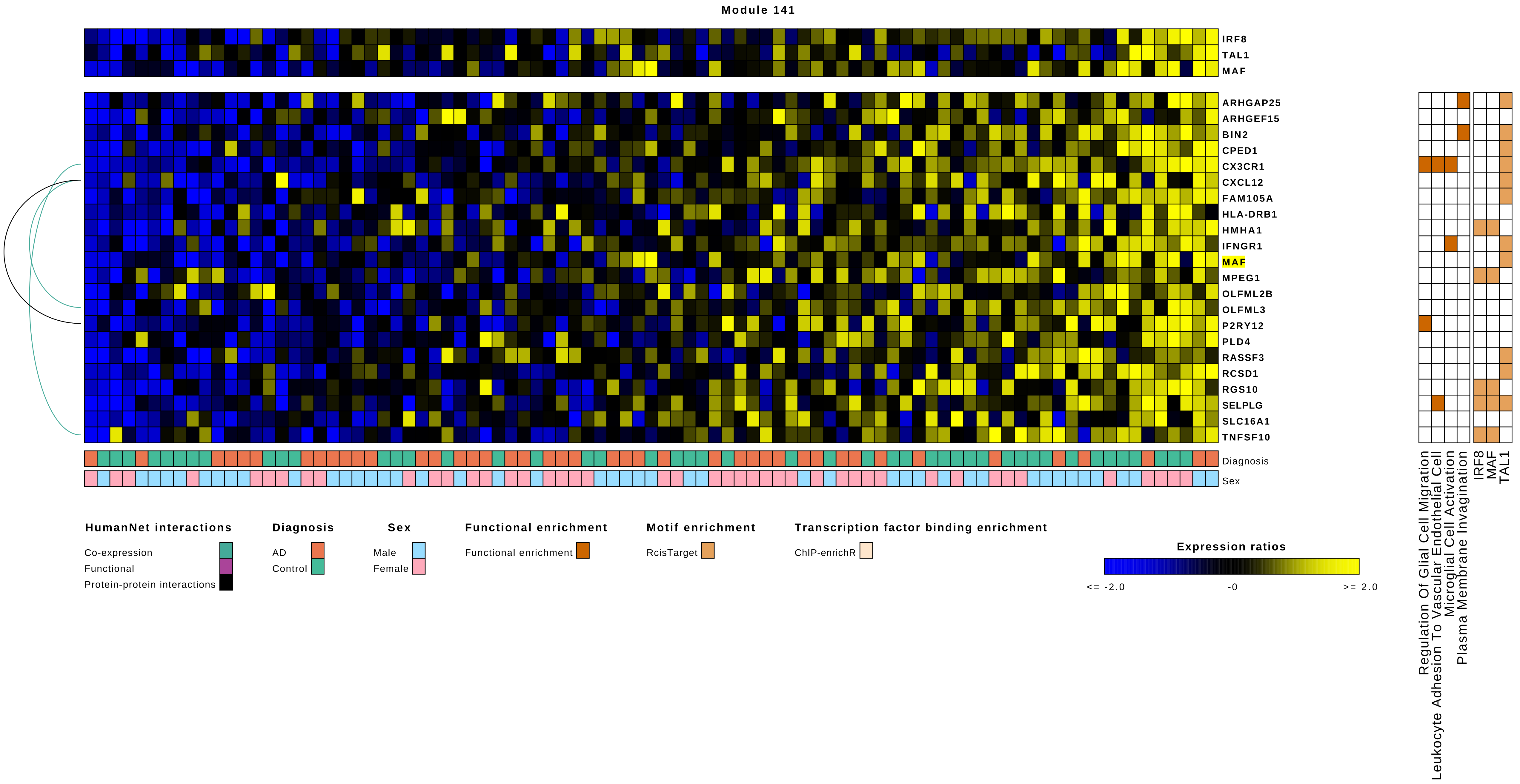
Module 141
IRF8
TAL1
MAF
ARHGAP25
ARHGEF15
BIN2
CPED1
CX3CR1
CXCL12
FAM105A
HLA-DRB1
HMHA1
IFNGR1
MAF
MPEG1
OLFML2B
OLFML3
P2RY12
PLD4
RASSF3
RCSD1
RGS10
SELPLG
SLC16A1
TNFSF10
Diagnosis
Sex
Regulation Of Glial Cell Migration
Leukocyte Adhesion To Vascular Endothelial Cell
Microglial Cell Activation
Plasma Membrane Invagination
IRF8
MAF
TAL1
HumanNet interactions
Co-expression
Functional
Protein-protein interactions
Diagnosis
AD
Control
Sex
Male
Female
Functional enrichment
Functional enrichment
Motif enrichment
RcisTarget
Transcription factor binding enrichment
ChIP-enrichR
Expression ratios
<= -2.0
-0
>= 2.0

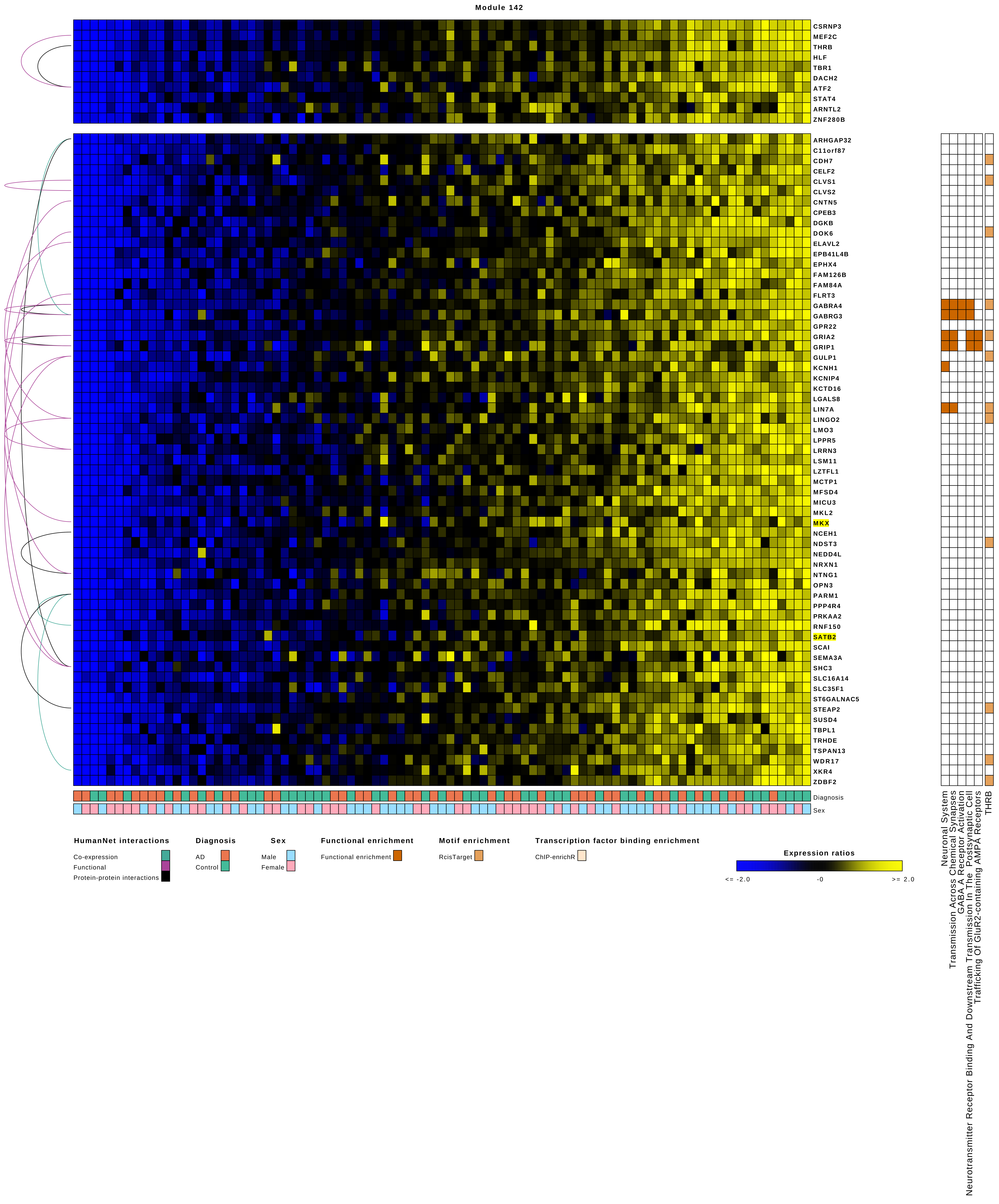
Module 142
CSRNP3
MEF2C
THRB
HLF
TBR1
DACH2
ATF2
STAT4
ARNTL2
ZNF280B
ARHGAP32
C11orf87
CDH7
CELF2
CLVS1
CLVS2
CNTN5
CPEB3
DGKB
DOK6
ELAVL2
EPB41L4B
EPHX4
FAM126B
FAM84A
FLRT3
GABRA4
GABRG3
GPR22
GRIA2
GRIP1
GULP1
KCNH1
KCNIP4
KCTD16
LGALS8
LIN7A
LINGO2
LMO3
LPPR5
LRRN3
LSM11
LZTFL1
MCTP1
MFSD4
MICU3
MKL2
MKX
NCEH1
NDST3
NEDD4L
NRXN1
NTNG1
OPN3
PARM1
PPP4R4
PRKAA2
RNF150
SATB2
SCAI
SEMA3A
SHC3
SLC16A14
SLC35F1
ST6GALNAC5
STEAP2
SUSD4
TBPL1
TRHDE
TSPAN13
WDR17
XKR4
ZDBF2
Diagnosis
Sex
Neuronal System
Transmission Across Chemical Synapses
GABA A Receptor Activation
Neurotransmitter Receptor Binding And Downstream Transmission In The Postsynaptic Cell
Trafficking Of GluR2-containing AMPA Receptors
THRB
HumanNet interactions
Co-expression
Functional
Protein-protein interactions
Diagnosis
AD
Control
Sex
Male
Female
Functional enrichment
Functional enrichment
Motif enrichment
RcisTarget
Transcription factor binding enrichment
ChIP-enrichR
Expression ratios
<= -2.0
-0
>= 2.0

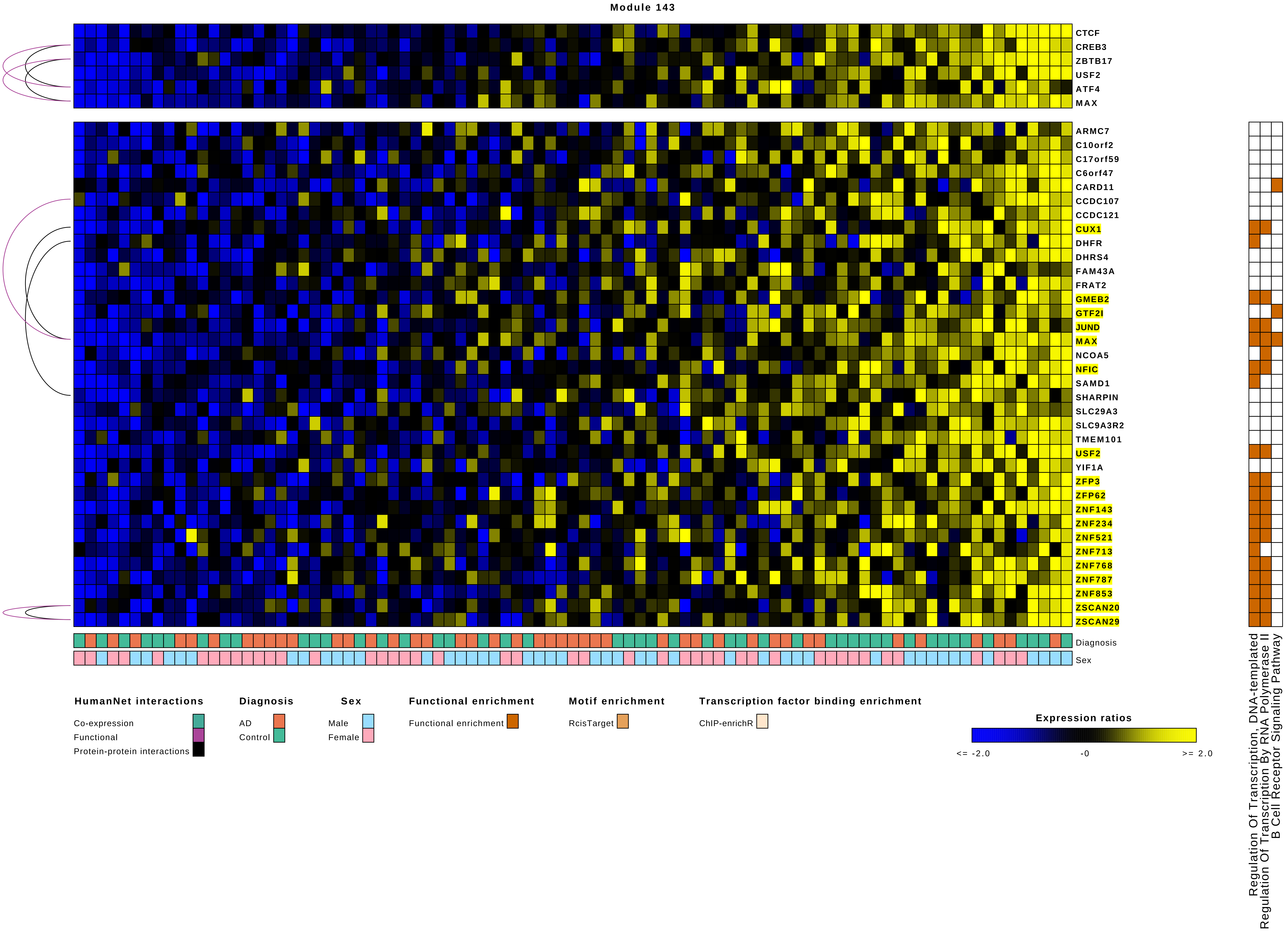
Module 143
CTCF
CREB3
ZBTB17
USF2
ATF4
MAX
ARMC7
C10orf2
C17orf59
C6orf47
CARD11
CCDC107
CCDC121
CUX1
DHFR
DHRS4
FAM43A
FRAT2
GMEB2
GTF2I
JUND
MAX
NCOA5
NFIC
SAMD1
SHARPIN
SLC29A3
SLC9A3R2
TMEM101
USF2
YIF1A
ZFP3
ZFP62
ZNF143
ZNF234
ZNF521
ZNF713
ZNF768
ZNF787
ZNF853
ZSCAN20
ZSCAN29
Diagnosis
Sex
Regulation Of Transcription, DNA-templated
Regulation Of Transcription By RNA Polymerase II
B Cell Receptor Signaling Pathway
HumanNet interactions
Co-expression
Functional
Protein-protein interactions
Diagnosis
AD
Control
Sex
Male
Female
Functional enrichment
Functional enrichment
Motif enrichment
RcisTarget
Transcription factor binding enrichment
ChIP-enrichR
Expression ratios
<= -2.0
-0
>= 2.0

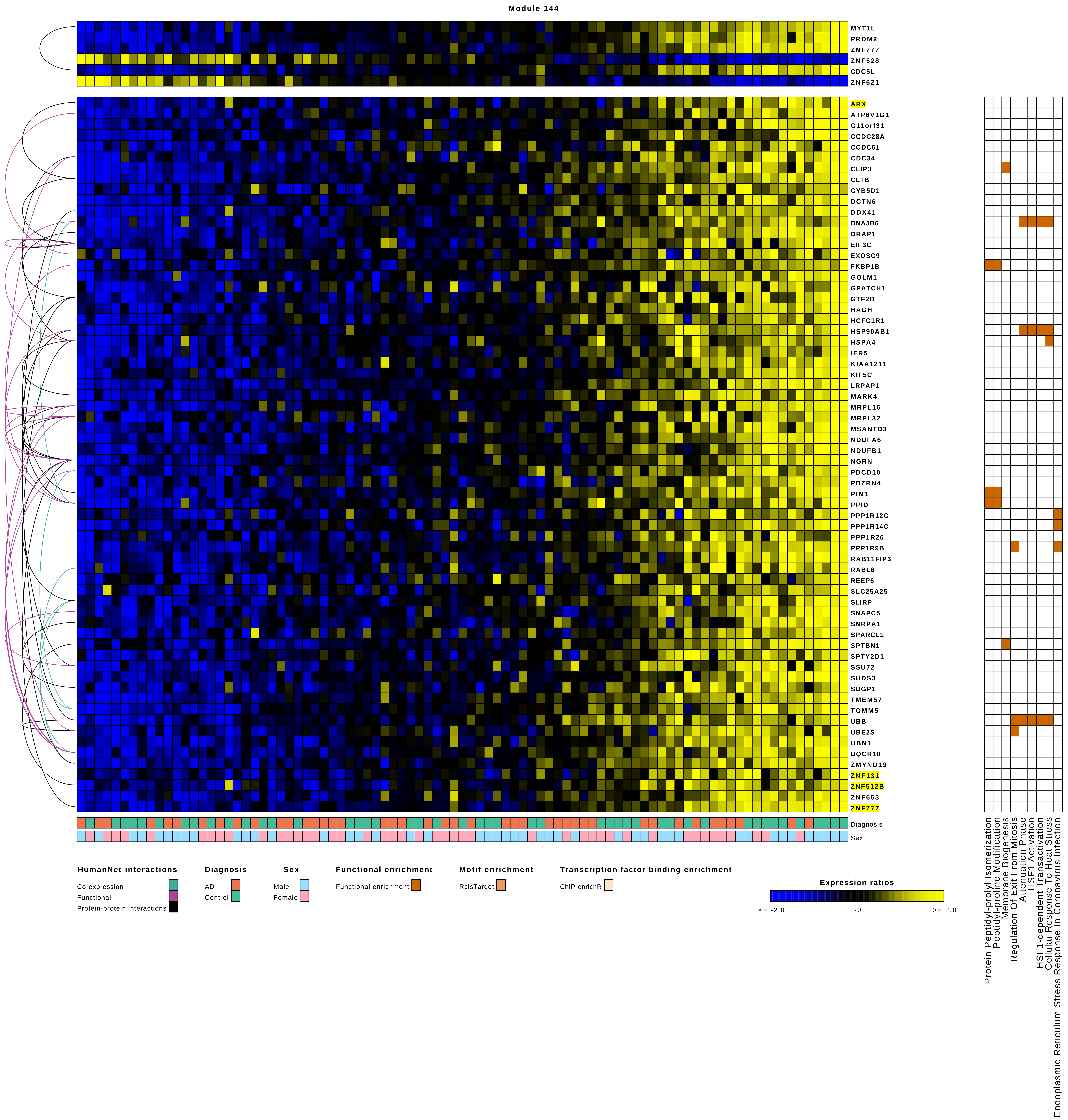

Module 144
MYT1L
PRDM2
ZNF777
ZNF528
CDC5L
ZNF621
ARX
ATP6V1G1
C11orf31
CCDC28A
CCDC51
CDC34
CLIP3
CLTB
CYB5D1
DCTN6
DDX41
DNAJB6
DRAP1
EIF3C
EXOSC9
FKBP1B
GOLM1
GPATCH1
GTF2B
HAGH
HCFC1R1
HSP90AB1
HSPA4
IER5
KIAA1211
KIF5C
LRPAP1
MARK4
MRPL16
MRPL32
MSANTD3
NDUFA6
NDUFB1
NGRN
PDCD10
PDZRN4
PIN1
PPID
PPP1R12C
PPP1R14C
PPP1R26
PPP1R9B
RAB11FIP3
RABL6
REEP6
SLC25A25
SLIRP
SNAPC5
SNRPA1
SPARCL1
SPTBN1
SPTY2D1
SSU72
SUDS3
SUGP1
TMEM57
TOMM5
UBB
UBE2S
UBN1
UQCR10
ZMYND19
ZNF131
ZNF512B
ZNF653
ZNF777
Diagnosis
Sex
Protein Peptidyl-prolyl Isomerization
Peptidyl-proline Modification
Membrane Biogenesis
Regulation Of Exit From Mitosis
Attenuation Phase
HSF1 Activation
HSF1-dependent Transactivation
Cellular Response To Heat Stress
Endoplasmic Reticulum Stress Response In Coronavirus Infection
HumanNet interactions
Co-expression
Functional
Protein-protein interactions
Diagnosis
AD
Control
Sex
Male
Female
Functional enrichment
Functional enrichment
Motif enrichment
RcisTarget
Transcription factor binding enrichment
ChIP-enrichR
Expression ratios
<= -2.0
-0
>= 2.0

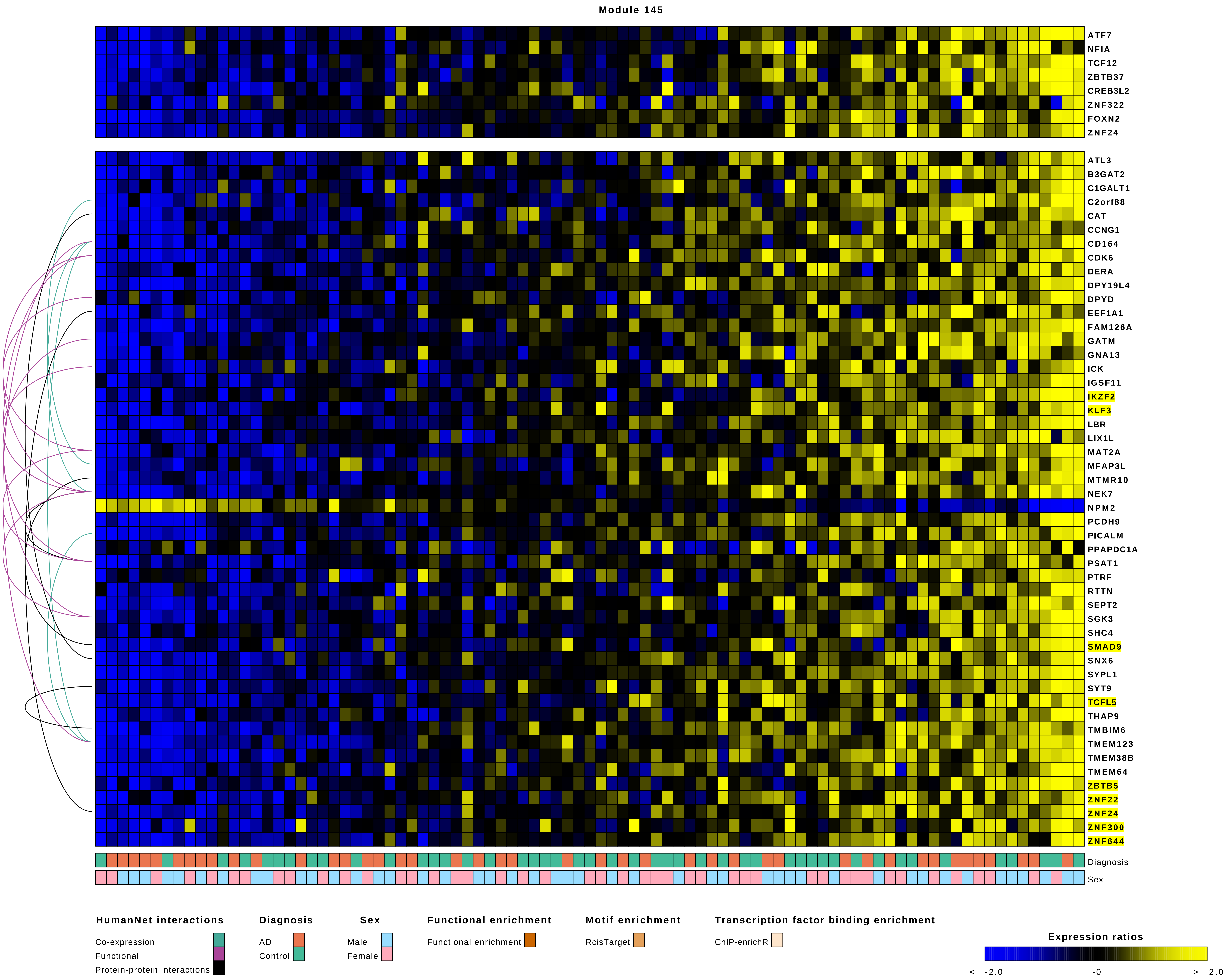
Module 145
ATF7
NFIA
TCF12
ZBTB37
CREB3L2
ZNF322
FOXN2
ZNF24
ATL3
B3GAT2
C1GALT1
C2orf88
CAT
CCNG1
CD164
CDK6
DERA
DPY19L4
DPYD
EEF1A1
FAM126A
GATM
GNA13
ICK
IGSF11
IKZF2
KLF3
LBR
LIX1L
MAT2A
MFAP3L
MTMR10
NEK7
NPM2
PCDH9
PICALM
PPAPDC1A
PSAT1
PTRF
RTTN
SEPT2
SGK3
SHC4
SMAD9
SNX6
SYPL1
SYT9
TCFL5
THAP9
TMBIM6
TMEM123
TMEM38B
TMEM64
ZBTB5
ZNF22
ZNF24
ZNF300
ZNF644
Diagnosis
Sex
HumanNet interactions
Co-expression
Functional
Protein-protein interactions
Diagnosis
AD
Control
Sex
Male
Female
Functional enrichment
Functional enrichment
Motif enrichment
RcisTarget
Transcription factor binding enrichment
ChIP-enrichR
Expression ratios
<= -2.0
-0
>= 2.0

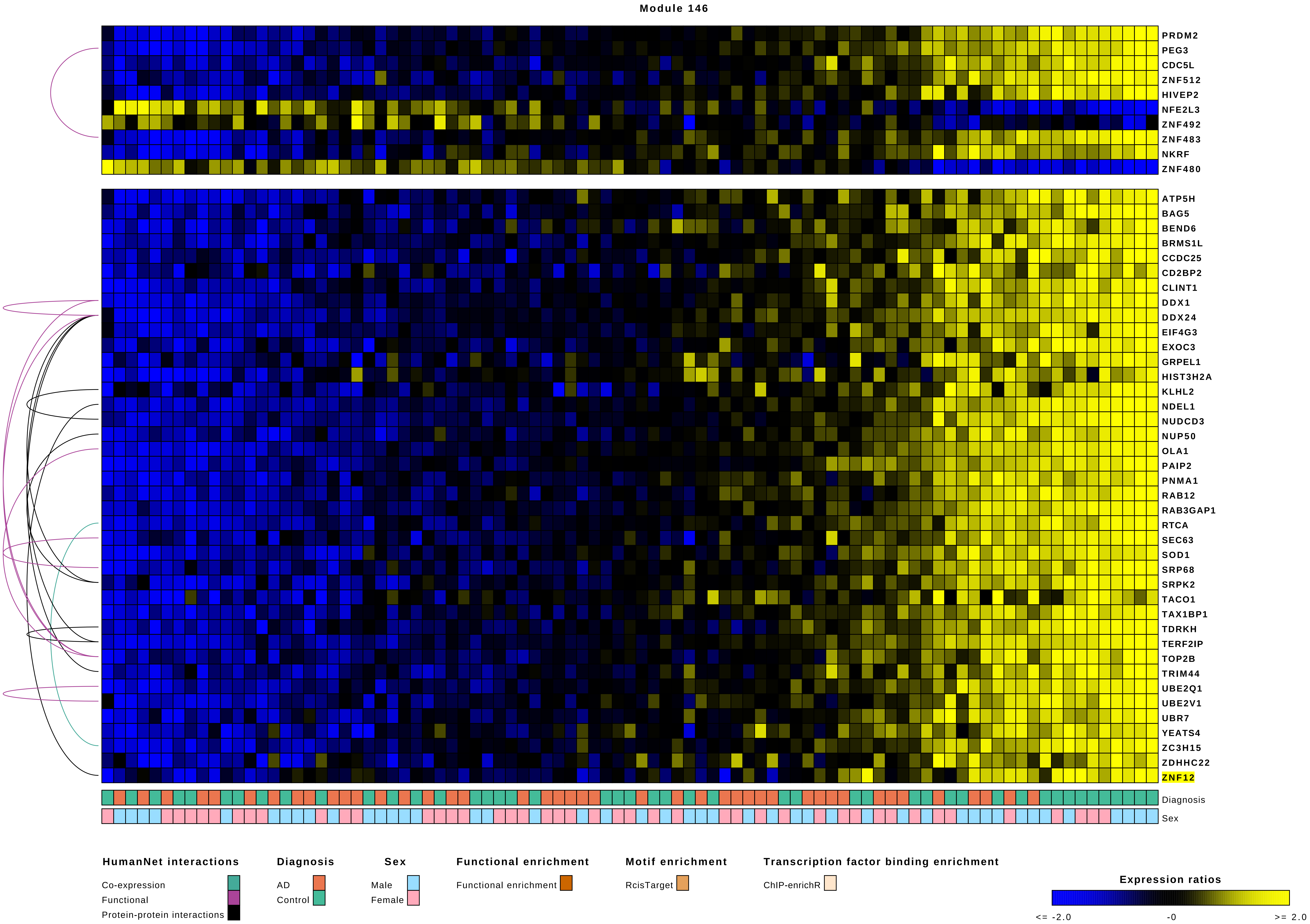

Module 146
PRDM2
PEG3
CDC5L
ZNF512
HIVEP2
NFE2L3
ZNF492
ZNF483
NKRF
ZNF480
ATP5H
BAG5
BEND6
BRMS1L
CCDC25
CD2BP2
CLINT1
DDX1
DDX24
EIF4G3
EXOC3
GRPEL1
HIST3H2A
KLHL2
NDEL1
NUDCD3
NUP50
OLA1
PAIP2
PNMA1
RAB12
RAB3GAP1
RTCA
SEC63
SOD1
SRP68
SRPK2
TACO1
TAX1BP1
TDRKH
TERF2IP
TOP2B
TRIM44
UBE2Q1
UBE2V1
UBR7
YEATS4
ZC3H15
ZDHHC22
ZNF12
Diagnosis
Sex
HumanNet interactions
Co-expression
Functional
Protein-protein interactions
Diagnosis
AD
Control
Sex
Male
Female
Functional enrichment
Functional enrichment
Motif enrichment
RcisTarget
Transcription factor binding enrichment
ChIP-enrichR
Expression ratios
<= -2.0
-0
>= 2.0

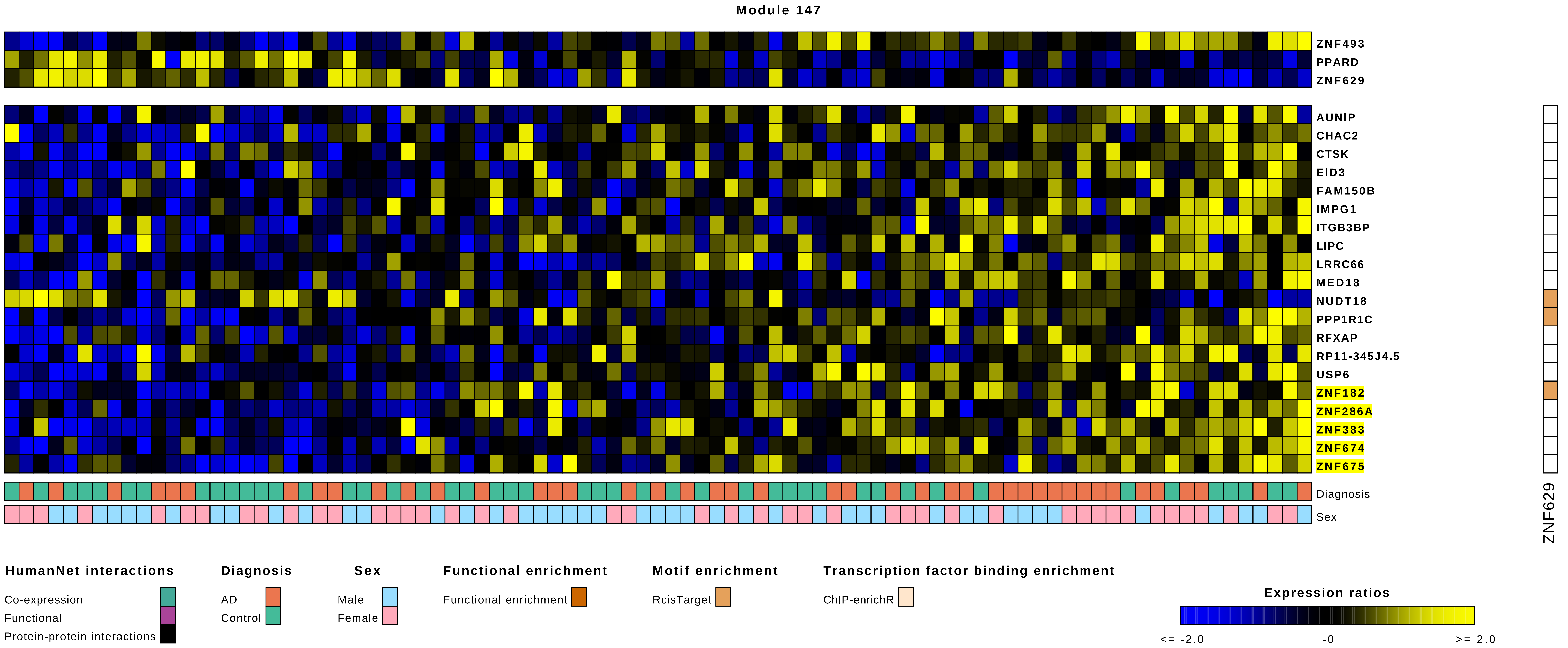
Module 147
ZNF493
PPARD
ZNF629
AUNIP
CHAC2
CTSK
EID3
FAM150B
IMPG1
ITGB3BP
LIPC
LRRC66
MED18
NUDT18
PPP1R1C
RFXAP
RP11-345J4.5
USP6
ZNF182
ZNF286A
ZNF383
ZNF674
ZNF675
Diagnosis
Sex
ZNF629
HumanNet interactions
Co-expression
Functional
Protein-protein interactions
Diagnosis
AD
Control
Sex
Male
Female
Functional enrichment
Functional enrichment
Motif enrichment
RcisTarget
Transcription factor binding enrichment
ChIP-enrichR
Expression ratios
<= -2.0
-0
>= 2.0

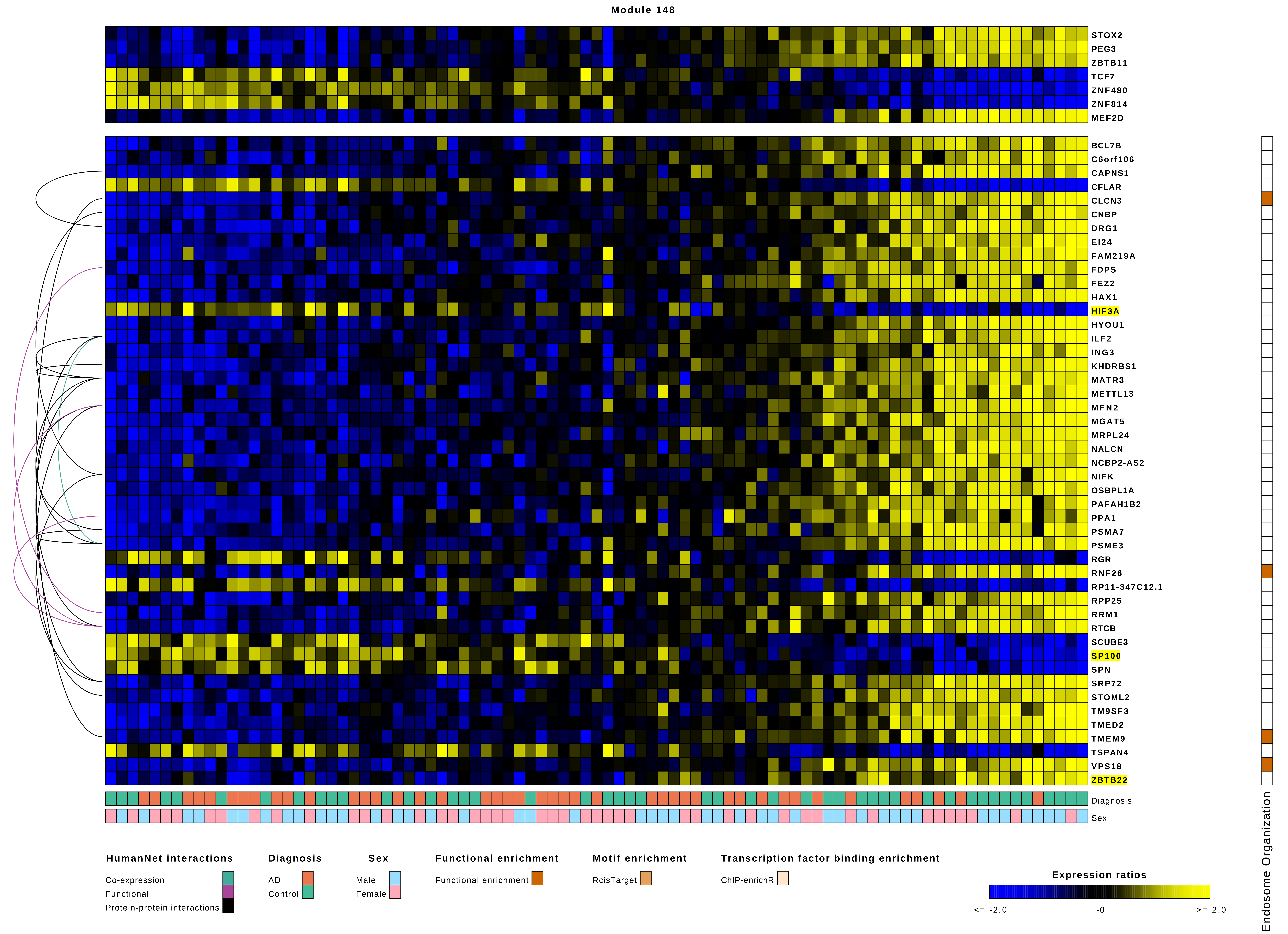
Module 148
STOX2
PEG3
ZBTB11
TCF7
ZNF480
ZNF814
MEF2D
BCL7B
C6orf106
CAPNS1
CFLAR
CLCN3
CNBP
DRG1
EI24
FAM219A
FDPS
FEZ2
HAX1
HIF3A
HYOU1
ILF2
ING3
KHDRBS1
MATR3
METTL13
MFN2
MGAT5
MRPL24
NALCN
NCBP2-AS2
NIFK
OSBPL1A
PAFAH1B2
PPA1
PSMA7
PSME3
RGR
RNF26
RP11-347C12.1
RPP25
RRM1
RTCB
SCUBE3
SP100
SPN
SRP72
STOML2
TM9SF3
TMED2
TMEM9
TSPAN4
VPS18
ZBTB22
Diagnosis
Sex
Endosome Organization
HumanNet interactions
Co-expression
Functional
Protein-protein interactions
Diagnosis
AD
Control
Sex
Male
Female
Functional enrichment
Functional enrichment
Motif enrichment
RcisTarget
Transcription factor binding enrichment
ChIP-enrichR
Expression ratios
<= -2.0
-0
>= 2.0

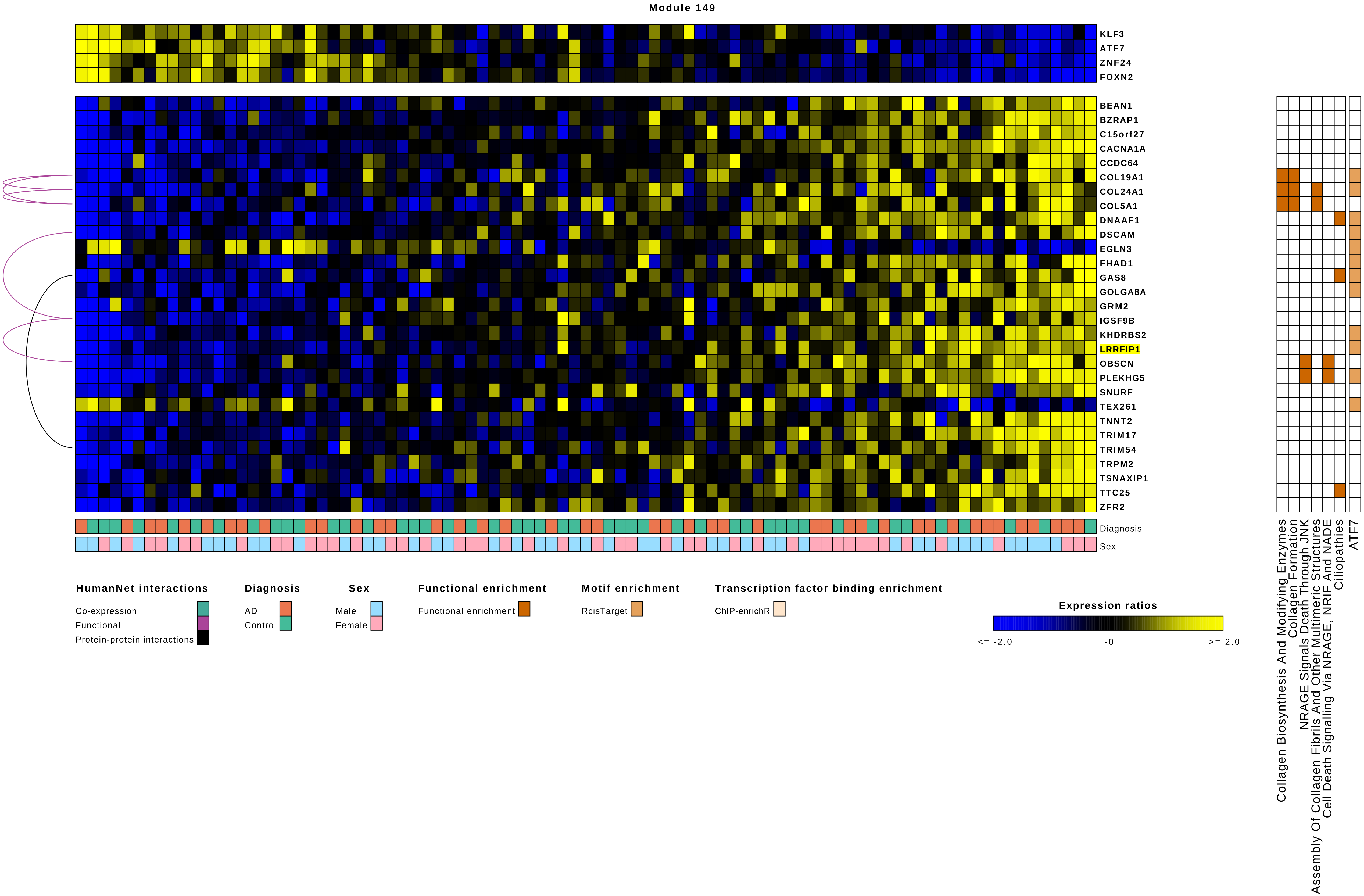

Module 149
KLF3
ATF7
ZNF24
FOXN2
BEAN1
BZRAP1
C15orf27
CACNA1A
CCDC64
COL19A1
COL24A1
COL5A1
DNAAF1
DSCAM
EGLN3
FHAD1
GAS8
GOLGA8A
GRM2
IGSF9B
KHDRBS2
LRRFIP1
OBSCN
PLEKHG5
SNURF
TEX261
TNNT2
TRIM17
TRIM54
TRPM2
TSNAXIP1
TTC25
ZFR2
Diagnosis
Sex
Collagen Biosynthesis And Modifying Enzymes
Collagen Formation
NRAGE Signals Death Through JNK
Assembly Of Collagen Fibrils And Other Multimeric Structures
Cell Death Signalling Via NRAGE, NRIF And NADE
Ciliopathies
ATF7
HumanNet interactions
Co-expression
Functional
Protein-protein interactions
Diagnosis
AD
Control
Sex
Male
Female
Functional enrichment
Functional enrichment
Motif enrichment
RcisTarget
Transcription factor binding enrichment
ChIP-enrichR
Expression ratios
<= -2.0
-0
>= 2.0

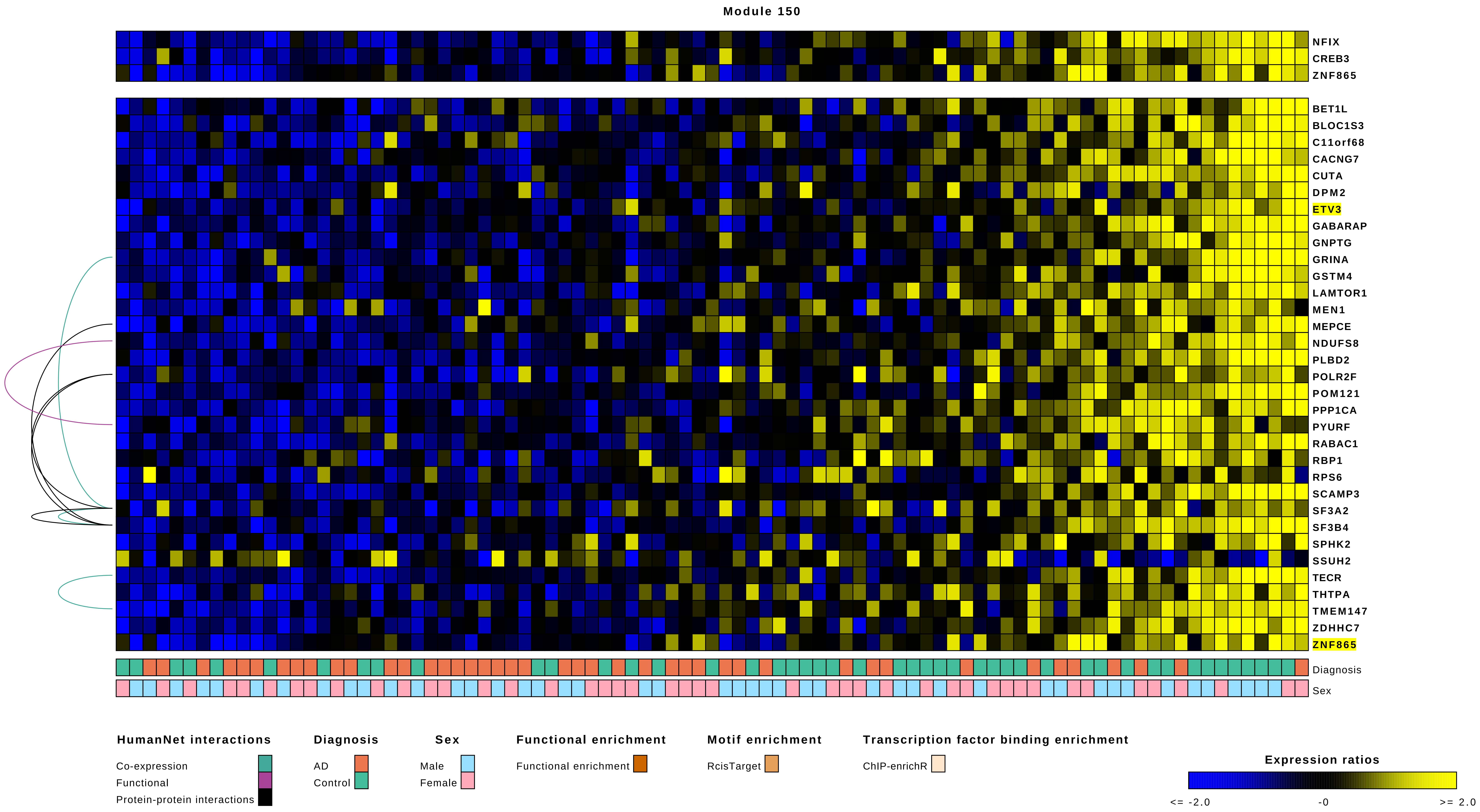

Module 150
NFIX
CREB3
ZNF865
BET1L
BLOC1S3
C11orf68
CACNG7
CUTA
DPM2
ETV3
GABARAP
GNPTG
GRINA
GSTM4
LAMTOR1
MEN1
MEPCE
NDUFS8
PLBD2
POLR2F
POM121
PPP1CA
PYURF
RABAC1
RBP1
RPS6
SCAMP3
SF3A2
SF3B4
SPHK2
SSUH2
TECR
THTPA
TMEM147
ZDHHC7
ZNF865
Diagnosis
Sex
HumanNet interactions
Co-expression
Functional
Protein-protein interactions
Diagnosis
AD
Control
Sex
Male
Female
Functional enrichment
Functional enrichment
Motif enrichment
RcisTarget
Transcription factor binding enrichment
ChIP-enrichR
Expression ratios
<= -2.0
-0
>= 2.0

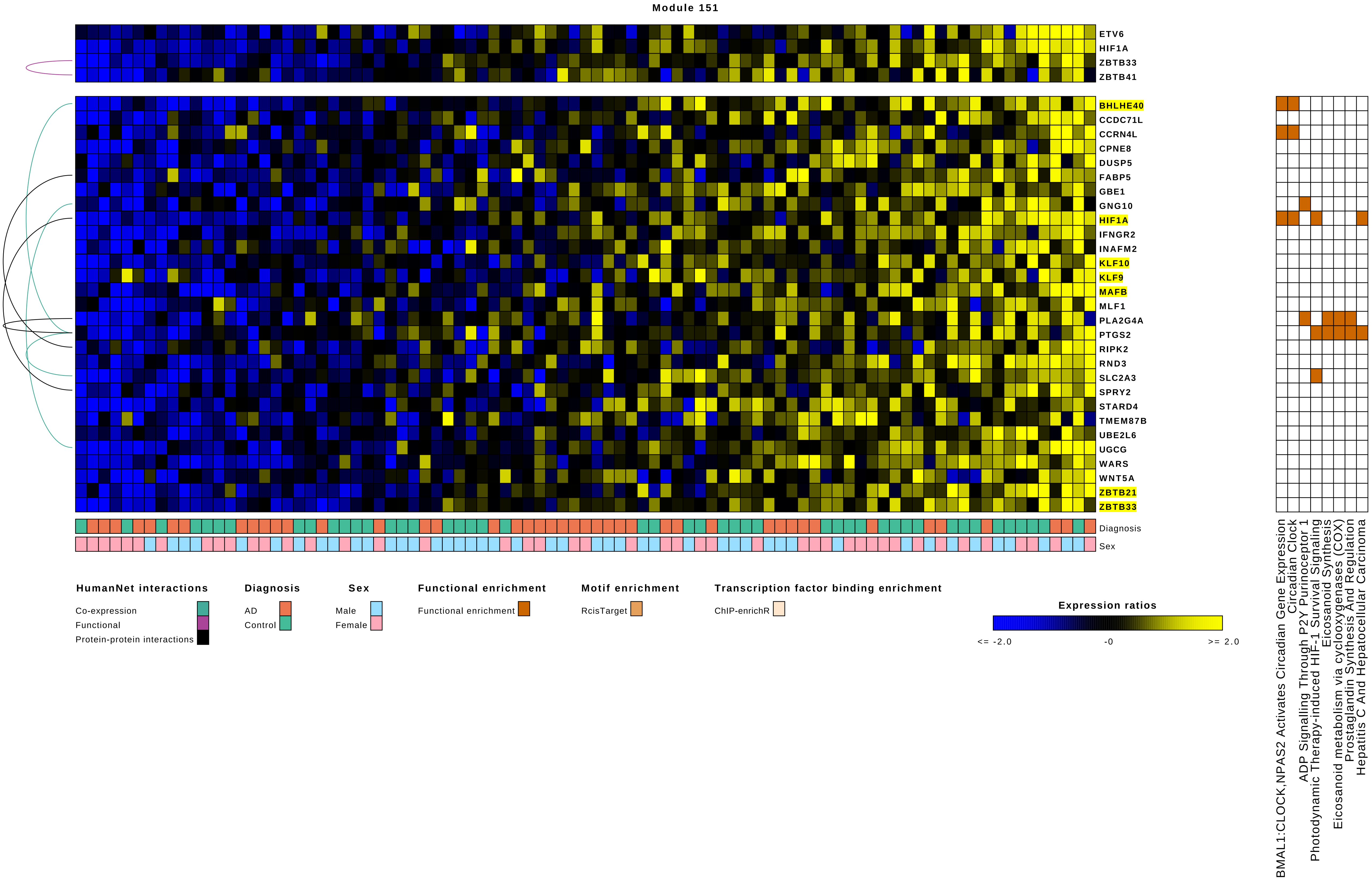
Module 151
ETV6
HIF1A
ZBTB33
ZBTB41
BHLHE40
CCDC71L
CCRN4L
CPNE8
DUSP5
FABP5
GBE1
GNG10
HIF1A
IFNGR2
INAFM2
KLF10
KLF9
MAFB
MLF1
PLA2G4A
PTGS2
RIPK2
RND3
SLC2A3
SPRY2
STARD4
TMEM87B
UBE2L6
UGCG
WARS
WNT5A
ZBTB21
ZBTB33
Diagnosis
Sex
BMAL1:CLOCK,NPAS2 Activates Circadian Gene Expression
Circadian Clock
ADP Signalling Through P2Y Purinoceptor 1
Photodynamic Therapy-induced HIF-1 Survival Signaling
Eicosanoid Synthesis
Eicosanoid metabolism via cyclooxygenases (COX)
Prostaglandin Synthesis And Regulation
Hepatitis C And Hepatocellular Carcinoma
HumanNet interactions
Co-expression
Functional
Protein-protein interactions
Diagnosis
AD
Control
Sex
Male
Female
Functional enrichment
Functional enrichment
Motif enrichment
RcisTarget
Transcription factor binding enrichment
ChIP-enrichR
Expression ratios
<= -2.0
-0
>= 2.0

Module 152

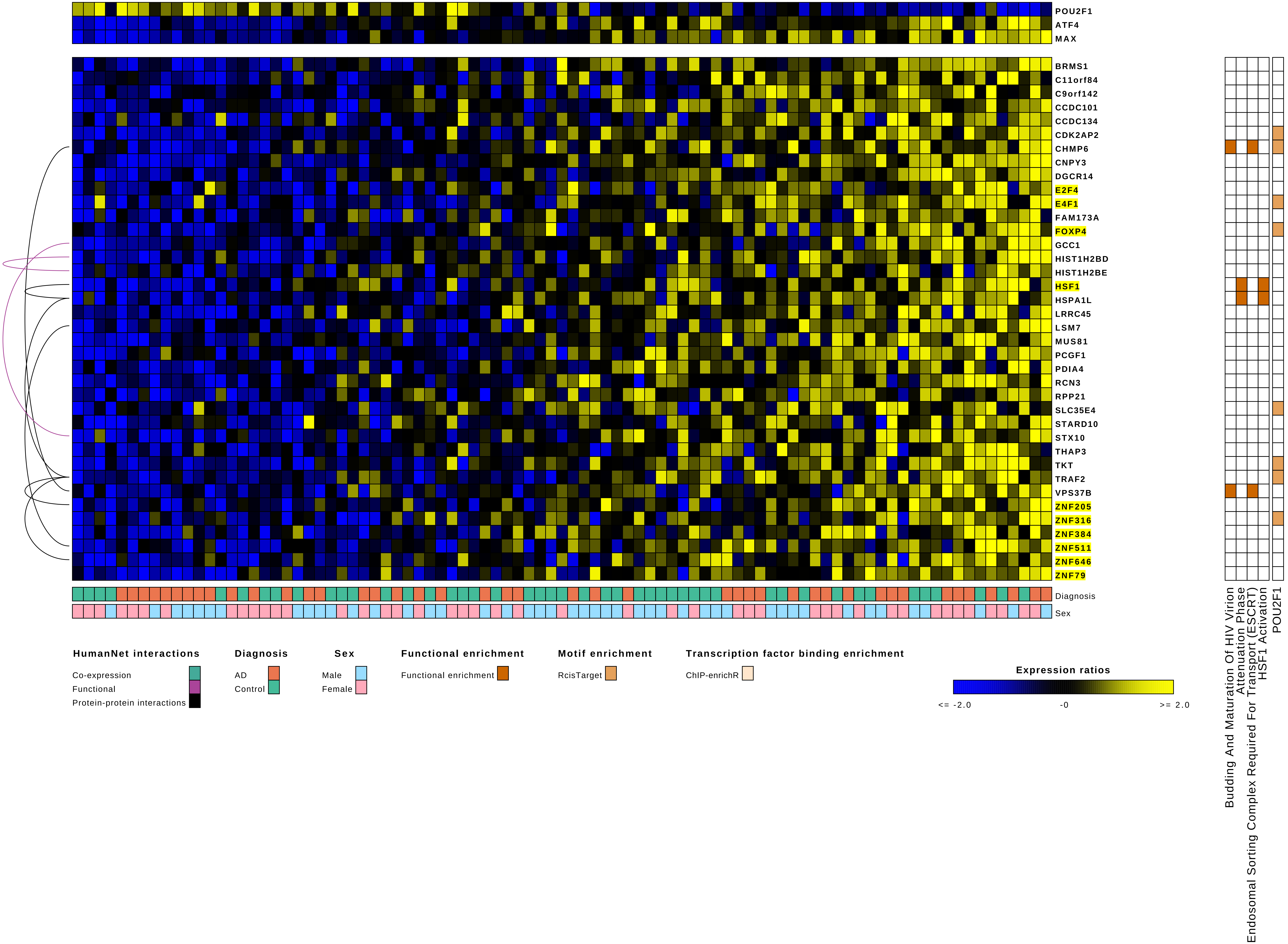
POU2F1
ATF4
MAX
BRMS1
C11orf84
C9orf142
CCDC101
CCDC134
CDK2AP2
CHMP6
CNPY3
DGCR14
E2F4
E4F1
FAM173A
FOXP4
GCC1
HIST1H2BD
HIST1H2BE
HSF1
HSPA1L
LRRC45
LSM7
MUS81
PCGF1
PDIA4
RCN3
RPP21
SLC35E4
STARD10
STX10
THAP3
TKT
TRAF2
VPS37B
ZNF205
ZNF316
ZNF384
ZNF511
ZNF646
ZNF79
Diagnosis
Sex
Budding And Maturation Of HIV Virion
Attenuation Phase
Endosomal Sorting Complex Required For Transport (ESCRT)
HSF1 Activation
POU2F1
HumanNet interactions
Co-expression
Functional
Protein-protein interactions
Diagnosis
AD
Control
Sex
Male
Female
Functional enrichment
Functional enrichment
Motif enrichment
RcisTarget
Transcription factor binding enrichment
ChIP-enrichR
Expression ratios
<= -2.0
-0
>= 2.0

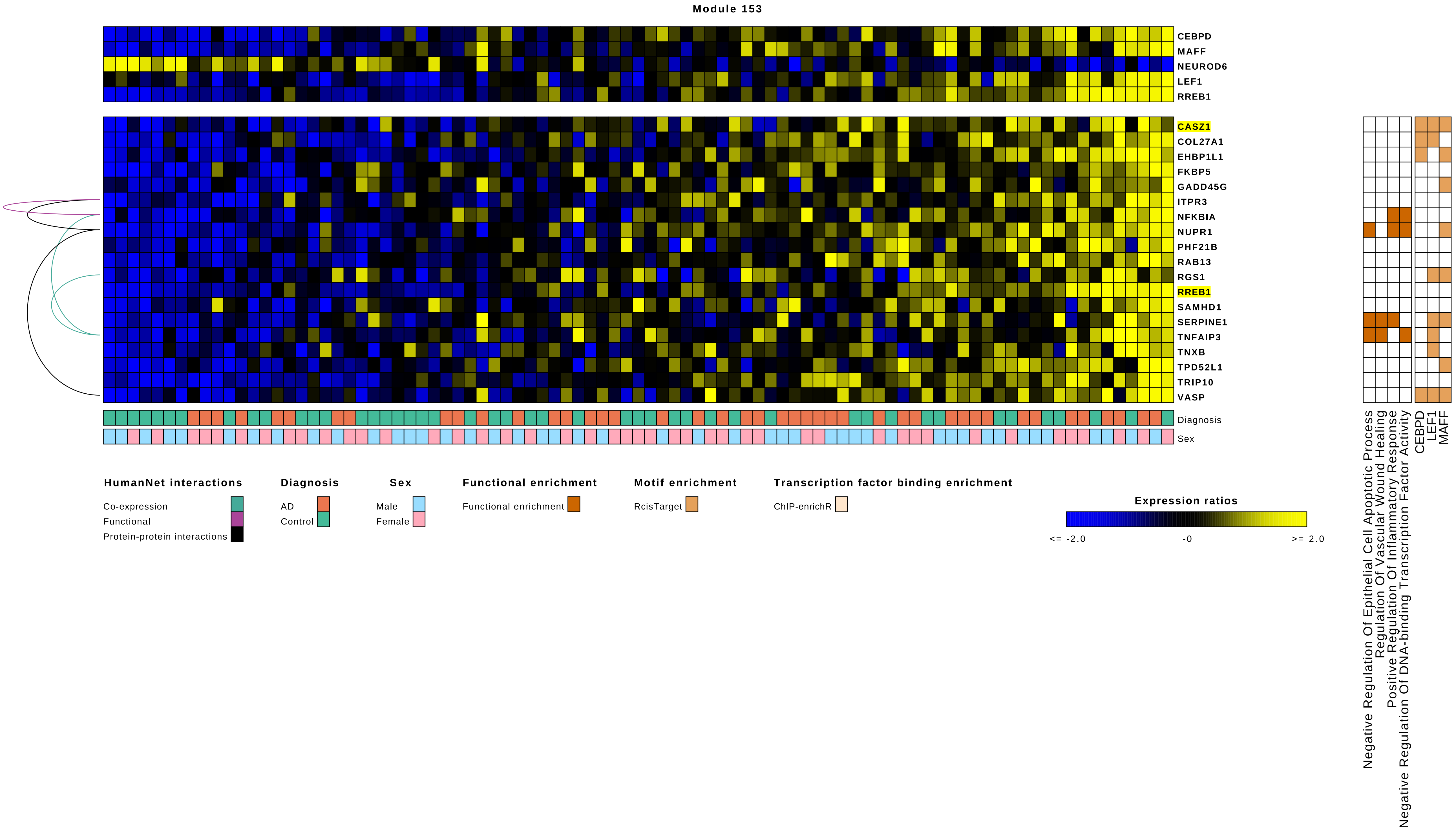
Module 153
CEBPD
MAFF
NEUROD6
LEF1
RREB1
CASZ1
COL27A1
EHBP1L1
FKBP5
GADD45G
ITPR3
NFKBIA
NUPR1
PHF21B
RAB13
RGS1
RREB1
SAMHD1
SERPINE1
TNFAIP3
TNXB
TPD52L1
TRIP10
VASP
Diagnosis
Sex
Negative Regulation Of Epithelial Cell Apoptotic Process
Regulation Of Vascular Wound Healing
Positive Regulation Of Inflammatory Response
Negative Regulation Of DNA-binding Transcription Factor Activity
CEBPD
LEF1
MAFF
HumanNet interactions
Co-expression
Functional
Protein-protein interactions
Diagnosis
AD
Control
Sex
Male
Female
Functional enrichment
Functional enrichment
Motif enrichment
RcisTarget
Transcription factor binding enrichment
ChIP-enrichR
Expression ratios
<= -2.0
-0
>= 2.0

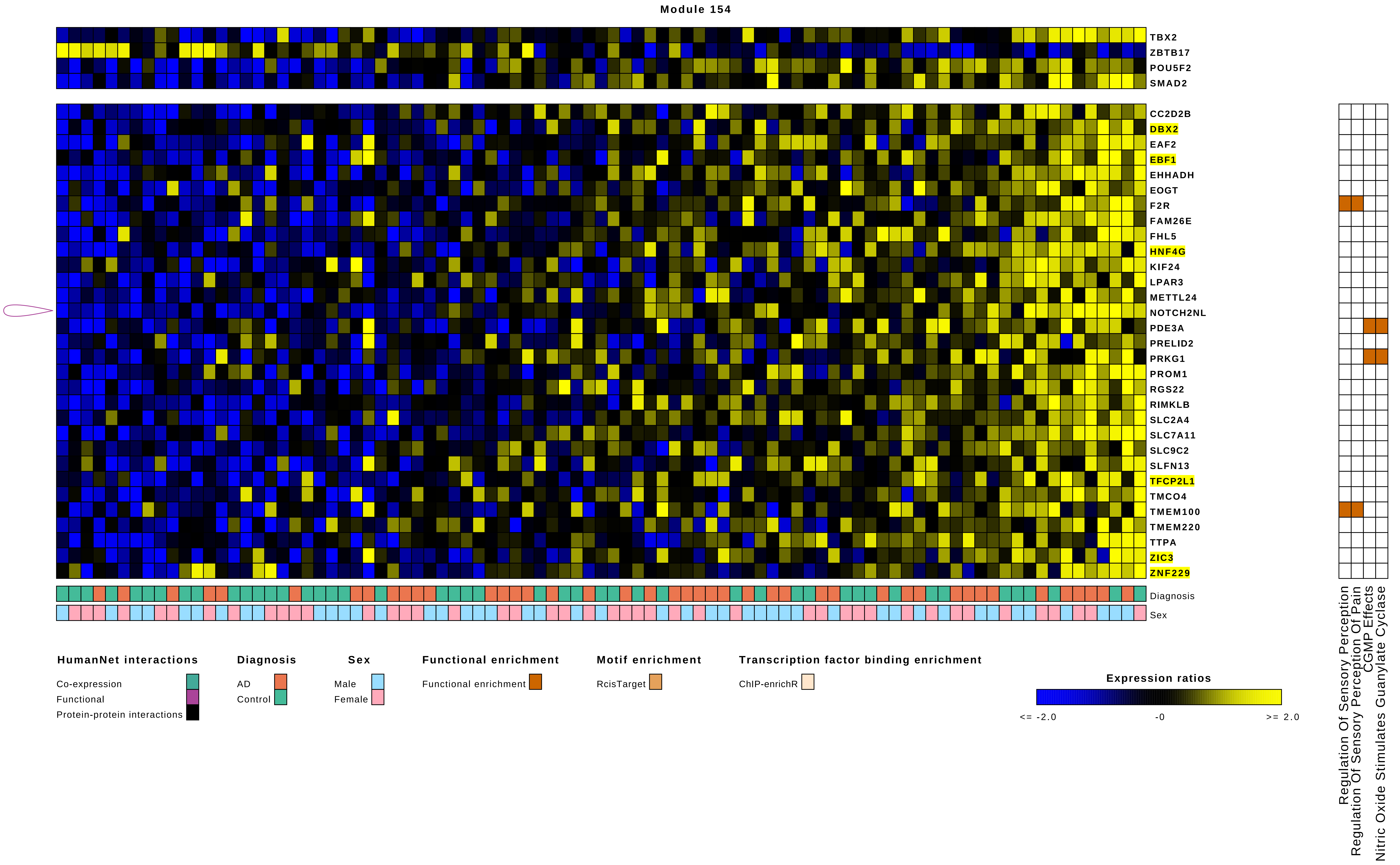
Module 154
TBX2
ZBTB17
POU5F2
SMAD2
CC2D2B
DBX2
EAF2
EBF1
EHHADH
EOGT
F2R
FAM26E
FHL5
HNF4G
KIF24
LPAR3
METTL24
NOTCH2NL
PDE3A
PRELID2
PRKG1
PROM1
RGS22
RIMKLB
SLC2A4
SLC7A11
SLC9C2
SLFN13
TFCP2L1
TMCO4
TMEM100
TMEM220
TTPA
ZIC3
ZNF229
Diagnosis
Sex
Regulation Of Sensory Perception
Regulation Of Sensory Perception Of Pain
CGMP Effects
Nitric Oxide Stimulates Guanylate Cyclase
HumanNet interactions
Co-expression
Functional
Protein-protein interactions
Diagnosis
AD
Control
Sex
Male
Female
Functional enrichment
Functional enrichment
Motif enrichment
RcisTarget
Transcription factor binding enrichment
ChIP-enrichR
Expression ratios
<= -2.0
-0
>= 2.0

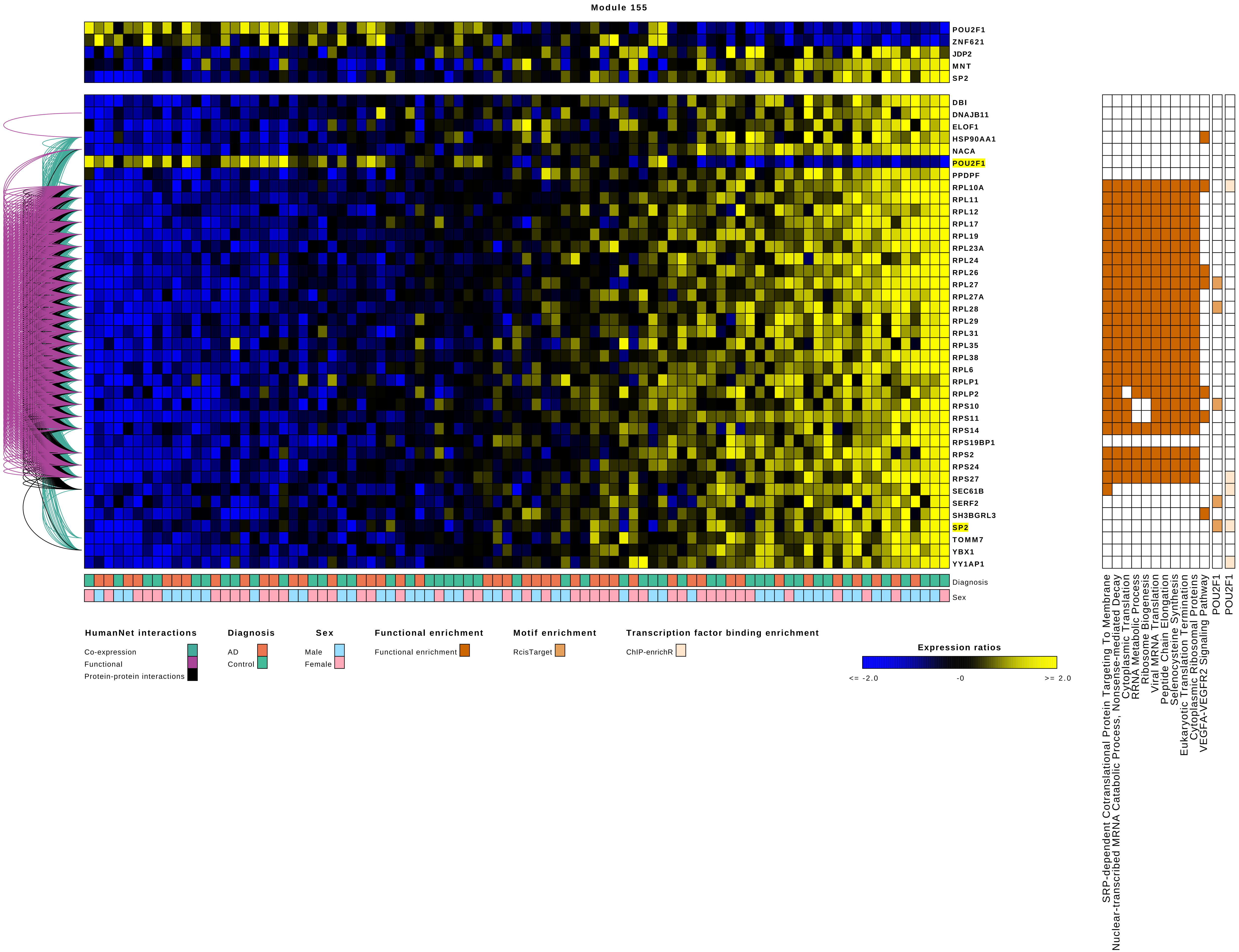
Module 155
POU2F1
ZNF621
JDP2
MNT
SP2
DBI
DNAJB11
ELOF1
HSP90AA1
NACA
POU2F1
PPDPF
RPL10A
RPL11
RPL12
RPL17
RPL19
RPL23A
RPL24
RPL26
RPL27
RPL27A
RPL28
RPL29
RPL31
RPL35
RPL38
RPL6
RPLP1
RPLP2
RPS10
RPS11
RPS14
RPS19BP1
RPS2
RPS24
RPS27
SEC61B
SERF2
SH3BGRL3
SP2
TOMM7
YBX1
YY1AP1
Diagnosis
Sex
HumanNet interactions
Co-expression
Functional
Protein-protein interactions
Diagnosis
AD
Control
Sex
Male
Female
Functional enrichment
Functional enrichment
Motif enrichment
RcisTarget
Transcription factor binding enrichment
ChIP-enrichR
Expression ratios
<= -2.0
-0
>= 2.0
SRP-dependent Cotranslational Protein Targeting To Membrane
Nuclear-transcribed MRNA Catabolic Process, Nonsense-mediated Decay
Cytoplasmic Translation
RRNA Metabolic Process
Ribosome Biogenesis
Viral MRNA Translation
Peptide Chain Elongation
Selenocysteine Synthesis
Eukaryotic Translation Termination
Cytoplasmic Ribosomal Proteins
VEGFA-VEGFR2 Signaling Pathway
POU2F1
POU2F1
