## Supplemental Figure 11 for "Comparative gene regulatory network analysis in Alzheimer’s disease and major depressive disorder identifies shared core regulatory circuits"

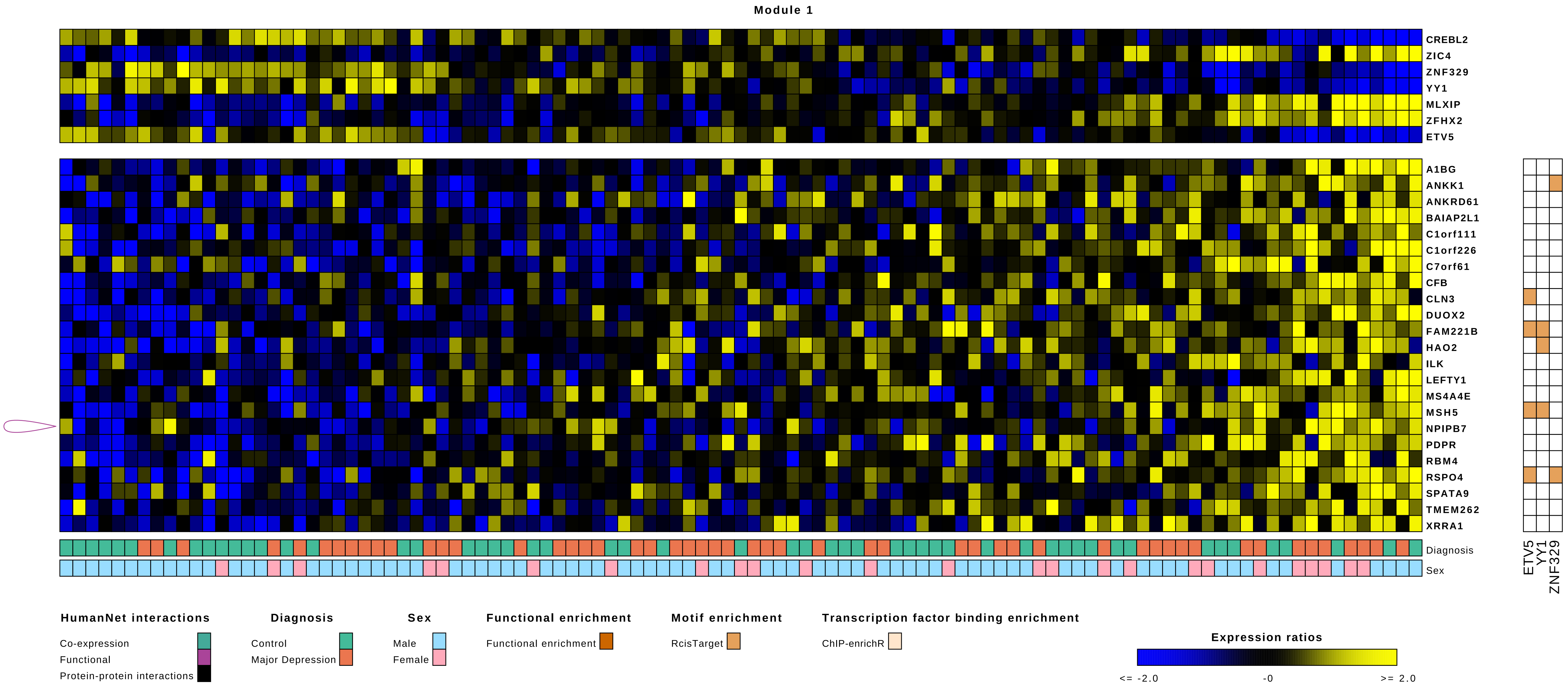
Module 1
CREBL2
ZIC4
ZNF329
YY1
MLXIP
ZFHX2
ETV5
A1BG
ANKK1
ANKRD61
BAIAP2L1
C1orf111
C1orf226
C7orf61
CFB
CLN3
DUOX2
FAM221B
HAO2
ILK
LEFTY1
MS4A4E
MSH5
NPIPB7
PDPR
RBM4
RSPO4
SPATA9
TMEM262
XRRA1
Diagnosis
Sex
ETV5
YY1
ZNF329
HumanNet interactions
Co-expression
Functional
Protein-protein interactions
Diagnosis
Control
Major Depression
Sex
Male
Female
Functional enrichment
Functional enrichment
Motif enrichment
RcisTarget
Transcription factor binding enrichment
ChIP-enrichR
Expression ratios
<= -2.0
-0
>= 2.0

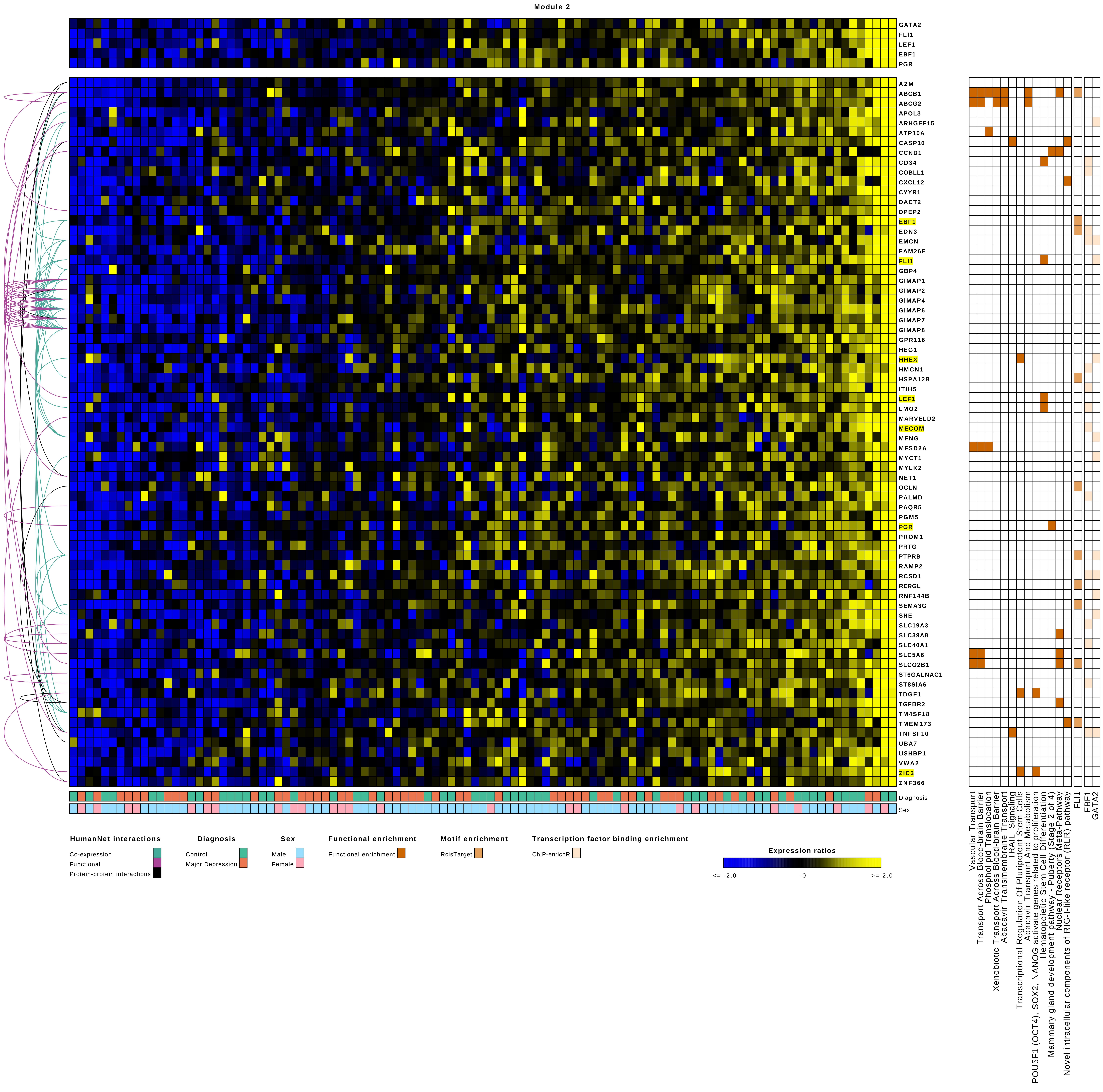
Module 2
GATA2
FLI1
LEF1
EBF1
PGR
A2M
ABCB1
ABCG2
APOL3
ARHGEF15
ATP10A
CASP10
CCND1
CD34
COBLL1
CXCL12
CYYR1
DACT2
DPEP2
EBF1
EDN3
EMCN
FAM26E
FLI1
GBP4
GIMAP1
GIMAP2
GIMAP4
GIMAP6
GIMAP7
GIMAP8
GPR116
HEG1
HHEX
HMCN1
HSPA12B
ITIH5
LEF1
LMO2
MARVELD2
MECOM
MFNG
MFSD2A
MYCT1
MYLK2
NET1
OCLN
PALMD
PAQR5
PGM5
PGR
PROM1
PRTG
PTPRB
RAMP2
RCSD1
RERGL
RNF144B
SEMA3G
SHE
SLC19A3
SLC39A8
SLC40A1
SLC5A6
SLCO2B1
ST6GALNAC1
ST8SIA6
TDGF1
TGFBR2
TM4SF18
TMEM173
TNFSF10
UBA7
USHBP1
VWA2
ZIC3
ZNF366
Diagnosis
Sex
Vascular Transport
Transport Across Blood-brain Barrier
Phospholipid Translocation
Xenobiotic Transport Across Blood-brain Barrier
Abacavir Transmembrane Transport
TRAIL_Signaling
Transcriptional Regulation Of Pluripotent Stem Cells
Abacavir Transport And Metabolism
POU5F1 (OCT4), SOX2, NANOG activate genes related to proliferation
Hematopoietic Stem Cell Differentiation
Mammary gland development pathway - Puberty (Stage 2 of 4)
Nuclear Receptors Meta-Pathway
Novel intracellular components of RIG-I-like receptor (RLR) pathway
FLI1
EBF1
GATA2
HumanNet interactions
Co-expression
Functional
Protein-protein interactions
Diagnosis
Control
Major Depression
Sex
Male
Female
Functional enrichment
Functional enrichment
Motif enrichment
RcisTarget
Transcription factor binding enrichment
ChIP-enrichR
Expression ratios
<= -2.0
-0
>= 2.0

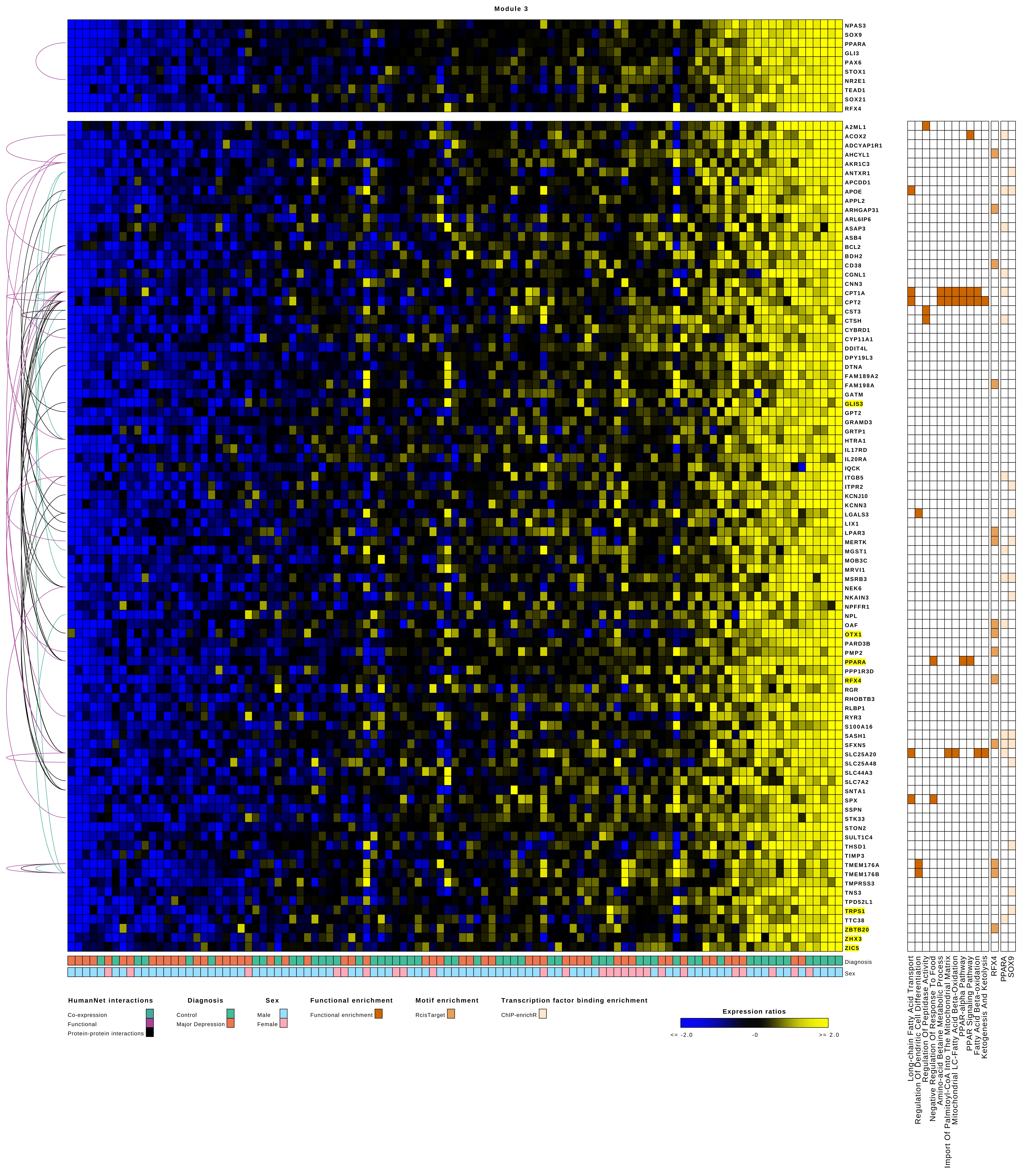
Module 3
NPAS3
SOX9
PPARA
GLI3
PAX6
STOX1
NR2E1
TEAD1
SOX21
RFX4
A2ML1
ACOX2
ADCYAP1R1
AHCYL1
AKR1C3
ANTXR1
APCDD1
APOE
APPL2
ARHGAP31
ARL6IP6
ASAP3
ASB4
BCL2
BDH2
CD38
CGNL1
CNN3
CPT1A
CPT2
CST3
CTSH
CYBRD1
CYP11A1
DDIT4L
DPY19L3
DTNA
FAM189A2
FAM198A
GATM
GLIS3
GPT2
GRAMD3
GRTP1
HTRA1
IL17RD
IL20RA
IQCK
ITGB5
ITPR2
KCNJ10
KCNN3
LGALS3
LIX1
LPAR3
MERTK
MGST1
MOB3C
MRVI1
MSRB3
NEK6
NKAIN3
NPFFR1
NPL
OAF
OTX1
PARD3B
PMP2
PPARA
PPP1R3D
RFX4
RGR
RHOBTB3
RLBP1
RYR3
S100A16
SASH1
SFXN5
SLC25A20
SLC25A48
SLC44A3
SLC7A2
SNTA1
SPX
SSPN
STK33
STON2
SULT1C4
THSD1
TIMP3
TMEM176A
TMEM176B
TMPRSS3
TNS3
TPD52L1
TRPS1
TTC38
ZBTB20
ZHX3
ZIC5
Diagnosis
Sex
Long-chain Fatty Acid Transport
Regulation Of Dendritic Cell Differentiation
Regulation Of Peptidase Activity
Negative Regulation Of Response To Food
Amino-acid Betaine Metabolic Process
Import Of Palmitoyl-CoA Into The Mitochondrial Matrix
Mitochondrial LC-Fatty Acid Beta-Oxidation
PPAR-alpha Pathway
PPAR Signaling Pathway
Fatty Acid Beta-oxidation
Ketogenesis And Ketolysis
RFX4
PPARA
SOX9
HumanNet interactions
Co-expression
Functional
Protein-protein interactions
Diagnosis
Control
Major Depression
Sex
Male
Female
Functional enrichment
Functional enrichment
Motif enrichment
RcisTarget
Transcription factor binding enrichment
ChIP-enrichR
Expression ratios
<= -2.0
-0
>= 2.0

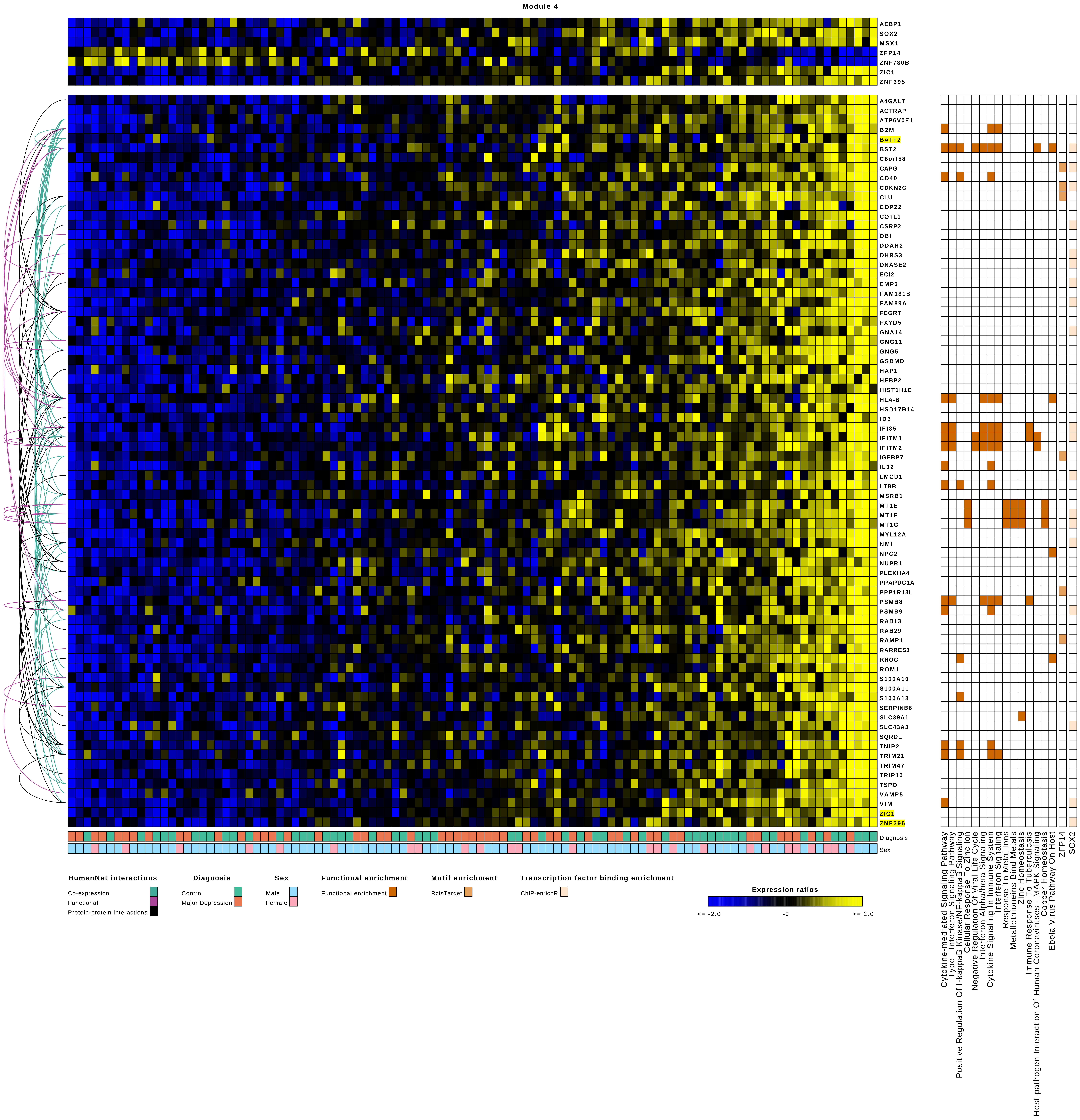

Module 4
AEBP1
SOX2
MSX1
ZFP14
ZNF780B
ZIC1
ZNF395
A4GALT
AGTRAP
ATP6V0E1
B2M
BATF2
BST2
C8orf58
CAPG
CD40
CDKN2C
CLU
COPZ2
COTL1
CSRP2
DBI
DDAH2
DHRS3
DNASE2
ECI2
EMP3
FAM181B
FAM89A
FCGRT
FXYD5
GNA14
GNG11
GNG5
GSDMD
HAP1
HEBP2
HIST1H1C
HLA-B
HSD17B14
ID3
IFI35
IFITM1
IFITM2
IGFBP7
IL32
LMCD1
LTBR
MSRB1
MT1E
MT1F
MT1G
MYL12A
NMI
NPC2
NUPR1
PLEKHA4
PPAPDC1A
PPP1R13L
PSMB8
PSMB9
RAB13
RAB29
RAMP1
RARRES3
RHOC
ROM1
S100A10
S100A11
S100A13
SERPINB6
SLC39A1
SLC43A3
SQRDL
TNIP2
TRIM21
TRIM47
TRIP10
TSPO
VAMP5
VIM
ZIC1
ZNF395
Diagnosis
Sex
Cytokine-mediated Signaling Pathway
Type I Interferon Signaling Pathway
Positive Regulation Of I-kappaB Kinase/NF-kappaB Signaling
Cellular Response To Zinc Ion
Negative Regulation Of Viral Life Cycle
Interferon Alpha/beta Signaling
Cytokine Signaling In Immune System
Interferon Signaling
Response To Metal Ions
Metallothioneins Bind Metals
Zinc Homeostasis
Immune Response To Tuberculosis
Host-pathogen Interaction Of Human Coronaviruses - MAPK Signaling
Copper Homeostasis
Ebola Virus Pathway On Host
ZFP14
SOX2
HumanNet interactions
Co-expression
Functional
Protein-protein interactions
Diagnosis
Control
Major Depression
Sex
Male
Female
Functional enrichment
Functional enrichment
Motif enrichment
RcisTarget
Transcription factor binding enrichment
ChIP-enrichR
Expression ratios
<= -2.0
-0
>= 2.0

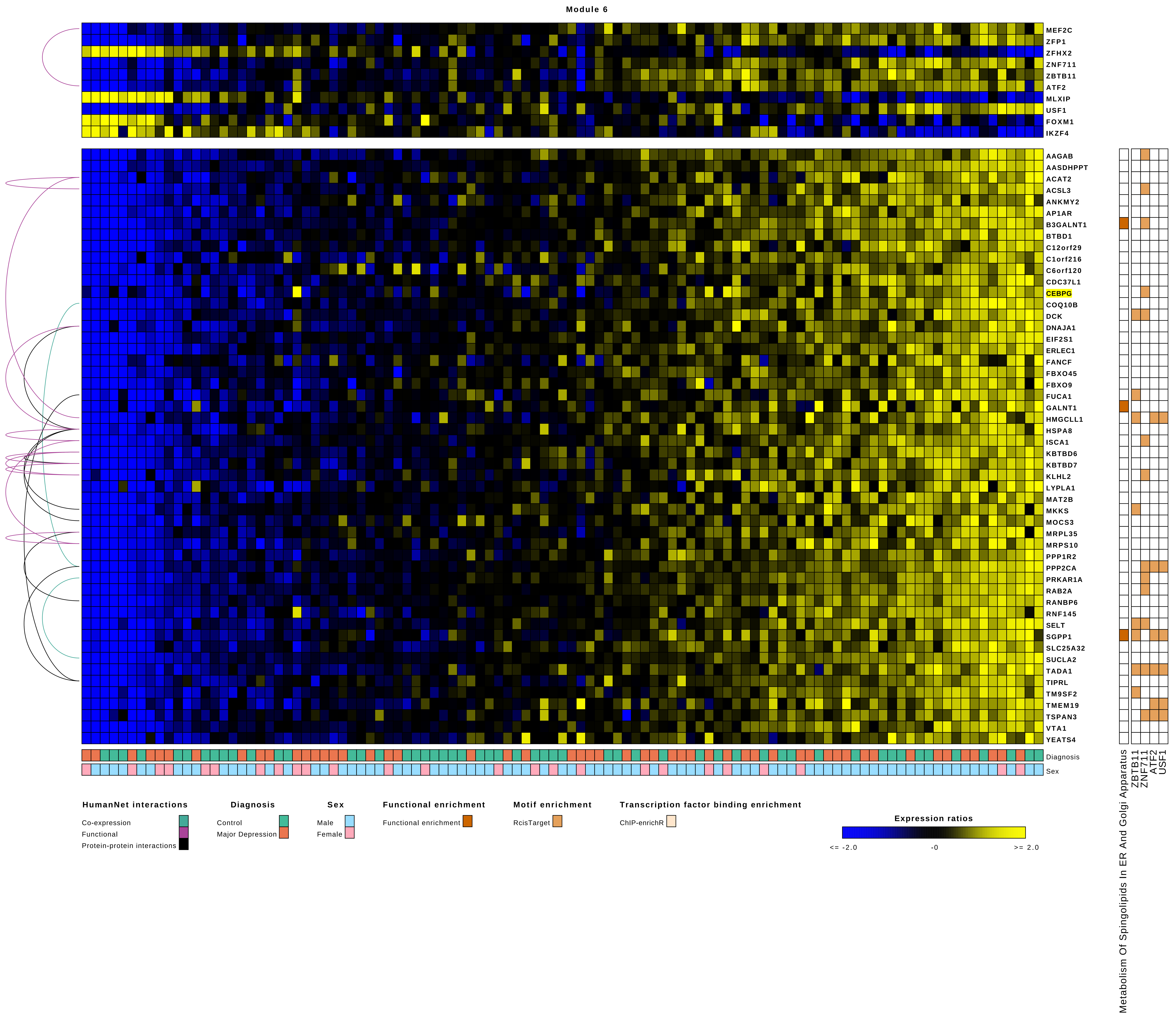

Module 6
MEF2C
ZFP1
ZFHX2
ZNF711
ZBTB11
ATF2
MLXIP
USF1
FOXM1
IKZF4
AAGAB
AASDHPPT
ACAT2
ACSL3
ANKMY2
AP1AR
B3GALNT1
BTBD1
C12orf29
C1orf216
C6orf120
CDC37L1
CEBPG
COQ10B
DCK
DNAJA1
EIF2S1
ERLEC1
FANCF
FBXO45
FBXO9
FUCA1
GALNT1
HMGCLL1
HSPA8
ISCA1
KBTBD6
KBTBD7
KLHL2
LYPLA1
MAT2B
MKKS
MOCS3
MRPL35
MRPS10
PPP1R2
PPP2CA
PRKAR1A
RAB2A
RANBP6
RNF145
SELT
SGPP1
SLC25A32
SUCLA2
TADA1
TIPRL
TM9SF2
TMEM19
TSPAN3
VTA1
YEATS4
Diagnosis
Sex
Metabolism Of Spingolipids In ER And Golgi Apparatus
ZBTB11
ZNF711
ATF2
USF1
HumanNet interactions
Co-expression
Functional
Protein-protein interactions
Diagnosis
Control
Major Depression
Sex
Male
Female
Functional enrichment
Functional enrichment
Motif enrichment
RcisTarget
Transcription factor binding enrichment
ChIP-enrichR
Expression ratios
<= -2.0
-0
>= 2.0

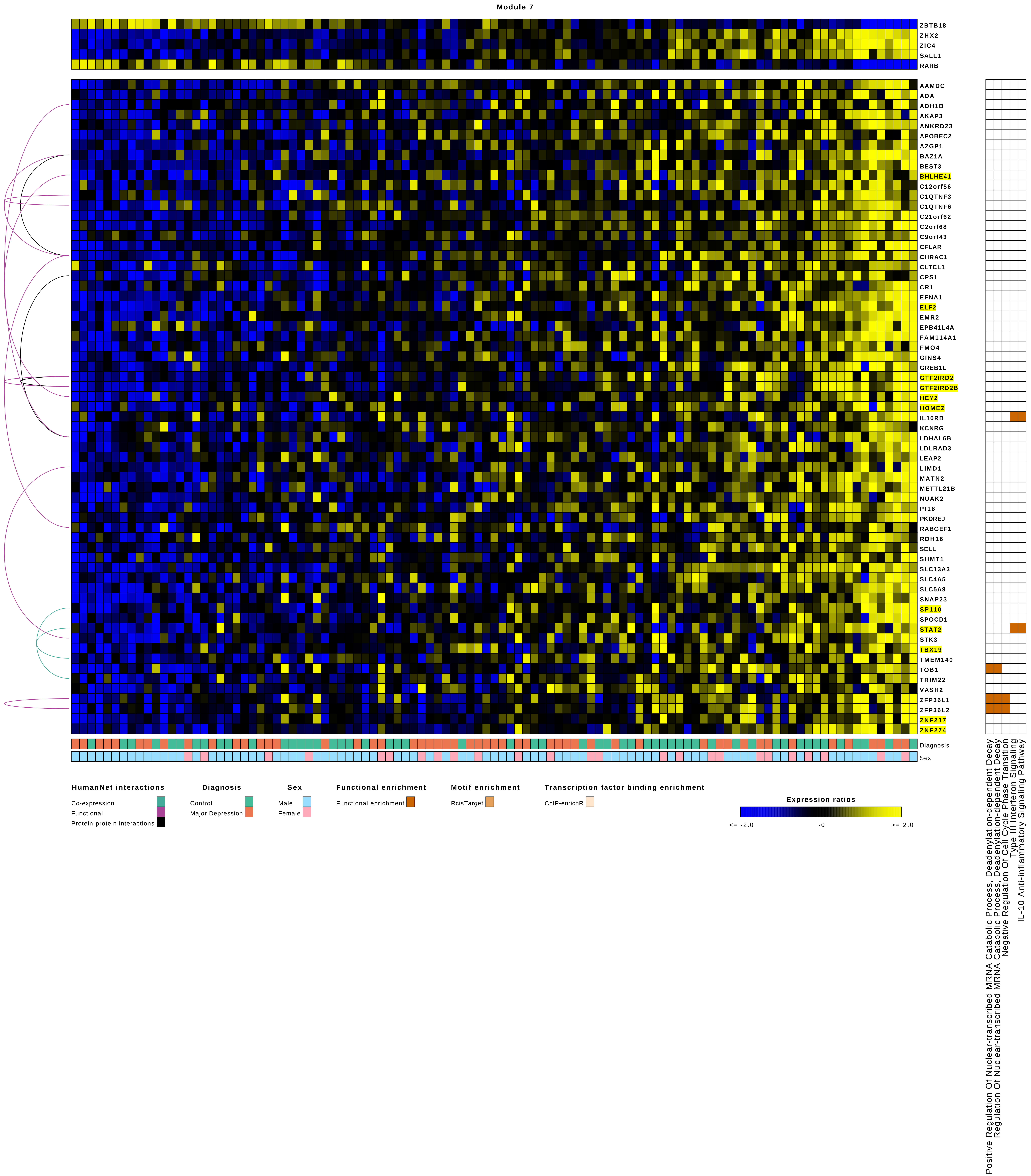

Module 7
ZBTB18
ZHX2
ZIC4
SALL1
RARB
AAMDC
ADA
ADH1B
AKAP3
ANKRD23
APOBEC2
AZGP1
BAZ1A
BEST3
BHLHE41
C12orf56
C1QTNF3
C1QTNF6
C21orf62
C2orf68
C9orf43
CFLAR
CHRAC1
CLTCL1
CPS1
CR1
EFNA1
ELF2
EMR2
EPB41L4A
FAM114A1
FMO4
GINS4
GREB1L
GTF2IRD2
GTF2IRD2B
HEY2
HOMEZ
IL10RB
KCNRG
LDHAL6B
LDLRAD3
LEAP2
LIMD1
MATN2
METTL21B
NUAK2
PI16
PKDREJ
RABGEF1
RDH16
SELL
SHMT1
SLC13A3
SLC4A5
SLC5A9
SNAP23
SP110
SPOCD1
STAT2
STK3
TBX19
TMEM140
TOB1
TRIM22
VASH2
ZFP36L1
ZFP36L2
ZNF217
ZNF274
Diagnosis
Sex
Positive Regulation Of Nuclear-transcribed MRNA Catabolic Process, Deadenylation-dependent Decay
Regulation Of Nuclear-transcribed MRNA Catabolic Process, Deadenylation-dependent Decay
Negative Regulation Of Cell Cycle Phase Transition
Type III Interferon Signaling
IL-10 Anti-inflammatory Signaling Pathway
HumanNet interactions
Co-expression
Functional
Protein-protein interactions
Diagnosis
Control
Major Depression
Sex
Male
Female
Functional enrichment
Functional enrichment
Motif enrichment
RcisTarget
Transcription factor binding enrichment
ChIP-enrichR
Expression ratios
<= -2.0
-0
>= 2.0

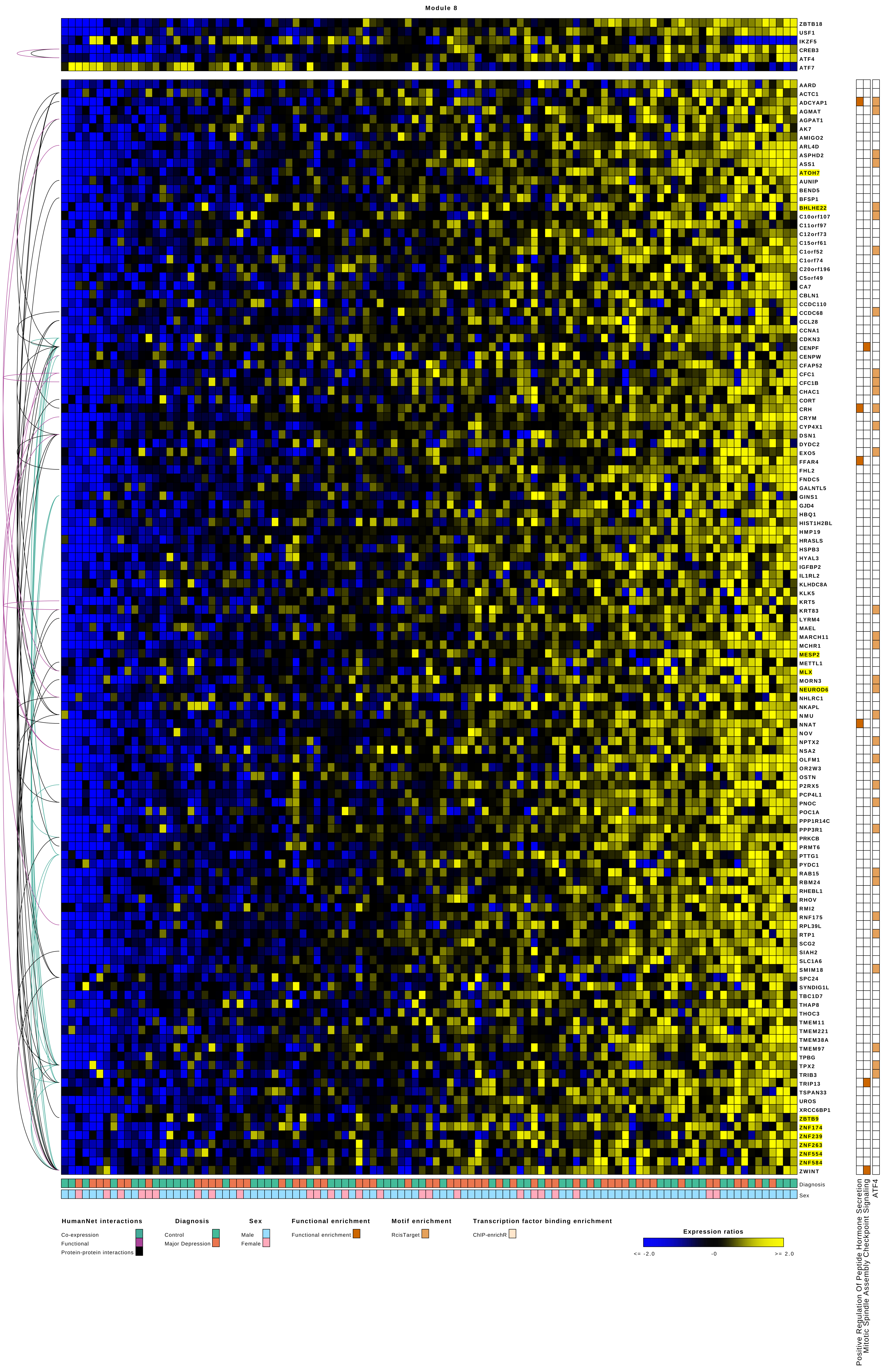

Module 8
ZBTB18
USF1
IKZF5
CREB3
ATF4
ATF7
AARD
ACTC1
ADCYAP1
AGMAT
AGPAT1
AK7
AMIGO2
ARL4D
ASPHD2
ASS1
ATOH7
AUNIP
BEND5
BFSP1
BHLHE22
C10orf107
C11orf97
C12orf73
C15orf61
C1orf52
C1orf74
C20orf196
C5orf49
CA7
CBLN1
CCDC110
CCDC68
CCL28
CCNA1
CDKN3
CENPF
CENPW
CFAP52
CFC1
CFC1B
CHAC1
CORT
CRH
CRYM
CYP4X1
DSN1
DYDC2
EXO5
FFAR4
FHL2
FNDC5
GALNTL5
GINS1
GJD4
HBQ1
HIST1H2BL
HMP19
HRASLS
HSPB3
HYAL3
IGFBP2
IL1RL2
KLHDC8A
KLK5
KRT5
KRT83
LYRM4
MAEL
MARCH11
MCHR1
MESP2
METTL1
MLX
MORN3
NEUROD6
NHLRC1
NKAPL
NMU
NNAT
NOV
NPTX2
NSA2
OLFM1
OR2W3
OSTN
P2RX5
PCP4L1
PNOC
POC1A
PPP1R14C
PPP3R1
PRKCB
PRMT6
PTTG1
PYDC1
RAB15
RBM24
RHEBL1
RHOV
RMI2
RNF175
RPL39L
RTP1
SCG2
SIAH2
SLC1A6
SMIM18
SPC24
SYNDIG1L
TBC1D7
THAP8
THOC3
TMEM11
TMEM221
TMEM38A
TMEM97
TPBG
TPX2
TRIB3
TRIP13
TSPAN33
UROS
XRCC6BP1
ZBTB9
ZNF174
ZNF239
ZNF263
ZNF554
ZNF584
ZWINT
Diagnosis
Sex
Positive Regulation Of Peptide Hormone Secretion
Mitotic Spindle Assembly Checkpoint Signaling
ATF4
HumanNet interactions
Co-expression
Functional
Protein-protein interactions
Diagnosis
Control
Major Depression
Sex
Male
Female
Functional enrichment
Functional enrichment
Motif enrichment
RcisTarget
Transcription factor binding enrichment
ChIP-enrichR
Expression ratios
<= -2.0
-0
>= 2.0

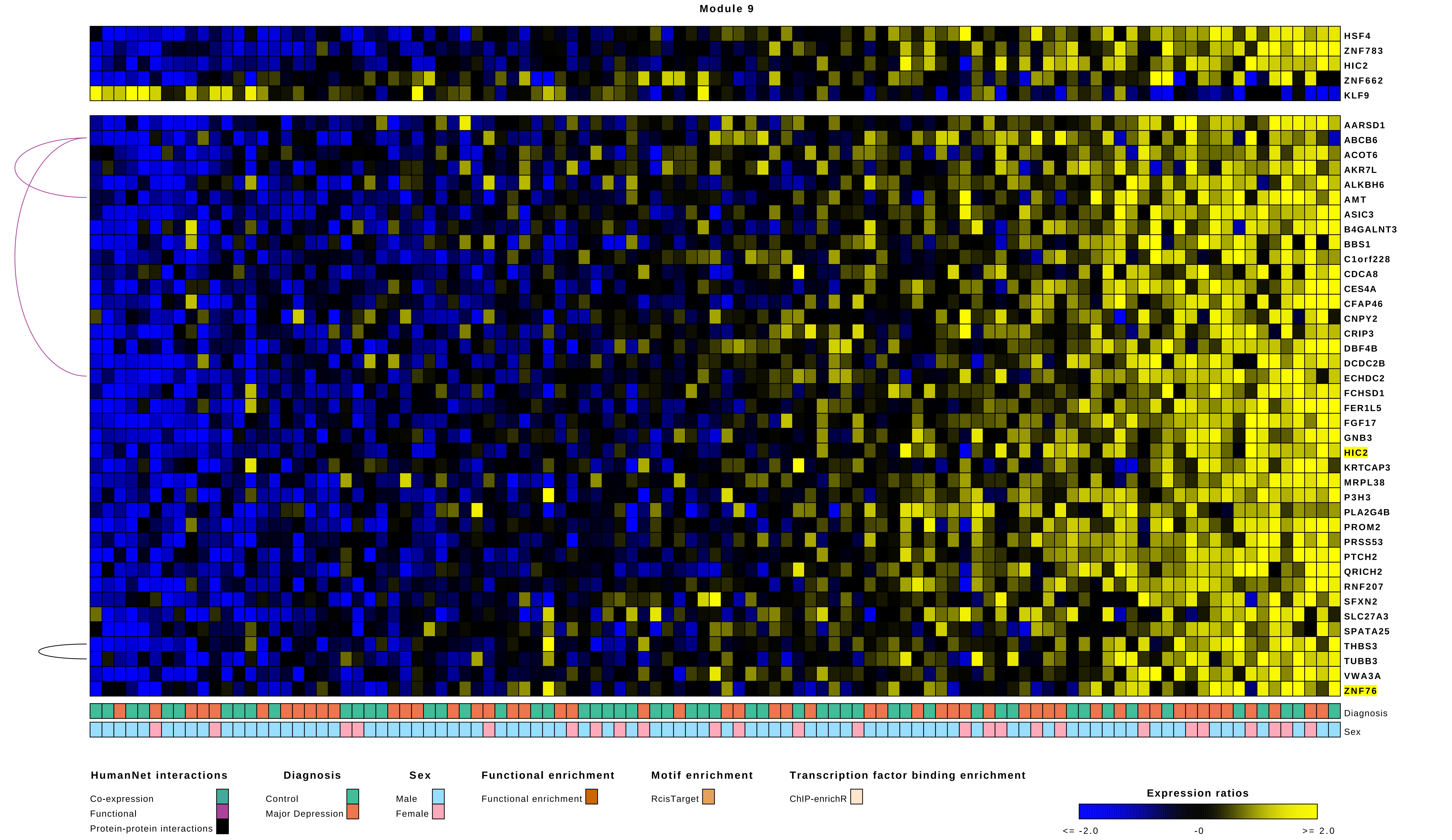
Module 9
HSF4
ZNF783
HIC2
ZNF662
KLF9
AARSD1
ABCB6
ACOT6
AKR7L
ALKBH6
AMT
ASIC3
B4GALNT3
BBS1
C1orf228
CDCA8
CES4A
CFAP46
CNPY2
CRIP3
DBF4B
DCDC2B
ECHDC2
FCHSD1
FER1L5
FGF17
GNB3
HIC2
KRTCAP3
MRPL38
P3H3
PLA2G4B
PROM2
PRSS53
PTCH2
QRICH2
RNF207
SFXN2
SLC27A3
SPATA25
THBS3
TUBB3
VWA3A
ZNF76
Diagnosis
Sex
HumanNet interactions
Co-expression
Functional
Protein-protein interactions
Diagnosis
Control
Major Depression
Sex
Male
Female
Functional enrichment
Functional enrichment
Motif enrichment
RcisTarget
Transcription factor binding enrichment
ChIP-enrichR
Expression ratios
<= -2.0
-0
>= 2.0

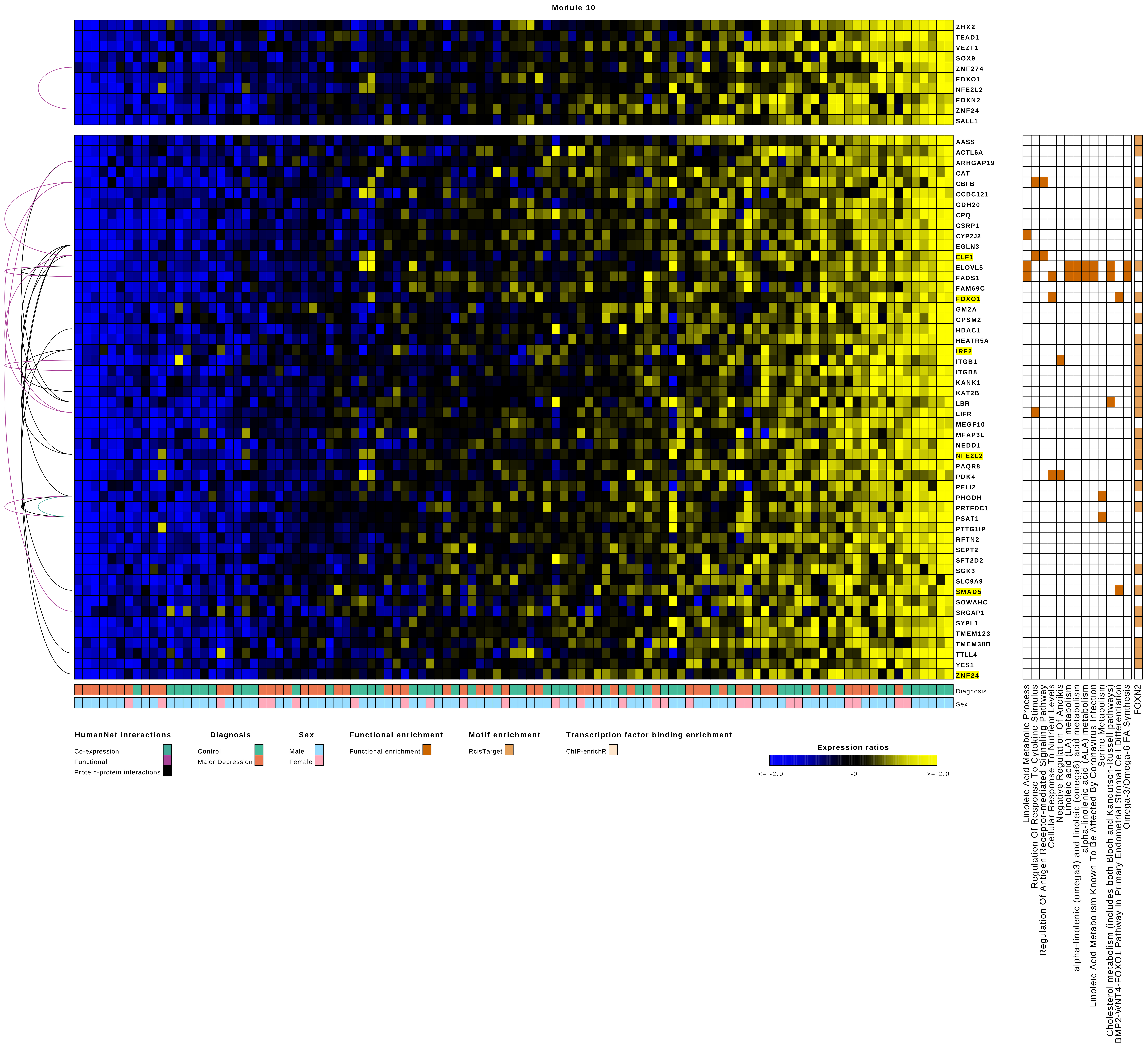
Module 10
ZHX2
TEAD1
VEZF1
SOX9
ZNF274
FOXO1
NFE2L2
FOXN2
ZNF24
SALL1
AASS
ACTL6A
ARHGAP19
CAT
CBFB
CCDC121
CDH20
CPQ
CSRP1
CYP2J2
EGLN3
ELF1
ELOVL5
FADS1
FAM69C
FOXO1
GM2A
GPSM2
HDAC1
HEATR5A
IRF2
ITGB1
ITGB8
KANK1
KAT2B
LBR
LIFR
MEGF10
MFAP3L
NEDD1
NFE2L2
PAQR8
PDK4
PELI2
PHGDH
PRTFDC1
PSAT1
PTTG1IP
RFTN2
SEPT2
SFT2D2
SGK3
SLC9A9
SMAD5
SOWAHC
SRGAP1
SYPL1
TMEM123
TMEM38B
TTLL4
YES1
ZNF24
Diagnosis
Sex
Linoleic Acid Metabolic Process
Regulation Of Response To Cytokine Stimulus
Regulation Of Antigen Receptor-mediated Signaling Pathway
Cellular Response To Nutrient Levels
Negative Regulation Of Anoikis
Linoleic acid (LA) metabolism
alpha-linolenic (omega3) and linoleic (omega6) acid metabolism
alpha-linolenic acid (ALA) metabolism
Linoleic Acid Metabolism Known To Be Affected By Coronavirus Infection
Serine Metabolism
Cholesterol metabolism (includes both Bloch and Kandutsch-Russell pathways)
BMP2-WNT4-FOXO1 Pathway In Primary Endometrial Stromal Cell Differentiation
Omega-3/Omega-6 FA Synthesis
FOXN2
HumanNet interactions
Co-expression
Functional
Protein-protein interactions
Diagnosis
Control
Major Depression
Sex
Male
Female
Functional enrichment
Functional enrichment
Motif enrichment
RcisTarget
Transcription factor binding enrichment
ChIP-enrichR
Expression ratios
<= -2.0
-0
>= 2.0

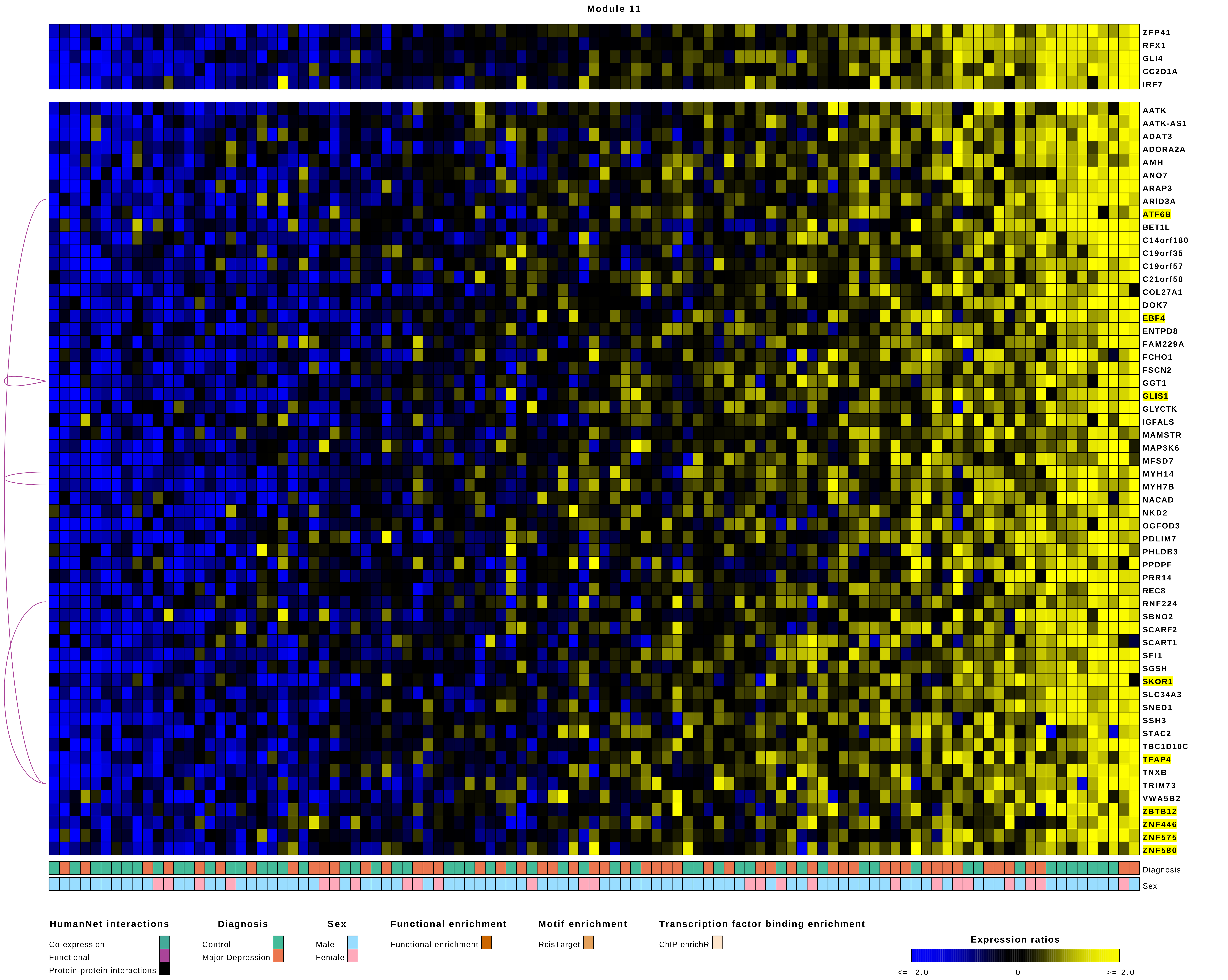
Module 11
ZFP41
RFX1
GLI4
CC2D1A
IRF7
AATK
AATK-AS1
ADAT3
ADORA2A
AMH
ANO7
ARAP3
ARID3A
ATF6B
BET1L
C14orf180
C19orf35
C19orf57
C21orf58
COL27A1
DOK7
EBF4
ENTPD8
FAM229A
FCHO1
FSCN2
GGT1
GLIS1
GLYCTK
IGFALS
MAMSTR
MAP3K6
MFSD7
MYH14
MYH7B
NACAD
NKD2
OGFOD3
PDLIM7
PHLDB3
PPDPF
PRR14
REC8
RNF224
SBNO2
SCARF2
SCART1
SFI1
SGSH
SKOR1
SLC34A3
SNED1
SSH3
STAC2
TBC1D10C
TFAP4
TNXB
TRIM73
VWA5B2
ZBTB12
ZNF446
ZNF575
ZNF580
Diagnosis
Sex
HumanNet interactions
Co-expression
Functional
Protein-protein interactions
Diagnosis
Control
Major Depression
Sex
Male
Female
Functional enrichment
Functional enrichment
Motif enrichment
RcisTarget
Transcription factor binding enrichment
ChIP-enrichR
Expression ratios
<= -2.0
-0
>= 2.0

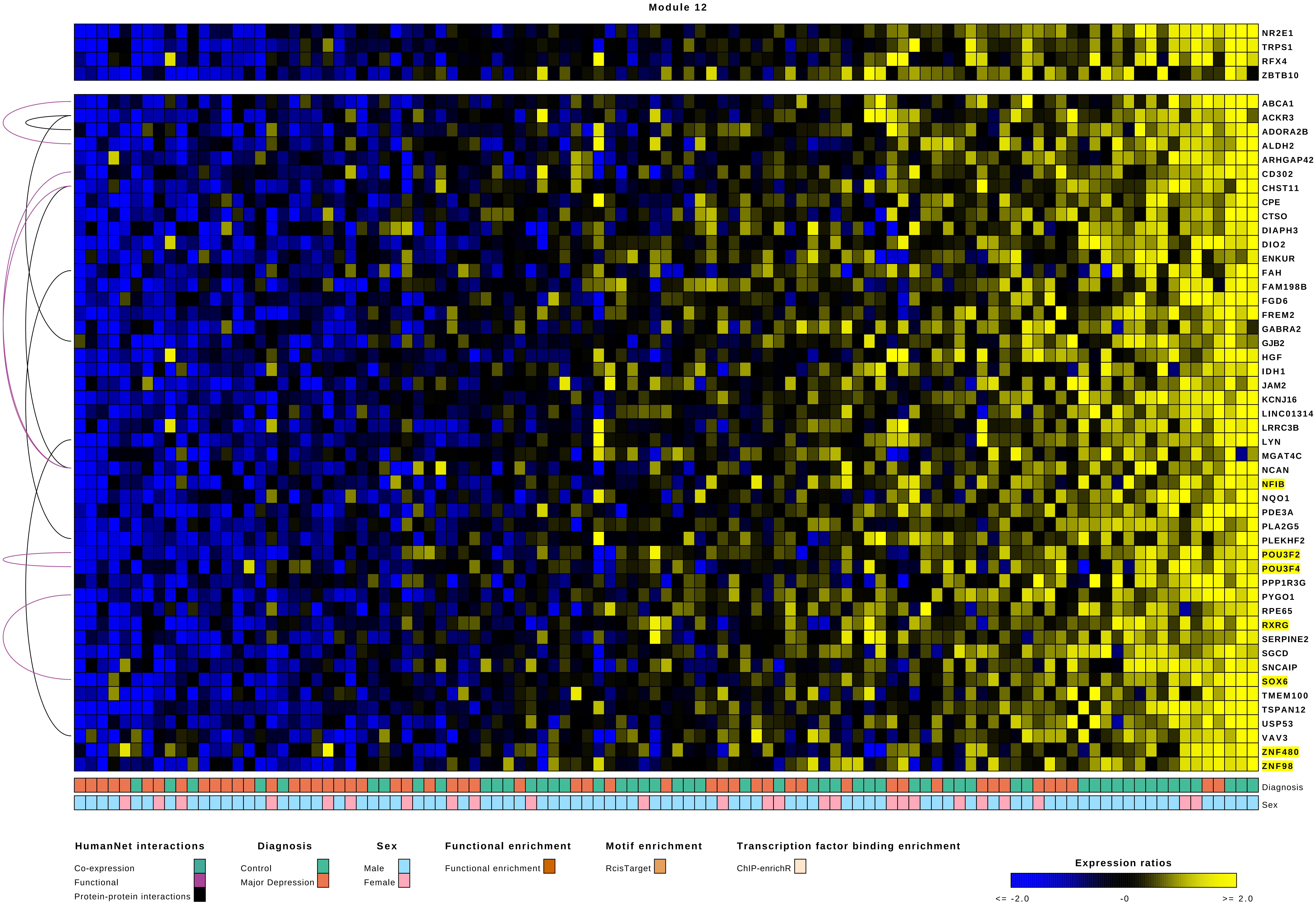
Module 12
NR2E1
TRPS1
RFX4
ZBTB10
ABCA1
ACKR3
ADORA2B
ALDH2
ARHGAP42
CD302
CHST11
CPE
CTSO
DIAPH3
DIO2
ENKUR
FAH
FAM198B
FGD6
FREM2
GABRA2
GJB2
HGF
IDH1
JAM2
KCNJ16
LINC01314
LRRC3B
LYN
MGAT4C
NCAN
NFIB
NQO1
PDE3A
PLA2G5
PLEKHF2
POU3F2
POU3F4
PPP1R3G
PYGO1
RPE65
RXRG
SERPINE2
SGCD
SNCAIP
SOX6
TMEM100
TSPAN12
USP53
VAV3
ZNF480
ZNF98
Diagnosis
Sex
HumanNet interactions
Co-expression
Functional
Protein-protein interactions
Diagnosis
Control
Major Depression
Sex
Male
Female
Functional enrichment
Functional enrichment
Motif enrichment
RcisTarget
Transcription factor binding enrichment
ChIP-enrichR
Expression ratios
<= -2.0
-0
>= 2.0

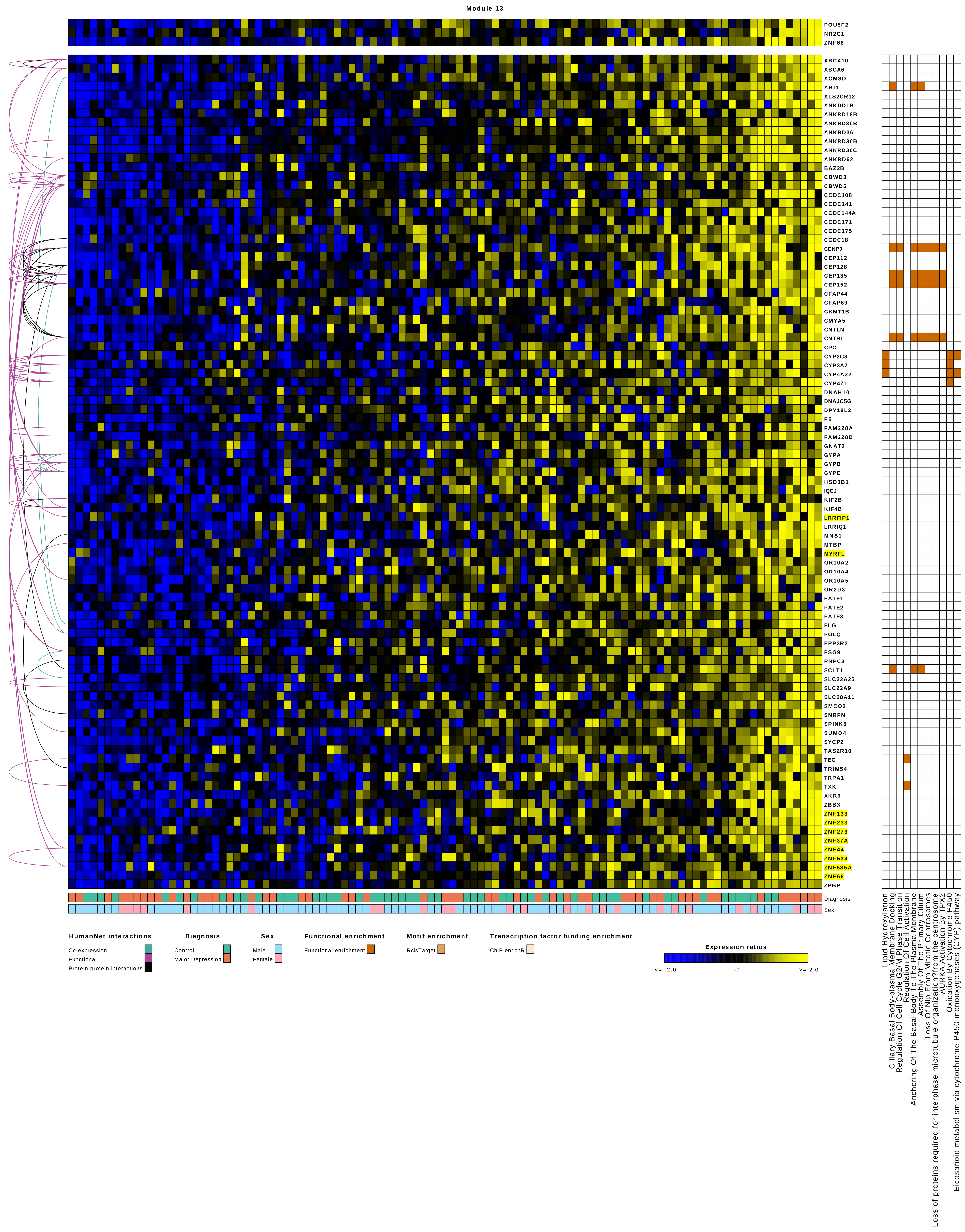
Module 13
POU5F2
NR2C1
ZNF66
ABCA10
ABCA6
ACMSD
AHI1
ALS2CR12
ANKDD1B
ANKRD18B
ANKRD30B
ANKRD36
ANKRD36B
ANKRD36C
ANKRD62
BAZ2B
CBWD3
CBWD5
CCDC108
CCDC141
CCDC144A
CCDC171
CCDC175
CCDC18
CENPJ
CEP112
CEP128
CEP135
CEP152
CFAP44
CFAP69
CKMT1B
CMYA5
CNTLN
CNTRL
CPO
CYP2C8
CYP3A7
CYP4A22
CYP4Z1
DNAH10
DNAJC5G
DPY19L2
F5
FAM228A
FAM228B
GNAT2
GYPA
GYPB
GYPE
HSD3B1
IQCJ
KIF2B
KIF4B
LRRFIP1
LRRIQ1
MNS1
MTBP
MYRFL
OR10A2
OR10A4
OR10A5
OR2D3
PATE1
PATE2
PATE3
PLG
POLQ
PPP3R2
PSG9
RNPC3
SCLT1
SLC22A25
SLC22A9
SLC38A11
SMCO2
SNRPN
SPINK5
SUMO4
SYCP2
TAS2R10
TEC
TRIM54
TRPA1
TXK
XKR6
ZBBX
ZNF133
ZNF233
ZNF273
ZNF37A
ZNF44
ZNF534
ZNF585A
ZNF66
ZPBP
Diagnosis
Sex
Lipid Hydroxylation
Ciliary Basal Body-plasma Membrane Docking
Regulation Of Cell Cycle G2/M Phase Transition
Regulation Of Cell Activation
Anchoring Of The Basal Body To The Plasma Membrane
Assembly Of The Primary Cilium
Loss Of Nlp From Mitotic Centrosomes
Loss of proteins required for interphase microtubule organization?from the centrosome
AURKA Activation By TPX2
Oxidation By Cytochrome P450
Eicosanoid metabolism via cytochrome P450 monooxygenases (CYP) pathway
HumanNet interactions
Co-expression
Functional
Protein-protein interactions
Diagnosis
Control
Major Depression
Sex
Male
Female
Functional enrichment
Functional enrichment
Motif enrichment
RcisTarget
Transcription factor binding enrichment
ChIP-enrichR
Expression ratios
<= -2.0
-0
>= 2.0

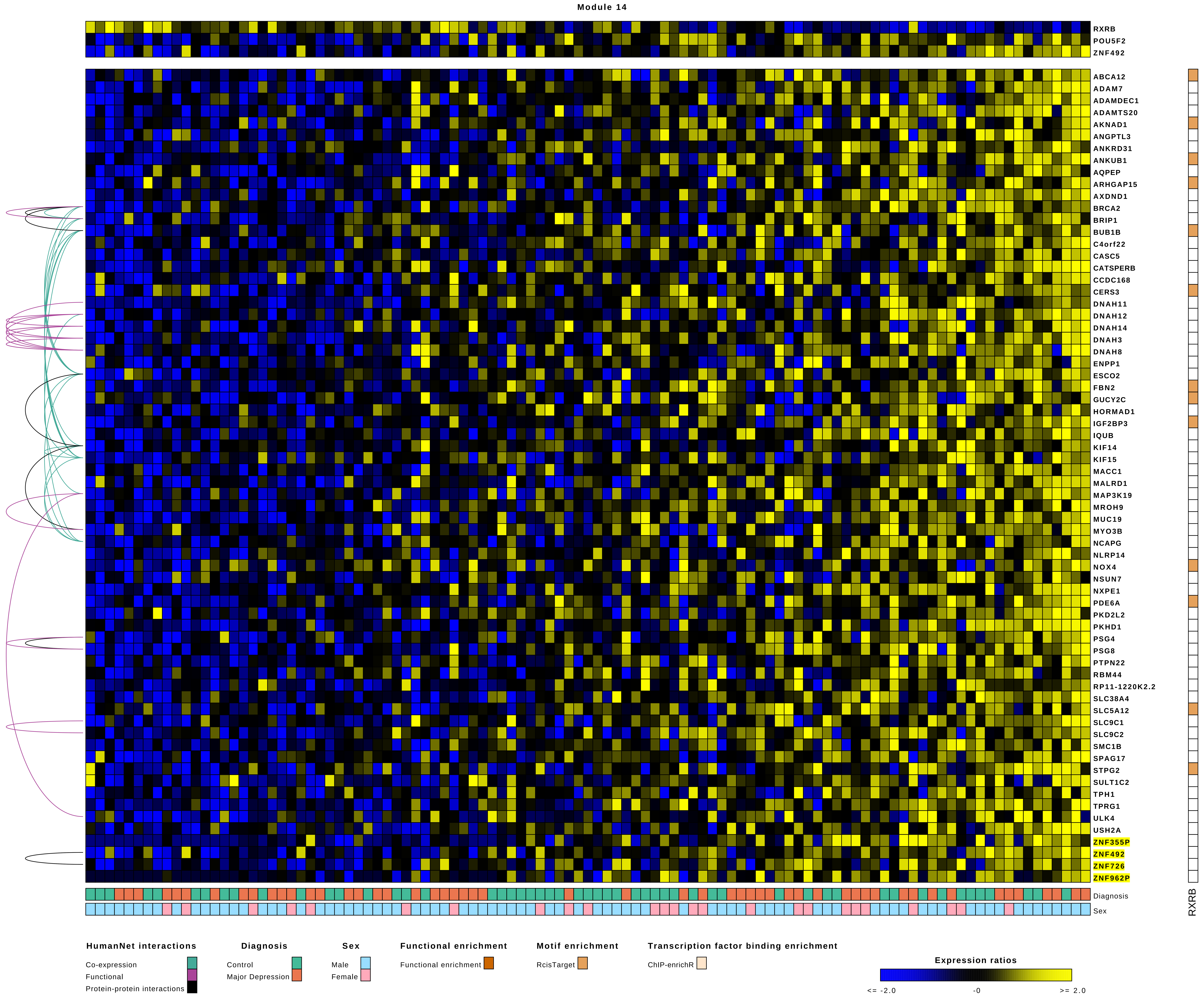

Module 14
RXRB
POU5F2
ZNF492
ABCA12
ADAM7
ADAMDEC1
ADAMTS20
AKNAD1
ANGPTL3
ANKRD31
ANKUB1
AQPEP
ARHGAP15
AXDND1
BRCA2
BRIP1
BUB1B
C4orf22
CASC5
CATSPERB
CCDC168
CERS3
DNAH11
DNAH12
DNAH14
DNAH3
DNAH8
ENPP1
ESCO2
FBN2
GUCY2C
HORMAD1
IGF2BP3
IQUB
KIF14
KIF15
MACC1
MALRD1
MAP3K19
MROH9
MUC19
MYO3B
NCAPG
NLRP14
NOX4
NSUN7
NXPE1
PDE6A
PKD2L2
PKHD1
PSG4
PSG8
PTPN22
RBM44
RP11-1220K2.2
SLC38A4
SLC5A12
SLC9C1
SLC9C2
SMC1B
SPAG17
STPG2
SULT1C2
TPH1
TPRG1
ULK4
USH2A
ZNF355P
ZNF492
ZNF726
ZNF962P
Diagnosis
Sex
RXRB
HumanNet interactions
Co-expression
Functional
Protein-protein interactions
Diagnosis
Control
Major Depression
Sex
Male
Female
Functional enrichment
Functional enrichment
Motif enrichment
RcisTarget
Transcription factor binding enrichment
ChIP-enrichR
Expression ratios
<= -2.0
-0
>= 2.0

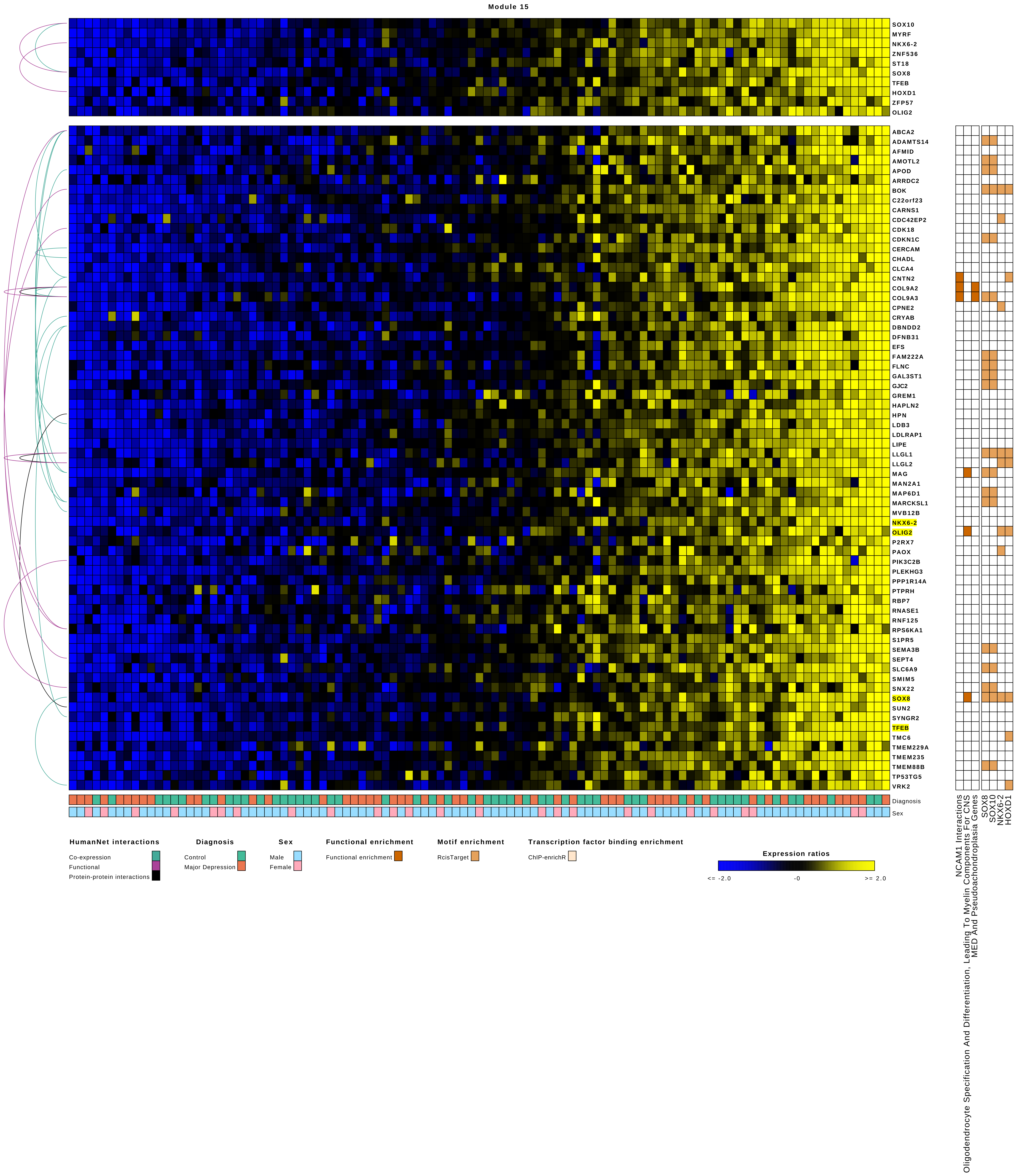
Module 15
SOX10
MYRF
NKX6-2
ZNF536
ST18
SOX8
TFEB
HOXD1
ZFP57
OLIG2
ABCA2
ADAMTS14
AFMID
AMOTL2
APOD
ARRDC2
BOK
C22orf23
CARNS1
CDC42EP2
CDK18
CDKN1C
CERCAM
CHADL
CLCA4
CNTN2
COL9A2
COL9A3
CPNE2
CRYAB
DBNDD2
DFNB31
EFS
FAM222A
FLNC
GAL3ST1
GJC2
GREM1
HAPLN2
HPN
LDB3
LDLRAP1
LIPE
LLGL1
LLGL2
MAG
MAN2A1
MAP6D1
MARCKSL1
MVB12B
NKX6-2
OLIG2
P2RX7
PAOX
PIK3C2B
PLEKHG3
PPP1R14A
PTPRH
RBP7
RNASE1
RNF125
RPS6KA1
S1PR5
SEMA3B
SEPT4
SLC6A9
SMIM5
SNX22
SOX8
SUN2
SYNGR2
TFEB
TMC6
TMEM229A
TMEM235
TMEM88B
TP53TG5
VRK2
Diagnosis
Sex
NCAM1 Interactions
Oligodendrocyte Specification And Differentiation, Leading To Myelin Components For CNS
MED And Pseudoachondroplasia Genes
SOX8
SOX10
NKX6-2
HOXD1
HumanNet interactions
Co-expression
Functional
Protein-protein interactions
Diagnosis
Control
Major Depression
Sex
Male
Female
Functional enrichment
Functional enrichment
Motif enrichment
RcisTarget
Transcription factor binding enrichment
ChIP-enrichR
Expression ratios
<= -2.0
-0
>= 2.0

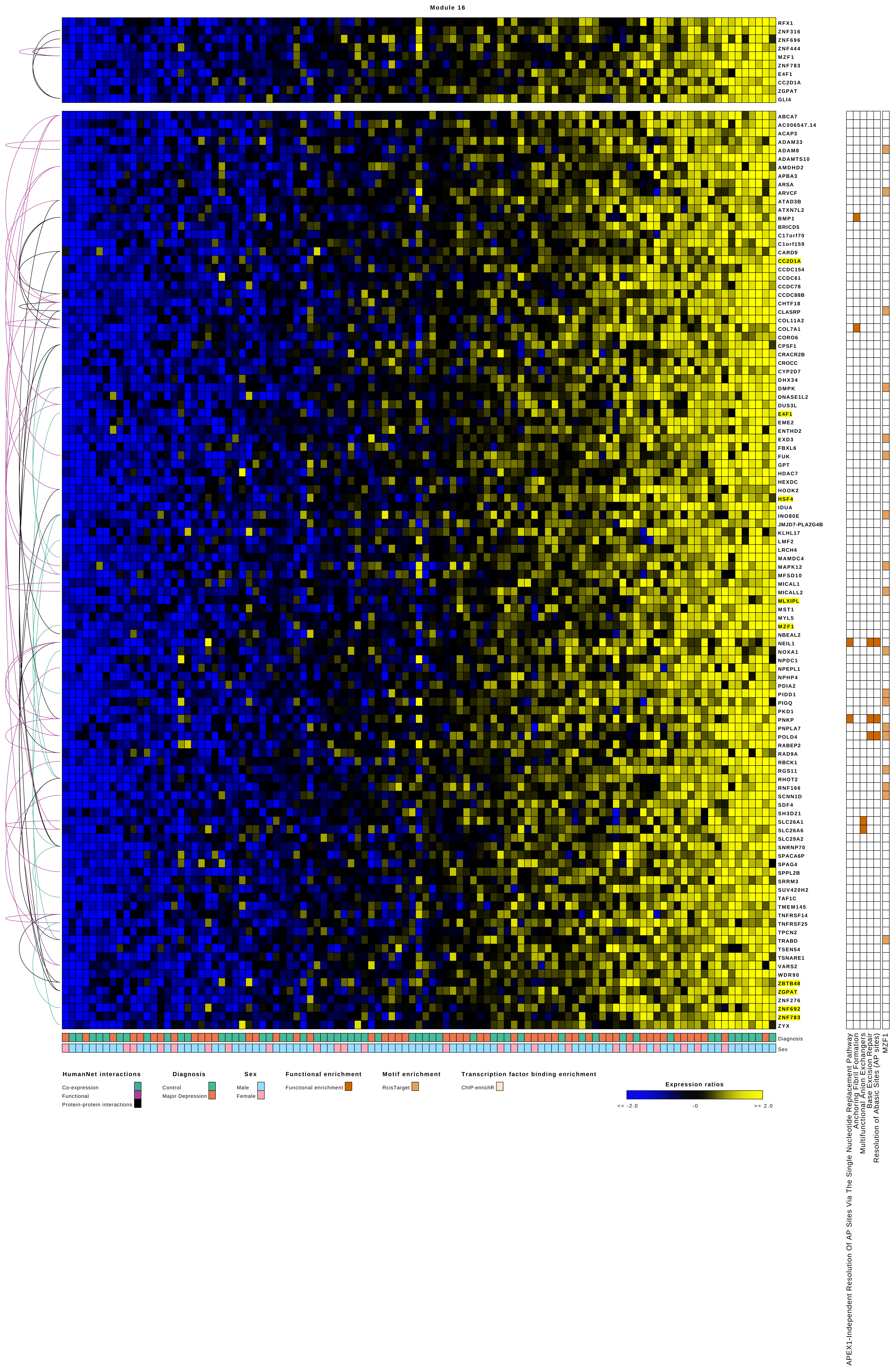

Module 16
RFX1
ZNF316
ZNF696
ZNF444
MZF1
ZNF783
E4F1
CC2D1A
ZGPAT
GLI4
ABCA7
AC006547.14
ACAP3
ADAM33
ADAM8
ADAMTS10
AMDHD2
APBA3
ARSA
ARVCF
ATAD3B
ATXN7L2
BMP1
BRICD5
C17orf70
C1orf159
CARD9
CC2D1A
CCDC154
CCDC61
CCDC78
CCDC88B
CHTF18
CLASRP
COL11A2
COL7A1
CORO6
CPSF1
CRACR2B
CROCC
CYP2D7
DHX34
DMPK
DNASE1L2
DUS3L
E4F1
EME2
ENTHD2
EXD3
FBXL6
FUK
GPT
HDAC7
HEXDC
HOOK2
HSF4
IDUA
INO80E
JMJD7-PLA2G4B
KLHL17
LMF2
LRCH4
MAMDC4
MAPK12
MFSD10
MICAL1
MICALL2
MLXIPL
MST1
MYL5
MZF1
NBEAL2
NEIL1
NOXA1
NPDC1
NPEPL1
NPHP4
PDIA2
PIDD1
PIGQ
PKD1
PNKP
PNPLA7
POLD4
RABEP2
RAD9A
RBCK1
RGS11
RHOT2
RNF166
SCNN1D
SDF4
SH3D21
SLC26A1
SLC26A6
SLC29A2
SNRNP70
SPACA6P
SPAG4
SPPL2B
SRRM3
SUV420H2
TAF1C
TMEM145
TNFRSF14
TNFRSF25
TPCN2
TRABD
TSEN54
TSNARE1
VARS2
WDR90
ZBTB48
ZGPAT
ZNF276
ZNF692
ZNF783
ZYX
Diagnosis
Sex
APEX1-Independent Resolution Of AP Sites Via The Single Nucleotide Replacement Pathway
Anchoring Fibril Formation
Multifunctional Anion Exchangers
Base Excision Repair
Resolution of Abasic Sites (AP sites)
MZF1
HumanNet interactions
Co-expression
Functional
Protein-protein interactions
Diagnosis
Control
Major Depression
Sex
Male
Female
Functional enrichment
Functional enrichment
Motif enrichment
RcisTarget
Transcription factor binding enrichment
ChIP-enrichR
Expression ratios
<= -2.0
-0
>= 2.0

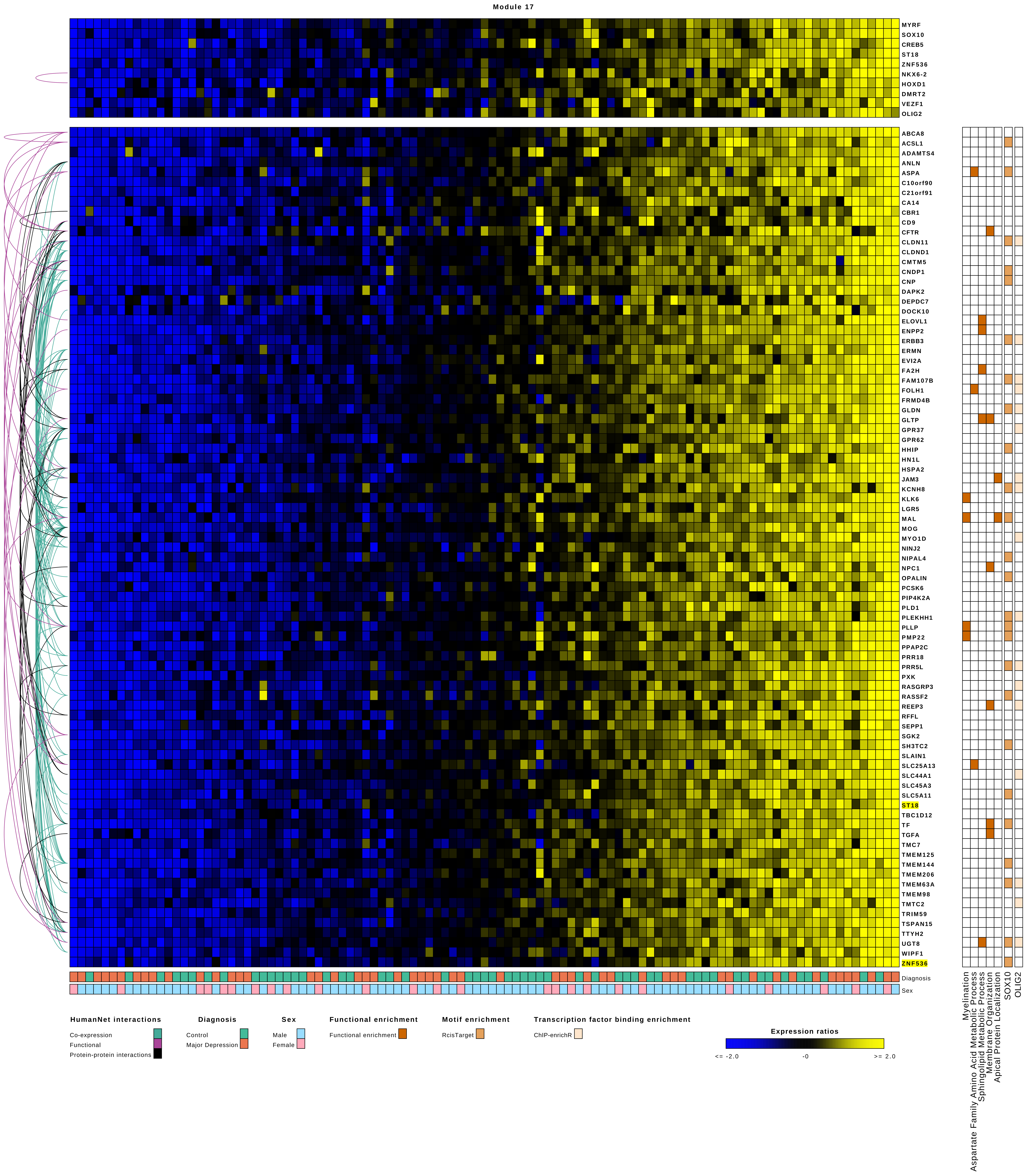
Module 17
MYRF
SOX10
CREB5
ST18
ZNF536
NKX6-2
HOXD1
DMRT2
VEZF1
OLIG2
ABCA8
ACSL1
ADAMTS4
ANLN
ASPA
C10orf90
C21orf91
CA14
CBR1
CD9
CFTR
CLDN11
CLDND1
CMTM5
CNDP1
CNP
DAPK2
DEPDC7
DOCK10
ELOVL1
ENPP2
ERBB3
ERMN
EVI2A
FA2H
FAM107B
FOLH1
FRMD4B
GLDN
GLTP
GPR37
GPR62
HHIP
HN1L
HSPA2
JAM3
KCNH8
KLK6
LGR5
MAL
MOG
MYO1D
NINJ2
NIPAL4
NPC1
OPALIN
PCSK6
PIP4K2A
PLD1
PLEKHH1
PLLP
PMP22
PPAP2C
PRR18
PRR5L
PXK
RASGRP3
RASSF2
REEP3
RFFL
SEPP1
SGK2
SH3TC2
SLAIN1
SLC25A13
SLC44A1
SLC45A3
SLC5A11
ST18
TBC1D12
TF
TGFA
TMC7
TMEM125
TMEM144
TMEM206
TMEM63A
TMEM98
TMTC2
TRIM59
TSPAN15
TTYH2
UGT8
WIPF1
ZNF536
Diagnosis
Sex
Myelination
Aspartate Family Amino Acid Metabolic Process
Sphingolipid Metabolic Process
Membrane Organization
Apical Protein Localization
SOX10
OLIG2
HumanNet interactions
Co-expression
Functional
Protein-protein interactions
Diagnosis
Control
Major Depression
Sex
Male
Female
Functional enrichment
Functional enrichment
Motif enrichment
RcisTarget
Transcription factor binding enrichment
ChIP-enrichR
Expression ratios
<= -2.0
-0
>= 2.0

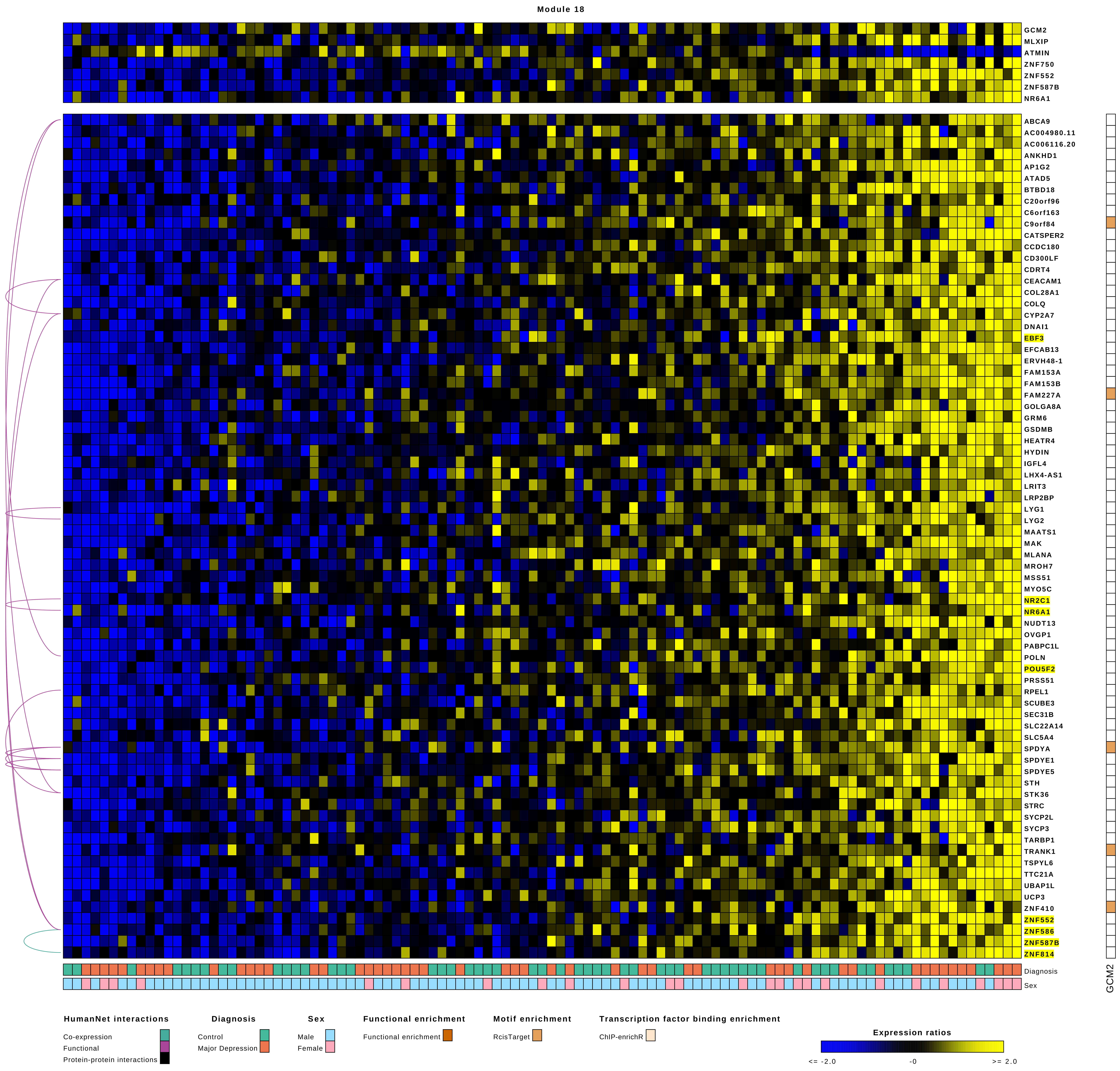

Module 18
GCM2
MLXIP
ATMIN
ZNF750
ZNF552
ZNF587B
NR6A1
ABCA9
AC004980.11
AC006116.20
ANKHD1
AP1G2
ATAD5
BTBD18
C20orf96
C6orf163
C9orf84
CATSPER2
CCDC180
CD300LF
CDRT4
CEACAM1
COL28A1
COLQ
CYP2A7
DNAI1
EBF3
EFCAB13
ERVH48-1
FAM153A
FAM153B
FAM227A
GOLGA8A
GRM6
GSDMB
HEATR4
HYDIN
IGFL4
LHX4-AS1
LRIT3
LRP2BP
LYG1
LYG2
MAATS1
MAK
MLANA
MROH7
MSS51
MYO5C
NR2C1
NR6A1
NUDT13
OVGP1
PABPC1L
POLN
POU5F2
PRSS51
RPEL1
SCUBE3
SEC31B
SLC22A14
SLC5A4
SPDYA
SPDYE1
SPDYE5
STH
STK36
STRC
SYCP2L
SYCP3
TARBP1
TRANK1
TSPYL6
TTC21A
UBAP1L
UCP3
ZNF410
ZNF552
ZNF586
ZNF587B
ZNF814
Diagnosis
Sex
GCM2
HumanNet interactions
Co-expression
Functional
Protein-protein interactions
Diagnosis
Control
Major Depression
Sex
Male
Female
Functional enrichment
Functional enrichment
Motif enrichment
RcisTarget
Transcription factor binding enrichment
ChIP-enrichR
Expression ratios
<= -2.0
-0
>= 2.0

Module 19

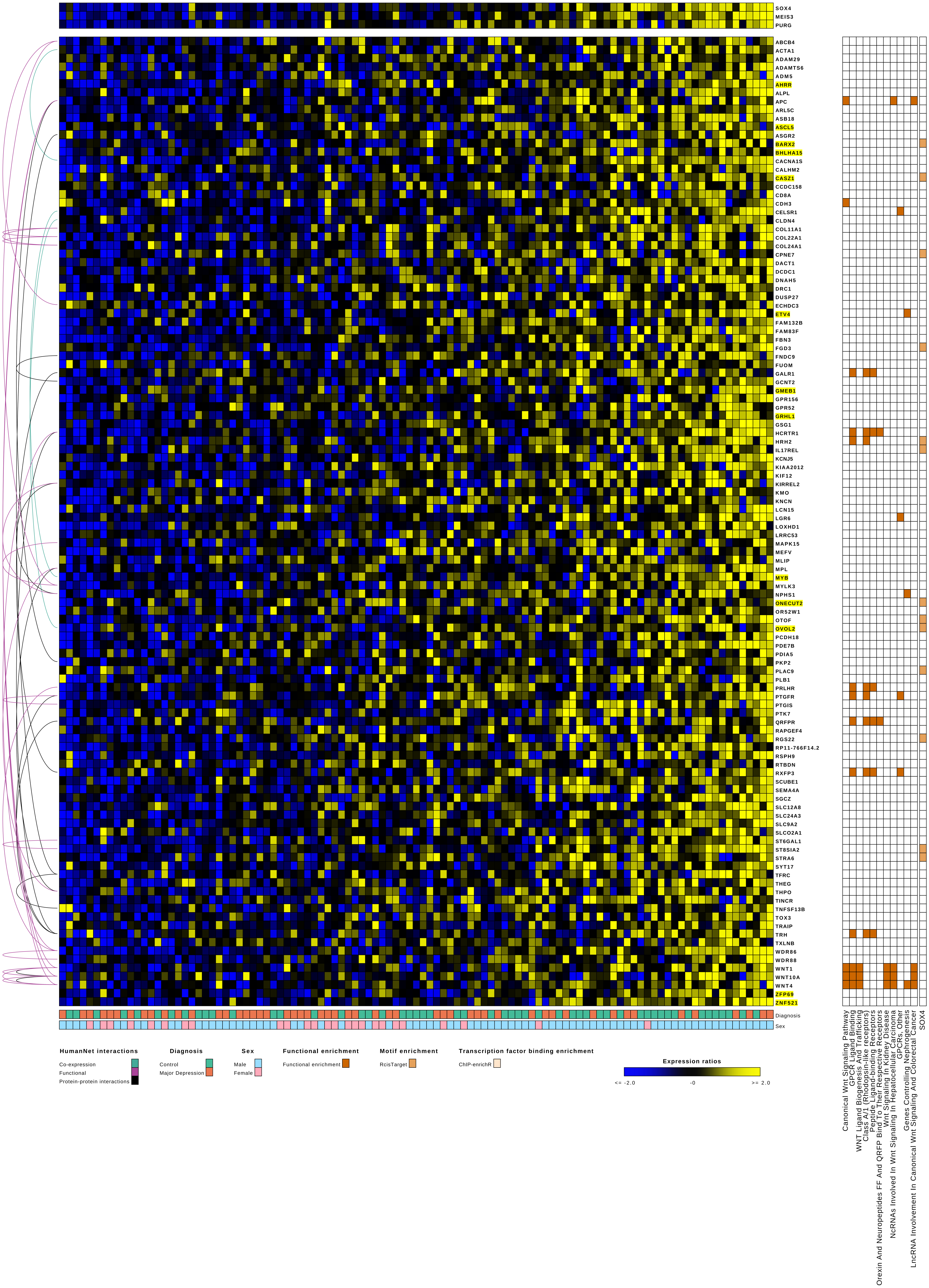
SOX4
MEIS3
PURG
ABCB4
ACTA1
ADAM29
ADAMTS6
ADM5
AHRR
ALPL
APC
ARL5C
ASB18
ASCL5
ASGR2
BARX2
BHLHA15
CACNA1S
CALHM2
CASZ1
CCDC158
CD8A
CDH3
CELSR1
CLDN4
COL11A1
COL22A1
COL24A1
CPNE7
DACT1
DCDC1
DNAH5
DRC1
DUSP27
ECHDC3
ETV4
FAM132B
FAM83F
FBN3
FGD3
FNDC9
FUOM
GALR1
GCNT2
GMEB1
GPR156
GPR52
GRHL1
GSG1
HCRTR1
HRH2
IL17REL
KCNJ5
KIAA2012
KIF12
KIRREL2
KMO
KNCN
LCN15
LGR6
LOXHD1
LRRC53
MAPK15
MEFV
MLIP
MPL
MYB
MYLK3
NPHS1
ONECUT2
OR52W1
OTOF
OVOL2
PCDH18
PDE7B
PDIA5
PKP2
PLAC9
PLB1
PRLHR
PTGFR
PTGIS
PTK7
QRFPR
RAPGEF4
RGS22
RP11-766F14.2
RSPH9
RTBDN
RXFP3
SCUBE1
SEMA4A
SGCZ
SLC12A8
SLC24A3
SLC9A2
SLCO2A1
ST6GAL1
ST8SIA2
STRA6
SYT17
TFRC
THEG
THPO
TINCR
TNFSF13B
TOX3
TRAIP
TRH
TXLNB
WDR86
WDR88
WNT1
WNT10A
WNT4
ZFP69
ZNF521
Diagnosis
Sex
Canonical Wnt Signaling Pathway
GPCR Ligand Binding
WNT Ligand Biogenesis And Trafficking
Class A/1 (Rhodopsin-like receptors)
Peptide Ligand-binding Receptors
Orexin And Neuropeptides FF And QRFP Bind To Their Respective Receptors
Wnt Signaling In Kidney Disease
NcRNAs Involved In Wnt Signaling In Hepatocellular Carcinoma
GPCRs, Other
Genes Controlling Nephrogenesis
LncRNA Involvement In Canonical Wnt Signaling And Colorectal Cancer
SOX4
HumanNet interactions
Co-expression
Functional
Protein-protein interactions
Diagnosis
Control
Major Depression
Sex
Male
Female
Functional enrichment
Functional enrichment
Motif enrichment
RcisTarget
Transcription factor binding enrichment
ChIP-enrichR
Expression ratios
<= -2.0
-0
>= 2.0

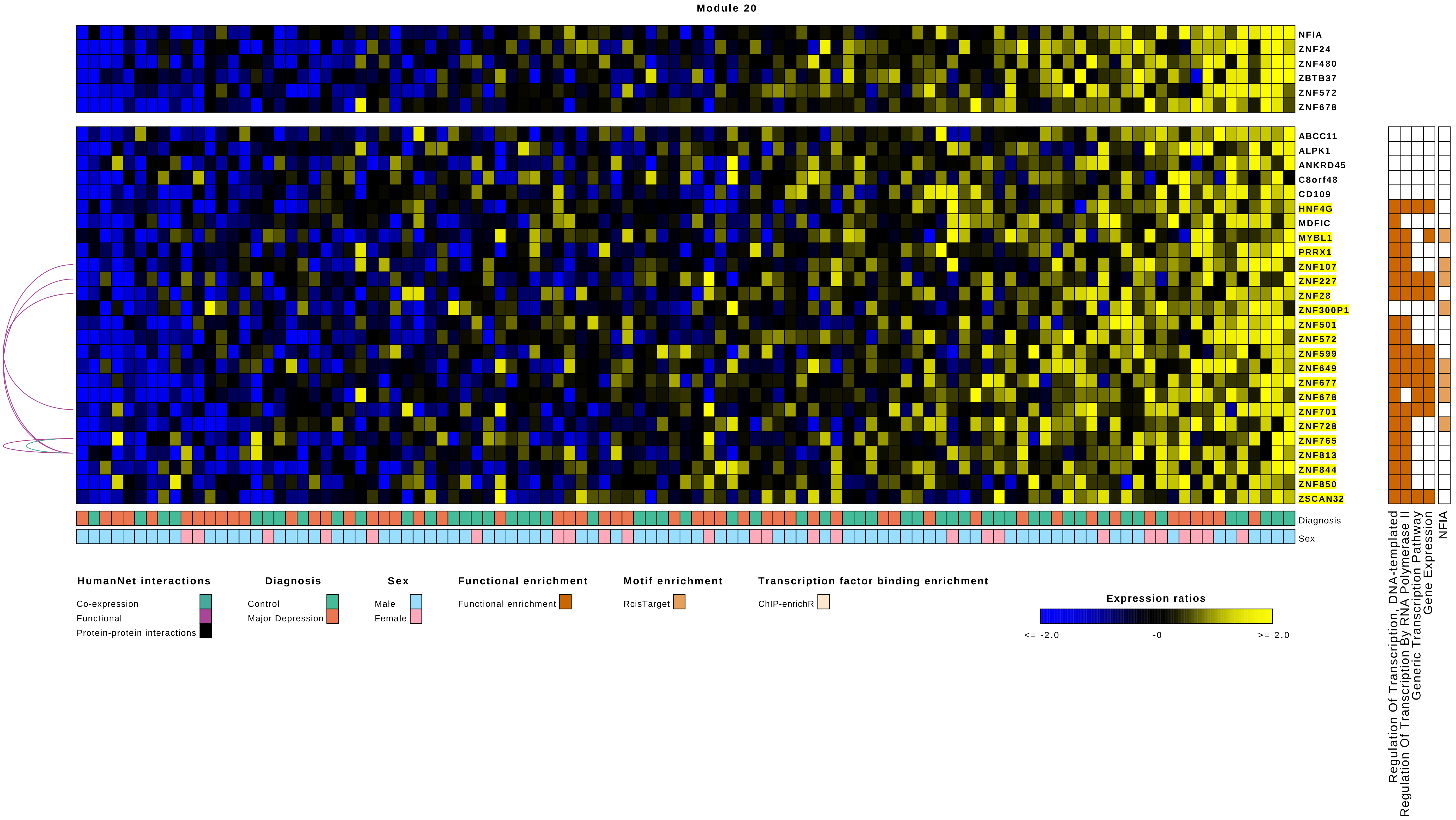

Module 20
NFIA
ZNF24
ZNF480
ZBTB37
ZNF572
ZNF678
ABCC11
ALPK1
ANKRD45
C8orf48
CD109
HNF4G
MDFIC
MYBL1
PRRX1
ZNF107
ZNF227
ZNF28
ZNF300P1
ZNF501
ZNF572
ZNF599
ZNF649
ZNF677
ZNF678
ZNF701
ZNF728
ZNF765
ZNF813
ZNF844
ZNF850
ZSCAN32
Diagnosis
Sex
Regulation Of Transcription, DNA-templated
Regulation Of Transcription By RNA Polymerase II
Generic Transcription Pathway
Gene Expression
NFIA
HumanNet interactions
Co-expression
Functional
Protein-protein interactions
Diagnosis
Control
Major Depression
Sex
Male
Female
Functional enrichment
Functional enrichment
Motif enrichment
RcisTarget
Transcription factor binding enrichment
ChIP-enrichR
Expression ratios
<= -2.0
-0
>= 2.0

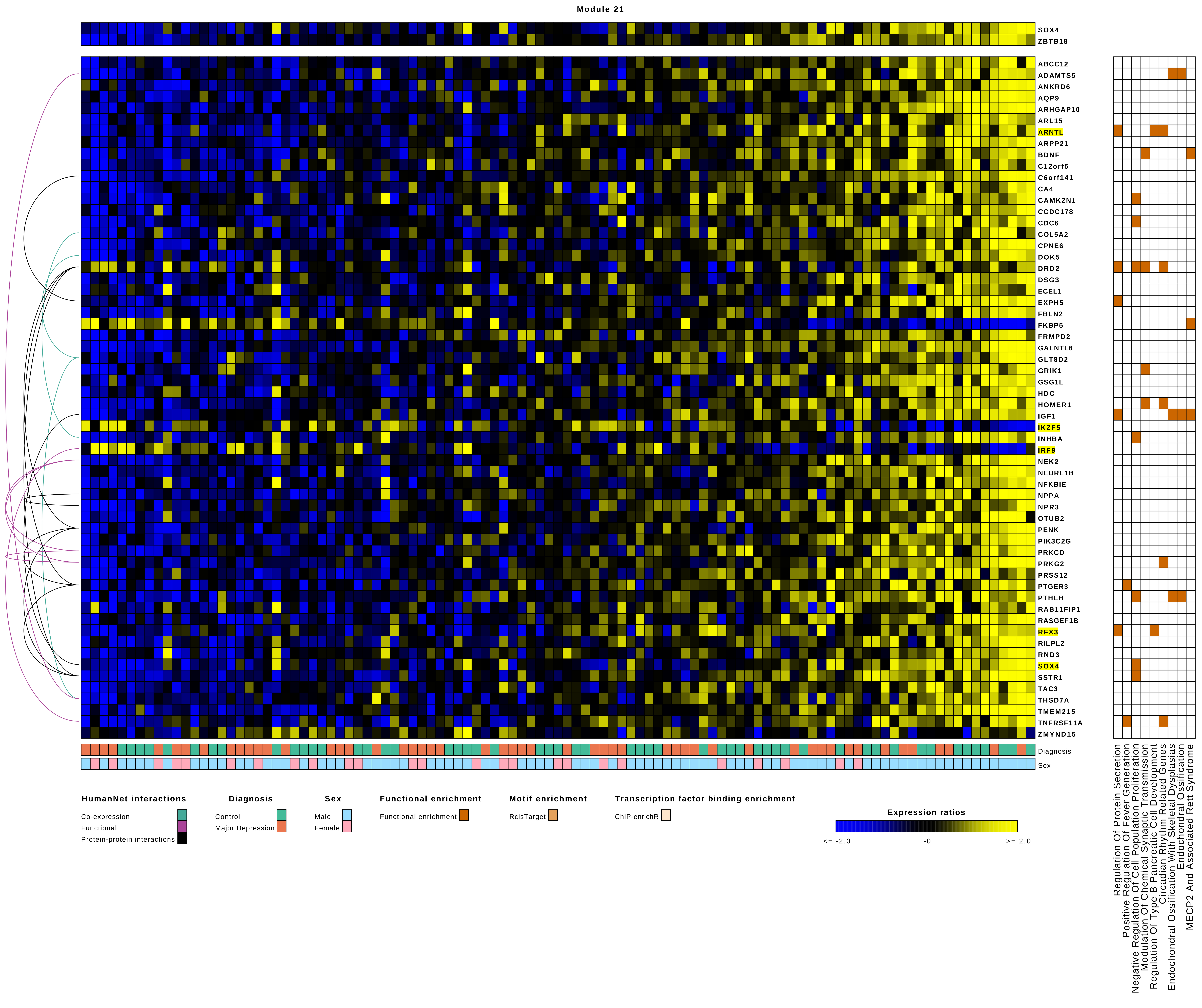
Module 21
SOX4
ZBTB18
ABCC12
ADAMTS5
ANKRD6
AQP9
ARHGAP10
ARL15
ARNTL
ARPP21
BDNF
C12orf5
C6orf141
CA4
CAMK2N1
CCDC178
CDC6
COL5A2
CPNE6
DOK5
DRD2
DSG3
ECEL1
EXPH5
FBLN2
FKBP5
FRMPD2
GALNTL6
GLT8D2
GRIK1
GSG1L
HDC
HOMER1
IGF1
IKZF5
INHBA
IRF9
NEK2
NEURL1B
NFKBIE
NPPA
NPR3
OTUB2
PENK
PIK3C2G
PRKCD
PRKG2
PRSS12
PTGER3
PTHLH
RAB11FIP1
RASGEF1B
RFX3
RILPL2
RND3
SOX4
SSTR1
TAC3
THSD7A
TMEM215
TNFRSF11A
ZMYND15
Diagnosis
Sex
Regulation Of Protein Secretion
Positive Regulation Of Fever Generation
Negative Regulation Of Cell Population Proliferation
Modulation Of Chemical Synaptic Transmission
Regulation Of Type B Pancreatic Cell Development
Circadian Rhythm Related Genes
Endochondral Ossification With Skeletal Dysplasias
Endochondral Ossification
MECP2 And Associated Rett Syndrome
HumanNet interactions
Co-expression
Functional
Protein-protein interactions
Diagnosis
Control
Major Depression
Sex
Male
Female
Functional enrichment
Functional enrichment
Motif enrichment
RcisTarget
Transcription factor binding enrichment
ChIP-enrichR
Expression ratios
<= -2.0
-0
>= 2.0

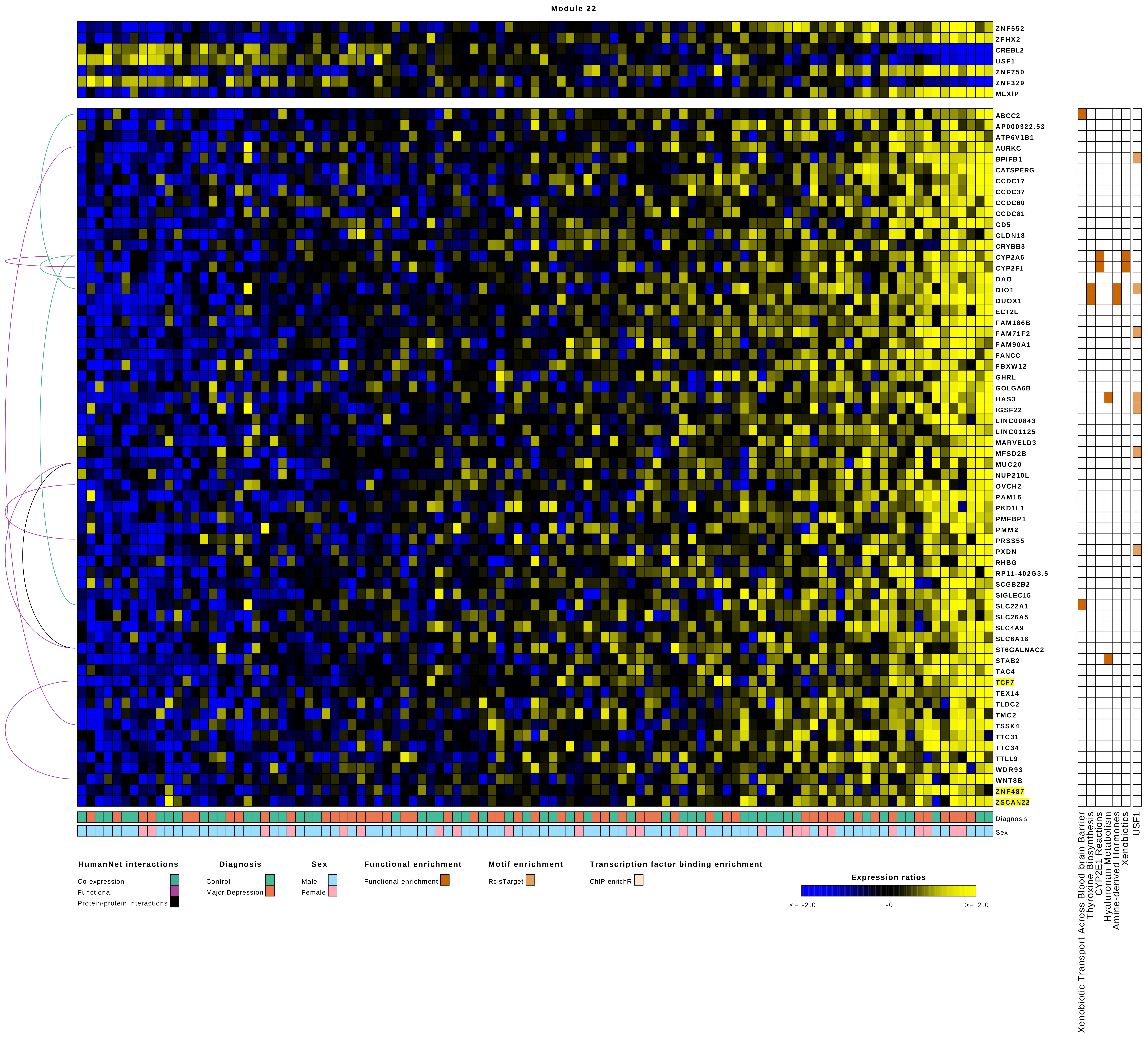
Module 22
ZNF552
ZFHX2
CREBL2
USF1
ZNF750
ZNF329
MLXIP
ABCC2
AP000322.53
ATP6V1B1
AURKC
BPIFB1
CATSPERG
CCDC17
CCDC37
CCDC60
CCDC81
CD5
CLDN18
CRYBB3
CYP2A6
CYP2F1
DAO
DIO1
DUOX1
ECT2L
FAM186B
FAM71F2
FAM90A1
FANCC
FBXW12
GHRL
GOLGA6B
HAS3
IGSF22
LINC00843
LINC01125
MARVELD3
MFSD2B
MUC20
NUP210L
OVCH2
PAM16
PKD1L1
PMFBP1
PMM2
PRSS55
PXDN
RHBG
RP11-402G3.5
SCGB2B2
SIGLEC15
SLC22A1
SLC26A5
SLC4A9
SLC6A16
ST6GALNAC2
STAB2
TAC4
TCF7
TEX14
TLDC2
TMC2
TSSK4
TTC31
TTC34
TTLL9
WDR93
WNT8B
ZNF487
ZSCAN22
Diagnosis
Sex
Xenobiotic Transport Across Blood-brain Barrier
Thyroxine Biosynthesis
CYP2E1 Reactions
Hyaluronan Metabolism
Amine-derived Hormones
Xenobiotics
USF1
HumanNet interactions
Co-expression
Functional
Protein-protein interactions
Diagnosis
Control
Major Depression
Sex
Male
Female
Functional enrichment
Functional enrichment
Motif enrichment
RcisTarget
Transcription factor binding enrichment
ChIP-enrichR
Expression ratios
<= -2.0
-0
>= 2.0

### Module 23

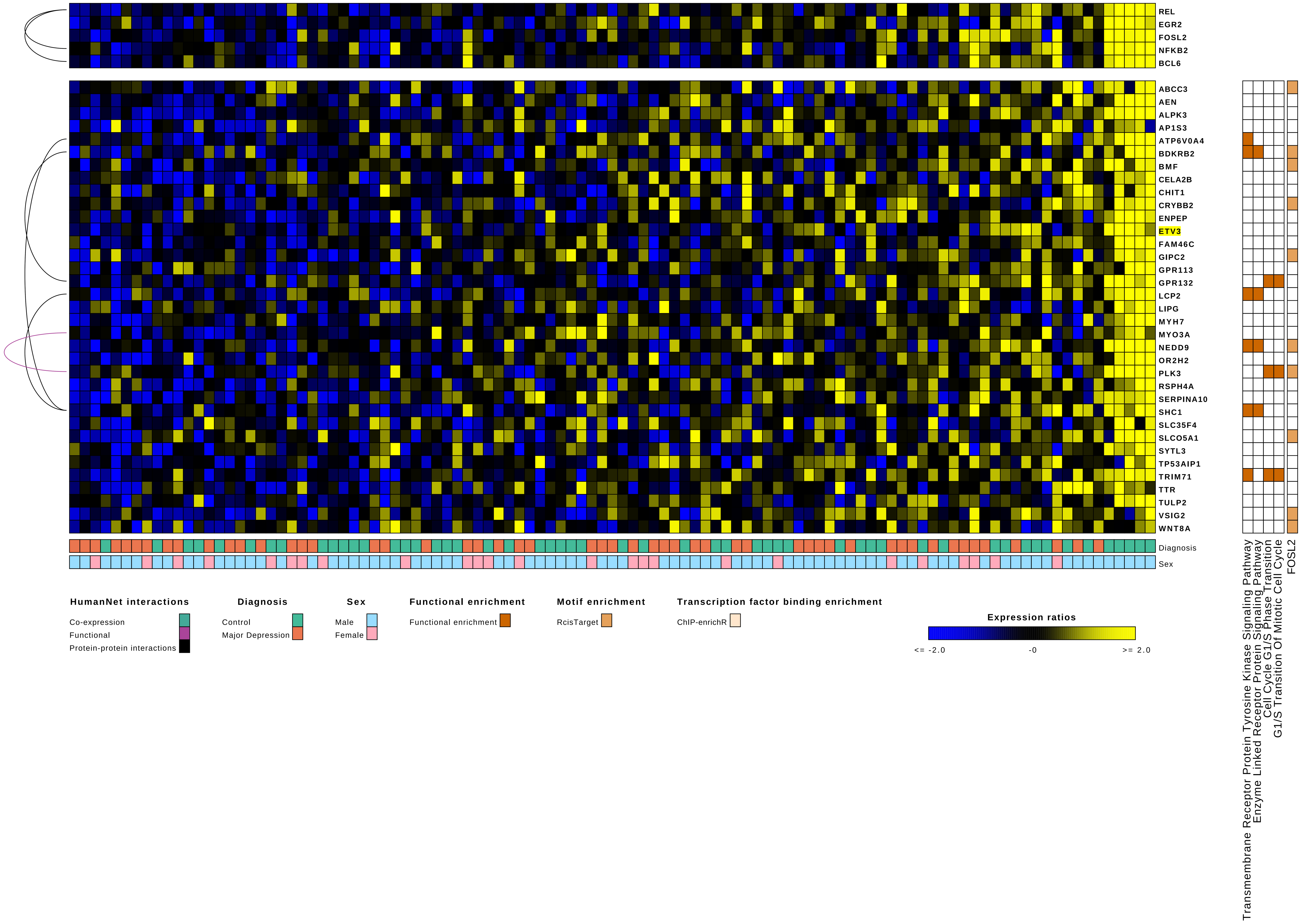

REL
EGR2
FOSL2
NFKB2
BCL6
ABCC3
AEN
ALPK3
AP1S3
ATP6V0A4
BDKRB2
BMF
CELA2B
CHIT1
CRYBB2
ENPEP
ETV3
FAM46C
GIPC2
GPR113
GPR132
LCP2
LIPG
MYH7
MYO3A
NEDD9
OR2H2
PLK3
RSPH4A
SERPINA10
SHC1
SLC35F4
SLCO5A1
SYTL3
TP53AIP1
TRIM71
TTR
TULP2
VSIG2
WNT8A
Diagnosis
Sex
Transmembrane Receptor Protein Tyrosine Kinase Signaling Pathway
Enzyme Linked Receptor Protein Signaling Pathway
Cell Cycle G1/S Phase Transition
G1/S Transition Of Mitotic Cell Cycle
FOSL2
HumanNet interactions
Co-expression
Functional
Protein-protein interactions
Diagnosis
Control
Major Depression
Sex
Male
Female
Functional enrichment
Functional enrichment
Motif enrichment
RcisTarget
Transcription factor binding enrichment
ChIP-enrichR
Expression ratios
<= -2.0
-0
>= 2.0

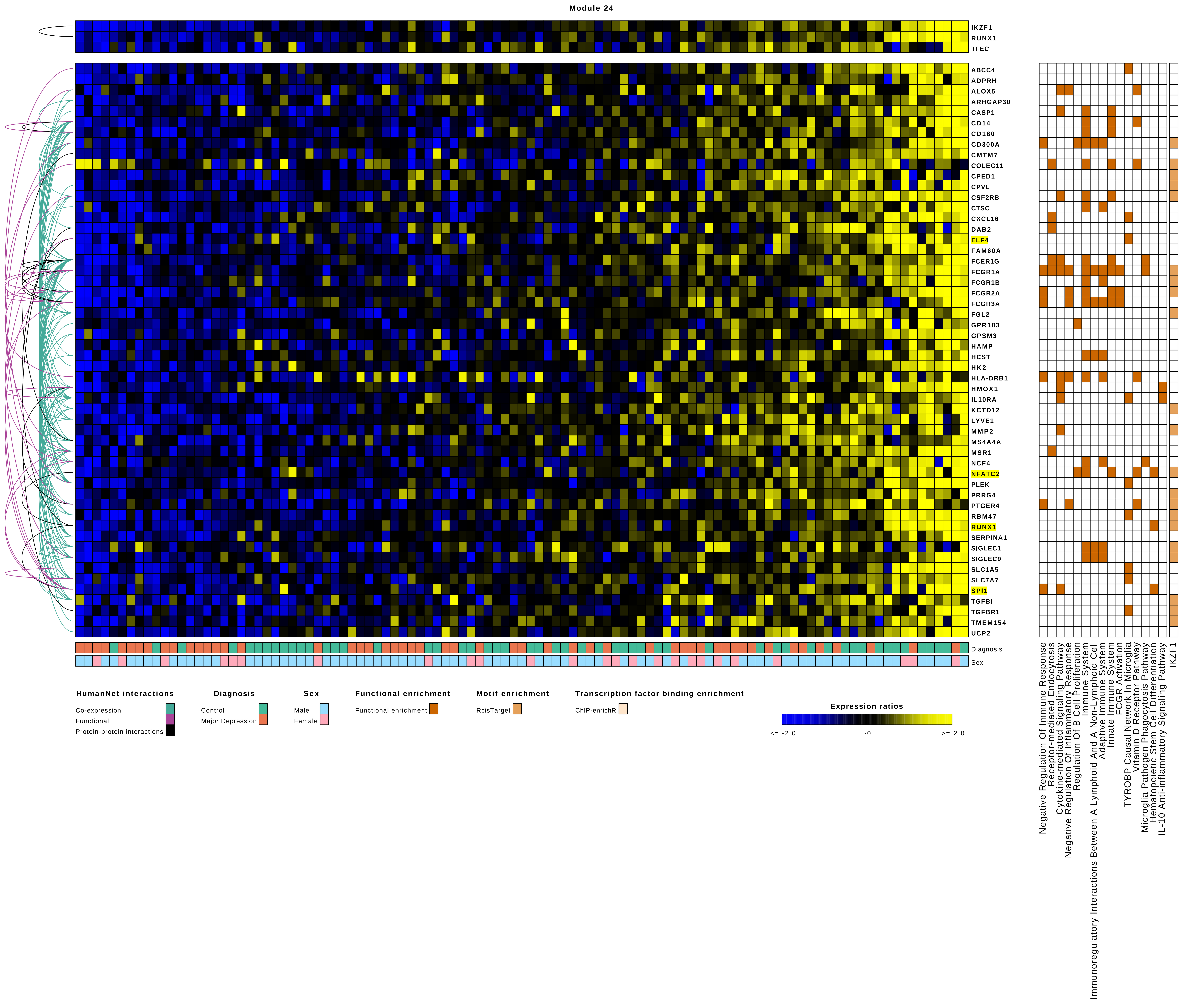
Module 24
IKZF1
RUNX1
TFEC
ABCC4
ADPRH
ALOX5
ARHGAP30
CASP1
CD14
CD180
CD300A
CMTM7
COLEC11
CPED1
CPVL
CSF2RB
CTSC
CXCL16
DAB2
ELF4
FAM60A
FCER1G
FCGR1A
FCGR1B
FCGR2A
FCGR3A
FGL2
GPR183
GPSM3
HAMP
HCST
HK2
HLA-DRB1
HMOX1
IL10RA
KCTD12
LYVE1
MMP2
MS4A4A
MSR1
NCF4
NFATC2
PLEK
PRRG4
PTGER4
RBM47
RUNX1
SERPINA1
SIGLEC1
SIGLEC9
SLC1A5
SLC7A7
SPI1
TGFBI
TGFBR1
TMEM154
UCP2
Diagnosis
Sex
Negative Regulation Of Immune Response
Receptor-mediated Endocytosis
Cytokine-mediated Signaling Pathway
Negative Regulation Of Inflammatory Response
Regulation Of B Cell Proliferation
Immune System
Immunoregulatory Interactions Between A Lymphoid And A Non-Lymphoid Cell
Adaptive Immune System
Innate Immune System
FCGR Activation
TYROBP Causal Network In Microglia
Vitamin D Receptor Pathway
Microglia Pathogen Phagocytosis Pathway
Hematopoietic Stem Cell Differentiation
IL-10 Anti-inflammatory Signaling Pathway
IKZF1
HumanNet interactions
Co-expression
Functional
Protein-protein interactions
Diagnosis
Control
Major Depression
Sex
Male
Female
Functional enrichment
Functional enrichment
Motif enrichment
RcisTarget
Transcription factor binding enrichment
ChIP-enrichR
Expression ratios
<= -2.0
-0
>= 2.0

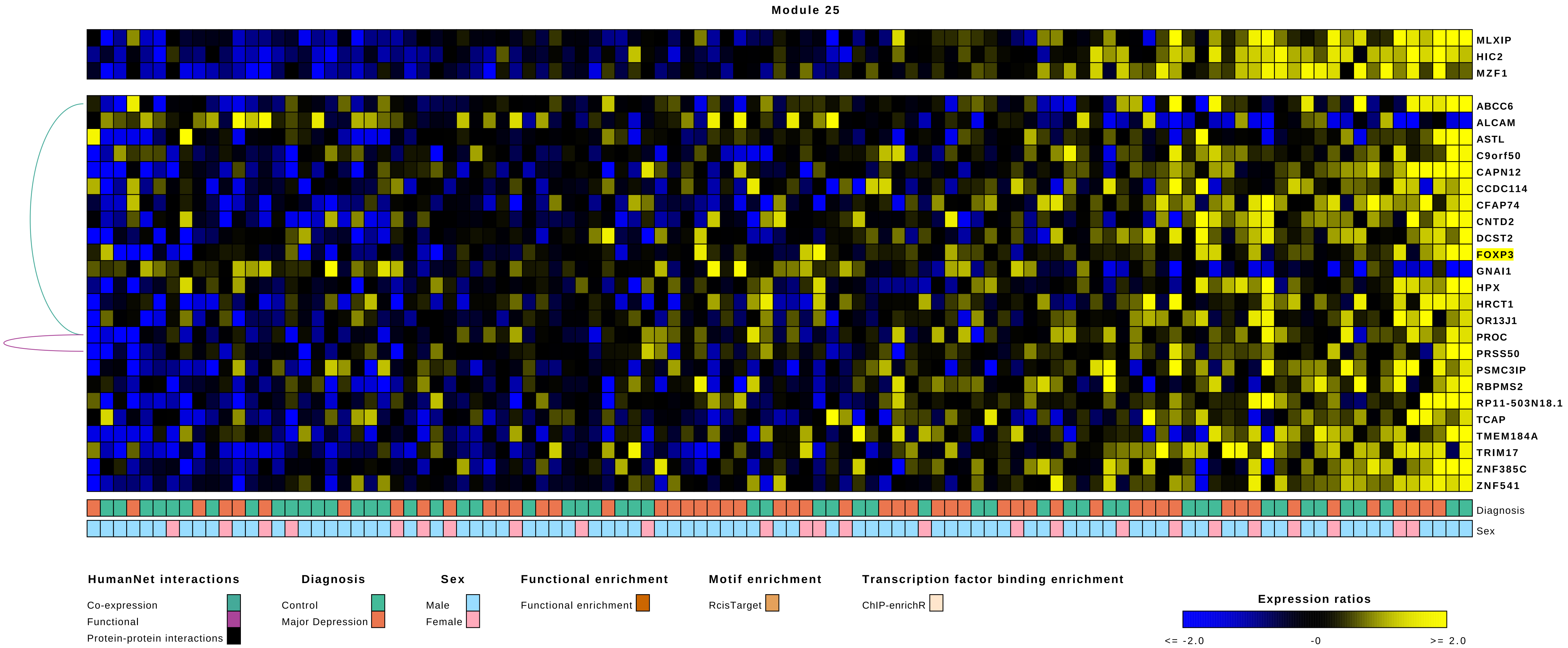

Module 25
MLXIP
HIC2
MZF1
ABCC6
ALCAM
ASTL
C9orf50
CAPN12
CCDC114
CFAP74
CNTD2
DCST2
FOXP3
GNAI1
HPX
HRCT1
OR13J1
PROC
PRSS50
PSMC3IP
RBPMS2
RP11-503N18.1
TCAP
TMEM184A
TRIM17
ZNF385C
ZNF541
Diagnosis
Sex
HumanNet interactions
Co-expression
Functional
Protein-protein interactions
Diagnosis
Control
Major Depression
Sex
Male
Female
Functional enrichment
Functional enrichment
Motif enrichment
RcisTarget
Transcription factor binding enrichment
ChIP-enrichR
Expression ratios
<= -2.0
-0
>= 2.0

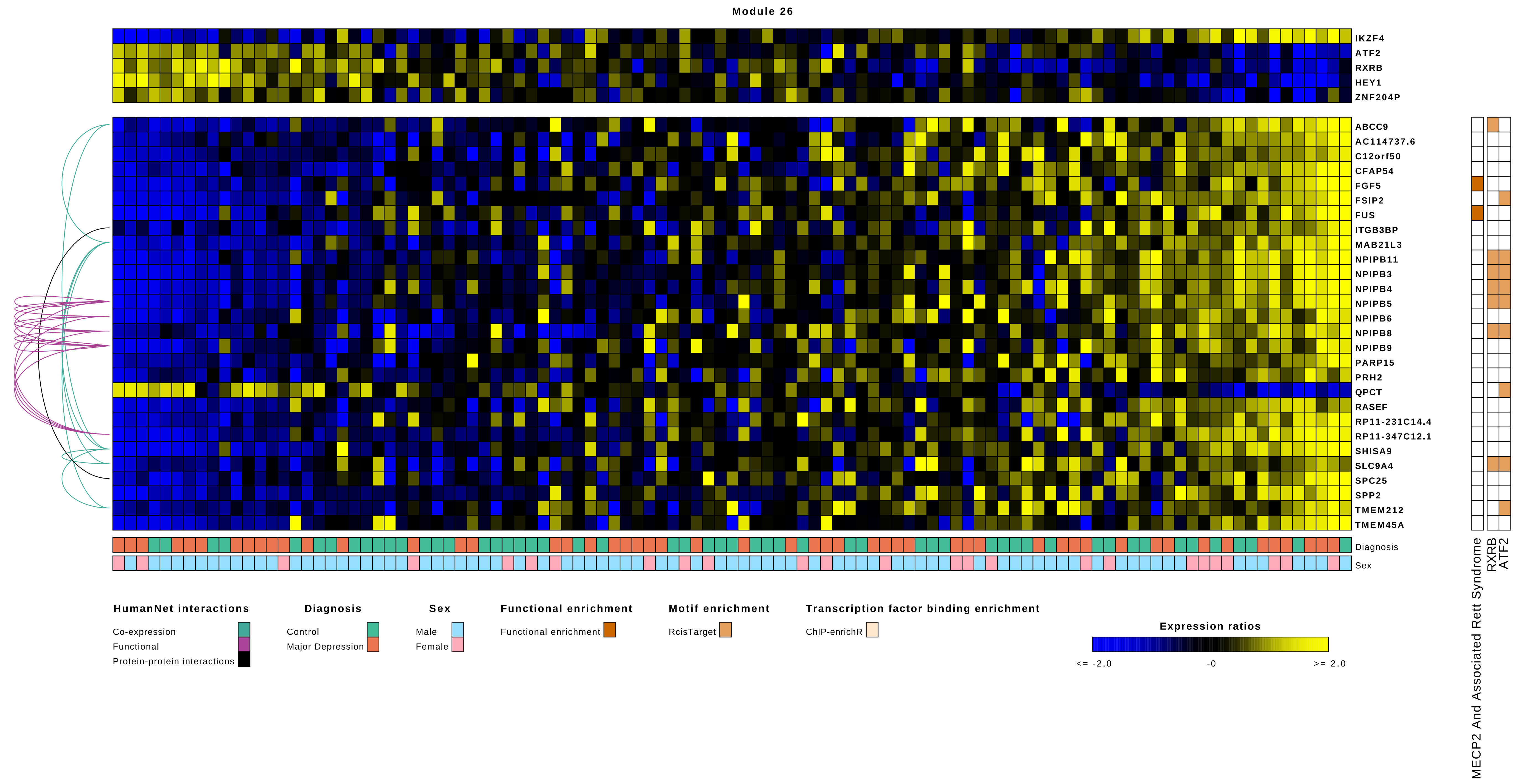

Module 26
IKZF4
ATF2
RXRB
HEY1
ZNF204P
ABCC9
AC114737.6
C12orf50
CFAP54
FGF5
FSIP2
FUS
ITGB3BP
MAB21L3
NPIPB11
NPIPB3
NPIPB4
NPIPB5
NPIPB6
NPIPB8
NPIPB9
PARP15
PRH2
QPCT
RASEF
RP11-231C14.4
RP11-347C12.1
SHISA9
SLC9A4
SPC25
SPP2
TMEM212
TMEM45A
Diagnosis
Sex
MECP2 And Associated Rett Syndrome
RXRB
ATF2
HumanNet interactions
Co-expression
Functional
Protein-protein interactions
Diagnosis
Control
Major Depression
Sex
Male
Female
Functional enrichment
Functional enrichment
Motif enrichment
RcisTarget
Transcription factor binding enrichment
ChIP-enrichR
Expression ratios
<= -2.0
-0
>= 2.0

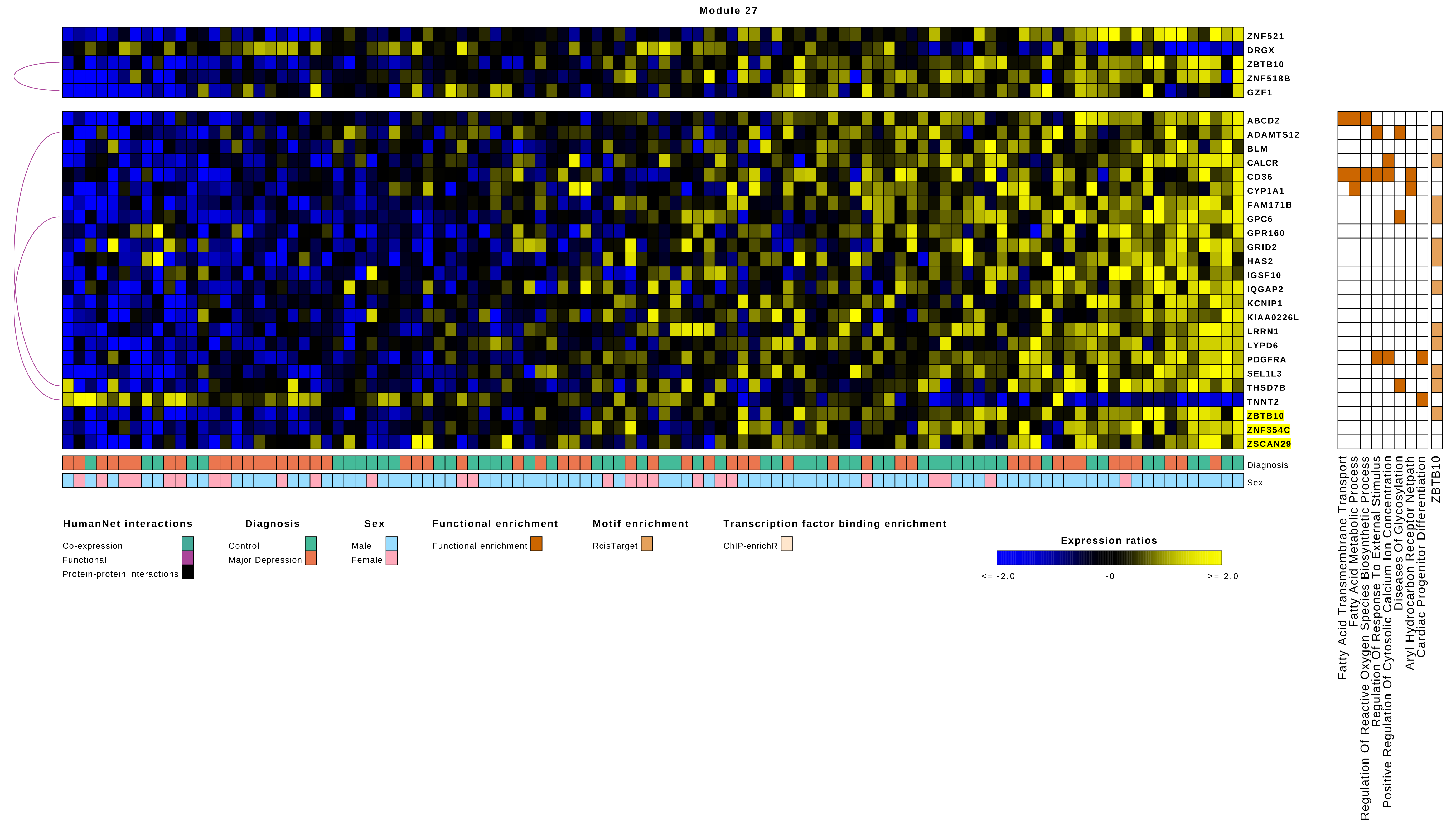
Module 27
ZNF521
DRGX
ZBTB10
ZNF518B
GZF1
ABCD2
ADAMTS12
BLM
CALCR
CD36
CYP1A1
FAM171B
GPC6
GPR160
GRID2
HAS2
IGSF10
IQGAP2
KCNIP1
KIAA0226L
LRRN1
LYPD6
PDGFRA
SEL1L3
THSD7B
TNNT2
ZBTB10
ZNF354C
ZSCAN29
Diagnosis
Sex
Fatty Acid Transmembrane Transport
Fatty Acid Metabolic Process
Regulation Of Reactive Oxygen Species Biosynthetic Process
Regulation Of Response To External Stimulus
Positive Regulation Of Cytosolic Calcium Ion Concentration
Diseases Of Glycosylation
Aryl Hydrocarbon Receptor Netpath
Cardiac Progenitor Differentiation
ZBTB10
HumanNet interactions
Co-expression
Functional
Protein-protein interactions
Diagnosis
Control
Major Depression
Sex
Male
Female
Functional enrichment
Functional enrichment
Motif enrichment
RcisTarget
Transcription factor binding enrichment
ChIP-enrichR
Expression ratios
<= -2.0
-0
>= 2.0

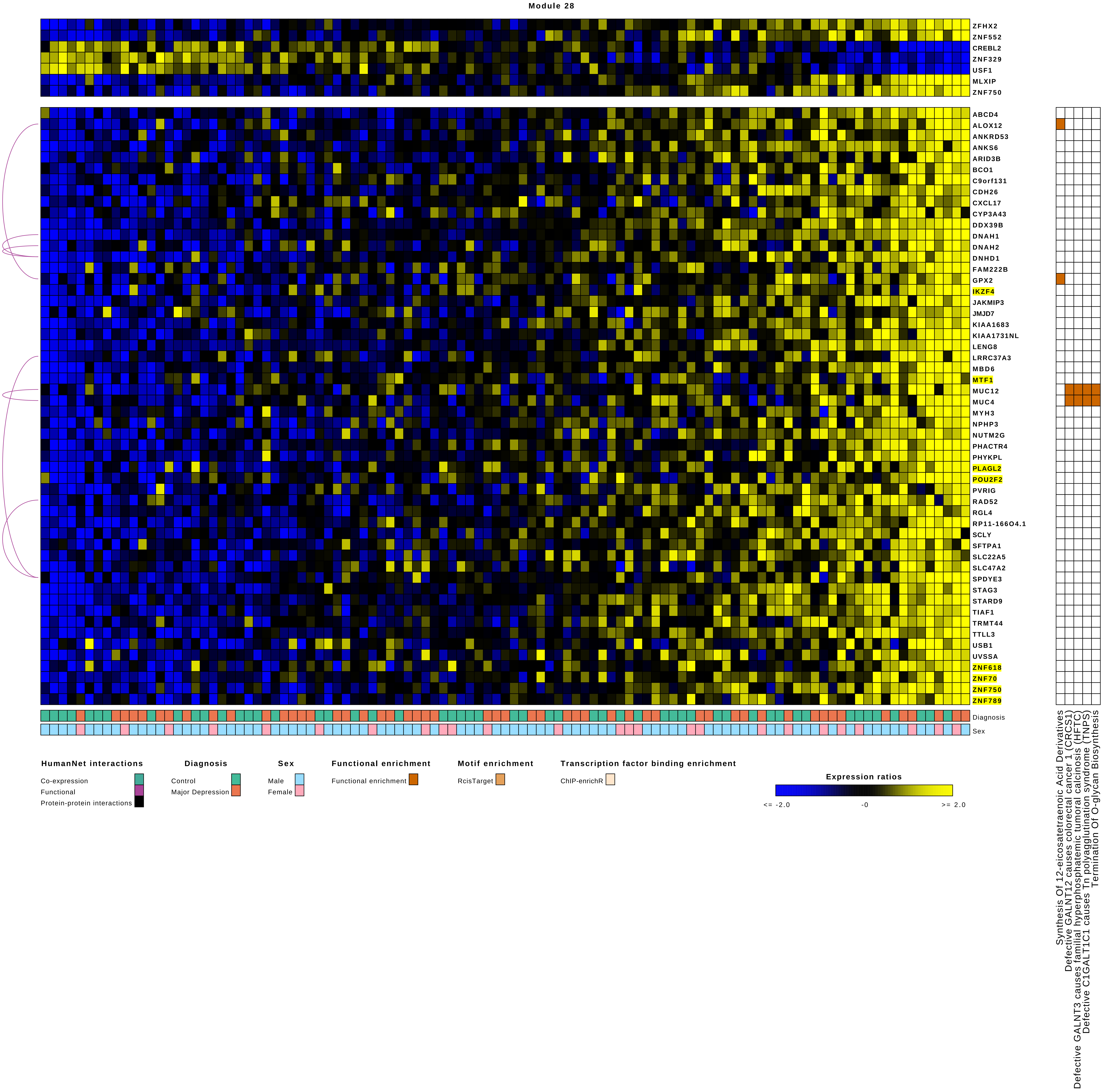

Module 28
ZFHX2
ZNF552
CREBL2
ZNF329
USF1
MLXIP
ZNF750
ABCD4
ALOX12
ANKRD53
ANKS6
ARID3B
BCO1
C9orf131
CDH26
CXCL17
CYP3A43
DDX39B
DNAH1
DNAH2
DNHD1
FAM222B
GPX2
IKZF4
JAKMIP3
JMJD7
KIAA1683
KIAA1731NL
LENG8
LRRC37A3
MBD6
MTF1
MUC12
MUC4
MYH3
NPHP3
NUTM2G
PHACTR4
PHYKPL
PLAGL2
POU2F2
PVRIG
RAD52
RGL4
RP11-166O4.1
SCLY
SFTPA1
SLC22A5
SLC47A2
SPDYE3
STAG3
STARD9
TIAF1
TRMT44
TTLL3
USB1
UVSSA
ZNF618
ZNF70
ZNF750
ZNF789
Diagnosis
Sex
Synthesis Of 12-eicosatetraenoic Acid Derivatives
Defective GALNT12 causes colorectal cancer 1 (CRCS1)
Defective GALNT3 causes familial hyperphosphatemic tumoral calcinosis (HFTC)
Defective C1GALT1C1 causes Tn polyagglutination syndrome (TNPS)
Termination Of O-glycan Biosynthesis
HumanNet interactions
Co-expression
Functional
Protein-protein interactions
Diagnosis
Control
Major Depression
Sex
Male
Female
Functional enrichment
Functional enrichment
Motif enrichment
RcisTarget
Transcription factor binding enrichment
ChIP-enrichR
Expression ratios
<= -2.0
-0
>= 2.0

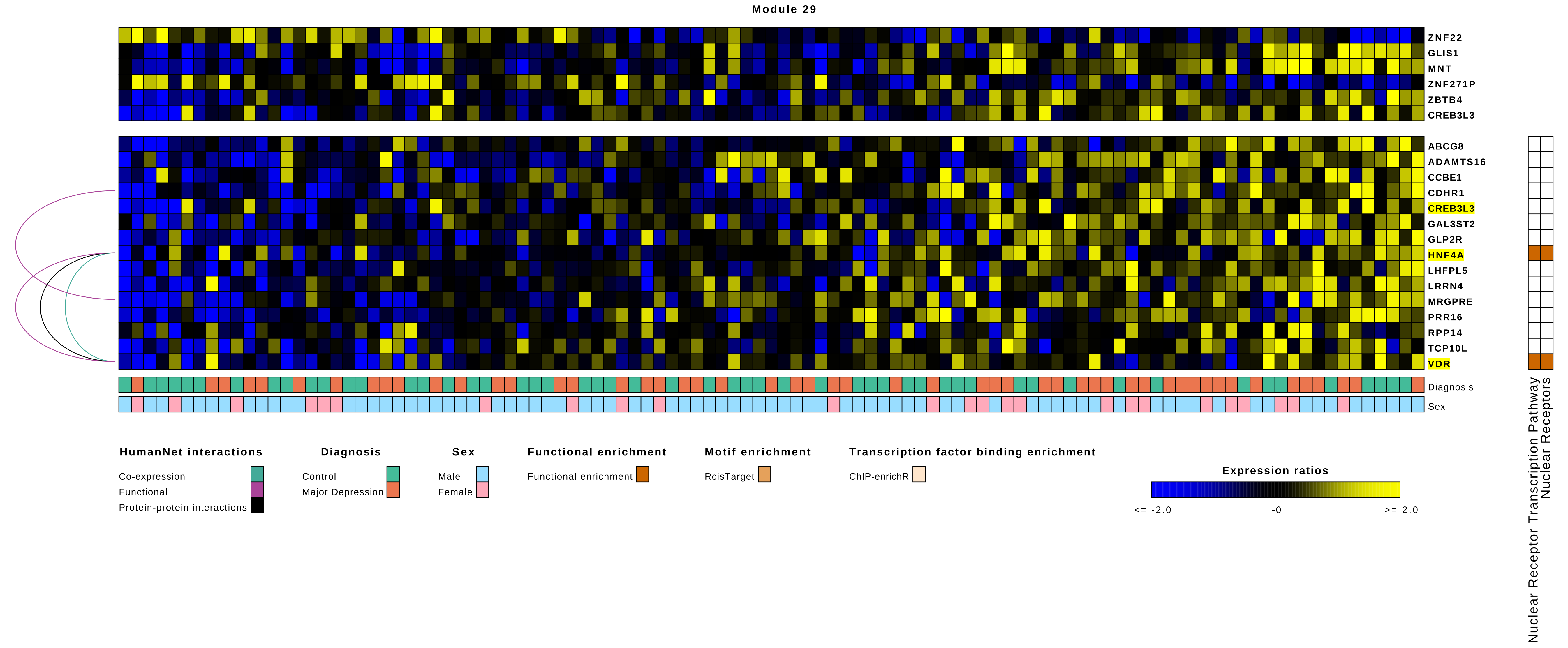
Module 29
ZNF22
GLIS1
MNT
ZNF271P
ZBTB4
CREB3L3
ABCG8
ADAMTS16
CCBE1
CDHR1
CREB3L3
GAL3ST2
GLP2R
HNF4A
LHFPL5
LRRN4
MRGPRE
PRR16
RPP14
TCP10L
VDR
Diagnosis
Sex
Nuclear Receptor Transcription Pathway
Nuclear Receptors
HumanNet interactions
Co-expression
Functional
Protein-protein interactions
Diagnosis
Control
Major Depression
Sex
Male
Female
Functional enrichment
Functional enrichment
Motif enrichment
RcisTarget
Transcription factor binding enrichment
ChIP-enrichR
Expression ratios
<= -2.0
-0
>= 2.0

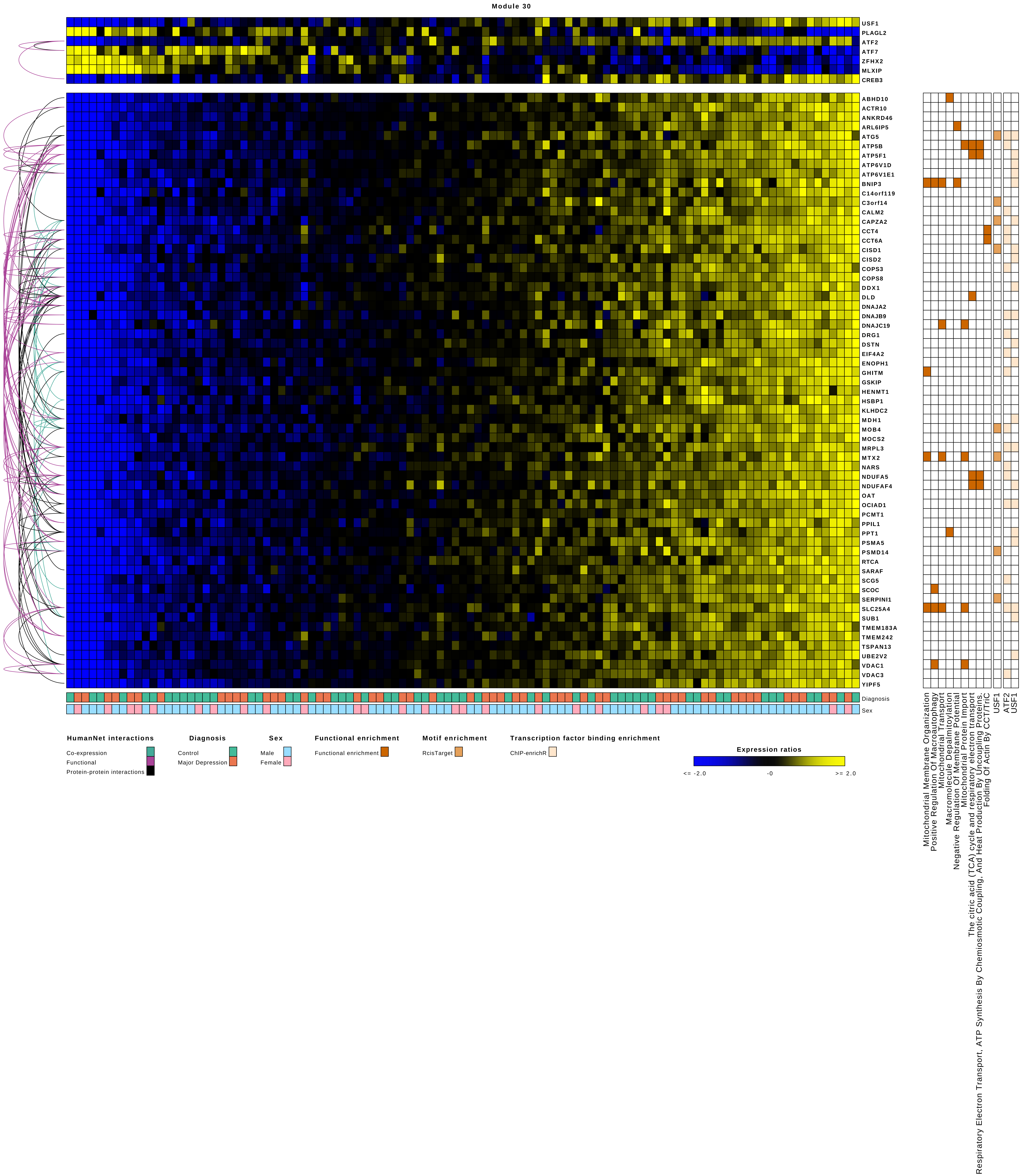

Module 30
USF1
PLAGL2
ATF2
ATF7
ZFHX2
MLXIP
CREB3
ABHD10
ACTR10
ANKRD46
ARL6IP5
ATG5
ATP5B
ATP5F1
ATP6V1D
ATP6V1E1
BNIP3
C14orf119
C3orf14
CALM2
CAPZA2
CCT4
CCT6A
CISD1
CISD2
COPS3
COPS8
DDX1
DLD
DNAJA2
DNAJB9
DNAJC19
DRG1
DSTN
EIF4A2
ENOPH1
GHITM
GSKIP
HENMT1
HSBP1
KLHDC2
MDH1
MOB4
MOCS2
MRPL3
MTX2
NARS
NDUFA5
NDUFAF4
OAT
OCIAD1
PCMT1
PPIL1
PPT1
PSMA5
PSMD14
RTCA
SARAF
SCG5
SCOC
SERPINI1
SLC25A4
SUB1
TMEM183A
TMEM242
TSPAN13
UBE2V2
VDAC1
VDAC3
YIPF5
Diagnosis
Sex
Mitochondrial Membrane Organization
Positive Regulation Of Macroautophagy
Mitochondrial Transport
Macromolecule Depalmitoylation
Negative Regulation Of Membrane Potential
Mitochondrial Protein Import
The citric acid (TCA) cycle and respiratory electron transport
Respiratory Electron Transport, ATP Synthesis By Chemiosmotic Coupling, And Heat Production By Uncoupling Proteins.
Folding Of Actin By CCT/TriC
USF1
ATF2
USF1
HumanNet interactions
Co-expression
Functional
Protein-protein interactions
Diagnosis
Control
Major Depression
Sex
Male
Female
Functional enrichment
Functional enrichment
Motif enrichment
RcisTarget
Transcription factor binding enrichment
ChIP-enrichR
Expression ratios
<= -2.0
-0
>= 2.0

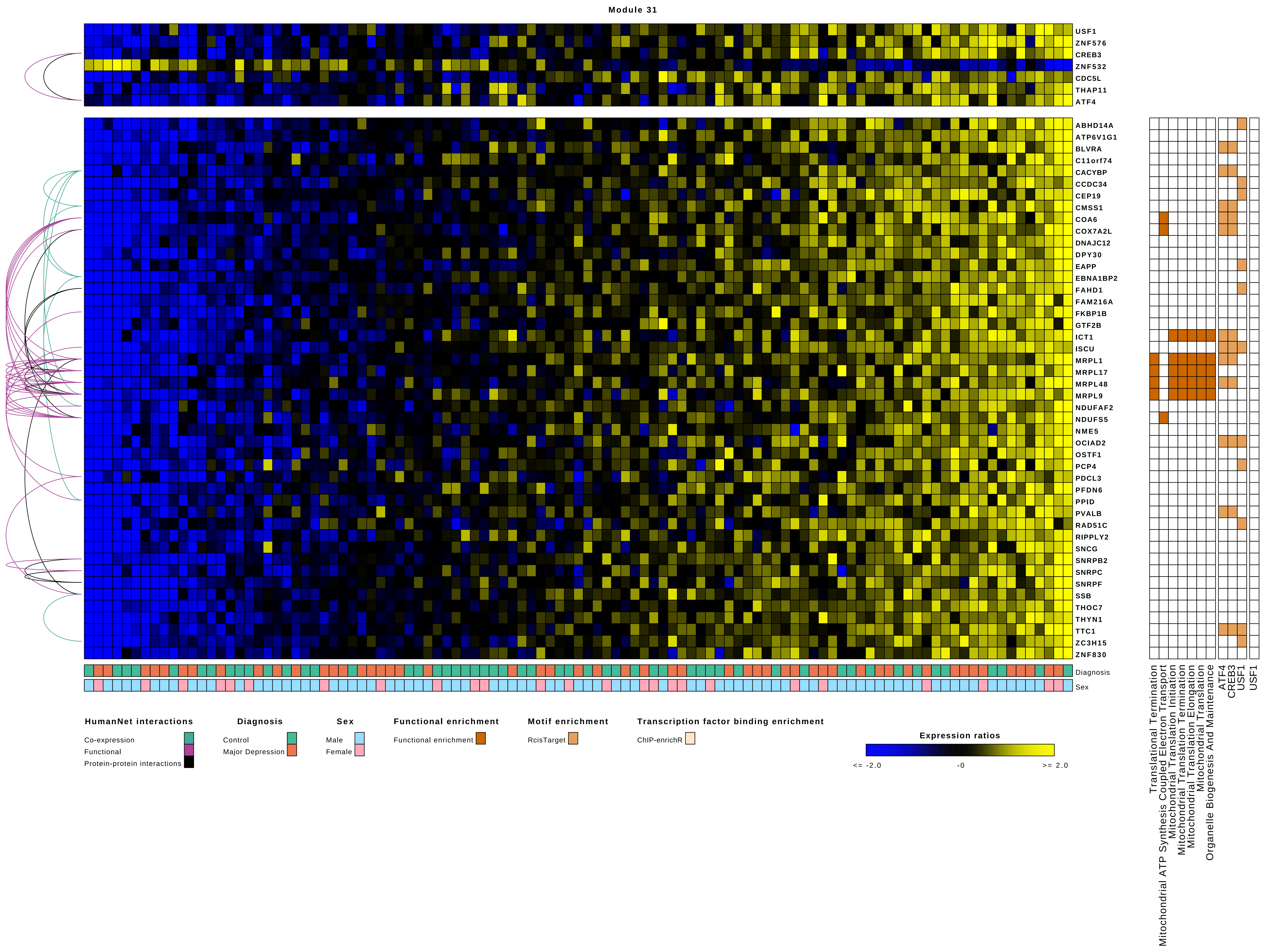

Module 31
USF1
ZNF576
CREB3
ZNF532
CDC5L
THAP11
ATF4
ABHD14A
ATP6V1G1
BLVRA
C11orf74
CACYBP
CCDC34
CEP19
CMSS1
COA6
COX7A2L
DNAJC12
DPY30
EAPP
EBNA1BP2
FAHD1
FAM216A
FKBP1B
GTF2B
ICT1
ISCU
MRPL1
MRPL17
MRPL48
MRPL9
NDUFAF2
NDUFS5
NME5
OCIAD2
OSTF1
PCP4
PDCL3
PFDN6
PPID
PVALB
RAD51C
RIPPLY2
SNCG
SNRPB2
SNRPC
SNRPF
SSB
THOC7
THYN1
TTC1
ZC3H15
ZNF830
Diagnosis
Sex
Translational Termination
Mitochondrial ATP Synthesis Coupled Electron Transport
Mitochondrial Translation Initiation
Mitochondrial Translation Termination
Mitochondrial Translation Elongation
Mitochondrial Translation
Organelle Biogenesis And Maintenance
ATF4
CREB3
USF1
USF1
HumanNet interactions
Co-expression
Functional
Protein-protein interactions
Diagnosis
Control
Major Depression
Sex
Male
Female
Functional enrichment
Functional enrichment
Motif enrichment
RcisTarget
Transcription factor binding enrichment
ChIP-enrichR
Expression ratios
<= -2.0
-0
>= 2.0

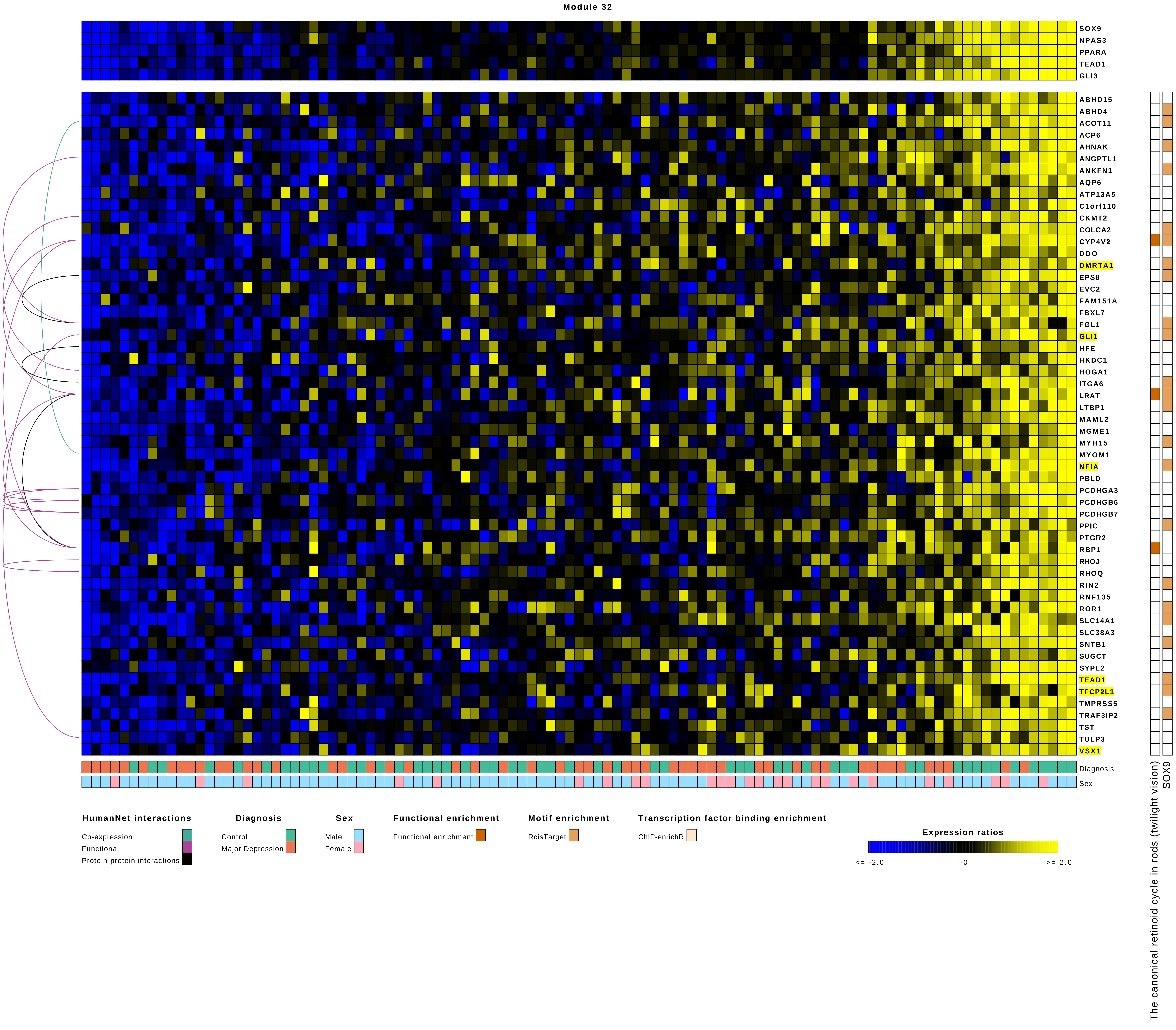

Module 32
SOX9
NPAS3
PPARA
TEAD1
GLI3
ABHD15
ABHD4
ACOT11
ACP6
AHNAK
ANGPTL1
ANKFN1
AQP6
ATP13A5
C1orf110
CKMT2
COLCA2
CYP4V2
DDO
DMRTA1
EPS8
EVC2
FAM151A
FBXL7
FGL1
GLI1
HFE
HKDC1
HOGA1
ITGA6
LRAT
LTBP1
MAML2
MGME1
MYH15
MYOM1
NFIA
PBLD
PCDHGA3
PCDHGB6
PCDHGB7
PPIC
PTGR2
RBP1
RHOJ
RHOQ
RIN2
RNF135
ROR1
SLC14A1
SLC38A3
SNTB1
SUGCT
SYPL2
TEAD1
TFCP2L1
TMPRSS5
TRAF3IP2
TST
TULP3
VSX1
Diagnosis
Sex
The canonical retinoid cycle in rods (twilight vision)
SOX9
HumanNet interactions
Co-expression
Functional
Protein-protein interactions
Diagnosis
Control
Major Depression
Sex
Male
Female
Functional enrichment
Functional enrichment
Motif enrichment
RcisTarget
Transcription factor binding enrichment
ChIP-enrichR
Expression ratios
<= -2.0
-0
>= 2.0

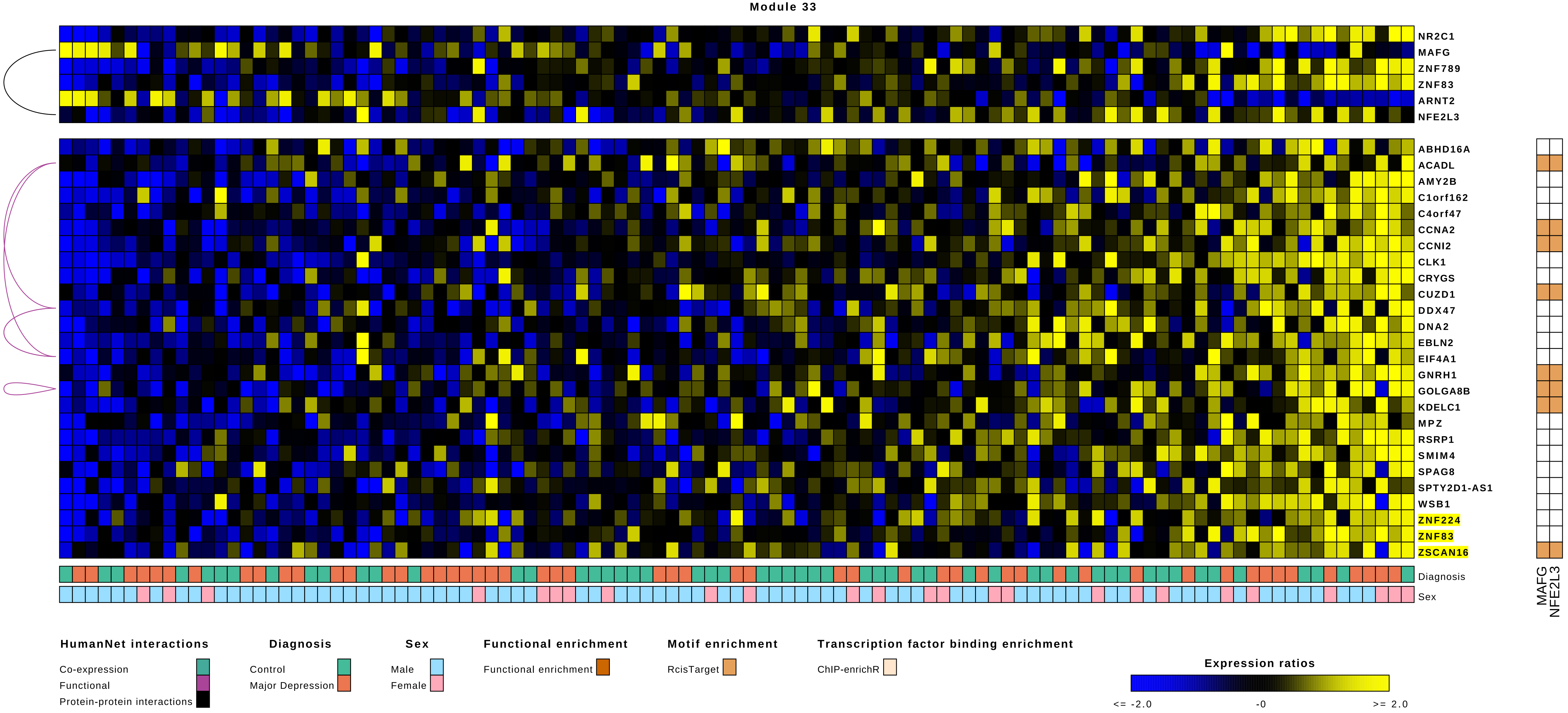

Module 33
NR2C1
MAFG
ZNF789
ZNF83
ARNT2
NFE2L3
ABHD16A
ACADL
AMY2B
C1orf162
C4orf47
CCNA2
CCNI2
CLK1
CRYGS
CUZD1
DDX47
DNA2
EBLN2
EIF4A1
GNRH1
GOLGA8B
KDELC1
MPZ
RSRP1
SMIM4
SPAG8
SPTY2D1-AS1
WSB1
ZNF224
ZNF83
ZSCAN16
Diagnosis
Sex
MAFG
NFE2L3
HumanNet interactions
Co-expression
Functional
Protein-protein interactions
Diagnosis
Control
Major Depression
Sex
Male
Female
Functional enrichment
Functional enrichment
Motif enrichment
RcisTarget
Transcription factor binding enrichment
ChIP-enrichR
Expression ratios
<= -2.0
-0
>= 2.0

Module 34

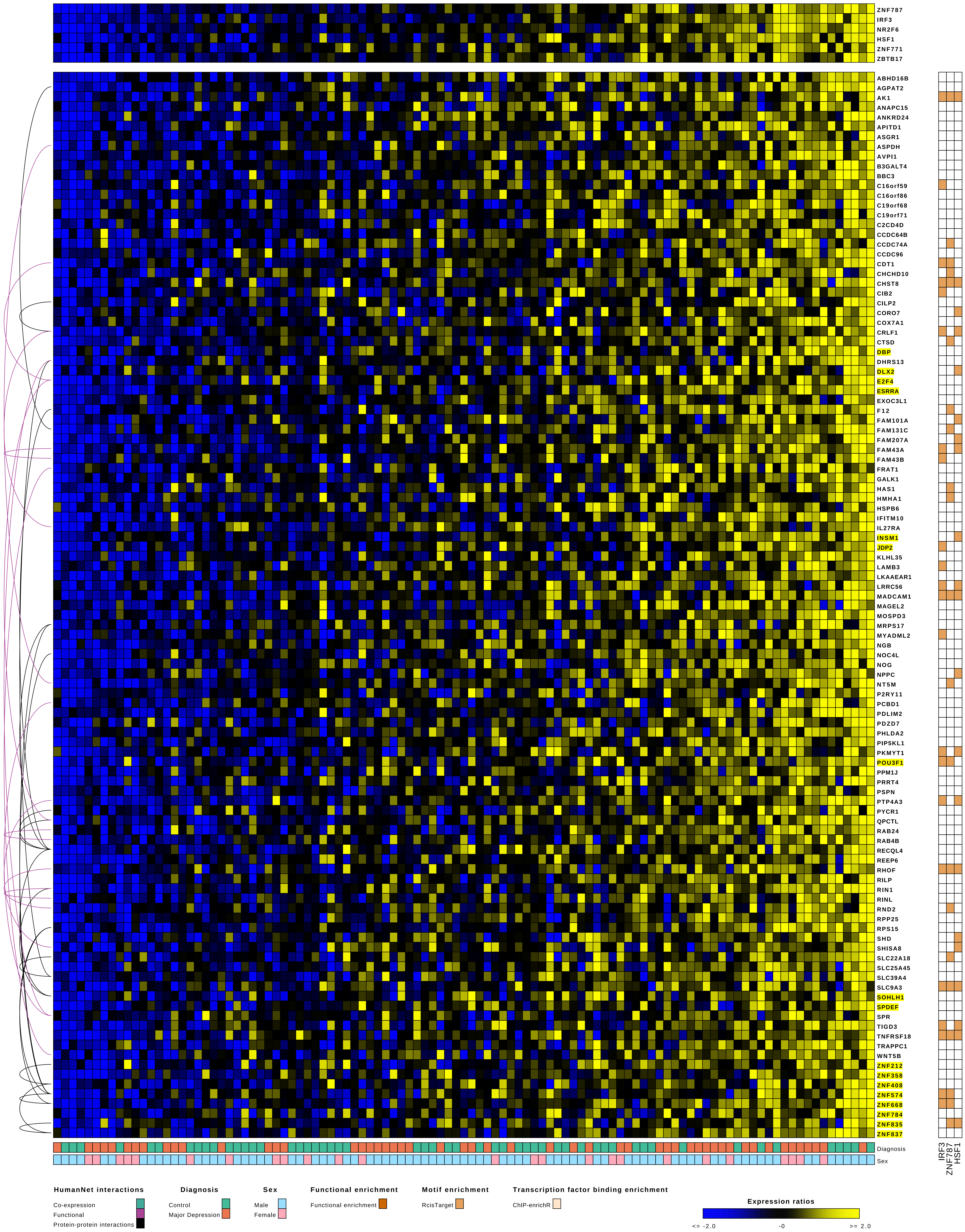

ZNF787
IRF3
NR2F6
HSF1
ZNF771
ZBTB17
ABHD16B
AGPAT2
AK1
ANAPC15
ANKRD24
APITD1
ASGR1
ASPDH
AVPI1
B3GALT4
BBC3
C16orf59
C16orf86
C19orf68
C19orf71
C2CD4D
CCDC64B
CCDC74A
CCDC96
CDT1
CHCHD10
CHST8
CIB2
CILP2
CORO7
COX7A1
CRLF1
CTSD
DBP
DHRS13
DLX2
E2F4
ESRRA
EXOC3L1
F12
FAM101A
FAM131C
FAM207A
FAM43A
FAM43B
FRAT1
GALK1
HAS1
HMHA1
HSPB6
IFITM10
IL27RA
INSM1
JDP2
KLHL35
LAMB3
LKAAEAR1
LRRC56
MADCAM1
MAGEL2
MOSPD3
MRPS17
MYADML2
NGB
NOC4L
NOG
NPPC
NT5M
P2RY11
PCBD1
PDLIM2
PDZD7
PHLDA2
PIP5KL1
PKMYT1
POU3F1
PPM1J
PRRT4
PSPN
PTP4A3
PYCR1
QPCTL
RAB24
RAB4B
RECQL4
REEP6
RHOF
RILP
RIN1
RINL
RND2
RPP25
RPS15
SHD
SHISA8
SLC22A18
SLC25A45
SLC39A4
SLC9A3
SOHLH1
SPDEF
SPR
TIGD3
TNFRSF18
TRAPPC1
WNT5B
ZNF212
ZNF358
ZNF408
ZNF574
ZNF668
ZNF784
ZNF835
ZNF837
Diagnosis
Sex
IRF3
ZNF787
HSF1
HumanNet interactions
Co-expression
Functional
Protein-protein interactions
Diagnosis
Control
Major Depression
Sex
Male
Female
Functional enrichment
Functional enrichment
Motif enrichment
RcisTarget
Transcription factor binding enrichment
ChIP-enrichR
Expression ratios
<= -2.0
-0
>= 2.0

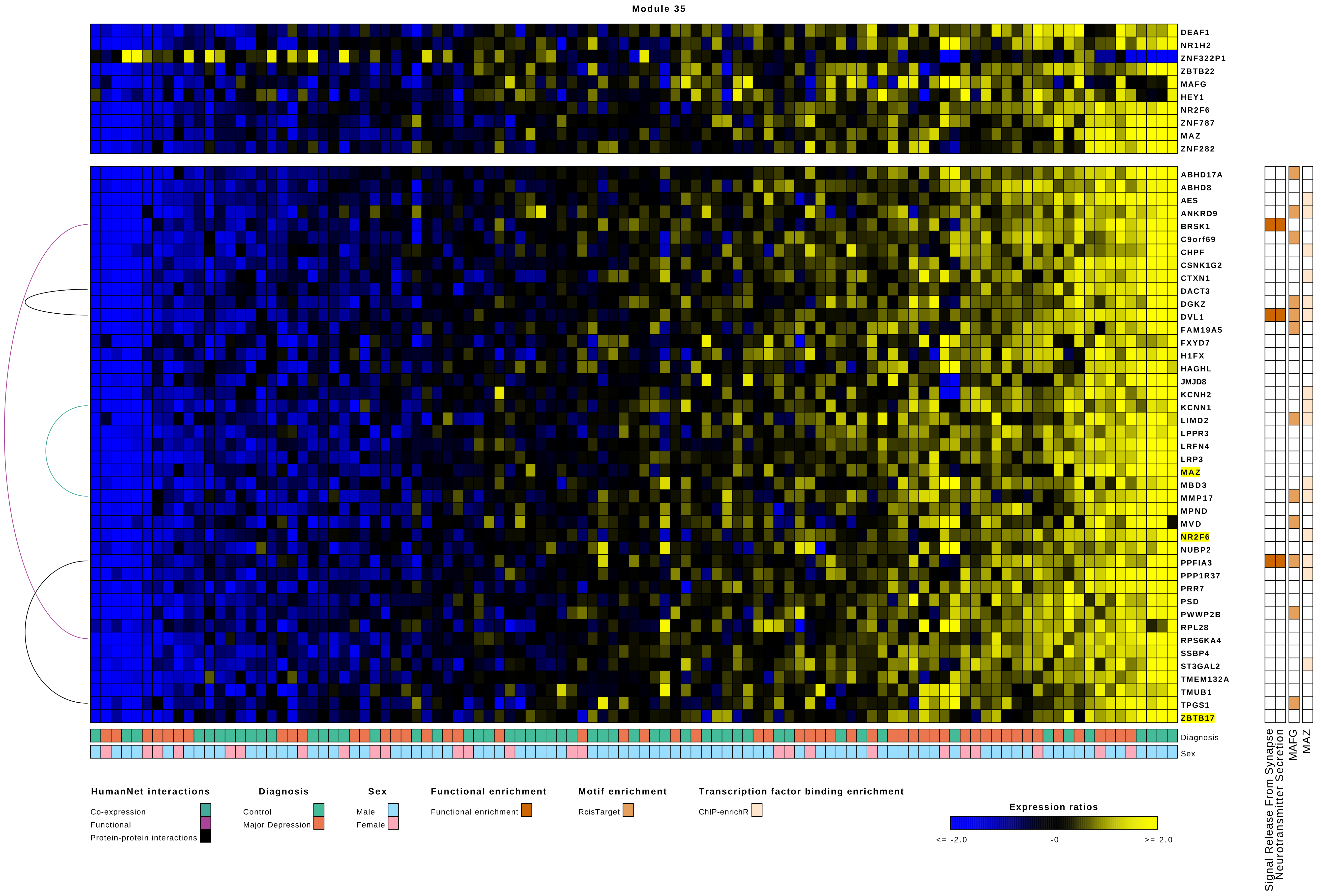

Module 35
DEAF1
NR1H2
ZNF322P1
ZBTB22
MAFG
HEY1
NR2F6
ZNF787
MAZ
ZNF282
ABHD17A
ABHD8
AES
ANKRD9
BRSK1
C9orf69
CHPF
CSNK1G2
CTXN1
DACT3
DGKZ
DVL1
FAM19A5
FXYD7
H1FX
HAGHL
JMJD8
KCNH2
KCNN1
LIMD2
LPPR3
LRFN4
LRP3
MAZ
MBD3
MMP17
MPND
MVD
NR2F6
NUBP2
PPFIA3
PPP1R37
PRR7
PSD
PWWP2B
RPL28
RPS6KA4
SSBP4
ST3GAL2
TMEM132A
TMUB1
TPGS1
ZBTB17
Diagnosis
Sex
Signal Release From Synapse
Neurotransmitter Secretion
MAFG
MAZ
HumanNet interactions
Co-expression
Functional
Protein-protein interactions
Diagnosis
Control
Major Depression
Sex
Male
Female
Functional enrichment
Functional enrichment
Motif enrichment
RcisTarget
Transcription factor binding enrichment
ChIP-enrichR
Expression ratios
<= -2.0
-0
>= 2.0

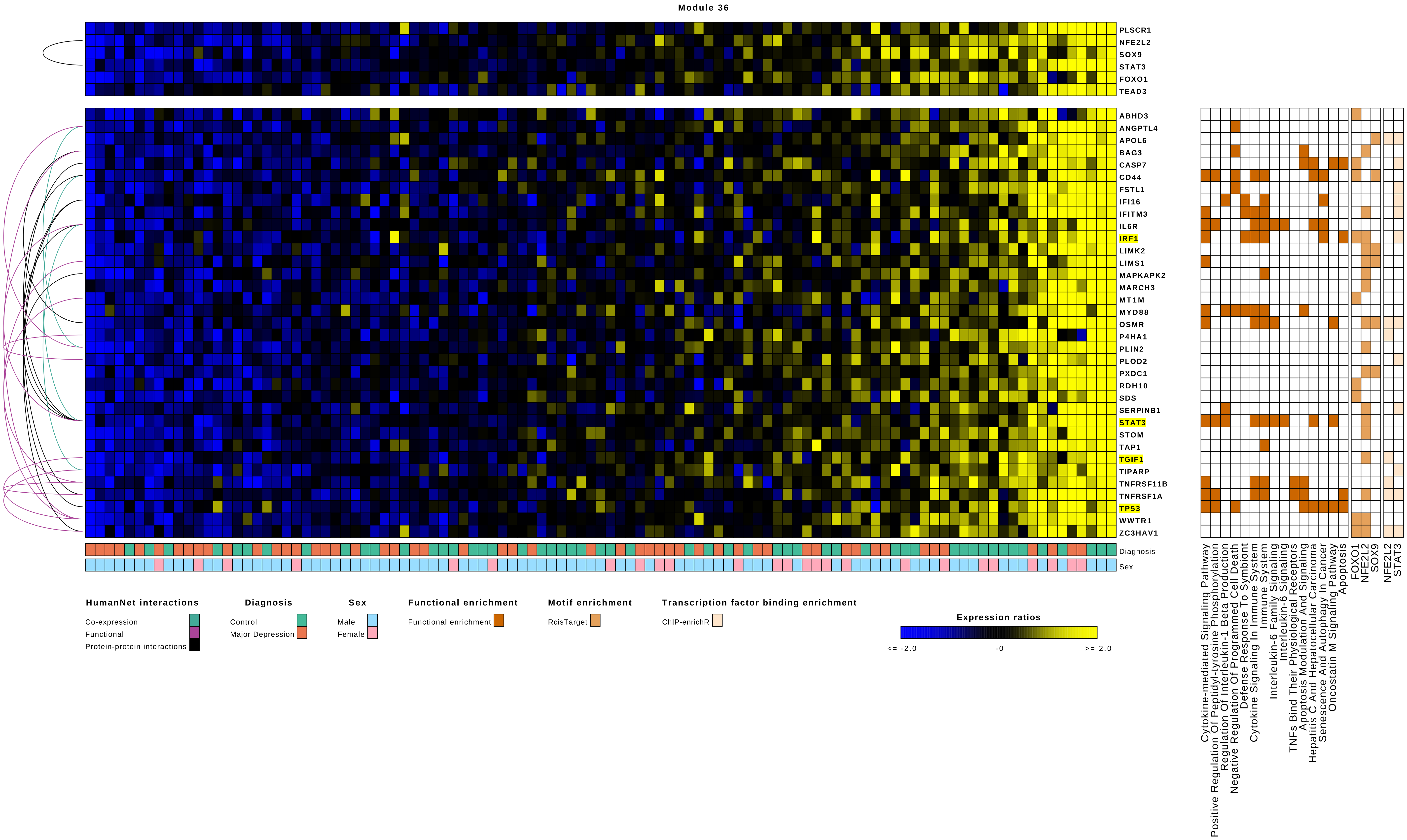

Module 36
PLSCR1
NFE2L2
SOX9
STAT3
FOXO1
TEAD3
ABHD3
ANGPTL4
APOL6
BAG3
CASP7
CD44
FSTL1
IFI16
IFITM3
IL6R
IRF1
LIMK2
LIMS1
MAPKAPK2
MARCH3
MT1M
MYD88
OSMR
P4HA1
PLIN2
PLOD2
PXDC1
RDH10
SDS
SERPINB1
STAT3
STOM
TAP1
TGIF1
TIPARP
TNFRSF11B
TNFRSF1A
TP53
WWTR1
ZC3HAV1
Diagnosis
Sex
Cytokine-mediated Signaling Pathway
Positive Regulation Of Peptidyl-tyrosine Phosphorylation
Regulation Of Interleukin-1 Beta Production
Negative Regulation Of Programmed Cell Death
Defense Response To Symbiont
Cytokine Signaling In Immune System
Immune System
Interleukin-6 Family Signaling
Interleukin-6 Signaling
TNFs Bind Their Physiological Receptors
Apoptosis Modulation And Signaling
Hepatitis C And Hepatocellular Carcinoma
Senescence And Autophagy In Cancer
Oncostatin M Signaling Pathway
Apoptosis
FOXO1
NFE2L2
SOX9
NFE2L2
STAT3
HumanNet interactions
Co-expression
Functional
Protein-protein interactions
Diagnosis
Control
Major Depression
Sex
Male
Female
Functional enrichment
Functional enrichment
Motif enrichment
RcisTarget
Transcription factor binding enrichment
ChIP-enrichR
Expression ratios
<= -2.0
-0
>= 2.0

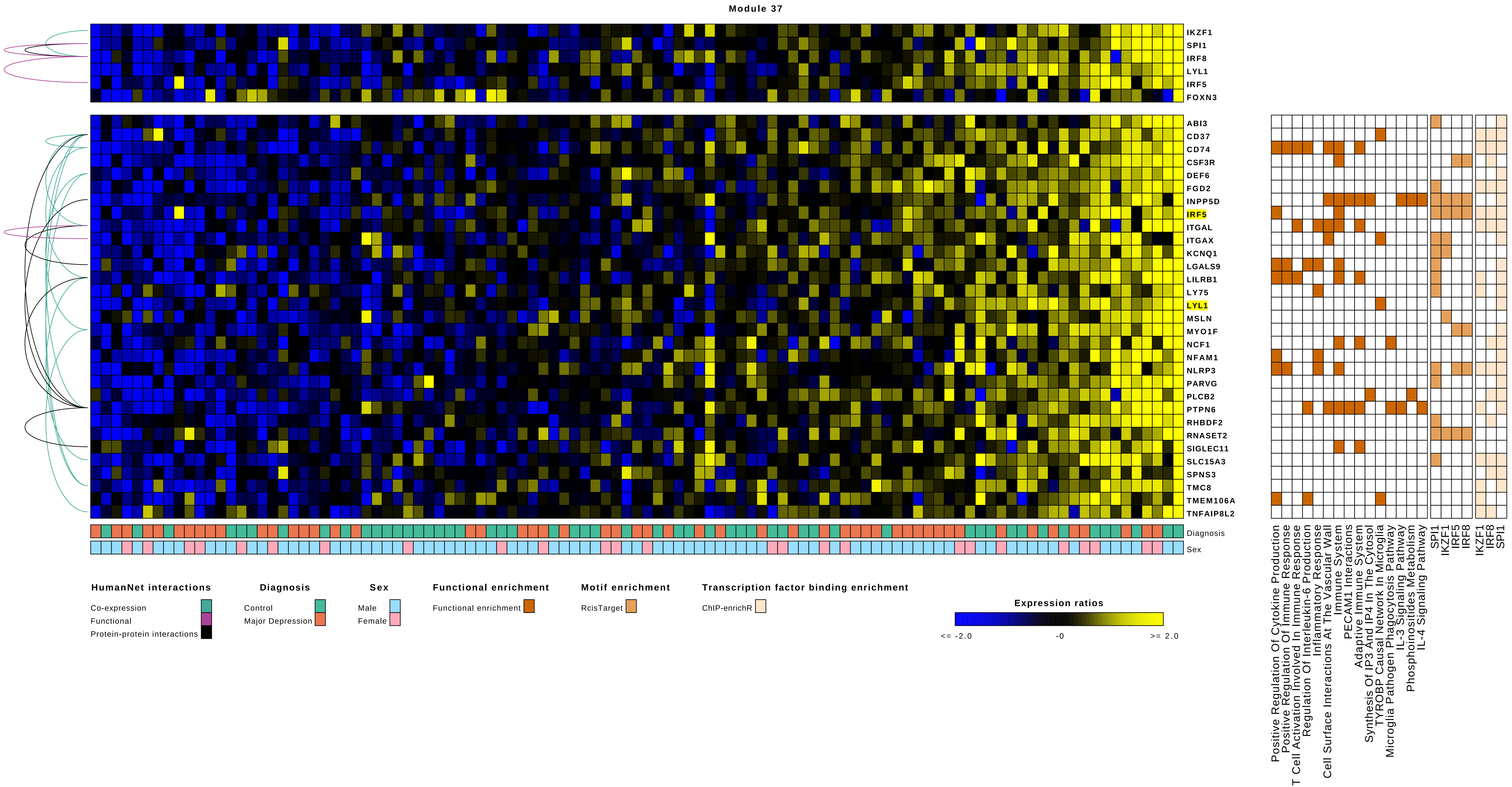

Module 37
IKZF1
SPI1
IRF8
LYL1
IRF5
FOXN3
ABI3
CD37
CD74
CSF3R
DEF6
FGD2
INPP5D
IRF5
ITGAL
ITGAX
KCNQ1
LGALS9
LILRB1
LY75
LYL1
MSLN
MYO1F
NCF1
NFAM1
NLRP3
PARVG
PLCB2
PTPN6
RHBDF2
RNASET2
SIGLEC11
SLC15A3
SPNS3
TMC8
TMEM106A
TNFAIP8L2
Diagnosis
Sex
HumanNet interactions
Co-expression
Functional
Protein-protein interactions
Diagnosis
Control
Major Depression
Sex
Male
Female
Functional enrichment
Functional enrichment
Motif enrichment
RcisTarget
Transcription factor binding enrichment
ChIP-enrichR
Expression ratios
<= -2.0
-0
>= 2.0
Positive Regulation Of Cytokine Production
Positive Regulation Of Immune Response
T Cell Activation Involved In Immune Response
Regulation Of Interleukin-6 Production
Inflammatory Response
Cell Surface Interactions At The Vascular Wall
Immune System
PECAM1 Interactions
Adaptive Immune System
Synthesis Of IP3 And IP4 In The Cytosol
TYROBP Causal Network In Microglia
Microglia Pathogen Phagocytosis Pathway
IL-3 Signaling Pathway
Phosphoinositides Metabolism
IL-4 Signaling Pathway
SPI1
IKZF1
IRF5
IRF8
IKZF1
IRF8
SPI1

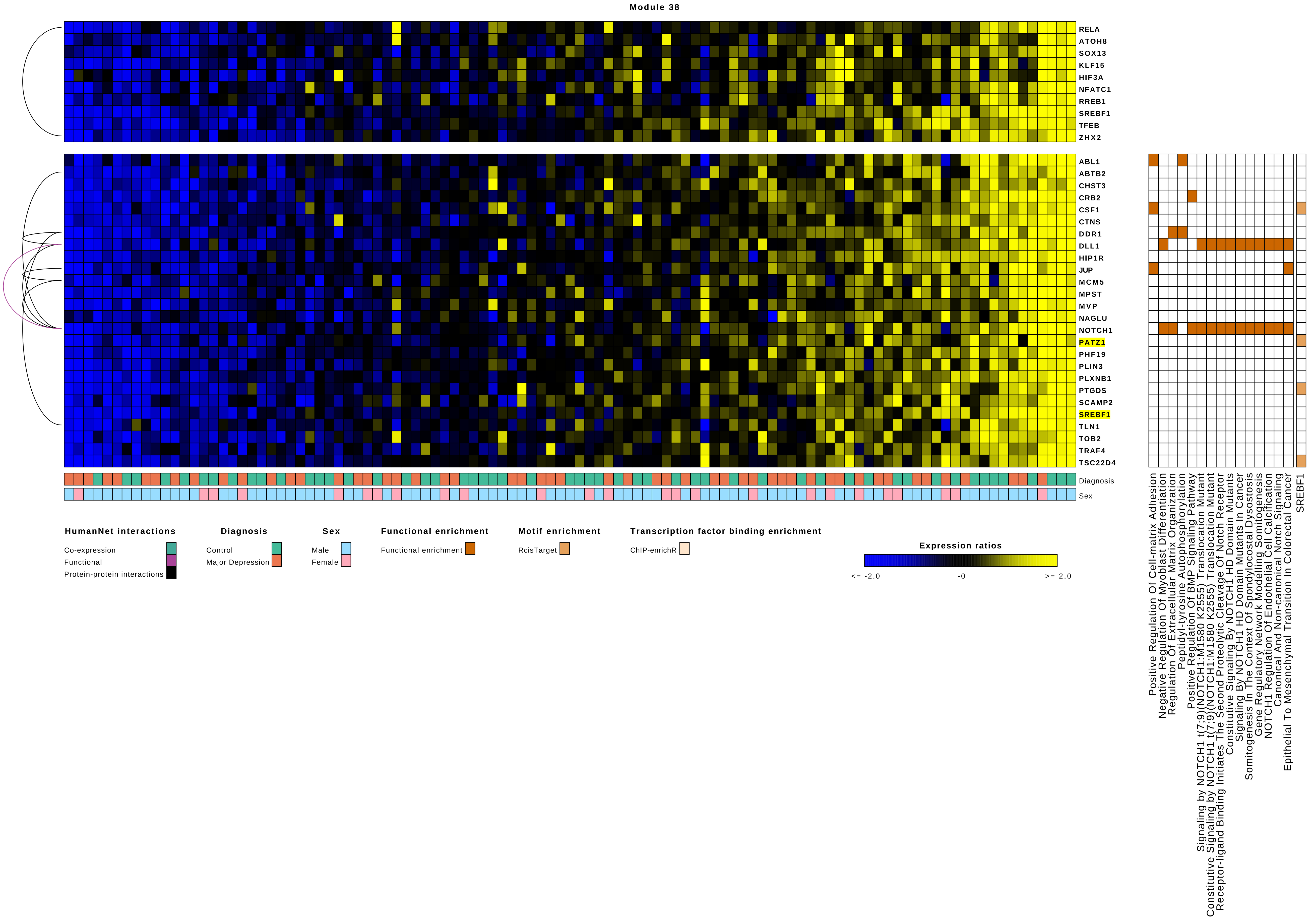

Module 38
RELA
ATOH8
SOX13
KLF15
HIF3A
NFATC1
RREB1
SREBF1
TFEB
ZHX2
ABL1
ABTB2
CHST3
CRB2
CSF1
CTNS
DDR1
DLL1
HIP1R
JUP
MCM5
MPST
MVP
NAGLU
NOTCH1
PATZ1
PHF19
PLIN3
PLXNB1
PTGDS
SCAMP2
SREBF1
TLN1
TOB2
TRAF4
TSC22D4
Diagnosis
Sex
Positive Regulation Of Cell-matrix Adhesion
Negative Regulation Of Myoblast Differentiation
Regulation Of Extracellular Matrix Organization
Peptidyl-tyrosine Autophosphorylation
Positive Regulation Of BMP Signaling Pathway
Signaling by NOTCH1 t(7;9)(NOTCH1:M1580 K2555) Translocation Mutant
Constitutive Signaling by NOTCH1 t(7;9)(NOTCH1:M1580 K2555) Translocation Mutant
Receptor-ligand Binding Initiates The Second Proteolytic Cleavage Of Notch Receptor
Constitutive Signaling By NOTCH1 HD Domain Mutants
Signaling By NOTCH1 HD Domain Mutants In Cancer
Somitogenesis In The Context Of Spondylocostal Dysostosis
Gene Regulatory Network Modelling Somitogenesis
NOTCH1 Regulation Of Endothelial Cell Calcification
Canonical And Non-canonical Notch Signaling
Epithelial To Mesenchymal Transition In Colorectal Cancer
SREBF1
HumanNet interactions
Co-expression
Functional
Protein-protein interactions
Diagnosis
Control
Major Depression
Sex
Male
Female
Functional enrichment
Functional enrichment
Motif enrichment
RcisTarget
Transcription factor binding enrichment
ChIP-enrichR
Expression ratios
<= -2.0
-0
>= 2.0

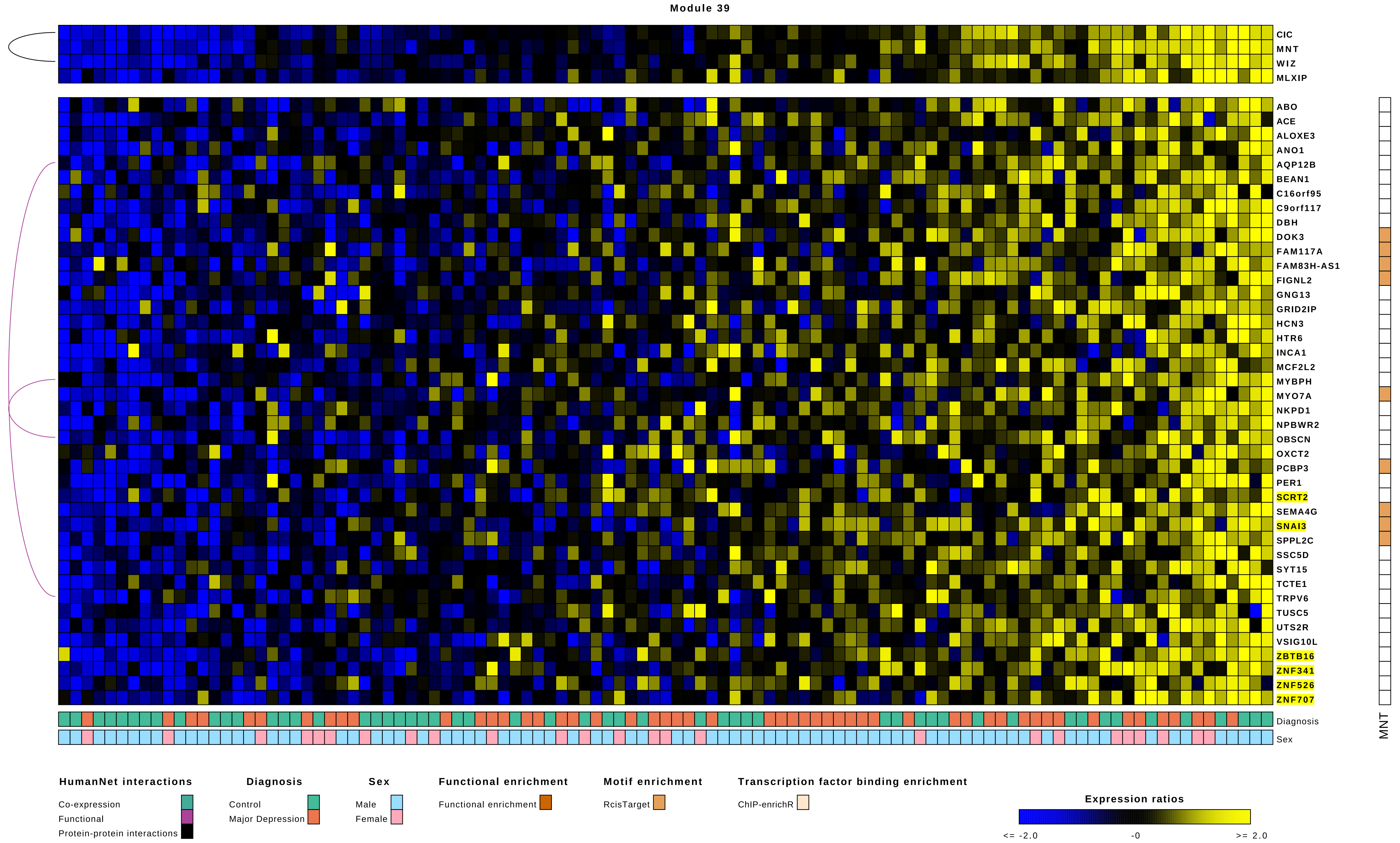
Module 39
CIC
MNT
WIZ
MLXIP
ABO
ACE
ALOXE3
ANO1
AQP12B
BEAN1
C16orf95
C9orf117
DBH
DOK3
FAM117A
FAM83H-AS1
FIGNL2
GNG13
GRID2IP
HCN3
HTR6
INCA1
MCF2L2
MYBPH
MYO7A
NKPD1
NPBWR2
OBSCN
OXCT2
PCBP3
PER1
SCRT2
SEMA4G
SNAI3
SPPL2C
SSC5D
SYT15
TCTE1
TRPV6
TUSC5
UTS2R
VSIG10L
ZBTB16
ZNF341
ZNF526
ZNF707
Diagnosis
Sex
MNT
HumanNet interactions
Co-expression
Functional
Protein-protein interactions
Diagnosis
Control
Major Depression
Sex
Male
Female
Functional enrichment
Functional enrichment
Motif enrichment
RcisTarget
Transcription factor binding enrichment
ChIP-enrichR
Expression ratios
<= -2.0
-0
>= 2.0

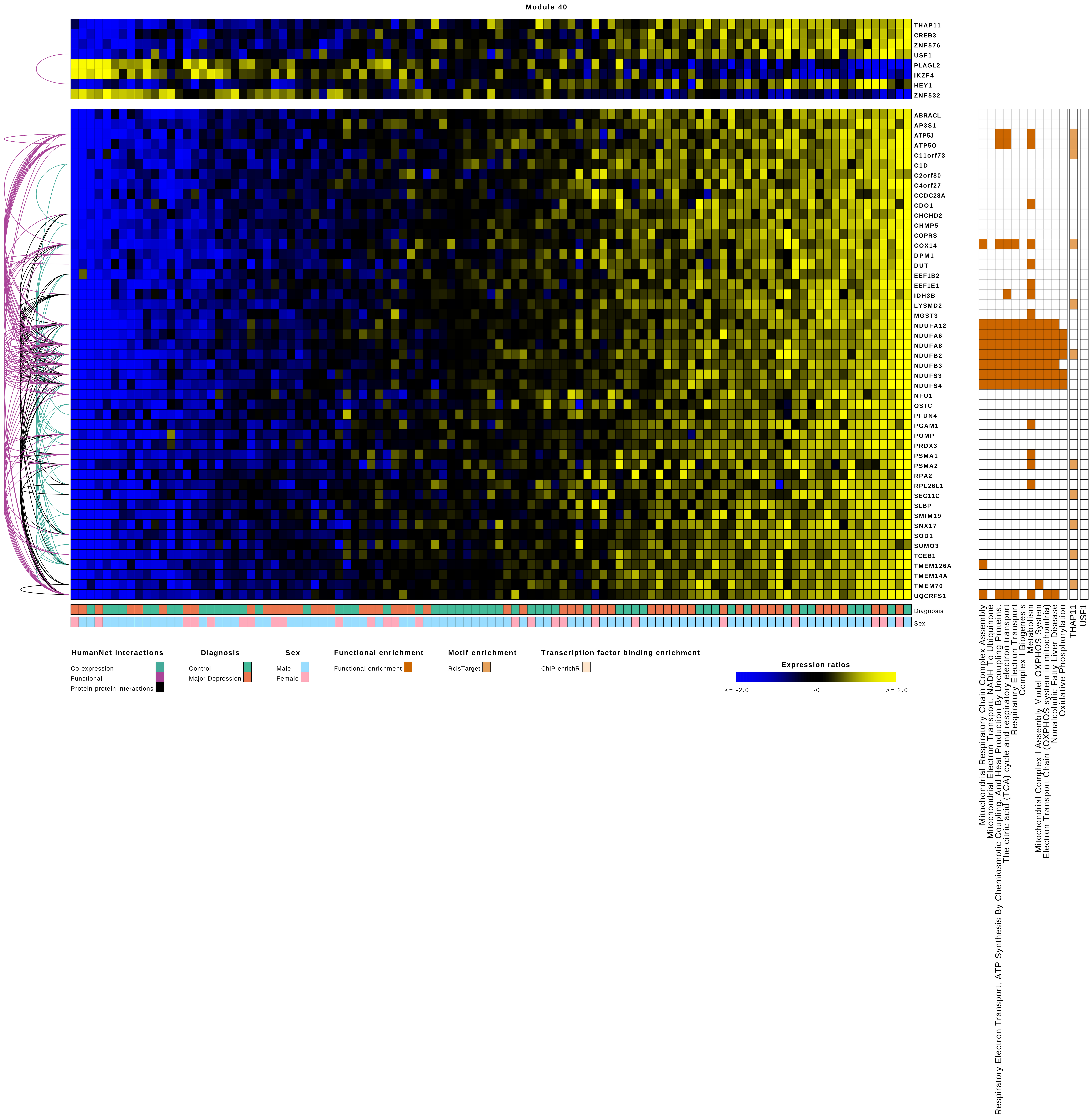
Module 40
THAP11
CREB3
ZNF576
USF1
PLAGL2
IKZF4
HEY1
ZNF532
ABRACL
AP3S1
ATP5J
ATP5O
C11orf73
C1D
C2orf80
C4orf27
CCDC28A
CDO1
CHCHD2
CHMP5
COPRS
COX14
DPM1
DUT
EEF1B2
EEF1E1
IDH3B
LYSMD2
MGST3
NDUFA12
NDUFA6
NDUFA8
NDUFB2
NDUFB3
NDUFS3
NDUFS4
NFU1
OSTC
PFDN4
PGAM1
POMP
PRDX3
PSMA1
PSMA2
RPA2
RPL26L1
SEC11C
SLBP
SMIM19
SNX17
SOD1
SUMO3
TCEB1
TMEM126A
TMEM14A
TMEM70
UQCRFS1
Diagnosis
Sex
HumanNet interactions
Co-expression
Functional
Protein-protein interactions
Diagnosis
Control
Major Depression
Sex
Male
Female
Functional enrichment
Functional enrichment
Motif enrichment
RcisTarget
Transcription factor binding enrichment
ChIP-enrichR
Expression ratios
<= -2.0
-0
>= 2.0
Mitochondrial Respiratory Chain Complex Assembly
Mitochondrial Electron Transport, NADH To Ubiquinone
Respiratory Electron Transport, ATP Synthesis By Chemiosmotic Coupling, And Heat Production By Uncoupling Proteins
The citric acid (TCA) cycle and respiratory electron transport
Respiratory Electron Transport
Complex I Biogenesis
Metabolism
Mitochondrial Complex I Assembly Model OXPHOS System
Electron Transport Chain (OXPHOS system in mitochondria)
Nonalcoholic Fatty Liver Disease
Oxidative Phosphorylation
THAP11
USF1

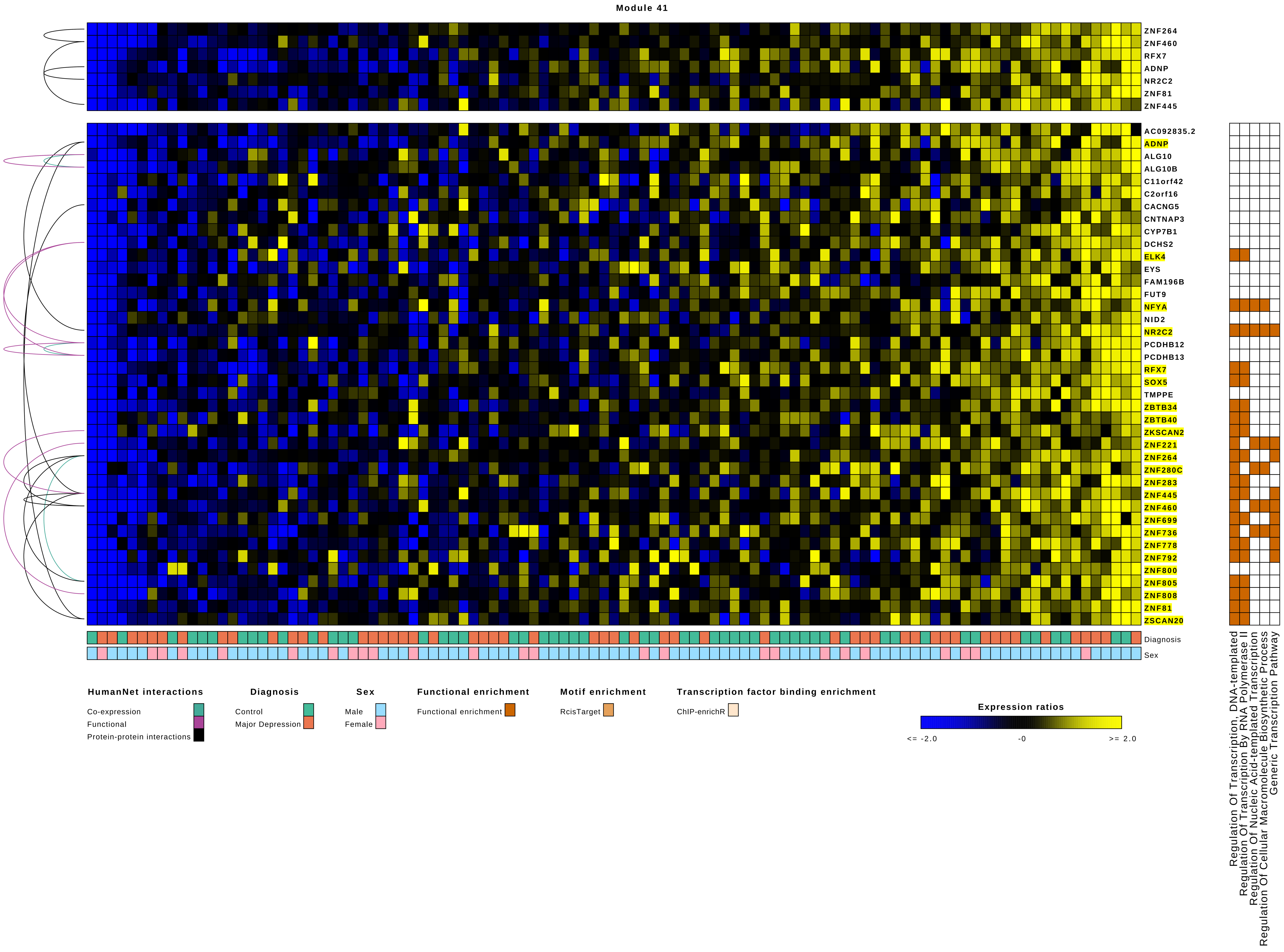

Module 41
ZNF264
ZNF460
RFX7
ADNP
NR2C2
ZNF81
ZNF445
AC092835.2
ADNP
ALG10
ALG10B
C11orf42
C2orf16
CACNG5
CNTNAP3
CYP7B1
DCHS2
ELK4
EYS
FAM196B
FUT9
NFYA
NID2
NR2C2
PCDHB12
PCDHB13
RFX7
SOX5
TMPPE
ZBTB34
ZBTB40
ZKSCAN2
ZNF221
ZNF264
ZNF280C
ZNF283
ZNF445
ZNF460
ZNF699
ZNF736
ZNF778
ZNF792
ZNF800
ZNF805
ZNF808
ZNF81
ZSCAN20
Diagnosis
Sex
Regulation Of Transcription, DNA-templated
Regulation Of Transcription By RNA Polymerase II
Regulation Of Nucleic Acid-templated Transcription
Regulation Of Cellular Macromolecule Biosynthetic Process
Generic Transcription Pathway
HumanNet interactions
Co-expression
Functional
Protein-protein interactions
Diagnosis
Control
Major Depression
Sex
Male
Female
Functional enrichment
Functional enrichment
Motif enrichment
RcisTarget
Transcription factor binding enrichment
ChIP-enrichR
Expression ratios
<= -2.0
-0
>= 2.0

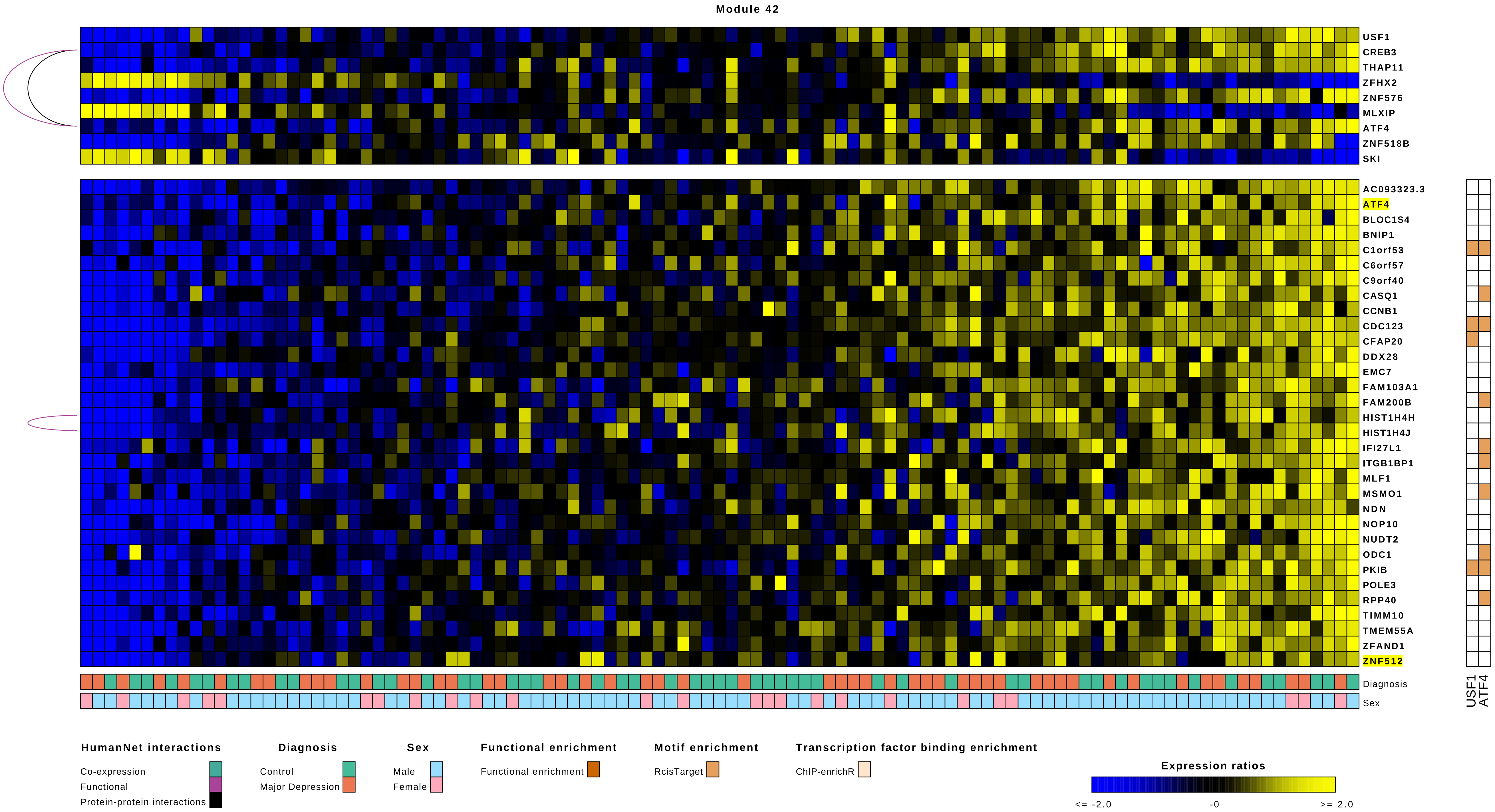
Module 42
USF1
CREB3
THAP11
ZFHX2
ZNF576
MLXIP
ATF4
ZNF518B
SKI
AC093323.3
ATF4
BLOC1S4
BNIP1
C1orf53
C6orf57
C9orf40
CASQ1
CCNB1
CDC123
CFAP20
DDX28
EMC7
FAM103A1
FAM200B
HIST1H4H
HIST1H4J
IFI27L1
ITGB1BP1
MLF1
MSMO1
NDN
NOP10
NUDT2
ODC1
PKIB
POLE3
RPP40
TIMM10
TMEM55A
ZFAND1
ZNF512
Diagnosis
Sex
USF1
ATF4
HumanNet interactions
Co-expression
Functional
Protein-protein interactions
Diagnosis
Control
Major Depression
Sex
Male
Female
Functional enrichment
Functional enrichment
Motif enrichment
RcisTarget
Transcription factor binding enrichment
ChIP-enrichR
Expression ratios
<= -2.0
-0
>= 2.0

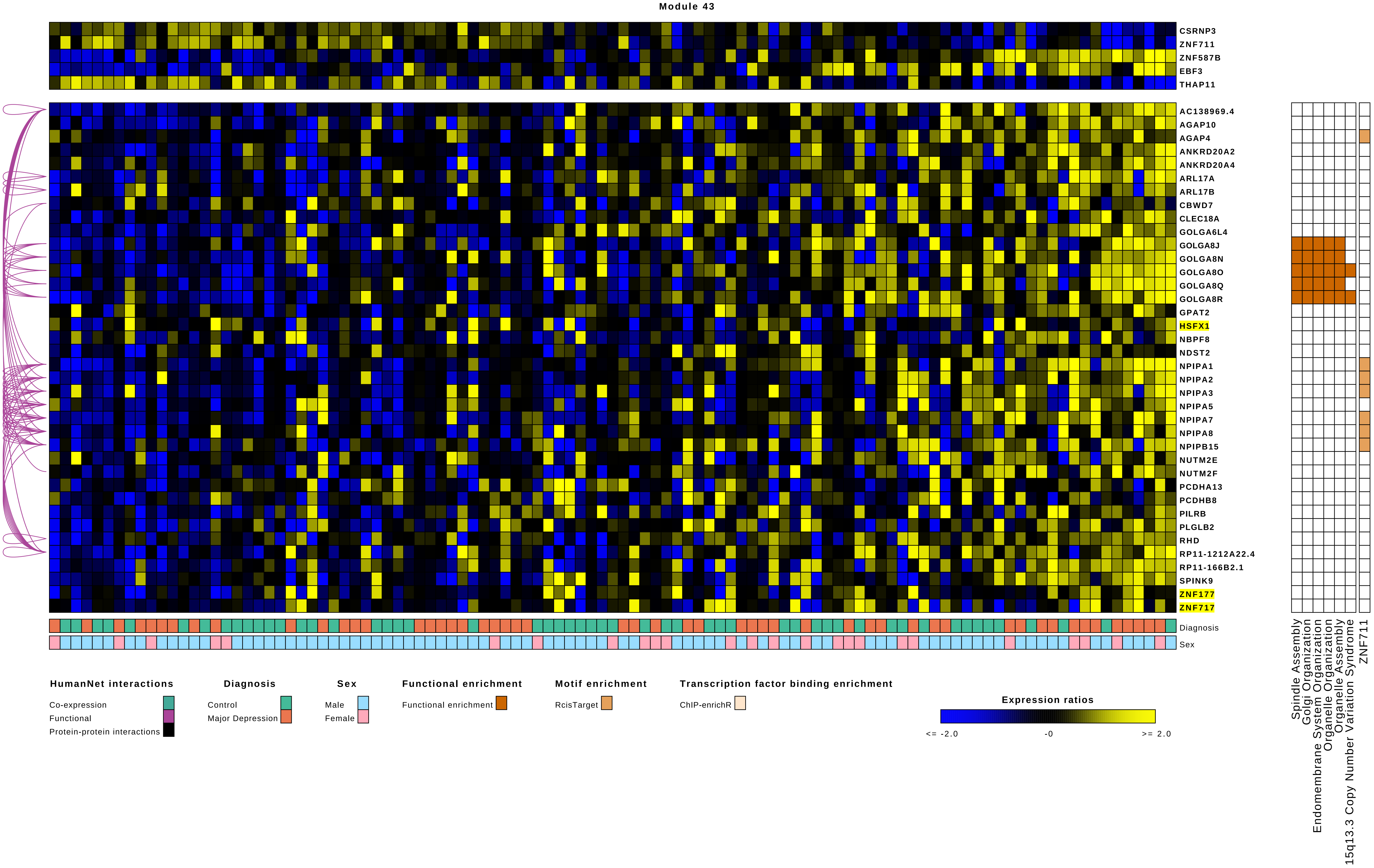

Module 43
CSRNP3
ZNF711
ZNF587B
EBF3
THAP11
AC138969.4
AGAP10
AGAP4
ANKRD20A2
ANKRD20A4
ARL17A
ARL17B
CBWD7
CLEC18A
GOLGA6L4
GOLGA8J
GOLGA8N
GOLGA8O
GOLGA8Q
GOLGA8R
GPAT2
HSFX1
NBPF8
NDST2
NPIPA1
NPIPA2
NPIPA3
NPIPA5
NPIPA7
NPIPA8
NPIPB15
NUTM2E
NUTM2F
PCDHA13
PCDHB8
PILRB
PLGLB2
RHD
RP11-1212A22.4
RP11-166B2.1
SPINK9
ZNF177
ZNF717
Diagnosis
Sex
Spindle Assembly
Golgi Organization
Endomembrane System Organization
Organelle Organization
Organelle Assembly
15q13.3 Copy Number Variation Syndrome
ZNF711
HumanNet interactions
Co-expression
Functional
Protein-protein interactions
Diagnosis
Control
Major Depression
Sex
Male
Female
Functional enrichment
Functional enrichment
Motif enrichment
RcisTarget
Transcription factor binding enrichment
ChIP-enrichR
Expression ratios
<= -2.0
-0
>= 2.0

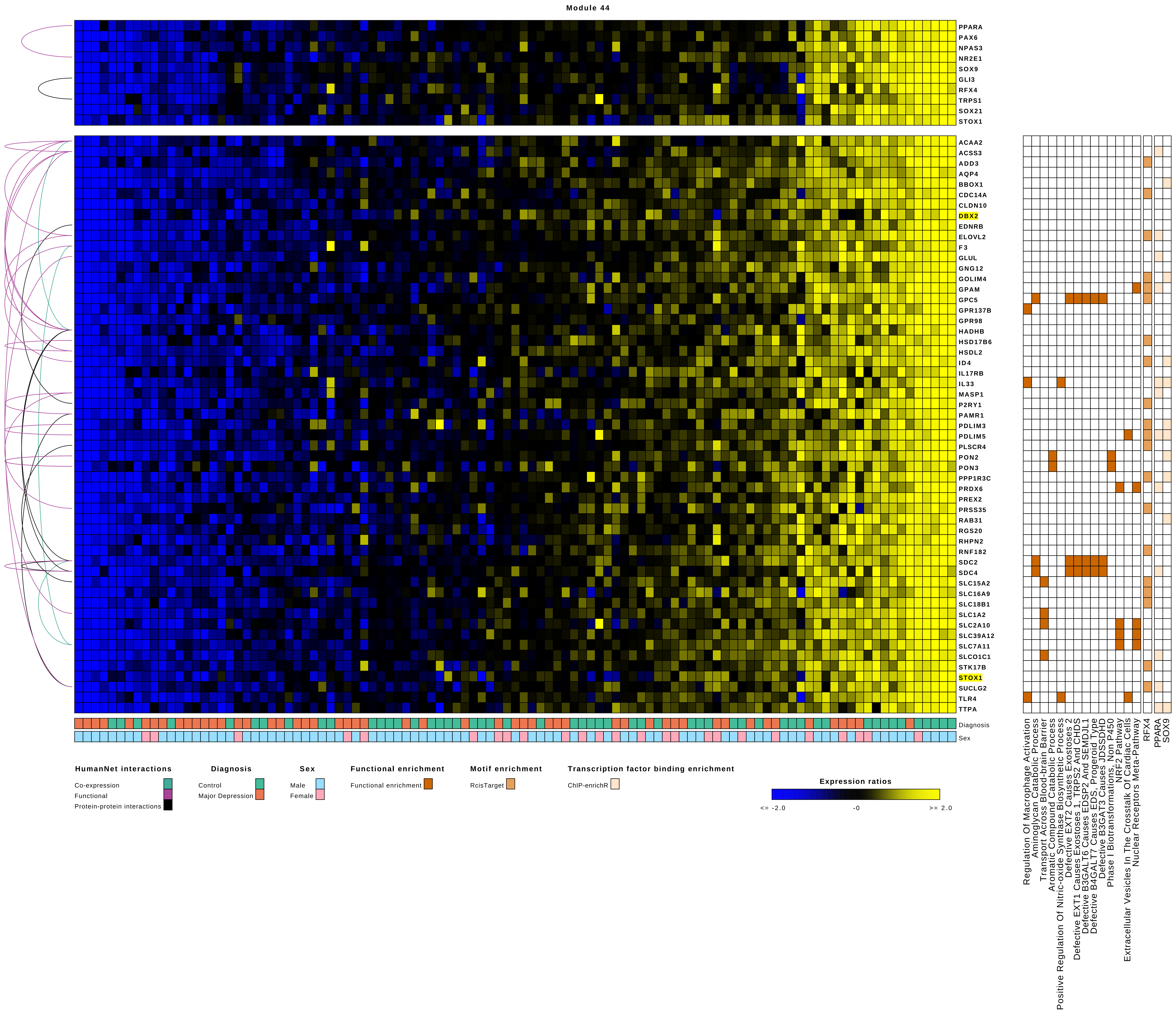
Module 44
PPARA
PAX6
NPAS3
NR2E1
SOX9
GLI3
RFX4
TRPS1
SOX21
STOX1
ACAA2
ACSS3
ADD3
AQP4
BBOX1
CDC14A
CLDN10
DBX2
EDNRB
ELOVL2
F3
GLUL
GNG12
GOLIM4
GPAM
GPC5
GPR137B
GPR98
HADHB
HSD17B6
HSDL2
ID4
IL17RB
IL33
MASP1
P2RY1
PAMR1
PDLIM3
PDLIM5
PLSCR4
PON2
PON3
PPP1R3C
PRDX6
PREX2
PRSS35
RAB31
RGS20
RHPN2
RNF182
SDC2
SDC4
SLC15A2
SLC16A9
SLC18B1
SLC1A2
SLC2A10
SLC39A12
SLC7A11
SLCO1C1
STK17B
STOX1
SUCLG2
TLR4
TTPA
Diagnosis
Sex
Regulation Of Macrophage Activation
Aminoglycan Catabolic Process
Transport Across Blood-brain Barrier
Aromatic Compound Catabolic Process
Positive Regulation Of Nitric-oxide Synthase Biosynthetic Process
Defective EXT2 Causes Exostoses 2
Defective EXT1 Causes Exostoses 1, TRPS2 And CHDS
Defective B3GALT6 Causes EDSP2 And SEMDJL1
Defective B4GALT7 Causes EDS, Progeroid Type
Defective B3GAT3 Causes JDSSDHD
Phase I Biotransformations, Non P450
NRF2 Pathway
Extracellular Vesicles In The Crosstalk Of Cardiac Cells
Nuclear Receptors Meta-Pathway
RFX4
PPARA
SOX9
HumanNet interactions
Co-expression
Functional
Protein-protein interactions
Diagnosis
Control
Major Depression
Sex
Male
Female
Functional enrichment
Functional enrichment
Motif enrichment
RcisTarget
Transcription factor binding enrichment
ChIP-enrichR
Expression ratios
<= -2.0
-0
>= 2.0

### Module 45

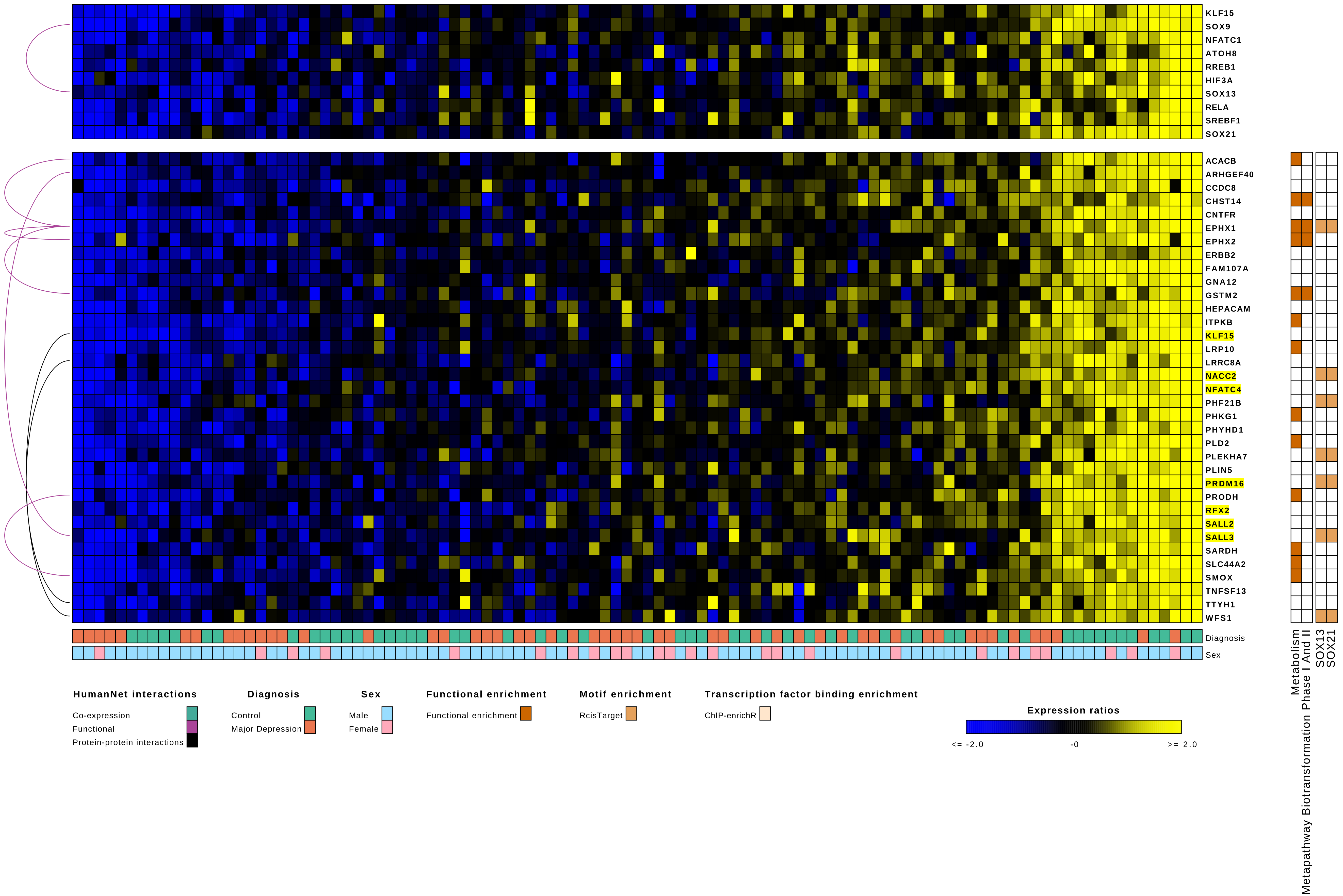
KLF15
SOX9
NFATC1
ATOH8
RREB1
HIF3A
SOX13
RELA
SREBF1
SOX21
ACACB
ARHGEF40
CCDC8
CHST14
CNTFR
EPHX1
EPHX2
ERBB2
FAM107A
GNA12
GSTM2
HEPACAM
ITPKB
KLF15
LRP10
LRRC8A
NACC2
NFATC4
PHF21B
PHKG1
PHYHD1
PLD2
PLEKHA7
PLIN5
PRDM16
PRODH
RFX2
SALL2
SALL3
SARDH
SLC44A2
SMOX
TNFSF13
TTYH1
WFS1
Diagnosis
Sex
Metapathway Biotransformation Phase I And II
Metabolism
SOX13
SOX21
HumanNet interactions
Co-expression
Functional
Protein-protein interactions
Diagnosis
Control
Major Depression
Sex
Male
Female
Functional enrichment
Functional enrichment
Motif enrichment
RcisTarget
Transcription factor binding enrichment
ChIP-enrichR
Expression ratios
<= -2.0
-0
>= 2.0

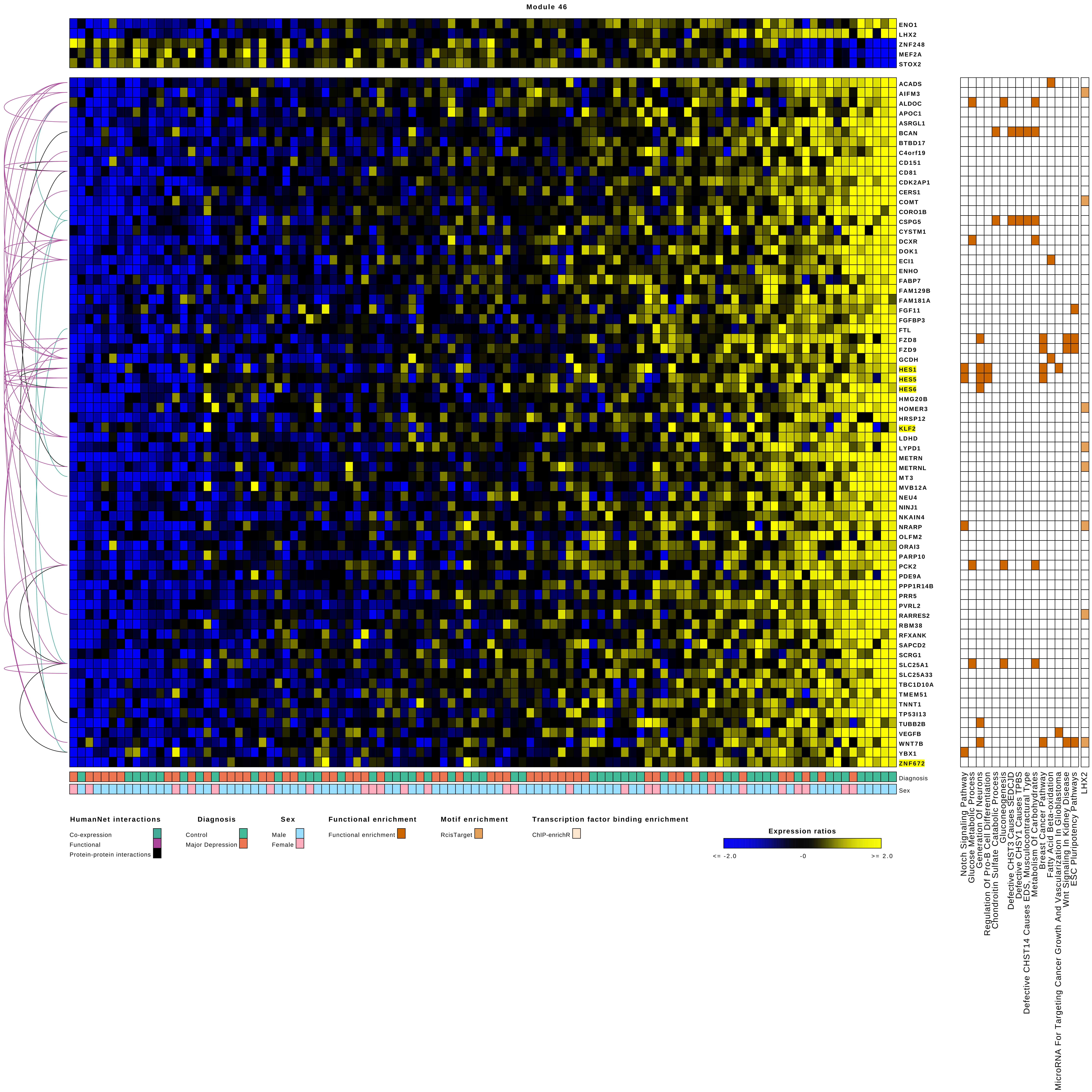
Module 46
ENO1
LHX2
ZNF248
MEF2A
STOX2
ACADS
AIFM3
ALDOC
APOC1
ASRGL1
BCAN
BTBD17
C4orf19
CD151
CD81
CDK2AP1
CERS1
COMT
CORO1B
CSPG5
CYSTM1
DCXR
DOK1
ECI1
ENHO
FABP7
FAM129B
FAM181A
FGF11
FGFBP3
FTL
FZD8
FZD9
GCDH
HES1
HES5
HES6
HMG20B
HOMER3
HRSP12
KLF2
LDHD
LYPD1
METRN
METRNL
MT3
MVB12A
NEU4
NINJ1
NKAIN4
NRARP
OLFM2
ORAI3
PARP10
PCK2
PDE9A
PPP1R14B
PRR5
PVRL2
RARRES2
RBM38
RFXANK
SAPCD2
SCRG1
SLC25A1
SLC25A33
TBC1D10A
TMEM51
TNNT1
TP53I13
TUBB2B
VEGFB
WNT7B
YBX1
ZNF672
Diagnosis
Sex
Notch Signaling Pathway
Glucose Metabolic Process
Generation Of Neurons
Regulation Of Pro-B Cell Differentiation
Chondroitin Sulfate Catabolic Process
Gluconeogenesis
Defective CHST3 Causes SEDCJD
Defective CHSY1 Causes TPBS
Defective CHST14 Causes EDS, Musculocontractural Type
Metabolism Of Carbohydrates
Breast Cancer Pathway
Fatty Acid Beta-oxidation
MicroRNA For Targeting Cancer Growth And Vascularization In Glioblastoma
Wnt Signaling In Kidney Disease
ESC Pluripotency Pathways
LHX2
HumanNet interactions
Co-expression
Functional
Protein-protein interactions
Diagnosis
Control
Major Depression
Sex
Male
Female
Functional enrichment
Functional enrichment
Motif enrichment
RcisTarget
Transcription factor binding enrichment
ChIP-enrichR
Expression ratios
<= -2.0
-0
>= 2.0

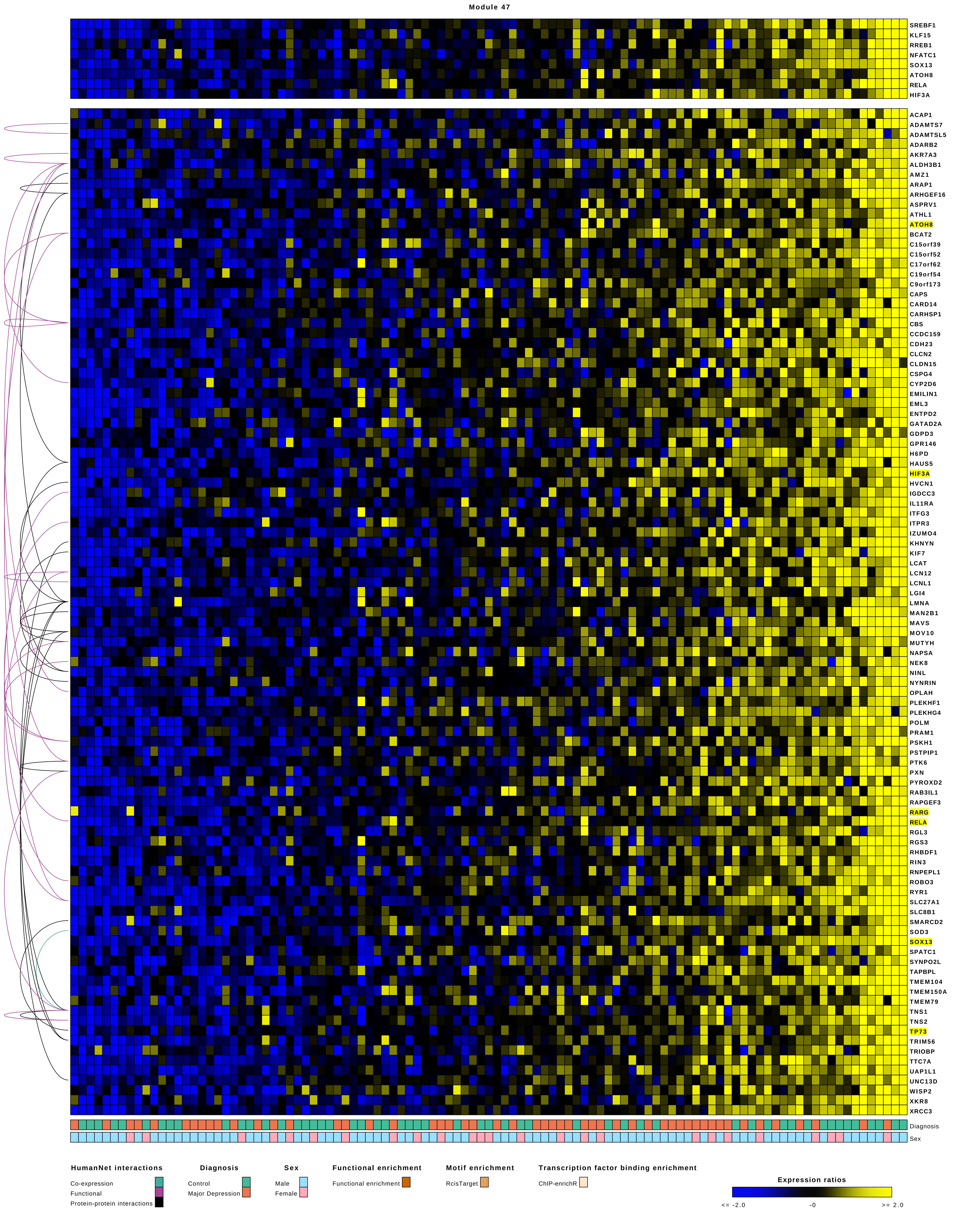

Module 47
SREBF1
KLF15
RREB1
NFATC1
SOX13
ATOH8
RELA
HIF3A
ACAP1
ADAMTS7
ADAMTSL5
ADARB2
AKR7A3
ALDH3B1
AMZ1
ARAP1
ARHGEF16
ASPRV1
ATHL1
ATOH8
BCAT2
C15orf39
C15orf52
C17orf62
C19orf54
C9orf173
CAPS
CARD14
CARHSP1
CBS
CCDC159
CDH23
CLCN2
CLDN15
CSPG4
CYP2D6
EMILIN1
EML3
ENTPD2
GATAD2A
GDPD3
GPR146
H6PD
HAUS5
HIF3A
HVCN1
IGDCC3
IL11RA
ITFG3
ITPR3
IZUMO4
KHNYN
KIF7
LCAT
LCN12
LCNL1
LGI4
LMNA
MAN2B1
MAVS
MOV10
MUTYH
NAPSA
NEK8
NINL
NYNRIN
OPLAH
PLEKHF1
PLEKHG4
POLM
PRAM1
PSKH1
PSTPIP1
PTK6
PXN
PYROXD2
RAB3IL1
RAPGEF3
RARG
RELA
RGL3
RGS3
RHBDF1
RIN3
RNPEPL1
ROBO3
RYR1
SLC27A1
SLC8B1
SMARCD2
SOD3
SOX13
SPATC1
SYNPO2L
TAPBPL
TMEM104
TMEM150A
TMEM79
TNS1
TNS2
TP73
TRIM56
TRIOBP
TTC7A
UAP1L1
UNC13D
WISP2
XKR8
XRCC3
Diagnosis
Sex
HumanNet interactions
Co-expression
Functional
Protein-protein interactions
Diagnosis
Control
Major Depression
Sex
Male
Female
Functional enrichment
Functional enrichment
Motif enrichment
RcisTarget
Transcription factor binding enrichment
ChIP-enrichR
Expression ratios
<= -2.0
-0
>= 2.0

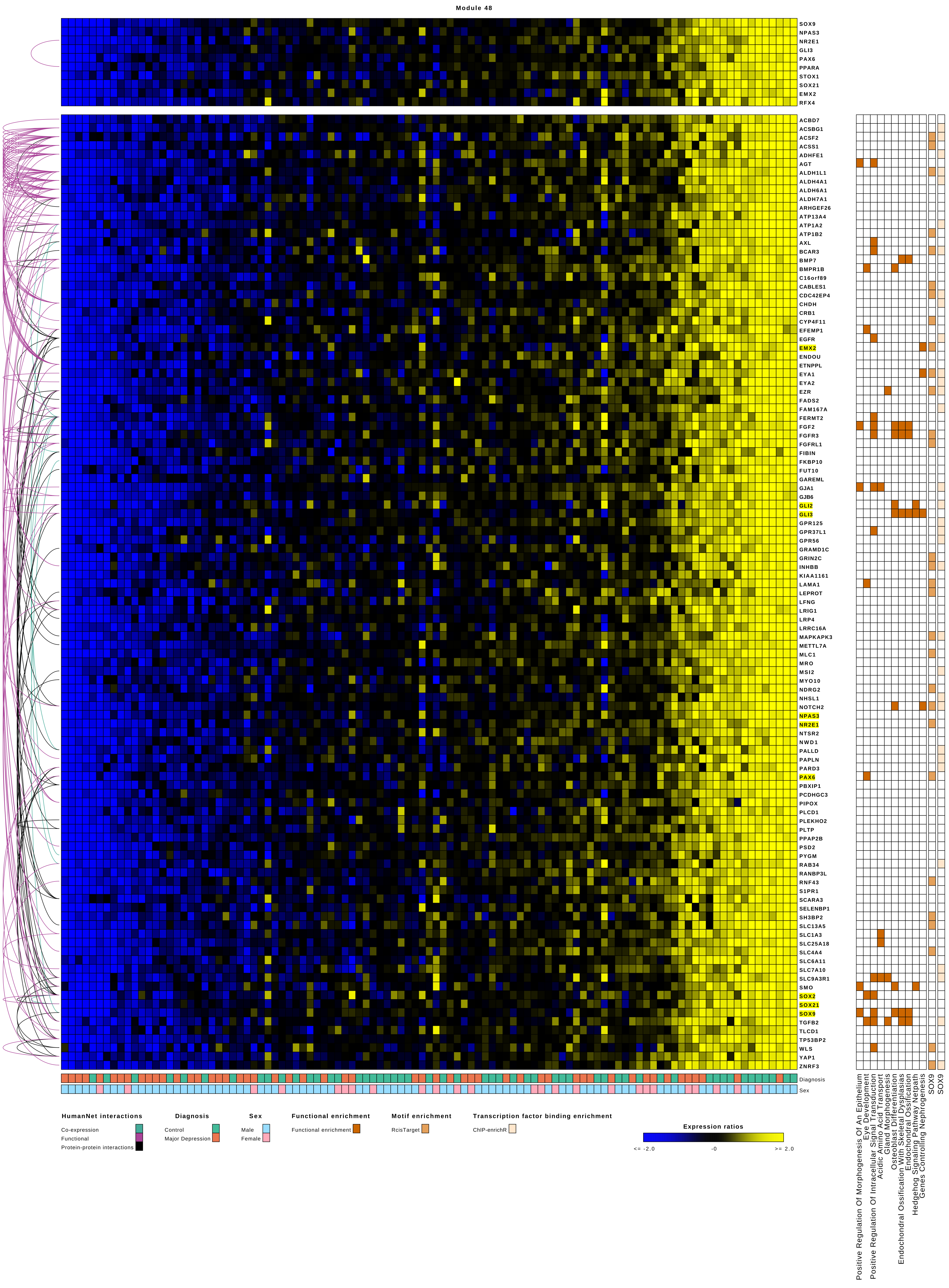

Module 48
SOX9
NPAS3
NR2E1
GLI3
PAX6
PPARA
STOX1
SOX21
EMX2
RFX4
ACBD7
ACSBG1
ACSF2
ACSS1
ADHFE1
AGT
ALDH1L1
ALDH4A1
ALDH6A1
ALDH7A1
ARHGEF26
ATP13A4
ATP1A2
ATP1B2
AXL
BCAR3
BMP7
BMPR1B
C16orf89
CABLES1
CDC42EP4
CHDH
CRB1
CYP4F11
EFEMP1
EGFR
EMX2
ENDOU
ETNPPL
EYA1
EYA2
EZR
FADS2
FAM167A
FERMT2
FGF2
FGFR3
FGFRL1
FIBIN
FKBP10
FUT10
GAREML
GJA1
GJB6
GLI2
GLI3
GPR125
GPR37L1
GPR56
GRAMD1C
GRIN2C
INHBB
KIAA1161
LAMA1
LEPROT
LFNG
LRIG1
LRP4
LRRC16A
MAPKAPK3
METTL7A
MLC1
MRO
MSI2
MYO10
NDRG2
NHSL1
NOTCH2
NPAS3
NR2E1
NTSR2
NWD1
PALLD
PAPLN
PARD3
PAX6
PBXIP1
PCDHGC3
PIPOX
PLCD1
PLEKHO2
PLTP
PPAP2B
PSD2
PYGM
RAB34
RANBP3L
RNF43
S1PR1
SCARA3
SELENBP1
SH3BP2
SLC13A5
SLC1A3
SLC25A18
SLC4A4
SLC6A11
SLC7A10
SLC9A3R1
SMO
SOX2
SOX21
SOX9
TGFB2
TLCD1
TP53BP2
WLS
YAP1
ZNRF3
Diagnosis
Sex
Positive Regulation Of Morphogenesis Of An Epithelium
Eye Development
Positive Regulation Of Intracellular Signal Transduction
Acidic Amino Acid Transport
Gland Morphogenesis
Osteoblast Differentiation
Endochondral Ossification With Skeletal Dysplasias
Endochondral Ossification
Hedgehog Signaling Pathway Netpath
Genes Controlling Nephrogenesis
SOX9
SOX9
HumanNet interactions
Co-expression
Functional
Protein-protein interactions
Diagnosis
Control
Major Depression
Sex
Male
Female
Functional enrichment
Functional enrichment
Motif enrichment
RcisTarget
Transcription factor binding enrichment
ChIP-enrichR
Expression ratios
<= -2.0
-0
>= 2.0

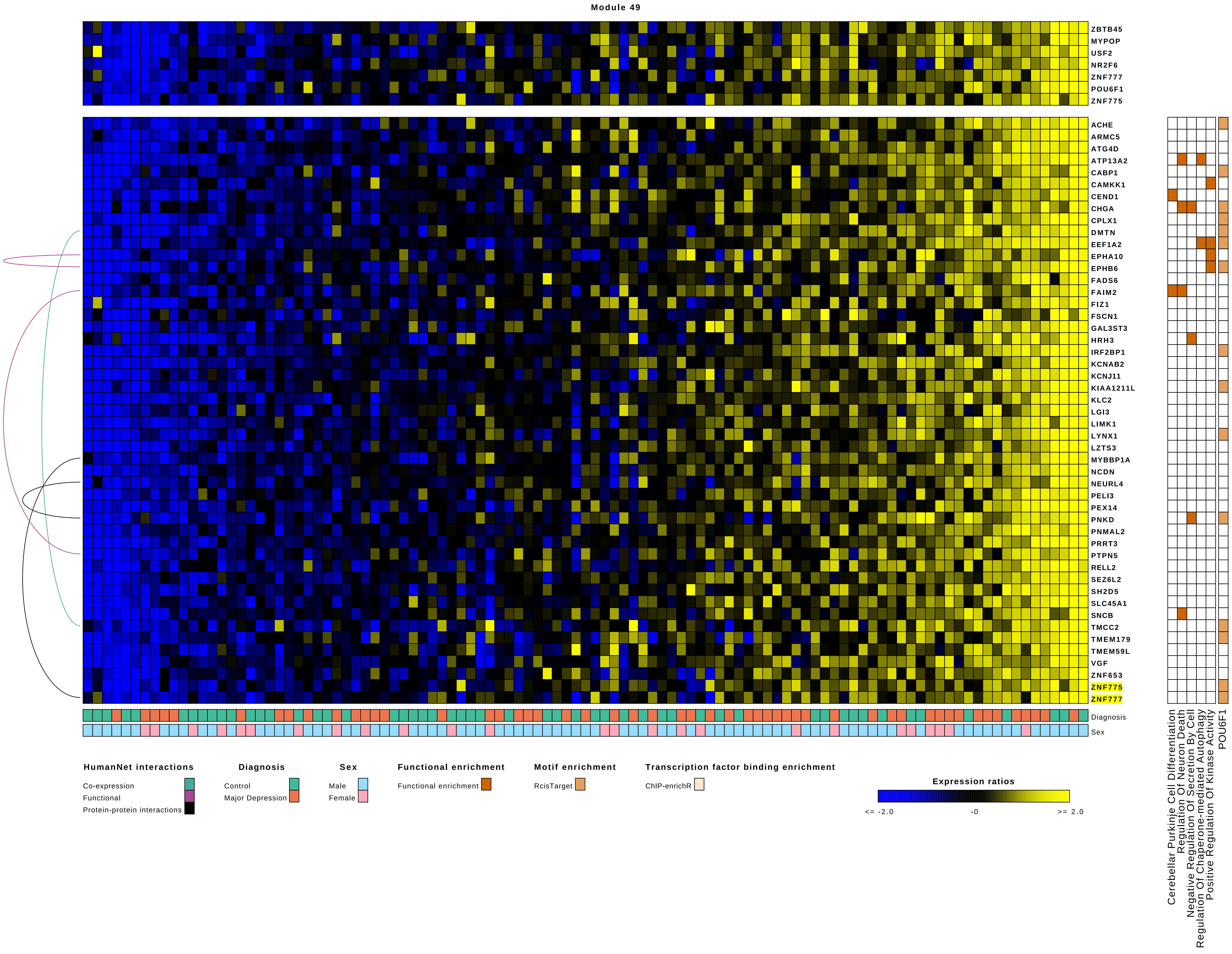
Module 49
ZBTB45
MYPOP
USF2
NR2F6
ZNF777
POU6F1
ZNF775
ACHE
ARMC5
ATG4D
ATP13A2
CABP1
CAMKK1
CEND1
CHGA
CPLX1
DMTN
EEF1A2
EPHA10
EPHB6
FADS6
FAIM2
FIZ1
FSCN1
GAL3ST3
HRH3
IRF2BP1
KCNAB2
KCNJ11
KIAA1211L
KLC2
LGI3
LIMK1
LYNX1
LZTS3
MYBBP1A
NCDN
NEURL4
PELI3
PEX14
PNKD
PNMAL2
PRRT3
PTPN5
RELL2
SEZ6L2
SH2D5
SLC45A1
SNCB
TMCC2
TMEM179
TMEM59L
VGF
ZNF653
ZNF775
ZNF777
Diagnosis
Sex
Cerebellar Purkinje Cell Differentiation
Regulation Of Neuron Death
Negative Regulation Of Secretion By Cell
Regulation Of Chaperone-mediated Autophagy
Positive Regulation Of Kinase Activity
POU6F1
HumanNet interactions
Co-expression
Functional
Protein-protein interactions
Diagnosis
Control
Major Depression
Sex
Male
Female
Functional enrichment
Functional enrichment
Motif enrichment
RcisTarget
Transcription factor binding enrichment
ChIP-enrichR
Expression ratios
<= -2.0
-0
>= 2.0

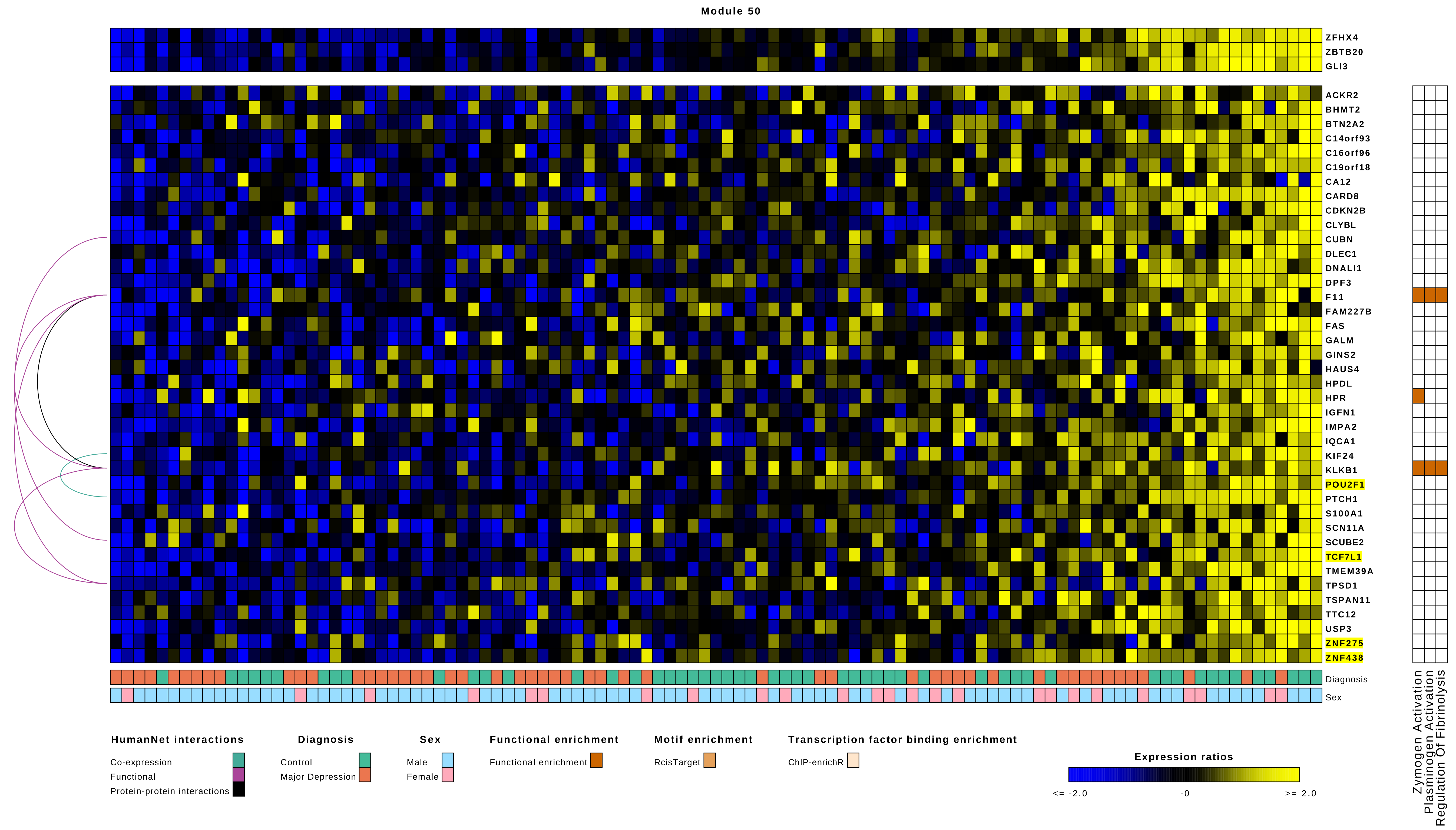
Module 50
ZFHX4
ZBTB20
GLI3
ACKR2
BHMT2
BTN2A2
C14orf93
C16orf96
C19orf18
CA12
CARD8
CDKN2B
CLYBL
CUBN
DLEC1
DNALI1
DPF3
F11
FAM227B
FAS
GALM
GINS2
HAUS4
HPDL
HPR
IGFN1
IMPA2
IQCA1
KIF24
KLKB1
POU2F1
PTCH1
S100A1
SCN11A
SCUBE2
TCF7L1
TMEM39A
TPSD1
TSPAN11
TTC12
USP3
ZNF275
ZNF438
Diagnosis
Sex
Zymogen Activation
Plasminogen Activation
Regulation Of Fibrinolysis
HumanNet interactions
Co-expression
Functional
Protein-protein interactions
Diagnosis
Control
Major Depression
Sex
Male
Female
Functional enrichment
Functional enrichment
Motif enrichment
RcisTarget
Transcription factor binding enrichment
ChIP-enrichR
Expression ratios
<= -2.0
-0
>= 2.0

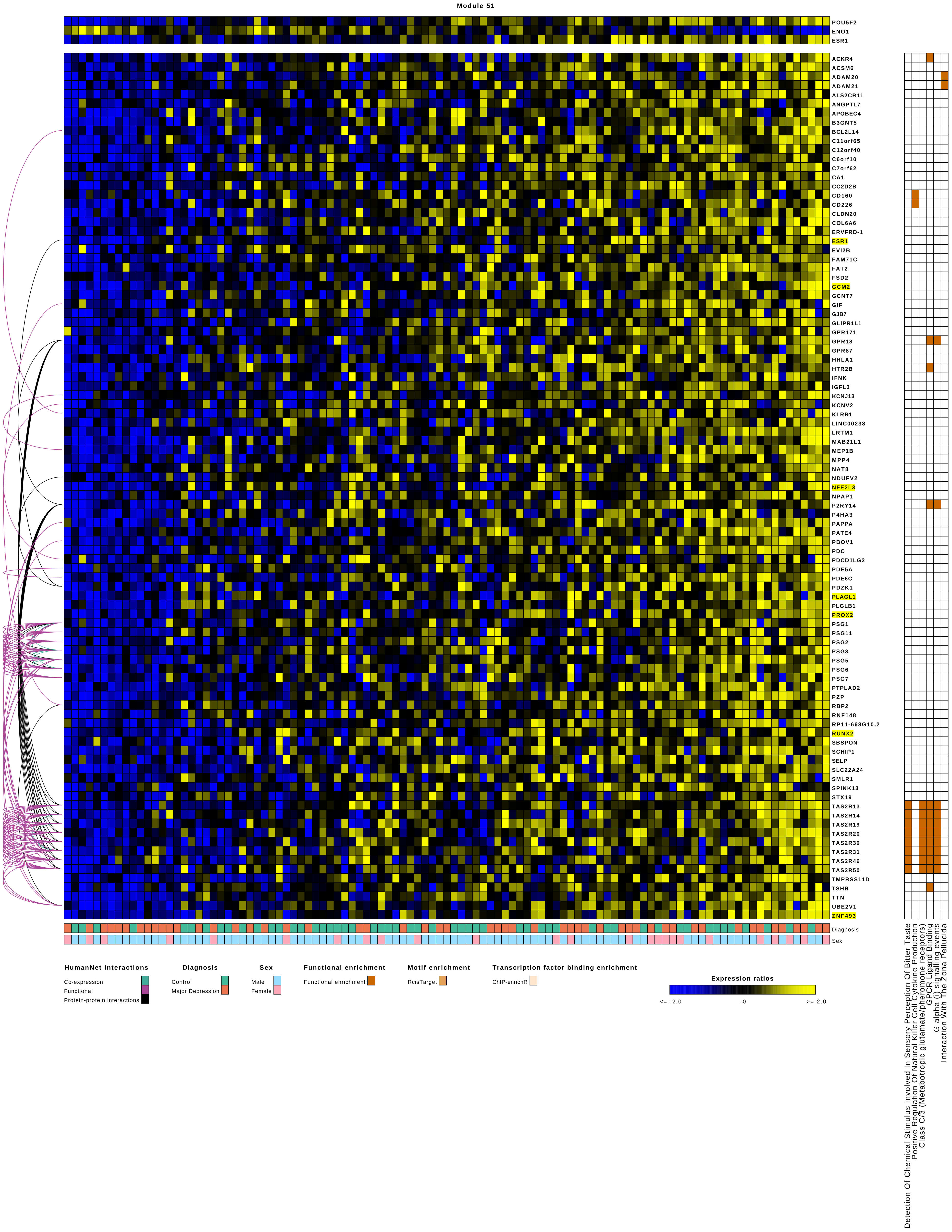
Module 51
POU5F2
ENO1
ESR1
ACKR4
ACSM6
ADAM20
ADAM21
ALS2CR11
ANGPTL7
APOBEC4
B3GNT5
BCL2L14
C11orf65
C12orf40
C6orf10
C7orf62
CA1
CC2D2B
CD160
CD226
CLDN20
COL6A6
ERVFRD-1
ESR1
EVI2B
FAM71C
FAT2
FSD2
GCM2
GCNT7
GIF
GJB7
GLIPR1L1
GPR171
GPR18
GPR87
HHLA1
HTR2B
IFNK
IGFL3
KCNJ13
KCNV2
KLRB1
LINC00238
LRTM1
MAB21L1
MEP1B
MPP4
NAT8
NDUFV2
NFE2L3
NPAP1
P2RY14
P4HA3
PAPPA
PATE4
PBOV1
PDC
PDCD1LG2
PDE5A
PDE6C
PDZK1
PLAGL1
PLGLB1
PROX2
PSG1
PSG11
PSG2
PSG3
PSG5
PSG6
PSG7
PTPLAD2
PZP
RBP2
RNF148
RP11-668G10.2
RUNX2
SBSPON
SCHIP1
SELP
SLC22A24
SMLR1
SPINK13
STX19
TAS2R13
TAS2R14
TAS2R19
TAS2R20
TAS2R30
TAS2R31
TAS2R46
TAS2R50
TMPRSS11D
TSHR
TTN
UBE2V1
ZNF493
Diagnosis
Sex
Detection Of Chemical Stimulus Involved In Sensory Perception Of Bitter Taste
Positive Regulation Of Natural Killer Cell Cytokine Production
Class C/3 (Metabotropic glutamate/pheromone receptors)
GPCR Ligand Binding
G alpha (i) signalling events
Interaction With The Zona Pellucida
HumanNet interactions
Co-expression
Functional
Protein-protein interactions
Diagnosis
Control
Major Depression
Sex
Male
Female
Functional enrichment
Functional enrichment
Motif enrichment
RcisTarget
Transcription factor binding enrichment
ChIP-enrichR
Expression ratios
<= -2.0
-0
>= 2.0

Module 52

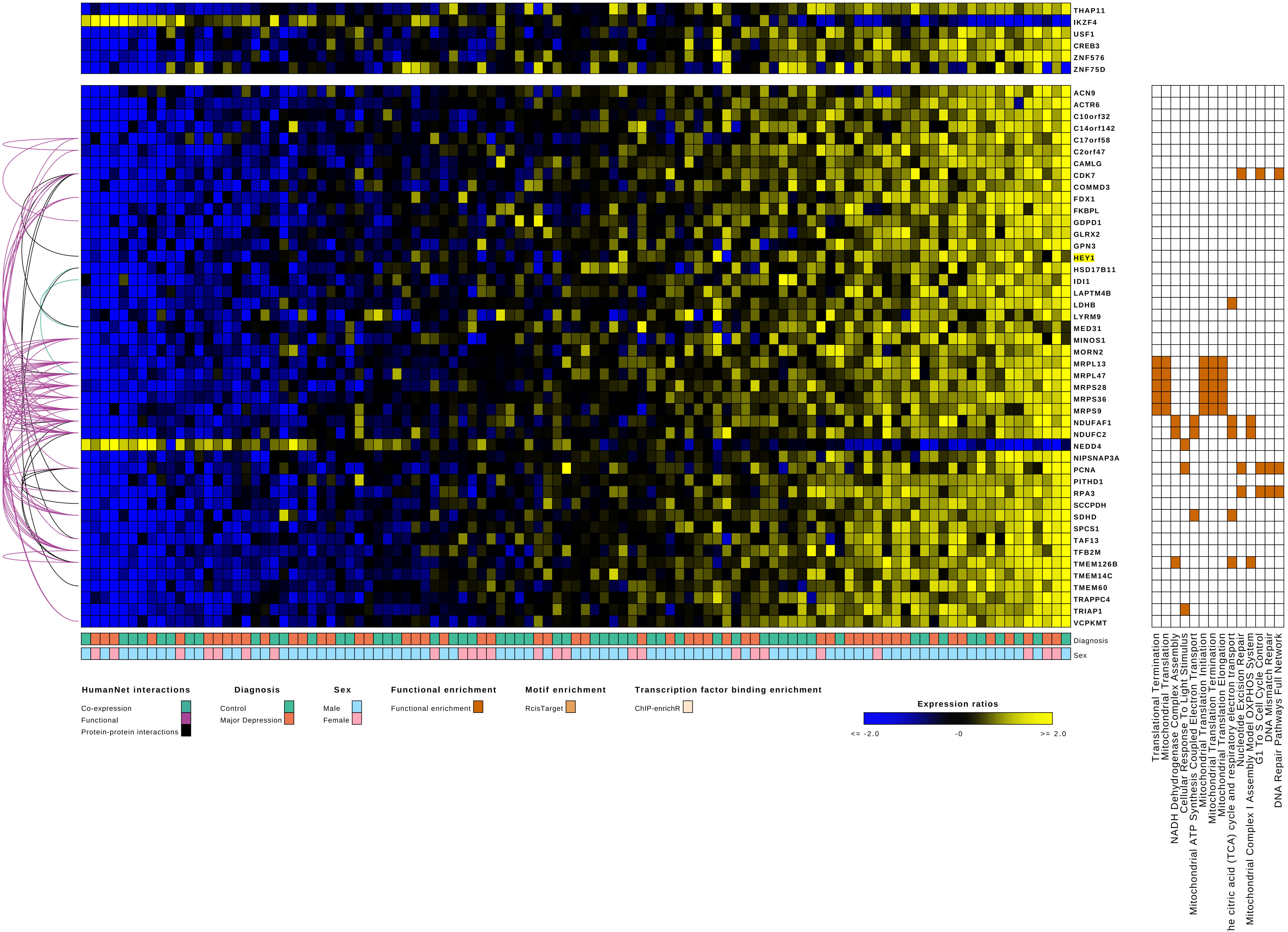
THAP11
IKZF4
USF1
CREB3
ZNF576
ZNF75D
ACN9
ACTR6
C10orf32
C14orf142
C17orf58
C2orf47
CAMLG
CDK7
COMMD3
FDX1
FKBPL
GDPD1
GLRX2
GPN3
HEY1
HSD17B11
IDI1
LAPTM4B
LDHB
LYRM9
MED31
MINOS1
MORN2
MRPL13
MRPL47
MRPS28
MRPS36
MRPS9
NDUFAF1
NDUFC2
NEDD4
NIPSNAP3A
PCNA
PITHD1
RPA3
SCCPDH
SDHD
SPCS1
TAF13
TFB2M
TMEM126B
TMEM14C
TMEM60
TRAPPC4
TRIAP1
VCPKMT
Diagnosis
Sex
Translational Termination
Mitochondrial Translation
NADH Dehydrogenase Complex Assembly
Cellular Response To Light Stimulus
Mitochondrial ATP Synthesis Coupled Electron Transport
Mitochondrial Translation Initiation
Mitochondrial Translation Termination
Mitochondrial Translation Elongation
The citric acid (TCA) cycle and respiratory electron transport
Nucleotide Excision Repair
Mitochondrial Complex I Assembly Model OXPHOS System
G1 To S Cell Cycle Control
DNA Mismatch Repair
DNA Repair Pathways Full Network
HumanNet interactions
Co-expression
Functional
Protein-protein interactions
Diagnosis
Control
Major Depression
Sex
Male
Female
Functional enrichment
Functional enrichment
Motif enrichment
RcisTarget
Transcription factor binding enrichment
ChIP-enrichR
Expression ratios
<= -2.0
-0
>= 2.0

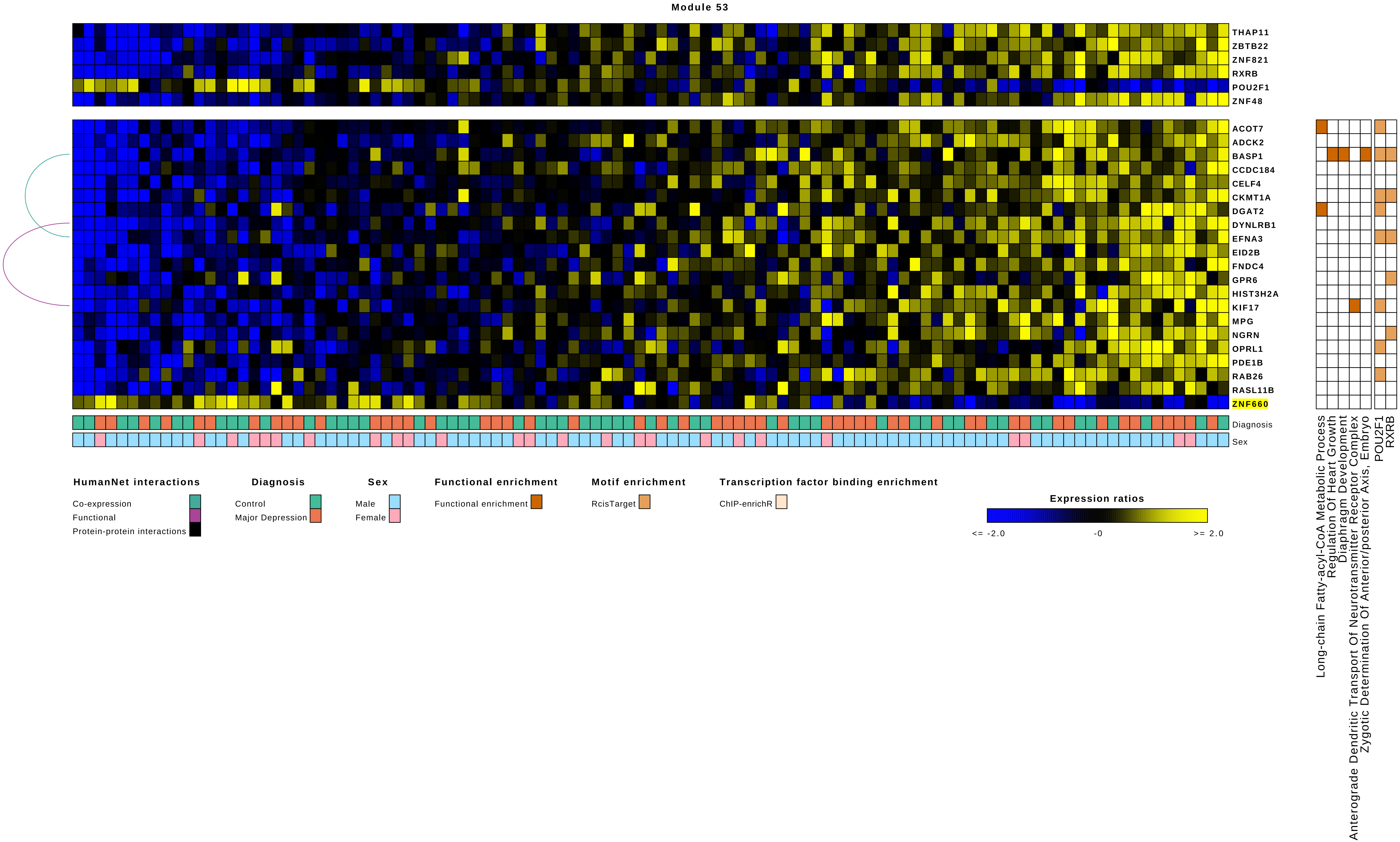

Module 53
THAP11
ZBTB22
ZNF821
RXRB
POU2F1
ZNF48
ACOT7
ADCK2
BASP1
CCDC184
CELF4
CKMT1A
DGAT2
DYNLRB1
EFNA3
EID2B
FNDC4
GPR6
HIST3H2A
KIF17
MPG
NGRN
OPRL1
PDE1B
RAB26
RASL11B
ZNF660
Diagnosis
Sex
Long-chain Fatty-acyl-CoA Metabolic Process
Regulation Of Heart Growth
Diaphragm Development
Anterograde Dendritic Transport Of Neurotransmitter Receptor Complex
Zygotic Determination Of Anterior/posterior Axis, Embryo
POU2F1
RXRB
HumanNet interactions
Co-expression
Functional
Protein-protein interactions
Diagnosis
Control
Major Depression
Sex
Male
Female
Functional enrichment
Functional enrichment
Motif enrichment
RcisTarget
Transcription factor binding enrichment
ChIP-enrichR
Expression ratios
<= -2.0
-0
>= 2.0

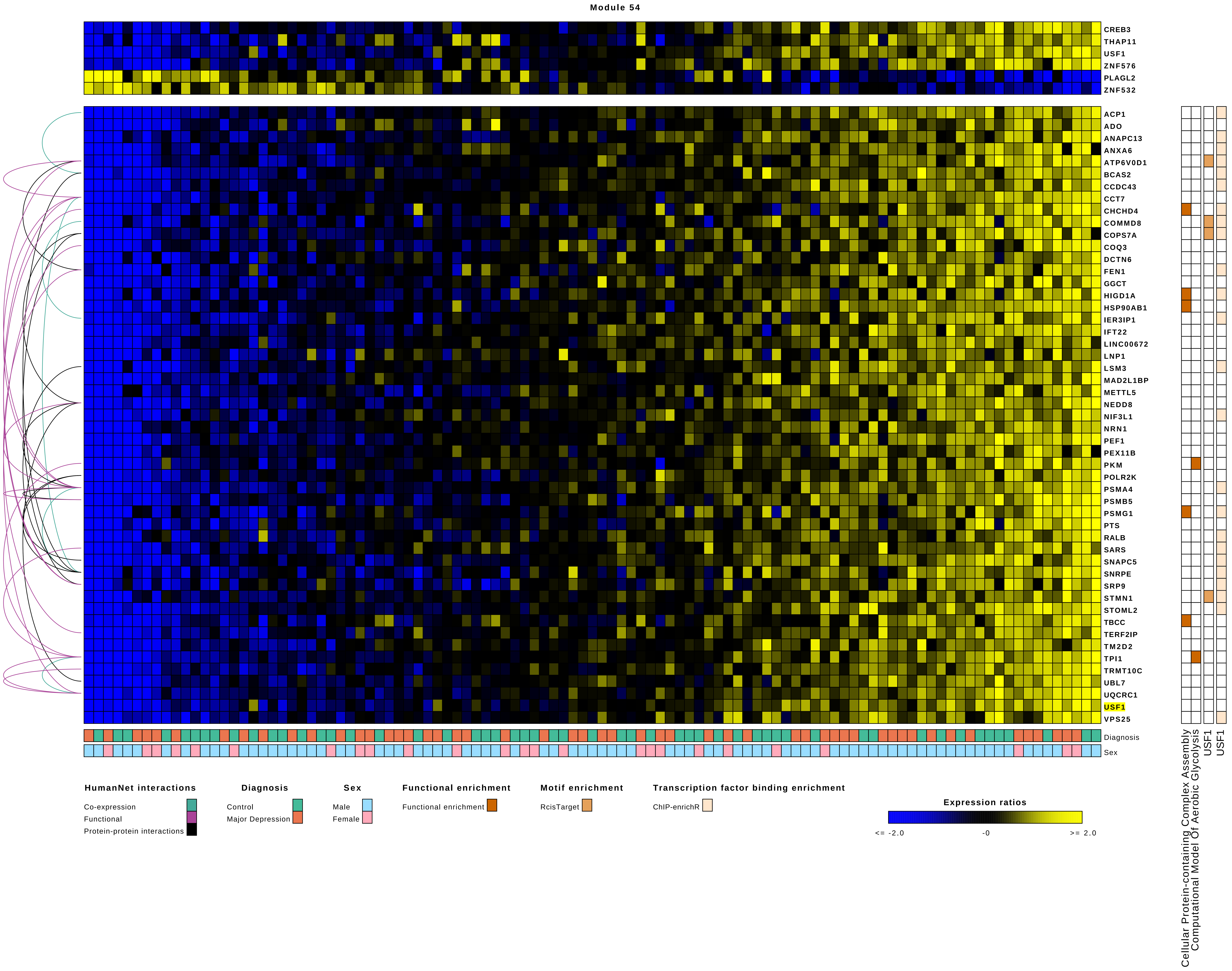

Module 54
CREB3
THAP11
USF1
ZNF576
PLAGL2
ZNF532
ACP1
ADO
ANAPC13
ANXA6
ATP6V0D1
BCAS2
CCDC43
CCT7
CHCHD4
COMMD8
COPS7A
COQ3
DCTN6
FEN1
GGCT
HIGD1A
HSP90AB1
IER3IP1
IFT22
LINC00672
LNP1
LSM3
MAD2L1BP
METTL5
NEDD8
NIF3L1
NRN1
PEF1
PEX11B
PKM
POLR2K
PSMA4
PSMB5
PSMG1
PTS
RALB
SARS
SNAPC5
SNRPE
SRP9
STMN1
STOML2
TBCC
TERF2IP
TM2D2
TPI1
TRMT10C
UBL7
UQCRC1
USF1
VPS25
Diagnosis
Sex
Cellular Protein-containing Complex Assembly
Computational Model Of Aerobic Glycolysis
USF1
USF1
HumanNet interactions
Co-expression
Functional
Protein-protein interactions
Diagnosis
Control
Major Depression
Sex
Male
Female
Functional enrichment
Functional enrichment
Motif enrichment
RcisTarget
Transcription factor binding enrichment
ChIP-enrichR
Expression ratios
<= -2.0
-0
>= 2.0

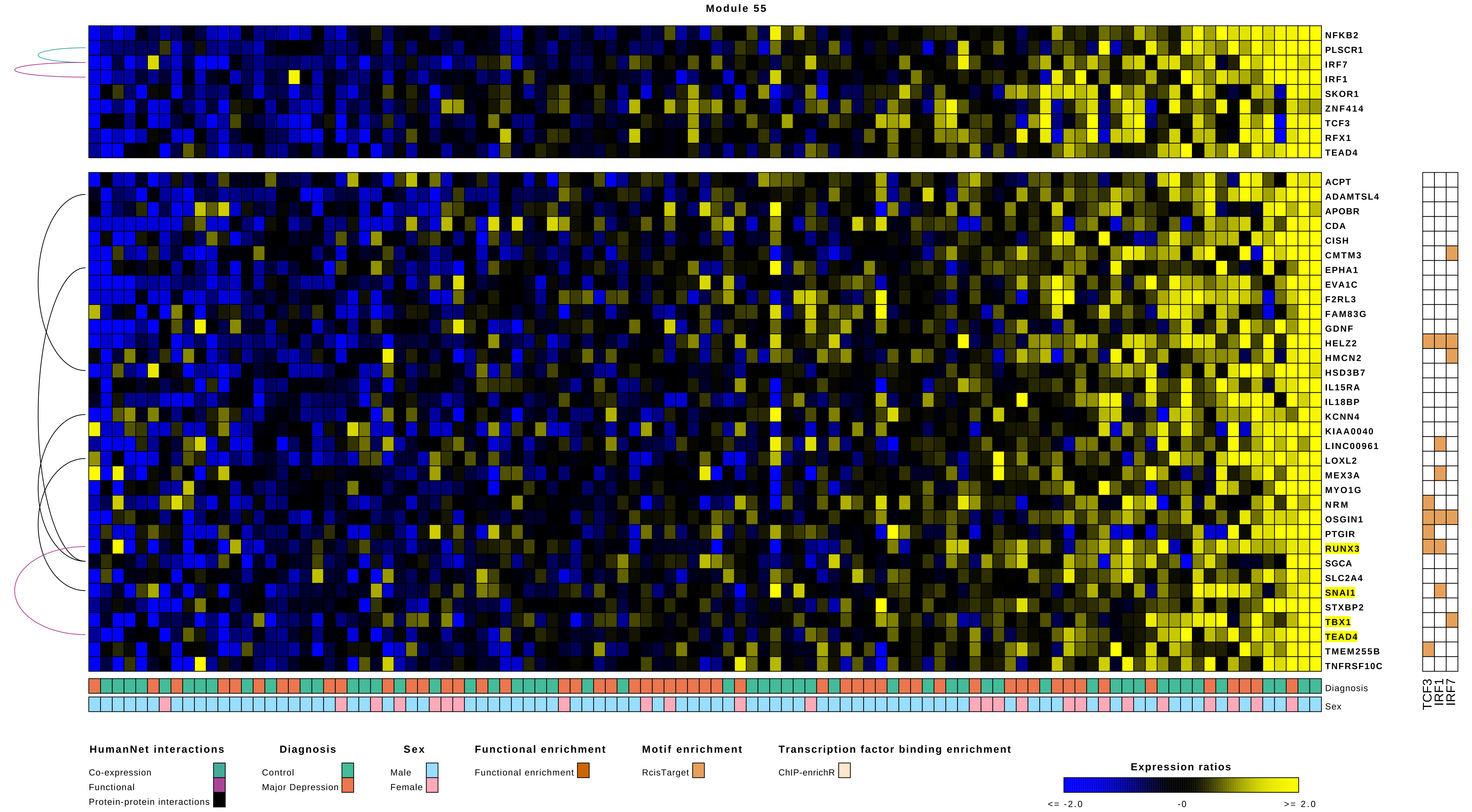
Module 55
NFKB2
PLSCR1
IRF7
IRF1
SKOR1
ZNF414
TCF3
RFX1
TEAD4
ACPT
ADAMTSL4
APOBR
CDA
CISH
CMTM3
EPHA1
EVA1C
F2RL3
FAM83G
GDNF
HELZ2
HMCN2
HSD3B7
IL15RA
IL18BP
KCNN4
KIAA0040
LINC00961
LOXL2
MEX3A
MYO1G
NRM
OSGIN1
PTGIR
RUNX3
SGCA
SLC2A4
SNAI1
STXBP2
TBX1
TEAD4
TMEM255B
TNFRSF10C
Diagnosis
Sex
TCF3
IRF1
IRF7
HumanNet interactions
Co-expression
Functional
Protein-protein interactions
Diagnosis
Control
Major Depression
Sex
Male
Female
Functional enrichment
Functional enrichment
Motif enrichment
RcisTarget
Transcription factor binding enrichment
ChIP-enrichR
Expression ratios
<= -2.0
-0
>= 2.0

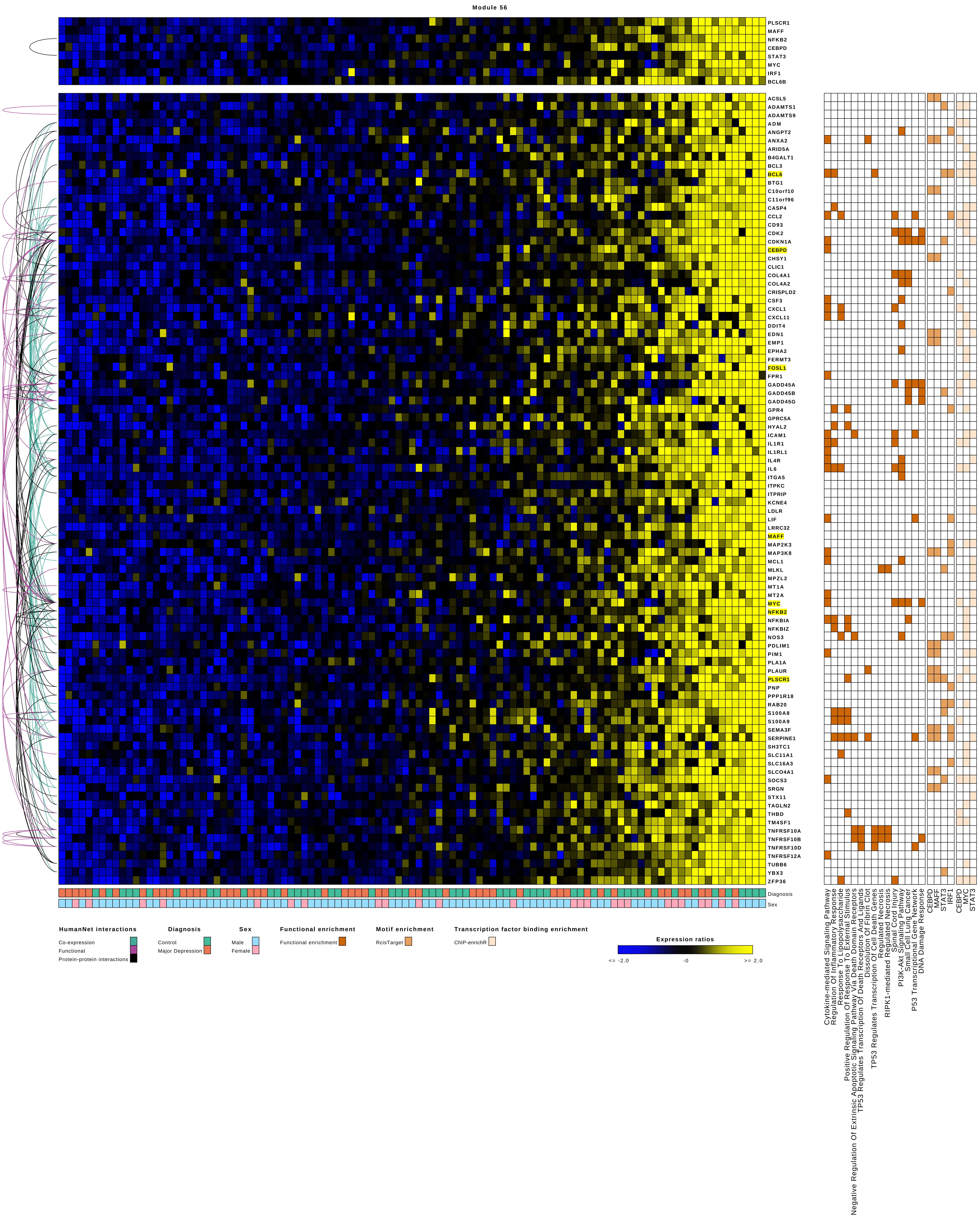
Module 56
PLSCR1
MAFF
NFKB2
CEBPD
STAT3
MYC
IRF1
BCL6B
ACSL5
ADAMTS1
ADAMTS9
ADM
ANGPT2
ANXA2
ARID5A
B4GALT1
BCL3
BCL6
BTG1
C10orf10
C11orf96
CASP4
CCL2
CD93
CDK2
CDKN1A
CEBPD
CHSY1
CLIC1
COL4A1
COL4A2
CRISPLD2
CSF3
CXCL1
CXCL11
DDIT4
EDN1
EMP1
EPHA2
FERMT3
FOSL1
FPR1
GADD45A
GADD45B
GADD45G
GPR4
GPRC5A
HYAL2
ICAM1
IL1R1
IL1RL1
IL4R
IL6
ITGA5
ITPKC
ITPRIP
KCNE4
LDLR
LIF
LRRC32
MAFF
MAP2K3
MAP3K8
MCL1
MLKL
MPZL2
MT1A
MT2A
MYC
NFKB2
NFKBIA
NFKBIZ
NOS3
PDLIM1
PIM1
PLA1A
PLAUR
PLSCR1
PNP
PPP1R18
RAB20
S100A8
S100A9
SEMA3F
SERPINE1
SH3TC1
SLC11A1
SLC16A3
SLCO4A1
SOCS3
SRGN
STX11
TAGLN2
THBD
TM4SF1
TNFRSF10A
TNFRSF10B
TNFRSF10D
TNFRSF12A
TUBB6
YBX3
ZFP36
Diagnosis
Sex
HumanNet interactions
Co-expression
Functional
Protein-protein interactions
Diagnosis
Control
Major Depression
Sex
Male
Female
Functional enrichment
Functional enrichment
Motif enrichment
RcisTarget
Transcription factor binding enrichment
ChIP-enrichR
Expression ratios
<= -2.0
-0
>= 2.0
Cytokine-mediated Signaling Pathway
Regulation Of Inflammatory Response
Response To Lipopolysaccharide
Positive Regulation Of Response To External Stimulus
Negative Regulation Of Extrinsic Apoptotic Signaling Pathway Via Death Domain Receptors
TP53 Regulates Transcription Of Death Receptors And Ligands
Dissolution Of Fibrin Clot
TP53 Regulates Transcription Of Cell Death Genes
Regulated Necrosis
RIPK1-mediated Regulated Necrosis
Spinal Cord Injury
PI3K-Akt Signaling Pathway
Small Cell Lung Cancer
P53 Transcriptional Gene Network
DNA Damage Response
CEBPD
MAFF
STAT3
IRF1
CEBPD
MYC
STAT3

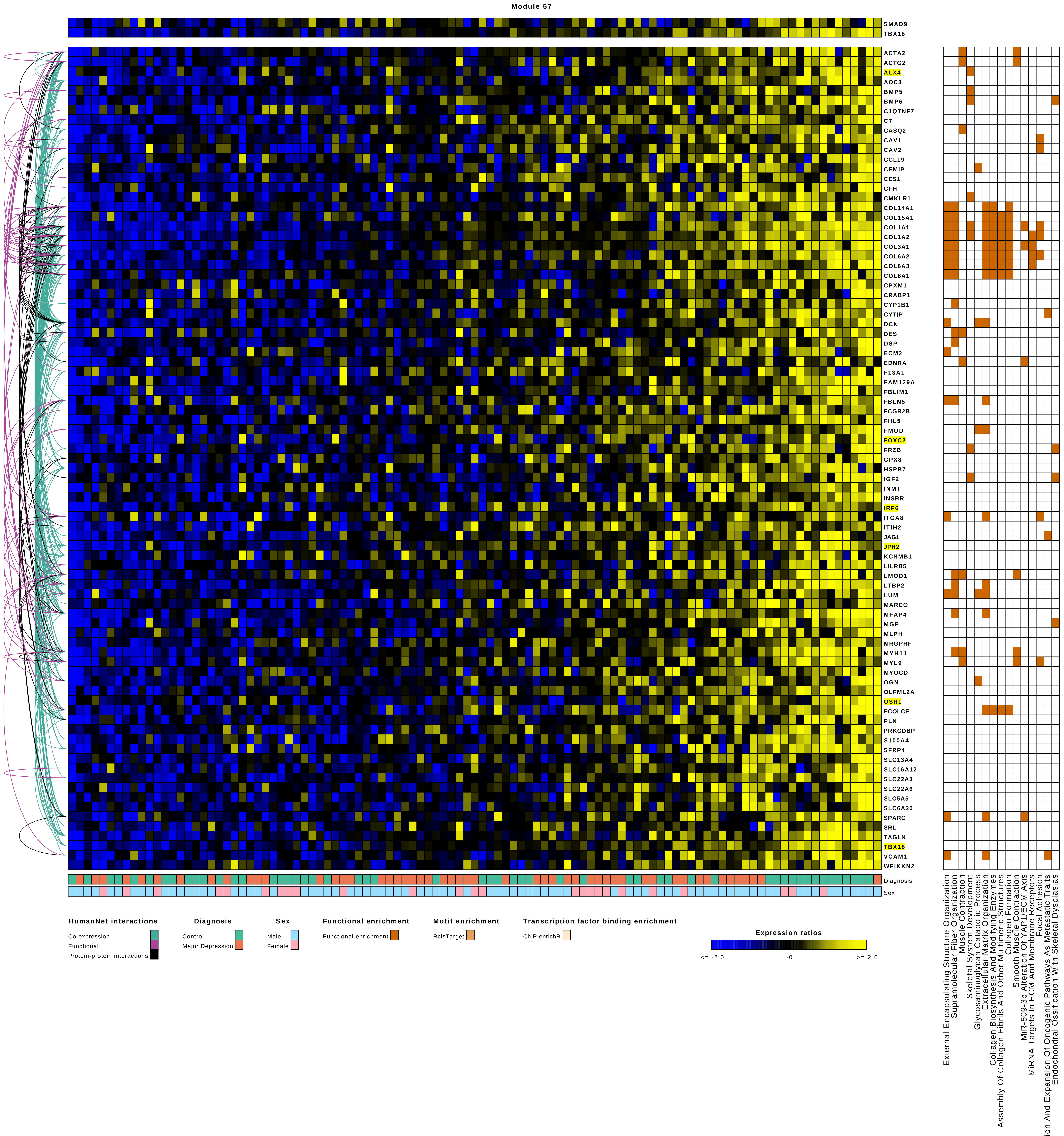
Module 57
SMAD9
TBX18
ACTA2
ACTG2
ALX4
AOC3
BMP5
BMP6
C1QTNF7
C7
CASQ2
CAV1
CAV2
CCL19
CEMIP
CES1
CFH
CMKLR1
COL14A1
COL15A1
COL1A1
COL1A2
COL3A1
COL6A2
COL6A3
COL8A1
CPXM1
CRABP1
CYP1B1
CYTIP
DCN
DES
DSP
ECM2
EDNRA
F13A1
FAM129A
FBLIM1
FBLN5
FCGR2B
FHL5
FMOD
FOXC2
FRZB
GPX8
HSPB7
IGF2
INMT
INSRR
IRF6
ITGA8
ITIH2
JAG1
JPH2
KCNMB1
LILRB5
LMOD1
LTBP2
LUM
MARCO
MFAP4
MGP
MLPH
MRGPRF
MYH11
MYL9
MYOCD
OGN
OLFML2A
OSR1
PCOLCE
PLN
PRKCDBP
S100A4
SFRP4
SLC13A4
SLC16A12
SLC22A3
SLC22A6
SLC5A5
SLC6A20
SPARC
SRL
TAGLN
TBX18
VCAM1
WFIKKN2
Diagnosis
Sex
External Encapsulating Structure Organization
Supramolecular Fiber Organization
Muscle Contraction
Skeletal System Development
Glycosaminoglycan Catabolic Process
Extracellular Matrix Organization
Collagen Biosynthesis And Modifying Enzymes
Assembly Of Collagen Fibrils And Other Multimeric Structures
Collagen Formation
Smooth Muscle Contraction
MiR-509-3p Alteration Of YAP1/ECM Axis
MiRNA Targets In ECM And Membrane Receptors
Focal Adhesion
Amplification And Expansion Of Oncogenic Pathways As Metastatic Traits
Endochondral Ossification With Skeletal Dysplasias
HumanNet interactions
Co-expression
Functional
Protein-protein interactions
Diagnosis
Control
Major Depression
Sex
Male
Female
Functional enrichment
Functional enrichment
Motif enrichment
RcisTarget
Transcription factor binding enrichment
ChIP-enrichR
Expression ratios
<= -2.0
-0
>= 2.0

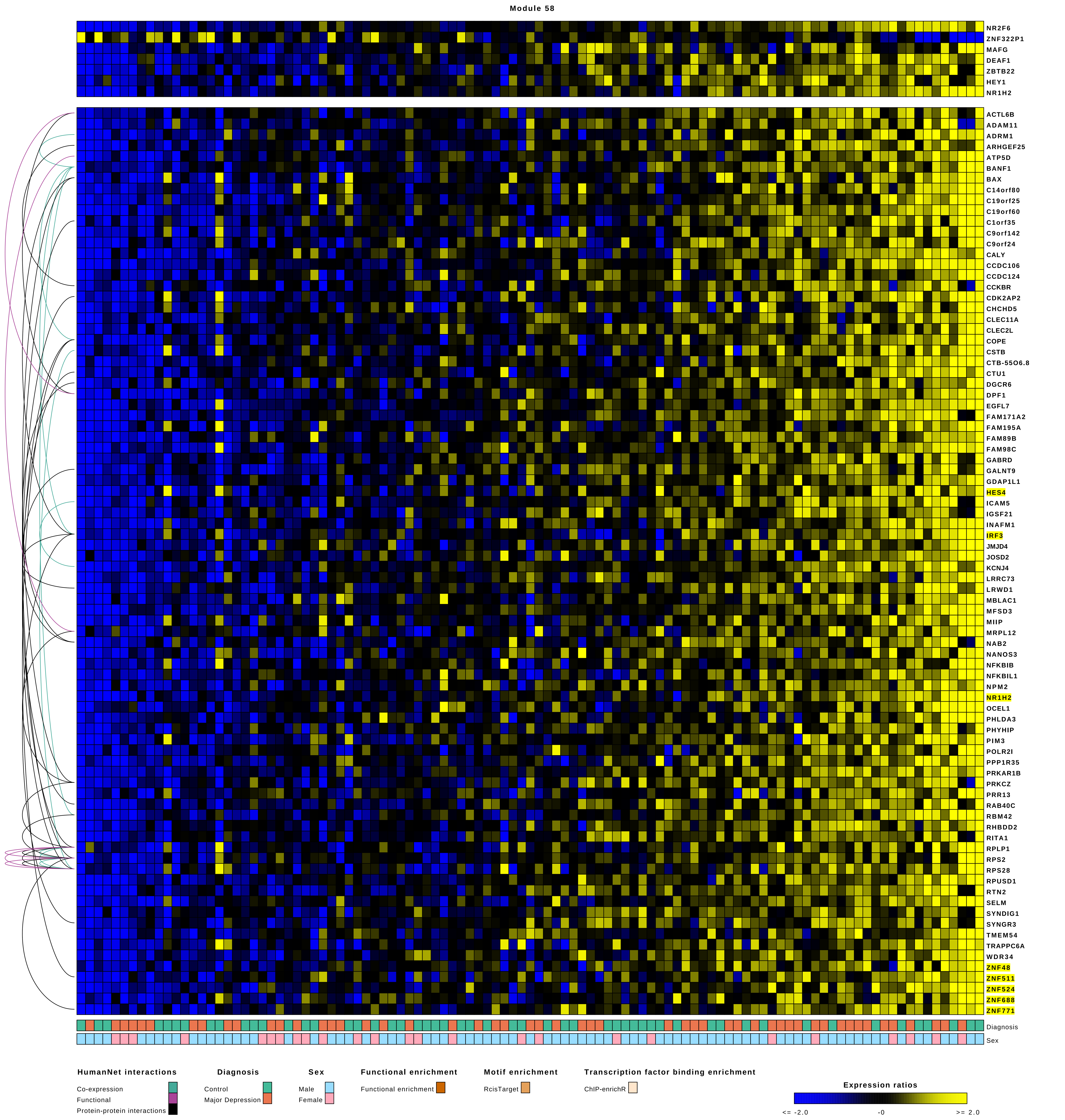
Module 58
NR2F6
ZNF322P1
MAFG
DEAF1
ZBTB22
HEY1
NR1H2
ACTL6B
ADAM11
ADRM1
ARHGEF25
ATP5D
BANF1
BAX
C14orf80
C19orf25
C19orf60
C1orf35
C9orf142
C9orf24
CALY
CCDC106
CCDC124
CCKBR
CDK2AP2
CHCHD5
CLEC11A
CLEC2L
COPE
CSTB
CTB-55O6.8
CTU1
DGCR6
DPF1
EGFL7
FAM171A2
FAM195A
FAM89B
FAM98C
GABRD
GALNT9
GDAP1L1
HES4
ICAM5
IGSF21
INAFM1
IRF3
JMJD4
JOSD2
KCNJ4
LRRC73
LRWD1
MBLAC1
MFSD3
MIIP
MRPL12
NAB2
NANOS3
NFKBIB
NFKBIL1
NPM2
NR1H2
OCEL1
PHLDA3
PHYHIP
PIM3
POLR2I
PPP1R35
PRKAR1B
PRKCZ
PRR13
RAB40C
RBM42
RHBDD2
RITA1
RPLP1
RPS2
RPS28
RPUSD1
RTN2
SELM
SYNDIG1
SYNGR3
TMEM54
TRAPPC6A
WDR34
ZNF48
ZNF511
ZNF524
ZNF688
ZNF771
Diagnosis
Sex
HumanNet interactions
Co-expression
Functional
Protein-protein interactions
Diagnosis
Control
Major Depression
Sex
Male
Female
Functional enrichment
Functional enrichment
Motif enrichment
RcisTarget
Transcription factor binding enrichment
ChIP-enrichR
Expression ratios
<= -2.0
-0
>= 2.0

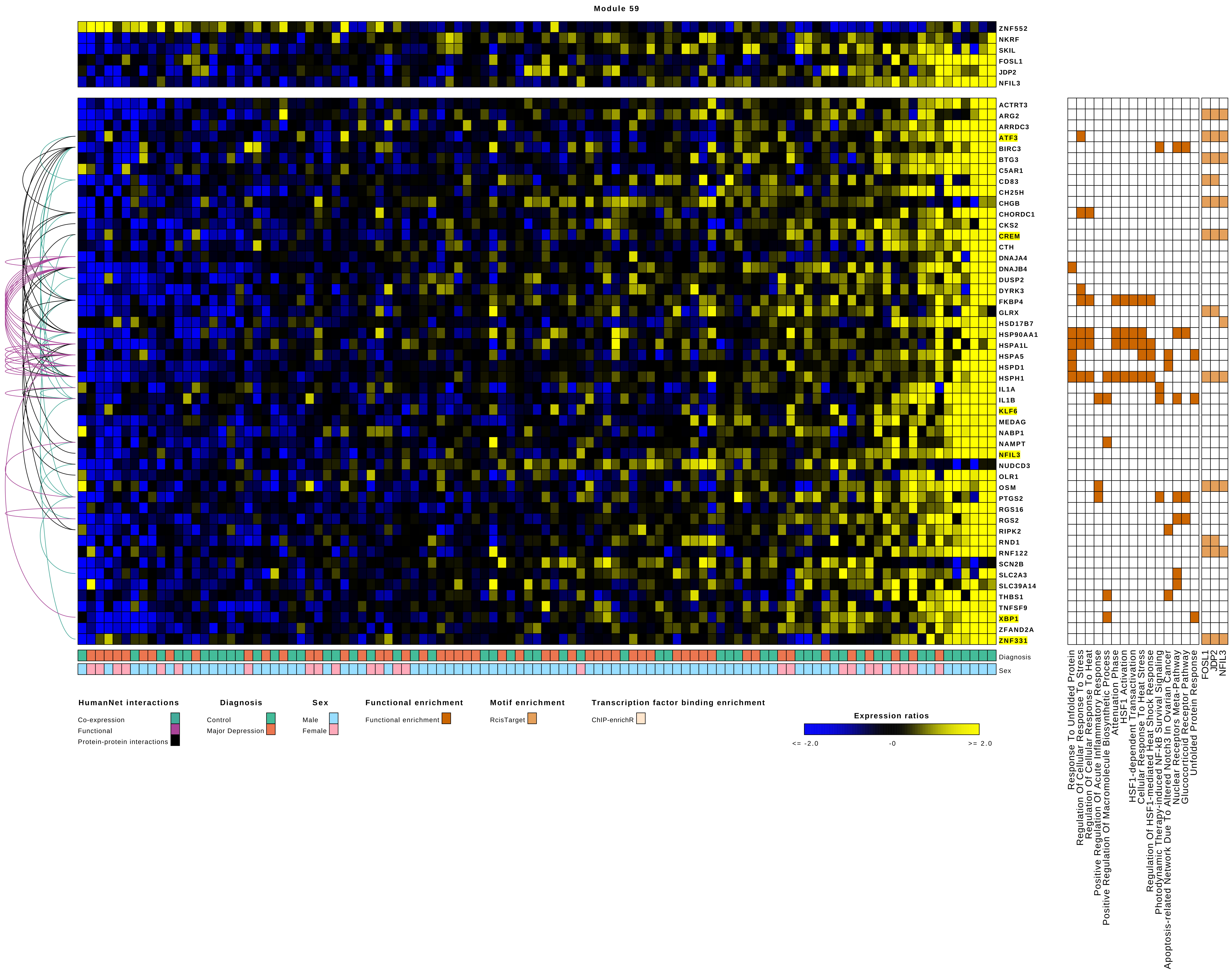

Module 59
ZNF552
NKRF
SKIL
FOSL1
JDP2
NFIL3
ACTRT3
ARG2
ARRDC3
ATF3
BIRC3
BTG3
C5AR1
CD83
CH25H
CHGB
CHORDC1
CKS2
CREM
CTH
DNAJA4
DNAJB4
DUSP2
DYRK3
FKBP4
GLRX
HSD17B7
HSP90AA1
HSPA1L
HSPA5
HSPD1
HSPH1
IL1A
IL1B
KLF6
MEDAG
NABP1
NAMPT
NFIL3
NUDCD3
OLR1
OSM
PTGS2
RGS16
RGS2
RIPK2
RND1
RNF122
SCN2B
SLC2A3
SLC39A14
THBS1
TNFSF9
XBP1
ZFAND2A
ZNF331
Diagnosis
Sex
Response To Unfolded Protein
Regulation Of Cellular Response To Stress
Regulation Of Cellular Response To Heat
Positive Regulation Of Acute Inflammatory Response
Positive Regulation Of Macromolecule Biosynthetic Process
Attenuation Phase
HSF1 Activation
HSF1-dependent Transactivation
Cellular Response To Heat Stress
Regulation Of HSF1-mediated Heat Shock Response
Photodynamic Therapy-induced NF-kB Survival Signaling
Apoptosis-related Network Due To Altered Notch3 In Ovarian Cancer
Nuclear Receptors Meta-Pathway
Glucocorticoid Receptor Pathway
Unfolded Protein Response
FOSL1
JDP2
NFIL3
HumanNet interactions
Co-expression
Functional
Protein-protein interactions
Diagnosis
Control
Major Depression
Sex
Male
Female
Functional enrichment
Functional enrichment
Motif enrichment
RcisTarget
Transcription factor binding enrichment
ChIP-enrichR
Expression ratios
<= -2.0
-0
>= 2.0

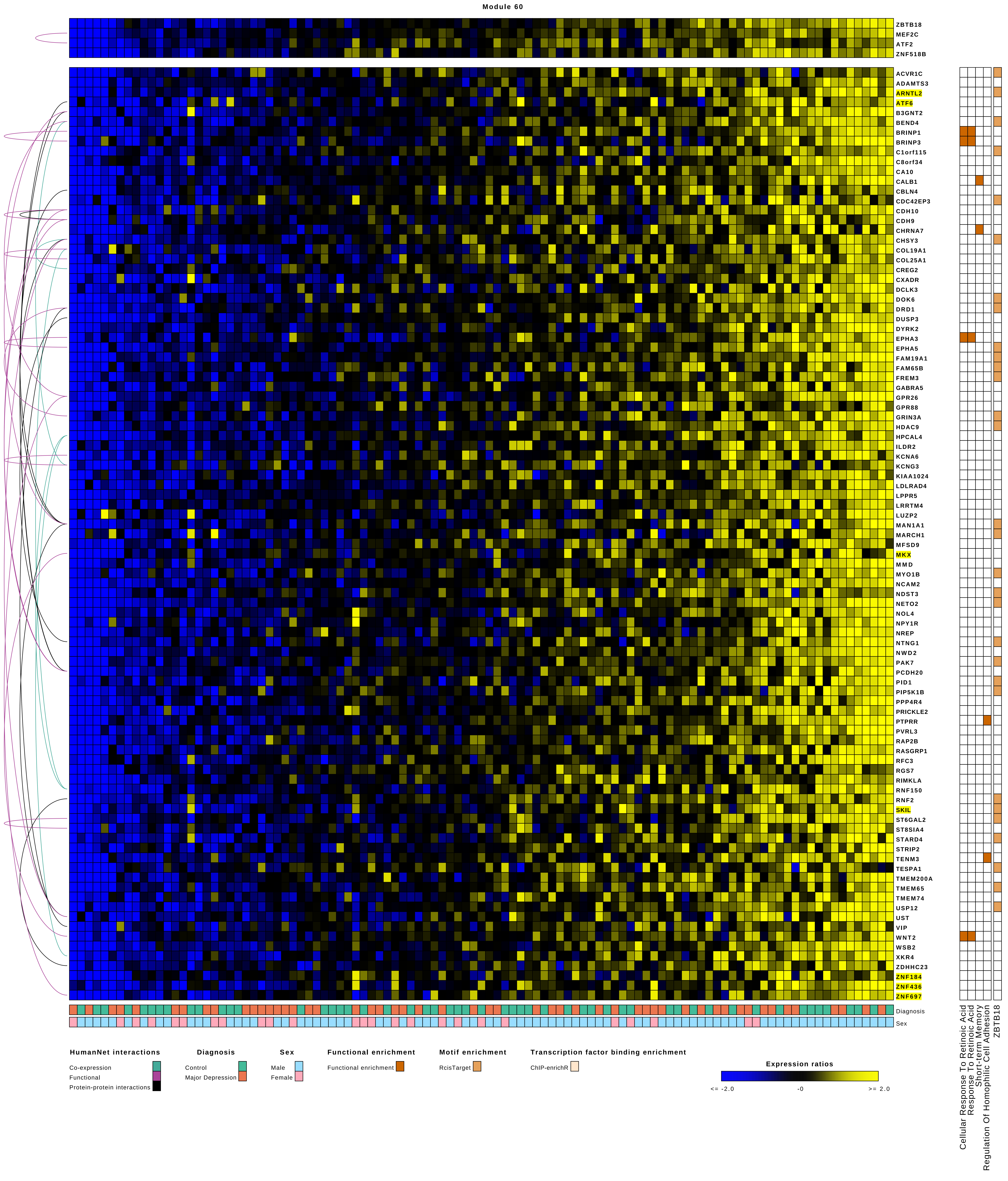
Module 60
ZBTB18
MEF2C
ATF2
ZNF518B
ACVR1C
ADAMTS3
ARNTL2
ATF6
B3GNT2
BEND4
BRINP1
BRINP3
C1orf115
C8orf34
CA10
CALB1
CBLN4
CDC42EP3
CDH10
CDH9
CHRNA7
CHSY3
COL19A1
COL25A1
CREG2
CXADR
DCLK3
DOK6
DRD1
DUSP3
DYRK2
EPHA3
EPHA5
FAM19A1
FAM65B
FREM3
GABRA5
GPR26
GPR88
GRIN3A
HDAC9
HPCAL4
ILDR2
KCNA6
KCNG3
KIAA1024
LDLRAD4
LPPR5
LRRTM4
LUZP2
MAN1A1
MARCH1
MFSD9
MKX
MMD
MYO1B
NCAM2
NDST3
NETO2
NOL4
NPY1R
NREP
NTNG1
NWD2
PAK7
PCDH20
PID1
PIP5K1B
PPP4R4
PRICKLE2
PTPRR
PVRL3
RAP2B
RASGRP1
RFC3
RGS7
RIMKLA
RNF150
RNF2
SKIL
ST6GAL2
ST8SIA4
STARD4
STRIP2
TENM3
TESPA1
TMEM200A
TMEM65
TMEM74
USP12
UST
VIP
WNT2
WSB2
XKR4
ZDHHC23
ZNF184
ZNF436
ZNF697
Diagnosis
Sex
Cellular Response To Retinoic Acid
Response To Retinoic Acid
Short-term Memory
Regulation Of Homophilic Cell Adhesion
ZBTB18
HumanNet interactions
Co-expression
Functional
Protein-protein interactions
Diagnosis
Control
Major Depression
Sex
Male
Female
Functional enrichment
Functional enrichment
Motif enrichment
RcisTarget
Transcription factor binding enrichment
ChIP-enrichR
Expression ratios
<= -2.0
-0
>= 2.0

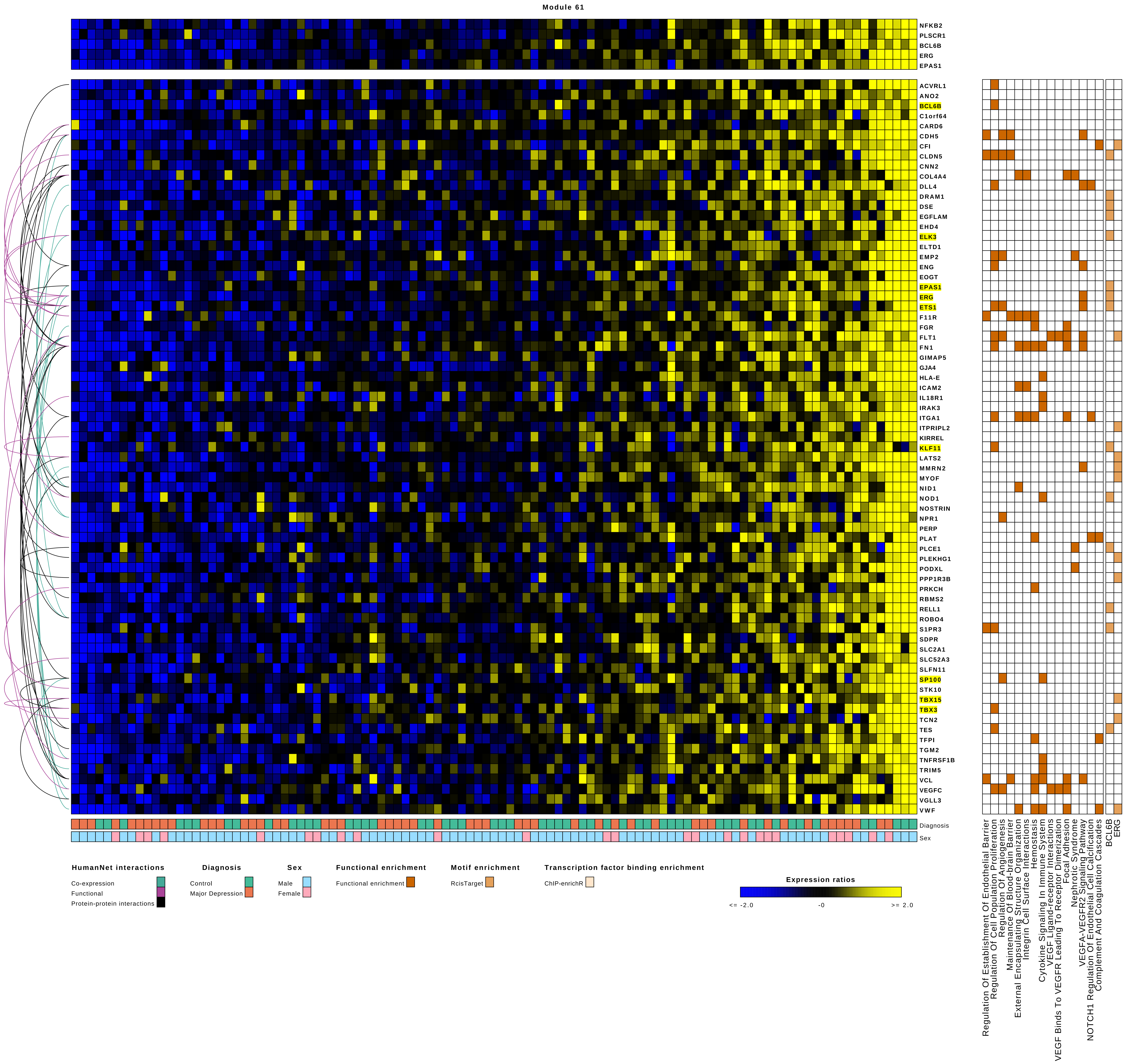
Module 61
NFKB2
PLSCR1
BCL6B
ERG
EPAS1
ACVRL1
ANO2
BCL6B
C1orf64
CARD6
CDH5
CFI
CLDN5
CNN2
COL4A4
DLL4
DRAM1
DSE
EGFLAM
EHD4
ELK3
ELTD1
EMP2
ENG
EOGT
EPAS1
ERG
ETS1
F11R
FGR
FLT1
FN1
GIMAP5
GJA4
HLA-E
ICAM2
IL18R1
IRAK3
ITGA1
ITPRIPL2
KIRREL
KLF11
LATS2
MMRN2
MYOF
NID1
NOD1
NOSTRIN
NPR1
PERP
PLAT
PLCE1
PLEKHG1
PODXL
PPP1R3B
PRKCH
RBMS2
RELL1
ROBO4
S1PR3
SDPR
SLC2A1
SLC52A3
SLFN11
SP100
STK10
TBX15
TBX3
TCN2
TES
TFPI
TGM2
TNFRSF1B
TRIM5
VCL
VEGFC
VGLL3
VWF
Diagnosis
Sex
Regulation Of Establishment Of Endothelial Barrier
Regulation Of Cell Population Proliferation
Regulation Of Angiogenesis
Maintenance Of Blood-brain Barrier
External Encapsulating Structure Organization
Integrin Cell Surface Interactions
Hemostasis
Cytokine Signaling In Immune System
VEGF Ligand-receptor Interactions
VEGF Binds To VEGFR Leading To Receptor Dimerization
Focal Adhesion
Nephrotic Syndrome
VEGFA-VEGFR2 Signaling Pathway
NOTCH1 Regulation Of Endothelial Cell Calcification
Complement And Coagulation Cascades
BCL6B
ERG
HumanNet interactions
Co-expression
Functional
Protein-protein interactions
Diagnosis
Control
Major Depression
Sex
Male
Female
Functional enrichment
Functional enrichment
Motif enrichment
RcisTarget
Transcription factor binding enrichment
ChIP-enrichR
Expression ratios
<= -2.0
-0
>= 2.0

Module 62

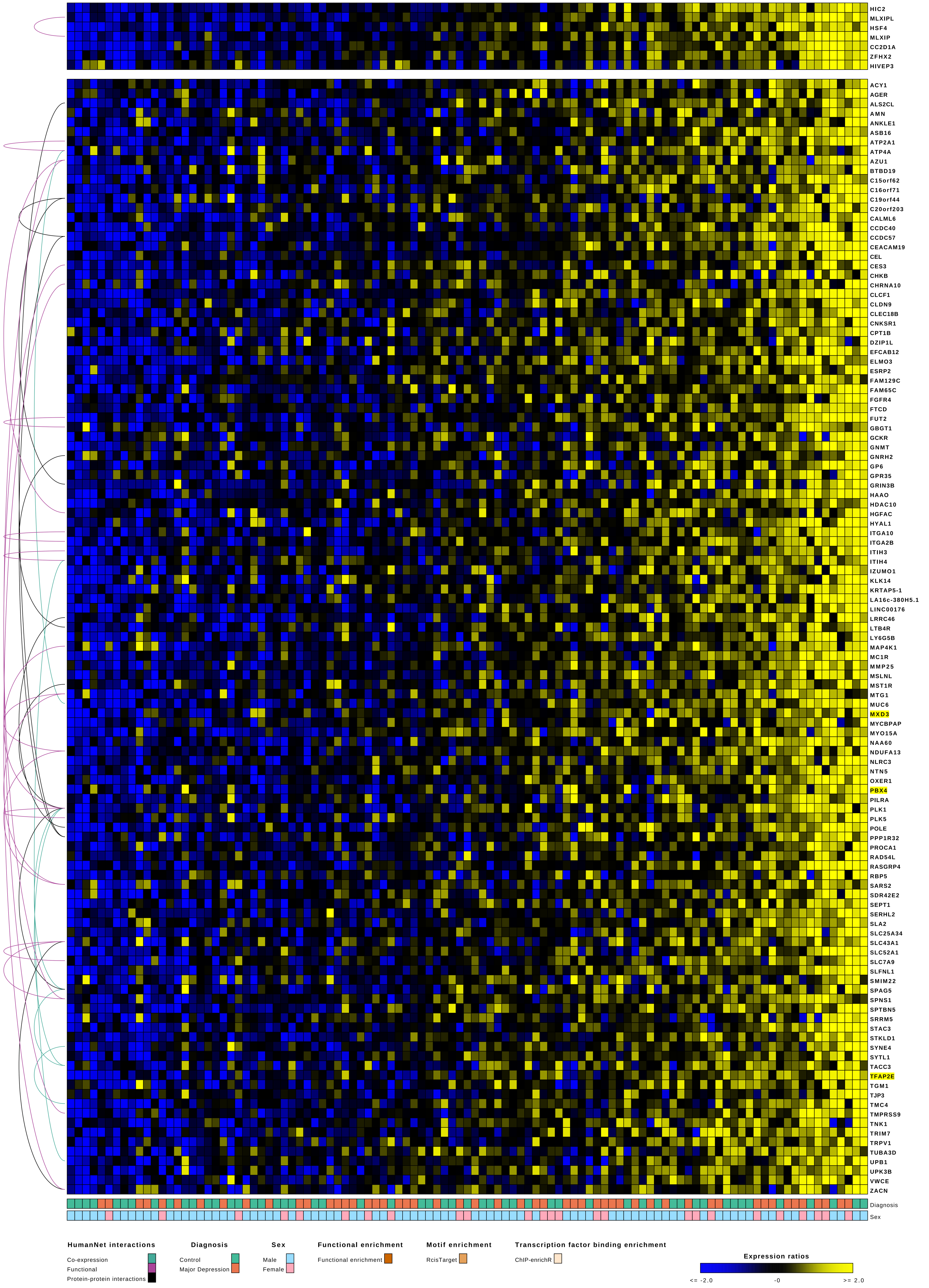

HIC2
MLXIPL
HSF4
MLXIP
CC2D1A
ZFHX2
HIVEP3
ACY1
AGER
ALS2CL
AMN
ANKLE1
ASB16
ATP2A1
ATP4A
AZU1
BTBD19
C15orf62
C16orf71
C19orf44
C20orf203
CALML6
CCDC40
CCDC57
CEACAM19
CEL
CES3
CHKB
CHRNA10
CLCF1
CLDN9
CLEC18B
CNKSR1
CPT1B
DZIP1L
EFCAB12
ELMO3
ESRP2
FAM129C
FAM65C
FGFR4
FTCD
FUT2
GBGT1
GCKR
GNMT
GNRH2
GP6
GPR35
GRIN3B
HAAO
HDAC10
HGFAC
HYAL1
ITGA10
ITGA2B
ITIH3
ITIH4
IZUMO1
KLK14
KRTAP5-1
LA16c-380H5.1
LINC00176
LRRC46
LTB4R
LY6G5B
MAP4K1
MC1R
MMP25
MSLNL
MST1R
MTG1
MUC6
MXD3
MYCBPAP
MYO15A
NAA60
NDUFA13
NLRC3
NTN5
OXER1
PBX4
PILRA
PLK1
PLK5
POLE
PPP1R32
PROCA1
RAD54L
RASGRP4
RBP5
SARS2
SDR42E2
SEPT1
SERHL2
SLA2
SLC25A34
SLC43A1
SLC52A1
SLC7A9
SLFNL1
SMIM22
SPAG5
SPNS1
SPTBN5
SRRM5
STAC3
STKLD1
SYNE4
SYTL1
TACC3
TFAP2E
TGM1
TJP3
TMC4
TMPRSS9
TNK1
TRIM7
TRPV1
TUBA3D
UPB1
UPK3B
VWCE
ZACN
Diagnosis
Sex
HumanNet interactions
Co-expression
Functional
Protein-protein interactions
Diagnosis
Control
Major Depression
Sex
Male
Female
Functional enrichment
Functional enrichment
Motif enrichment
RcisTarget
Transcription factor binding enrichment
ChIP-enrichR
Expression ratios
<= -2.0
-0
>= 2.0

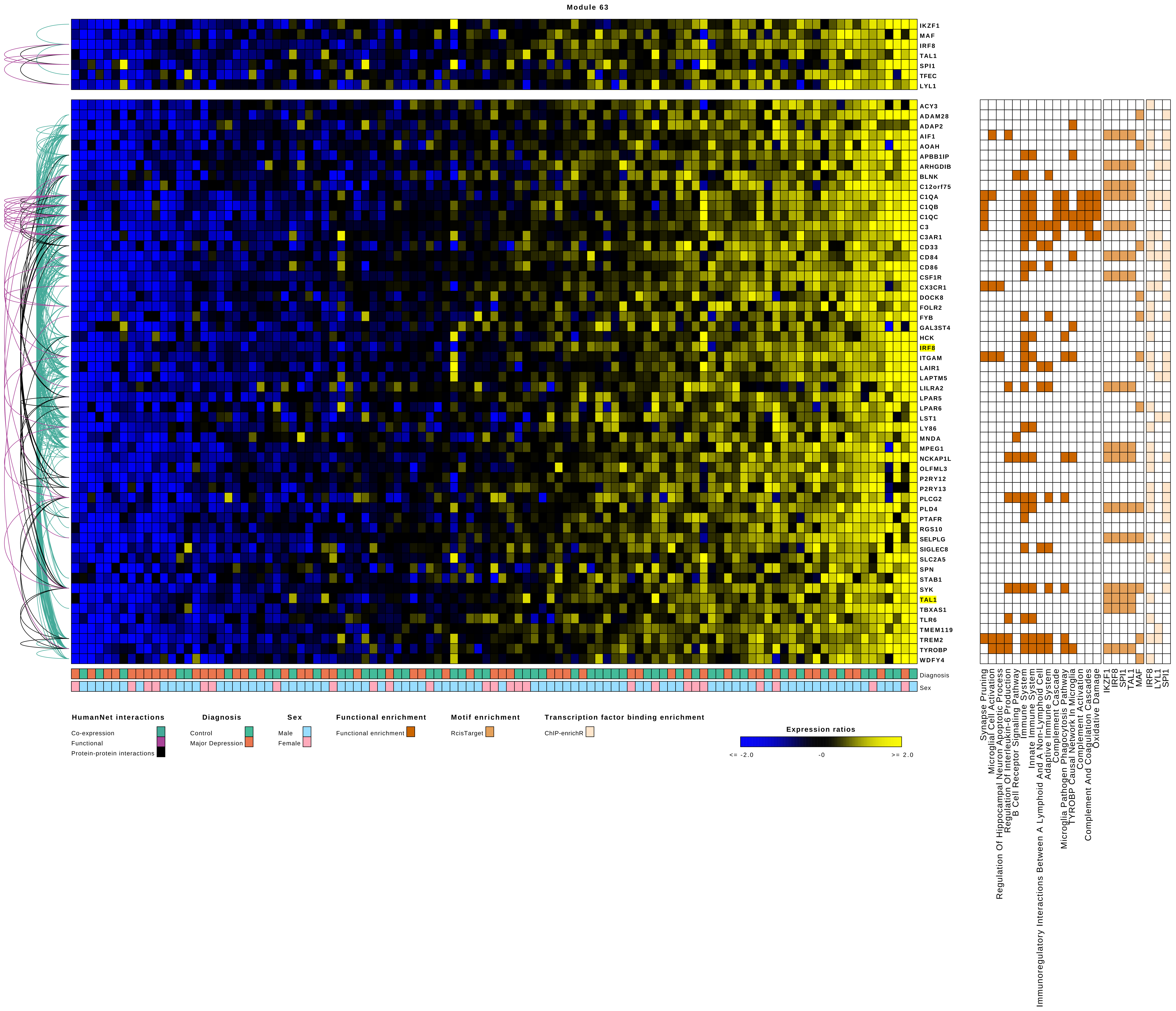

Module 63
IKZF1
MAF
IRF8
TAL1
SPI1
TFEC
LYL1
ACY3
ADAM28
ADAP2
AIF1
AOAH
APBB1IP
ARHGDIB
BLNK
C12orf75
C1QA
C1QB
C1QC
C3
C3AR1
CD33
CD84
CD86
CSF1R
CX3CR1
DOCK8
FOLR2
FYB
GAL3ST4
HCK
IRF8
ITGAM
LAIR1
LAPTM5
LILRA2
LPAR5
LPAR6
LST1
LY86
MNDA
MPEG1
NCKAP1L
OLFML3
P2RY12
P2RY13
PLCG2
PLD4
PTAFR
RGS10
SELPLG
SIGLEC8
SLC2A5
SPN
STAB1
SYK
TAL1
TBXAS1
TLR6
TMEM119
TREM2
TYROBP
WDFY4
Diagnosis
Sex
Synapse Pruning
Microglial Cell Activation
Regulation Of Hippocampal Neuron Apoptotic Process
Regulation Of Interleukin-6 Production
B Cell Receptor Signaling Pathway
Immune System
Innate Immune System
Immunoregulatory Interactions Between A Lymphoid And A Non-Lymphoid Cell
Adaptive Immune System
Complement Cascade
Microglia Pathogen Phagocytosis Pathway
TYROBP Causal Network In Microglia
Complement Activation
Complement And Coagulation Cascades
Oxidative Damage
IKZF1
IRF8
SPI1
TAL1
MAF
IRF8
LYL1
SPI1
HumanNet interactions
Co-expression
Functional
Protein-protein interactions
Diagnosis
Control
Major Depression
Sex
Male
Female
Functional enrichment
Functional enrichment
Motif enrichment
RcisTarget
Transcription factor binding enrichment
ChIP-enrichR
Expression ratios
<= -2.0
-0
>= 2.0

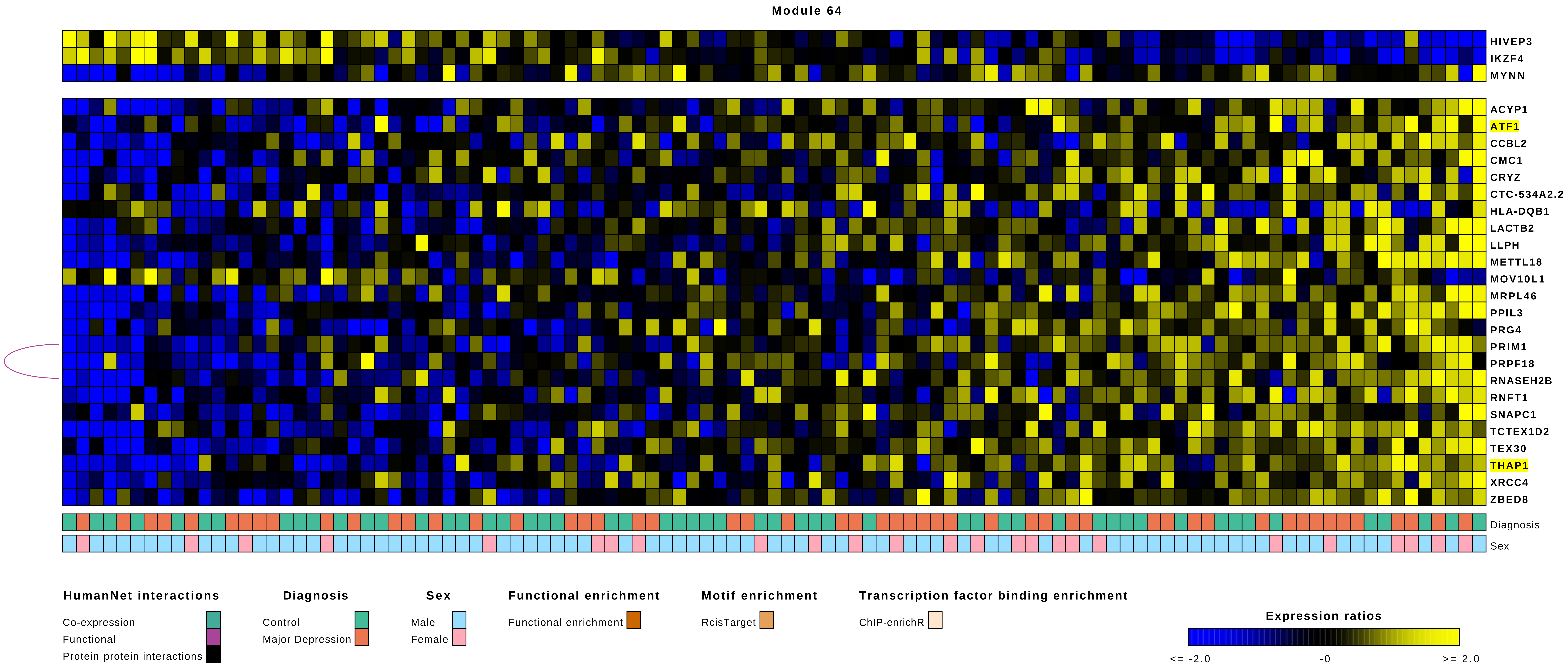

Module 64
HIVEP3
IKZF4
MYNN
ACYP1
ATF1
CCBL2
CMC1
CRYZ
CTC-534A2.2
HLA-DQB1
LACTB2
LLPH
METTL18
MOV10L1
MRPL46
PPIL3
PRG4
PRIM1
PRPF18
RNASEH2B
RNFT1
SNAPC1
TCTEX1D2
TEX30
THAP1
XRCC4
ZBED8
Diagnosis
Sex
HumanNet interactions
Co-expression
Functional
Protein-protein interactions
Diagnosis
Control
Major Depression
Sex
Male
Female
Functional enrichment
Functional enrichment
Motif enrichment
RcisTarget
Transcription factor binding enrichment
ChIP-enrichR
Expression ratios
<= -2.0
-0
>= 2.0

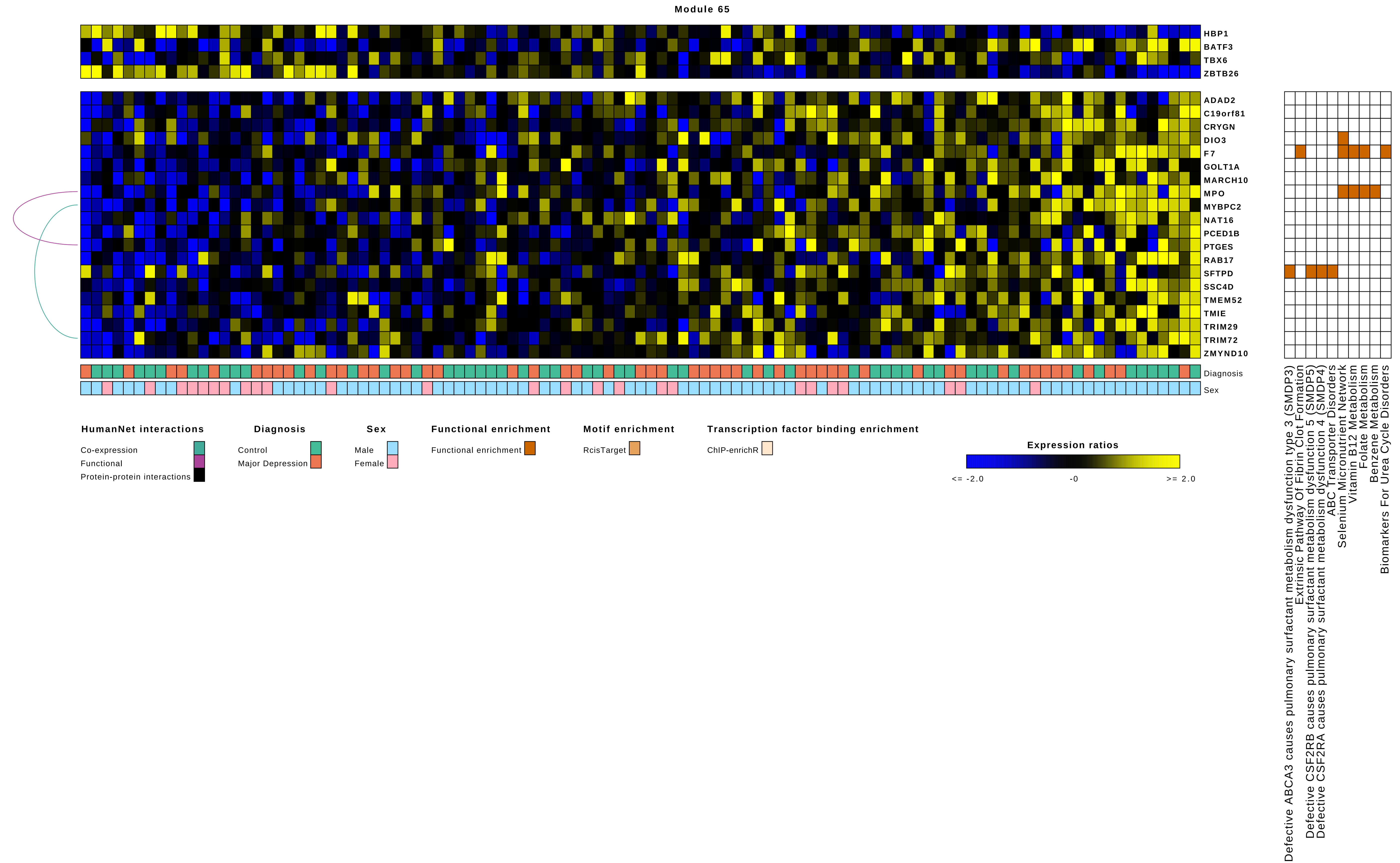
Module 65
HBP1
BATF3
TBX6
ZBTB26
ADAD2
C19orf81
CRYGN
DIO3
F7
GOLT1A
MARCH10
MPO
MYBPC2
NAT16
PCED1B
PTGES
RAB17
SFTPD
SSC4D
TMEM52
TMIE
TRIM29
TRIM72
ZMYND10
Diagnosis
Sex
Defective ABCA3 causes pulmonary surfactant metabolism dysfunction type 3 (SMDP3)
Extrinsic Pathway Of Fibrin Clot Formation
Defective CSF2RB causes pulmonary surfactant metabolism dysfunction 5 (SMDP5)
Defective CSF2RA causes pulmonary surfactant metabolism dysfunction 4 (SMDP4)
ABC Transporter Disorders
Selenium Micronutrient Network
Vitamin B12 Metabolism
Folate Metabolism
Benzene Metabolism
Biomarkers For Urea Cycle Disorders
HumanNet interactions
Co-expression
Functional
Protein-protein interactions
Diagnosis
Control
Major Depression
Sex
Male
Female
Functional enrichment
Functional enrichment
Motif enrichment
RcisTarget
Transcription factor binding enrichment
ChIP-enrichR
Expression ratios
<= -2.0
-0
>= 2.0

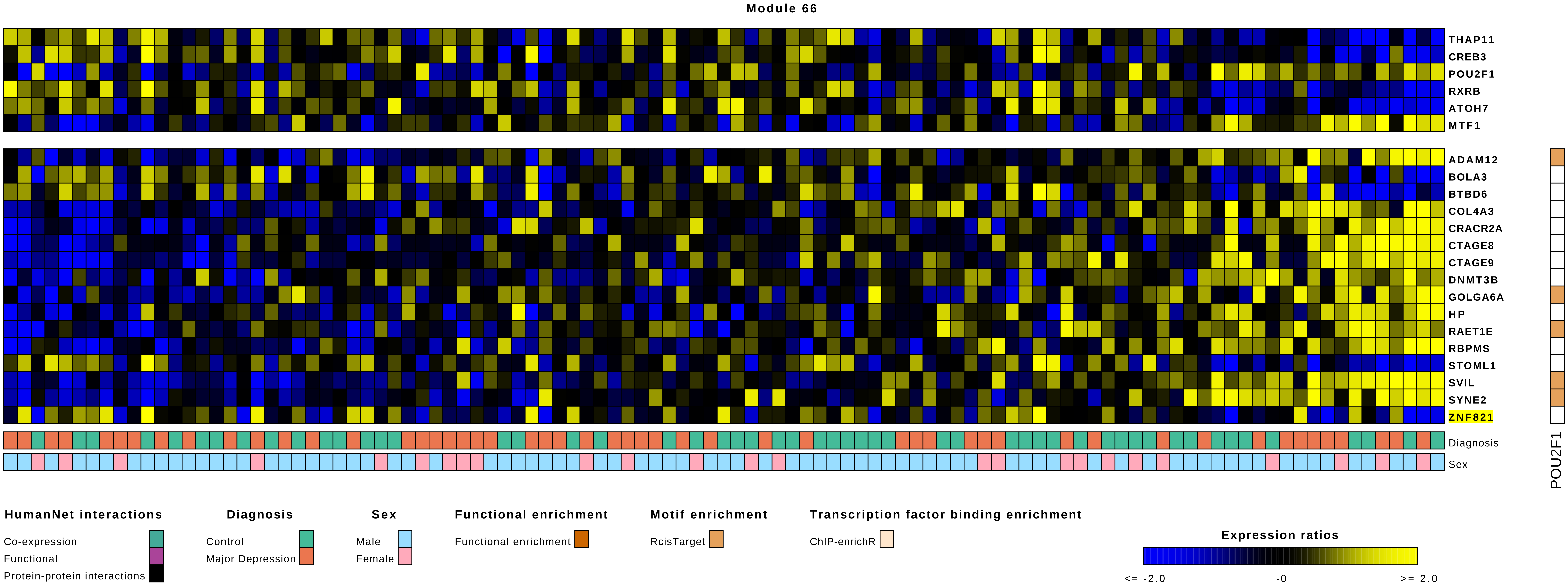
Module 66
THAP11
CREB3
POU2F1
RXRB
ATOH7
MTF1
ADAM12
BOLA3
BTBD6
COL4A3
CRACR2A
CTAGE8
CTAGE9
DNMT3B
GOLGA6A
HP
RAET1E
RBPMS
STOML1
SVIL
SYNE2
ZNF821
Diagnosis
Sex
POU2F1
HumanNet interactions
Co-expression
Functional
Protein-protein interactions
Diagnosis
Control
Major Depression
Sex
Male
Female
Functional enrichment
Functional enrichment
Motif enrichment
RcisTarget
Transcription factor binding enrichment
ChIP-enrichR
Expression ratios
<= -2.0
-0
>= 2.0

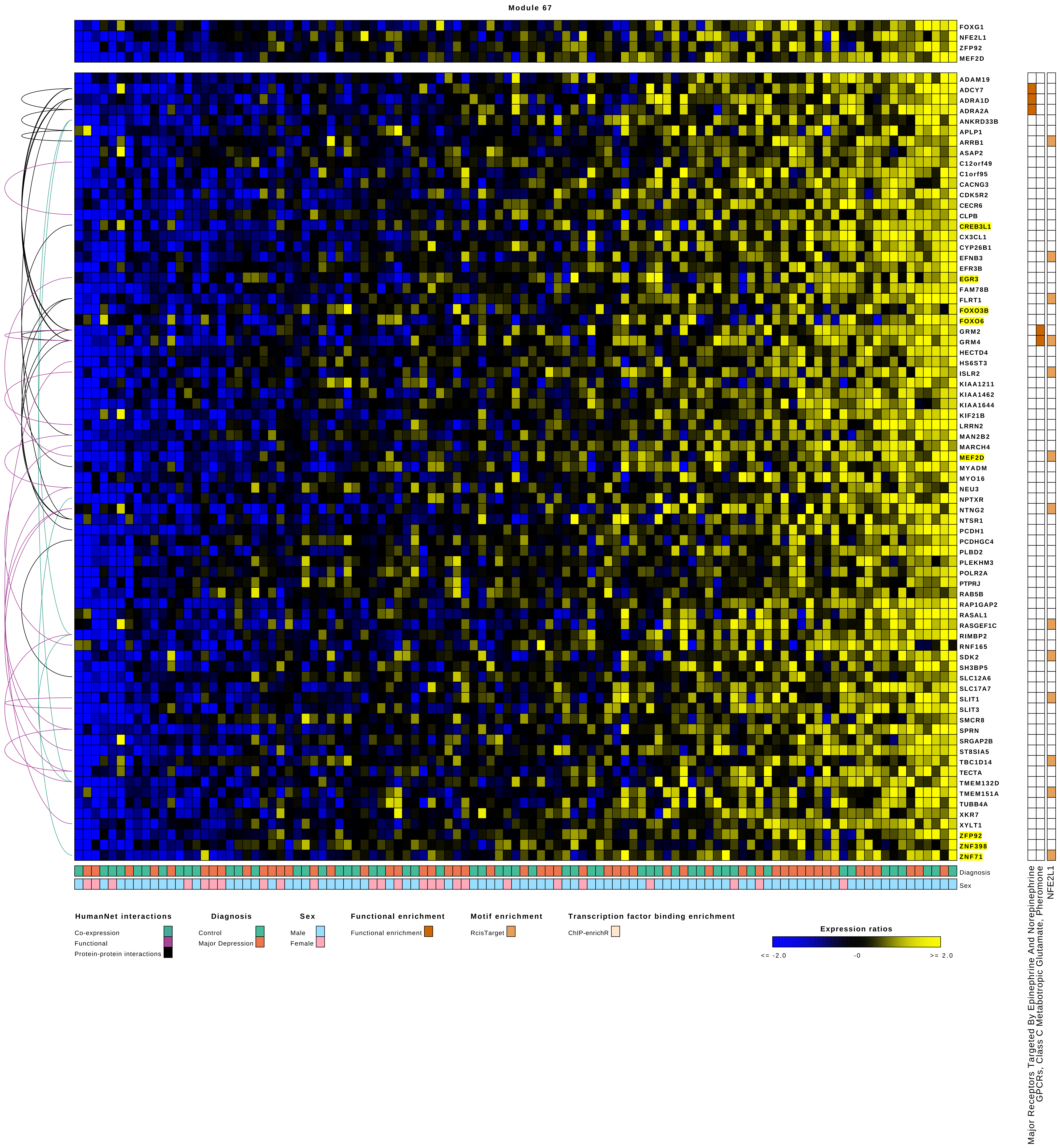
Module 67
FOXG1
NFE2L1
ZFP92
MEF2D
ADAM19
ADCY7
ADRA1D
ADRA2A
ANKRD33B
APLP1
ARRB1
ASAP2
C12orf49
C1orf95
CACNG3
CDK5R2
CECR6
CLPB
CREB3L1
CX3CL1
CYP26B1
EFNB3
EFR3B
EGR3
FAM78B
FLRT1
FOXO3B
FOXO6
GRM2
GRM4
HECTD4
HS6ST3
ISLR2
KIAA1211
KIAA1462
KIAA1644
KIF21B
LRRN2
MAN2B2
MARCH4
MEF2D
MYADM
MYO16
NEU3
NPTXR
NTNG2
NTSR1
PCDH1
PCDHGC4
PLBD2
PLEKHM3
POLR2A
PTPRJ
RAB5B
RAP1GAP2
RASAL1
RASGEF1C
RIMBP2
RNF165
SDK2
SH3BP5
SLC12A6
SLC17A7
SLIT1
SLIT3
SMCR8
SPRN
SRGAP2B
ST8SIA5
TBC1D14
TECTA
TMEM132D
TMEM151A
TUBB4A
XKR7
XYLT1
ZFP92
ZNF398
ZNF71
Diagnosis
Sex
Major Receptors Targeted By Epinephrine And Norepinephrine
GPCRs, Class C Metabotropic Glutamate, Pheromone
NFE2L1
HumanNet interactions
Co-expression
Functional
Protein-protein interactions
Diagnosis
Control
Major Depression
Sex
Male
Female
Functional enrichment
Functional enrichment
Motif enrichment
RcisTarget
Transcription factor binding enrichment
ChIP-enrichR
Expression ratios
<= -2.0
-0
>= 2.0

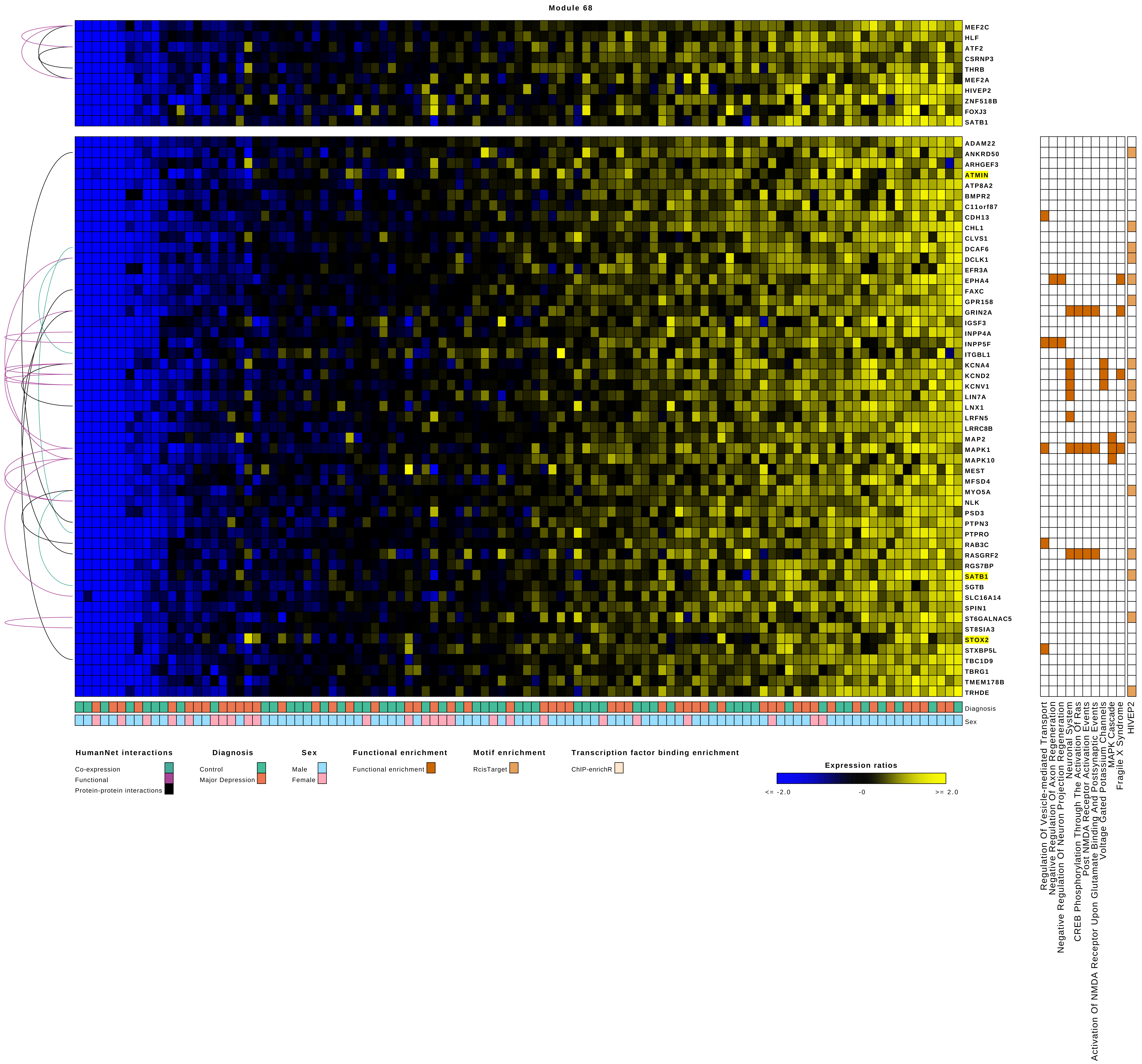

Module 68
MEF2C
HLF
ATF2
CSRNP3
THRB
MEF2A
HIVEP2
ZNF518B
FOXJ3
SATB1
ADAM22
ANKRD50
ARHGEF3
ATMIN
ATP8A2
BMPR2
C11orf87
CDH13
CHL1
CLVS1
DCAF6
DCLK1
EFR3A
EPHA4
FAXC
GPR158
GRIN2A
IGSF3
INPP4A
INPP5F
ITGBL1
KCNA4
KCND2
KCNV1
LIN7A
LNX1
LRFN5
LRRC8B
MAP2
MAPK1
MAPK10
MEST
MFSD4
MYO5A
NLK
PSD3
PTPN3
PTPRO
RAB3C
RASGRF2
RGS7BP
SATB1
SGTB
SLC16A14
SPIN1
ST6GALNAC5
ST8SIA3
STOX2
STXBP5L
TBC1D9
TBRG1
TMEM178B
TRHDE
Diagnosis
Sex
Regulation Of Vesicle-mediated Transport
Negative Regulation Of Axon Regeneration
Negative Regulation Of Neuron Projection Regeneration
Neuronal System
CREB Phosphorylation Through The Activation Of Ras
Post NMDA Receptor Activation Events
Activation Of NMDA Receptor Upon Glutamate Binding And Postsynaptic Events
Voltage Gated Potassium Channels
MAPK Cascade
Fragile X Syndrome
HIVEP2
HumanNet interactions
Co-expression
Functional
Protein-protein interactions
Diagnosis
Control
Major Depression
Sex
Male
Female
Functional enrichment
Functional enrichment
Motif enrichment
RcisTarget
Transcription factor binding enrichment
ChIP-enrichR
Expression ratios
<= -2.0
-0
>= 2.0

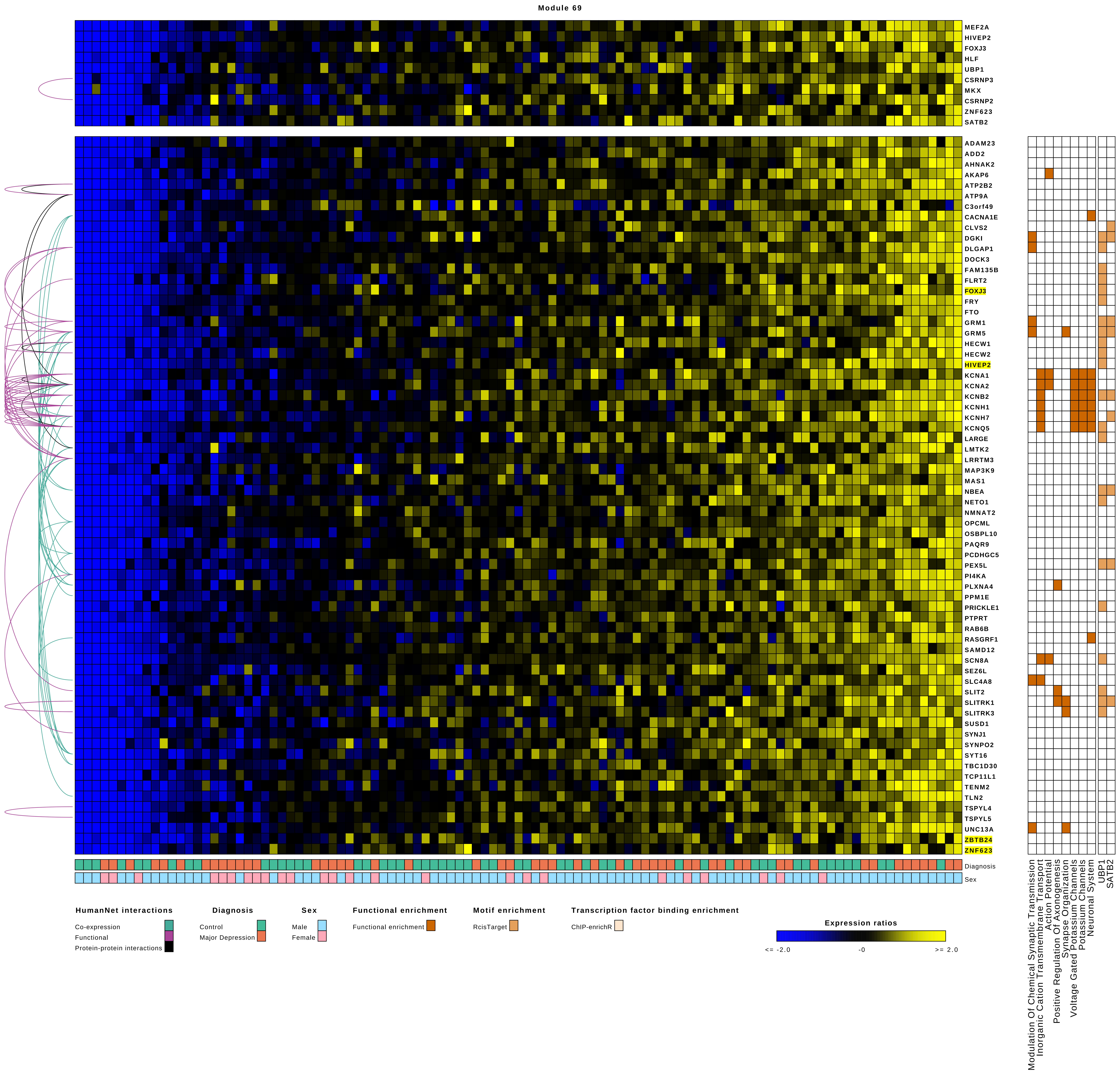
Module 69
MEF2A
HIVEP2
FOXJ3
HLF
UBP1
CSRNP3
MKX
CSRNP2
ZNF623
SATB2
ADAM23
ADD2
AHNAK2
AKAP6
ATP2B2
ATP9A
C3orf49
CACNA1E
CLVS2
DGKI
DLGAP1
DOCK3
FAM135B
FLRT2
FOXJ3
FRY
FTO
GRM1
GRM5
HECW1
HECW2
HIVEP2
KCNA1
KCNA2
KCNB2
KCNH1
KCNH7
KCNQ5
LARGE
LMTK2
LRRTM3
MAP3K9
MAS1
NBEA
NETO1
NMNAT2
OPCML
OSBPL10
PAQR9
PCDHGC5
PEX5L
PI4KA
PLXNA4
PPM1E
PRICKLE1
PTPRT
RAB6B
RASGRF1
SAMD12
SCN8A
SEZ6L
SLC4A8
SLIT2
SLITRK1
SLITRK3
SUSD1
SYNJ1
SYNPO2
SYT16
TBC1D30
TCP11L1
TENM2
TLN2
TSPYL4
TSPYL5
UNC13A
ZBTB24
ZNF623
Diagnosis
Sex
Modulation Of Chemical Synaptic Transmission
Inorganic Cation Transmembrane Transport
Action Potential
Positive Regulation Of Axonogenesis
Synapse Organization
Voltage Gated Potassium Channels
Potassium Channels
Neuronal System
UBP1
SATB2
HumanNet interactions
Co-expression
Functional
Protein-protein interactions
Diagnosis
Control
Major Depression
Sex
Male
Female
Functional enrichment
Functional enrichment
Motif enrichment
RcisTarget
Transcription factor binding enrichment
ChIP-enrichR
Expression ratios
<= -2.0
-0
>= 2.0

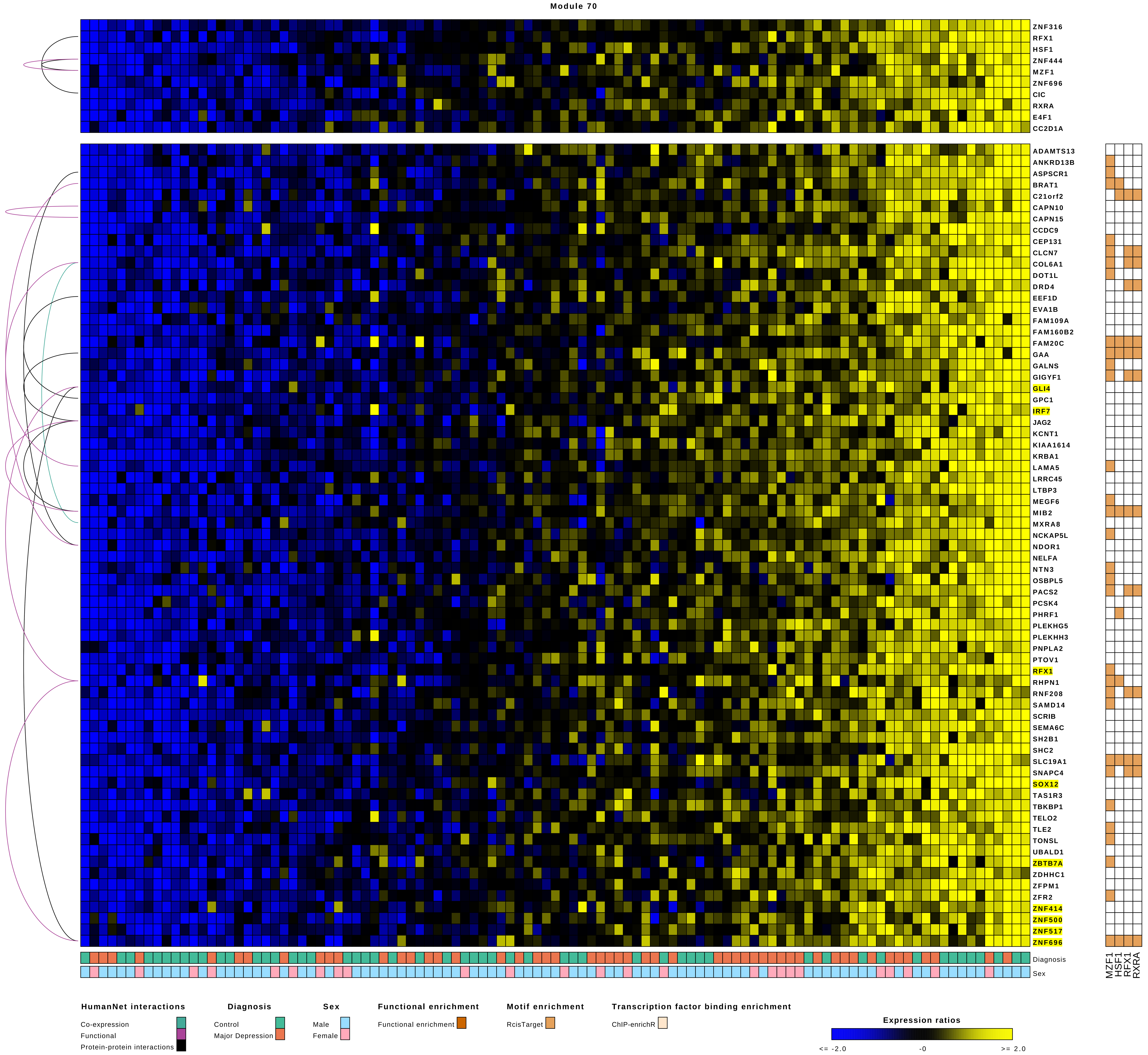

Module 70
ZNF316
RFX1
HSF1
ZNF444
MZF1
ZNF696
CIC
RXRA
E4F1
CC2D1A
ADAMTS13
ANKRD13B
ASPSCR1
BRAT1
C21orf2
CAPN10
CAPN15
CCDC9
CEP131
CLCN7
COL6A1
DOT1L
DRD4
EEF1D
EVA1B
FAM109A
FAM160B2
FAM20C
GAA
GALNS
GIGYF1
GLI4
GPC1
IRF7
JAG2
KCNT1
KIAA1614
KRBA1
LAMA5
LRRC45
LTBP3
MEGF6
MIB2
MXRA8
NCKAP5L
NDOR1
NELFA
NTN3
OSBPL5
PACS2
PCSK4
PHRF1
PLEKHG5
PLEKHH3
PNPLA2
PTOV1
RFX1
RHPN1
RNF208
SAMD14
SCRIB
SEMA6C
SH2B1
SHC2
SLC19A1
SNAPC4
SOX12
TAS1R3
TBKBP1
TELO2
TLE2
TONSL
UBALD1
ZBTB7A
ZDHHC1
ZFPM1
ZFR2
ZNF414
ZNF500
ZNF517
ZNF696
Diagnosis
Sex
MZF1
HSF1
RFX1
RXRA
HumanNet interactions
Co-expression
Functional
Protein-protein interactions
Diagnosis
Control
Major Depression
Sex
Male
Female
Functional enrichment
Functional enrichment
Motif enrichment
RcisTarget
Transcription factor binding enrichment
ChIP-enrichR
Expression ratios
<= -2.0
-0
>= 2.0

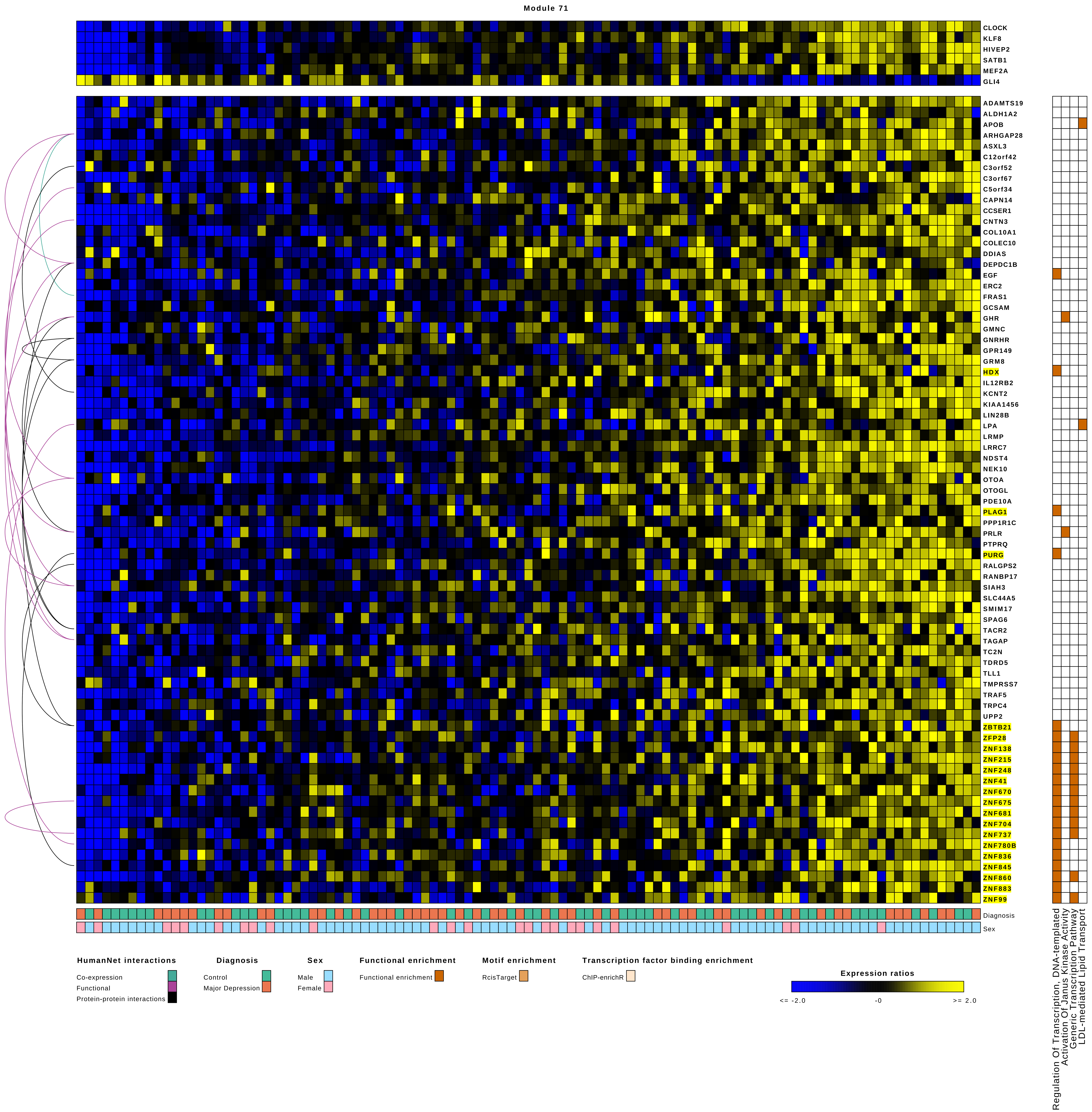
Module 71
CLOCK
KLF8
HIVEP2
SATB1
MEF2A
GLI4
ADAMTS19
ALDH1A2
APOB
ARHGAP28
ASXL3
C12orf42
C3orf52
C3orf67
C5orf34
CAPN14
CCSER1
CNTN3
COL10A1
COLEC10
DDIAS
DEPDC1B
EGF
ERC2
FRAS1
GCSAM
GHR
GMNC
GNRHR
GPR149
GRM8
HDX
IL12RB2
KCNT2
KIAA1456
LIN28B
LPA
LRMP
LRRC7
NDST4
NEK10
OTOA
OTOGL
PDE10A
PLAG1
PPP1R1C
PRLR
PTPRQ
PURG
RALGPS2
RANBP17
SIAH3
SLC44A5
SMIM17
SPAG6
TACR2
TAGAP
TC2N
TDRD5
TLL1
TMPRSS7
TRAF5
TRPC4
UPP2
ZBTB21
ZFP28
ZNF138
ZNF215
ZNF248
ZNF41
ZNF670
ZNF675
ZNF681
ZNF704
ZNF737
ZNF780B
ZNF836
ZNF845
ZNF860
ZNF883
ZNF99
Diagnosis
Sex
Regulation Of Transcription, DNA-templated
Activation Of Janus Kinase Activity
Generic Transcription Pathway
LDL-mediated Lipid Transport
HumanNet interactions
Co-expression
Functional
Protein-protein interactions
Diagnosis
Control
Major Depression
Sex
Male
Female
Functional enrichment
Functional enrichment
Motif enrichment
RcisTarget
Transcription factor binding enrichment
ChIP-enrichR
Expression ratios
<= -2.0
-0
>= 2.0

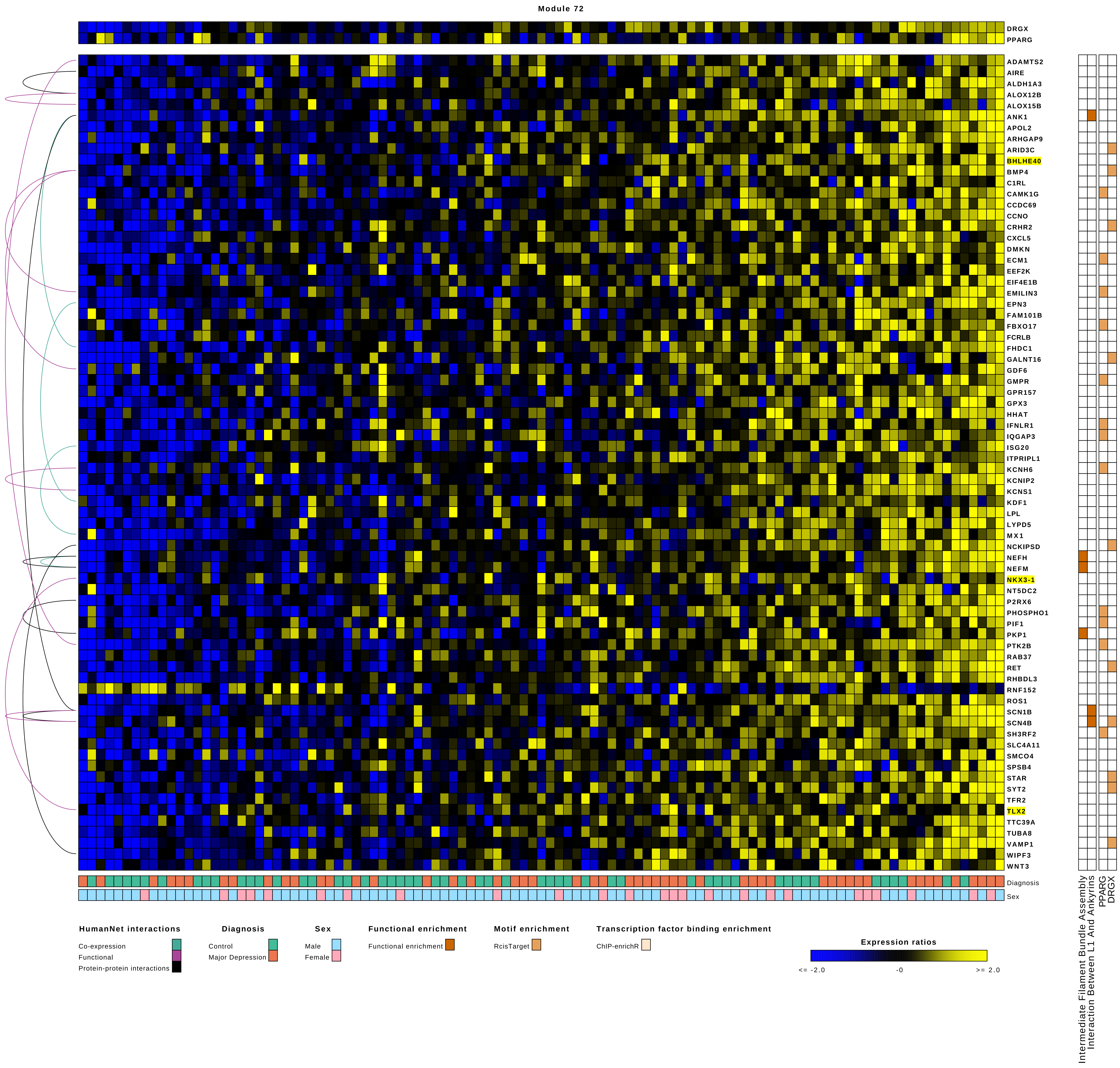
Module 72
DRGX
PPARG
ADAMTS2
AIRE
ALDH1A3
ALOX12B
ALOX15B
ANK1
APOL2
ARHGAP9
ARID3C
BHLHE40
BMP4
C1RL
CAMK1G
CCDC69
CCNO
CRHR2
CXCL5
DMKN
ECM1
EEF2K
EIF4E1B
EMILIN3
EPN3
FAM101B
FBXO17
FCRLB
FHDC1
GALNT16
GDF6
GMPR
GPR157
GPX3
HHAT
IFNLR1
IQGAP3
ISG20
ITPRIPL1
KCNH6
KCNIP2
KCNS1
KDF1
LPL
LYPD5
MX1
NCKIPSD
NEFH
NEFM
NKX3-1
NT5DC2
P2RX6
PHOSPHO1
PIF1
PKP1
PTK2B
RAB37
RET
RHBDL3
RNF152
ROS1
SCN1B
SCN4B
SH3RF2
SLC4A11
SMCO4
SPSB4
STAR
SYT2
TFR2
TLX2
TTC39A
TUBA8
VAMP1
WIPF3
WNT3
Diagnosis
Sex
Intermediate Filament Bundle Assembly
Interaction Between L1 And Ankyrins
PPARG
DRGX
HumanNet interactions
Co-expression
Functional
Protein-protein interactions
Diagnosis
Control
Major Depression
Sex
Male
Female
Functional enrichment
Functional enrichment
Motif enrichment
RcisTarget
Transcription factor binding enrichment
ChIP-enrichR
Expression ratios
<= -2.0
-0
>= 2.0

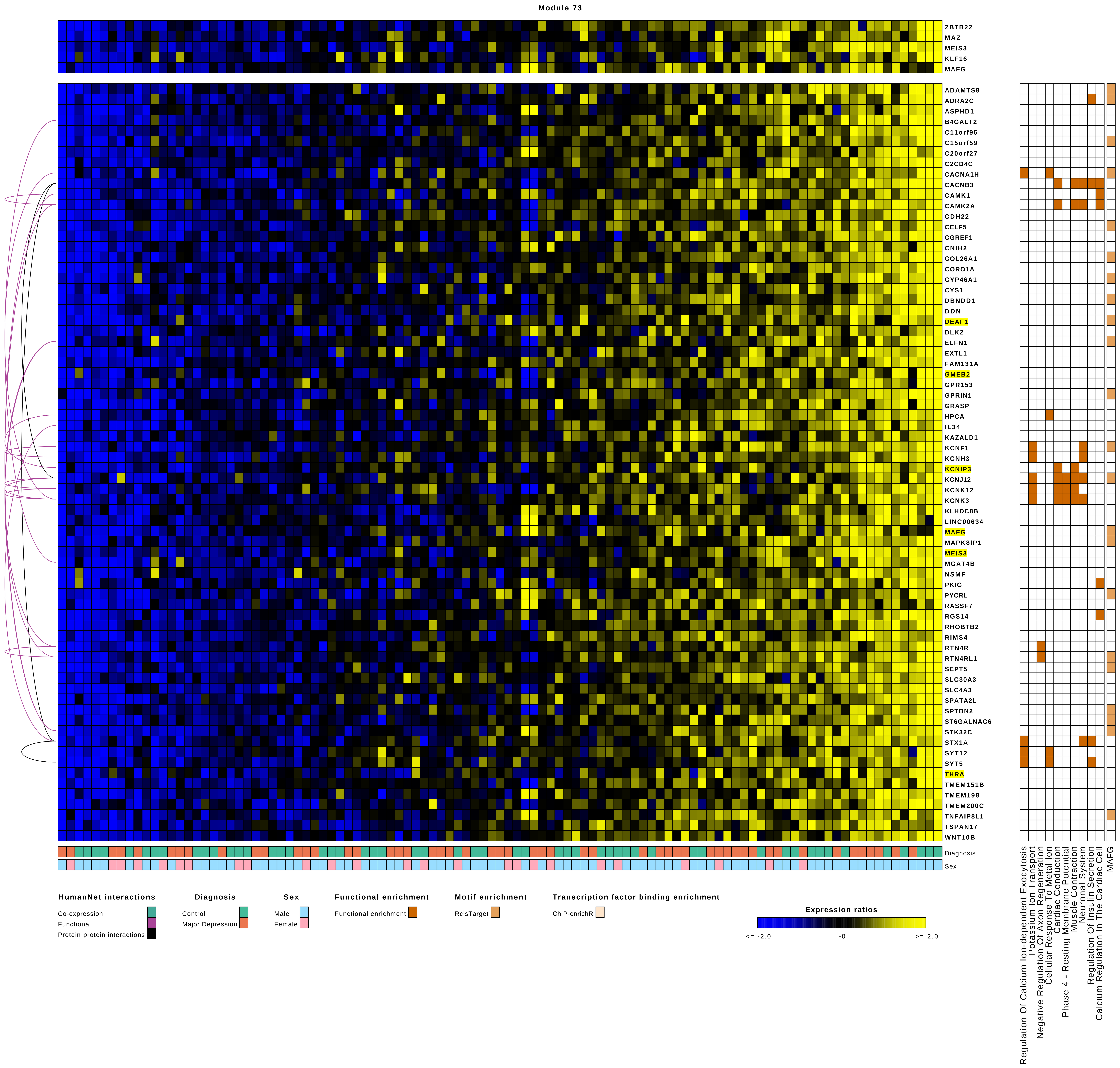

Module 73
ZBTB22
MAZ
MEIS3
KLF16
MAFG
ADAMTS8
ADRA2C
ASPHD1
B4GALT2
C11orf95
C15orf59
C20orf27
C2CD4C
CACNA1H
CACNB3
CAMK1
CAMK2A
CDH22
CELF5
CGREF1
CNIH2
COL26A1
CORO1A
CYP46A1
CYS1
DBNDD1
DDN
DEAF1
DLK2
ELFN1
EXTL1
FAM131A
GMEB2
GPR153
GPRIN1
GRASP
HPCA
IL34
KAZALD1
KCNF1
KCNH3
KCNIP3
KCNJ12
KCNK12
KCNK3
KLHDC8B
LINC00634
MAFG
MAPK8IP1
MEIS3
MGAT4B
NSMF
PKIG
PYCRL
RASSF7
RGS14
RHOBTB2
RIMS4
RTN4R
RTN4RL1
SEPT5
SLC30A3
SLC4A3
SPATA2L
SPTBN2
ST6GALNAC6
STK32C
STX1A
SYT12
SYT5
THRA
TMEM151B
TMEM198
TMEM200C
TNFAIP8L1
TSPAN17
WNT10B
Diagnosis
Sex
Regulation Of Calcium Ion-dependent Exocytosis
Potassium Ion Transport
Negative Regulation Of Axon Regeneration
Cellular Response To Metal Ion
Cardiac Conduction
Phase 4 - Resting Membrane Potential
Muscle Contraction
Neuronal System
Regulation Of Insulin Secretion
Calcium Regulation In The Cardiac Cell
MAFG
HumanNet interactions
Co-expression
Functional
Protein-protein interactions
Diagnosis
Control
Major Depression
Sex
Male
Female
Functional enrichment
Functional enrichment
Motif enrichment
RcisTarget
Transcription factor binding enrichment
ChIP-enrichR
Expression ratios
<= -2.0
-0
>= 2.0

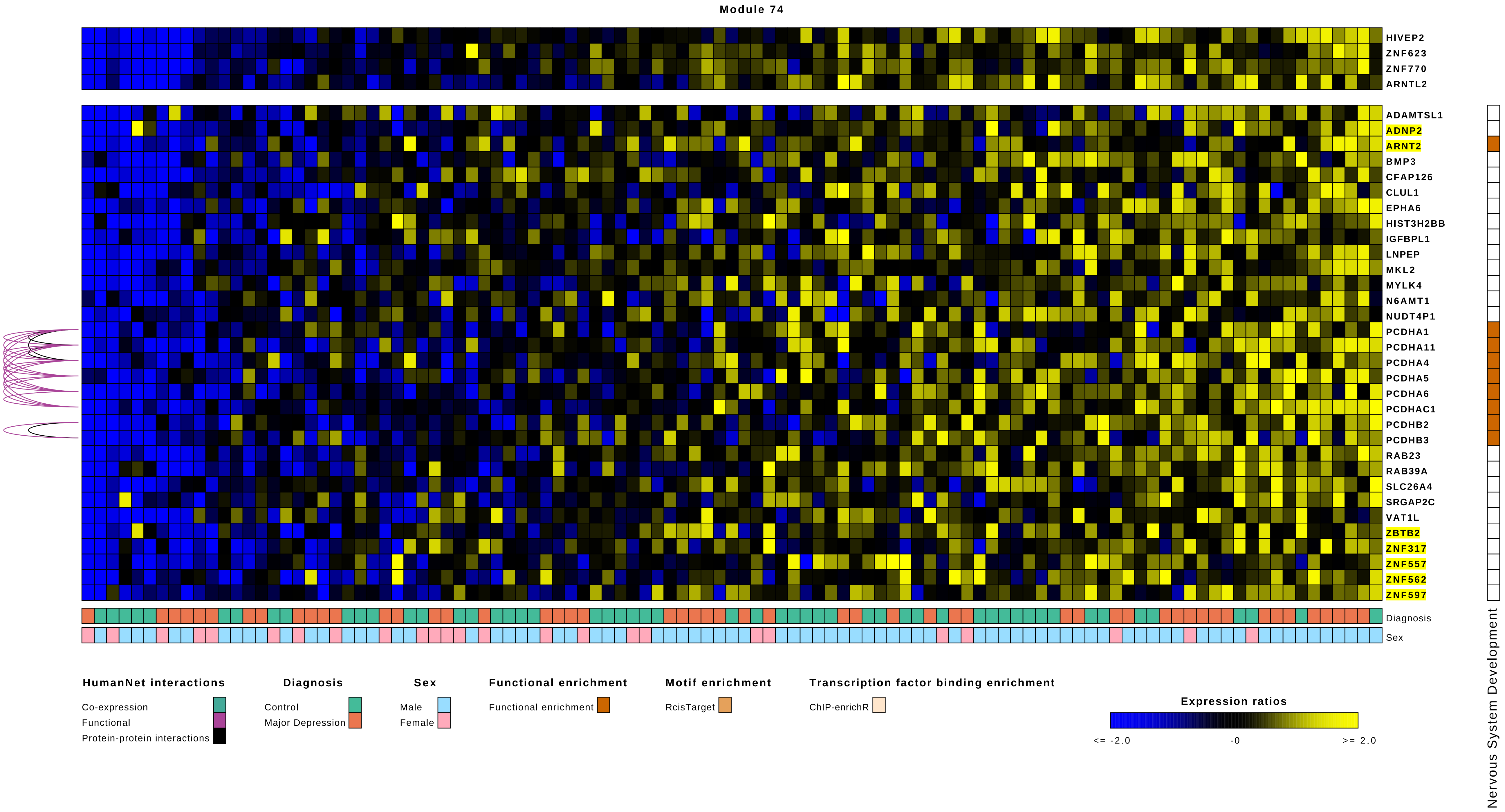

Module 74
HIVEP2
ZNF623
ZNF770
ARNTL2
ADAMTSL1
ADNP2
ARNT2
BMP3
CFAP126
CLUL1
EPHA6
HIST3H2BB
IGFBPL1
LNPEP
MKL2
MYLK4
N6AMT1
NUDT4P1
PCDHA1
PCDHA11
PCDHA4
PCDHA5
PCDHA6
PCDHAC1
PCDHB2
PCDHB3
RAB23
RAB39A
SLC26A4
SRGAP2C
VAT1L
ZBTB2
ZNF317
ZNF557
ZNF562
ZNF597
Diagnosis
Sex
Nervous System Development
HumanNet interactions
Co-expression
Functional
Protein-protein interactions
Diagnosis
Control
Major Depression
Sex
Male
Female
Functional enrichment
Functional enrichment
Motif enrichment
RcisTarget
Transcription factor binding enrichment
ChIP-enrichR
Expression ratios
<= -2.0
-0
>= 2.0

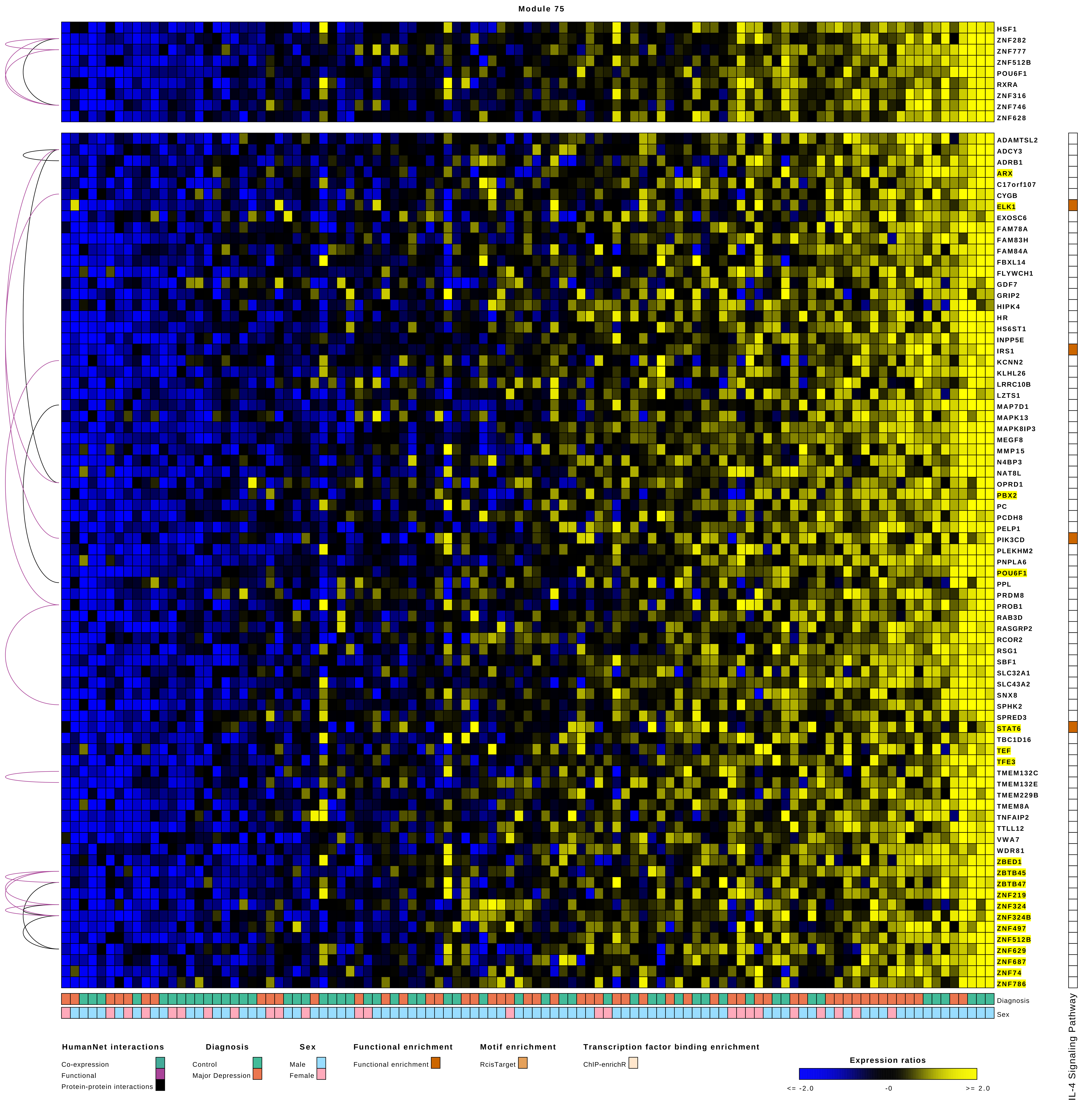
Module 75
HSF1
ZNF282
ZNF777
ZNF512B
POU6F1
RXRA
ZNF316
ZNF746
ZNF628
ADAMTSL2
ADCY3
ADRB1
ARX
C17orf107
CYGB
ELK1
EXOSC6
FAM78A
FAM83H
FAM84A
FBXL14
FLYWCH1
GDF7
GRIP2
HIPK4
HR
HS6ST1
INPP5E
IRS1
KCNN2
KLHL26
LRRC10B
LZTS1
MAP7D1
MAPK13
MAPK8IP3
MEGF8
MMP15
N4BP3
NAT8L
OPRD1
PBX2
PC
PCDH8
PELP1
PIK3CD
PLEKHM2
PNPLA6
POU6F1
PPL
PRDM8
PROB1
RAB3D
RASGRP2
RCOR2
RSG1
SBF1
SLC32A1
SLC43A2
SNX8
SPHK2
SPRED3
STAT6
TBC1D16
TEF
TFE3
TMEM132C
TMEM132E
TMEM229B
TMEM8A
TNFAIP2
TTLL12
VWA7
WDR81
ZBED1
ZBTB45
ZBTB47
ZNF219
ZNF324
ZNF324B
ZNF497
ZNF512B
ZNF629
ZNF687
ZNF74
ZNF786
Diagnosis
Sex
IL-4 Signaling Pathway
HumanNet interactions
Co-expression
Functional
Protein-protein interactions
Diagnosis
Control
Major Depression
Sex
Male
Female
Functional enrichment
Functional enrichment
Motif enrichment
RcisTarget
Transcription factor binding enrichment
ChIP-enrichR
Expression ratios
<= -2.0
-0
>= 2.0

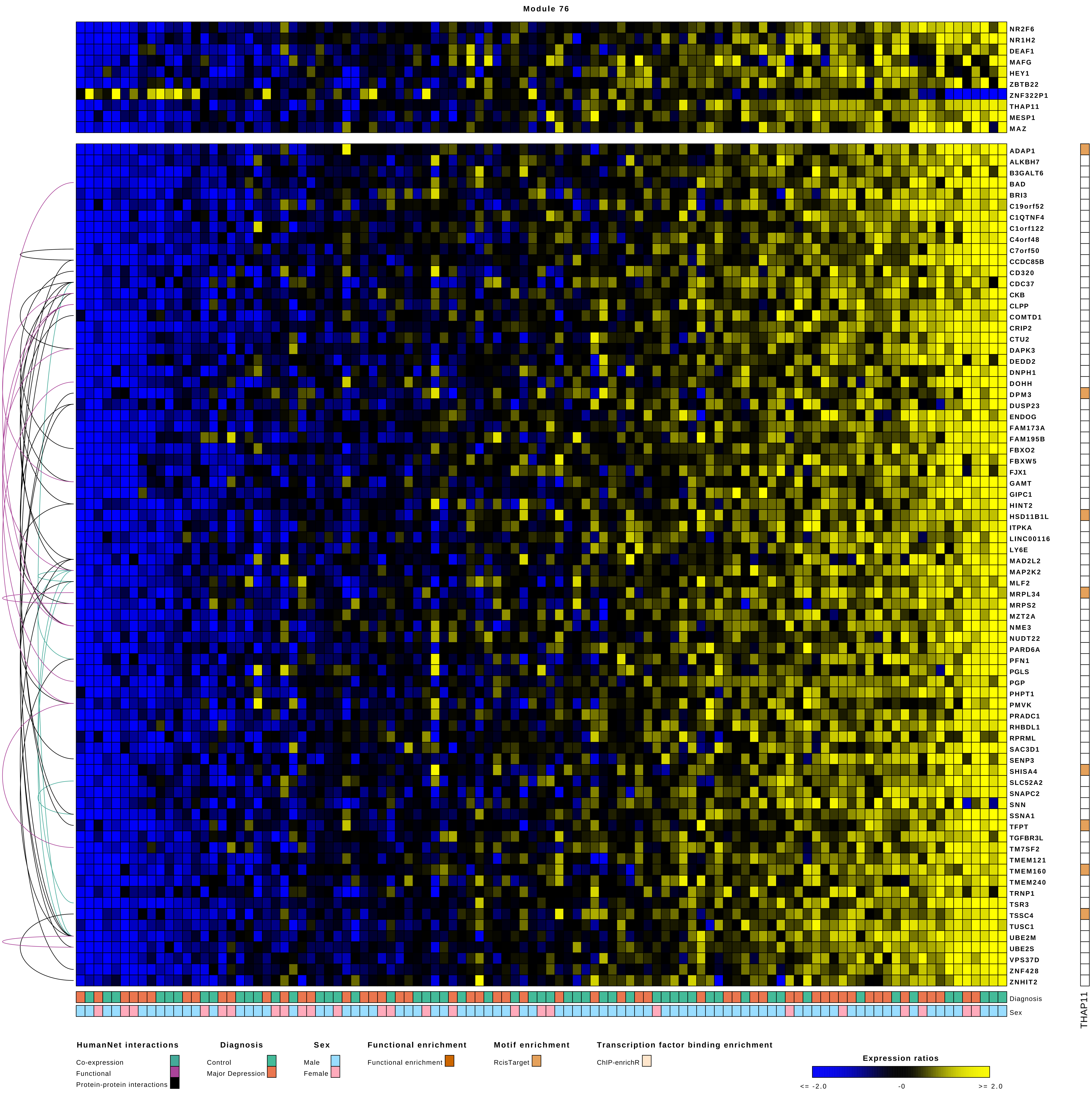
Module 76
NR2F6
NR1H2
DEAF1
MAFG
HEY1
ZBTB22
ZNF322P1
THAP11
MESP1
MAZ
ADAP1
ALKBH7
B3GALT6
BAD
BRI3
C19orf52
C1QTNF4
C1orf122
C4orf48
C7orf50
CCDC85B
CD320
CDC37
CKB
CLPP
COMTD1
CRIP2
CTU2
DAPK3
DEDD2
DNPH1
DOHH
DPM3
DUSP23
ENDOG
FAM173A
FAM195B
FBXO2
FBXW5
FJX1
GAMT
GIPC1
HINT2
HSD11B1L
ITPKA
LINC00116
LY6E
MAD2L2
MAP2K2
MLF2
MRPL34
MRPS2
MZT2A
NME3
NUDT22
PARD6A
PFN1
PGLS
PGP
PHPT1
PMVK
PRADC1
RHBDL1
RPRML
SAC3D1
SENP3
SHISA4
SLC52A2
SNAPC2
SNN
SSNA1
TFPT
TGFBR3L
TM7SF2
TMEM121
TMEM160
TMEM240
TRNP1
TSR3
TSSC4
TUSC1
UBE2M
UBE2S
VPS37D
ZNF428
ZNHIT2
Diagnosis
Sex
THAP11
HumanNet interactions
Co-expression
Functional
Protein-protein interactions
Diagnosis
Control
Major Depression
Sex
Male
Female
Functional enrichment
Functional enrichment
Motif enrichment
RcisTarget
Transcription factor binding enrichment
ChIP-enrichR
Expression ratios
<= -2.0
-0
>= 2.0

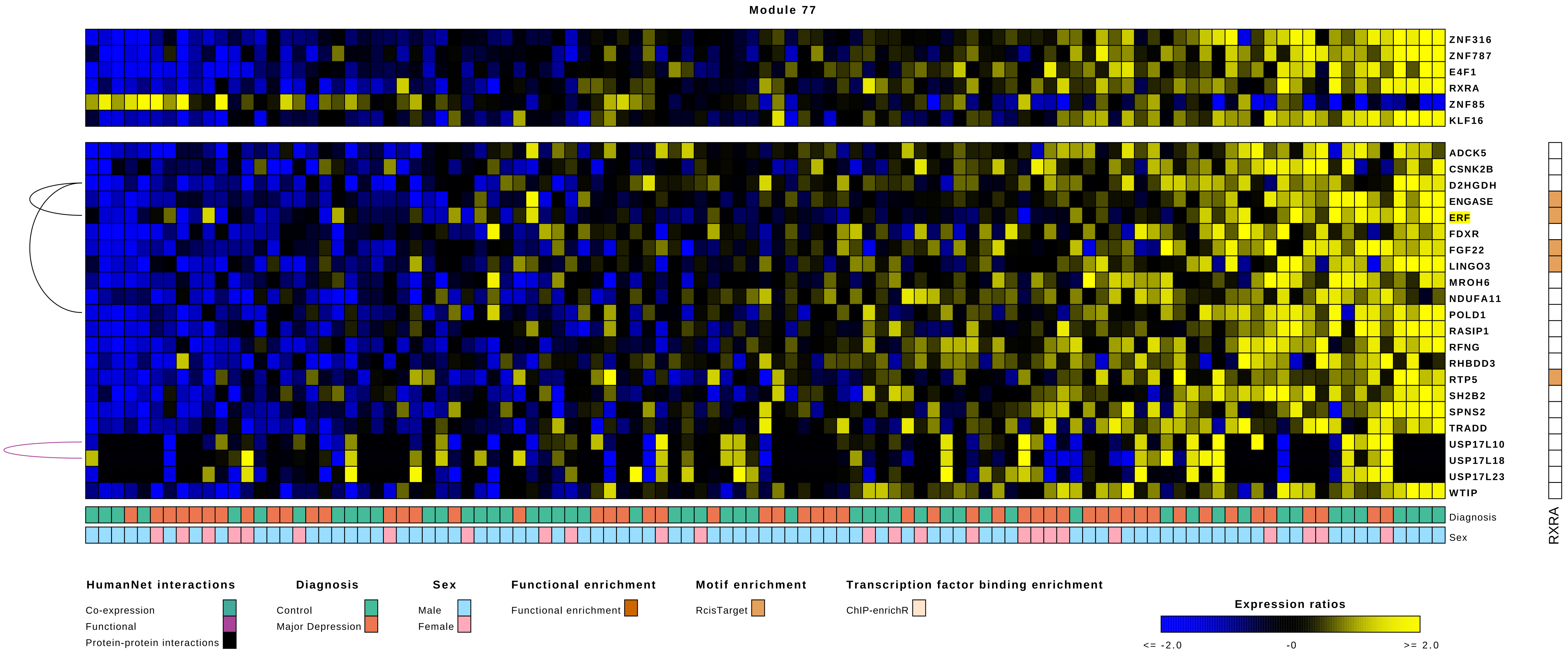

Module 77
ZNF316
ZNF787
E4F1
RXRA
ZNF85
KLF16
ADCK5
CSNK2B
D2HGDH
ENGASE
ERF
FDXR
FGF22
LINGO3
MROH6
NDUFA11
POLD1
RASIP1
RFNG
RHBDD3
RTP5
SH2B2
SPNS2
TRADD
USP17L10
USP17L18
USP17L23
WTIP
Diagnosis
Sex
RXRA
HumanNet interactions
Co-expression
Functional
Protein-protein interactions
Diagnosis
Control
Major Depression
Sex
Male
Female
Functional enrichment
Functional enrichment
Motif enrichment
RcisTarget
Transcription factor binding enrichment
ChIP-enrichR
Expression ratios
<= -2.0
-0
>= 2.0

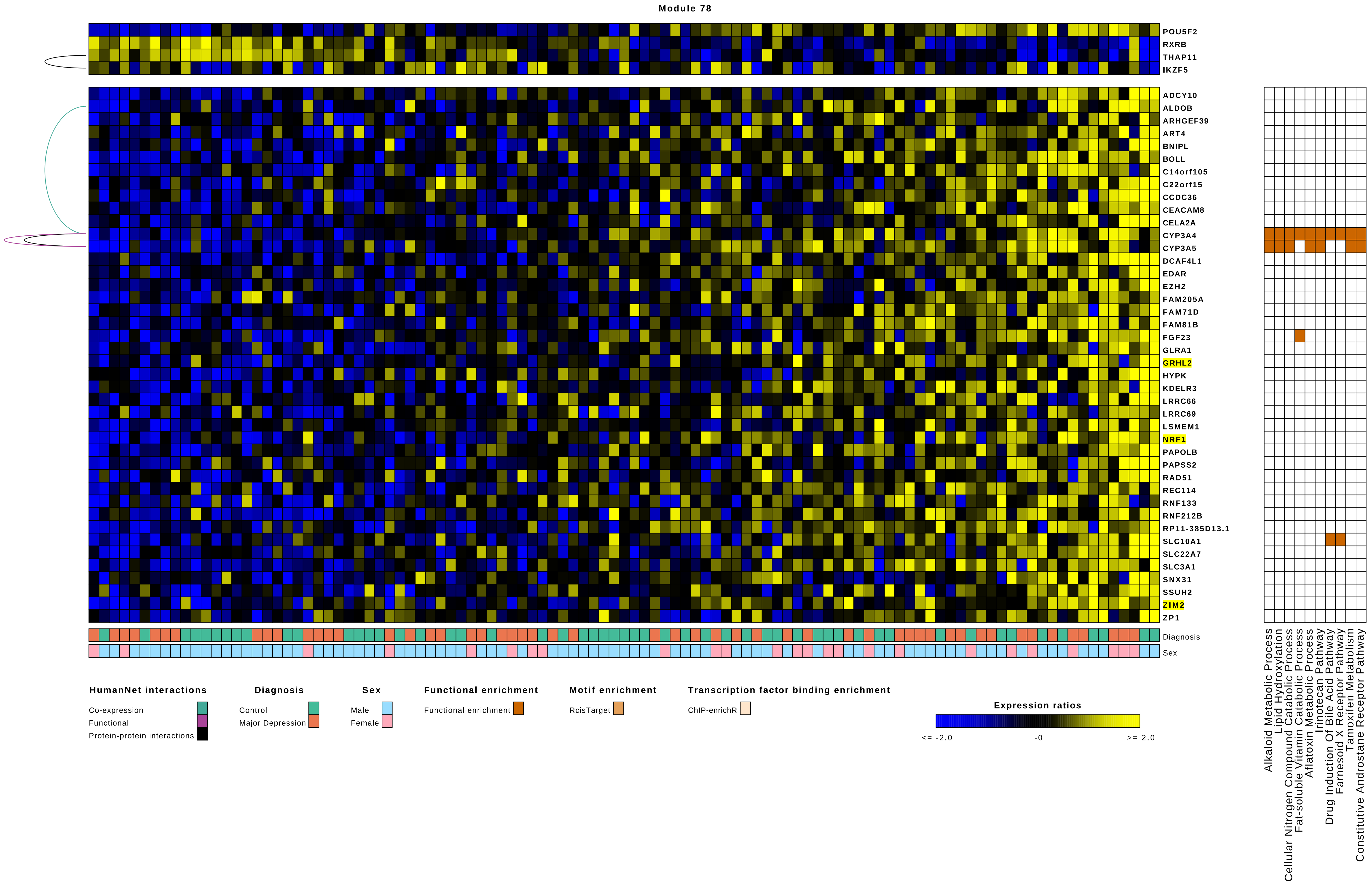

Module 78
POU5F2
RXRB
THAP11
IKZF5
ADCY10
ALDOB
ARHGEF39
ART4
BNIPL
BOLL
C14orf105
C22orf15
CCDC36
CEACAM8
CELA2A
CYP3A4
CYP3A5
DCAF4L1
EDAR
EZH2
FAM205A
FAM71D
FAM81B
FGF23
GLRA1
GRHL2
HYPK
KDELR3
LRRC66
LRRC69
LSMEM1
NRF1
PAPOLB
PAPSS2
RAD51
REC114
RNF133
RNF212B
RP11-385D13.1
SLC10A1
SLC22A7
SLC3A1
SNX31
SSUH2
ZIM2
ZP1
Diagnosis
Sex
Alkaloid Metabolic Process
Lipid Hydroxylation
Cellular Nitrogen Compound Catabolic Process
Fat-soluble Vitamin Catabolic Process
Aflatoxin Metabolic Process
Irinotecan Pathway
Drug Induction Of Bile Acid Pathway
Farnesoid X Receptor Pathway
Tamoxifen Metabolism
Constitutive Androstane Receptor Pathway
HumanNet interactions
Co-expression
Functional
Protein-protein interactions
Diagnosis
Control
Major Depression
Sex
Male
Female
Functional enrichment
Functional enrichment
Motif enrichment
RcisTarget
Transcription factor binding enrichment
ChIP-enrichR
Expression ratios
<= -2.0
-0
>= 2.0

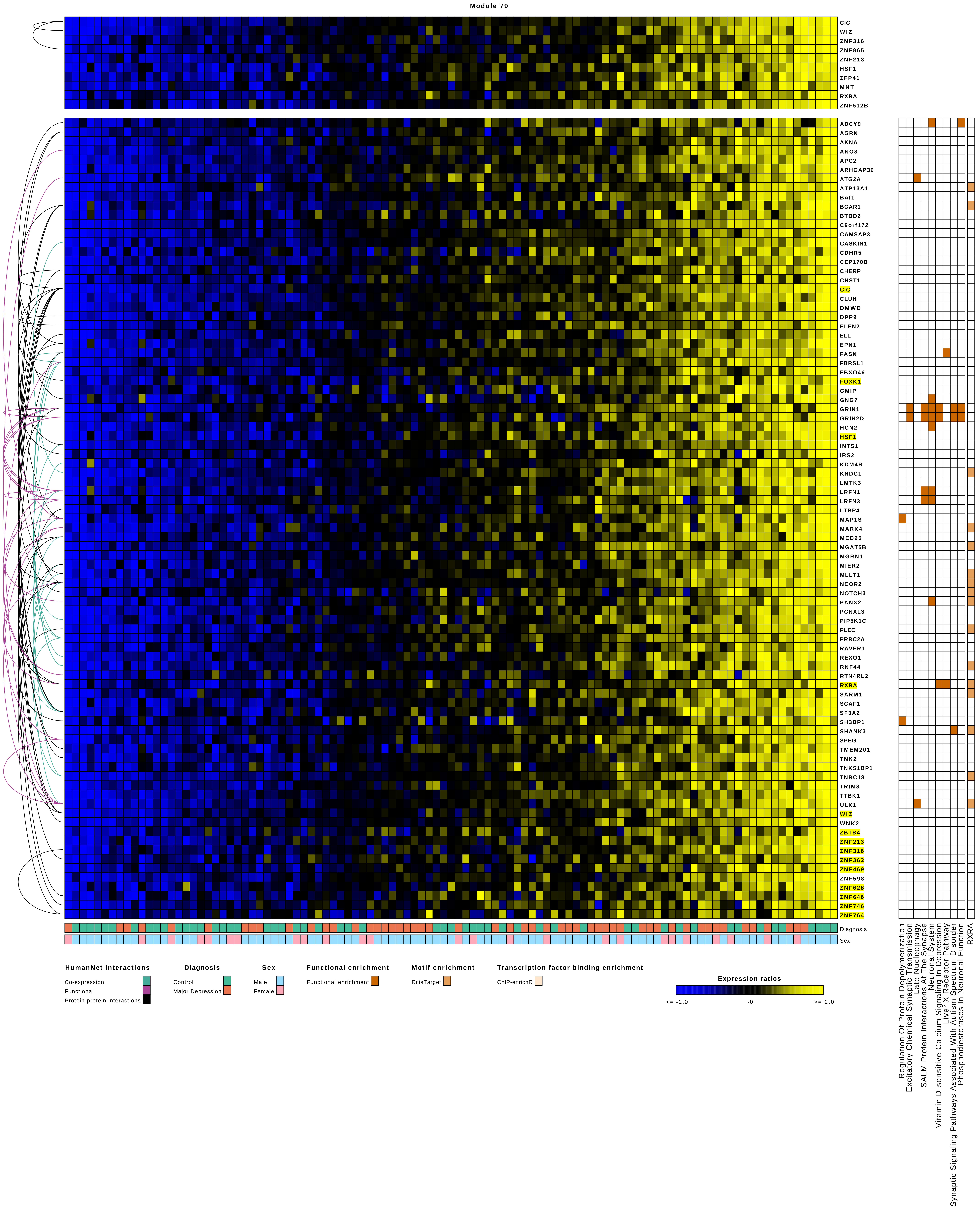
Module 79
CIC
WIZ
ZNF316
ZNF865
ZNF213
HSF1
ZFP41
MNT
RXRA
ZNF512B
ADCY9
AGRN
AKNA
ANO8
APC2
ARHGAP39
ATG2A
ATP13A1
BAI1
BCAR1
BTBD2
C9orf172
CAMSAP3
CASKIN1
CDHR5
CEP170B
CHERP
CHST1
CIC
CLUH
DMWD
DPP9
ELFN2
ELL
EPN1
FASN
FBRSL1
FBXO46
FOXK1
GMIP
GNG7
GRIN1
GRIN2D
HCN2
HSF1
INTS1
IRS2
KDM4B
KNDC1
LMTK3
LRFN1
LRFN3
LTBP4
MAP1S
MARK4
MED25
MGAT5B
MGRN1
MIER2
MLLT1
NCOR2
NOTCH3
PANX2
PCNXL3
PIP5K1C
PLEC
PRRC2A
RAVER1
REXO1
RNF44
RTN4RL2
RXRA
SARM1
SCAF1
SF3A2
SH3BP1
SHANK3
SPEG
TMEM201
TNK2
TNKS1BP1
TNRC18
TRIM8
TTBK1
ULK1
WIZ
WNK2
ZBTB4
ZNF213
ZNF316
ZNF362
ZNF469
ZNF598
ZNF628
ZNF646
ZNF746
ZNF764
Diagnosis
Sex
Regulation Of Protein Depolymerization
Excitatory Chemical Synaptic Transmission
Late Nucleophagy
SALM Protein Interactions At The Synapse
Neuronal System
Vitamin D-sensitive Calcium Signaling In Depression
Liver X Receptor Pathway
Synaptic Signaling Pathways Associated With Autism Spectrum Disorder
Phosphodiesterases In Neuronal Function
RXRA
HumanNet interactions
Co-expression
Functional
Protein-protein interactions
Diagnosis
Control
Major Depression
Sex
Male
Female
Functional enrichment
Functional enrichment
Motif enrichment
RcisTarget
Transcription factor binding enrichment
ChIP-enrichR
Expression ratios
<= -2.0
-0
>= 2.0

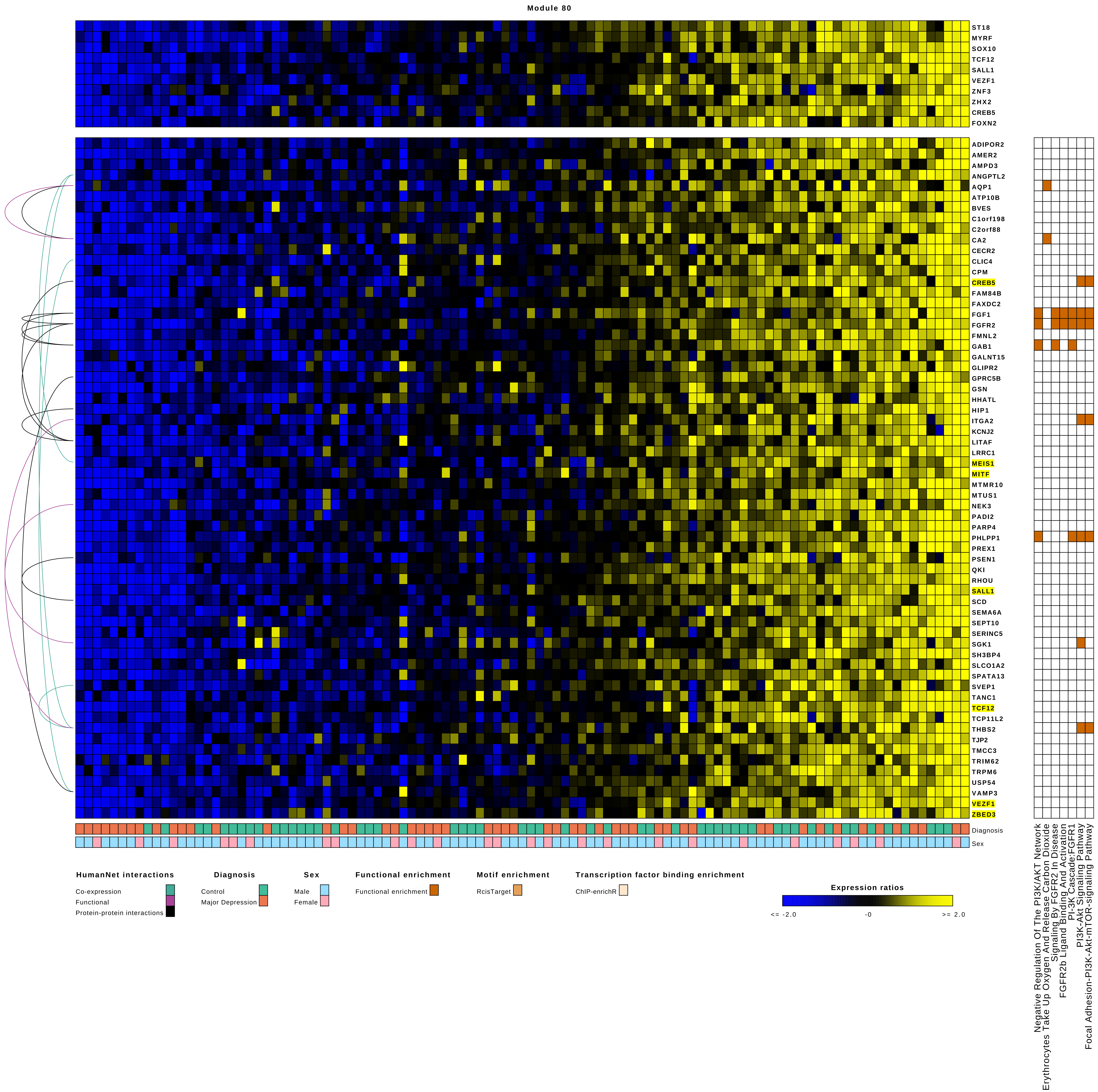

Module 80
ST18
MYRF
SOX10
TCF12
SALL1
VEZF1
ZNF3
ZHX2
CREB5
FOXN2
ADIPOR2
AMER2
AMPD3
ANGPTL2
AQP1
ATP10B
BVES
C1orf198
C2orf88
CA2
CECR2
CLIC4
CPM
CREB5
FAM84B
FAXDC2
FGF1
FGFR2
FMNL2
GAB1
GALNT15
GLIPR2
GPRC5B
GSN
HHATL
HIP1
ITGA2
KCNJ2
LITAF
LRRC1
MEIS1
MITF
MTMR10
MTUS1
NEK3
PADI2
PARP4
PHLPP1
PREX1
PSEN1
QKI
RHOU
SALL1
SCD
SEMA6A
SEPT10
SERINC5
SGK1
SH3BP4
SLCO1A2
SPATA13
SVEP1
TANC1
TCF12
TCP11L2
THBS2
TJP2
TMCC3
TRIM62
TRPM6
USP54
VAMP3
VEZF1
ZBED3
Diagnosis
Sex
Negative Regulation Of The PI3K/AKT Network
Erythrocytes Take Up Oxygen And Release Carbon Dioxide
Signaling By FGFR2 In Disease
FGFR2b Ligand Binding And Activation
PI-3K Cascade:FGFR1
PI3K-Akt Signaling Pathway
Focal Adhesion-PI3K-Akt-mTOR-signaling Pathway
HumanNet interactions
Co-expression
Functional
Protein-protein interactions
Diagnosis
Control
Major Depression
Sex
Male
Female
Functional enrichment
Functional enrichment
Motif enrichment
RcisTarget
Transcription factor binding enrichment
ChIP-enrichR
Expression ratios
<= -2.0
-0
>= 2.0

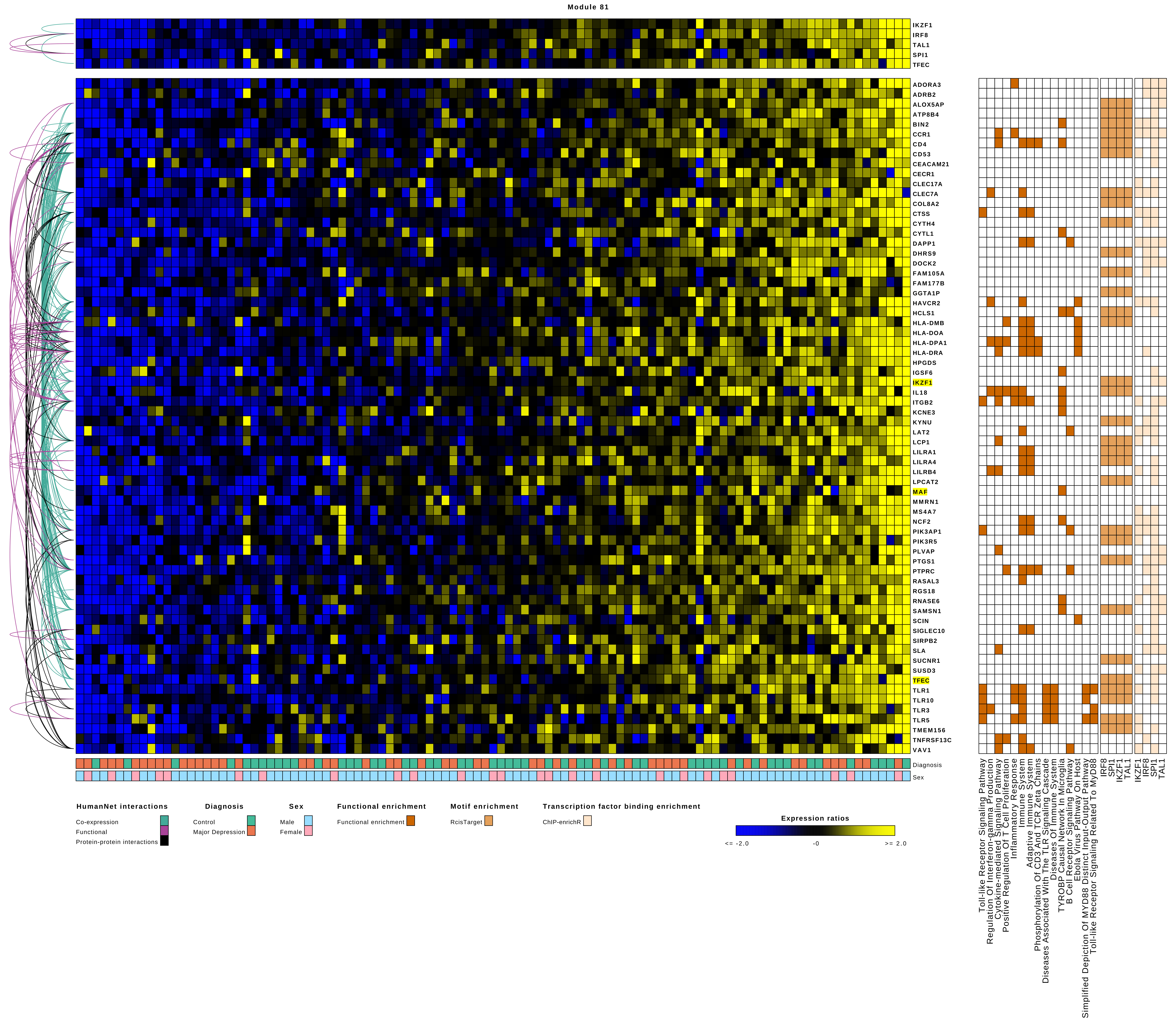
Module 81
IKZF1
IRF8
TAL1
SPI1
TFEC
ADORA3
ADRB2
ALOX5AP
ATP8B4
BIN2
CCR1
CD4
CD53
CEACAM21
CECR1
CLEC17A
CLEC7A
COL8A2
CTSS
CYTH4
CYTL1
DAPP1
DHRS9
DOCK2
FAM105A
FAM177B
GGTA1P
HAVCR2
HCLS1
HLA-DMB
HLA-DOA
HLA-DPA1
HLA-DRA
HPGDS
IGSF6
IKZF1
IL18
ITGB2
KCNE3
KYNU
LAT2
LCP1
LILRA1
LILRA4
LILRB4
LPCAT2
MAF
MMRN1
MS4A7
NCF2
PIK3AP1
PIK3R5
PLVAP
PTGS1
PTPRC
RASAL3
RGS18
RNASE6
SAMSN1
SCIN
SIGLEC10
SIRPB2
SLA
SUCNR1
SUSD3
TFEC
TLR1
TLR10
TLR3
TLR5
TMEM156
TNFRSF13C
VAV1
Diagnosis
Sex
Toll-like Receptor Signaling Pathway
Regulation Of Interferon-gamma Production
Cytokine-mediated Signaling Pathway
Positive Regulation Of T Cell Proliferation
Inflammatory Response
Immune System
Adaptive Immune System
Phosphorylation Of CD3 And TCR Zeta Chains
Diseases Associated With The TLR Signaling Cascade
Diseases Of Immune System
TYROBP Causal Network In Microglia
B Cell Receptor Signaling Pathway
Ebola Virus Pathway On Host
Simplified Depiction Of MYD88 Distinct Input-Output Pathway
Toll-like Receptor Signaling Related To MyD88
IRF8
SPI1
IKZF1
TAL1
IKZF1
IRF8
SPI1
TAL1
HumanNet interactions
Co-expression
Functional
Protein-protein interactions
Diagnosis
Control
Major Depression
Sex
Male
Female
Functional enrichment
Functional enrichment
Motif enrichment
RcisTarget
Transcription factor binding enrichment
ChIP-enrichR
Expression ratios
<= -2.0
-0
>= 2.0

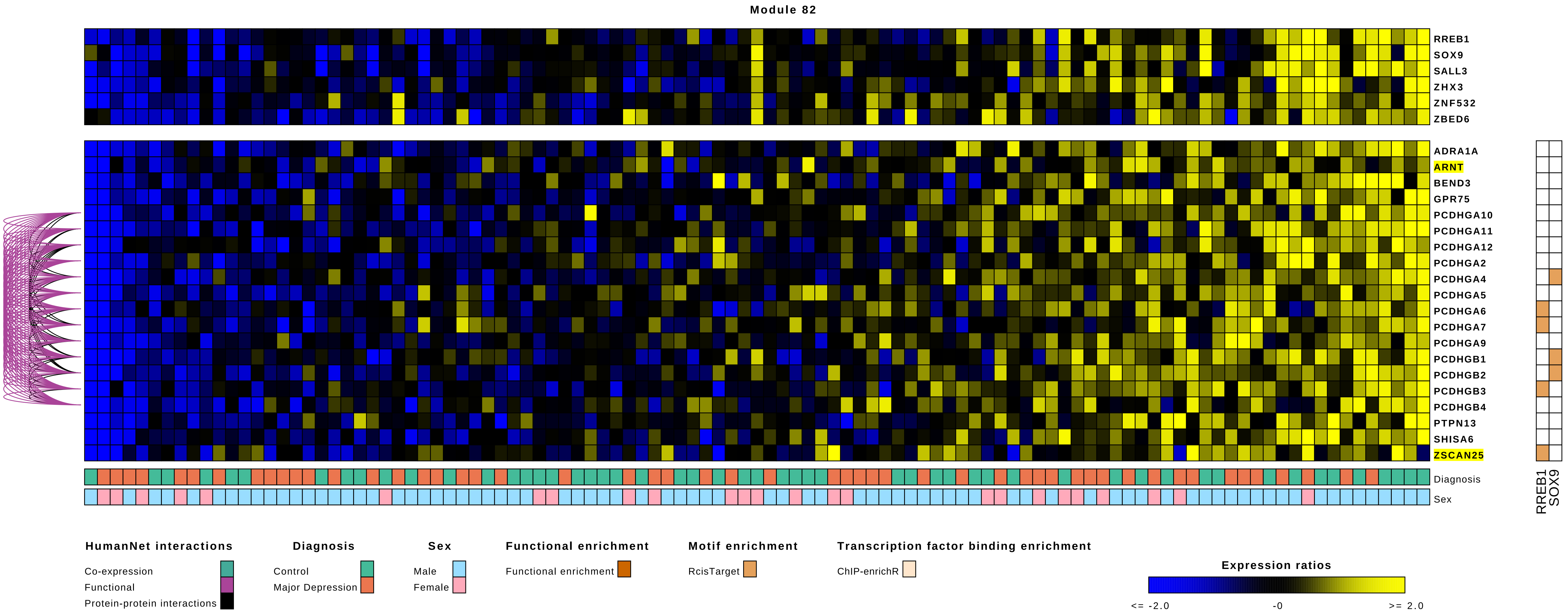

Module 82
RREB1
SOX9
SALL3
ZHX3
ZNF532
ZBED6
ADRA1A
ARNT
BEND3
GPR75
PCDHGA10
PCDHGA11
PCDHGA12
PCDHGA2
PCDHGA4
PCDHGA5
PCDHGA6
PCDHGA7
PCDHGA9
PCDHGB1
PCDHGB2
PCDHGB3
PCDHGB4
PTPN13
SHISA6
ZSCAN25
Diagnosis
Sex
RREB1
SOX9
HumanNet interactions
Co-expression
Functional
Protein-protein interactions
Diagnosis
Control
Major Depression
Sex
Male
Female
Functional enrichment
Functional enrichment
Motif enrichment
RcisTarget
Transcription factor binding enrichment
ChIP-enrichR
Expression ratios
<= -2.0
-0
>= 2.0

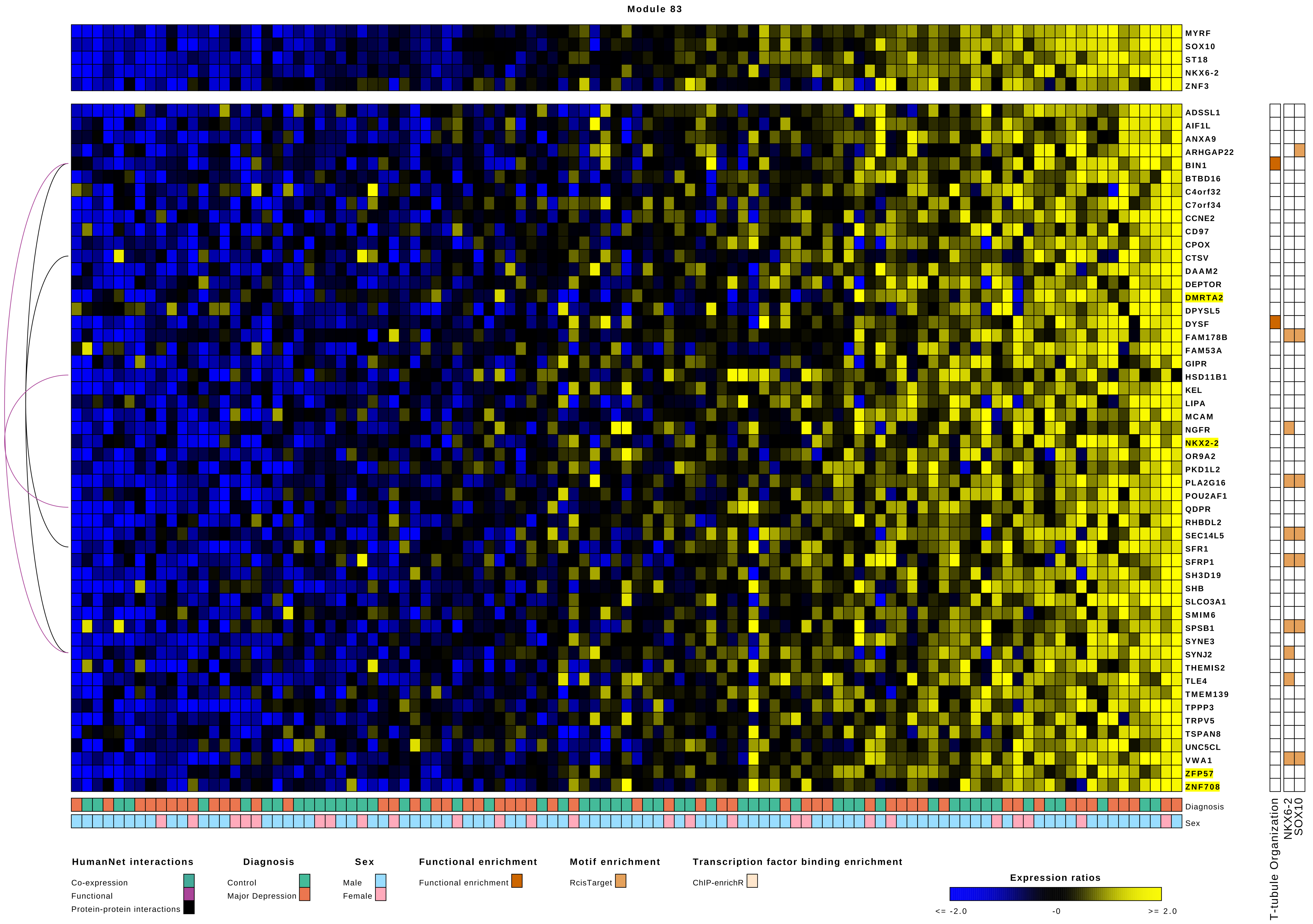

Module 83
MYRF
SOX10
ST18
NKX6-2
ZNF3
ADSSL1
AIF1L
ANXA9
ARHGAP22
BIN1
BTBD16
C4orf32
C7orf34
CCNE2
CD97
CPOX
CTSV
DAAM2
DEPTOR
DMRTA2
DPYSL5
DYSF
FAM178B
FAM53A
GIPR
HSD11B1
KEL
LIPA
MCAM
NGFR
NKX2-2
OR9A2
PKD1L2
PLA2G16
POU2AF1
QDPR
RHBDL2
SEC14L5
SFR1
SFRP1
SH3D19
SHB
SLCO3A1
SMIM6
SPSB1
SYNE3
SYNJ2
THEMIS2
TLE4
TMEM139
TPPP3
TRPV5
TSPAN8
UNC5CL
VWA1
ZFP57
ZNF708
Diagnosis
Sex
T-tubule Organization
NKX6-2
SOX10
HumanNet interactions
Co-expression
Functional
Protein-protein interactions
Diagnosis
Control
Major Depression
Sex
Male
Female
Functional enrichment
Functional enrichment
Motif enrichment
RcisTarget
Transcription factor binding enrichment
ChIP-enrichR
Expression ratios
<= -2.0
-0
>= 2.0

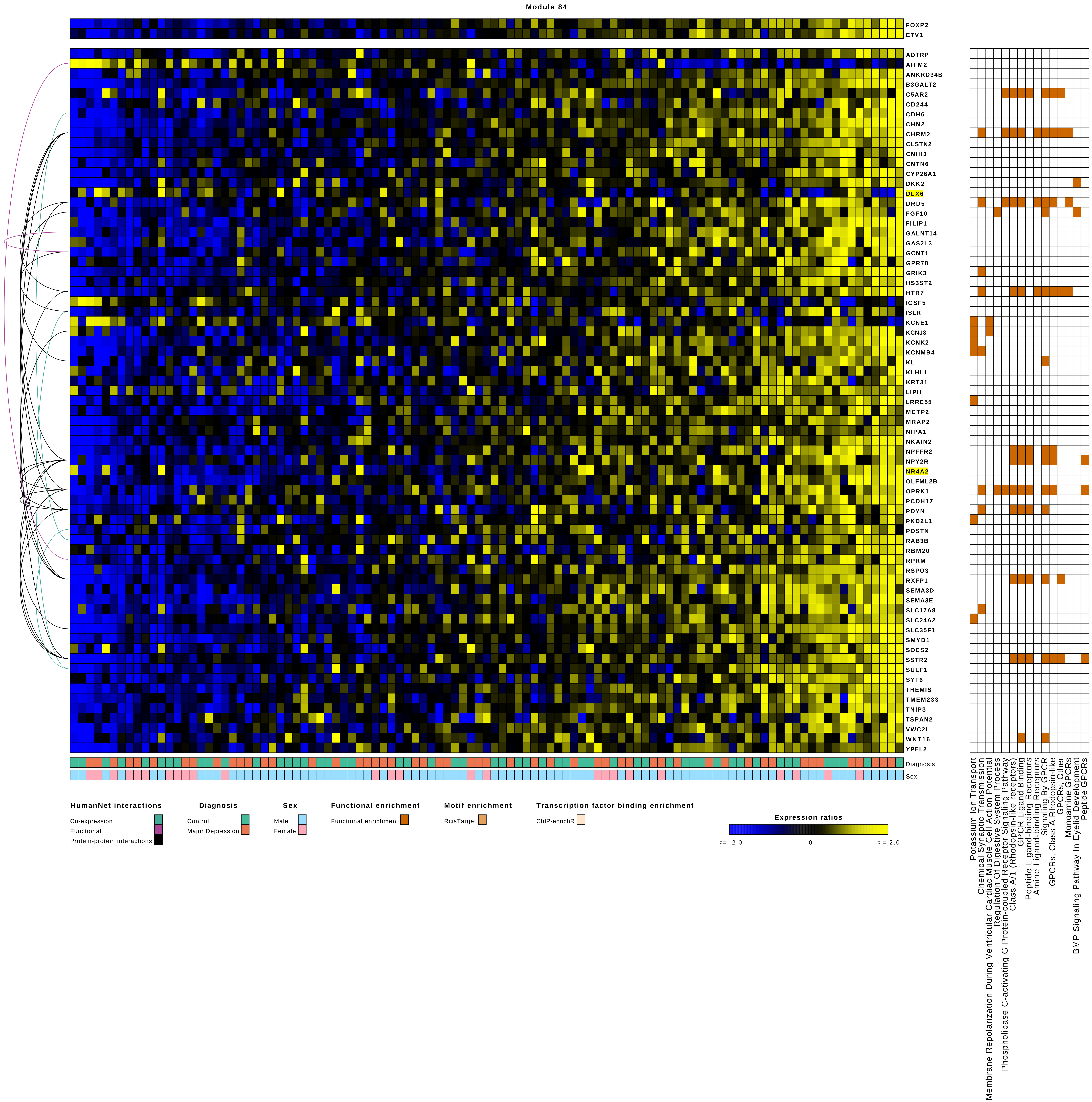

Module 84
FOXP2
ETV1
ADTRP
AIFM2
ANKRD34B
B3GALT2
C5AR2
CD244
CDH6
CHN2
CHRM2
CLSTN2
CNIH3
CNTN6
CYP26A1
DKK2
DLX6
DRD5
FGF10
FILIP1
GALNT14
GAS2L3
GCNT1
GPR78
GRIK3
HS3ST2
HTR7
IGSF5
ISLR
KCNE1
KCNJ8
KCNK2
KCNMB4
KL
KLHL1
KRT31
LIPH
LRRC55
MCTP2
MRAP2
NIPA1
NKAIN2
NPFFR2
NPY2R
NR4A2
OLFML2B
OPRK1
PCDH17
PDYN
PKD2L1
POSTN
RAB3B
RBM20
RPRM
RSPO3
RXFP1
SEMA3D
SEMA3E
SLC17A8
SLC24A2
SLC35F1
SMYD1
SOCS2
SSTR2
SULF1
SYT6
THEMIS
TMEM233
TNIP3
TSPAN2
VWC2L
WNT16
YPEL2
Diagnosis
Sex
Potassium Ion Transport
Chemical Synaptic Transmission
Membrane Repolarization During Ventricular Cardiac Muscle Cell Action Potential
Regulation Of Digestive System Process
Phospholipase C-activating G Protein-coupled Receptor Signaling Pathway
Class A/1 (Rhodopsin-like receptors)
GPCR Ligand Binding
Peptide Ligand-binding Receptors
Amine Ligand-binding Receptors
Signaling By GPCR
GPCRs, Class A Rhodopsin-like
GPCRs, Other
Monoamine GPCRs
BMP Signaling Pathway In Eyelid Development
Peptide GPCRs
HumanNet interactions
Co-expression
Functional
Protein-protein interactions
Diagnosis
Control
Major Depression
Sex
Male
Female
Functional enrichment
Functional enrichment
Motif enrichment
RcisTarget
Transcription factor binding enrichment
ChIP-enrichR
Expression ratios
<= -2.0
-0
>= 2.0

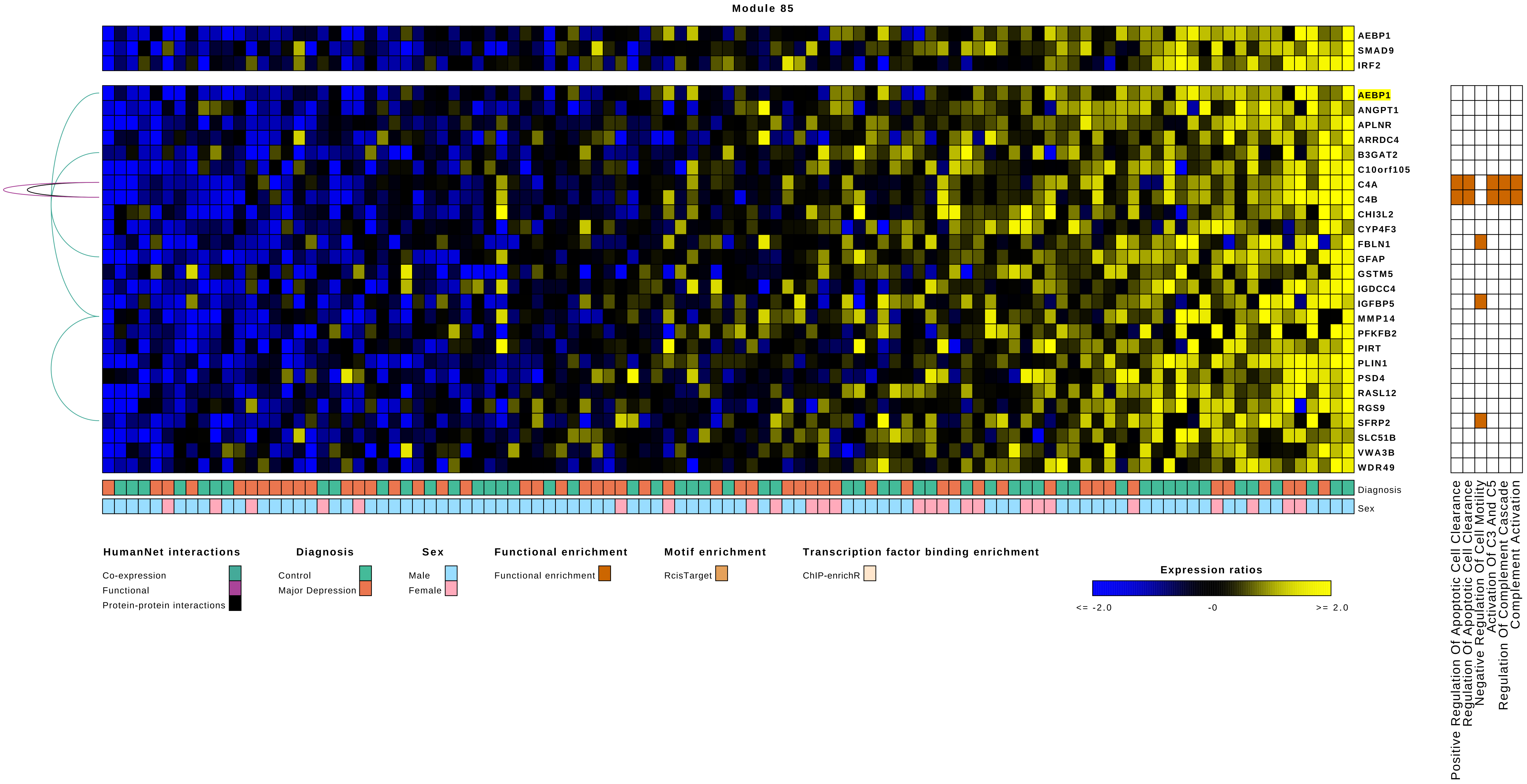
Module 85
AEBP1
SMAD9
IRF2
AEBP1
ANGPT1
APLNR
ARRDC4
B3GAT2
C10orf105
C4A
C4B
CHI3L2
CYP4F3
FBLN1
GFAP
GSTM5
IGDCC4
IGFBP5
MMP14
PFKFB2
PIRT
PLIN1
PSD4
RASL12
RGS9
SFRP2
SLC51B
VWA3B
WDR49
Diagnosis
Sex
Positive Regulation Of Apoptotic Cell Clearance
Regulation Of Apoptotic Cell Clearance
Negative Regulation Of Cell Motility
Activation Of C3 And C5
Regulation Of Complement Cascade
Complement Activation
HumanNet interactions
Co-expression
Functional
Protein-protein interactions
Diagnosis
Control
Major Depression
Sex
Male
Female
Functional enrichment
Functional enrichment
Motif enrichment
RcisTarget
Transcription factor binding enrichment
ChIP-enrichR
Expression ratios
<= -2.0
-0
>= 2.0

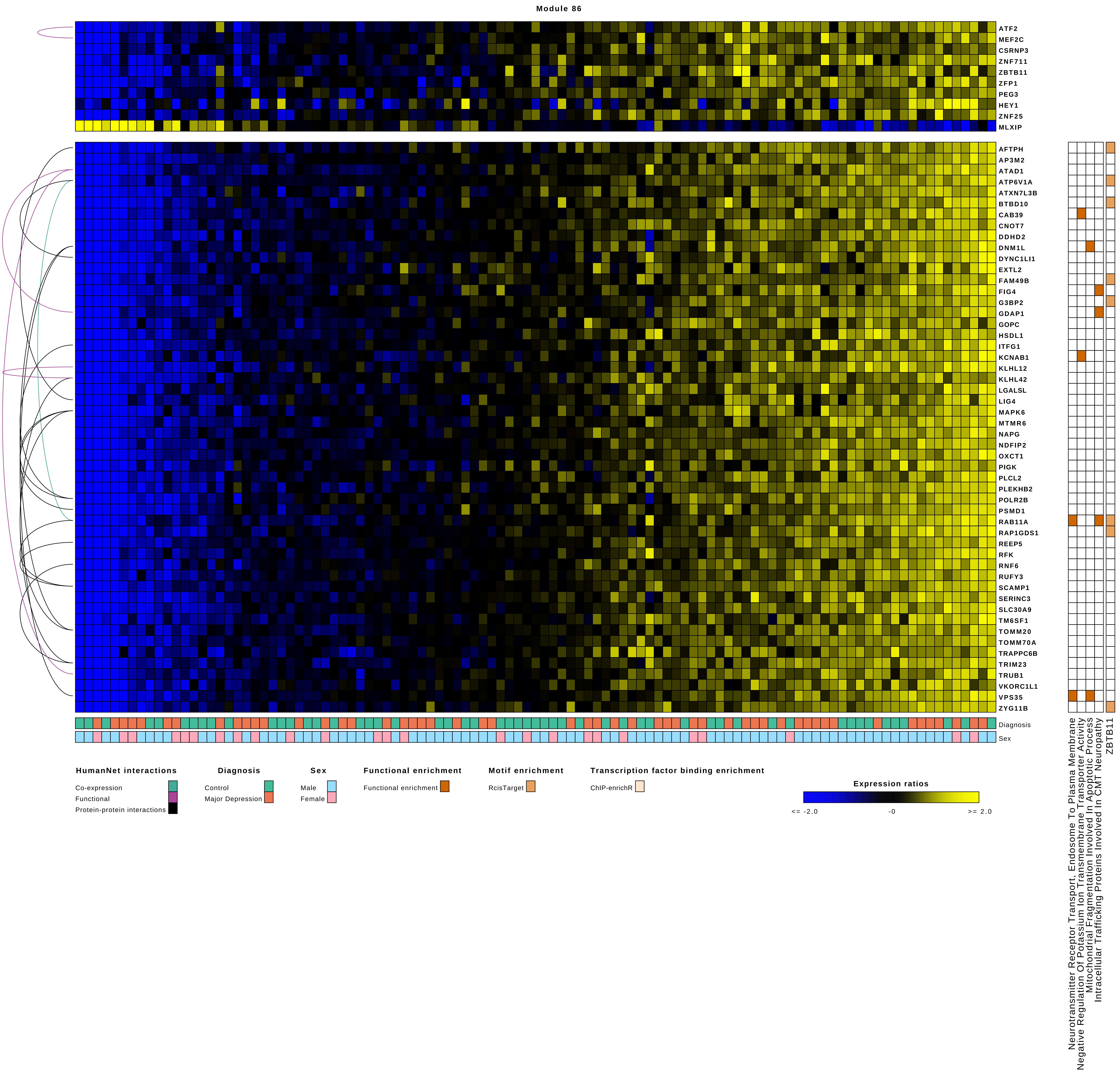
Module 86
ATF2
MEF2C
CSRNP3
ZNF711
ZBTB11
ZFP1
PEG3
HEY1
ZNF25
MLXIP
AFTPH
AP3M2
ATAD1
ATP6V1A
ATXN7L3B
BTBD10
CAB39
CNOT7
DDHD2
DNM1L
DYNC1LI1
EXTL2
FAM49B
FIG4
G3BP2
GDAP1
GOPC
HSDL1
ITFG1
KCNAB1
KLHL12
KLHL42
LGALSL
LIG4
MAPK6
MTMR6
NAPG
NDFIP2
OXCT1
PIGK
PLCL2
PLEKHB2
POLR2B
PSMD1
RAB11A
RAP1GDS1
REEP5
RFK
RNF6
RUFY3
SCAMP1
SERINC3
SLC30A9
TM6SF1
TOMM20
TOMM70A
TRAPPC6B
TRIM23
TRUB1
VKORC1L1
VPS35
ZYG11B
Diagnosis
Sex
Neurotransmitter Receptor Transport, Endosome To Plasma Membrane
Negative Regulation Of Potassium Ion Transmembrane Transporter Activity
Mitochondrial Fragmentation Involved In Apoptotic Process
Intracellular Trafficking Proteins Involved In CMT Neuropathy
ZBTB11
HumanNet interactions
Co-expression
Functional
Protein-protein interactions
Diagnosis
Control
Major Depression
Sex
Male
Female
Functional enrichment
Functional enrichment
Motif enrichment
RcisTarget
Transcription factor binding enrichment
ChIP-enrichR
Expression ratios
<= -2.0
-0
>= 2.0

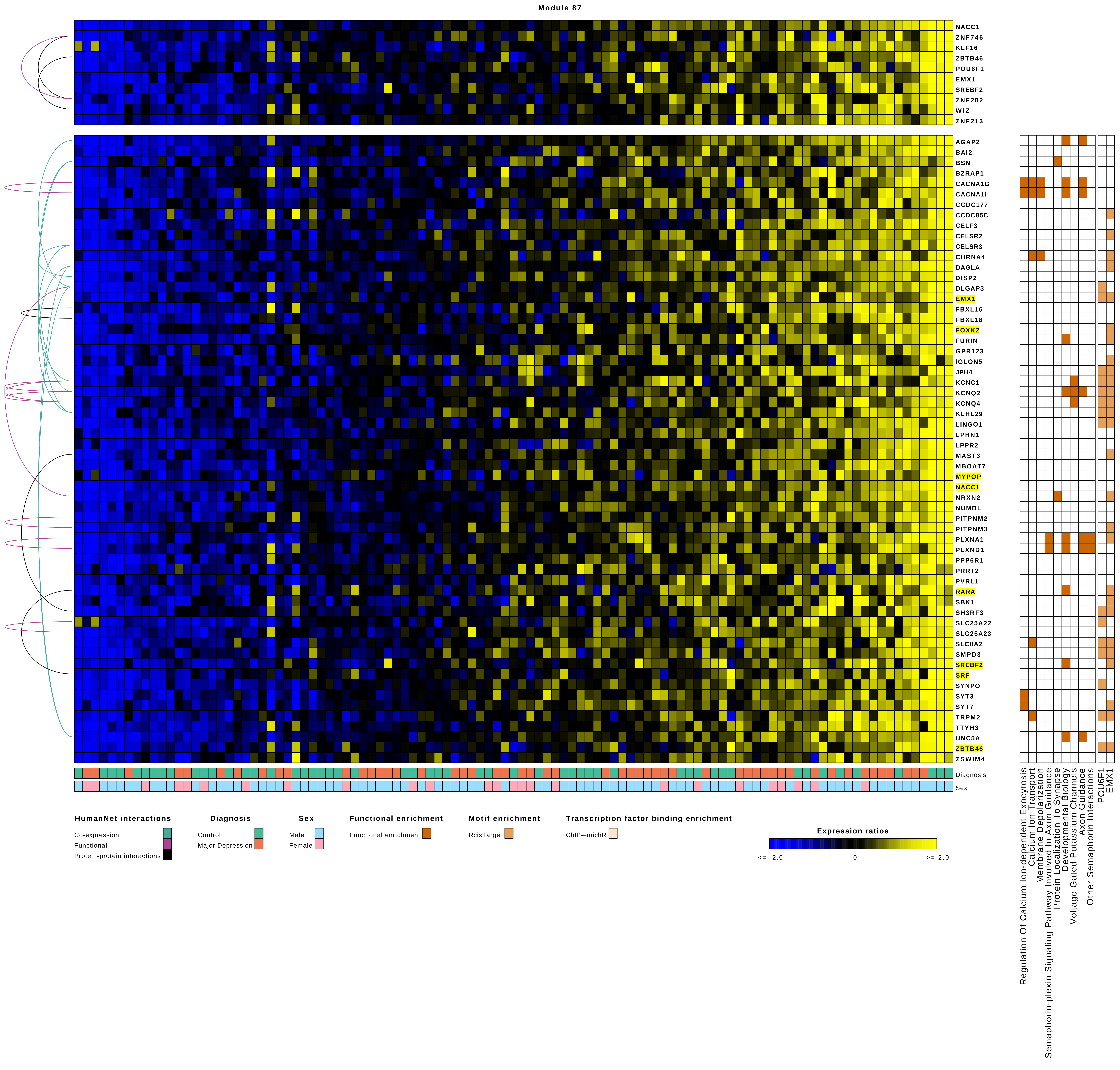

Module 87
NACC1
ZNF746
KLF16
ZBTB46
POU6F1
EMX1
SREBF2
ZNF282
WIZ
ZNF213
AGAP2
BAI2
BSN
BZRAP1
CACNA1G
CACNA1I
CCDC177
CCDC85C
CELF3
CELSR2
CELSR3
CHRNA4
DAGLA
DISP2
DLGAP3
EMX1
FBXL16
FBXL18
FOXK2
FURIN
GPR123
IGLON5
JPH4
KCNC1
KCNQ2
KCNQ4
KLHL29
LINGO1
LPHN1
LPPR2
MAST3
MBOAT7
MYPOP
NACC1
NRXN2
NUMBL
PITPNM2
PITPNM3
PLXNA1
PLXND1
PPP6R1
PRRT2
PVRL1
RARA
SBK1
SH3RF3
SLC25A22
SLC25A23
SLC8A2
SMPD3
SREBF2
SRF
SYNPO
SYT3
SYT7
TRPM2
TTYH3
UNC5A
ZBTB46
ZSWIM4
Diagnosis
Sex
Regulation Of Calcium Ion-dependent Exocytosis
Calcium Ion Transport
Membrane Depolarization
Semaphorin-plexin Signaling Pathway Involved In Axon Guidance
Protein Localization To Synapse
Developmental Biology
Voltage Gated Potassium Channels
Axon Guidance
Other Semaphorin Interactions
POU6F1
EMX1
HumanNet interactions
Co-expression
Functional
Protein-protein interactions
Diagnosis
Control
Major Depression
Sex
Male
Female
Functional enrichment
Functional enrichment
Motif enrichment
RcisTarget
Transcription factor binding enrichment
ChIP-enrichR
Expression ratios
<= -2.0
-0
>= 2.0

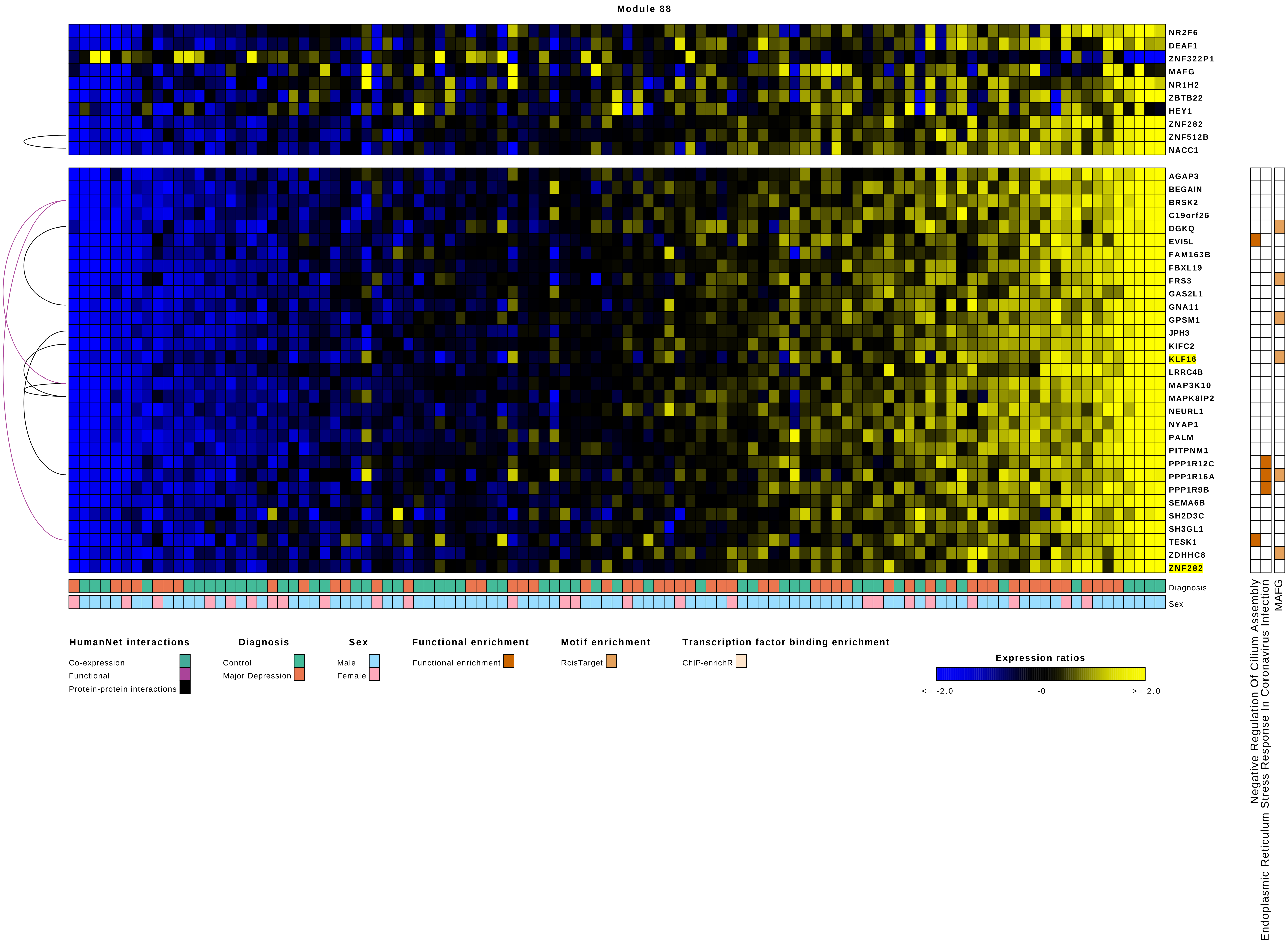
Module 88
NR2F6
DEAF1
ZNF322P1
MAFG
NR1H2
ZBTB22
HEY1
ZNF282
ZNF512B
NACC1
AGAP3
BEGAIN
BRSK2
C19orf26
DGKQ
EVI5L
FAM163B
FBXL19
FRS3
GAS2L1
GNA11
GPSM1
JPH3
KIFC2
KLF16
LRRC4B
MAP3K10
MAPK8IP2
NEURL1
NYAP1
PALM
PITPNM1
PPP1R12C
PPP1R16A
PPP1R9B
SEMA6B
SH2D3C
SH3GL1
TESK1
ZDHHC8
ZNF282
Diagnosis
Sex
Negative Regulation Of Cilium Assembly
Endoplasmic Reticulum Stress Response In Coronavirus Infection
MAFG
HumanNet interactions
Co-expression
Functional
Protein-protein interactions
Diagnosis
Control
Major Depression
Sex
Male
Female
Functional enrichment
Functional enrichment
Motif enrichment
RcisTarget
Transcription factor binding enrichment
ChIP-enrichR
Expression ratios
<= -2.0
-0
>= 2.0

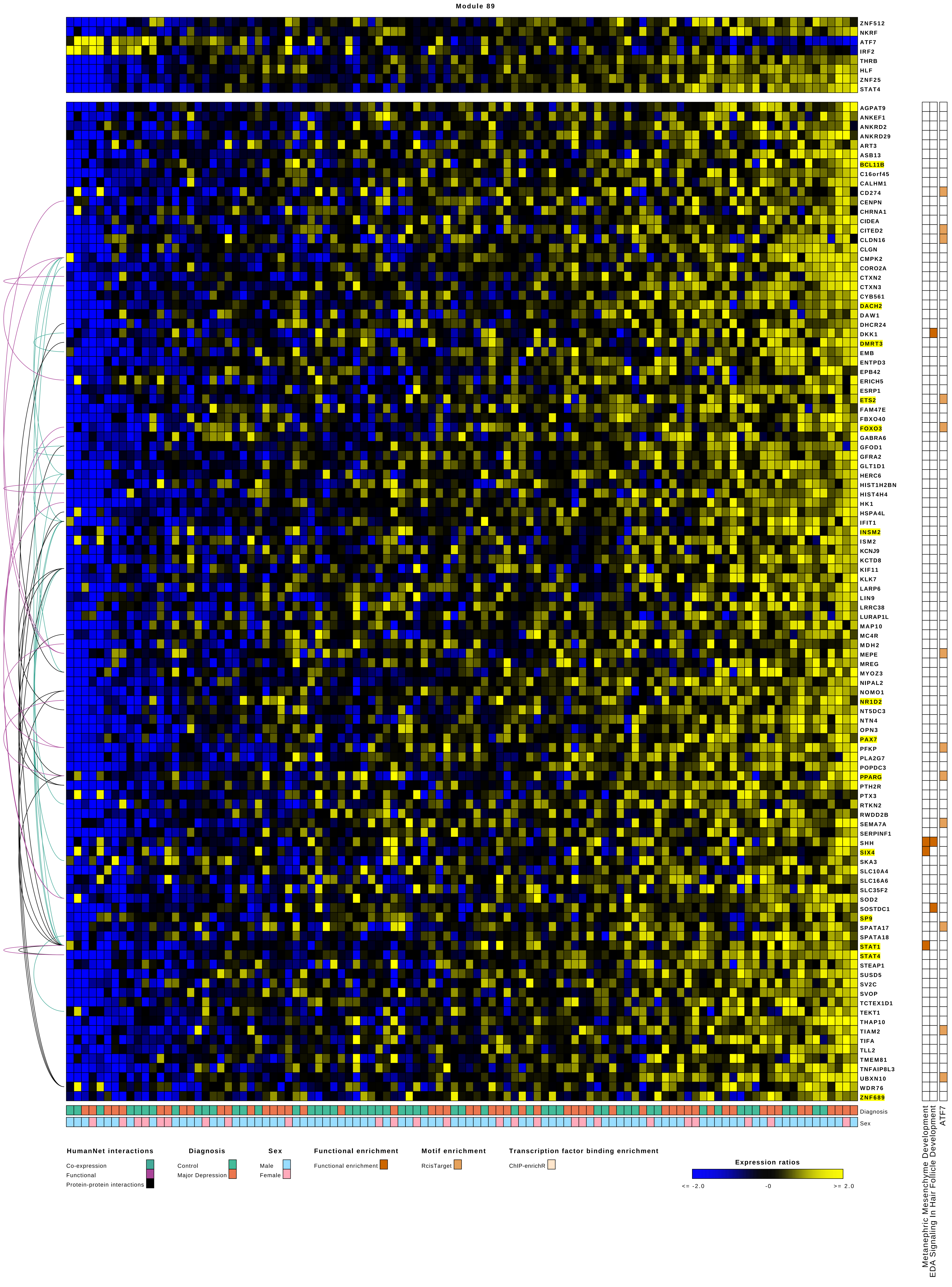
Module 89
ZNF512
NKRF
ATF7
IRF2
THRB
HLF
ZNF25
STAT4
AGPAT9
ANKEF1
ANKRD2
ANKRD29
ART3
ASB13
BCL11B
C16orf45
CALHM1
CD274
CENPN
CHRNA1
CIDEA
CITED2
CLDN16
CLGN
CMPK2
CORO2A
CTXN2
CTXN3
CYB561
DACH2
DAW1
DHCR24
DKK1
DMRT3
EMB
ENTPD3
EPB42
ERICH5
ESRP1
ETS2
FAM47E
FBXO40
FOXO3
GABRA6
GFOD1
GFRA2
GLT1D1
HERC6
HIST1H2BN
HIST4H4
HK1
HSPA4L
IFIT1
INSM2
ISM2
KCNJ9
KCTD8
KIF11
KLK7
LARP6
LIN9
LRRC38
LURAP1L
MAP10
MC4R
MDH2
MEPE
MREG
MYOZ3
NIPAL2
NOMO1
NR1D2
NT5DC3
NTN4
OPN3
PAX7
PFKP
PLA2G7
POPDC3
PPARG
PTH2R
PTX3
RTKN2
RWDD2B
SEMA7A
SERPINF1
SHH
SIX4
SKA3
SLC10A4
SLC16A6
SLC35F2
SOD2
SOSTDC1
SP9
SPATA17
SPATA18
STAT1
STAT4
STEAP1
SUSD5
SV2C
SVOP
TCTEX1D1
TEKT1
THAP10
TIAM2
TIFA
TLL2
TMEM81
TNFAIP8L3
UBXN10
WDR76
ZNF689
Diagnosis
Sex
Metanephric Mesenchyme Development
EDA Signaling In Hair Follicle Development
ATF7
HumanNet interactions
Co-expression
Functional
Protein-protein interactions
Diagnosis
Control
Major Depression
Sex
Male
Female
Functional enrichment
Functional enrichment
Motif enrichment
RcisTarget
Transcription factor binding enrichment
ChIP-enrichR
Expression ratios
<= -2.0
-0
>= 2.0

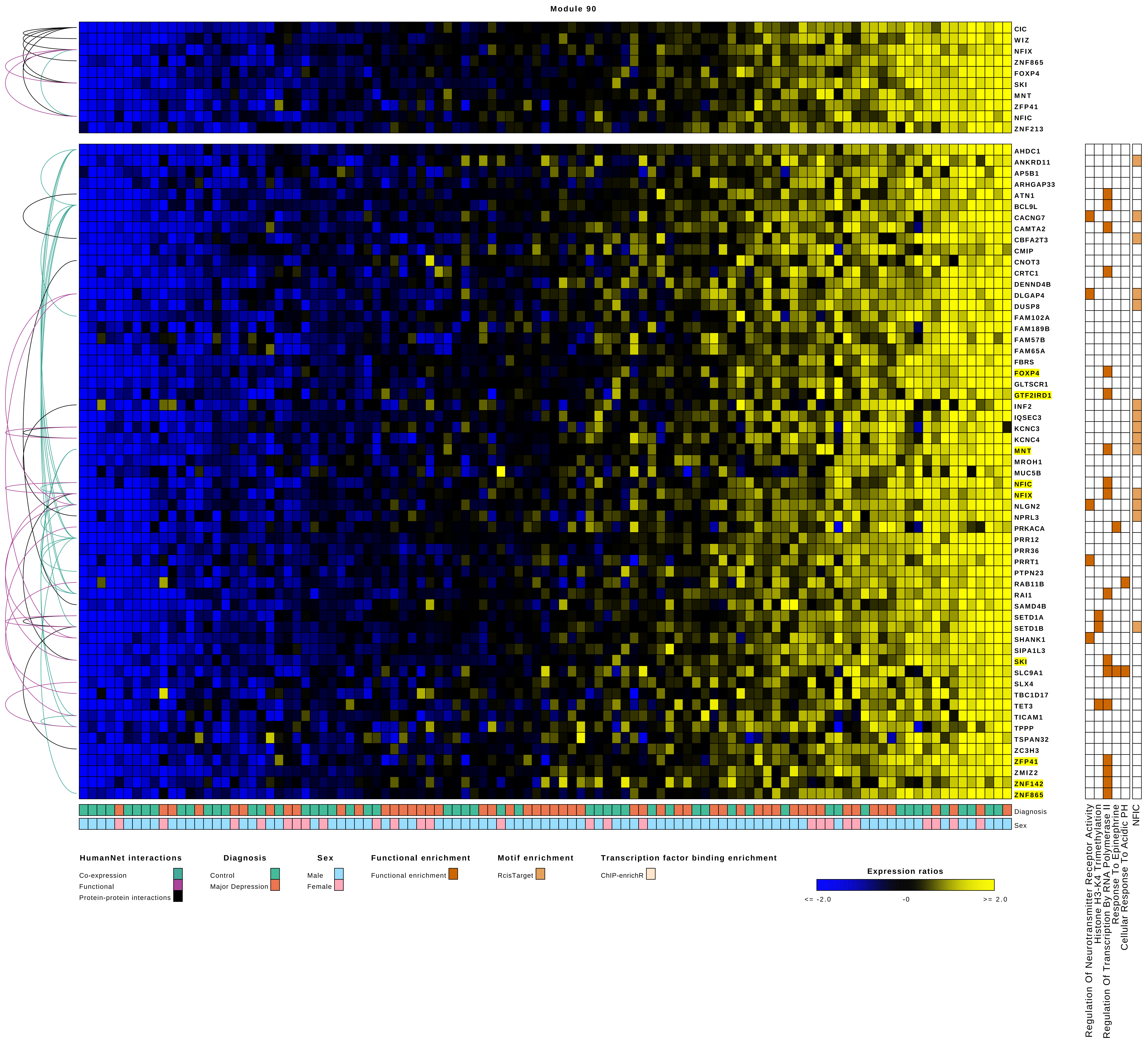

Module 90
CIC
WIZ
NFIX
ZNF865
FOXP4
SKI
MNT
ZFP41
NFIC
ZNF213
AHDC1
ANKRD11
AP5B1
ARHGAP33
ATN1
BCL9L
CACNG7
CAMTA2
CBFA2T3
CMIP
CNOT3
CRTC1
DENND4B
DLGAP4
DUSP8
FAM102A
FAM189B
FAM57B
FAM65A
FBRS
FOXP4
GLTSCR1
GTF2IRD1
INF2
IQSEC3
KCNC3
KCNC4
MNT
MROH1
MUC5B
NFIC
NFIX
NLGN2
NPRL3
PRKACA
PRR12
PRR36
PRRT1
PTPN23
RAB11B
RAI1
SAMD4B
SETD1A
SETD1B
SHANK1
SIPA1L3
SKI
SLC9A1
SLX4
TBC1D17
TET3
TICAM1
TPPP
TSPAN32
ZC3H3
ZFP41
ZMIZ2
ZNF142
ZNF865
Diagnosis
Sex
Regulation Of Neurotransmitter Receptor Activity
Histone H3-K4 Trimethylation
Regulation Of Transcription By RNA Polymerase II
Response To Epinephrine
Cellular Response To Acidic PH
NFIC
HumanNet interactions
Co-expression
Functional
Protein-protein interactions
Diagnosis
Control
Major Depression
Sex
Male
Female
Functional enrichment
Functional enrichment
Motif enrichment
RcisTarget
Transcription factor binding enrichment
ChIP-enrichR
Expression ratios
<= -2.0
-0
>= 2.0

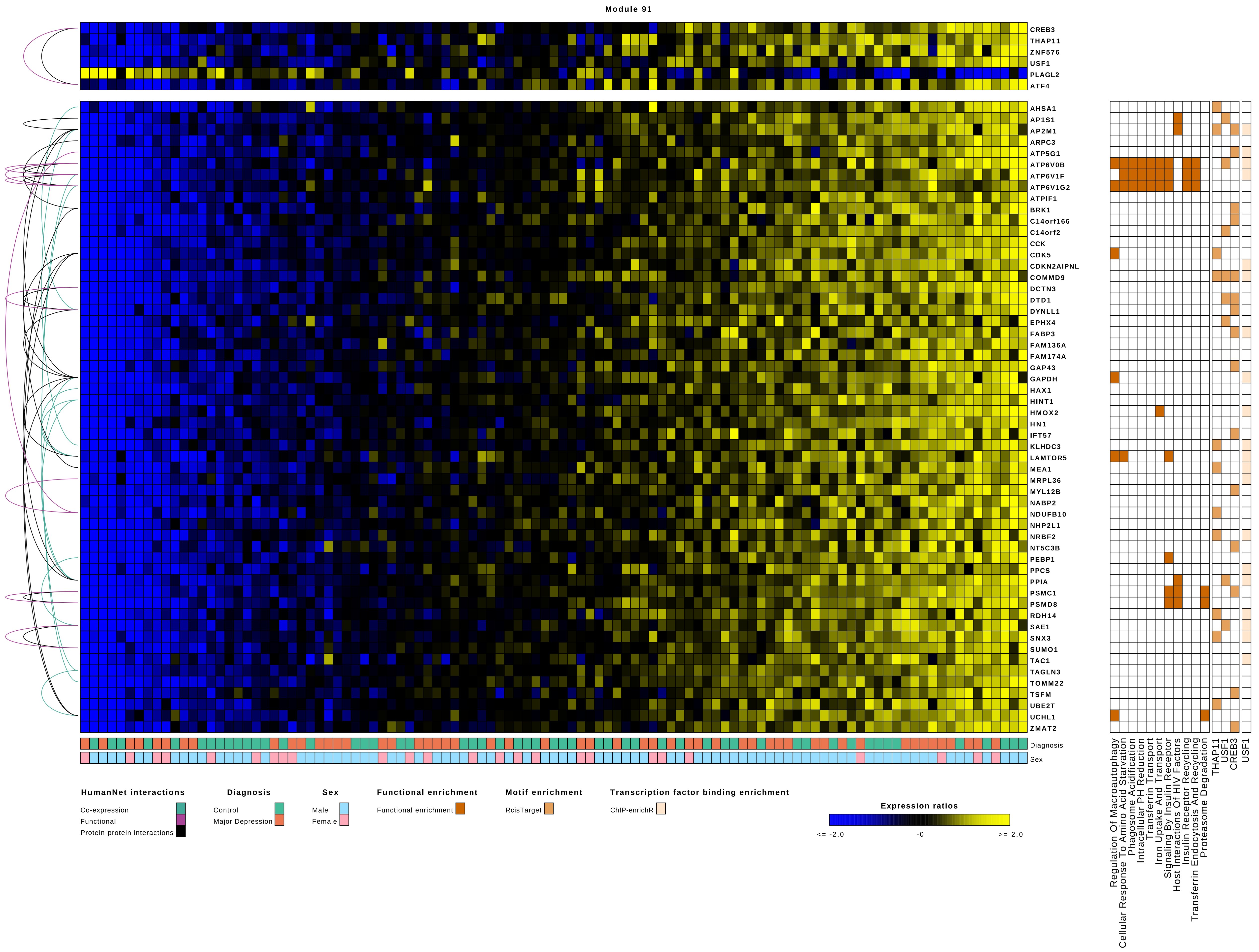
Module 91
CREB3
THAP11
ZNF576
USF1
PLAGL2
ATF4
AHSA1
AP1S1
AP2M1
ARPC3
ATP5G1
ATP6V0B
ATP6V1F
ATP6V1G2
ATPIF1
BRK1
C14orf166
C14orf2
CCK
CDK5
CDKN2AIPNL
COMMD9
DCTN3
DTD1
DYNLL1
EPHX4
FABP3
FAM136A
FAM174A
GAP43
GAPDH
HAX1
HINT1
HMOX2
HN1
IFT57
KLHDC3
LAMTOR5
MEA1
MRPL36
MYL12B
NABP2
NDUFB10
NHP2L1
NRBF2
NT5C3B
PEBP1
PPCS
PPIA
PSMC1
PSMD8
RDH14
SAE1
SNX3
SUMO1
TAC1
TAGLN3
TOMM22
TSFM
UBE2T
UCHL1
ZMAT2
Diagnosis
Sex
Regulation Of Macroautophagy
Cellular Response To Amino Acid Starvation
Phagosome Acidification
Intracellular PH Reduction
Transferrin Transport
Iron Uptake And Transport
Signaling By Insulin Receptor
Host Interactions Of HIV Factors
Insulin Receptor Recycling
Transferrin Endocytosis And Recycling
Proteasome Degradation
THAP11
USF1
CREB3
USF1
HumanNet interactions
Co-expression
Functional
Protein-protein interactions
Diagnosis
Control
Major Depression
Sex
Male
Female
Functional enrichment
Functional enrichment
Motif enrichment
RcisTarget
Transcription factor binding enrichment
ChIP-enrichR
Expression ratios
<= -2.0
-0
>= 2.0

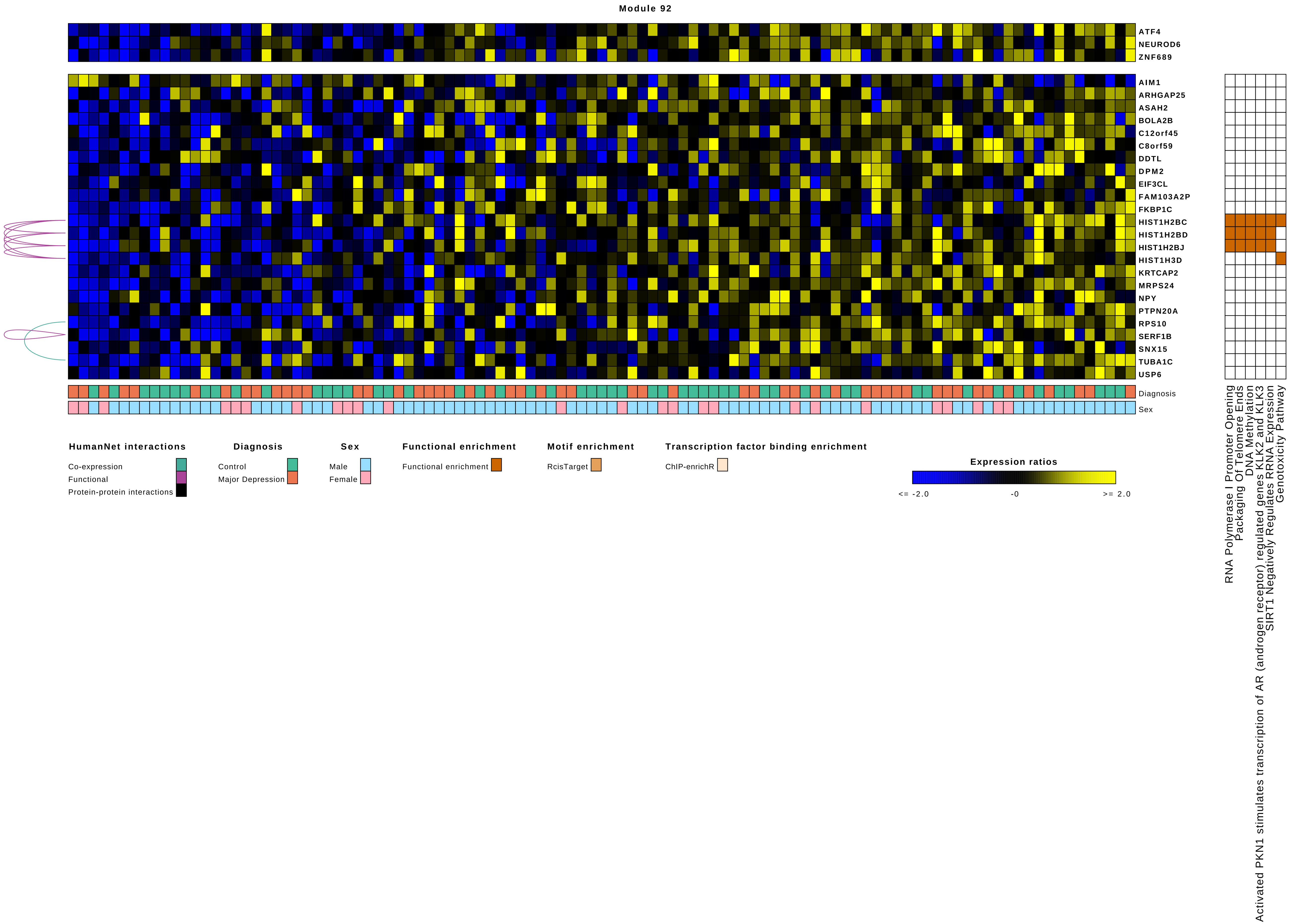
Module 92
ATF4
NEUROD6
ZNF689
AIM1
ARHGAP25
ASAH2
BOLA2B
C12orf45
C8orf59
DDTL
DPM2
EIF3CL
FAM103A2P
FKBP1C
HIST1H2BC
HIST1H2BD
HIST1H2BJ
HIST1H3D
KRTCAP2
MRPS24
NPY
PTPN20A
RPS10
SERF1B
SNX15
TUBA1C
USP6
Diagnosis
Sex
RNA Polymerase I Promoter Opening
Packaging Of Telomere Ends
DNA Methylation
Activated PKN1 stimulates transcription of AR (androgen receptor) regulated genes KLK2 and KLK3
SIRT1 Negatively Regulates RRNA Expression
Genotoxicity Pathway
HumanNet interactions
Co-expression
Functional
Protein-protein interactions
Diagnosis
Control
Major Depression
Sex
Male
Female
Functional enrichment
Functional enrichment
Motif enrichment
RcisTarget
Transcription factor binding enrichment
ChIP-enrichR
Expression ratios
<= -2.0
-0
>= 2.0

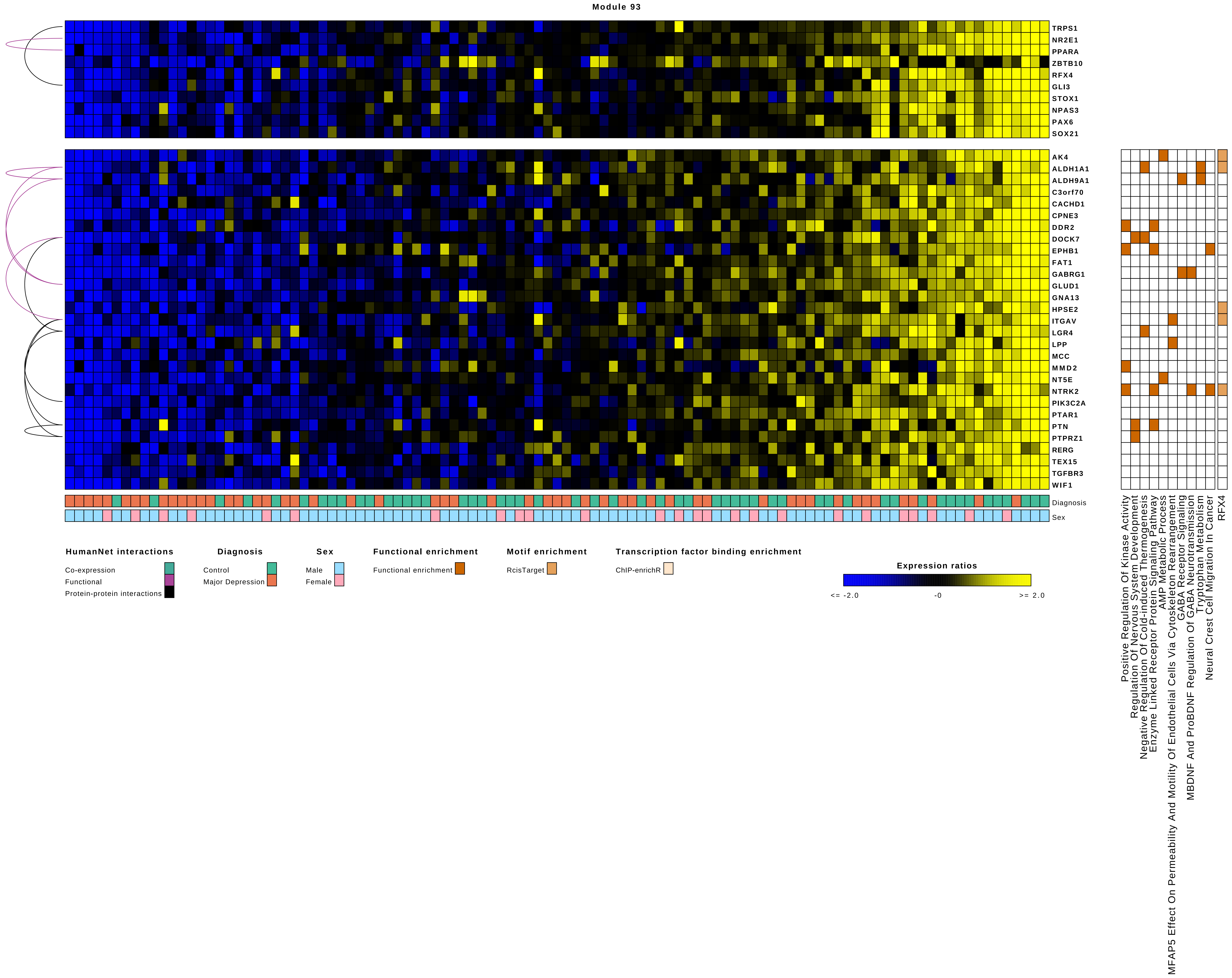

Module 93
TRPS1
NR2E1
PPARA
ZBTB10
RFX4
GLI3
STOX1
NPAS3
PAX6
SOX21
AK4
ALDH1A1
ALDH9A1
C3orf70
CACHD1
CPNE3
DDR2
DOCK7
EPHB1
FAT1
GABRG1
GLUD1
GNA13
HPSE2
ITGAV
LGR4
LPP
MCC
MMD2
NT5E
NTRK2
PIK3C2A
PTAR1
PTN
PTPRZ1
RERG
TEX15
TGFBR3
WIF1
Diagnosis
Sex
Positive Regulation Of Kinase Activity
Regulation Of Nervous System Development
Negative Regulation Of Cold-induced Thermogenesis
Enzyme Linked Receptor Protein Signaling Pathway
AMP Metabolic Process
MFAP5 Effect On Permeability And Motility Of Endothelial Cells Via Cytoskeleton Rearrangement
GABA Receptor Signaling
MBDNF And ProBDNF Regulation Of GABA Neurotransmission
Tryptophan Metabolism
Neural Crest Cell Migration In Cancer
RFX4
HumanNet interactions
Co-expression
Functional
Protein-protein interactions
Diagnosis
Control
Major Depression
Sex
Male
Female
Functional enrichment
Functional enrichment
Motif enrichment
RcisTarget
Transcription factor binding enrichment
ChIP-enrichR
Expression ratios
<= -2.0
-0
>= 2.0

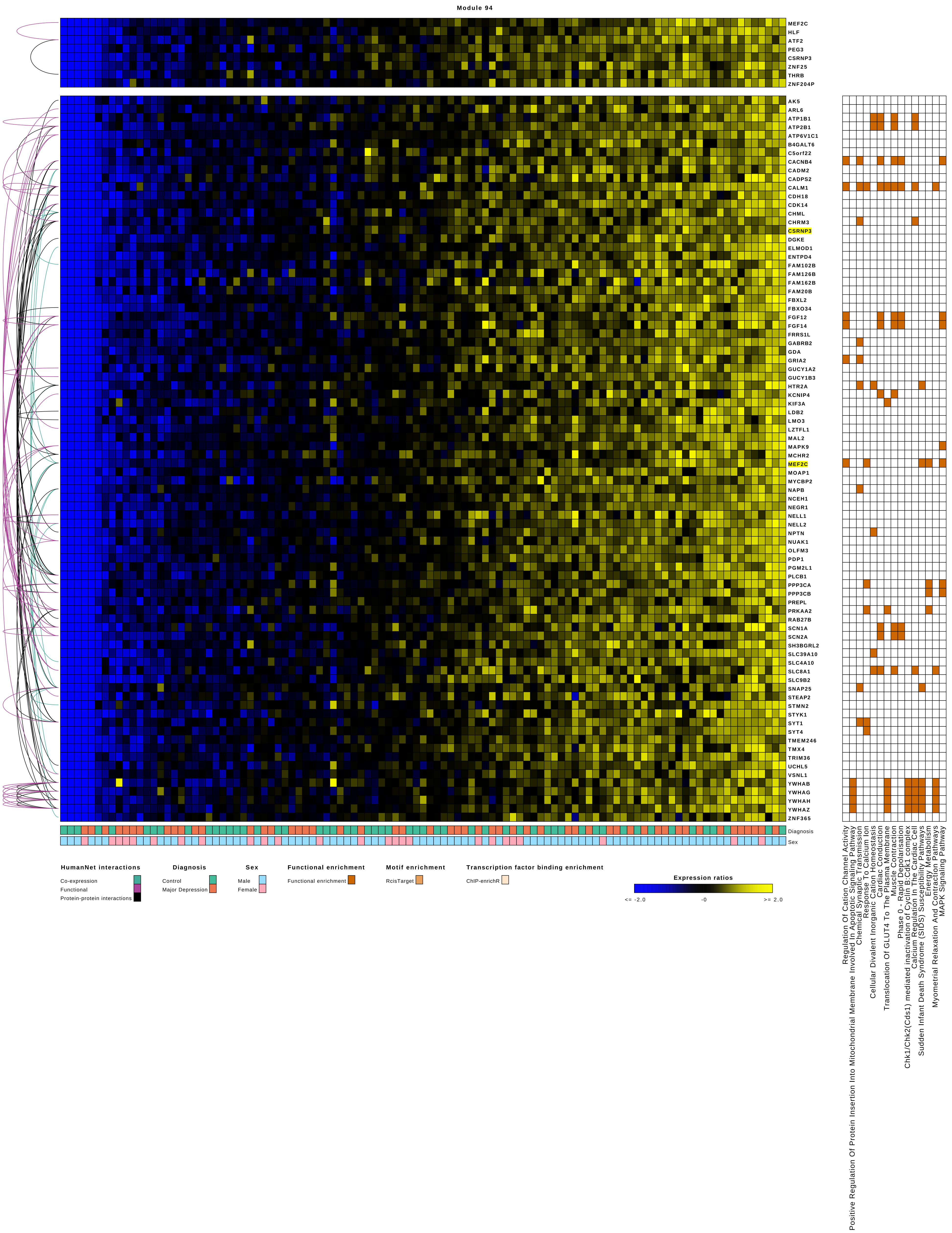
Module 94
MEF2C
HLF
ATF2
PEG3
CSRNP3
ZNF25
THRB
ZNF204P
AK5
ARL6
ATP1B1
ATP2B1
ATP6V1C1
B4GALT6
C5orf22
CACNB4
CADM2
CADPS2
CALM1
CDH18
CDK14
CHML
CHRM3
CSRNP3
DGKE
ELMOD1
ENTPD4
FAM102B
FAM126B
FAM162B
FAM20B
FBXL2
FBXO34
FGF12
FGF14
FRRS1L
GABRB2
GDA
GRIA2
GUCY1A2
GUCY1B3
HTR2A
KCNIP4
KIF3A
LDB2
LMO3
LZTFL1
MAL2
MAPK9
MCHR2
MEF2C
MOAP1
MYCBP2
NAPB
NCEH1
NEGR1
NELL1
NELL2
NPTN
NUAK1
OLFM3
PDP1
PGM2L1
PLCB1
PPP3CA
PPP3CB
PREPL
PRKAA2
RAB27B
SCN1A
SCN2A
SH3BGRL2
SLC39A10
SLC4A10
SLC8A1
SLC9B2
SNAP25
STEAP2
STMN2
STYK1
SYT1
SYT4
TMEM246
TMX4
TRIM36
UCHL5
VSNL1
YWHAB
YWHAG
YWHAH
YWHAZ
ZNF365
Diagnosis
Sex
HumanNet interactions
Co-expression
Functional
Protein-protein interactions
Diagnosis
Control
Major Depression
Sex
Male
Female
Functional enrichment
Functional enrichment
Motif enrichment
RcisTarget
Transcription factor binding enrichment
ChIP-enrichR
Expression ratios
<= -2.0
-0
>= 2.0
Regulation Of Cation Channel Activity
Positive Regulation Of Protein Insertion Into Mitochondrial Membrane Involved In Apoptotic Signaling Pathway
Chemical Synaptic Transmission
Response To Calcium Ion
Cellular Divalent Inorganic Cation Homeostasis
Cardiac Conduction
Translocation Of GLUT4 To The Plasma Membrane
Muscle Contraction
Phase 0 - Rapid Depolarisation
Chk1/Chk2(Cds1) mediated inactivation of Cyclin B:Cdk1 complex
Calcium Regulation In The Cardiac Cell
Sudden Infant Death Syndrome (SIDS) Susceptibility Pathways
Energy Metabolism
Myometrial Relaxation And Contraction Pathways
MAPK Signaling Pathway

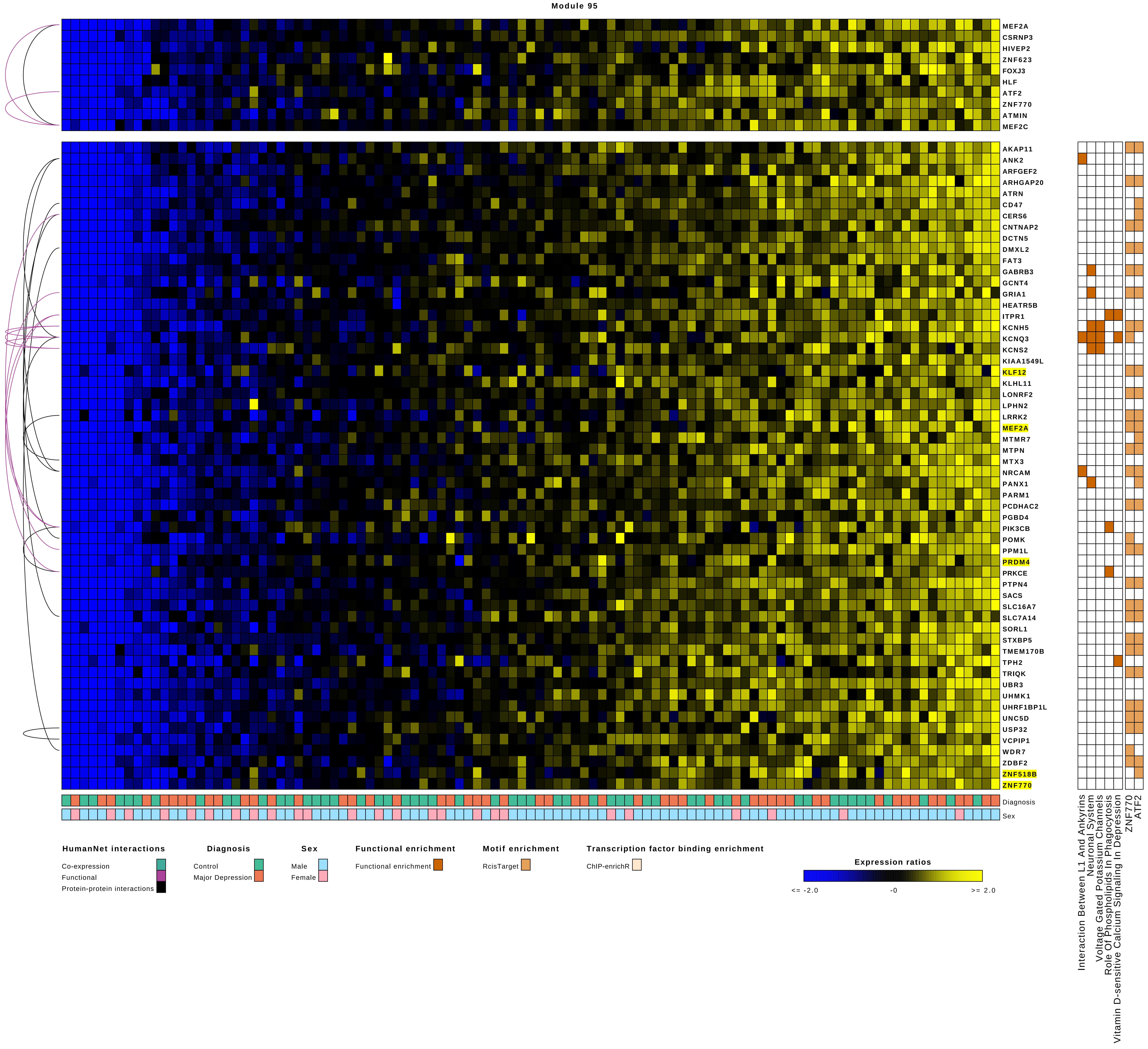
Module 95
MEF2A
CSRNP3
HIVEP2
ZNF623
FOXJ3
HLF
ATF2
ZNF770
ATMIN
MEF2C
AKAP11
ANK2
ARFGEF2
ARHGAP20
ATRN
CD47
CERS6
CNTNAP2
DCTN5
DMXL2
FAT3
GABRB3
GCNT4
GRIA1
HEATR5B
ITPR1
KCNH5
KCNQ3
KCNS2
KIAA1549L
KLF12
KLHL11
LONRF2
LPHN2
LRRK2
MEF2A
MTMR7
MTPN
MTX3
NRCAM
PANX1
PARM1
PCDHAC2
PGBD4
PIK3CB
POMK
PPM1L
PRDM4
PRKCE
PTPN4
SACS
SLC16A7
SLC7A14
SORL1
STXBP5
TMEM170B
TPH2
TRIQK
UBR3
UHMK1
UHRF1BP1L
UNC5D
USP32
VCPIP1
WDR7
ZDBF2
ZNF518B
ZNF770
Diagnosis
Sex
Interaction Between L1 And Ankyrins
Neuronal System
Voltage Gated Potassium Channels
Role Of Phospholipids In Phagocytosis
Vitamin D-sensitive Calcium Signaling In Depression
ZNF770
ATF2
HumanNet interactions
Co-expression
Functional
Protein-protein interactions
Diagnosis
Control
Major Depression
Sex
Male
Female
Functional enrichment
Functional enrichment
Motif enrichment
RcisTarget
Transcription factor binding enrichment
ChIP-enrichR
Expression ratios
<= -2.0
-0
>= 2.0

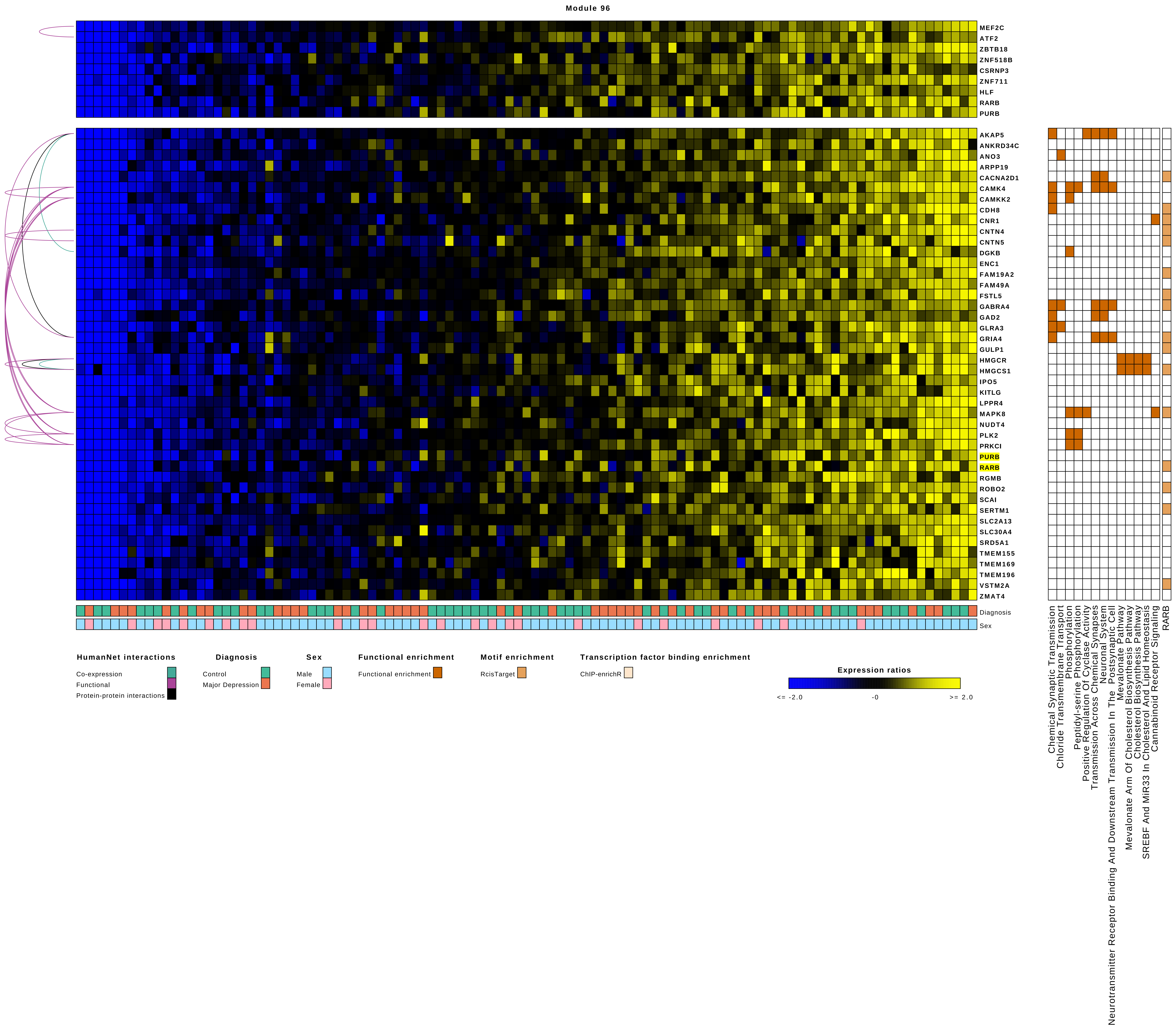

Module 96
MEF2C
ATF2
ZBTB18
ZNF518B
CSRNP3
ZNF711
HLF
RARB
PURB
AKAP5
ANKRD34C
ANO3
ARPP19
CACNA2D1
CAMK4
CAMKK2
CDH8
CNR1
CNTN4
CNTN5
DGKB
ENC1
FAM19A2
FAM49A
FSTL5
GABRA4
GAD2
GLRA3
GRIA4
GULP1
HMGCR
HMGCS1
IPO5
KITLG
LPPR4
MAPK8
NUDT4
PLK2
PRKCI
PURB
RARB
RGMB
ROBO2
SCAI
SERTM1
SLC2A13
SLC30A4
SRD5A1
TMEM155
TMEM169
TMEM196
VSTM2A
ZMAT4
Diagnosis
Sex
HumanNet interactions
Co-expression
Functional
Protein-protein interactions
Diagnosis
Control
Major Depression
Sex
Male
Female
Functional enrichment
Functional enrichment
Motif enrichment
RcisTarget
Transcription factor binding enrichment
ChIP-enrichR
Expression ratios
<= -2.0
-0
>= 2.0
Chemical Synaptic Transmission
Chloride Transmembrane Transport
Phosphorylation
Peptidyl-serine Phosphorylation
Positive Regulation Of Cyclase Activity
Transmission Across Chemical Synapses
Neuronal System
Neurotransmitter Receptor Binding And Downstream Transmission In The Postsynaptic Cell
Mevalonate Pathway
Mevalonate Arm Of Cholesterol Biosynthesis Pathway
Cholesterol Biosynthesis Pathway
SREBF And MiR33 In Cholesterol And Lipid Homeostasis
Cannabinoid Receptor Signaling
RARB

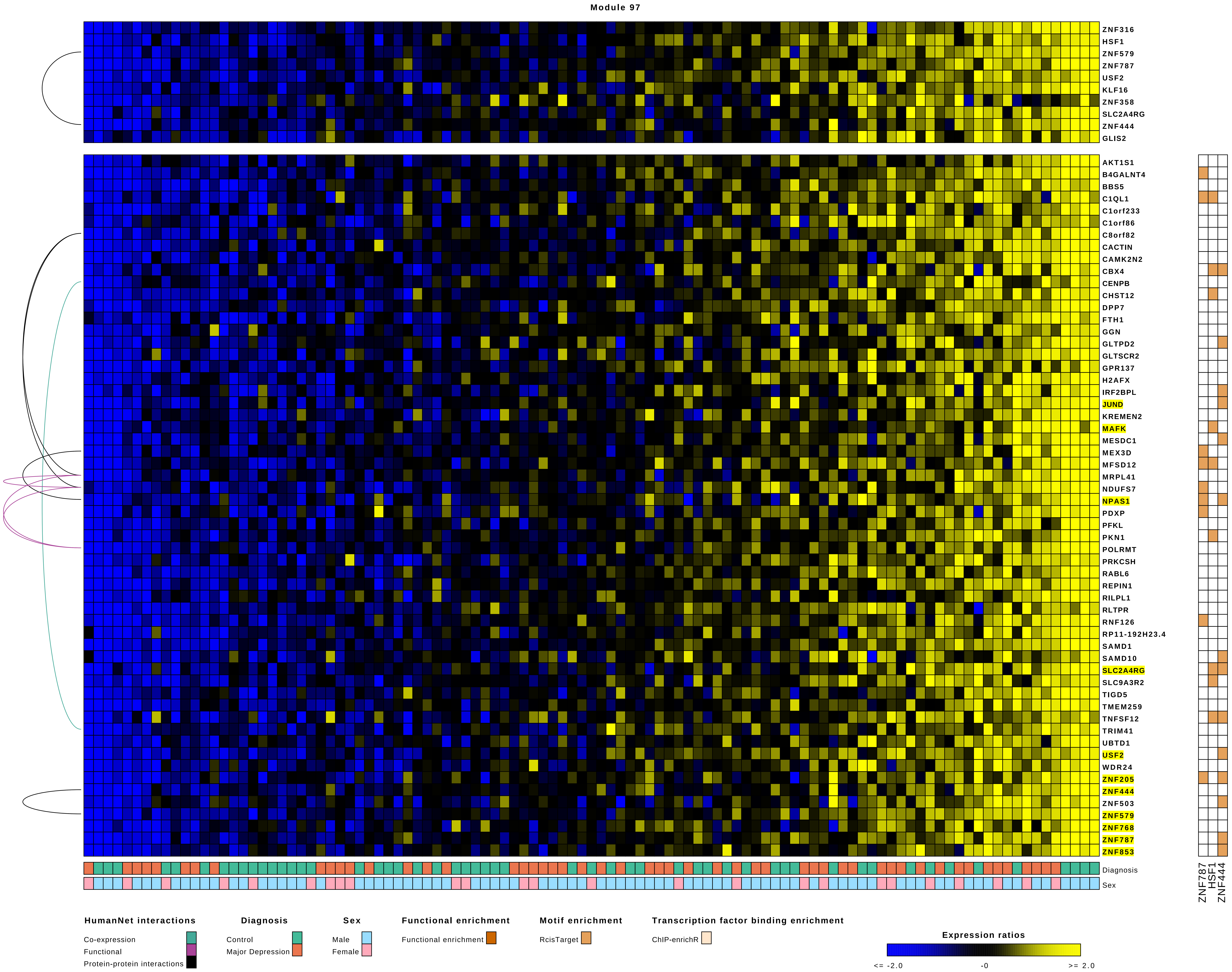

Module 97
ZNF316
HSF1
ZNF579
ZNF787
USF2
KLF16
ZNF358
SLC2A4RG
ZNF444
GLIS2
AKT1S1
B4GALNT4
BBS5
C1QL1
C1orf233
C1orf86
C8orf82
CACTIN
CAMK2N2
CBX4
CENPB
CHST12
DPP7
FTH1
GGN
GLTPD2
GLTSCR2
GPR137
H2AFX
IRF2BPL
JUND
KREMEN2
MAFK
MESDC1
MEX3D
MFSD12
MRPL41
NDUFS7
NPAS1
PDXP
PFKL
PKN1
POLRMT
PRKCSH
RABL6
REPIN1
RILPL1
RLTPR
RNF126
RP11-192H23.4
SAMD1
SAMD10
SLC2A4RG
SLC9A3R2
TIGD5
TMEM259
TNFSF12
TRIM41
UBTD1
USF2
WDR24
ZNF205
ZNF444
ZNF503
ZNF579
ZNF768
ZNF787
ZNF853
Diagnosis
Sex
ZNF787
HSF1
ZNF444
HumanNet interactions
Co-expression
Functional
Protein-protein interactions
Diagnosis
Control
Major Depression
Sex
Male
Female
Functional enrichment
Functional enrichment
Motif enrichment
RcisTarget
Transcription factor binding enrichment
ChIP-enrichR
Expression ratios
<= -2.0
-0
>= 2.0

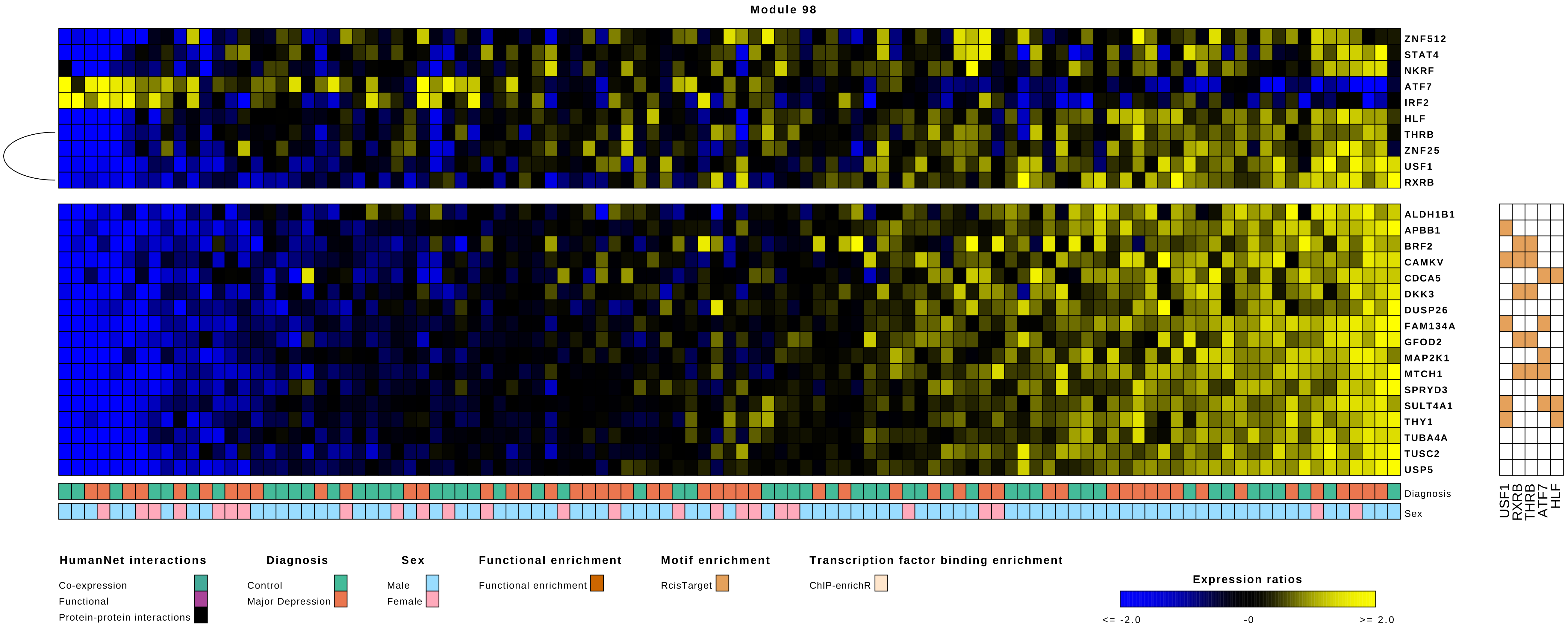

Module 98
ZNF512
STAT4
NKRF
ATF7
IRF2
HLF
THRB
ZNF25
USF1
RXRB
ALDH1B1
APBB1
BRF2
CAMKV
CDCA5
DKK3
DUSP26
FAM134A
GFOD2
MAP2K1
MTCH1
SPRYD3
SULT4A1
THY1
TUBA4A
TUSC2
USP5
Diagnosis
Sex
USF1
RXRB
THRB
ATF7
HLF
HumanNet interactions
Co-expression
Functional
Protein-protein interactions
Diagnosis
Control
Major Depression
Sex
Male
Female
Functional enrichment
Functional enrichment
Motif enrichment
RcisTarget
Transcription factor binding enrichment
ChIP-enrichR
Expression ratios
<= -2.0
-0
>= 2.0

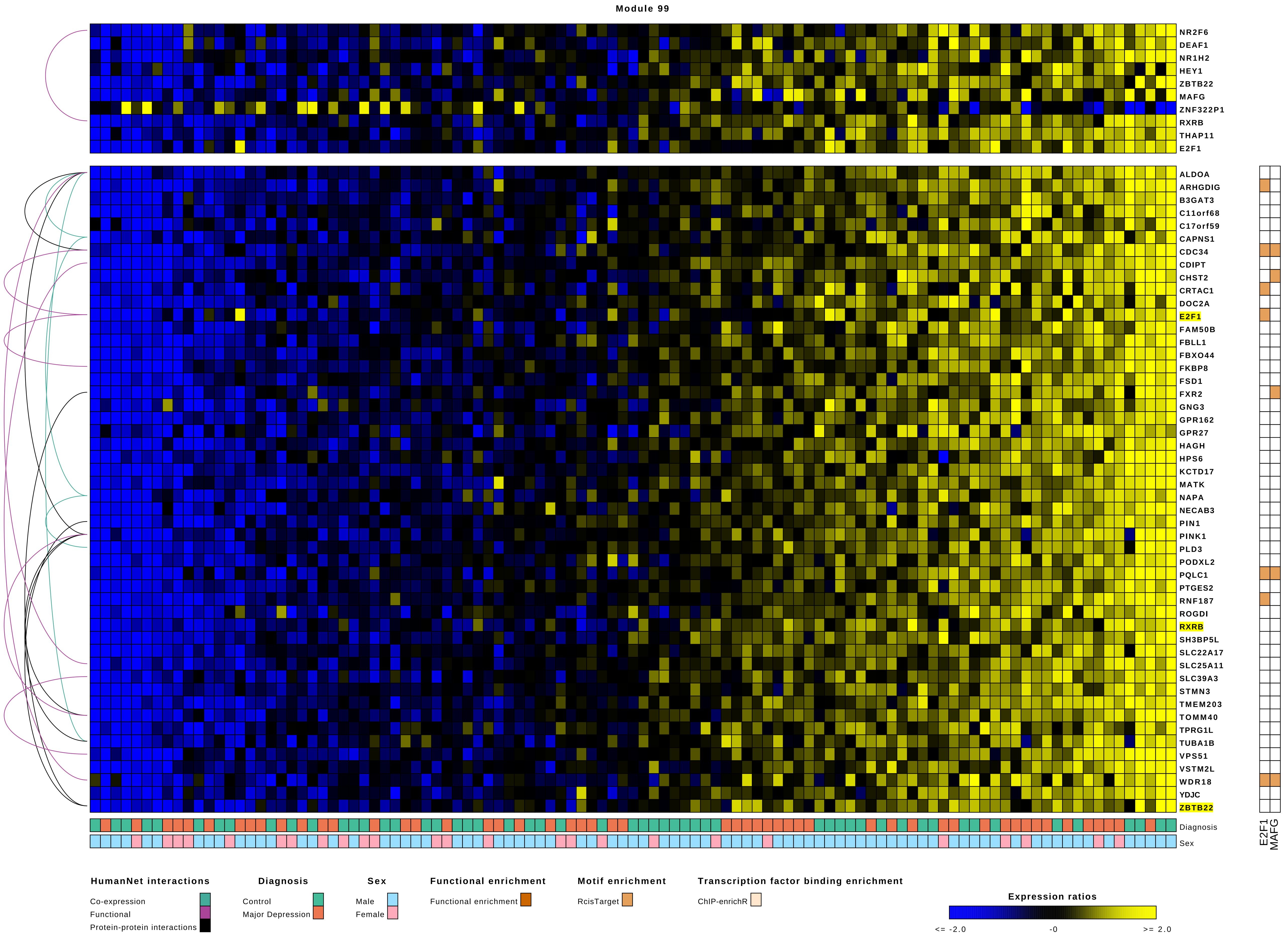
Module 99
NR2F6
DEAF1
NR1H2
HEY1
ZBTB22
MAFG
ZNF322P1
RXRB
THAP11
E2F1
ALDOA
ARHGDIG
B3GAT3
C11orf68
C17orf59
CAPNS1
CDC34
CDIPT
CHST2
CRTAC1
DOC2A
E2F1
FAM50B
FBLL1
FBXO44
FKBP8
FSD1
FXR2
GNG3
GPR162
GPR27
HAGH
HPS6
KCTD17
MATK
NAPA
NECAB3
PIN1
PINK1
PLD3
PODXL2
PQLC1
PTGES2
RNF187
ROGDI
RXRB
SH3BP5L
SLC22A17
SLC25A11
SLC39A3
STMN3
TMEM203
TOMM40
TPRG1L
TUBA1B
VPS51
VSTM2L
WDR18
YDJC
ZBTB22
Diagnosis
Sex
E2F1
MAFG
HumanNet interactions
Co-expression
Functional
Protein-protein interactions
Diagnosis
Control
Major Depression
Sex
Male
Female
Functional enrichment
Functional enrichment
Motif enrichment
RcisTarget
Transcription factor binding enrichment
ChIP-enrichR
Expression ratios
<= -2.0
-0
>= 2.0

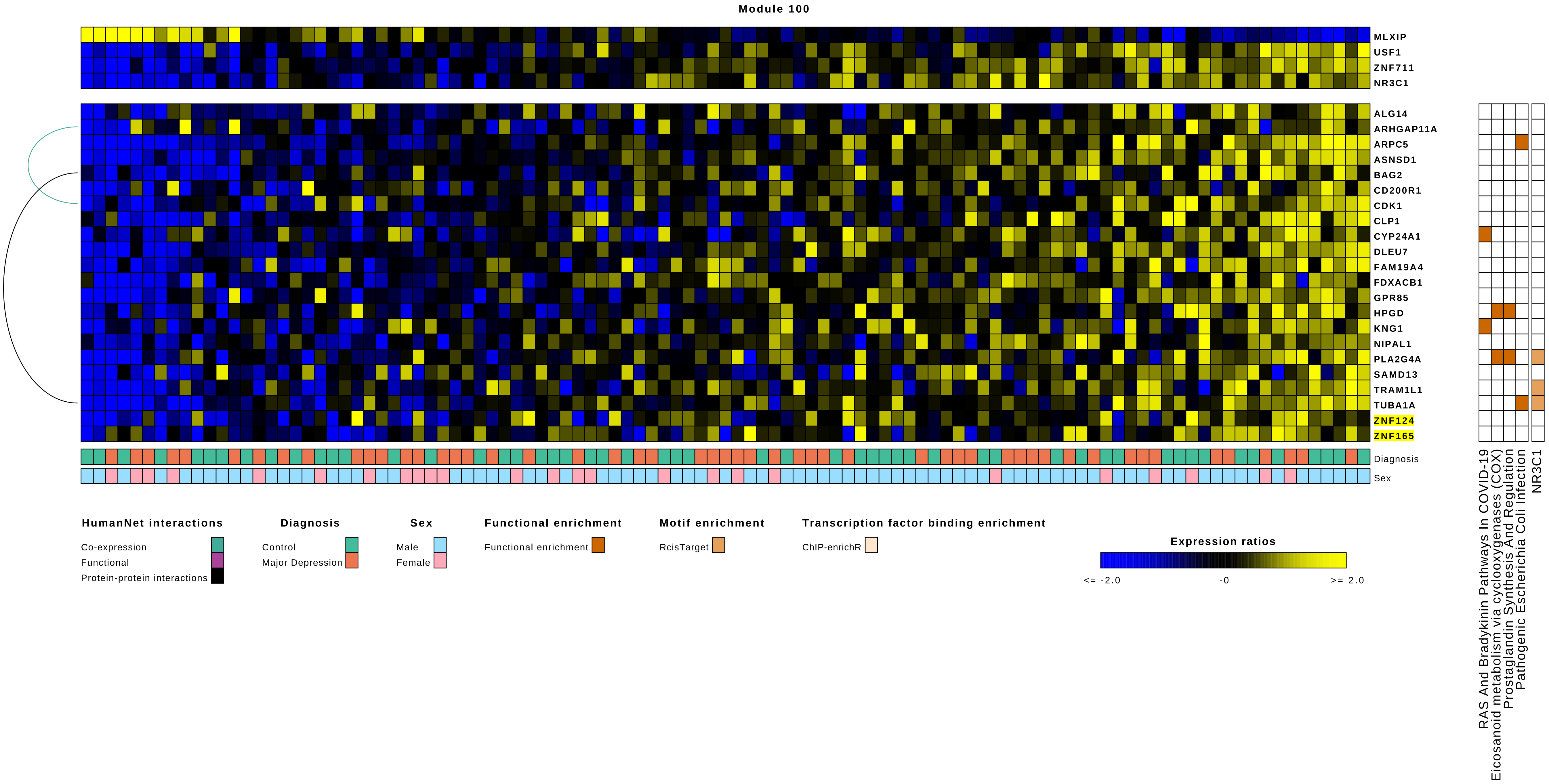
Module 100
MLXIP
USF1
ZNF711
NR3C1
ALG14
ARHGAP11A
ARPC5
ASNSD1
BAG2
CD200R1
CDK1
CLP1
CYP24A1
DLEU7
FAM19A4
FDXACB1
GPR85
HPGD
KNG1
NIPAL1
PLA2G4A
SAMD13
TRAM1L1
TUBA1A
ZNF124
ZNF165
Diagnosis
Sex
RAS And Bradykinin Pathways In COVID-19
Eicosanoid metabolism via cyclooxygenases (COX)
Prostaglandin Synthesis And Regulation
Pathogenic Escherichia Coli Infection
NR3C1
HumanNet interactions
Co-expression
Functional
Protein-protein interactions
Diagnosis
Control
Major Depression
Sex
Male
Female
Functional enrichment
Functional enrichment
Motif enrichment
RcisTarget
Transcription factor binding enrichment
ChIP-enrichR
Expression ratios
<= -2.0
-0
>= 2.0

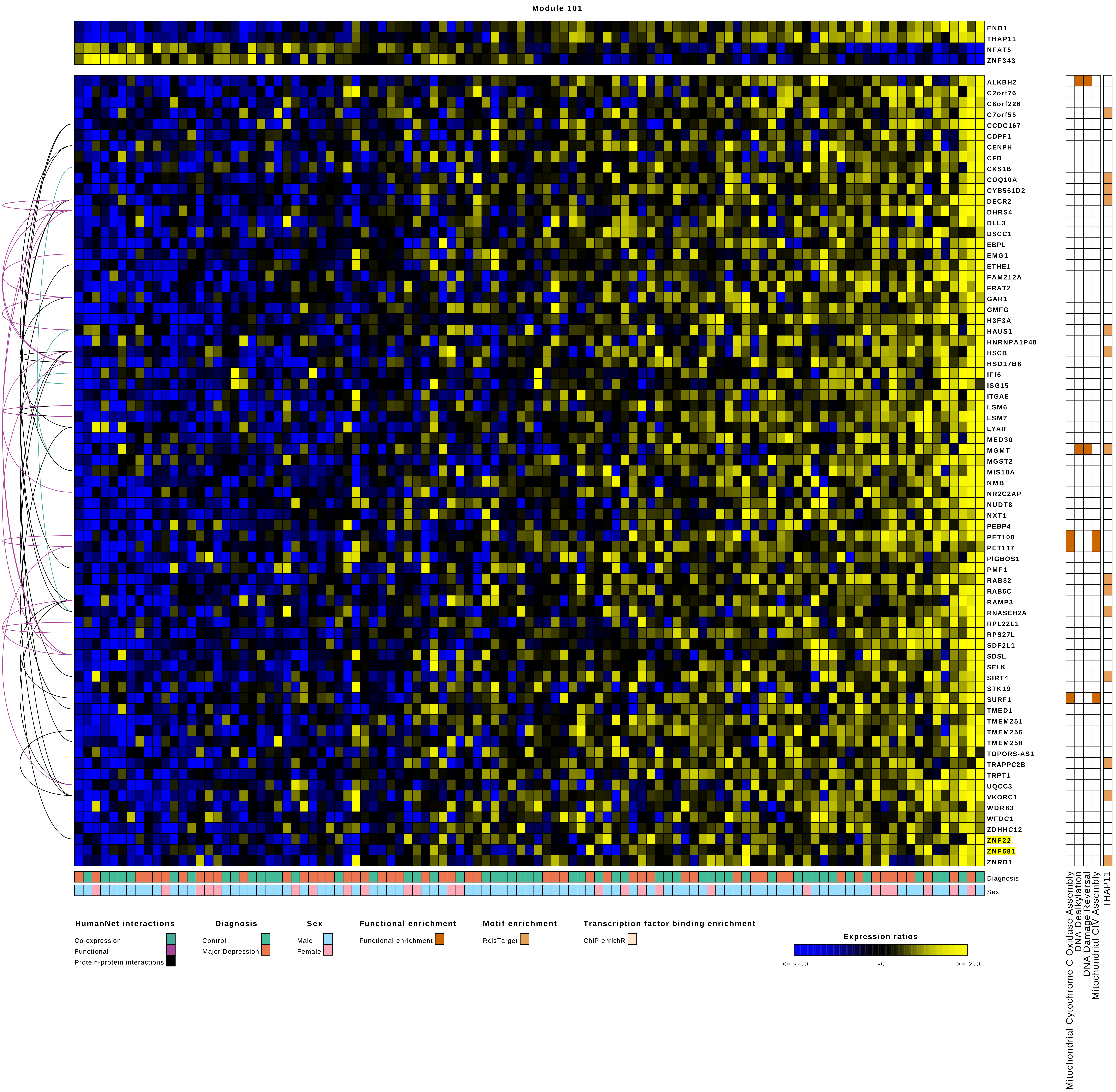
Module 101
ENO1
THAP11
NFAT5
ZNF343
ALKBH2
C2orf76
C6orf226
C7orf55
CCDC167
CDPF1
CENPH
CFD
CKS1B
COQ10A
CYB561D2
DECR2
DHRS4
DLL3
DSCC1
EBPL
EMG1
ETHE1
FAM212A
FRAT2
GAR1
GMFG
H3F3A
HAUS1
HNRNPA1P48
HSCB
HSD17B8
IFI6
ISG15
ITGAE
LSM6
LSM7
LYAR
MED30
MGMT
MGST2
MIS18A
NMB
NR2C2AP
NUDT8
NXT1
PEBP4
PET100
PET117
PIGBOS1
PMF1
RAB32
RAB5C
RAMP3
RNASEH2A
RPL22L1
RPS27L
SDF2L1
SDSL
SELK
SIRT4
STK19
SURF1
TMED1
TMEM251
TMEM256
TMEM258
TOPORS-AS1
TRAPPC2B
TRPT1
UQCC3
VKORC1
WDR83
WFDC1
ZDHHC12
ZNF22
ZNF581
ZNRD1
Diagnosis
Sex
Mitochondrial Cytochrome C Oxidase Assembly
DNA Dealkylation
DNA Damage Reversal
Mitochondrial CIV Assembly
THAP11
HumanNet interactions
Co-expression
Functional
Protein-protein interactions
Diagnosis
Control
Major Depression
Sex
Male
Female
Functional enrichment
Functional enrichment
Motif enrichment
RcisTarget
Transcription factor binding enrichment
ChIP-enrichR
Expression ratios
<= -2.0
-0
>= 2.0

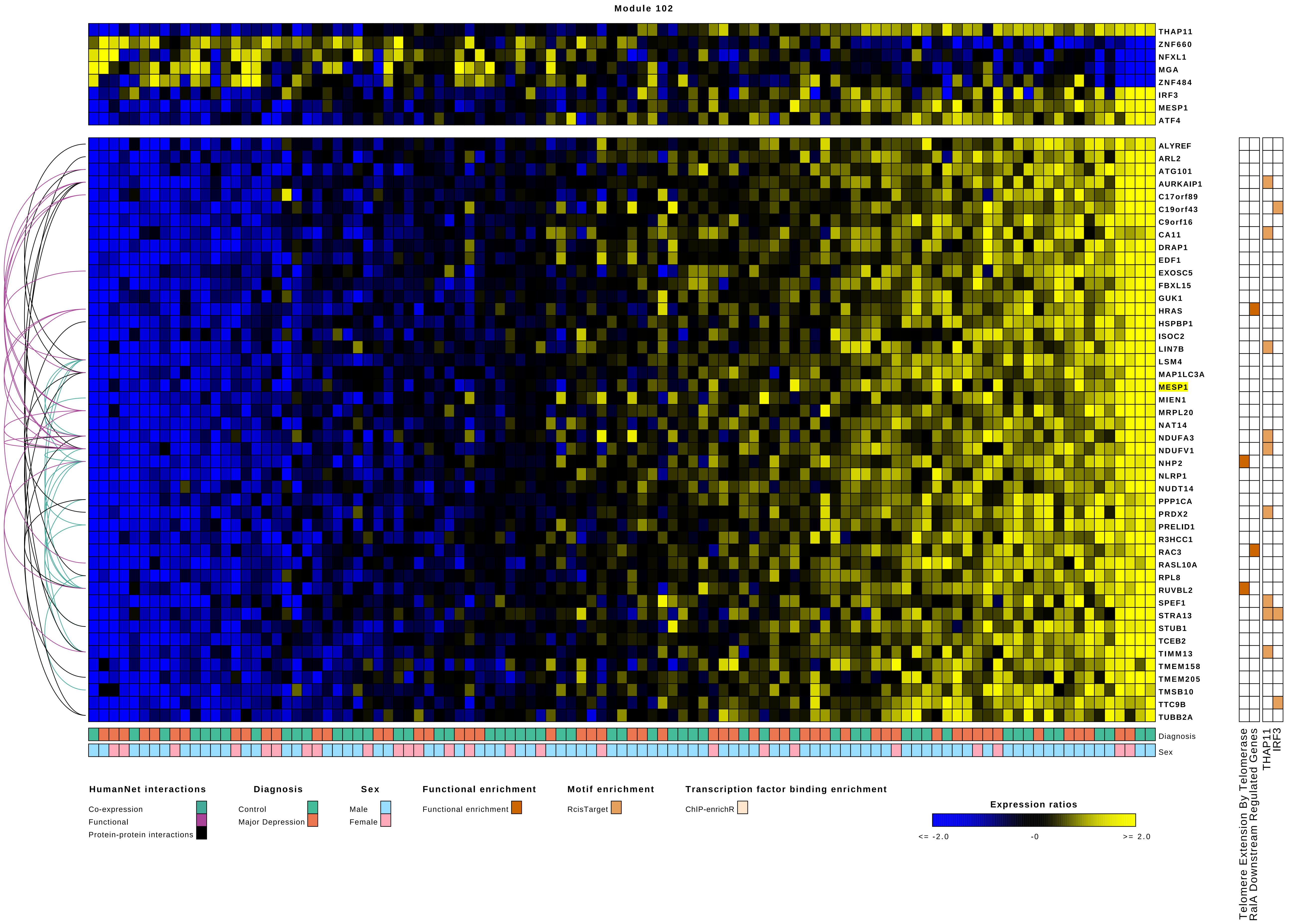
Module 102
THAP11
ZNF660
NFXL1
MGA
ZNF484
IRF3
MESP1
ATF4
ALYREF
ARL2
ATG101
AURKAIP1
C17orf89
C19orf43
C9orf16
CA11
DRAP1
EDF1
EXOSC5
FBXL15
GUK1
HRAS
HSPBP1
ISOC2
LIN7B
LSM4
MAP1LC3A
MESP1
MIEN1
MRPL20
NAT14
NDUFA3
NDUFV1
NHP2
NLRP1
NUDT14
PPP1CA
PRDX2
PRELID1
R3HCC1
RAC3
RASL10A
RPL8
RUVBL2
SPEF1
STRA13
STUB1
TCEB2
TIMM13
TMEM158
TMEM205
TMSB10
TTC9B
TUBB2A
Diagnosis
Sex
Telomere Extension By Telomerase
RalA Downstream Regulated Genes
THAP11
IRF3
HumanNet interactions
Co-expression
Functional
Protein-protein interactions
Diagnosis
Control
Major Depression
Sex
Male
Female
Functional enrichment
Functional enrichment
Motif enrichment
RcisTarget
Transcription factor binding enrichment
ChIP-enrichR
Expression ratios
<= -2.0
-0
>= 2.0

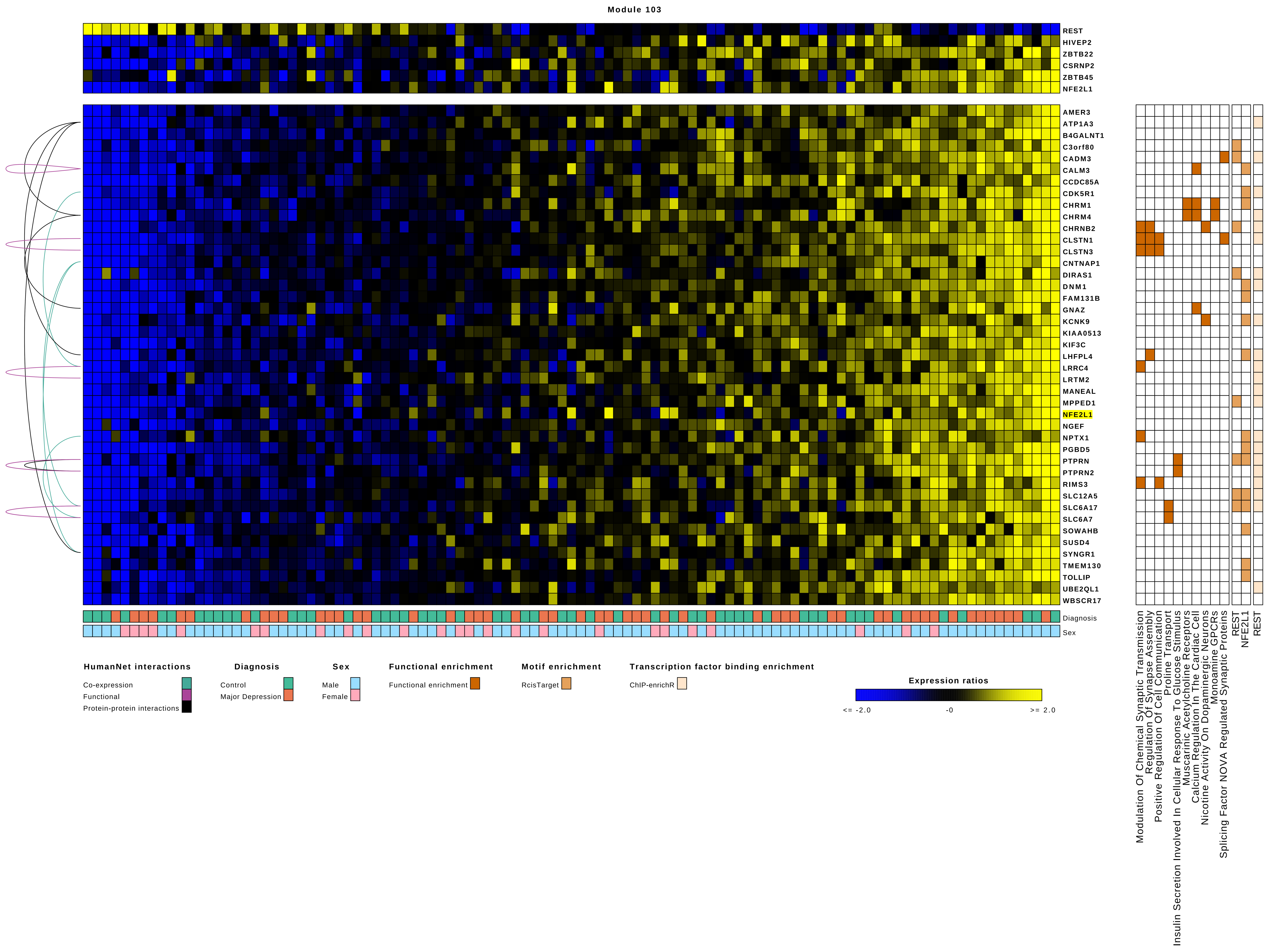

Module 103
REST
HIVEP2
ZBTB22
CSRNP2
ZBTB45
NFE2L1
AMER3
ATP1A3
B4GALNT1
C3orf80
CADM3
CALM3
CCDC85A
CDK5R1
CHRM1
CHRM4
CHRNB2
CLSTN1
CLSTN3
CNTNAP1
DIRAS1
DNM1
FAM131B
GNAZ
KCNK9
KIAA0513
KIF3C
LHFPL4
LRRC4
LRTM2
MANEAL
MPPED1
NFE2L1
NGEF
NPTX1
PGBD5
PTPRN
PTPRN2
RIMS3
SLC12A5
SLC6A17
SLC6A7
SOWAHB
SUSD4
SYNGR1
TMEM130
TOLLIP
UBE2QL1
WBSCR17
Diagnosis
Sex
Modulation Of Chemical Synaptic Transmission
Regulation Of Synapse Assembly
Positive Regulation Of Cell Communication
Proline Transport
Insulin Secretion Involved In Cellular Response To Glucose Stimulus
Muscarinic Acetylcholine Receptors
Calcium Regulation In The Cardiac Cell
Nicotine Activity On Dopaminergic Neurons
Monoamine GPCRs
Splicing Factor NOVA Regulated Synaptic Proteins
REST
NFE2L1
REST
HumanNet interactions
Co-expression
Functional
Protein-protein interactions
Diagnosis
Control
Major Depression
Sex
Male
Female
Functional enrichment
Functional enrichment
Motif enrichment
RcisTarget
Transcription factor binding enrichment
ChIP-enrichR
Expression ratios
<= -2.0
-0
>= 2.0

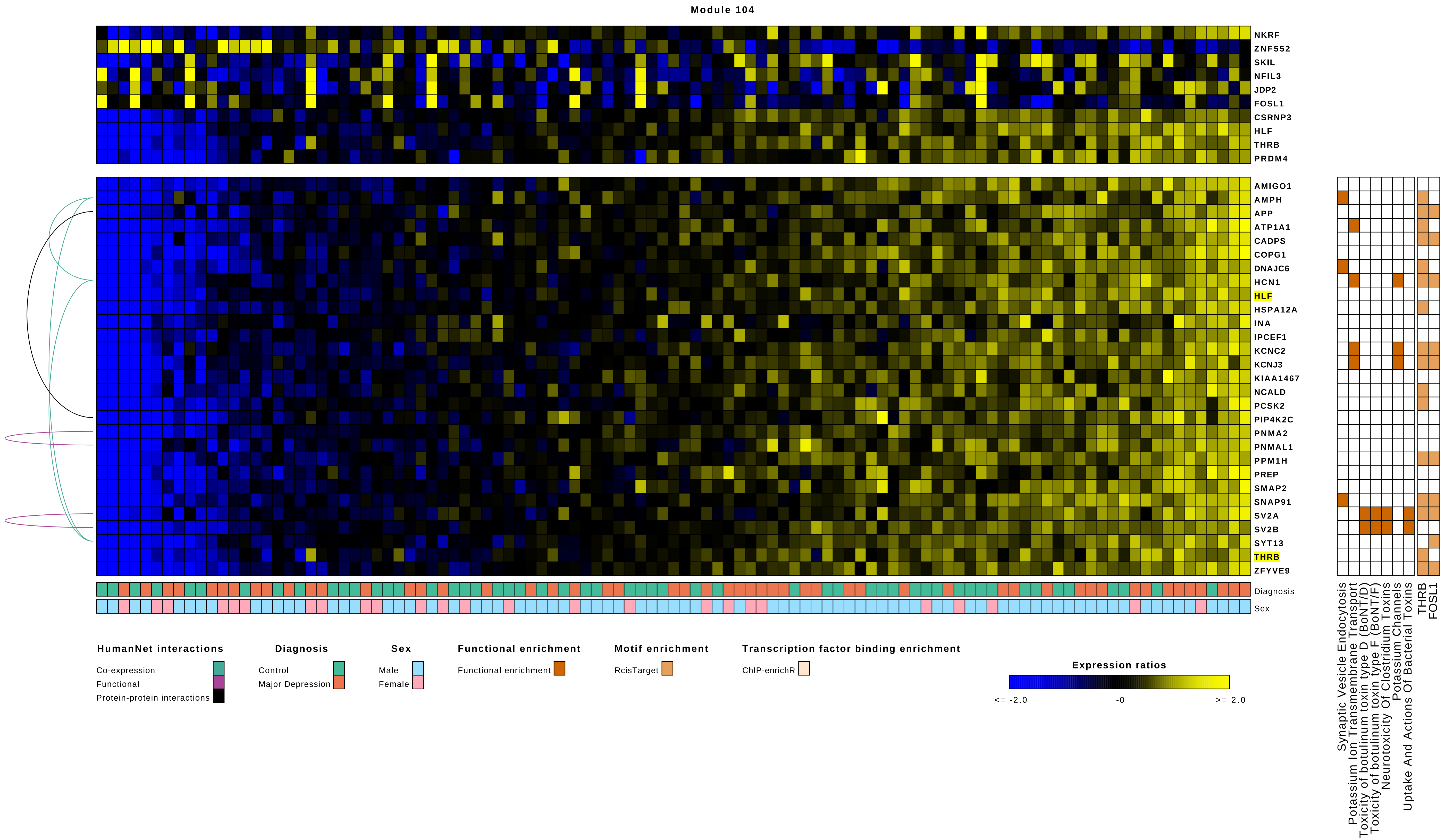
Module 104
NKRF
ZNF552
SKIL
NFIL3
JDP2
FOSL1
CSRNP3
HLF
THRB
PRDM4
AMIGO1
AMPH
APP
ATP1A1
CADPS
COPG1
DNAJC6
HCN1
HLF
HSPA12A
INA
IPCEF1
KCNC2
KCNJ3
KIAA1467
NCALD
PCSK2
PIP4K2C
PNMA2
PNMAL1
PPM1H
PREP
SMAP2
SNAP91
SV2A
SV2B
SYT13
THRB
ZFYVE9
Diagnosis
Sex
Synaptic Vesicle Endocytosis
Potassium Ion Transmembrane Transport
Toxicity of botulinum toxin type D (BoNT/D)
Toxicity of botulinum toxin type F (BoNT/F)
Neurotoxicity Of Clostridium Toxins
Potassium Channels
Uptake And Actions Of Bacterial Toxins
THRB
FOSL1
HumanNet interactions
Co-expression
Functional
Protein-protein interactions
Diagnosis
Control
Major Depression
Sex
Male
Female
Functional enrichment
Functional enrichment
Motif enrichment
RcisTarget
Transcription factor binding enrichment
ChIP-enrichR
Expression ratios
<= -2.0
-0
>= 2.0

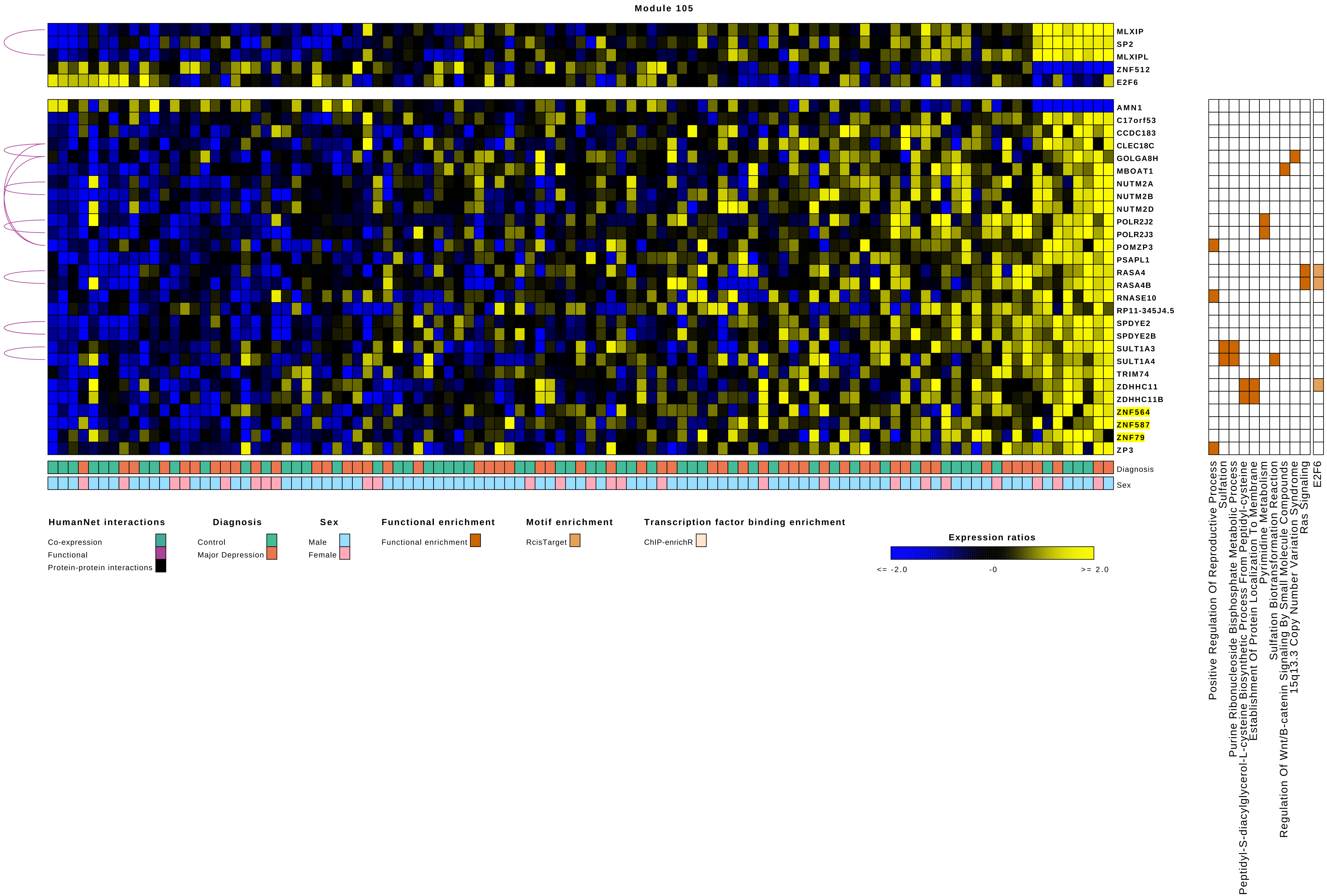
Module 105
MLXIP
SP2
MLXIPL
ZNF512
E2F6
AMN1
C17orf53
CCDC183
CLEC18C
GOLGA8H
MBOAT1
NUTM2A
NUTM2B
NUTM2D
POLR2J2
POLR2J3
POMZP3
PSAPL1
RASA4
RASA4B
RNASE10
RP11-345J4.5
SPDYE2
SPDYE2B
SULT1A3
SULT1A4
TRIM74
ZDHHC11
ZDHHC11B
ZNF564
ZNF587
ZNF79
ZP3
Diagnosis
Sex
Positive Regulation Of Reproductive Process
Sulfation
Purine Ribonucleoside Bisphosphate Metabolic Process
Peptidyl-S-diacylglycerol-L-cysteine Biosynthetic Process From Peptidyl-cysteine
Establishment Of Protein Localization To Membrane
Pyrimidine Metabolism
Sulfation Biotransformation Reaction
Regulation Of Wnt/B-catenin Signaling By Small Molecule Compounds
15q13.3 Copy Number Variation Syndrome
Ras Signaling
E2F6
HumanNet interactions
Co-expression
Functional
Protein-protein interactions
Diagnosis
Control
Major Depression
Sex
Male
Female
Functional enrichment
Functional enrichment
Motif enrichment
RcisTarget
Transcription factor binding enrichment
ChIP-enrichR
Expression ratios
<= -2.0
-0
>= 2.0

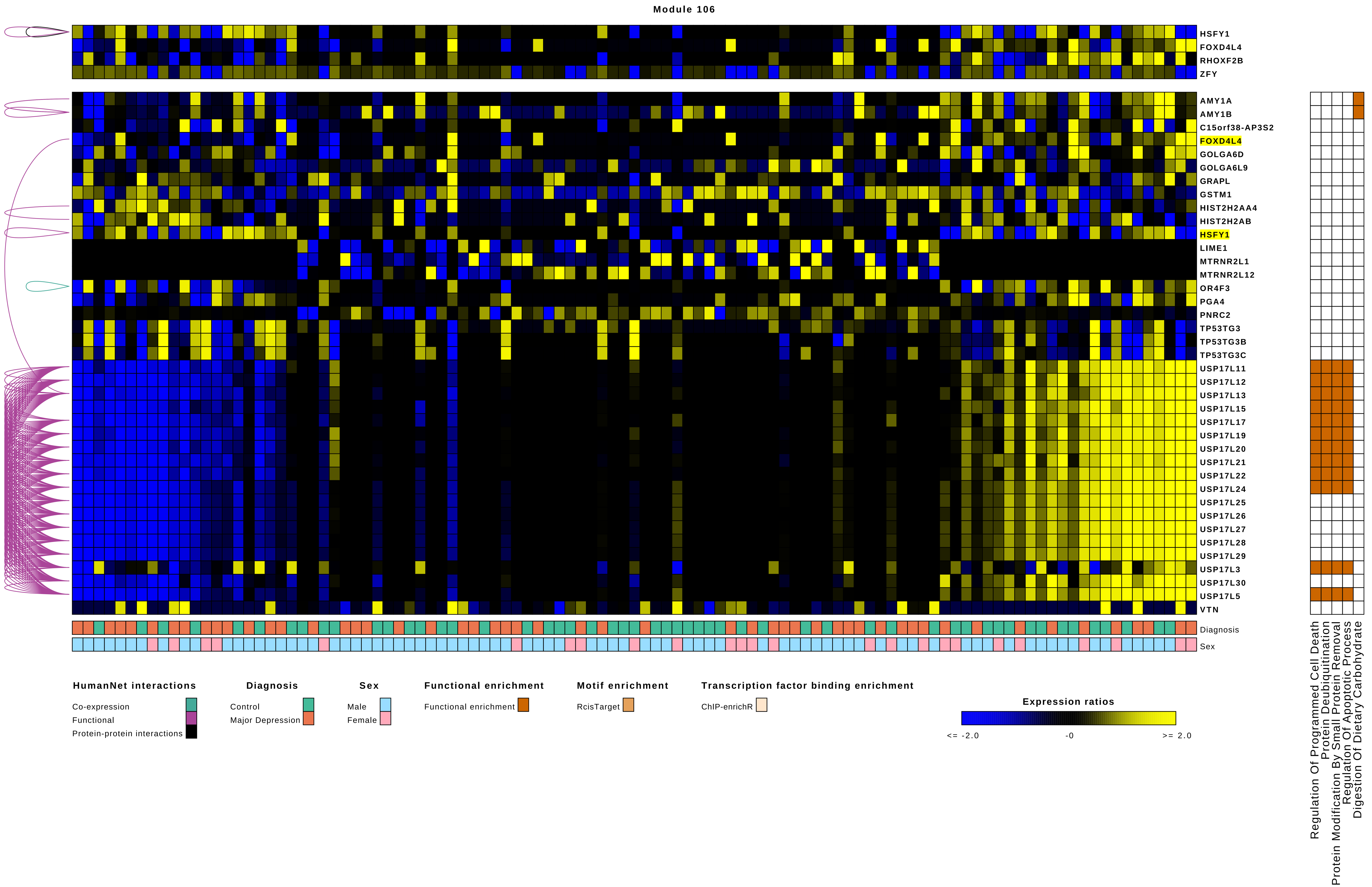
Module 106
HSFY1
FOXD4L4
RHOXF2B
ZFY
AMY1A
AMY1B
C15orf38-AP3S2
FOXD4L4
GOLGA6D
GOLGA6L9
GRAPL
GSTM1
HIST2H2AA4
HIST2H2AB
HSFY1
LIME1
MTRNR2L1
MTRNR2L12
OR4F3
PGA4
PNRC2
TP53TG3
TP53TG3B
TP53TG3C
USP17L11
USP17L12
USP17L13
USP17L15
USP17L17
USP17L19
USP17L20
USP17L21
USP17L22
USP17L24
USP17L25
USP17L26
USP17L27
USP17L28
USP17L29
USP17L3
USP17L30
USP17L5
VTN
Diagnosis
Sex
Regulation Of Programmed Cell Death
Protein Deubiquitination
Protein Modification By Small Protein Removal
Regulation Of Apoptotic Process
Digestion Of Dietary Carbohydrate
HumanNet interactions
Co-expression
Functional
Protein-protein interactions
Diagnosis
Control
Major Depression
Sex
Male
Female
Functional enrichment
Functional enrichment
Motif enrichment
RcisTarget
Transcription factor binding enrichment
ChIP-enrichR
Expression ratios
<= -2.0
-0
>= 2.0

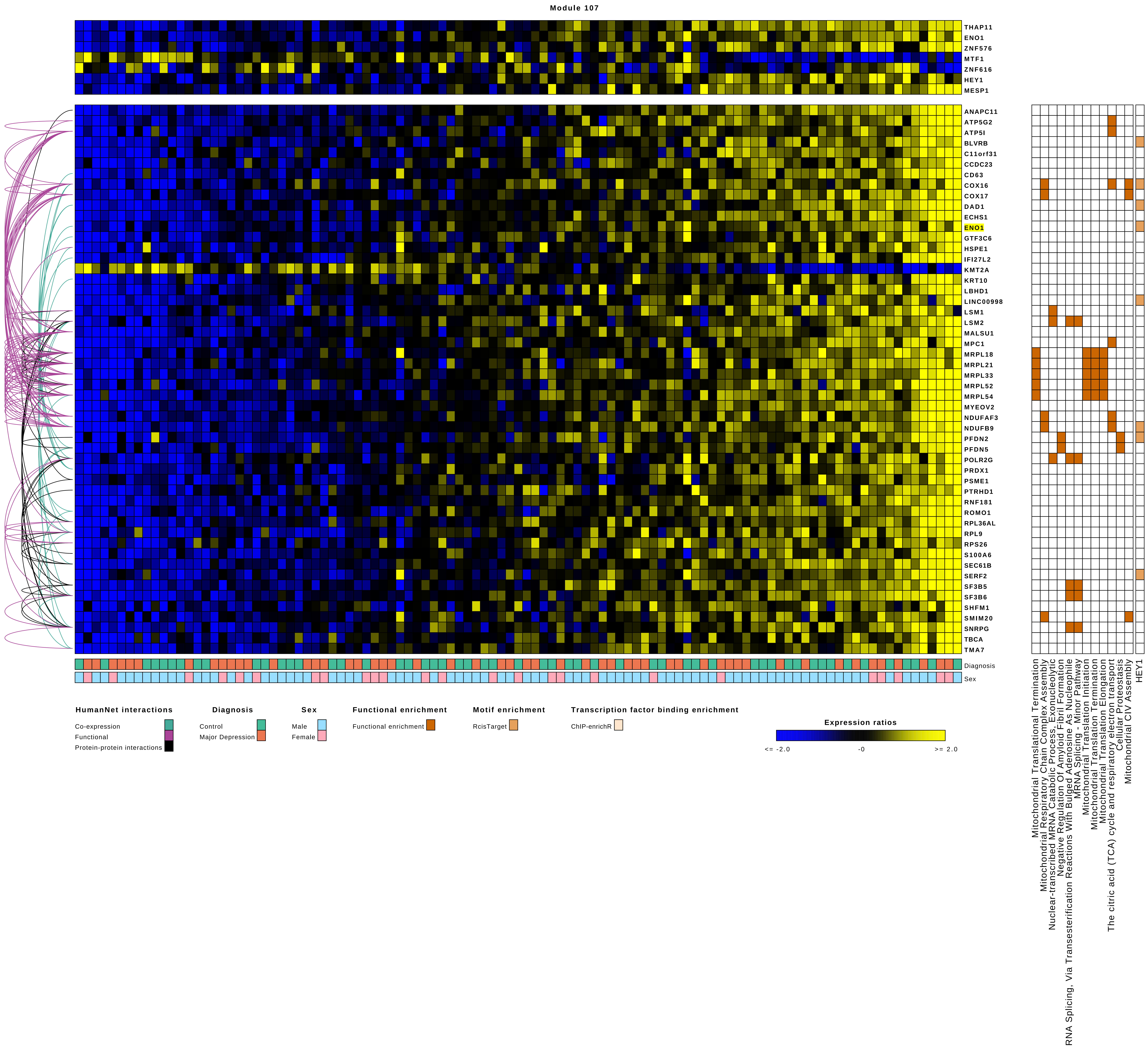
Module 107
THAP11
ENO1
ZNF576
MTF1
ZNF616
HEY1
MESP1
ANAPC11
ATP5G2
ATP5I
BLVRB
C11orf31
CCDC23
CD63
COX16
COX17
DAD1
ECHS1
ENO1
GTF3C6
HSPE1
IFI27L2
KMT2A
KRT10
LBHD1
LINC00998
LSM1
LSM2
MALSU1
MPC1
MRPL18
MRPL21
MRPL33
MRPL52
MRPL54
MYEOV2
NDUFAF3
NDUFB9
PFDN2
PFDN5
POLR2G
PRDX1
PSME1
PTRHD1
RNF181
ROMO1
RPL36AL
RPL9
RPS26
S100A6
SEC61B
SERF2
SF3B5
SF3B6
SHFM1
SMIM20
SNRPG
TBCA
TMA7
Diagnosis
Sex
Mitochondrial Translational Termination
Mitochondrial Respiratory Chain Complex Assembly
Nuclear-transcribed MRNA Catabolic Process, Exonucleolytic
Negative Regulation Of Amyloid Fibril Formation
RNA Splicing, Via Transesterification Reactions With Bulged Adenosine As Nucleophile
MRNA Splicing - Minor Pathway
Mitochondrial Translation Initiation
Mitochondrial Translation Termination
Mitochondrial Translation Elongation
The citric acid (TCA) cycle and respiratory electron transport
Cellular Proteostasis
Mitochondrial CIV Assembly
HEY1
HumanNet interactions
Co-expression
Functional
Protein-protein interactions
Diagnosis
Control
Major Depression
Sex
Male
Female
Functional enrichment
Functional enrichment
Motif enrichment
RcisTarget
Transcription factor binding enrichment
ChIP-enrichR
Expression ratios
<= -2.0
-0
>= 2.0

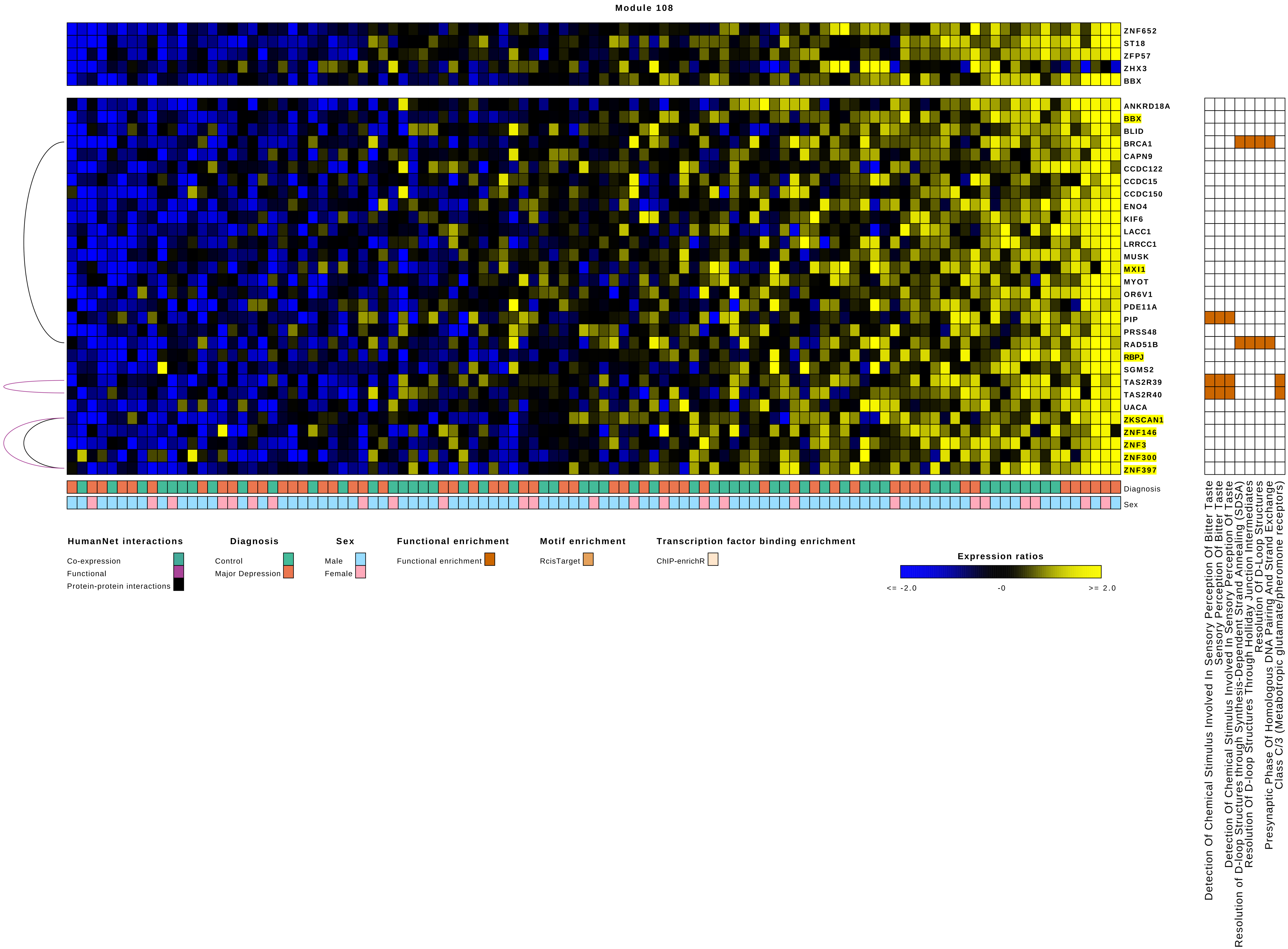
Module 108
ZNF652
ST18
ZFP57
ZHX3
BBX
ANKRD18A
BBX
BLID
BRCA1
CAPN9
CCDC122
CCDC15
CCDC150
ENO4
KIF6
LACC1
LRRCC1
MUSK
MXI1
MYOT
OR6V1
PDE11A
PIP
PRSS48
RAD51B
RBPJ
SGMS2
TAS2R39
TAS2R40
UACA
ZKSCAN1
ZNF146
ZNF3
ZNF300
ZNF397
Diagnosis
Sex
Detection Of Chemical Stimulus Involved In Sensory Perception Of Bitter Taste
Sensory Perception Of Bitter Taste
Detection Of Chemical Stimulus Involved In Sensory Perception Of Taste
Resolution of D-loop Structures through Synthesis-Dependent Strand Annealing (SDSA)
Resolution Of D-loop Structures Through Holliday Junction Intermediates
Resolution Of D-Loop Structures
Presynaptic Phase Of Homologous DNA Pairing And Strand Exchange
Class C/3 (Metabotropic glutamate/pheromone receptors)
HumanNet interactions
Co-expression
Functional
Protein-protein interactions
Diagnosis
Control
Major Depression
Sex
Male
Female
Functional enrichment
Functional enrichment
Motif enrichment
RcisTarget
Transcription factor binding enrichment
ChIP-enrichR
Expression ratios
<= -2.0
-0
>= 2.0

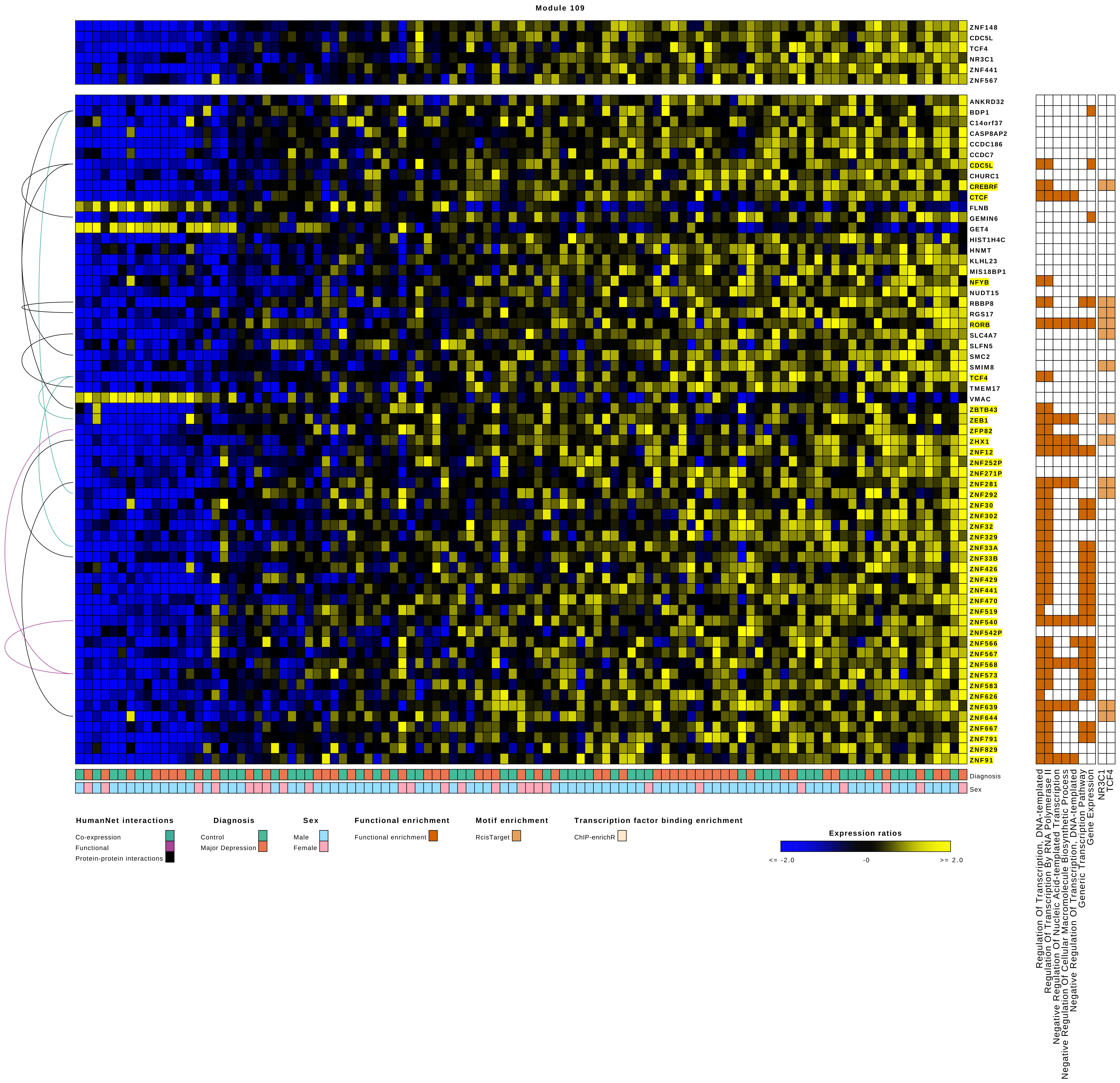
Module 109
ZNF148
CDC5L
TCF4
NR3C1
ZNF441
ZNF567
ANKRD32
BDP1
C14orf37
CASP8AP2
CCDC186
CCDC7
CDC5L
CHURC1
CREBRF
CTCF
FLNB
GEMIN6
GET4
HIST1H4C
HNMT
KLHL23
MIS18BP1
NFYB
NUDT15
RBBP8
RGS17
RORB
SLC4A7
SLFN5
SMC2
SMIM8
TCF4
TMEM17
VMAC
ZBTB43
ZEB1
ZFP82
ZHX1
ZNF12
ZNF252P
ZNF271P
ZNF281
ZNF292
ZNF30
ZNF302
ZNF32
ZNF329
ZNF33A
ZNF33B
ZNF426
ZNF429
ZNF441
ZNF470
ZNF519
ZNF540
ZNF542P
ZNF566
ZNF567
ZNF568
ZNF573
ZNF583
ZNF626
ZNF639
ZNF644
ZNF667
ZNF791
ZNF829
ZNF91
Diagnosis
Sex
Regulation Of Transcription, DNA-templated
Regulation Of Transcription By RNA Polymerase II
Negative Regulation Of Nucleic Acid-templated Transcription
Negative Regulation Of Cellular Macromolecule Biosynthetic Process
Negative Regulation Of Transcription, DNA-templated
Generic Transcription Pathway
Gene Expression
NR3C1
TCF4
HumanNet interactions
Co-expression
Functional
Protein-protein interactions
Diagnosis
Control
Major Depression
Sex
Male
Female
Functional enrichment
Functional enrichment
Motif enrichment
RcisTarget
Transcription factor binding enrichment
ChIP-enrichR
Expression ratios
<= -2.0
-0
>= 2.0

### Module 110

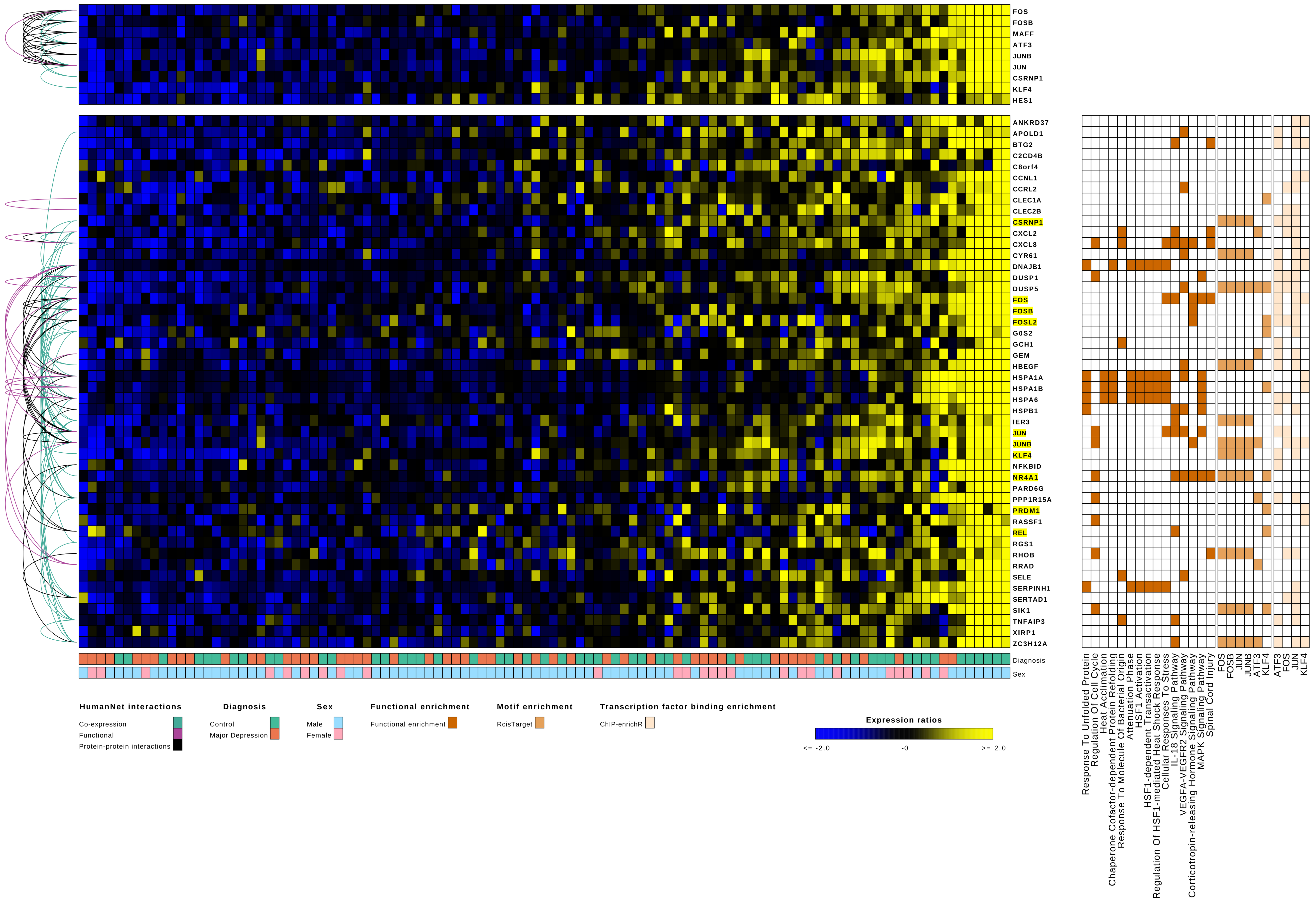
FOS
FOSB
MAFF
ATF3
JUNB
JUN
CSRNP1
KLF4
HES1
ANKRD37
APOLD1
BTG2
C2CD4B
C8orf4
CCNL1
CCRL2
CLEC1A
CLEC2B
CSRNP1
CXCL2
CXCL8
CYR61
DNAJB1
DUSP1
DUSP5
FOS
FOSB
FOSL2
G0S2
GCH1
GEM
HBEGF
HSPA1A
HSPA1B
HSPA6
HSPB1
IER3
JUN
JUNB
KLF4
NFKBID
NR4A1
PARD6G
PPP1R15A
PRDM1
RASSF1
REL
RGS1
RHOB
RRAD
SELE
SERPINH1
SERTAD1
SIK1
TNFAIP3
XIRP1
ZC3H12A
Diagnosis
Sex
Response To Unfolded Protein
Regulation Of Cell Cycle
Heat Acclimation
Chaperone Cofactor-dependent Protein Refolding
Response To Molecule Of Bacterial Origin
Attenuation Phase
HSF1 Activation
HSF1-dependent Transactivation
Regulation Of HSF1-mediated Heat Shock Response
Cellular Responses To Stress
IL-18 Signaling Pathway
VEGFA-VEGFR2 Signaling Pathway
Corticotropin-releasing Hormone Signaling Pathway
MAPK Signaling Pathway
Spinal Cord Injury
FOS
FOSB
JUN
JUNB
ATF3
KLF4
ATF3
FOS
JUN
KLF4
HumanNet interactions
Co-expression
Functional
Protein-protein interactions
Diagnosis
Control
Major Depression
Sex
Male
Female
Functional enrichment
Functional enrichment
Motif enrichment
RcisTarget
Transcription factor binding enrichment
ChIP-enrichR
Expression ratios
<= -2.0
-0
>= 2.0

### Module 111

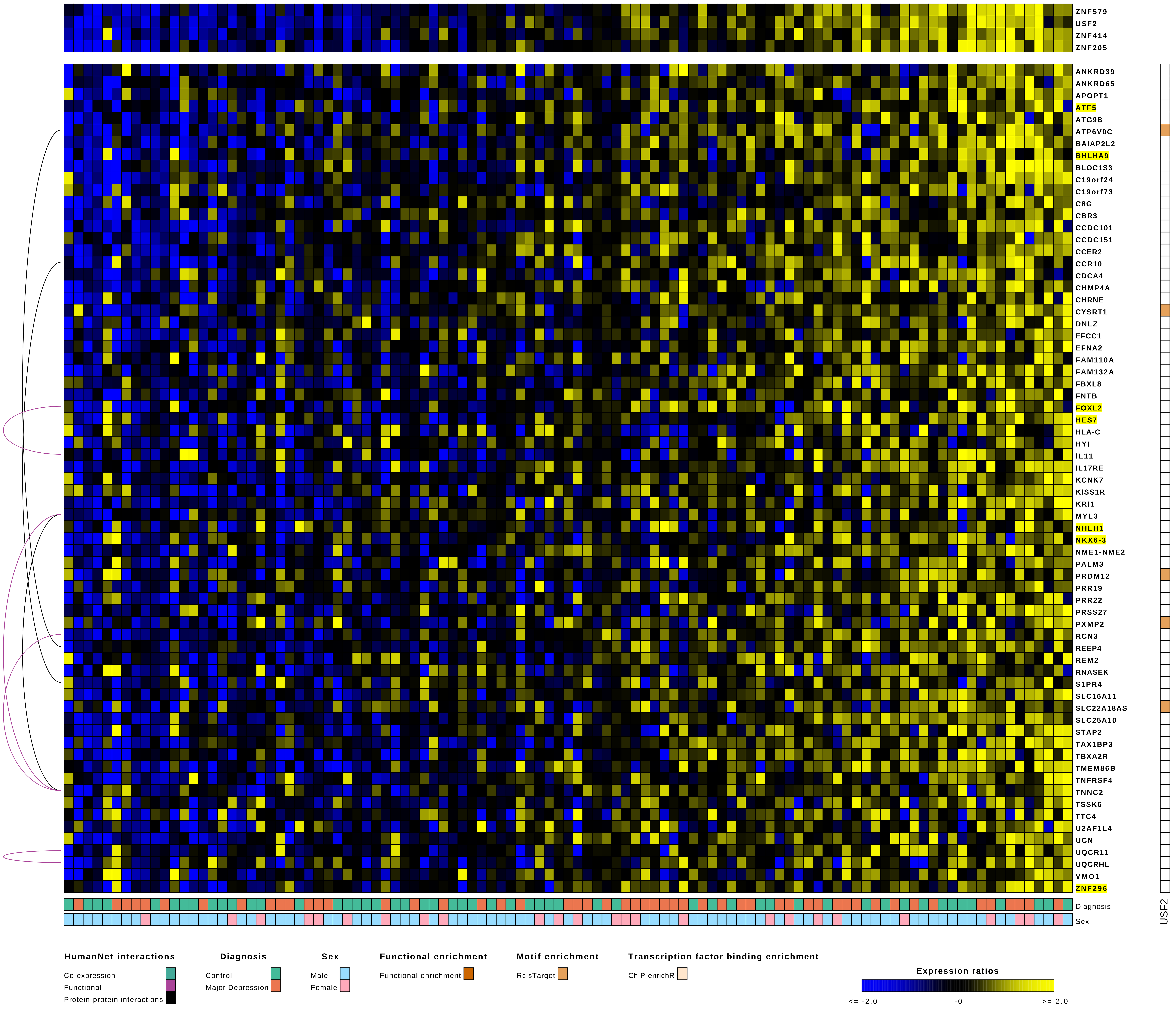

ZNF579
USF2
ZNF414
ZNF205
ANKRD39
ANKRD65
APOPT1
ATF5
ATG9B
ATP6V0C
BAIAP2L2
BHLHA9
BLOC1S3
C19orf24
C19orf73
C8G
CBR3
CCDC101
CCDC151
CCER2
CCR10
CDCA4
CHMP4A
CHRNE
CYSRT1
DNLZ
EFCC1
EFNA2
FAM110A
FAM132A
FBXL8
FNTB
FOXL2
HES7
HLA-C
HYI
IL11
IL17RE
KCNK7
KISS1R
KRI1
MYL3
NHLH1
NKX6-3
NME1-NME2
PALM3
PRDM12
PRR19
PRR22
PRSS27
PXMP2
RCN3
REEP4
REM2
RNASEK
S1PR4
SLC16A11
SLC22A18AS
SLC25A10
STAP2
TAX1BP3
TBXA2R
TMEM86B
TNFRSF4
TNNC2
TSSK6
TTC4
U2AF1L4
UCN
UQCR11
UQCRHL
VMO1
ZNF296
Diagnosis
Sex
USF2
HumanNet interactions
Co-expression
Functional
Protein-protein interactions
Diagnosis
Control
Major Depression
Sex
Male
Female
Functional enrichment
Functional enrichment
Motif enrichment
RcisTarget
Transcription factor binding enrichment
ChIP-enrichR
Expression ratios
<= -2.0
-0
>= 2.0

Module 112

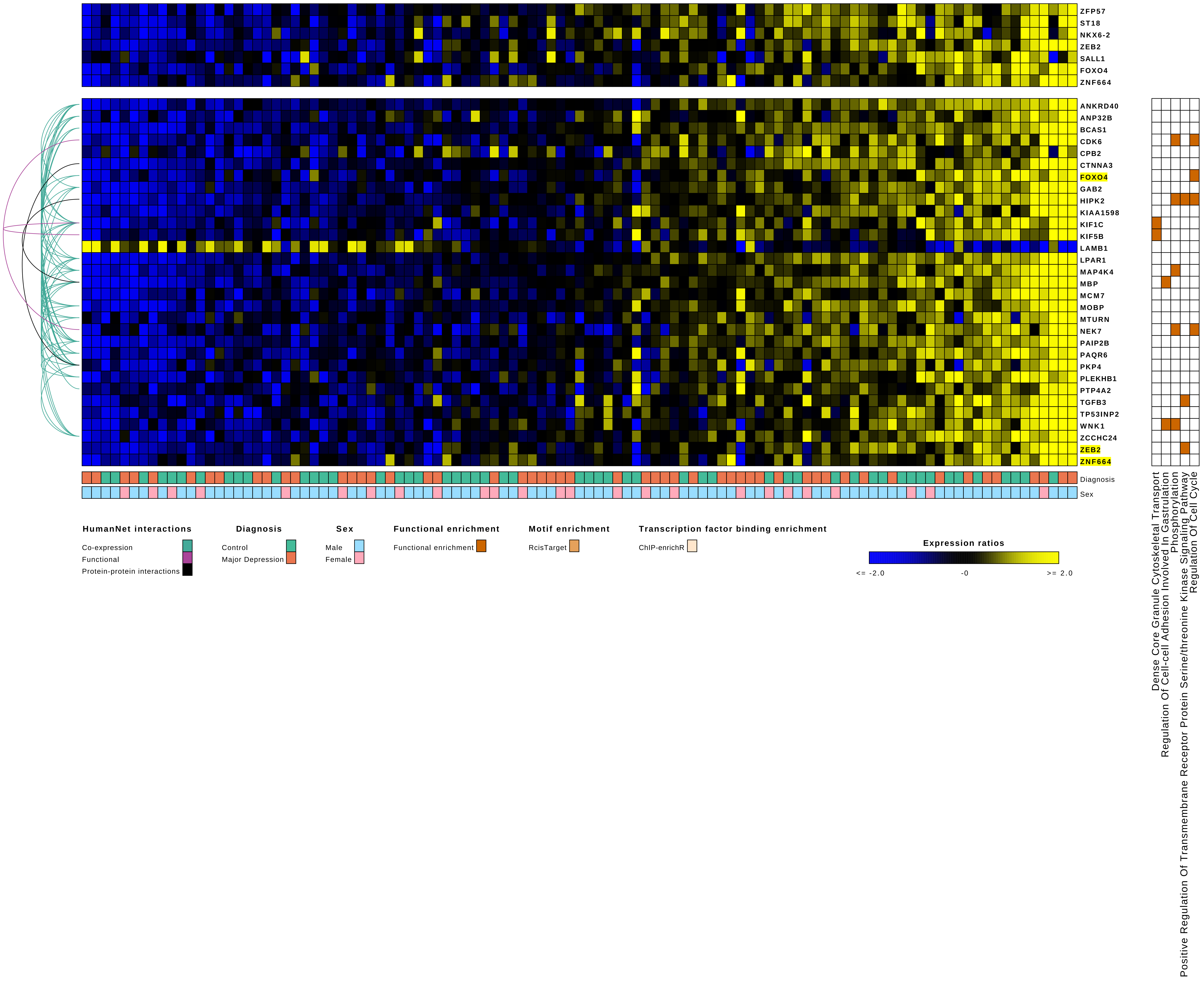

ZFP57
ST18
NKX6-2
ZEB2
SALL1
FOXO4
ZNF664
ANKRD40
ANP32B
BCAS1
CDK6
CPB2
CTNNA3
FOXO4
GAB2
HIPK2
KIAA1598
KIF1C
KIF5B
LAMB1
LPAR1
MAP4K4
MBP
MCM7
MOBP
MTURN
NEK7
PAIP2B
PAQR6
PKP4
PLEKHB1
PTP4A2
TGFB3
TP53INP2
WNK1
ZCCHC24
ZEB2
ZNF664
Diagnosis
Sex
Dense Core Granule Cytoskeletal Transport
Regulation Of Cell-cell Adhesion Involved In Gastrulation
Phosphorylation
Positive Regulation Of Transmembrane Receptor Protein Serine/threonine Kinase Signaling Pathway
Regulation Of Cell Cycle
HumanNet interactions
Co-expression
Functional
Protein-protein interactions
Diagnosis
Control
Major Depression
Sex
Male
Female
Functional enrichment
Functional enrichment
Motif enrichment
RcisTarget
Transcription factor binding enrichment
ChIP-enrichR
Expression ratios
<= -2.0
-0
>= 2.0

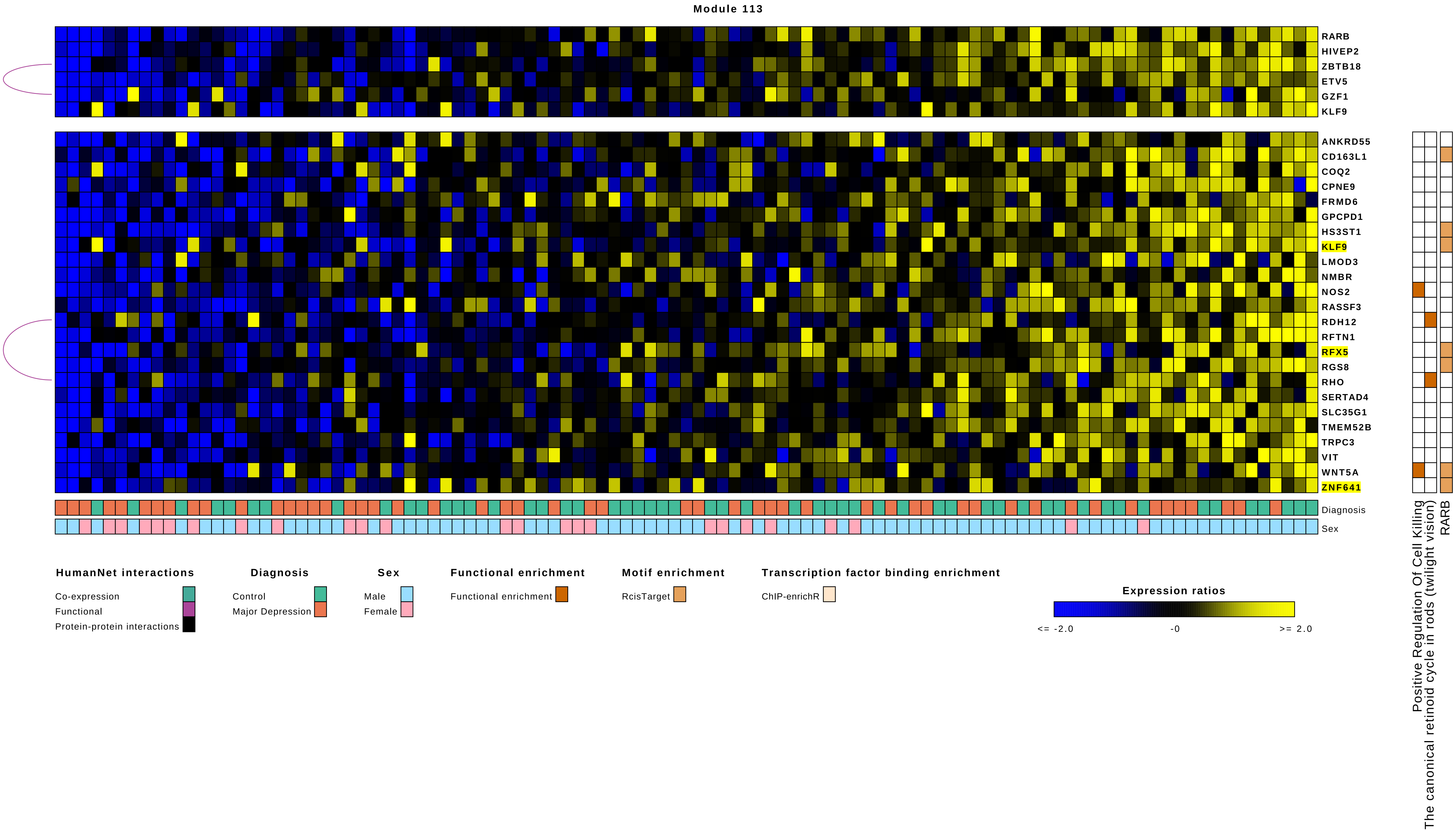
Module 113
RARB
HIVEP2
ZBTB18
ETV5
GZF1
KLF9
ANKRD55
CD163L1
COQ2
CPNE9
FRMD6
GPCPD1
HS3ST1
KLF9
LMOD3
NMBR
NOS2
RASSF3
RDH12
RFTN1
RFX5
RGS8
RHO
SERTAD4
SLC35G1
TMEM52B
TRPC3
VIT
WNT5A
ZNF641
Diagnosis
Sex
Positive Regulation Of Cell Killing
The canonical retinoid cycle in rods (twilight vision)
RARB
HumanNet interactions
Co-expression
Functional
Protein-protein interactions
Diagnosis
Control
Major Depression
Sex
Male
Female
Functional enrichment
Functional enrichment
Motif enrichment
RcisTarget
Transcription factor binding enrichment
ChIP-enrichR
Expression ratios
<= -2.0
-0
>= 2.0

Module 114

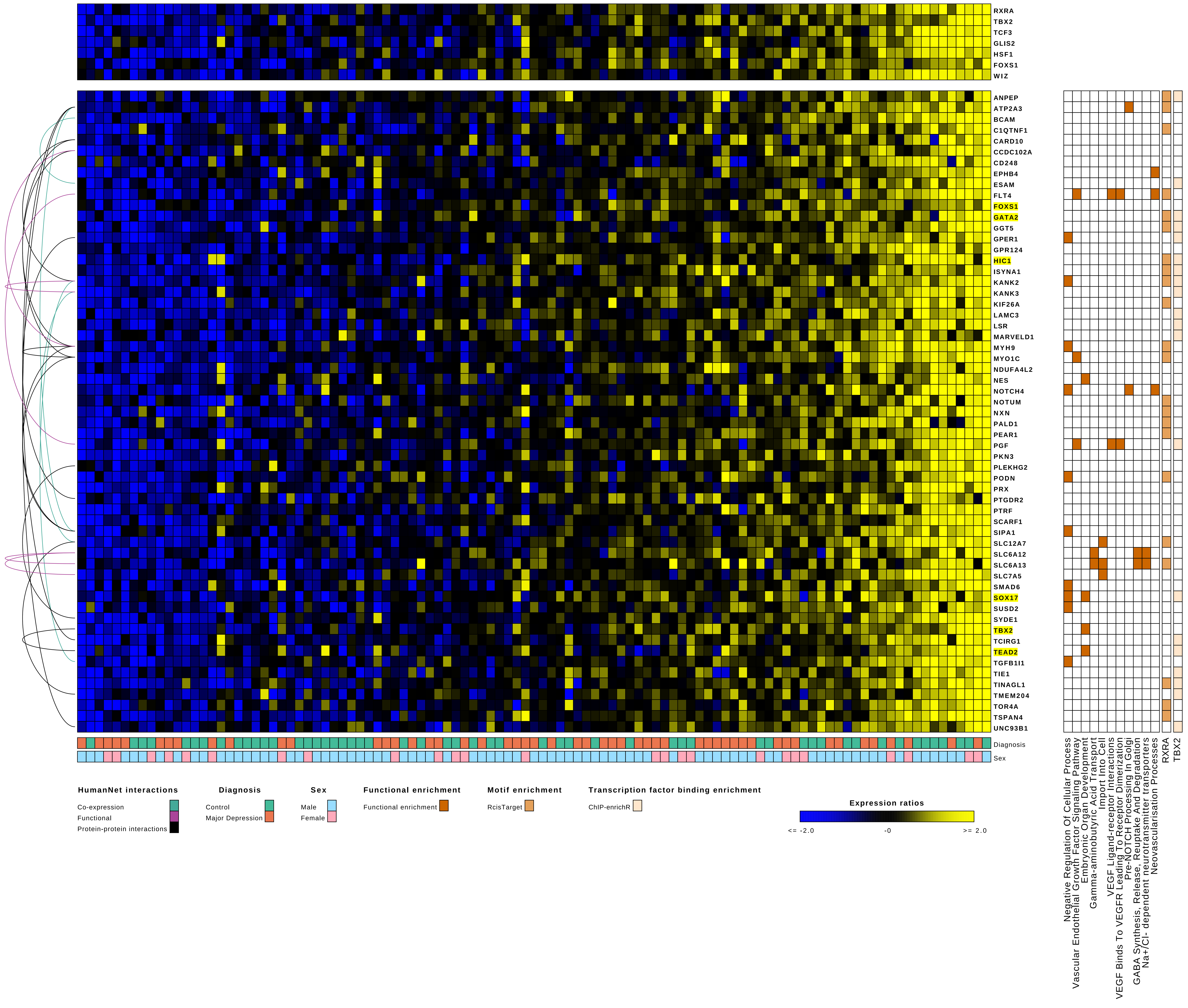

RXRA
TBX2
TCF3
GLIS2
HSF1
FOXS1
WIZ
ANPEP
ATP2A3
BCAM
C1QTNF1
CARD10
CCDC102A
CD248
EPHB4
ESAM
FLT4
FOXS1
GATA2
GGT5
GPER1
GPR124
HIC1
ISYNA1
KANK2
KANK3
KIF26A
LAMC3
LSR
MARVELD1
MYH9
MYO1C
NDUFA4L2
NES
NOTCH4
NOTUM
NXN
PALD1
PEAR1
PGF
PKN3
PLEKHG2
PODN
PRX
PTGDR2
PTRF
SCARF1
SIPA1
SLC12A7
SLC6A12
SLC6A13
SLC7A5
SMAD6
SOX17
SUSD2
SYDE1
TBX2
TCIRG1
TEAD2
TGFB1I1
TIE1
TINAGL1
TMEM204
TOR4A
TSPAN4
UNC93B1
Diagnosis
Sex
Negative Regulation Of Cellular Process
Vascular Endothelial Growth Factor Signaling Pathway
Embryonic Organ Development
Gamma-aminobutyric Acid Transport
Import Into Cell
VEGF Ligand-receptor Interactions
VEGF Binds To VEGFR Leading To Receptor Dimerization
Pre-NOTCH Processing In Golgi
GABA Synthesis, Release, Reuptake And Degradation
Na+/Cl- dependent neurotransmitter transporters
Neovascularisation Processes
RXRA
TBX2
HumanNet interactions
Co-expression
Functional
Protein-protein interactions
Diagnosis
Control
Major Depression
Sex
Male
Female
Functional enrichment
Functional enrichment
Motif enrichment
RcisTarget
Transcription factor binding enrichment
ChIP-enrichR
Expression ratios
<= -2.0
-0
>= 2.0

Module 115

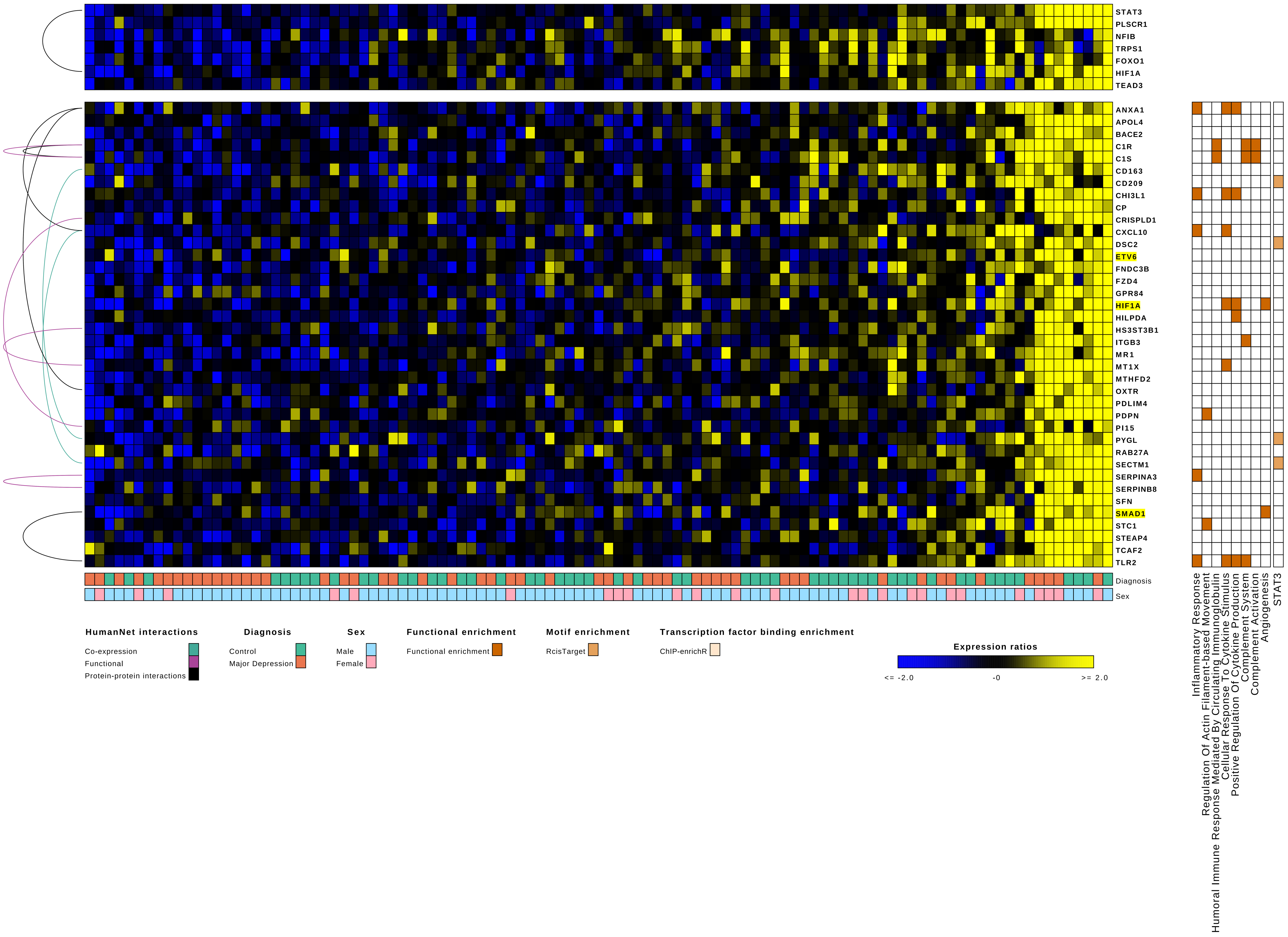
STAT3
PLSCR1
NFIB
TRPS1
FOXO1
HIF1A
TEAD3
ANXA1
APOL4
BACE2
C1R
C1S
CD163
CD209
CHI3L1
CP
CRISPLD1
CXCL10
DSC2
ETV6
FNDC3B
FZD4
GPR84
HIF1A
HILPDA
HS3ST3B1
ITGB3
MR1
MT1X
MTHFD2
OXTR
PDLIM4
PDPN
PI15
PYGL
RAB27A
SECTM1
SERPINA3
SERPINB8
SFN
SMAD1
STC1
STEAP4
TCAF2
TLR2
Diagnosis
Sex
Inflammatory Response
Regulation Of Actin Filament-based Movement
Humoral Immune Response Mediated By Circulating Immunoglobulin
Cellular Response To Cytokine Stimulus
Positive Regulation Of Cytokine Production
Complement System
Complement Activation
Angiogenesis
STAT3
HumanNet interactions
Co-expression
Functional
Protein-protein interactions
Diagnosis
Control
Major Depression
Sex
Male
Female
Functional enrichment
Functional enrichment
Motif enrichment
RcisTarget
Transcription factor binding enrichment
ChIP-enrichR
Expression ratios
<= -2.0
-0
>= 2.0

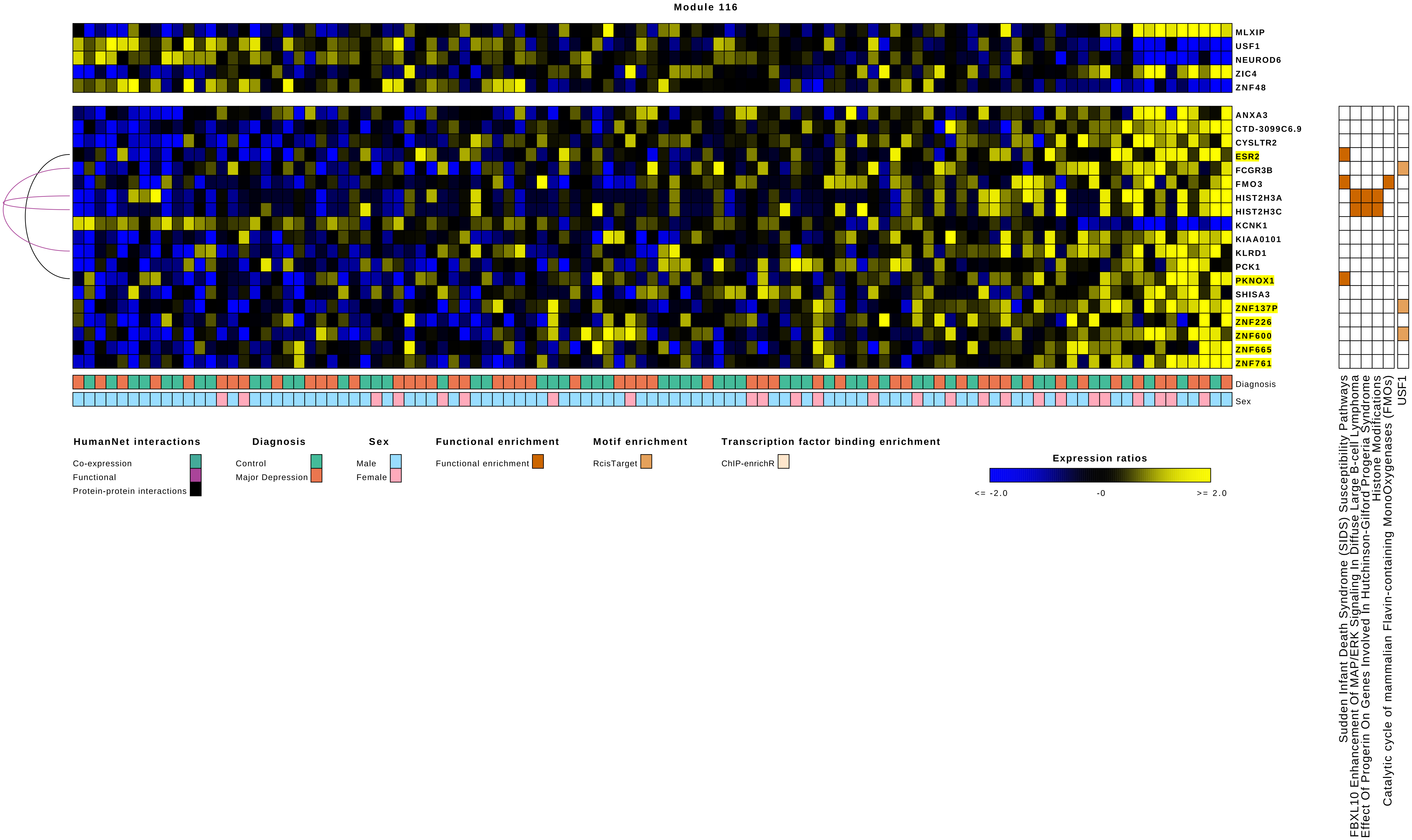
Module 116
MLXIP
USF1
NEUROD6
ZIC4
ZNF48
ANXA3
CTD-3099C6.9
CYSLTR2
ESR2
FCGR3B
FMO3
HIST2H3A
HIST2H3C
KCNK1
KIAA0101
KLRD1
PCK1
PKNOX1
SHISA3
ZNF137P
ZNF226
ZNF600
ZNF665
ZNF761
Diagnosis
Sex
Sudden Infant Death Syndrome (SIDS) Susceptibility Pathways
FBXL10 Enhancement Of MAP/ERK Signaling In Diffuse Large B-cell Lymphoma
Effect Of Progerin On Genes Involved In Hutchinson-Gilford Progeria Syndrome
Histone Modifications
Catalytic cycle of mammalian Flavin-containing MonoOxygenases (FMOs)
USF1
HumanNet interactions
Co-expression
Functional
Protein-protein interactions
Diagnosis
Control
Major Depression
Sex
Male
Female
Functional enrichment
Functional enrichment
Motif enrichment
RcisTarget
Transcription factor binding enrichment
ChIP-enrichR
Expression ratios
<= -2.0
-0
>= 2.0

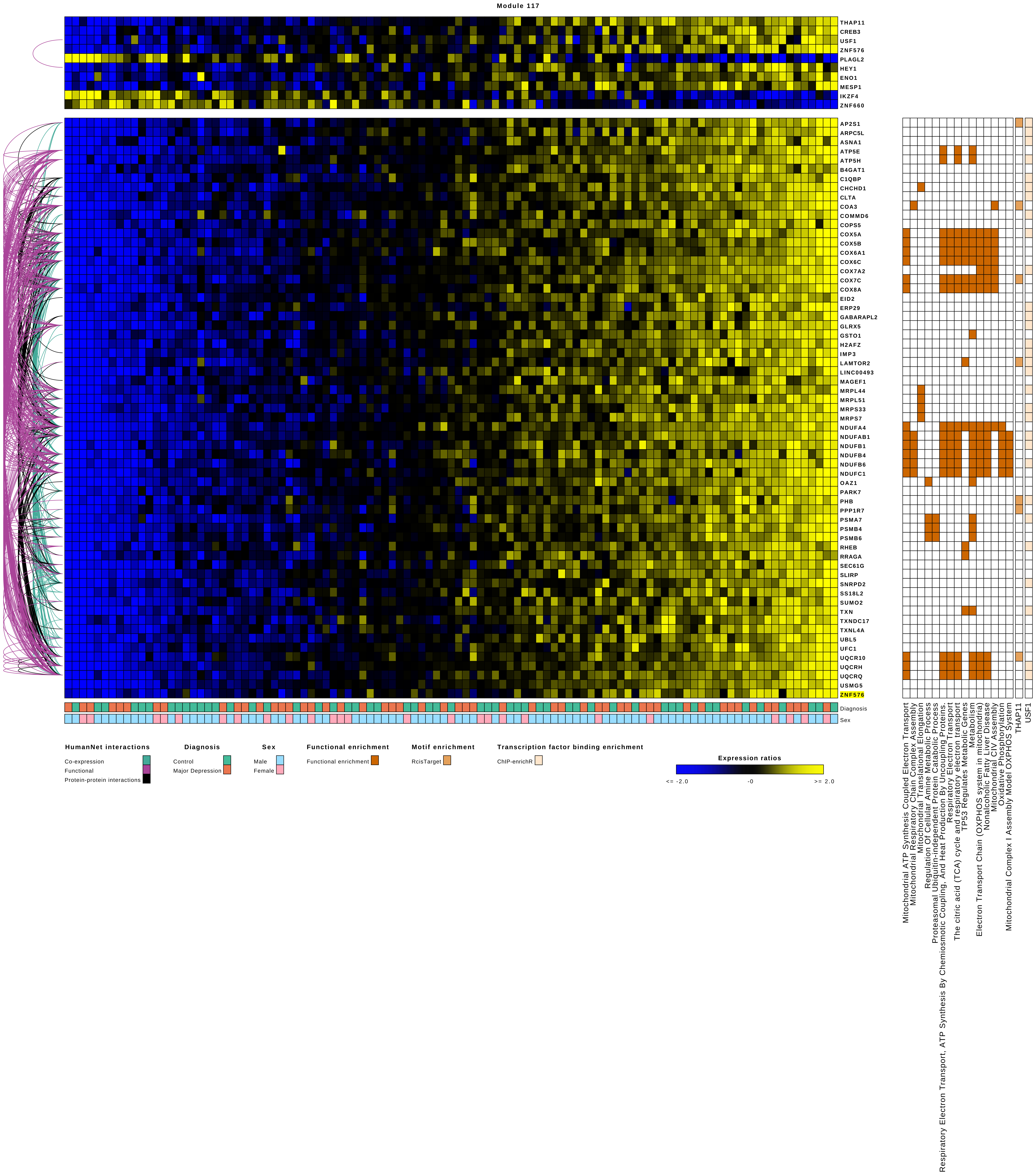
Module 117
THAP11
CREB3
USF1
ZNF576
PLAGL2
HEY1
ENO1
MESP1
IKZF4
ZNF660
AP2S1
ARPC5L
ASNA1
ATP5E
ATP5H
B4GAT1
C1QBP
CHCHD1
CLTA
COA3
COMMD6
COPS5
COX5A
COX5B
COX6A1
COX6C
COX7A2
COX7C
COX8A
EID2
ERP29
GABARAPL2
GLRX5
GSTO1
H2AFZ
IMP3
LAMTOR2
LINC00493
MAGEF1
MRPL44
MRPL51
MRPS33
MRPS7
NDUFA4
NDUFAB1
NDUFB1
NDUFB4
NDUFB6
NDUFC1
OAZ1
PARK7
PHB
PPP1R7
PSMA7
PSMB4
PSMB6
RHEB
RRAGA
SEC61G
SLIRP
SNRPD2
SS18L2
SUMO2
TXN
TXNDC17
TXNL4A
UBL5
UFC1
UQCR10
UQCRH
UQCRQ
USMG5
ZNF576
Diagnosis
Sex
Mitochondrial ATP Synthesis Coupled Electron Transport
Mitochondrial Respiratory Chain Complex Assembly
Mitochondrial Translational Elongation
Regulation Of Cellular Amine Metabolic Process
Proteasomal Ubiquitin-independent Protein Catabolic Process
Respiratory Electron Transport, ATP Synthesis By Chemiosmotic Coupling, And Heat Production By Uncoupling Proteins.
Respiratory Electron Transport
The citric acid (TCA) cycle and respiratory electron transport
TP53 Regulates Metabolic Genes
Metabolism
Electron Transport Chain (OXPHOS system in mitochondria)
Nonalcoholic Fatty Liver Disease
Mitochondrial CIV Assembly
Oxidative Phosphorylation
Mitochondrial Complex I Assembly Model OXPHOS System
THAP11
USF1
HumanNet interactions
Co-expression
Functional
Protein-protein interactions
Diagnosis
Control
Major Depression
Sex
Male
Female
Functional enrichment
Functional enrichment
Motif enrichment
RcisTarget
Transcription factor binding enrichment
ChIP-enrichR
Expression ratios
<= -2.0
-0
>= 2.0

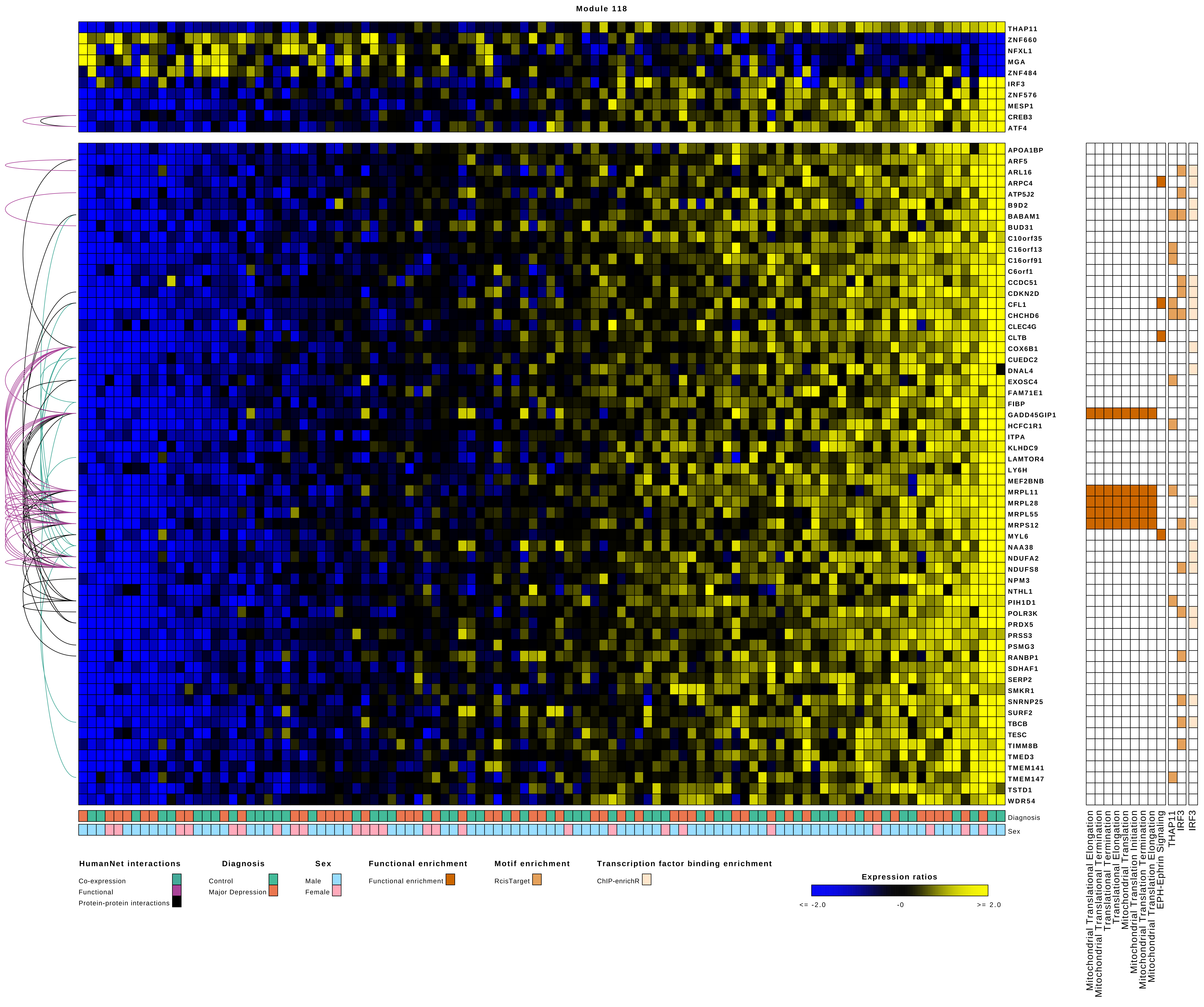

Module 118
THAP11
ZNF660
NFXL1
MGA
ZNF484
IRF3
ZNF576
MESP1
CREB3
ATF4
APOA1BP
ARF5
ARL16
ARPC4
ATP5J2
B9D2
BABAM1
BUD31
C10orf35
C16orf13
C16orf91
C6orf1
CCDC51
CDKN2D
CFL1
CHCHD6
CLEC4G
CLTB
COX6B1
CUEDC2
DNAL4
EXOSC4
FAM71E1
FIBP
GADD45GIP1
HCFC1R1
ITPA
KLHDC9
LAMTOR4
LY6H
MEF2BNB
MRPL11
MRPL28
MRPL55
MRPS12
MYL6
NAA38
NDUFA2
NDUFS8
NPM3
NTHL1
PIH1D1
POLR3K
PRDX5
PRSS3
PSMG3
RANBP1
SDHAF1
SERP2
SMKR1
SNRNP25
SURF2
TBCB
TESC
TIMM8B
TMED3
TMEM141
TMEM147
TSTD1
WDR54
Diagnosis
Sex
Mitochondrial Translational Elongation
Mitochondrial Translational Termination
Translational Termination
Translational Elongation
Mitochondrial Translation
Mitochondrial Translation Initiation
Mitochondrial Translation Termination
Mitochondrial Translation Elongation
EPH-Ephrin Signaling
THAP11
IRF3
IRF3
HumanNet interactions
Co-expression
Functional
Protein-protein interactions
Diagnosis
Control
Major Depression
Sex
Male
Female
Functional enrichment
Functional enrichment
Motif enrichment
RcisTarget
Transcription factor binding enrichment
ChIP-enrichR
Expression ratios
<= -2.0
-0
>= 2.0

Module 119

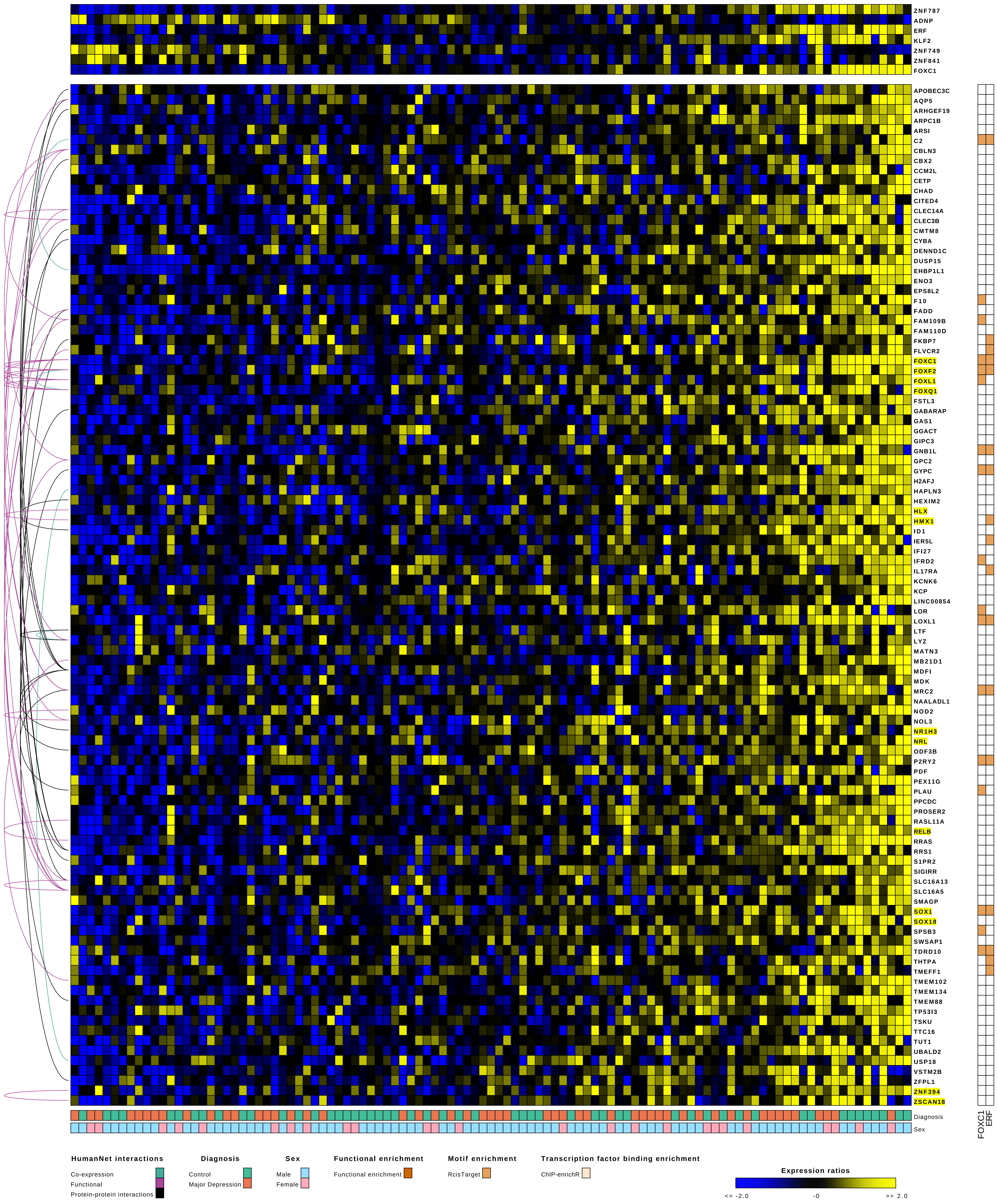

ZNF787
ADNP
ERF
KLF2
ZNF749
ZNF841
FOXC1
APOBEC3C
AQP5
ARHGEF19
ARPC1B
ARSI
C2
CBLN3
CBX2
CCM2L
CETP
CHAD
CITED4
CLEC14A
CLEC3B
CMTM8
CYBA
DENND1C
DUSP15
EHBP1L1
ENO3
EPS8L2
F10
FADD
FAM109B
FAM110D
FKBP7
FLVCR2
FOXC1
FOXF2
FOXL1
FOXQ1
FSTL3
GABARAP
GAS1
GGACT
GIPC3
GNB1L
GPC2
GYPC
H2AFJ
HAPLN3
HEXIM2
HLX
HMX1
ID1
IER5L
IFI27
IFRD2
IL17RA
KCNK6
KCP
LINC00854
LOR
LOXL1
LTF
LYZ
MATN3
MB21D1
MDFI
MDK
MRC2
NAALADL1
NOD2
NOL3
NR1H3
NRL
ODF3B
P2RY2
PDF
PEX11G
PLAU
PPCDC
PROSER2
RASL11A
RELB
RRAS
RRS1
S1PR2
SIGIRR
SLC16A13
SLC16A5
SMAGP
SOX1
SOX18
SPSB3
SWSAP1
TDRD10
THTPA
TMEFF1
TMEM102
TMEM134
TMEM88
TP53I3
TSKU
TTC16
TUT1
UBALD2
USP18
VSTM2B
ZFPL1
ZNF394
ZSCAN18
Diagnosis
Sex
FOXC1
ERF
HumanNet interactions
Co-expression
Functional
Protein-protein interactions
Diagnosis
Control
Major Depression
Sex
Male
Female
Functional enrichment
Functional enrichment
Motif enrichment
RcisTarget
Transcription factor binding enrichment
ChIP-enrichR
Expression ratios
<= -2.0
-0
>= 2.0

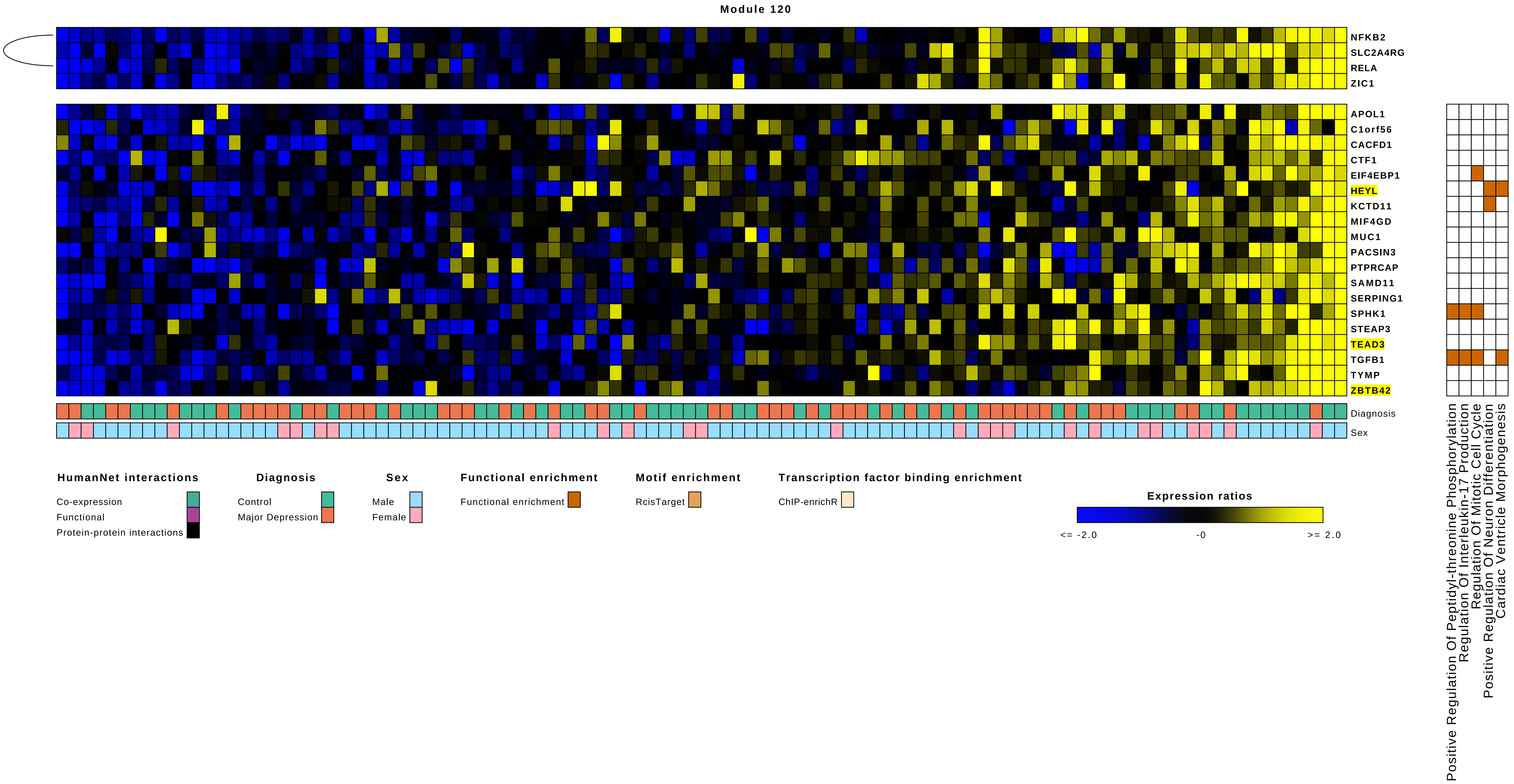
Module 120
NFKB2
SLC2A4RG
RELA
ZIC1
APOL1
C1orf56
CACFD1
CTF1
EIF4EBP1
HEYL
KCTD11
MIF4GD
MUC1
PACSIN3
PTPRCAP
SAMD11
SERPING1
SPHK1
STEAP3
TEAD3
TGFB1
TYMP
ZBTB42
Diagnosis
Sex
Positive Regulation Of Peptidyl-threonine Phosphorylation
Regulation Of Interleukin-17 Production
Regulation Of Mitotic Cell Cycle
Positive Regulation Of Neuron Differentiation
Cardiac Ventricle Morphogenesis
HumanNet interactions
Co-expression
Functional
Protein-protein interactions
Diagnosis
Control
Major Depression
Sex
Male
Female
Functional enrichment
Functional enrichment
Motif enrichment
RcisTarget
Transcription factor binding enrichment
ChIP-enrichR
Expression ratios
<= -2.0
-0
>= 2.0

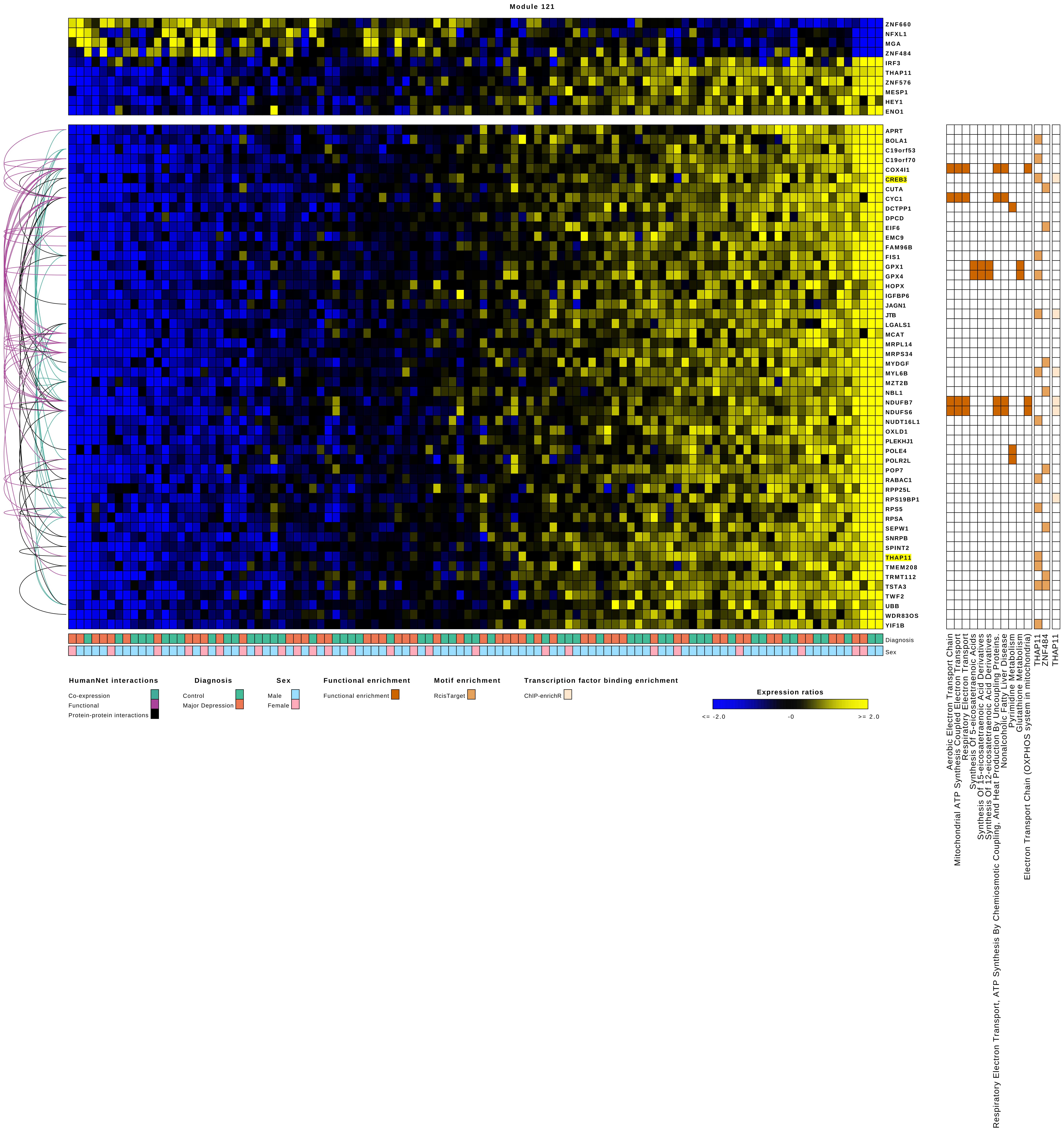
Module 121
ZNF660
NFXL1
MGA
ZNF484
IRF3
THAP11
ZNF576
MESP1
HEY1
ENO1
APRT
BOLA1
C19orf53
C19orf70
COX4I1
CREB3
CUTA
CYC1
DCTPP1
DPCD
EIF6
EMC9
FAM96B
FIS1
GPX1
GPX4
HOPX
IGFBP6
JAGN1
JTB
LGALS1
MCAT
MRPL14
MRPS34
MYDGF
MYL6B
MZT2B
NBL1
NDUFB7
NDUFS6
NUDT16L1
OXLD1
PLEKHJ1
POLE4
POLR2L
POP7
RABAC1
RPP25L
RPS19BP1
RPS5
RPSA
SEPW1
SNRPB
SPINT2
THAP11
TMEM208
TRMT112
TSTA3
TWF2
UBB
WDR83OS
YIF1B
Diagnosis
Sex
HumanNet interactions
Co-expression
Functional
Protein-protein interactions
Diagnosis
Control
Major Depression
Sex
Male
Female
Functional enrichment
Functional enrichment
Motif enrichment
RcisTarget
Transcription factor binding enrichment
ChIP-enrichR
Expression ratios
<= -2.0
-0
>= 2.0
Aerobic Electron Transport Chain
Mitochondrial ATP Synthesis Coupled Electron Transport
Respiratory Electron Transport
Synthesis Of 5-eicosatetraenoic Acids
Synthesis Of 15-eicosatetraenoic Acid Derivatives
Synthesis Of 12-eicosatetraenoic Acid Derivatives
Respiratory Electron Transport, ATP Synthesis By Chemiosmotic Coupling, And Heat Production By Uncoupling Proteins.
Nonalcoholic Fatty Liver Disease
Pyrimidine Metabolism
Glutathione Metabolism
Electron Transport Chain (OXPHOS system in mitochondria)
THAP11
ZNF484
THAP11

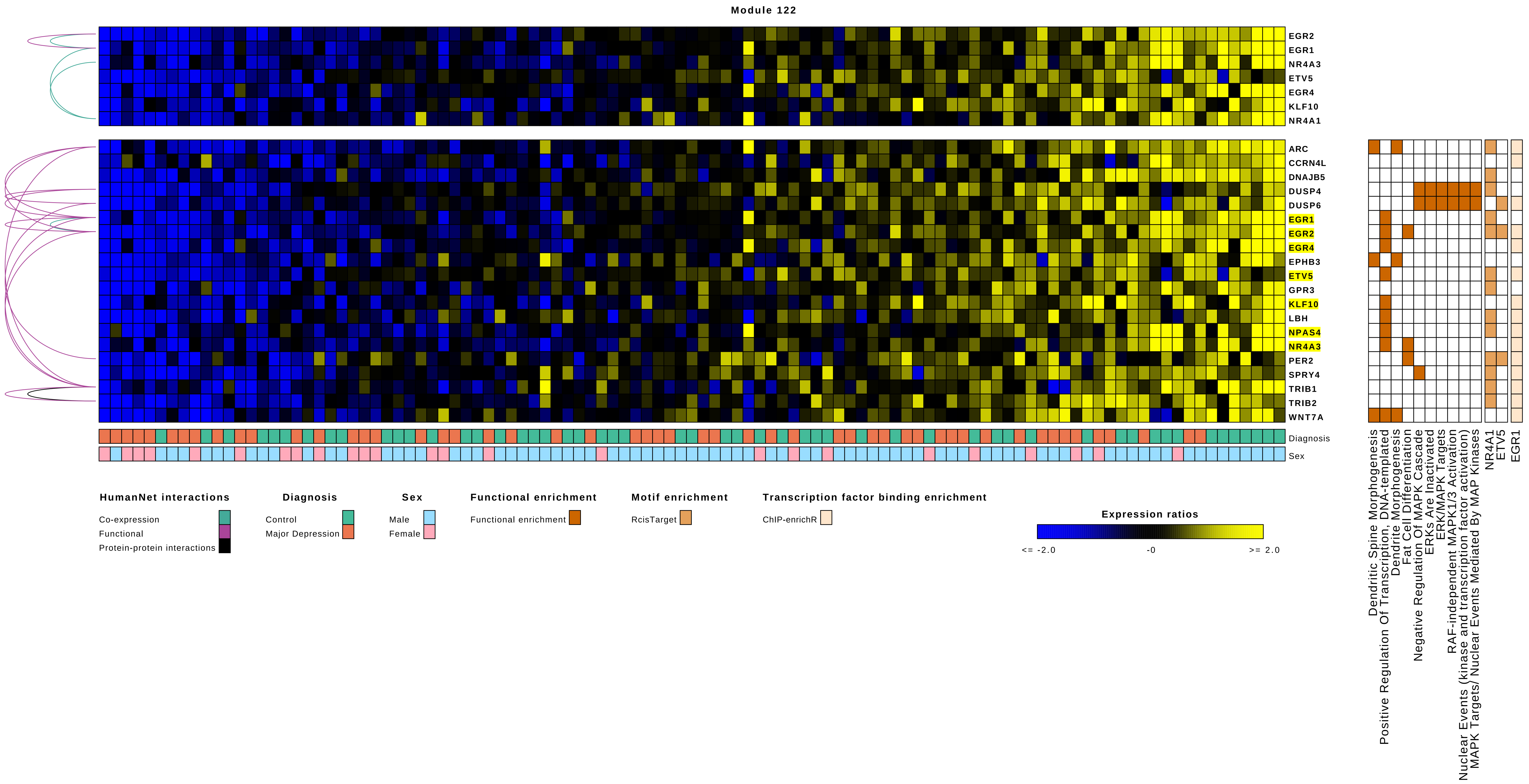
Module 122
EGR2
EGR1
NR4A3
ETV5
EGR4
KLF10
NR4A1
ARC
CCRN4L
DNAJB5
DUSP4
DUSP6
EGR1
EGR2
EGR4
EPHB3
ETV5
GPR3
KLF10
LBH
NPAS4
NR4A3
PER2
SPRY4
TRIB1
TRIB2
WNT7A
Diagnosis
Sex
Dendritic Spine Morphogenesis
Positive Regulation Of Transcription, DNA-templated
Dendrite Morphogenesis
Fat Cell Differentiation
Negative Regulation Of MAPK Cascade
ERKs Are Inactivated
ERK/MAPK Targets
RAF-independent MAPK1/3 Activation
Nuclear Events (kinase and transcription factor activation)
MAPK Targets/ Nuclear Events Mediated By MAP Kinases
NR4A1
ETV5
EGR1
HumanNet interactions
Co-expression
Functional
Protein-protein interactions
Diagnosis
Control
Major Depression
Sex
Male
Female
Functional enrichment
Functional enrichment
Motif enrichment
RcisTarget
Transcription factor binding enrichment
ChIP-enrichR
Expression ratios
<= -2.0
-0
>= 2.0

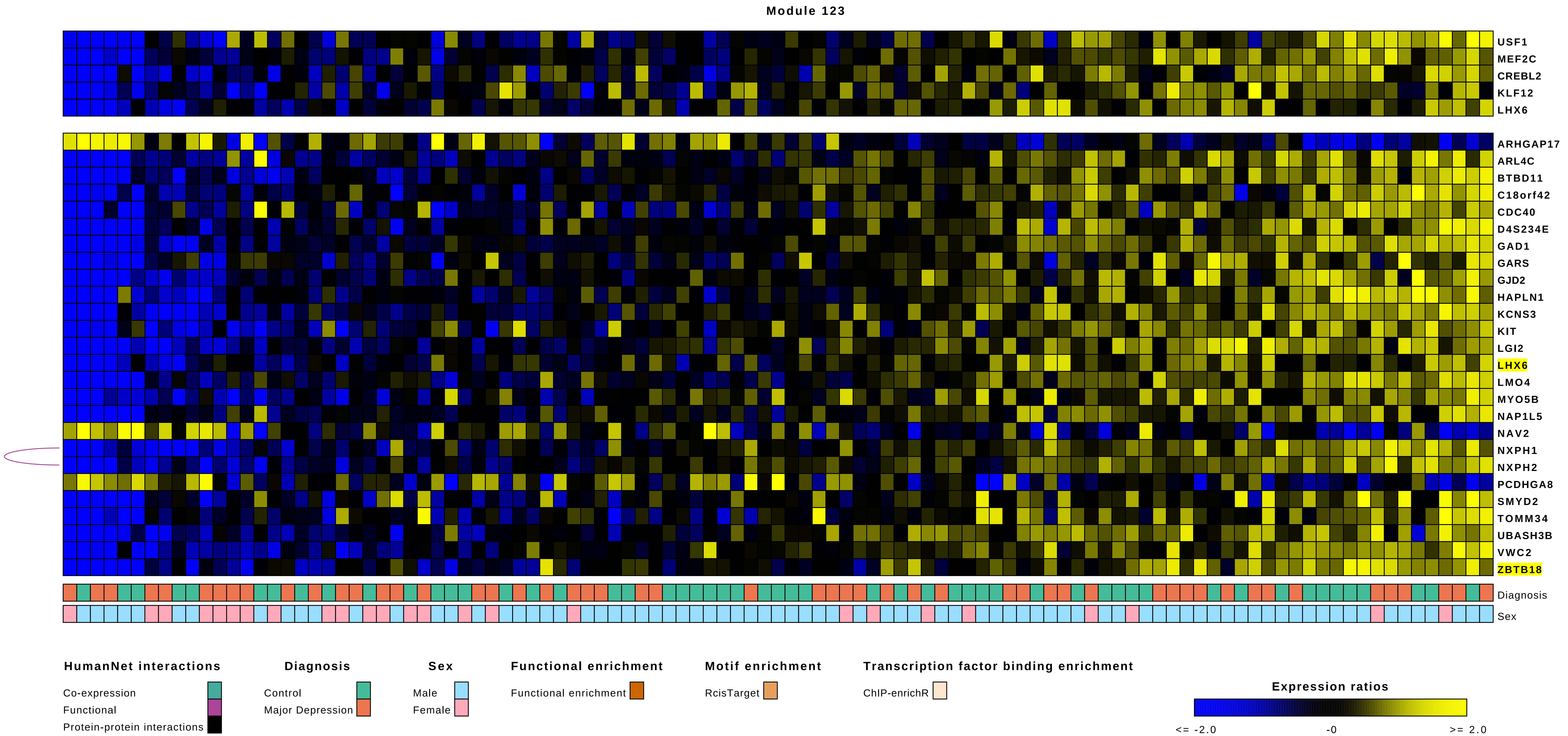

Module 123
USF1
MEF2C
CREBL2
KLF12
LHX6
ARHGAP17
ARL4C
BTBD11
C18orf42
CDC40
D4S234E
GAD1
GARS
GJD2
HAPLN1
KCNS3
KIT
LGI2
LHX6
LMO4
MYO5B
NAP1L5
NAV2
NXPH1
NXPH2
PCDHGA8
SMYD2
TOMM34
UBASH3B
VWC2
ZBTB18
Diagnosis
Sex
HumanNet interactions
Co-expression
Functional
Protein-protein interactions
Diagnosis
Control
Major Depression
Sex
Male
Female
Functional enrichment
Functional enrichment
Motif enrichment
RcisTarget
Transcription factor binding enrichment
ChIP-enrichR
Expression ratios
<= -2.0
-0
>= 2.0

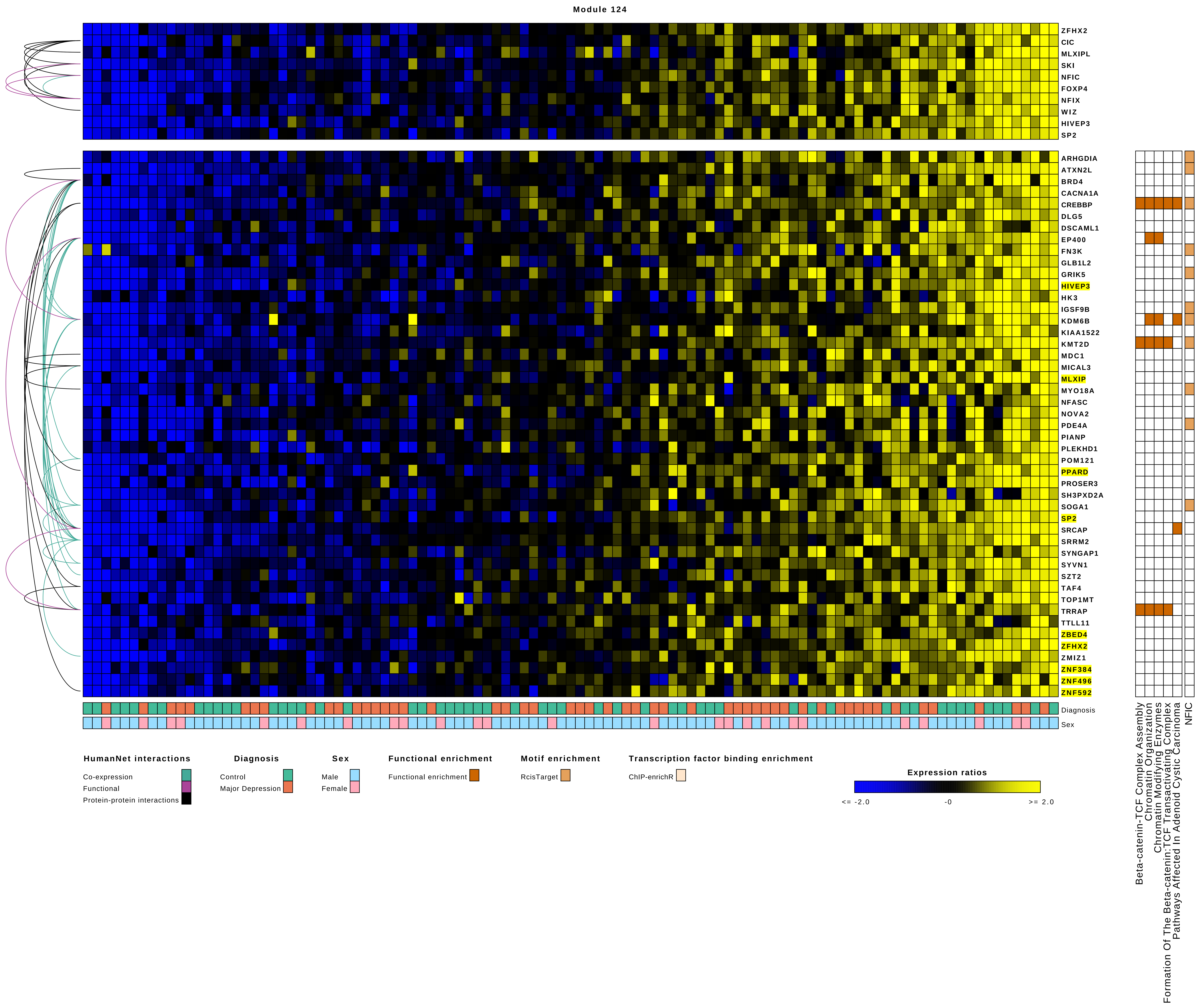
Module 124
ZFHX2
CIC
MLXIPL
SKI
NFIC
FOXP4
NFIX
WIZ
HIVEP3
SP2
ARHGDIA
ATXN2L
BRD4
CACNA1A
CREBBP
DLG5
DSCAML1
EP400
FN3K
GLB1L2
GRIK5
HIVEP3
HK3
IGSF9B
KDM6B
KIAA1522
KMT2D
MDC1
MICAL3
MLXIP
MYO18A
NFASC
NOVA2
PDE4A
PIANP
PLEKHD1
POM121
PPARD
PROSER3
SH3PXD2A
SOGA1
SP2
SRCAP
SRRM2
SYNGAP1
SYVN1
SZT2
TAF4
TOP1MT
TRRAP
TTLL11
ZBED4
ZFHX2
ZMIZ1
ZNF384
ZNF496
ZNF592
Diagnosis
Sex
Beta-catenin-TCF Complex Assembly
Chromatin Organization
Chromatin Modifying Enzymes
Formation Of The Beta-catenin:TCF Transactivating Complex
Pathways Affected In Adenoid Cystic Carcinoma
NFIC
HumanNet interactions
Co-expression
Functional
Protein-protein interactions
Diagnosis
Control
Major Depression
Sex
Male
Female
Functional enrichment
Functional enrichment
Motif enrichment
RcisTarget
Transcription factor binding enrichment
ChIP-enrichR
Expression ratios
<= -2.0
-0
>= 2.0

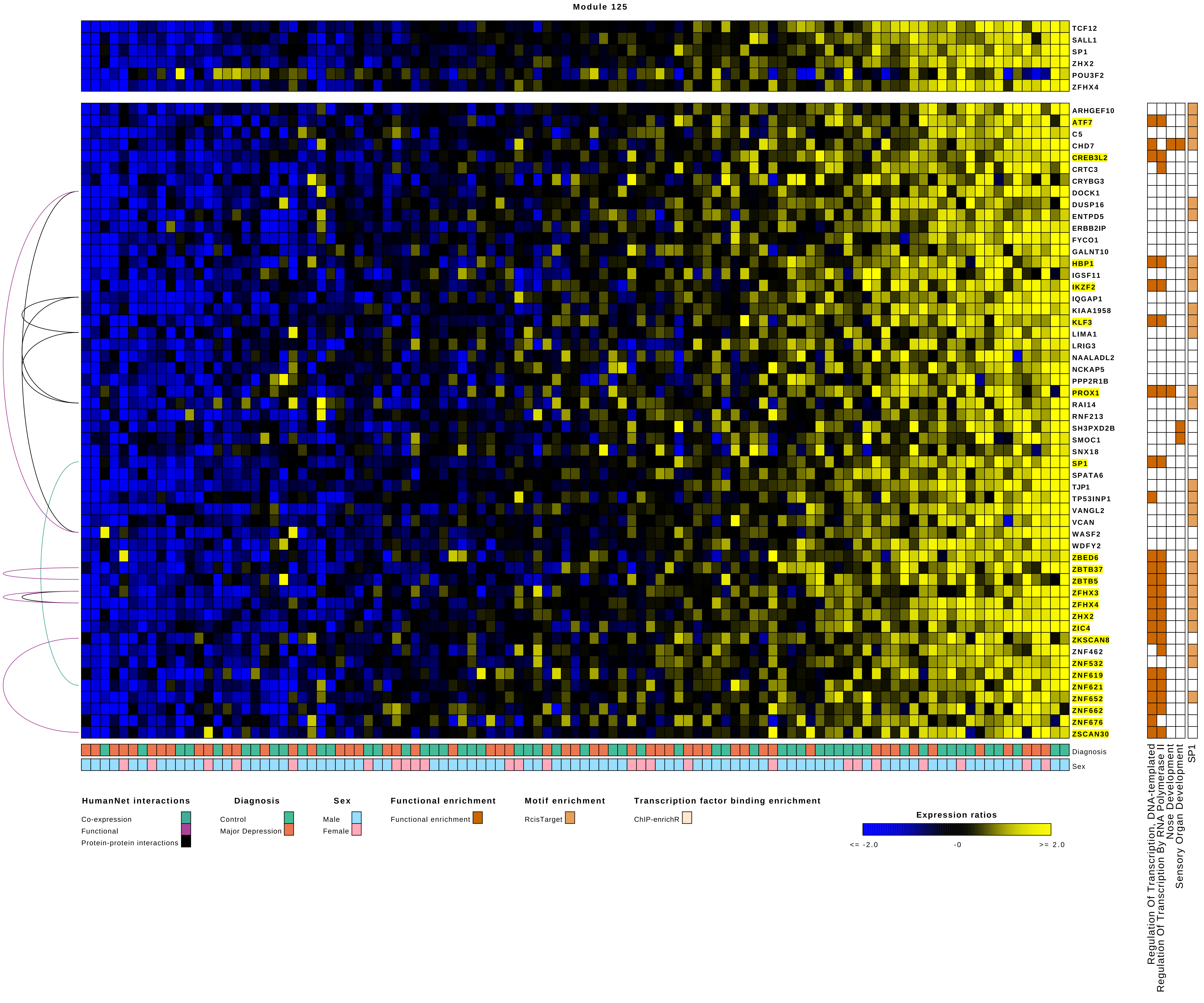
Module 125
TCF12
SALL1
SP1
ZHX2
POU3F2
ZFHX4
ARHGEF10
ATF7
C5
CHD7
CREB3L2
CRTC3
CRYBG3
DOCK1
DUSP16
ENTPD5
ERBB2IP
FYCO1
GALNT10
HBP1
IGSF11
IKZF2
IQGAP1
KIAA1958
KLF3
LIMA1
LRIG3
NAALADL2
NCKAP5
PPP2R1B
PROX1
RAI14
RNF213
SH3PXD2B
SMOC1
SNX18
SP1
SPATA6
TJP1
TP53INP1
VANGL2
VCAN
WASF2
WDFY2
ZBED6
ZBTB37
ZBTB5
ZFHX3
ZFHX4
ZHX2
ZIC4
ZKSCAN8
ZNF462
ZNF532
ZNF619
ZNF621
ZNF652
ZNF662
ZNF676
ZSCAN30
Diagnosis
Sex
Regulation Of Transcription, DNA-templated
Regulation Of Transcription By RNA Polymerase II
Nose Development
Sensory Organ Development
SP1
HumanNet interactions
Co-expression
Functional
Protein-protein interactions
Diagnosis
Control
Major Depression
Sex
Male
Female
Functional enrichment
Functional enrichment
Motif enrichment
RcisTarget
Transcription factor binding enrichment
ChIP-enrichR
Expression ratios
<= -2.0
-0
>= 2.0

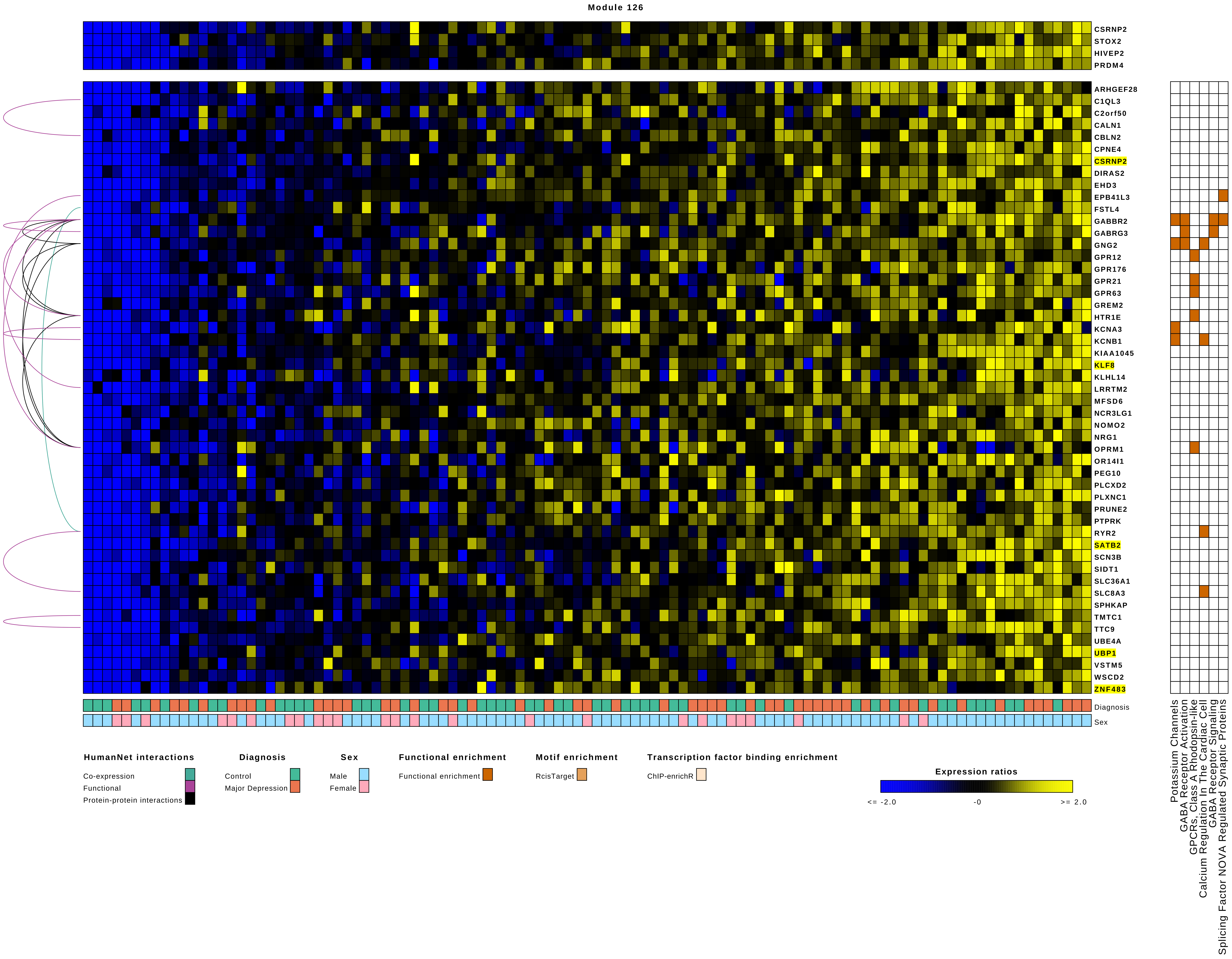
Module 126
CSRNP2
STOX2
HIVEP2
PRDM4
ARHGEF28
C1QL3
C2orf50
CALN1
CBLN2
CPNE4
CSRNP2
DIRAS2
EHD3
EPB41L3
FSTL4
GABBR2
GABRG3
GNG2
GPR12
GPR176
GPR21
GPR63
GREM2
HTR1E
KCNA3
KCNB1
KIAA1045
KLF8
KLHL14
LRRTM2
MFSD6
NCR3LG1
NOMO2
NRG1
OPRM1
OR14I1
PEG10
PLCXD2
PLXNC1
PRUNE2
PTPRK
RYR2
SATB2
SCN3B
SIDT1
SLC36A1
SLC8A3
SPHKAP
TMTC1
TTC9
UBE4A
UBP1
VSTM5
WSCD2
ZNF483
Diagnosis
Sex
Potassium Channels
GABA Receptor Activation
GPCRs, Class A Rhodopsin-like
Calcium Regulation In The Cardiac Cell
GABA Receptor Signaling
Splicing Factor NOVA Regulated Synaptic Proteins
HumanNet interactions
Co-expression
Functional
Protein-protein interactions
Diagnosis
Control
Major Depression
Sex
Male
Female
Functional enrichment
Functional enrichment
Motif enrichment
RcisTarget
Transcription factor binding enrichment
ChIP-enrichR
Expression ratios
<= -2.0
-0
>= 2.0

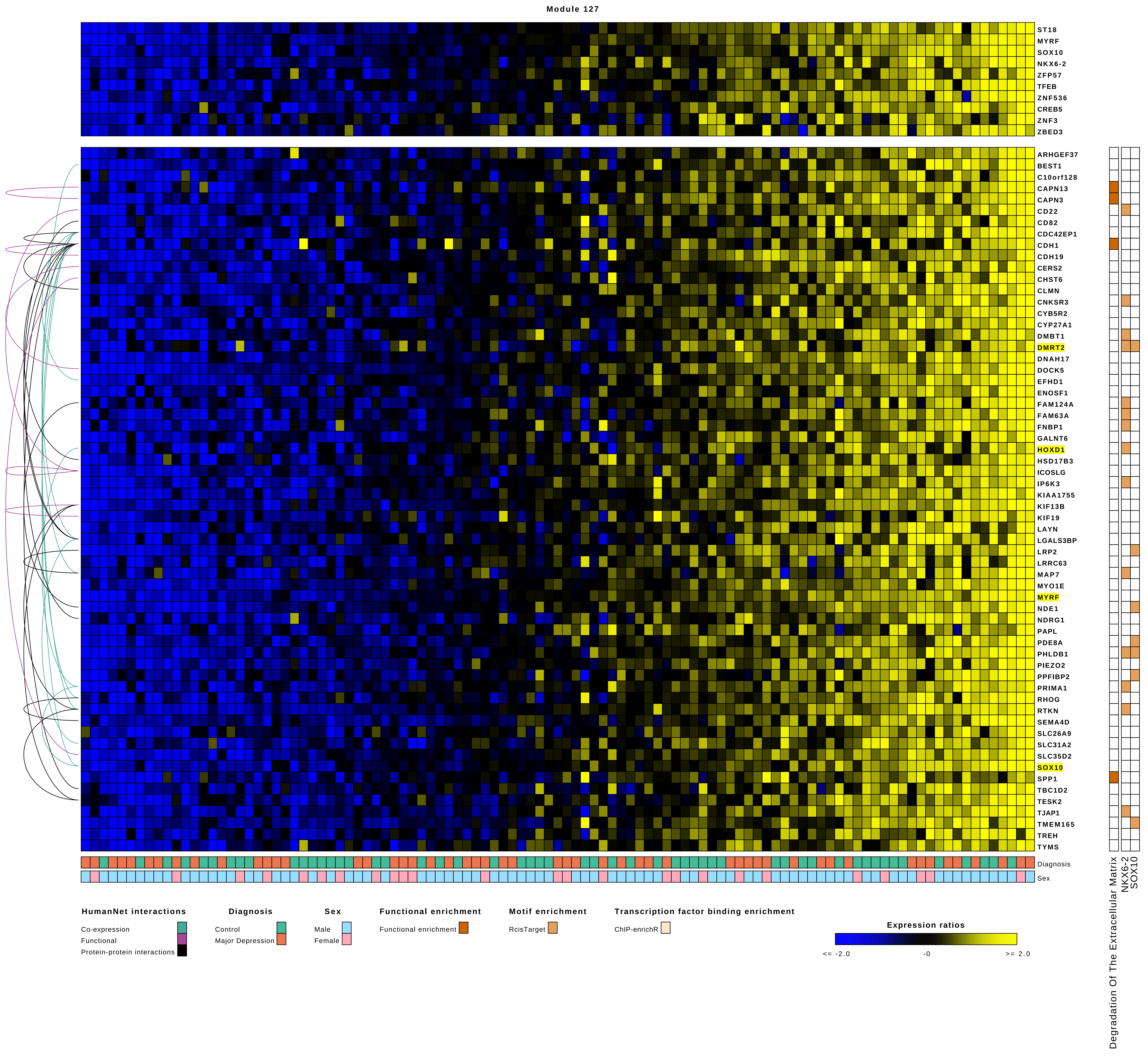

Module 127
ST18
MYRF
SOX10
NKX6-2
ZFP57
TFEB
ZNF536
CREB5
ZNF3
ZBED3
ARHGEF37
BEST1
C10orf128
CAPN13
CAPN3
CD22
CD82
CDC42EP1
CDH1
CDH19
CERS2
CHST6
CLMN
CNKSR3
CYB5R2
CYP27A1
DMBT1
DMRT2
DNAH17
DOCK5
EFHD1
ENOSF1
FAM124A
FAM63A
FNBP1
GALNT6
HOXD1
HSD17B3
ICOSLG
IP6K3
KIAA1755
KIF13B
KIF19
LAYN
LGALS3BP
LRP2
LRRC63
MAP7
MYO1E
MYRF
NDE1
NDRG1
PAPL
PDE8A
PHLDB1
PIEZO2
PPFIBP2
PRIMA1
RHOG
RTKN
SEMA4D
SLC26A9
SLC31A2
SLC35D2
SOX10
SPP1
TBC1D2
TESK2
TJAP1
TMEM165
TREH
TYMS
Diagnosis
Sex
Degradation Of The Extracellular Matrix
NKX6-2
SOX10
HumanNet interactions
Co-expression
Functional
Protein-protein interactions
Diagnosis
Control
Major Depression
Sex
Male
Female
Functional enrichment
Functional enrichment
Motif enrichment
RcisTarget
Transcription factor binding enrichment
ChIP-enrichR
Expression ratios
<= -2.0
-0
>= 2.0

### Module 128

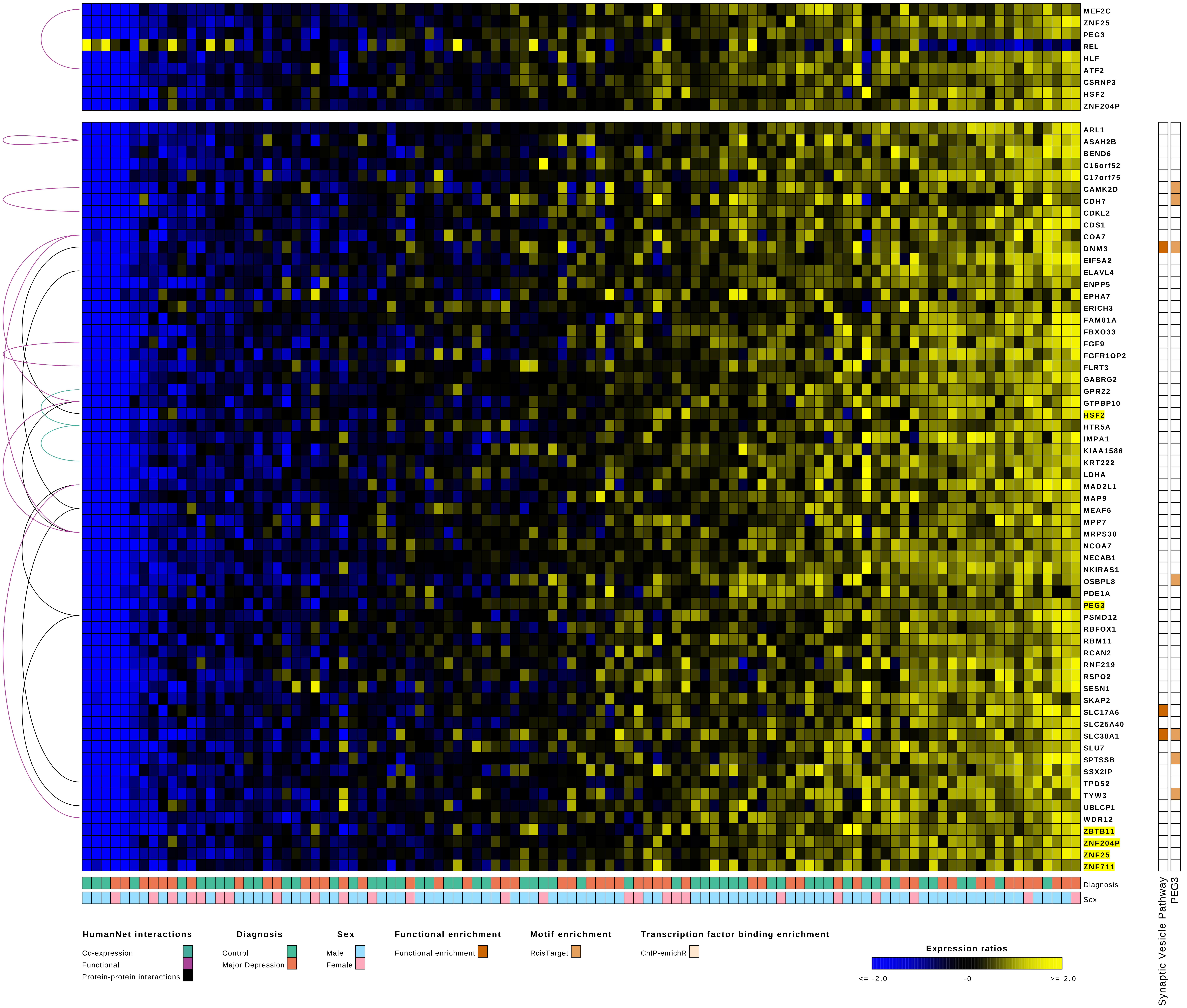
MEF2C
ZNF25
PEG3
REL
HLF
ATF2
CSRNP3
HSF2
ZNF204P
ARL1
ASAH2B
BEND6
C16orf52
C17orf75
CAMK2D
CDH7
CDKL2
CDS1
COA7
DNM3
EIF5A2
ELAVL4
ENPP5
EPHA7
ERICH3
FAM81A
FBXO33
FGF9
FGFR1OP2
FLRT3
GABRG2
GPR22
GTPBP10
HSF2
HTR5A
IMPA1
KIAA1586
KRT222
LDHA
MAD2L1
MAP9
MEAF6
MPP7
MRPS30
NCOA7
NECAB1
NKIRAS1
OSBPL8
PDE1A
PEG3
PSMD12
RBFOX1
RBM11
RCAN2
RNF219
RSPO2
SESN1
SKAP2
SLC17A6
SLC25A40
SLC38A1
SLU7
SPTSSB
SSX2IP
TPD52
TYW3
UBLCP1
WDR12
ZBTB11
ZNF204P
ZNF25
ZNF711
Diagnosis
Sex
Synaptic Vesicle Pathway
PEG3
HumanNet interactions
Co-expression
Functional
Protein-protein interactions
Diagnosis
Control
Major Depression
Sex
Male
Female
Functional enrichment
Functional enrichment
Motif enrichment
RcisTarget
Transcription factor binding enrichment
ChIP-enrichR
Expression ratios
<= -2.0
-0
>= 2.0

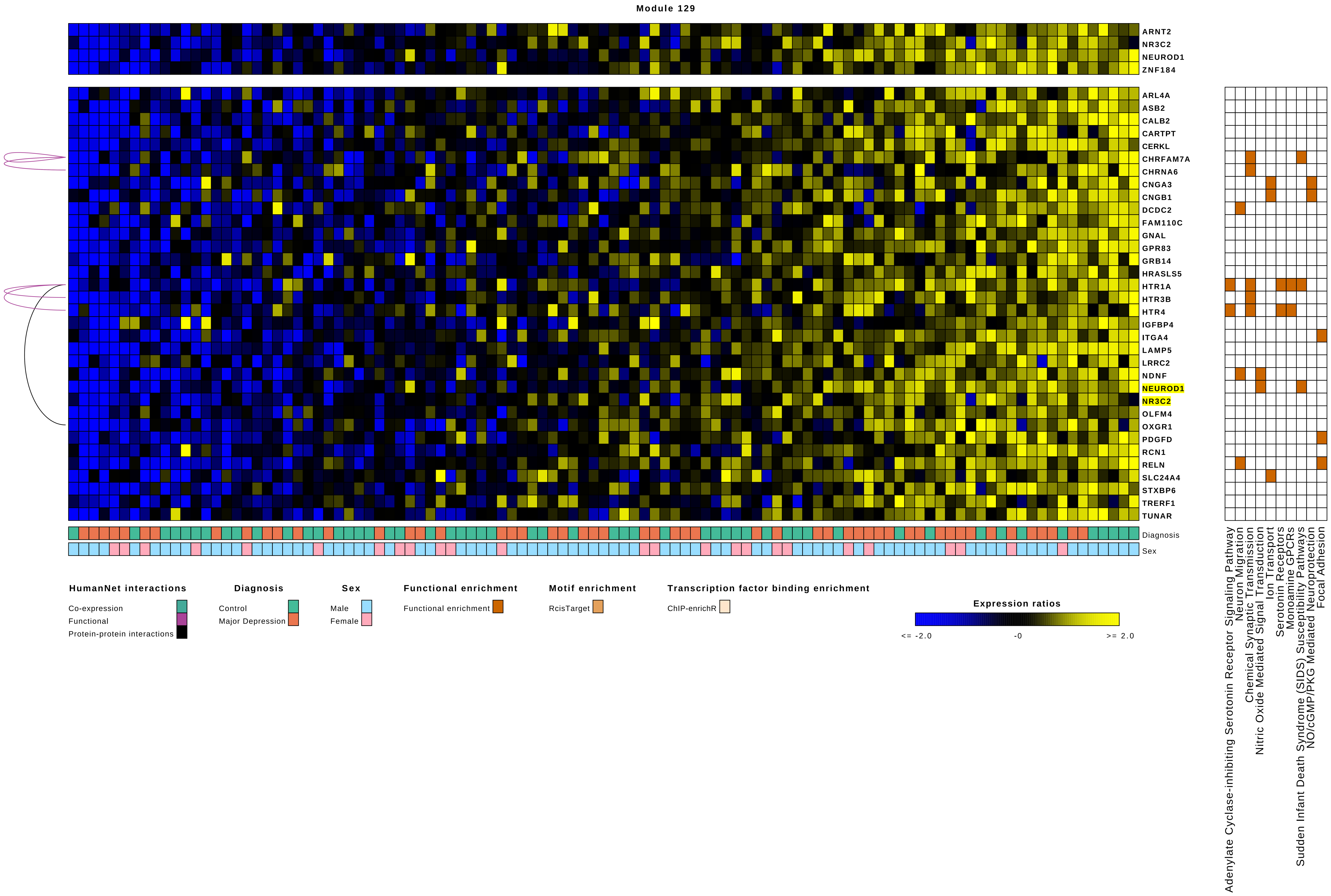

Module 129
ARNT2
NR3C2
NEUROD1
ZNF184
ARL4A
ASB2
CALB2
CARTPT
CERKL
CHRFAM7A
CHRNA6
CNGA3
CNGB1
DCDC2
FAM110C
GNAL
GPR83
GRB14
HRASLS5
HTR1A
HTR3B
HTR4
IGFBP4
ITGA4
LAMP5
LRRC2
NDNF
NEUROD1
NR3C2
OLFM4
OXGR1
PDGFD
RCN1
RELN
SLC24A4
STXBP6
TRERF1
TUNAR
Diagnosis
Sex
Adenylate Cyclase-inhibiting Serotonin Receptor Signaling Pathway
Neuron Migration
Chemical Synaptic Transmission
Nitric Oxide Mediated Signal Transduction
Ion Transport
Serotonin Receptors
Monoamine GPCRs
Sudden Infant Death Syndrome (SIDS) Susceptibility Pathways
NO/cGMP/PKG Mediated Neuroprotection
Focal Adhesion
HumanNet interactions
Co-expression
Functional
Protein-protein interactions
Diagnosis
Control
Major Depression
Sex
Male
Female
Functional enrichment
Functional enrichment
Motif enrichment
RcisTarget
Transcription factor binding enrichment
ChIP-enrichR
Expression ratios
<= -2.0
-0
>= 2.0

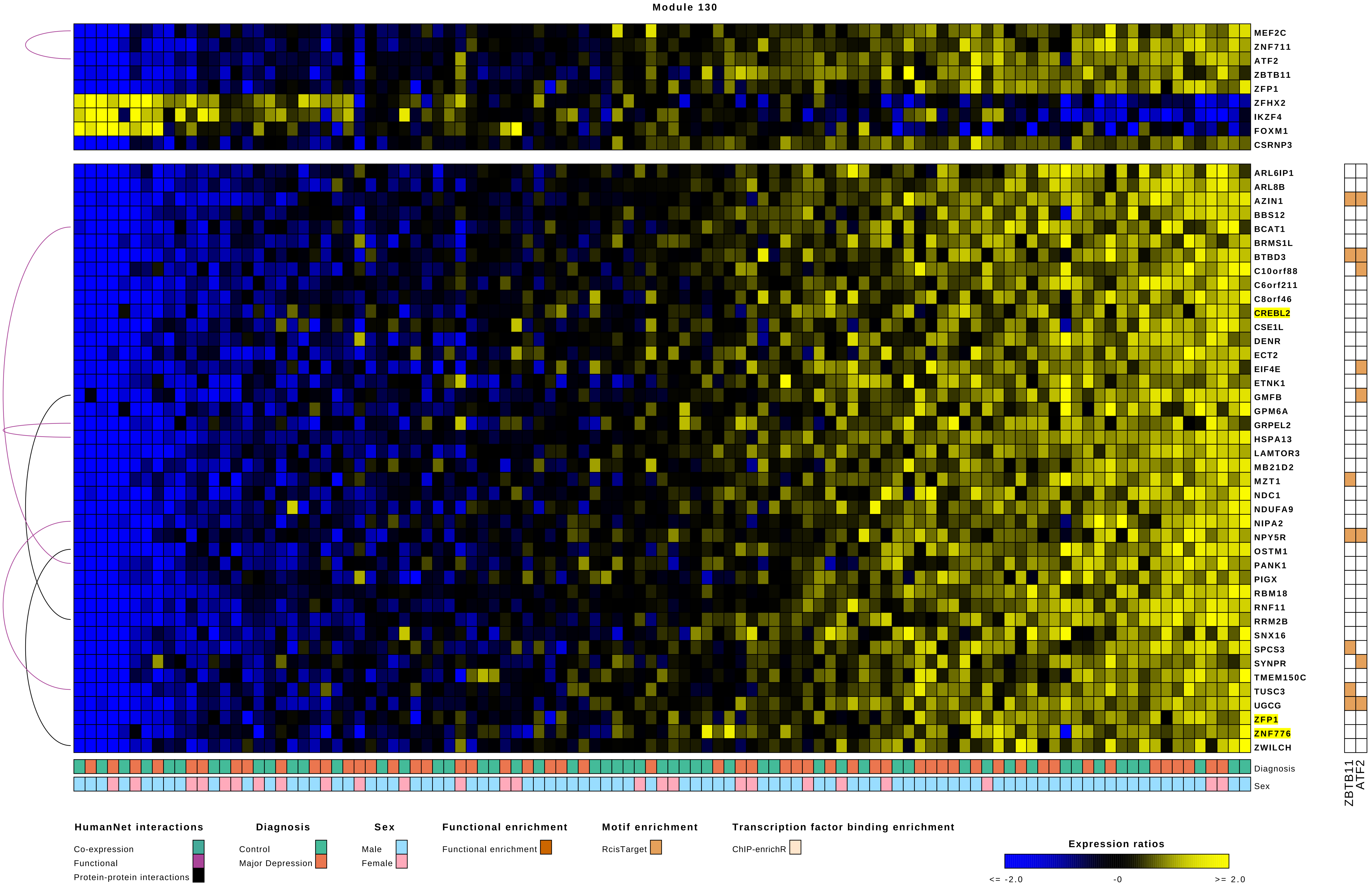
Module 130
MEF2C
ZNF711
ATF2
ZBTB11
ZFP1
ZFHX2
IKZF4
FOXM1
CSRNP3
ARL6IP1
ARL8B
AZIN1
BBS12
BCAT1
BRMS1L
BTBD3
C10orf88
C6orf211
C8orf46
CREBL2
CSE1L
DENR
ECT2
EIF4E
ETNK1
GMFB
GPM6A
GRPEL2
HSPA13
LAMTOR3
MB21D2
MZT1
NDC1
NDUFA9
NIPA2
NPY5R
OSTM1
PANK1
PIGX
RBM18
RNF11
RRM2B
SNX16
SPCS3
SYNPR
TMEM150C
TUSC3
UGCG
ZFP1
ZNF776
ZWILCH
Diagnosis
Sex
ZBTB11
ATF2
HumanNet interactions
Co-expression
Functional
Protein-protein interactions
Diagnosis
Control
Major Depression
Sex
Male
Female
Functional enrichment
Functional enrichment
Motif enrichment
RcisTarget
Transcription factor binding enrichment
ChIP-enrichR
Expression ratios
<= -2.0
-0
>= 2.0

### Module 131

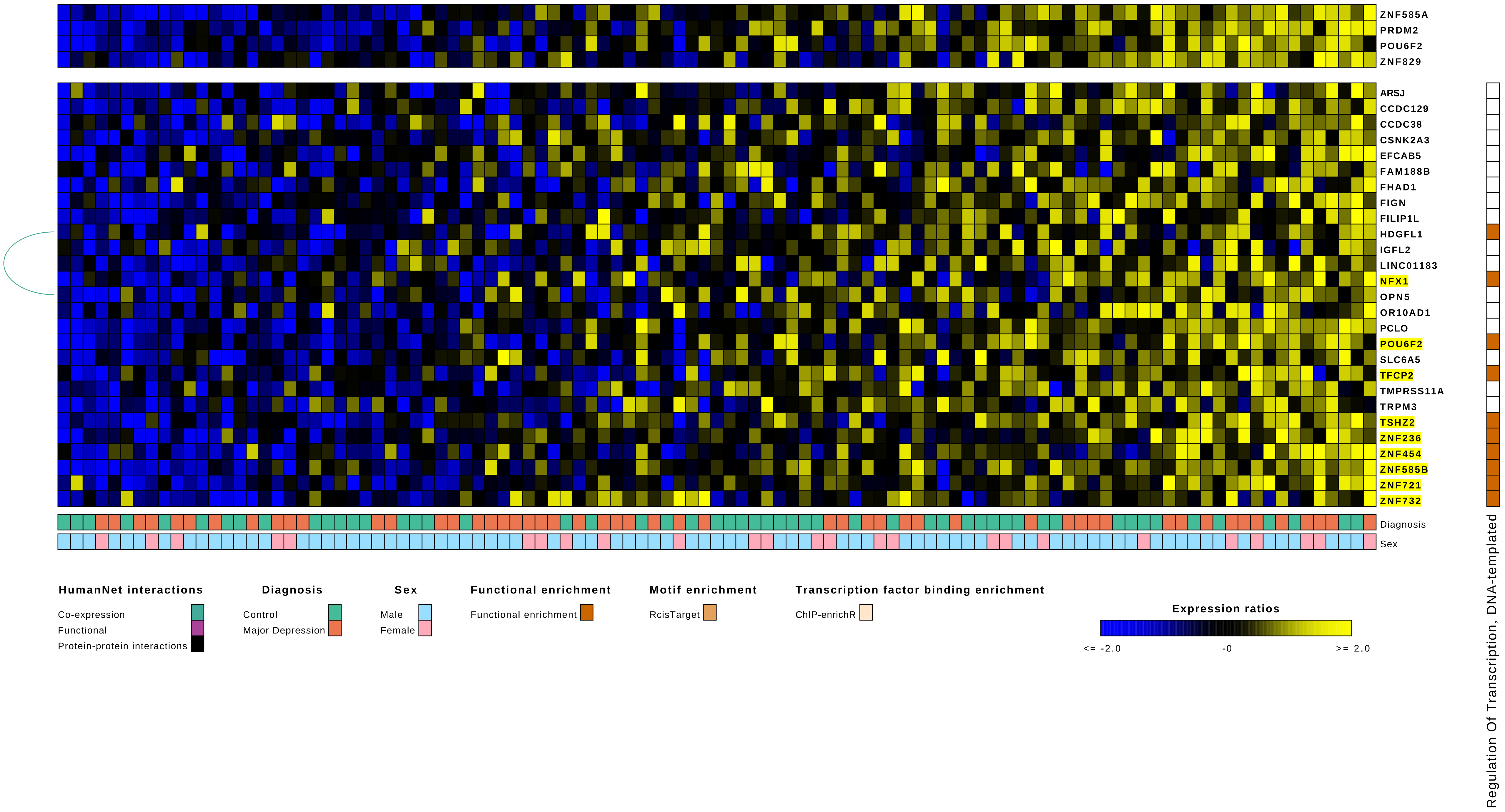

ZNF585A
PRDM2
POU6F2
ZNF829
ARSJ
CCDC129
CCDC38
CSNK2A3
EFCAB5
FAM188B
FHAD1
FIGN
FILIP1L
HDGFL1
IGFL2
LINC01183
NFX1
OPN5
OR10AD1
PCLO
POU6F2
SLC6A5
TFCP2
TMPRSS11A
TRPM3
TSHZ2
ZNF236
ZNF454
ZNF585B
ZNF721
ZNF732
Diagnosis
Sex
Regulation Of Transcription, DNA-templated
HumanNet interactions
Co-expression
Functional
Protein-protein interactions
Diagnosis
Control
Major Depression
Sex
Male
Female
Functional enrichment
Functional enrichment
Motif enrichment
RcisTarget
Transcription factor binding enrichment
ChIP-enrichR
Expression ratios
<= -2.0
-0
>= 2.0

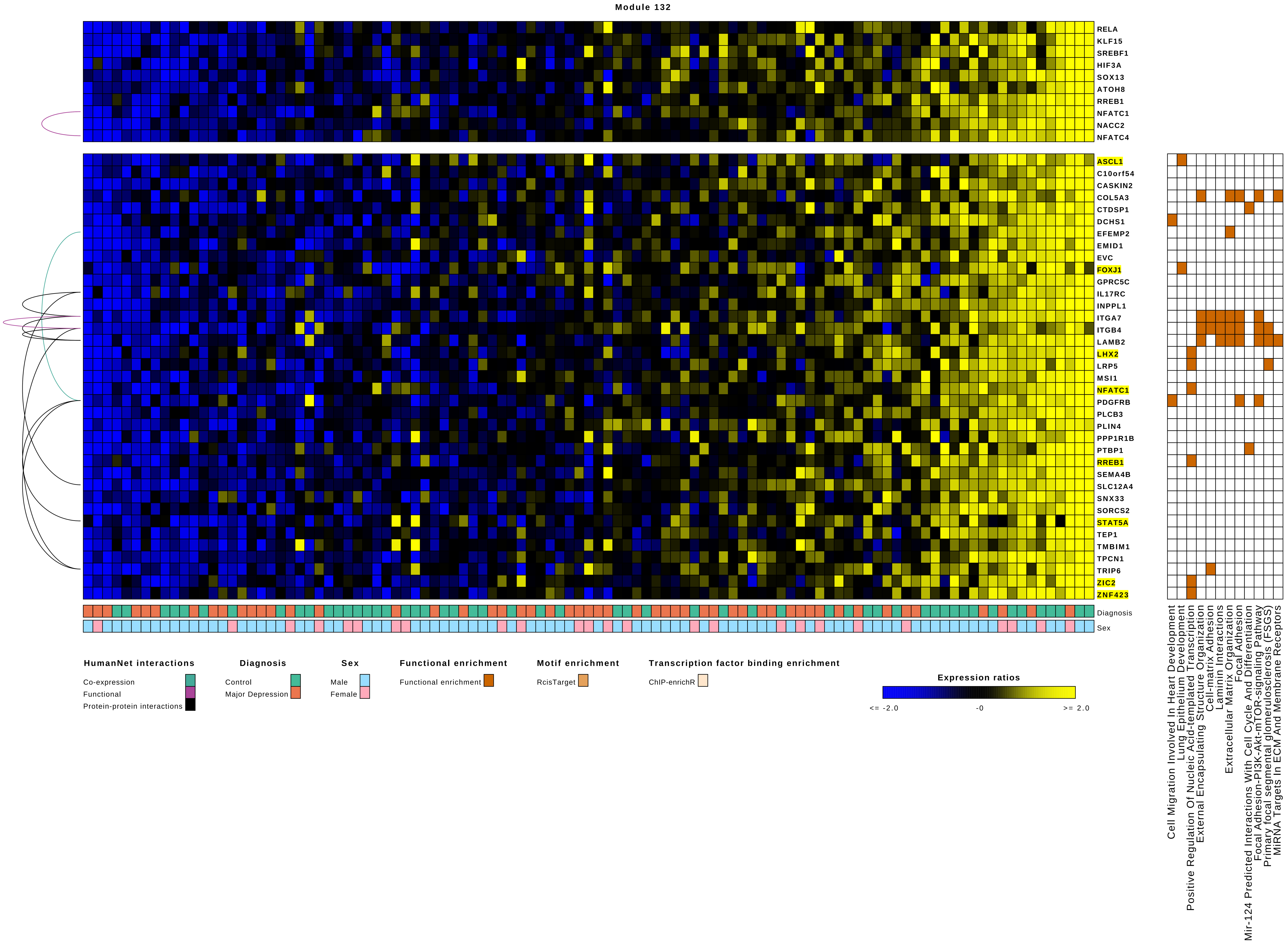
Module 132
RELA
KLF15
SREBF1
HIF3A
SOX13
ATOH8
RREB1
NFATC1
NACC2
NFATC4
ASCL1
C10orf54
CASKIN2
COL5A3
CTDSP1
DCHS1
EFEMP2
EMID1
EVC
FOXJ1
GPRC5C
IL17RC
INPPL1
ITGA7
ITGB4
LAMB2
LHX2
LRP5
MSI1
NFATC1
PDGFRB
PLCB3
PLIN4
PPP1R1B
PTBP1
RREB1
SEMA4B
SLC12A4
SNX33
SORCS2
STAT5A
TEP1
TMBIM1
TPCN1
TRIP6
ZIC2
ZNF423
Diagnosis
Sex
Cell Migration Involved In Heart Development
Lung Epithelium Development
Positive Regulation Of Nucleic Acid-templated Transcription
External Encapsulating Structure Organization
Cell-matrix Adhesion
Laminin Interactions
Extracellular Matrix Organization
Focal Adhesion
Mir-124 Predicted Interactions With Cell Cycle And Differentiation
Focal Adhesion-PI3K-Akt-mTOR-signaling Pathway
Primary focal segmental glomerulosclerosis (FSGS)
MiRNA Targets In ECM And Membrane Receptors
HumanNet interactions
Co-expression
Functional
Protein-protein interactions
Diagnosis
Control
Major Depression
Sex
Male
Female
Functional enrichment
Functional enrichment
Motif enrichment
RcisTarget
Transcription factor binding enrichment
ChIP-enrichR
Expression ratios
<= -2.0
-0
>= 2.0

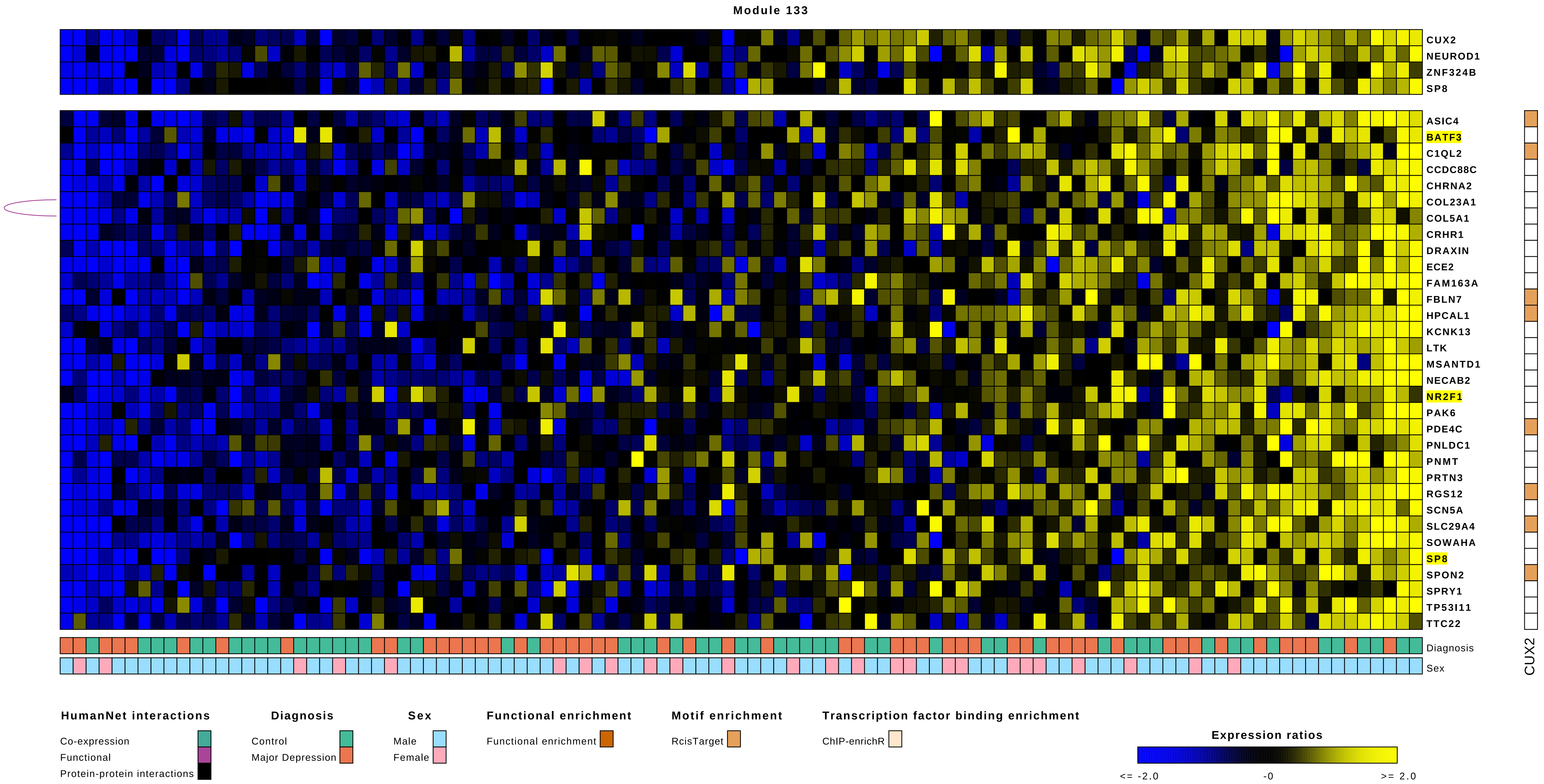
Module 133
CUX2
NEUROD1
ZNF324B
SP8
ASIC4
BATF3
C1QL2
CCDC88C
CHRNA2
COL23A1
COL5A1
CRHR1
DRAXIN
ECE2
FAM163A
FBLN7
HPCAL1
KCNK13
LTK
MSANTD1
NECAB2
NR2F1
PAK6
PDE4C
PNLDC1
PNMT
PRTN3
RGS12
SCN5A
SLC29A4
SOWAHA
SP8
SPON2
SPRY1
TP53I11
TTC22
Diagnosis
Sex
CUX2
HumanNet interactions
Co-expression
Functional
Protein-protein interactions
Diagnosis
Control
Major Depression
Sex
Male
Female
Functional enrichment
Functional enrichment
Motif enrichment
RcisTarget
Transcription factor binding enrichment
ChIP-enrichR
Expression ratios
<= -2.0
-0
>= 2.0

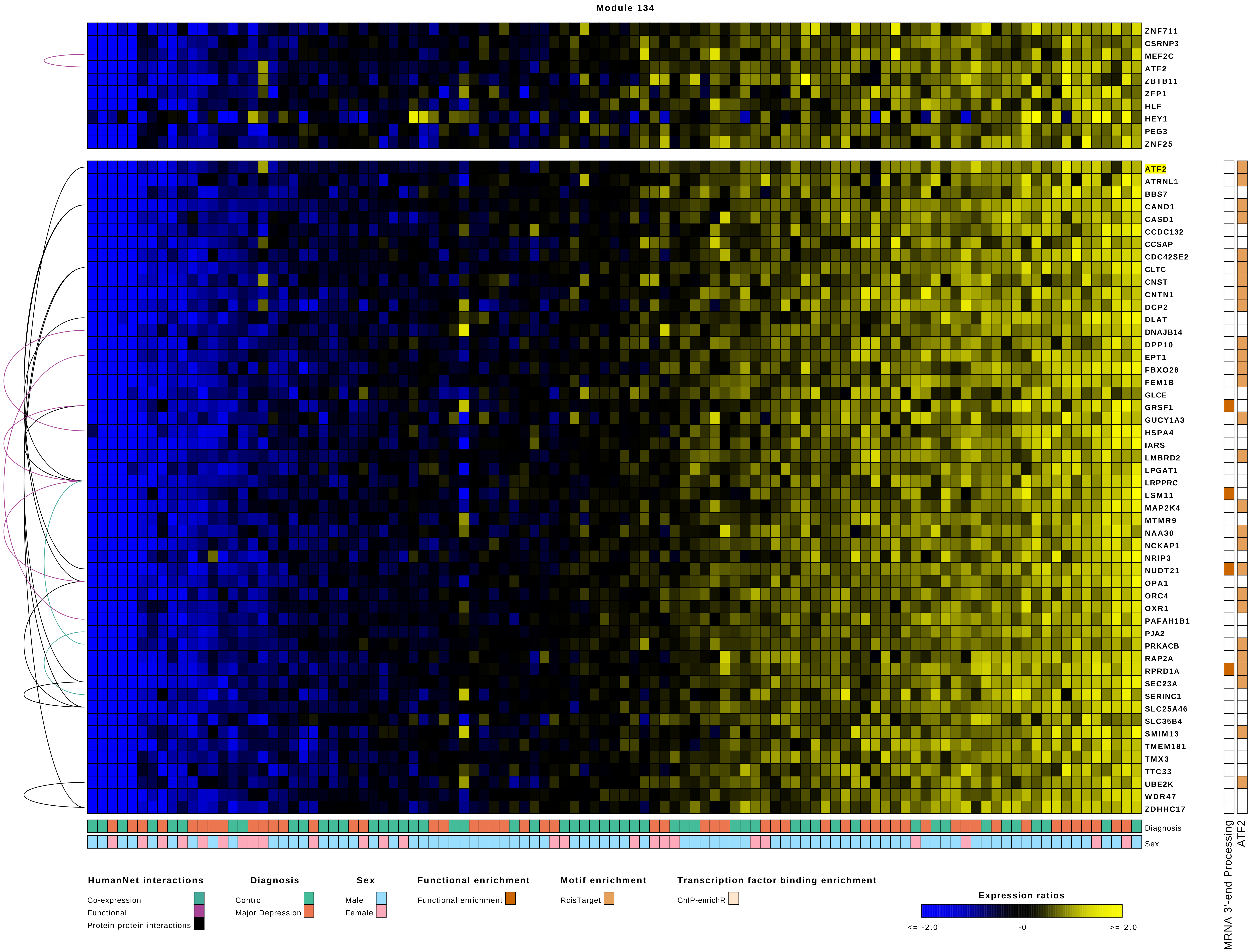

Module 134
ZNF711
CSRNP3
MEF2C
ATF2
ZBTB11
ZFP1
HLF
HEY1
PEG3
ZNF25
ATF2
ATRNL1
BBS7
CAND1
CASD1
CCDC132
CCSAP
CDC42SE2
CLTC
CNST
CNTN1
DCP2
DLAT
DNAJB14
DPP10
EPT1
FBXO28
FEM1B
GLCE
GRSF1
GUCY1A3
HSPA4
IARS
LMBRD2
LPGAT1
LRPPRC
LSM11
MAP2K4
MTMR9
NAA30
NCKAP1
NRIP3
NUDT21
OPA1
ORC4
OXR1
PAFAH1B1
PJA2
PRKACB
RAP2A
RPRD1A
SEC23A
SERINC1
SLC25A46
SLC35B4
SMIM13
TMEM181
TMX3
TTC33
UBE2K
WDR47
ZDHHC17
Diagnosis
Sex
MRNA 3'-end Processing
ATF2
HumanNet interactions
Co-expression
Functional
Protein-protein interactions
Diagnosis
Control
Major Depression
Sex
Male
Female
Functional enrichment
Functional enrichment
Motif enrichment
RcisTarget
Transcription factor binding enrichment
ChIP-enrichR
Expression ratios
<= -2.0
-0
>= 2.0

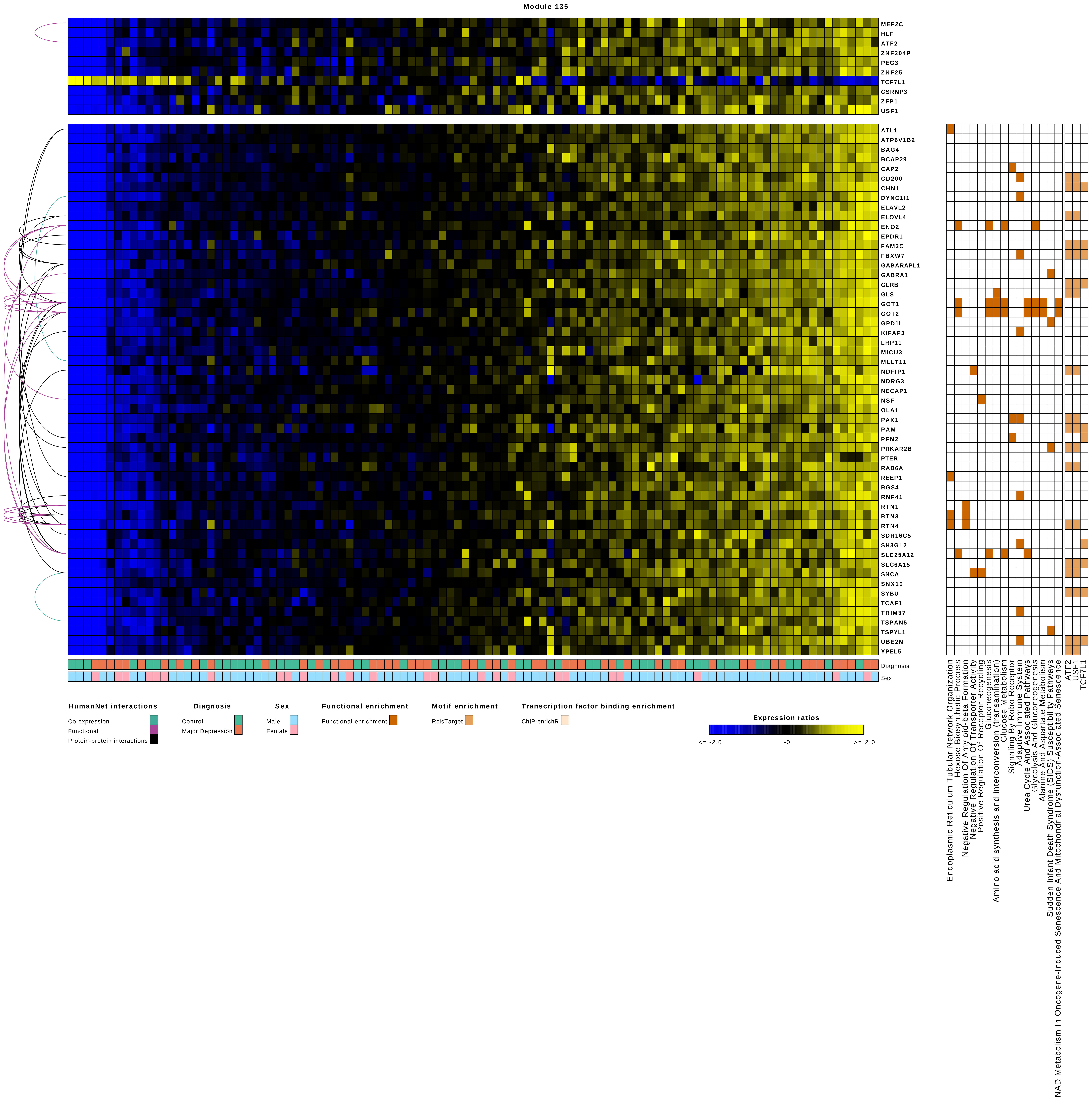
Module 135
MEF2C
HLF
ATF2
ZNF204P
PEG3
ZNF25
TCF7L1
CSRNP3
ZFP1
USF1
ATL1
ATP6V1B2
BAG4
BCAP29
CAP2
CD200
CHN1
DYNC1I1
ELAVL2
ELOVL4
ENO2
EPDR1
FAM3C
FBXW7
GABARAPL1
GABRA1
GLRB
GLS
GOT1
GOT2
GPD1L
KIFAP3
LRP11
MICU3
MLLT11
NDFIP1
NDRG3
NECAP1
NSF
OLA1
PAK1
PAM
PFN2
PRKAR2B
PTER
RAB6A
REEP1
RGS4
RNF41
RTN1
RTN3
RTN4
SDR16C5
SH3GL2
SLC25A12
SLC6A15
SNCA
SNX10
SYBU
TCAF1
TRIM37
TSPAN5
TSPYL1
UBE2N
YPEL5
Diagnosis
Sex
HumanNet interactions
Co-expression
Functional
Protein-protein interactions
Diagnosis
Control
Major Depression
Sex
Male
Female
Functional enrichment
Functional enrichment
Motif enrichment
RcisTarget
Transcription factor binding enrichment
ChIP-enrichR
Expression ratios
<= -2.0
-0
>= 2.0
Endoplasmic Reticulum Tubular Network Organization
Hexose Biosynthetic Process
Negative Regulation Of Amyloid-beta Formation
Negative Regulation Of Transporter Activity
Positive Regulation Of Receptor Recycling
Gluconeogenesis
Amino acid synthesis and interconversion (transamination)
Glucose Metabolism
Signaling By Robo Receptor
Adaptive Immune System
Urea Cycle And Associated Pathways
Glycolysis And Gluconeogenesis
Alanine And Aspartate Metabolism
Sudden Infant Death Syndrome (SIDS) Susceptibility Pathways
NAD Metabolism In Oncogene-Induced Senescence And Mitochondrial Dysfunction-Associated Senescence
ATF2
USF1
TCF7L1

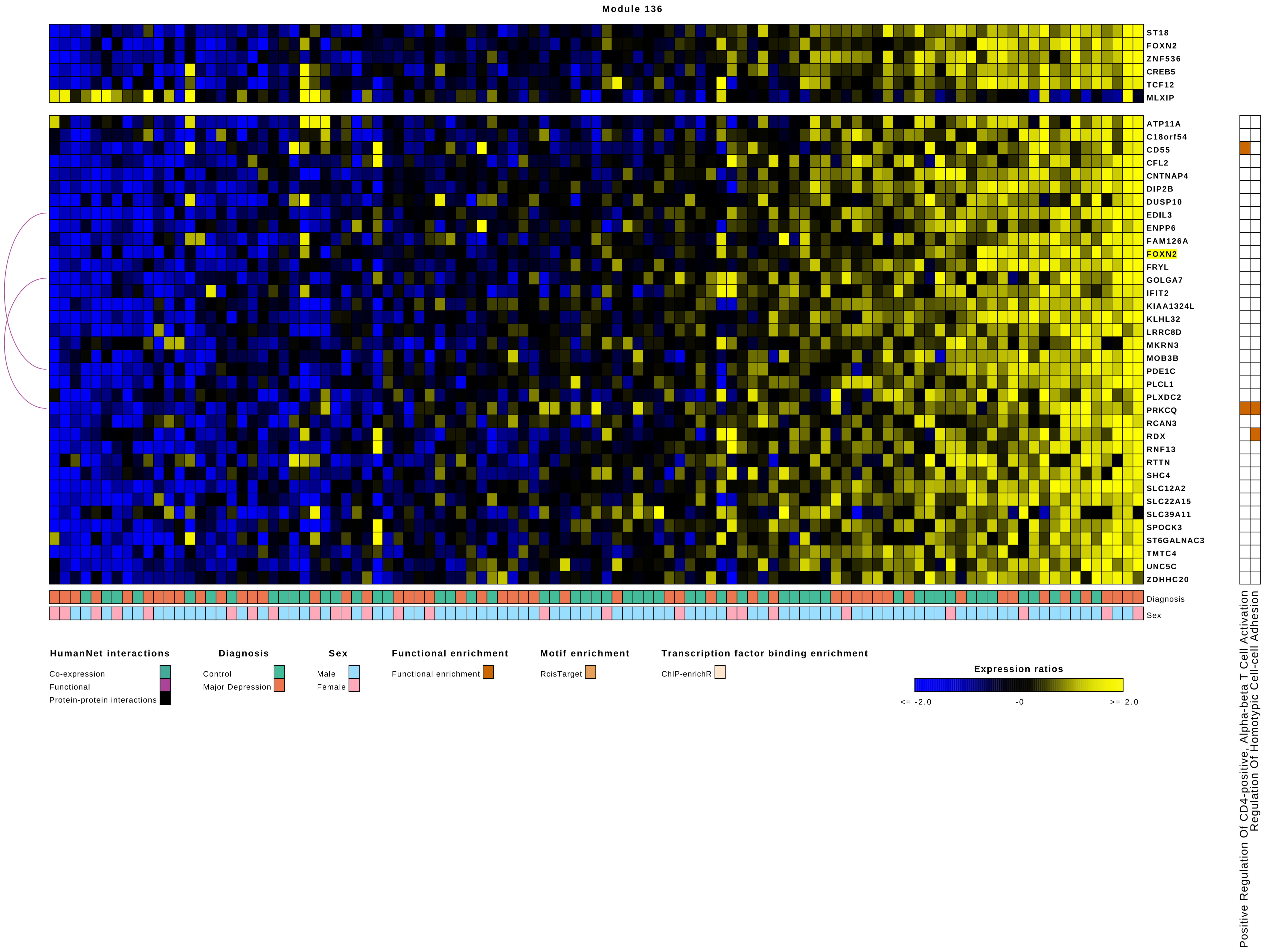
Module 136
ST18
FOXN2
ZNF536
CREB5
TCF12
MLXIP
ATP11A
C18orf54
CD55
CFL2
CNTNAP4
DIP2B
DUSP10
EDIL3
ENPP6
FAM126A
FOXN2
FRYL
GOLGA7
IFIT2
KIAA1324L
KLHL32
LRRC8D
MKRN3
MOB3B
PDE1C
PLCL1
PLXDC2
PRKCQ
RCAN3
RDX
RNF13
RTTN
SHC4
SLC12A2
SLC22A15
SLC39A11
SPOCK3
ST6GALNAC3
TMTC4
UNC5C
ZDHHC20
Diagnosis
Sex
HumanNet interactions
Co-expression
Functional
Protein-protein interactions
Diagnosis
Control
Major Depression
Sex
Male
Female
Functional enrichment
Functional enrichment
Motif enrichment
RcisTarget
Transcription factor binding enrichment
ChIP-enrichR
Expression ratios
<= -2.0
-0
>= 2.0
Positive Regulation Of CD4-positive, Alpha-beta T Cell Activation
Regulation Of Homotypic Cell-cell Adhesion

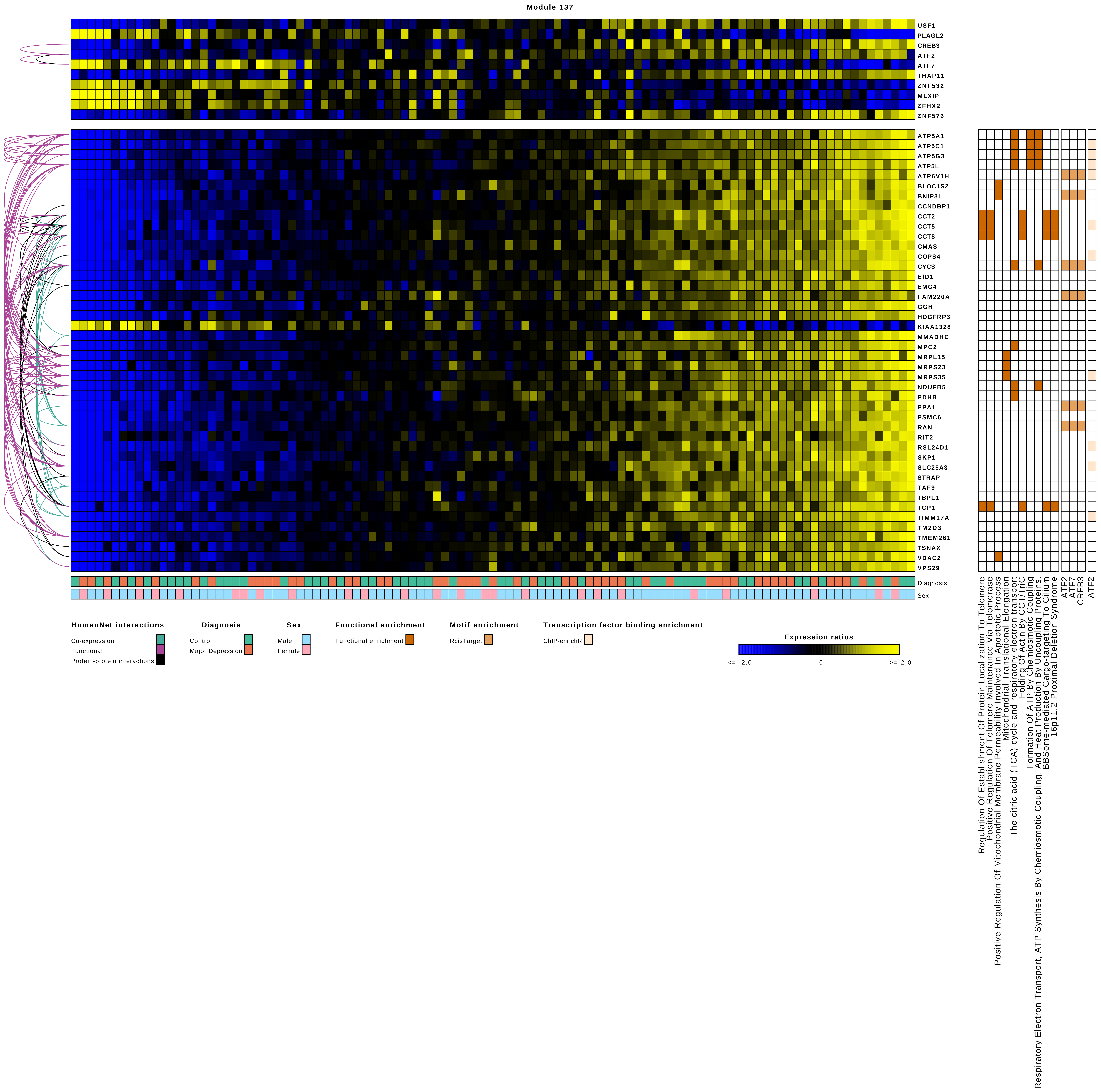

Module 137
USF1
PLAGL2
CREB3
ATF2
ATF7
THAP11
ZNF532
MLXIP
ZFHX2
ZNF576
ATP5A1
ATP5C1
ATP5G3
ATP5L
ATP6V1H
BLOC1S2
BNIP3L
CCNDBP1
CCT2
CCT5
CCT8
CMAS
COPS4
CYCS
EID1
EMC4
FAM220A
GGH
HDGFRP3
KIAA1328
MMADHC
MPC2
MRPL15
MRPS23
MRPS35
NDUFB5
PDHB
PPA1
PSMC6
RAN
RIT2
RSL24D1
SKP1
SLC25A3
STRAP
TAF9
TBPL1
TCP1
TIMM17A
TM2D3
TMEM261
TSNAX
VDAC2
VPS29
Diagnosis
Sex
Regulation Of Establishment Of Protein Localization To Telomere
Positive Regulation Of Telomere Maintenance Via Telomerase
Positive Regulation Of Mitochondrial Membrane Permeability Involved In Apoptotic Process
Mitochondrial Translational Elongation
The citric acid (TCA) cycle and respiratory electron transport
Folding Of Actin By CCT/TriC
Formation Of ATP By Chemiosmotic Coupling
Respiratory Electron Transport, ATP Synthesis By Chemiosmotic Coupling, And Heat Production By Uncoupling Proteins.
BBSome-mediated Cargo-targeting To Cilium
16p11.2 Proximal Deletion Syndrome
ATF2
ATF7
CREB3
ATF2
HumanNet interactions
Co-expression
Functional
Protein-protein interactions
Diagnosis
Control
Major Depression
Sex
Male
Female
Functional enrichment
Functional enrichment
Motif enrichment
RcisTarget
Transcription factor binding enrichment
ChIP-enrichR
Expression ratios
<= -2.0
-0
>= 2.0

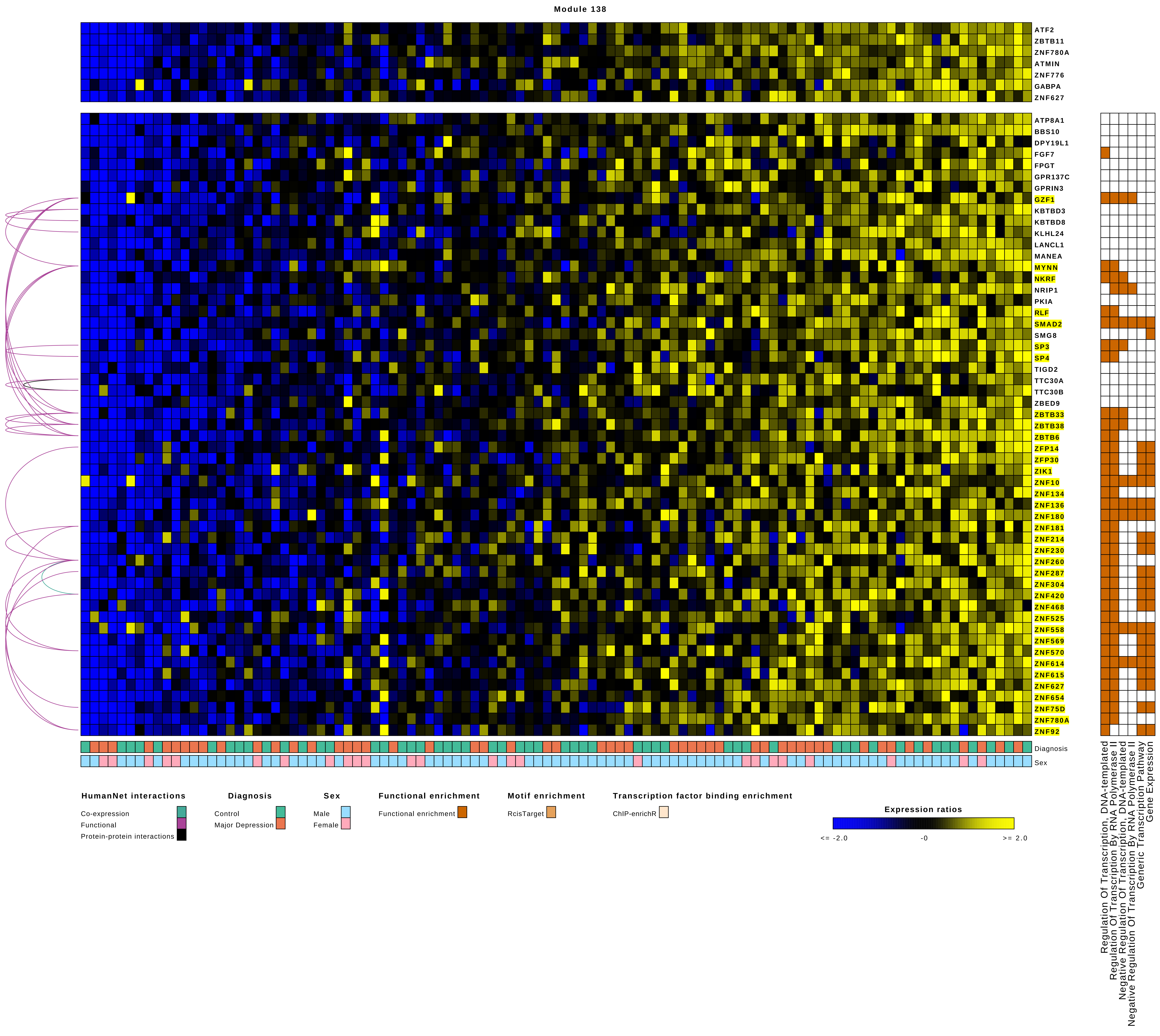

Module 138
ATF2
ZBTB11
ZNF780A
ATMIN
ZNF776
GABPA
ZNF627
ATP8A1
BBS10
DPY19L1
FGF7
FPGT
GPR137C
GPRIN3
GZF1
KBTBD3
KBTBD8
KLHL24
LANCL1
MANEA
MYNN
NKRF
NRIP1
PKIA
RLF
SMAD2
SMG8
SP3
SP4
TIGD2
TTC30A
TTC30B
ZBED9
ZBTB33
ZBTB38
ZBTB6
ZFP14
ZFP30
ZIK1
ZNF10
ZNF134
ZNF136
ZNF180
ZNF181
ZNF214
ZNF230
ZNF260
ZNF287
ZNF304
ZNF420
ZNF468
ZNF525
ZNF558
ZNF569
ZNF570
ZNF614
ZNF615
ZNF627
ZNF654
ZNF75D
ZNF780A
ZNF92
Diagnosis
Sex
HumanNet interactions
Co-expression
Functional
Protein-protein interactions
Diagnosis
Control
Major Depression
Sex
Male
Female
Functional enrichment
Functional enrichment
Motif enrichment
RcisTarget
Transcription factor binding enrichment
ChIP-enrichR
Expression ratios
<= -2.0
-0
>= 2.0
Regulation Of Transcription, DNA-templated
Regulation Of Transcription By RNA Polymerase II
Negative Regulation Of Transcription, DNA-templated
Negative Regulation Of Transcription By RNA Polymerase II
Generic Transcription Pathway
Gene Expression

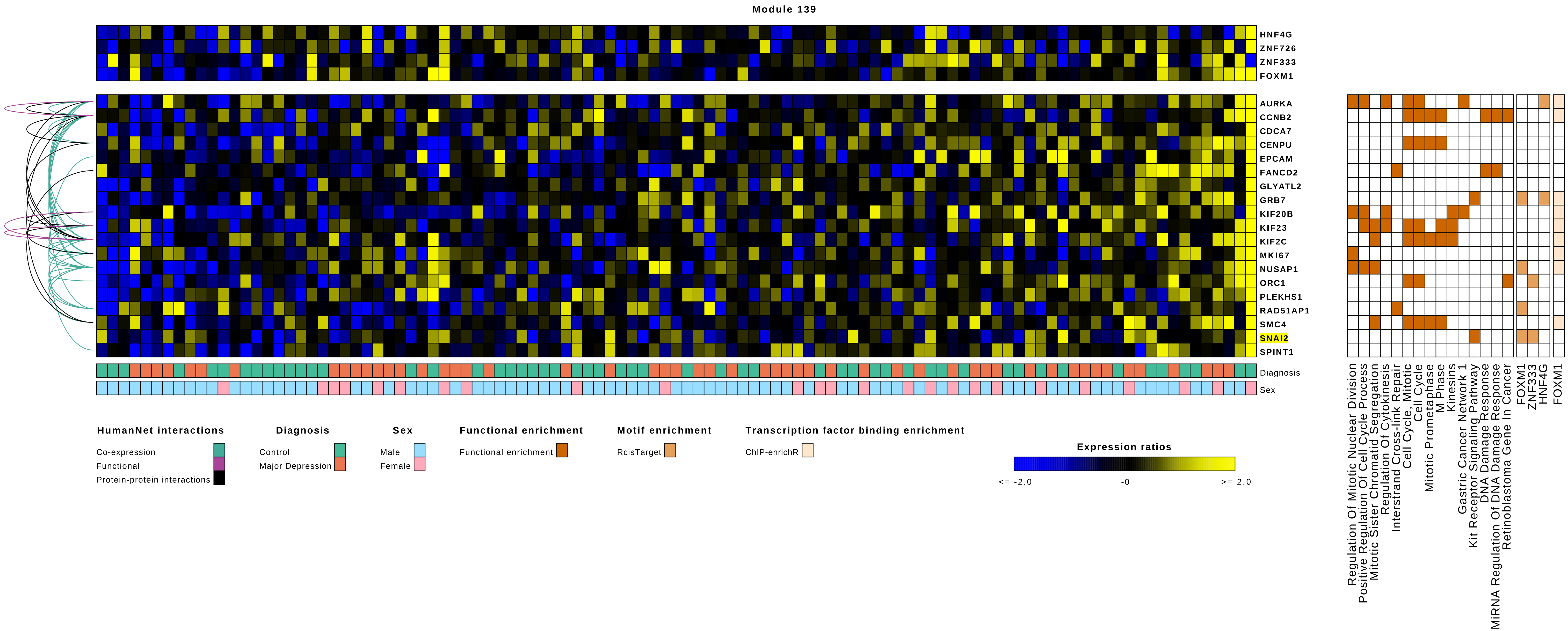

Module 139
HNF4G
ZNF726
ZNF333
FOXM1
AURKA
CCNB2
CDCA7
CENPU
EPCAM
FANCD2
GLYATL2
GRB7
KIF20B
KIF23
KIF2C
MKI67
NUSAP1
ORC1
PLEKHS1
RAD51AP1
SMC4
SNAI2
SPINT1
Diagnosis
Sex
HumanNet interactions
Co-expression
Functional
Protein-protein interactions
Diagnosis
Control
Major Depression
Sex
Male
Female
Functional enrichment
Functional enrichment
Motif enrichment
RcisTarget
Transcription factor binding enrichment
ChIP-enrichR
Expression ratios
<= -2.0
-0
>= 2.0
Regulation Of Mitotic Nuclear Division
Positive Regulation Of Cell Cycle Process
Mitotic Sister Chromatid Segregation
Regulation Of Cytokinesis
Interstrand Cross-link Repair
Cell Cycle, Mitotic
Cell Cycle
Mitotic Prometaphase
M Phase
Kinesins
Gastric Cancer Network 1
Kit Receptor Signaling Pathway
DNA Damage Response
MiRNA Regulation Of DNA Damage Response
Retinoblastoma Gene In Cancer
FOXM1
ZNF333
HNF4G
FOXM1

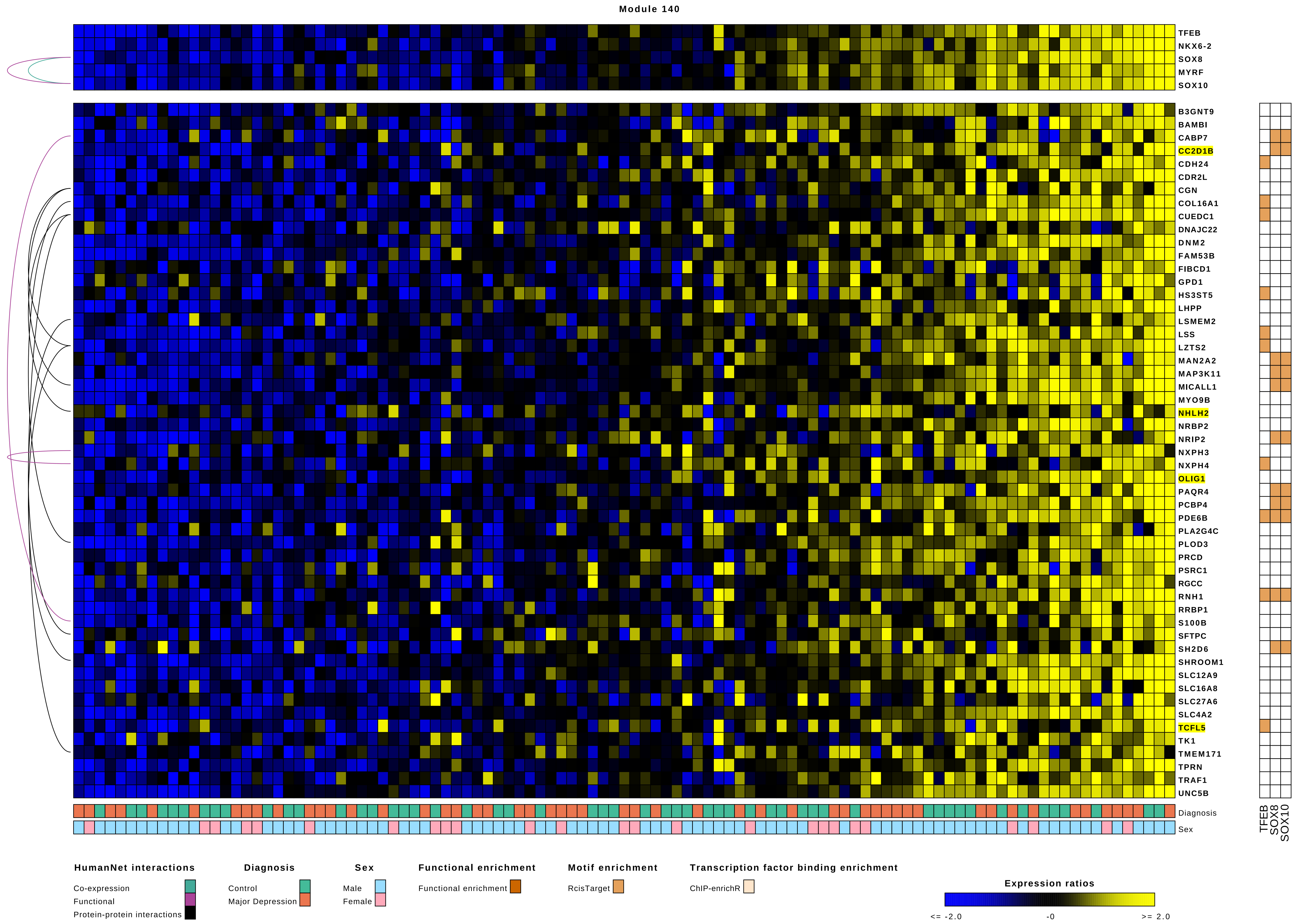
Module 140
TFEB
NKX6-2
SOX8
MYRF
SOX10
B3GNT9
BAMBI
CABP7
CC2D1B
CDH24
CDR2L
CGN
COL16A1
CUEDC1
DNAJC22
DNM2
FAM53B
FIBCD1
GPD1
HS3ST5
LHPP
LSMEM2
LSS
LZTS2
MAN2A2
MAP3K11
MICALL1
MYO9B
NHLH2
NRBP2
NRIP2
NXPH3
NXPH4
OLIG1
PAQR4
PCBP4
PDE6B
PLA2G4C
PLOD3
PRCD
PSRC1
RGCC
RNH1
RRBP1
S100B
SFTPC
SH2D6
SHROOM1
SLC12A9
SLC16A8
SLC27A6
SLC4A2
TCFL5
TK1
TMEM171
TPRN
TRAF1
UNC5B
Diagnosis
Sex
TFEB
SOX8
SOX10
HumanNet interactions
Co-expression
Functional
Protein-protein interactions
Diagnosis
Control
Major Depression
Sex
Male
Female
Functional enrichment
Functional enrichment
Motif enrichment
RcisTarget
Transcription factor binding enrichment
ChIP-enrichR
Expression ratios
<= -2.0
-0
>= 2.0

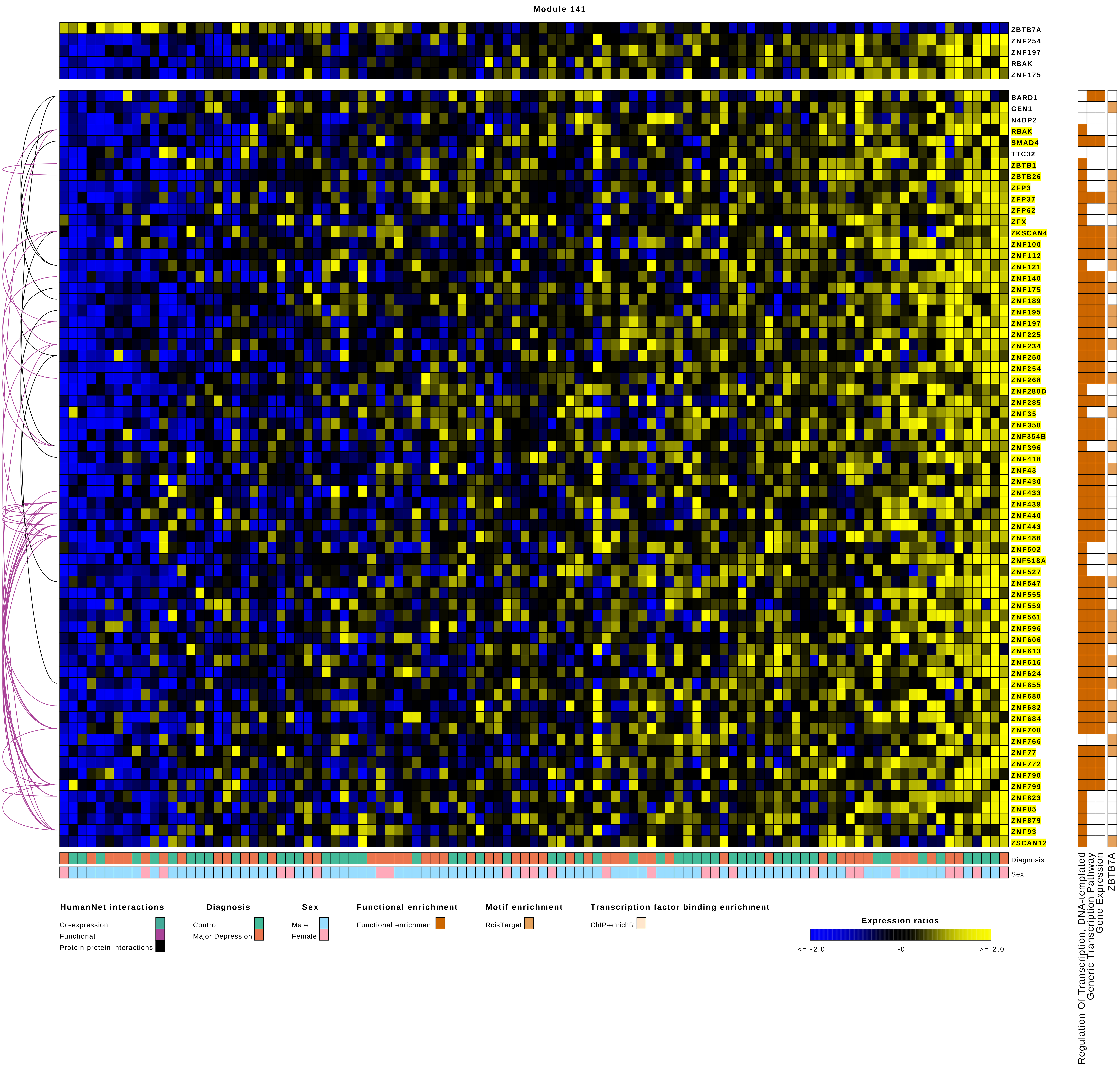

Module 141
ZBTB7A
ZNF254
ZNF197
RBAK
ZNF175
BARD1
GEN1
N4BP2
RBAK
SMAD4
TTC32
ZBTB1
ZBTB26
ZFP3
ZFP37
ZFP62
ZFX
ZKSCAN4
ZNF100
ZNF112
ZNF121
ZNF140
ZNF175
ZNF189
ZNF195
ZNF197
ZNF225
ZNF234
ZNF250
ZNF254
ZNF268
ZNF280D
ZNF285
ZNF35
ZNF350
ZNF354B
ZNF396
ZNF418
ZNF43
ZNF430
ZNF433
ZNF439
ZNF440
ZNF443
ZNF486
ZNF502
ZNF518A
ZNF527
ZNF547
ZNF555
ZNF559
ZNF561
ZNF596
ZNF606
ZNF613
ZNF616
ZNF624
ZNF655
ZNF680
ZNF682
ZNF684
ZNF700
ZNF766
ZNF77
ZNF772
ZNF790
ZNF799
ZNF823
ZNF85
ZNF879
ZNF93
ZSCAN12
Diagnosis
Sex
Regulation Of Transcription, DNA-templated
Generic Transcription Pathway
Gene Expression
ZBTB7A
HumanNet interactions
Co-expression
Functional
Protein-protein interactions
Diagnosis
Control
Major Depression
Sex
Male
Female
Functional enrichment
Functional enrichment
Motif enrichment
RcisTarget
Transcription factor binding enrichment
ChIP-enrichR
Expression ratios
<= -2.0
-0
>= 2.0

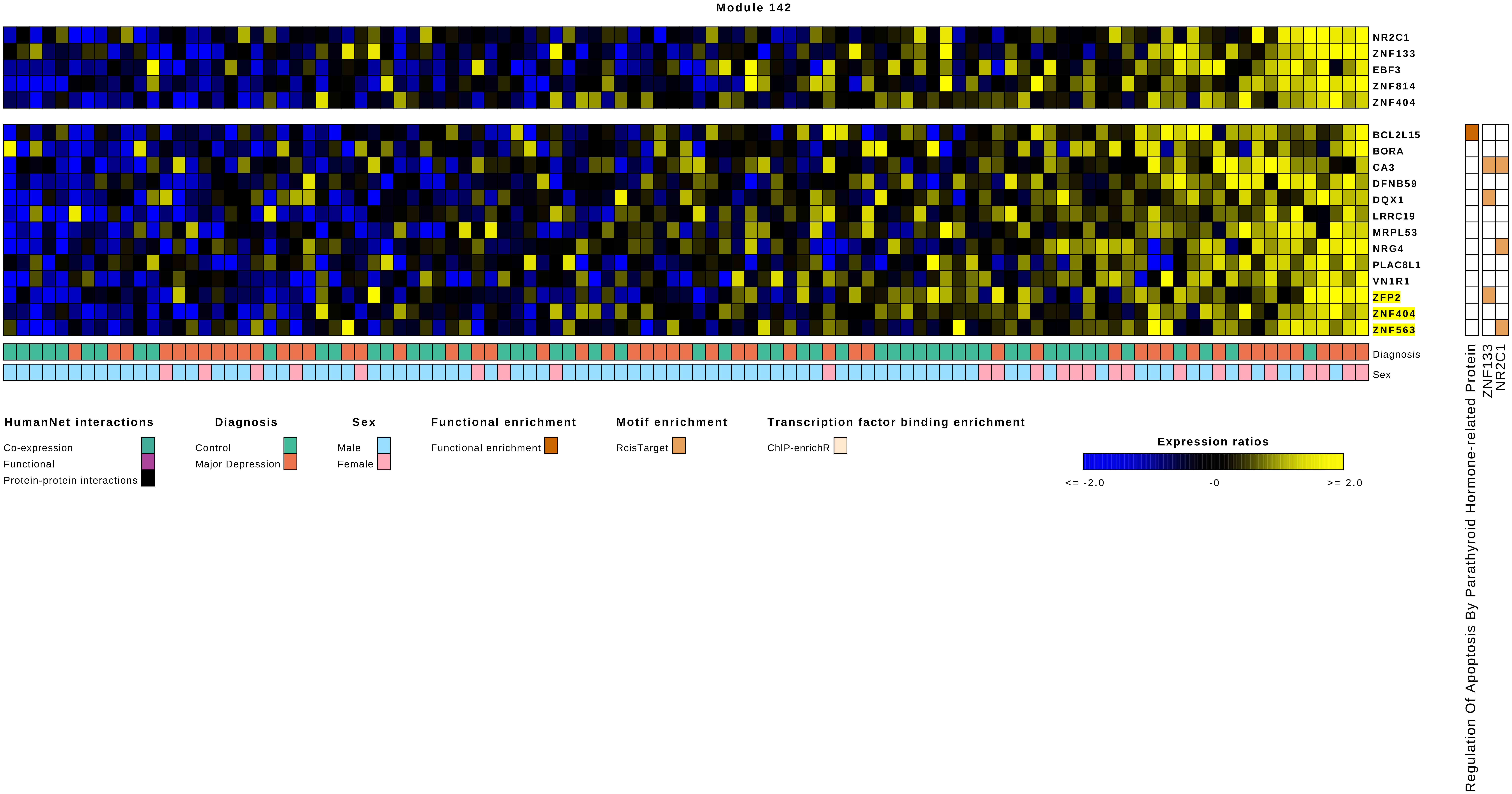

Module 142
NR2C1
ZNF133
EBF3
ZNF814
ZNF404
BCL2L15
BORA
CA3
DFNB59
DQX1
LRRC19
MRPL53
NRG4
PLAC8L1
VN1R1
ZFP2
ZNF404
ZNF563
Diagnosis
Sex
Regulation Of Apoptosis By Parathyroid Hormone-related Protein
ZNF133
NR2C1
HumanNet interactions
Co-expression
Functional
Protein-protein interactions
Diagnosis
Control
Major Depression
Sex
Male
Female
Functional enrichment
Functional enrichment
Motif enrichment
RcisTarget
Transcription factor binding enrichment
ChIP-enrichR
Expression ratios
<= -2.0
-0
>= 2.0

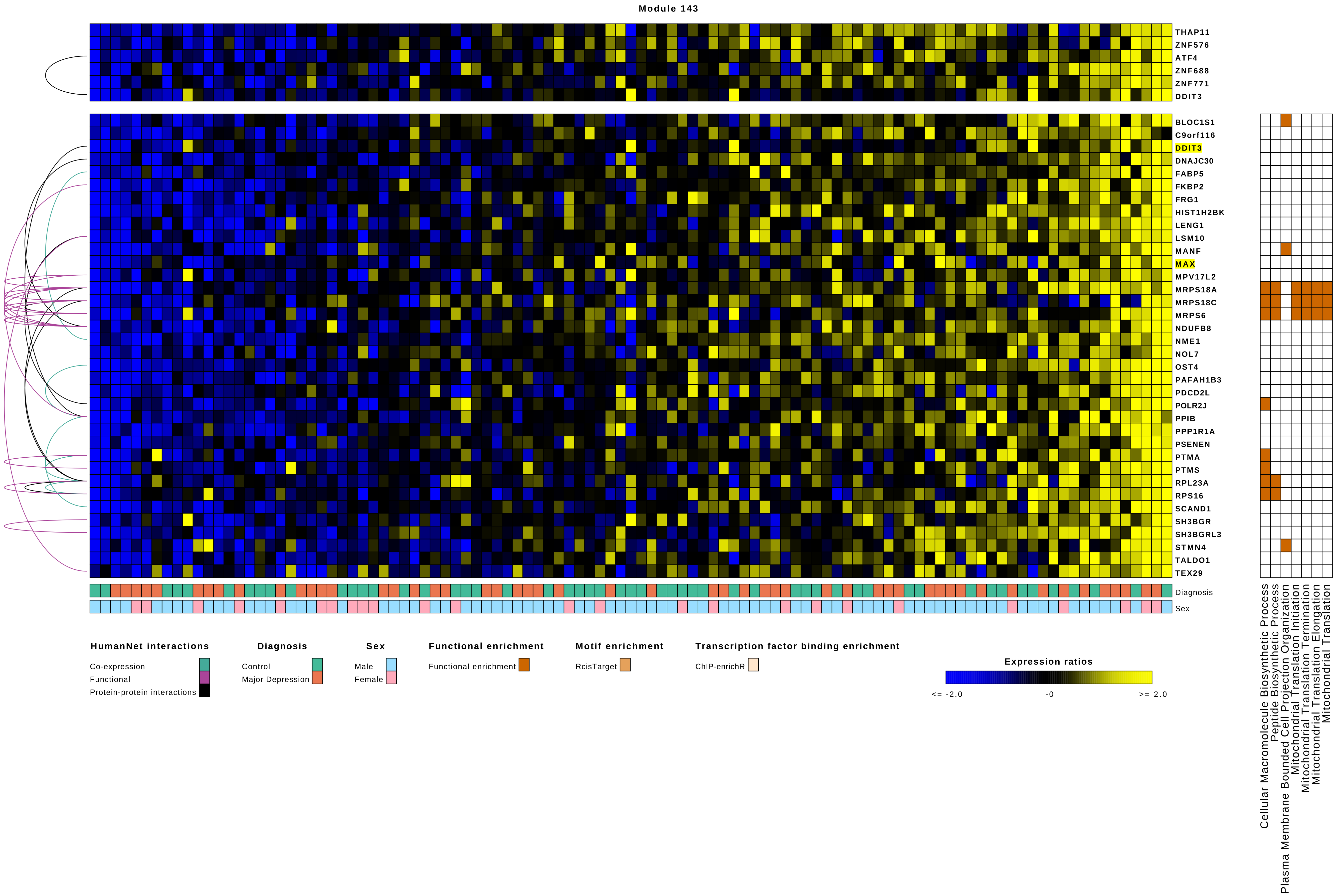

Module 143
THAP11
ZNF576
ATF4
ZNF688
ZNF771
DDIT3
BLOC1S1
C9orf116
DDIT3
DNAJC30
FABP5
FKBP2
FRG1
HIST1H2BK
LENG1
LSM10
MANF
MAX
MPV17L2
MRPS18A
MRPS18C
MRPS6
NDUFB8
NME1
NOL7
OST4
PAFAH1B3
PDCD2L
POLR2J
PPIB
PPP1R1A
PSENEN
PTMA
PTMS
RPL23A
RPS16
SCAND1
SH3BGR
SH3BGRL3
STMN4
TALDO1
TEX29
Diagnosis
Sex
Cellular Macromolecule Biosynthetic Process
Peptide Biosynthetic Process
Plasma Membrane Bounded Cell Projection Organization
Mitochondrial Translation Initiation
Mitochondrial Translation Termination
Mitochondrial Translation Elongation
Mitochondrial Translation
HumanNet interactions
Co-expression
Functional
Protein-protein interactions
Diagnosis
Control
Major Depression
Sex
Male
Female
Functional enrichment
Functional enrichment
Motif enrichment
RcisTarget
Transcription factor binding enrichment
ChIP-enrichR
Expression ratios
<= -2.0
-0
>= 2.0

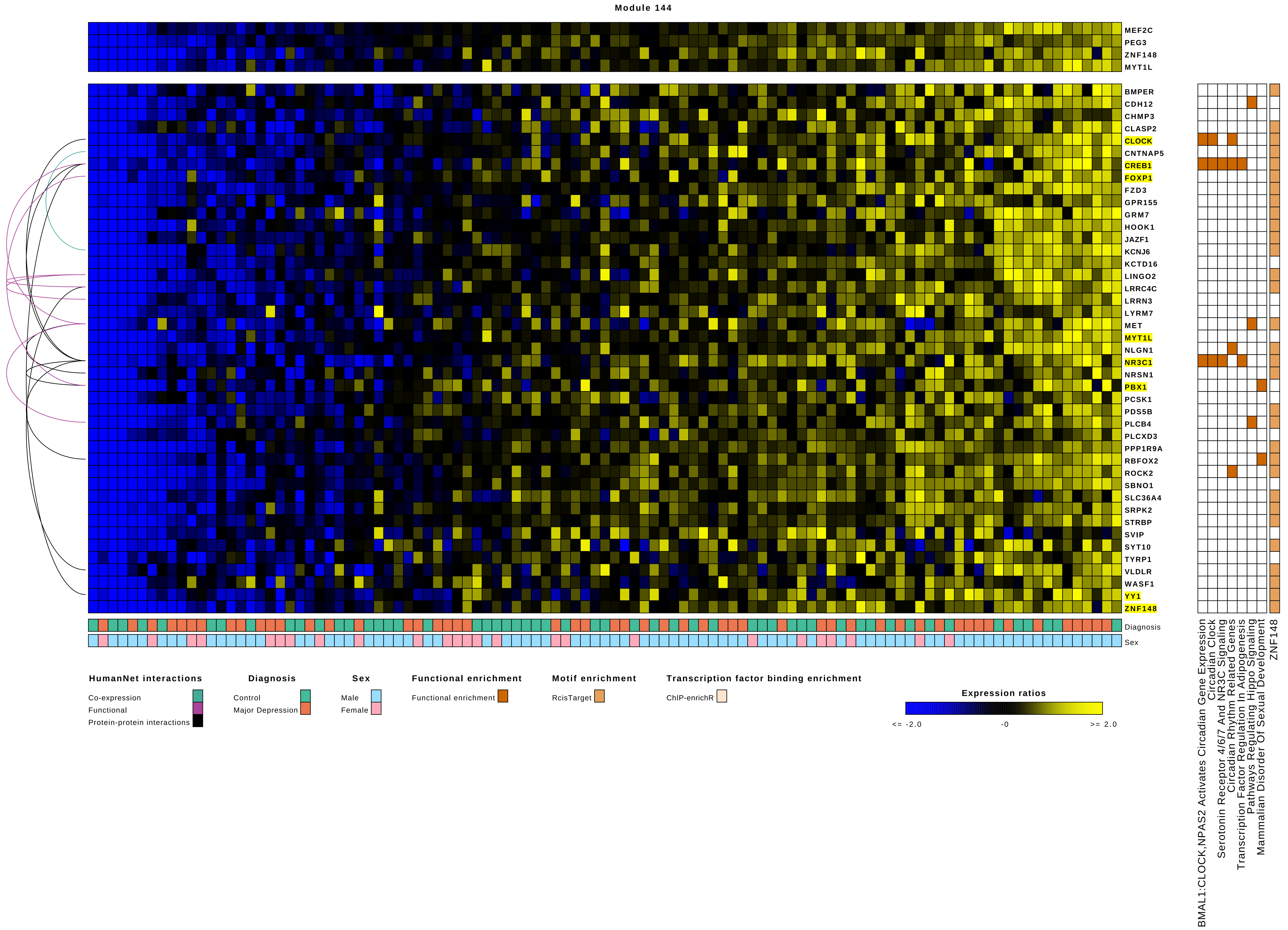

Module 144
MEF2C
PEG3
ZNF148
MYT1L
BMPER
CDH12
CHMP3
CLASP2
CLOCK
CNTNAP5
CREB1
FOXP1
FZD3
GPR155
GRM7
HOOK1
JAZF1
KCNJ6
KCTD16
LINGO2
LRRC4C
LRRN3
LYRM7
MET
MYT1L
NLGN1
NR3C1
NRSN1
PBX1
PCSK1
PDS5B
PLCB4
PLCXD3
PPP1R9A
RBFOX2
ROCK2
SBNO1
SLC36A4
SRPK2
STRBP
SVIP
SYT10
TYRP1
VLDLR
WASF1
YY1
ZNF148
Diagnosis
Sex
BMAL1:CLOCK,NPAS2 Activates Circadian Gene Expression
Circadian Clock
Serotonin Receptor 4/6/7 And NR3C Signaling
Circadian Rhythm Related Genes
Transcription Factor Regulation In Adipogenesis
Pathways Regulating Hippo Signaling
Mammalian Disorder Of Sexual Development
ZNF148
HumanNet interactions
Co-expression
Functional
Protein-protein interactions
Diagnosis
Control
Major Depression
Sex
Male
Female
Functional enrichment
Functional enrichment
Motif enrichment
RcisTarget
Transcription factor binding enrichment
ChIP-enrichR
Expression ratios
<= -2.0
-0
>= 2.0

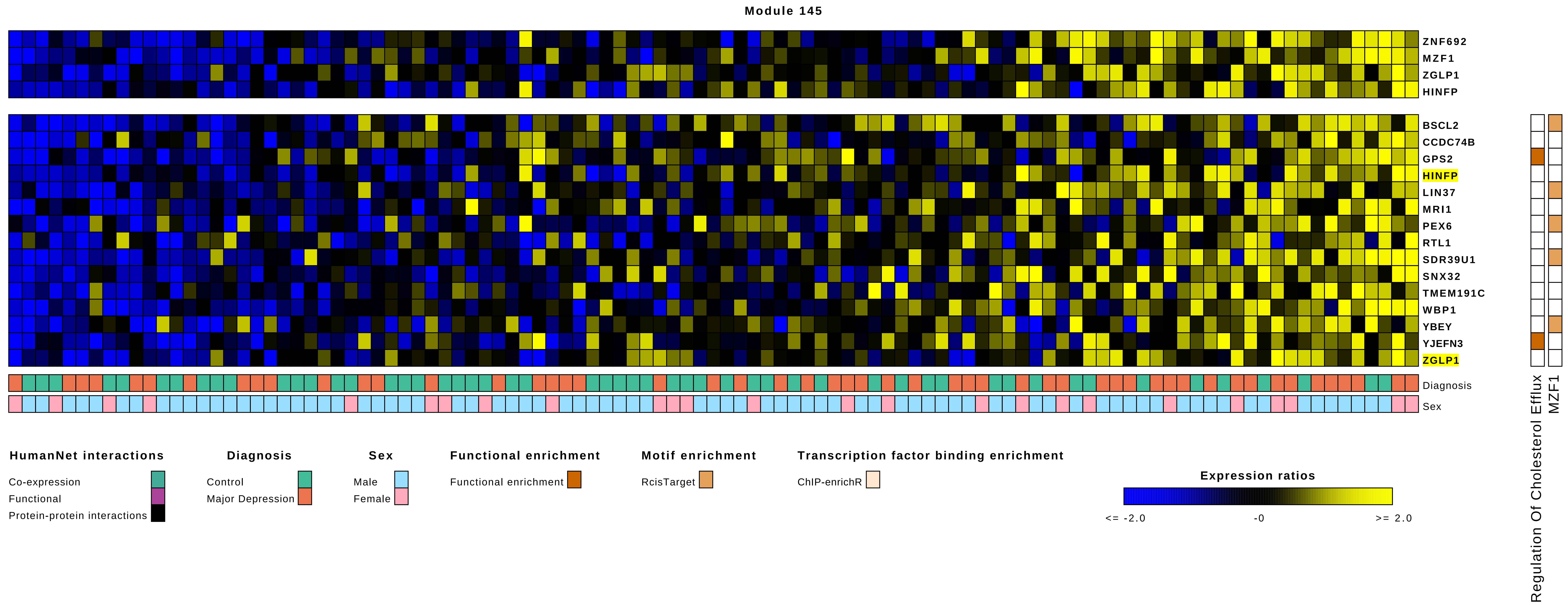
Module 145
ZNF692
MZF1
ZGLP1
HINFP
BSCL2
CCDC74B
GPS2
HINFP
LIN37
MRI1
PEX6
RTL1
SDR39U1
SNX32
TMEM191C
WBP1
YBEY
YJEFN3
ZGLP1
Diagnosis
Sex
Regulation Of Cholesterol Efflux
MZF1
HumanNet interactions
Co-expression
Functional
Protein-protein interactions
Diagnosis
Control
Major Depression
Sex
Male
Female
Functional enrichment
Functional enrichment
Motif enrichment
RcisTarget
Transcription factor binding enrichment
ChIP-enrichR
Expression ratios
<= -2.0
-0
>= 2.0

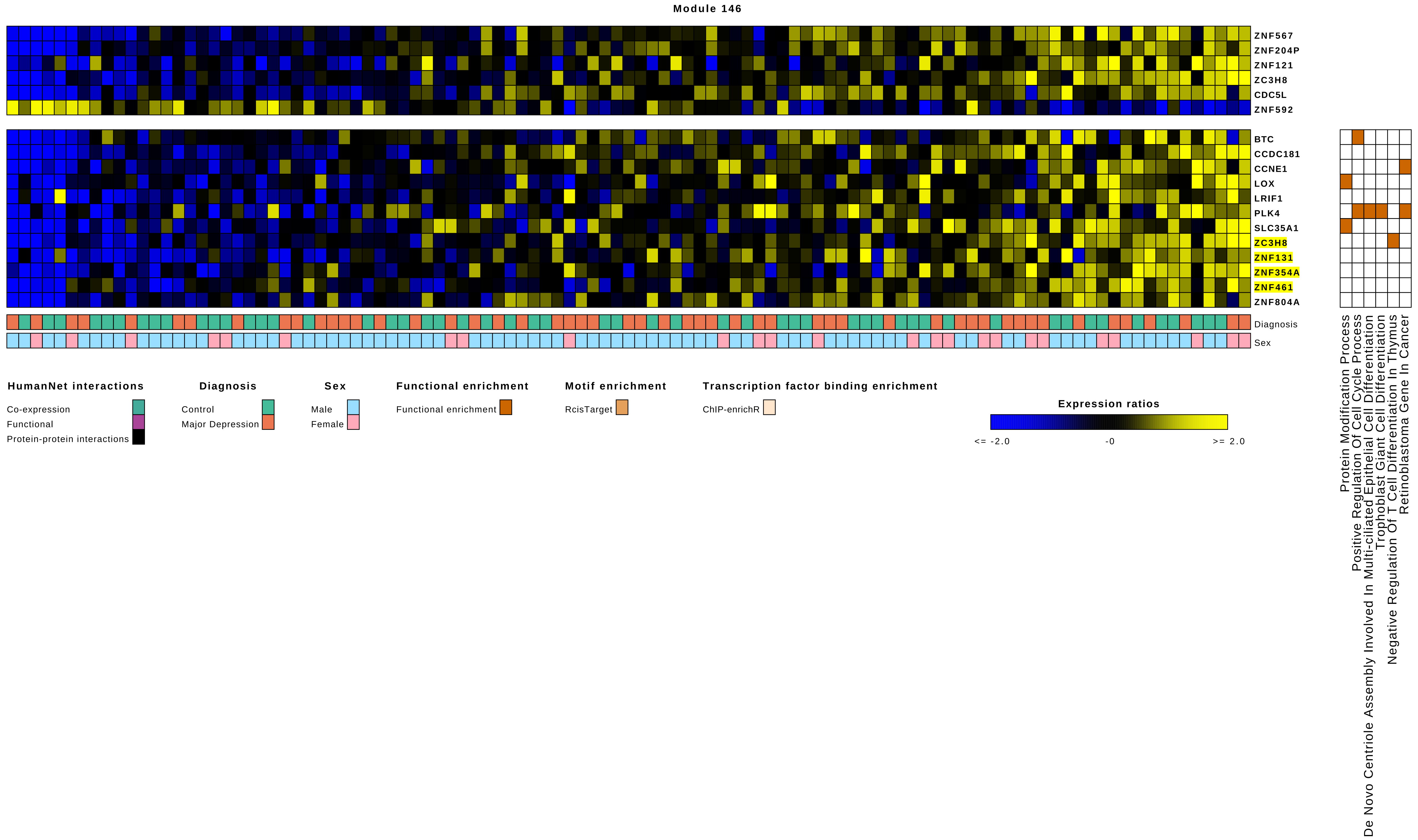
Module 146
ZNF567
ZNF204P
ZNF121
ZC3H8
CDC5L
ZNF592
BTC
CCDC181
CCNE1
LOX
LRIF1
PLK4
SLC35A1
ZC3H8
ZNF131
ZNF354A
ZNF461
ZNF804A
Diagnosis
Sex
Protein Modification Process
Positive Regulation Of Cell Cycle Process
De Novo Centriole Assembly Involved In Multi-ciliated Epithelial Cell Differentiation
Trophoblast Giant Cell Differentiation
Negative Regulation Of T Cell Differentiation In Thymus
Retinoblastoma Gene In Cancer
HumanNet interactions
Co-expression
Functional
Protein-protein interactions
Diagnosis
Control
Major Depression
Sex
Male
Female
Functional enrichment
Functional enrichment
Motif enrichment
RcisTarget
Transcription factor binding enrichment
ChIP-enrichR
Expression ratios
<= -2.0
-0
>= 2.0

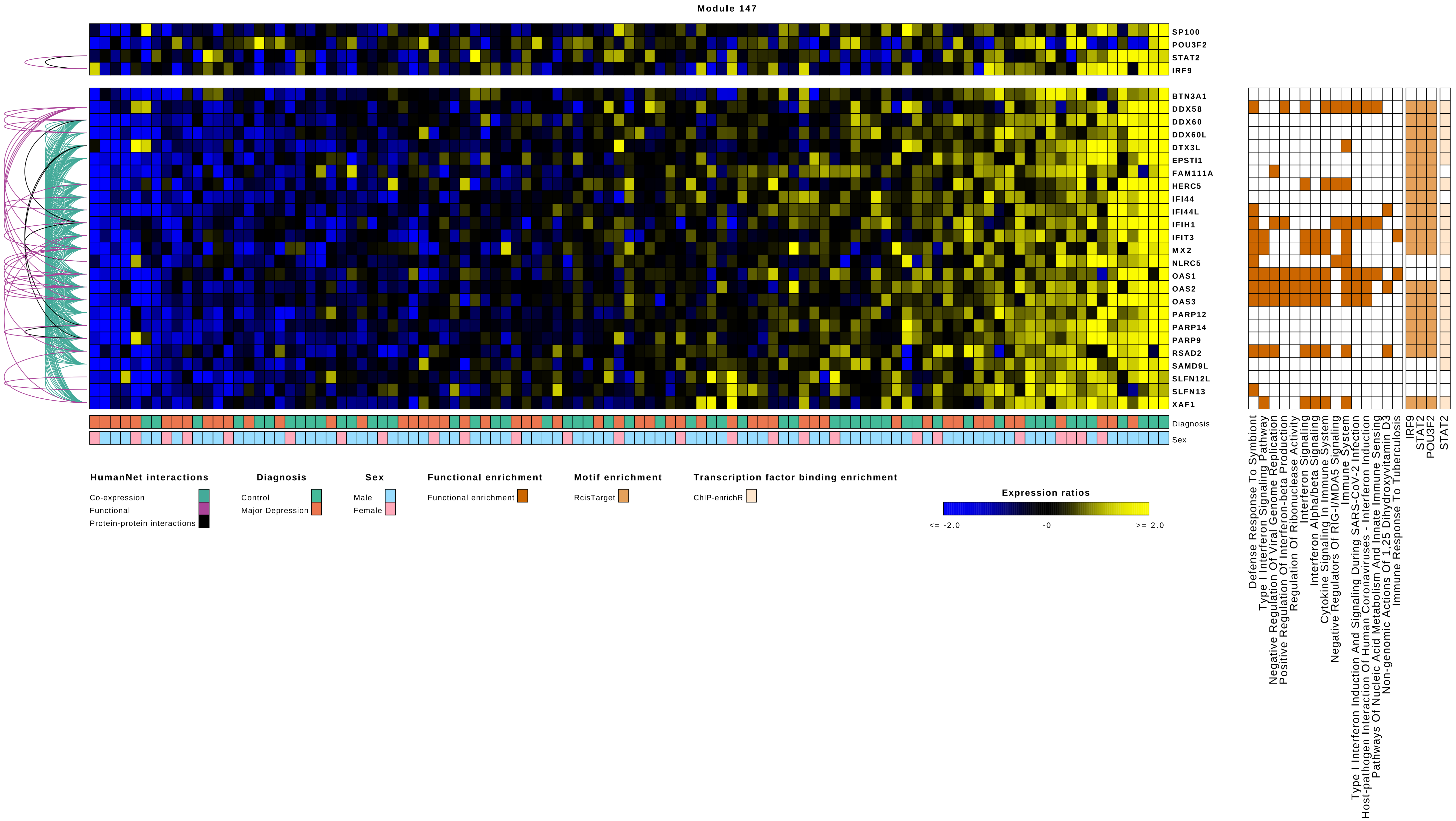
Module 147
SP100
POU3F2
STAT2
IRF9
BTN3A1
DDX58
DDX60
DDX60L
DTX3L
EPSTI1
FAM111A
HERC5
IFI44
IFI44L
IFIH1
IFIT3
MX2
NLRC5
OAS1
OAS2
OAS3
PARP12
PARP14
PARP9
RSAD2
SAMD9L
SLFN12L
SLFN13
XAF1
Diagnosis
Sex
HumanNet interactions
Co-expression
Functional
Protein-protein interactions
Diagnosis
Control
Major Depression
Sex
Male
Female
Functional enrichment
Functional enrichment
Motif enrichment
RcisTarget
Transcription factor binding enrichment
ChIP-enrichR
Expression ratios
<= -2.0
-0
>= 2.0
Defense Response To Symbiont
Type I Interferon Signaling Pathway
Negative Regulation Of Viral Genome Replication
Positive Regulation Of Interferon-beta Production
Regulation Of Ribonuclease Activity
Interferon Signaling
Interferon Alpha/beta Signaling
Cytokine Signaling In Immune System
Negative Regulators Of RIG-I/MDA5 Signaling
Immune System
Type I Interferon Induction And Signaling During SARS-CoV-2 Infection
Host-pathogen Interaction Of Human Coronaviruses - Interferon Induction
Pathways Of Nucleic Acid Metabolism And Innate Immune Sensing
Non-genomic Actions Of 1,25 Dihydroxyvitamin D3
Immune Response To Tuberculosis
IRF9
STAT2
POU3F2
STAT2

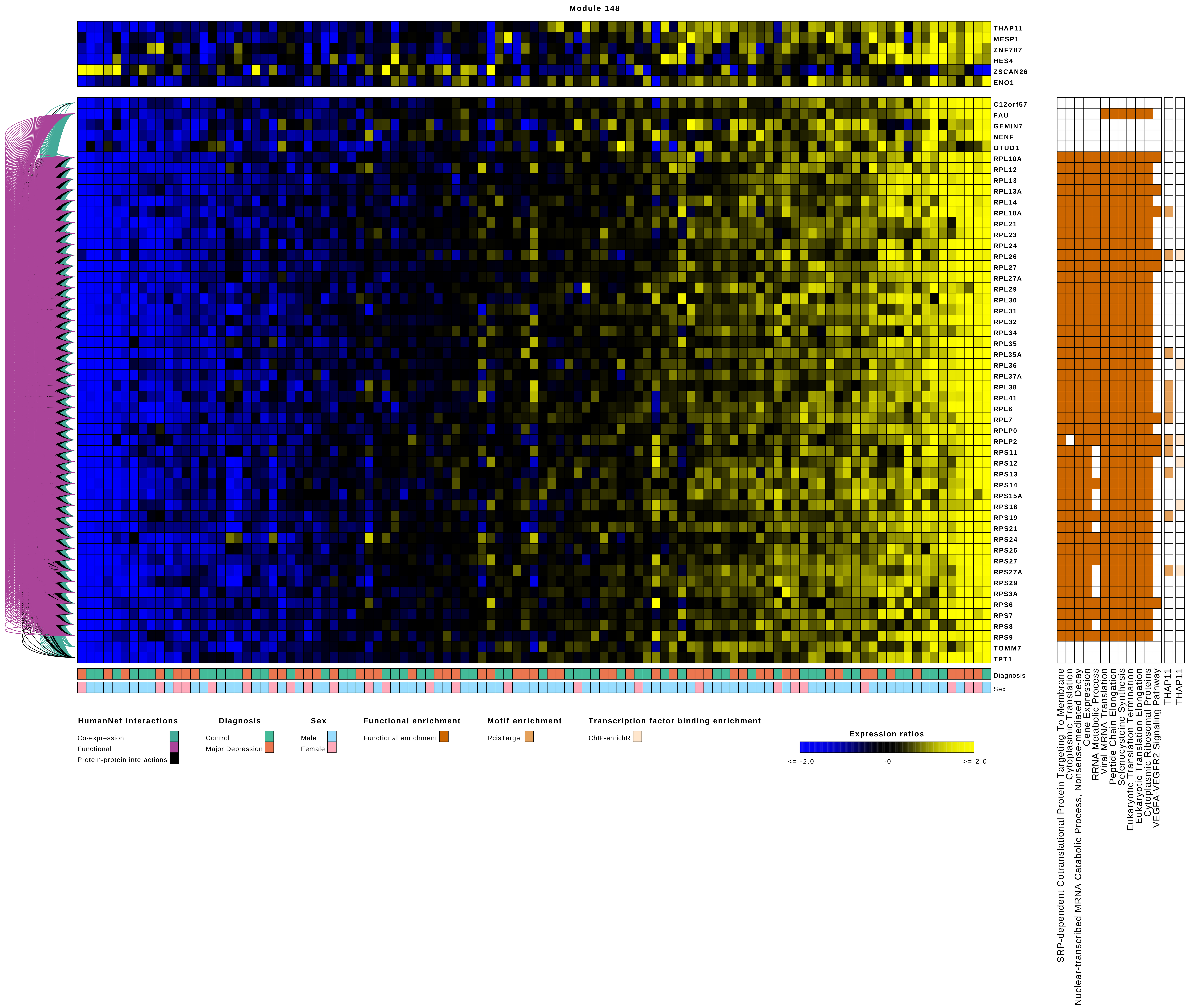
Module 148
THAP11
MESP1
ZNF787
HES4
ZSCAN26
ENO1
C12orf57
FAU
GEMIN7
NENF
OTUD1
RPL10A
RPL12
RPL13
RPL13A
RPL14
RPL18A
RPL21
RPL23
RPL24
RPL26
RPL27
RPL27A
RPL29
RPL30
RPL31
RPL32
RPL34
RPL35
RPL35A
RPL36
RPL37A
RPL38
RPL41
RPL6
RPL7
RPLP0
RPLP2
RPS11
RPS12
RPS13
RPS14
RPS15A
RPS18
RPS19
RPS21
RPS24
RPS25
RPS27
RPS27A
RPS29
RPS3A
RPS6
RPS7
RPS8
RPS9
TOMM7
TPT1
Diagnosis
Sex
SRP-dependent Cotranslational Protein Targeting To Membrane
Cytoplasmic Translation
Nuclear-transcribed MRNA Catabolic Process, Nonsense-mediated Decay
Gene Expression
RRNA Metabolic Process
Viral MRNA Translation
Peptide Chain Elongation
Selenocysteine Synthesis
Eukaryotic Translation Termination
Eukaryotic Translation Elongation
Cytoplasmic Ribosomal Proteins
VEGFA-VEGFR2 Signaling Pathway
THAP11
THAP11
HumanNet interactions
Co-expression
Functional
Protein-protein interactions
Diagnosis
Control
Major Depression
Sex
Male
Female
Functional enrichment
Functional enrichment
Motif enrichment
RcisTarget
Transcription factor binding enrichment
ChIP-enrichR
Expression ratios
<= -2.0
-0
>= 2.0

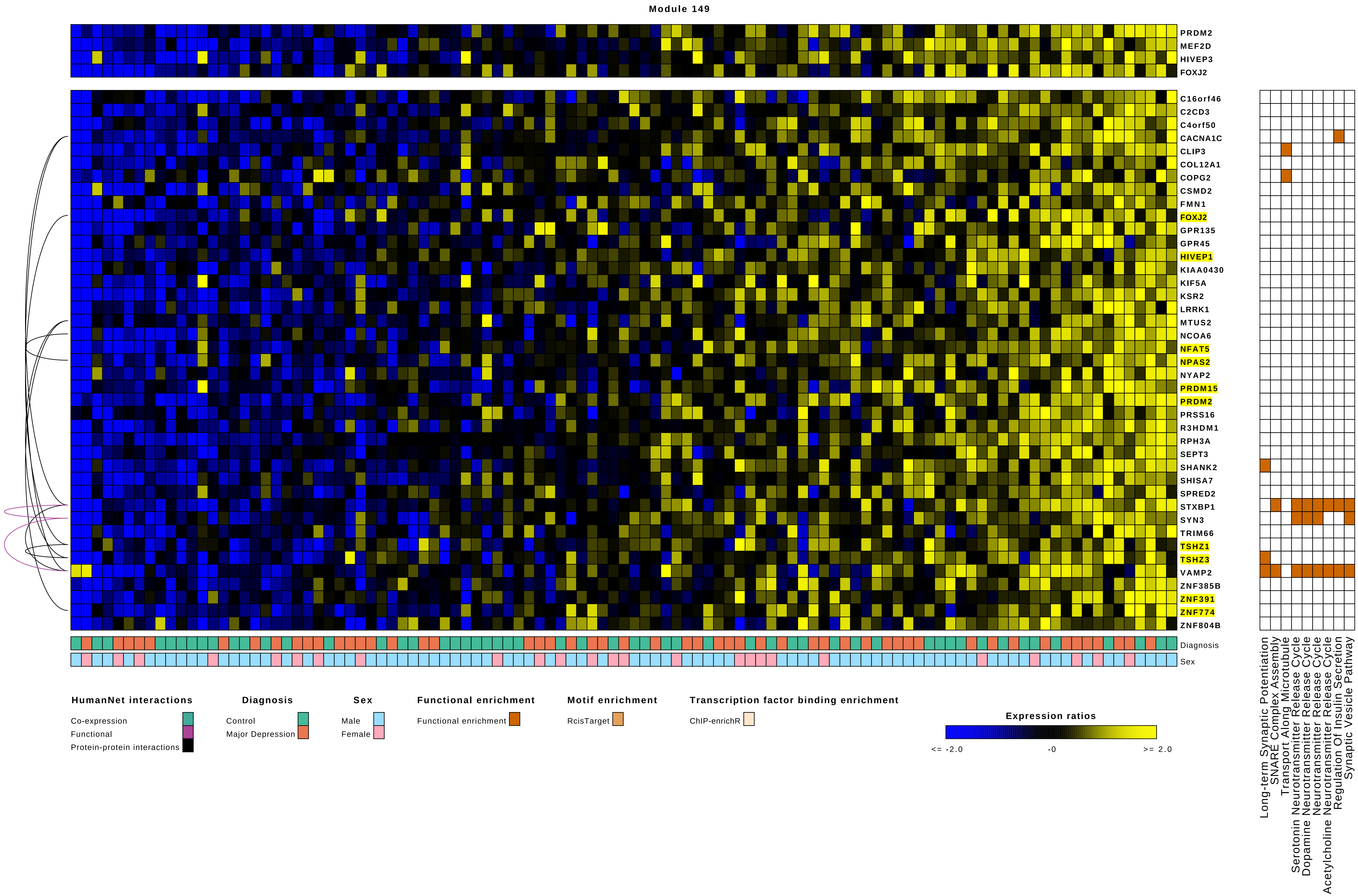

Module 149
PRDM2
MEF2D
HIVEP3
FOXJ2
C16orf46
C2CD3
C4orf50
CACNA1C
CLIP3
COL12A1
COPG2
CSMD2
FMN1
FOXJ2
GPR135
GPR45
HIVEP1
KIAA0430
KIF5A
KSR2
LRRK1
MTUS2
NCOA6
NFAT5
NPAS2
NYAP2
PRDM15
PRDM2
PRSS16
R3HDM1
RPH3A
SEPT3
SHANK2
SHISA7
SPRED2
STXBP1
SYN3
TRIM66
TSHZ1
TSHZ3
VAMP2
ZNF385B
ZNF391
ZNF774
ZNF804B
Diagnosis
Sex
Long-term Synaptic Potentiation
SNARE Complex Assembly
Transport Along Microtubule
Serotonin Neurotransmitter Release Cycle
Dopamine Neurotransmitter Release Cycle
Neurotransmitter Release Cycle
Acetylcholine Neurotransmitter Release Cycle
Regulation Of Insulin Secretion
Synaptic Vesicle Pathway
HumanNet interactions
Co-expression
Functional
Protein-protein interactions
Diagnosis
Control
Major Depression
Sex
Male
Female
Functional enrichment
Functional enrichment
Motif enrichment
RcisTarget
Transcription factor binding enrichment
ChIP-enrichR
Expression ratios
<= -2.0
-0
>= 2.0

**Module 150**

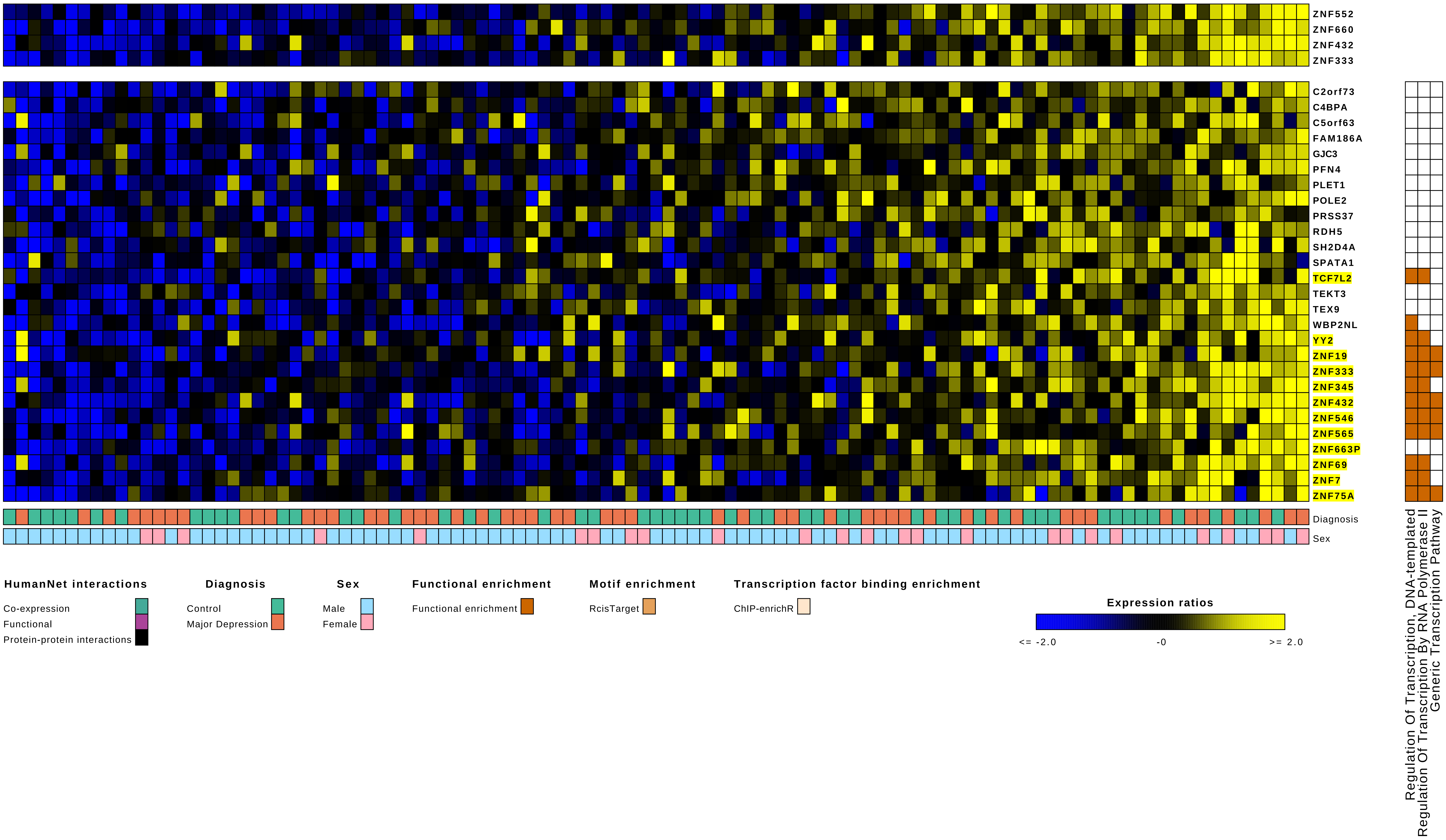
ZNF552
ZNF660
ZNF432
ZNF333
C2orf73
C4BPA
C5orf63
FAM186A
GJC3
PFN4
PLET1
POLE2
PRSS37
RDH5
SH2D4A
SPATA1
TCF7L2
TEKT3
TEX9
WBP2NL
YY2
ZNF19
ZNF333
ZNF345
ZNF432
ZNF546
ZNF565
ZNF663P
ZNF69
ZNF7
ZNF75A
Diagnosis
Sex
Regulation Of Transcription, DNA-templated
Regulation Of Transcription By RNA Polymerase II
Generic Transcription Pathway
HumanNet interactions
Co-expression
Functional
Protein-protein interactions
Diagnosis
Control
Major Depression
Sex
Male
Female
Functional enrichment
Functional enrichment
Motif enrichment
RcisTarget
Transcription factor binding enrichment
ChIP-enrichR
Expression ratios
<= -2.0
-0
>= 2.0

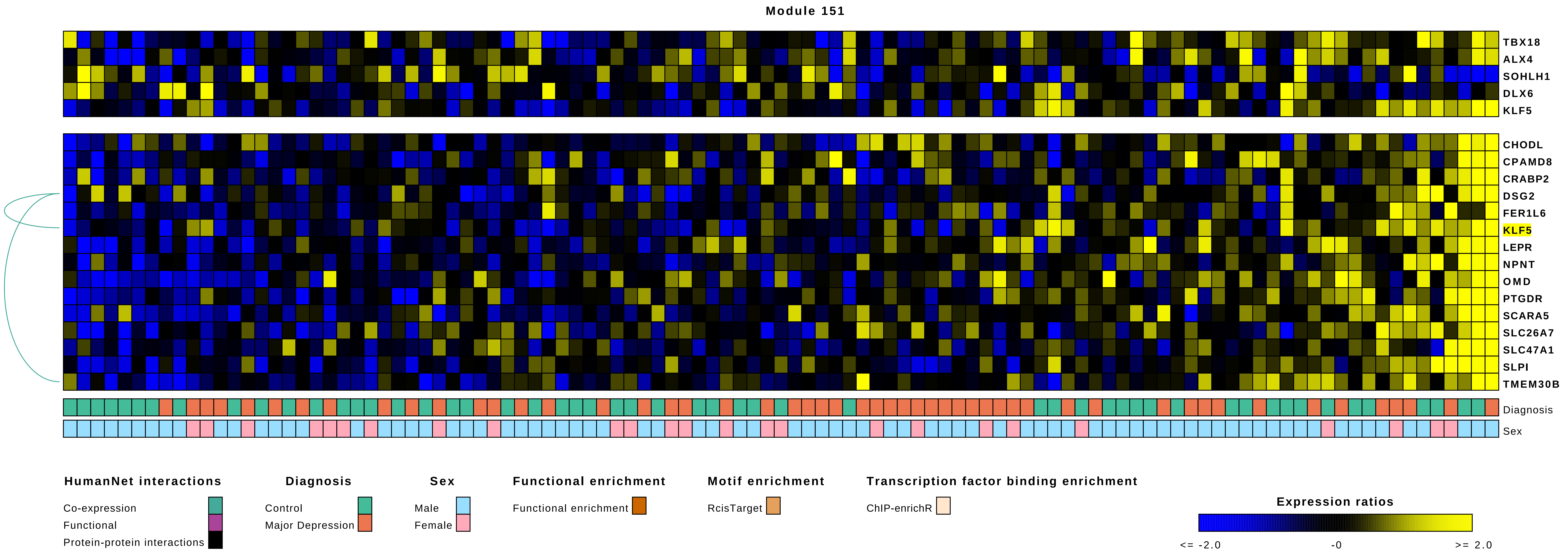
Module 151
TBX18
ALX4
SOHLH1
DLX6
KLF5
CHODL
CPAMD8
CRABP2
DSG2
FER1L6
KLF5
LEPR
NPNT
OMD
PTGDR
SCARA5
SLC26A7
SLC47A1
SLPI
TMEM30B
Diagnosis
Sex
HumanNet interactions
Co-expression
Functional
Protein-protein interactions
Diagnosis
Control
Major Depression
Sex
Male
Female
Functional enrichment
Functional enrichment
Motif enrichment
RcisTarget
Transcription factor binding enrichment
ChIP-enrichR
Expression ratios
<= -2.0
-0
>= 2.0

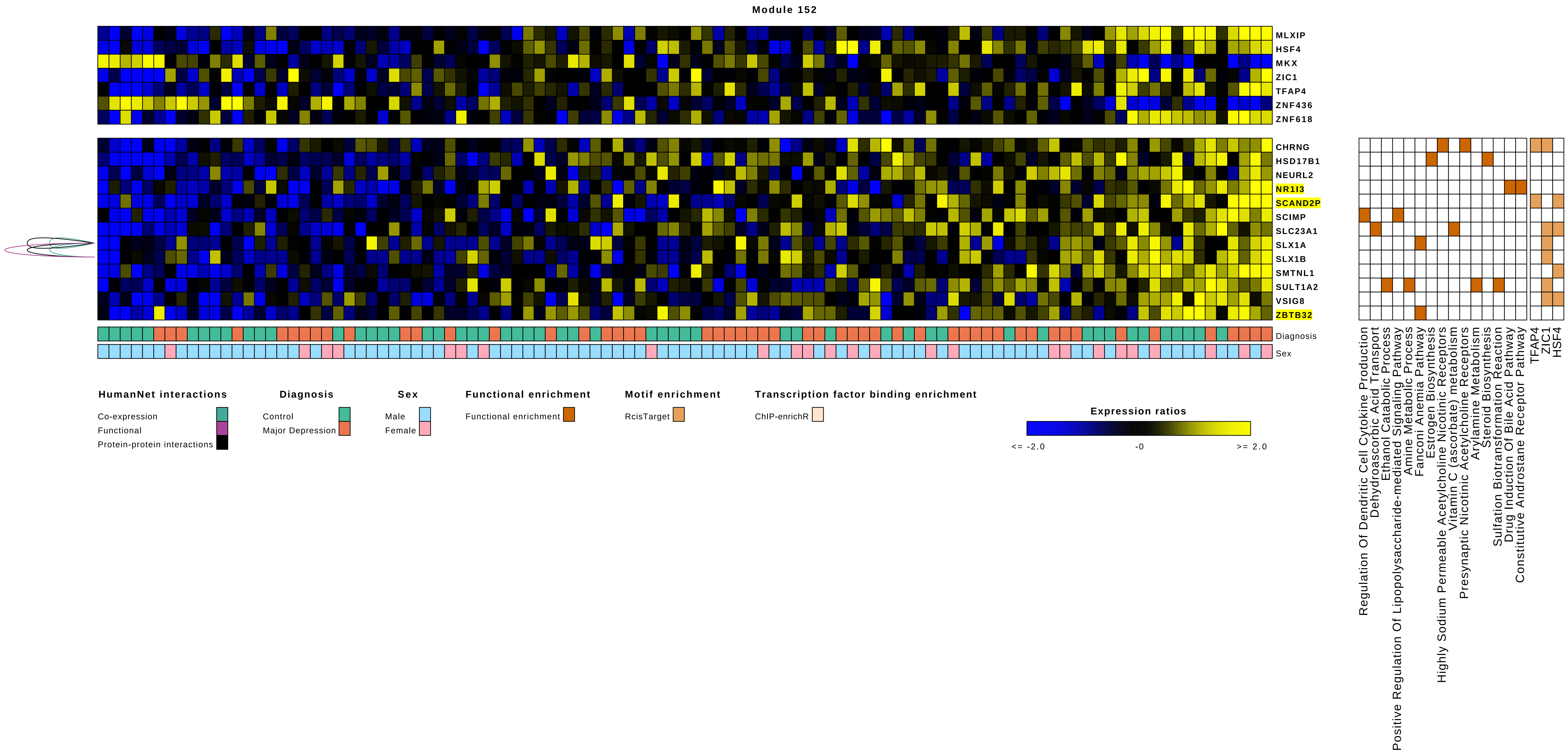

Module 152
MLXIP
HSF4
MKX
ZIC1
TFAP4
ZNF436
ZNF618
CHRNG
HSD17B1
NEURL2
NR1I3
SCAND2P
SCIMP
SLC23A1
SLX1A
SLX1B
SMTNL1
SULT1A2
VSIG8
ZBTB32
Diagnosis
Sex
Regulation Of Dendritic Cell Cytokine Production
Dehydroascorbic Acid Transport
Ethanol Catabolic Process
Positive Regulation Of Lipopolysaccharide-mediated Signaling Pathway
Amine Metabolic Process
Fanconi Anemia Pathway
Estrogen Biosynthesis
Highly Sodium Permeable Acetylcholine Nicotinic Receptors
Vitamin C (ascorbate) metabolism
Presynaptic Nicotinic Acetylcholine Receptors
Arylamine Metabolism
Steroid Biosynthesis
Sulfation Biotransformation Reaction
Drug Induction Of Bile Acid Pathway
Constitutive Androstane Receptor Pathway
TFAP4
ZIC1
HSF4
HumanNet interactions
Co-expression
Functional
Protein-protein interactions
Diagnosis
Control
Major Depression
Sex
Male
Female
Functional enrichment
Functional enrichment
Motif enrichment
RcisTarget
Transcription factor binding enrichment
ChIP-enrichR
Expression ratios
<= -2.0
-0
>= 2.0

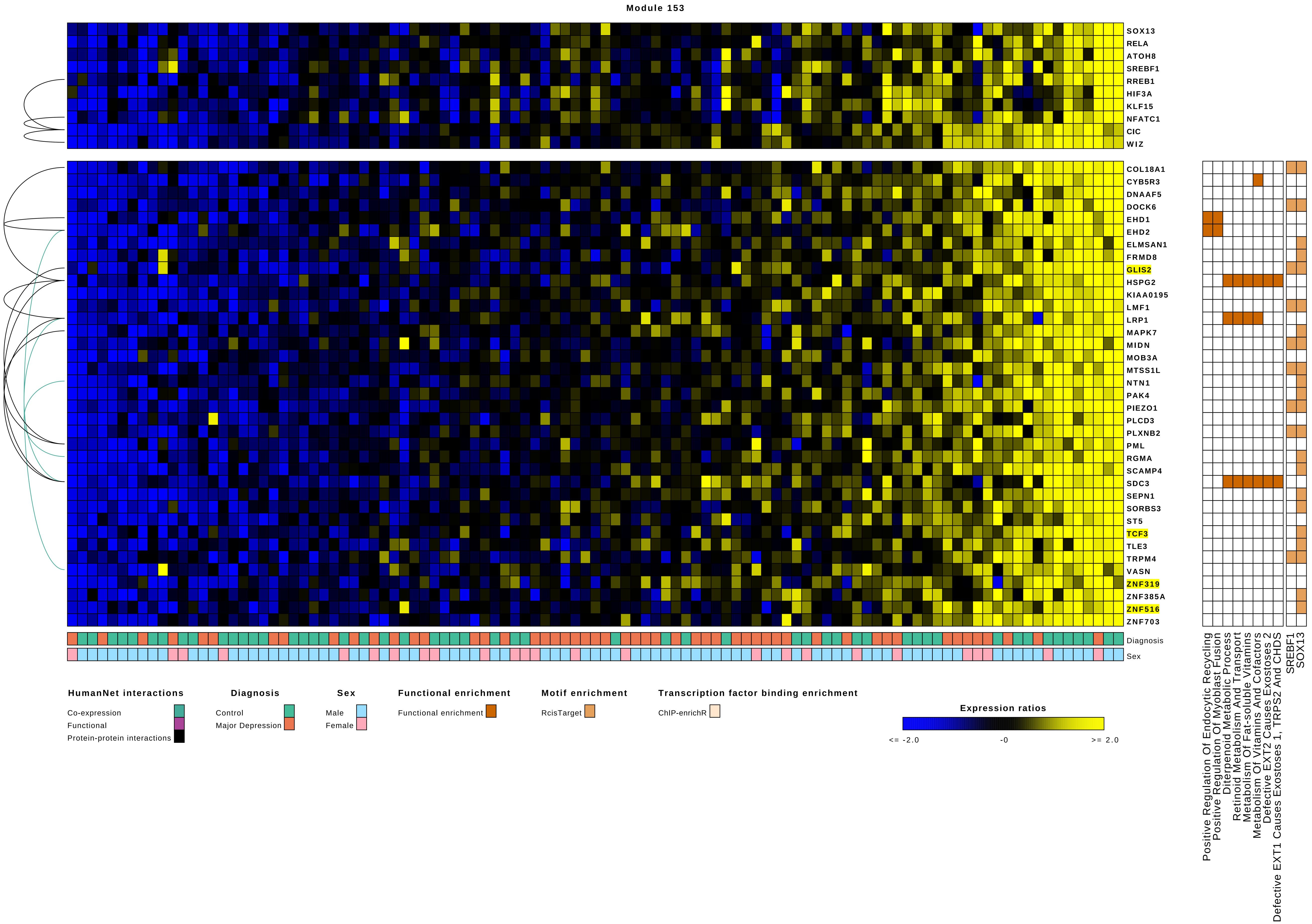

Module 153
SOX13
RELA
ATOH8
SREBF1
RREB1
HIF3A
KLF15
NFATC1
CIC
WIZ
COL18A1
CYB5R3
DNAAF5
DOCK6
EHD1
EHD2
ELMSAN1
FRMD8
GLIS2
HSPG2
KIAA0195
LMF1
LRP1
MAPK7
MIDN
MOB3A
MTSS1L
NTN1
PAK4
PIEZO1
PLCD3
PLXNB2
PML
RGMA
SCAMP4
SDC3
SEPN1
SORBS3
ST5
TCF3
TLE3
TRPM4
VASN
ZNF319
ZNF385A
ZNF516
ZNF703
Diagnosis
Sex
Positive Regulation Of Endocytic Recycling
Positive Regulation Of Myoblast Fusion
Diterpenoid Metabolic Process
Retinoid Metabolism And Transport
Metabolism Of Fat-soluble Vitamins
Metabolism Of Vitamins And Cofactors
Defective EXT2 Causes Exostoses 2
Defective EXT1 Causes Exostoses 1, TRPS2 And CHDS
SREBF1
SOX13
HumanNet interactions
Co-expression
Functional
Protein-protein interactions
Diagnosis
Control
Major Depression
Sex
Male
Female
Functional enrichment
Functional enrichment
Motif enrichment
RcisTarget
Transcription factor binding enrichment
ChIP-enrichR
Expression ratios
<= -2.0
-0
>= 2.0

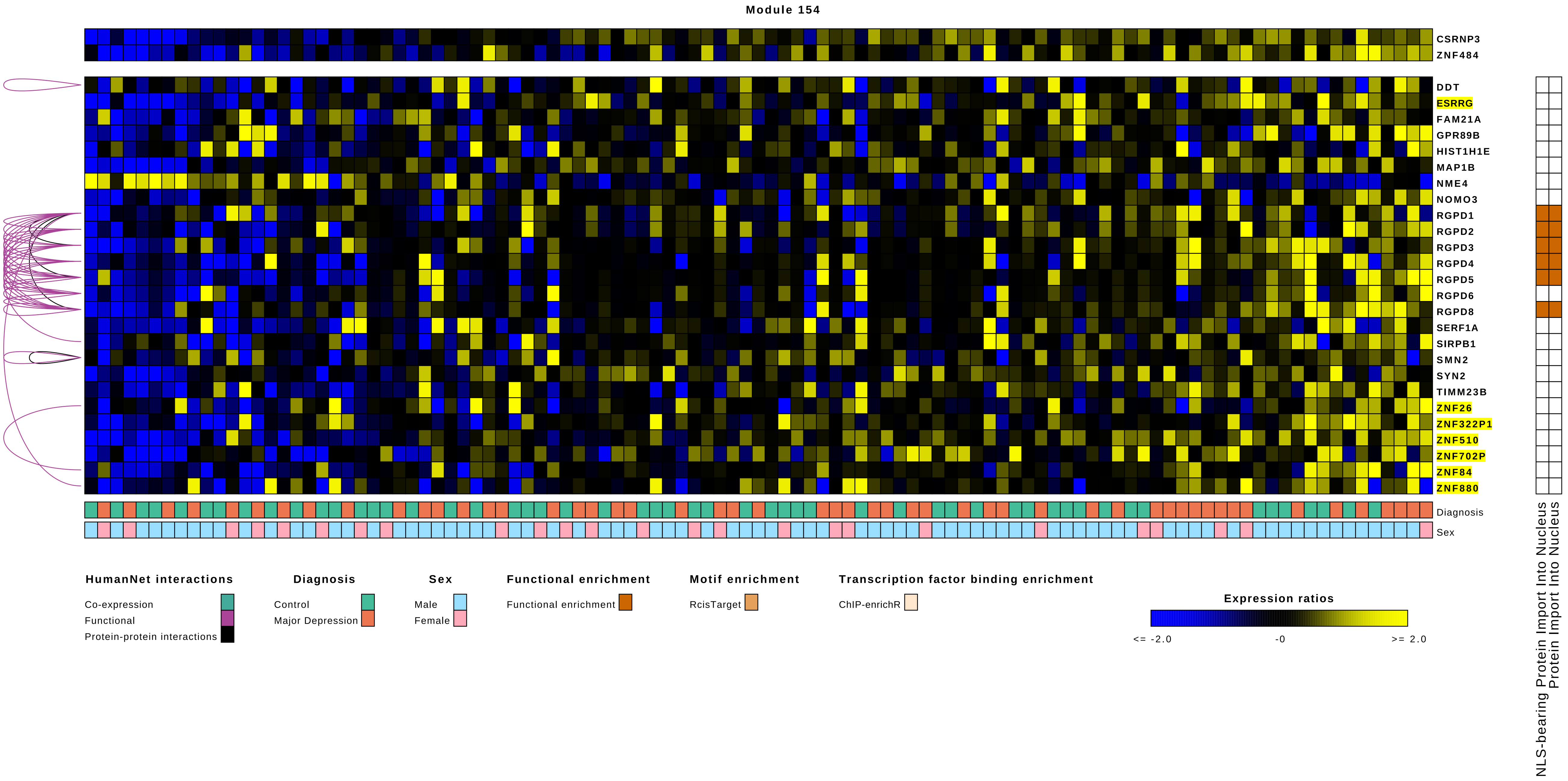
Module 154
CSRNP3
ZNF484
DDT
ESRRG
FAM21A
GPR89B
HIST1H1E
MAP1B
NME4
NOMO3
RGPD1
RGPD2
RGPD3
RGPD4
RGPD5
RGPD6
RGPD8
SERF1A
SIRPB1
SMN2
SYN2
TIMM23B
ZNF26
ZNF322P1
ZNF510
ZNF702P
ZNF84
ZNF880
Diagnosis
Sex
NLS-bearing Protein Import Into Nucleus
Protein Import Into Nucleus
HumanNet interactions
Co-expression
Functional
Protein-protein interactions
Diagnosis
Control
Major Depression
Sex
Male
Female
Functional enrichment
Functional enrichment
Motif enrichment
RcisTarget
Transcription factor binding enrichment
ChIP-enrichR
Expression ratios
<= -2.0
-0
>= 2.0

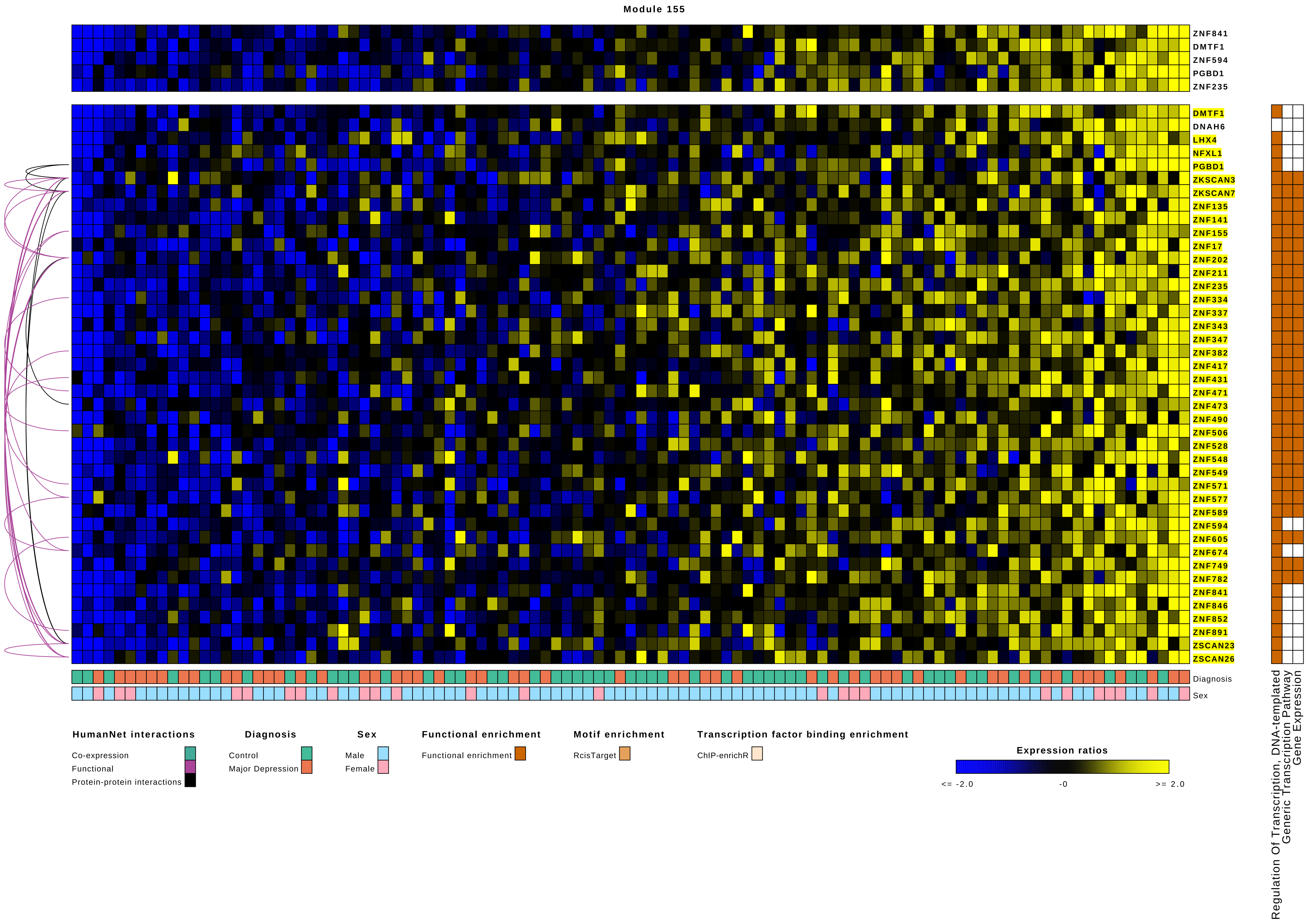
Module 155
ZNF841
DMTF1
ZNF594
PGBD1
ZNF235
DMTF1
DNAH6
LHX4
NFXL1
PGBD1
ZKSCAN3
ZKSCAN7
ZNF135
ZNF141
ZNF155
ZNF17
ZNF202
ZNF211
ZNF235
ZNF334
ZNF337
ZNF343
ZNF347
ZNF382
ZNF417
ZNF431
ZNF471
ZNF473
ZNF490
ZNF506
ZNF528
ZNF548
ZNF549
ZNF571
ZNF577
ZNF589
ZNF594
ZNF605
ZNF674
ZNF749
ZNF782
ZNF841
ZNF846
ZNF852
ZNF891
ZSCAN23
ZSCAN26
Diagnosis
Sex
Regulation Of Transcription, DNA-templated
Generic Transcription Pathway
Gene Expression
HumanNet interactions
Co-expression
Functional
Protein-protein interactions
Diagnosis
Control
Major Depression
Sex
Male
Female
Functional enrichment
Functional enrichment
Motif enrichment
RcisTarget
Transcription factor binding enrichment
ChIP-enrichR
Expression ratios
<= -2.0
-0
>= 2.0

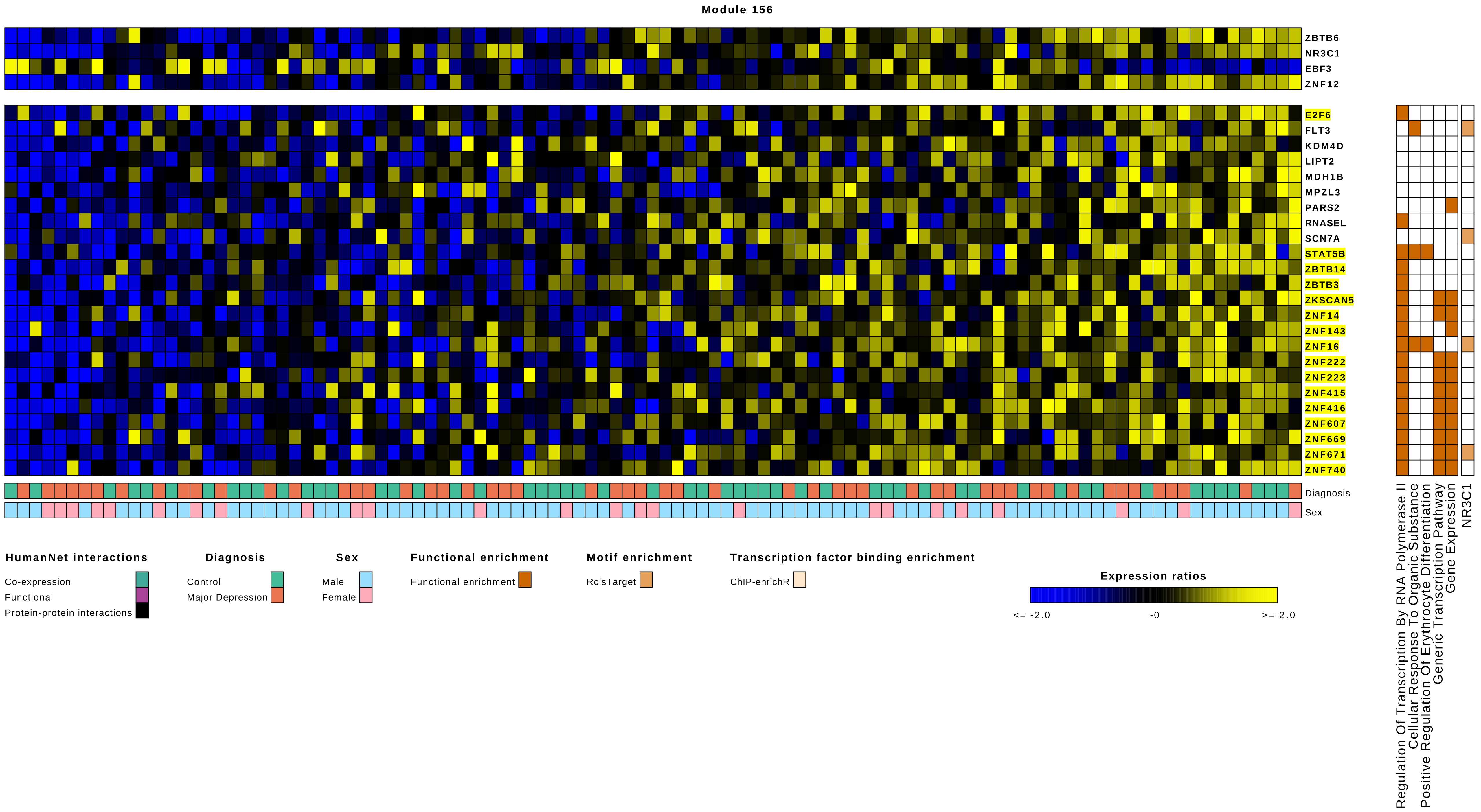

Module 156
ZBTB6
NR3C1
EBF3
ZNF12
E2F6
FLT3
KDM4D
LIPT2
MDH1B
MPZL3
PARS2
RNASEL
SCN7A
STAT5B
ZBTB14
ZBTB3
ZKSCAN5
ZNF14
ZNF143
ZNF16
ZNF222
ZNF223
ZNF415
ZNF416
ZNF607
ZNF669
ZNF671
ZNF740
Diagnosis
Sex
Regulation Of Transcription By RNA Polymerase II
Cellular Response To Organic Substance
Positive Regulation Of Erythrocyte Differentiation
Generic Transcription Pathway
Gene Expression
NR3C1
HumanNet interactions
Co-expression
Functional
Protein-protein interactions
Diagnosis
Control
Major Depression
Sex
Male
Female
Functional enrichment
Functional enrichment
Motif enrichment
RcisTarget
Transcription factor binding enrichment
ChIP-enrichR
Expression ratios
<= -2.0
-0
>= 2.0
